# Delineation of the SUMO-Modified Proteome Reveals Regulatory Functions Throughout Meiosis

**DOI:** 10.1101/828442

**Authors:** Nikhil Bhagwat, Shannon Owens, Masaru Ito, Jay Boinapalli, Philip Poa, Alexander Ditzel, Srujan Kopparapu, Meghan Mahalawat, Owen R. Davies, Sean R. Collins, Jeffrey R. Johnson, Nevan J. Krogan, Neil Hunter

## Abstract

Protein modification by SUMO helps orchestrate the elaborate events of meiosis to faithfully produce haploid gametes. To date, only a handful of meiotic SUMO targets have been identified. Here we delineate a multidimensional SUMO-modified meiotic proteome in budding yeast, identifying 2747 conjugation sites in 775 targets, and defining their relative levels and dynamics. Modified sites cluster in disordered regions and only a minority match consensus motifs. Target identities and modification dynamics imply that SUMOylation regulates all levels of chromosome organization and each step of homologous recombination. Execution-point analysis confirms these inferences, revealing functions for SUMO in S-phase, the initiation of recombination, chromosome synapsis and crossing over. K15-linked SUMO chains become prominent as chromosomes synapse and recombine, consistent with roles in these processes. SUMO also modifies ubiquitin, forming hybrid oligomers with potential to modulate ubiquitin signaling. We conclude that SUMO plays diverse and unanticipated roles in regulating meiotic chromosome metabolism.

## INTRODUCTION

Meiosis precisely halves the chromosome complement enabling parents to contribute equally to their progeny while maintaining a stable ploidy through successive generations (Hunter, 2015). Ploidy reduction occurs by appending two rounds of chromosome segregation to a single round of replication to produce haploid gametes from diploid germline cells (**Figure 1A**). The key events of meiosis that ensure accurate segregation include the connection of homologous chromosomes by crossovers (Hunter, 2006), monopolar orientation of sister-kinetochores during meiosis I (Watanabe, 2012), and the stepwise loss of sister-chromatid cohesion, first from chromosome arms at anaphase I and then from sister-centromeres at anaphase II (McNicoll et al., 2013). Crossover formation is the culmination of a complex series of interdependent chromosomal events during meiotic prophase I that include programmed homologous recombination, and the intimate pairing and synapsis of homologs (Zickler and Kleckner, 2015).

**Figure 1.**
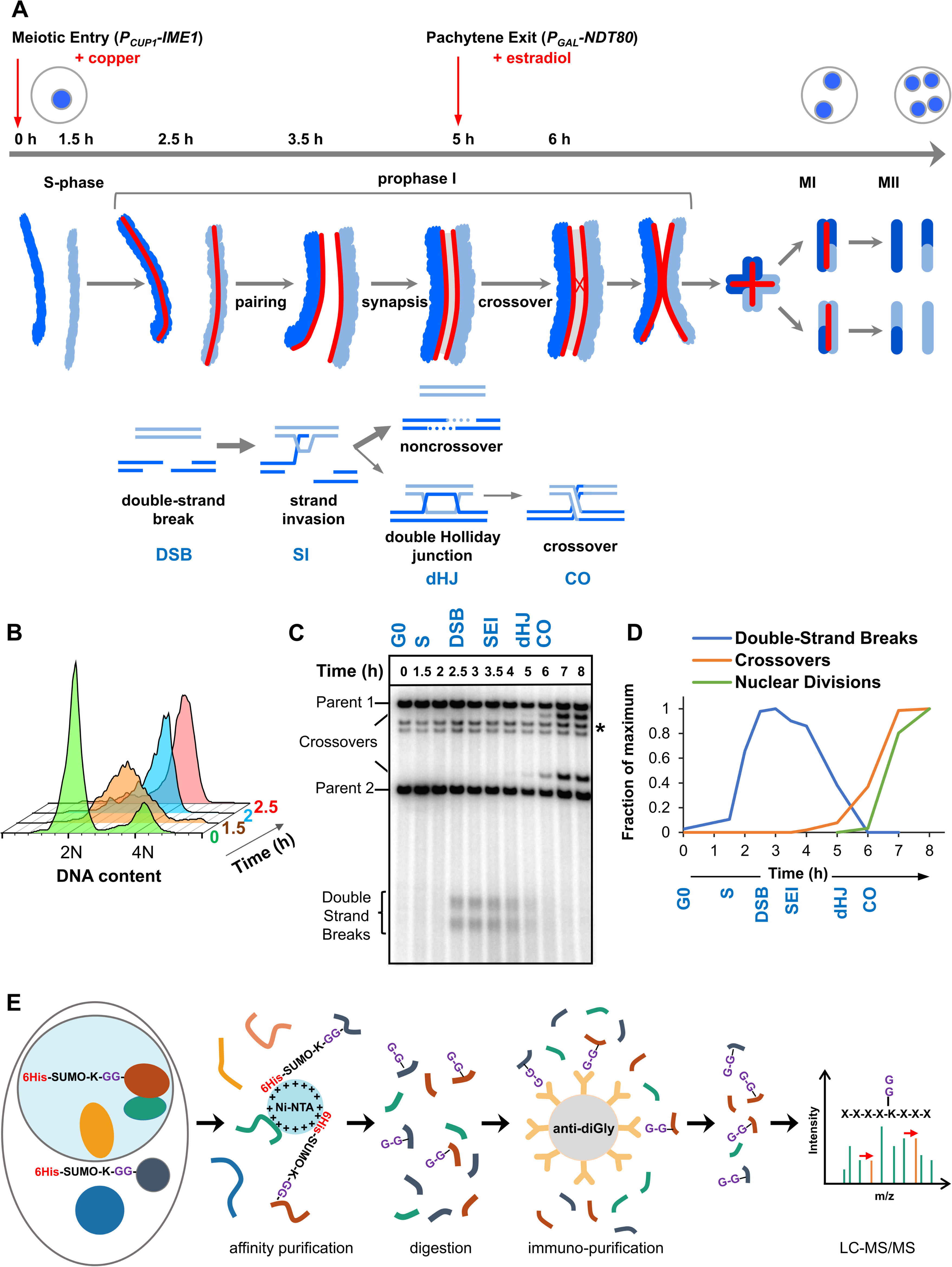
Experimental approach for profiling SUMOylation during meiosis. (A) Cell synchronization regimen showing the timing of nuclear (top), chromosomal (middle) and recombination (bottom) events. Samples were collected at the indicated timepoints (h, hours). Red arrows denote induction points of *IME1* and *NDT80* expression. (B) Flow cytometry data illustrating the progression of the meiotic S-phase following *P_CUP1_-IME1* induction. (C) Southern blot image showing the progression of recombination at the *HIS4::LEU2* hotspot. (D) Timing and synchrony of meiotic cultures. Levels of DSBs and crossovers at *HIS4::LEU2*, and nuclear divisions (MI±MII) were normalized to 1. The timing of SEIs and dHJs was determined using 2D gels (**Supplemental Information,** Figure 1). (E) Regimen for purification of peptides harboring K-ε-GG SUMO remnants (G-G-). Ni-NTA, nickel-nitrilotriacetic acid resin; anti-diGly, anti-K-ε-GG antibody beads).

Meiotic recombination is initiated by programmed DNA double-strand breaks (DSBs), some ∼200-300 DSBs per nucleus in budding yeast, mouse and human (Lam and Keeney, 2014). Ensuing recombinational interactions promote chromosome pairing and the assembly of synaptonemal complexes (SCs), densely-packed transverse filaments with a zipper-like morphology that connect homologs during the pachytene stage (Fraune et al., 2012; von Wettstein et al., 1984; Zickler and Kleckner, 1999). Within the context of the SCs, selected recombinational interactions mature into crossovers in such a way that each pair of chromosomes attains at least one crossover despite a low number of events per nucleus. Homologs then desynapse and prepare for the meiosis-I division. The connections created by crossovers enable the stable bipolar orientation of homologs on the meiosis-I spindle, and thus accurate segregation during meiosis I.

Orchestrating the elaborate events of meiosis are regulatory networks that function at the transcriptional, post-transcriptional, translational and post-translational levels (Bose et al., 2014; Brar et al., 2012; Cahoon and Hawley, 2016; Cheng et al., 2018; Crichton et al., 2014; Gao and Colaiacovo, 2018; Govin and Berger, 2009; Gray and Cohen, 2016; Jin and Neiman, 2016; Nottke et al., 2017; Otto and Brar, 2018; Tresenrider and Unal, 2018). The post-translational, SUMO (Small Ubiquitin-like MOdifier) modification system (SMS) is now recognized as an essential regulator of meiotic prophase (Cheng et al., 2007; de Carvalho and Colaiacovo, 2006; Lake and Hawley, 2013; Nottke et al., 2017; Rodriguez and Pangas, 2015; Sakaguchi et al., 2007; Vujin and Zetka, 2017; Watts and Hoffmann, 2011). Like ubiquitin, SUMO is conjugated to lysine (K) side-chains on target proteins via a cascade of enzymes that activate (E1) and conjugate (E2) SUMO, and provide target specificity (E3 ligases)(Dohmen, 2004; Gareau and Lima, 2010; Jentsch and Psakhye, 2013; Johnson, 2004; Zhao, 2018). SUMOylation can also be reversed by the action of dedicated proteases (Kunz et al., 2018). The consequences of SUMOylation are varied and target specific (Zhao, 2018), but include conformational changes, creating and masking binding interfaces to mediate protein interactions, and competing with other lysine modifications such as ubiquitylation and acetylation (Almedawar et al., 2012; Flotho and Melchior, 2013; Liebelt and Vertegaal, 2016; Papouli et al., 2005; Steinacher and Schar, 2005).

To date, only a handful of meiotic SUMO conjugates have been identified and studied in any detail. In budding yeast these include the SC component Ecm11 (Humphryes et al., 2013; Zavec et al., 2007), SUMO E2 conjugase, Ubc9 (Klug et al., 2013), core recombination factor, Rad52 (Sacher et al., 2006), chromosome-axis protein Red1 (Cheng et al., 2013; Eichinger and Jentsch, 2010; Lin et al., 2010; Zhang et al., 2014) and type-II topoisomerase, Top2 (Zhang et al., 2014). Also, in *C. elegans*, SUMO targets components of the chromosome congression/segregation ring complexes (Davis-Roca et al., 2018; Pelisch et al., 2017).

This paucity of examples underlines how understanding meiotic SUMOylation has been impeded by inefficient piecemeal approaches to identifying targets and mapping sites of SUMO conjugation. To overcome this impediment, we developed an efficient mass spectrometry (MS) regimen to map SUMO-conjugation sites proteome-wide during meiosis in budding yeast. In combination with label-free quantitation (LFQ) and highly synchronous meiotic time courses this approach allows the extent of SUMOylation at protein and site levels to be monitored during the key stages of meiotic prophase I. The resulting MS datasets provide a comprehensive and unprecedented view of the SUMO landscape, revealing dynamic waves of modification coincident with the major landmarks of meiosis. Functional classes of SUMO targets imply roles in basic cellular functions including metabolism, chromatin organization, transcription, ribosome biogenesis and translation. In addition, meiosis-specific aspects of chromosome metabolism are strongly represented by SUMOylated proteins pointing to roles in regulating recombination, chromosome synapsis and segregation. These inferences were explored by acutely inactivating *de novo* SUMOylation at different times during meiotic prophase. This analysis reveals distinct execution points for SUMO modification, and identifies roles in the onset of S-phase, DSB formation, crossing over and chromosome synapsis. Together, our analysis delineates a diverse and dynamic meiotic SUMO-modified proteome and provides a rich resource towards a mechanistic understanding how SUMO regulates the complex events of meiosis.

## RESULTS AND DISCUSSION

### Cell Synchronization and Purification of SUMO-Modified Peptides

Standard meiotic time-courses in budding yeast have relatively poor temporal resolution of the key events of meiotic prophase. To sharpen culture synchrony, we employed the method of Berchowitz et al. in which cells synchronize in G0 before meiosis is triggered by inducing expression of the master meiotic regulator, *IME1*, which is under control of the copper-inducible *CUP1* promoter (*P_CUP1_-IME1*; **Figure 1A**)(Berchowitz et al., 2013). 5 hours after induction of meiosis, cells were synchronized for a second time during pachytene, when homologs are fully synapsed. This was achieved by reversibly arresting cells using an estradiol inducible *NDT80* gene (*P_GAL_-NDT80*)(Benjamin et al., 2003), which encodes a meiosis-specific transcription factor required for progression beyond pachytene. Upon *P_GAL_-NDT80* expression, double-Holliday junction intermediates (dHJs) are rapidly resolved into crossovers, SCs disassemble and cells progress to MI (Allers and Lichten, 2001; Chu and Herskowitz, 1998; Clyne et al., 2003; Sourirajan and Lichten, 2008).

Cell samples from synchronized cultures were harvested and processed for SUMO proteomics at six different time-points that capture the key transitions of meiotic prophase I (**Figure 1A–D** and **Supplemental Information Figure 1**). Cells prior to *P_CUP1_-IME1* induction were sampled as a pre-meiotic control (**G0**). 1.5 hours after *P_CUP1_-IME1* induction (**S**), cells were in meiotic S-phase, but DSB formation had not begun. By 2.5 hours (**DSB**), DNA replication was complete and DSB formation was ongoing. 3.5 hours (**SI**), captures the events of DNA strand-invasion and accompanying SC formation. By 5 hours (**dHJ**), cells were arrested in pachytene with fully synapsed chromosomes and unresolved dHJs. *P_GAL_-NDT80* expression was then induced and cells harvested one hour later (**CO**), as dHJs were being resolved into crossovers but meiotic divisions had not yet begun.

To obtain the highest quality peptide samples for SUMO proteomics, we overcame three major impediments: (i) proteases are hyper-activated in meiotic cells (Klar and Halvorson, 1975); (ii) the stoichiometry of SUMOylation is typically very low (the “SUMO paradox”)(Hay, 2005); (iii) the native branched SUMO remnant produced by trypsin digestion (K-ε-GGIQE) is not amenable to efficient MS-based identification (Wohlschlegel et al., 2006). Thus, strains were generated in which the native *SMT3* locus was engineered to express hexa-histidine tagged Smt3 with an I96K mutation (6His-Smt3-I96K; **Figure 1E**)(Wohlschlegel et al., 2006; Xu et al., 2010). This construct enabled a two-step purification regimen in which SUMO-conjugated proteins were initially enriched using immobilized metal-affinity chromatography under denaturing conditions (6M guanidine), thereby limiting proteolysis. Eluted samples were then split and digested with LysC, or a combination of LysC plus GluC, to yield peptides with di-glycine branched SUMO-remnants that are amenable to further affinity purification using anti-di-glycyl-lysine antibodies (Xu et al., 2010). Eluted peptides were subjected to LC-MS/MS over a 90-minute acetonitrile gradient on a Q-Exactive Orbitrap (Thermo Scientific) with data-dependent acquisition; and data were processed using MaxQuant and Perseus software (Max Planck Institute)(Cox et al., 2014; Cox and Mann, 2008; Tyanova et al., 2016). To obtain biological replicates with high correlation scores (*r* ≥ 0.8 Pearson correlations, **Supplemental Information Figure 2A**) for quantitative analysis of SUMOylation dynamics, samples were collected from three independent time courses, processed in parallel, and then LC-MS/MS was performed in tandem with a randomized sample order and identical run conditions. In order to maximize the identification of target proteins and their conjugation sites, data from additional time courses were included in a separate analysis (Supplemental Information, Key MaxQuant Tables and EMBL-EBI PRIDE Archive accession number PXD012418).

**Figure 2.**
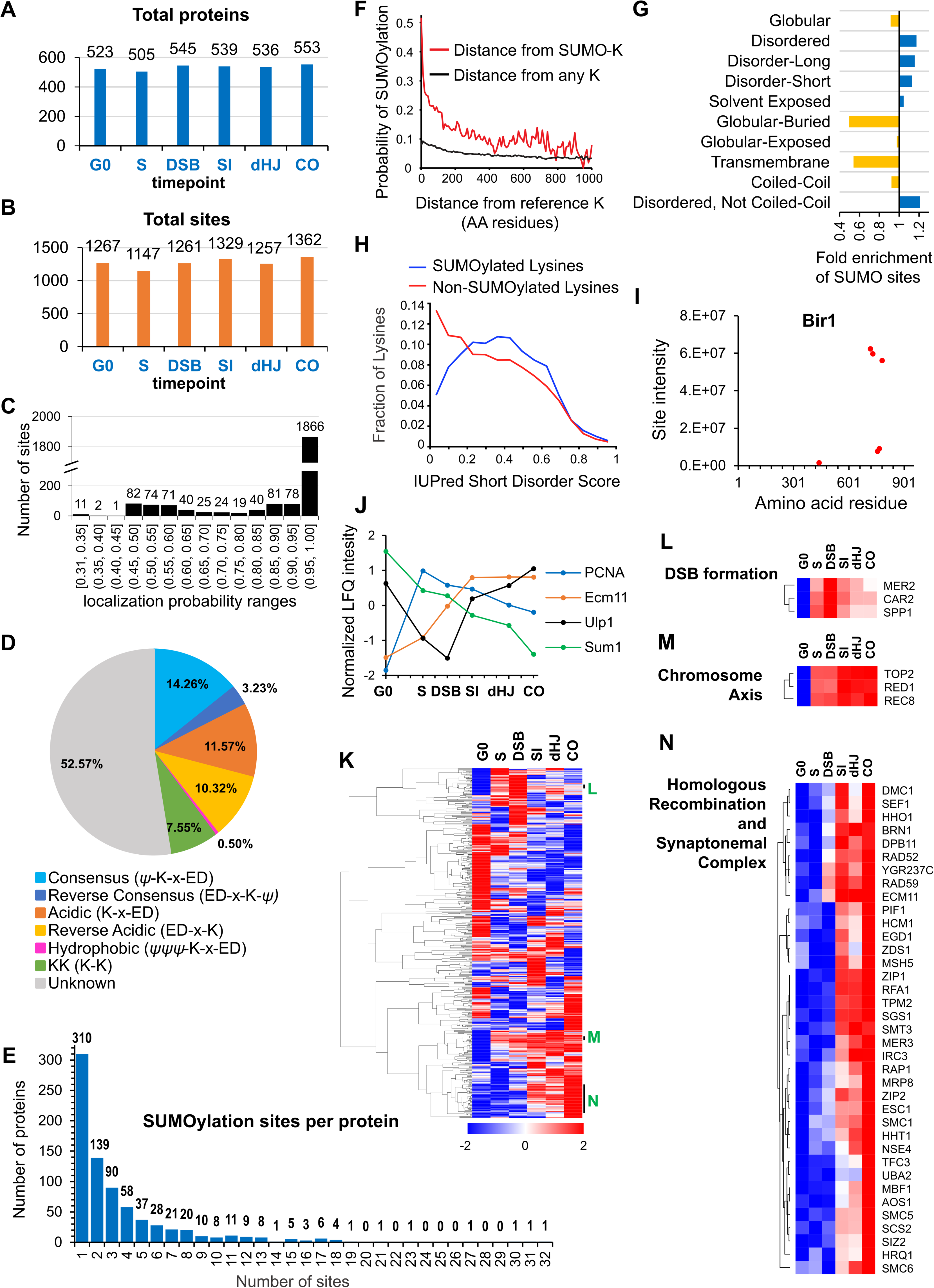
Characteristics of SUMOylation sites and temporal LFQ profiles of SUMOylated proteins. (A) (B) Total numbers of proteins (A) and SUMO sites (B) identified from each time point in the triplicate experiments used for quantitative analysis. (C) Localization probabilities for identified SUMO sites as calculated by MaxQuant (D) Proportions of identified SUMOylation sites that conform to indicated consensus sequences. Ψ, hydrophobic amino acid; x: any amino acid. (E) Distribution of SUMOylation sites per protein. (F) Probability of a lysine being SUMOylated as a function of its distance from either a SUMOylated lysine (red line) or any lysine. (G) Enrichment or depletion of SUMOylation sites relative to protein secondary structure. (H) Distributions of all lysines (red line) and SUMOylated lysines (blue line) relative to IUPred short disorder score. (I) Site intensity profile of Bir1 illustrating site clustering. (J) Normalized temporal LFQ profiles for PCNA, Ecm11, Ulp1 and Sum1 highlighting the diverse dynamics of SUMOylation during meiosis. (K) Hierarchical clustering of normalized LFQ profiles. (L) (M) (N) Clustering of SUMO targets involved in DSB formation (L), chromosome axes (M), and recombination and synapsis (N).

### Key Features of the SUMO-Modified Meiotic Proteome

Parallel analysis of samples digested separately with LysC and LyC + GluC enhanced peptide coverage, thereby increasing the number of SUMOylated proteins identified by 11% and the number of conjugation sites mapped by 27%. Combined analysis of all samples identified 2747 SUMOylation sites in 775 proteins. By comparison, the most comprehensive analysis of SUMO targets in vegetative yeast to date identified 244 targets and 257 sites (Esteras et al., 2017). Of these targets, 166 were also identified in our analysis indicating that meiotic and mitotic SUMO targets show significant overlap (**Supplemental Information Table 1** summarizes previous yeast SUMO MS/MS studies in vegetative cells).

For individual time-point samples in the three parallel time courses, between 505 and 553 target proteins, and 1147 and 1362 sites were identified, with a majority of proteins being identified in multiple time points (**Figure 2A** and **2B**). The relatively narrow numerical range of proteins identified across samples is one indication that sample preparation was consistent over the entire set. Also, of the 2414 conjugation sites identified in a single MaxQuant run, 1866 could be assigned with high confidence, with localization probabilities of ≥0.96 (**Figure 2C**). Moreover, a majority of lower confidence sites remain viable for functional analysis because decreased probabilities typically stemmed from ambiguity between two adjacent lysines in the same peptide.

Ubc9 binds directly to consensus SUMOylation peptide, ψ-K-x-E/D (ψ = large hydrophobic residue), to favor modification at these sites (Bernier-Villamor et al., 2002; Rodriguez et al., 2001; Sampson et al., 2001). However, only 14.26% of identified conjugation sites conformed to this consensus, with an additional 0.50% displaying the hydrophobic variant (ψψψ -K-x-E/D) and 3.23% having the reverse consensus sequence (E/D-x-K-ψ; **Figure 2D**). Around a fifth of sites comprised partial-consensus acidic (K-x-E/D, 11.57%) and reverse-acidic (E/D-x-K, 10.32%) motifs. Di-lysine was the only other recognizable motif (7.55%). Thus, a majority of modified sites (52.57%) did not have a recognizable motif. Identified sites were compared to those predicted by GPS-SUMO software (Zhao et al., 2014). Even when the search threshold setting was “low”, less than 19% (512 of 2745) of identified sites were predicted, emphasizing the limited utility of such software and the value of our proteomics dataset (**Supplemental Information Table 2**).

Two or more conjugation site were identified in 465 (60%) of the 775 SUMOylated proteins and the distribution of site numbers had a long tail, with 178 (23%) proteins containing five or more sites and three proteins with ≥30 sites (Ulp1, Red1 and Sir4; **Figure 2E**). When multiple conjugation sites were present in a single protein, they tended to cluster, with a 47% probability of adjacent SUMO sites being less than five residues apart (**Figure 2F**). This clustering effect implies that target features such as local secondary structure, solvent exposure and targeting by E3 ligases are more important determinants of SUMOylation site specificity than a consensus conjugation motif, which is absent from a majority of sites. Consistently, SUMO sites were depleted from regions of predicted globular, buried or transmembrane structure, and were enriched in regions of moderate disorder (**Figure 2G** and **H**). For each target protein identified in our analysis, we generated a diagram detailing the locations of SUMO sites relative to non-SUMOylated lysines, predicted SIMs, PFAM domains and protein secondary structure (**Supplemental Information, Protein Diagrams**).

### SUMOylation Dynamics and Target Identities

#### SUMOylation Site Intensities and Dynamics

Label-free quantification (LFQ)(Cox et al., 2014) further betrayed the immense complexity of meiotic SUMOylation (**Figure 2I–N**). First, for individual targets with multiple conjugation sites, cumulative site intensities (across all time points) provides a readout of relative site usage and site clustering (**Figure 2I**; also see examples below). This analysis also has predictive value for identifying functionally important sites, as exemplified by the much higher intensities of sites with known physiological roles in the SUMO E2 ligase, Ubc9 (K153, **Figure 3C**), and the SC component Ecm11 (K5, **Figure 5L**, and discussed further below; (Klug et al., 2013; Leung et al., 2015)). Second, intensity profiles revealed that different sites within a single target can have distinct SUMOylation dynamics, e.g., the profile of histone H4-K77 is distinct from those of the other SUMOylated H4 sites (**Figure 4F**). Third, normalized LFQ profiles showed that different targets can have radically different SUMOylation dynamics (**Figure 2J**). However, hierarchical analysis of LFQ profiles rendered clusters of functionally related proteins (e.g. DSB formation and HR), and physically interacting proteins (e.g. chromosome axis and SC) (**Figure 4 K–N**).

**Figure 3.**
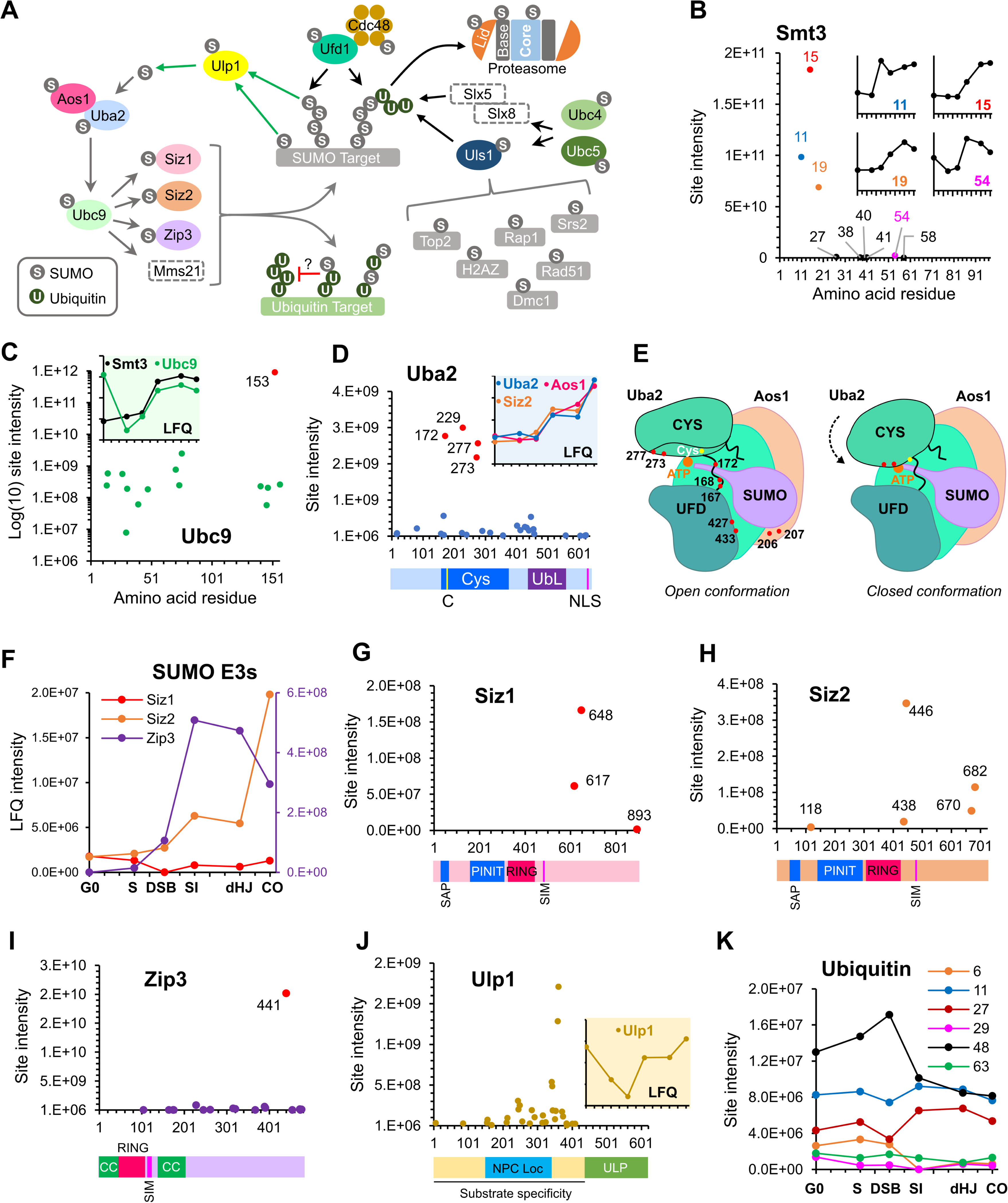
SUMOylation of the SUMO and Ubiquitin-proteasome machinery. (A) Summary of targets in the SUMO and ubiquitin-proteasome systems and the relationships between these factors. Dashed rectangles indicate SUMOylation was not detected. Note that although SUMOylation of Mms21 was not detected, the associated Smc5/6 complex was modified. (B) Intensity plot of SUMOylated sites on Smt3. Insets show temporal profiles of the four most prominent SUMO-chain linkages. (C) Intensity plot of SUMOylated sites on Ubc9 SUMOylation sites. Inset shows normalized LFQ profiles of Ubc9 and Smt3. (D) SUMO-site intensities on Uba2. The cartoon below shows key domains along the peptide backbone. C: active-site cysteine, NLS: nuclear localization sequence. Inset shows normalized LFQ profiles of Uba2, Aos1 and Siz2 (E) Cartoon of the Aos1-Uba2 structure highlighting the locations of SUMOylation sites. (F) LFQ profiles of the SUMO E3 ligases. Zip3 is plotted on the right-hand side Y-axis. (G) (H) (I) Site intensities of Siz1 (G), Siz2 (H) and Zip3 (I) plotted over their respective domain structures. SAP, SAF-A/B, Acinus and PIAS domain; PINIT, PINIT motif-containing domain; RING: SpRING domain; SIM, SUMO interacting motif; CC, coiled coil. (J) Ulp1 SUMO-site intensities plotted over its domain structure. NPC Loc, nuclear pore complex localization domain; ULP, Ubiquitin-like protease domain. Inset shows th normalized LFQ profile. (K) Intensity profiles of SUMOylation sites on Ubiquitin.

**Figure 4.**
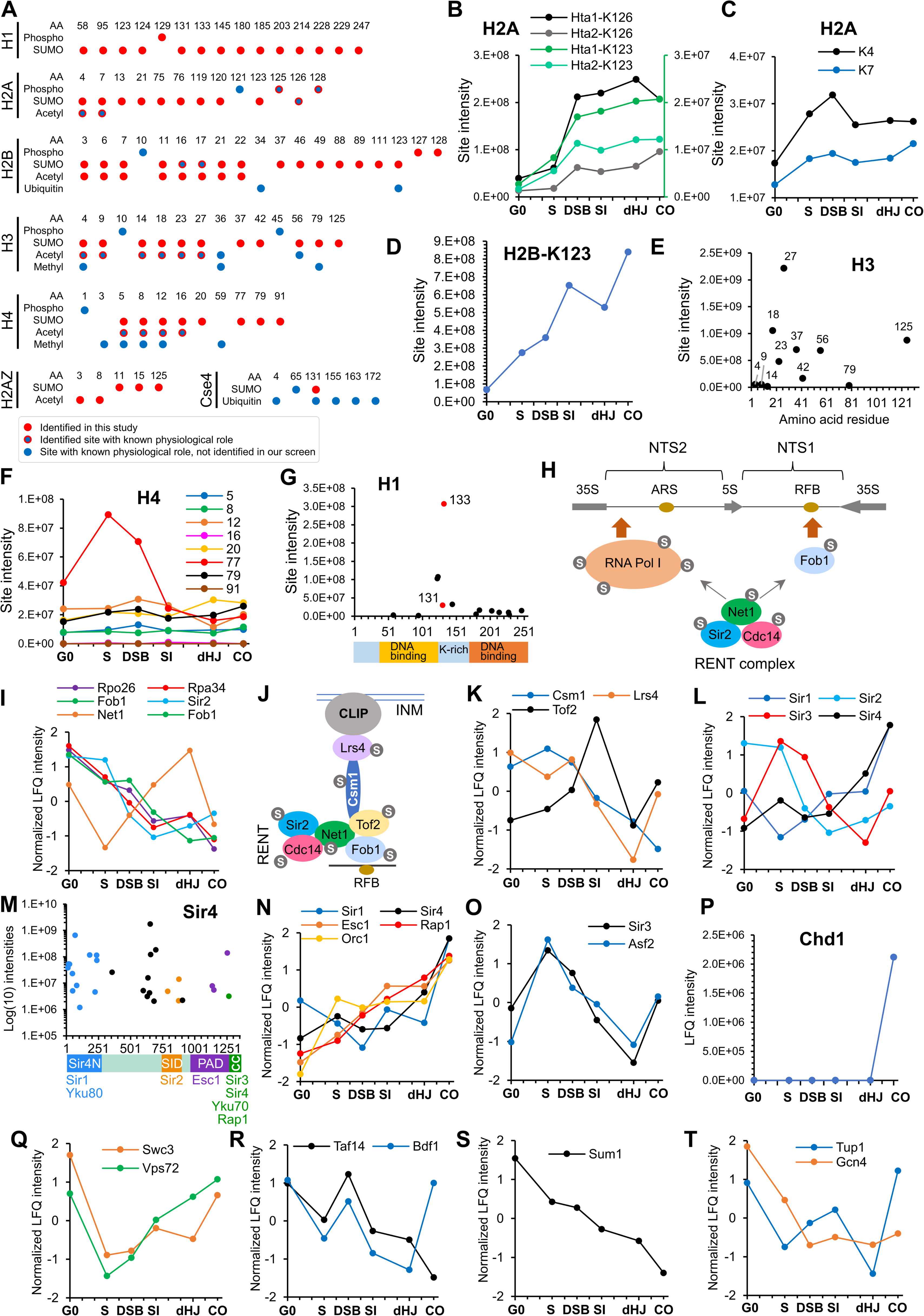
SUMOylation of chromatin and associated – Histones, SIR proteins, chromatin remodelers and transcription factors. (A) Chart summarizing histone SUMOylation, phosphorylation and acetylation sites identified in this study, together with previously identified modifications. (B) Temporal profiles of SUMOylation on K123 and K126 of histones H2A1 and H2A2. The K123 profiles are plotted on the right-hand side Y-axis. (C) Temporal profiles of K4 and K7 SUMOylation on H2A. (D) Temporal profiles of H2B-K123 SUMOylation. (E) SUMO site intensities on H3. (F) Temporal profiles of H4 SUMOylation sites. (G) Site intensities of H1 SUMOylation sites plotted along its domain structure. Two sites that are co-modified with phosphorylation are indicated in red. (H) Summary of SUMOylation proteins involved in rDNA silencing. (I) Normalized LFQ profiles of the RENT complex. NTS1/2, non-transcribed sequences; RFB, replication fork barrier. (J) Cartoon illustrating the recruitment of rDNA loci to the inner nuclear membrane (INM) via RENT, cohibin (Csm1, Lrs4) and CLIP (chromosome linkage inner-nuclear membrane proteins) complexes. INM, inner-nuclear membrane. (K) Normalized LFQ profiles of Cohibin and Tof2. (L) Normalized LFQ profiles of the SIR proteins. (M) Site intensity plot of Sir4 plotted along its domain structure. (N) Normalized LFQ profiles of Sir4 and its binding partners. (O) Normalized LFQ profiles of Sir3 and its binding partner Asf2. (P) LFQ profile of Chd1. (Q) Normalized LFQ profiles of SWR1-complex components Swc3 and Vps72 (R) Normalized LFQ profiles of Bdf1 and Taf14. (S) Normalized LFQ profile of Sum1. (T) Normalized LFQ profiles of Tup1 and Gcn4.

**Figure 5.**
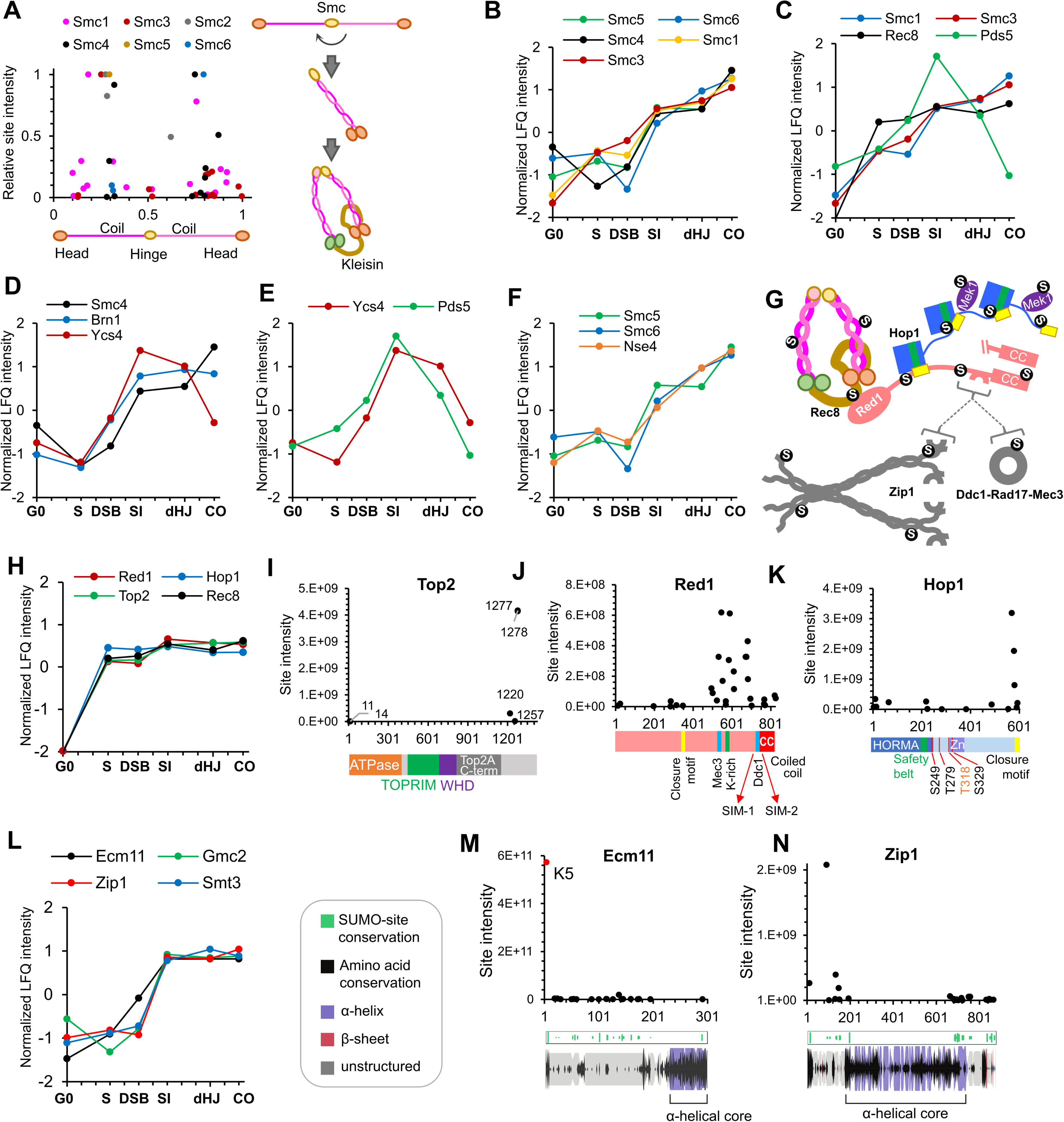
SUMOylation of homolog axes and synaptonemal complex. (A) Relative positions of SUMO sites on the SMC proteins. Adjacent cartoon illustrates how SMCs are assembled into ring structures. (B–F) Normalized LFQ profiles of the SMCs (B; Smc2 did not have measurable LFQ intensities); components of Rec8-cohesin (C); condensin (D); HEAT-repeat proteins Ycs4 and Pds5 (E); and components of theSmc5/6 complex (F). (G) Summary of SUMOylated axis proteins and pertinent interaction partners. Arcs in Red1 and Zip1 indicate SIMs. (H) Normalized LFQ profiles of axis components Red1, Hop1, Top2 and Rec8. (I) Site intensities of Top2 plotted along its domain structure. TOPRIM, Topoisomerase-primase domain; WHD, winged-helix domain. (J) Site intensities of Red1 plotted along its domain structure. (K) Site intensities of Hop1 plotted along its domain structure. S249, T279 T318 and S329 are key phosphorylation sites. (L) Normalized LFQ profiles of SC central region components Ecm11, Zip1 and Gmc2 relative to SUMO (Smt3). (M) (N) Site intensity plots for Ecm11 (M) and Zip1 (N) plotted along predicted secondary structures.

#### Gene Ontology and Network Analysis

Gene Ontology (GO) analysis of targets showed strong enrichment for nuclear processes (**Supplemental Information Figure 2B** and **2C**), especially those associated with chromosome metabolism and ribosome biogenesis, as previously seen for mitotically cycling cells (Albuquerque et al., 2013; Albuquerque et al., 2015; Esteras et al., 2017; Hendriks et al., 2014; Srikumar et al., 2013a). However, cytosolic processes were also enriched due the strong representation of glycolytic enzymes and other carbohydrate metabolic processes. Membrane-associated processes were notably underrepresented, although this could in part reflect their low expression, hydrophobicity and lower density of Lys-C cutting sites (Vit and Petrak, 2017). Network analysis revealed large clusters of physically interacting proteins consistent with the group SUMOylation paradigm advanced by Jentsch and colleagues (Jentsch and Psakhye, 2013; Psakhye and Jentsch, 2012). The most striking example is the ribosome with almost all of the 40S and 60S subunits being SUMOylated, for a total of 84 proteins and 527 conjugation sites (**Supplemental Information, Figure 3** and **Table 3**).

### Functional Classes of Meiotic SUMO Targets

Targets involved in all aspect of meiotic chromosome metabolism were SUMOylated, including DNA replication and repair, the DNA damage response; chromatin, transcription, telomeres, homologous recombination, synapsis and chromosome segregation. Below, we discus subsets of these targets pertinent for regulation the SUMO and ubiquitin modification systems, chromatin, chromosome structure and homologous recombination.

#### The SUMO Machinery

##### SUMO (Smt3)

A principal target of meiotic SUMOylation was the SUMO machinery (**Figure 3A**). SUMO itself was the fifth largest contributor to total LFQ signal intensity and all 9 lysines on Smt3 were identified as acceptor sites (**Figure 3B**). K15 was the dominant linkage with K11 and K19 also making significant contributions, which is consonant with previous studies indicating that SUMO chains are linked primarily via the flexible N-terminal extension (Klug et al., 2013). SUMO chains are essential for meiosis and have been implicated in SC formation and inter-homolog recombination (Cheng et al., 2006; Lin et al., 2010). Consistently, SUMO chains increased sharply after DSB formation and peaked in pachytene (**Figure 3B** and inset in **C**). The next most abundant linkage, K54, had the same profile, suggesting a possible role for K54-linked chains in meiotic prophase. The most prominent meiotic substrate of poly-SUMOylation is the SC central element protein Ecm11 (Leung et al., 2015) and the LFQ profiles of SUMO chains and Ecm11-SUMO were closely matched (**Figure 6N**).

**Figure 6.**
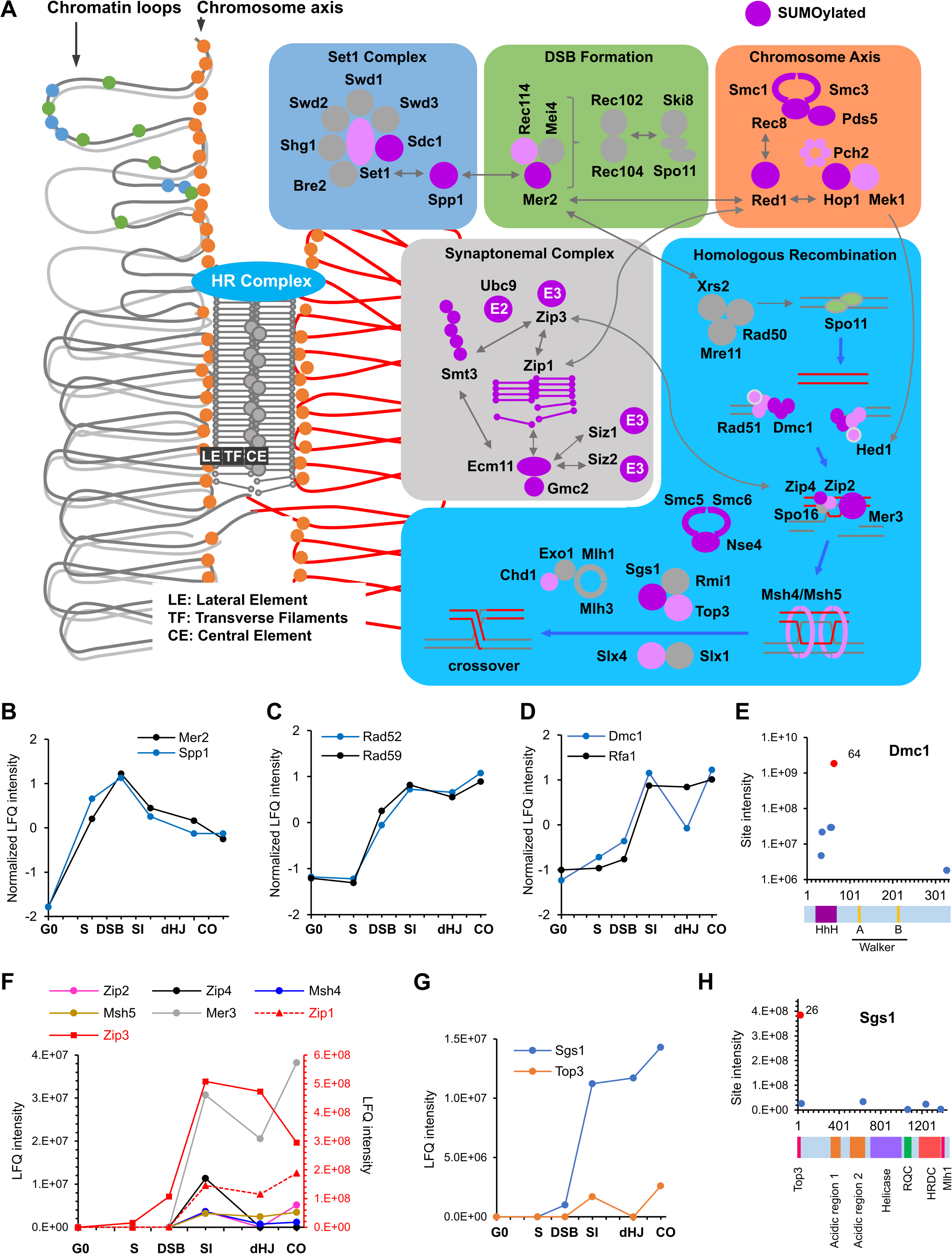
SUMOylation of the recombination machinery. (A) Summary of SUMOylated meiotic HR factors and their functional integration with other pertinent SUMO targets in chromosome axes and SCs. (B–D) Normalized LFQ profiles of DSB formation proteins Mer2 and Spp1 (B); mediator and strans annealing proteins Rad52 and Rad59 (C); and the strand exchange factors Dmc1 and Rfa1 (D). (E) Site intensities of Dmc1 plotted along its domain structure. HhH: Helix-hairpin-helix. (F) LFQ profiles of the ZMM proteins (SUMOylation of Spo16 was not detected). Zip1 and Zip3 are plotted on the right-hand side Y-axis in red. (G) Normalized LFQ profiles of helicases Sgs1 and Top3. (H) Sgs1 site intensities mapped along its domain structure. Top3, Top3 interacting domain; RQC, RecQ C-terminal domain; HRDC, helicase and RNaseD C-terminal domain; Mlh1, Mlh1 interacting domain.

##### Ubc9 (E2)

Ubc9 was the largest single contributor to total LFQ signal intensity. 15/17 lysines in Ubc9 were identified as SUMO acceptor sites (**Figure 3C**). K153 accounted for the vast majority of the signal with a 365-fold higher intensity than the next most abundant site. Klug et al. showed that K153-SUMO inhibits conjugase activity and converts Ubc9 into a cofactor that enhances chain formation by unmodified Ubc9 (Klug et al., 2013). This activity is particularly important during meiosis where it facilitates SC formation. Ubc9-K153-SUMO was high in G0, dropped in S phase and then increased again at the time of SC formation. High levels of Ubc9-K153-SUMO in G0 seem paradoxical because SUMO chains were relatively low at this time (**Figure 3C**, inset). These observations suggest that Ubc9-K153-SUMO is not the sole regulator of SUMO chain formation. Alternatively or in addition, SUMO chains may be less stable during G0.

Whether SUMOylation at other sites of Ubc9 plays a function in meiosis is unknown. Knipscheer et al. showed that SUMOylation of human Ubc9 at K14 regulates target discrimination (Knipscheer et al., 2008). K76 SUMOylation also has the potential to alter target discrimination as this residue lies in a basic patch that mediates Ubc9 specificity for acidic and phosphorylation-dependent consensus conjugation sites (Mohideen et al., 2009). Intriguingly, K76 SUMOylation was highest at G0, but diminished by the time of SC formation after which K153 completely dominated.

##### Aos1 ^SAE1^-Uba2 ^SAE2^ (E1)

27 acceptor sites (57% of all lysines) were identified on Uba2^SAE2^, the catalytic subunit of the E1 activating enzyme, Aos1 ^SAE1^-Uba2 ^SAE2^ (**Figure 3D**). LFQ identified four prominent sites, K172, 229, 273 and 277, with intensities ∼4-fold higher than the next most intense site). SUMOylation of K172 (or adjacent K167 and K168) is expected to be inhibitory because it lies in the crossover loop involved in the conformational changes that accompany activation (Olsen et al., 2010)(**Figure 3E**). Although a triple (167,168,172) K-R mutant was previously shown to have negligible impact on the growth and genotoxin sensitivity of vegetative cells (Albuquerque et al., 2015), its effects on meiosis are unknown. K273 and K277 lie proximal to the adenylate in the closed conformation of the E1 and are also predicted to be inhibitory. SUMOylation at both sites rises after G0 and then remains relatively constant until the final crossover time point, when levels spike sharply. Also notable is a cluster of eight acceptor sites that mapped around the hinge of the ubiquitin-fold domain (residues 400-460) and could also influence conformational changes involved in activation. The diverse intensity-profiles of individual sites suggest that Uba2 regulation may be complex (**Supplemental Information Figure 4**). Finally, in Aos1^SAE1^, a cluster of four acceptor sites, centered on the intense K207 site, lies close to an insertion loop that could influence adenylation and transfer of SUMO to Ubc9. Aos1-K207 SUMOylation rose steadily after DSB formation.

##### E3 ligases, Siz1, Siz2 and Zip3

SUMOylation of an E3 can enhance apparent catalytic activity by providing SUMO *in cis* to interact with the “backside” of Ubc9 and thereby stimulate its activity (Cappadocia et al., 2015). SUMO conjugation could also modulate E3 activity in other ways. Three of the four known yeast SUMO E3s, Siz1, Siz2 and Zip3, were modified in meiosis and showed diverse SUMOylation profiles (**Figure 3F**; modification of Mms21 was not detected). Siz1 SUMOylation on three C-terminal tail sites, dominated by K648 (**Figure 3G**), was relatively high in G0 and S-phase, dipped during DSB formation and returned thereafter. This profile may reflect known substrates of Siz1 such as PCNA, the septin Cdc3, and transcriptional repressors Sum1 and Tup1, all of which were SUMOylated during meiosis (**Supplemental Information Table 3**)(Johnson and Gupta, 2001; Papouli et al., 2005; Pfander et al., 2005; Srikumar et al., 2013b). Mid-prophase targets of Siz1 also include the SC component Ecm11 (Leung et al., 2015).

Ecm11 can also be targeted by Siz2, which was SUMOylated at five sites. While two acceptor sites reside in the C-terminal tail of Siz2, two other sites lie between the SP-CTD and the SIM, including the most intense site K446 (**Figure 3H**). Modification at K438 or K446 could provide a well-positioned “backside SUMO^B^” *in cis* to stimulate Siz2 activity (Streich and Lima, 2016). Distinct from Siz1, SUMOylation of Siz2 was relatively low during S phase, but rose after DSB formation and showed a large increase during CO formation (**Figure 3F**), closely matching the profiles of Aos1 and Uba2 (**Figure 3D**, inset), and the silencing factor, Esc1 (see below).

Zip3 is a meiosis-specific SUMO ligase that promotes the chromosomal localization of the ZMM group of pro-crossover factors, and helps locally couple SC formation to prospective CO sites (Cheng et al., 2006) (Agarwal and Roeder, 2000; Chua and Roeder, 1998; Shinohara et al., 2008). Zip3 also helps recruit Ubc9 and 26S proteasomes to meiotic chromosomes, reminiscent of the SUMO-Ubiquitin axis described in mammals (Ahuja et al., 2017; Hooker and Roeder, 2006; Rao et al., 2017). Zip3 comprises an N-terminal RING domain followed by an essential SIM, a short region of putative coiled-coil and a disordered serine-rich tail. The 17 acceptor sites mapped in Zip3 were scattered throughout the coiled-coil and tail (**Figure 3I**). The one exception, K105 lies adjacent to the SIM, with potential for auto-regulation. Sites within the coiled-coil (K165, 175, 176) could influence Zip3 oligomerization. SUMOylation at K441 accounted for almost 90% of the total intensity. Intriguingly, this site appears to be part of a compound modification motif, 441-KRSNSTQ-447, comprising a consensus site for the DNA damage-response kinases Mec1/Tel1 (S/T-Q), phosphorylation of which could prime Cdc7-mediated phosphorylation of upstream serines to create an acidic consensus site for SUMOylation, K-x-S(P). Consistent with this idea, phosphorylation of these sites was identified in our dataset, with co-modification being detected on ∼2/3rds of the peptides (**Supplemental Information Table 3**). Serrentino et al. previously showed that the four S/T-Q sites of Zip3 are required for its efficient accumulation at recombination sites and full crossover function (Serrentino et al., 2013). Zip3-K441-SUMO is possibly the downstream effector of Mec1/Tel1 phosphorylation.

##### SUMO isopeptidase Ulp1

The essential Ulp1 isopeptidase matures the SUMO pro-peptide and de-SUMOylates a subset of substrates (Li and Hochstrasser, 1999; Zhao, 2018). In mitotically cycling cells, Ulp1 regulates myriad processes including DSB repair, 2µm copy number, mRNA surveillance, silencing, nuclear import and export of pre-60S ribosomes, but its roles in meiosis have not been characterized (Dobson et al., 2005; Lewis et al., 2007; Palancade et al., 2007; Panse et al., 2006; Stade et al., 2002; Zhao et al., 2004). A remarkable 32 acceptor-sites were mapped on Ulp1, more than any other protein (**Figure 3J**). Sites were found across the N-terminal region of the protein but excluded from the essential C-terminal catalytic domain (Li and Hochstrasser, 2003; Mossessova and Lima, 2000; Panse et al., 2003). Most sites locate in the overlapping regions required for tethering of Ulp1 to nuclear pores (144–346) and for substrate specificity (amino acids 1-417)(Li and Hochstrasser, 2003; Panse et al., 2003), pointing to roles for Ulp1 SUMOylation in these processes. The overall signal for Ulp1-SUMO dipped around the time of DSB formation but increased again with synapsis, suggesting global changes in protein de-SUMOylation at these times (**Figure 3J**, inset).

#### Cross Talk With Ubiquitin and Autophagy

SUMO targets also revealed cross talk with the ubiquitin-proteasome system (UPS; **Figure 3A**). UPS targets included Ubc4 and Ubc5, paralogous E2 conjugases involved in protein quality control, stress response and cell-cycle regulation by the APC/C (Finley et al., 2012). Notably, Ubc4/5 work with SUMO-targeted ubiquitin ligases (STUbLs) that target poly-SUMOylated substrates (Sriramachandran and Dohmen, 2014; Uzunova et al., 2007). SUMO also modified five ubiquitin E3 ligases, Ubr1, Ufo1, Uls1, Gid7 and Ufd4, involved in a variety processes including the N-end rule, catabolite-degradation of fructose-1,6-bisphosphatase, transcription, cell cycle and genome maintenance (Bao et al., 2015; Baranes-Bachar et al., 2008; Finley et al., 2012; Kramarz et al., 2017; Lin et al., 2015; Sriramachandran and Dohmen, 2014; Varshavsky, 1997). Notable is Uls1, a Swi2/Snf2-family DNA translocase and putative STUbL (Sriramachandran and Dohmen, 2014). Uls1 is thought to compete with a second STUbL, Slx5-Slx8, to displace from DNA poly-SUMOylated proteins that have been rendered defective or inactive (Tan et al., 2013; Wei et al., 2017). Inferred Uls1 targets, all of which were SUMOylated in meiosis, include Top2 (Wei et al., 2017), H2A.Z (Takahashi et al., 2017), Rap1 (Lescasse et al., 2013), Srs2 (Kramarz et al., 2017), Rad51 (Chi et al., 2011) and possibly Dmc1 (Dresser et al., 1997)(**Figure 3A. Supplemental Information Table 3**). The ubiquitin-dependent segregase, Cdc48/p97, can target proteins co-modified by ubiquitin and SUMO (Bergink et al., 2013; Nie et al., 2012). Both Cdc48 and Ufd1, the cofactor implicated in SUMO binding, were SUMOylated (**Figure 3A, Supplemental Information table 3**).

Intriguingly, ubiquitin itself was SUMOylated on six lysines that are also sites for ubiquitin chain formation (K6, 11, 27, 29, 48 and 63; **Figure 3K**). Clear differences in site intensities were detected, with K48 and K11 being the most abundant followed by K27. Interestingly, K48 SUMOylation, which has the potential to modulate targeting to proteasomes, peaked during DSB formation and dipped thereafter. Ubiquitin is expressed from four loci in budding yeast, as a head-to-tail poly-ubiquitin precursor from the *UBI4* locus, and as ubiquitin fusions to the ribosomal proteins Rps31 and Rpl40A/Rpl40B. Whether ubiquitin SUMOylation reflects modification of ubiquitin precursors, free ubiquitin and/or ubiquitin conjugates is unclear. However, this observation further corroborates the evidence for a unique class of mixed ubiquitin-SUMO chains with potential for novel signaling functions (Esteras et al., 2017; Hendriks et al., 2014).

Autophagy is particularly important for the initiation of meiosis (Wen et al., 2016), but also functions in nuclear architecture and chromosome segregation (Matsuhara and Yamamoto, 2016). Three autophagy factors were SUMOylated: the ubiquitin-like Atg8 protein that undergoes lipidation via conjugation to phosphatidylethanolamine, its cognate E1 enzyme Atg7, and the dual receptor Cue5, which simultaneously binds ubiquitylated cargo and Atg8 (**Supplemental Information Table 3**). These conjugates raise the possibility that SUMO modulates the targeting of aggregated proteins and inactivated proteasomes to phagophores (Wen and Klionsky, 2016).

#### Chromatin

##### Histones

SUMOylation of histones is generally implicated in transcriptional repression by recruiting HDACs, or by competing with activating marks such as acetylation and ubiquitylation (Cubenas-Potts and Matunis, 2013; Nathan et al., 2006; Shiio and Eisenman, 2003). However, mechanistic details and roles of SUMOylation at individual sites remain largely unexplored. We found that the four core histones, H2A variant Htz1 (H2A.Z), linker histone H1, and the H3-like centromeric histone, Cse4, were SUMOylated in meiosis (**Figure 4A**). In many cases, mapped sites are also known targets of other lysine modifications suggesting antagonistic relationships. However, we also found specific examples of obligate co-modification with phosphorylation or acetylation, pointing to more complex, potentially cooperative interactions between SUMO and other histone marks.

###### Histone H2A

Mec1/Tel1-mediated H2A-S128 phosphorylation (γ-H2A) plays important roles in the DNA damage response (Downs et al., 2000; Morrison et al., 2004; Redon et al., 2003; van Attikum et al., 2004). In yeast meiosis, γ-H2A is initially triggered in S-phase (Cheng et al., 2013) and contributes to a checkpoint that prevents re-replication (Najor et al., 2016). In our dataset, Hta1 (H2A1) K123 SUMOylation was only observed on a peptide that was also phosphorylated at either at T125 or S128. Similarly, Hta2 (H2A2) K123 SUMOylation was always coincident with S128 phosphorylation. Although the K123 site does not conform to a canonical phospho-dependent SUMOylation motif (PDSM, ΨKxExxSP), S128 phosphorylation is suitably positioned to enhance its conjugation (Hietakangas et al., 2006). Unlike K123, the much more abundant K126 SUMOylation (on either Hta1 or Hta2) was not always coincident with phosphorylation. Thus, SUMOylation of K123, but not K126 appears to be phospho-dependent. Analogous to S128 phosphorylation (Cheng et al., 2013), K123 and K126 SUMOylation appeared at the onset of S-phase, increased with DSB formation and then persisted throughout prophase (**Figure 4B**). These observations suggest a compound role for T125/128 phosphorylation and K123 SUMOylation in S-phase and/or recombination.

SUMOylation of H2A at K4 or K7 was always coincident with acetylation of K7 or K4, respectively (**Figure 4A**), suggesting that N-terminal SUMOylation is stimulated by pre-existing acetylation, as seen for human H3 (Hendriks et al., 2014). This subset of H2A modifications showed distinct dynamics relative to those in the C-terminus, with peak intensities in the S-phase and DSB samples (**Figure 4C**), when gene promoters are being targeted for DSB induction. Notably, H2A K5/K8 acetylation leads to Swr1-dependent incorporation of H2A.Z at promoters and activation of transcription (Altaf et al., 2010). Possibly, this co-modification event facilitates chromatin changes that modulate DSB formation.

###### Histone H2B

N-terminal SUMOylation of H2B (K6, 7, 16, 17) was previously inferred to compete with acetylation to repress transcription (Nathan et al., 2006). SUMOylation of these sites was also detected in meiosis, both alone and in combination with acetylation (**Figure 4A**). K123 was the most prominent H2B SUMOylation site, which climbed in intensity throughout prophase I (**Figure 4D**). Ubiquitination at this site has pleiotropic roles in transcriptional activation and elongation, telomeric silencing, DNA replication and DNA repair (Fuchs and Oren, 2014). Notably, in meiosis, H2B K123 and the Bre1 ubiquitin ligase are required for efficient DSB formation (Yamashita et al., 2004), likely due to the role of H2B-K123 ubiquitylation in stimulating Set1-mediated H3-K4 trimethylation, which in turn helps recruit the DSB machinery to chromatin (Borde et al., 2009; Lam and Keeney, 2014). Set1 also facilitates progression of meiotic S-phase and the expression of middle meiotic genes (Sollier et al., 2004). We suggest that H2B K123 SUMOylation may compete with ubiquitylation to down-regulate H3-K4 methylation and thereby repress middle meiotic genes and attenuate DSB formation in later stages of meiotic prophase.

###### Histone H3

Extensive SUMOylation and acetylation of the N-terminus of H3 was also detected (**Figure 4A**). This included H3-K4 suggesting that SUMOylation of a second histone site might also negatively regulate DSB formation. However, SUMOylation at K4, 9 and 14 was much less intense than at the adjacent sites, K18, 23 and 27 (**Figure 4E**). Acetylation of H3, especially at K14, is involved in Gcn5-dependent transcriptional activation and SUMOylation may modulate this activity (Grant et al., 1999; Suka et al., 2001). Dot1-catalyzed H3-K79 methylation facilitates meiotic checkpoint activation, promotes DSB formation when H3-K4 is mutated, and helps prevent re-replication in combination with γ-H2A (Bani Ismail et al., 2014; Najor et al., 2016; Ontoso et al., 2013). SUMOylation was also detected at H3-K79 suggesting antagonism of these activities.

###### Histone H4

SUMOylation of H4 functions in transcriptional repression (Nathan et al., 2006; Shiio and Eisenman, 2003). We found extensive SUMOylation of the H4 N-terminus including K5, 16 and 20, which was always coincident with acetylation (**Figure 4A and F**). Interestingly, the most abundantly SUMOylated site, K77 showed a distinct transient peak in the S-phase and DSB time points (**Figure 4F**). K77 and the adjacent K79 are known sites of acetylation and their mutation disrupts silencing of telomeric and rDNA loci (Hyland et al., 2005). However, coincident SUMOylation and acetylation at K77 and K79 was not observed pointing to an antagonistic relationship at these sites. Possibly, K77/79 SUMOylation facilitates replication of transcriptionally repressed regions of the genome.

###### Histone H1

H1 (Hho1) is a poorly conserved linker histone (Millan-Arino et al., 2016) that binds DNA at nucleosome entry and exit points, and functions in chromatin compaction, silencing of rDNA, and suppression of homologous recombination (Downs et al., 2003; Georgieva et al., 2012; Levy et al., 2008; Schafer et al., 2008). In budding yeast meiosis, H1 is inferred to help repress early meiotic genes, is subsequently depleted during prophase, and finally accumulates in mature spores to facilitate chromatin compaction (Bryant et al., 2012). Metazoan H1 is phosphorylated in S-phase and mitosis by Cdk2, which leads to decompaction and destabilization of chromatin (Hale et al., 2006; Roth and Allis, 1992; Sarg et al., 2006). Mammalian H1 undergoes K63-linked polyubiquitination in response to DNA damage, which mediates the propagation of ubiquitin-mediated damage signaling (Thorslund et al., 2015). We found H1 to be extensively SUMOylated during meiosis, with levels increasing throughout meiosis. In particular, SUMO sites were scattered throughout the second DNA binding domain; and a prominent cluster was detected in the lysine-rich region that connects the DNA-binding domains (**Figure 4G**). Two of the highest intensity SUMO sites in this region, K131 and K133, were always present on a peptide co-modified with phosphorylation at S129, a CDK site (S-P). As with H2A, these SUMO sites do not conform to the canonical PDSM consensus but exhibit apparent dependence on phosphorylation. These observations point to a cooperative function for SUMO and phosphorylation in regulating H1 dynamics on meiotic chromatin.

In summary, histones are extensively SUMOylated during meiosis suggesting competitive and cooperative functions with other modifications to regulate gene expression, S-phase, DSB formation, recombination and chromatin compaction.

##### Histone modifiers and the SIR complex

Dynamic changes in meiotic histone modifications were accompanied by extensive SUMOylation of histone modifying enzymes, including acetyltransferases (SAGA, NuA3, Nua4 and ADA complexes), deactylases (Set3C, Hda1, Rpd3S, Rpd3L, SIR, RENT and Sum1-Rfm1-Hst1 complexes), the COMPASS methytransferase and the JmjC demthylase (**Supplemental Information Table 3**). Here we focus on the Sir (Silent Information Regulator) proteins, given their central roles in meiotic chromosome metabolism. Sir proteins are implicated in the pachytene-checkpoint response, crossover interference and global patterning of DSBs, including suppression of DSB formation in the rDNA array and sub-telomere regions (Mieczkowski et al., 2007; Subramanian and Hochwagen, 2014; Vader et al., 2011; Zhang et al., 2014).

The H4-K16Ac deacetylase Sir2 is recruited to rDNA as part of the RENT complex (regulator of nucleolar silencing and telophase exit), Cdc14-Net1-Sir2 (Gartenberg and Smith, 2016). Net1 is the key scaffolding protein for both assembly of RENT and its recruitment to the two rDNA nontranscribed-spacers via interactions with Pol I and Fob1 (**Figure 4H**). These proteins represent a heavily SUMOylated cohort, with 16 sites on Net1, eight on Fob1, three on Sir2, a single site on Cdc14, and 20 sites across the nine subunits of RNA Pol I (Supplemental Information Table 3). Net1 SUMOylation intensity diminished during S-phase, increased again through the dHJ/synapsis time point, and then dropped again during CO formation (**Figure 4I**). This could reflect dynamic changes associated with rDNA replication and perhaps the release of Sir2 from the nucleolus to perform nuclear functions, such as the synapsis checkpoint and crossover interference. In vegetative cells, localization of Sir2 to the nucleolus is dependent on its SUMOylation (Hannan et al., 2015). However, SUMOylation of Sir2, Fob1 and two prominently SUMOylated components of RNA Pol I (Rpa34 and Rpo26) all decreased as meiosis progressed (**Figure 4I**). Additionally, nucleolar localization of Sir2 during meiosis requires Dot1 (San-Segundo and Roeder, 2000), which is required for pachytene checkpoint arrest. These considerations suggest that the regulation of Sir2 dynamics may be distinct in meiotic and vegetative cells.

rDNA stability also requires anchoring to the nuclear periphery, which involves another cohort of SUMOylated proteins that couple Fob1 to the membrane associated CLIP complex (Gartenberg and Smith, 2016). These include the Net1 paralog, Tof2 (with 15 acceptor sites) and the cohibin complex Csm1-Lrs4 (3 and 17 sites, respectively; **Figure 4J and Supplemental Information Table 3**). During mid meiosis, cohibin relocalizes to kinetochores and functions with meiosis-specific Mam1 as the monopolin complex, which mediates monopolar attachment of sister-kinetochores to the MI spindle (Rabitsch et al., 2003). Perhaps reflecting these dynamics, SUMOylation of Csm1 and Lrs4 was high through the G0, S and DSB time points, but then declined (**Figure 4K**). Mam1 was also SUMOylated on three sites, possibly functioning to stabilize monopolin (**Supplemental Information Table 3**). In addition, Csm1-K139-SUMO could influence partner interactions as it lies close to the binding sites for the isopeptidase Ulp2 (which helps maintain silencing)(Liang et al., 2017), and Mam1.

To silence expression at locations other than the rDNA, Sir2 partners with Sir1, 3 and 4, which showed heterogeneous meiotic SUMO profiles suggesting distinct regulation (**Figure 4L**). Siz2-dependent SUMOylation of Sir2 disrupts its interaction with scaffold protein Sir4, causing Sir2 to redistribute from telomeres to the nucleolus (Hannan et al., 2015). In meiosis, Sir2 SUMOylation was relatively high in G0 and S, but dipped at the DSB time point and then remained relatively low suggesting that Sir2 largely redistributes to the nucleolus after S phase (**Figure 4L**). Sir4 was one of the most abundantly SUMOylated meiotic proteins with 30 acceptor sites in three apparent clusters that may differentially affect binding to DNA and histone-tails, and its interactions with partners such as Ku70/80, Rap1, Esc1, Sir2 and Sir3 (Gartenberg and Smith, 2016)(**Figure 4M**). Sir4 SUMOylation increased throughout meiotic prophase roughly in parallel with partners that help recruit it to locations on chromatin (Sir1, Orc1 and Rap1), and to the nuclear envelope (Esc1; **Figure 4N**).

Sir3 binds nucleosomes to facilitate the assembly of silent chromatin domains (Gartenberg and Smith, 2016). In contrast to Sir4, SUMOylation of Sir3 showed a sharp peak during in S phase, was very low in the dHJ/synapsis time point and then increased again as crossovers formed (**Figure 4L** and **4O**). Interestingly, this profile matches closely with that of Asf2, a poorly characterized Sir4 paralog whose overexpression disrupts silencing (Buchberger et al., 2008; Le et al., 1997). Our data suggest that Sir3 may partner Asf2 during meiosis. The general spike in SUMOylation of the SIR complex and its partners at the CO time point suggests late roles, perhaps in preparing chromosomes for segregation.

##### Chromatin remodelers

Chromatin remodelers have well characterized roles in DSB repair in mitotically cycling cells (Seeber et al., 2013), but their roles in meiotic HR are less well characterized. Analysis in *Schizosaccharomyces pombe* invokes roles for multiple remodelers in meiotic HR, including Ino80, Swr1 and Fun30, and the Nap1 and Hir complex histone chaperones (Storey et al., 2018). Notably, in budding yeast, Chd1 was recently shown to act at the dHJ resolution step to facilitate MutLγ-dependent crossing over (Wild et al., 2019). All four families of chromatin remodelers in budding yeast (SWI/SNF, ISWI, CHD, INO80/SWRI), covering eight separate complexes (Ino80, Rsc, Swr1, Swi/Snf, Isw1a, Isw1b, Chrac and Paf1), and the Hir complex were SUMOylated on multiple subunits during meiosis (**Supplemental Information Table 3**). Consistent with a role for SUMOylation in activating the resolution function of Chd1, modification at an N-terminal cluster of four sites occurred only after *P_CUP1_-NDT80* expression, as dHJs were being resolved into crossovers (**Figure 4P**). Cox et al. showed that interaction between mammalian INO80 subunits, INO80E and TFPT, is mediated by SUMO (Cox et al., 2017). We detected SUMOylation on 11/15 yeast INO80 subunits. 10/14 SWR complex subunits were SUMOylated with a U-shaped SUMOylation profile (exemplified by Swc3 and Vps72; **Figure 4Q**). The drop in SUMOylation during S-phase is reminiscent of transcription factors such as Gcn4, suggesting that SUMOylation may suppress their activities. By contrast, bromodomain factor Bdf1, the SWR subunit with an affinity for acetylated H4 (Matangkasombut and Buratowski, 2003), showed a sharp transient SUMOylation peak during DSB formation (**Figure 4R**). Taf14 – a subunit shared between TFIID, TFIIF, INO80, Swi/Snf, and NuA3m with an affinity for acetylated H3 – showed a similar SUMOylation peak suggesting roles in DSB formation or processing.

##### Transcription factors

A majority of SUMOylated transcription factors (31 out of 37) showed a sharp decline in modification level during the transition from G0 to S, consistent with a repressive role for SUMO prior to meiotic entry (**Supplemental Information Table 3**). Sum1, a component of the Sum1-Rfm1-Hst1 complex that represses middle-meiotic genes prior to Ndt80 expression (Winter, 2012), showed a more progressive decline of SUMOylation suggesting a role in anticipation of Ndt80-dependent induction (**Figure 4S**). Coordinated SUMOylation of the Tup1-Ssn6 repressor and Gcn4 transcription factor hastens removal of Gcn4 to enable repression (Ng et al., 2015; Rosonina et al., 2012; Texari et al., 2013). Gcn4 is inferred to influence around a third of meiotic recombination hotspots and deletion of *GCN4* reduces recombination (Abdullah and Borts, 2001). SUMOylation of Tup1 and Gcn4 declined after G0 (**Figure 4T**). Notably, Gcn4-SUMO was lowest at the time of DSB formation and remained diminished thereafter suggesting that SUMO may influence the global DSB landscape via modification of transcription factors.

#### SMCs, Homolog Axes and Synaptonemal Complex

##### SMC (Structural Maintenance of Chromosomes) complexes

SMC complexes are fundamental to meiotic chromosome metabolism, mediating sister-chromatid cohesion, the organization of chromosomes into linear arrays of chromatin loops, chromosome compaction and segregation, and regulating all aspects of recombination (Hunter, 2015; Jessberger, 2002; Zickler and Kleckner, 2015). Mapping meiotic SUMOylation sites onto the six SMC proteins revealed a strong bias for modification of the coiled-coil regions (**Figure 5A**). Helical projections showed that the majority of the sites are solvent exposed and therefore unlikely to alter underlying coiled-coil structures (**Supplemental Information Figure 5**). Given this general propensity and similar SUMOylation profiles (**Figure 5B**), it’s possible that SUMOylation of all SMC proteins has a common function. SUMO could compete or synergize with acetylation to regulate interactions between SMC coiled coils (Kulemzina et al., 2016); or could influence folding at the elbow region (Burmann et al., 2019).

###### Cohesin

All components of meiotic cohesin were SUMOylated on multiple sites: Smc1 (16 sites), Smc3 (14), Pds5 (6), Irr1/Scc3 (3) and the meiosis-specific kleisin Rec8 (16) (**Supplemental Information Table 3**). Although cohesin components other than Rec8 are not meiosis-specific, SUMOylation was largely undetectable in G0 (**Figure 5C**). Rec8-SUMO increased rapidly by S-phase, continued to increase through DSB and SI time points and remained relatively high thereafter. SUMOylations of Smc1 and Smc3 were less intense than Rec8 and although low levels were detected in S phase, big increases were seen later, after DSB formation (**Figure 5B** and **5C**). These profiles are consonant with studies in mitotically cycling cells indicating that SUMOylation is essential for establishment of cohesion in S phase (Almedawar et al., 2012), but also point to later roles. For example, Rec8 SUMOylation could influence its interactions with the HEAT repeat proteins, Pds5 and Irr1/Scc3.

Intriguingly, Pds5 somehow protects the mitotic Kleisin, Mcd1/Scc1 from poly-SUMOylation and ensuing Slx5/8-dependent degradation (D’Ambrosio and Lavoie, 2014) consistent with the observation that overexpression of the Ulp2 SUMO isopeptidase suppresses *pds5* mutant phenotypes (Stead et al., 2003). However, SUMOylation of Mcd1/Scc1 is also important for damage-induced cohesion in mitotic cells (McAleenan et al., 2012; Wu et al., 2012). Pds5 and Rec8 may share similar relationships during meiosis. Rising SUMOylation of Smc1/3 at the SI timepoint was accompanied by transient peaks of Pds5 and Irr1/Scc3 modification (**Figure 5C**; and not shown), which declined thereafter. Pds5-SUMO may be associated with cohesin dynamics occurring as HR ensues and axes mature. The six acceptor sites mapped on Pds5 were dominated by K1103, which is predicted to lie close to the interface with Rec8 and Smc3, with potential to influence ring opening (Hons et al., 2016; Ouyang and Yu, 2017). Finally, the cohesin loader complex, Scc2-Scc4, had a single low intensity site in Scc2, K76, which maps to the interaction interface.

###### Condensin

SUMOylation of all condensin subunits was detected in meiosis and, like cohesin, the Kleisin subunit, Brn1, was the most intense (**Figure 5D** and **Supplemental Information Table 3**). 10 sites were mapped in Brn1 but a single site predominated, K628, which has potential to influence interaction with Ycg1 (Kschonsak et al., 2017). Four other sites (K421, 439, 445 and 457) clustered around the safety-belt loop predicted to trap DNA (Kschonsak et al., 2017). Unlike cohesin, significant SUMOylation of condensin subunits was detected in G0 and then increased steeply after DSB formation, consistent with roles after meiotic S-phase (Yu and Koshland, 2003). The modification profile of Brn1 is similar to proteins involved in SC formation (Zip1, Zip3, Smt3, Ubc9, Ecm11 and Gmc2); and unlike that of Rec8, which shows high levels of SUMOylation already in S-phase, and closely matches the profiles of other axis proteins (Hop1, Red1 and Top2, see below and **Figure 5G**). These observations are consistent with later loading and function of condensin in meiotic prophase. During mitosis, condensin is targeted to the rDNA in part via Ycs4 SUMOylation, which is promoted by the FEAR kinase, Cdc14 (D’Amours et al., 2004). In meiotic prophase, Ycs4-SUMO peaked at the SI time point with a profile analogous to that of Pds5, perhaps reflecting a similar function in regulating condensin dynamics in response to DSBs and nascent synapsis (**Figure 5D and 5E**).

###### Smc5/6 complex

The third eukaryotic SMC complex facilitates DNA replication, repair and recombination and interacts with a dedicated SUMO E3 ligase, Nse2/Mms21, via the Smc5 subunit, and with the Siz1/2 ligases via Nse5 (Bustard et al., 2016; Uhlmann, 2016). In meiosis, Smc5/6 facilitates JM formation and resolution likely through regulation of the Mus81-Mms4 and Sgs1-Top3-Rmi1 complexes (Copsey et al., 2013; Lilienthal et al., 2013; Xaver et al., 2013).

SUMOylation of Smc5, Smc6 and Kleisin Nse4, but not the other five subunits of the complex, was detected in meiosis. Like cohesin, the Smc5/6 complex was not SUMOylated in G0, increased in S phase, as expected, dipped as DSBs formed and then increased thereafter consistent with its roles throughout meiotic prophase (**Figure 5F**)(Copsey et al., 2013; Lilienthal et al., 2013; Xaver et al., 2013).

##### Axis Components Hop1, Red1 and Top2

Red1 and Hop1 augment cohesin-based homolog axes and play essential roles in HR, synapsis and checkpoint control (Hunter, 2006; Subramanian and Hochwagen, 2014; Zickler and Kleckner, 2015). Red1 appears to be recruited to axes principally via an interaction with Rec8, while Hop1 interacts with Red1 (**Figure 5G**)(Smith and Roeder, 1997; Sun et al., 2015; West et al., 2019; West et al., 2018; Woltering et al., 2000). Type-II topoisomerase Top2 is an additional axial component implicated in loop-axis organization and crossover interference (Klein et al., 1992; Moens and Earnshaw, 1989; Zhang et al., 2014). The physical and functional interactions between Rec8, Red1, Hop1 and Top2 were reflected in their SUMOylation profiles, which emerged as a tightly clustered cohort (**Figure 5H**).

In somatic cells, SUMOylation of Top2 has been implicated in multiple aspects of centromere function as well as the removal of abortive cleavage complexes (Cubenas-Potts and Matunis, 2013; Wei et al., 2018; Zhao, 2018). In meiosis, Top2-SUMO facilitates crossover interference together with Red1-SUMO, Sir2 and the STUbL Slx5-Slx8 (Zhang et al., 2014). Our analysis identified two relatively weak N-terminal SUMO sites in addition to the previously mapped C-terminal sites (**Figure 5I**). In Red1, a remarkable 32 conjugated sites were identified, 20 of which clustered into a broad domain (492-702) surrounding a lysine-rich patch previously shown to be important for Red1 SUMOylation, SC assembly and crossover interference (**Figure 5J** and **Supplemental Information Table 3**)(Eichinger and Jentsch, 2010; Zhang et al., 2014). These include sites that overlap regions shown to interact with Mec3 and Ddc1, components of the 9-1-1 checkpoint-sensor clamp (Eichinger and Jentsch, 2010; Zhang et al., 2014), as well as two SUMO-interaction motifs inferred to mediate interaction with the SC transverse-filament protein, Zip1 (Lin et al., 2010). Additional sites were located at the Red1 closure motif, which is bound by Hop1; and in the C-terminal coiled-coil motif that mediates Red1 oligomerization (West et al., 2019; West et al., 2018). Thus, SUMO may influence all aspects of Red1 function.

Hop1 oligomerizes via interactions between the N-terminal HORMA domain and a C-terminal closure motif and thereby activates the checkpoint kinase Mek1 (**Figure 5G** and **5K**)(West et al., 2018). The HORMA domain also anchors Hop1 to homolog axes by interacting with a closure motif in Red1. 17 SUMO-conjugation sites were scattered throughout Hop1, including an N-terminal cluster and sites located around the “safety belt”, which wraps the closure motifs. However, the highest-intensity sites were found immediately adjacent to the closure motif, suggesting effects on oligomerization (**Figure 5K**).

##### Synaptonemal Complex Components Zip1, Ecm11 and Gmc2

SUMO has well documented roles in SC assembly: it localizes to the SC central region and SUMO chains are required for SC formation (Cheng et al., 2006; Voelkel-Meiman et al., 2013). Consistently, SUMO chain formation was concurrent with the SUMOylation of SC central region proteins, Zip1, Ecm11 and Gmc2 (**Figure 5L**). The transverse filament protein, Zip1, activates SUMOylation of Ecm11, which then stimulates Zip1 to oligomerize (Leung et al., 2015). 20 of the 21 conjugation sites in Ecm11 mapped throughout the predicted unstructured N-terminal region, but SUMOylation at K5 was over four orders of magnitude more intense than at any other site (**Figure 5M** and **Supplemental Information Table 3**). K5 was previously shown to be responsible for most Ecm11 SUMOylation and is essential for its function, underscoring the value of our LFQ analysis for identifying functionally relevant conjugation sites (Humphryes et al., 2013; Zavec et al., 2008). Gmc2, the partner of Ecm11, was also SUMOylated at 6 sites (**Supplemental Information Table 3**).

The 23 sites mapped in Zip1 included two clusters at the boundaries between the central helical core and the disordered N- and C-terminal regions (**Figure 5N**). In the context of the bifurcated tetramer model of Dunce et al. (Dunce et al., 2018), SUMO site locations point to possible roles in self-assembly of the Zip1 zipper-like lattice, both at the αN-end head-to-head assembly and the αC-end tetramization regions. Consistent with this possibility, residues 21-163, encompassing the most intense Zip1 SUMO sites, are important for SC assembly and Ecm11 SUMOylation (Voelkel-Meiman et al., 2016). The most N-terminal site, K13, lies in a region important for crossing over and recruitment of the E3 ligase, Zip3 (Voelkel-Meiman et al., 2019). A C-terminal cluster of SUMO sites (K833, 841, 847, 854 and 867) overlaps with a region of Zip1 that interacts with Red1 and includes a SIM that is required for Zip1-Red1 interaction (Zip1 residues 853–864) (Lin et al., 2010). Thus, SUMOylation of Zip1 may modulate its oligomerization, axis association and interaction with Zip3 and perhaps other HR factors.

#### Homologous Recombination

Meiotic HR is physically and functional linked to homolog axes and synaptonemal complexes (summarized in **Figure 6A**)(Zickler and Kleckner, 2015). Our analysis implies that each step of meiotic HR is regulated by SUMO.

##### DSB formation and processing

Spo11-catalyzed DSBs occur in narrow hotspots located in gene promoters marked by H3K4 trimethylation (Pan et al., 2011). Hotspots map to chromatin loops, while the DSB machinery localizes to the axes (Blat et al., 2002; Pan et al., 2011; Panizza et al., 2011). Through the formation of tethered loop-axis complexes, loop sequences become axis associated and DSB formation is triggered (Blat et al., 2002; Lam and Keeney, 2014). 10 factors that assemble into four complexes are essential for DSB formation, *viz.* Spo11-Ski8, Rec102-Rec104, Mer2^IHO1^-Rec114-Mei4 and Mre11-Rad50-Xrs2^NBS1^ (MRX)(Lam and Keeney, 2014). DSB formation is also strongly, through not completely, dependent on axis associated factors Hop1, Red1 and Rec8-cohesin, all of which were heavily SUMOylated, as described above (**Figure 5**). Intriguingly, two key regulatory DSB factors, Spp1 and Mer2, were SUMOylated at multiple sites (**Figure 6B**). Spp1 reads H3K4 trimethylation marks using its PHD finger and recruits DSB hotspots to the axis via an interaction with Mer2 (Acquaviva et al., 2013; Adam et al., 2018; Sommermeyer et al., 2013). Mer2^IHO1^ is recruited to axes via Red1 and Hop1, where its cell cycle and replication-dependent phosphorylation leads to association with other DSB factors, Rec114-Mei4 and Xrs2, to trigger DSB formation only on replicated sister chromatids (Lam and Keeney, 2014). Our data revealed sharp SUMOylation peaks for both Spp1 and Mer2 at the DSB time point (**Figure 6B**), suggesting roles in initiating HR. The 13 SUMO sites in Mer2 do not overlap with known phosphorylation sites suggesting parallel regulation. Low intensity SUMO sites were also identified on Rec114 and two subunits of the H3K4 methyltranferase (**Figure 6A** and **Supplemental Information Table 3**).

Processing of DSB ends to liberate Spo11-oligo complexes and produce long 3’ single-stranded tails involves Sae2^CtIP^, the endonuclease and 3’-5’ exonuclease activities of the MRX complex, and the 5’-3’ exonuclease Exo1 (Lam and Keeney, 2014; Zakharyevich et al., 2010). Only SUMOylation of Sae2^CtIP^ was detected in meiosis, at a site previously shown to enhance Sae2^CtIP^ solubility and facilitate its DSB resection function (Sarangi et al., 2015). However, Sae2-SUMO signal was weak and only detected in G0 and S time point samples.

##### Homology search and DNA strand exchange

Mediators help the strand-exchange proteins, Rad51 and Dmc1, displacing RPA from single-stranded DNA and thereby assemble into nucleoprotein filaments (Brown and Bishop, 2014). Mediators include Rad52, which was SUMOylated at 10 sites, including those previously shown to be important in vegetative cells (K10/11 and 220). During meiosis, Rad52 mediates assembly of Rad51, but not Dmc1, and is important for post-invasion strand-annealing steps during dHJ formation and non-crossover formation (Gasior et al., 1998; Lao et al., 2008). The SUMOylation profiles of Rad52 and its paralog, Rad59 were tightly matched, rising at the time of DSB formation and continuing to rise during strand invasion, before rising again at the time of crossing over (**Figure 6C**). Diverse effects of Rad52 SUMOylation have been reported in mitotically cycling cells (Dhingra and Zhao, 2019), including both enhancing and disfavoring Rad52-Rad51 interaction and HR; and promoting interaction with Rad59 to favor Rad51-indpendent repair such as single-strand annealing (SSA). Analysis of Rad52/Rad59 SUMOylation in the context of meiotic HR may help disentangle these seemingly contradictory functions.

Rfa1, the large subunit of the trimeric RPA complex, was also SUMOylated at five sites, K68, 170, 180, 411, 427 (**Supplemental Information Table 3**). The latter four sites enhance DNA-damage induced checkpoint signaling in vegetative cells by strengthening interaction with the DNA helicase, Sgs1 (Dhingra et al., 2019). Rfa1-SUMO levels were low until the SI time point, during which they increased dramatically and then remained high (**Figure 6D**). This profile argues against an early role in Dmc1/Rad51 assembly and suggests that D-loop associated Rfa1 becomes SUMOylated, perhaps to regulate post-invasion steps including DNA synthesis, and strand annealing during dHJ formation and noncrossover formation.

Dmc1 and Rad51 cooperate during meiosis to bias strand-exchange between homologs (Brown and Bishop, 2014; Cloud et al., 2012; Lao et al., 2013). Dmc1 catalyzes the majority of strand-exchange in concert with its essential accessory factor Hop1-Mnd1, while Rad51 plays an essential support function. In fact, efficient inter-homolog bias involves inhibition of Rad51 strand-exchange activity by a meiosis-specific factor Hed1 (Busygina et al., 2008). Dmc1, Rad51, Hop1, Mnd1 and Hed1 were all SUMOylated during meiosis (**Supplemental Information Table 3**). Dmc1 was the most prominent of these, with a profile similar to that of Rfa1, except that levels dipped after strand-exchange, at the dHJ time point, before rising again as crossovers formed (**Figure 6D**). Five of the six SUMO sites in Dmc1 clustered in the N-terminal domain (**Figure 6E**), which is conserved in Rad51 and implicated in filament subunit interactions, DNA binding and ATPase activation (Galkin et al., 2006; Zhang et al., 2005).

##### Crossover/noncrossover differentiation

The ZMMs (Zip1, Zip2, Zip3, Zip4, Spo16, Msh4-Msh5 and Mer3) are conserved meiosis-specific pro-crossover factors that locally couple HR to synapsis (Hunter, 2015). All ZMMs other than Spo16 were SUMOylated coincident with strand invasion and synapsis (**Figure 6F**). Intensities varied by at least an order of magnitude, with Zip1 and Zip3 (discussed above) being the most prominent (red scale in **Figure 6F**). Both subunits of the HJ-binding MutSγ complex, Msh4-Msh5, were SUMOylated for a total of 11 sites, with the most intense sites located in the ABC ATPase domains that interface to form two composite ATPase sites (Warren et al., 2007). The DNA helicase Mer3 promotes dHJ formation while limiting the extent of heteroduplex the (Borner et al., 2004; Duroc et al., 2017). Mer3 SUMO sites were located predominantly in the disordered the N- and C-termini that may mediate protein-protein interactions and/or protein stability (**Supplemental Information, Protein Diagrams**).

##### Joint molecule (JM) resolution

The conserved endonuclease MutLγ (Mlh1-Mlh3) works with a second DNA mismatch-repair factor Exo1, and the chromatin remodeler Chd1 to promote crossover-biased resolution of dHJs (Wild et al., 2019; Zakharyevich et al., 2010; Zakharyevich et al., 2012). As described above, Chd1 was SUMOylated specifically at the time of JM resolution (**Figure 4P**), but modification of MutLγ and Exo1 was not detected. The structure-selective endonuclease Mus81-Mms4^EME1^ and the Smc5/6 complex define a second JM resolvase pathway (**Figure 5F**). While modification of Mus81-Mms4^EME1^ was not detected, SUMOylation of Smc5/6 components continued to rise during resolution (discussed above and **Figure 5F**). The STR complex, Sgs1^BLM^-Top3-Rmi1, is a potent decatenase that functions as a noncrossover-specific dHJ “dissolavse”, but is also a general mediator of JM formation and facilittaes MutLγ-Exo1 dependent crossing over (Hunter, 2015). In vegetative cells, of Sgs1 SUMOylation fosters interaction with Top3, facilitates JM resolution and suppresses crossing over (Bermudez-Lopez et al., 2016; Bonner et al., 2016). Both Sgs1 and Top3, but not Rmi1, were SUMOylated during meiosis (**Figure 6G** and **Supplemental Table 3**). Sgs1-SUMO rose dramatically with strand exchange and rose again during crossover formation. By comparison, Top3-SUMO was very weak and only detected in the SI and CO time points. Contrary to the inference that modification of Sgs1 in vegetative cells occurs primarily at three consensus sites (K175, 621, and 831)(Bermudez-Lopez et al., 2016), meiotic SUMOylation occurred at six non- and partial-consensus sites, dominated by K26 (**Figure 6H**). K26 and a second SUMO site, K35, lie in the Top3-interaction domain of Sgs1 and may influence their interaction.

### SUMOylation is required throughout meiotic prophase I

The diversity of targets identified by our analysis points to roles for SUMOylation throughout meiotic prophase I and at each step of HR. To test this assertion and define execution points for SUMOylation we employed an auxin-induced degron (AID) allele of the E1 subunit Aos1 to acutely block *de novo* SUMOylation at four key transition points (**Figure 7A** and **7B**; very similar results were obtained using a Uba2-AID degron allele, **Supplemental Information Figure 7**). In each case, meiotic cultures were split at the appropriate time point, and auxin was added to one sub-culture to induce degradation of Aos1-AID. When Aos1-AID degradation was induced 30 mins after entry into meiosis (condition 1, **Figure 7B and 7C**), the onset of S-phase, as assessed by FACS analysis, was delayed by ≥30 mins and meiotic divisions were completely blocked (**Figure 7D and 7E**). Identified SUMOylation targets involved in DNA replication, cell cycle and metabolic regulation could be responsible for these phenotypes (**Supplemental Information Table 3**).

**Figure 7.**
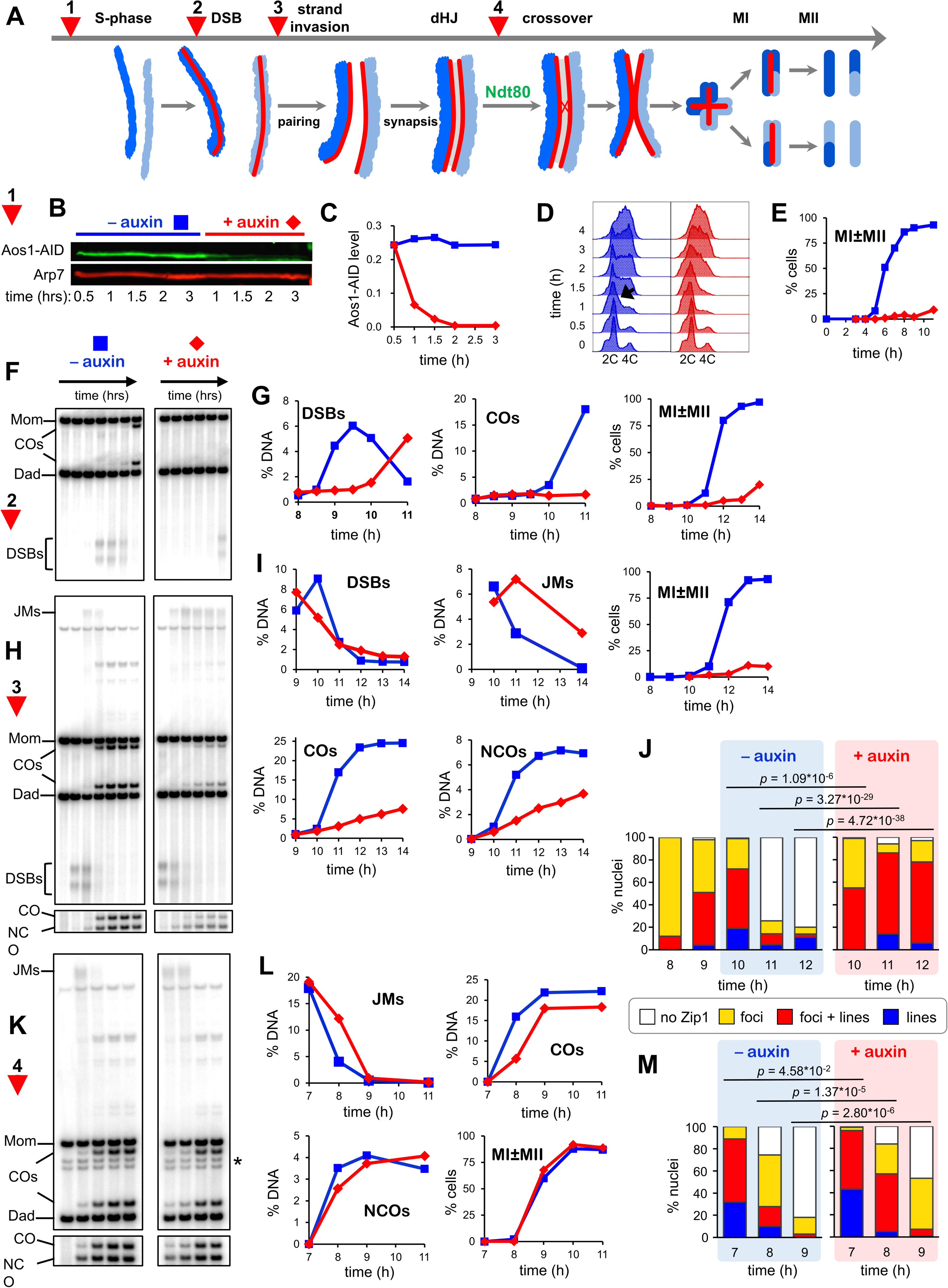
SUMOylation functions throughout meiosis. (A) Cartoon showing the four points (experiments 1-4) at which *de novo* SUMOylation was acutely inactivated. (B) Immunoblot of Aos1-AID with and without addition of auxin at 30 mins (experiment 1). Arp7 was used as loading control. (C) Quantification of the blot shown in (B). (D) Flow cytometry analysis of S-phase progression for experiment 1. The black arrow indicates the onset of S-phase in control cells. (E) Meiotic nuclear divisions (MI ± MII) for experiment 1. (F) 1D gel Southern blot image for analysis of DSBs and crossovers in experiment 2. (G) Quantification of DSBs, crossovers and meiotic divisions for experiment 2. (H) 1D gel Southern blot images for experiment 3. The top gel was used to quantify DSBs and crossovers (COs). The bottom gel was used to quantify non-crossover products (NCOs). (I) Quantification of DSBs, joint molecules (JMs), COs, NCOs and meiotic divisions for experiment 3. JMs were quantified from 2D gel Southern analysis (Supplemental Information Figure 6). (J) Quantification of synapsis (Zip1-staining classes) for experiment 3 (Supplemental Information Figure 8). (K) 1D gel Southern blot images for experiment 4. The top gel was used to quantify DSBs and crossovers (COs). The bottom gel was used to quantify non-crossover products (NCOs). (L) Quantification of DSBs, joint molecules (JMs), COs, NCOs and meiotic divisions for experiment 3. JMs were quantified from 2D gel Southern analysis (Supplemental Information Figure 6). (M) Quantification of synapsis (Zip1-staining classes) for experiment 4.

To uncover post S-phase functions of SUMO that could account for the block to meiotic divisions, cells were synchronized using an analog-sensitive allele of the Cdc7 kinase (*cdc7-as*; condition 2, **Figure 7F,G**)(Wan et al., 2006). Treatment of *cdc7-as* cells with the ATP analog PP1 causes meiotic cultures to arrest after S-phase, but prior to the initiation of recombination by DSB formation. The DNA events of HR were monitored using Southern blot assays at a well-characterized DSB hotspot (**Supplemental Information Figure 8A–E**); and synapsis was analyzed by immunostaining chromosome spreads for Zip1 (**Supplemental Information Figure 8F**). Degradation of Aos1-AID immediately following release from *cdc7-as* arrest blocked DSB formation for ∼2 hrs indicating an unanticipated role for SUMO in the initiation of HR. Identified targets that could affect DSB formation include cohesin, Hop1, Red1, Spp1, Mer2, Rec114 and Dbf4 (discussed above; **Supplemental Information Table 3**).

Condition 2 also resulted in a complete block to crossing over and meiotic divisions (**Figure 7G**), suggesting additional roles for SUMO in HR and/or the progression of meiosis. Therefore, we also determined the effects of degrading Aos1-AID later, just after DSBs were formed following release from *cdc7-as* arrest (condition 3, **Figure 7H,I**). In control cells (no auxin), initial DSB levels were high and continued to rise for one hour, before being repaired to yield high levels of crossovers and noncrossovers. When Aos1-AID was degraded, initial DSB levels were also high, but instead of continuing to rise, levels immediately decreased. These observations are consistent with the role for SUMO in DSB formation defined above, with later degradation of Aos1-AID preventing only later forming DSBs. JMs appeared to form efficiently following Aos1-AID degradation, but reached peak levels with a ∼1 hr delay relative to control cells suggesting slower formation (Figure 7I and Supplemental Information Figure 6). Subsequent resolution of JMs was severely delayed and cells again failed to divide. Consistent with a JM resolution defect, crossover and non-crossover products were reduced by 70% and 48%, respectively (**Figure 7I**).

For condition 3, we also analyzed SC formation by quantifying four classes of Zip1 staining nuclei: no staining, foci only, partial synapsis with both lines and foci, and full synapsis with extensive linear staining (**Figure 7J** and **Supplemental Figure 8F**. Both synapsis and, unexpectedly, de-synapsis were defective when Aos1-AID was degraded. Without auxin, synapsis levels peaked at 10 hrs (1 hr after the culture was split), with partial synapsis in 53% of nuclei and full SCs in 19%. De-synapsis rapidly ensued and by 11 hrs, 75% of cells had no Zip1 staining. By comparison, Aos1-AID degradation appeared to stall synapsis, with levels remaining largely unchanged between 9 and 10 hrs (*P=* 0.09, *G*-test; **Figure 7J**). However, by 11 hrs, very high levels of synapsis were achieved; 72% of cells had partial synapsis and 14% had full SCs. At 12 hrs, synapsis levels were almost unchanged suggesting defective de-synapsis. Thus, *de novo* SUMOylation has both a post-DSB function to promote the timely formation of JMs and SCs, and a post-synapsis function in JM resolution and de-synapsis. These roles of SUMO may be mediated by the numerous targets identified in processes such as HR, SC formation, the DNA damage response, and transcription (**Figures 5 and 6; Supplemental Information Table 3**).

Finally, late roles of *de novo* SUMOylation were determined by degrading Aos1-AID as cells were released form pachytene arrest (condition 4; **Figure 7A**), using the *P_GAL_-NDT80* allele (see above and **Figure 1A**)(Benjamin et al., 2003). In contrast to the other conditions, meiotic divisions occurred efficiently when Aos1-AID was degraded at this late stage, suggesting that *de novo* SUMOylation is not essential for meiotic divisions (**Figure 7K,L**). However, when a degron allele of Uba2 was degraded, a reproducible delay in MI was observed and spore viability was reduced to 76% compared to 91% in the no auxin control (**Supplementary Information Figure 7**). This may be a consequence of more acute inactivation of SUMOylation due to faster and more complete degradation, and/or the fact that Uba2 is the catalytic subunit of E1. Although divisions occurred efficiently when Aos1-AID/Uba2-AID were degraded, JM resolution and product formation were delayed by ∼30 minutes, and final crossover levels were reduced by ∼18% (**Figure 7K,L** and **Supplemental Information Figure 7**). These HR defects were accompanied by a delay in de-synapsis (**Figure 7M** and **Supplemental Information Figure 7**). Without auxin, only 25% of cells still had partial or full synapsis one hour after *NDT80-IN* expression, compared to 57% when Aos1-AID was degraded. After two hours, SCs had completely disassembled in 82% of control cells without auxin, compared to 47% following Aos1-AID degradation. Late defects caused by E1 inactivation could reflect SUMO targets such as Sgs1, Top3, Slx4, Smc5/6, Chd1, the ZMMs and components of SCs.

Collectively, real-time inactivation of the SUMO E1 enzyme confirms that SUMOylation regulates the major transitions of meiotic prophase and identifies roles in S-phase, DSB formation, the formation and resolution of joint molecules, synapsis and de-synapsis, and the progression of meiotic prophase I. In summary, our mass spectrometry dataset marks a breakthrough for the field, delineating a diverse and dynamic meiotic SUMO-modified proteome and providing a rich resource for functional analyses, to identify pertinent targets and define regulatory networks. With this unique dataset in hand, major advances in understanding how SUMO regulates meiotic prophase are anticipated.

## Supporting information

Key MaxQuant Tables

Supplemental Information - Protein Diagrams

## EXPERIMENTAL PROCEDURES

Extended methods are described in the Supplemental Information.

## SUPPLEMENTAL INFORMATION

Supplemental Information includes 8 figures, 4 tables, protein diagrams and extended methods.

## ACKNOWLEDGEMENTS

We thank Chris Lima (MSKCC), Kevin Corbett (UC San Diego), Katrin Karbstein (The Scripps Research Institute), Ilan Attali and James Berger (Johns Hopkins University School of Medicine), Mike Botchan (UC Berkeley), Wolf Heyer (UC Davis) and members of the Hunter Lab for support and discussions. Anthony Herren and Brett Phinney (UC Davis Proteomics Core) provided dedicated proteomics services. pFA6a-kanMX6-PGAL1-HBH was a gift from Peter Kaiser and Hongwei Zhang (UC Irvine). This work was supported by NIH NIGMS grant GM074223 to N.H. S.O. was supported by NIH NIGMS T32 Training Program in Molecular and Cellular Biology 5T32GM007377 and an F31 Ruth L. Kirschstein National Research Service Award 1F31GM125106. M.I. was supported by a JSPS postdoctoral fellowship. N.H. is an Investigator of the Howard Hughes Medical Institute.

## AUTHOR CONTRIBUTIONS

N.B. and N.H. conceived the study and designed the experiments. S.O. and M.I. performed experiments in Figure 7. P.P. and N.B. constructed strains for proteomics analysis. J.B., P.P., A.D., S.K., M.M. and N.B. performed proteomics experiments. J.J. helped optimize proteomics protocols. O.D. performed protein structure analysis. S.C analyzed SUMO site distribution and protein secondary structure and generated protein diagrams. N.B. and N.H. wrote the manuscript with inputs and edits from all authors.

## DECLARATION OF INTERESTS

The authors declare no competing interests.

## Notes

https://www.ebi.ac.uk/pride/archive/projects/PXD012418

## REFERENCES

Abdullah, M.F., and Borts, R.H. (2001). Meiotic recombination frequencies are affected by nutritional states in Saccharomycescerevisiae. Proceedings of the National Academy of Sciences of the United States of America 98, 14524–14529.

Acquaviva, L., Szekvolgyi, L., Dichtl, B., Dichtl, B.S., de La Roche Saint Andre, C., Nicolas, A., and Geli, V. (2013). The COMPASS subunit Spp1 links histone methylation to initiation of meiotic recombination. Science 339, 215–218.

Adam, C., Guerois, R., Citarella, A., Verardi, L., Adolphe, F., Beneut, C., Sommermeyer, V., Ramus, C., Govin, J., Coute, Y., et al. (2018). The PHD finger protein Spp1 has distinct functions in the Set1 and the meiotic DSB formation complexes. PLoS genetics 14, e1007223.

Agarwal, S., and Roeder, G.S. (2000). Zip3 provides a link between recombination enzymes and synaptonemal complex proteins. Cell 102, 245–255.

Ahuja, J.S., Sandhu, R., Mainpal, R., Lawson, C., Henley, H., Hunt, P.A., Yanowitz, J.L., and Borner, G.V. (2017). Control of meiotic pairing and recombination by chromosomally tethered 26S proteasome. Science 355, 408–411.

Albuquerque, C.P., Wang, G., Lee, N.S., Kolodner, R.D., Putnam, C.D., and Zhou, H. (2013). Distinct SUMO ligases cooperate with Esc2 and Slx5 to suppress duplication-mediated genome rearrangements. PLoS genetics 9, e1003670.

Albuquerque, C.P., Yeung, E., Ma, S., Fu, T., Corbett, K.D., and Zhou, H. (2015). A Chemical and Enzymatic Approach to Study Site-Specific Sumoylation. PloS one 10, e0143810.

Allers, T., and Lichten, M. (2001). Differential timing and control of noncrossover and crossover recombination during meiosis. Cell 106, 47–57.

Almedawar, S., Colomina, N., Bermudez-Lopez, M., Pocino-Merino, I., and Torres-Rosell, J. (2012). A SUMO-dependent step during establishment of sister chromatid cohesion. Current biology : CB 22, 1576–1581.

Altaf, M., Auger, A., Monnet-Saksouk, J., Brodeur, J., Piquet, S., Cramet, M., Bouchard, N., Lacoste, N., Utley, R.T., Gaudreau, L., et al. (2010). NuA4-dependent acetylation of nucleosomal histones H4 and H2A directly stimulates incorporation of H2A.Z by the SWR1 complex. The Journal of biological chemistry 285, 15966–15977.

Bani Ismail, M., Shinohara, M., and Shinohara, A. (2014). Dot1-dependent histone H3K79 methylation promotes the formation of meiotic double-strand breaks in the absence of histone H3K4 methylation in budding yeast. PloS one 9, e96648.

Bao, X., Johnson, J.L., and Rao, H. (2015). Rad25 protein is targeted for degradation by the Ubc4-Ufd4 pathway. The Journal of biological chemistry 290, 8606–8612.

Baranes-Bachar, K., Khalaila, I., Ivantsiv, Y., Lavut, A., Voloshin, O., and Raveh, D. (2008). New interacting partners of the F-box protein Ufo1 of yeast. Yeast 25, 733–743.

Benjamin, K.R., Zhang, C., Shokat, K.M., and Herskowitz, I. (2003). Control of landmark events in meiosis by the CDK Cdc28 and the meiosis-specific kinase Ime2. Genes & development 17, 1524–1539.

Berchowitz, L.E., Gajadhar, A.S., van Werven, F.J., De Rosa, A.A., Samoylova, M.L., Brar, G.A., Xu, Y., Xiao, C., Futcher, B., Weissman, J.S., et al. (2013). A developmentally regulated translational control pathway establishes the meiotic chromosome segregation pattern. Genes & development 27, 2147–2163.

Bergink, S., Ammon, T., Kern, M., Schermelleh, L., Leonhardt, H., and Jentsch, S. (2013). Role of Cdc48/p97 as a SUMO-targeted segregase curbing Rad51-Rad52 interaction. Nature cell biology 15, 526–532.

Bermudez-Lopez, M., Villoria, M.T., Esteras, M., Jarmuz, A., Torres-Rosell, J., Clemente-Blanco, A., and Aragon, L. (2016). Sgs1’s roles in DNA end resection, HJ dissolution, and crossover suppression require a two-step SUMO regulation dependent on Smc5/6. Genes & development 30, 1339–1356.

Bernier-Villamor, V., Sampson, D.A., Matunis, M.J., and Lima, C.D. (2002). Structural basis for E2-mediated SUMO conjugation revealed by a complex between ubiquitin-conjugating enzyme Ubc9 and RanGAP1. Cell 108, 345–356.

Blat, Y., Protacio, R.U., Hunter, N., and Kleckner, N. (2002). Physical and functional interactions among basic chromosome organizational features govern early steps of meiotic chiasma formation. Cell 111, 791–802.

Bonner, J.N., Choi, K., Xue, X., Torres, N.P., Szakal, B., Wei, L., Wan, B., Arter, M., Matos, J., Sung, P., et al. (2016). Smc5/6 Mediated Sumoylation of the Sgs1-Top3-Rmi1 Complex Promotes Removal of Recombination Intermediates. Cell reports 16, 368–378.

Borde, V., Robine, N., Lin, W., Bonfils, S., Geli, V., and Nicolas, A. (2009). Histone H3 lysine 4 trimethylation marks meiotic recombination initiation sites. The EMBO journal 28, 99–111.

Borner, G.V., Kleckner, N., and Hunter, N. (2004). Crossover/noncrossover differentiation, synaptonemal complex formation, and regulatory surveillance at the leptotene/zygotene transition of meiosis. Cell 117, 29–45.

Bose, R., Manku, G., Culty, M., and Wing, S.S. (2014). Ubiquitin-proteasome system in spermatogenesis. Adv Exp Med Biol 759, 181–213.

Brar, G.A., Yassour, M., Friedman, N., Regev, A., Ingolia, N.T., and Weissman, J.S. (2012). High-resolution view of the yeast meiotic program revealed by ribosome profiling. Science 335, 552–557.

Brown, M.S., and Bishop, D.K. (2014). DNA strand exchange and RecA homologs in meiosis. Cold Spring Harbor perspectives in biology 7, a016659.

Bryant, J.M., Govin, J., Zhang, L., Donahue, G., Pugh, B.F., and Berger, S.L. (2012). The linker histone plays a dual role during gametogenesis in Saccharomyces cerevisiae. Molecular and cellular biology 32, 2771–2783.

Buchberger, J.R., Onishi, M., Li, G., Seebacher, J., Rudner, A.D., Gygi, S.P., and Moazed, D. (2008). Sir3-nucleosome interactions in spreading of silent chromatin in Saccharomyces cerevisiae. Molecular and cellular biology 28, 6903–6918.

Burmann, F., Lee, B.G., Than, T., Sinn, L., O’Reilly, F.J., Yatskevich, S., Rappsilber, J., Hu, B., Nasmyth, K., and Lowe, J. (2019). A folded conformation of MukBEF and cohesin. Nature structural & molecular biology 26, 227–236.

Bustard, D.E., Ball, L.G., and Cobb, J.A. (2016). Non-Smc element 5 (Nse5) of the Smc5/6 complex interacts with SUMO pathway components. Biol Open 5, 777–785.

Busygina, V., Sehorn, M.G., Shi, I.Y., Tsubouchi, H., Roeder, G.S., and Sung, P. (2008). Hed1 regulates Rad51-mediated recombination via a novel mechanism. Genes & development 22, 786–795.

Cahoon, C.K., and Hawley, R.S. (2016). Regulating the construction and demolition of the synaptonemal complex. Nature structural & molecular biology 23, 369–377.

Cappadocia, L., Pichler, A., and Lima, C.D. (2015). Structural basis for catalytic activation by the human ZNF451 SUMO E3 ligase. Nature structural & molecular biology 22, 968–975.

Cheng, C.H., Lin, F.M., Lo, Y.H., and Wang, T.F. (2007). Tying SUMO modifications to dynamic behaviors of chromosomes during meiotic prophase of Saccharomyces cerevisiae. J Biomed Sci 14, 481–490.

Cheng, C.H., Lo, Y.H., Liang, S.S., Ti, S.C., Lin, F.M., Yeh, C.H., Huang, H.Y., and Wang, T.F. (2006). SUMO modifications control assembly of synaptonemal complex and polycomplex in meiosis of Saccharomyces cerevisiae. Genes & development 20, 2067–2081.

Cheng, Y.H., Chuang, C.N., Shen, H.J., Lin, F.M., and Wang, T.F. (2013). Three distinct modes of Mec1/ATR and Tel1/ATM activation illustrate differential checkpoint targeting during budding yeast early meiosis. Molecular and cellular biology 33, 3365–3376.

Cheng, Z., Otto, G.M., Powers, E.N., Keskin, A., Mertins, P., Carr, S.A., Jovanovic, M., and Brar, G.A. (2018). Pervasive, Coordinated Protein-Level Changes Driven by Transcript Isoform Switching during Meiosis. Cell 172, 910–923 e916.

Chi, P., Kwon, Y., Visnapuu, M.L., Lam, I., Santa Maria, S.R., Zheng, X., Epshtein, A., Greene, E.C., Sung, P., and Klein, H.L. (2011). Analyses of the yeast Rad51 recombinase A265V mutant reveal different in vivo roles of Swi2-like factors. Nucleic acids research 39, 6511–6522.

Chu, S., and Herskowitz, I. (1998). Gametogenesis in yeast is regulated by a transcriptional cascade dependent on Ndt80. Molecular cell 1, 685–696.

Chua, P.R., and Roeder, G.S. (1998). Zip2, a meiosis-specific protein required for the initiation of chromosome synapsis. Cell 93, 349–359.

Cloud, V., Chan, Y.L., Grubb, J., Budke, B., and Bishop, D.K. (2012). Rad51 is an accessory factor for Dmc1-mediated joint molecule formation during meiosis. Science 337, 1222–1225.

Clyne, R.K., Katis, V.L., Jessop, L., Benjamin, K.R., Herskowitz, I., Lichten, M., and Nasmyth, K. (2003). Polo-like kinase Cdc5 promotes chiasmata formation and cosegregation of sister centromeres at meiosis I. Nature cell biology 5, 480–485.

Copsey, A., Jordan, P.W., Tang, S., Blitzblau, H., Chan, A.C., Newcombe, S., Newnham, L., Li, A., Arumugam, P., Hochwagen, A., et al. (2013). The Smc5-Smc6 Complex Promotes Chromosome Resolution at Meiosis I By Regulating Meiotic Cohesin and Promoting Joint Molecule Resolution. PLoS genetics 9, e1004071.

Cox, E., Hwang, W., Uzoma, I., Hu, J., Guzzo, C.M., Jeong, J., Matunis, M.J., Qian, J., Zhu, H., and Blackshaw, S. (2017). Global Analysis of SUMO-Binding Proteins Identifies SUMOylation as a Key Regulator of the INO80 Chromatin Remodeling Complex. Mol Cell Proteomics 16, 812–823.

Cox, J., Hein, M.Y., Luber, C.A., Paron, I., Nagaraj, N., and Mann, M. (2014). Accurate proteome-wide label-free quantification by delayed normalization and maximal peptide ratio extraction, termed MaxLFQ. Mol Cell Proteomics 13, 2513–2526.

Cox, J., and Mann, M. (2008). MaxQuant enables high peptide identification rates, individualized p.p.b.-range mass accuracies and proteome-wide protein quantification. Nat Biotechnol 26, 1367–1372.

Crichton, J.H., Playfoot, C.J., and Adams, I.R. (2014). The role of chromatin modifications in progression through mouse meiotic prophase. J Genet Genomics 41, 97–106.

Cubenas-Potts, C., and Matunis, M.J. (2013). SUMO: a multifaceted modifier of chromatin structure and function. Dev Cell 24, 1–12.

D’Ambrosio, L.M., and Lavoie, B.D. (2014). Pds5 Prevents the PolySUMO-Dependent Separation of Sister Chromatids. Current Biology 24, 361–371.

D’Amours, D., Stegmeier, F., and Amon, A. (2004). Cdc14 and condensin control the dissolution of cohesin-independent chromosome linkages at repeated DNA. Cell 117, 455–469.

Davis-Roca, A.C., Divekar, N.S., Ng, R.K., and Wignall, S.M. (2018). Dynamic SUMO remodeling drives a series of critical events during the meiotic divisions in C. elegans. PLoS genetics 14, e1007626.

de Carvalho, C.E., and Colaiacovo, M.P. (2006). SUMO-mediated regulation of synaptonemal complex formation during meiosis. Genes & development 20, 1986–1992.

Dhingra, N., Wei, L., and Zhao, X. (2019). Replication protein A (RPA) sumoylation positively influences the DNA damage checkpoint response in yeast. The Journal of biological chemistry 294, 2690–2699.

Dhingra, N., and Zhao, X. (2019). Intricate SUMO-based control of the homologous recombination machinery. Genes & development 33, 1346–1354.

Dobson, M.J., Pickett, A.J., Velmurugan, S., Pinder, J.B., Barrett, L.A., Jayaram, M., and Chew, J.S. (2005). The 2 microm plasmid causes cell death in Saccharomyces cerevisiae with a mutation in Ulp1 protease. Molecular and cellular biology 25, 4299–4310.

Dohmen, R.J. (2004). SUMO protein modification. Biochimica et biophysica acta 1695, 113–131.

Downs, J.A., Kosmidou, E., Morgan, A., and Jackson, S.P. (2003). Suppression of homologous recombination by the Saccharomyces cerevisiae linker histone. Molecular cell 11, 1685–1692.

Downs, J.A., Lowndes, N.F., and Jackson, S.P. (2000). A role for Saccharomyces cerevisiae histone H2A in DNA repair. Nature 408, 1001–1004.

Dresser, M.E., Ewing, D.J., Conrad, M.N., Dominguez, A.M., Barstead, R., Jiang, H., and Kodadek, T. (1997). DMC1 functions in a Saccharomyces cerevisiae meiotic pathway that is largely independent of the RAD51 pathway. Genetics 147, 533–544.

Dunce, J.M., Dunne, O.M., Ratcliff, M., Millan, C., Madgwick, S., Uson, I., and Davies, O.R. (2018). Structural basis of meiotic chromosome synapsis through SYCP1 self-assembly. Nature structural & molecular biology 25, 557–569.

Duroc, Y., Kumar, R., Ranjha, L., Adam, C., Guerois, R., Md Muntaz, K., Marsolier-Kergoat, M.C., Dingli, F., Laureau, R., Loew, D., et al. (2017). Concerted action of the MutLbeta heterodimer and Mer3 helicase regulates the global extent of meiotic gene conversion. eLife 6.

Eichinger, C.S., and Jentsch, S. (2010). Synaptonemal complex formation and meiotic checkpoint signaling are linked to the lateral element protein Red1. Proceedings of the National Academy of Sciences of the United States of America 107, 11370–11375.

Esteras, M., Liu, I.C., Snijders, A.P., Jarmuz, A., and Aragon, L. (2017). Identification of SUMO conjugation sites in the budding yeast proteome. Microb Cell 4, 331–341.

Finley, D., Ulrich, H.D., Sommer, T., and Kaiser, P. (2012). The ubiquitin-proteasome system of Saccharomyces cerevisiae. Genetics 192, 319–360.

Flotho, A., and Melchior, F. (2013). Sumoylation: a regulatory protein modification in health and disease. Annual review of biochemistry 82, 357–385.

Fraune, J., Schramm, S., Alsheimer, M., and Benavente, R. (2012). The mammalian synaptonemal complex: protein components, assembly and role in meiotic recombination. Experimental cell research 318, 1340–1346.

Fuchs, G., and Oren, M. (2014). Writing and reading H2B monoubiquitylation. Biochimica et biophysica acta 1839, 694–701.

Galkin, V.E., Wu, Y., Zhang, X.P., Qian, X., He, Y., Yu, X., Heyer, W.D., Luo, Y., and Egelman, E.H. (2006). The Rad51/RadA N-terminal domain activates nucleoprotein filament ATPase activity. Structure 14, 983–992.

Gao, J., and Colaiacovo, M.P. (2018). Zipping and Unzipping: Protein Modifications Regulating Synaptonemal Complex Dynamics. Trends in genetics : TIG 34, 232–245.

Gareau, J.R., and Lima, C.D. (2010). The SUMO pathway: emerging mechanisms that shape specificity, conjugation and recognition. Nature reviews. Molecular cell biology 11, 861–871.

Gartenberg, M.R., and Smith, J.S. (2016). The Nuts and Bolts of Transcriptionally Silent Chromatin in Saccharomyces cerevisiae. Genetics 203, 1563-+.

Gasior, S.L., Wong, A.K., Kora, Y., Shinohara, A., and Bishop, D.K. (1998). Rad52 associates with RPA and functions with rad55 and rad57 to assemble meiotic recombination complexes. Genes & development 12, 2208–2221.

Georgieva, M., Roguev, A., Balashev, K., Zlatanova, J., and Miloshev, G. (2012). Hho1p, the linker histone of Saccharomyces cerevisiae, is important for the proper chromatin organization in vivo. Biochimica et biophysica acta 1819, 366–374.

Govin, J., and Berger, S.L. (2009). Genome reprogramming during sporulation. Int J Dev Biol 53, 425–432.

Grant, P.A., Eberharter, A., John, S., Cook, R.G., Turner, B.M., and Workman, J.L. (1999). Expanded lysine acetylation specificity of Gcn5 in native complexes. The Journal of biological chemistry 274, 5895–5900.

Gray, S., and Cohen, P.E. (2016). Control of Meiotic Crossovers: From Double-Strand Break Formation to Designation. Annual review of genetics 50, 175–210.

Hale, T.K., Contreras, A., Morrison, A.J., and Herrera, R.E. (2006). Phosphorylation of the linker histone H1 by CDK regulates its binding to HP1alpha. Molecular cell 22, 693–699.

Hannan, A., Abraham, N.M., Goyal, S., Jamir, I., Priyakumar, U.D., and Mishra, K. (2015). Sumoylation of Sir2 differentially regulates transcriptional silencing in yeast. Nucleic acids research 43, 10213–10226.

Hay, R.T. (2005). SUMO: a history of modification. Molecular cell 18, 1–12.

Hendriks, I.A., D’Souza, R.C., Yang, B., Verlaan-de Vries, M., Mann, M., and Vertegaal, A.C. (2014). Uncovering global SUMOylation signaling networks in a site-specific manner. Nature structural & molecular biology 21, 927–936.

Hietakangas, V., Anckar, J., Blomster, H.A., Fujimoto, M., Palvimo, J.J., Nakai, A., and Sistonen, L. (2006). PDSM, a motif for phosphorylation-dependent SUMO modification. Proceedings of the National Academy of Sciences of the United States of America 103, 45–50.

Hons, M.T., Huis In ’t Veld, P.J., Kaesler, J., Rombaut, P., Schleiffer, A., Herzog, F., Stark, H., and Peters, J.M. (2016). Topology and structure of an engineered human cohesin complex bound to Pds5B. Nature communications 7, 12523.

Hooker, G.W., and Roeder, G.S. (2006). A Role for SUMO in meiotic chromosome synapsis. Current biology : CB 16, 1238–1243.

Humphryes, N., Leung, W.K., Argunhan, B., Terentyev, Y., Dvorackova, M., and Tsubouchi, H. (2013). The Ecm11-Gmc2 complex promotes synaptonemal complex formation through assembly of transverse filaments in budding yeast. PLoS genetics 9, e1003194.

Hunter, N. (2006). Meiotic Recombination. In Molecular Genetics of Recombination, A. Aguilera, and R. Rothstein, eds. (Heidelberg: Springer-Verlag), pp. 381–442.

Hunter, N. (2015). Meiotic Recombination: The Essence of Heredity. Cold Spring Harbor perspectives in biology 7.

Hyland, E.M., Cosgrove, M.S., Molina, H., Wang, D., Pandey, A., Cottee, R.J., and Boeke, J.D. (2005). Insights into the role of histone H3 and histone H4 core modifiable residues in Saccharomyces cerevisiae. Molecular and cellular biology 25, 10060–10070.

Jentsch, S., and Psakhye, I. (2013). Control of nuclear activities by substrate-selective and protein-group SUMOylation. Annual review of genetics 47, 167–186.

Jessberger, R. (2002). The many functions of SMC proteins in chromosome dynamics. Nature reviews. Molecular cell biology 3, 767–778.

Jin, L., and Neiman, A.M. (2016). Post-transcriptional regulation in budding yeast meiosis. Curr Genet 62, 313–315.

Johnson, E.S. (2004). Protein modification by SUMO. Annual review of biochemistry 73, 355–382.

Johnson, E.S., and Gupta, A.A. (2001). An E3-like factor that promotes SUMO conjugation to the yeast septins. Cell 106, 735–744.

Klar, A.J., and Halvorson, H.O. (1975). Proteinase activities of Saccharomyces cerevisiae during sporulation. J Bacteriol 124, 863–869.

Klein, F., Laroche, T., Cardenas, M.E., Hofmann, J.F., Schweizer, D., and Gasser, S.M. (1992). Localization of RAP1 and topoisomerase II in nuclei and meiotic chromosomes of yeast. The Journal of cell biology 117, 935–948.

Klug, H., Xaver, M., Chaugule, V.K., Koidl, S., Mittler, G., Klein, F., and Pichler, A. (2013). Ubc9 sumoylation controls SUMO chain formation and meiotic synapsis in Saccharomyces cerevisiae. Molecular cell 50, 625–636.

Knipscheer, P., Flotho, A., Klug, H., Olsen, J.V., van Dijk, W.J., Fish, A., Johnson, E.S., Mann, M., Sixma, T.K., and Pichler, A. (2008). Ubc9 sumoylation regulates SUMO target discrimination. Molecular cell 31, 371–382.

Kramarz, K., Mucha, S., Litwin, I., Barg-Wojas, A., Wysocki, R., and Dziadkowiec, D. (2017). DNA Damage Tolerance Pathway Choice Through Uls1 Modulation of Srs2 SUMOylation in Saccharomyces cerevisiae. Genetics 206, 513–525.

Kschonsak, M., Merkel, F., Bisht, S., Metz, J., Rybin, V., Hassler, M., and Haering, C.H. (2017). Structural Basis for a Safety-Belt Mechanism That Anchors Condensin to Chromosomes. Cell 171, 588–600 e524.

Kulemzina, I., Ang, K., Zhao, X., Teh, J.T., Verma, V., Suranthran, S., Chavda, A.P., Huber, R.G., Eisenhaber, B., Eisenhaber, F., et al. (2016). A Reversible Association between Smc Coiled Coils Is Regulated by Lysine Acetylation and Is Required for Cohesin Association with the DNA. Molecular cell 63, 1044–1054.

Kunz, K., Piller, T., and Muller, S. (2018). SUMO-specific proteases and isopeptidases of the SENP family at a glance. Journal of cell science 131.

Lake, C.M., and Hawley, R.S. (2013). RNF212 marks the spot. Nature genetics 45, 228–229.

Lam, I., and Keeney, S. (2014). Mechanism and regulation of meiotic recombination initiation. Cold Spring Harbor perspectives in biology 7, a016634.

Lao, J.P., Cloud, V., Huang, C.C., Grubb, J., Thacker, D., Lee, C.Y., Dresser, M.E., Hunter, N., and Bishop, D.K. (2013). Meiotic crossover control by concerted action of Rad51-Dmc1 in homolog template bias and robust homeostatic regulation. PLoS genetics 9, e1003978.

Lao, J.P., Oh, S.D., Shinohara, M., Shinohara, A., and Hunter, N. (2008). Rad52 promotes postinvasion steps of meiotic double-strand-break repair. Molecular cell 29, 517–524.

Le, S., Davis, C., Konopka, J.B., and Sternglanz, R. (1997). Two new S-phase-specific genes from Saccharomyces cerevisiae. Yeast 13, 1029–1042.

Lescasse, R., Pobiega, S., Callebaut, I., and Marcand, S. (2013). End-joining inhibition at telomeres requires the translocase and polySUMO-dependent ubiquitin ligase Uls1. The EMBO journal 32, 805–815.

Leung, W.K., Humphryes, N., Afshar, N., Argunhan, B., Terentyev, Y., Tsubouchi, T., and Tsubouchi, H. (2015). The synaptonemal complex is assembled by a polySUMOylation-driven feedback mechanism in yeast. The Journal of cell biology 211, 785–793.

Levy, A., Eyal, M., Hershkovits, G., Salmon-Divon, M., Klutstein, M., and Katcoff, D.J. (2008). Yeast linker histone Hho1p is required for efficient RNA polymerase I processivity and transcriptional silencing at the ribosomal DNA. Proceedings of the National Academy of Sciences of the United States of America 105, 11703–11708.

Lewis, A., Felberbaum, R., and Hochstrasser, M. (2007). A nuclear envelope protein linking nuclear pore basket assembly, SUMO protease regulation, and mRNA surveillance. The Journal of cell biology 178, 813–827.

Li, S.J., and Hochstrasser, M. (1999). A new protease required for cell-cycle progression in yeast. Nature 398, 246–251.

Li, S.J., and Hochstrasser, M. (2003). The Ulp1 SUMO isopeptidase: distinct domains required for viability, nuclear envelope localization, and substrate specificity. The Journal of cell biology 160, 1069–1081.

Liang, J., Singh, N., Carlson, C.R., Albuquerque, C.P., Corbett, K.D., and Zhou, H.L. (2017). Recruitment of a SUMO isopeptidase to rDNA stabilizes silencing complexes by opposing SUMO targeted ubiquitin ligase activity. Genes & development 31, 802–815.

Liebelt, F., and Vertegaal, A.C. (2016). Ubiquitin-dependent and independent roles of SUMO in proteostasis. Am J Physiol Cell Physiol 311, C284–296.

Lilienthal, I., Kanno, T., and Sjogren, C. (2013). Inhibition of the Smc5/6 complex during meiosis perturbs joint molecule formation and resolution without significantly changing crossover or non-crossover levels. PLoS genetics 9, e1003898.

Lin, F.M., Lai, Y.J., Shen, H.J., Cheng, Y.H., and Wang, T.F. (2010). Yeast axial-element protein, Red1, binds SUMO chains to promote meiotic interhomologue recombination and chromosome synapsis. The EMBO journal 29, 586–596.

Lin, K.W., McDonald, K.R., Guise, A.J., Chan, A., Cristea, I.M., and Zakian, V.A. (2015). Proteomics of yeast telomerase identified Cdc48-Npl4-Ufd1 and Ufd4 as regulators of Est1 and telomere length. Nature communications 6, 8290.

Matangkasombut, O., and Buratowski, S. (2003). Different sensitivities of bromodomain factors 1 and 2 to histone H4 acetylation. Molecular cell 11, 353–363.

Matsuhara, H., and Yamamoto, A. (2016). Autophagy is required for efficient meiosis progression and proper meiotic chromosome segregation in fission yeast. Genes to cells : devoted to molecular & cellular mechanisms 21, 65–87.

McAleenan, A., Cordon-Preciado, V., Clemente-Blanco, A., Liu, I.C., Sen, N., Leonard, J., Jarmuz, A., and Aragon, L. (2012). SUMOylation of the alpha-kleisin subunit of cohesin is required for DNA damage-induced cohesion. Current biology : CB 22, 1564–1575.

McNicoll, F., Stevense, M., and Jessberger, R. (2013). Cohesin in gametogenesis. Curr Top Dev Biol 102, 1–34.

Mieczkowski, P.A., Dominska, M., Buck, M.J., Lieb, J.D., and Petes, T.D. (2007). Loss of a histone deacetylase dramatically alters the genomic distribution of Spo11p-catalyzed DNA breaks in Saccharomyces cerevisiae. Proceedings of the National Academy of Sciences of the United States of America 104, 3955–3960.

Millan-Arino, L., Izquierdo-Bouldstridge, A., and Jordan, A. (2016). Specificities and genomic distribution of somatic mammalian histone H1 subtypes. Biochimica et biophysica acta 1859, 510–519.

Moens, P.B., and Earnshaw, W.C. (1989). Anti-topoisomerase II recognizes meiotic chromosome cores. Chromosoma 98, 317–322.

Mohideen, F., Capili, A.D., Bilimoria, P.M., Yamada, T., Bonni, A., and Lima, C.D. (2009). A molecular basis for phosphorylation-dependent SUMO conjugation by the E2 UBC9. Nature structural & molecular biology 16, 945–952.

Morrison, A.J., Highland, J., Krogan, N.J., Arbel-Eden, A., Greenblatt, J.F., Haber, J.E., and Shen, X. (2004). INO80 and gamma-H2AX interaction links ATP-dependent chromatin remodeling to DNA damage repair. Cell 119, 767–775.

Mossessova, E., and Lima, C.D. (2000). Ulp1-SUMO crystal structure and genetic analysis reveal conserved interactions and a regulatory element essential for cell growth in yeast. Molecular cell 5, 865–876.

Najor, N.A., Weatherford, L., and Brush, G.S. (2016). Prevention of DNA Rereplication Through a Meiotic Recombination Checkpoint Response. G3 6, 3869–3881.

Nathan, D., Ingvarsdottir, K., Sterner, D.E., Bylebyl, G.R., Dokmanovic, M., Dorsey, J.A., Whelan, K.A., Krsmanovic, M., Lane, W.S., Meluh, P.B., et al. (2006). Histone sumoylation is a negative regulator in Saccharomyces cerevisiae and shows dynamic interplay with positive-acting histone modifications. Genes & development 20, 966–976.

Ng, C.H., Akhter, A., Yurko, N., Burgener, J.M., Rosonina, E., and Manley, J.L. (2015). Sumoylation controls the timing of Tup1-mediated transcriptional deactivation. Nature communications 6, 6610.

Nie, M., Aslanian, A., Prudden, J., Heideker, J., Vashisht, A.A., Wohlschlegel, J.A., Yates, J.R., 3rd, and Boddy, M.N. (2012). Dual recruitment of Cdc48 (p97)-Ufd1-Npl4 ubiquitin-selective segregase by small ubiquitin-like modifier protein (SUMO) and ubiquitin in SUMO-targeted ubiquitin ligase-mediated genome stability functions. The Journal of biological chemistry 287, 29610–29619.

Nottke, A.C., Kim, H.M., and Colaiacovo, M.P. (2017). Wrestling with Chromosomes: The Roles of SUMO During Meiosis. Adv Exp Med Biol 963, 185–196.

Olsen, S.K., Capili, A.D., Lu, X., Tan, D.S., and Lima, C.D. (2010). Active site remodelling accompanies thioester bond formation in the SUMO E1. Nature 463, 906–912.

Ontoso, D., Acosta, I., van Leeuwen, F., Freire, R., and San-Segundo, P.A. (2013). Dot1-dependent histone H3K79 methylation promotes activation of the Mek1 meiotic checkpoint effector kinase by regulating the Hop1 adaptor. PLoS genetics 9, e1003262.

Otto, G.M., and Brar, G.A. (2018). Seq-ing answers: uncovering the unexpected in global gene regulation. Curr Genet.

Ouyang, Z., and Yu, H. (2017). Releasing the cohesin ring: A rigid scaffold model for opening the DNA exit gate by Pds5 and Wapl. BioEssays : news and reviews in molecular, cellular and developmental biology 39.

Palancade, B., Liu, X., Garcia-Rubio, M., Aguilera, A., Zhao, X., and Doye, V. (2007). Nucleoporins prevent DNA damage accumulation by modulating Ulp1-dependent sumoylation processes. Molecular biology of the cell 18, 2912–2923.

Pan, J., Sasaki, M., Kniewel, R., Murakami, H., Blitzblau, H.G., Tischfield, S.E., Zhu, X., Neale, M.J., Jasin, M., Socci, N.D., et al. (2011). A hierarchical combination of factors shapes the genome-wide topography of yeast meiotic recombination initiation. Cell 144, 719–731.

Panizza, S., Mendoza, M.A., Berlinger, M., Huang, L., Nicolas, A., Shirahige, K., and Klein, F. (2011). Spo11-accessory proteins link double-strand break sites to the chromosome axis in early meiotic recombination. Cell 146, 372–383.

Panse, V.G., Kressler, D., Pauli, A., Petfalski, E., Gnadig, M., Tollervey, D., and Hurt, E. (2006). Formation and nuclear export of preribosomes are functionally linked to the small-ubiquitin-related modifier pathway. Traffic 7, 1311–1321.

Panse, V.G., Kuster, B., Gerstberger, T., and Hurt, E. (2003). Unconventional tethering of Ulp1 to the transport channel of the nuclear pore complex by karyopherins. Nature cell biology 5, 21–27.

Papouli, E., Chen, S., Davies, A.A., Huttner, D., Krejci, L., Sung, P., and Ulrich, H.D. (2005). Crosstalk between SUMO and ubiquitin on PCNA is mediated by recruitment of the helicase Srs2p. Molecular cell 19, 123–133.

Pelisch, F., Tammsalu, T., Wang, B., Jaffray, E.G., Gartner, A., and Hay, R.T. (2017). A SUMO-Dependent Protein Network Regulates Chromosome Congression during Oocyte Meiosis. Molecular cell 65, 66–77.

Pfander, B., Moldovan, G.L., Sacher, M., Hoege, C., and Jentsch, S. (2005). SUMO-modified PCNA recruits Srs2 to prevent recombination during S phase. Nature 436, 428–433.

Psakhye, I., and Jentsch, S. (2012). Protein group modification and synergy in the SUMO pathway as exemplified in DNA repair. Cell 151, 807–820.

Rabitsch, K.P., Petronczki, M., Javerzat, J.P., Genier, S., Chwalla, B., Schleiffer, A., Tanaka, T.U., and Nasmyth, K. (2003). Kinetochore recruitment of two nucleolar proteins is required for homolog segregation in meiosis I. Developmental cell 4, 535–548.

Rao, H.B., Qiao, H., Bhatt, S.K., Bailey, L.R., Tran, H.D., Bourne, S.L., Qiu, W., Deshpande, A., Sharma, A.N., Beebout, C.J., et al. (2017). A SUMO-ubiquitin relay recruits proteasomes to chromosome axes to regulate meiotic recombination. Science 355, 403–407.

Redon, C., Pilch, D.R., Rogakou, E.P., Orr, A.H., Lowndes, N.F., and Bonner, W.M. (2003). Yeast histone 2A serine 129 is essential for the efficient repair of checkpoint-blind DNA damage. EMBO reports 4, 678–684.

Rodriguez, A., and Pangas, S.A. (2015). Regulation of germ cell function by SUMOylation. Cell Tissue Res.

Rodriguez, M.S., Dargemont, C., and Hay, R.T. (2001). SUMO-1 conjugation in vivo requires both a consensus modification motif and nuclear targeting. The Journal of biological chemistry 276, 12654–12659.

Rosonina, E., Duncan, S.M., and Manley, J.L. (2012). Sumoylation of transcription factor Gcn4 facilitates its Srb10-mediated clearance from promoters in yeast. Genes & development 26, 350–355.

Roth, S.Y., and Allis, C.D. (1992). Chromatin condensation: does histone H1 dephosphorylation play a role? Trends in biochemical sciences 17, 93–98.

Sacher, M., Pfander, B., Hoege, C., and Jentsch, S. (2006). Control of Rad52 recombination activity by double-strand break-induced SUMO modification. Nature cell biology 8, 1284–1290.

Sakaguchi, K., Koshiyama, A., and Iwabata, K. (2007). Meiosis and small ubiquitin-related modifier (SUMO)-conjugating enzyme, Ubc9. Febs J 274, 3519–3531.

Sampson, D.A., Wang, M., and Matunis, M.J. (2001). The small ubiquitin-like modifier-1 (SUMO-1) consensus sequence mediates Ubc9 binding and is essential for SUMO-1 modification. The Journal of biological chemistry 276, 21664–21669.

San-Segundo, P.A., and Roeder, G.S. (2000). Role for the silencing protein Dot1 in meiotic checkpoint control. Molecular biology of the cell 11, 3601–3615.

Sarangi, P., Steinacher, R., Altmannova, V., Fu, Q., Paull, T.T., Krejci, L., Whitby, M.C., and Zhao, X. (2015). Sumoylation influences DNA break repair partly by increasing the solubility of a conserved end resection protein. PLoS genetics 11, e1004899.

Sarg, B., Helliger, W., Talasz, H., Forg, B., and Lindner, H.H. (2006). Histone H1 phosphorylation occurs site-specifically during interphase and mitosis: identification of a novel phosphorylation site on histone H1. The Journal of biological chemistry 281, 6573–6580.

Schafer, G., McEvoy, C.R., and Patterton, H.G. (2008). The Saccharomyces cerevisiae linker histone Hho1p is essential for chromatin compaction in stationary phase and is displaced by transcription. Proceedings of the National Academy of Sciences of the United States of America 105, 14838–14843.

Seeber, A., Hauer, M., and Gasser, S.M. (2013). Nucleosome remodelers in double-strand break repair. Current opinion in genetics & development 23, 174–184.

Serrentino, M.E., Chaplais, E., Sommermeyer, V., and Borde, V. (2013). Differential association of the conserved SUMO ligase Zip3 with meiotic double-strand break sites reveals regional variations in the outcome of meiotic recombination. PLoS genetics 9, e1003416.

Shiio, Y., and Eisenman, R.N. (2003). Histone sumoylation is associated with transcriptional repression. Proceedings of the National Academy of Sciences of the United States of America 100, 13225–13230.

Shinohara, M., Oh, S.D., Hunter, N., and Shinohara, A. (2008). Crossover assurance and crossover interference are distinctly regulated by the ZMM proteins during yeast meiosis. Nature genetics 40, 299–309.

Smith, A.V., and Roeder, G.S. (1997). The yeast Red1 protein localizes to the cores of meiotic chromosomes. The Journal of cell biology 136, 957–967.

Sollier, J., Lin, W., Soustelle, C., Suhre, K., Nicolas, A., Geli, V., and de La Roche Saint-Andre, C. (2004). Set1 is required for meiotic S-phase onset, double-strand break formation and middle gene expression. The EMBO journal 23, 1957–1967.

Sommermeyer, V., Beneut, C., Chaplais, E., Serrentino, M.E., and Borde, V. (2013). Spp1, a member of the Set1 Complex, promotes meiotic DSB formation in promoters by tethering histone H3K4 methylation sites to chromosome axes. Molecular cell 49, 43–54.

Sourirajan, A., and Lichten, M. (2008). Polo-like kinase Cdc5 drives exit from pachytene during budding yeast meiosis. Genes & development 22, 2627–2632.

Srikumar, T., Lewicki, M.C., Costanzo, M., Tkach, J.M., van Bakel, H., Tsui, K., Johnson, E.S., Brown, G.W., Andrews, B.J., Boone, C., et al. (2013a). Global analysis of SUMO chain function reveals multiple roles in chromatin regulation. The Journal of cell biology 201, 145–163.

Srikumar, T., Lewicki, M.C., and Raught, B. (2013b). A global S. cerevisiae small ubiquitin-related modifier (SUMO) system interactome. Mol Syst Biol 9, 668.

Sriramachandran, A.M., and Dohmen, R.J. (2014). SUMO-targeted ubiquitin ligases. Biochimica et biophysica acta 1843, 75–85.

Stade, K., Vogel, F., Schwienhorst, I., Meusser, B., Volkwein, C., Nentwig, B., Dohmen, R.J., and Sommer, T. (2002). A lack of SUMO conjugation affects cNLS-dependent nuclear protein import in yeast. The Journal of biological chemistry 277, 49554–49561.

Stead, K., Aguilar, C., Hartman, T., Drexel, M., Meluh, P., and Guacci, V. (2003). Pds5p regulates the maintenance of sister chromatid cohesion and is sumoylated to promote the dissolution of cohesion. The Journal of cell biology 163, 729–741.

Steinacher, R., and Schar, P. (2005). Functionality of human thymine DNA glycosylase requires SUMO-regulated changes in protein conformation. Current biology : CB 15, 616–623.

Storey, A.J., Wang, H.P., Protacio, R.U., Davidson, M.K., Tackett, A.J., and Wahls, W.P. (2018). Chromatin-mediated regulators of meiotic recombination revealed by proteomics of a recombination hotspot. Epigenetics Chromatin 11, 64.

Streich, F.C., Jr., and Lima, C.D. (2016). Capturing a substrate in an activated RING E3/E2-SUMO complex. Nature 536, 304–308.

Subramanian, V.V., and Hochwagen, A. (2014). The meiotic checkpoint network: step-by-step through meiotic prophase. Cold Spring Harbor perspectives in biology 6, a016675.

Suka, N., Suka, Y., Carmen, A.A., Wu, J., and Grunstein, M. (2001). Highly specific antibodies determine histone acetylation site usage in yeast heterochromatin and euchromatin. Molecular cell 8, 473–479.

Sun, X., Huang, L., Markowitz, T.E., Blitzblau, H.G., Chen, D., Klein, F., and Hochwagen, A. (2015). Transcription dynamically patterns the meiotic chromosome-axis interface. eLife 4.

Takahashi, D., Orihara, Y., Kitagawa, S., Kusakabe, M., Shintani, T., Oma, Y., and Harata, M. (2017). Quantitative regulation of histone variant H2A.Z during cell cycle by ubiquitin proteasome system and SUMO-targeted ubiquitin ligases. Biosci Biotechnol Biochem 81, 1557–1560.

Tan, W., Wang, Z., and Prelich, G. (2013). Physical and Genetic Interactions Between Uls1 and the Slx5-Slx8 SUMO-Targeted Ubiquitin Ligase. G3 3, 771–780.

Texari, L., Dieppois, G., Vinciguerra, P., Contreras, M.P., Groner, A., Letourneau, A., and Stutz, F. (2013). The nuclear pore regulates GAL1 gene transcription by controlling the localization of the SUMO protease Ulp1. Molecular cell 51, 807–818.

Thorslund, T., Ripplinger, A., Hoffmann, S., Wild, T., Uckelmann, M., Villumsen, B., Narita, T., Sixma, T.K., Choudhary, C., Bekker-Jensen, S., et al. (2015). Histone H1 couples initiation and amplification of ubiquitin signalling after DNA damage. Nature 527, 389–393.

Tresenrider, A., and Unal, E. (2018). One-two punch mechanism of gene repression: a fresh perspective on gene regulation. Curr Genet 64, 581–588.

Tyanova, S., Temu, T., Sinitcyn, P., Carlson, A., Hein, M.Y., Geiger, T., Mann, M., and Cox, J. (2016). The Perseus computational platform for comprehensive analysis of (prote)omics data. Nature methods 13, 731–740.

Uhlmann, F. (2016). SMC complexes: from DNA to chromosomes. Nature reviews. Molecular cell biology 17, 399–412.

Uzunova, K., Gottsche, K., Miteva, M., Weisshaar, S.R., Glanemann, C., Schnellhardt, M., Niessen, M., Scheel, H., Hofmann, K., Johnson, E.S., et al. (2007). Ubiquitin-dependent proteolytic control of SUMO conjugates. The Journal of biological chemistry.

Vader, G., Blitzblau, H.G., Tame, M.A., Falk, J.E., Curtin, L., and Hochwagen, A. (2011). Protection of repetitive DNA borders from self-induced meiotic instability. Nature 477, 115–U141.

van Attikum, H., Fritsch, O., Hohn, B., and Gasser, S.M. (2004). Recruitment of the INO80 complex by H2A phosphorylation links ATP-dependent chromatin remodeling with DNA double-strand break repair. Cell 119, 777–788.

Varshavsky, A. (1997). The N-end rule pathway of protein degradation. Genes to cells : devoted to molecular & cellular mechanisms 2, 13–28.

Vit, O., and Petrak, J. (2017). Integral membrane proteins in proteomics. How to break open the black box? J Proteomics 153, 8–20.

Voelkel-Meiman, K., Cheng, S.Y., Morehouse, S.J., and MacQueen, A.J. (2016). Synaptonemal Complex Proteins of Budding Yeast Define Reciprocal Roles in MutSgamma-Mediated Crossover Formation. Genetics 203, 1091–1103.

Voelkel-Meiman, K., Cheng, S.Y., Parziale, M., Morehouse, S.J., Feil, A., Davies, O.R., de Muyt, A., Borde, V., and MacQueen, A.J. (2019). Crossover recombination and synapsis are linked by adjacent regions within the N terminus of the Zip1 synaptonemal complex protein. PLoS genetics 15, e1008201.

Voelkel-Meiman, K., Taylor, L.F., Mukherjee, P., Humphryes, N., Tsubouchi, H., and Macqueen, (2013). SUMO localizes to the central element of synaptonemal complex and is required for the full synapsis of meiotic chromosomes in budding yeast. PLoS genetics 9, e1003837.

von Wettstein, D., Rasmussen, S.W., and Holm, P.B. (1984). The synaptonemal complex in genetic segregation. Annu Rev Genet 18, 331–413.

Vujin, A., and Zetka, M. (2017). The proteasome enters the meiotic prophase fray. BioEssays : news and reviews in molecular, cellular and developmental biology.

Wan, L., Zhang, C., Shokat, K.M., and Hollingsworth, N.M. (2006). Chemical inactivation of cdc7 kinase in budding yeast results in a reversible arrest that allows efficient cell synchronization prior to meiotic recombination. Genetics 174, 1767–1774.

Warren, J.J., Pohlhaus, T.J., Changela, A., Iyer, R.R., Modrich, P.L., and Beese, L.S. (2007). Structure of the human MutSalpha DNA lesion recognition complex. Molecular cell 26, 579–592.

Watanabe, Y. (2012). Geometry and force behind kinetochore orientation: lessons from meiosis. Nature reviews. Molecular cell biology 13, 370–382.

Watts, F.Z., and Hoffmann, E. (2011). SUMO meets meiosis: an encounter at the synaptonemal complex: SUMO chains and sumoylated proteins suggest that heterogeneous and complex interactions lie at the centre of the synaptonemal complex. BioEssays : news and reviews in molecular, cellular and developmental biology 33, 529–537.

Wei, B., Huang, C., Liu, B., Wang, Y., Xia, N., Fan, Q., Chen, G.Q., and Cheng, J. (2018). Mitotic Phosphorylation of SENP3 Regulates DeSUMOylation of Chromosome-Associated Proteins and Chromosome Stability. Cancer research 78, 2171–2178.

Wei, Y., Diao, L.X., Lu, S., Wang, H.T., Suo, F., Dong, M.Q., and Du, L.L. (2017). SUMO-Targeted DNA Translocase Rrp2 Protects the Genome from Top2-Induced DNA Damage. Molecular cell 66, 581–596 e586.

Wen, F.P., Guo, Y.S., Hu, Y., Liu, W.X., Wang, Q., Wang, Y.T., Yu, H.Y., Tang, C.M., Yang, J., Zhou, T., et al. (2016). Distinct temporal requirements for autophagy and the proteasome in yeast meiosis. Autophagy 12, 671–688.

Wen, X., and Klionsky, D.J. (2016). An overview of macroautophagy in yeast. Journal of molecular biology 428, 1681–1699.

West, A.M., Rosenberg, S.C., Ur, S.N., Lehmer, M.K., Ye, Q., Hagemann, G., Caballero, I., Uson, I., MacQueen, A.J., Herzog, F., et al. (2019). A conserved filamentous assembly underlies the structure of the meiotic chromosome axis. eLife 8.

West, A.M.V., Komives, E.A., and Corbett, K.D. (2018). Conformational dynamics of the Hop1 HORMA domain reveal a common mechanism with the spindle checkpoint protein Mad2. Nucleic acids research 46, 279–292.

Wild, P., Susperregui, A., Piazza, I., Dorig, C., Oke, A., Arter, M., Yamaguchi, M., Hilditch, A.T., Vuina, K., Chan, K.C., et al. (2019). Network Rewiring of Homologous Recombination Enzymes during Mitotic Proliferation and Meiosis. Molecular cell.

Winter, E. (2012). The Sum1/Ndt80 transcriptional switch and commitment to meiosis in Saccharomyces cerevisiae. Microbiology and molecular biology reviews : MMBR 76, 1–15.

Wohlschlegel, J.A., Johnson, E.S., Reed, S.I., and Yates, J.R., 3rd (2006). Improved identification of SUMO attachment sites using C-terminal SUMO mutants and tailored protease digestion strategies. J Proteome Res 5, 761–770.

Woltering, D., Baumgartner, B., Bagchi, S., Larkin, B., Loidl, J., de los Santos, T., and Hollingsworth, N.M. (2000). Meiotic segregation, synapsis, and recombination checkpoint functions require physical interaction between the chromosomal proteins Red1p and Hop1p. Molecular and cellular biology 20, 6646–6658.

Wu, N., Kong, X., Ji, Z., Zeng, W., Potts, P.R., Yokomori, K., and Yu, H. (2012). Scc1 sumoylation by Mms21 promotes sister chromatid recombination through counteracting Wapl. Genes & development 26, 1473–1485.

Xaver, M., Huang, L., Chen, D., and Klein, F. (2013). Smc5/6-mms21 prevents and eliminates inappropriate recombination intermediates in meiosis. PLoS genetics 9, e1004067.

Xu, G., Paige, J.S., and Jaffrey, S.R. (2010). Global analysis of lysine ubiquitination by ubiquitin remnant immunoaffinity profiling. Nat Biotechnol 28, 868–873.

Yamashita, K., Shinohara, M., and Shinohara, A. (2004). Rad6-Bre1-mediated histone H2B ubiquitylation modulates the formation of double-strand breaks during meiosis. Proc Natl Acad Sci U S A 101, 11380–11385.

Yu, H.G., and Koshland, D.E. (2003). Meiotic condensin is required for proper chromosome compaction, SC assembly, and resolution of recombination-dependent chromosome linkages. The Journal of cell biology 163, 937–947.

Zakharyevich, K., Ma, Y., Tang, S., Hwang, P.Y., Boiteux, S., and Hunter, N. (2010). Temporally and biochemically distinct activities of Exo1 during meiosis: double-strand break resection and resolution of double Holliday junctions. Molecular cell 40, 1001–1015.

Zakharyevich, K., Tang, S., Ma, Y., and Hunter, N. (2012). Delineation of joint molecule resolution pathways in meiosis identifies a crossover-specific resolvase. Cell 149, 334–347.

Zavec, A.B., Comino, A., Lenassi, M., and Komel, R. (2007). Ecm11 protein of yeast Saccharomyces cerevisiae is regulated by sumoylation during meiosis. FEMS Yeast Res.

Zavec, A.B., Comino, A., Lenassi, M., and Komel, R. (2008). Ecm11 protein of yeast Saccharomyces cerevisiae is regulated by sumoylation during meiosis. FEMS Yeast Res 8, 64–70.

Zhang, L., Wang, S., Yin, S., Hong, S., Kim, K.P., and Kleckner, N. (2014). Topoisomerase II mediates meiotic crossover interference. Nature 511, 551–556.

Zhang, X.P., Lee, K.I., Solinger, J.A., Kiianitsa, K., and Heyer, W.D. (2005). Gly-103 in the N-terminal domain of Saccharomyces cerevisiae Rad51 protein is critical for DNA binding. The Journal of biological chemistry 280, 26303–26311.

Zhao, Q., Xie, Y., Zheng, Y., Jiang, S., Liu, W., Mu, W., Liu, Z., Zhao, Y., Xue, Y., and Ren, J. (2014). GPS-SUMO: a tool for the prediction of sumoylation sites and SUMO-interaction motifs. Nucleic acids research 42, W325–330.

Zhao, X. (2018). SUMO-Mediated Regulation of Nuclear Functions and Signaling Processes. Molecular cell 71, 409–418.

Zhao, X., Wu, C.Y., and Blobel, G. (2004). Mlp-dependent anchorage and stabilization of a desumoylating enzyme is required to prevent clonal lethality. The Journal of cell biology 167, 605–611.

Zickler, D., and Kleckner, N. (1999). Meiotic chromosomes: integrating structure and function. Annual review of genetics 33, 603–754.

Zickler, D., and Kleckner, N. (2015). Recombination, pairing, and synapsis of homologs during meiosis. Cold Spring Harbor perspectives in biology 7, pii: a016626. doi: 016610.011101/cshperspect.a016626.

