## Supplemental Information for "Delineation of the SUMO-Modified Proteome Reveals Regulatory Functions Throughout Meiosis"

**This file includes:**

Methods

Supplemental Figures 1-7

Supplemental Tables 1 and 2

### METHODS

### EXPERIMENTAL MODEL

#### Yeast strains and plasmids for SUMO proteomics:

The yeast SUMO gene, *SMT3* was N-terminally tagged as described previously (Booher and Kaiser, 2008), with some modifications. Plasmid pFA6a-kanMX6-$P_{GAL1}$-HBH (a gift from Dr. Peter Kaiser) was modified by replacing the HBH tag module with a 6His-Strep-tagII module. A PCR product containing *kanMX6*-$P_{GAL1}$-*6His-StrepII*, with 70bp homologies to the 5’ region of *SMT3* was used to transform haploid *GAL*+ yeast. Transformants were selected on YP-Gal + Geneticin (G418) solid media and genotyped by PCR. To restore the native *SMT3* promoter, these strains were transformed with $P_{SMT3}$ and selected for growth on YPD. Strains were then crossed to introduce $Cu^{2+}$-inducible versions of *IME1* and *IME4* (a gift from Folkert van Werven and Angelika Amon) and an estradiol-inducible *NDT80* gene (Benjamin et al., 2003a), which allowed for inducible, highly synchronous meiotic entry and pachytene exit, respectively. To generate the I96K mutation in Smt3, strains were transformed with a PCR product generated from *pFA6a-3HA-kanMX6* (Longtine et al., 1998) with PCR primers Smt3 KGG FW (*AAGATTTGGACATGGAGGATAACGATATTATTGAGGCTCACAGAGAACAG*___**AAA**___*GGTGGTGCTACGTATTAGCGGATCCCCGGGTTAATTAA*) and R1 Smt3 (*CTATGGTATTTTTTGGTGGGTGGAGGGAAGGGAGAGGTTTGTTGGCGTTCTTTAGGCATTGTTAAGAGTCGAATTCGAGCTCGTTTAAAC*). Transformants were screened by PCR genotyping and validated by sequencing. Integrity of the *HIS4::LEU2* hotspot was validated by Southern blotting. Strains of opposite mating types were mated to generate the diploid strain used in the proteomic experiments (Strain table, below)

#### Yeast strains for DNA physical assays

Full genotypes are shown in the strain table. The Auxin-Induced Degron (AID) system designed for use during meiosis has been described (Tang et al., 2015). Minimal AID fusions to Aos1 and Uba2 were constructed as described using plasmid p7aid-9m as a template for PCR-mediated allele replacement (Morawska and Ulrich, 2013; Tang et al., 2015). The estradiol-inducible *GAL4-ER IN-NDT80* system has been described (Benjamin et al., 2003b; Carlile and Amon, 2008; Louvion et al., 1993; Tang et al., 2015).

### METHOD DETAILS

#### Meiotic time courses for proteomic analysis

Synchronous meiotic time courses were performed as described (Owens et al., 2018) with the following modifications. Saturated 5 mL overnight YPD cultures were diluted 1:1000 into six 300 ml SPS in 2 L flasks and shaken overnight at 325 rpm. At 18 hours, cells were collected by centrifugation, washed with sterilize water and divided into twelve 2.8L baffled flasks, each containing 350 mL SPM. These cultures were then shaken at 350 rpm for two hours before adding $CuSO_4$ to a final concentration of 25 μM. Cells were collected at 0 (G0), 1.5 (premeiotic S-phase, S), 2.5 (double-strand breaks, DSB), 3.5 (strand invasion and synapsis, SI), and 5 hours after $CuSO_4$ addition (double Holliday junction formation and pachytene, dHJ). At the 5-hour time point, estradiol was added to a final concentration of 1 μM and a final sample was collected at 6 hours (crossing over and SC disassembly, CO). For each timepoint, cells from all twelve flasks were pooled, pelleted by centrifugation, washed with milliQ water, pelleted again and then flash frozen in liquid nitrogen. Frozen pellets were stored at -80℃ until all the samples had been collected, and then processed in parallel. For all quantitative analysis of temporal changes in SUMOylation, a triplicate set of time courses was collected and processed in parallel to minimize variation.

#### Cell lysis and protein purification for proteomics

Each 20-25 g cell pellet was resuspended in 60 mL fresh ice-cold 0.25 M NaOH, 1% β-mercaptoethanol lysis solution and incubated on ice for 20 minutes. Proteins were precipitated with 10 mL 100% trichloroacetic acid on ice for 20 minutes and pelleted by centrifugation at 5000 x g for 15 minutes. The pellet was broken up, thoroughly washed twice with 5 ml ice-cold acetone and air dried. Proteins were dissolved in 6 M Guanidine buffer (6 M Guanidine, 100 mM Tris pH 8.0, 500 mM NaCl, 10 mM imidazole pH 8.0) by vigorous shaking at 30˚C. Lysates were clarified by centrifugation at 10,000 x g for 30 minutes and filtration through a Whatman filter paper. A 1 mL HisTrap FF column mounted on an Äkta Avant FPLC system (GE Lifesciences) was equilibrated with 5 mL of 6 M Guanidine buffer and then the cell lysate was passed through, followed by washes with 20 mL 6 M Guanidine buffer, 10 mL each of Urea buffer pH 6.3 (6 M urea, 100 mM tris pH 6.3, 500 mM NaCl) and Urea buffer pH 8.0 (6 M urea, 100 mM Ammonium Bicarbonate pH 8.0, 150 mM NaCl, 20 mM Imidazole). Bound proteins were eluted with 0.5 M Imidazole (6 M urea, 100 mM Ammonium Bicarbonate pH 8.0, 150 mM NaCl, 500 mM Imidazole pH 8.0) and collected in 2 mL fractions. Fractions covering the elution peak were pooled and final protein concentrations were determined by the Bradford assay. Proteins were reduced with 4 mM TCEP (Pierce) for 30 minutes at room temperature, alkylated with 10 mM iodoacetamide (Sigma) for 30 minutes in the dark and then quenched with 10 mM DTT. A total of 2 mg protein from each sample was transferred to a new tube, samples were diluted with elution buffer so that all samples had the same final volume and then diluted with twice that volume of 100 mM ammonium bicarbonate pH 8.0. Samples were digested with 0.4 AU Lys-C (Wako Chemicals) overnight at 37˚C on a shaker at 225-250 rpm. The next day, samples were split and one half was diluted with 100 mM ammonium bicarbonate to 0.8 M urea and digested for another 6 hours with 10 µg Glu-C. Digestion was stopped by acidification with 10% trifluoroacetic acid (TFA) to pH ≤ 3, and samples were the desalted on Sep-Pak tC18 reversed phase columns (Waters). The desalted samples were lyophilized for at least 48 hours and then dissolved in 500 µL 1X immunoaffinity purification buffer (IAP buffer, Cell Signaling

Technologies). Samples were sonicated for 30 minutes at room temperature in a water-bath sonicator and then clarified by centrifugation. Peptides containing di-glycyl lysine (K-ε-GG) residues were enriched using the UbiScan kit (Cell Signaling technologies) according to the manufacturer's instructions with some modifications. Each tube of immunoaffinity beads was equilibrated with the IAP buffer and then split evenly into six tubes. Each set of six tubes was used for one set of timepoints (i.e., one tube per time course) to reduce variation. Peptide solutions were incubated with UbiScan beads at 4˚C for 90 minutes, washed twice with chilled IAP buffer and twice with chilled HPLC grade water (Fisher). Bound peptides were then eluted twice with 55 µL 0.15% TFA for 5-10 minutes at room temperature, eluates were pooled, flash frozen and then dried by vacuum centrifugation.

**Mass spectrometry**

K-ε-GG enriched peptide mixtures were analyzed with a Q Exactive Orbitrap tandem MS system (Thermo Scientific) with an upstream in-line Proxeon Easy-nLCII HPLC system (Thermo Scientific). Peptides were resuspended in 0.2% TFA and loaded on to a 25 mm Magic C18 RPLC column and eluted over a 90-minute acetonitrile gradient at 300 nl/min. MS1 spectra were sampled with a top-20 cutoff and 5s dynamic exclusion and subjected to high-energy collision dissociation to obtain MS2 spectra. For the triplicate set that was used for quantitative analysis, all samples were run back to back on the same LC column, with the same instrument settings to minimize variation. Additionally, the run-order of samples was randomized to eliminate any effect on the final data.

**Time course conditions for SUMO execution-point analysis**

***Auxin-Induced Degradation of SUMO E1 Proteins***

As described (Tang et al., 2015), 50 μM $CuSO_4$ (diluted from 100 mM stock) was added to induce expression of *$P_{CUP1}$-OsTIR1* and 2 mM auxin (dilution from a 2 M stock of 3-indoleacetic acid, Sigma I3750, dissolved in DMSO) was added to trigger Aos1-AID or Uba2-AID degradation.

***cdc7-as3* Time Courses**

Cells were incubated in SPM as previously described (Oh et al., 2009). After 30 minutes, PP1 (Tocris Bioscience 1397, 10 mM stock dissolved in DMSO) was added for a final concentration of 3.5 μM. This concentration is sufficient to block DSB formation. After 6.5 hours in PP1, cells were released from arrest by washing nine times in an equal volume of sporulation media over a 90-minute period.

***To Assay DSB Formation Following E1 Degradation***

Cells were cultured as for *cdc7-as3* time courses. At 5.5 hours, cultures were split and at 6 hours copper (1:2000 dilution of 100 mM CuSO4 stock for a final concentration of 50 μM) and auxin (1:1000 dilution of 2 M stock for a final concentration of 2 mM) were added to one sub-culture. At 6.5 hours, cultures were washed as described above with media containing copper and auxin. At 8 hours, cultures were placed in fresh flasks and at 8.5 hours, auxin was added again (1:2000 dilution of 2 M stock). Cell samples were then collected to assay protein depletion, meiotic divisions, and recombination intermediates.

***To Assay Recombination Intermediates Following E1 Degradation***

Cells were cultured as for *cdc7-as3* time courses. After washes were completed at 8 hours, cultures were split and at 9 hours, copper and auxin were added to one subculture. At 9.5 hours, auxin was added again (1:2000 dilution of 2 M stock). Cell samples were then collected to assay protein depletion, meiotic divisions, and recombination intermediates.

#### *IN-NDT80 Time Courses*

Cells were incubated in SPM as previously described (Oh et al., 2009). At 6.5 hrs, copper was added, and at 7 hours auxin was added to one subculture, and 1 μM estradiol (Sigma E2758 in DMSO) was added to both subcultures to induce *IN-NDT80.* At 7.5 hours, auxin was added again (1:2000 dilution of 2 M stock). Cell samples were collected to assay protein depletion, meiotic divisions, and recombination intermediates.

#### *Southern Blot Analysis of the DNA Events of Meiosis*

Detailed protocols for meiotic time courses and DNA physical assays at the *HIS4::LEU2* recombination hotspot have been described (Oh et al., 2009). Error bars show averages (±SD) from three experiments.

#### *Immunblotting*

Whole cell extracts were prepared using a TCA extraction method, as described (Johnson and Blobel, 1999; Tang et al., 2015). Following SDS-PAGE and Western blotting, an anti-c-Myc mouse monoclonal antibody (Roche; 11667149001) was used to detect Aos1-AID-9myc and Uba2-AID-9Myc. Anti-Arp7 goat polyclonal antibody (Santa Cruz Biotechnology; y-C20) was used to detect Arp7 as a loading control. Donkey anti-goat (IRDye 680; LI-COR Biosciences; 926-68074) and donkey anti-mouse (IRDye 800; LI-COR Biosciences; 926-32212) were used as secondary antibodies. Membranes were imaged using a LI-COR Odyssey system.

#### *Surface Spreading of Meiotic Nuclei, Immunofluorescence and Quantification of Synapsis*

Surface spreading of meiotic nuclei was performed as described by (Grubb et al., 2015). During spheroplasting, 20 μl of dithiothreitol (DTT) was used instead of the published 40 μl. Fixation was achieved with 4% PFA/sucrose solution. Spreads from Figure 7J,M were stained with anti-

Zip1 guinea pig polyclonal antibody (a gift from Scott Keeney, 1:500 dilution), and then with goat anti-guinea pig polyclonal secondary antibody (Alexa Fluor 555; ThermoFisher; A-21435, 1:200 dilution). Spreads from Figure S6 were stained with anti Zip1 goat polyclonal antibody (Santa Cruz Biotechnology; y-N16, 1:50 dilution), and then with donkey anti-goat polyclonal secondary antibody (Alexa Fluor 555; ThermoFisher; A-21432, 1:1000 dilution). Slides were mounted in antifade with added DAPI (Prolong Gold, Invitrogen; P36930) and images were captured using a Zeiss AxioPlan II microscope, Hamamatsu ORCA-ER CCD camera and Volocity software. Synapsis was quantified by characterizing four classes of Zip1 staining pattern: no Zip1, foci only, a mixture of foci and lines, and lines only (as previously described by Chen et al., 2015).

#### *FACS Analysis*

Cell pellets from 200 μl of meiotic culture were fixed in 70% ethanol. Cells were then washed in 1 ml of 50 mM sodium citrate (pH 7.5) and resuspended in 1 ml of 50 mM sodium citrate (pH 7.5) with 130 μg RNaseA and incubated at 37°C for 1 hour. 0.52 mg of Proteinase K was added and samples were incubated at 65°C for an additional hour. 100 μl of sodium citrate (pH 7.5) containing a 1:10,000 dilution of SYBR Green (Invitrogen S7563) and 25μl of 10% Triton-X 100 were added, and cells were sonicated with a probe sonicator on setting 1.5 for ten seconds. Cells were scanned on a FACScan (BD Biosciences) and data was acquired with CellQuest Pro, using the FL1 (green) detector to trigger doublet discrimination. Data was analyzed using FlowJo to gate live, single-cell events measuring SYBR Green signal. Histograms were plotted modally, or as the percent of the maximum.

### QUANTIFICATION AND STATISTICAL ANALYSIS

#### Processing of Mass Spectrometry Data

Raw files were searched with MaxQuant 1.6.1 (Max Planck Institute)(Cox and Mann, 2008). Samples from complementary digests for the same replicate were processed under the same experiment heading. Two digestion (Lys-C only and Lys-C + Glu-C) strategies were processed as separate groups. For all samples, a maximum of five missed cleavages and potential modifications were allowed per peptide. Two separate searches were run. Protein N-terminal acetylation, methionine oxidation and di-glycyl lysine were set as variable modifications, and carbamidomethyl cysteine was set as fixed modification in both runs. In addition, one search included serine, threonine and tyrosine phosphorylations, and the other had lysine acetylation as variable modifications. For peptides, a minimum length of 6 and maximum mass of 5200 Da were allowed. Label-free quantification was enabled with default settings (Cox et al., 2014), and was applied to all variable modifications. Standard orbitrap settings were used under the "instrument" parameter. An *S. cerevisiae* proteome downloaded from Uniprot was used for searches. Search parameters were adjusted to search for second peptides and dependent peptides, and to match identifications between samples. Defaults were used for all settings unless specified otherwise. Data tables generated by MaxQuant were used for further analysis.

**Data Analysis**

All proteomics data were processed with Perseus (Max Planck Institute)(Tyanova et al., 2016). Text files titled "proteinGroups" and "GlyGly (K)Sites" were loaded into a Perseus workspace. Protein and modification matrices were filtered to remove reverse matches and potential modifications. SUMOylated proteins were identified by filtering protein matrices for "GlyGly" modification. The list of Uniprot IDs of SUMOylated proteins was loaded on Panther Gene Ontology search (http://www.pantherdb.org/) and a statistical over-representation test was run for GO-slim Biological Processes and GO-slim cellular Component, with *S. cerevisiae* proteome as the background. To identify SUMOylation consensus sequence motifs, the motif configuration file of Perseus was edited to add the known SUMOylation consensus sequences, and motifs were added to the GlyGly-site matrix with “sequence window” as the search target. To calculate the number of proteins and SUMO sites identified at each timepoint, protein and SUMO-site matrices from the triplicate dataset were exported to Microsoft Excel and the entries in identification type columns were converted to numbers. Numbers for each triplicate sample were added up and rows containing valid values were counted to give number of identified proteins and sites respectively.

**Quantitative Proteomics**

Quantitative analysis was carried out in Perseus, with label-free quantitation generated by MaxQuant. For proteins, the “LFQ” values were used. For SUMO sites, “intensity *x_y*” values were used, where *x* is the timepoint and *y* is the replicate number. To analyze the temporal changes in SUMOylation states of proteins and SUMO sites, a new categorical annotation called “time” was applied to the protein and site matrices, and individual replicates were labeled to identify the timepoint they belonged to. This categorical annotation was used to average the quantitative data for triplicate samples at each timepoint. These averages were used for further analysis. To visualize the changes in protein SUMOylation profiles across the time course, the averaged LFQ profiles were loaded into Morpheus (https://software.broadinstitute.org/morpheus) and a similarity matrix was generated. To generate hierarchical clustering, proteins were filtered to isolate those that were quantifiable for at least one timepoint. LFQ values were transformed to Log (2). Invalid values were imputed from a separate normal distribution for each column. All values were then normalized by Z-scoring to fit them in a range of -2 to +2. These normalized LFQs were loaded into Morpheus and the rows were subjected hierarchical clustering by 1- Pearson correlation to generate clustering tree and heatmap.

The cumulative intensities of all identified sites were retrieved from the “intensity” values from the GlyGly site matrix of a full search of all (63) samples.

### Identifying Types of Secondary Structure That Are Enriched for SUMOylation Sites

For all proteins with at least one identified SUMOylation site, each lysine residue was scored for whether or not it was predicted to reside in different structural contexts. Globular domains, long disordered regions, and short disordered regions were predicted using IUPred (Dosztanyi, 2018) with default settings. An additional “Disordered” class was defined as any region predicted to be disordered, that is not inside a predicted globular domain. Solvent exposed and buried regions were predicted using ACCpro from the Scratch package (Magnan and Baldi, 2014). “Globular-Exposed” and “Globular-Buried” classes were then defined based on the combined IUPred and ACCpro predictions. Coiled-coil regions were predicted using ncoils with default parameters (Lupas et al., 1991). Transmembrane regions were predicted using HMMTOP version 2.1 with default settings (Tusnady and Simon, 1998). “Disordered Not CC” was defined from the Disordered and coiled coil predictions. “Disordered Terminus” was defined as a disordered region that extends to the N- or C-terminus of the protein. “Disorder in 100 AA of Terminus” was defined as being in a Disordered Not CC region and being within 100 amino acids of either the N- or C-terminus of the protein. For each structure class, the fold-enrichment was computed as the percentage of detected lysine SUMOylation sites falling within the structure class divided by the percentage of all lysines (in proteins with at least one detected SUMOylation site) falling within the same structure class.

### Analysis of SUMOylation Site Clustering

For all proteins with at least one identified SUMOylation site, the fraction of lysines detected as SUMOylated was computed as a function of the distance in amino acids from either a SUMOylated lysine (distance from SUMO-K in Figure 3F) or from a lysine (distance from K). To compute these curves, for every SUMOylation site in our data set we computed the distance to each other lysine in the protein and annotated whether or not the lysine was detected as a SUMOylation site. We then pooled the data for all SUMOylation sites and computed the fraction of lysines that were detected as SUMOylation sites as a function of distance in amino acids (rounded to the nearest ten amino acids). We then completed an analogous computation using all lysines rather than just the SUMOylation sites.

**Local Disorder Analysis Around SUMOylation Sites**

We measured the distribution of quantitative "Short Disorder" scores from IUPred for lysines detected as SUMOylation sites and for lysines that were not detected as SUMOylation sites.

**Diagrams of SUMOylation Sites Relative to Protein Domains and Secondary Structure**

For all proteins with at least one identified SUMOylation site, we generated an image annotating the positions of all lysines (upper track, black lines), all detected SUMOylation sites (upper track, red lines with residue numbers annotated), all predicted SUMO-interacting motifs (SIMs) defined using the GPS-SUMO algorithm with either the high (upper track, dark gray rectangles), medium (upper track, gray rectangles), or low (upper track, light gray rectangles) score threshold (Zhao et al., 2014). In the middle track, protein domains detected with HMMER version 3.1b2 and Pfam-A version 29 set of models are shown (Eddy, 1998; Finn et al., 2016). In the bottom track, globular domains predicted by IUPred (blue rectangles), coiled-coil regions predicted by ncoils (red rectangles), and transmembrane regions predicted by HMMTOP (black rectangles) are shown. The short disorder score from IUPred is also indicated by the width of light red region.

### DATA AVAILABILITY

https://www.ebi.ac.uk/pride/archive/projects/PXD012418

**KEY RESOURCES TABLE**

| Reagent or resource | Source | Identifier |
|---|---|---|
| Histrap FF 1ml | GE Healthcare | 17-5319-01 |
| Lysyl Endopeptidase (Lys-C) | Wako Chemicals | 125-02541 |
| Glu-C, sequencing grade | Promega | V1651 |
| Sep-Pak tC18 1cc vac cartridge 50mg | Waters | WAT054960 |
| PTMScan Ubiquitin Remnant Motif (K-ε-GG) | Cell Signaling technology | 5562S |
| PP1 | Tocris | 1397 |
| 3-Indoleacetic acid (Auxin) | Sigma-Aldrich | I3750-25G-A |
| β-Estradiol | Sigma-Aldrich | E2758 |
| Anti-c-Myc mouse monoclonal antibody | Roche | 11667149001, RRID: AB_390912 |
| Anti-Arp7 goat polyclonal antibody | Santa Cruz Biotechnology | y-C20, RRID: AB_671730 |
| Donkey anti-goat, IRDye 680 | LI-COR Biosciences | 926-68074, RRID: AB_10956736 |
| Donkey anti-mouse, IRDye 800 | LI-COR Biosciences | 926-32212, RRID: AB_621847 |
| Goat anti-guinea pig, Alexa Fluor 555 | Thermo Fisher | A-21435, RRID: AB_2535856 |
| Zip1 goat polyclonal antibody | Santa Cruz Biotechnology | y-N16, RRID:AB_794 259 |
| Donkey anti-goat, Alexa Fluor 555 | Thermo Fisher | A-21432, RRID: AB_141788 |

**Strain Table**

| Strain number | Mating type | Genotype | Experiment |
|---|---|---|---|
| **NHY7590** | *a/ α* | NHY7067 x NHY7083 | SUMO Proteomics |
| **NHY10438** | *a/ α* | NHY10416 x NHY10425 | Aos1-AID Condition 1 |
| **NHY10439** | *a/ α* | NHY10419 x NHY10422 | Aos1-AID Condition 1 |
| **NHY10406** | *a/ α* | NHY10318 x NHY10394 | Aos1-AID Condition 2,3 |
| **NHY10407** | *a/ α* | NHY10319 x NHY10395 | Aos1-AID Condition 2,3 |
| **NHY10408** | *a/ α* | NHY10316 x NHY10396 | Aos1-AID Condition 2,3 |
| **NHY10413** | *a/ α* | NHY10402 x NHY10410 | Aos1-AID Condition 4 |
| **NHY10414** | *a/ α* | NHY10403 x NHY10411 | Aos1-AID Condition 4 |
| **NHY10415** | *a/ α* | NHY10398 x NHY10412 | Aos1-AID Condition 4 |
| **NHY10440** | *a/ α* | NHY10428 x NHY10436 | Uba2-AID Condition 1 |
| **NHY10441** | *a/ α* | NHY10431 x NHY10434 | Uba2-AID Condition 1 |
| **NHY10371** | *a/ α* | NHY10320 x NHY10366 | Uba2-AID Condition 2,3 |
| **NHY10372** | *a/ α* | NHY10321 x NHY10368 | Uba2-AID Condition 2,3 |
| **NHY10382** | *a/ α* | *NHY10380 x NHY10381* | Uba2-AID Condition 4 |
| **NHY10383** | *a/ α* | *NHY10378 x NHY10381* | Uba2-AID Condition 4 |
| **NHY7067** | *a* | *ho LYS2 HIS4::LEU2-(BamHI; +ori) HIS6-StrepII-Smt3-KGG::KanMX ura3::pGPD1-GAL4-ER::URA3 pGAL-Ndt80::TRP1 irt1::pCUP1-IME1::NatMX ime4::pCUP1-IME4::NatMX* | |
| **NHY7083** | *α* | *ho his4-X::LEU2-(NgoMIV; +ori)--URA3 HIS6-StrepII-Smt3-KGG::KanMX ura3::pGPD1-GAL4-ER::URA3 pGAL-Ndt80::TRP1 irt1::pCUP1-IME1::NatMX ime4::pCUP1-IME4::NatMX* | |
| **NHY10416** | *α* | *ho::hisG leu2::hisG ura3(∆Sma-Pst) HIS4::LEU2-(BamHI; +ori) pCUP1-1-OsTIR1-9Myc-URA3 AOS1-AID-9MYC::HphMX* | |
| **NHY10419** | *a* | *ho::hisG leu2::hisG ura3(∆Sma-Pst) HIS4::LEU2-(BamHI; +ori) pCUP1-1-OsTIR1-9Myc-URA3 AOS1-AID-9MYC::HphMX* | |
| **NHY10422** | *α* | *ho::hisG leu2::hisG ura3(∆Sma-Pst) his4-X::LEU2-* | |

| | | |
|---|---|---|
| | | *(NgoMIV; +ori)--URA3 lys2::HphMX::pCUP1-1-OsTIR1-9myc AOS1-AID-9MYC::HphMX* |
| **NHY10425** | *a* | *ho::hisG leu2::hisG ura3(∆Sma-Pst) his4-X::LEU2-(NgoMIV; +ori)--URA3 lys2::HphMX::pCUP1-1-OsTIR1-9myc AOS1-AID-9MYC::HphMX* |
| **NHY10316** | *α* | *ho leu2::hisG arg4-Nsp lys2 ura3(∆Sma-Pst) HIS4::LEU2-(BamHI; +ori) pCUP1-1-OsTIR1-9Myc-URA3 cdc7-as3-MYC mcm5-bob1 AOS1-AID-9MYC::HphMX* |
| **NHY10318** | *a* | *ho leu2::hisG arg4-Nsp lys2 ura3(∆Sma-Pst) HIS4::LEU2-(BamHI; +ori) pCUP1-1-OsTIR1-9Myc-URA3 cdc7-as3-MYC mcm5-bob1 AOS1-AID-9MYC::HphMX* |
| **NHY10319** | *a* | *ho leu2::hisG arg4-Nsp lys2 ura3(∆Sma-Pst) HIS4::LEU2-(BamHI; +ori) pCUP1-1-OsTIR1-9Myc-URA3 cdc7-as3-MYC mcm5-bob1 AOS1-AID-9MYC::HphMX* |
| **NHY10394** | *α* | *ho::hisG leu2::hisG ura3(∆Sma-Pst) his4-X::LEU2-(NgoMIV; +ori)--URA3 lys2::HphMX::pCUP1-1-OsTIR1-9myc cdc7-as3-MYC mcm5-bob1 AOS1-AID-9MYC::HphMX* |
| **NHY10395** | *α* | *ho::hisG leu2::hisG ura3(∆Sma-Pst) his4-X::LEU2-(NgoMIV; +ori)--URA3 lys2::HphMX::pCUP1-1-OsTIR1-9myc cdc7-as3-MYC mcm5-bob1 AOS1-AID-9MYC::HphMX* |
| **NHY10396** | *a* | *ho::hisG leu2::hisG ura3(∆Sma-Pst) his4-X::LEU2-(NgoMIV; +ori)--URA3 lys2::HphMX::pCUP1-1-OsTIR1-9myc cdc7-as3-MYC mcm5-bob1 AOS1-AID-9MYC::HphMX* |
| **NHY10398** | *α* | *ho::hisG leu2::hisG ura3(∆Sma-Pst) HIS4::LEU2-(BamHI; +ori) HphMX::PGAL1-NDT80 pCUP1-1-OsTIR1-9Myc-URA3 AOS1-AID-9MYC::HphMX* |
| **NHY10402** | *a* | *ho::hisG leu2::hisG ura3(∆Sma-Pst) HIS4::LEU2-(BamHI; +ori) HphMX::PGAL1-NDT80 pCUP1-1-OsTIR1-9Myc-URA3 AOS1-AID-9MYC::HphMX* |
| **NHY10403** | *a* | *ho::hisG leu2::hisG ura3(∆Sma-Pst) HIS4::LEU2-(BamHI; +ori) HphMX::PGAL1-NDT80 pCUP1-1-OsTIR1-9Myc-URA3 AOS1-AID-9MYC::HphMX* |
| **NHY10410** | *α* | *ho::hisG leu2::hisG ura3(∆Sma-Pst) his4-X::LEU2-(NgoMIV; +ori)--URA3 Hygro::PGAL1-NDT80 ura3:PGPD1-GAL4(848)-ER:URA3 lys2::HphMX::pCUP1-1-OsTIR1-9myc AOS1-AID-9MYC::HphMX* |
| **NHY10411** | *α* | *ho::hisG leu2::hisG ura3(∆Sma-Pst) his4-X::LEU2-(NgoMIV; +ori)--URA3 HphMX::PGAL1-NDT80 ura3:PGPD1-GAL4(848)-ER:URA3 lys2::HphMX::pCUP1-1-OsTIR1-9myc AOS1-AID-9MYC::HphMX* |
| **NHY10412** | *a* | *ho::hisG leu2::hisG ura3(∆Sma-Pst) his4-X::LEU2-(NgoMIV; +ori)--URA3 Hygro::PGAL1-NDT80* |

| | | |
|---|---|---|
| | | *ura3:PGPD1-GAL4(848)-ER:URA3 lys2::HphMX::pCUP1-1-OsTIR1-9myc AOS1-AID-9MYC::HphMX* |
| **NHY10428** | *α* | *ho::hisG leu2::hisG ura3(ΔSma-Pst) HIS4::LEU2-(BamHI; +ori) pCUP1-1-OsTIR1-9Myc-URA3 UBA2-AID-9MYC::HphMX* |
| **NHY10431** | *a* | *ho::hisG leu2::hisG ura3(ΔSma-Pst) HIS4::LEU2-(BamHI; +ori) pCUP1-1-OsTIR1-9Myc-URA3 UBA2-AID-9MYC::HphMX* |
| **NHY10434** | *α* | *ho::hisG leu2::hisG ura3(ΔSma-Pst) his4-X::LEU2-(NgoMIV; +ori)--URA3 lys2::HphMX::pCUP1-1-OsTIR1-9myc UBA1-AID-9MYC::HphMX* |
| **NHY10436** | *a* | *ho::hisG leu2::hisG ura3(ΔSma-Pst) his4-X::LEU2-(NgoMIV; +ori)--URA3 lys2::HphMX::pCUP1-1-OsTIR1-9myc UBA1-AID-9MYC::HphMX* |
| **NHY10320** | *a* | *ho leu2::hisG arg4-Nsp lys2 ura3(ΔSma-Pst) HIS4::LEU2-(BamHI; +ori) pCUP1-1-OsTIR1-9Myc-URA3 cdc7-as3-MYC mcm5-bob1 UBA2-AID-9MYC::HphMX* |
| **NHY10321** | *a* | *ho leu2::hisG arg4-Nsp lys2 ura3(ΔSma-Pst) HIS4::LEU2-(BamHI; +ori) pCUP1-1-OsTIR1-9Myc-URA3 cdc7-as3-MYC mcm5-bob1 UBA2-AID-9MYC::HphMX* |
| **NHY10366** | *α* | *ho::hisG leu2::hisG ura3(ΔSma-Pst) his4-X::LEU2-(NgoMIV; +ori)--URA3 lys2::HphMX::pCUP1-1-OsTIR1-9myc cdc7-as3-MYC mcm5-bob1 UBA2-AID-9MYC::HphMX* |
| **NHY10368** | *α* | *ho::hisG leu2::hisG ura3(ΔSma-Pst) his4-X::LEU2-(NgoMIV; +ori)--URA3 lys2::HphMX::pCUP1-1-OsTIR1-9myc cdc7-as3-MYC mcm5-bob1 UBA2-AID-9MYC::HphMX* |
| **NHY10377** | *a* | *ho::hisG leu2::hisG ura3(ΔSma-Pst) HIS4::LEU2-(BamHI; +ori) HphMX::PGAL1-NDT80 pCUP1-1-OsTIR1-9Myc-URA3 UBA2-AID-9MYC::HphMX* |
| **NHY10380** | *α* | *ho::hisG leu2::hisG ura3(ΔSma-Pst) HIS4::LEU2-(BamHI; +ori) HphMX::PGAL1-NDT80 pCUP1-1-OsTIR1-9Myc-URA3 UBA2-AID-9MYC::HphMX* |
| **NHY10381** | *a* | *ho::hisG leu2::hisG ura3(ΔSma-Pst) his4-X::LEU2-(NgoMIV; +ori)--URA3 HphMX::PGAL1-NDT80 ura3:PGPD1-GAL4(848)-ER:URA3 lys2::HphMX::pCUP1-1-OsTIR1-9myc UBA2-AID-9MYC::HphMX* |

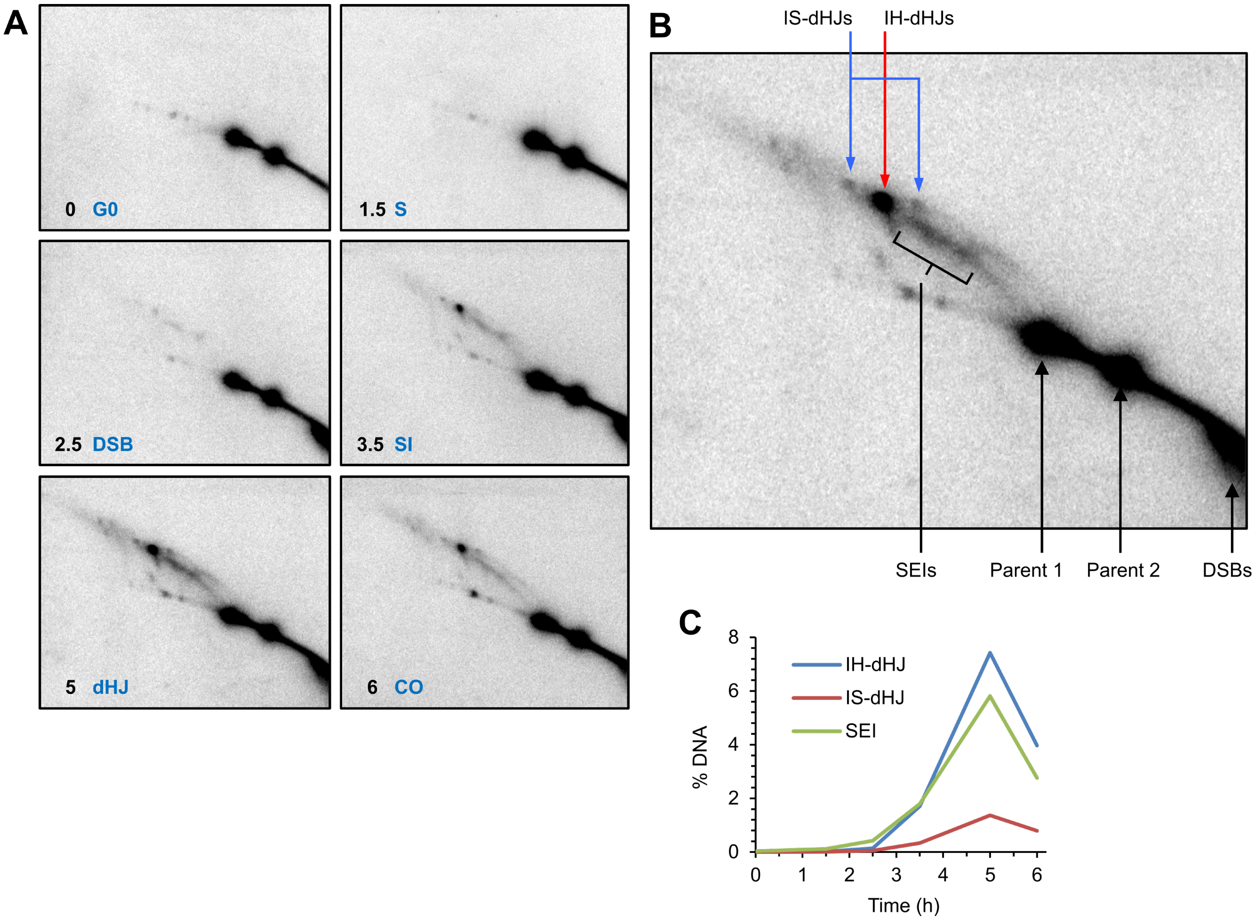

**Supplemental Figure 1 (related to Figure 1). Two-dimensional Southern analysis of recombination intermediates.**

**(A)** 2D gels Southern images representative of each timepoint collected for proteomics analysis. G0: G0 phase, S: S-phase, DSB: double-stranded break formation, SI: strand invasion, dHJ: double-Holliday junction formation, CO: crossover formation. **(B)** Single panel highlighting specific recombination intermediate. IH-dHJ: inter-homolog double-Holliday junction, IS-dHJ: inter-sister double-Holliday junction, SEI: single-end invasion, DSB: double-stranded break **(C)** Quantification of SEIs and dHJs over the entire meiotic time course.

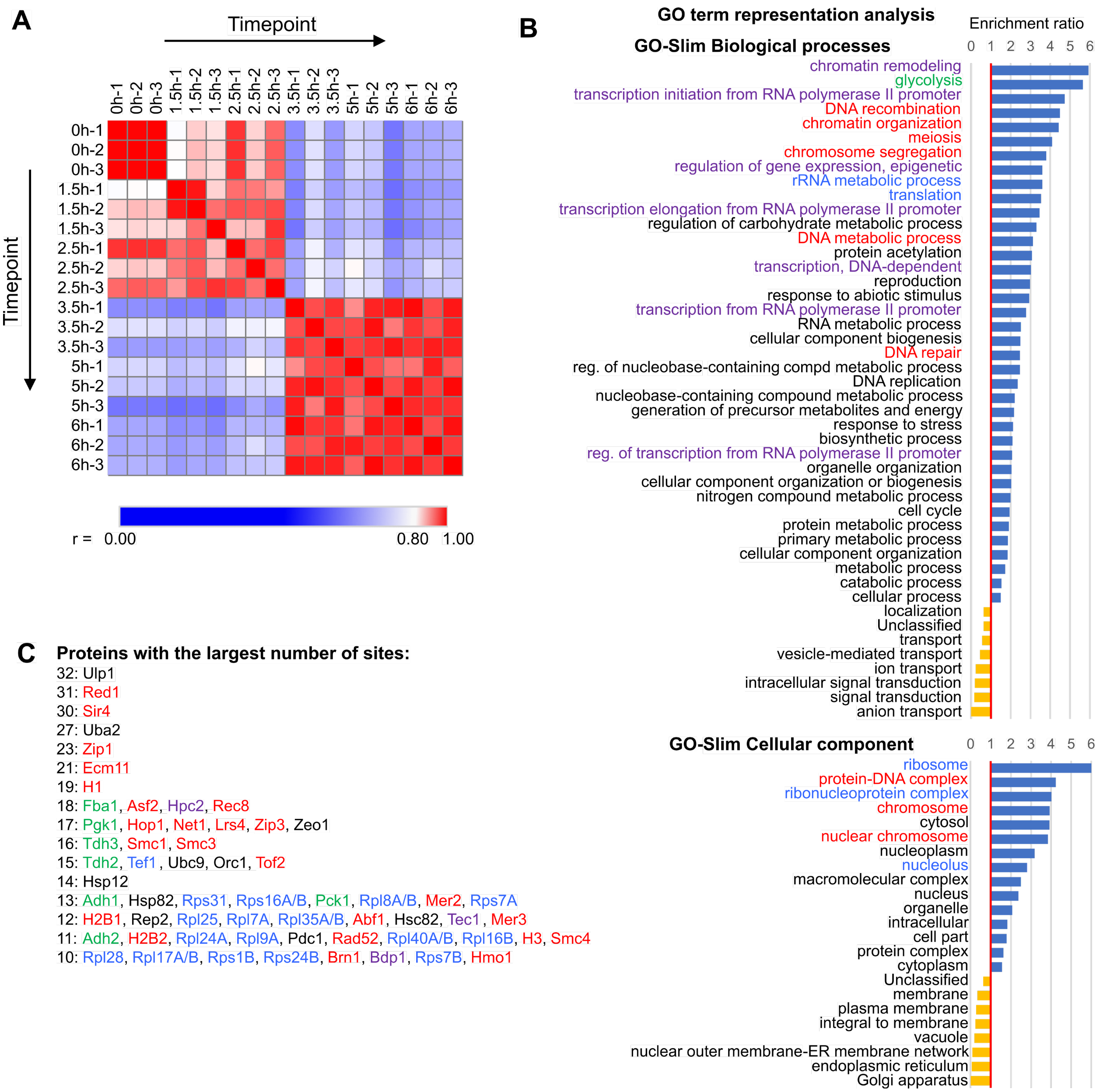

**Supplemental Figure 2 (related to Figure 2). (A)** Similarity matrix of samples used for label-free quantification. Heatmap representing the value of Pearson correlation coefficient (r) of protein LFQ (scale at the bottom). Labels indicate timepoint (hours) and replicate serial number. **(B)** Over-representation analysis of GO-Slim Biological processes and GO-Slim Cellular component showing categories that are significantly over- (blue bars) or under-represented (gold bars). Related processes are highlighted with colored text. **(C)** Proteins in descending order of SUMOylation site number, color-coded to match the GO categories that they represent.

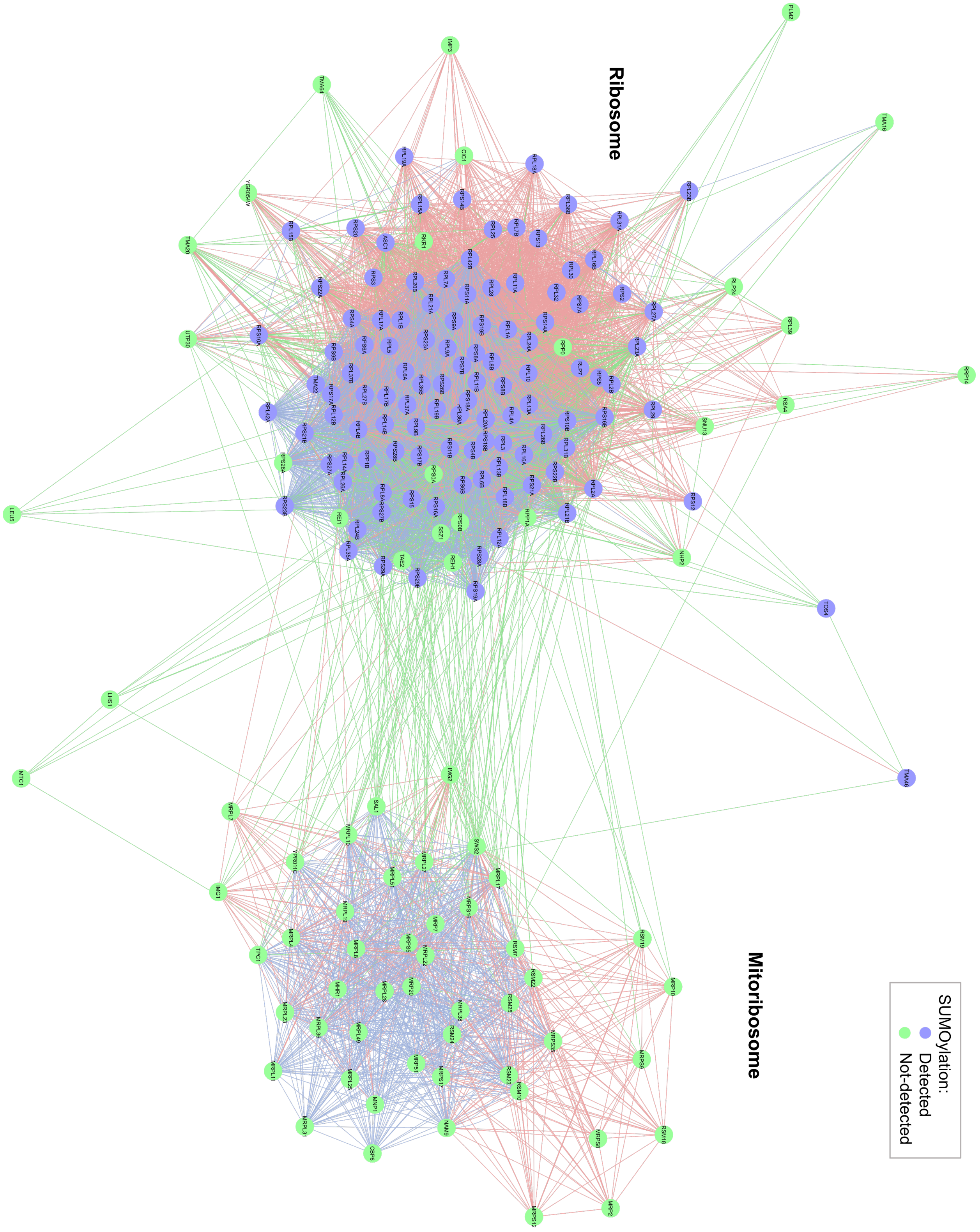

**Supplemental Figure 3 (related to Figure 2).** Network diagram showing SUMOylated members of the ribosome. Mito-ribosome components were not SUMOylated.

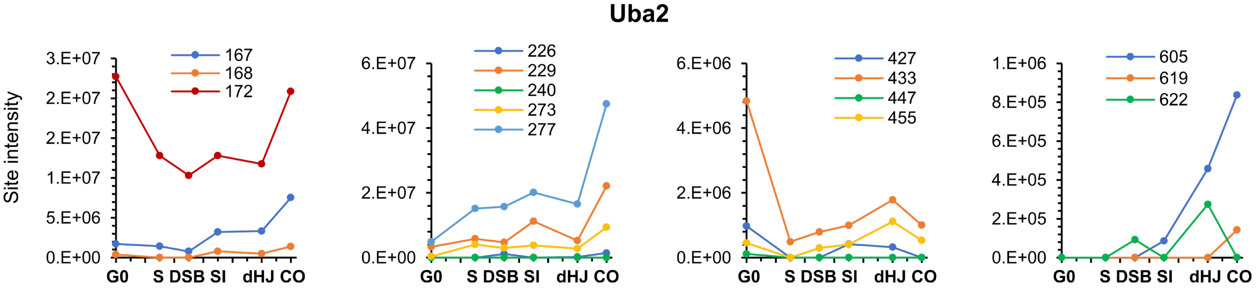

**Supplemental Figure 4 (related to Figure 4).** Diverse temporal profiles of SUMOylation sites on Uba2.

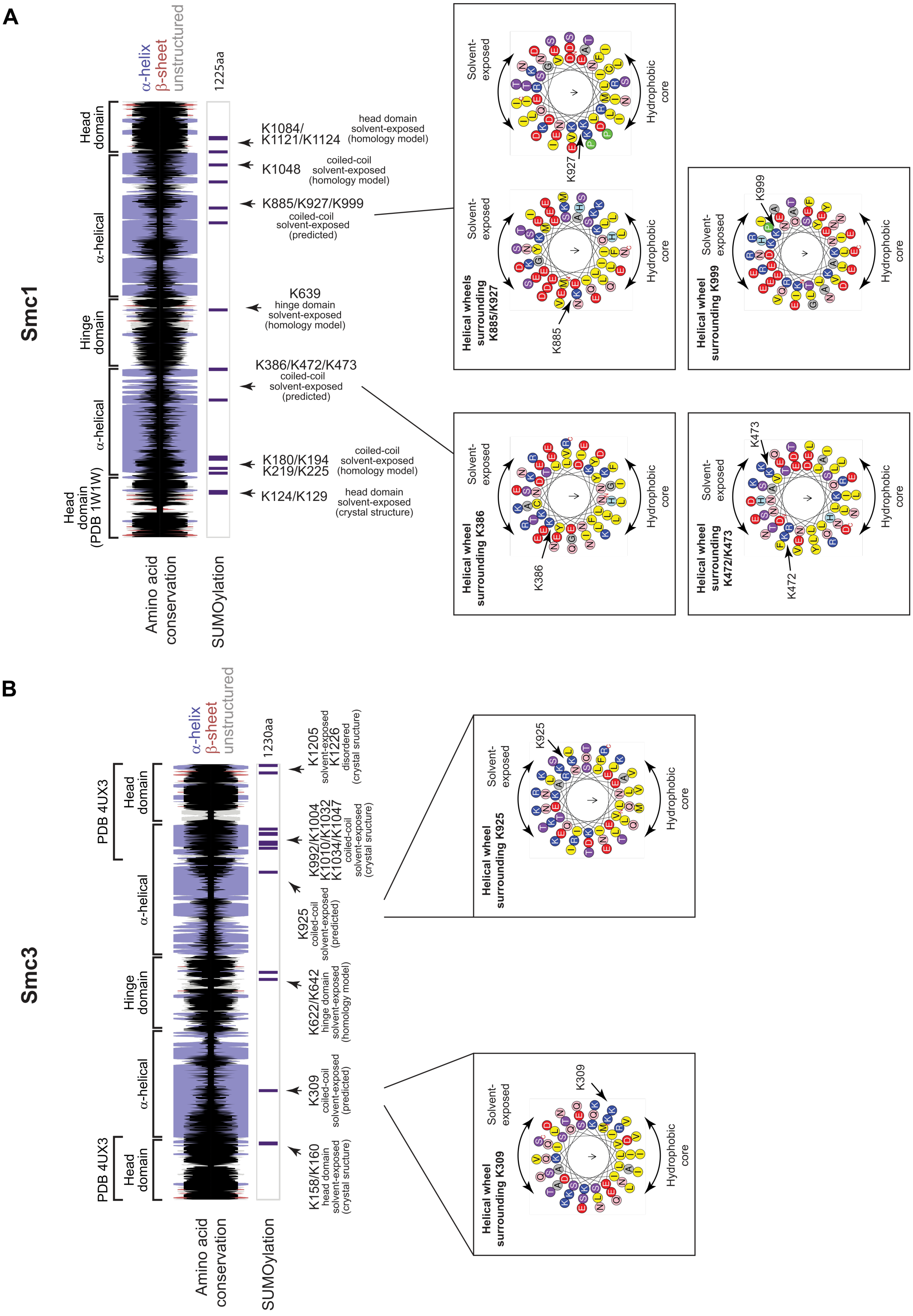

A
Smc1
α-helix
β-sheet
unstructured
1225aa
Head domain
α-helical
Hinge domain
α-helical
Head domain (PDB 1W1W)
Amino acid conservation
SUMOylation
K1084/ K1121/K1124
head domain solvent-exposed (homology model)
K1048
coiled-coil solvent-exposed (homology model)
K885/K927/K999
coiled-coil solvent-exposed (predicted)
K639
hinge domain solvent-exposed (homology model)
K386/K472/K473
coiled-coil solvent-exposed (predicted)
K180/K194 K219/K225
coiled-coil solvent-exposed (homology model)
K124/K129
head domain solvent-exposed (crystal structure)
Helical wheels surrounding K885/K927
Helical wheel surrounding K999
Helical wheel surrounding K386
Helical wheel surrounding K472/K473
Solvent-exposed
Hydrophobic core
K927
K885
K999
K386
K473
K472
B
Smc3
α-helix
β-sheet
unstructured
1230aa
PDB 4UX3
Head domain
α-helical
Hinge domain
α-helical
PDB 4UX3
Head domain
Amino acid conservation
SUMOylation
K1205 K1226
solvent-exposed disordered (crystal structure)
K992/K1004 K1010/K1032 K1034/K1047
coiled-coil solvent-exposed (crystal structure)
K925
coiled-coil solvent-exposed (predicted)
K622/K642
hinge domain solvent-exposed (homology model)
K309
coiled-coil solvent-exposed (predicted)
K158/K160
head domain solvent-exposed (crystal structure)
Helical wheel surrounding K925
Helical wheel surrounding K309
Solvent-exposed
Hydrophobic core
K925
K309

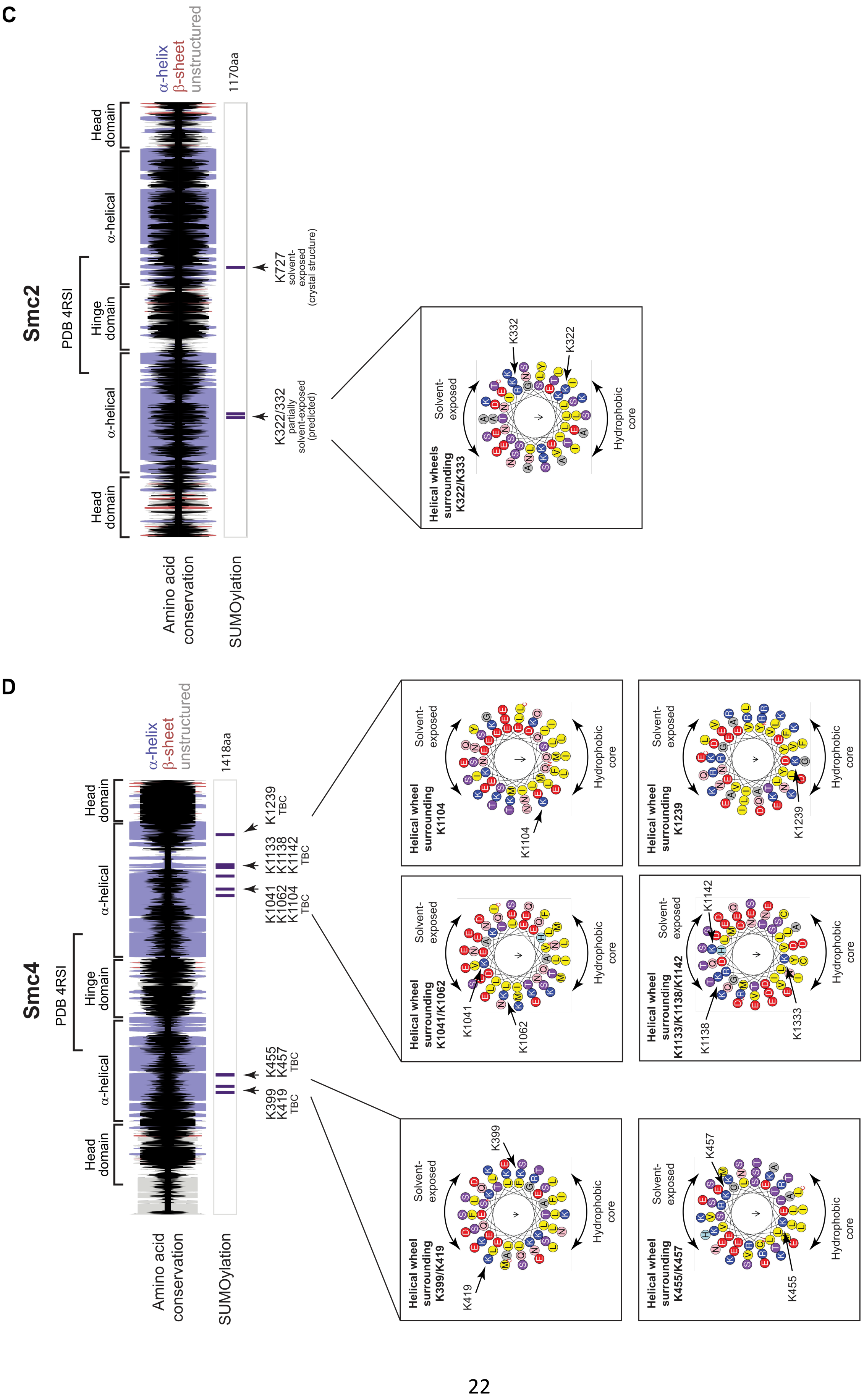
C
Smc2
PDB 4RSI
Head domain
α-helical
Hinge domain
α-helical
Head domain
α-helix
β-sheet
unstructured
1170aa
Amino acid conservation
SUMOylation
K727
solvent-exposed
(crystal structure)
K322/332
partially
solvent-exposed
(predicted)
Helical wheels surrounding K322/K333
K332
K322
Solvent-exposed
Hydrophobic core
D
Smc4
PDB 4RSI
Head domain
α-helical
Hinge domain
α-helical
Head domain
α-helix
β-sheet
unstructured
1418aa
Amino acid conservation
SUMOylation
K399
K419
TBC
K455
K457
TBC
K1041
K1062
K1104
TBC
K1133
K1138
K1142
TBC
K1239
TBC
Helical wheel surrounding K399/K419
K399
K419
Helical wheel surrounding K455/K457
K457
K455
Helical wheel surrounding K1041/K1062
K1041
K1062
Helical wheel surrounding K1133/K1138/K1142
K1142
K1138
K1333
Helical wheel surrounding K1104
K1104
Helical wheel surrounding K1239
K1239
Solvent-exposed
Hydrophobic core

**E**

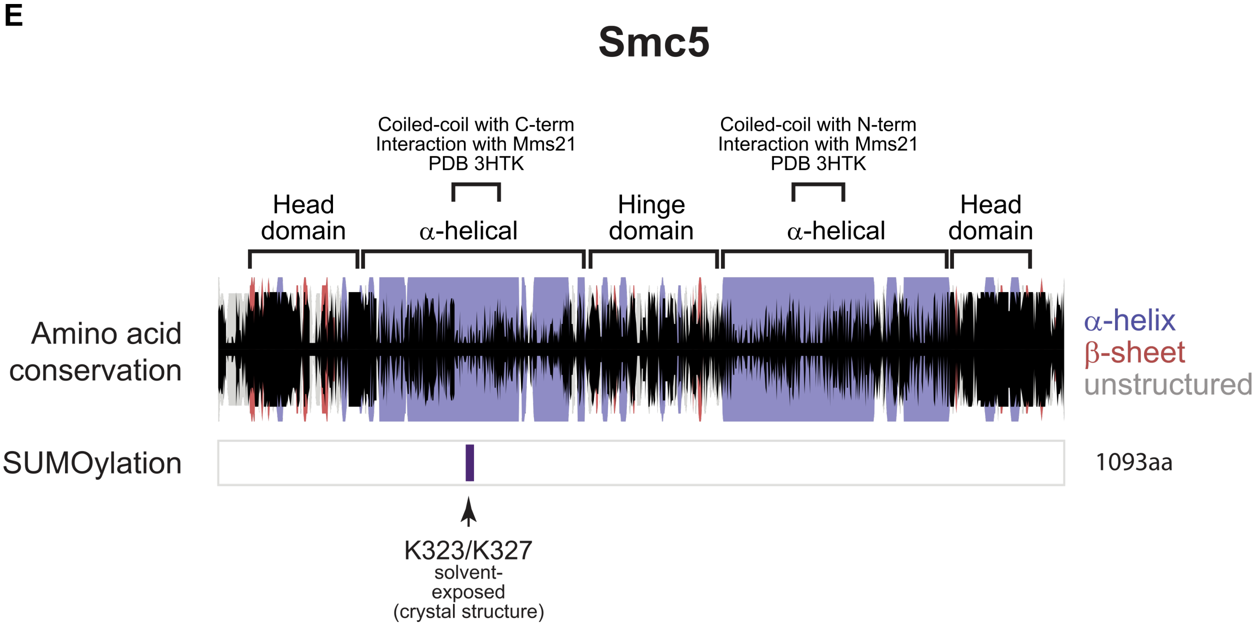

**Supplemental Figure 5 (related to Figure 5). Secondary structure and helical projections of SMC proteins.** Cartoons showing secondary structure predictions (α-helix: purple bar, β-sheet: red bar, unstructured: grey bar) and amino acid conservation (black bar) for SMC proteins: **(A)** Smc1 **(B)** Smc3 **(C)** Smc2 **(D)** Smc4 **(E)** Smc5. Width of the bar represents the level of AA conservation. SUMO sites are shown below. Insets show helical projections where SUMO sites lie on α-helix**.**

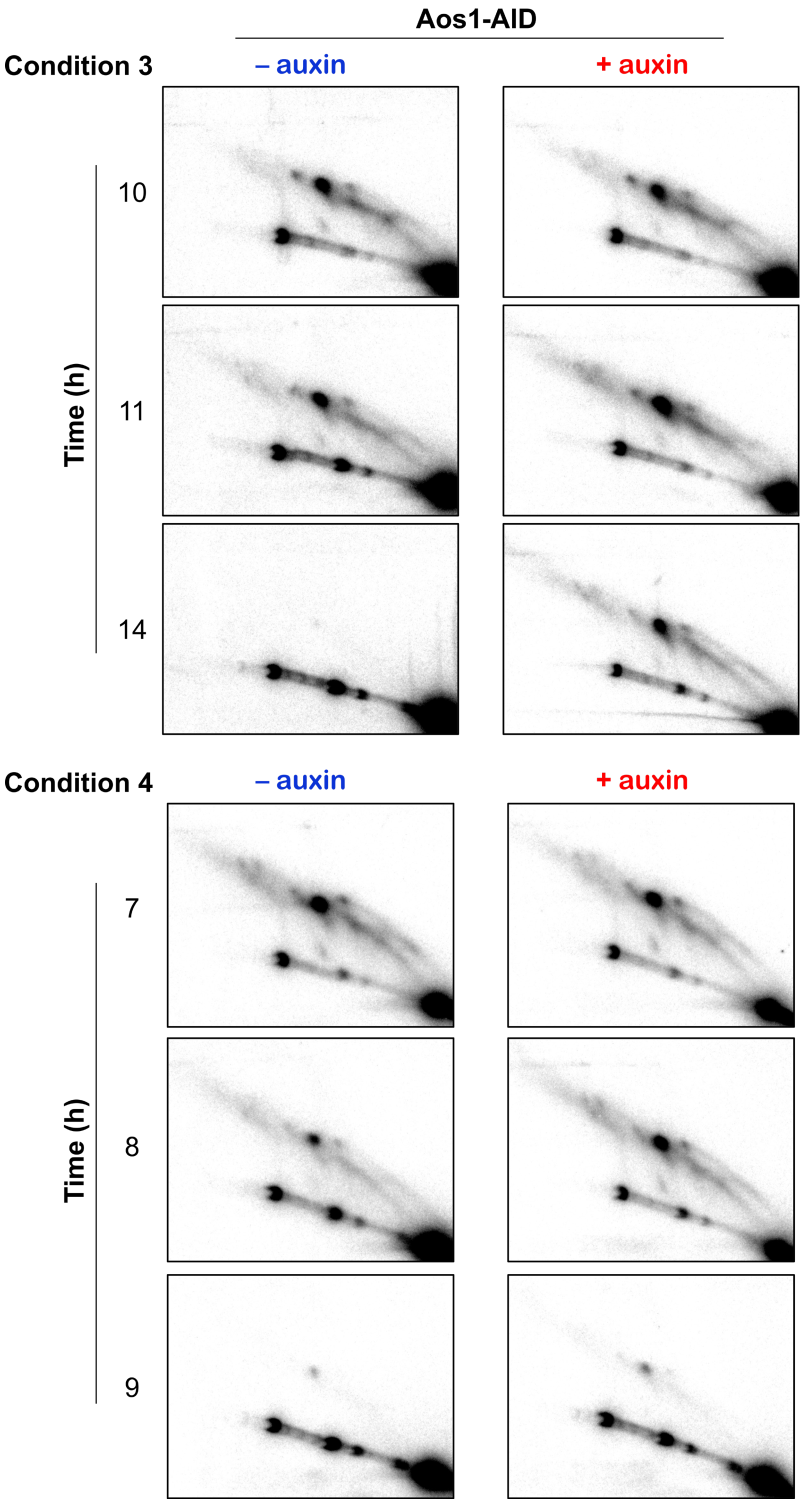

**Supplemental Figure 6 (related to Figure 7).** Southern blot images of joint-molecule analysis for experiments 3 and 4, with or without the addition of auxin to degrade Aos1-AID. Quantification is shown in **Figure 6**.

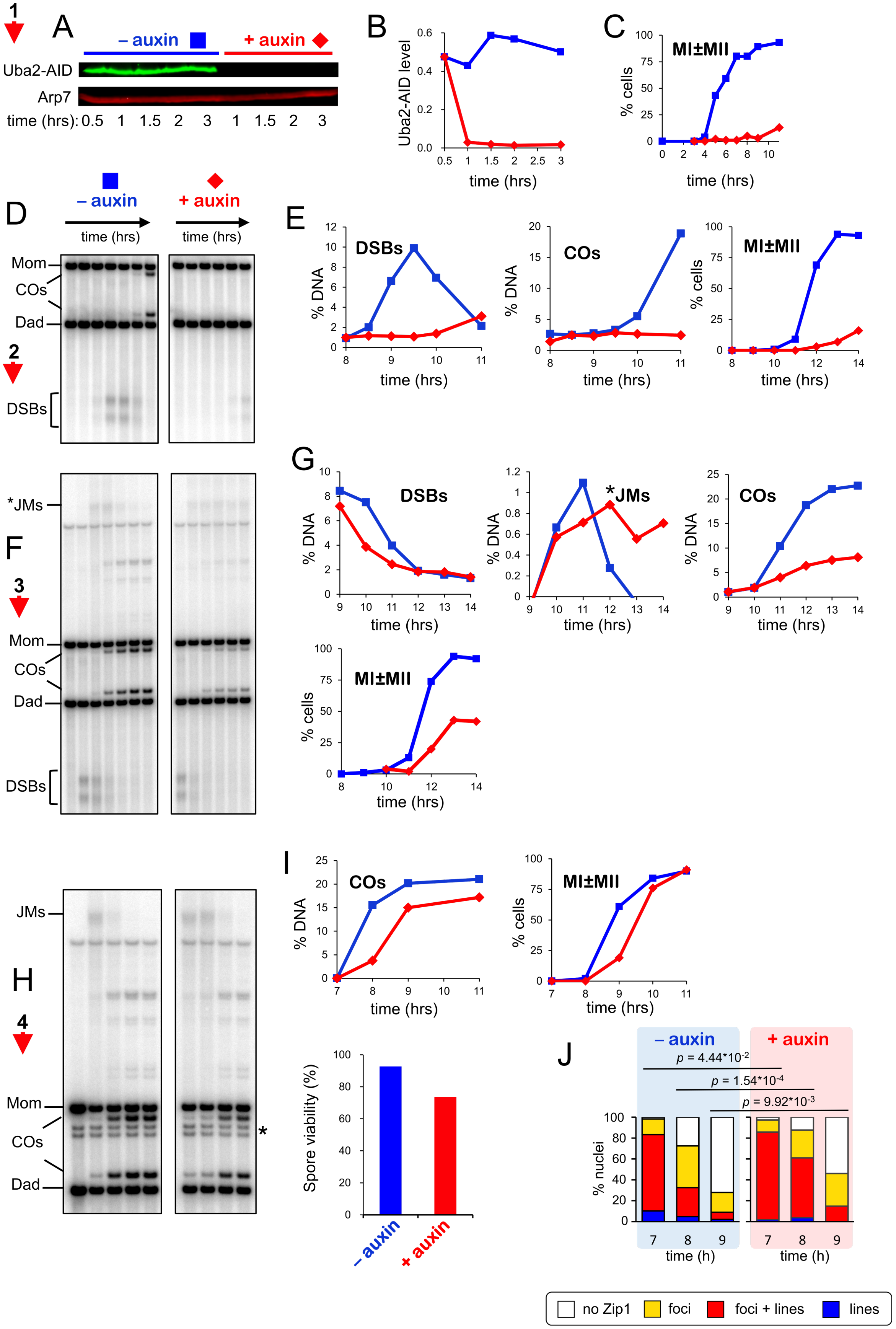

A
– auxin
+ auxin
Uba2-AID
Arp7
time (hrs): 0.5 1 1.5 2 3 1 1.5 2 3
B
Uba2-AID level
time (hrs)
C
MI±MII
% cells
time (hrs)
D
– auxin
+ auxin
time (hrs)
Mom
COs
Dad
DSBs
E
DSBs
COs
MI±MII
% DNA
% cells
time (hrs)
F
*JMs
Mom
COs
Dad
DSBs
G
DSBs
*JMs
COs
MI±MII
% DNA
% cells
time (hrs)
H
JMs
Mom
COs
Dad
I
COs
MI±MII
% DNA
% cells
time (hrs)
Spore viability (%)
– auxin
+ auxin
J
– auxin
+ auxin
p = 4.44*10-2
p = 1.54*10-4
p = 9.92*10-3
% nuclei
time (h)
no Zip1
foci
foci + lines
lines

**Supplemental Figure 7 (related to Figure 7). SUMO execution-point analysis using the Uba2-AID degron allele. (A)** Immunoblot showing the degradation of Uba2-AID after addition of auxin. **(B)** Quantification of the blot shown in (A). **(C)** Meiotic nuclear divisions (MI ± MII) for experiment 1. **(D)** 1D gel Southern blot image for analysis of DSBs and crossovers in experiment 2. **(E).** Quantification of DSBs, crossovers and meiotic divisions for experiment 2. **(F)** 1D gel Southern blot images for experiment 3. The top gel was used to quantify DSBs and crossovers (COs). The bottom gel was used to quantify non-crossover products (NCOs). **(G)** Quantification of DSBs, joint molecules (JMs), COs, NCOs and meiotic divisions for experiment 3. *JMs were quantified from the 1D Southern analysis. **(H)** 1D gel Southern blot image for experiment 4. **(I)** Quantification of COs and meiotic divisions for experiment 4. Spore viability was also analyzed and shown to be statistically lower following Uba2-AID degradation at the time of *IN-NDT80* expression (91.5% vs 76.4%, $p < 0.04$) **(J)** Quantification of synapsis (Zip1-staining classes) for experiment 4 indicates that SC disassembly is delayed.

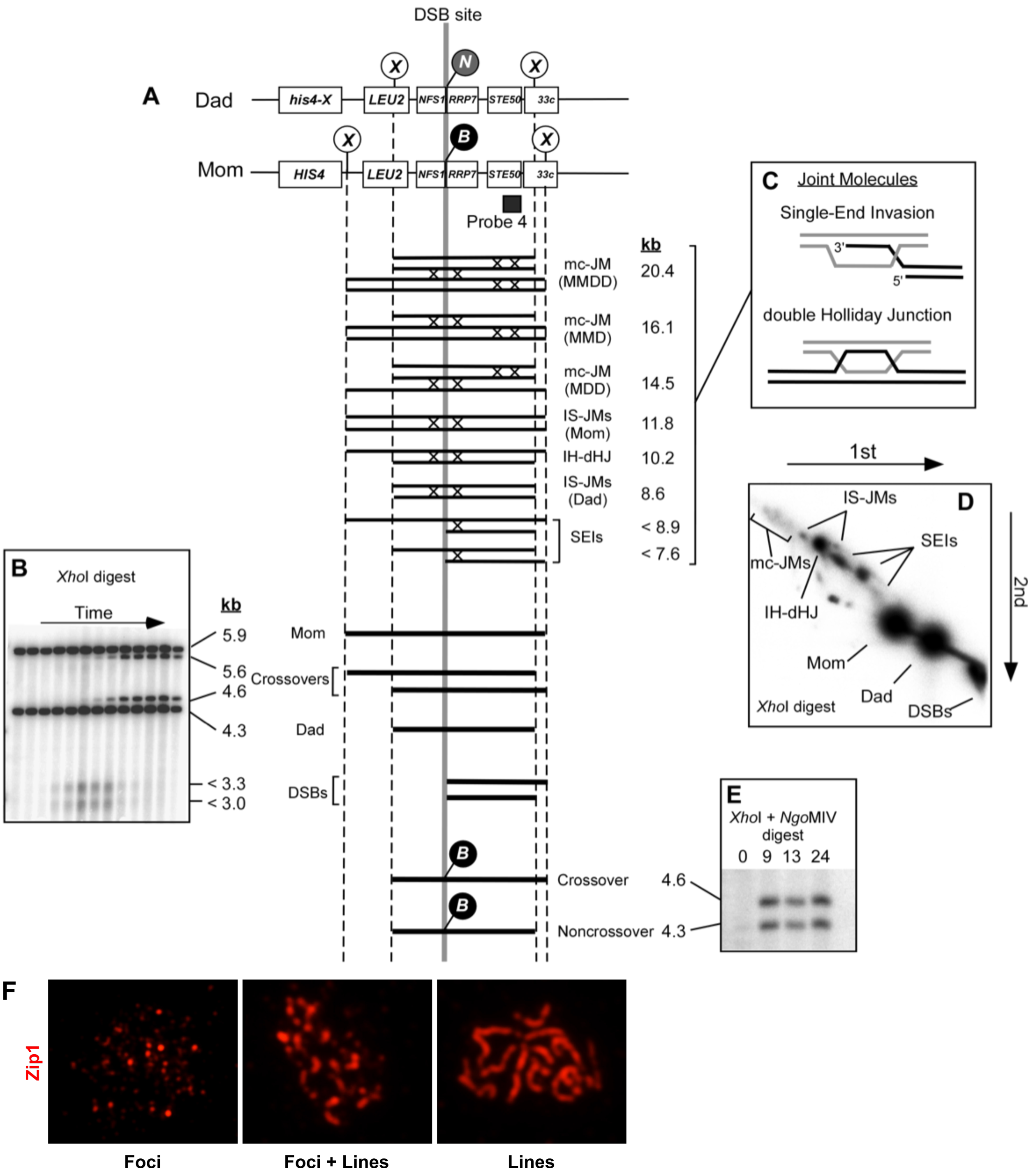

**Supplemental Figure 8 (related to Figure 7). Analysis of the meiotic recombination and synapsis. (A–E)** Physical assay for monitoring recombination at the *HIS4::LEU2* locus. **(A)** Map of the locus showing diagnostic restriction sites, hybridization probe, detectable DNA species and their molecular weights. X, XhoI site; B, BamHI site; N, NgoMIV site; DSB, double-strand break; IH-dHJ, interhomolog double-Holliday junction; IS-JM, inter-sister joint molecule; mc-JM, multi-chromatid joint molecule; SEI, single-end invasion. **(B)** Southern blot image of 1D gel analysis of DSBs and crossovers. **(C)** Inferred structures of SEI and dHJ intermediates. **(D)** Southern blot image of native/native 2D gel analysis of recombination intermediates. **(E)** 1D Southern blot image of 1D gel crossover/non-crossover analysis. **(F)** Representative immunofluorescence images of Zip1 immunostained nuclear spreads illustrating the different classes of Zip1 staining scored in Figure 7 and Supplemental Figure 7.

**Supplemental Table 1**

| Study | SUMOylated proteins reported | Overlap with this study |
|---|---|---|
| (Esteras et al., 2017) | 244 | 166 |
| (Lewicki et al., 2015) | 195 | 126 |
| (Albuquerque et al., 2015) | 31 | 26 |
| (Albuquerque et al., 2013) | 176 | 131 |
| (Hannich et al., 2005) | 150 | 93 |
| (Denison et al., 2005) | 251 | 151 |
| (Panse et al., 2004) | 151 | 51 |
| (Wohlschlegel et al., 2004) | 271 | 163 |

**Supplemental Table 2**

| | | **Number identified by SUMOsp** (Zhao et al., 2014) **(prediction threshold)** | | |
|---|---|---|---|---|
| **Lysine class** | **Number in dataset** | **(High)** | **(Medium)** | **(Low)** |
| **SUMOylated** | 2745 | 321 | 428 | 512 |
| **non-SUMOylated** | 29847 | 528 | 1105 | 1778 |

**References**

Albuquerque, C.P., Wang, G., Lee, N.S., Kolodner, R.D., Putnam, C.D., and Zhou, H. (2013). Distinct SUMO ligases cooperate with Esc2 and Slx5 to suppress duplication-mediated genome rearrangements. PLoS Genet *9*, e1003670.

Albuquerque, C.P., Yeung, E., Ma, S., Fu, T., Corbett, K.D., and Zhou, H.L. (2015). A Chemical and Enzymatic Approach to Study Site-Specific Sumoylation. Plos One *10*.

Benjamin, K.R., Zhang, C., Shokat, K.M., and Herskowitz, I. (2003a). Control of landmark events in meiosis by the CDK Cdc28 and the meiosis-specific kinase Ime2. Genes Dev *17*, 1524-1539.

Benjamin, K.R., Zhang, C., Shokat, K.M., and Herskowitz, I. (2003b). Control of landmark events in meiosis by the CDK Cdc28 and the meiosis-specific kinase Ime2. Genes & development *17*, 1524-1539.

Booher, K.R., and Kaiser, P. (2008). A PCR-based strategy to generate yeast strains expressing endogenous levels of amino-terminal epitope-tagged proteins. Biotechnol J *3*, 524-529.

Carlile, T.M., and Amon, A. (2008). Meiosis I is established through division-specific translational control of a cyclin. Cell *133*, 280-291.

Cox, J., Hein, M.Y., Luber, C.A., Paron, I., Nagaraj, N., and Mann, M. (2014). Accurate proteome-wide label-free quantification by delayed normalization and maximal peptide ratio extraction, termed MaxLFQ. Mol Cell Proteomics *13*, 2513-2526.

Cox, J., and Mann, M. (2008). MaxQuant enables high peptide identification rates, individualized p.p.b.-range mass accuracies and proteome-wide protein quantification. Nat Biotechnol *26*, 1367-1372.

Denison, C., Rudner, A.D., Gerber, S.A., Bakalarski, C.E., Moazed, D., and Gygi, S.P. (2005). A proteomic strategy for gaining insights into protein sumoylation in yeast. Mol Cell Proteomics *4*, 246-254.

Dosztanyi, Z. (2018). Prediction of protein disorder based on IUPred. Protein Sci *27*, 331-340.

Eddy, S.R. (1998). Profile hidden Markov models. Bioinformatics *14*, 755-763.

Esteras, M., Liu, I.C., Snijders, A.P., Jarmuz, A., and Aragon, L. (2017). Identification of SUMO conjugation sites in the budding yeast proteome. Microb Cell *4*, 331-341.

Finn, R.D., Coggill, P., Eberhardt, R.Y., Eddy, S.R., Mistry, J., Mitchell, A.L., Potter, S.C., Punta, M., Qureshi, M., Sangrador-Vegas, A*., et al.* (2016). The Pfam protein families database: towards a more sustainable future. Nucleic Acids Res *44*, D279-285.

Grubb, J., Brown, M.S., and Bishop, D.K. (2015). Surface Spreading and Immunostaining of Yeast Chromosomes. J Vis Exp, e53081.

Hannich, J.T., Lewis, A., Kroetz, M.B., Li, S.J., Heide, H., Emili, A., and Hochstrasser, M. (2005). Defining the SUMO-modified proteome by multiple approaches in Saccharomyces cerevisiae. J Biol Chem *280*, 4102-4110.

Johnson, E.S., and Blobel, G. (1999). Cell cycle-regulated attachment of the ubiquitin-related protein SUMO to the yeast septins. J Cell Biol *147*, 981-994.

Lewicki, M.C., Srikumar, T., Johnson, E., and Raught, B. (2015). The S. cereuisiae SUMO stress response is a conjugation-deconjugation cycle that targets the transcription machinery. J Proteomics *118*, 39-48.

Longtine, M.S., McKenzie, A., 3rd, Demarini, D.J., Shah, N.G., Wach, A., Brachat, A., Philippsen, P., and Pringle, J.R. (1998). Additional modules for versatile and economical PCR-based gene deletion and modification in Saccharomyces cerevisiae. Yeast *14*, 953-961.

Louvion, J.F., Havaux-Copf, B., and Picard, D. (1993). Fusion of GAL4-VP16 to a steroid-binding domain provides a tool for gratuitous induction of galactose-responsive genes in yeast. Gene *131*, 129-134.

Lupas, A., Van Dyke, M., and Stock, J. (1991). Predicting coiled coils from protein sequences. Science *252*, 1162-1164.

Magnan, C.N., and Baldi, P. (2014). SSpro/ACCpro 5: almost perfect prediction of protein secondary structure and relative solvent accessibility using profiles, machine learning and structural similarity. Bioinformatics *30*, 2592-2597.

Morawska, M., and Ulrich, H.D. (2013). An expanded tool kit for the auxin-inducible degron system in budding yeast. Yeast *30*, 341-351.

Oh, S.D., Jessop, L., Lao, J.P., Allers, T., Lichten, M., and Hunter, N. (2009). Stabilization and electrophoretic analysis of meiotic recombination intermediates in Saccharomyces cerevisiae. Methods Mol Biol *557*, 209-234.

Owens, S., Tang, S., and Hunter, N. (2018). Monitoring Recombination During Meiosis in Budding Yeast. Methods Enzymol *601*, 275-307.

Panse, V.G., Hardeland, U., Werner, T., Kuster, B., and Hurt, E. (2004). A proteome-wide approach identifies sumoylated substrate proteins in yeast. J Biol Chem *279*, 41346-41351.

Tang, S., Wu, M.K.Y., Zhang, R., and Hunter, N. (2015). Pervasive and essential roles of the Top3-Rmi1 decatenase orchestrate recombination and facilitate chromosome segregation in meiosis. Mol Cell *57*, 607-621.

Tusnady, G.E., and Simon, I. (1998). Principles governing amino acid composition of integral membrane proteins: application to topology prediction. J Mol Biol *283*, 489-506.

Tyanova, S., Temu, T., Sinitcyn, P., Carlson, A., Hein, M.Y., Geiger, T., Mann, M., and Cox, J. (2016). The Perseus computational platform for comprehensive analysis of (prote)omics data. Nat Methods *13*, 731-740.

Wohlschlegel, J.A., Johnson, E.S., Reed, S.I., and Yates, J.R., 3rd (2004). Global analysis of protein sumoylation in Saccharomyces cerevisiae. J Biol Chem *279*, 45662-45668.

Zhao, Q., Xie, Y., Zheng, Y., Jiang, S., Liu, W., Mu, W., Liu, Z., Zhao, Y., Xue, Y., and Ren, J. (2014). GPS-SUMO: a tool for the prediction of sumoylation sites and SUMO-interaction motifs. Nucleic Acids Res *42*, W325-330.
