## Supplemental Information - Protein Diagrams for "Delineation of the SUMO-Modified Proteome Reveals Regulatory Functions Throughout Meiosis"

#### Protein Diagram Legend:

##### Upper Track

- SUMOylated lysine
- Lysine

*Predicted SUMO Interaction Motifs*

- High threshold
- Medium threshold
- Low threshold

##### MiddleTrack

- PFAM domains

##### Lower Track

*Protein Secondary Structure*

- Globular Domains
- Disorder Score
- Coiled-coil
- Transmembrane

##### Horizontal axis

Amino acid residue number

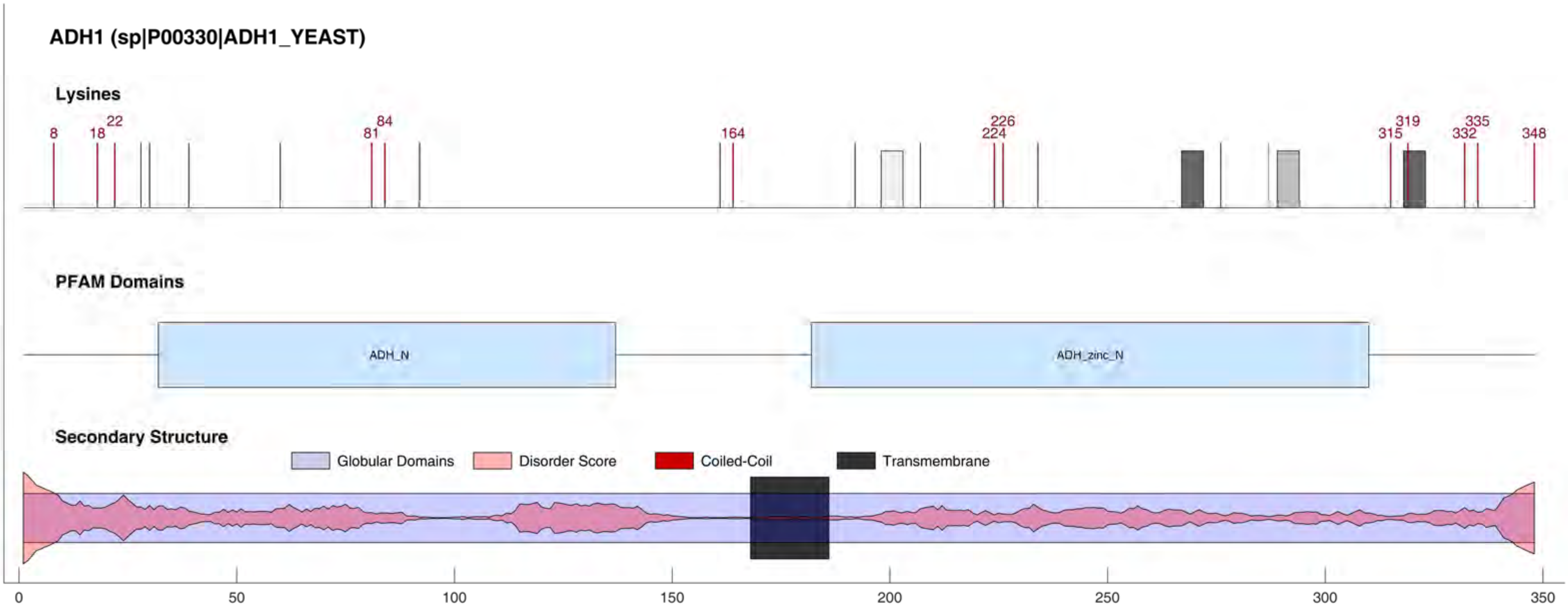

ADH1 (sp|P00330|ADH1_YEAST)
Lysines
8
18
22
81
84
164
224
226
315
319
332
335
348
PFAM Domains
ADH_N
ADH_zinc_N
Secondary Structure
Globular Domains
Disorder Score
Coiled-Coil
Transmembrane
0
50
100
150
200
250
300
350

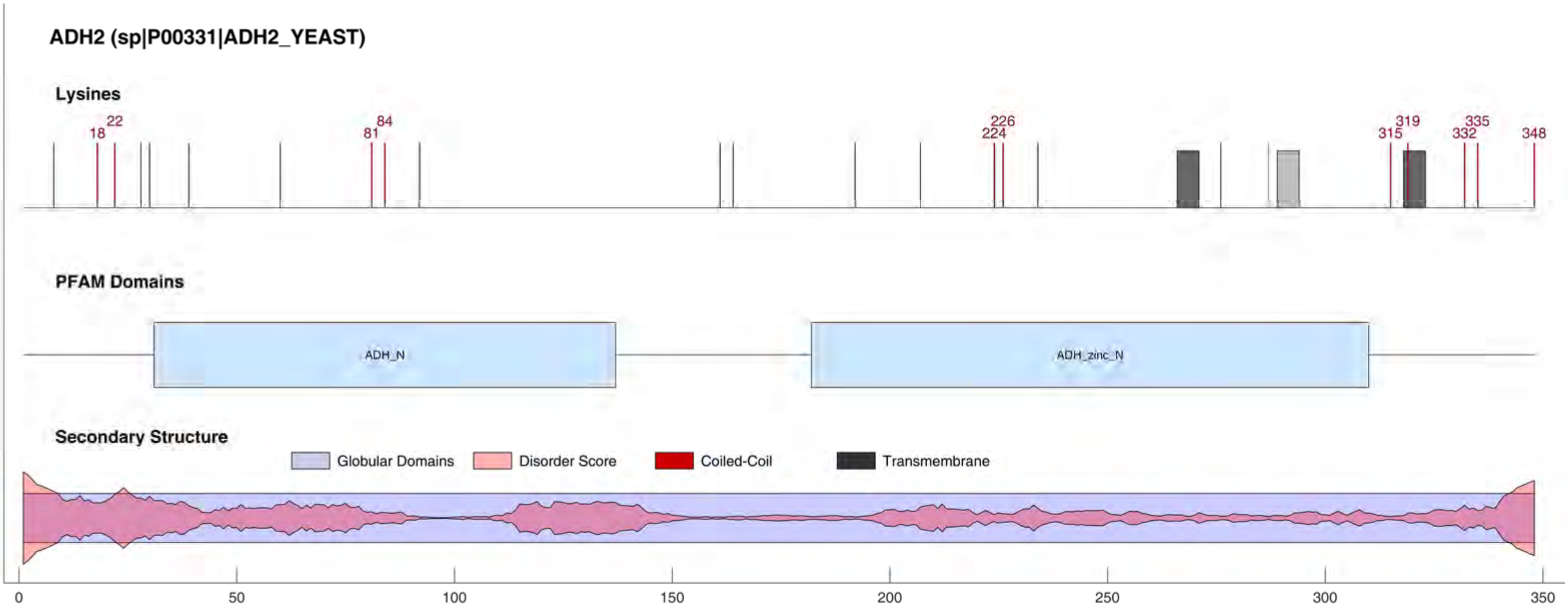
ADH2 (sp|P00331|ADH2_YEAST)
Lysines
18
22
81
84
224
226
315
319
332
335
348
PFAM Domains
ADH_N
ADH_zinc_N
Secondary Structure
Globular Domains
Disorder Score
Coiled-Coil
Transmembrane
0
50
100
150
200
250
300
350

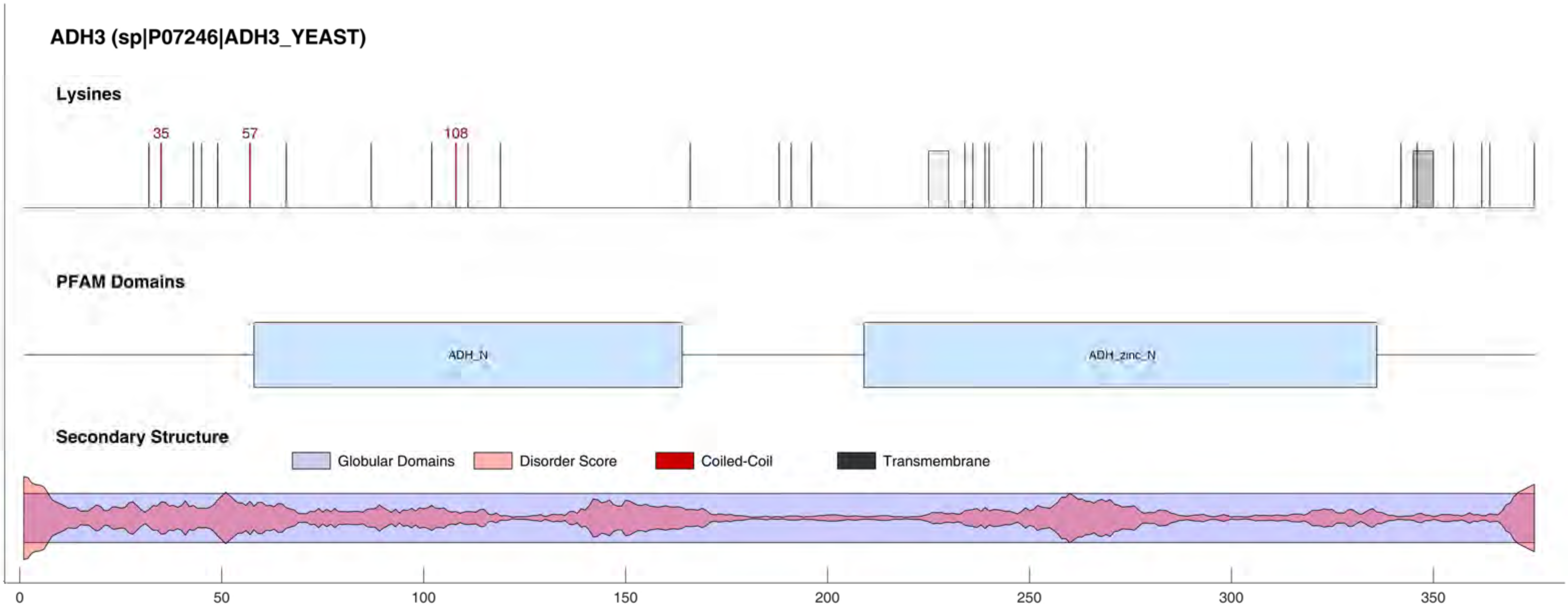
ADH3 (sp|P07246|ADH3_YEAST)
Lysines
35
57
108
PFAM Domains
ADH_N
ADH_zinc_N
Secondary Structure
Globular Domains
Disorder Score
Coiled-Coil
Transmembrane
0
50
100
150
200
250
300
350

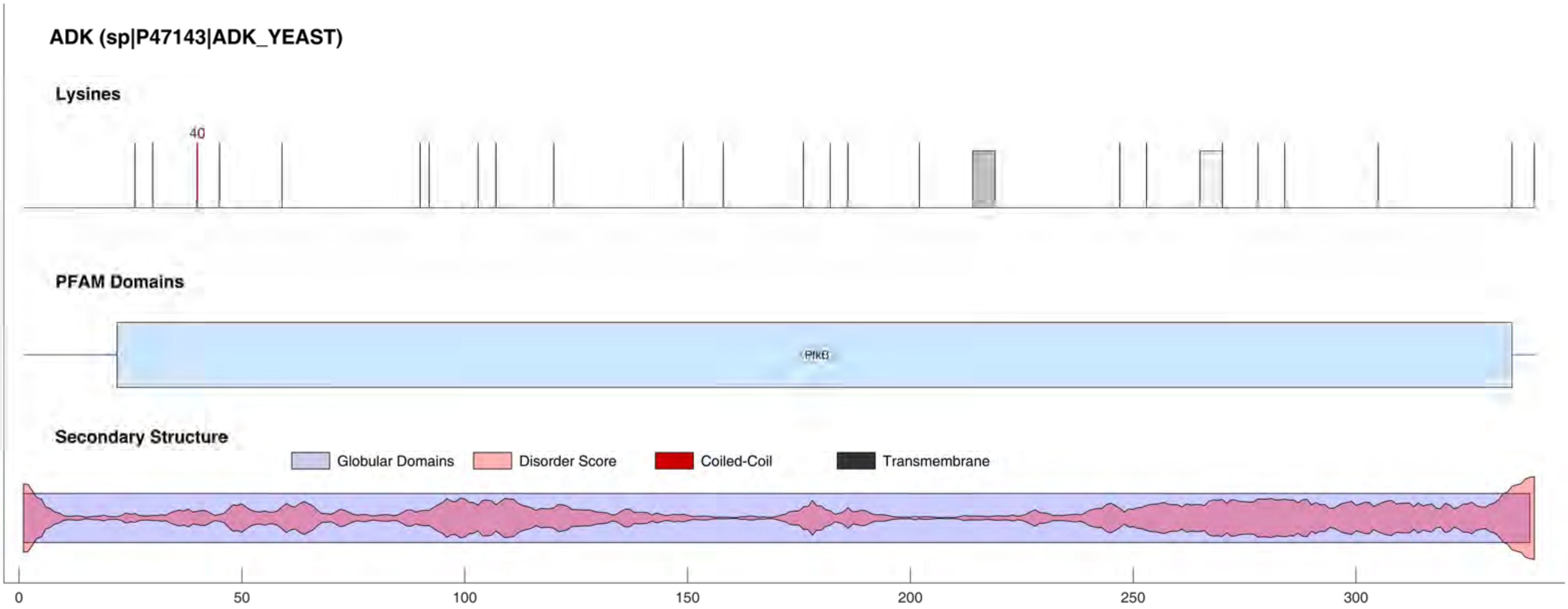

ADK (sp|P47143|ADK_YEAST)
Lysines
40
PFAM Domains
PfkB
Secondary Structure
Globular Domains
Disorder Score
Coiled-Coil
Transmembrane
0
50
100
150
200
250
300

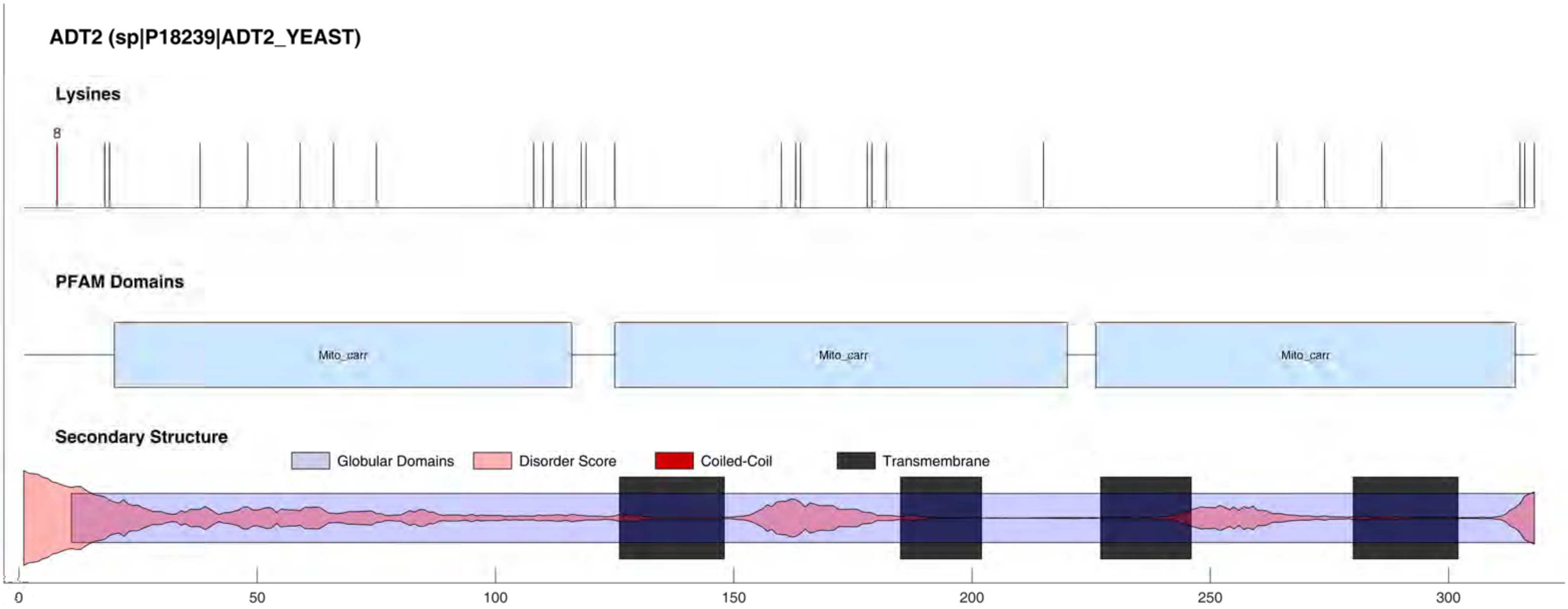

ADT2 (sp|P18239|ADT2_YEAST)
Lysines
8
PFAM Domains
Mito_carr
Mito_carr
Mito_carr
Secondary Structure
Globular Domains
Disorder Score
Coiled-Coil
Transmembrane
0
50
100
150
200
250
300

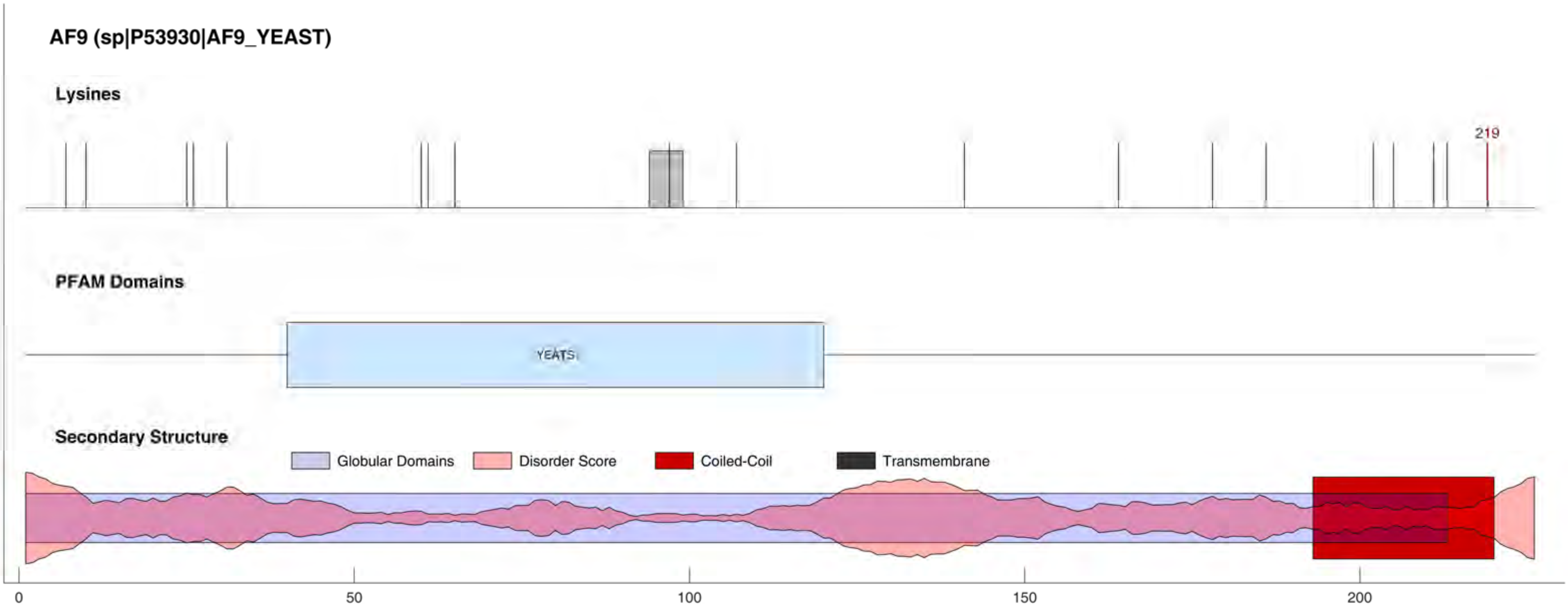
AF9 (sp|P53930|AF9_YEAST)
Lysines
219
PFAM Domains
YEATS
Secondary Structure
Globular Domains
Disorder Score
Coiled-Coil
Transmembrane
0
50
100
150
200

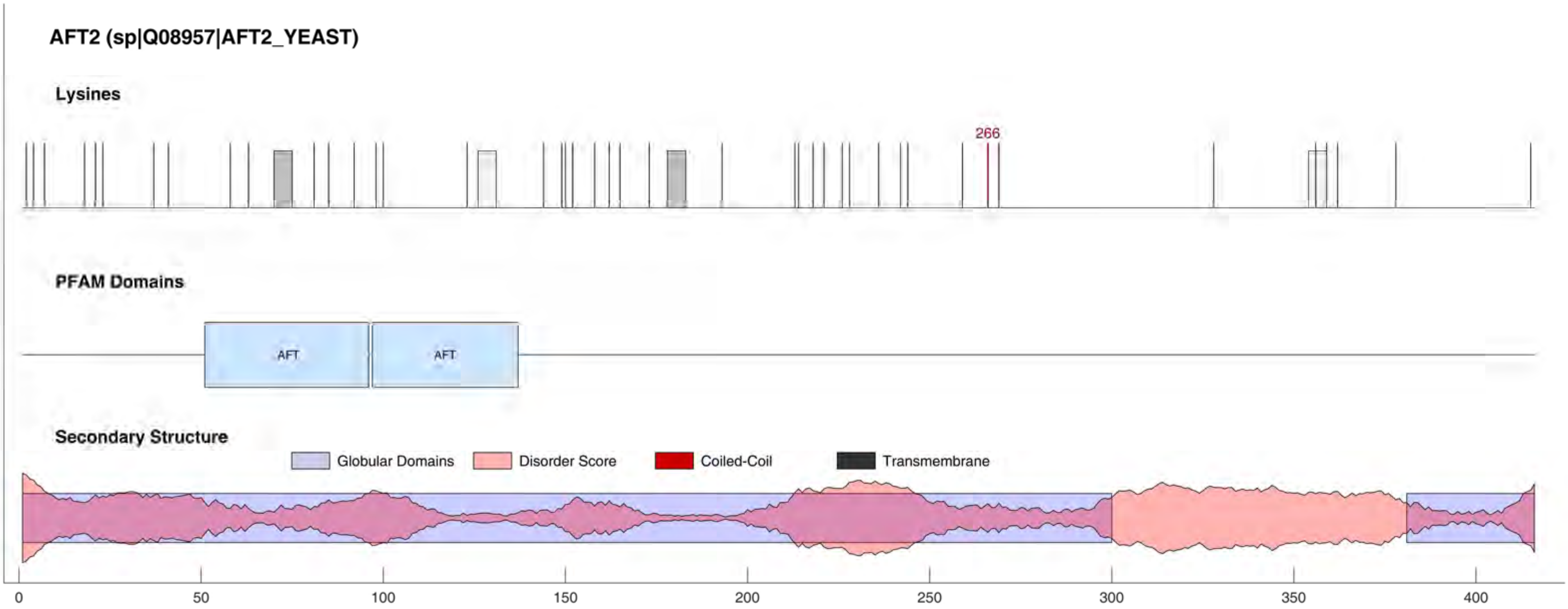

AFT2 (sp|Q08957|AFT2_YEAST)
Lysines
266
PFAM Domains
AFT
AFT
Secondary Structure
Globular Domains
Disorder Score
Coiled-Coil
Transmembrane
0
50
100
150
200
250
300
350
400

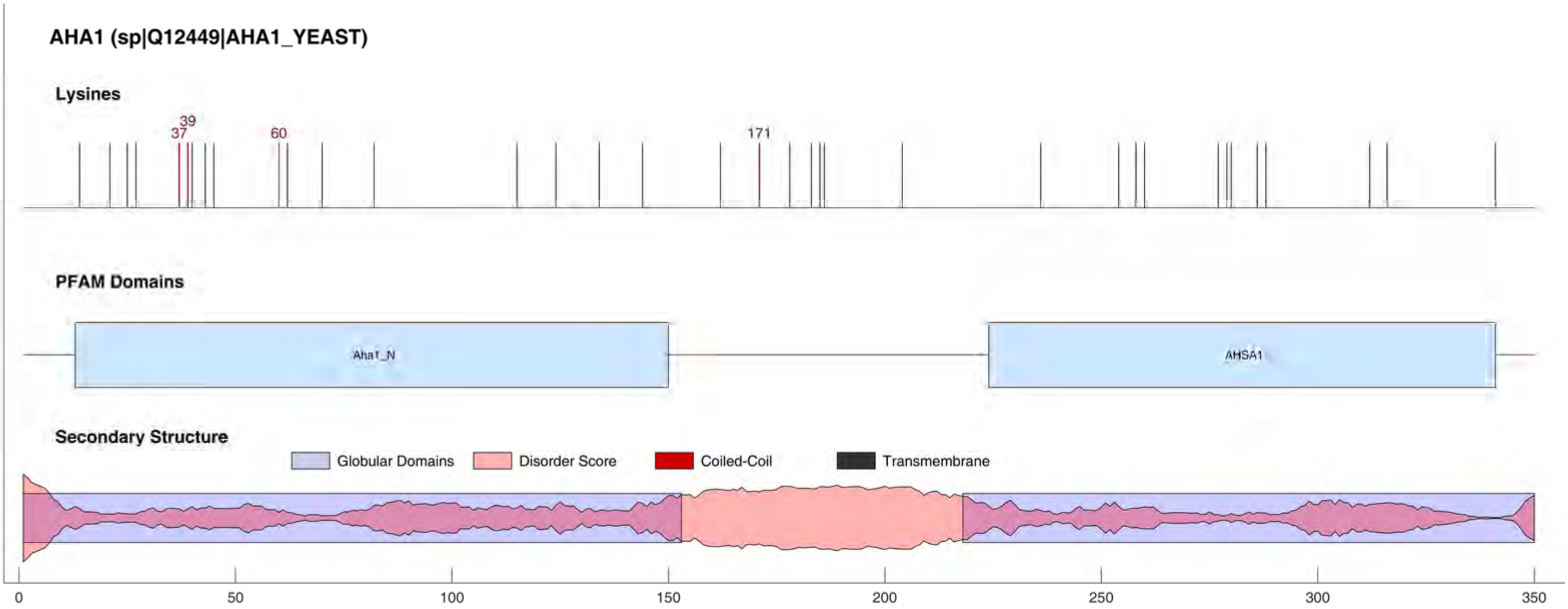
AHA1 (sp|Q12449|AHA1_YEAST)
Lysines
37
39
60
171
PFAM Domains
Aha1_N
AHSA1
Secondary Structure
Globular Domains
Disorder Score
Coiled-Coil
Transmembrane
0
50
100
150
200
250
300
350

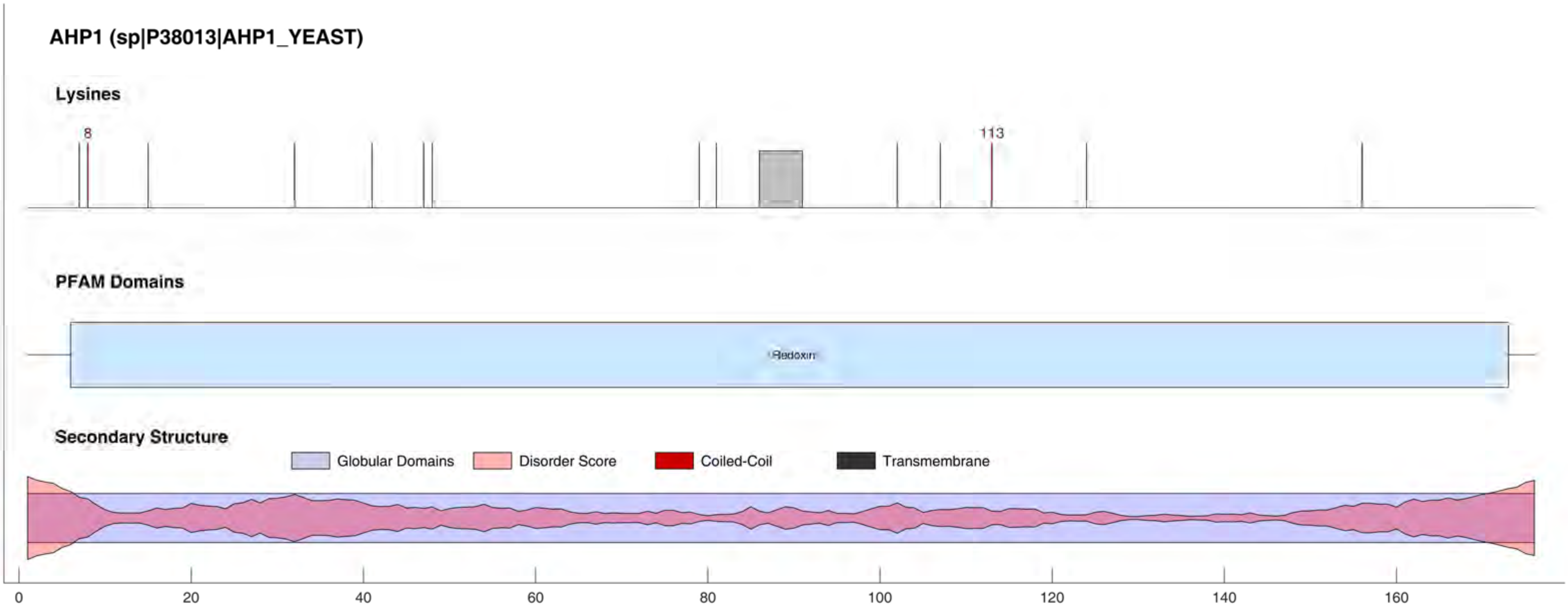
AHP1 (sp|P38013|AHP1_YEAST)
Lysines
8
113
PFAM Domains
Redoxin
Secondary Structure
Globular Domains
Disorder Score
Coiled-Coil
Transmembrane
0
20
40
60
80
100
120
140
160

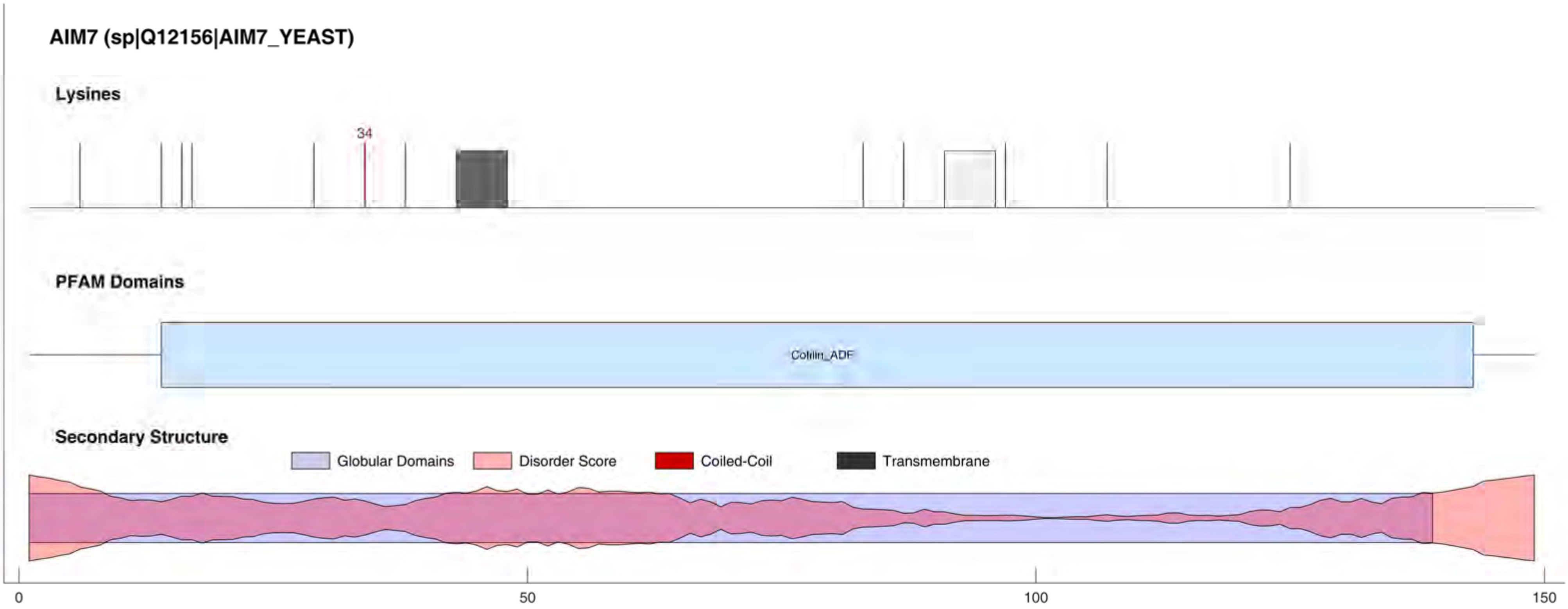
AIM7 (sp|Q12156|AIM7_YEAST)
Lysines
34
PFAM Domains
Cofilin_ADF
Secondary Structure
Globular Domains
Disorder Score
Coiled-Coil
Transmembrane
0
50
100
150

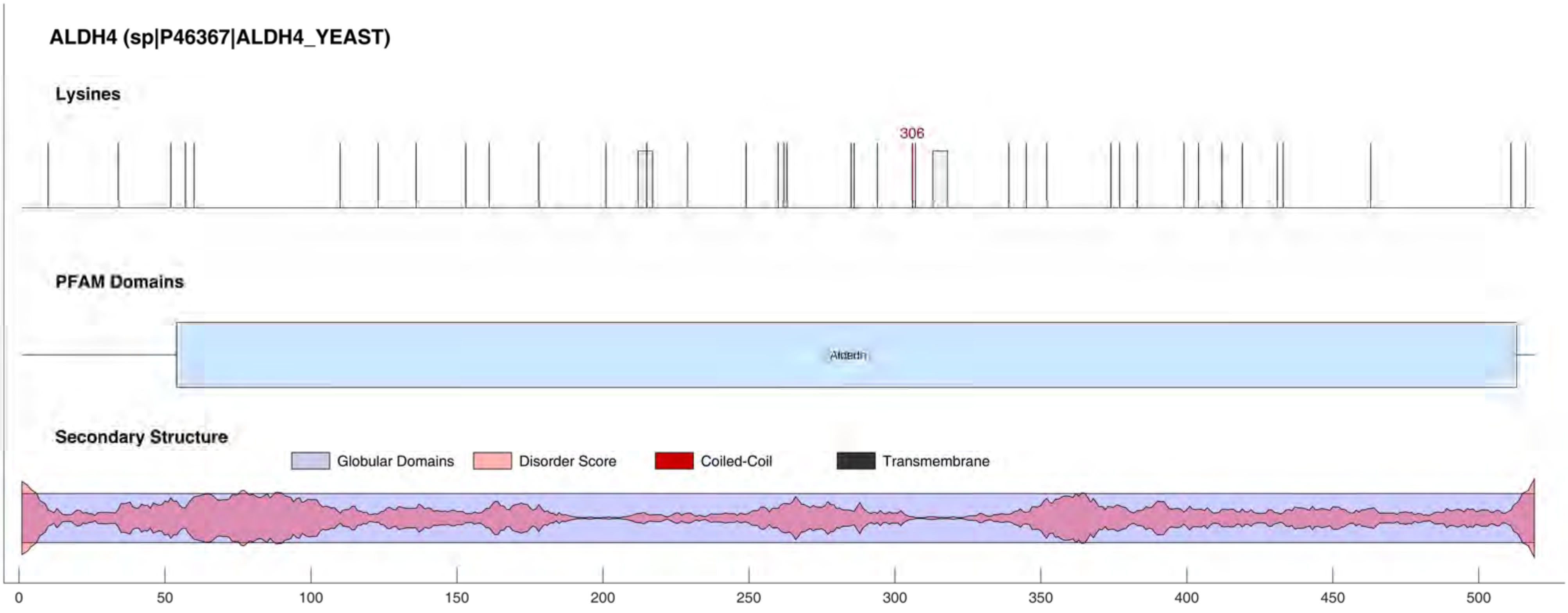
ALDH4 (sp|P46367|ALDH4_YEAST)
Lysines
306
PFAM Domains
Aldedh
Secondary Structure
Globular Domains
Disorder Score
Coiled-Coil
Transmembrane
0
50
100
150
200
250
300
350
400
450
500

**ALDH6 (sp|P54115|ALDH6_YEAST)**

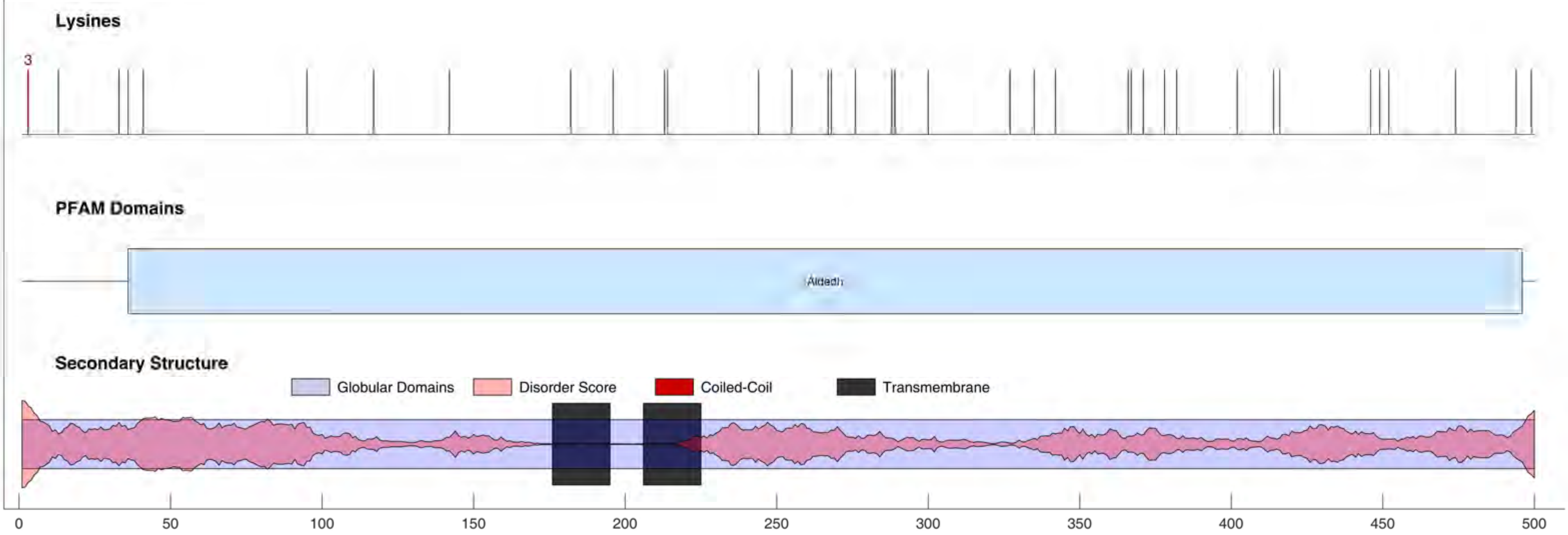

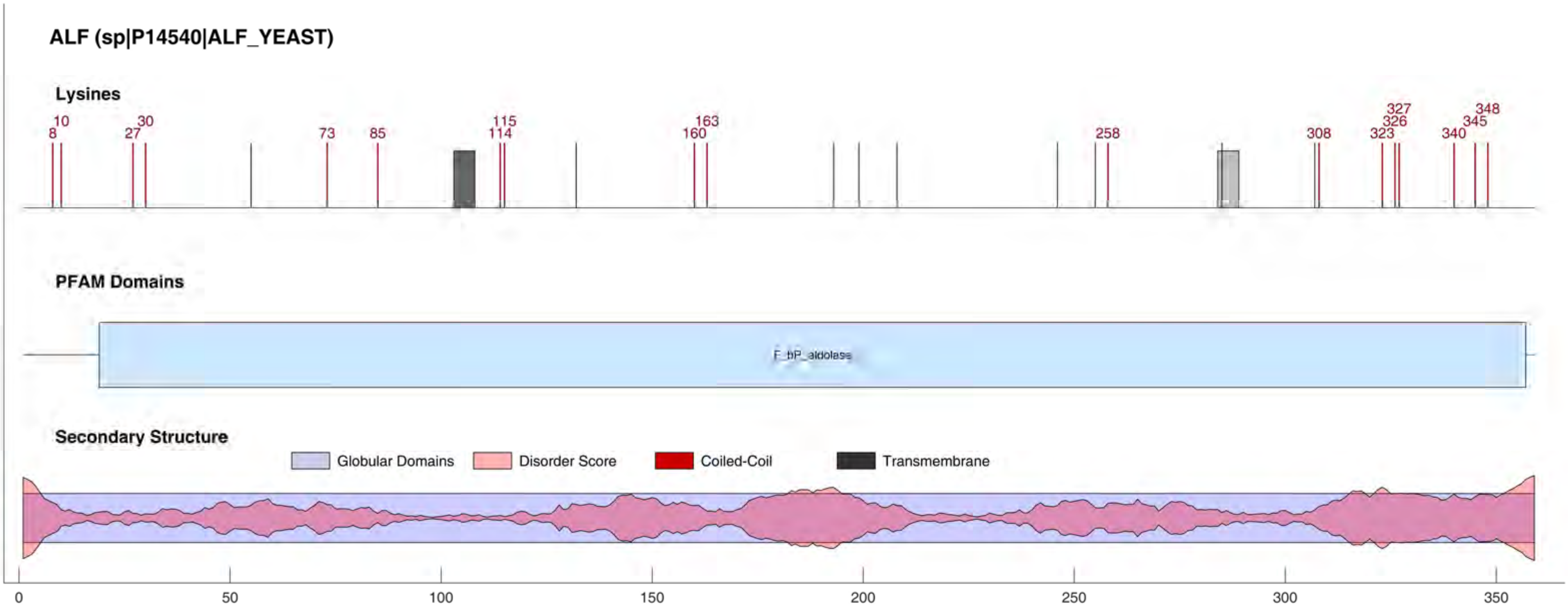
ALF (sp|P14540|ALF_YEAST)
Lysines
8
10
27
30
73
85
114
115
160
163
258
308
323
326
327
340
345
348
PFAM Domains
F_bP_aldolase
Secondary Structure
Globular Domains
Disorder Score
Coiled-Coil
Transmembrane
0
50
100
150
200
250
300
350

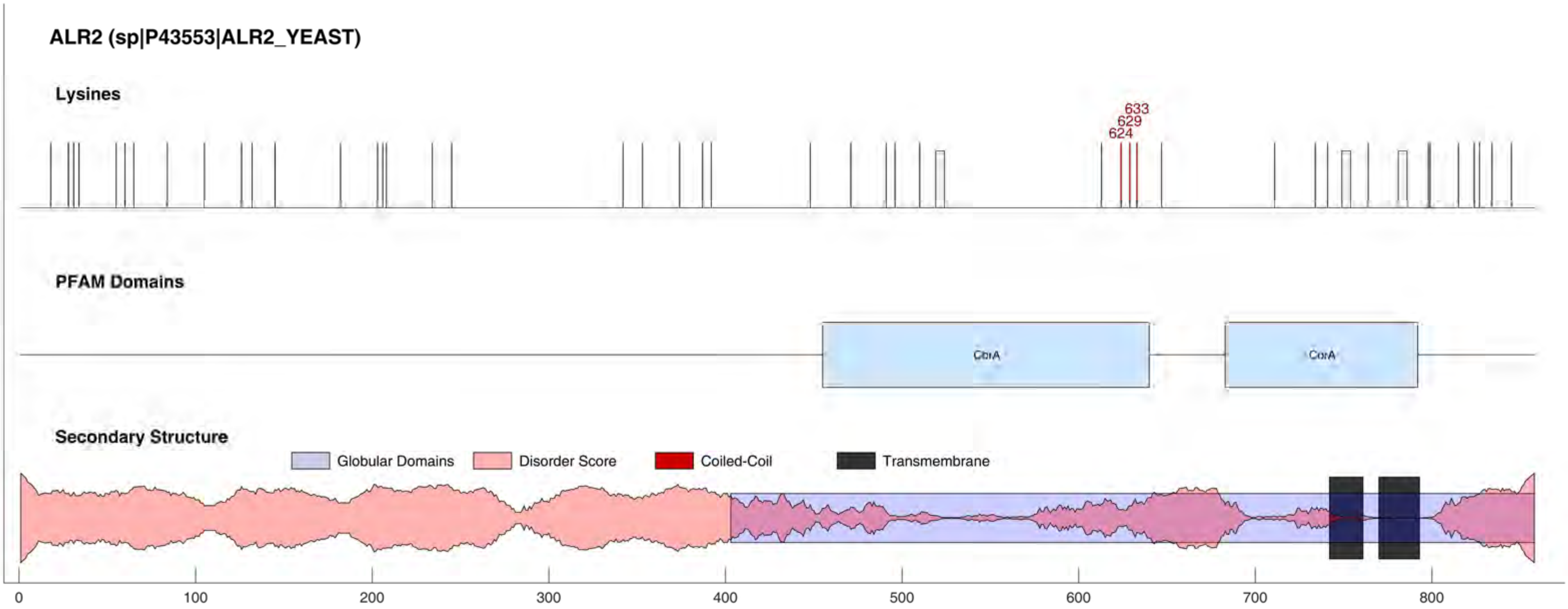
ALR2 (sp|P43553|ALR2_YEAST)
Lysines
633
629
624
PFAM Domains
CorA
CorA
Secondary Structure
Globular Domains
Disorder Score
Coiled-Coil
Transmembrane
0
100
200
300
400
500
600
700
800

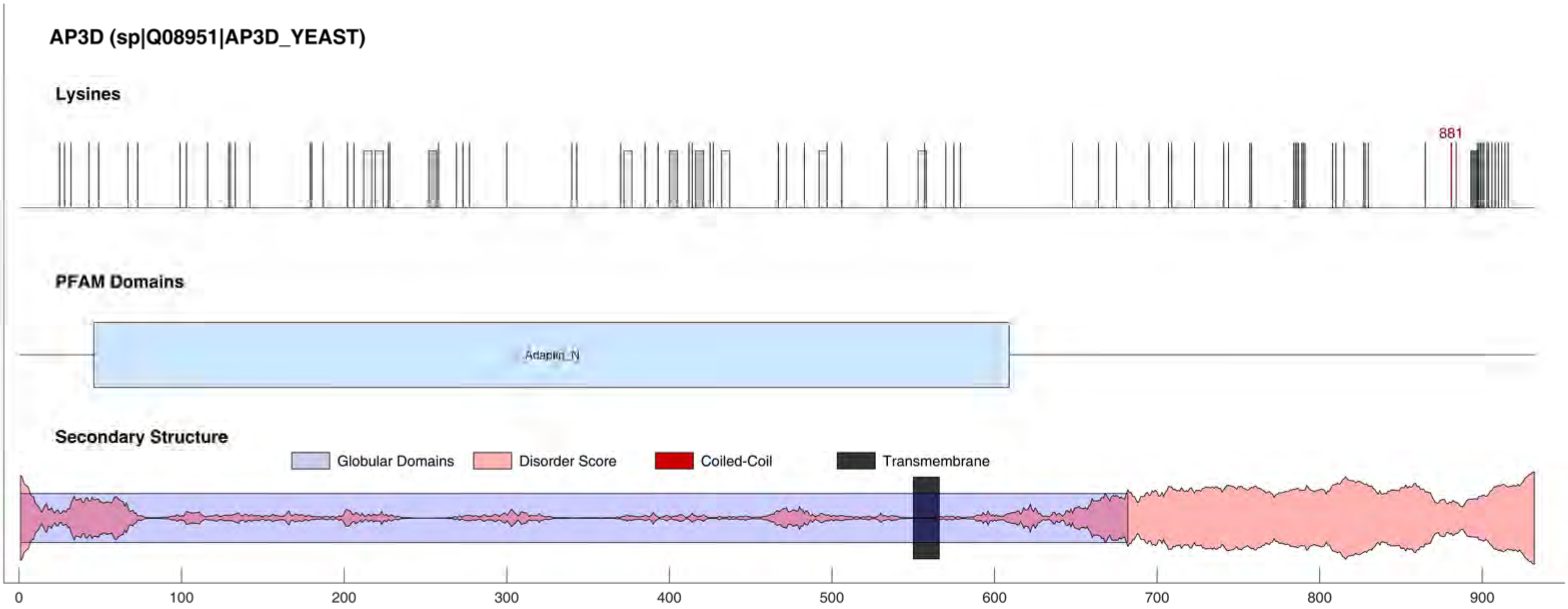
AP3D (sp|Q08951|AP3D_YEAST)
Lysines
881
PFAM Domains
Adaptin_N
Secondary Structure
Globular Domains
Disorder Score
Coiled-Coil
Transmembrane
0
100
200
300
400
500
600
700
800
900

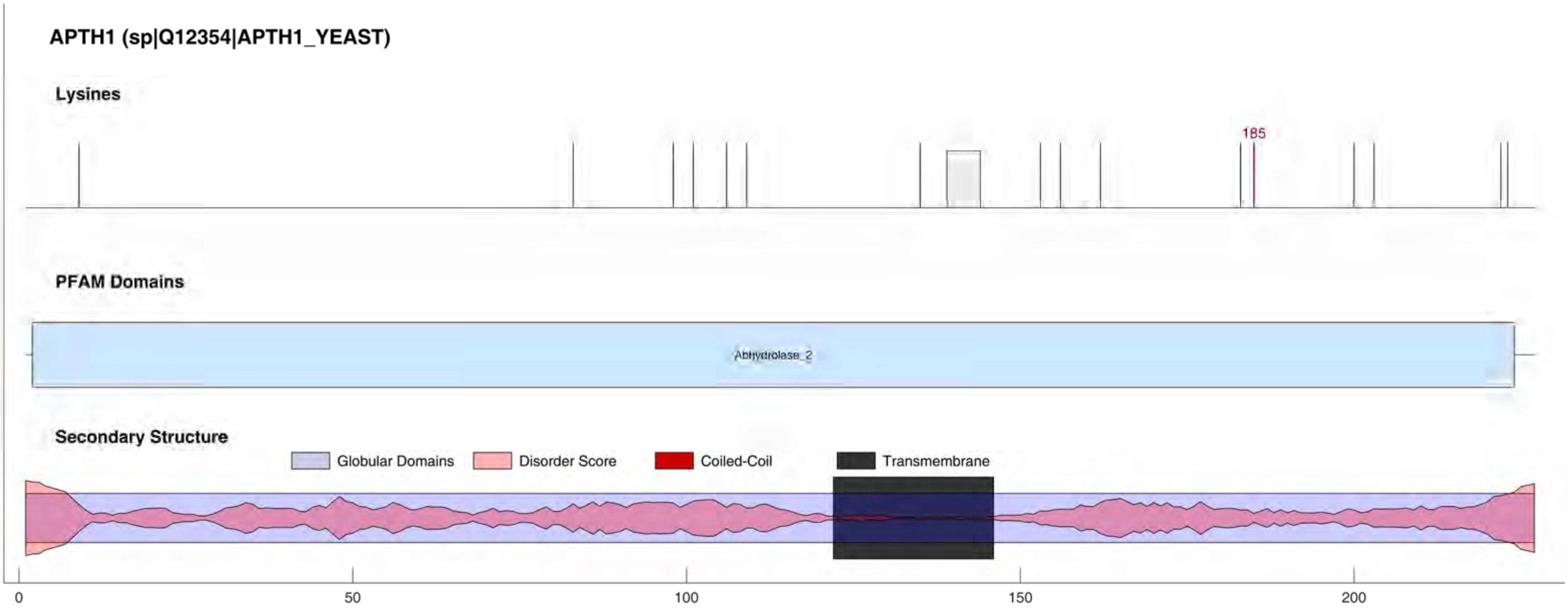
APTH1 (sp|Q12354|APTH1_YEAST)
Lysines
185
PFAM Domains
Abhydrolase_2
Secondary Structure
Globular Domains
Disorder Score
Coiled-Coil
Transmembrane
0
50
100
150
200

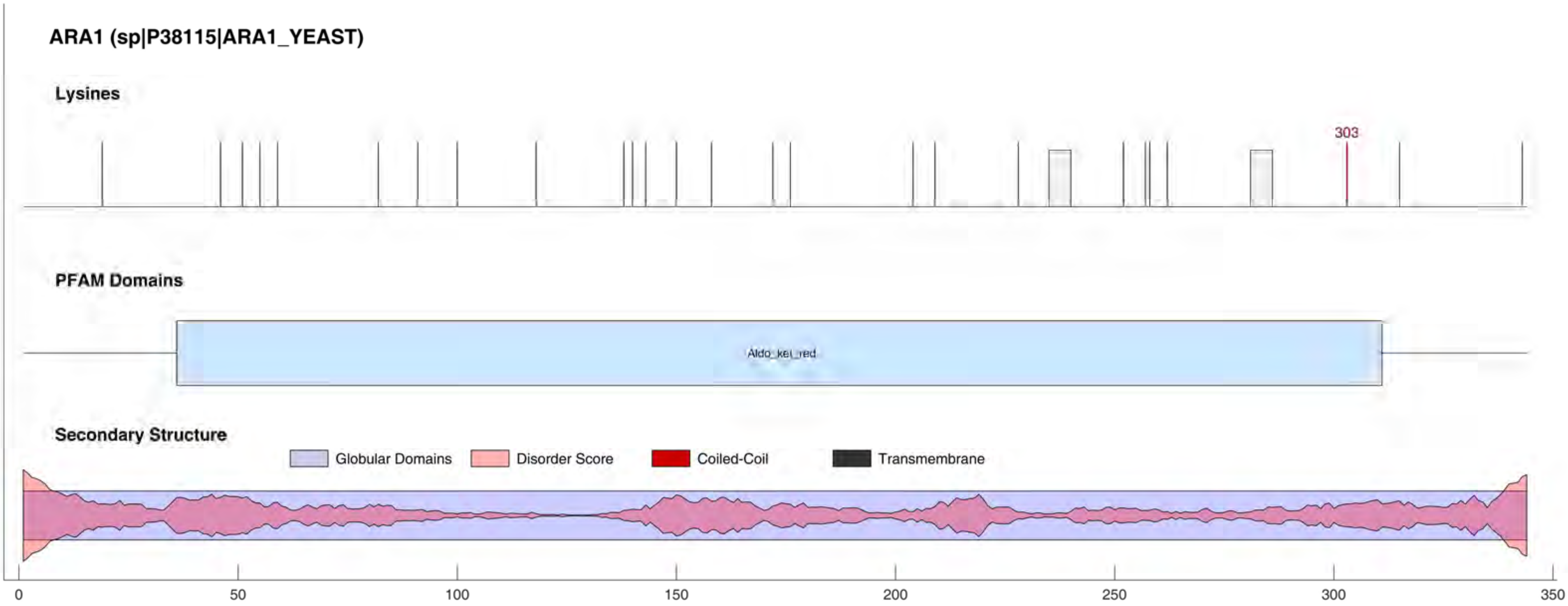
ARA1 (sp|P38115|ARA1_YEAST)
Lysines
303
PFAM Domains
Aldo_ket_red
Secondary Structure
Globular Domains
Disorder Score
Coiled-Coil
Transmembrane
0
50
100
150
200
250
300
350

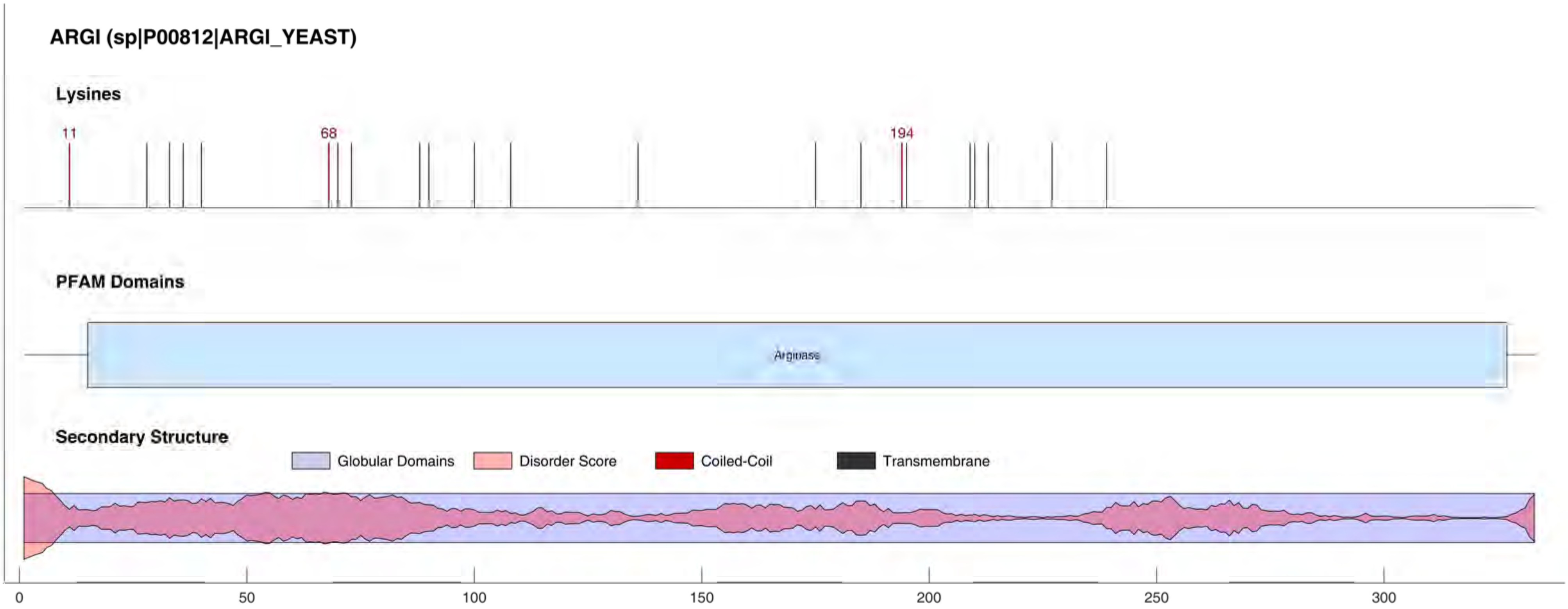
ARGI (sp|P00812|ARGI_YEAST)
Lysines
11
68
194
PFAM Domains
Arginase
Secondary Structure
Globular Domains
Disorder Score
Coiled-Coil
Transmembrane
0
50
100
150
200
250
300

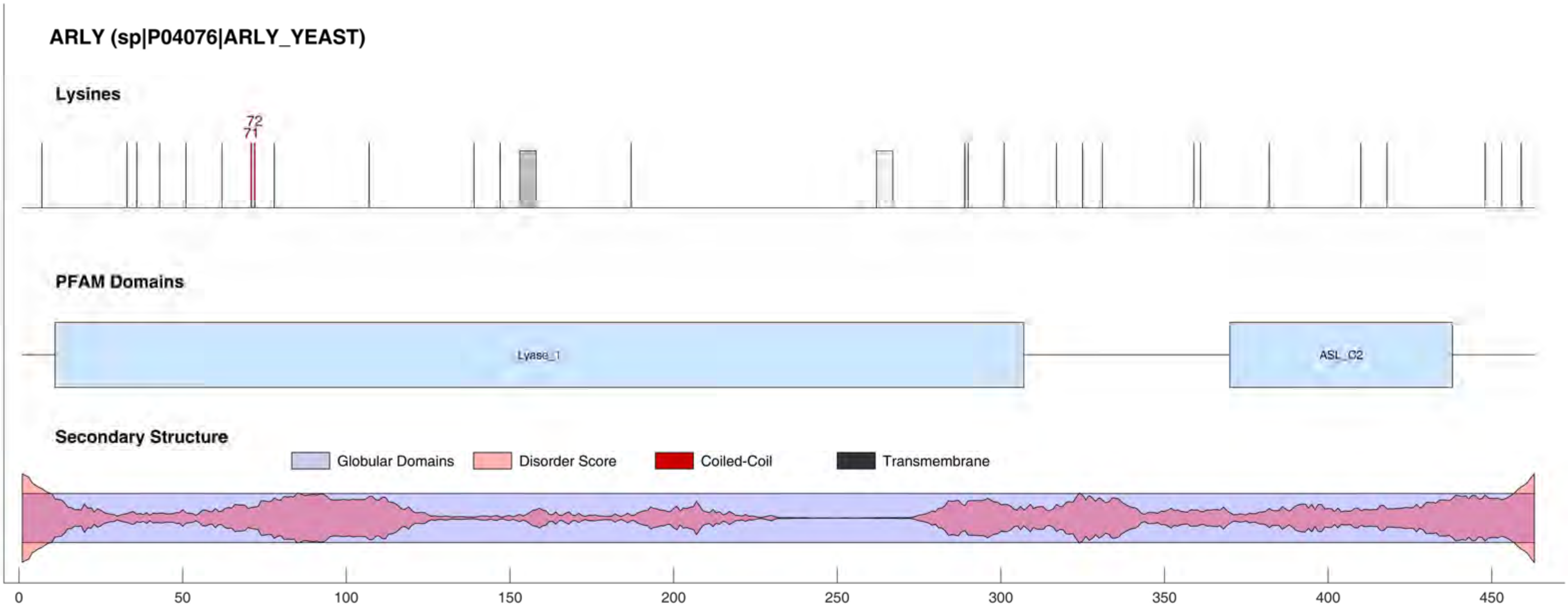
ARLY (sp|P04076|ARLY_YEAST)
Lysines
72
71
PFAM Domains
Lyase_1
ASL_C2
Secondary Structure
Globular Domains
Disorder Score
Coiled-Coil
Transmembrane
0
50
100
150
200
250
300
350
400
450

#### ARMT1 (sp|Q04371|ARMT1_YEAST)

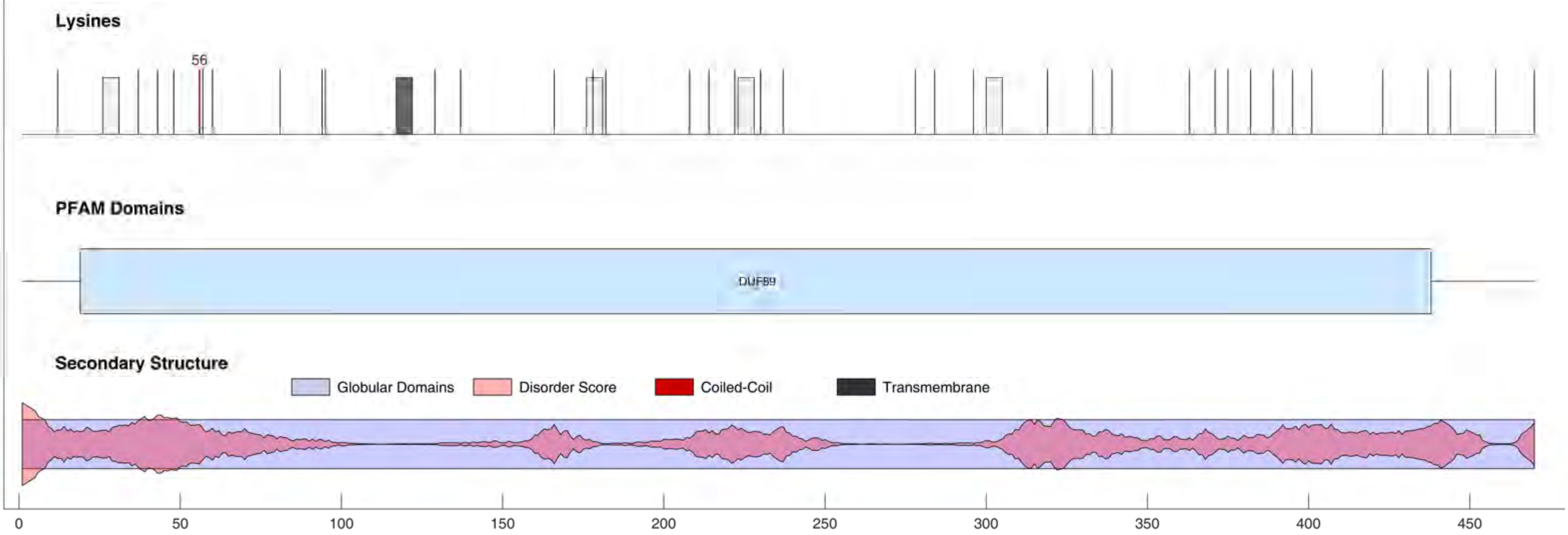

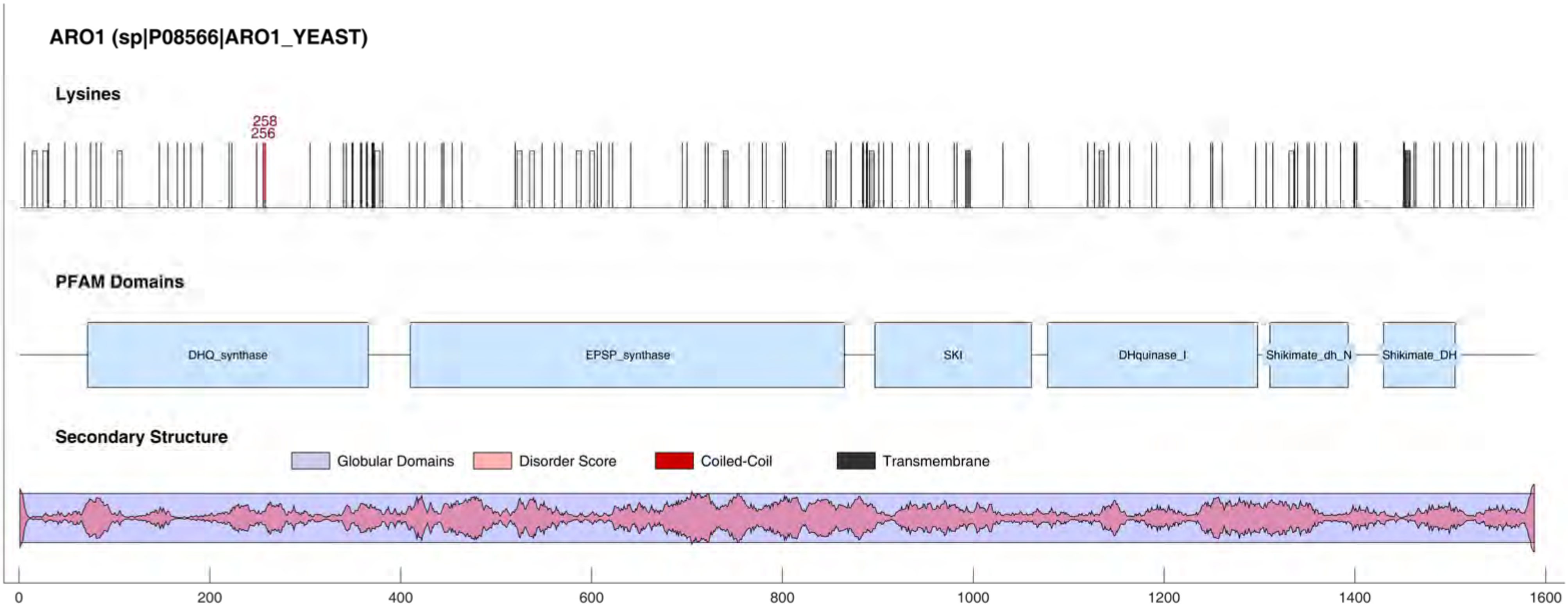
ARO1 (sp|P08566|ARO1_YEAST)
Lysines
258
256
PFAM Domains
DHQ_synthase
EPSP_synthase
SKI
DHquinase_I
Shikimate_dh_N
Shikimate_DH
Secondary Structure
Globular Domains
Disorder Score
Coiled-Coil
Transmembrane
0
200
400
600
800
1000
1200
1400
1600

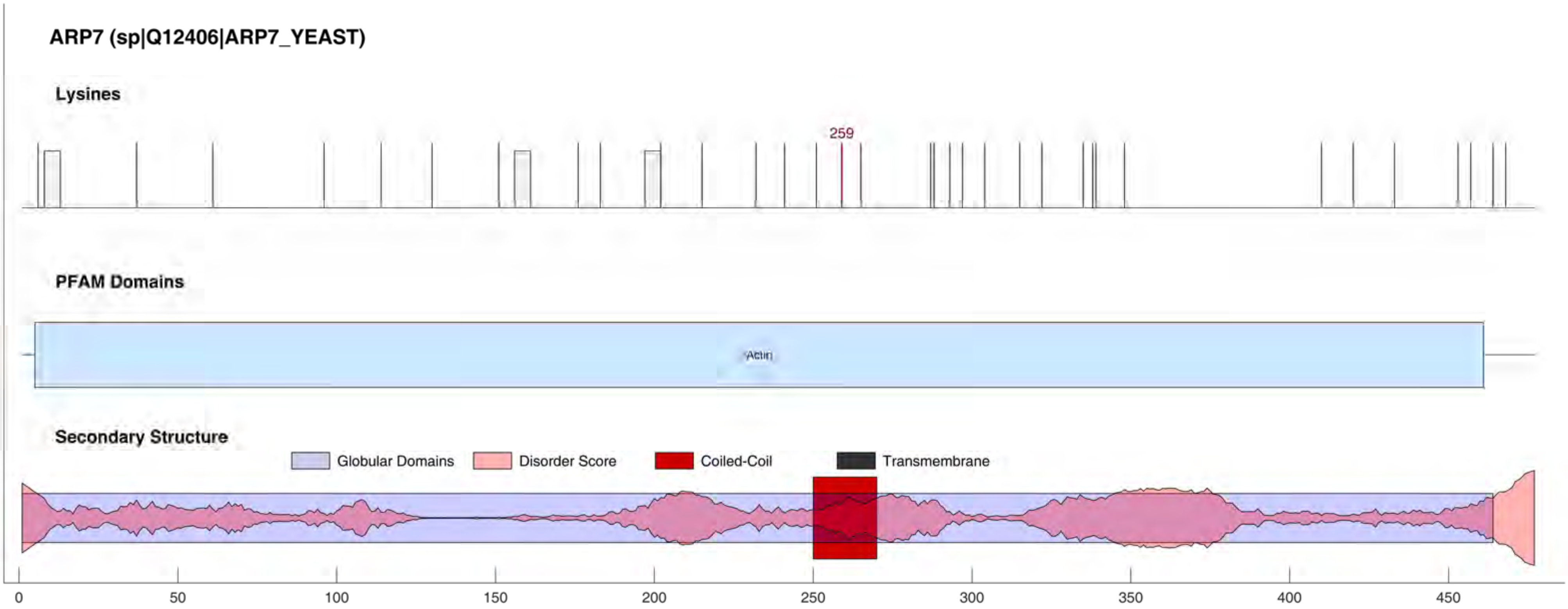
ARP7 (sp|Q12406|ARP7_YEAST)
Lysines
259
PFAM Domains
Actin
Secondary Structure
Globular Domains
Disorder Score
Coiled-Coil
Transmembrane
0
50
100
150
200
250
300
350
400
450

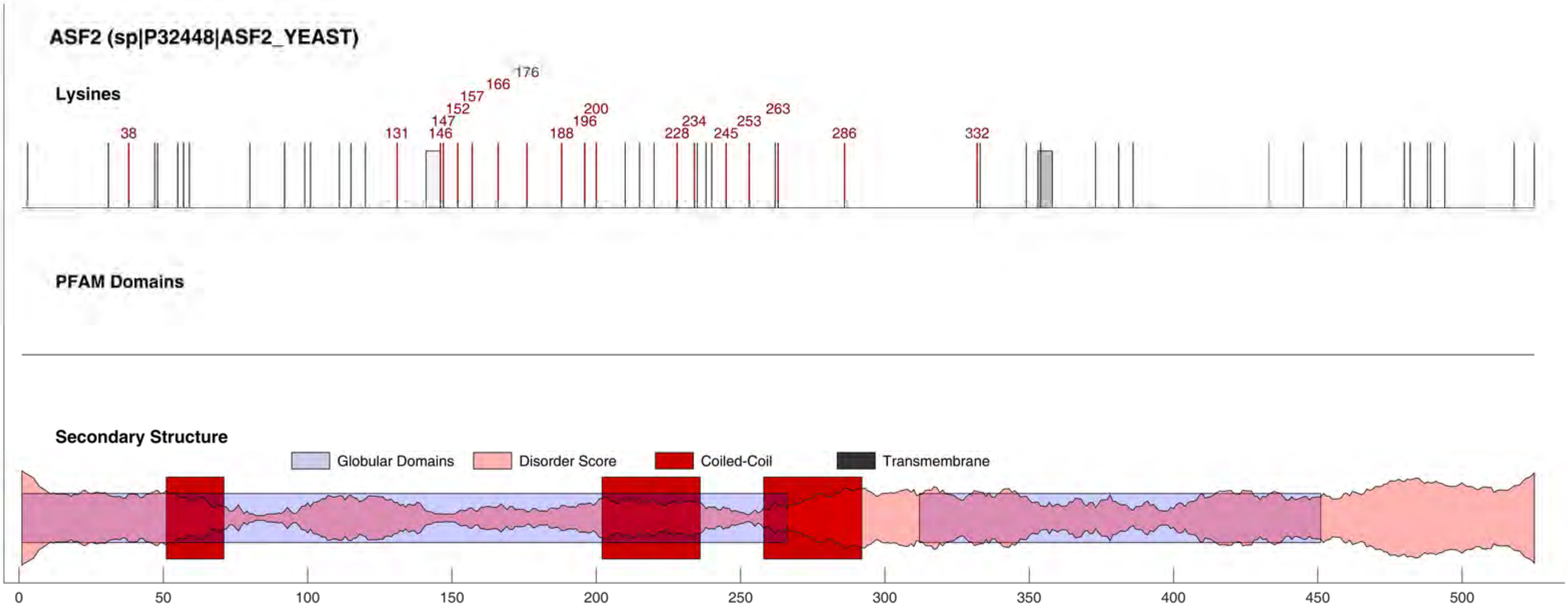
ASF2 (sp|P32448|ASF2_YEAST)
Lysines
38
131
146
147
152
157
166
176
188
196
200
228
234
245
253
263
286
332
PFAM Domains
Secondary Structure
Globular Domains
Disorder Score
Coiled-Coil
Transmembrane
0
50
100
150
200
250
300
350
400
450
500

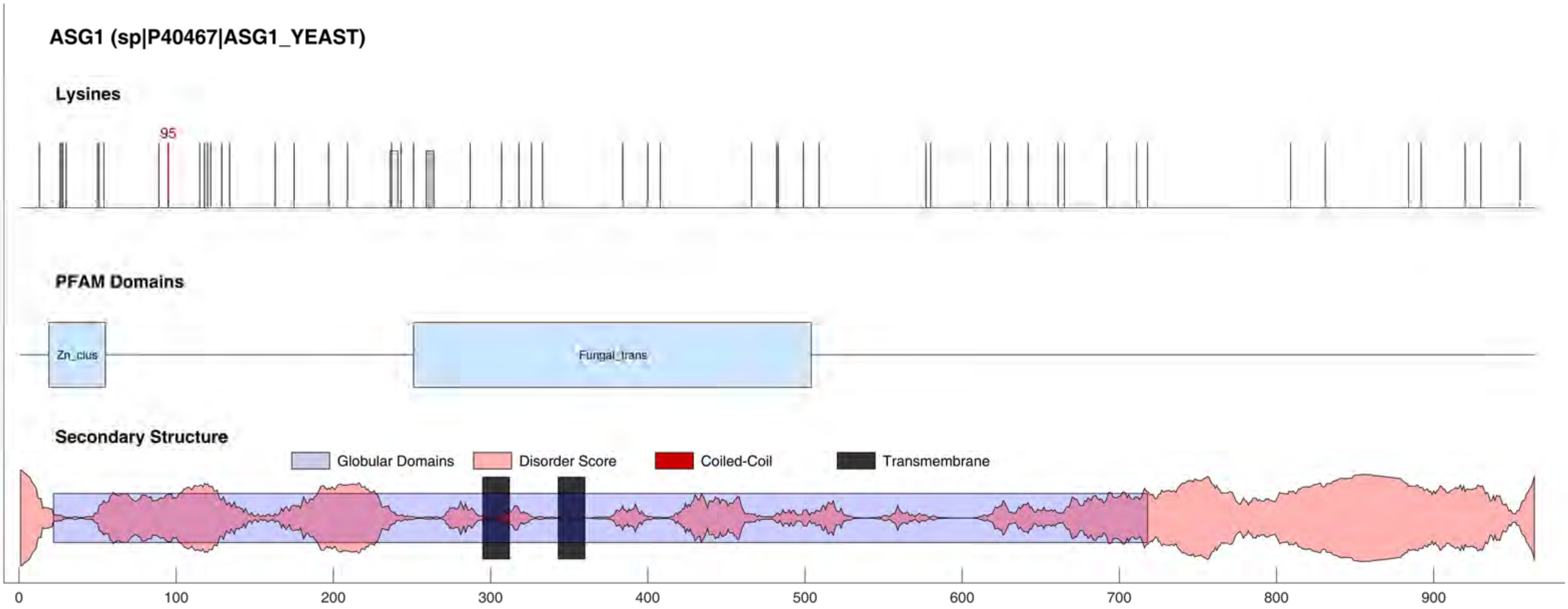

ASG1 (sp|P40467|ASG1_YEAST)
Lysines
95
PFAM Domains
Zn_clus
Fungal_trans
Secondary Structure
Globular Domains
Disorder Score
Coiled-Coil
Transmembrane
0
100
200
300
400
500
600
700
800
900

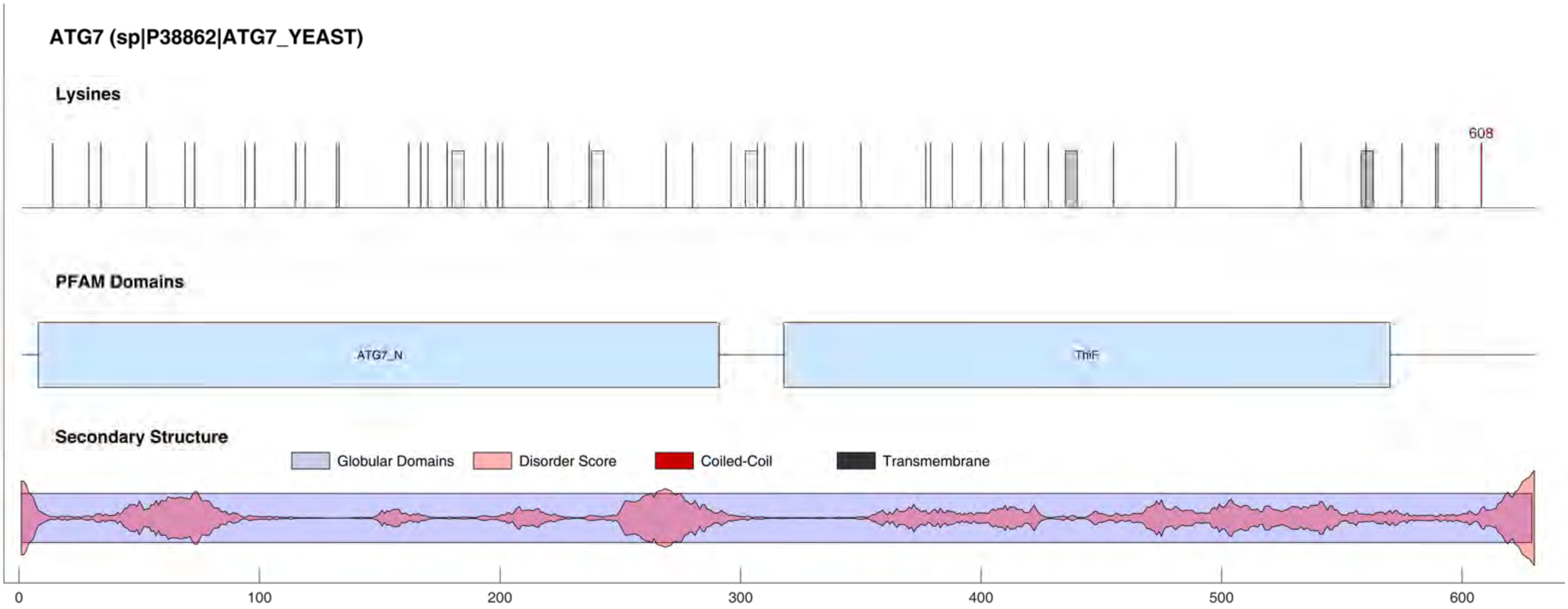
ATG7 (sp|P38862|ATG7_YEAST)
Lysines
608
PFAM Domains
ATG7_N
ThiF
Secondary Structure
Globular Domains
Disorder Score
Coiled-Coil
Transmembrane
0
100
200
300
400
500
600

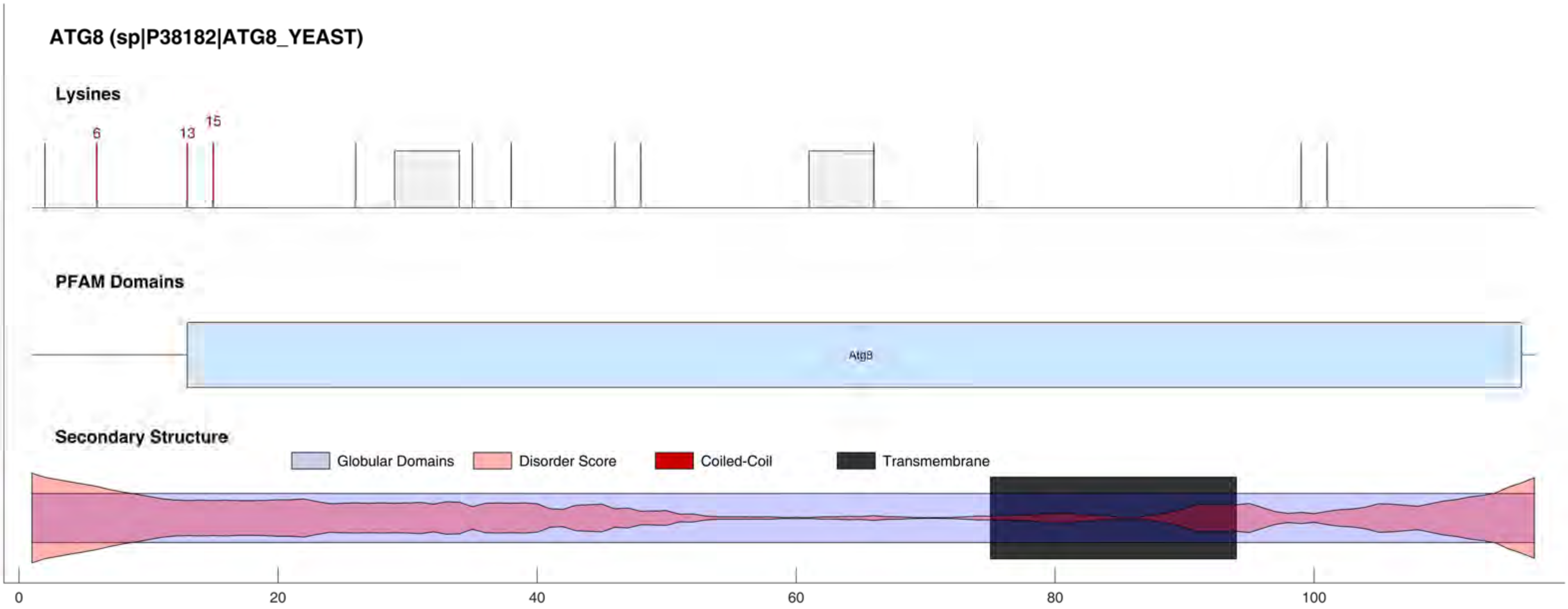
ATG8 (sp|P38182|ATG8_YEAST)
Lysines
6
13
15
PFAM Domains
Atg8
Secondary Structure
Globular Domains
Disorder Score
Coiled-Coil
Transmembrane
0
20
40
60
80
100

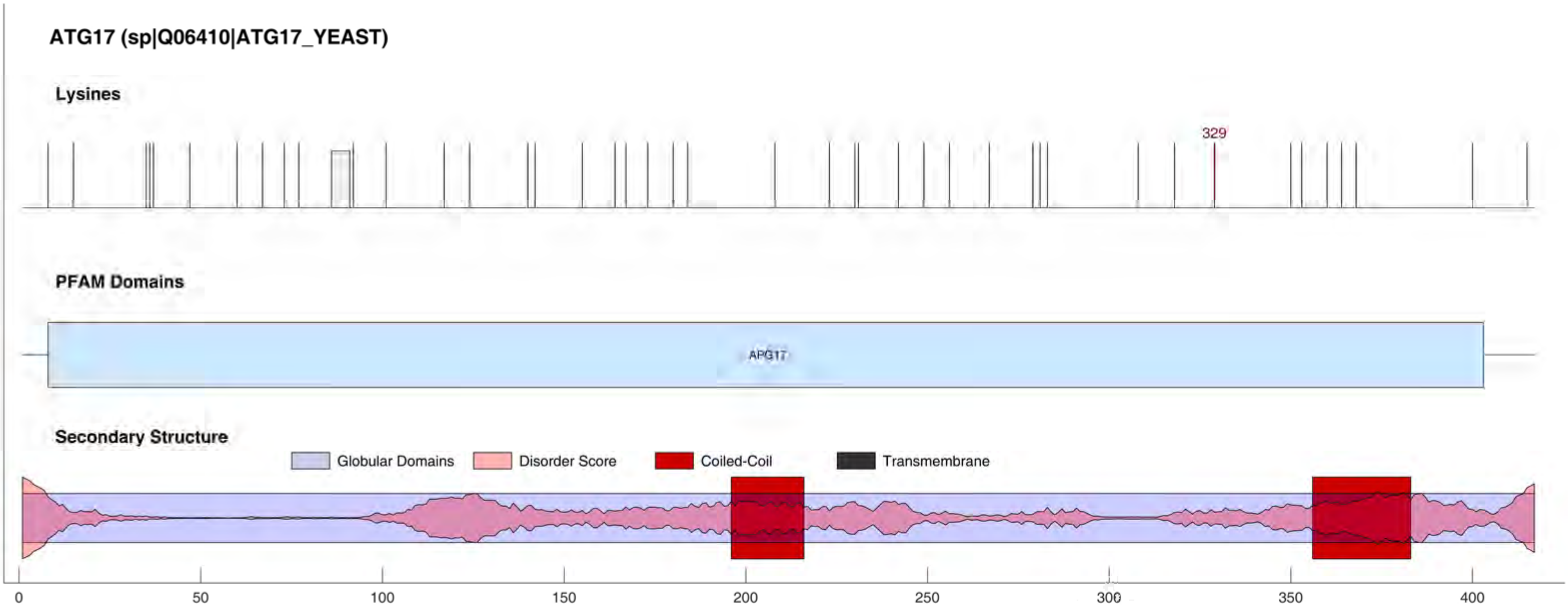
ATG17 (sp|Q06410|ATG17_YEAST)
Lysines
329
PFAM Domains
APG17
Secondary Structure
Globular Domains
Disorder Score
Coiled-Coil
Transmembrane
0
50
100
150
200
250
300
350
400

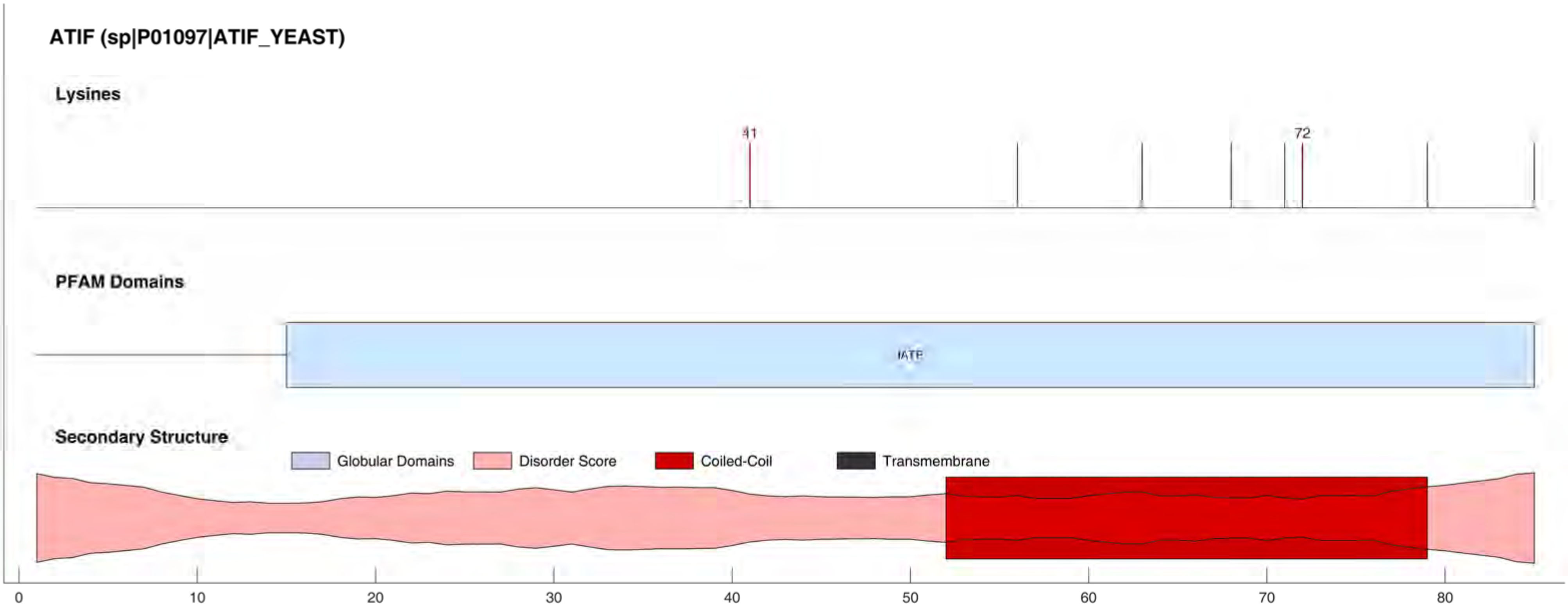
ATIF (sp|P01097|ATIF_YEAST)
Lysines
41
72
PFAM Domains
IATP
Secondary Structure
Globular Domains
Disorder Score
Coiled-Coil
Transmembrane
0
10
20
30
40
50
60
70
80

ATP7 (sp|P30902|ATP7_YEAST)

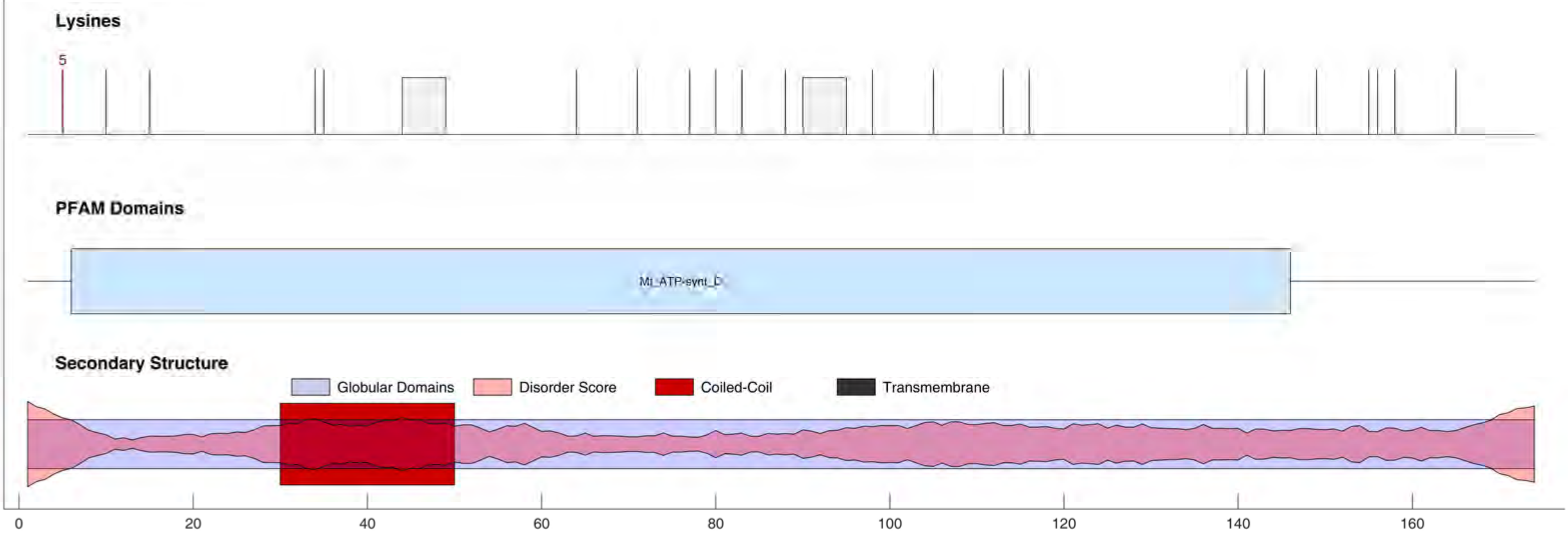

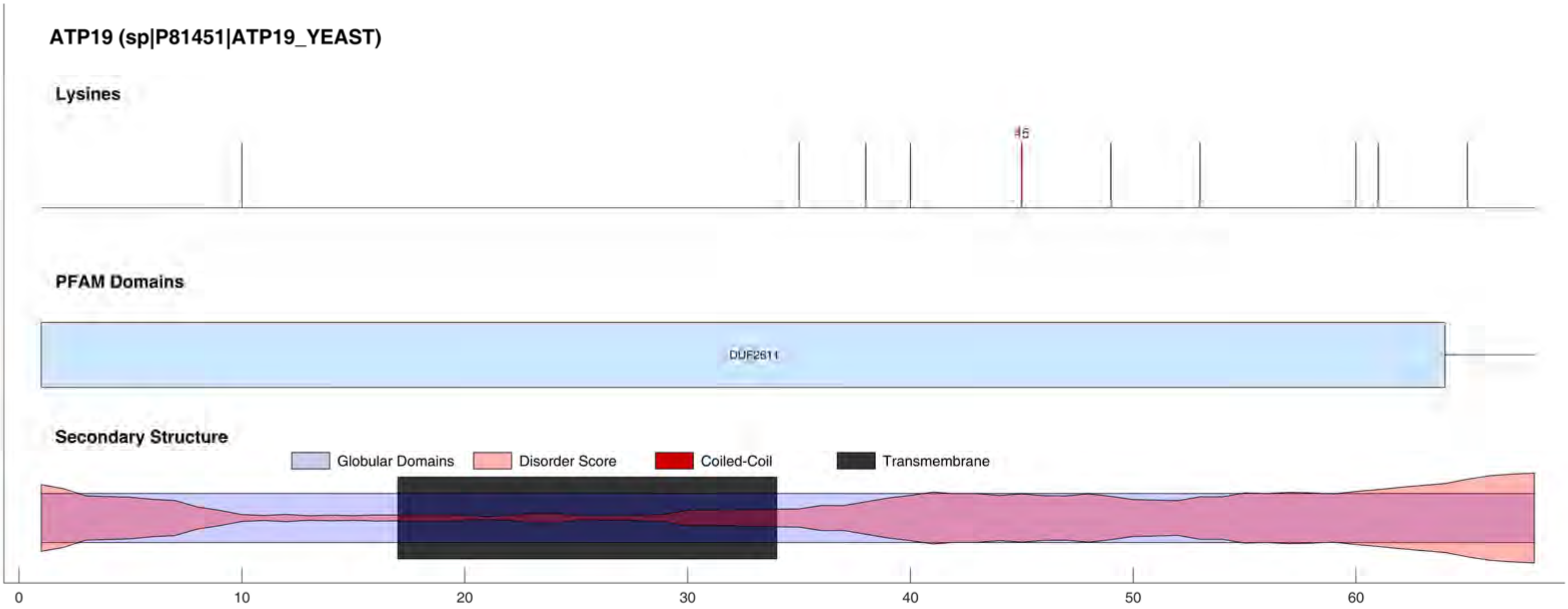
ATP19 (sp|P81451|ATP19_YEAST)
Lysines
45
PFAM Domains
DUF2611
Secondary Structure
Globular Domains
Disorder Score
Coiled-Coil
Transmembrane
0
10
20
30
40
50
60

#### ATPK (sp|Q06405|ATPK_YEAST)

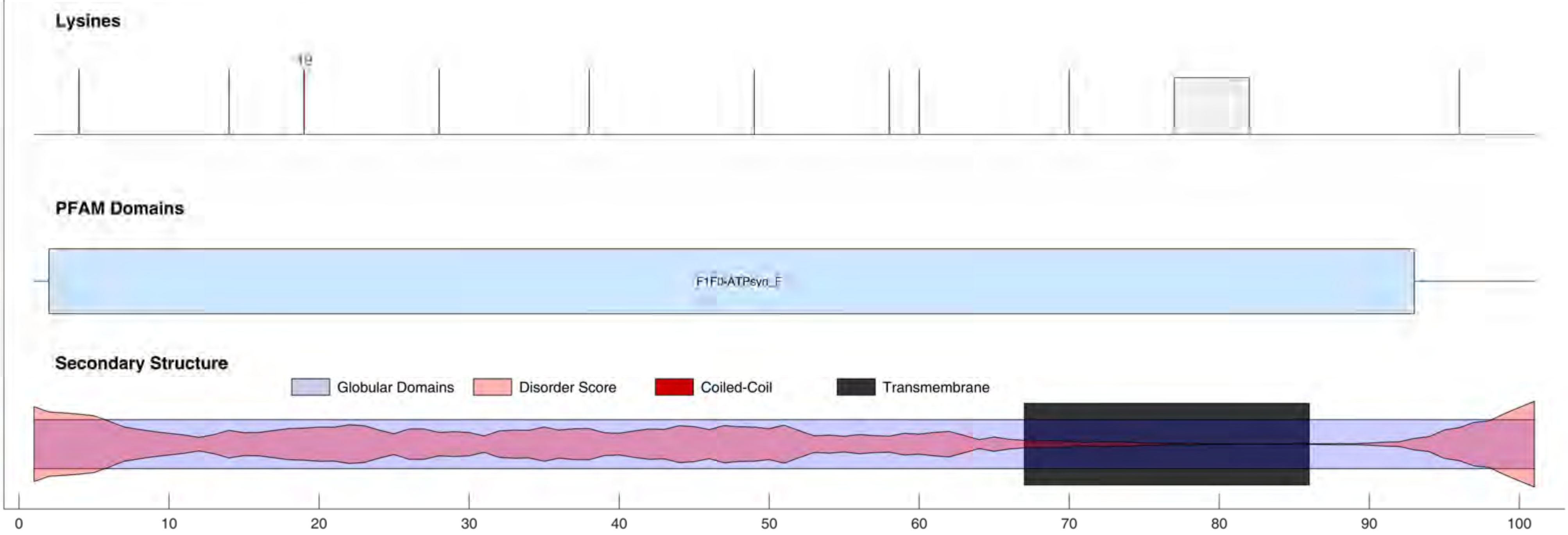

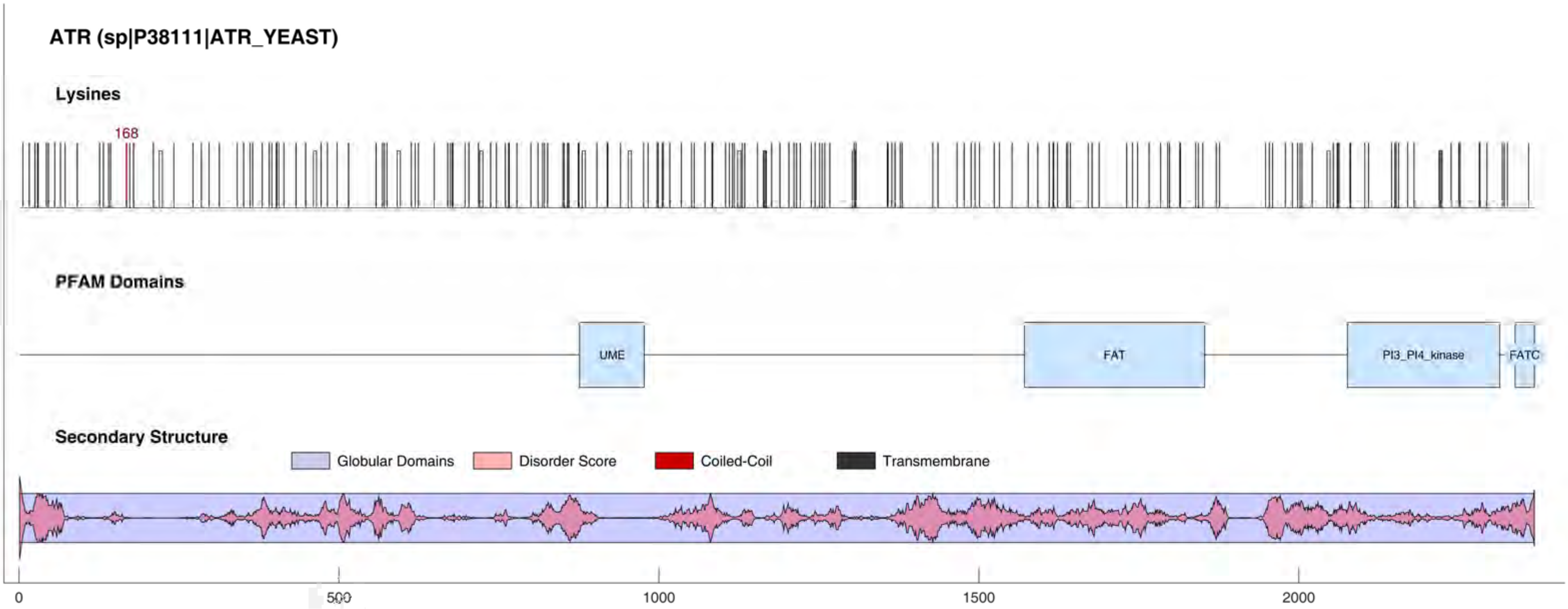
ATR (sp|P38111|ATR_YEAST)
Lysines
168
PFAM Domains
UME
FAT
PI3_PI4_kinase
FATC
Secondary Structure
Globular Domains
Disorder Score
Coiled-Coil
Transmembrane
0
500
1000
1500
2000

#### ATX1 (sp|P38636|ATX1_YEAST)

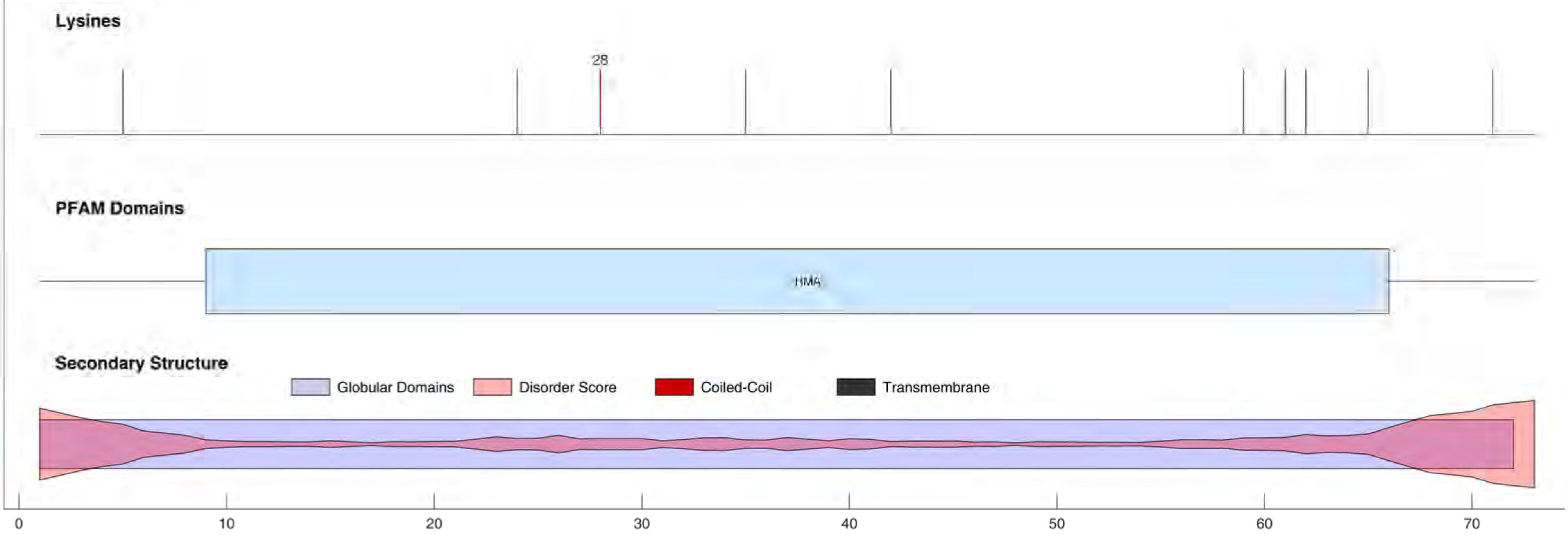

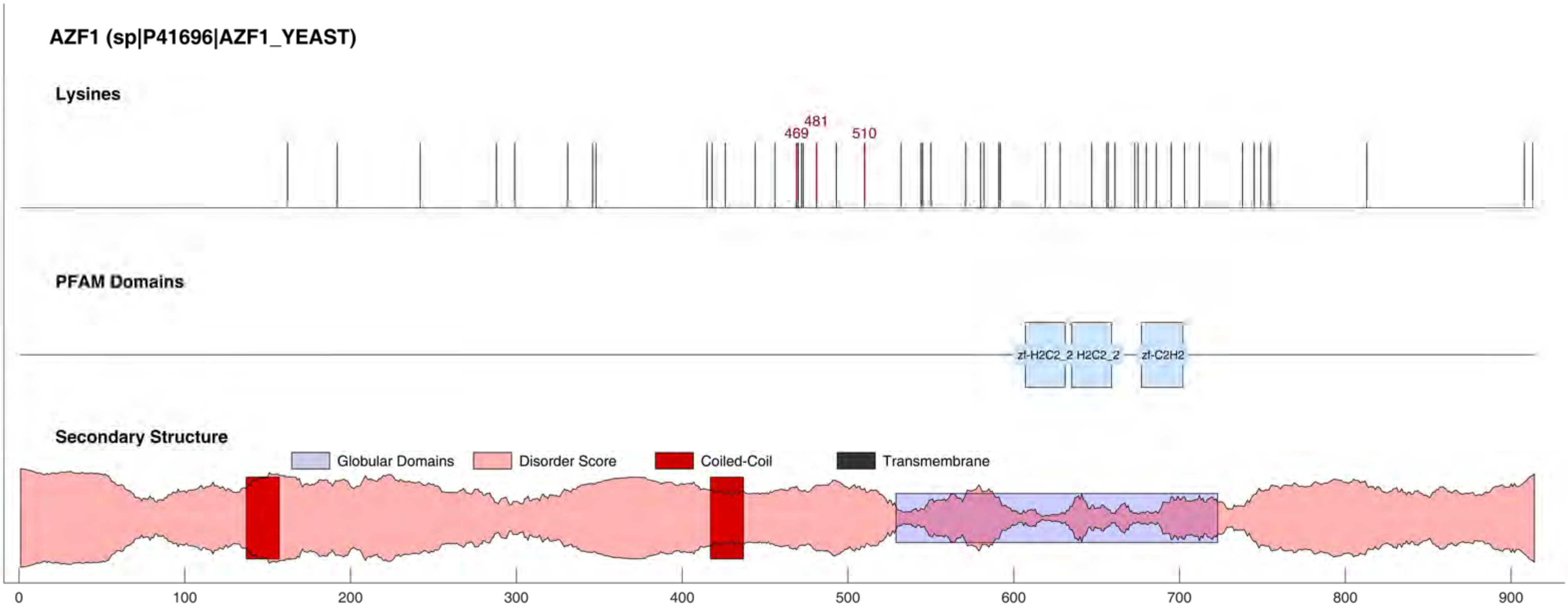
AZF1 (sp|P41696|AZF1_YEAST)
Lysines
469
481
510
PFAM Domains
zf-H2C2_2
H2C2_2
zf-C2H2
Secondary Structure
Globular Domains
Disorder Score
Coiled-Coil
Transmembrane
0
100
200
300
400
500
600
700
800
900

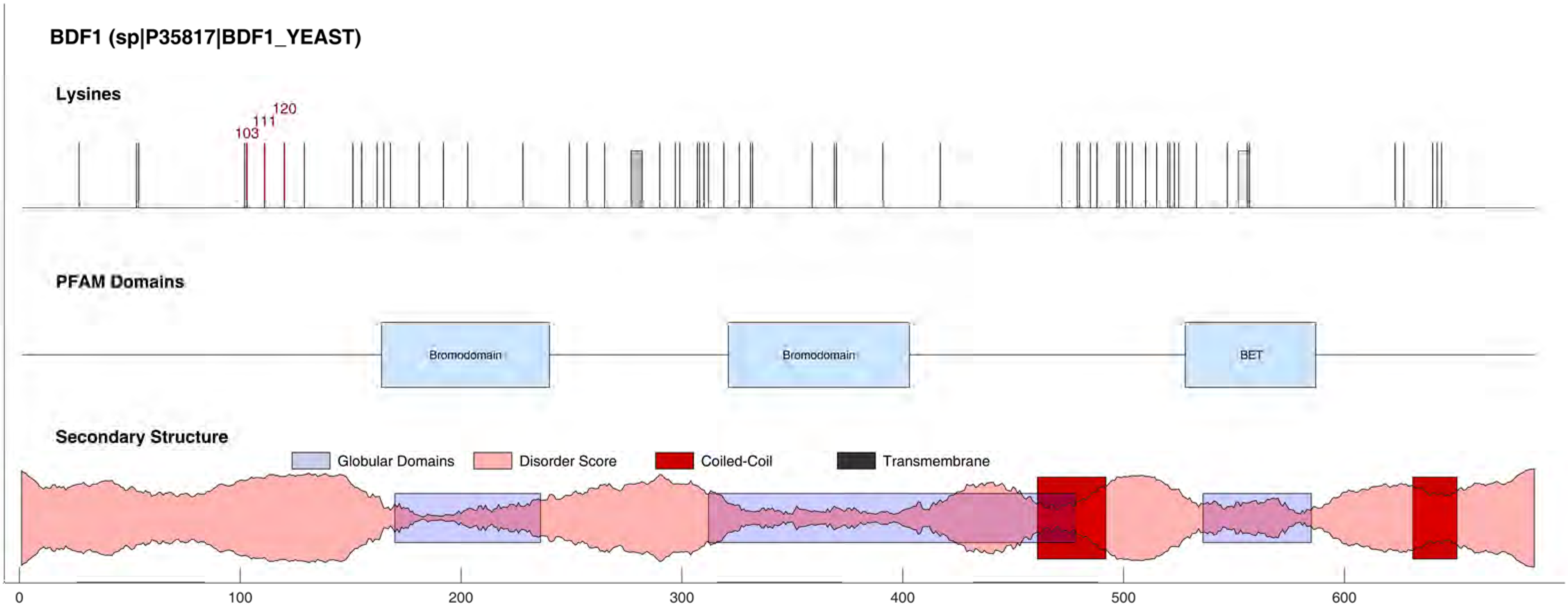
BDF1 (sp|P35817|BDF1_YEAST)
Lysines
103
111
120
PFAM Domains
Bromodomain
Bromodomain
BET
Secondary Structure
Globular Domains
Disorder Score
Coiled-Coil
Transmembrane
0
100
200
300
400
500
600

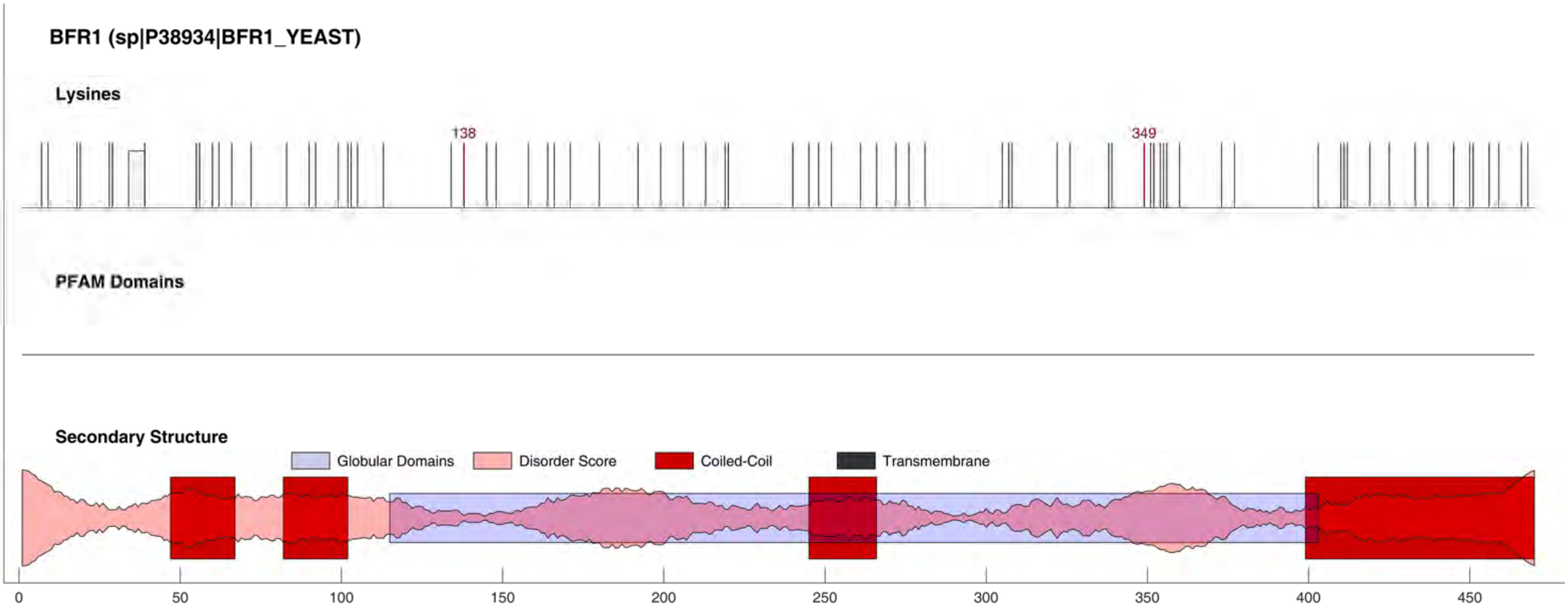
BFR1 (sp|P38934|BFR1_YEAST)
Lysines
138
349
PFAM Domains
Secondary Structure
Globular Domains
Disorder Score
Coiled-Coil
Transmembrane
0
50
100
150
200
250
300
350
400
450

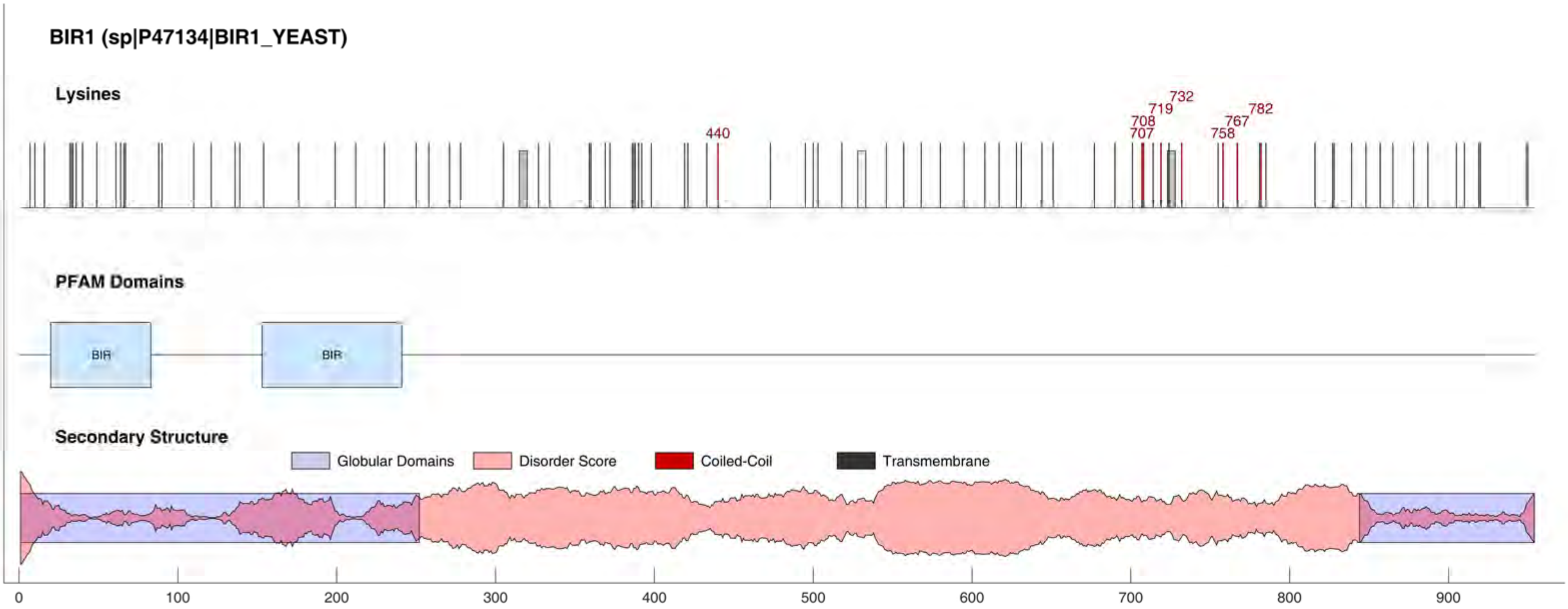
BIR1 (sp|P47134|BIR1_YEAST)
Lysines
440
707
708
719
732
758
767
782
PFAM Domains
BIR
BIR
Secondary Structure
Globular Domains
Disorder Score
Coiled-Coil
Transmembrane
0
100
200
300
400
500
600
700
800
900

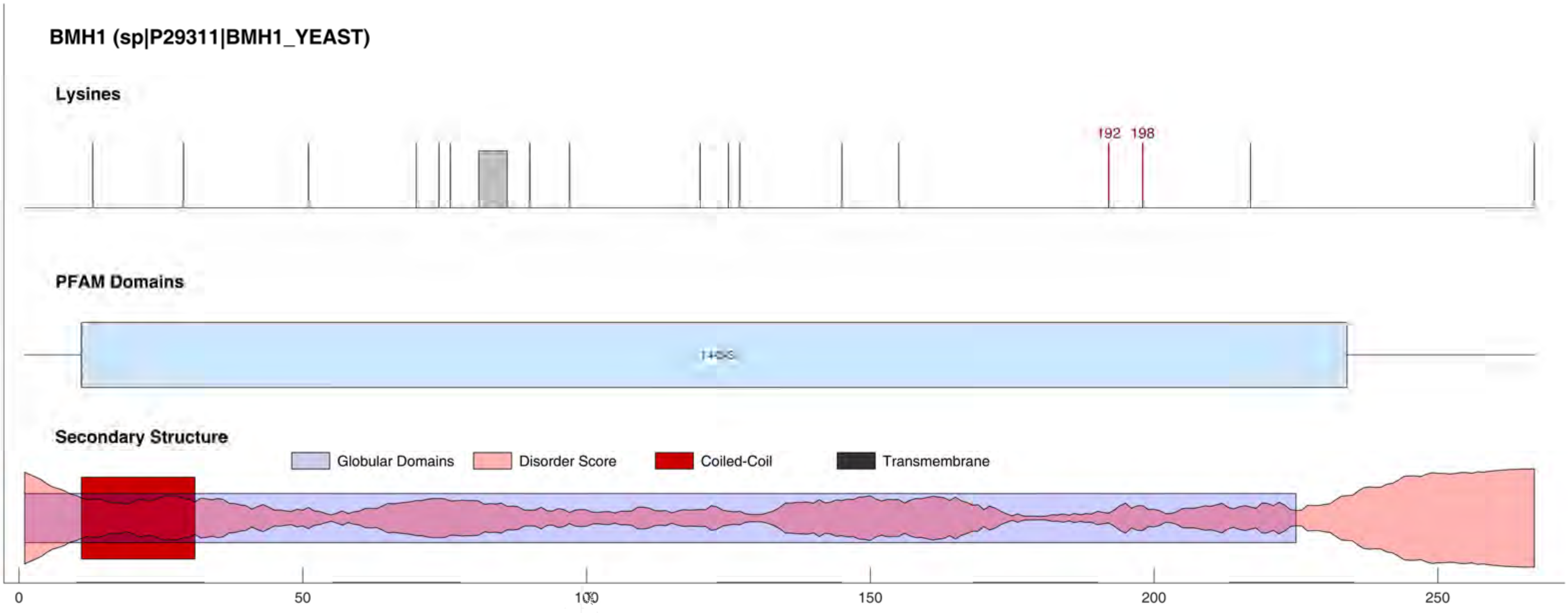
BMH1 (sp|P29311|BMH1_YEAST)
Lysines
192
198
PFAM Domains
Secondary Structure
Globular Domains
Disorder Score
Coiled-Coil
Transmembrane
0
50
100
150
200
250

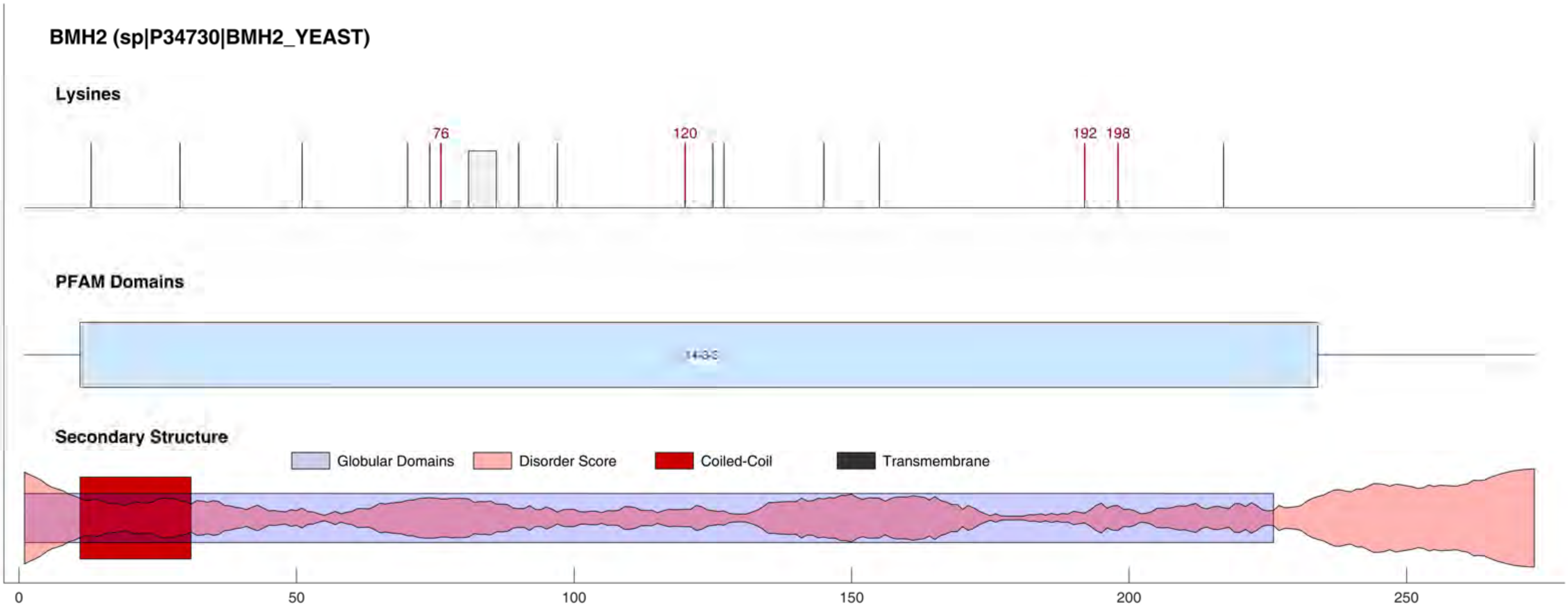
BMH2 (sp|P34730|BMH2_YEAST)
Lysines
76
120
192
198
PFAM Domains
14-3-3
Secondary Structure
Globular Domains
Disorder Score
Coiled-Coil
Transmembrane
0
50
100
150
200
250

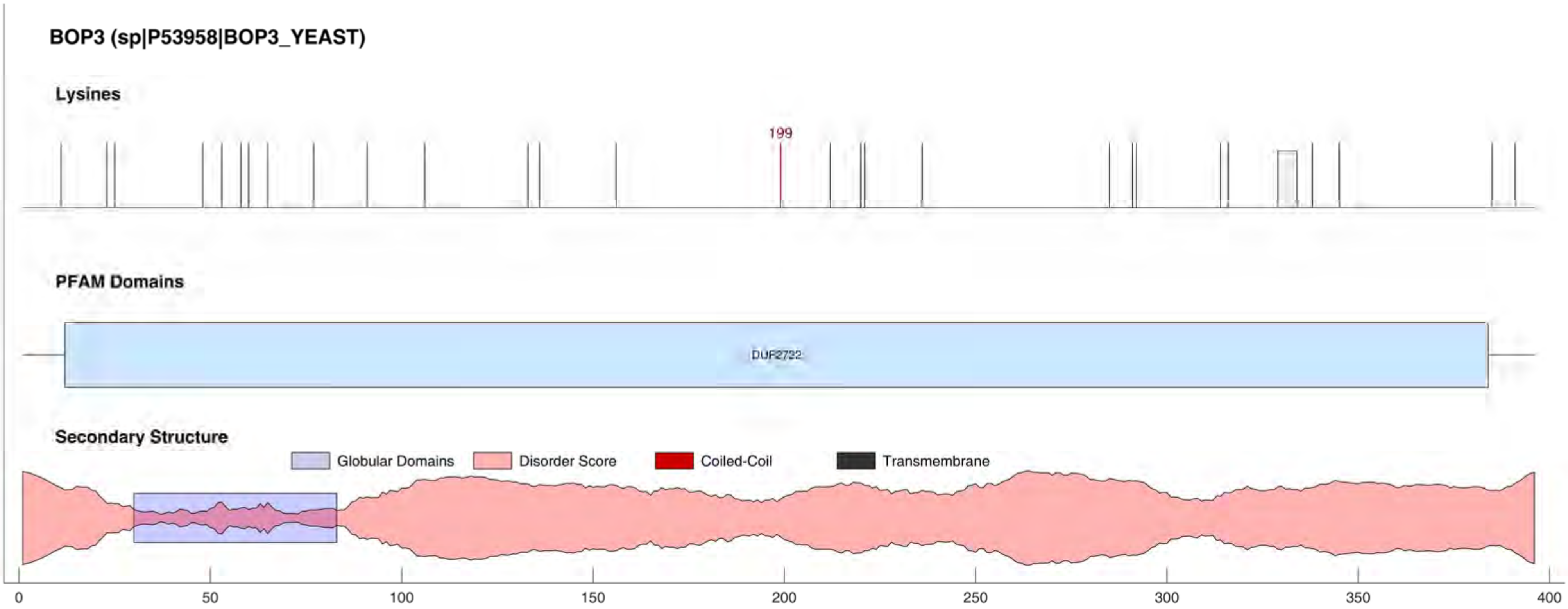
BOP3 (sp|P53958|BOP3_YEAST)
Lysines
199
PFAM Domains
DUF2722
Secondary Structure
Globular Domains
Disorder Score
Coiled-Coil
Transmembrane
0
50
100
150
200
250
300
350
400

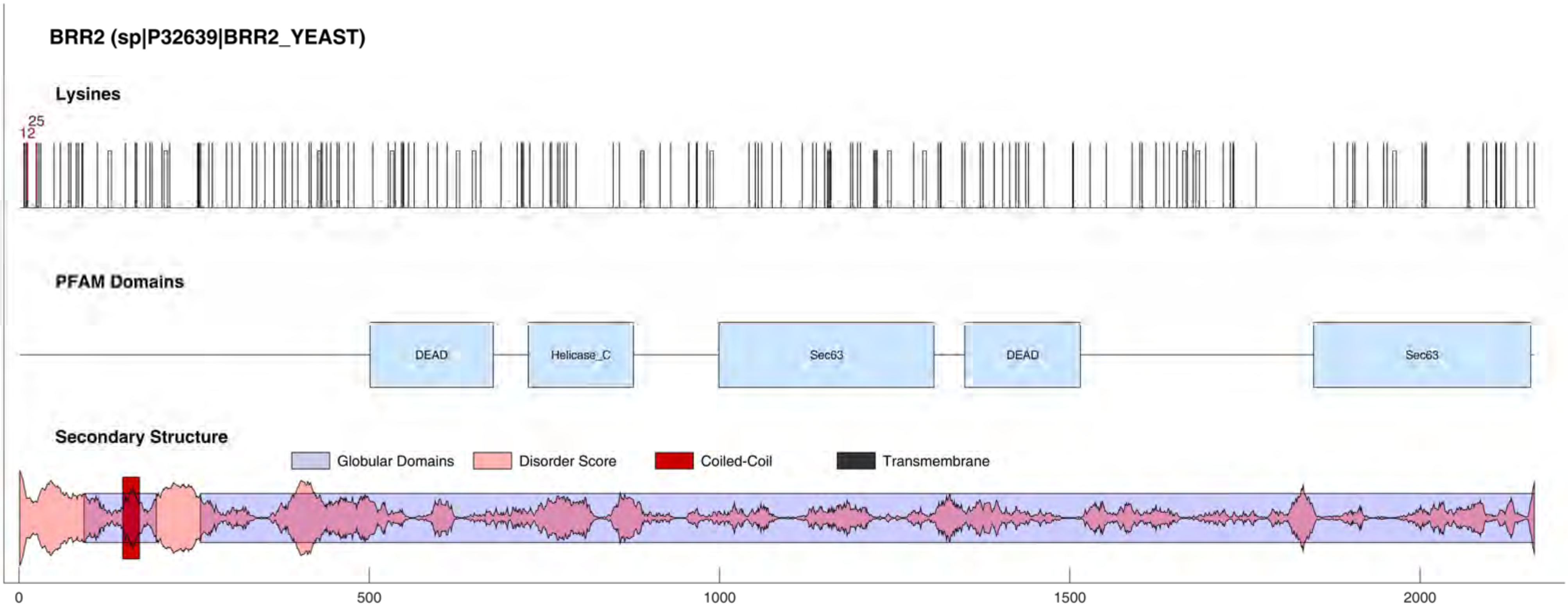
BRR2 (sp|P32639|BRR2_YEAST)
Lysines
25
12
PFAM Domains
DEAD
Helicase_C
Sec63
DEAD
Sec63
Secondary Structure
Globular Domains
Disorder Score
Coiled-Coil
Transmembrane
0
500
1000
1500
2000

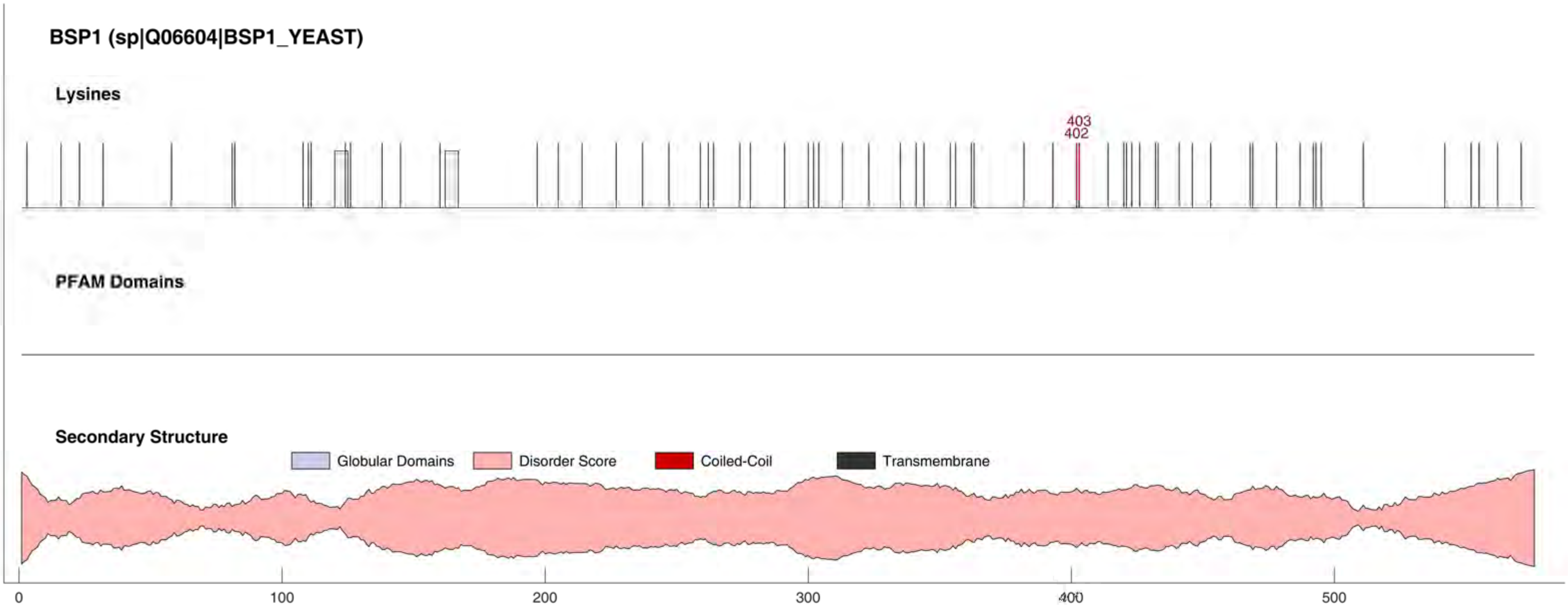

BSP1 (sp|Q06604|BSP1_YEAST)
Lysines
403
402
PFAM Domains
Secondary Structure
Globular Domains
Disorder Score
Coiled-Coil
Transmembrane
0
100
200
300
400
500

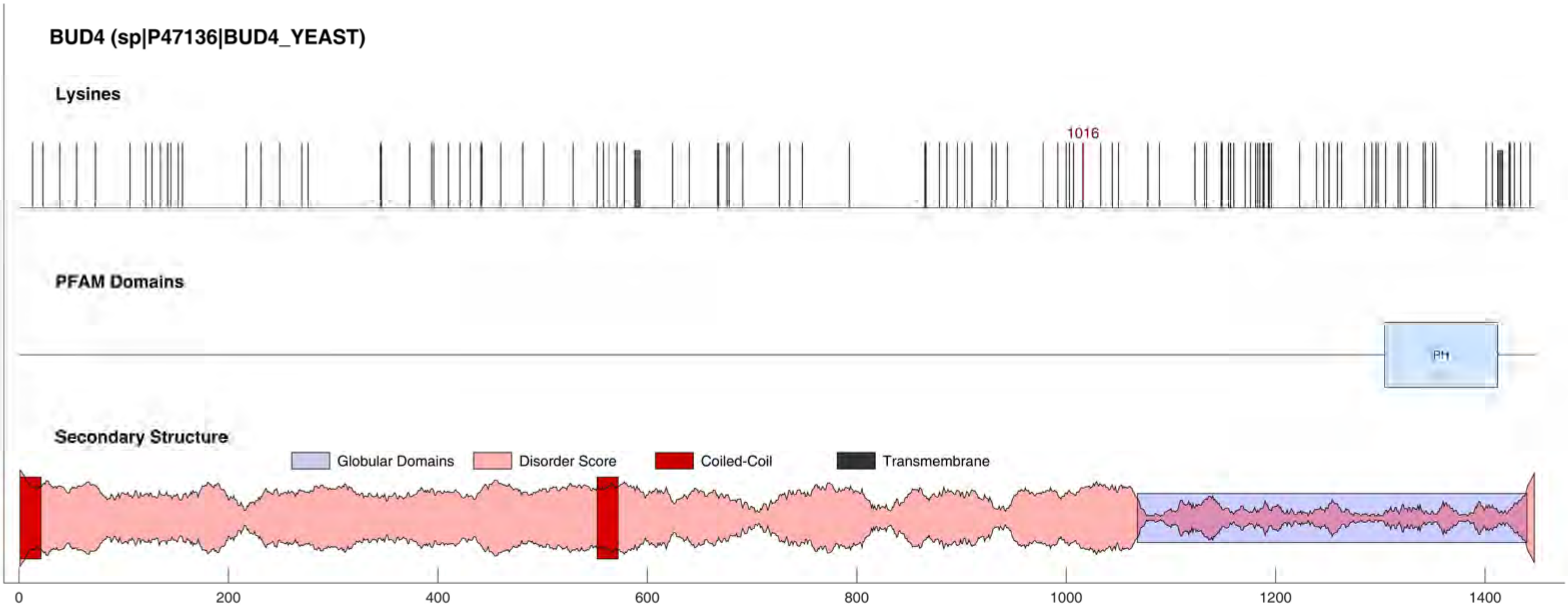
BUD4 (sp|P47136|BUD4_YEAST)
Lysines
1016
PFAM Domains
PH
Secondary Structure
Globular Domains
Disorder Score
Coiled-Coil
Transmembrane
0
200
400
600
800
1000
1200
1400

#### BUD6 (sp|P41697|BUD6_YEAST)

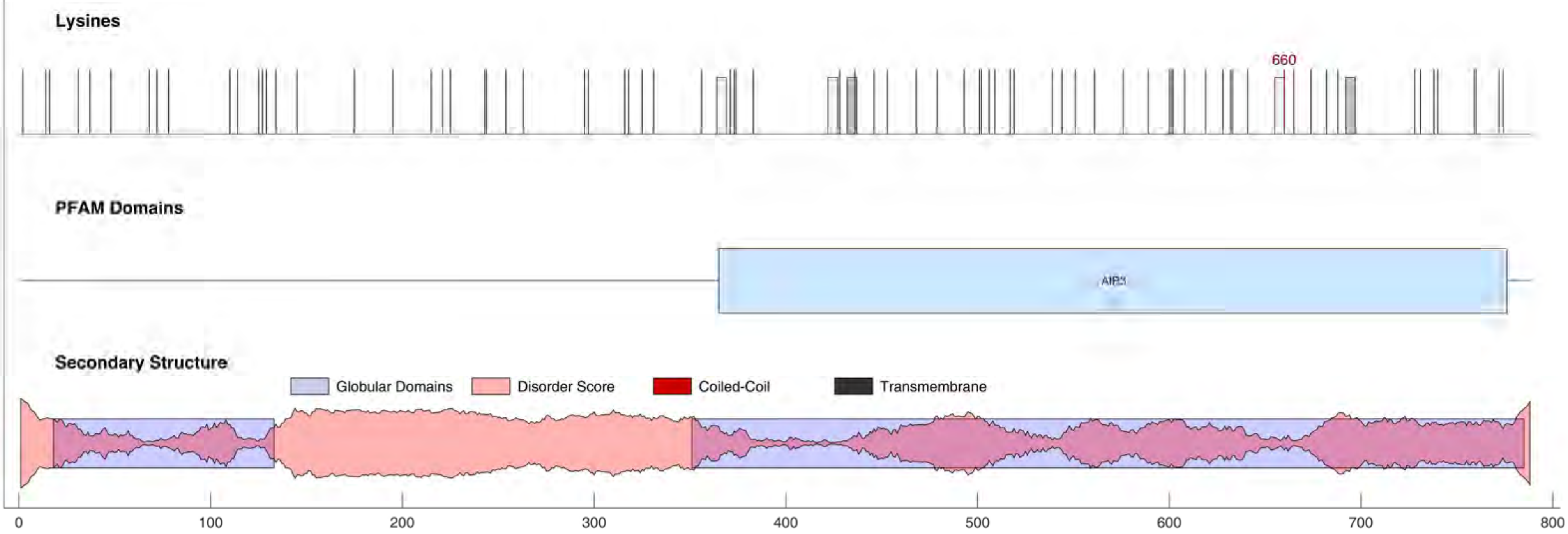

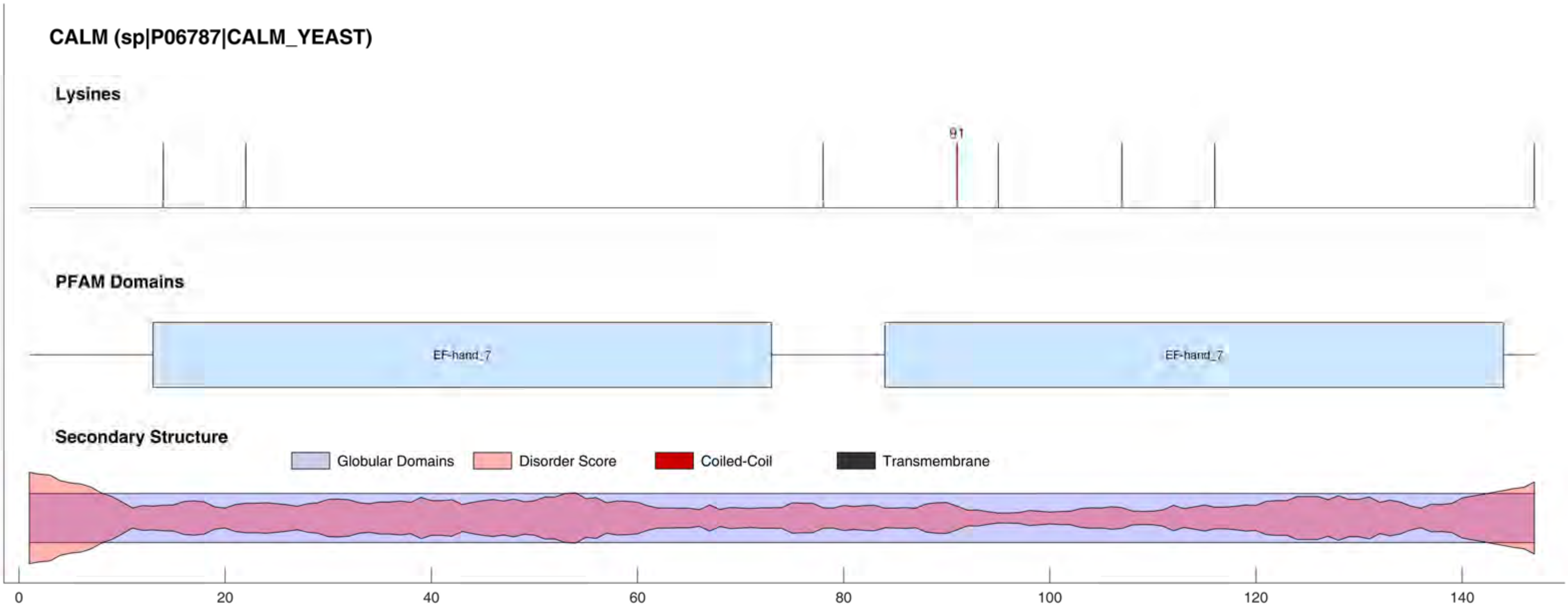

CALM (sp|P06787|CALM_YEAST)
Lysines
91
PFAM Domains
EF-hand_7
EF-hand_7
Secondary Structure
Globular Domains
Disorder Score
Coiled-Coil
Transmembrane
0
20
40
60
80
100
120
140

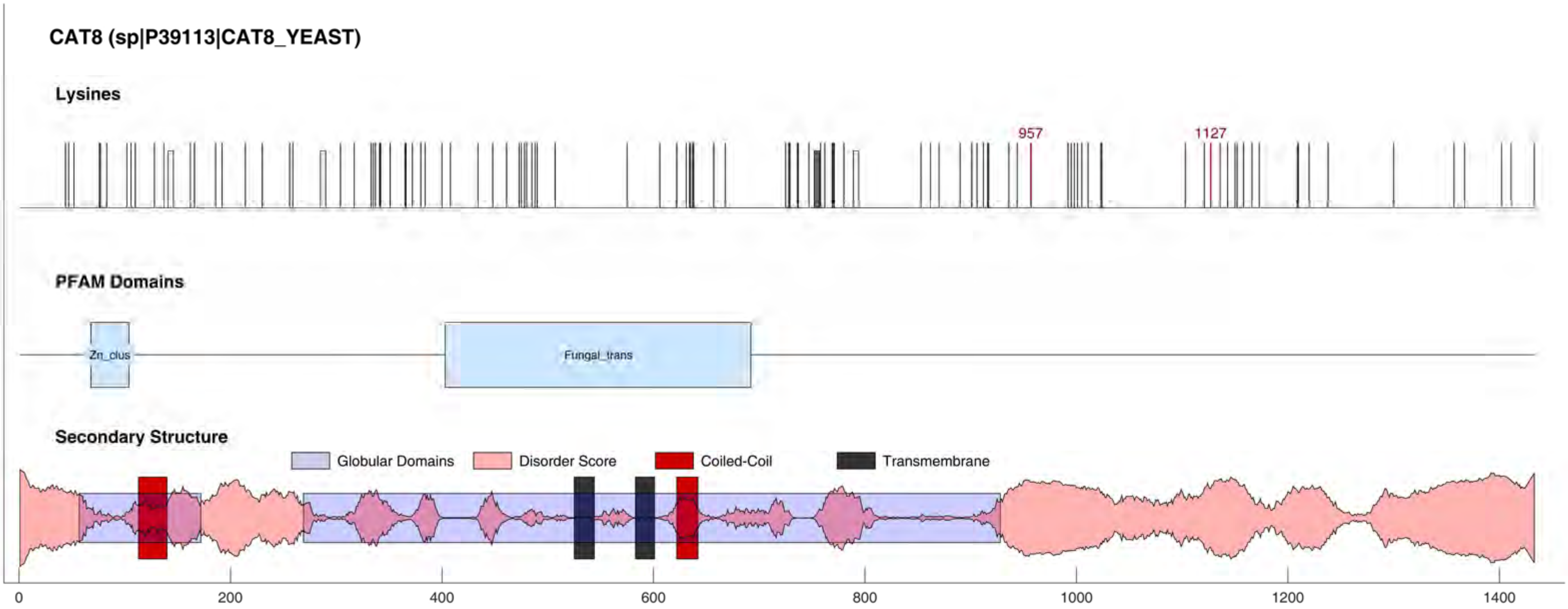

CAT8 (sp|P39113|CAT8_YEAST)
Lysines
957
1127
PFAM Domains
Zn_clus
Fungal_trans
Secondary Structure
Globular Domains
Disorder Score
Coiled-Coil
Transmembrane
0
200
400
600
800
1000
1200
1400

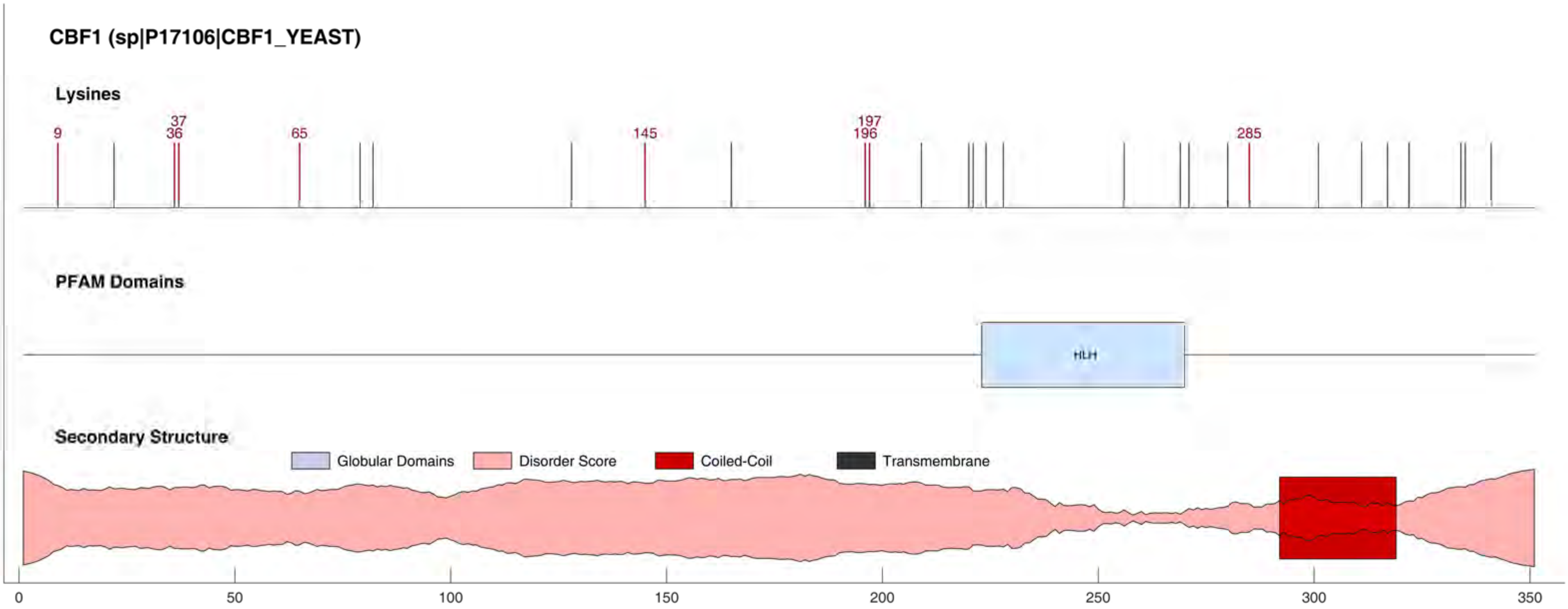
CBF1 (sp|P17106|CBF1_YEAST)
Lysines
9
37
36
65
145
197
196
285
PFAM Domains
HLH
Secondary Structure
Globular Domains
Disorder Score
Coiled-Coil
Transmembrane
0
50
100
150
200
250
300
350

#### CBF3A (sp|P32504|CBF3A_YEAST)

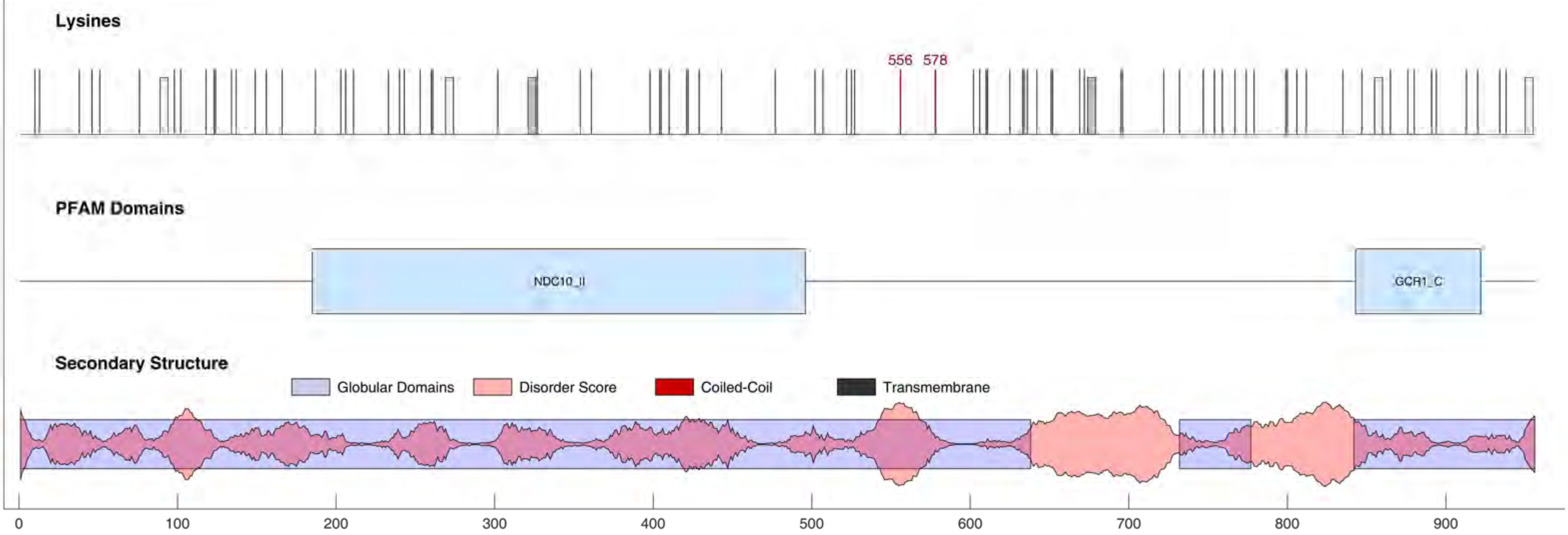

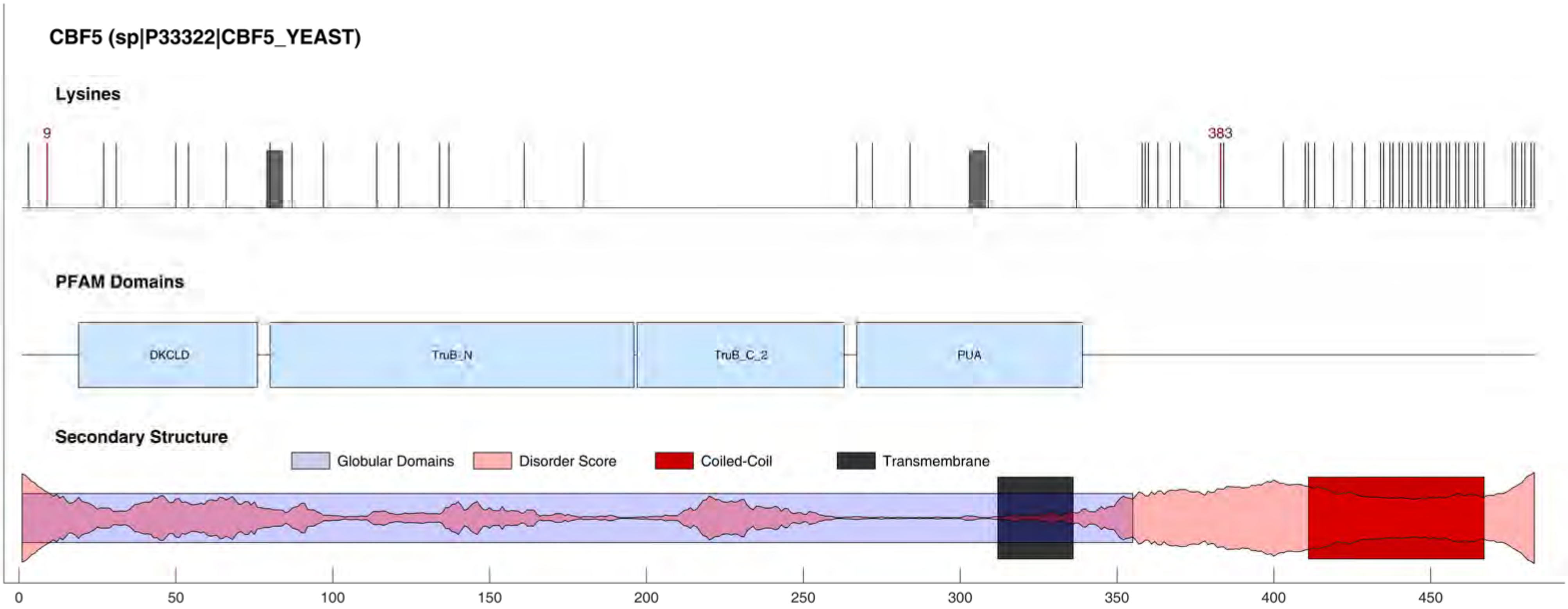
CBF5 (sp|P33322|CBF5_YEAST)
Lysines
9
383
PFAM Domains
DKCLD
TruB_N
TruB_C_2
PUA
Secondary Structure
Globular Domains
Disorder Score
Coiled-Coil
Transmembrane
0
50
100
150
200
250
300
350
400
450

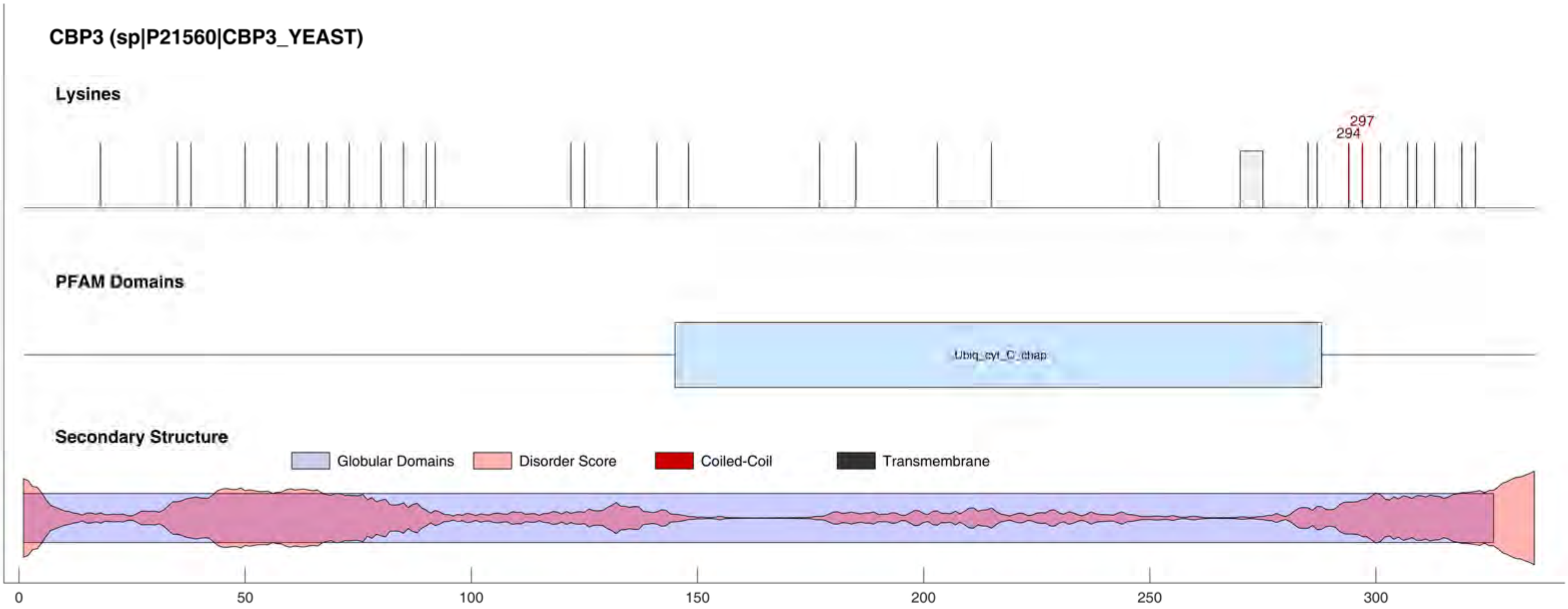

CBP3 (sp|P21560|CBP3_YEAST)
Lysines
297
294
PFAM Domains
Ubiq_cyt_C_chap
Secondary Structure
Globular Domains
Disorder Score
Coiled-Coil
Transmembrane
0
50
100
150
200
250
300

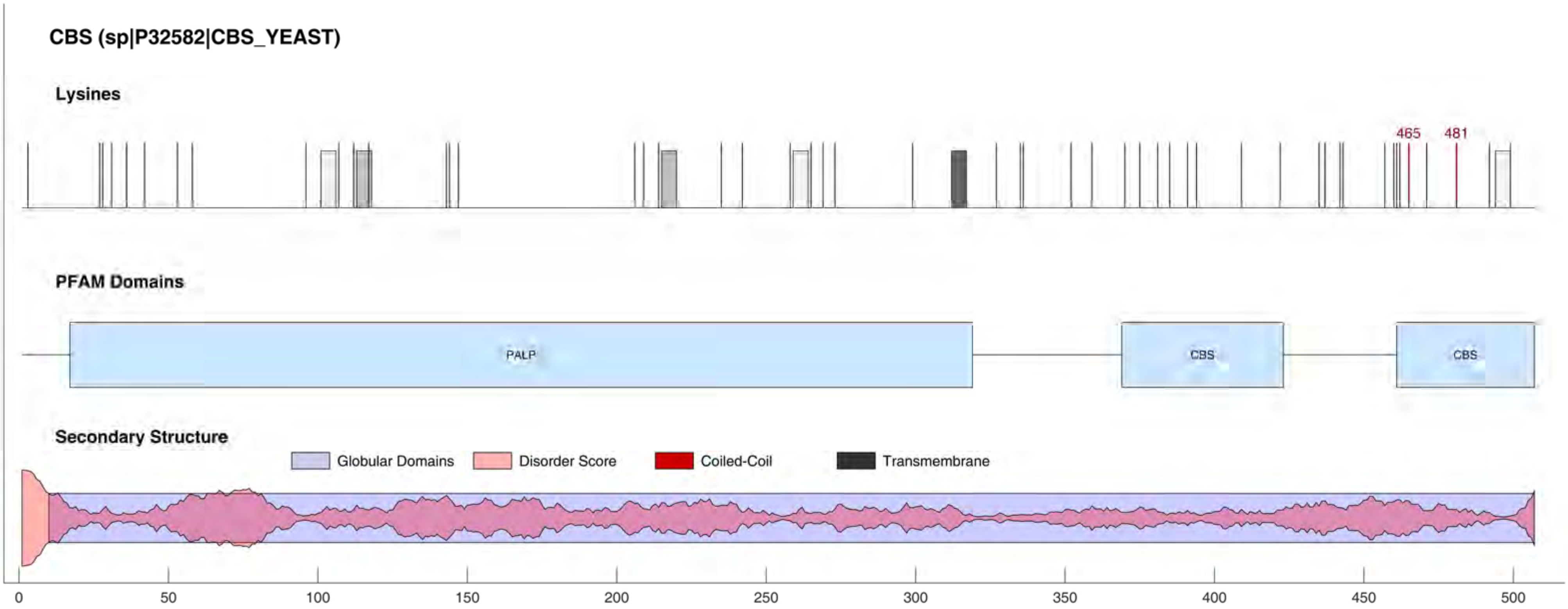
CBS (sp|P32582|CBS_YEAST)
Lysines
465
481
PFAM Domains
PALP
CBS
CBS
Secondary Structure
Globular Domains
Disorder Score
Coiled-Coil
Transmembrane
0
50
100
150
200
250
300
350
400
450
500

#### CCM1 (sp|P48237|CCM1_YEAST)

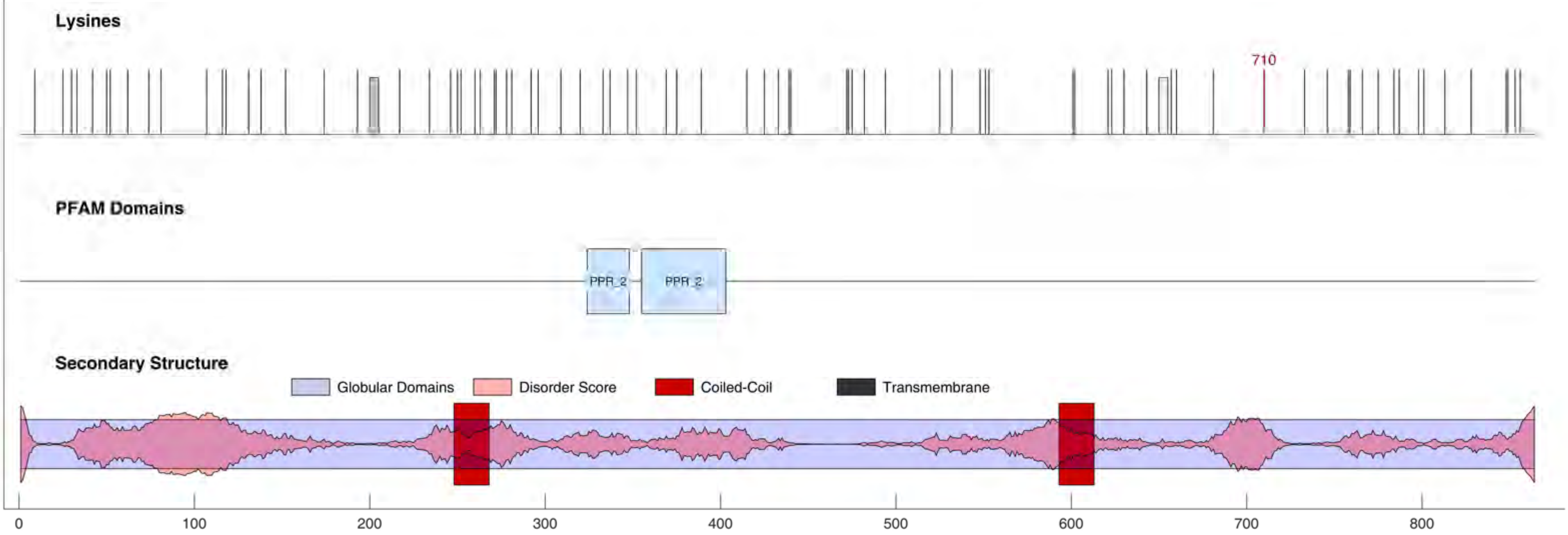

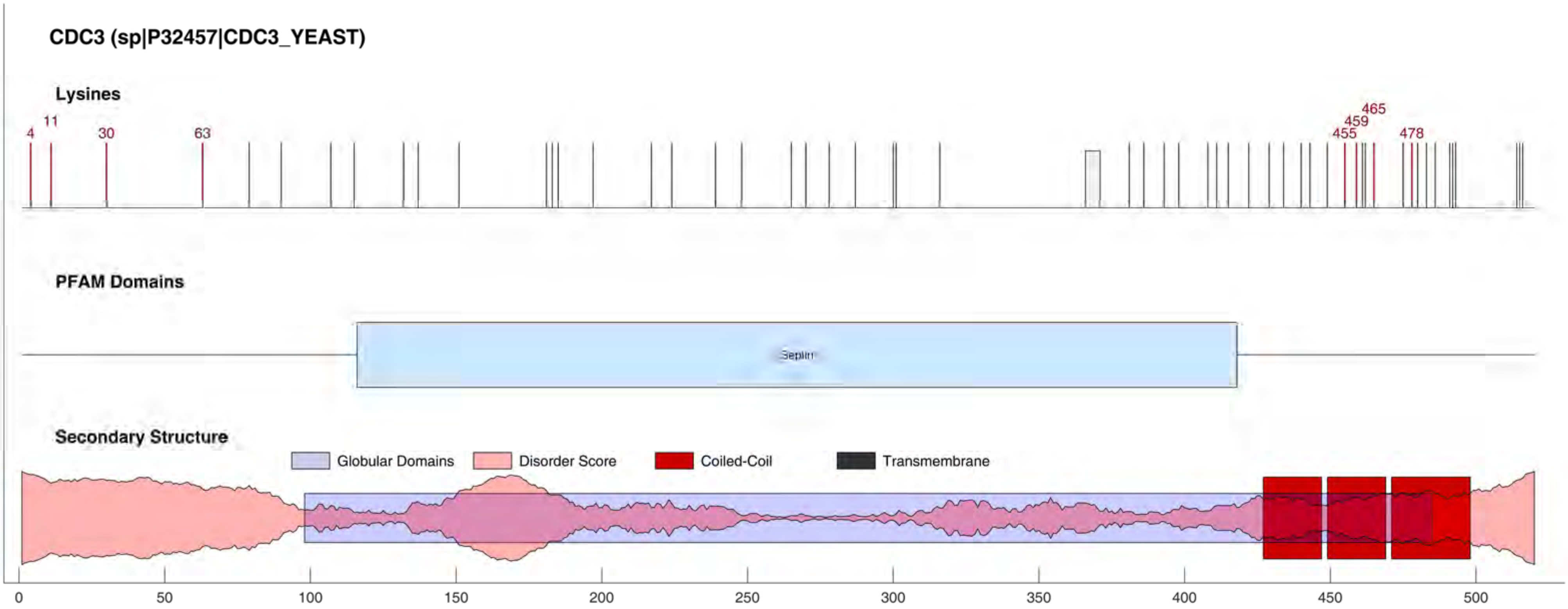
CDC3 (sp|P32457|CDC3_YEAST)
Lysines
4
11
30
63
455
459
465
478
PFAM Domains
Septin
Secondary Structure
Globular Domains
Disorder Score
Coiled-Coil
Transmembrane
0
50
100
150
200
250
300
350
400
450
500

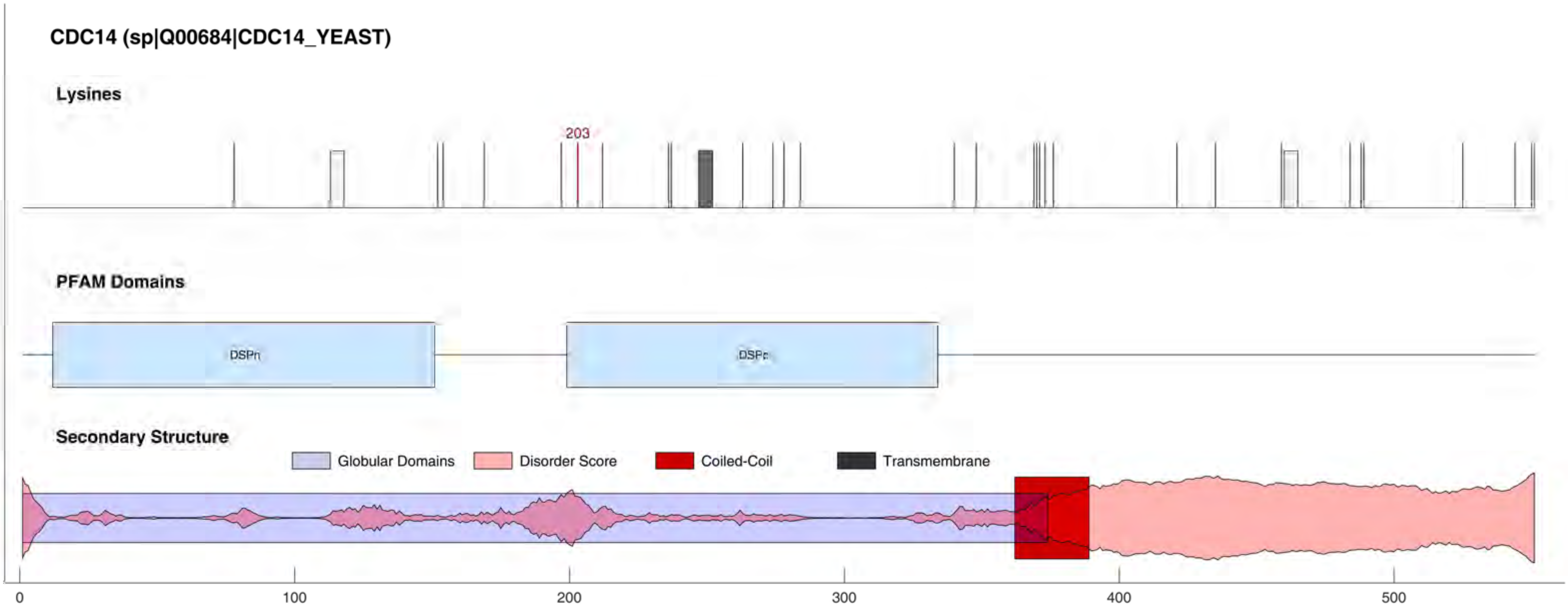
CDC14 (sp|Q00684|CDC14_YEAST)
Lysines
203
PFAM Domains
DSPn
DSPc
Secondary Structure
Globular Domains
Disorder Score
Coiled-Coil
Transmembrane
0
100
200
300
400
500

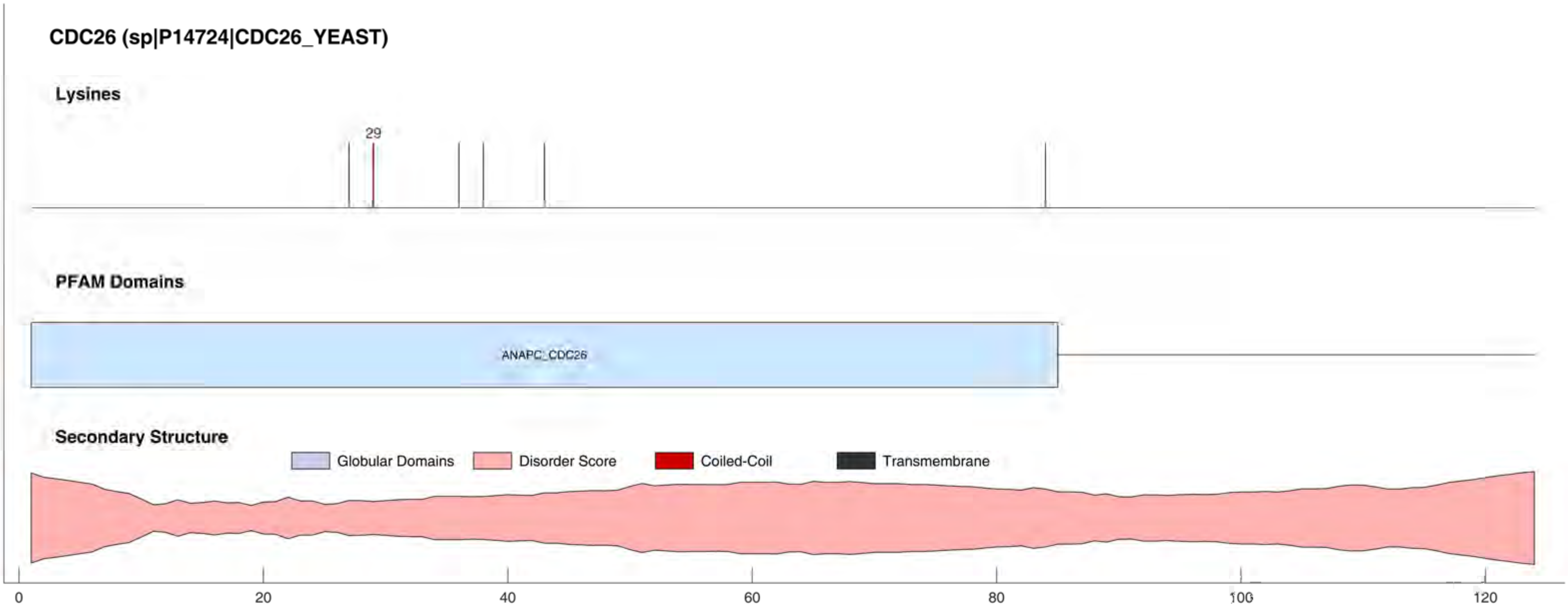

CDC26 (sp|P14724|CDC26_YEAST)
Lysines
29
PFAM Domains
ANAPC_CDC26
Secondary Structure
Globular Domains
Disorder Score
Coiled-Coil
Transmembrane
0
20
40
60
80
100
120

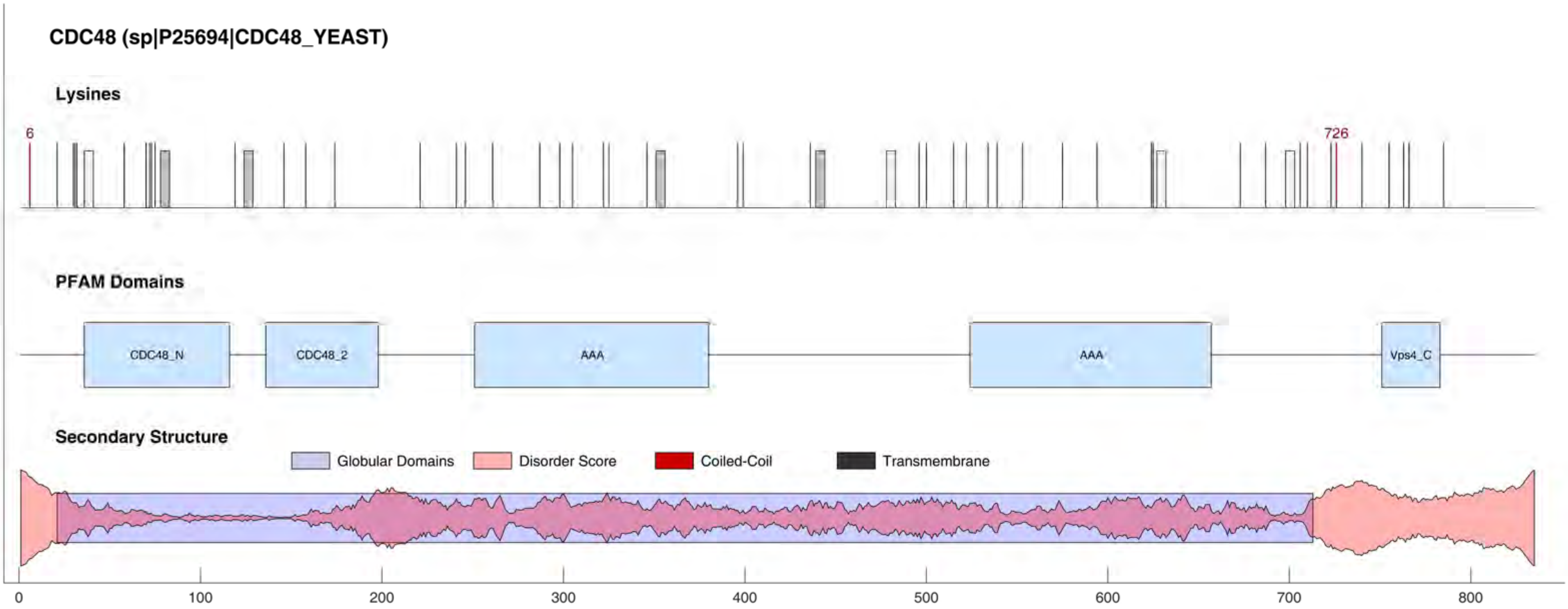
CDC48 (sp|P25694|CDC48_YEAST)
Lysines
6
726
PFAM Domains
CDC48_N
CDC48_2
AAA
AAA
Vps4_C
Secondary Structure
Globular Domains
Disorder Score
Coiled-Coil
Transmembrane
0
100
200
300
400
500
600
700
800

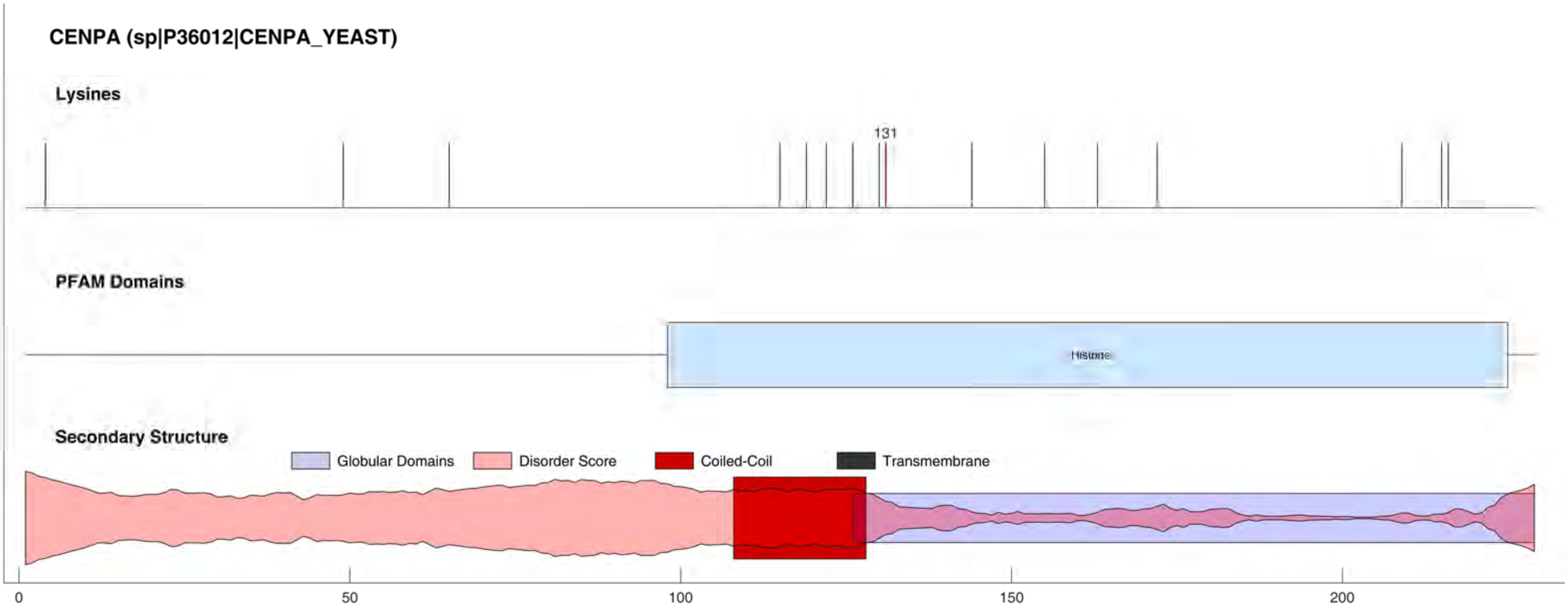
CENPA (sp|P36012|CENPA_YEAST)
Lysines
131
PFAM Domains
Histone
Secondary Structure
Globular Domains
Disorder Score
Coiled-Coil
Transmembrane
0
50
100
150
200

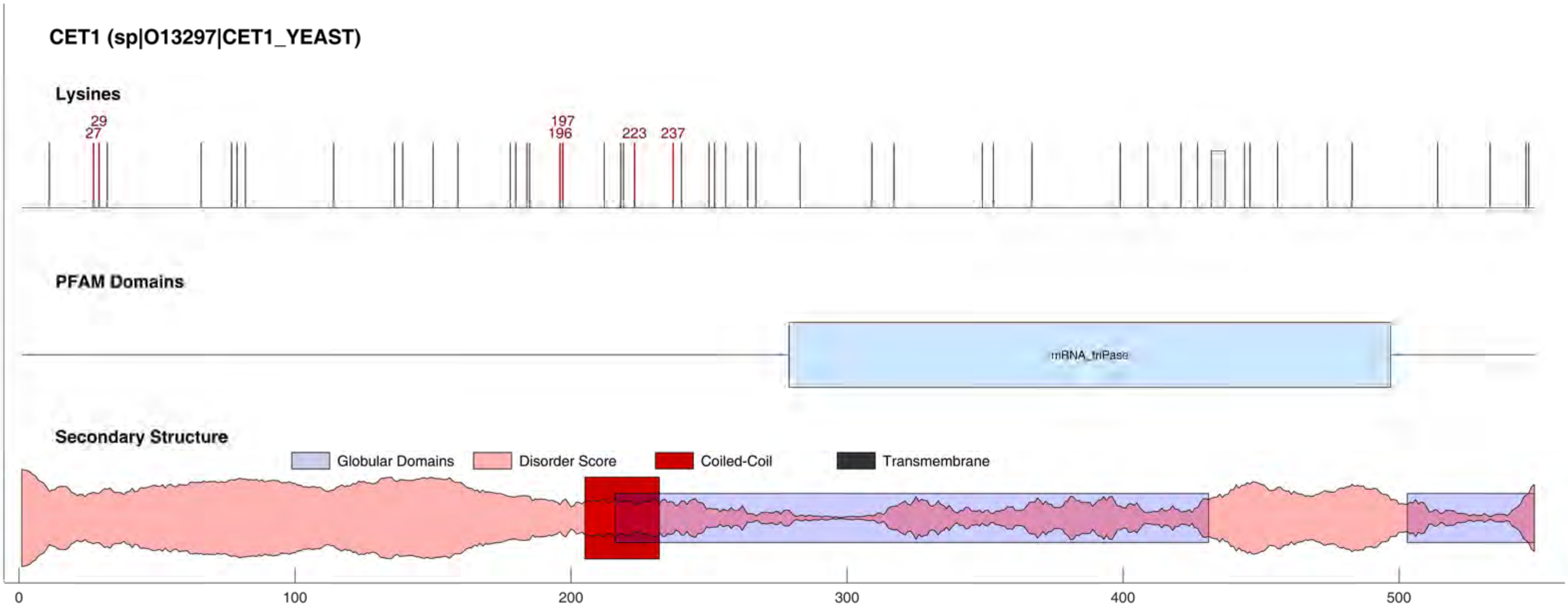
CET1 (sp|O13297|CET1_YEAST)
Lysines
29
27
197
196
223
237
PFAM Domains
mRNA_triPase
Secondary Structure
Globular Domains
Disorder Score
Coiled-Coil
Transmembrane
0
100
200
300
400
500

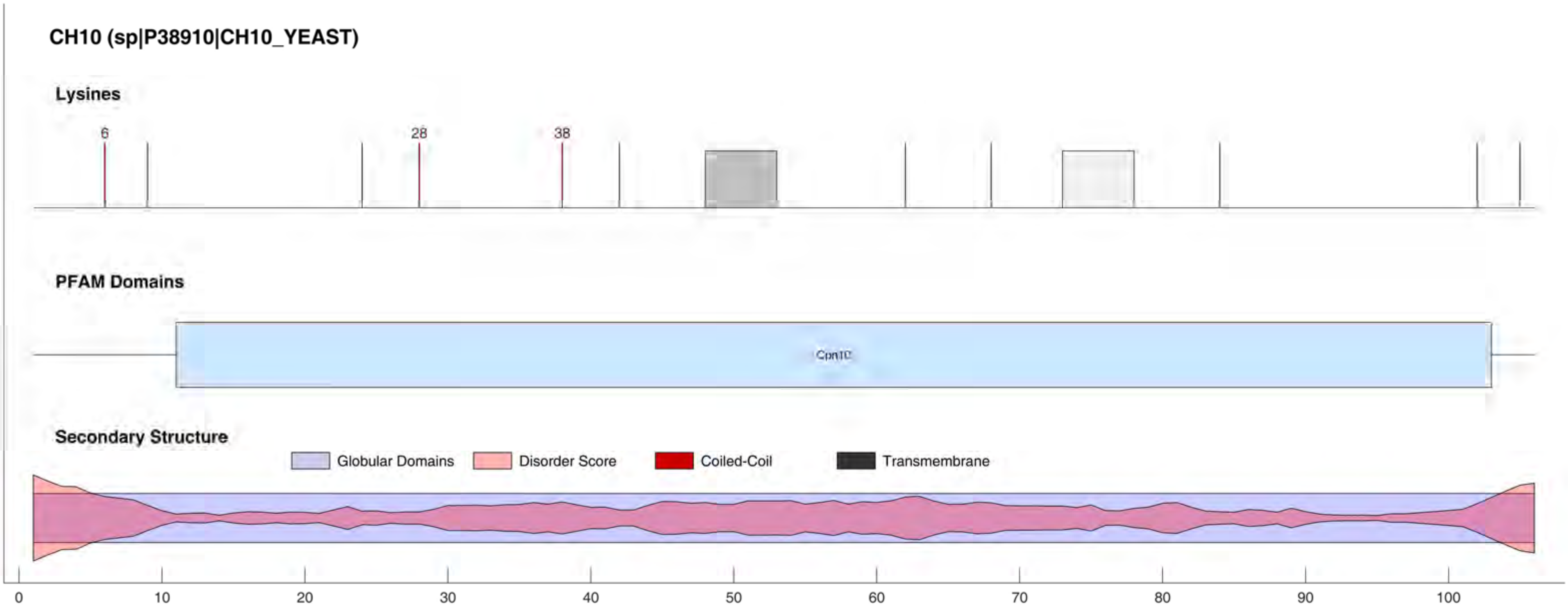

CH10 (sp|P38910|CH10_YEAST)
Lysines
6
28
38
PFAM Domains
Cpn10
Secondary Structure
Globular Domains
Disorder Score
Coiled-Coil
Transmembrane
0
10
20
30
40
50
60
70
80
90
100

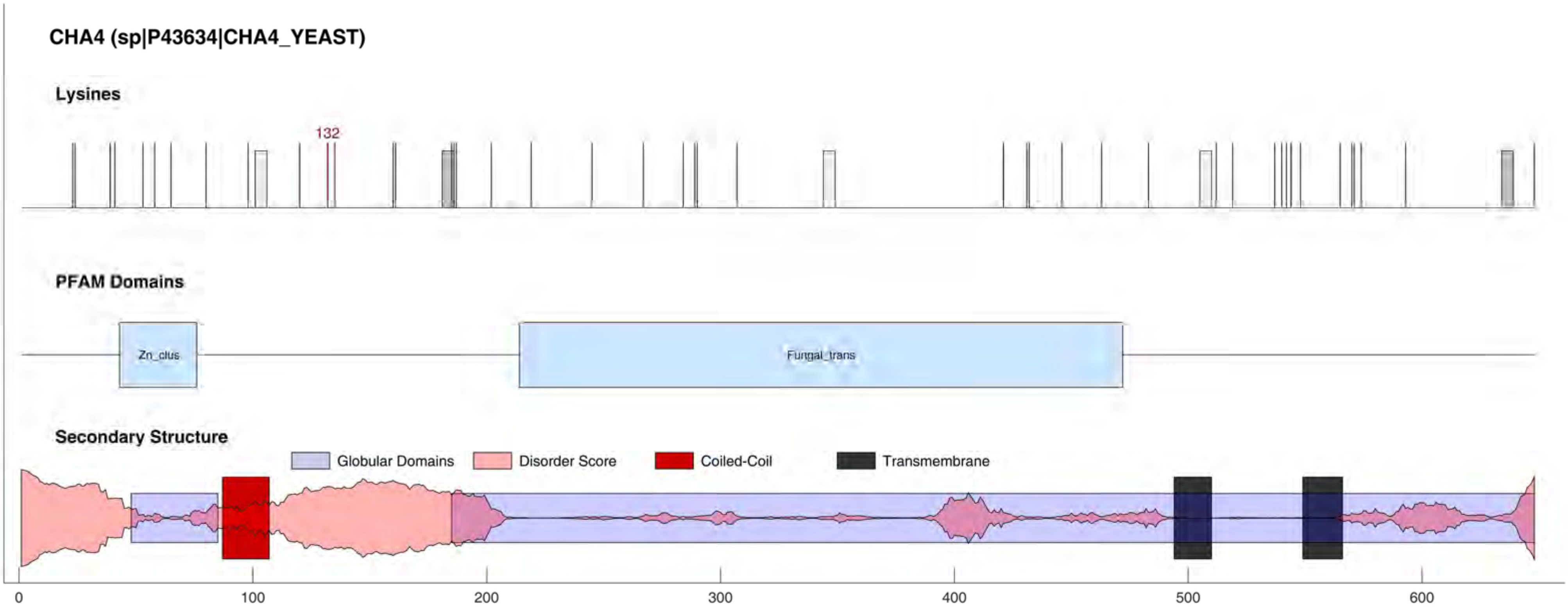

CHA4 (sp|P43634|CHA4_YEAST)
Lysines
132
PFAM Domains
Zn_clus
Fungal_trans
Secondary Structure
Globular Domains
Disorder Score
Coiled-Coil
Transmembrane
0
100
200
300
400
500
600

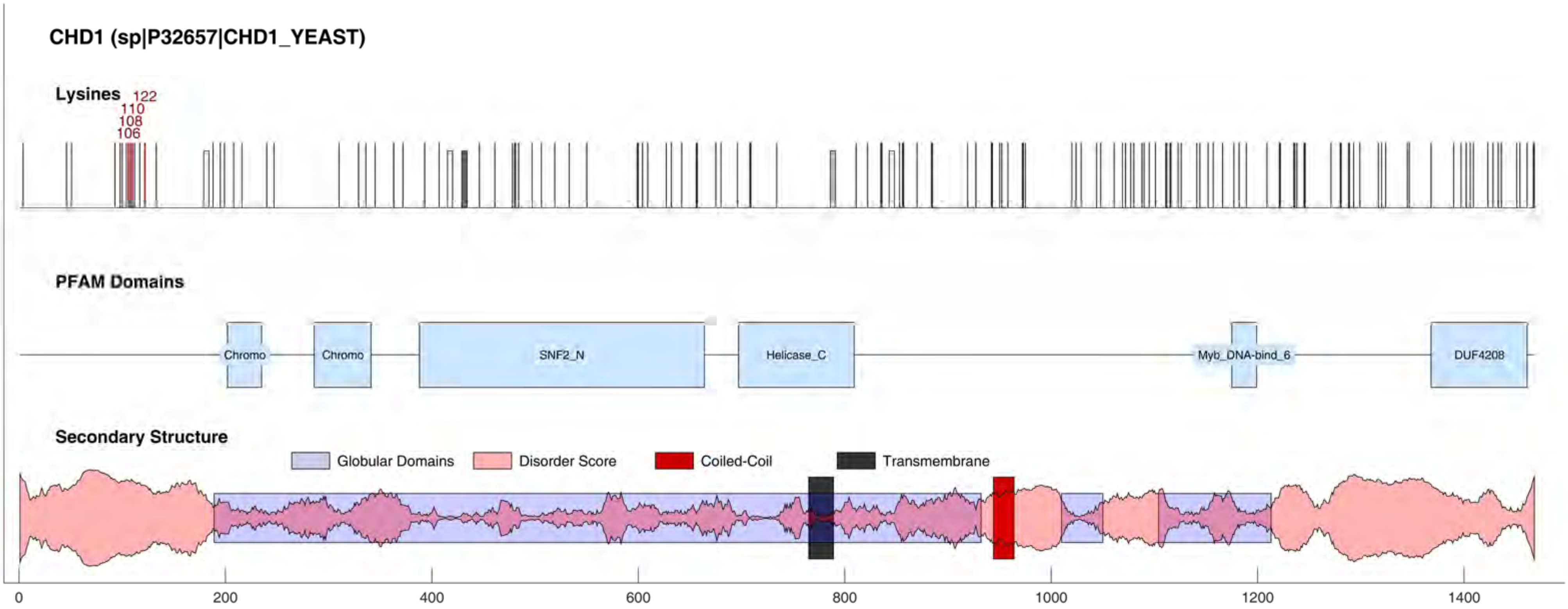
CHD1 (sp|P32657|CHD1_YEAST)
Lysines
122
110
108
106
PFAM Domains
Chromo
Chromo
SNF2_N
Helicase_C
Myb_DNA-bind_6
DUF4208
Secondary Structure
Globular Domains
Disorder Score
Coiled-Coil
Transmembrane
0
200
400
600
800
1000
1200
1400

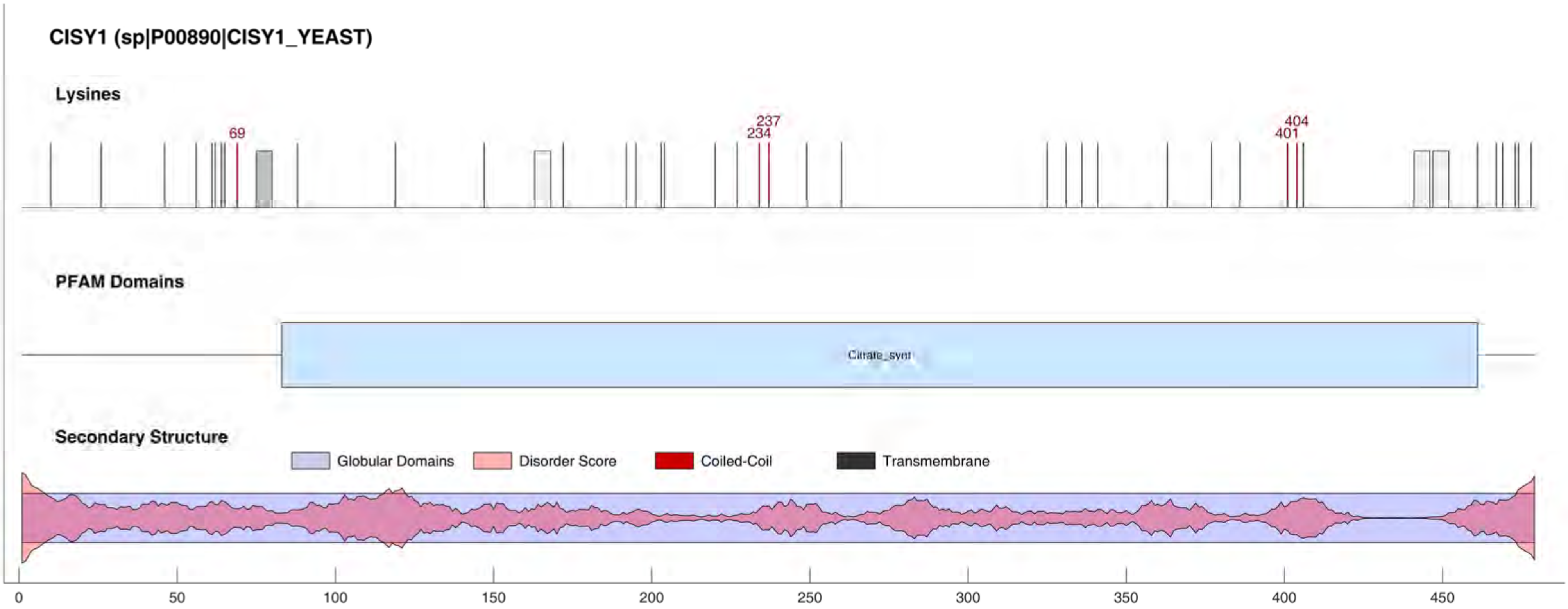
CISY1 (sp|P00890|CISY1_YEAST)
Lysines
69
234
237
401
404
PFAM Domains
Citrate_synt
Secondary Structure
Globular Domains
Disorder Score
Coiled-Coil
Transmembrane
0
50
100
150
200
250
300
350
400
450

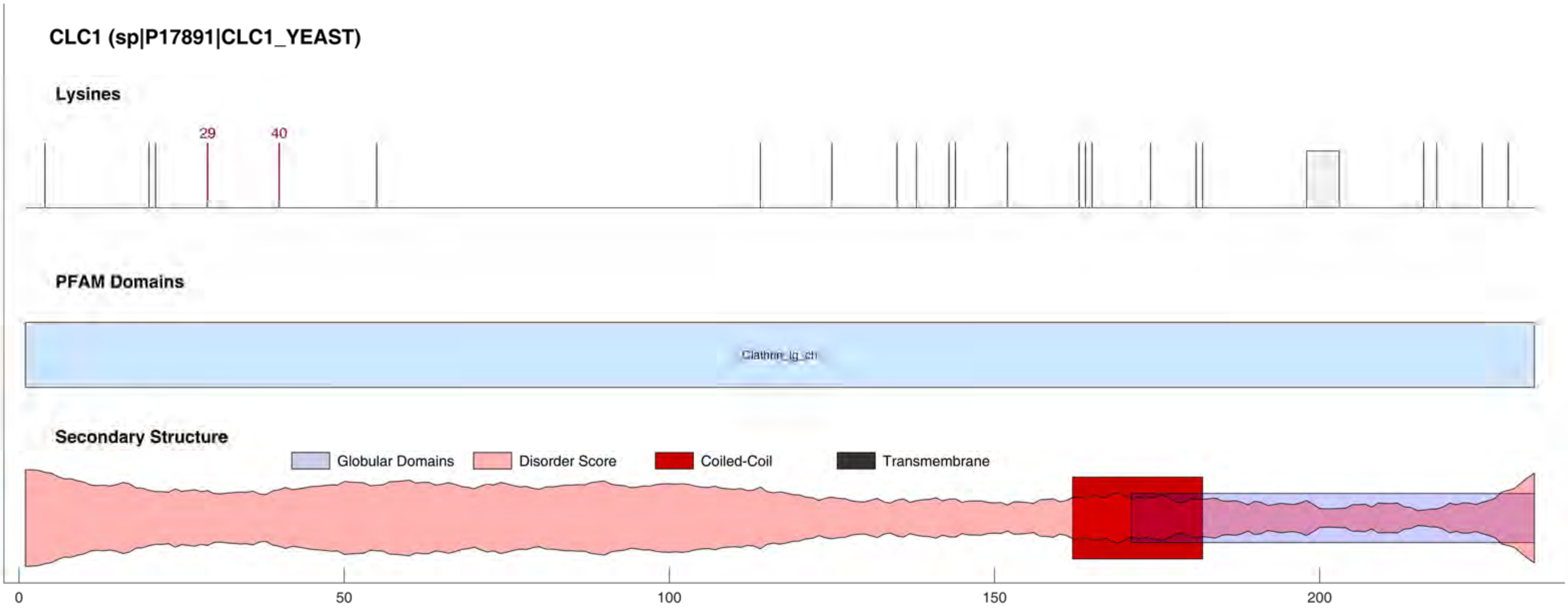
CLC1 (sp|P17891|CLC1_YEAST)
Lysines
29
40
PFAM Domains
Clathrin_lg_ch
Secondary Structure
Globular Domains
Disorder Score
Coiled-Coil
Transmembrane
0
50
100
150
200

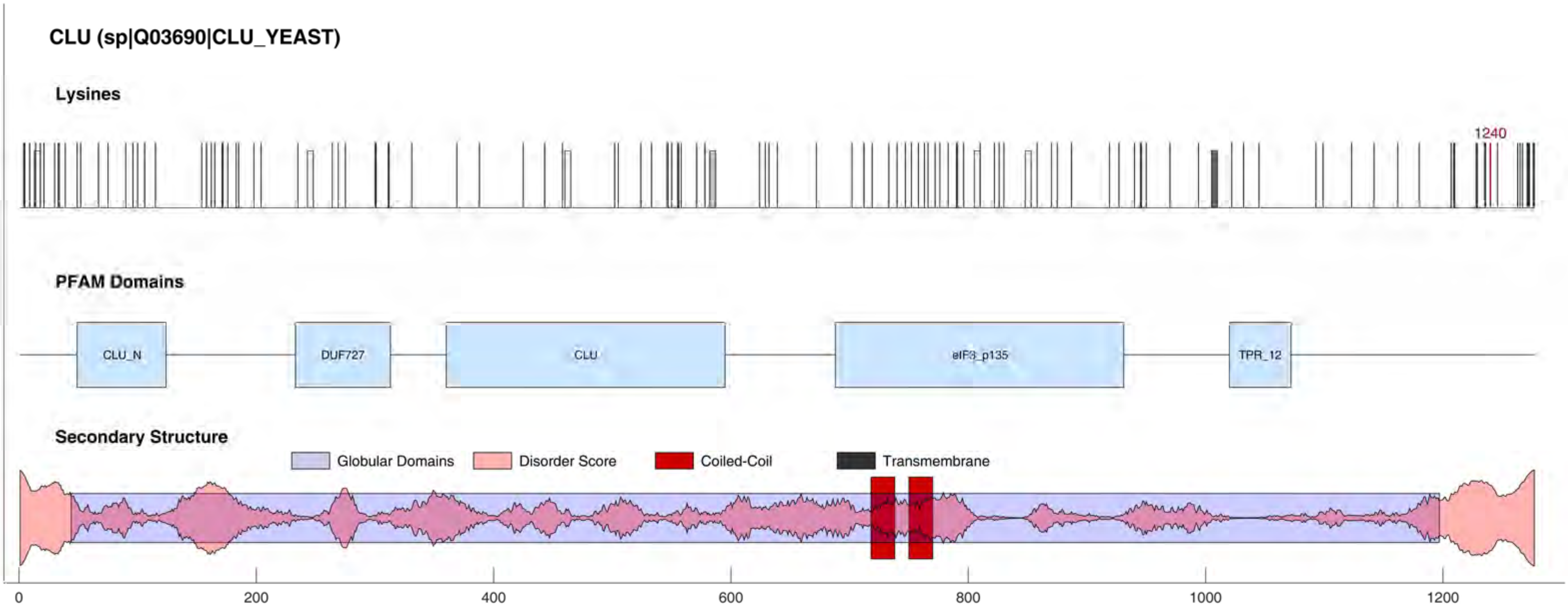

CLU (sp|Q03690|CLU_YEAST)
Lysines
1240
PFAM Domains
CLU_N
DUF727
CLU
eIF3_p135
TPR_12
Secondary Structure
Globular Domains
Disorder Score
Coiled-Coil
Transmembrane
0
200
400
600
800
1000
1200

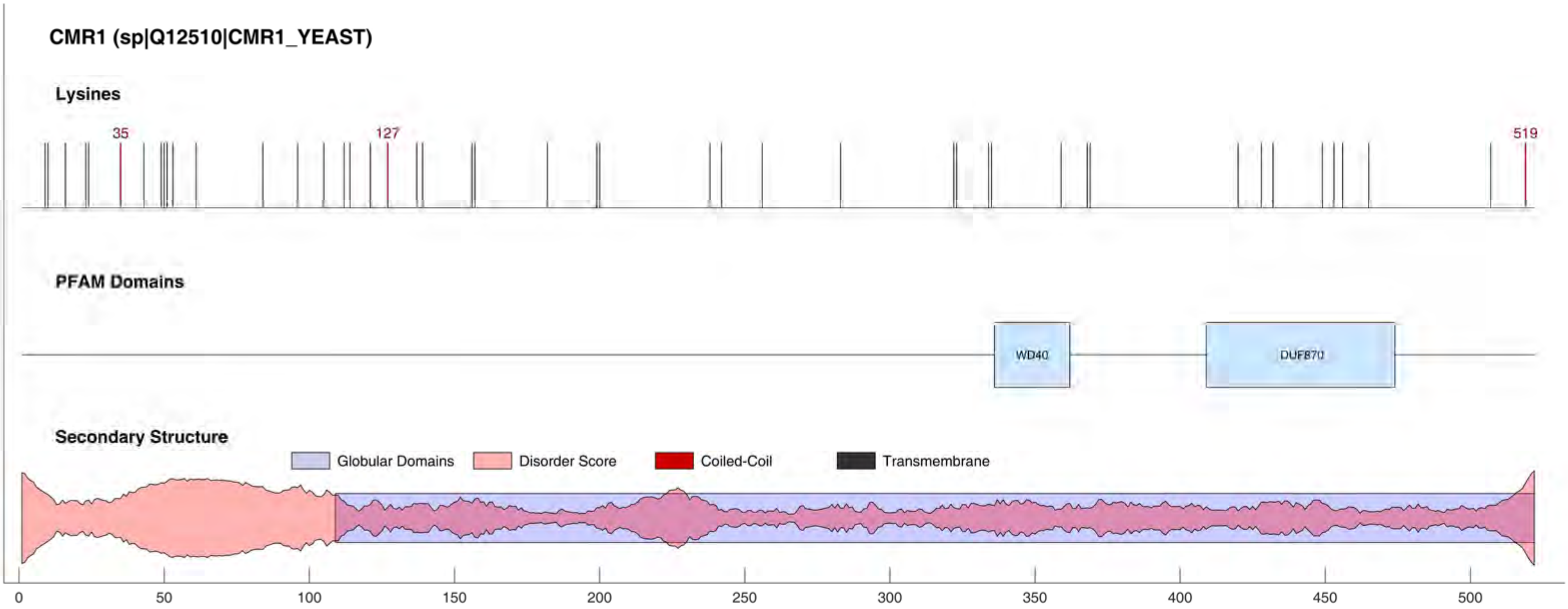

CMR1 (sp|Q12510|CMR1_YEAST)
Lysines
35
127
519
PFAM Domains
WD40
DUF870
Secondary Structure
Globular Domains
Disorder Score
Coiled-Coil
Transmembrane
0
50
100
150
200
250
300
350
400
450
500

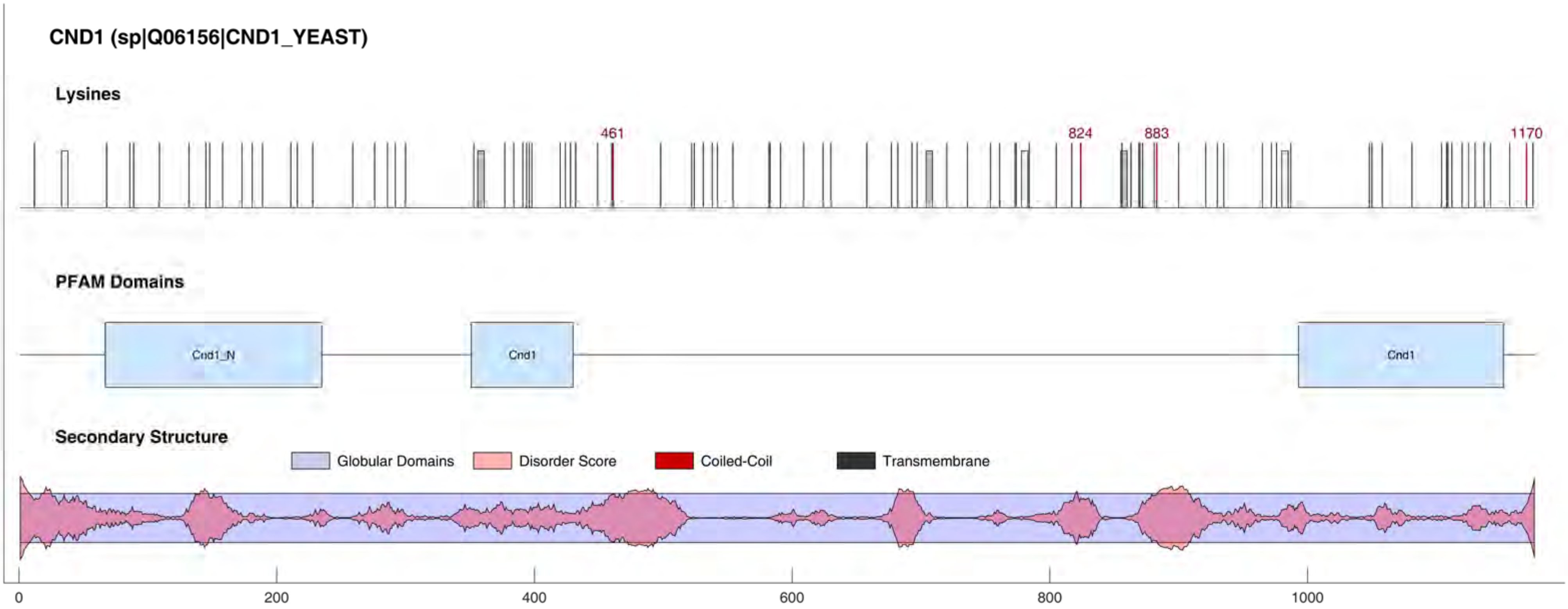

CND1 (sp|Q06156|CND1_YEAST)
Lysines
461
824
883
1170
PFAM Domains
Cnd1_N
Cnd1
Cnd1
Secondary Structure
Globular Domains
Disorder Score
Coiled-Coil
Transmembrane
0
200
400
600
800
1000

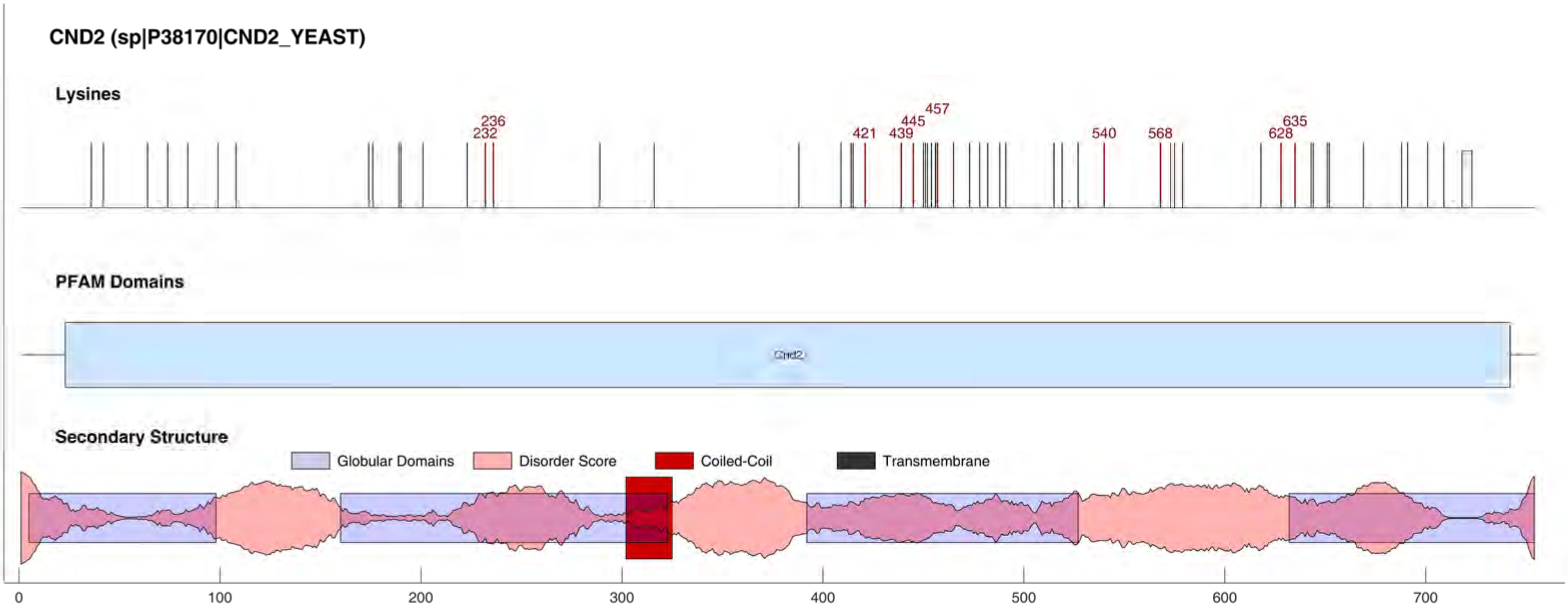
CND2 (sp|P38170|CND2_YEAST)
Lysines
236
232
421
439
445
457
540
568
628
635
PFAM Domains
Cnd2
Secondary Structure
Globular Domains
Disorder Score
Coiled-Coil
Transmembrane
0
100
200
300
400
500
600
700

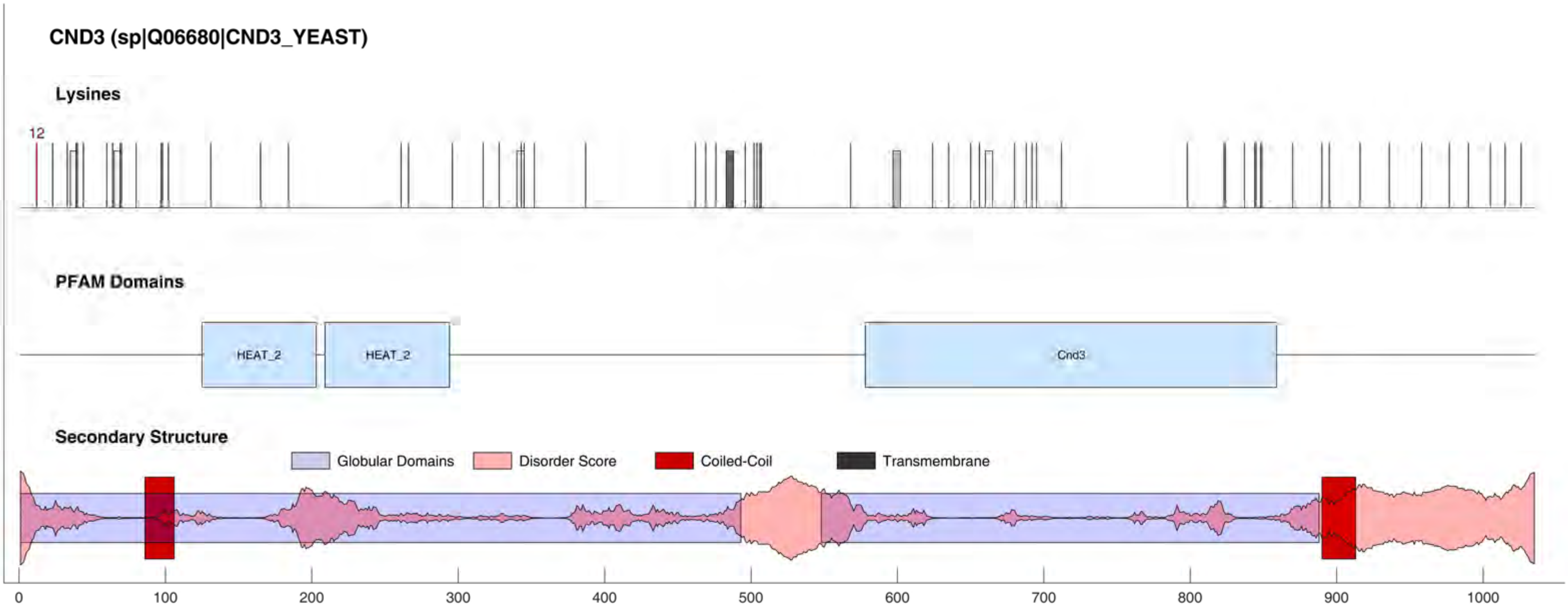
CND3 (sp|Q06680|CND3_YEAST)
Lysines
12
PFAM Domains
HEAT_2
HEAT_2
Cnd3
Secondary Structure
Globular Domains
Disorder Score
Coiled-Coil
Transmembrane
0
100
200
300
400
500
600
700
800
900
1000

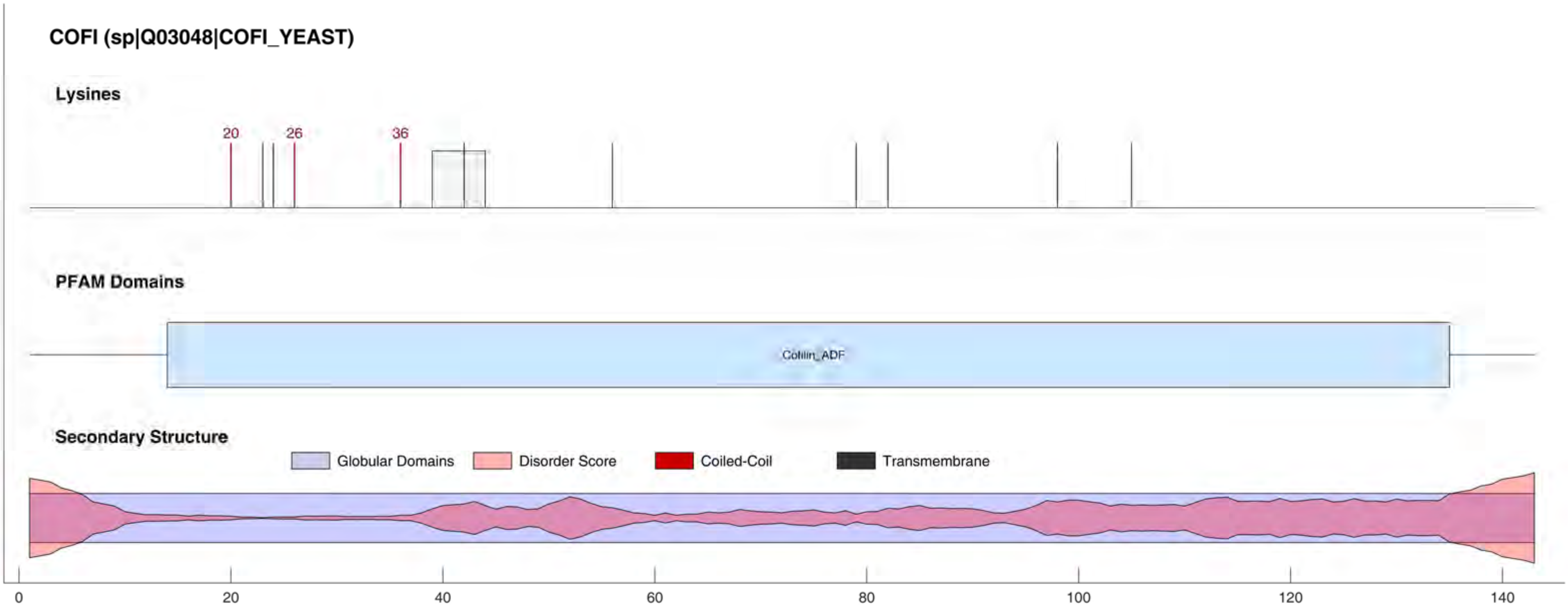
COFI (sp|Q03048|COFI_YEAST)
Lysines
20
26
36
PFAM Domains
Cofilin_ADF
Secondary Structure
Globular Domains
Disorder Score
Coiled-Coil
Transmembrane
0
20
40
60
80
100
120
140

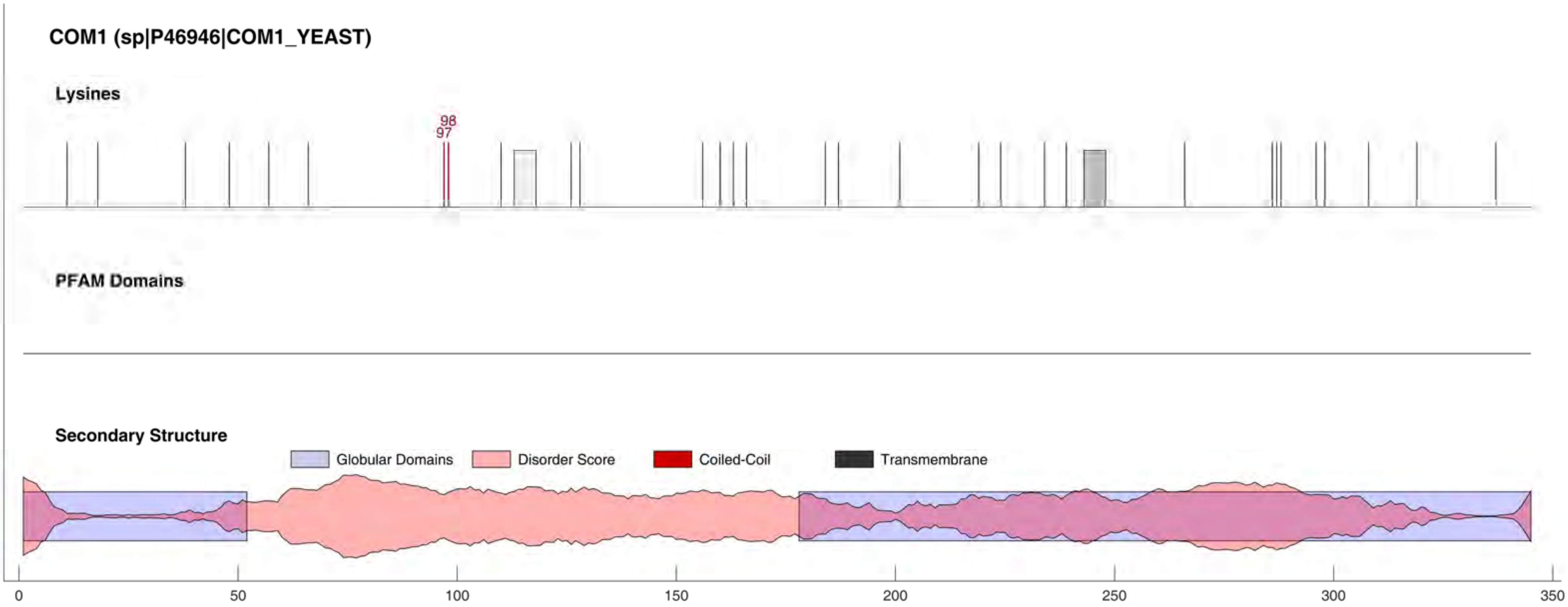
COM1 (sp|P46946|COM1_YEAST)
Lysines
98
97
PFAM Domains
Secondary Structure
Globular Domains
Disorder Score
Coiled-Coil
Transmembrane
0
50
100
150
200
250
300
350

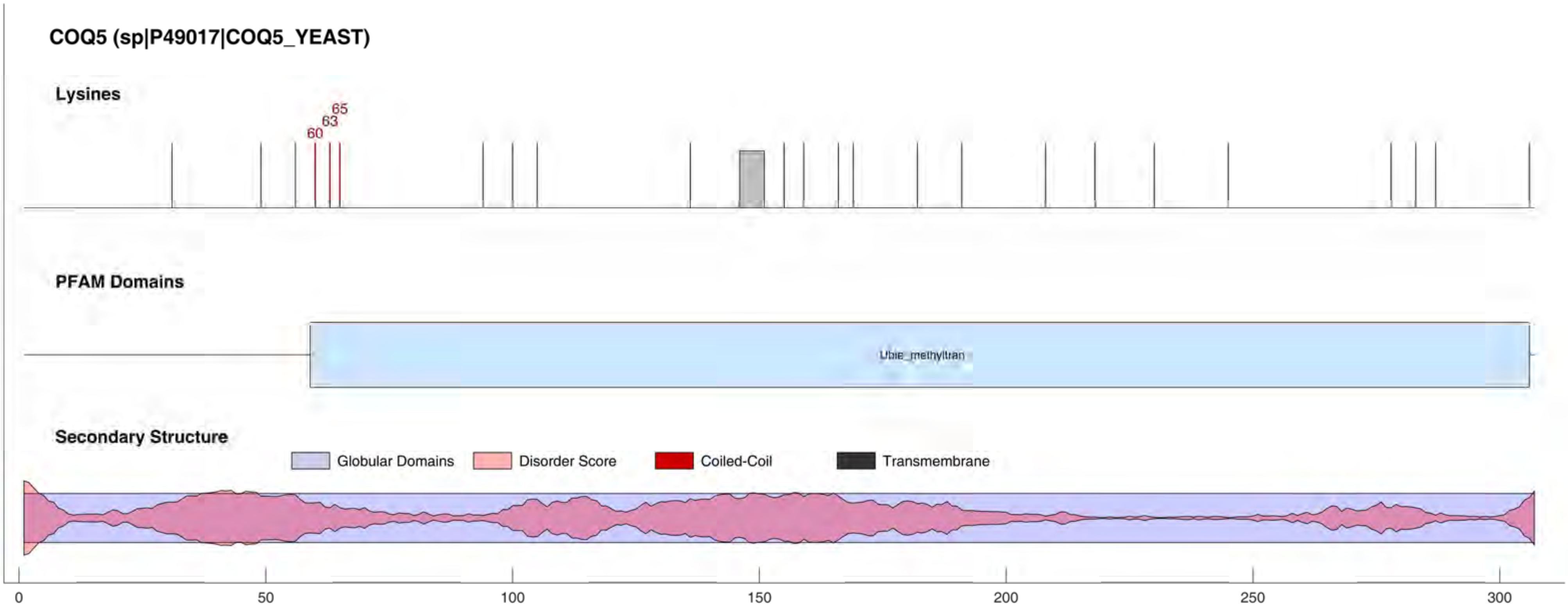

COQ5 (sp|P49017|COQ5_YEAST)
Lysines
60
63
65
PFAM Domains
Ubie_methyltran
Secondary Structure
Globular Domains
Disorder Score
Coiled-Coil
Transmembrane
0
50
100
150
200
250
300

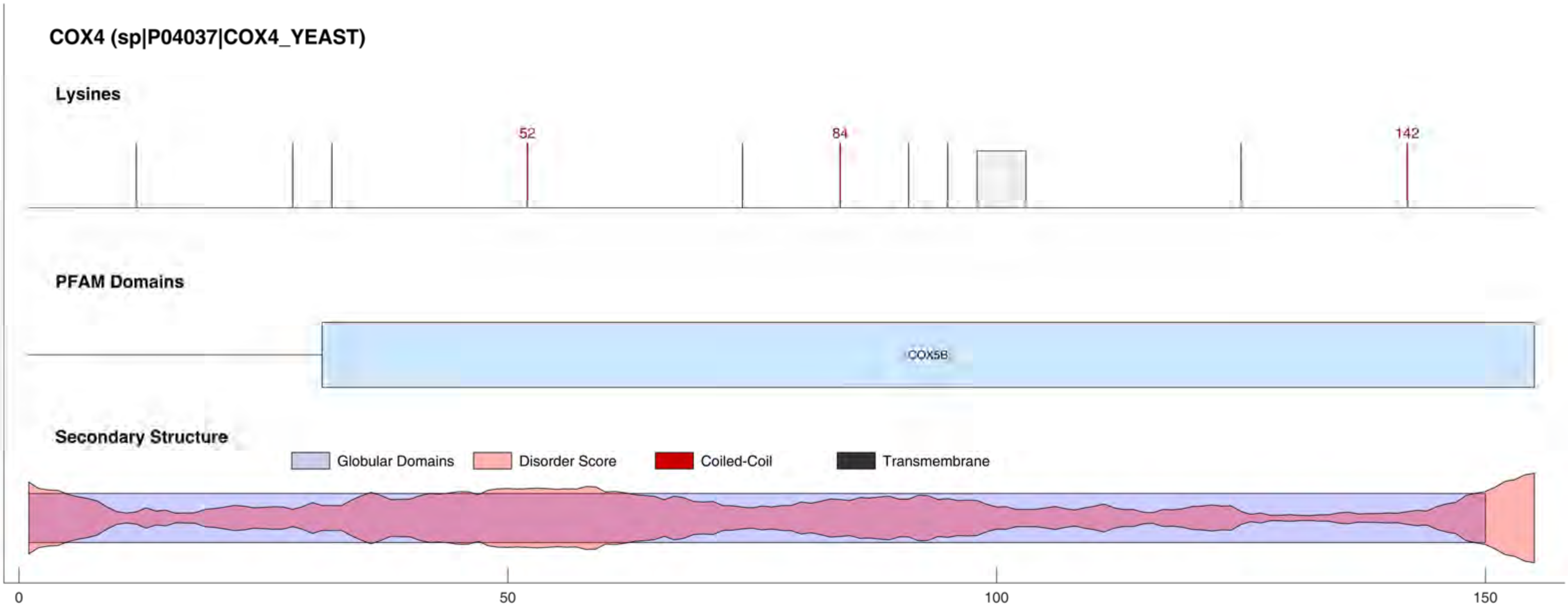
COX4 (sp|P04037|COX4_YEAST)
Lysines
52
84
142
PFAM Domains
COX5B
Secondary Structure
Globular Domains
Disorder Score
Coiled-Coil
Transmembrane
0
50
100
150

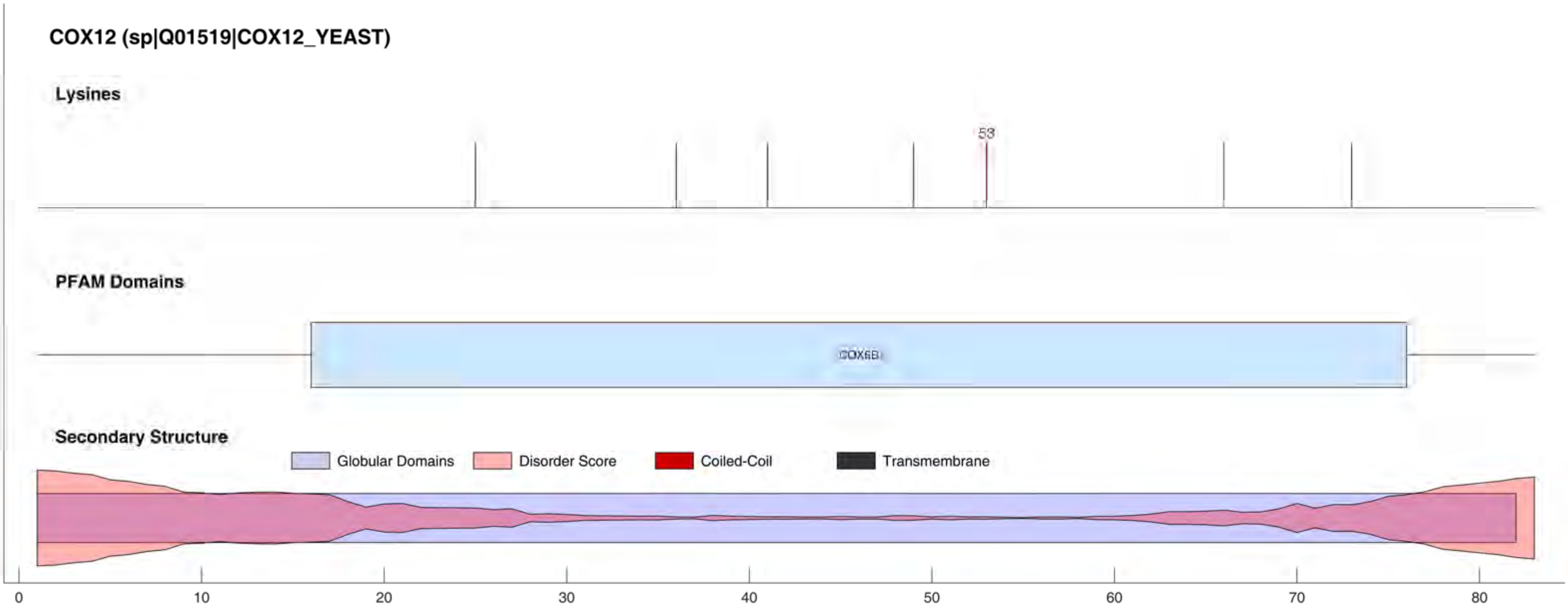
COX12 (sp|Q01519|COX12_YEAST)
Lysines
53
PFAM Domains
COX6B
Secondary Structure
Globular Domains
Disorder Score
Coiled-Coil
Transmembrane
0
10
20
30
40
50
60
70
80

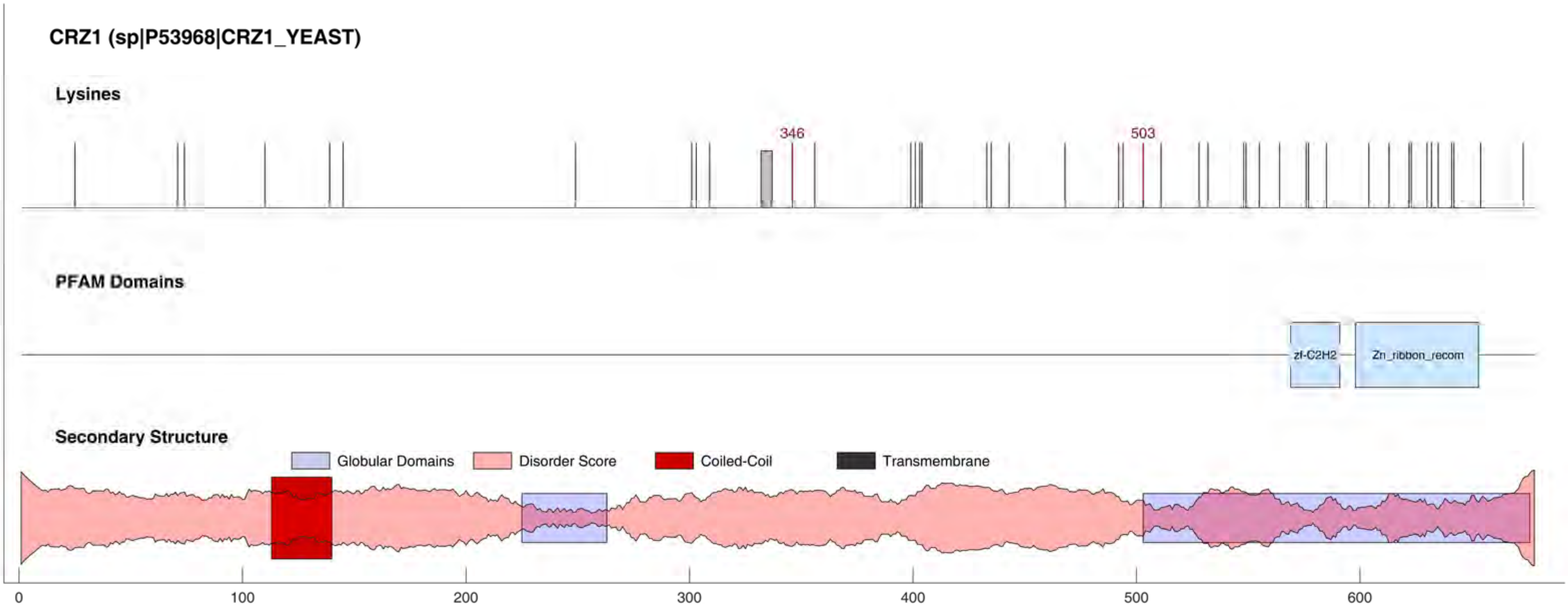
CRZ1 (sp|P53968|CRZ1_YEAST)
Lysines
346
503
PFAM Domains
zf-C2H2
Zn_ribbon_recom
Secondary Structure
Globular Domains
Disorder Score
Coiled-Coil
Transmembrane
0
100
200
300
400
500
600

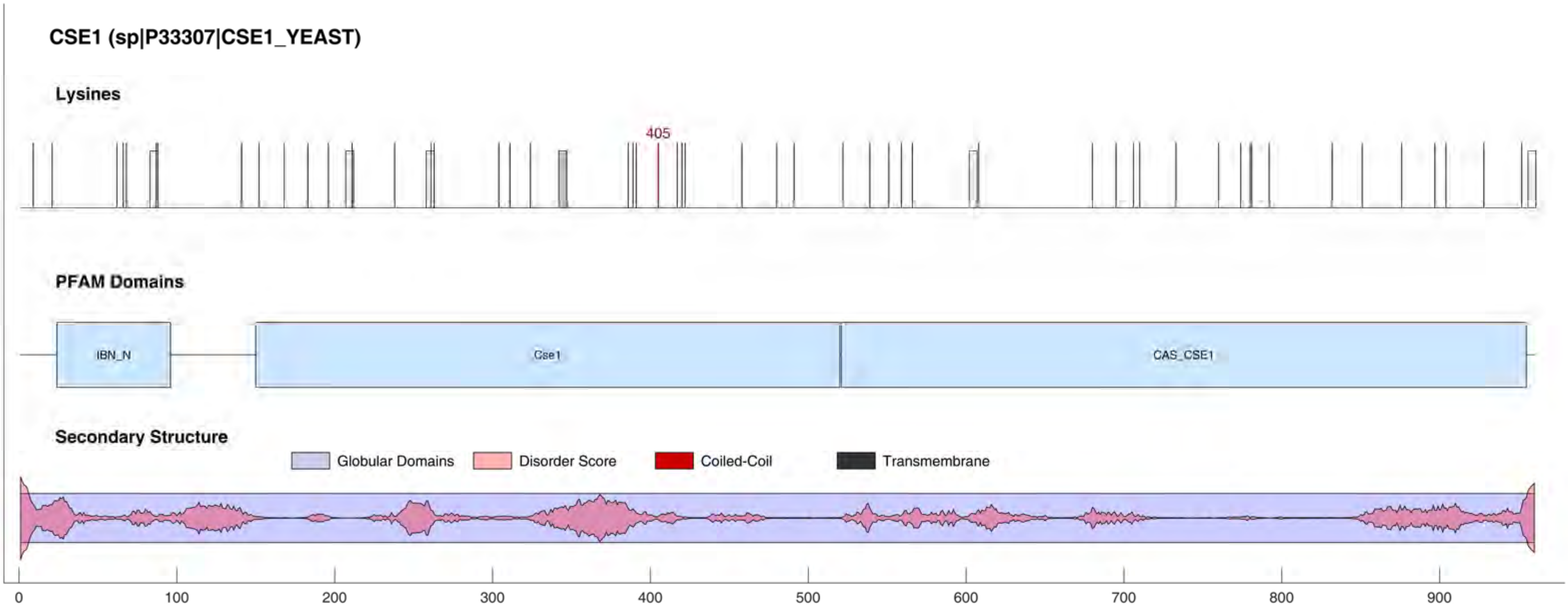
CSE1 (sp|P33307|CSE1_YEAST)
Lysines
405
PFAM Domains
IBN_N
Cse1
CAS_CSE1
Secondary Structure
Globular Domains
Disorder Score
Coiled-Coil
Transmembrane
0
100
200
300
400
500
600
700
800
900

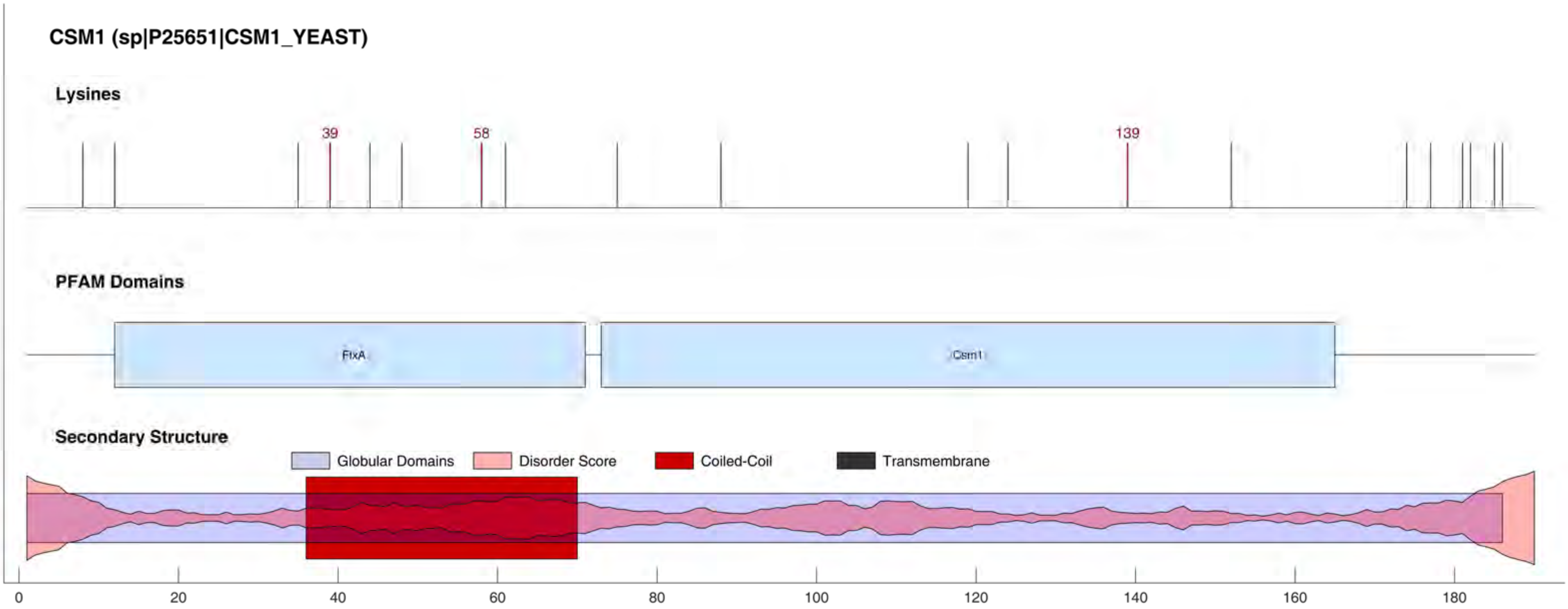

CSM1 (sp|P25651|CSM1_YEAST)
Lysines
39
58
139
PFAM Domains
FtxA
Csm1
Secondary Structure
Globular Domains
Disorder Score
Coiled-Coil
Transmembrane
0
20
40
60
80
100
120
140
160
180

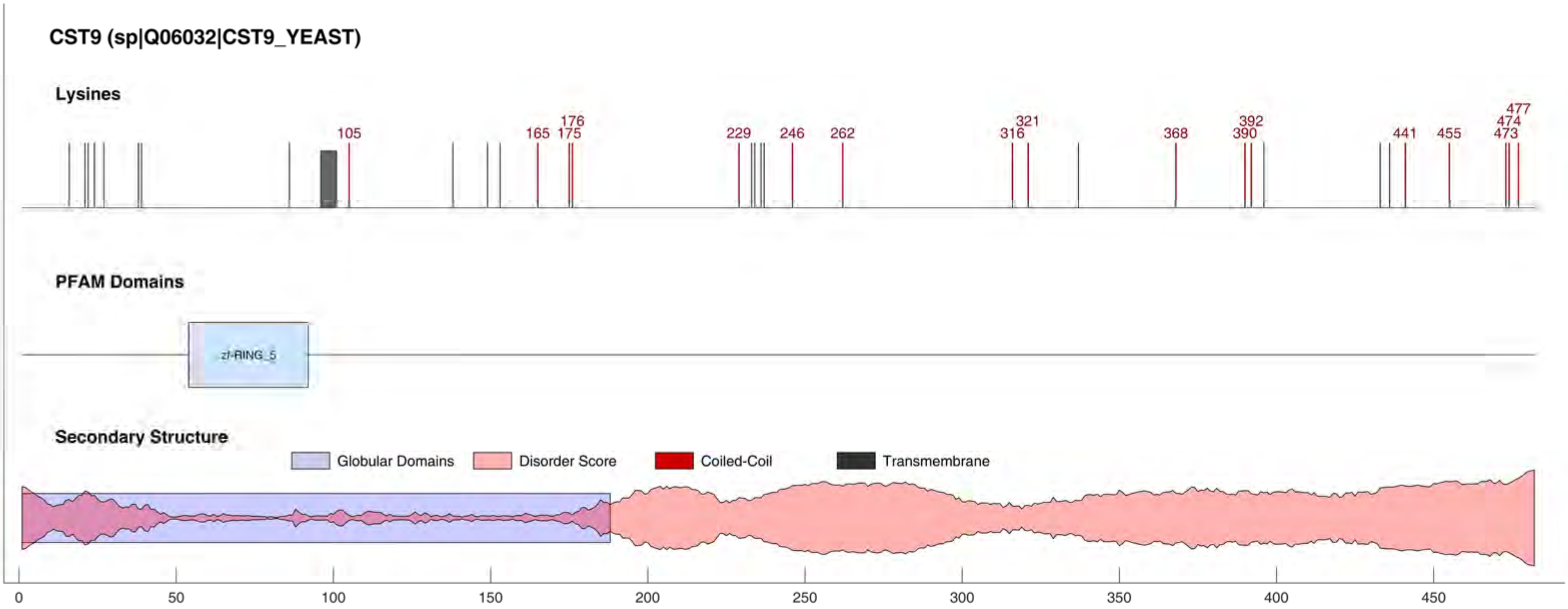

CST9 (sp|Q06032|CST9_YEAST)
Lysines
105
165
175
176
229
246
262
316
321
368
390
392
441
455
473
474
477
PFAM Domains
zf-RING_5
Secondary Structure
Globular Domains
Disorder Score
Coiled-Coil
Transmembrane
0
50
100
150
200
250
300
350
400
450

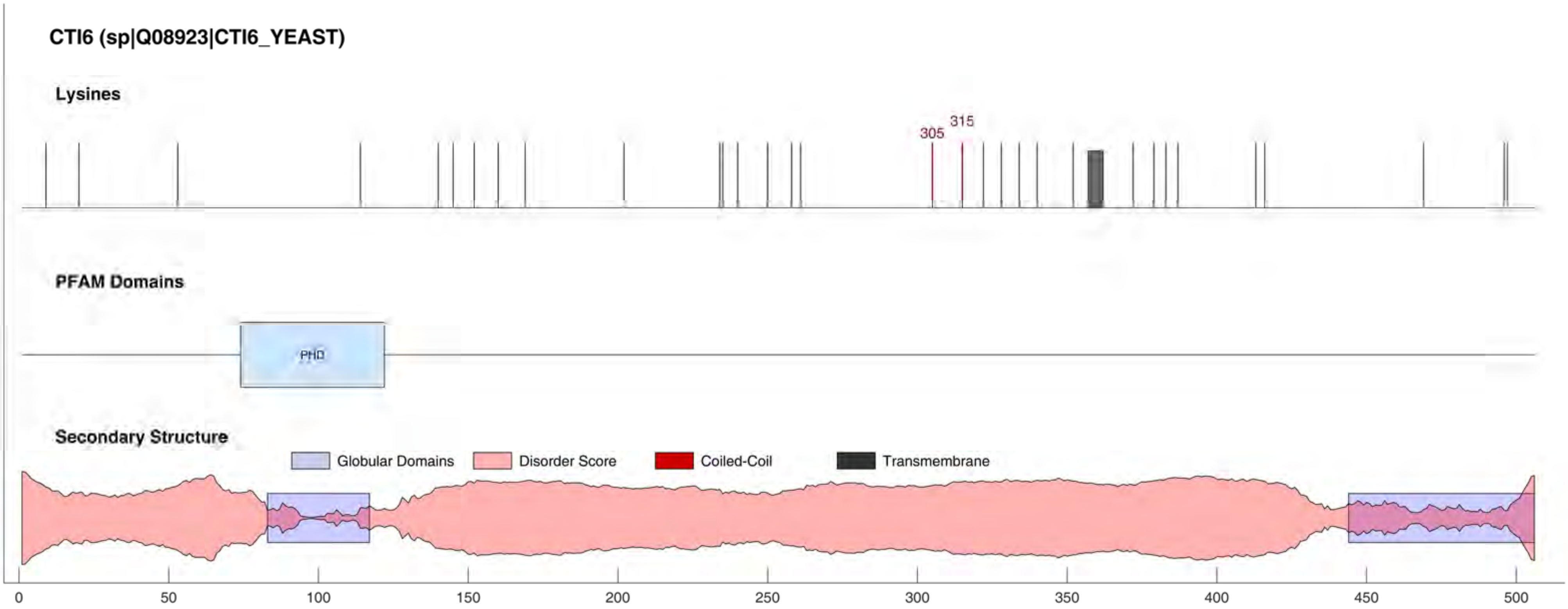

CTI6 (sp|Q08923|CTI6_YEAST)
Lysines
305
315
PFAM Domains
PHD
Secondary Structure
Globular Domains
Disorder Score
Coiled-Coil
Transmembrane
0
50
100
150
200
250
300
350
400
450
500

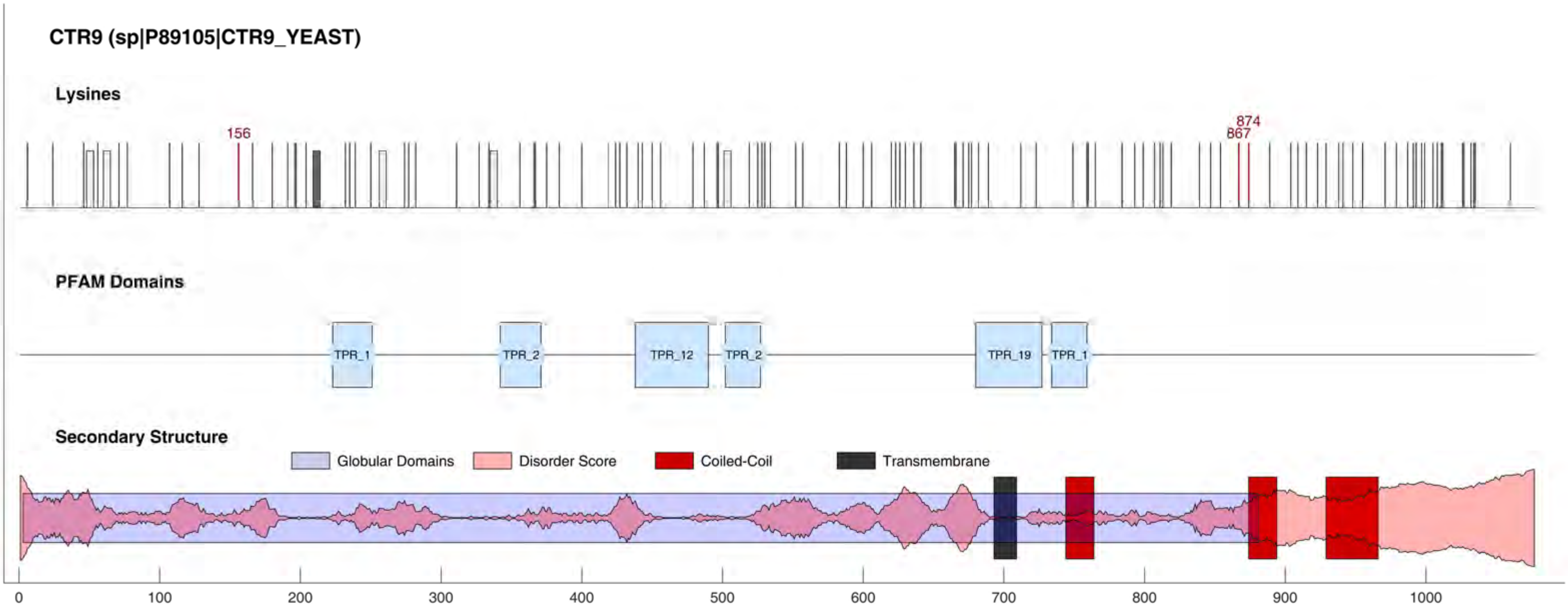
CTR9 (sp|P89105|CTR9_YEAST)
Lysines
156
874
867
PFAM Domains
TPR_1
TPR_2
TPR_12
TPR_2
TPR_19
TPR_1
Secondary Structure
Globular Domains
Disorder Score
Coiled-Coil
Transmembrane
0
100
200
300
400
500
600
700
800
900
1000

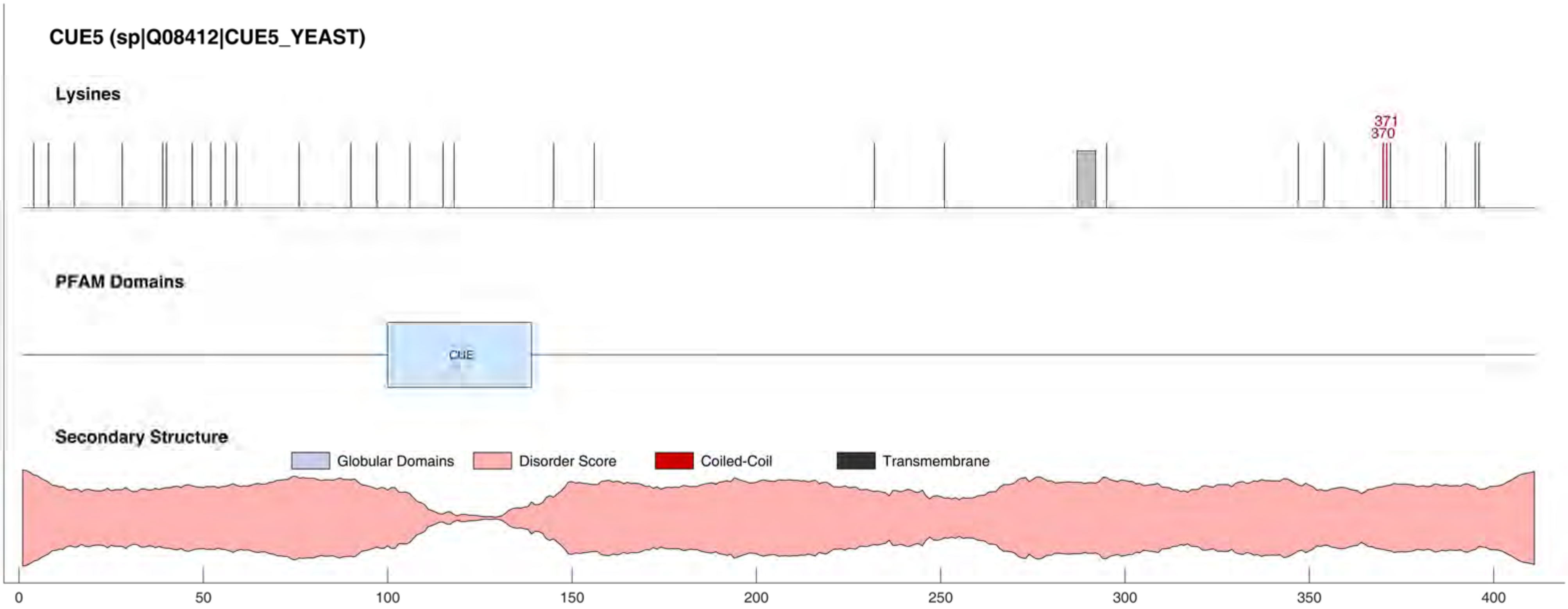

CUE5 (sp|Q08412|CUE5_YEAST)
Lysines
371
370
PFAM Domains
CUE
Secondary Structure
Globular Domains
Disorder Score
Coiled-Coil
Transmembrane
0
50
100
150
200
250
300
350
400

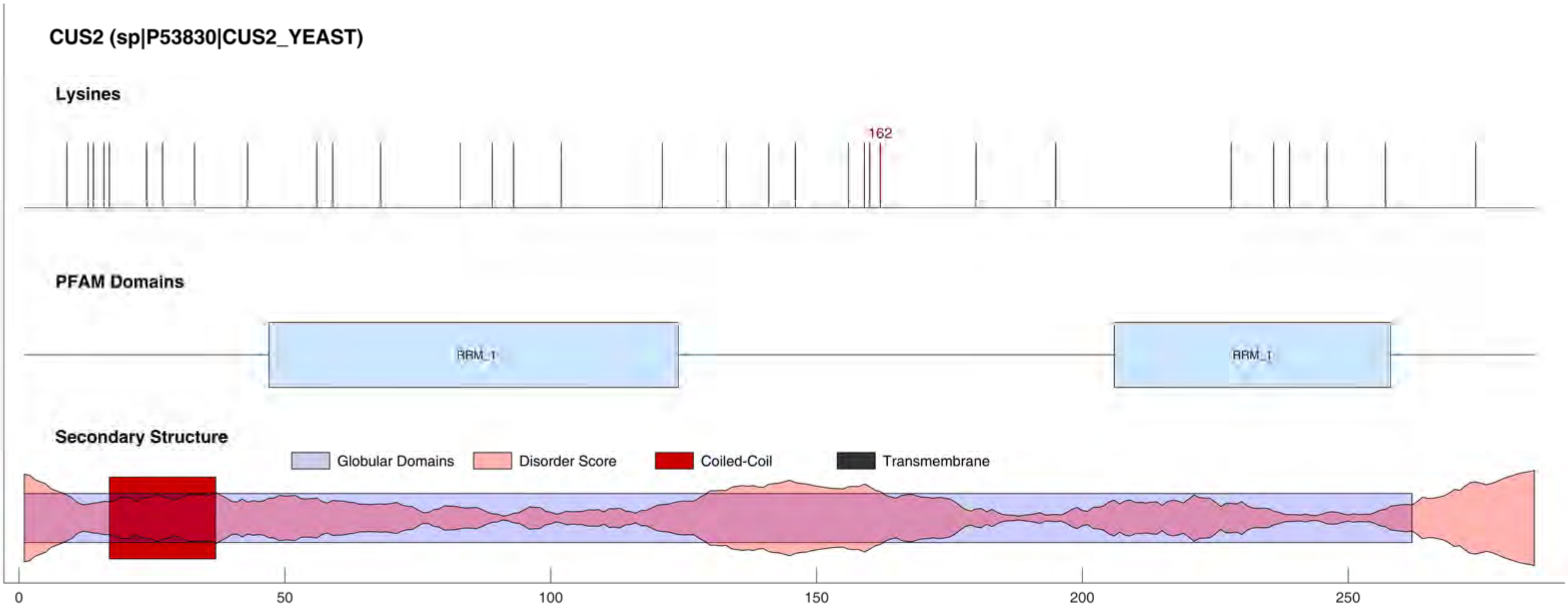

CUS2 (sp|P53830|CUS2_YEAST)
Lysines
162
PFAM Domains
RRM_1
RRM_1
Secondary Structure
Globular Domains
Disorder Score
Coiled-Coil
Transmembrane
0
50
100
150
200
250

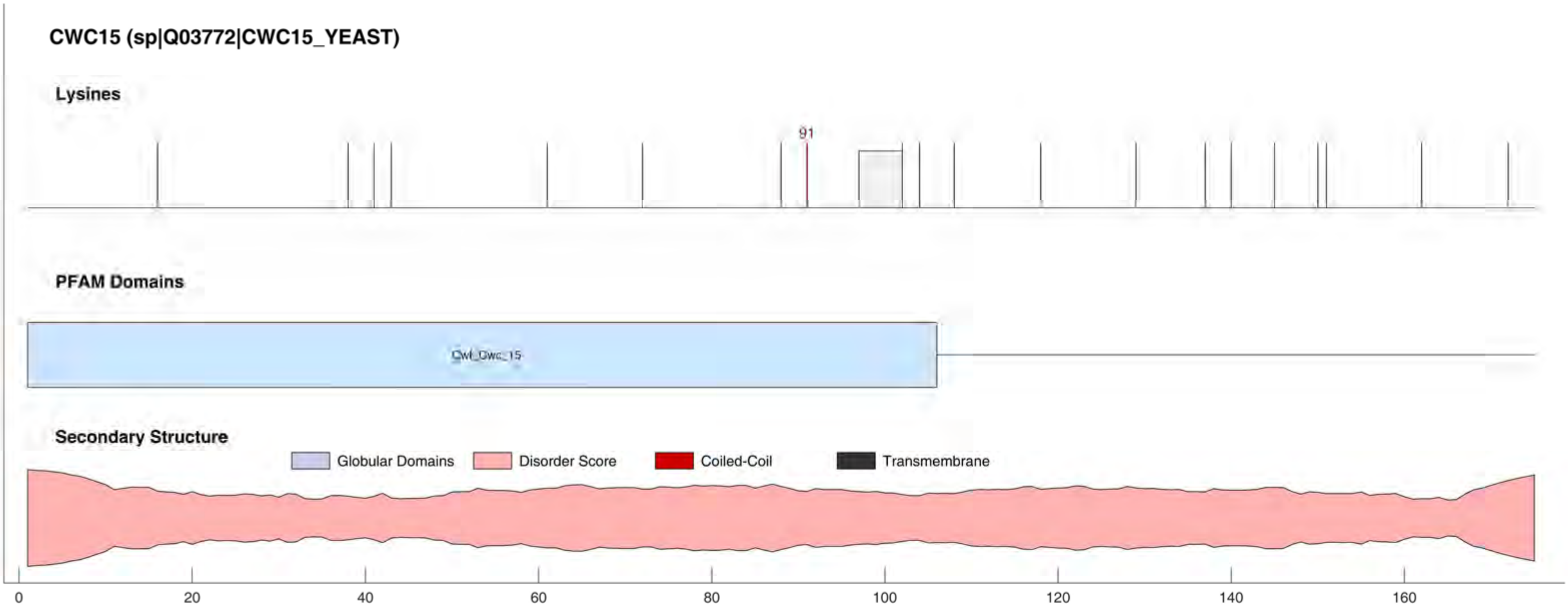
CWC15 (sp|Q03772|CWC15_YEAST)
Lysines
91
PFAM Domains
Cwf_Cwc_15
Secondary Structure
Globular Domains
Disorder Score
Coiled-Coil
Transmembrane
0
20
40
60
80
100
120
140
160

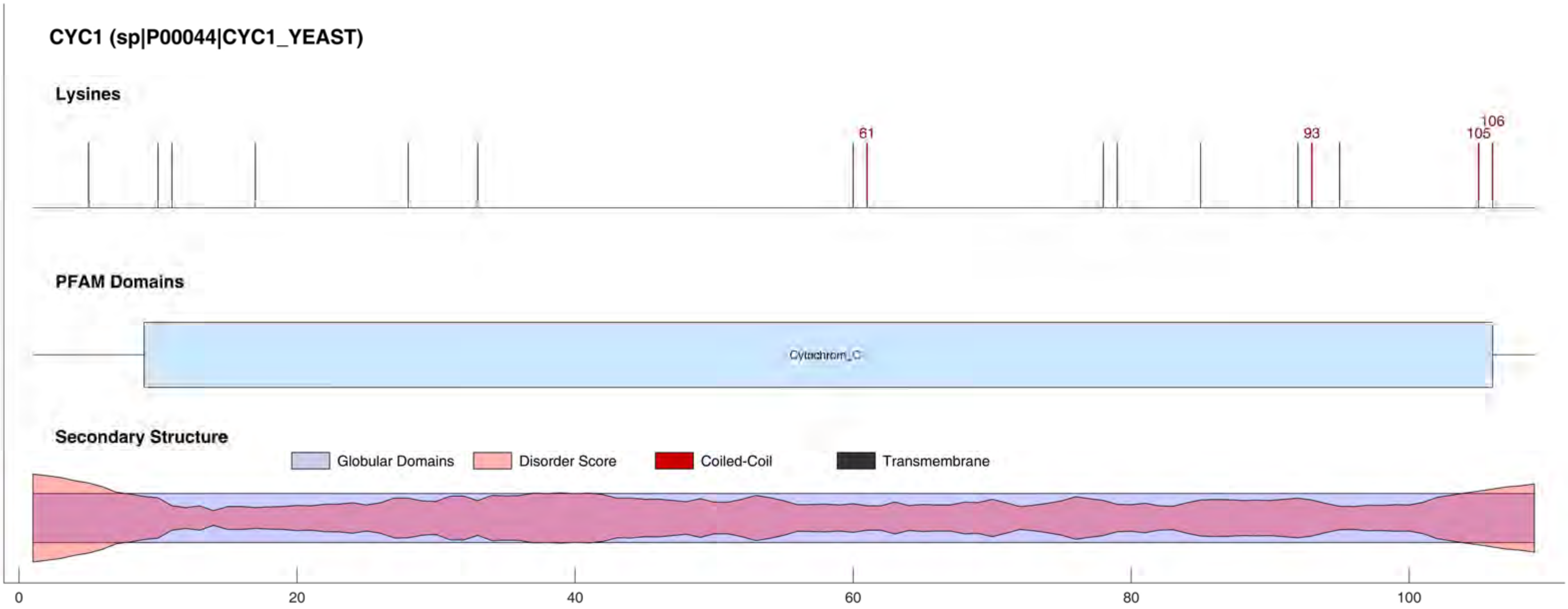

CYC1 (sp|P00044|CYC1_YEAST)
Lysines
61
93
105
106
PFAM Domains
Cytochrom_C
Secondary Structure
Globular Domains
Disorder Score
Coiled-Coil
Transmembrane
0
20
40
60
80
100

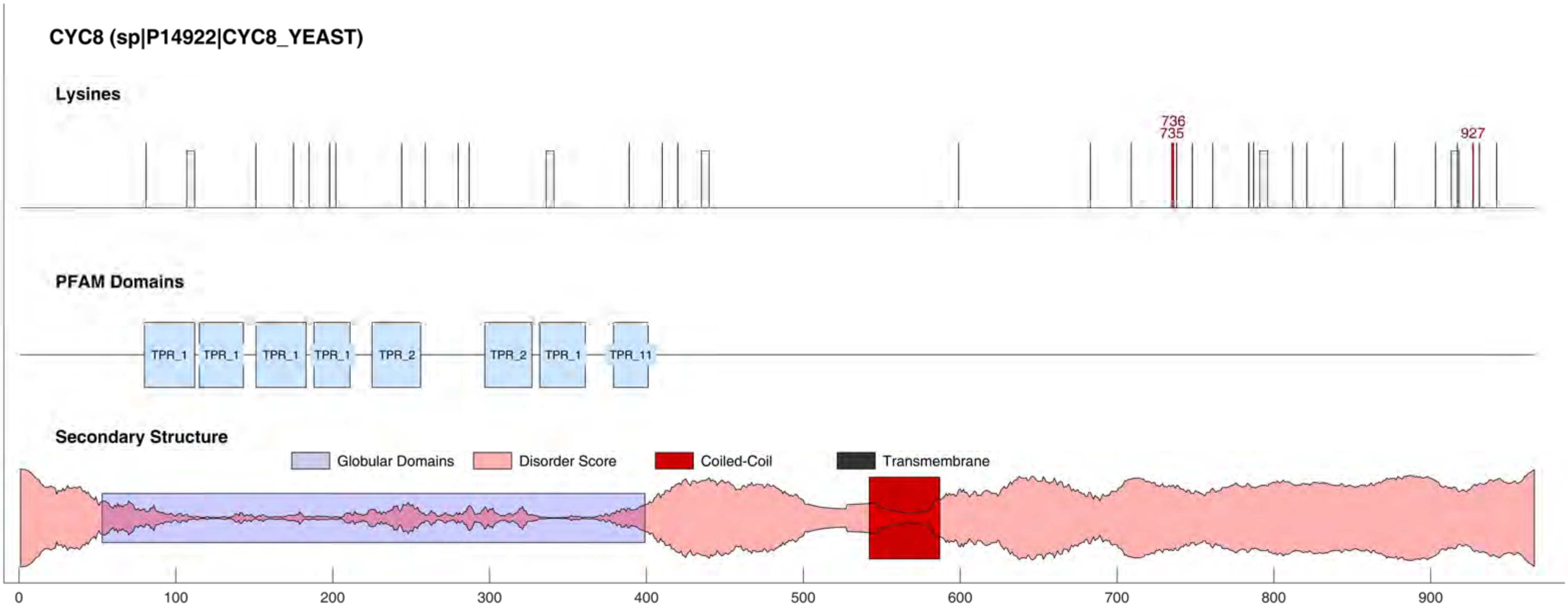

CYC8 (sp|P14922|CYC8_YEAST)
Lysines
736
735
927
PFAM Domains
TPR_1
TPR_1
TPR_1
TPR_1
TPR_2
TPR_2
TPR_1
TPR_11
Secondary Structure
Globular Domains
Disorder Score
Coiled-Coil
Transmembrane
0
100
200
300
400
500
600
700
800
900

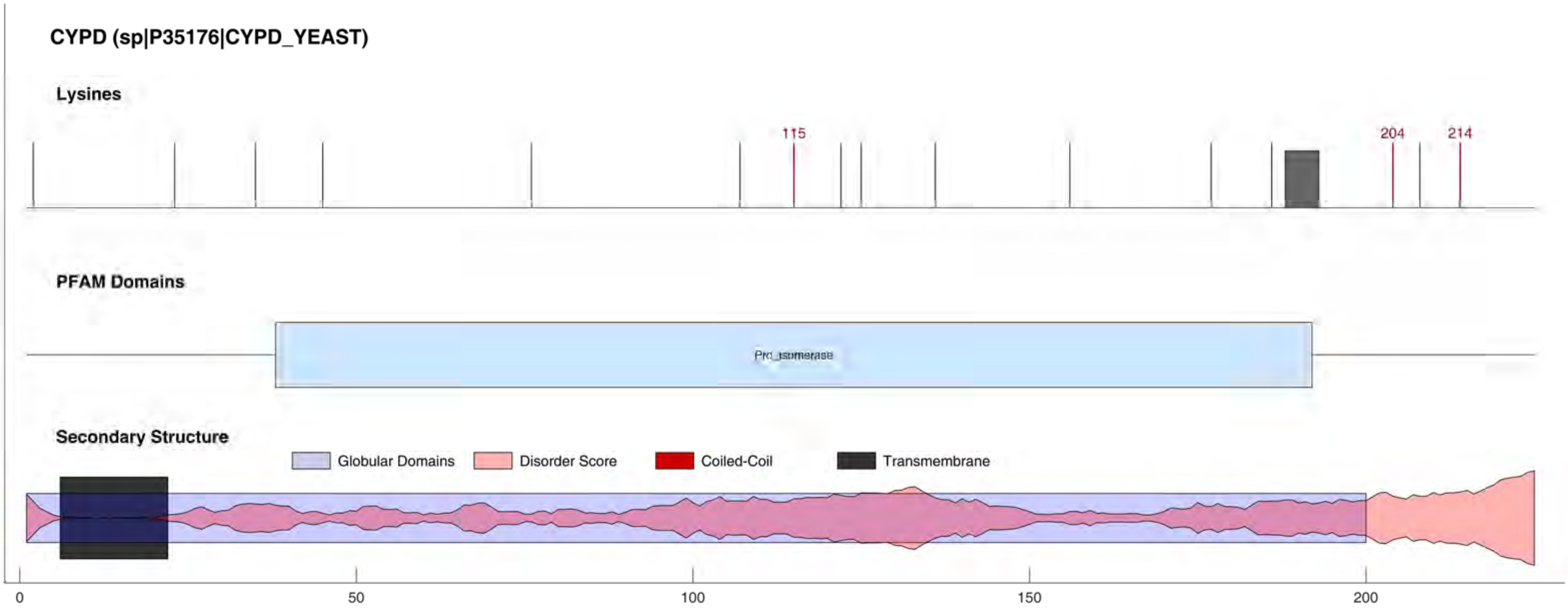
CYPD (sp|P35176|CYPD_YEAST)
Lysines
115
204
214
PFAM Domains
Pro_isomerase
Secondary Structure
Globular Domains
Disorder Score
Coiled-Coil
Transmembrane
0
50
100
150
200

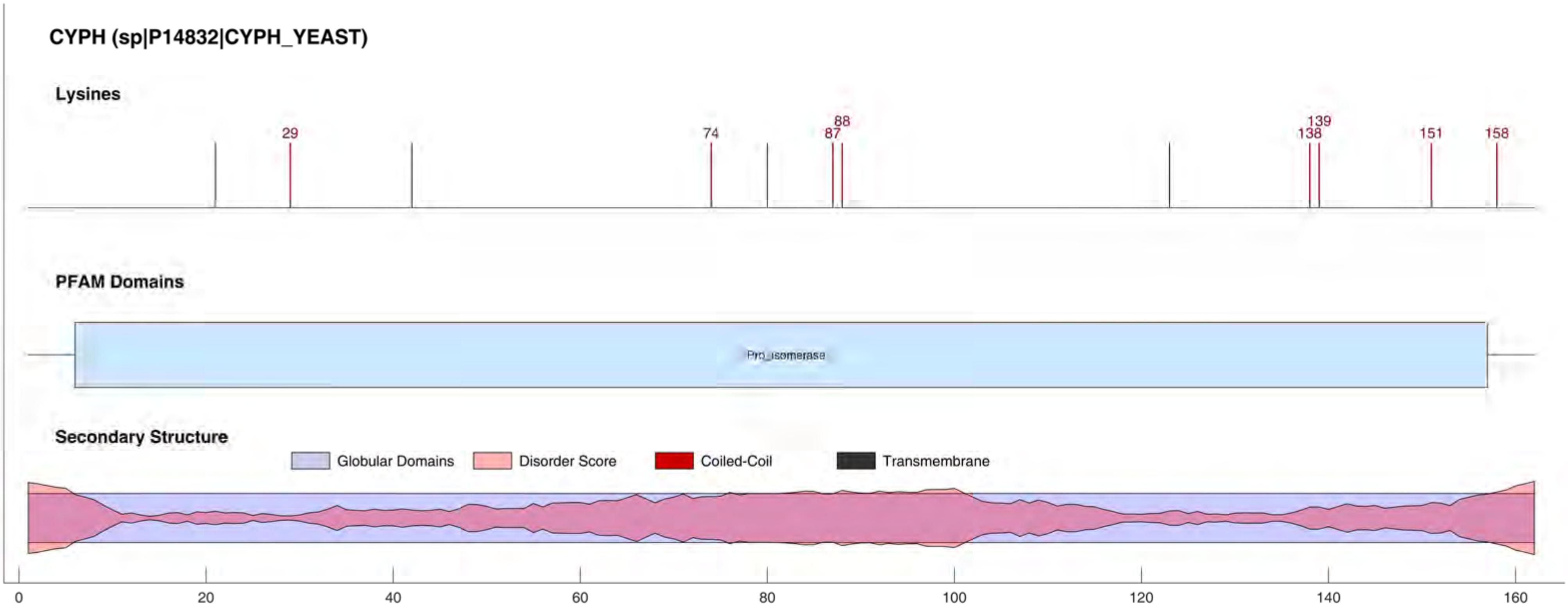
CYPH (sp|P14832|CYPH_YEAST)
Lysines
29
74
88
87
139
138
151
158
PFAM Domains
Pro_isomerase
Secondary Structure
Globular Domains
Disorder Score
Coiled-Coil
Transmembrane
0
20
40
60
80
100
120
140
160

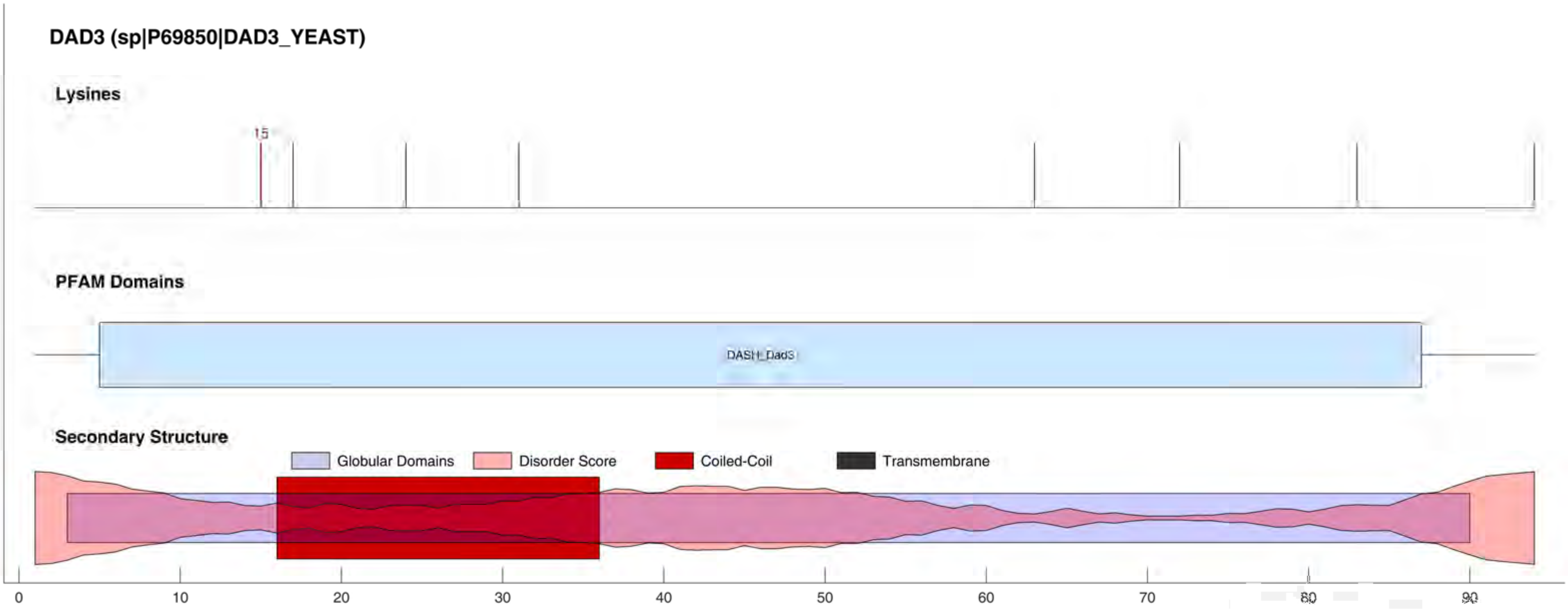
DAD3 (sp|P69850|DAD3_YEAST)
Lysines
15
PFAM Domains
DASH_Dad3
Secondary Structure
Globular Domains
Disorder Score
Coiled-Coil
Transmembrane
0
10
20
30
40
50
60
70
80
90

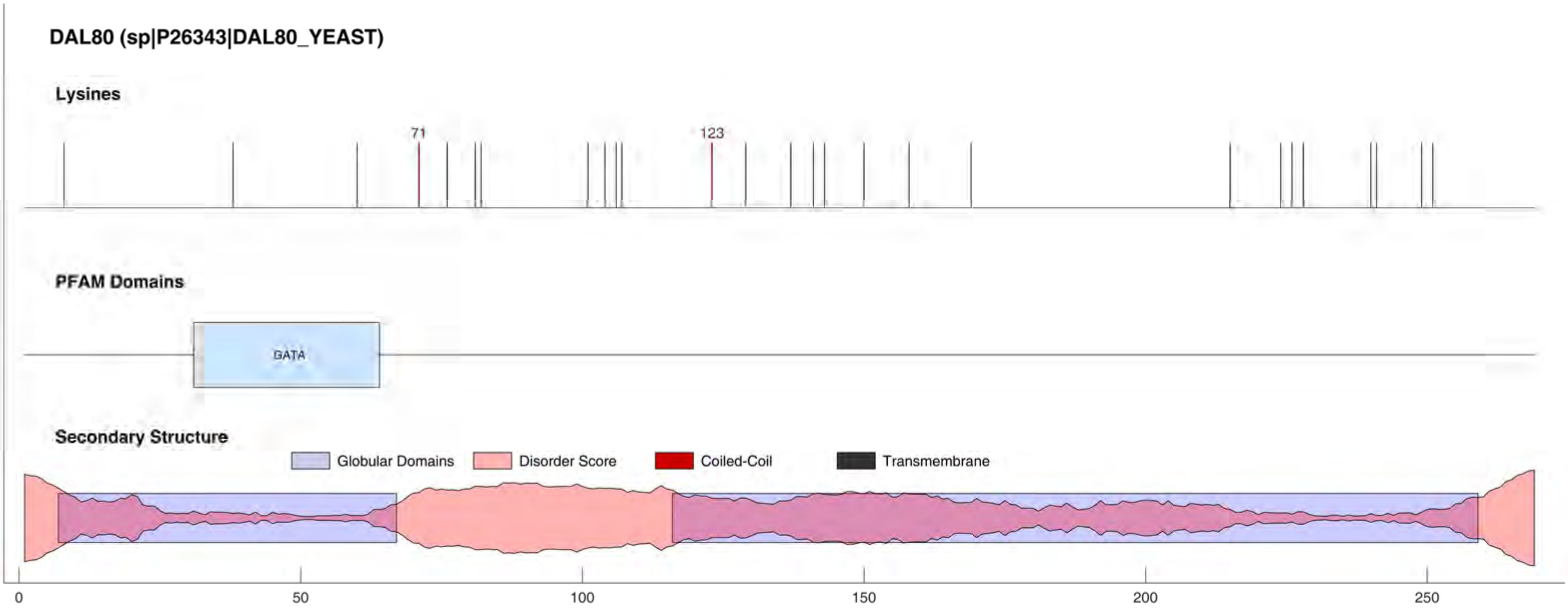

DAL80 (sp|P26343|DAL80_YEAST)
Lysines
71
123
PFAM Domains
GATA
Secondary Structure
Globular Domains
Disorder Score
Coiled-Coil
Transmembrane
0
50
100
150
200
250

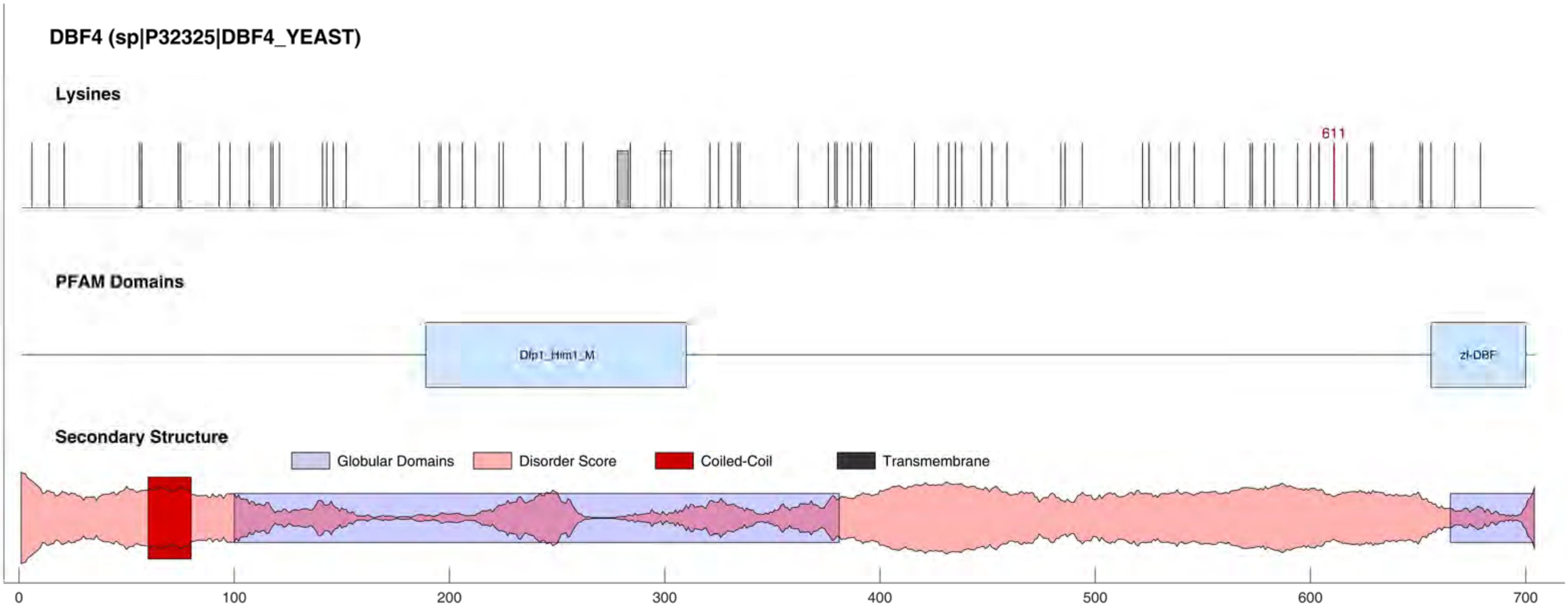
DBF4 (sp|P32325|DBF4_YEAST)
Lysines
611
PFAM Domains
Dfp1_Him1_M
zf-DBF
Secondary Structure
Globular Domains
Disorder Score
Coiled-Coil
Transmembrane
0
100
200
300
400
500
600
700

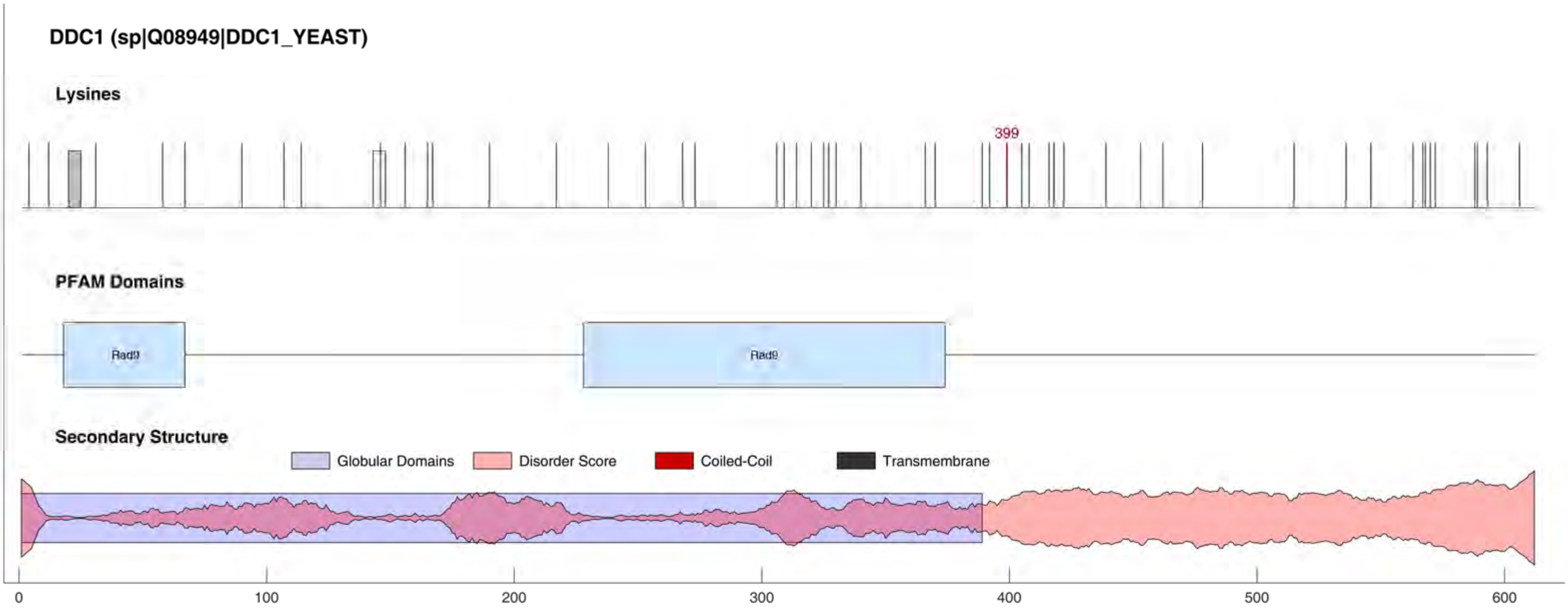
DDC1 (sp|Q08949|DDC1_YEAST)
Lysines
399
PFAM Domains
Rad9
Rad9
Secondary Structure
Globular Domains
Disorder Score
Coiled-Coil
Transmembrane
0
100
200
300
400
500
600

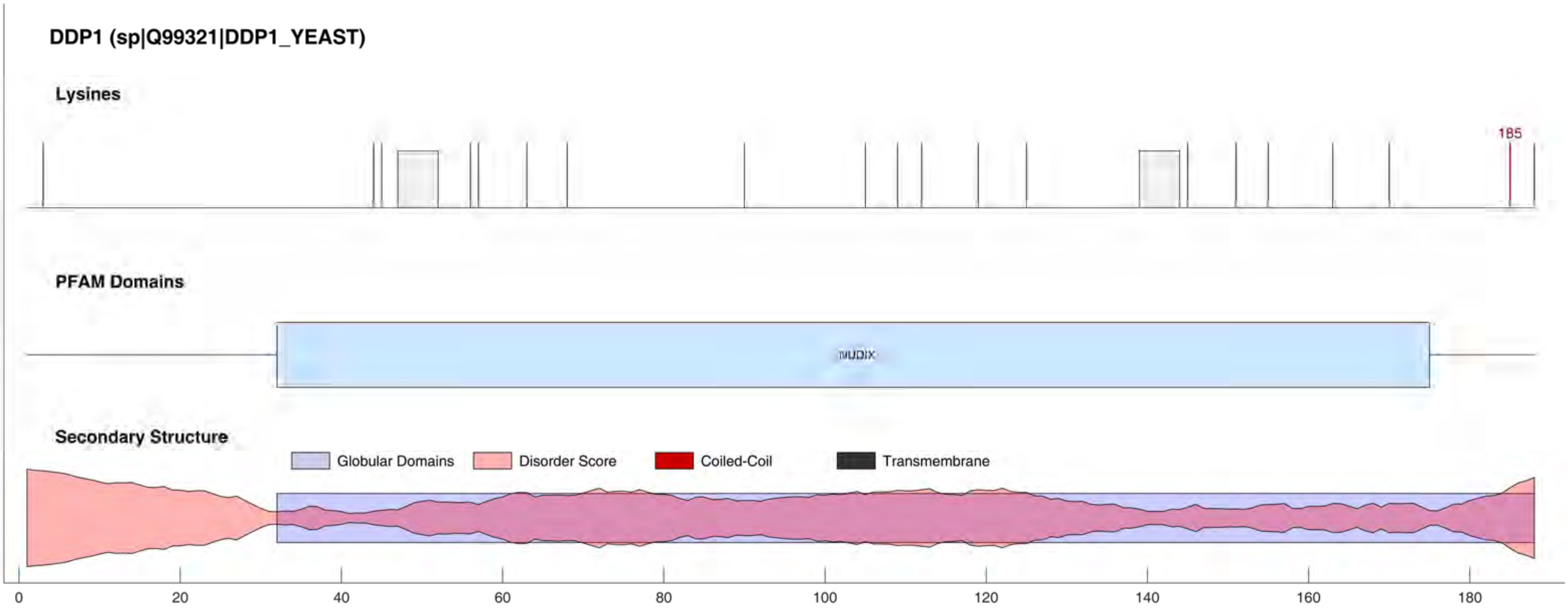
DDP1 (sp|Q99321|DDP1_YEAST)
Lysines
185
PFAM Domains
NUDIX
Secondary Structure
Globular Domains
Disorder Score
Coiled-Coil
Transmembrane
0
20
40
60
80
100
120
140
160
180

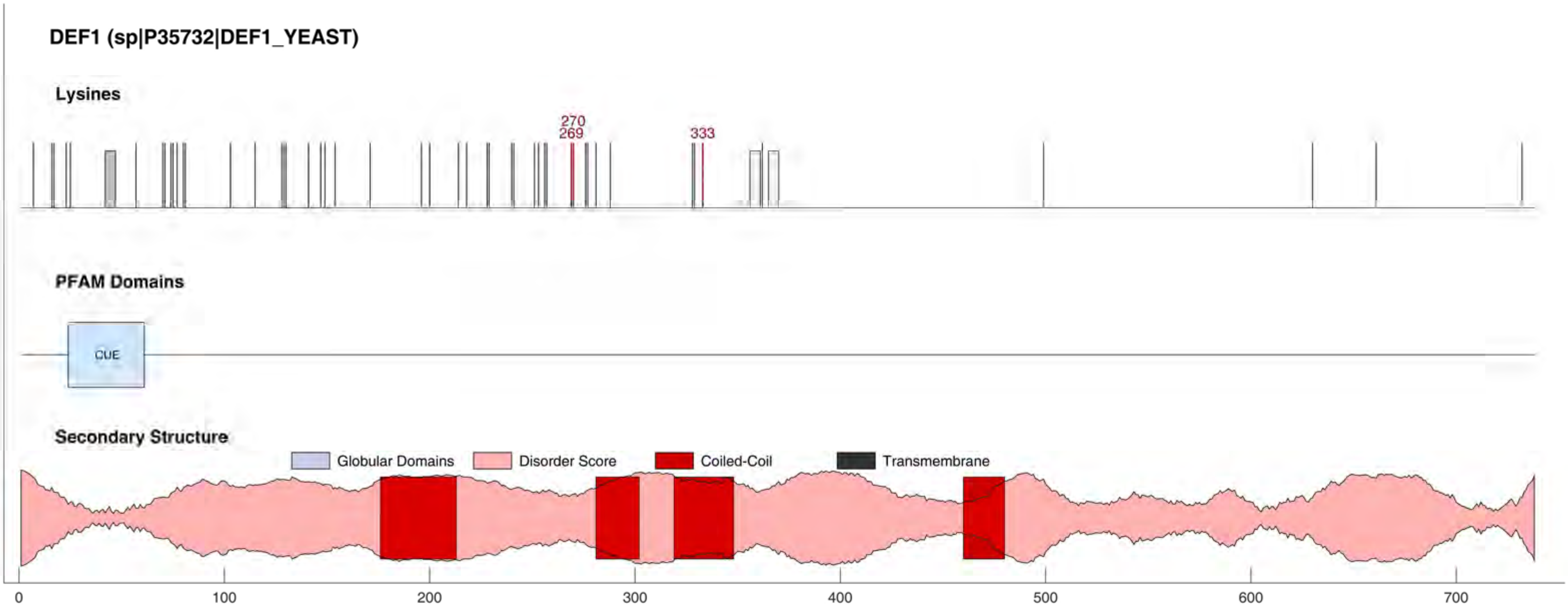
DEF1 (sp|P35732|DEF1_YEAST)
Lysines
270
269
333
PFAM Domains
CUE
Secondary Structure
Globular Domains
Disorder Score
Coiled-Coil
Transmembrane
0
100
200
300
400
500
600
700

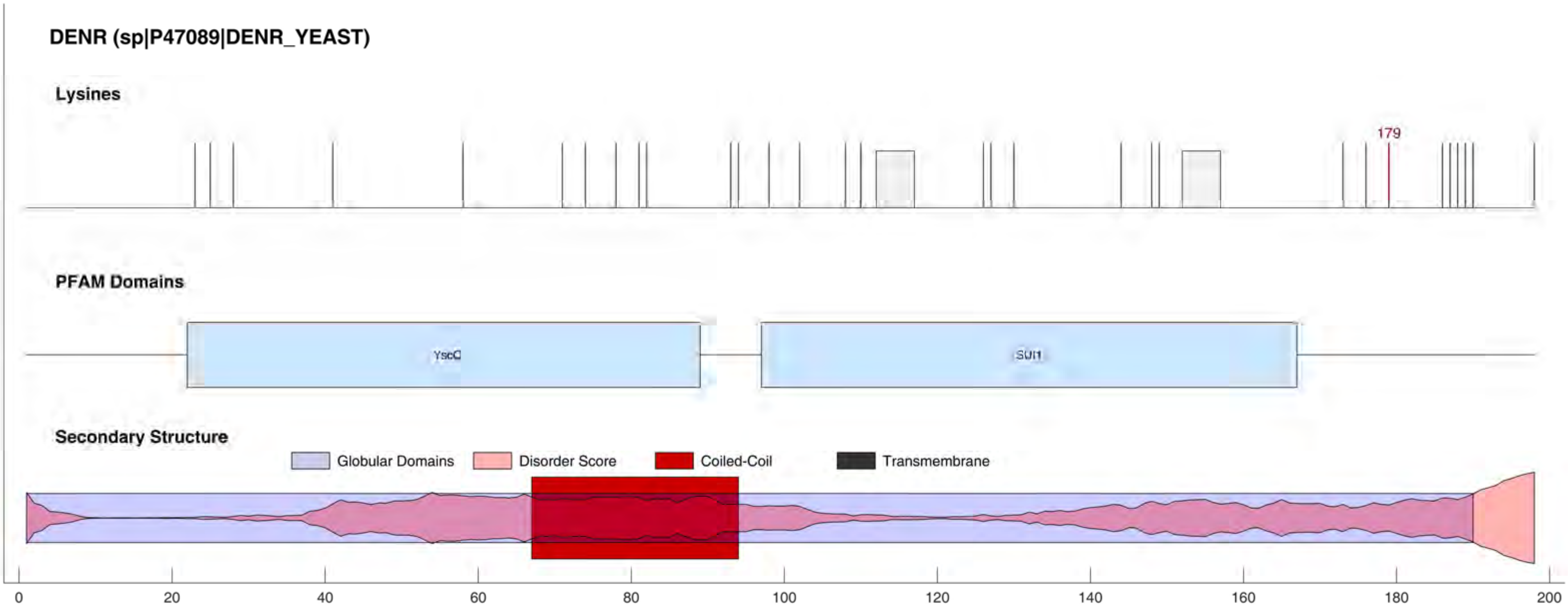

DENR (sp|P47089|DENR_YEAST)
Lysines
179
PFAM Domains
YscO
SUI1
Secondary Structure
Globular Domains
Disorder Score
Coiled-Coil
Transmembrane
0
20
40
60
80
100
120
140
160
180
200

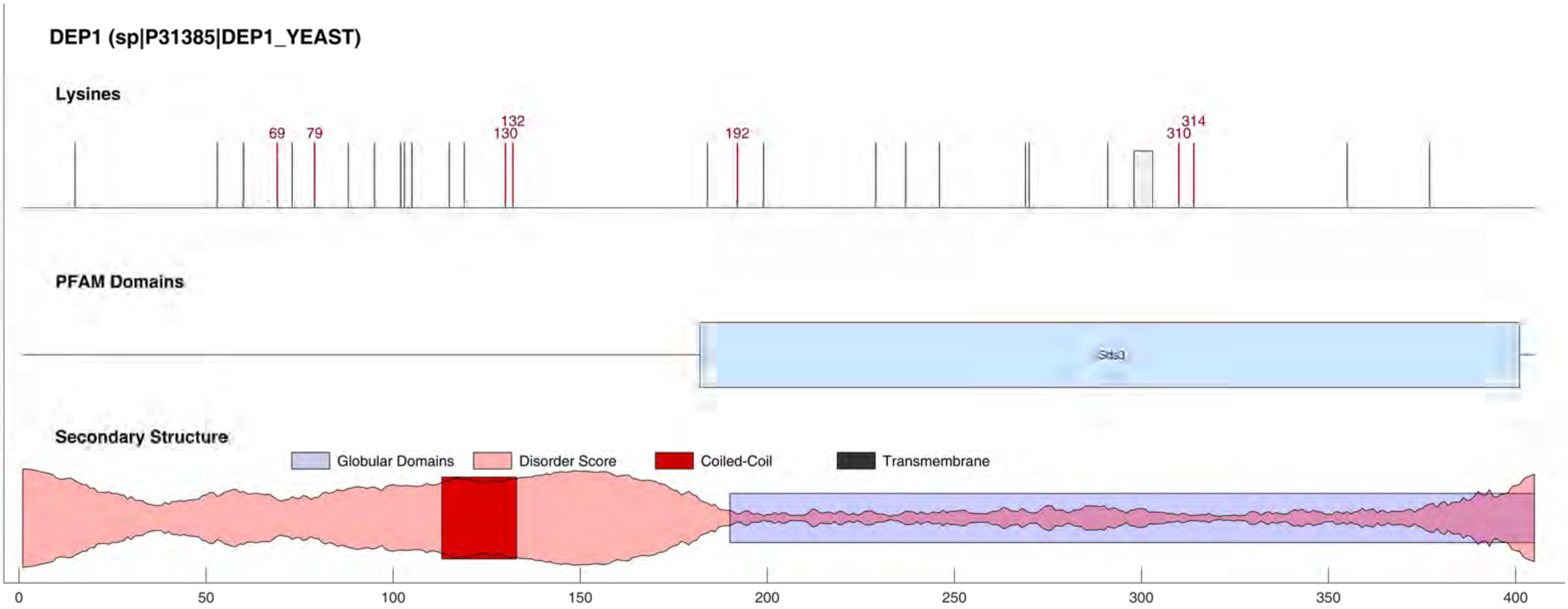

DEP1 (sp|P31385|DEP1_YEAST)
Lysines
69
79
132
130
192
314
310
PFAM Domains
Sds3
Secondary Structure
Globular Domains
Disorder Score
Coiled-Coil
Transmembrane
0
50
100
150
200
250
300
350
400

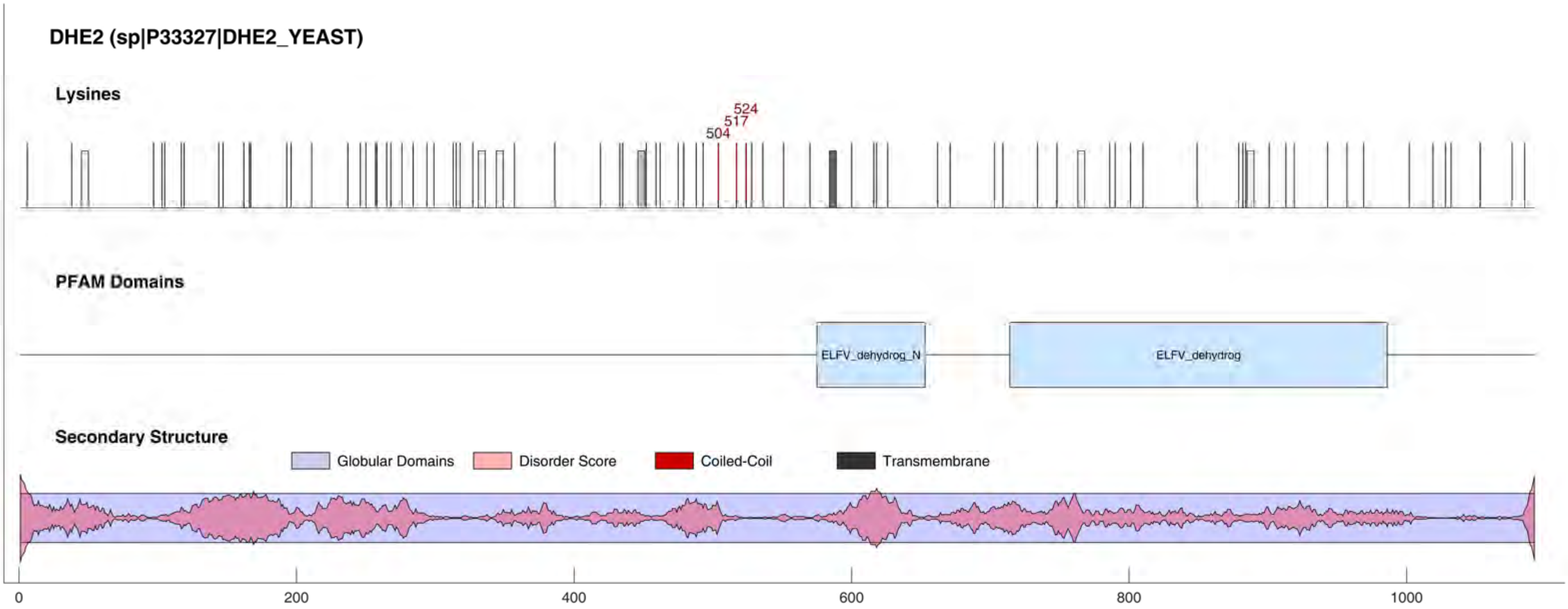
DHE2 (sp|P33327|DHE2_YEAST)
Lysines
524
517
504
PFAM Domains
ELFV_dehydrog_N
ELFV_dehydrog
Secondary Structure
Globular Domains
Disorder Score
Coiled-Coil
Transmembrane
0
200
400
600
800
1000

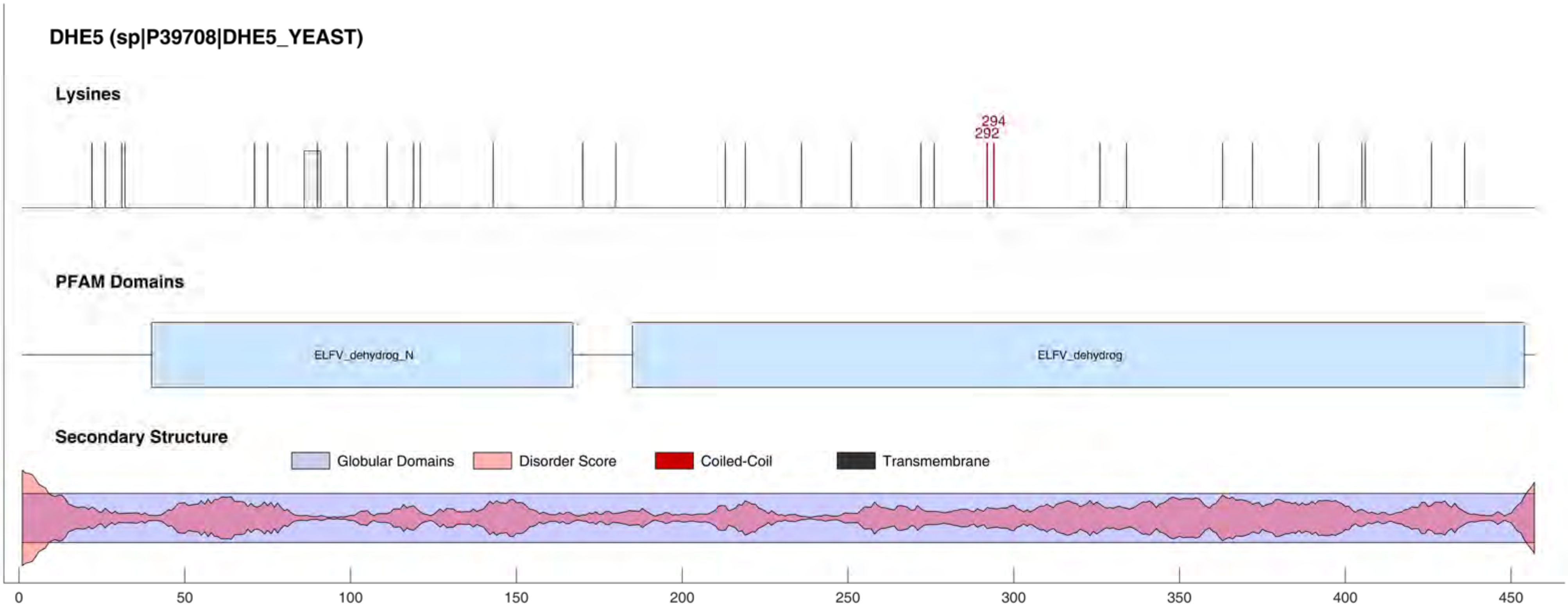
DHE5 (sp|P39708|DHE5_YEAST)
Lysines
294
292
PFAM Domains
ELFV_dehydrog_N
ELFV_dehydrog
Secondary Structure
Globular Domains
Disorder Score
Coiled-Coil
Transmembrane
0
50
100
150
200
250
300
350
400
450

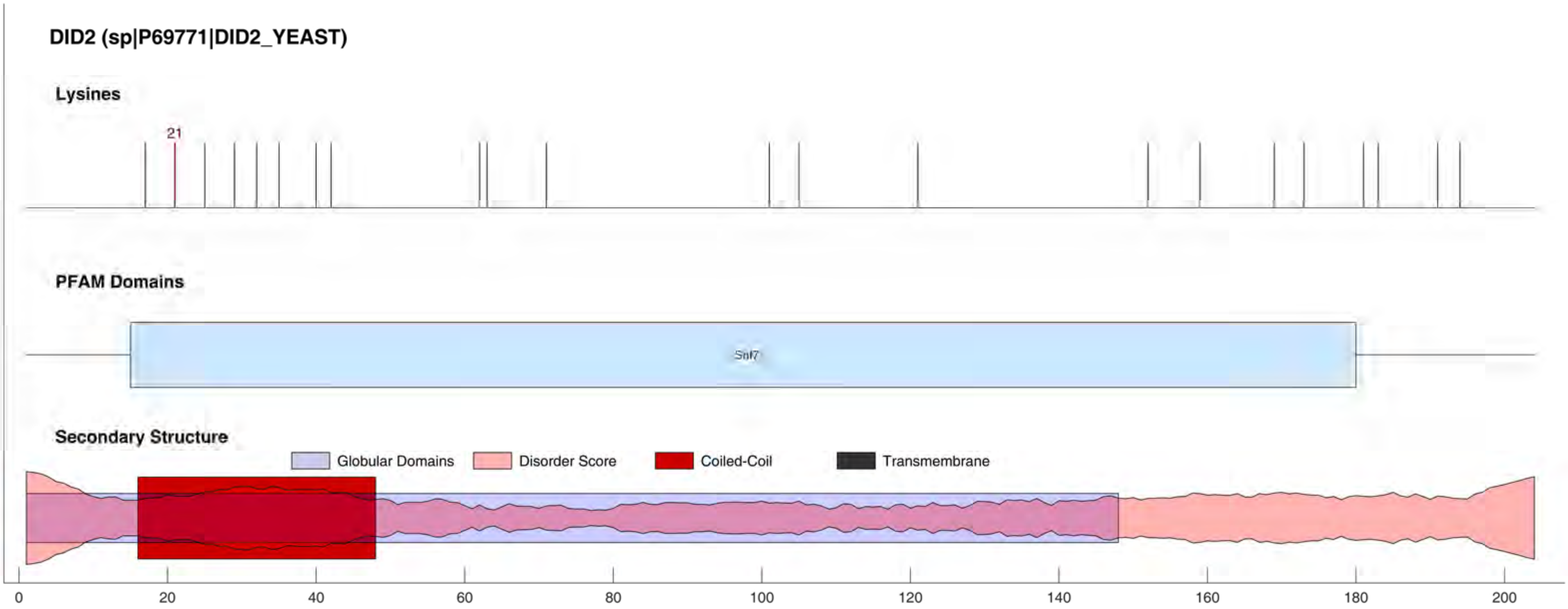
DID2 (sp|P69771|DID2_YEAST)
Lysines
21
PFAM Domains
Snf7
Secondary Structure
Globular Domains
Disorder Score
Coiled-Coil
Transmembrane
0
20
40
60
80
100
120
140
160
180
200

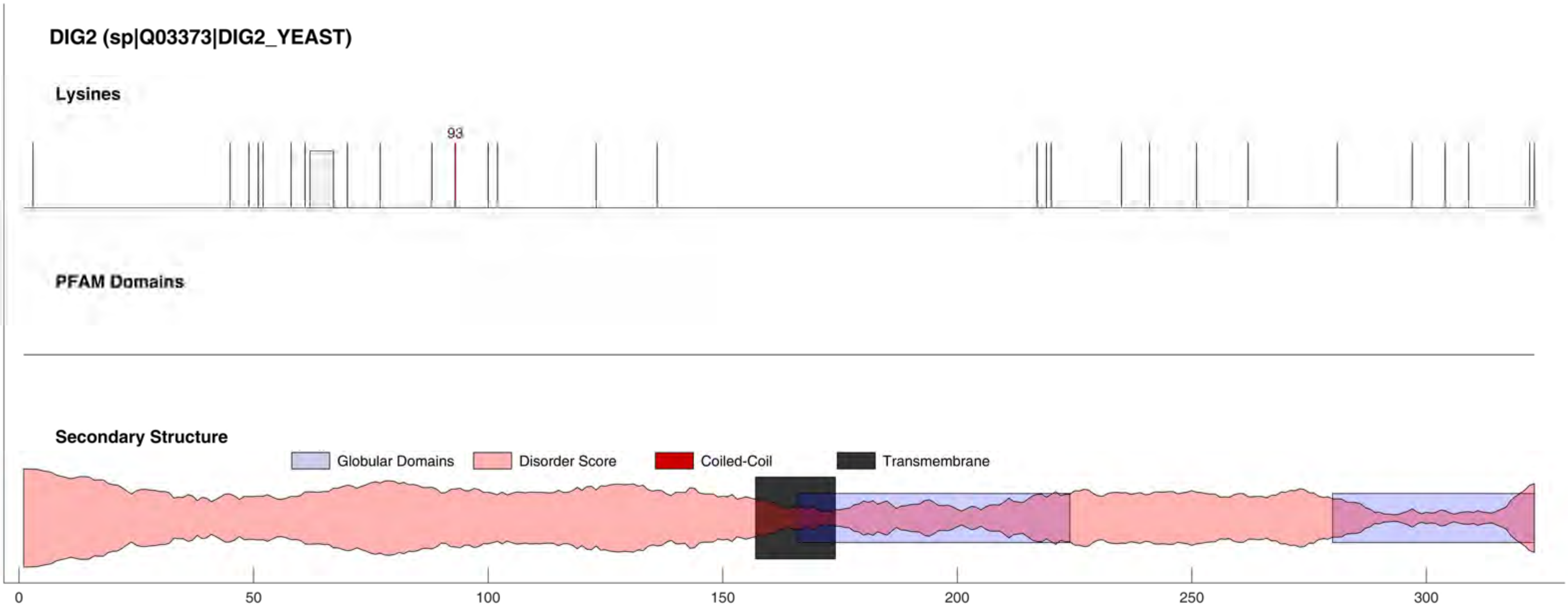
DIG2 (sp|Q03373|DIG2_YEAST)
Lysines
93
PFAM Domains
Secondary Structure
Globular Domains
Disorder Score
Coiled-Coil
Transmembrane
0
50
100
150
200
250
300

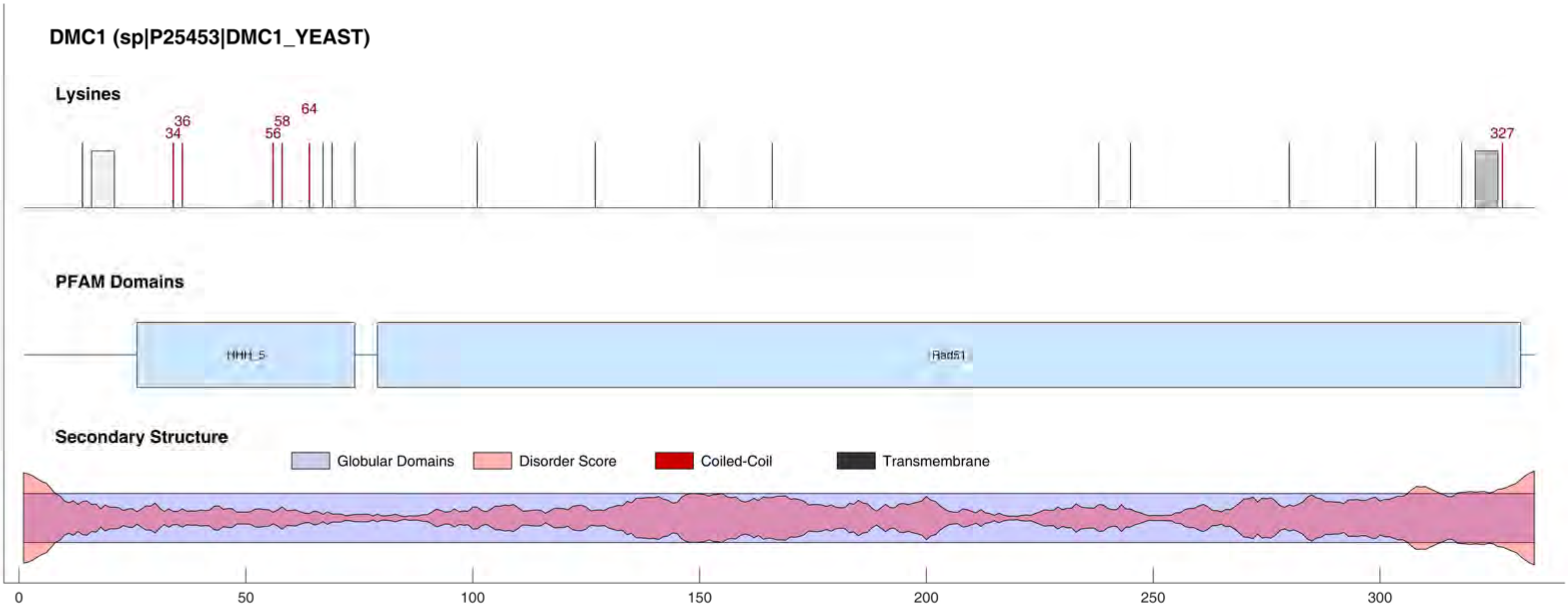
DMC1 (sp|P25453|DMC1_YEAST)
Lysines
34
36
56
58
64
327
PFAM Domains
HHH_5
Rad51
Secondary Structure
Globular Domains
Disorder Score
Coiled-Coil
Transmembrane
0
50
100
150
200
250
300

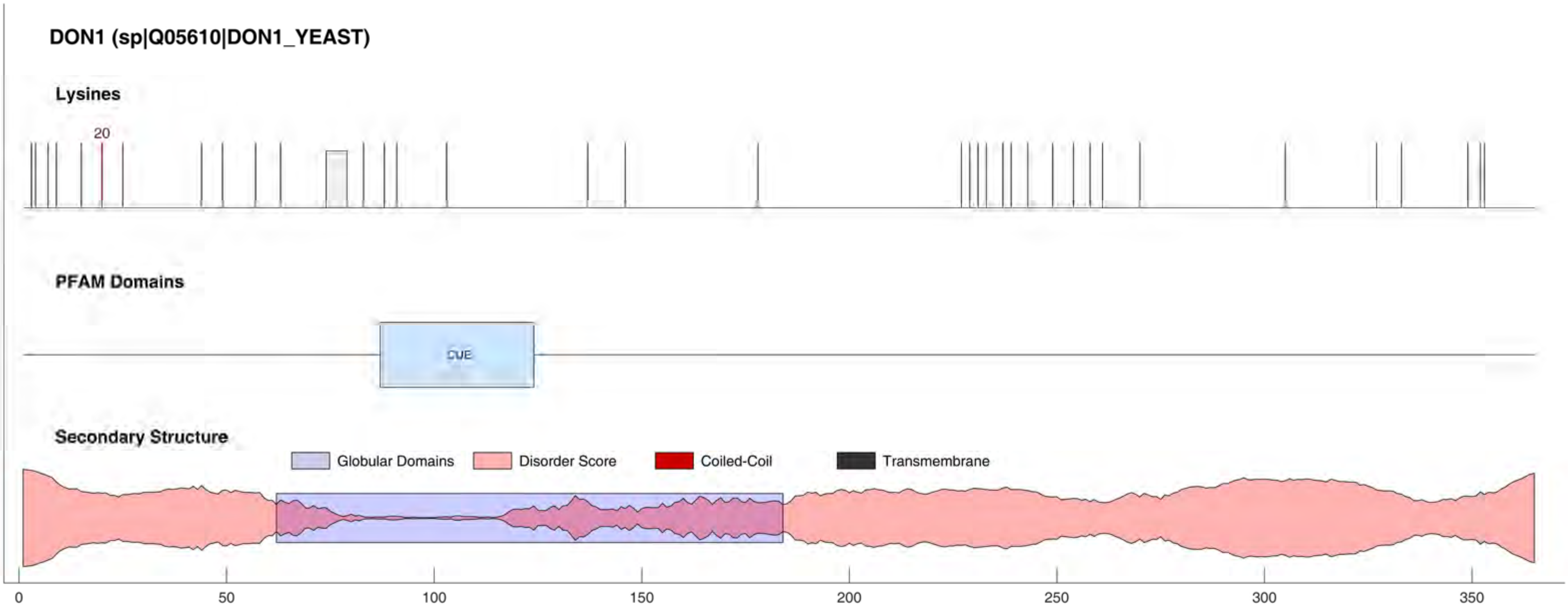
DON1 (sp|Q05610|DON1_YEAST)
Lysines
20
PFAM Domains
CUE
Secondary Structure
Globular Domains
Disorder Score
Coiled-Coil
Transmembrane
0
50
100
150
200
250
300
350

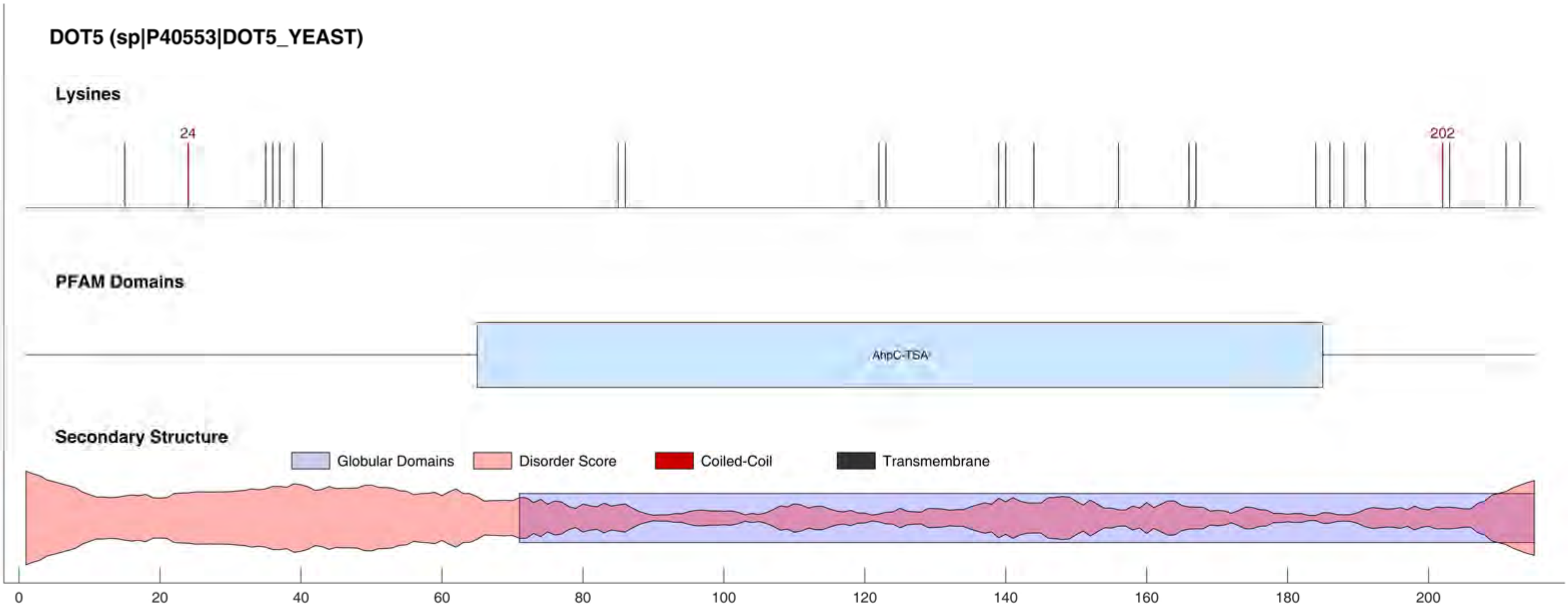
DOT5 (sp|P40553|DOT5_YEAST)
Lysines
24
202
PFAM Domains
AhpC-TSA
Secondary Structure
Globular Domains
Disorder Score
Coiled-Coil
Transmembrane
0
20
40
60
80
100
120
140
160
180
200

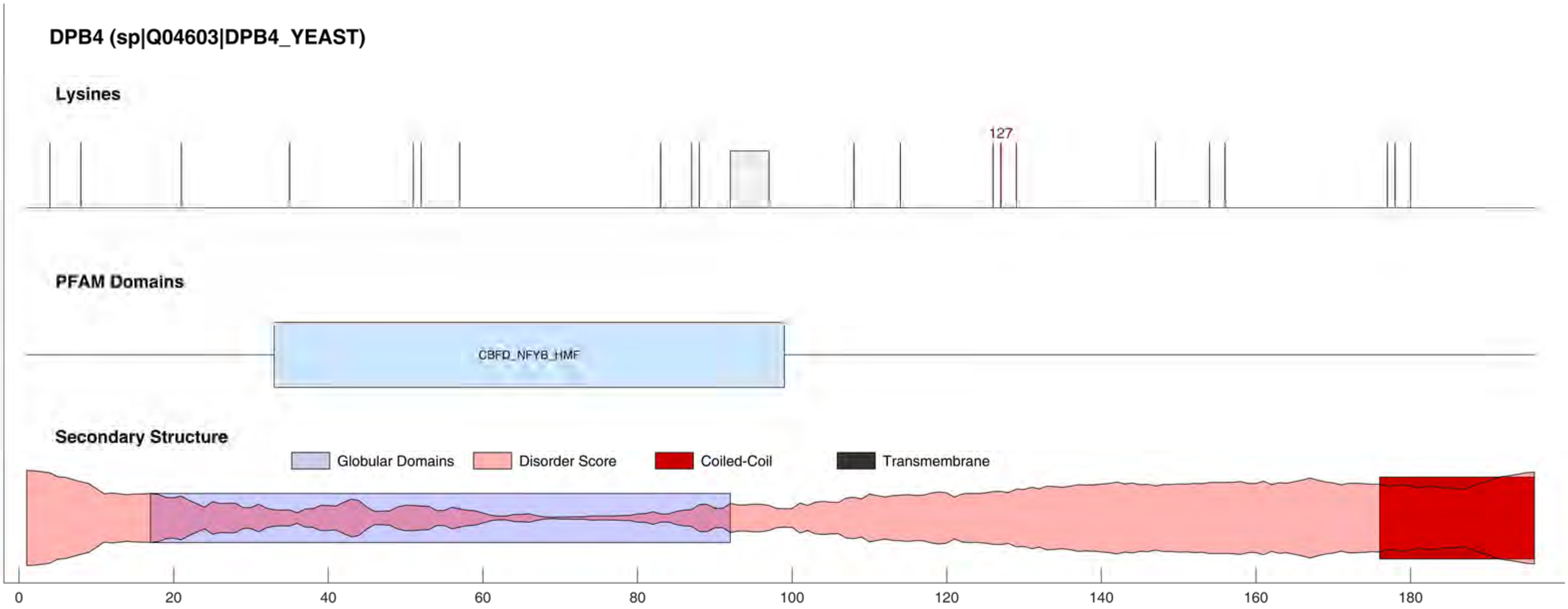
DPB4 (sp|Q04603|DPB4_YEAST)
Lysines
127
PFAM Domains
CBFD_NFYB_HMF
Secondary Structure
Globular Domains
Disorder Score
Coiled-Coil
Transmembrane
0
20
40
60
80
100
120
140
160
180

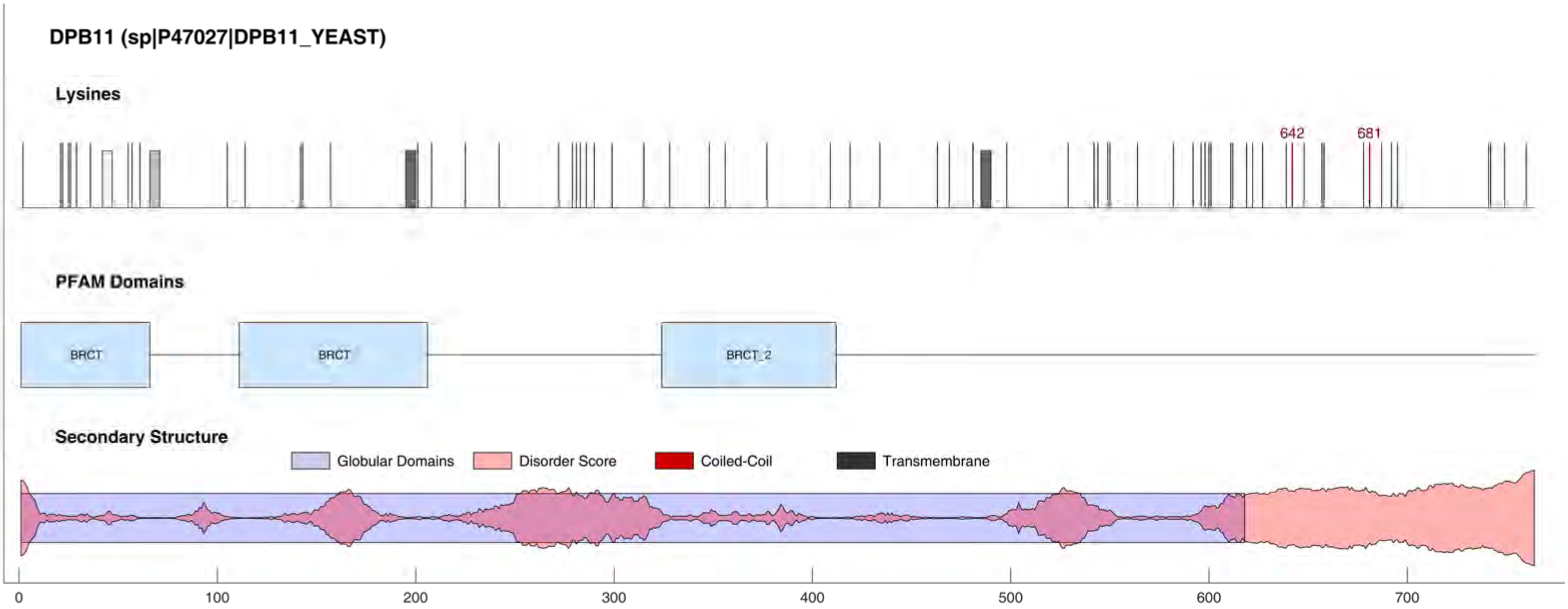
DPB11 (sp|P47027|DPB11_YEAST)
Lysines
642
681
PFAM Domains
BRCT
BRCT
BRCT_2
Secondary Structure
Globular Domains
Disorder Score
Coiled-Coil
Transmembrane
0
100
200
300
400
500
600
700

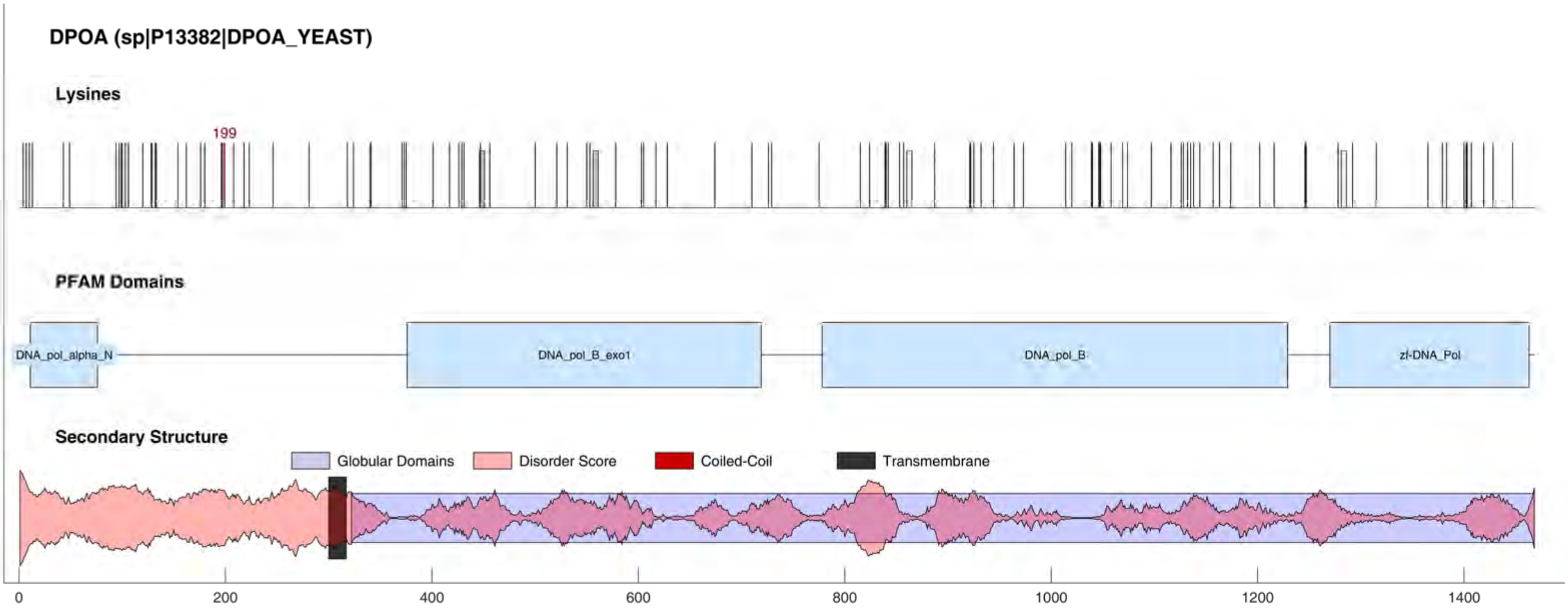

DPOA (sp|P13382|DPOA_YEAST)
Lysines
199
PFAM Domains
DNA_pol_alpha_N
DNA_pol_B_exo1
DNA_pol_B
zf-DNA_Pol
Secondary Structure
Globular Domains
Disorder Score
Coiled-Coil
Transmembrane
0
200
400
600
800
1000
1200
1400

#### DPOA2 (sp|P38121|DPOA2_YEAST)

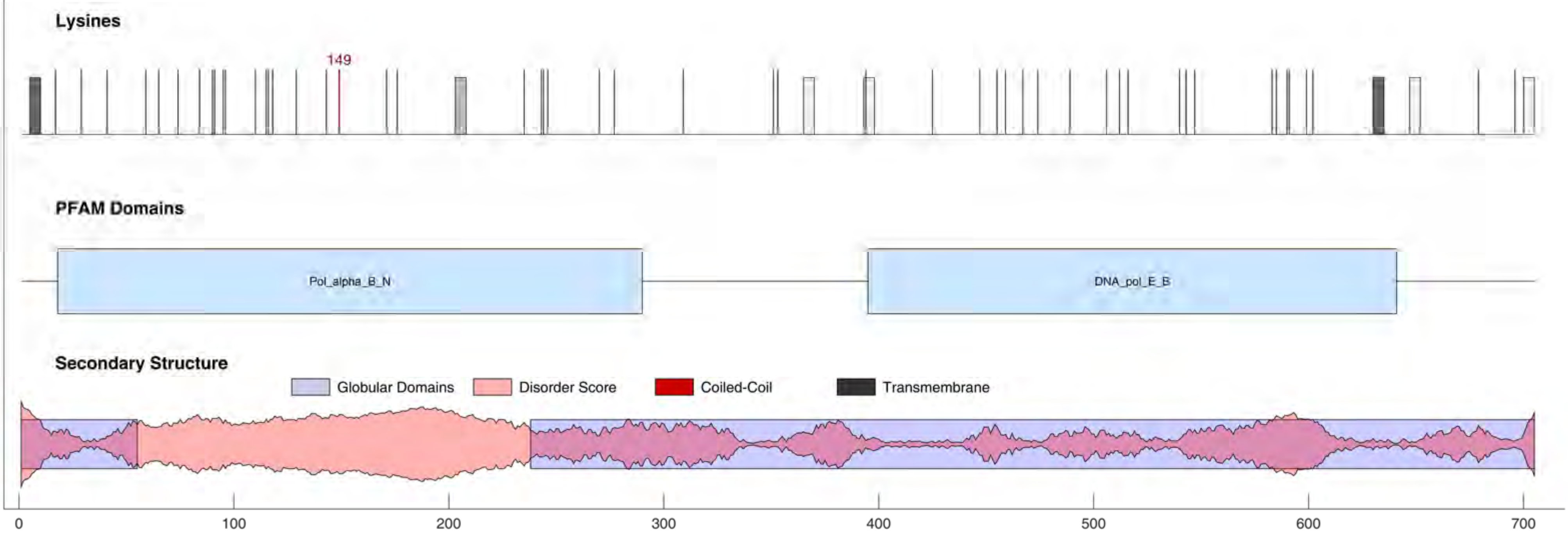

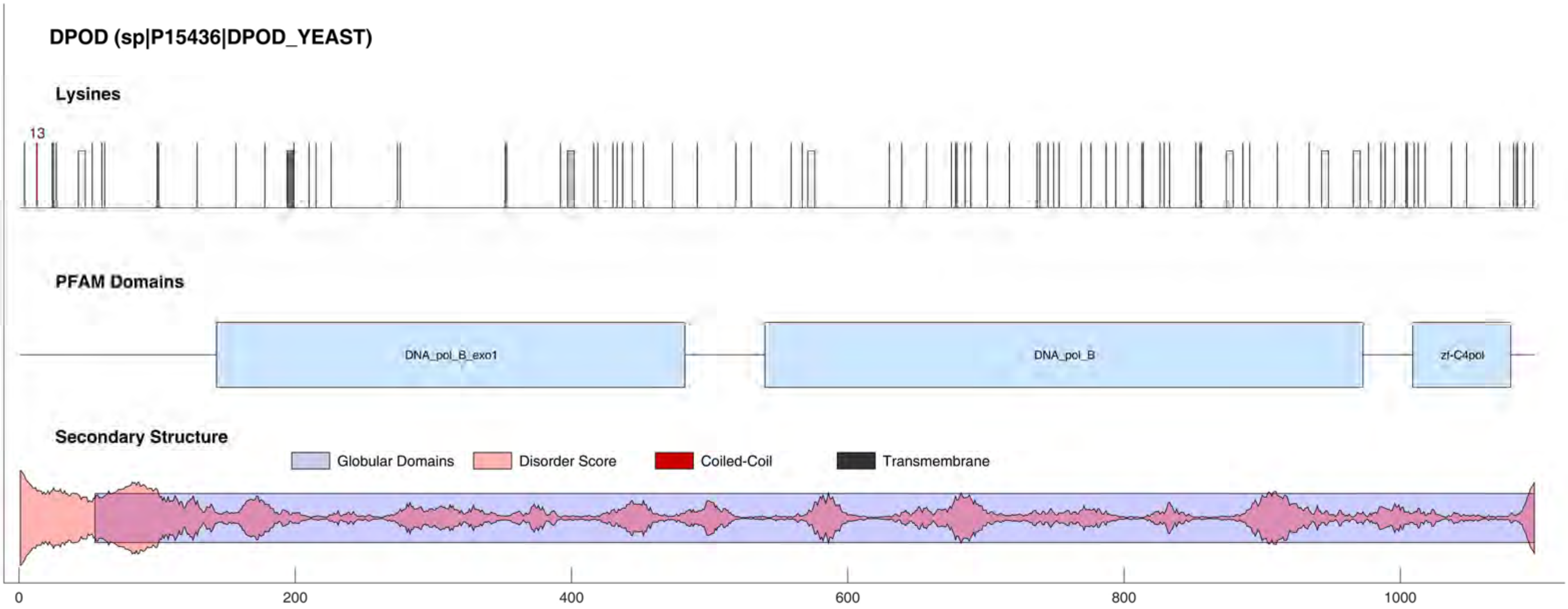
DPOD (sp|P15436|DPOD_YEAST)
Lysines
13
PFAM Domains
DNA_pol_B_exo1
DNA_pol_B
zf-C4pol
Secondary Structure
Globular Domains
Disorder Score
Coiled-Coil
Transmembrane
0
200
400
600
800
1000

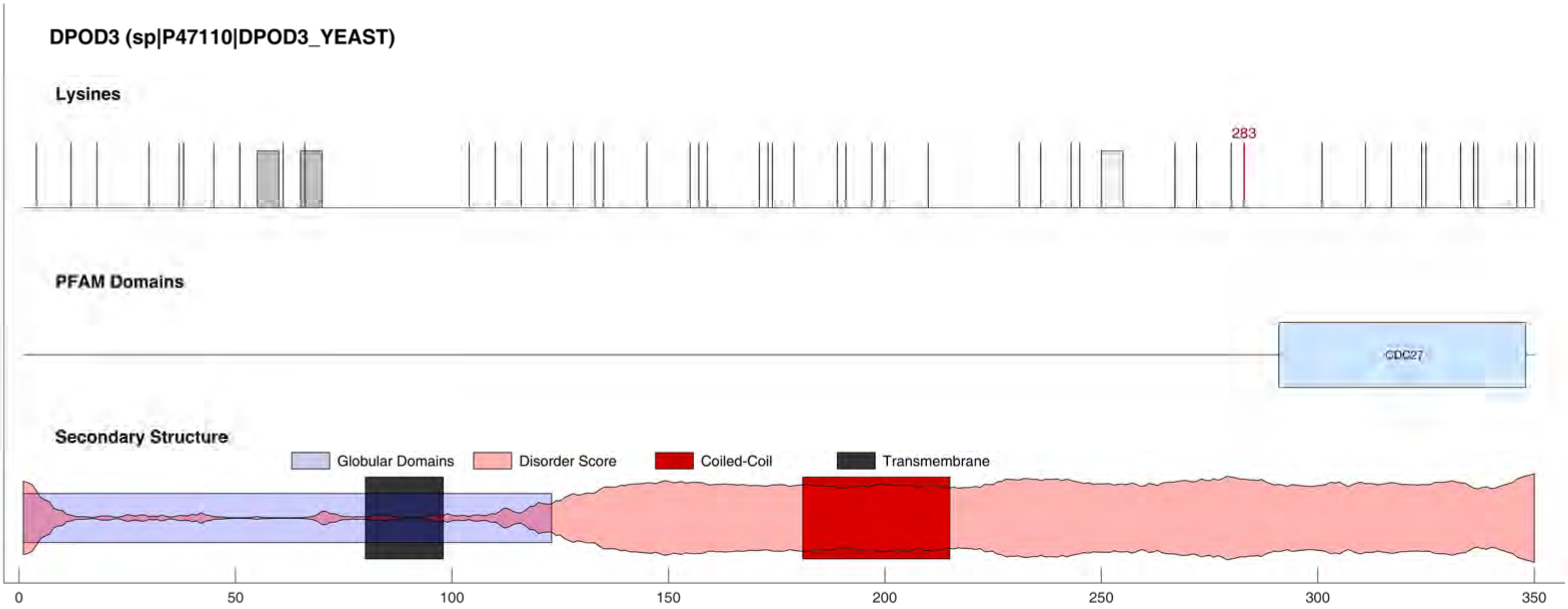
DPOD3 (sp|P47110|DPOD3_YEAST)
Lysines
283
PFAM Domains
CDC27
Secondary Structure
Globular Domains
Disorder Score
Coiled-Coil
Transmembrane
0
50
100
150
200
250
300
350

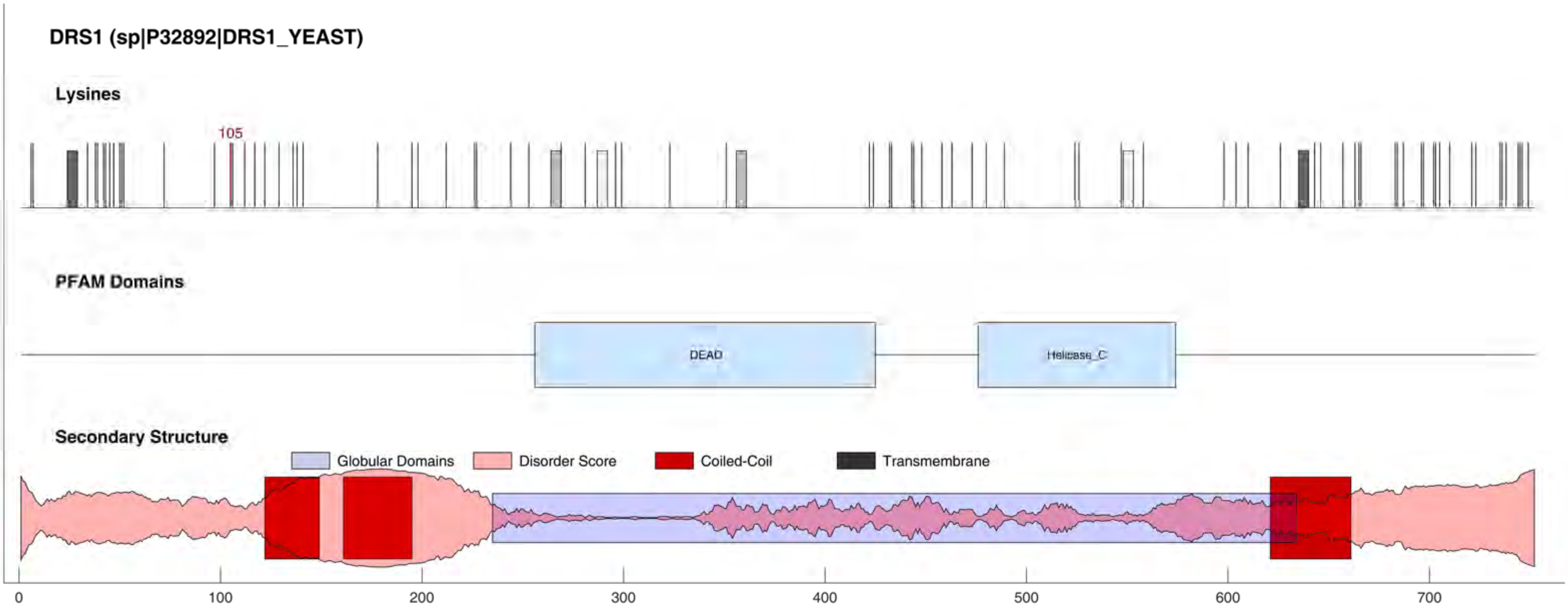

DRS1 (sp|P32892|DRS1_YEAST)
Lysines
105
PFAM Domains
DEAD
Helicase_C
Secondary Structure
Globular Domains
Disorder Score
Coiled-Coil
Transmembrane
0
100
200
300
400
500
600
700

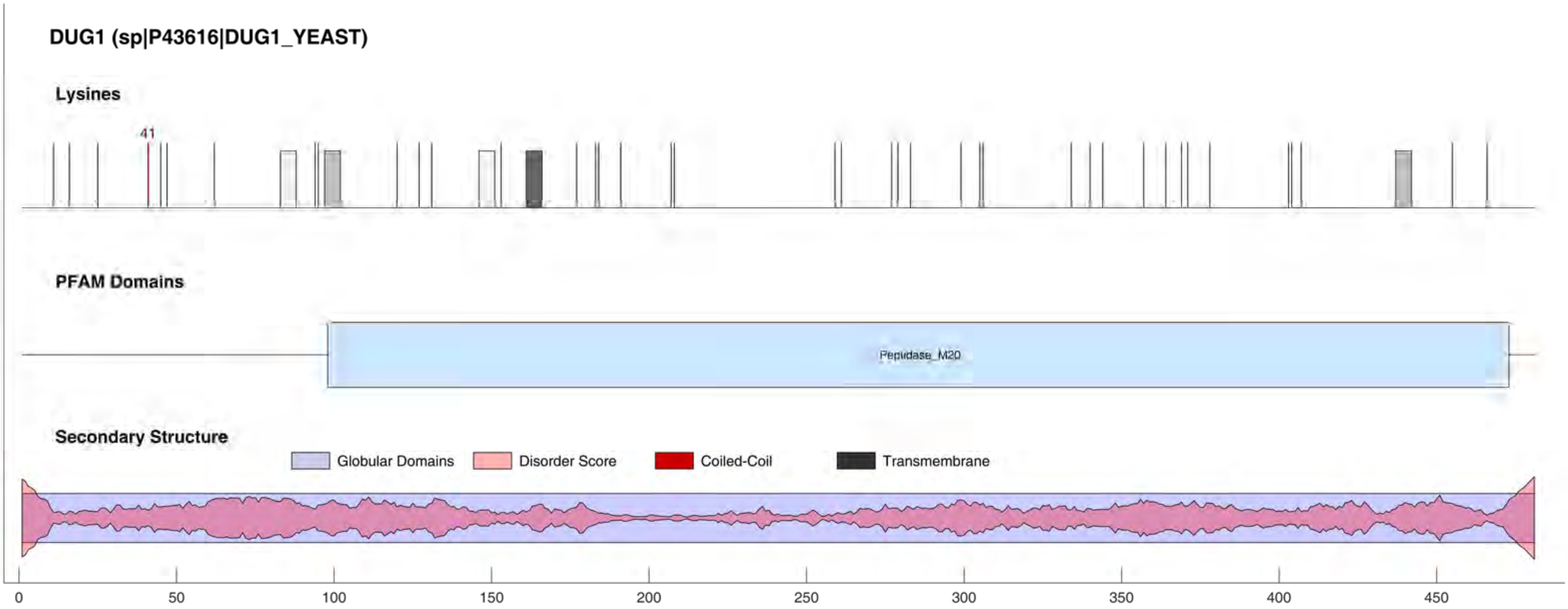
DUG1 (sp|P43616|DUG1_YEAST)
Lysines
41
PFAM Domains
Pepidase_M20
Secondary Structure
Globular Domains
Disorder Score
Coiled-Coil
Transmembrane
0
50
100
150
200
250
300
350
400
450

#### DYL1 (sp|Q02647|DYL1_YEAST)

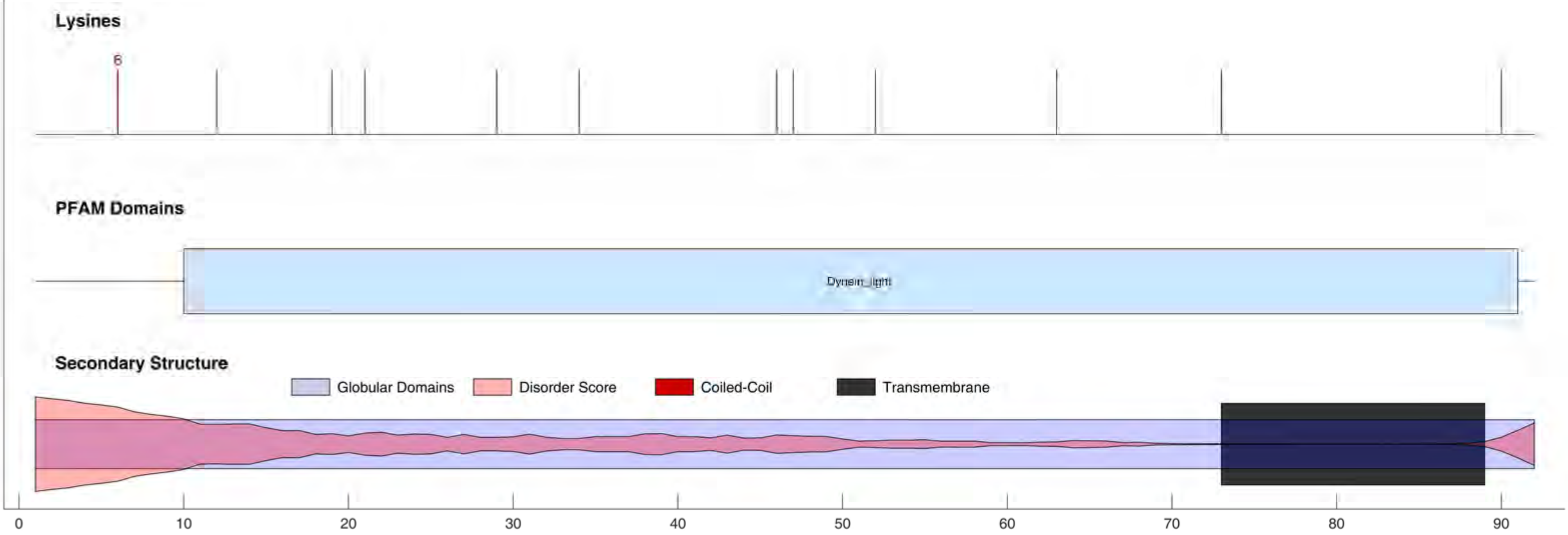

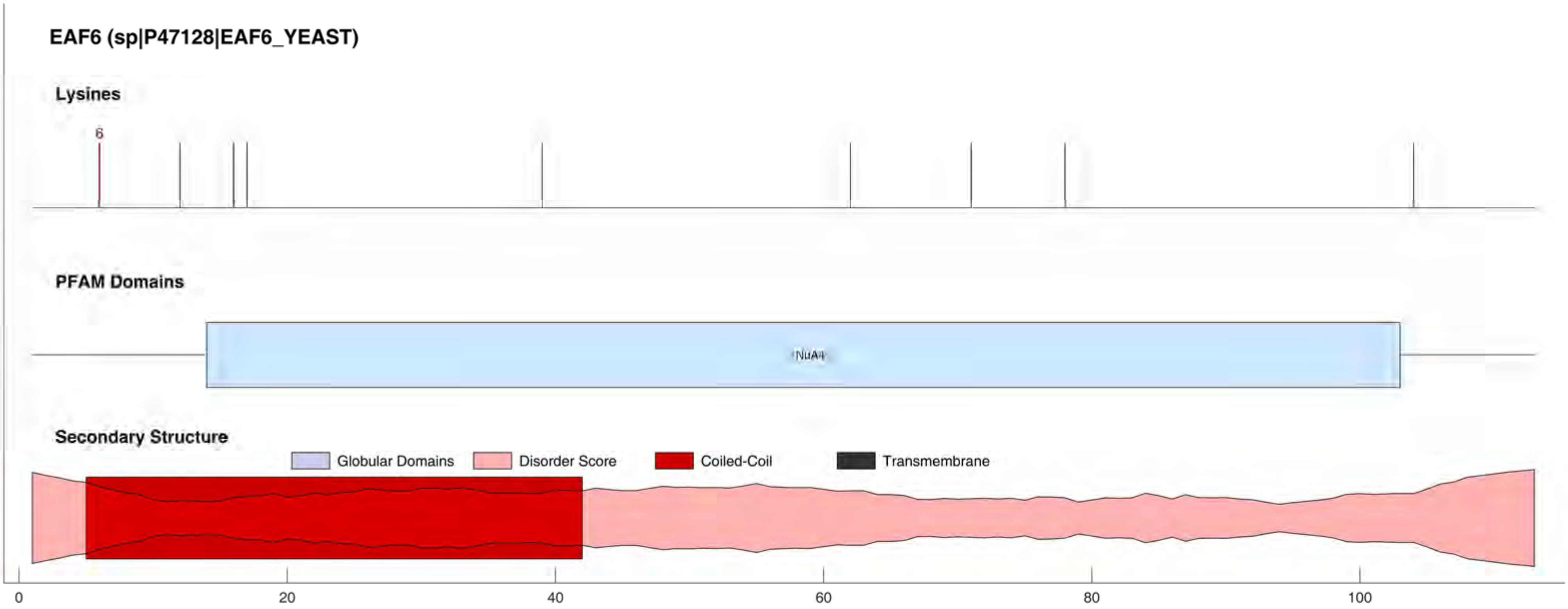
EAF6 (sp|P47128|EAF6_YEAST)
Lysines
6
PFAM Domains
NuA4
Secondary Structure
Globular Domains
Disorder Score
Coiled-Coil
Transmembrane
0
20
40
60
80
100

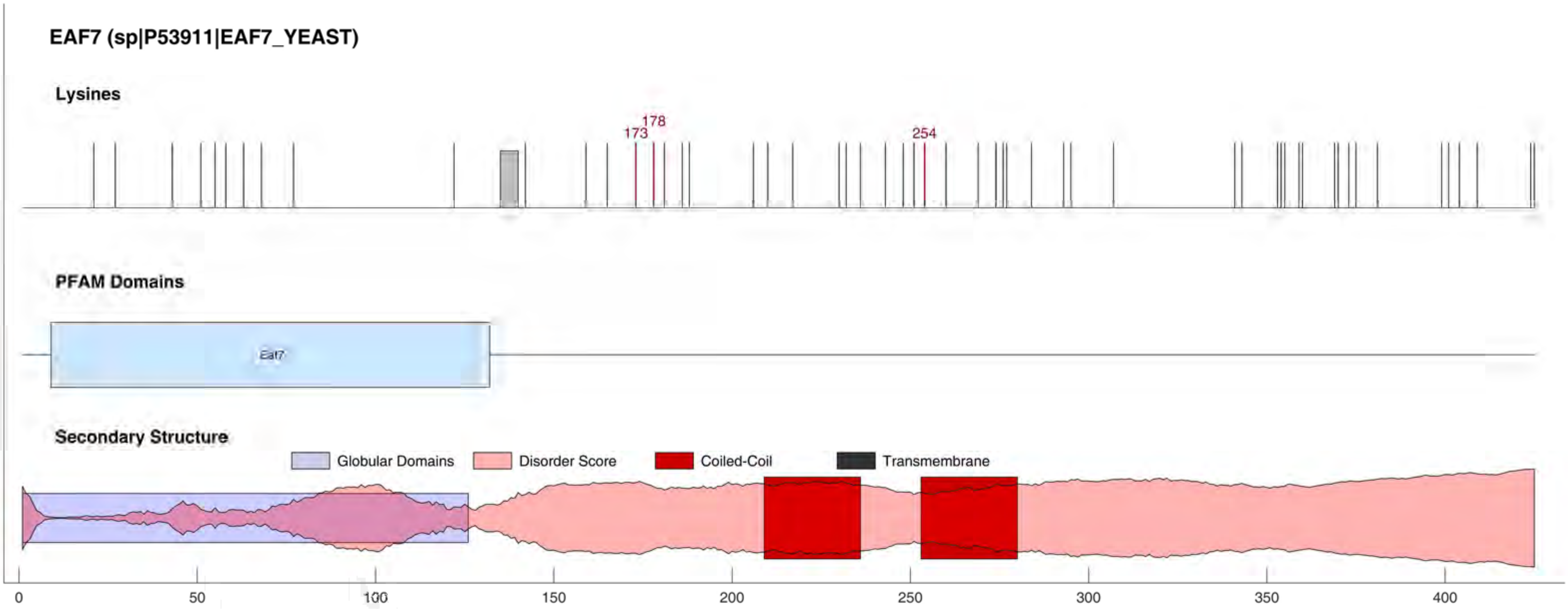
EAF7 (sp|P53911|EAF7_YEAST)
Lysines
178
173
254
PFAM Domains
Eaf7
Secondary Structure
Globular Domains
Disorder Score
Coiled-Coil
Transmembrane
0
50
100
150
200
250
300
350
400

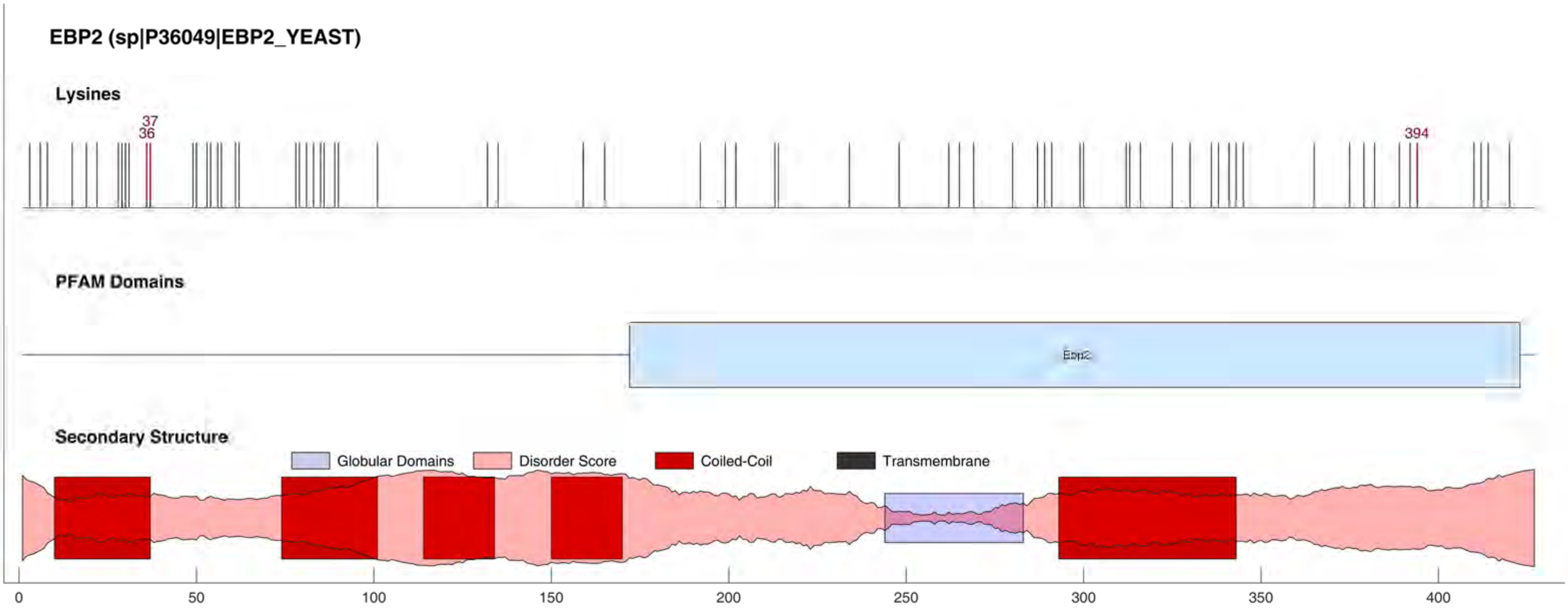

EBP2 (sp|P36049|EBP2_YEAST)
Lysines
37
36
394
PFAM Domains
Ebp2
Secondary Structure
Globular Domains
Disorder Score
Coiled-Coil
Transmembrane
0
50
100
150
200
250
300
350
400

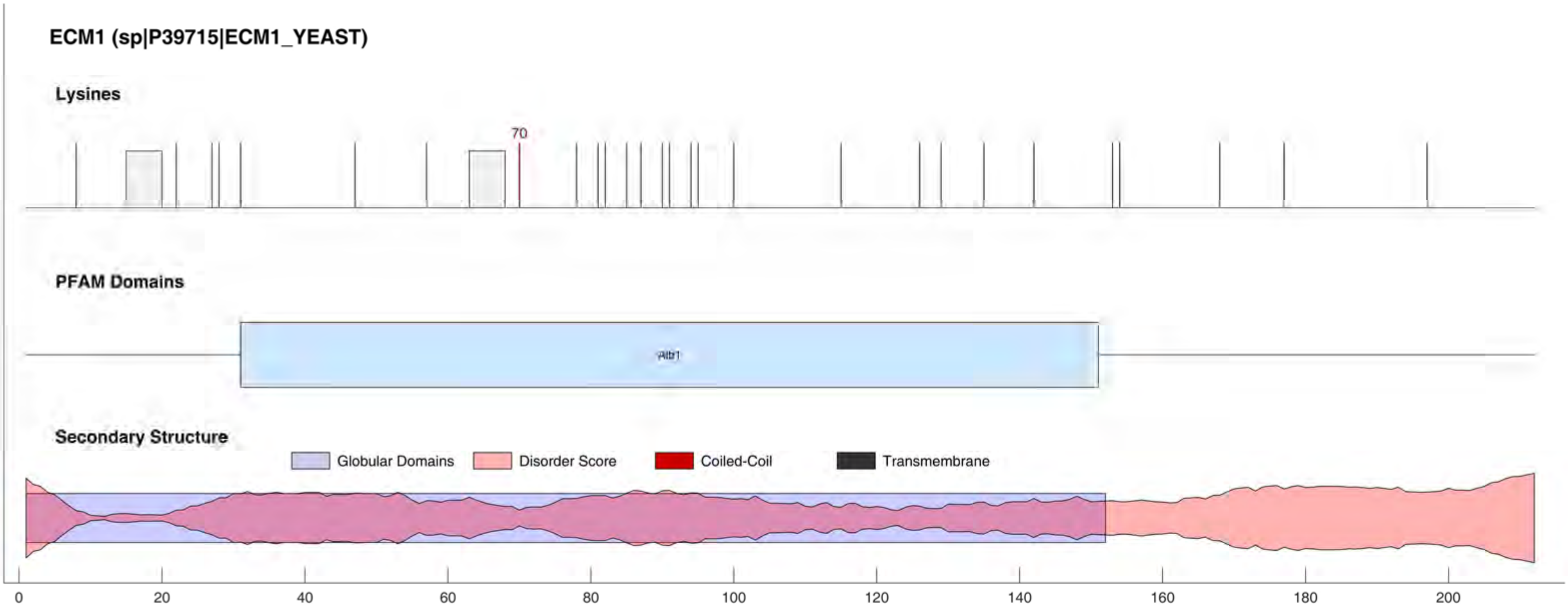

ECM1 (sp|P39715|ECM1_YEAST)
Lysines
70
PFAM Domains
Secondary Structure
Globular Domains
Disorder Score
Coiled-Coil
Transmembrane
0
20
40
60
80
100
120
140
160
180
200

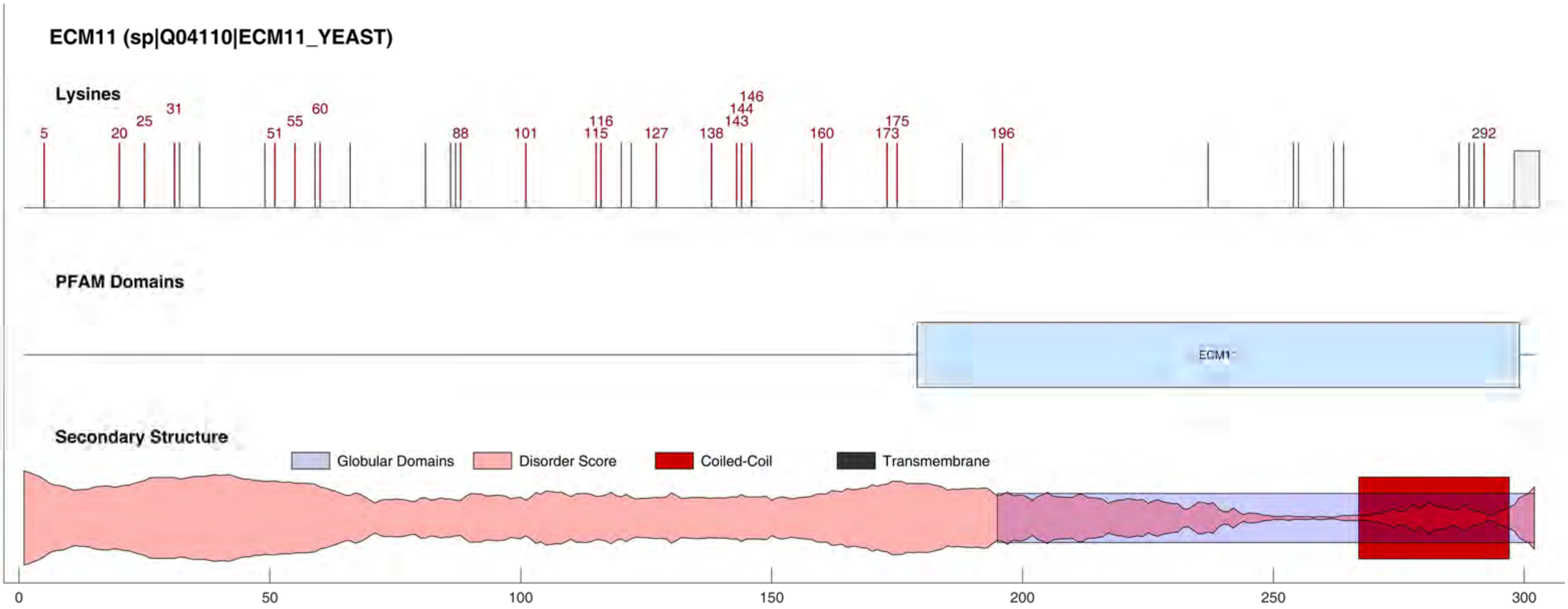
ECM11 (sp|Q04110|ECM11_YEAST)
Lysines
5
20
25
31
51
55
60
88
101
115
116
127
138
143
144
146
160
173
175
196
292
PFAM Domains
ECM1
Secondary Structure
Globular Domains
Disorder Score
Coiled-Coil
Transmembrane
0
50
100
150
200
250
300

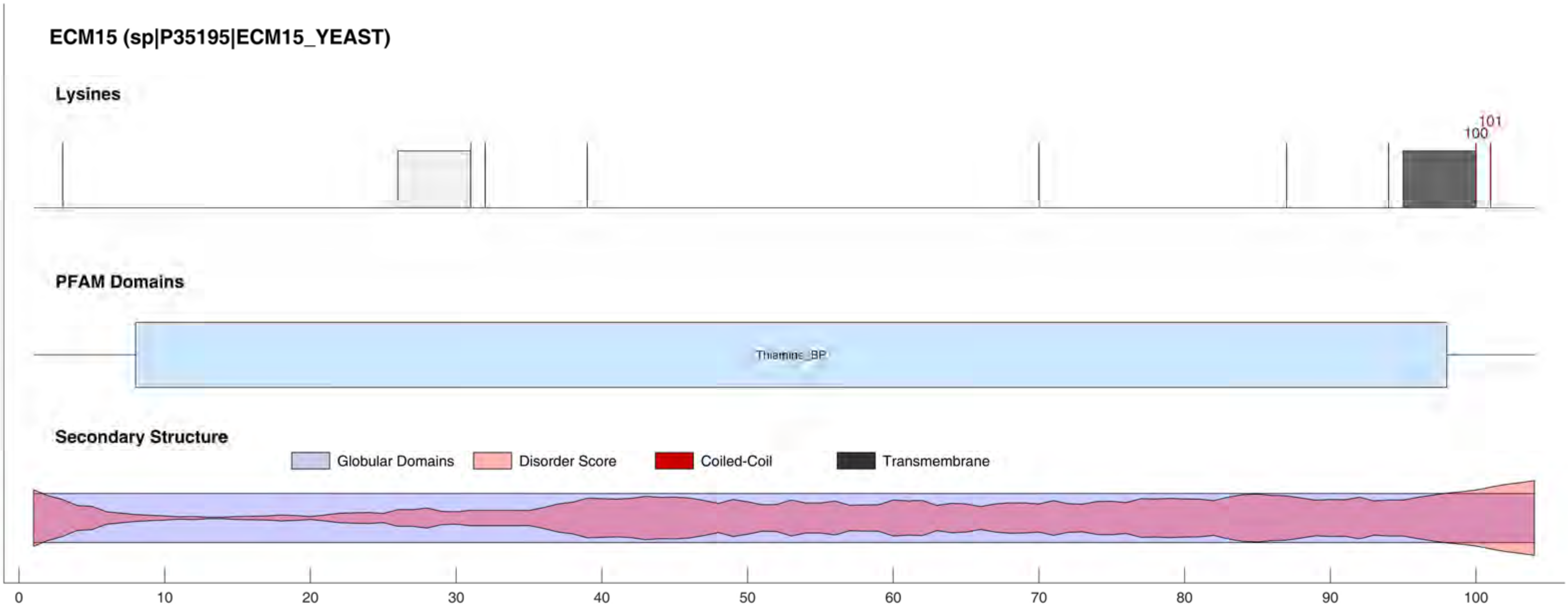

ECM15 (sp|P35195|ECM15_YEAST)
Lysines
100
101
PFAM Domains
Thiamine_BP
Secondary Structure
Globular Domains
Disorder Score
Coiled-Coil
Transmembrane
0
10
20
30
40
50
60
70
80
90
100

EDE1 (sp|P34216|EDE1_YEAST)

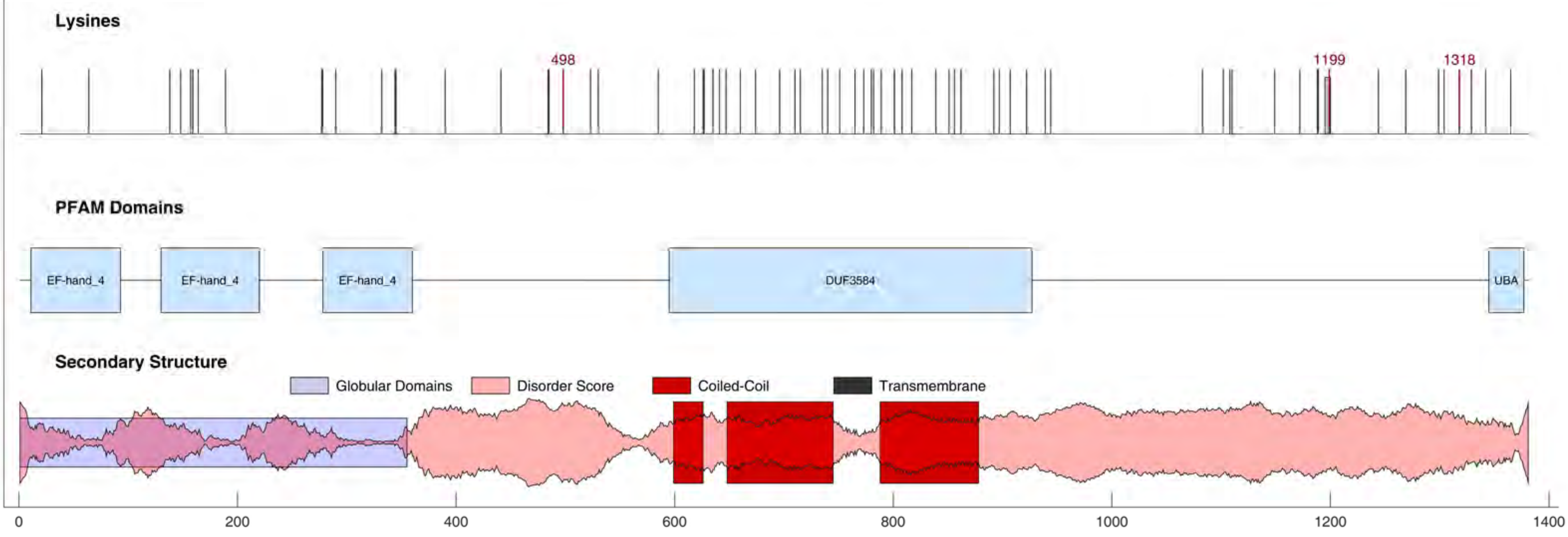

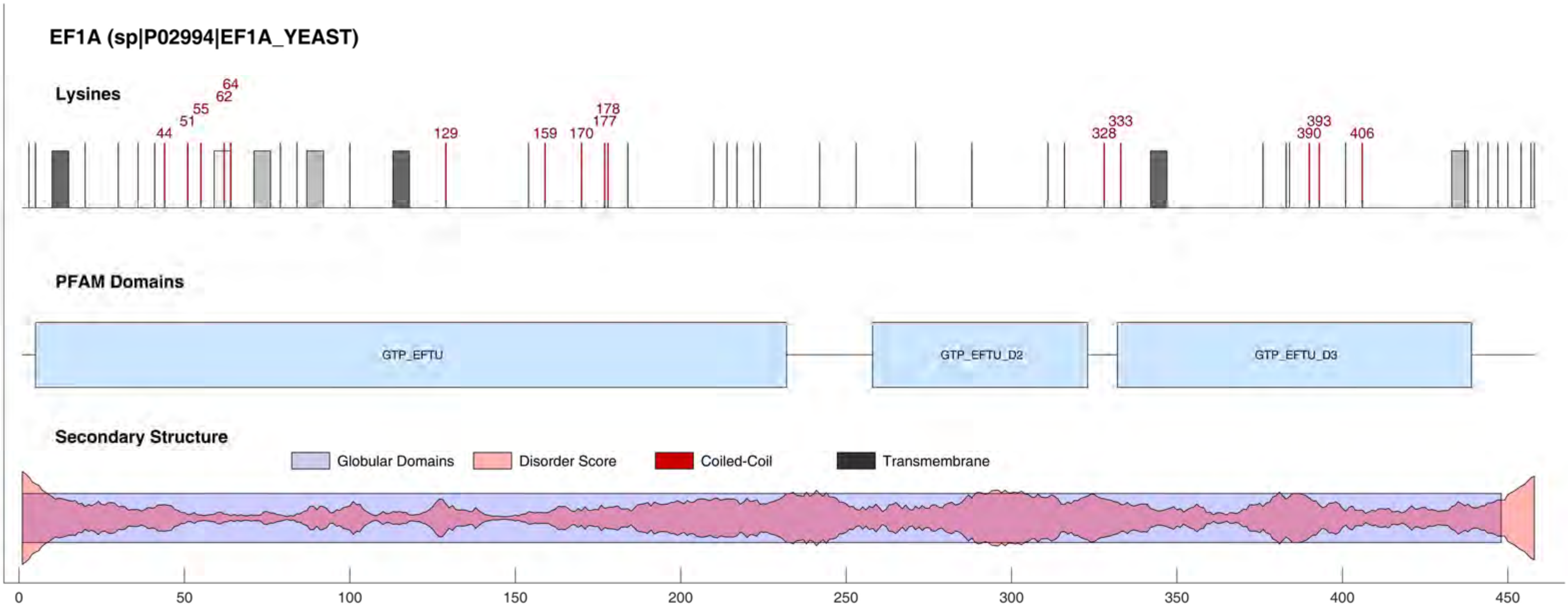
EF1A (sp|P02994|EF1A_YEAST)
Lysines
44
51
55
62
64
129
159
170
177
178
328
333
390
393
406
PFAM Domains
GTP_EFTU
GTP_EFTU_D2
GTP_EFTU_D3
Secondary Structure
Globular Domains
Disorder Score
Coiled-Coil
Transmembrane
0
50
100
150
200
250
300
350
400
450

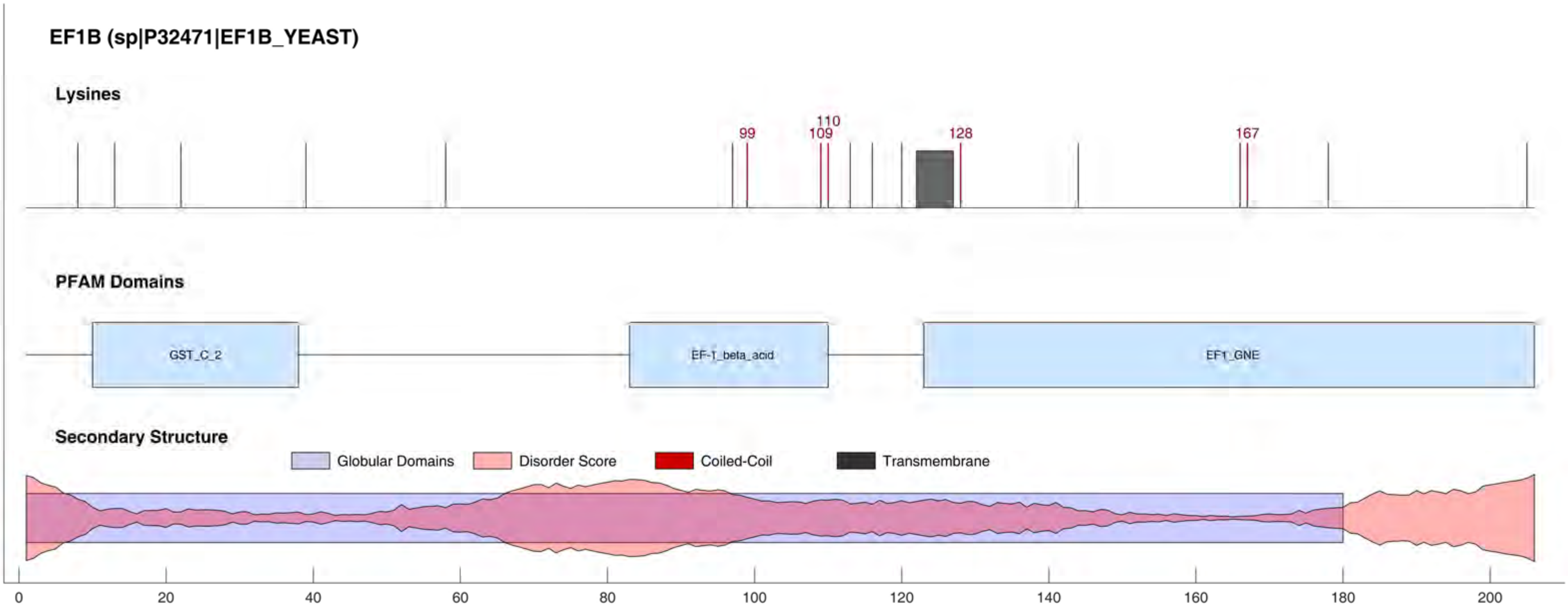
EF1B (sp|P32471|EF1B_YEAST)
Lysines
99
110
109
128
167
PFAM Domains
GST_C_2
EF-1_beta_acid
EF1_GNE
Secondary Structure
Globular Domains
Disorder Score
Coiled-Coil
Transmembrane
0
20
40
60
80
100
120
140
160
180
200

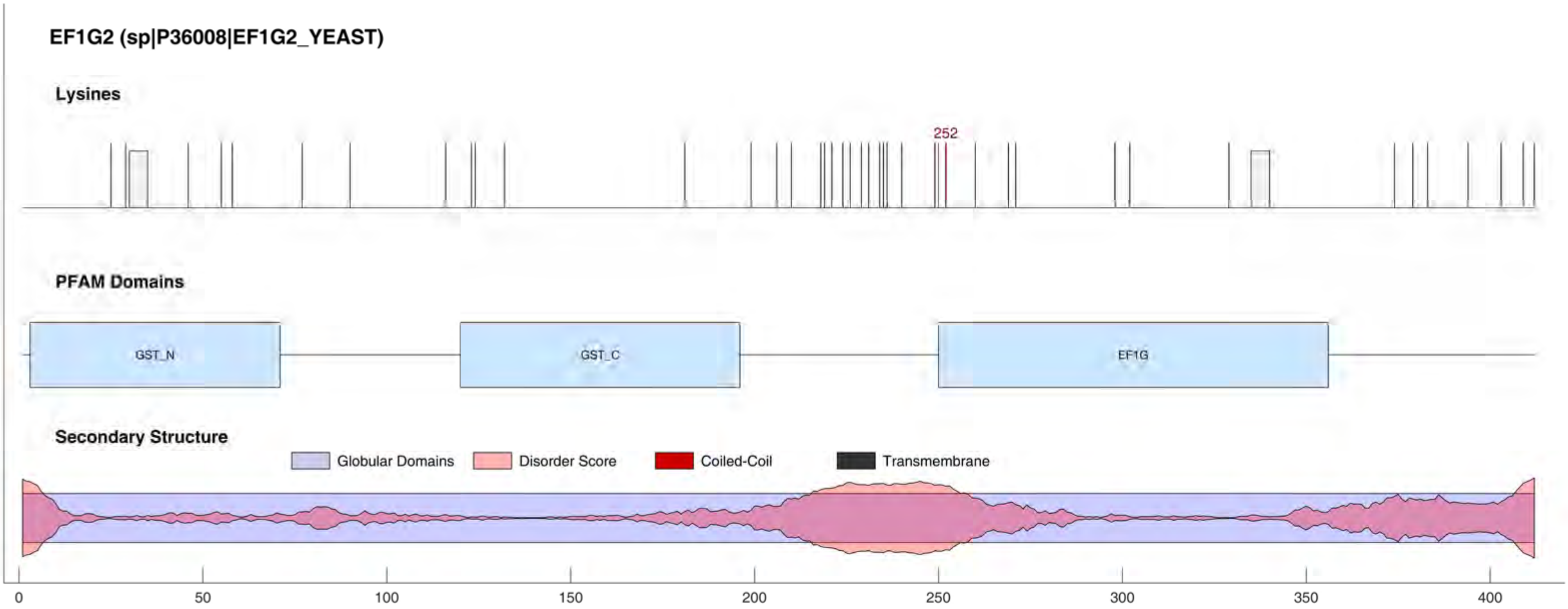
EF1G2 (sp|P36008|EF1G2_YEAST)
Lysines
252
PFAM Domains
GST_N
GST_C
EF1G
Secondary Structure
Globular Domains
Disorder Score
Coiled-Coil
Transmembrane
0
50
100
150
200
250
300
350
400

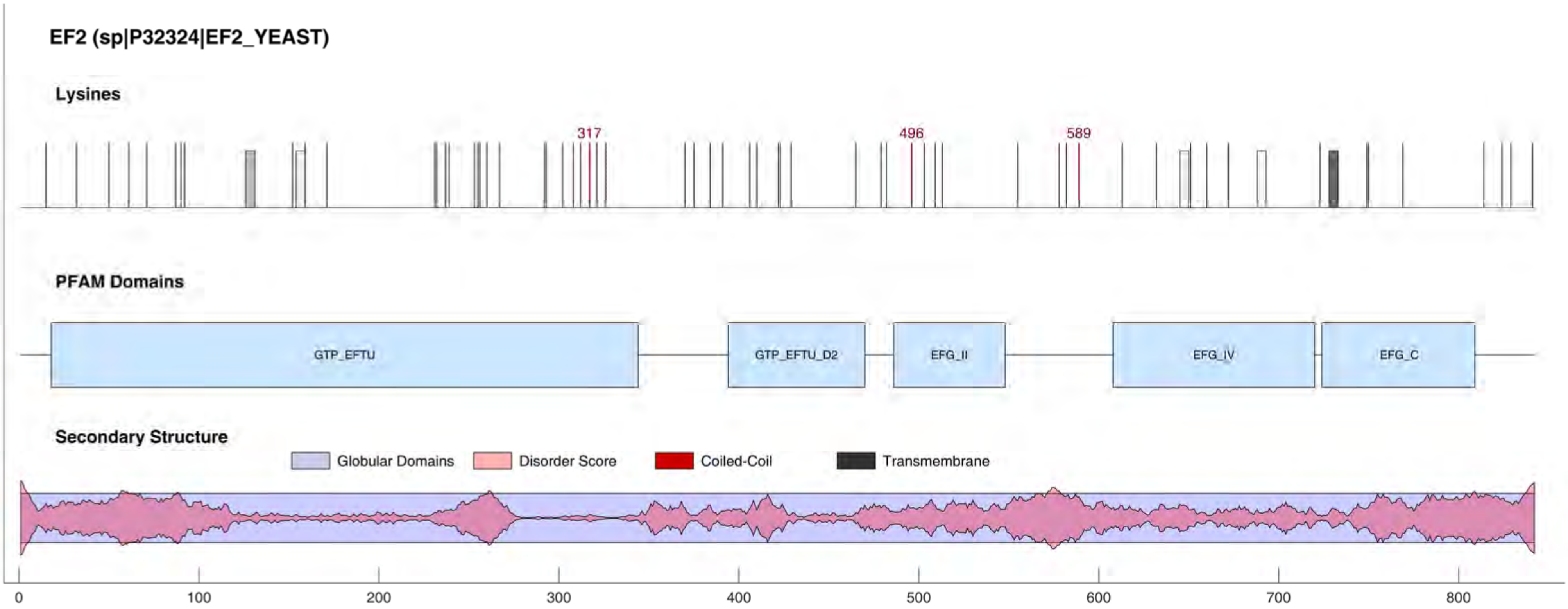

EF2 (sp|P32324|EF2_YEAST)
Lysines
317
496
589
PFAM Domains
GTP_EFTU
GTP_EFTU_D2
EFG_II
EFG_IV
EFG_C
Secondary Structure
Globular Domains
Disorder Score
Coiled-Coil
Transmembrane
0
100
200
300
400
500
600
700
800

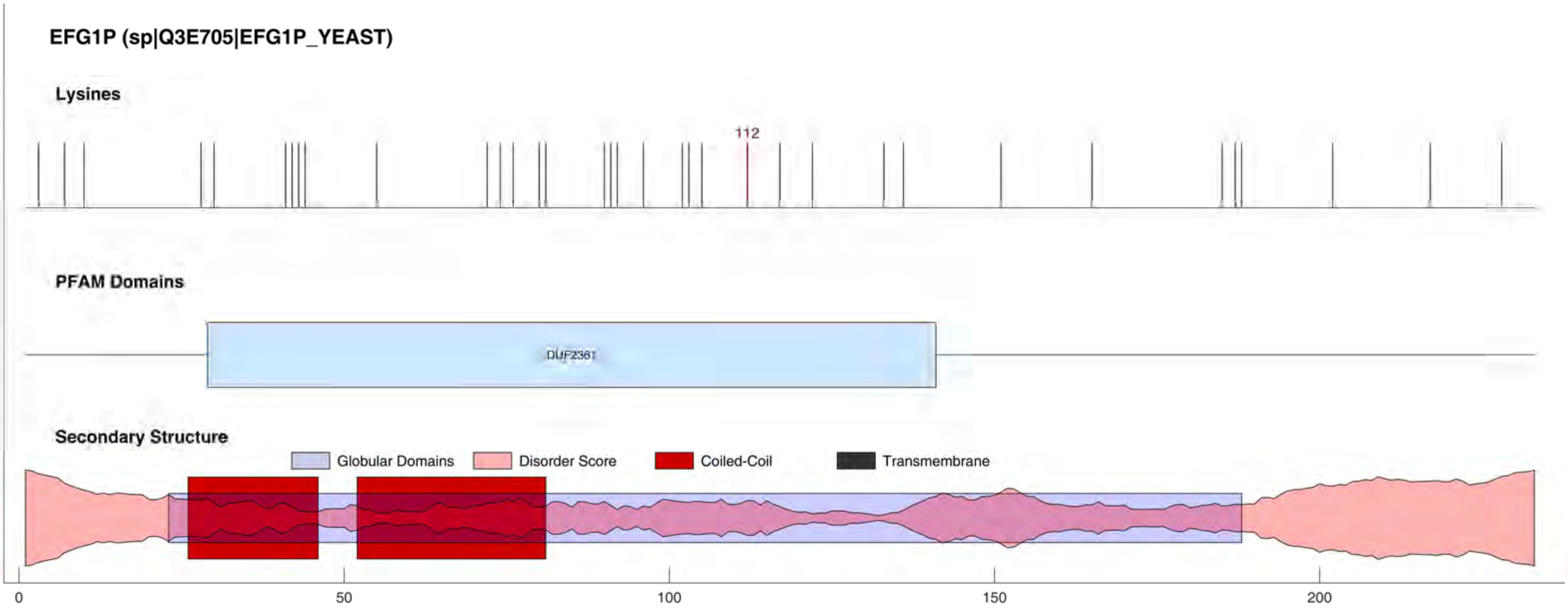
EFG1P (sp|Q3E705|EFG1P_YEAST)
Lysines
112
PFAM Domains
DUF2361
Secondary Structure
Globular Domains
Disorder Score
Coiled-Coil
Transmembrane
0
50
100
150
200

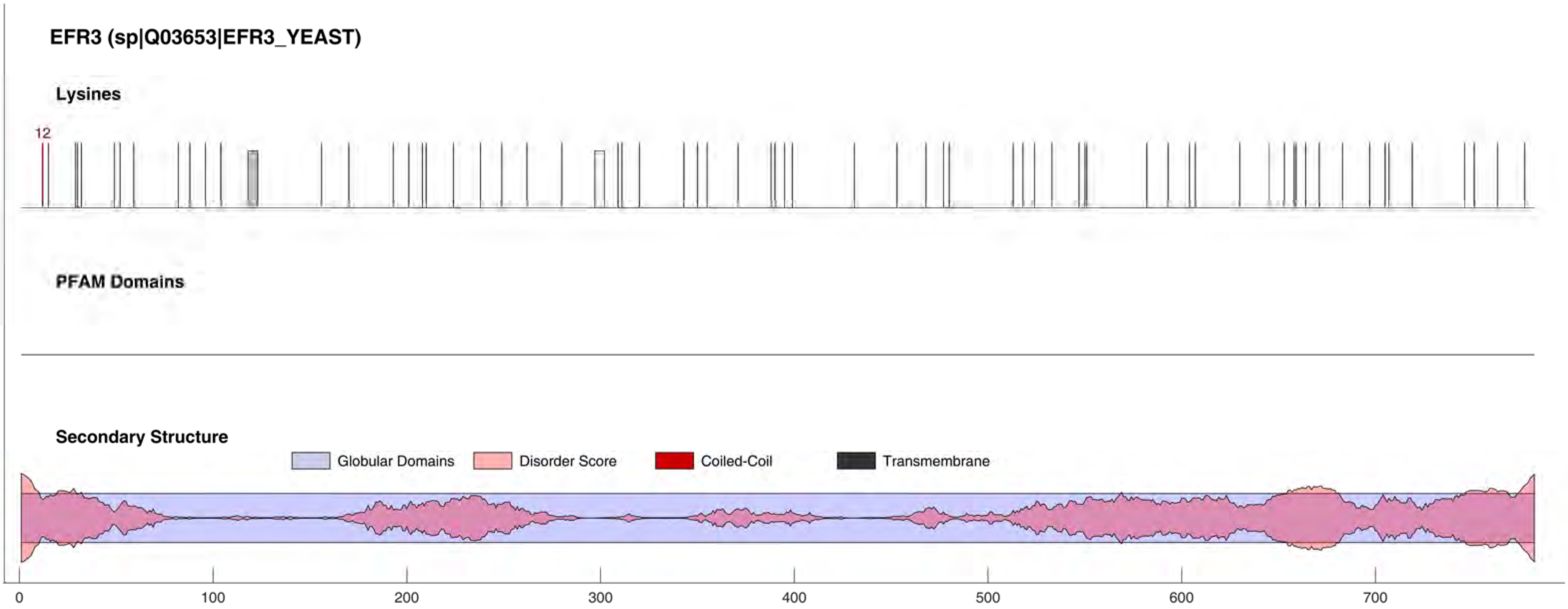

EFR3 (sp|Q03653|EFR3_YEAST)
Lysines
12
PFAM Domains
Secondary Structure
Globular Domains
Disorder Score
Coiled-Coil
Transmembrane
0
100
200
300
400
500
600
700

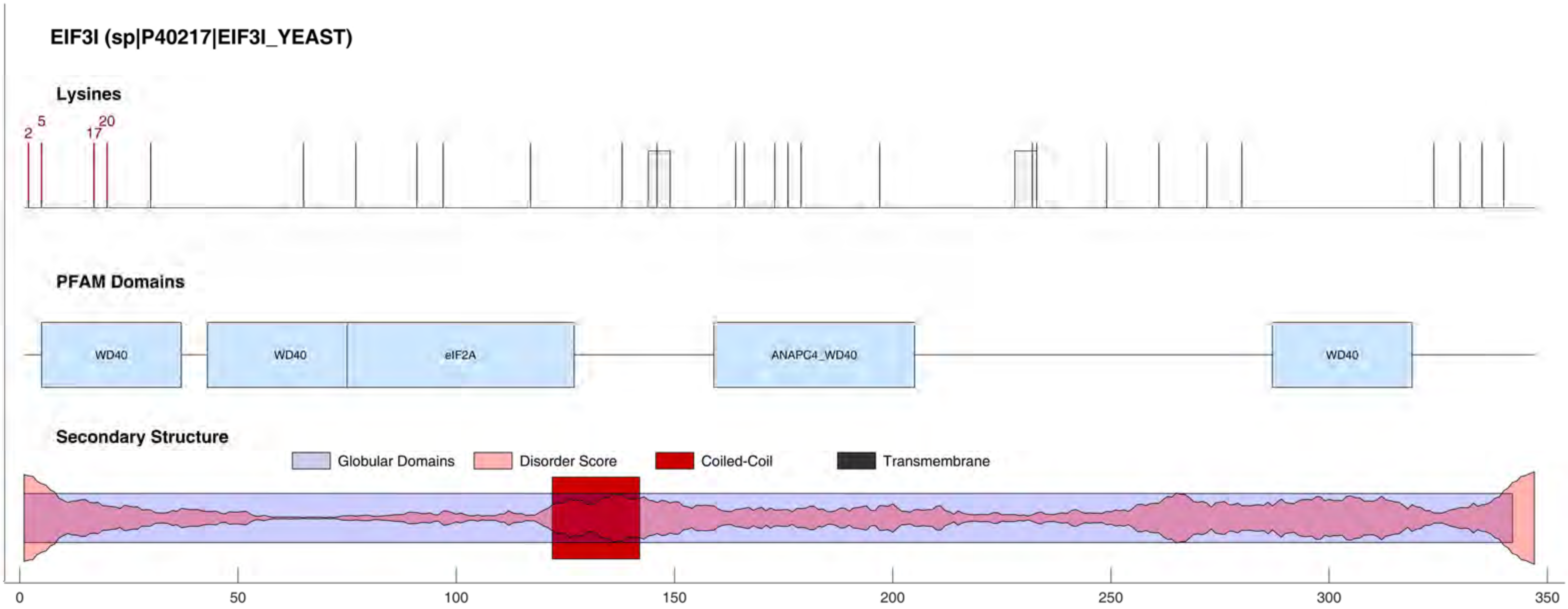
EIF3I (sp|P40217|EIF3I_YEAST)
Lysines
2
5
17
20
PFAM Domains
WD40
WD40
eIF2A
ANAPC4_WD40
WD40
Secondary Structure
Globular Domains
Disorder Score
Coiled-Coil
Transmembrane
0
50
100
150
200
250
300
350

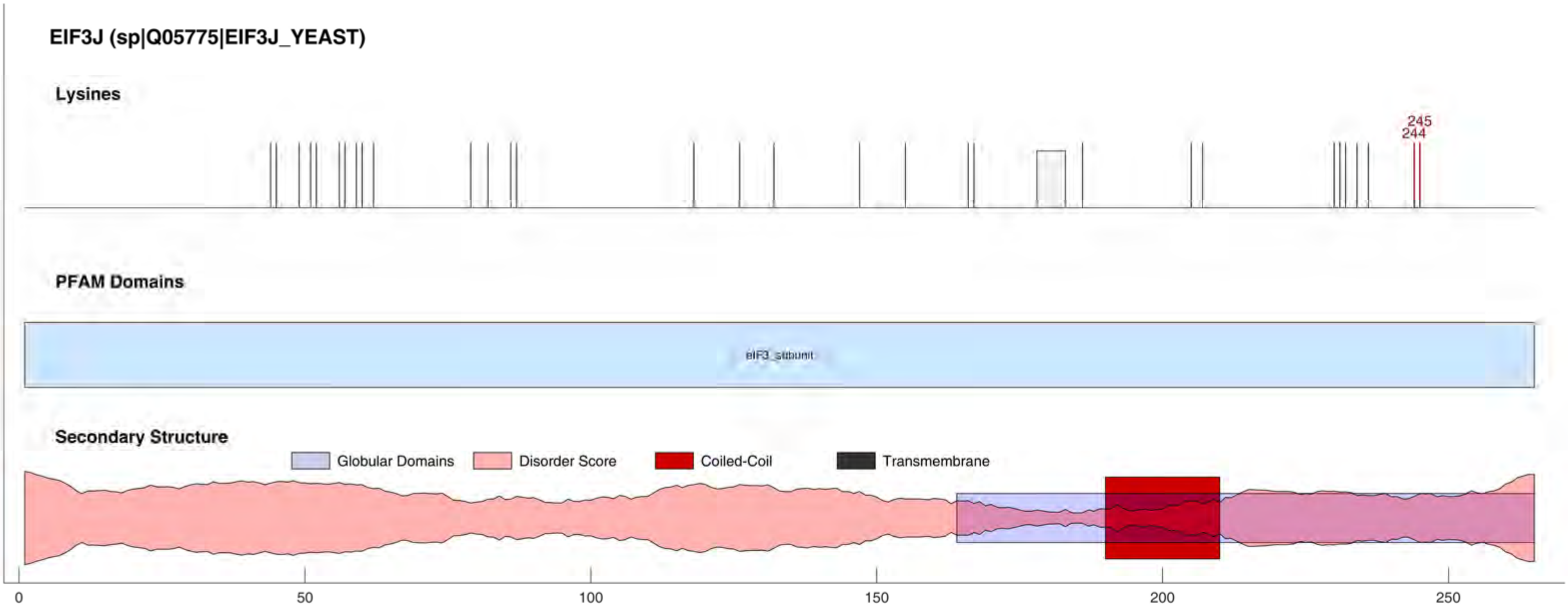
EIF3J (sp|Q05775|EIF3J_YEAST)
Lysines
245
244
PFAM Domains
eIF3_subunit
Secondary Structure
Globular Domains
Disorder Score
Coiled-Coil
Transmembrane
0
50
100
150
200
250

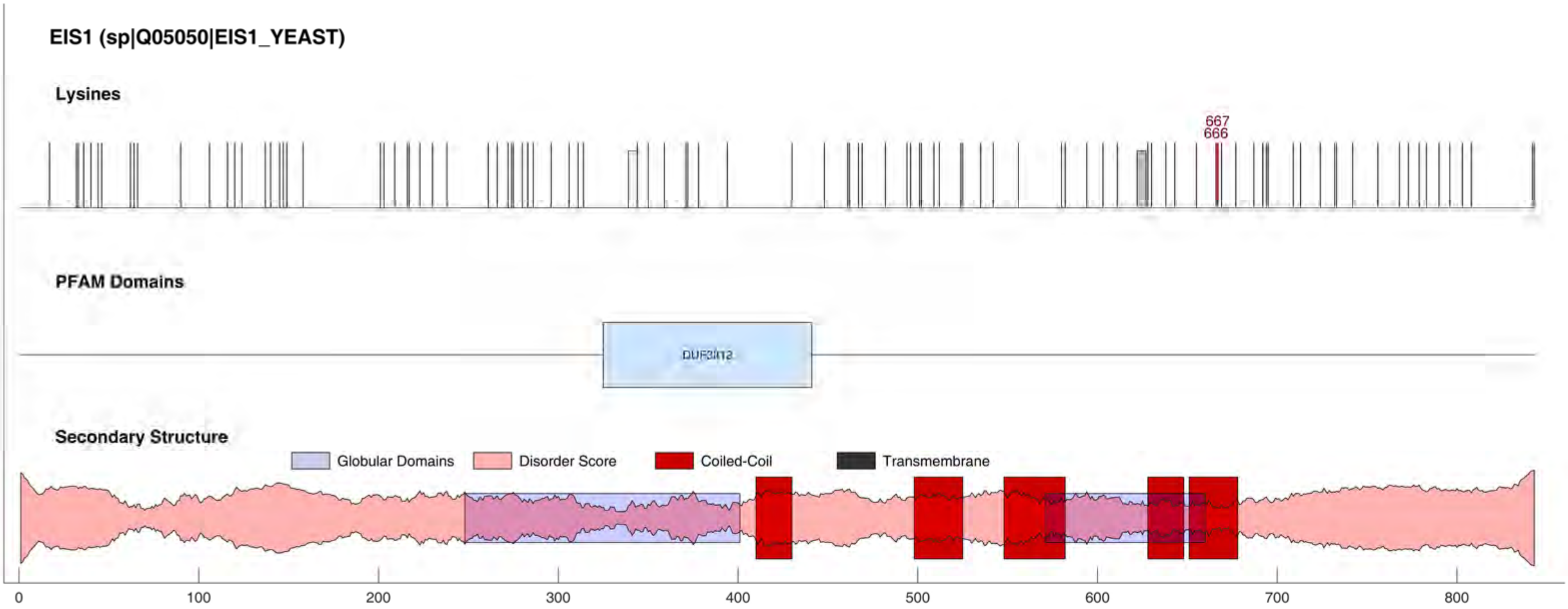
EIS1 (sp|Q05050|EIS1_YEAST)
Lysines
667
666
PFAM Domains
DUF3112
Secondary Structure
Globular Domains
Disorder Score
Coiled-Coil
Transmembrane
0
100
200
300
400
500
600
700
800

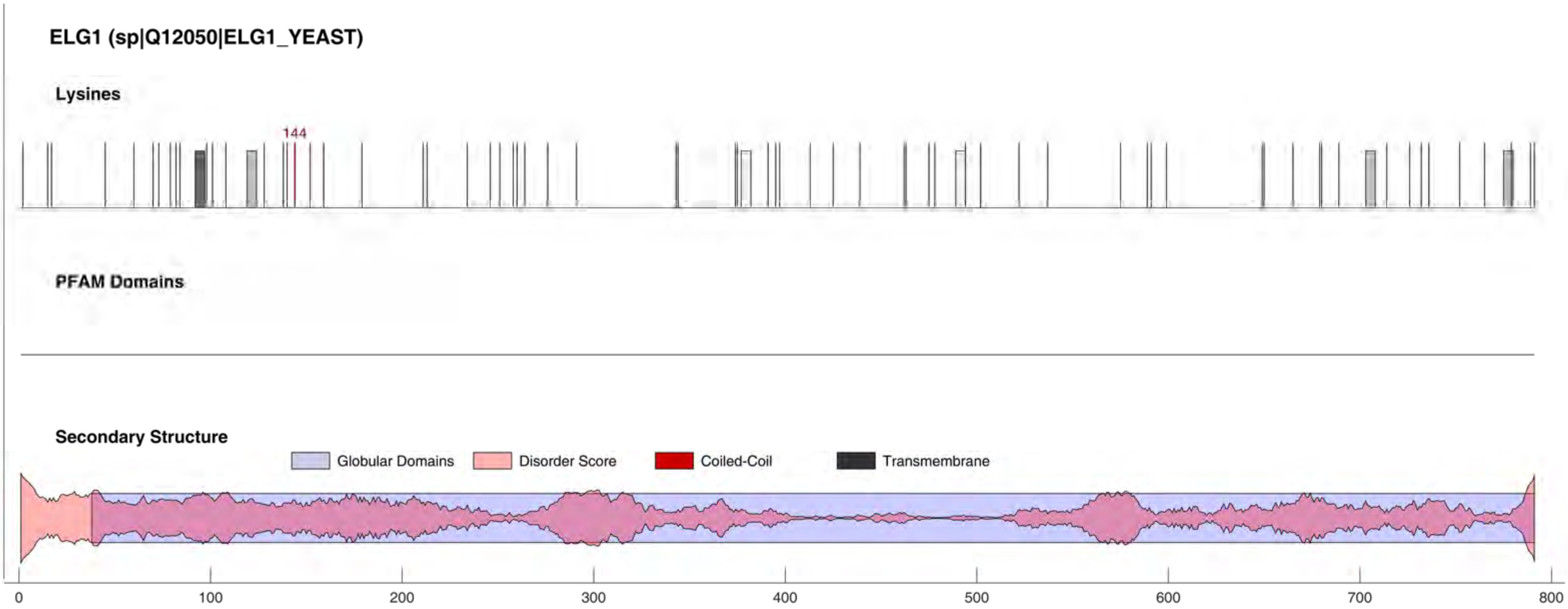
ELG1 (sp|Q12050|ELG1_YEAST)
Lysines
144
PFAM Domains
Secondary Structure
Globular Domains
Disorder Score
Coiled-Coil
Transmembrane
0
100
200
300
400
500
600
700
800

#### ELP1 (sp|Q06706|ELP1_YEAST)

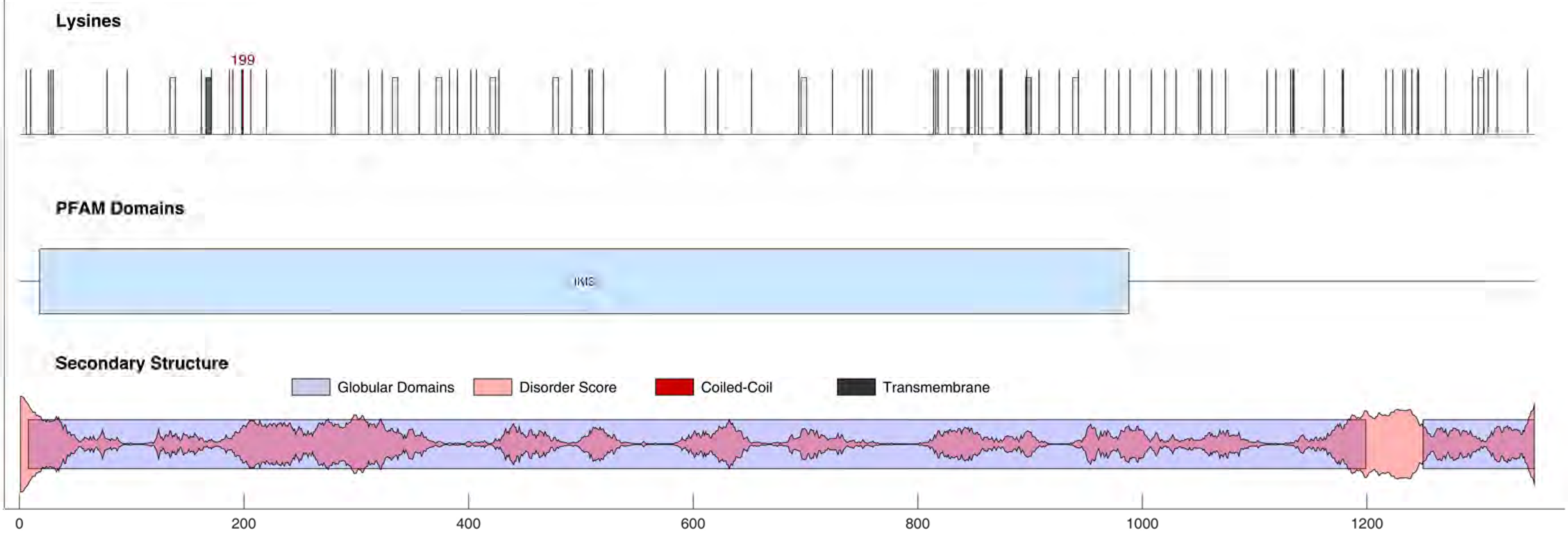

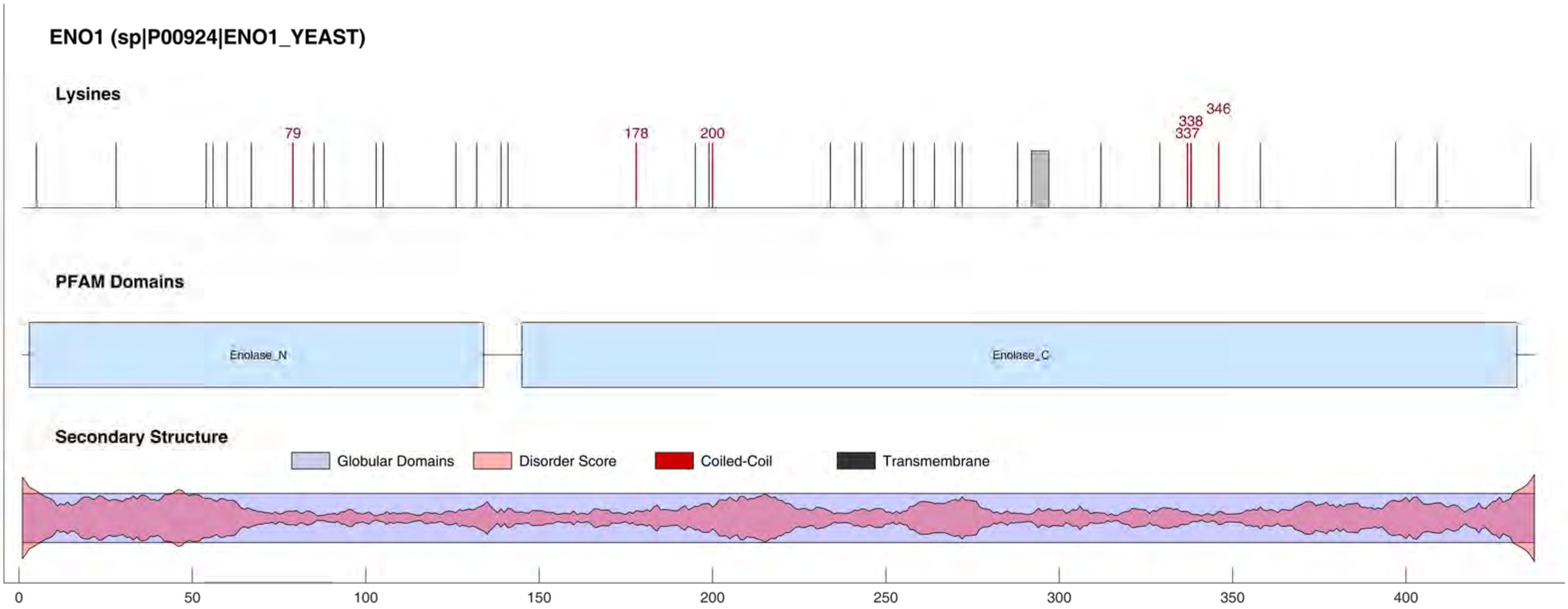

ENO1 (sp|P00924|ENO1_YEAST)
Lysines
79
178
200
337
338
346
PFAM Domains
Enolase_N
Enolase_C
Secondary Structure
Globular Domains
Disorder Score
Coiled-Coil
Transmembrane
0
50
100
150
200
250
300
350
400

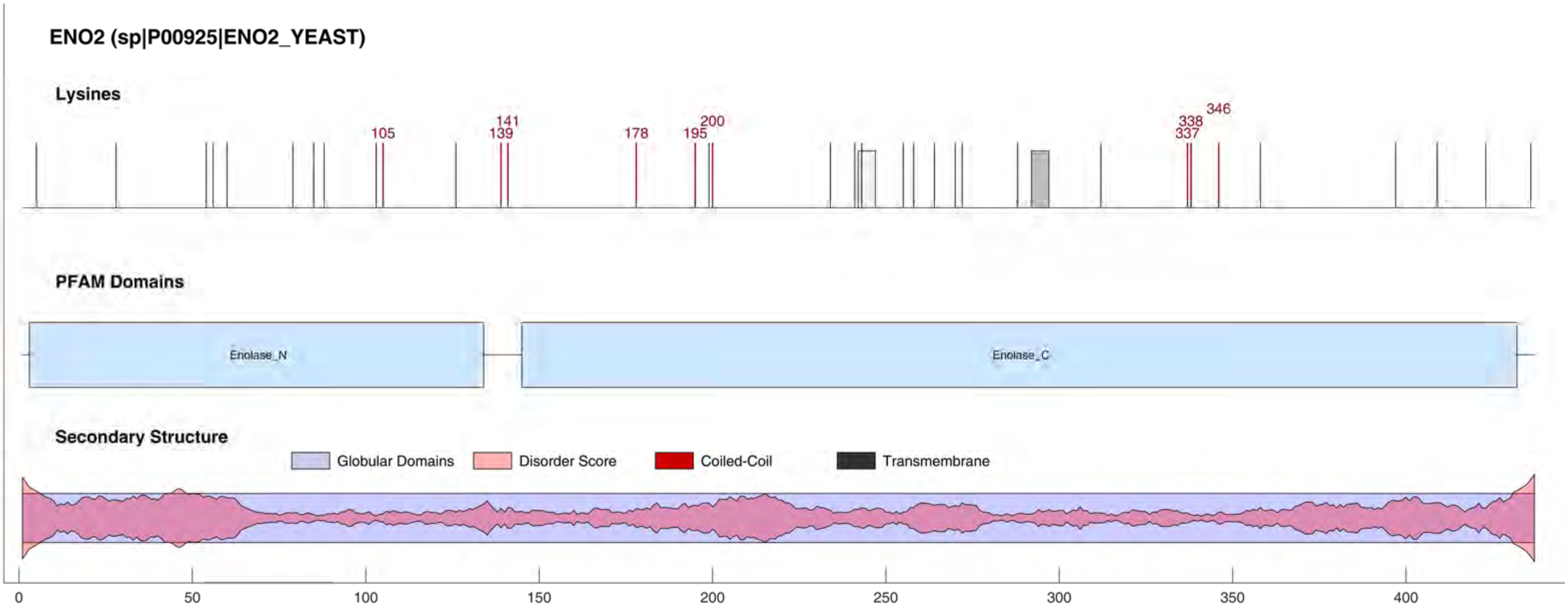
ENO2 (sp|P00925|ENO2_YEAST)
Lysines
105
141
139
178
195
200
338
337
346
PFAM Domains
Enolase_N
Enolase_C
Secondary Structure
Globular Domains
Disorder Score
Coiled-Coil
Transmembrane
0
50
100
150
200
250
300
350
400

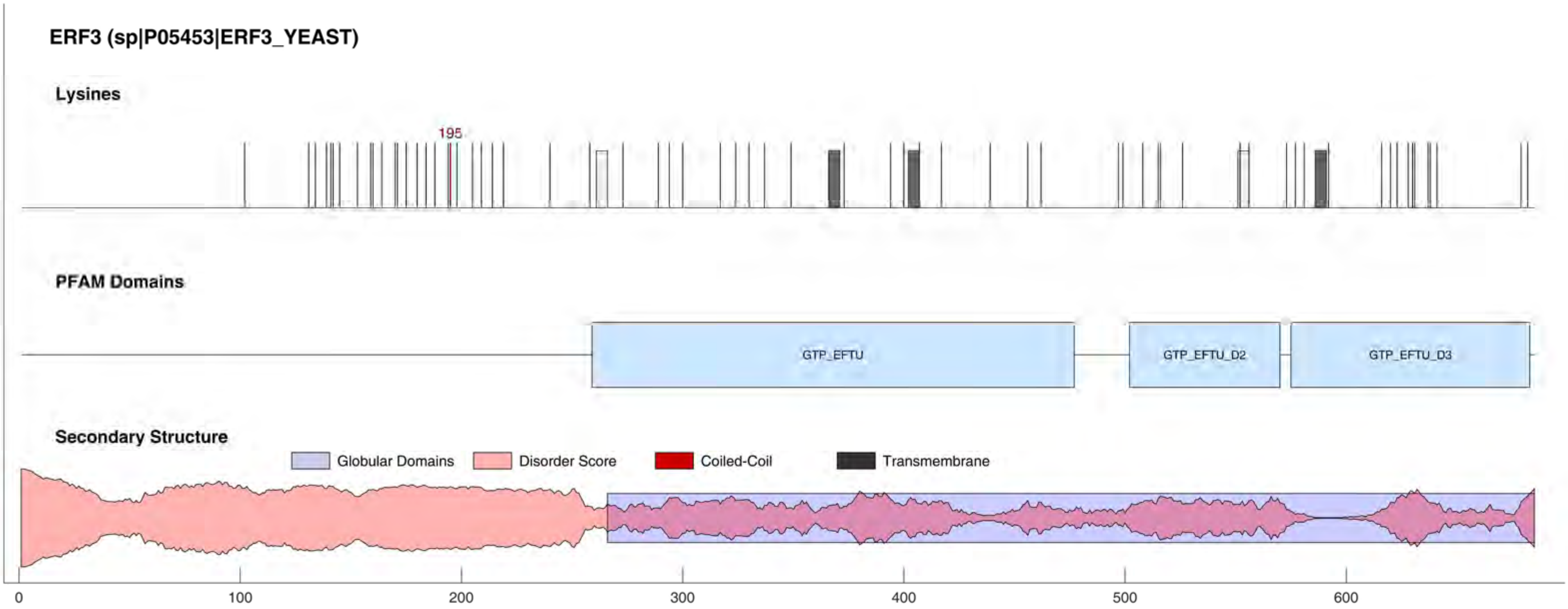
ERF3 (sp|P05453|ERF3_YEAST)
Lysines
195
PFAM Domains
GTP_EFTU
GTP_EFTU_D2
GTP_EFTU_D3
Secondary Structure
Globular Domains
Disorder Score
Coiled-Coil
Transmembrane
0
100
200
300
400
500
600

#### ESC1 (sp|Q03661|ESC1_YEAST)

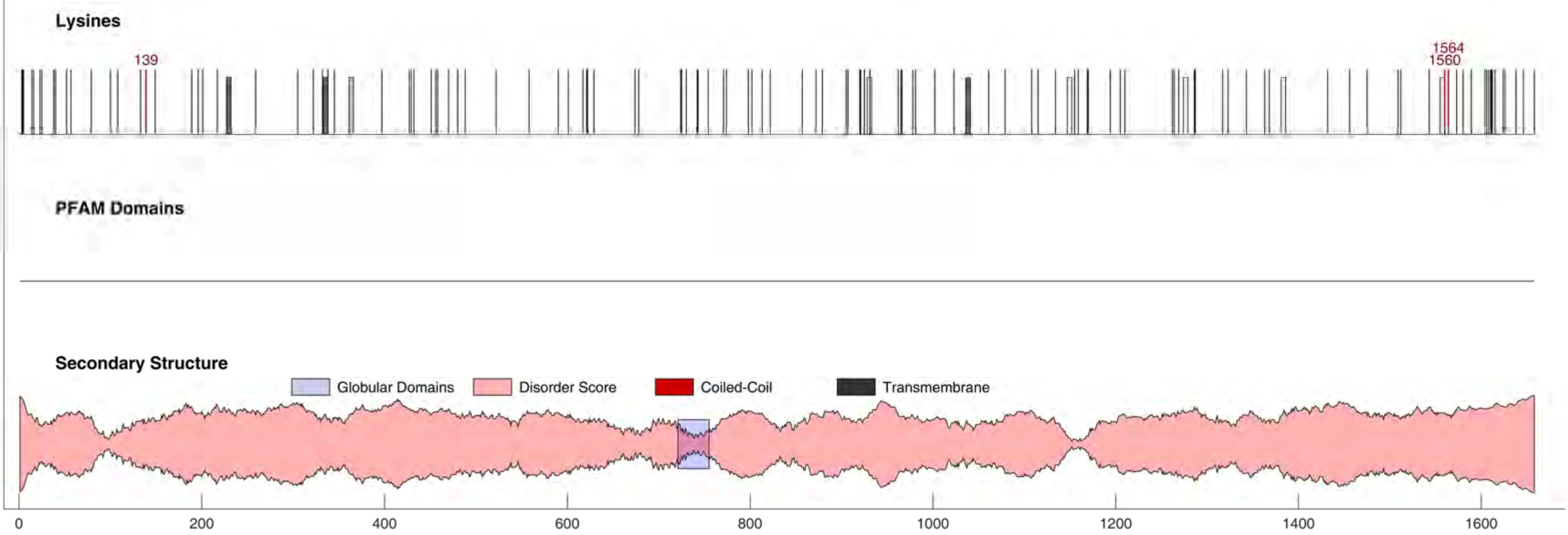

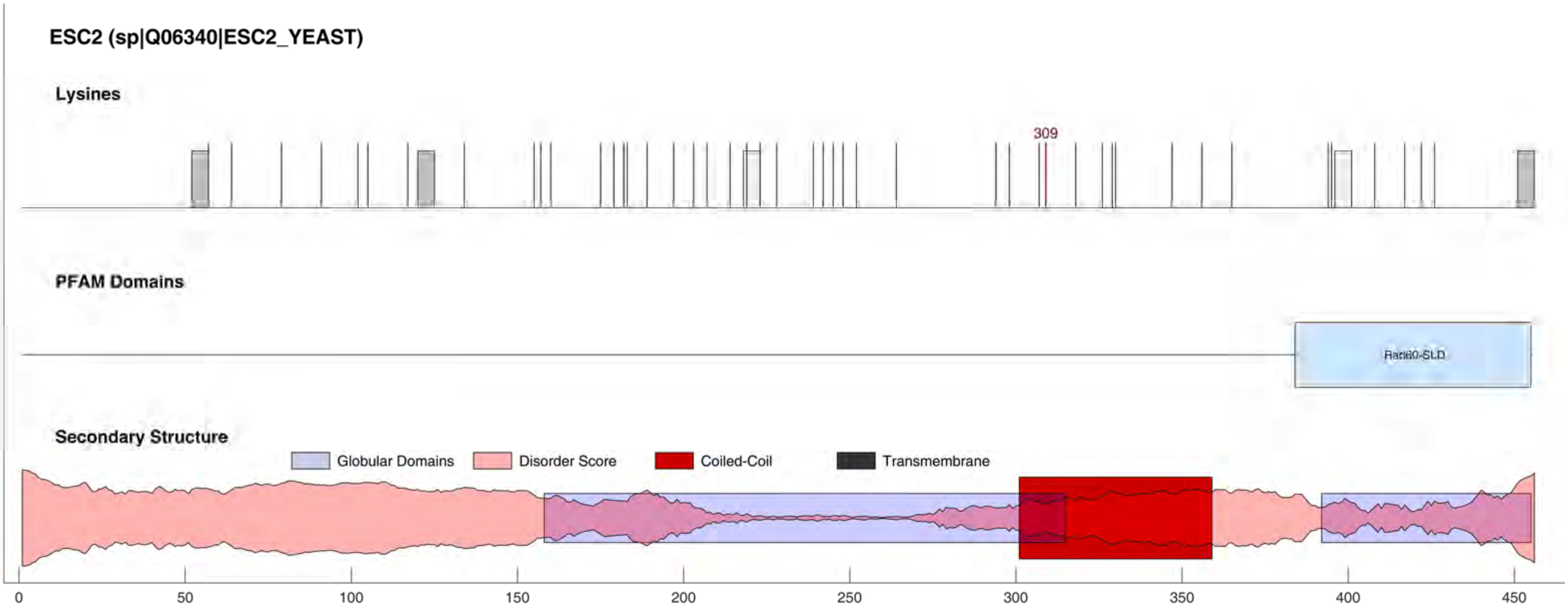
ESC2 (sp|Q06340|ESC2_YEAST)
Lysines
309
PFAM Domains
Rad60-SLD
Secondary Structure
Globular Domains
Disorder Score
Coiled-Coil
Transmembrane
0
50
100
150
200
250
300
350
400
450

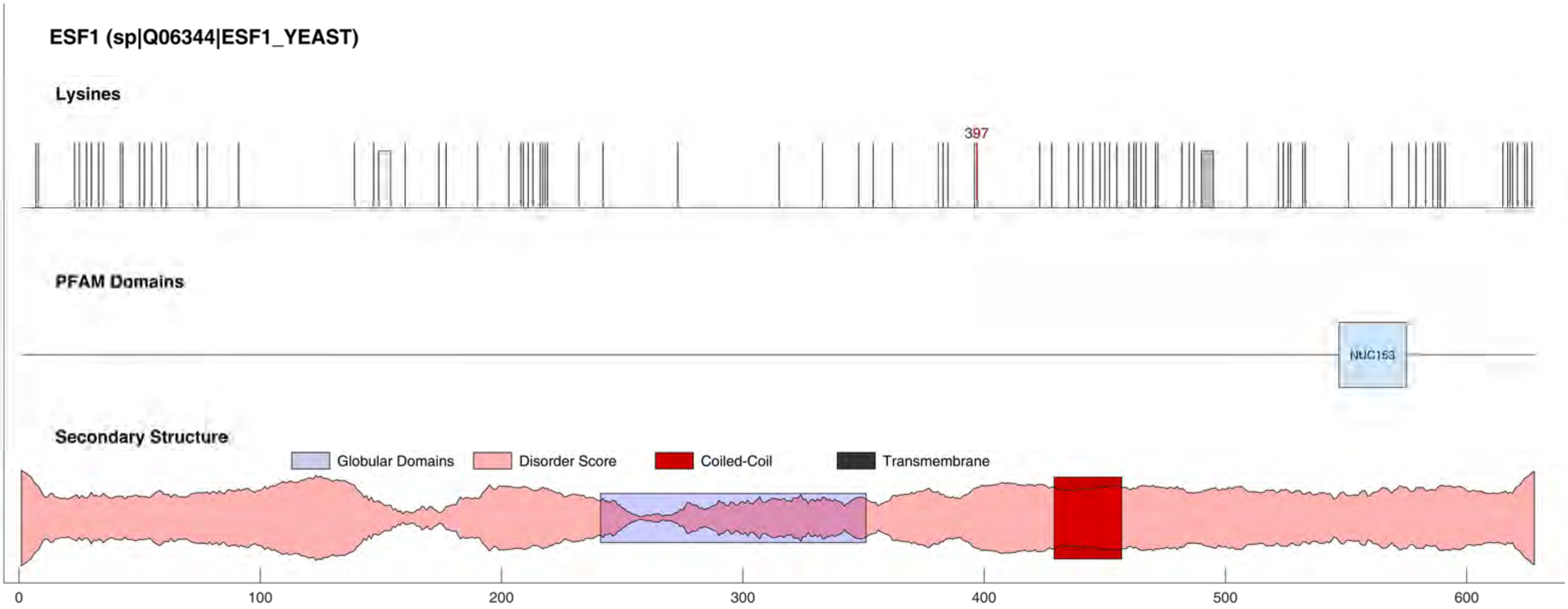
ESF1 (sp|Q06344|ESF1_YEAST)
Lysines
397
PFAM Domains
NUC153
Secondary Structure
Globular Domains
Disorder Score
Coiled-Coil
Transmembrane
0
100
200
300
400
500
600

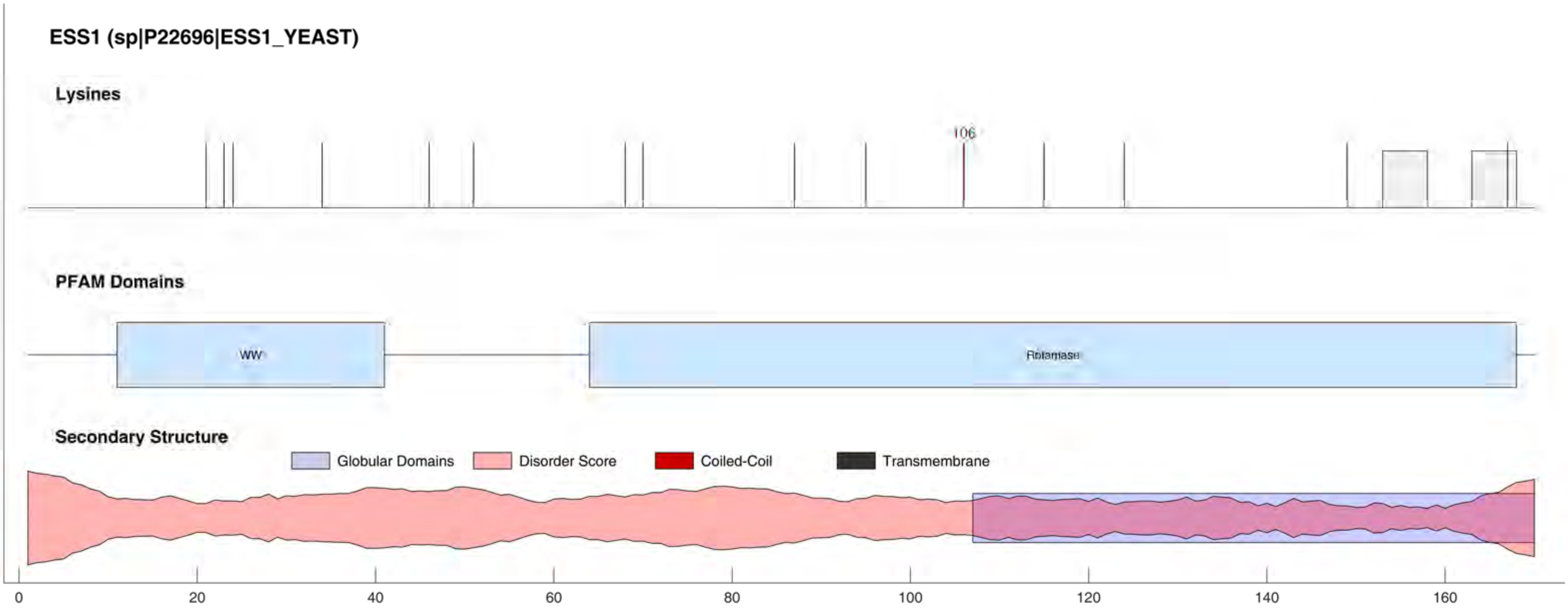
ESS1 (sp|P22696|ESS1_YEAST)
Lysines
106
PFAM Domains
WW
Rotamase
Secondary Structure
Globular Domains
Disorder Score
Coiled-Coil
Transmembrane
0
20
40
60
80
100
120
140
160

**EXO70 (sp|P19658|EXO70_YEAST)**

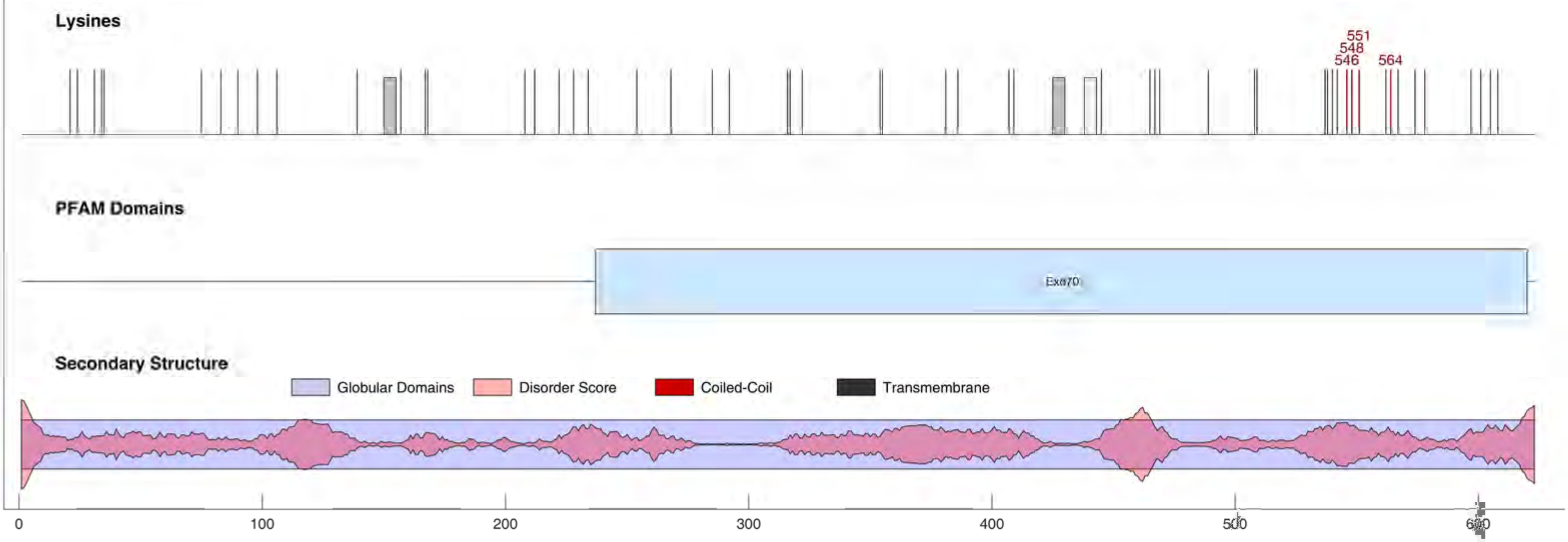

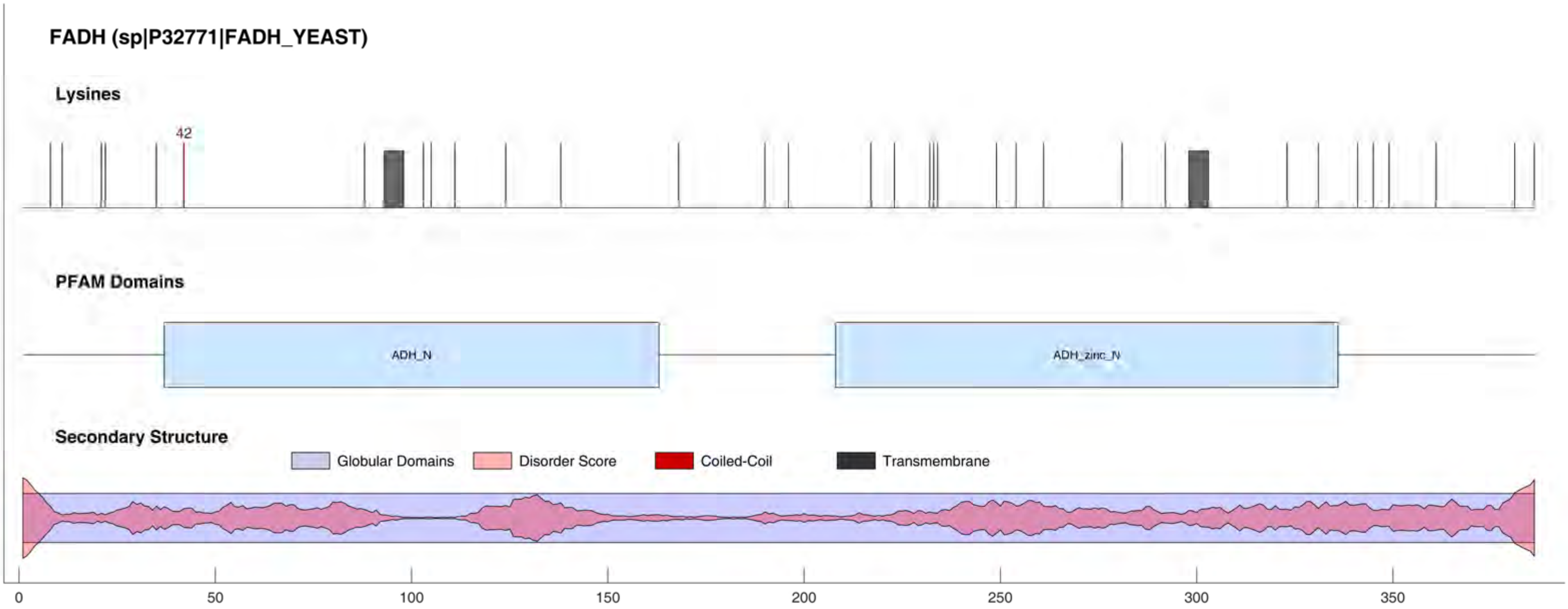

FADH (sp|P32771|FADH_YEAST)
Lysines
42
PFAM Domains
ADH_N
ADH_zinc_N
Secondary Structure
Globular Domains
Disorder Score
Coiled-Coil
Transmembrane
0
50
100
150
200
250
300
350

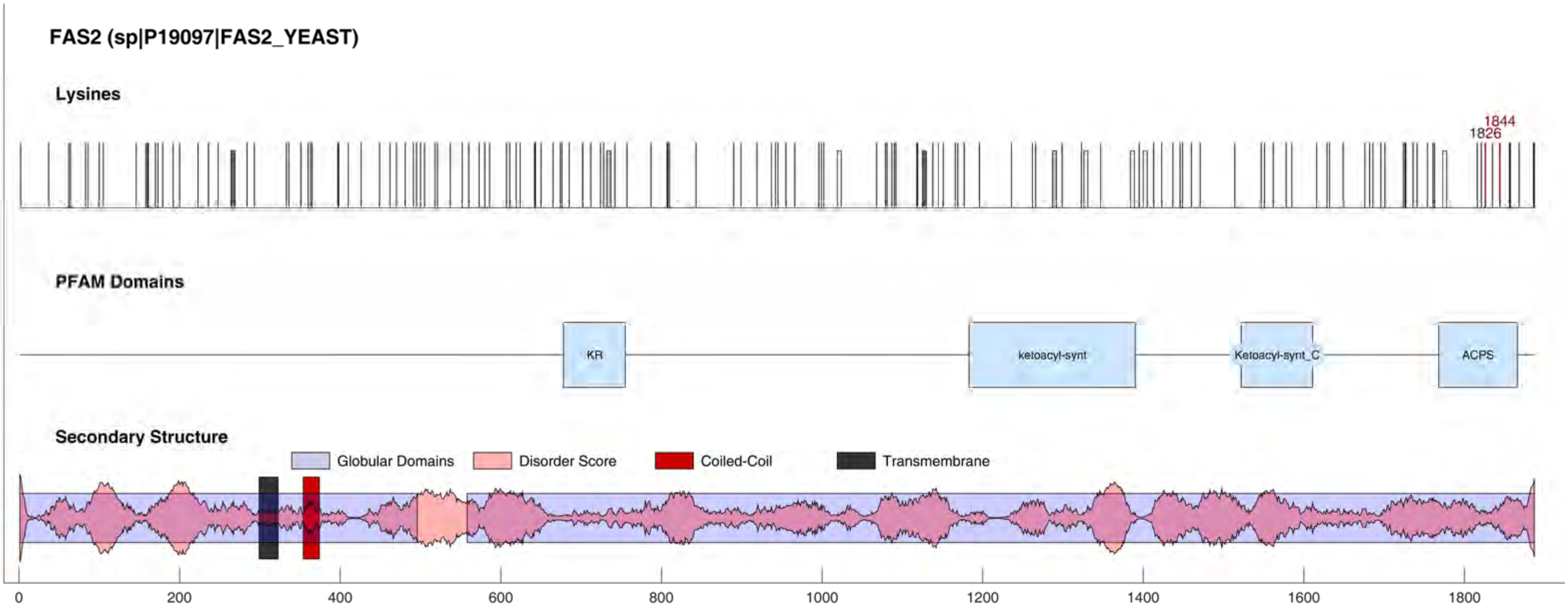
FAS2 (sp|P19097|FAS2_YEAST)
Lysines
1844
1826
PFAM Domains
KR
ketoacyl-synt
Ketoacyl-synt_C
ACPS
Secondary Structure
Globular Domains
Disorder Score
Coiled-Coil
Transmembrane
0
200
400
600
800
1000
1200
1400
1600
1800

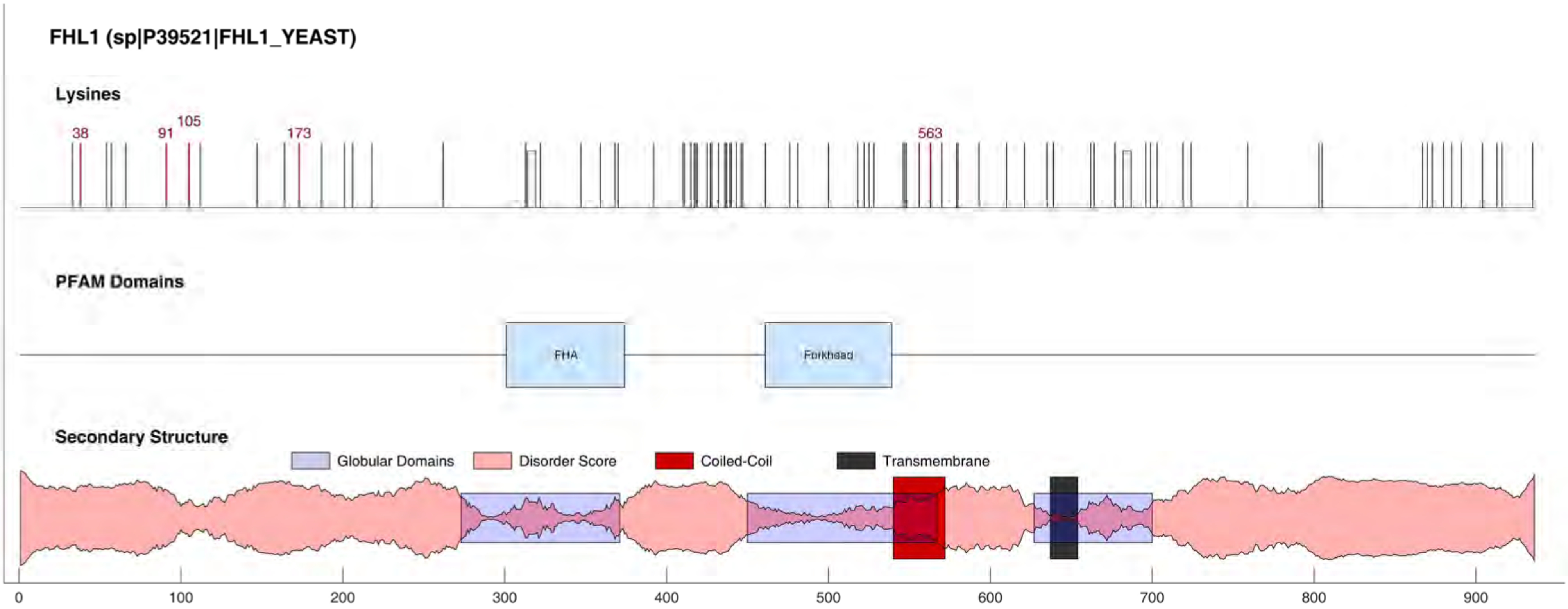

FHL1 (sp|P39521|FHL1_YEAST)
Lysines
38
91
105
173
563
PFAM Domains
FHA
Forkhead
Secondary Structure
Globular Domains
Disorder Score
Coiled-Coil
Transmembrane
0
100
200
300
400
500
600
700
800
900

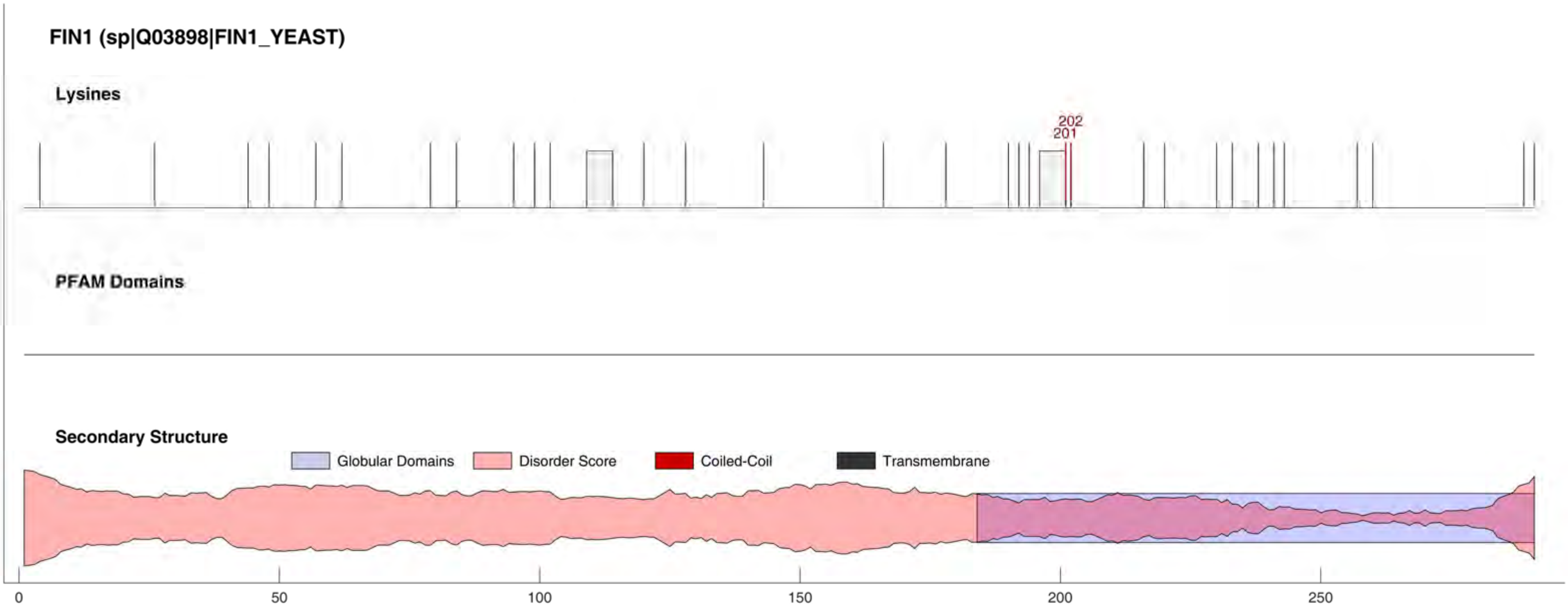

FIN1 (sp|Q03898|FIN1_YEAST)
Lysines
202
201
PFAM Domains
Secondary Structure
Globular Domains
Disorder Score
Coiled-Coil
Transmembrane
0
50
100
150
200
250

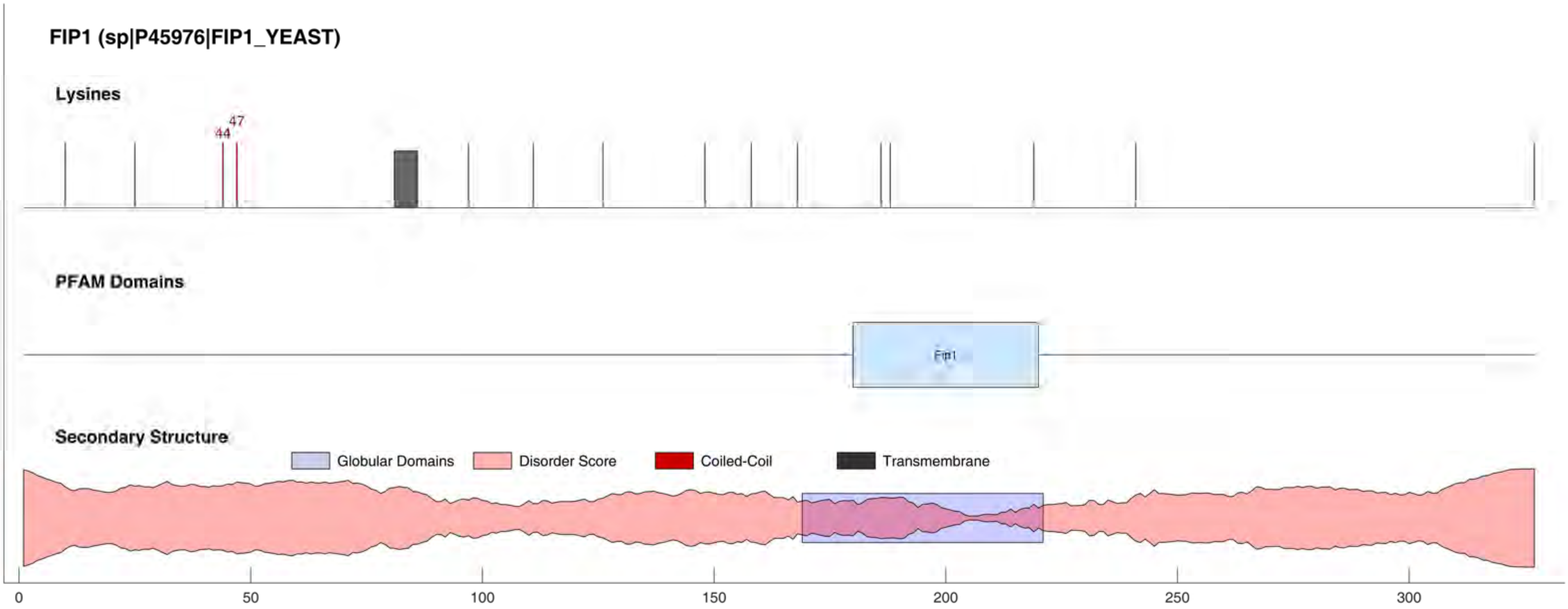

FIP1 (sp|P45976|FIP1_YEAST)
Lysines
44
47
PFAM Domains
Fip1
Secondary Structure
Globular Domains
Disorder Score
Coiled-Coil
Transmembrane
0
50
100
150
200
250
300

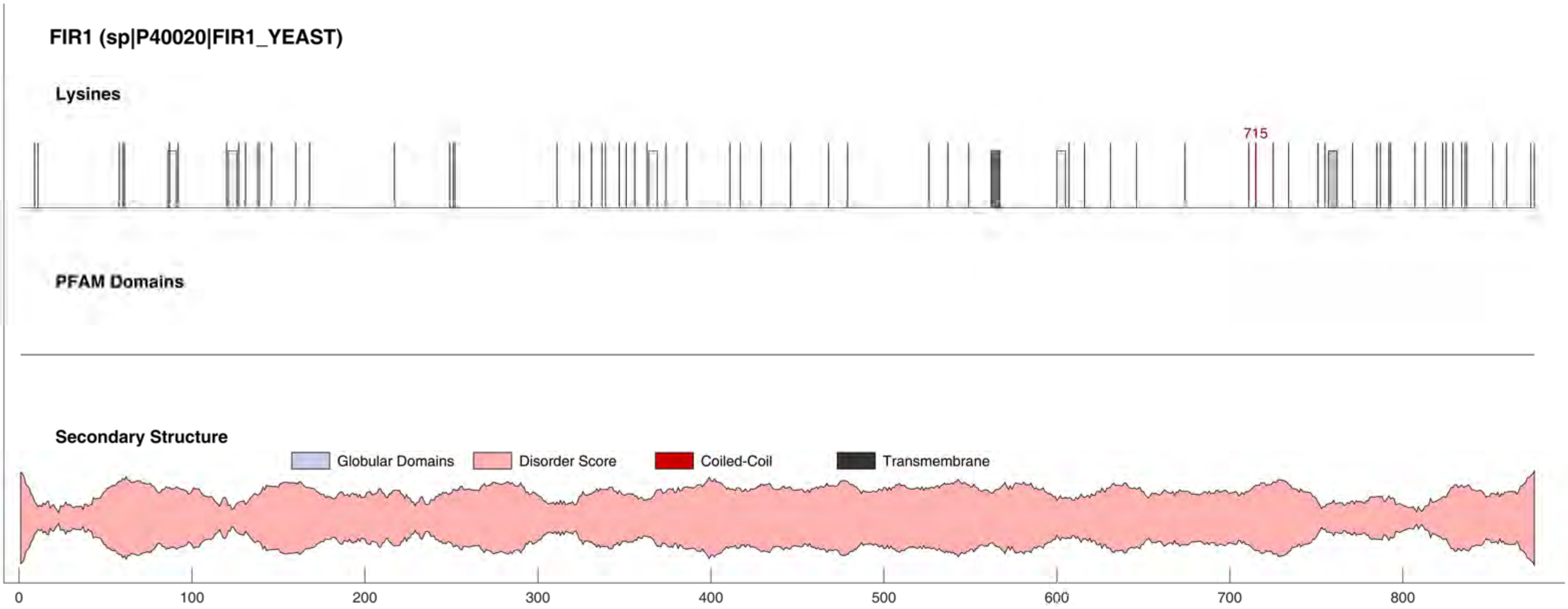
FIR1 (sp|P40020|FIR1_YEAST)
Lysines
715
PFAM Domains
Secondary Structure
Globular Domains
Disorder Score
Coiled-Coil
Transmembrane
0
100
200
300
400
500
600
700
800

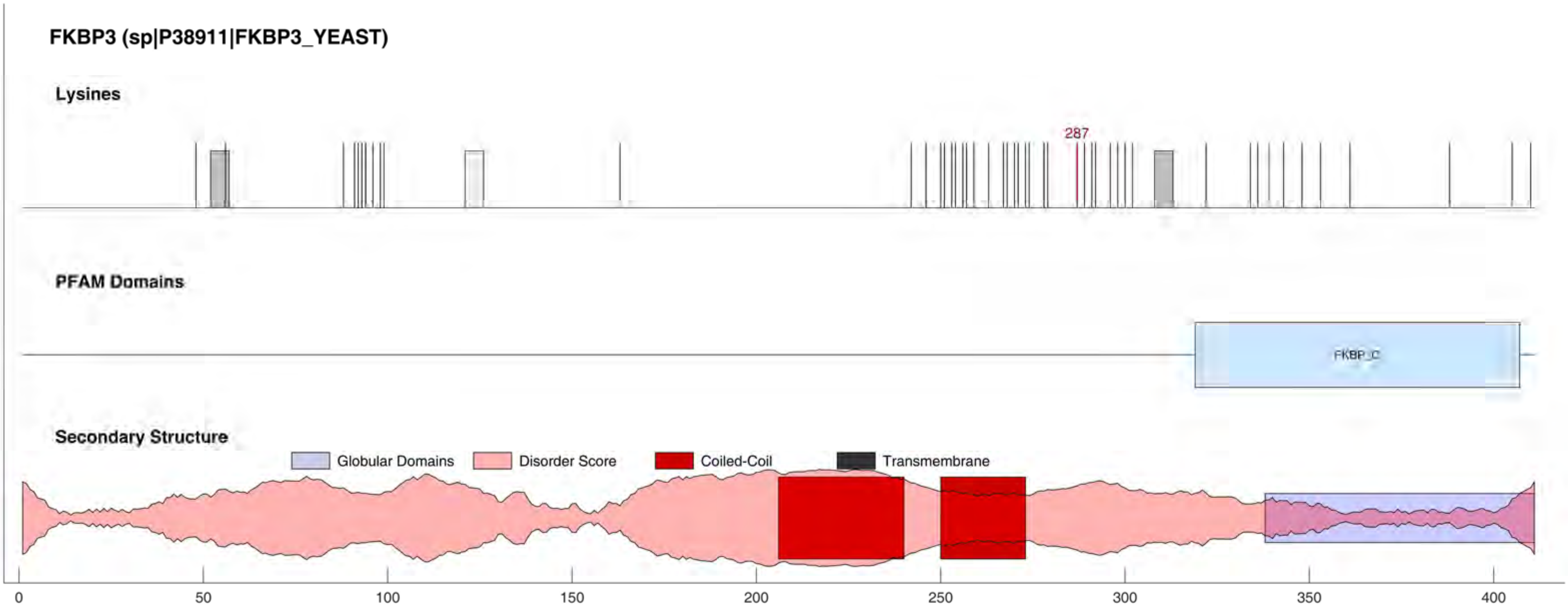

FKBP3 (sp|P38911|FKBP3_YEAST)
Lysines
287
PFAM Domains
FKBP_C
Secondary Structure
Globular Domains
Disorder Score
Coiled-Coil
Transmembrane
0
50
100
150
200
250
300
350
400

#### FKH1 (sp|P40466|FKH1_YEAST)

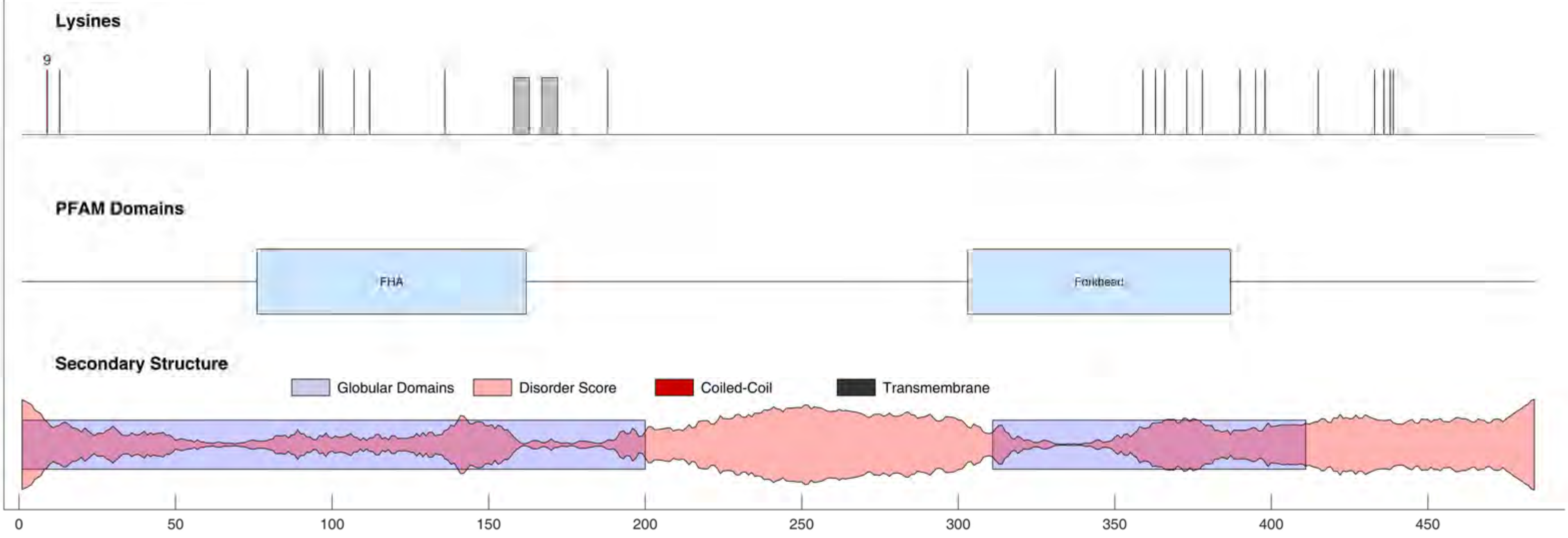

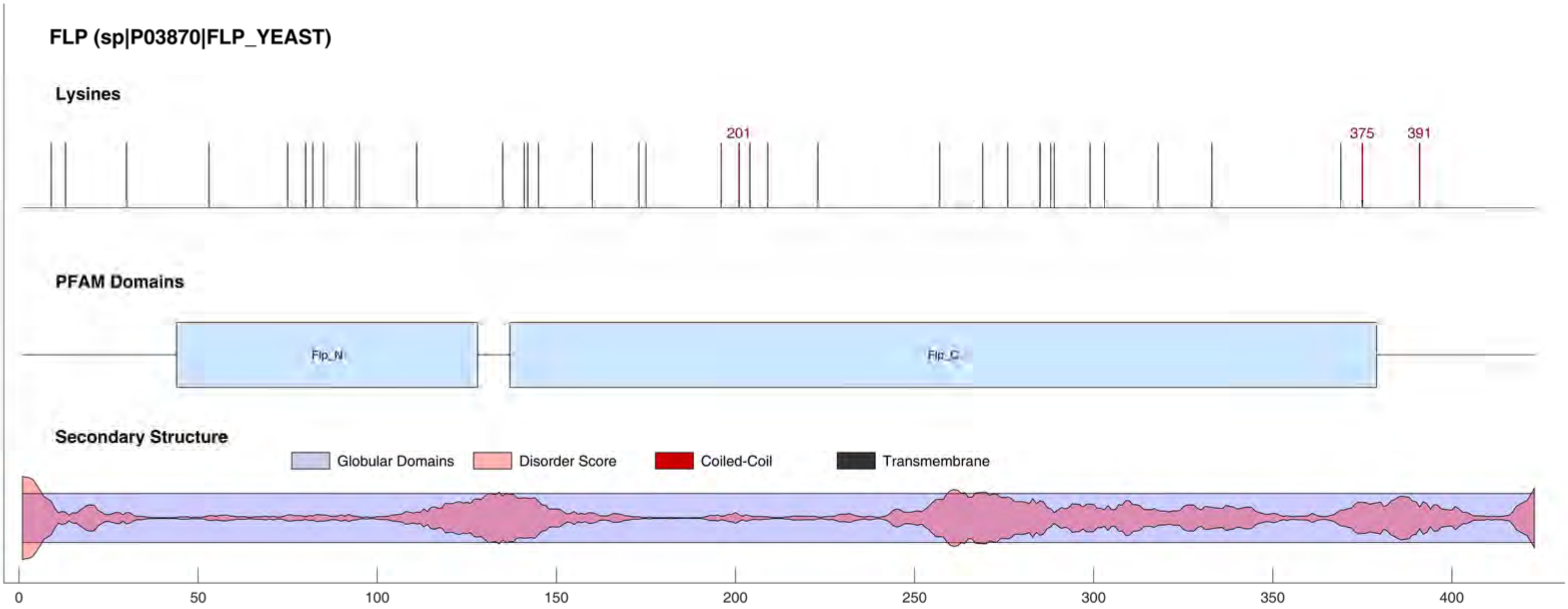

FLP (sp|P03870|FLP_YEAST)
Lysines
201
375
391
PFAM Domains
Flp_N
Flp_C
Secondary Structure
Globular Domains
Disorder Score
Coiled-Coil
Transmembrane
0
50
100
150
200
250
300
350
400

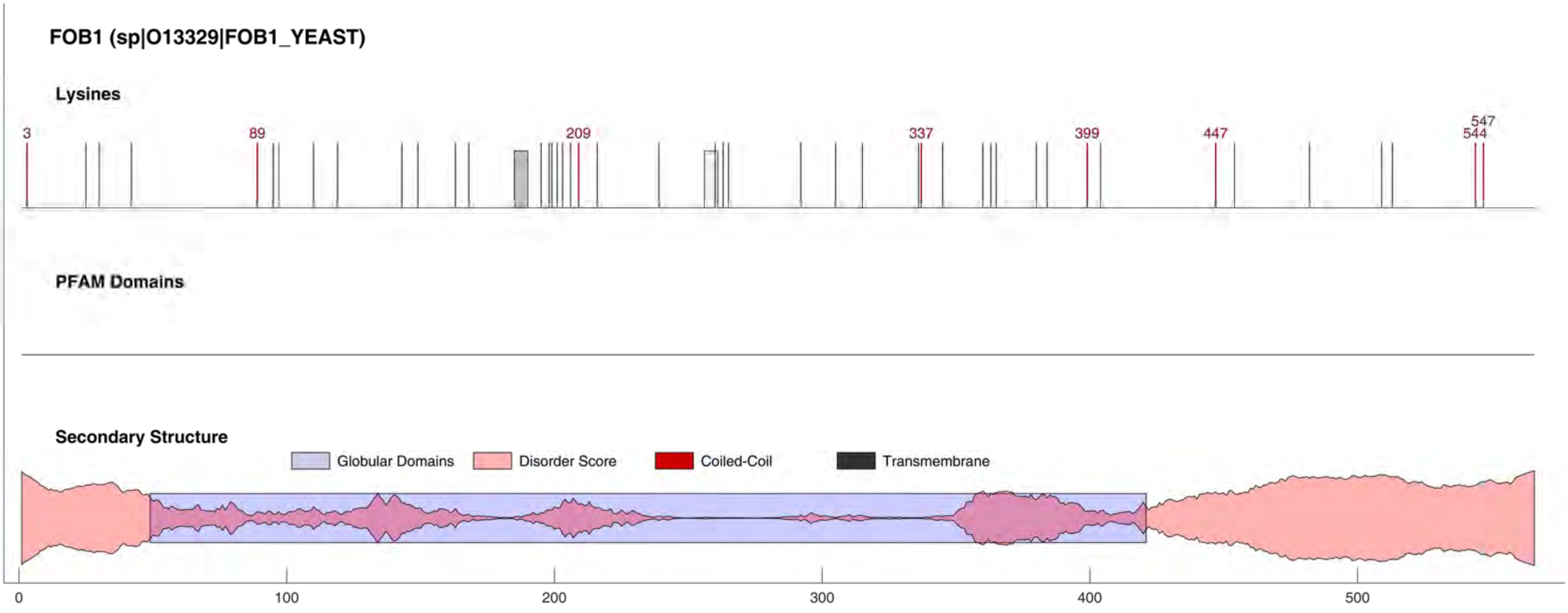
FOB1 (sp|O13329|FOB1_YEAST)
Lysines
3
89
209
337
399
447
547
544
PFAM Domains
Secondary Structure
Globular Domains
Disorder Score
Coiled-Coil
Transmembrane
0
100
200
300
400
500

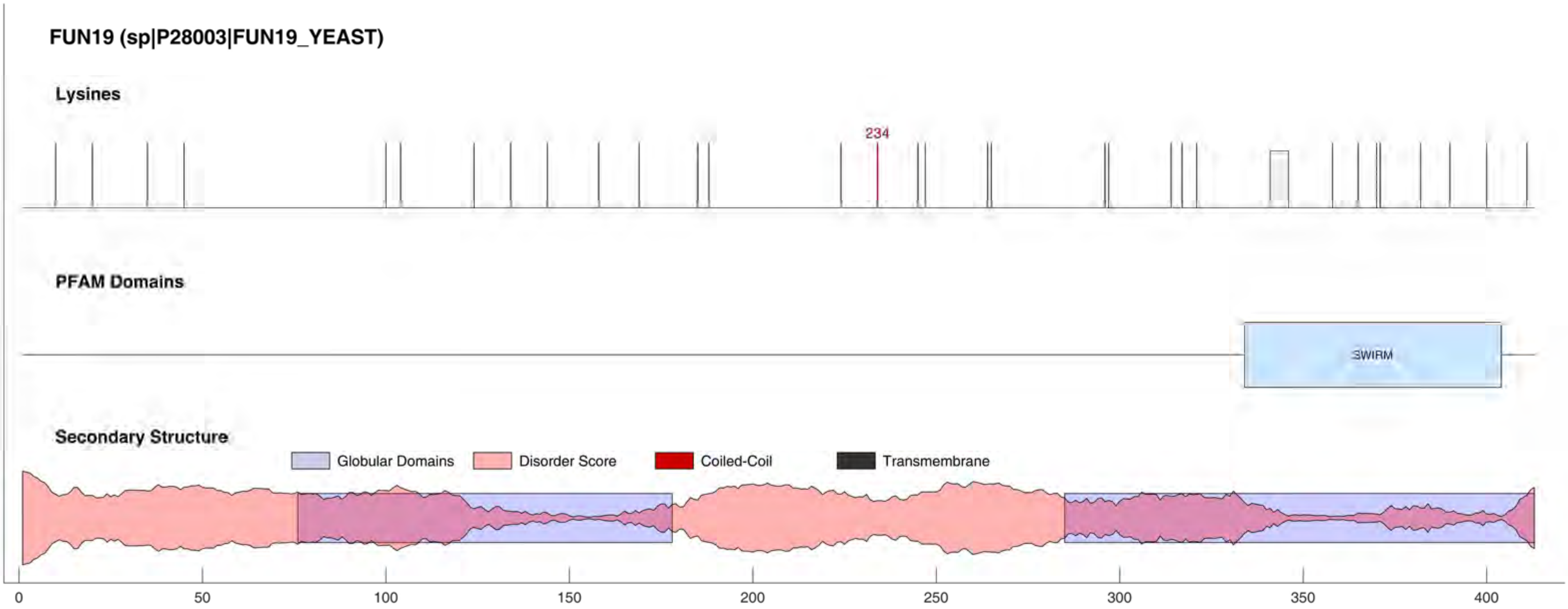
FUN19 (sp|P28003|FUN19_YEAST)
Lysines
234
PFAM Domains
SWIRM
Secondary Structure
Globular Domains
Disorder Score
Coiled-Coil
Transmembrane
0
50
100
150
200
250
300
350
400

#### FYV7 (sp|Q12247|FYV7_YEAST)

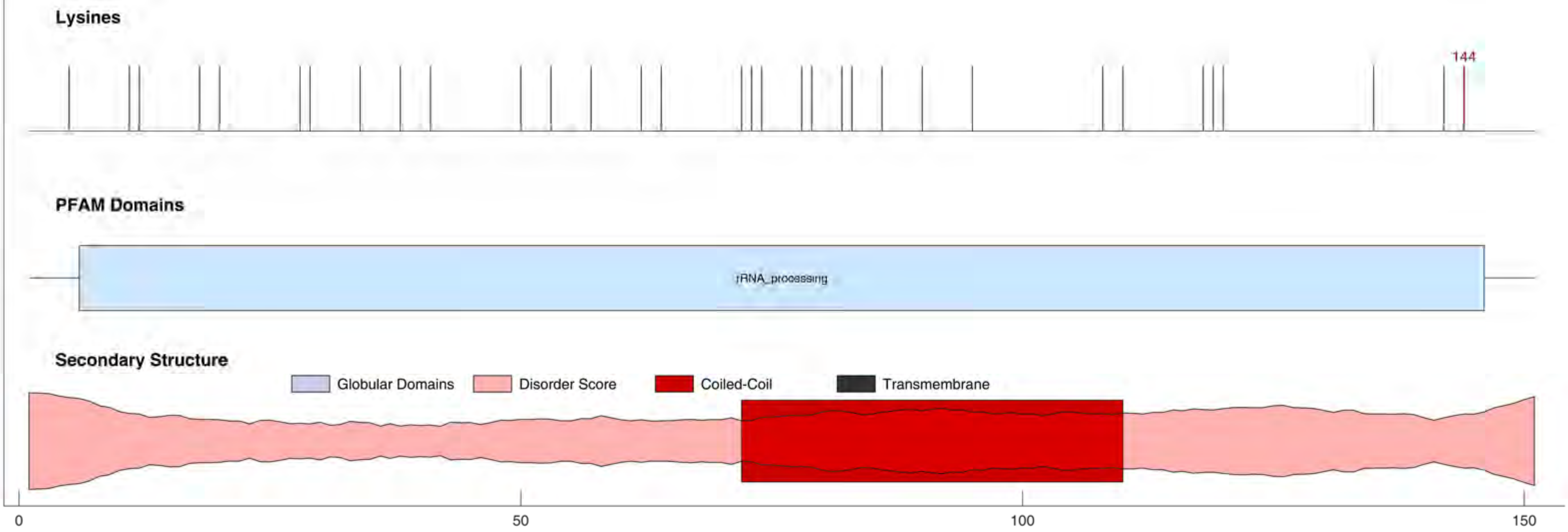

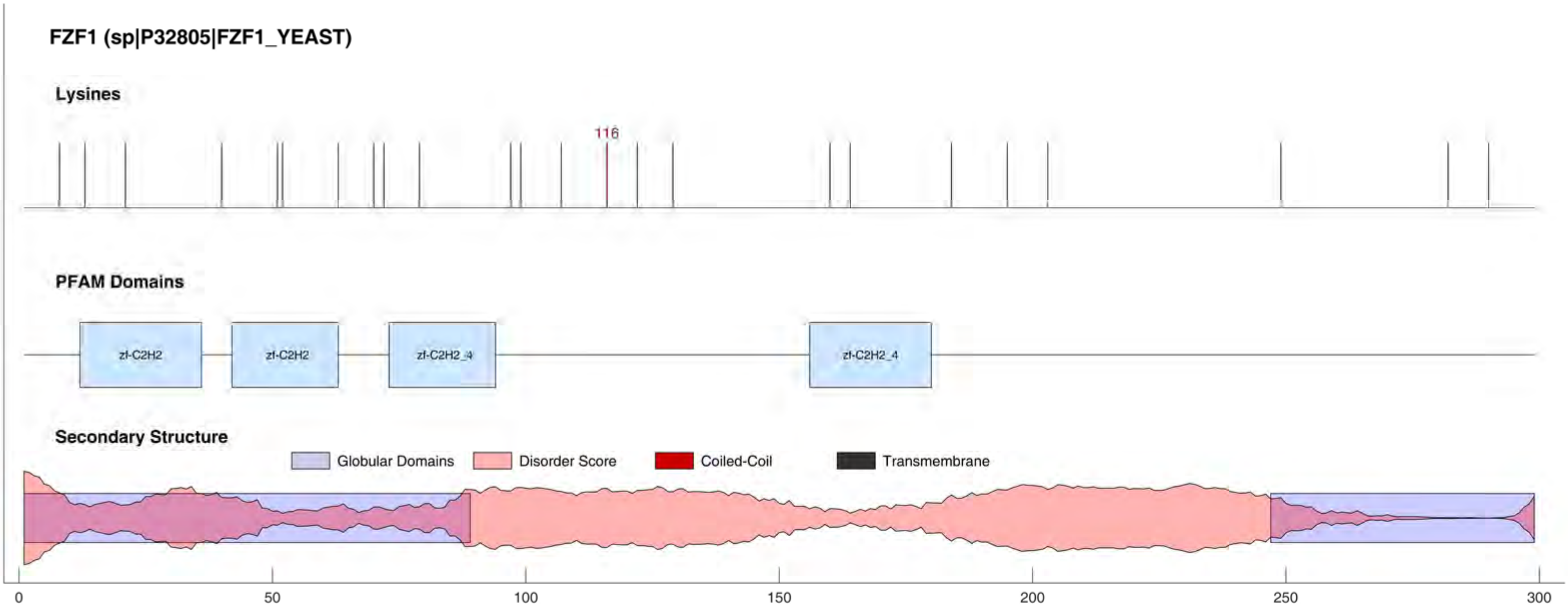

FZF1 (sp|P32805|FZF1_YEAST)
Lysines
116
PFAM Domains
zf-C2H2
zf-C2H2
zf-C2H2_4
zf-C2H2_4
Secondary Structure
Globular Domains
Disorder Score
Coiled-Coil
Transmembrane
0
50
100
150
200
250
300

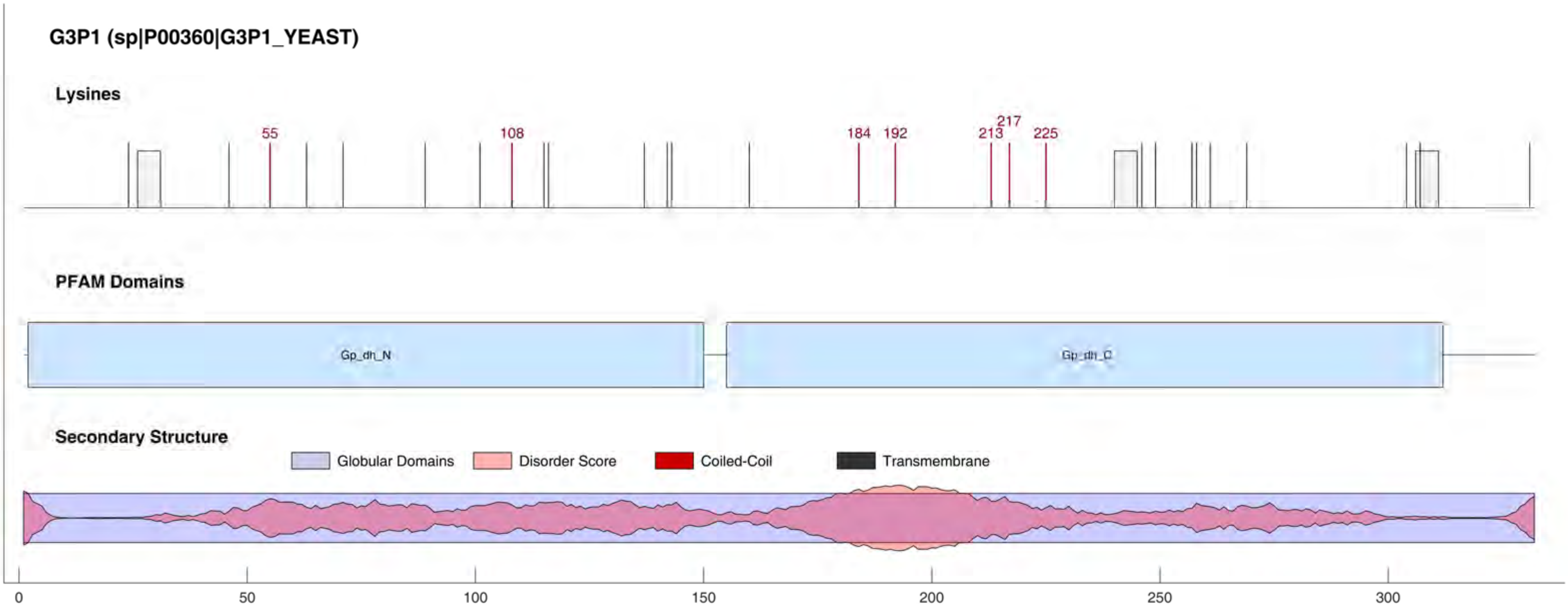
G3P1 (sp|P00360|G3P1_YEAST)
Lysines
55
108
184
192
213
217
225
PFAM Domains
Gp_dh_N
Gp_dh_C
Secondary Structure
Globular Domains
Disorder Score
Coiled-Coil
Transmembrane
0
50
100
150
200
250
300

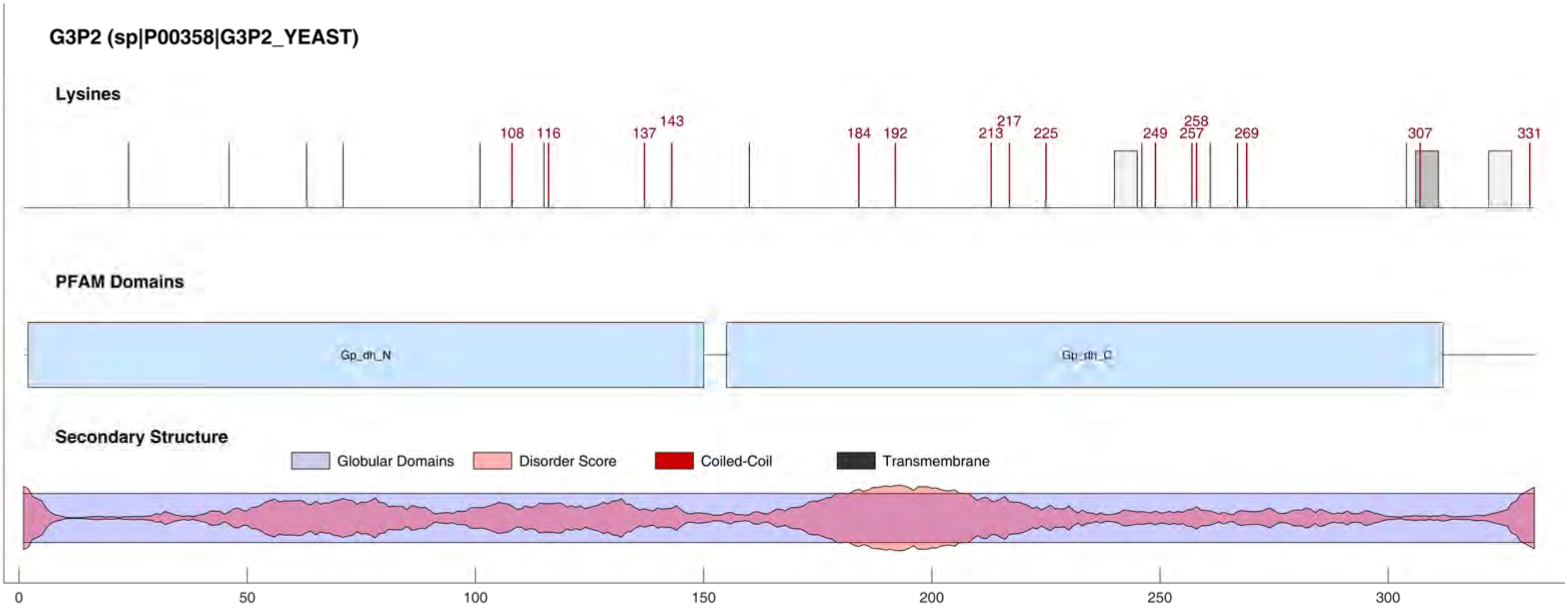
G3P2 (sp|P00358|G3P2_YEAST)
Lysines
108
116
137
143
184
192
213
217
225
249
257
258
269
307
331
PFAM Domains
Gp_dh_N
Gp_dh_C
Secondary Structure
Globular Domains
Disorder Score
Coiled-Coil
Transmembrane
0
50
100
150
200
250
300

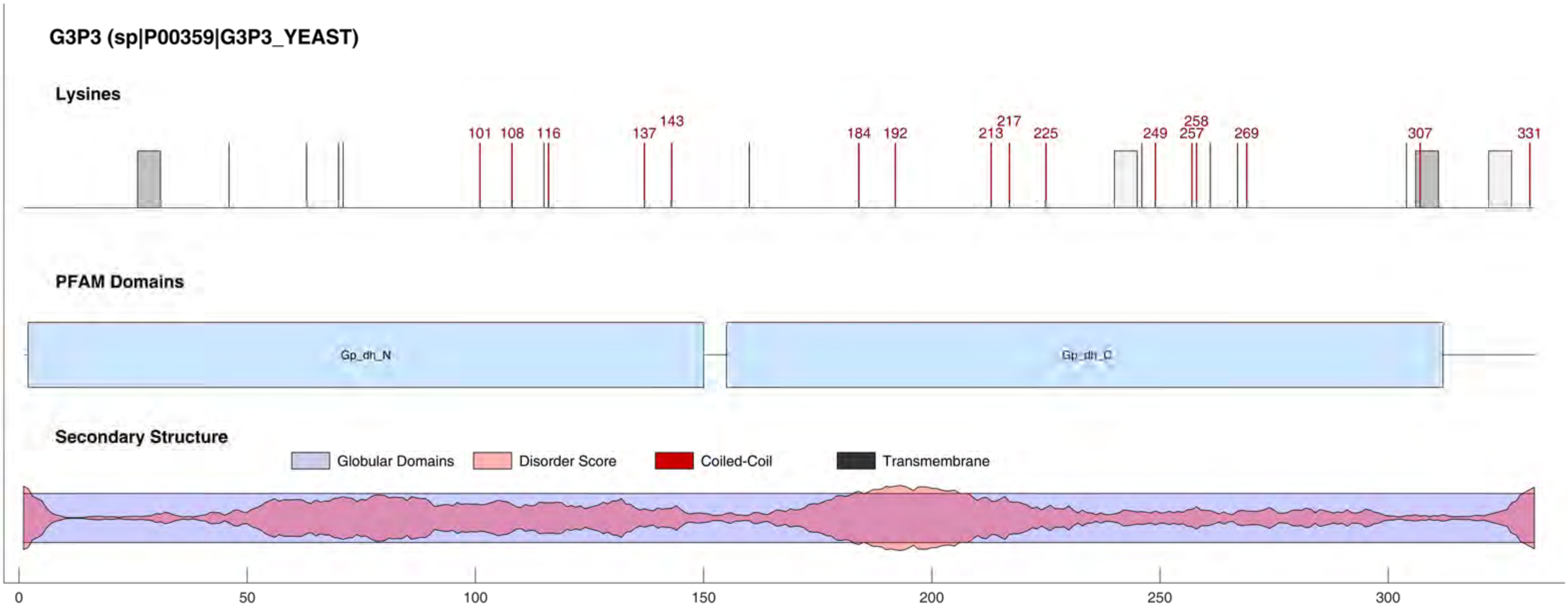
G3P3 (sp|P00359|G3P3_YEAST)
Lysines
101
108
116
137
143
184
192
213
217
225
249
257
258
269
307
331
PFAM Domains
Gp_dh_N
Gp_dh_C
Secondary Structure
Globular Domains
Disorder Score
Coiled-Coil
Transmembrane
0
50
100
150
200
250
300

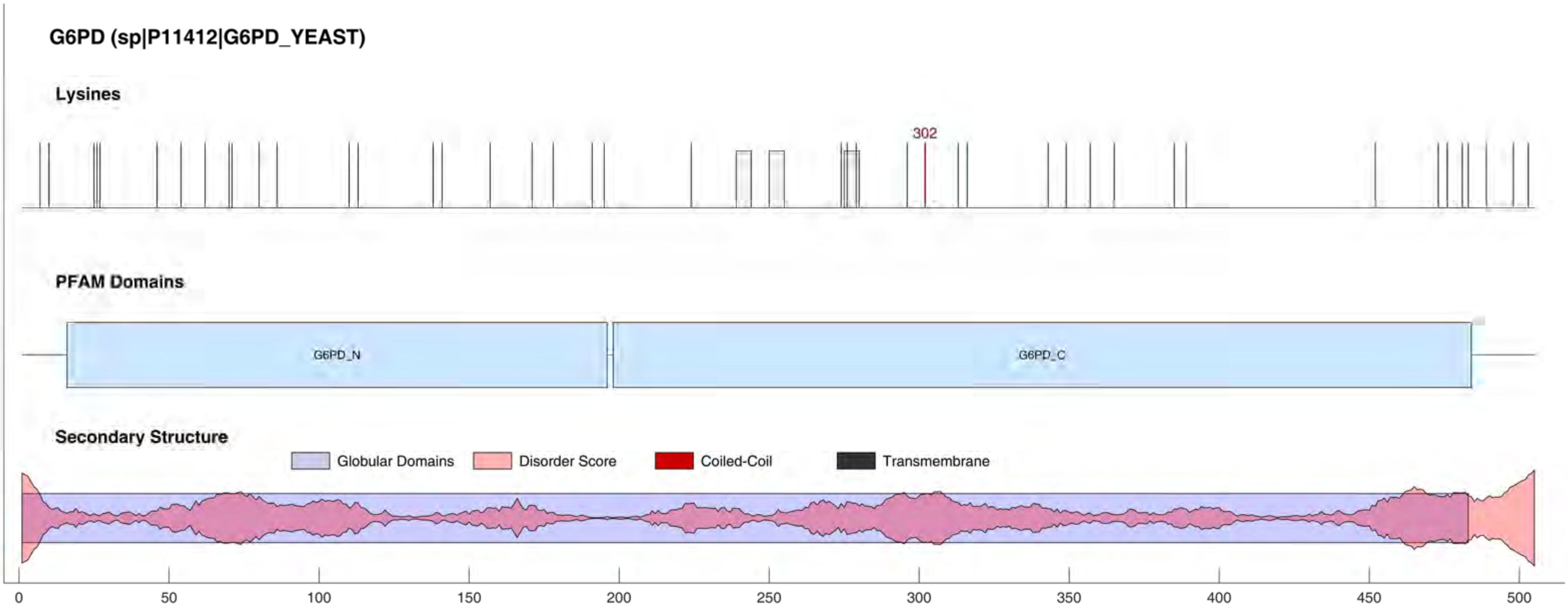
G6PD (sp|P11412|G6PD_YEAST)
Lysines
302
PFAM Domains
G6PD_N
G6PD_C
Secondary Structure
Globular Domains
Disorder Score
Coiled-Coil
Transmembrane
0
50
100
150
200
250
300
350
400
450
500

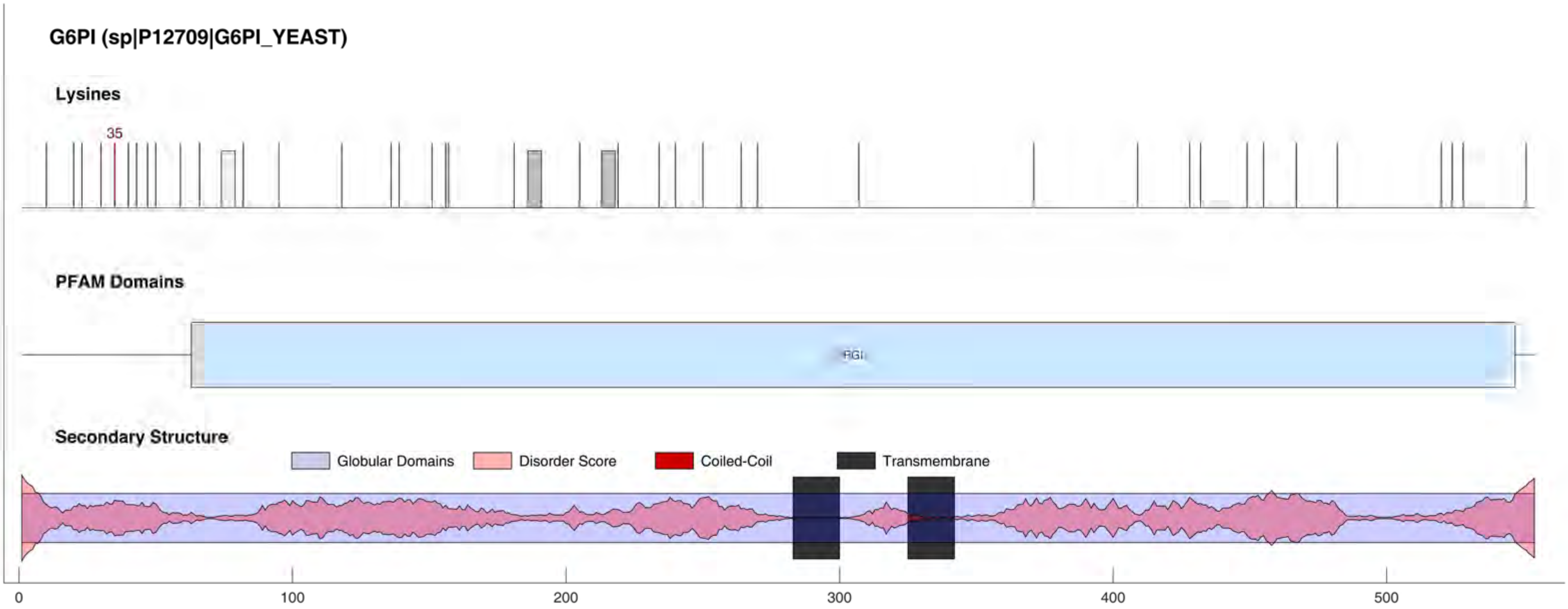

G6PI (sp|P12709|G6PI_YEAST)
Lysines
35
PFAM Domains
PGI
Secondary Structure
Globular Domains
Disorder Score
Coiled-Coil
Transmembrane
0
100
200
300
400
500

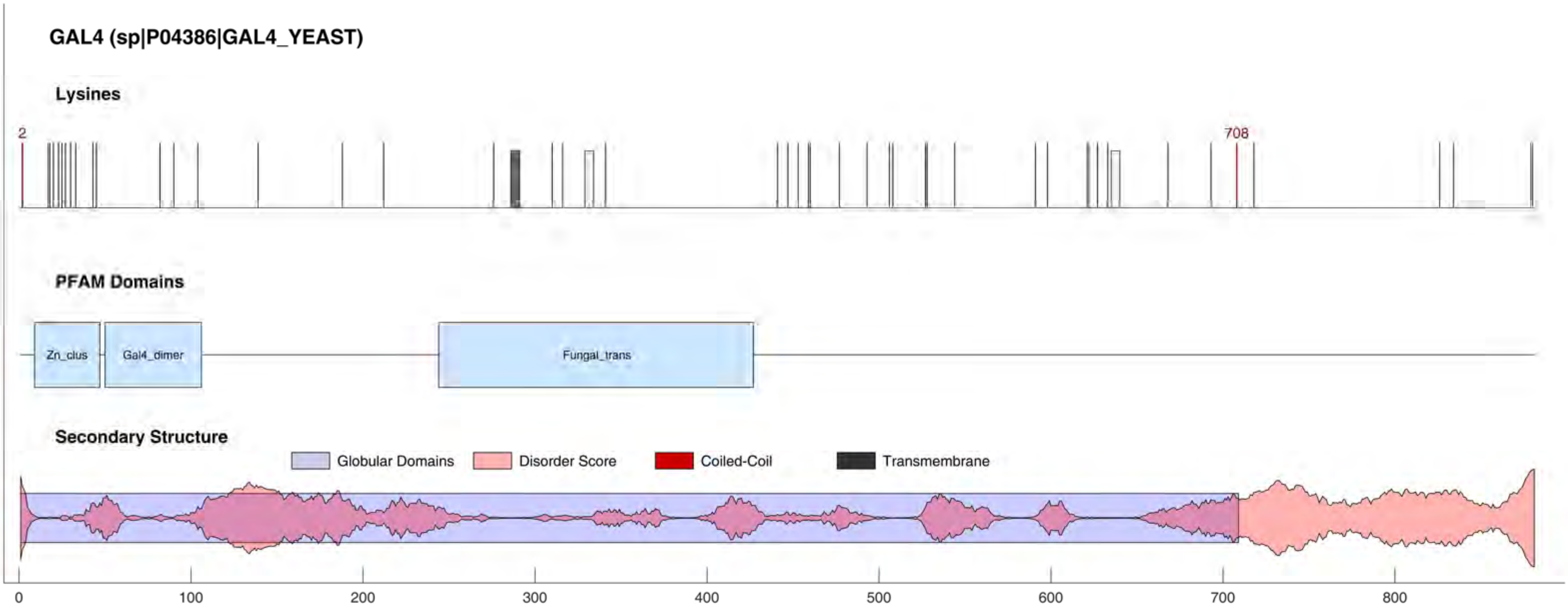

GAL4 (sp|P04386|GAL4_YEAST)
Lysines
2
708
PFAM Domains
Zn_clus
Gal4_dimer
Fungal_trans
Secondary Structure
Globular Domains
Disorder Score
Coiled-Coil
Transmembrane
0
100
200
300
400
500
600
700
800

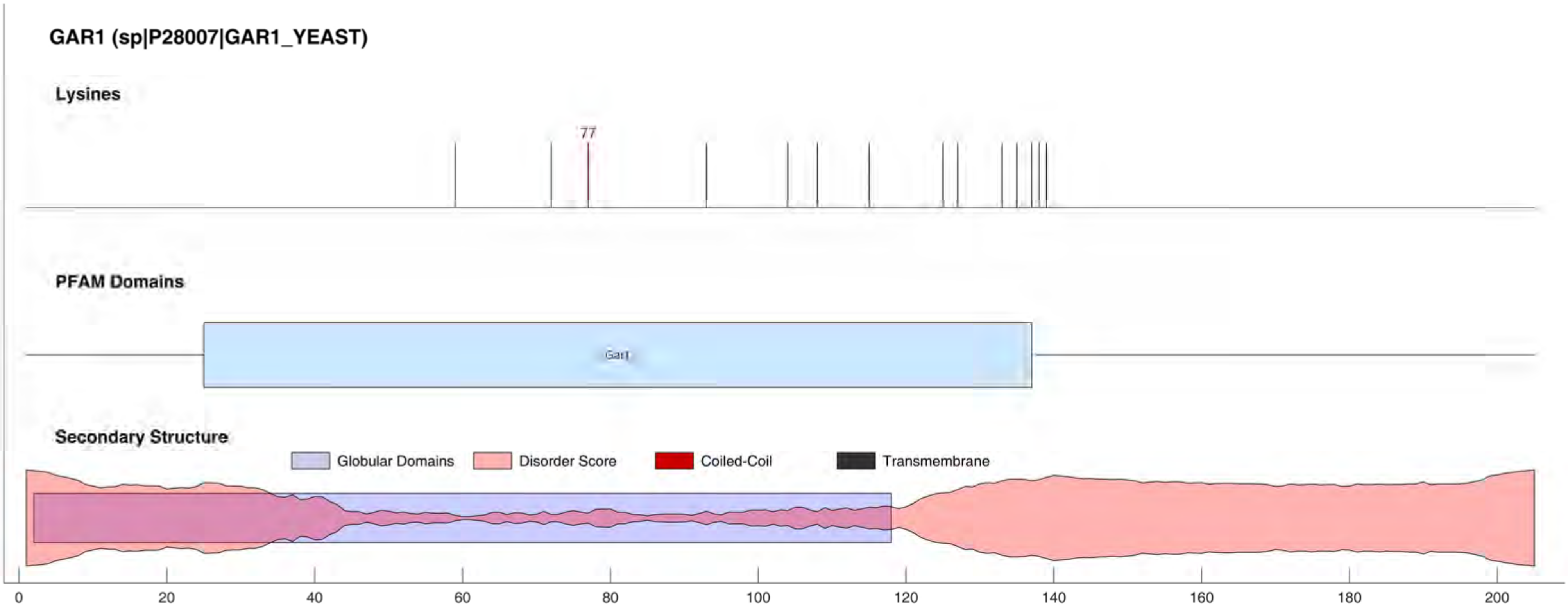

GAR1 (sp|P28007|GAR1_YEAST)
Lysines
77
PFAM Domains
Gar1
Secondary Structure
Globular Domains
Disorder Score
Coiled-Coil
Transmembrane
0
20
40
60
80
100
120
140
160
180
200

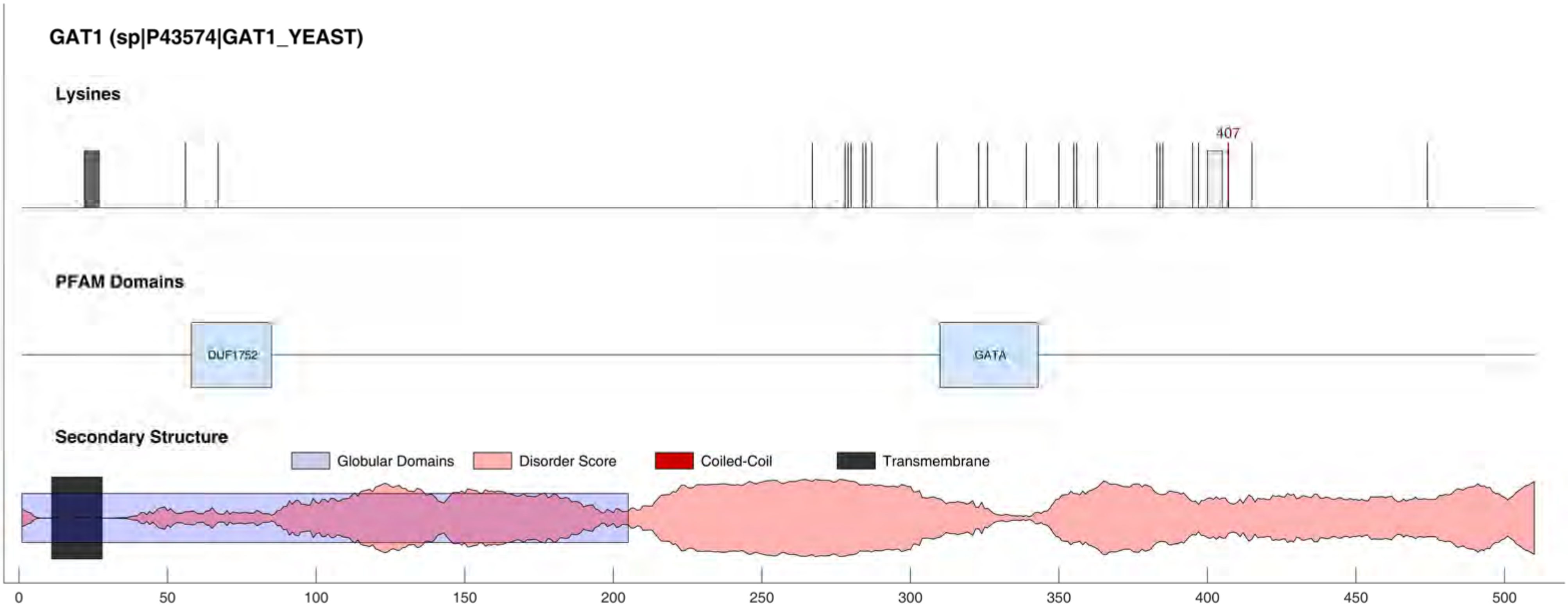
GAT1 (sp|P43574|GAT1_YEAST)
Lysines
407
PFAM Domains
DUF1752
GATA
Secondary Structure
Globular Domains
Disorder Score
Coiled-Coil
Transmembrane
0
50
100
150
200
250
300
350
400
450
500

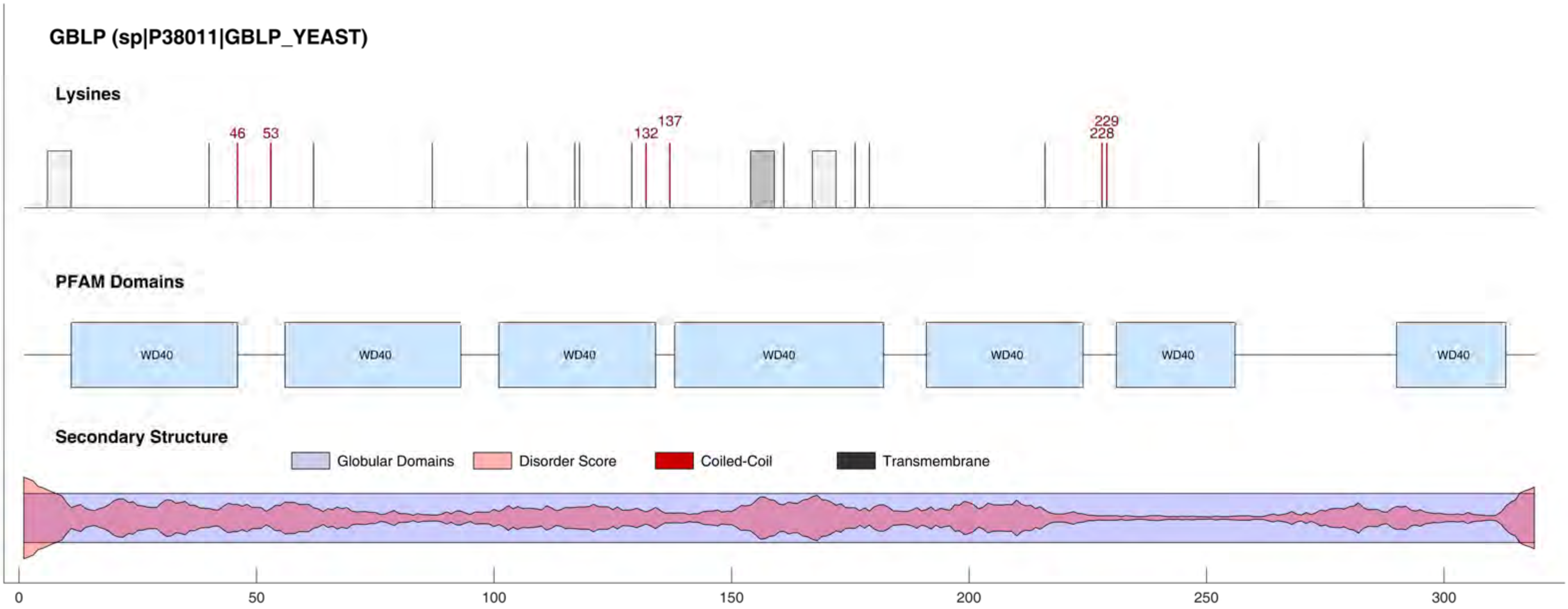
GBLP (sp|P38011|GBLP_YEAST)
Lysines
46
53
132
137
228
229
PFAM Domains
WD40
WD40
WD40
WD40
WD40
WD40
WD40
Secondary Structure
Globular Domains
Disorder Score
Coiled-Coil
Transmembrane
0
50
100
150
200
250
300

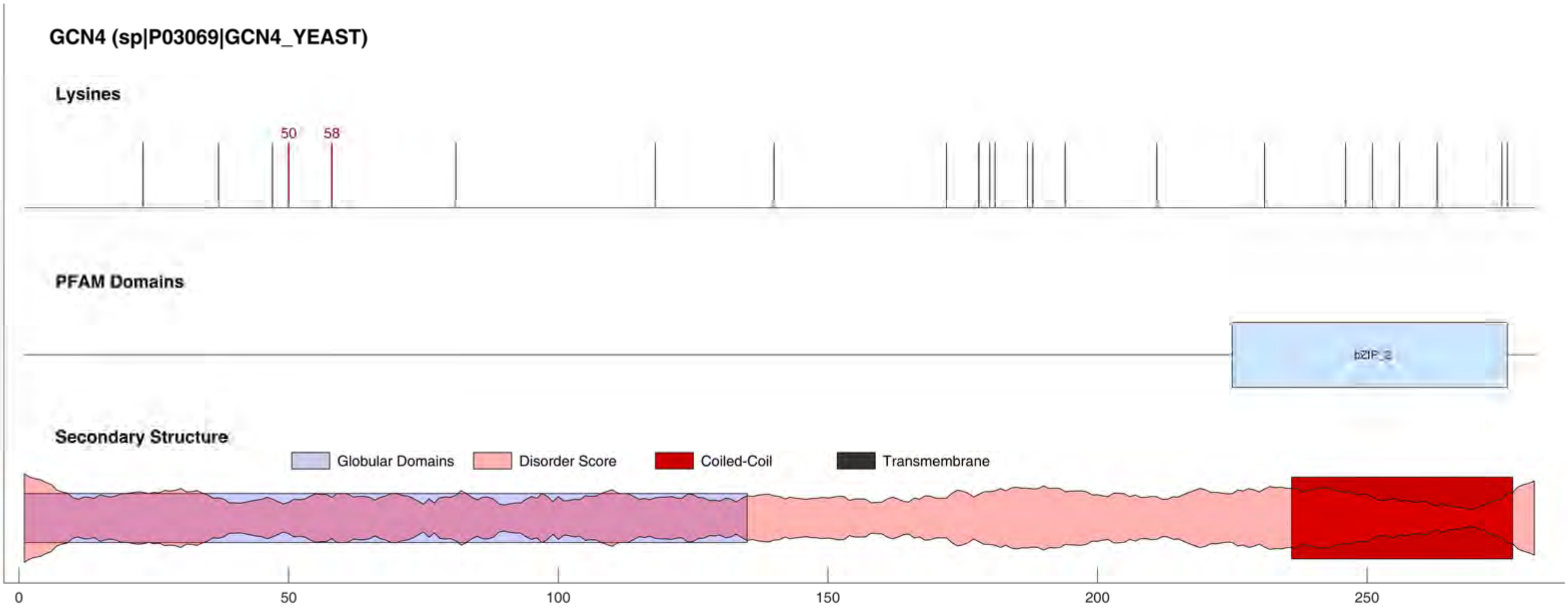
GCN4 (sp|P03069|GCN4_YEAST)
Lysines
50
58
PFAM Domains
bZIP_2
Secondary Structure
Globular Domains
Disorder Score
Coiled-Coil
Transmembrane
0
50
100
150
200
250

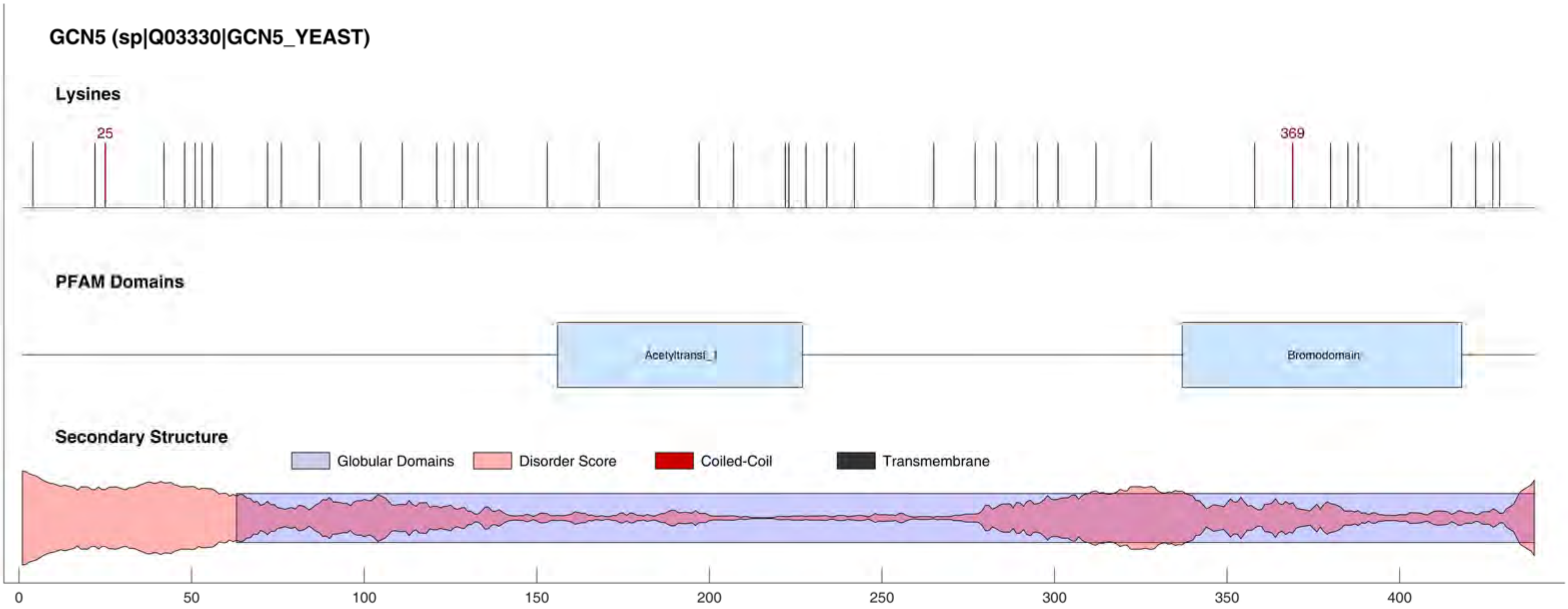
GCN5 (sp|Q03330|GCN5_YEAST)
Lysines
25
369
PFAM Domains
Acetyltransf_1
Bromodomain
Secondary Structure
Globular Domains
Disorder Score
Coiled-Coil
Transmembrane
0
50
100
150
200
250
300
350
400

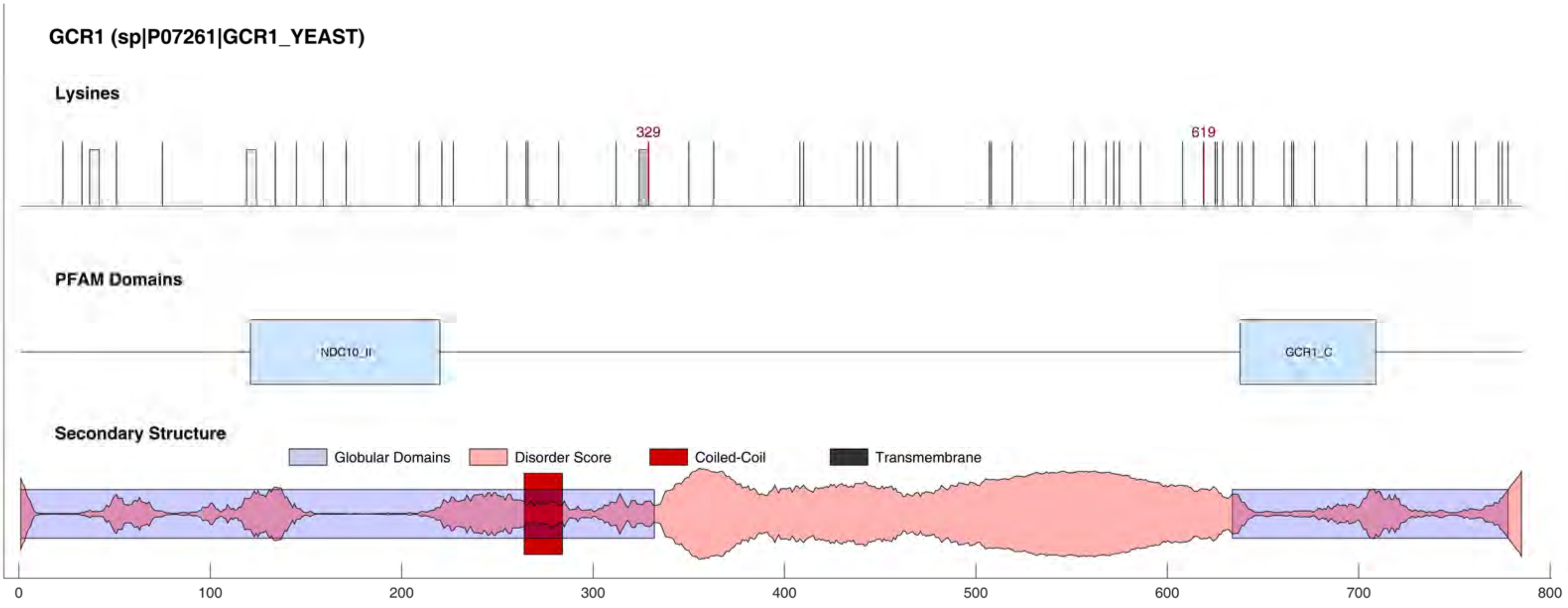

GCR1 (sp|P07261|GCR1_YEAST)
Lysines
329
619
PFAM Domains
NDC10_II
GCR1_C
Secondary Structure
Globular Domains
Disorder Score
Coiled-Coil
Transmembrane
0
100
200
300
400
500
600
700
800

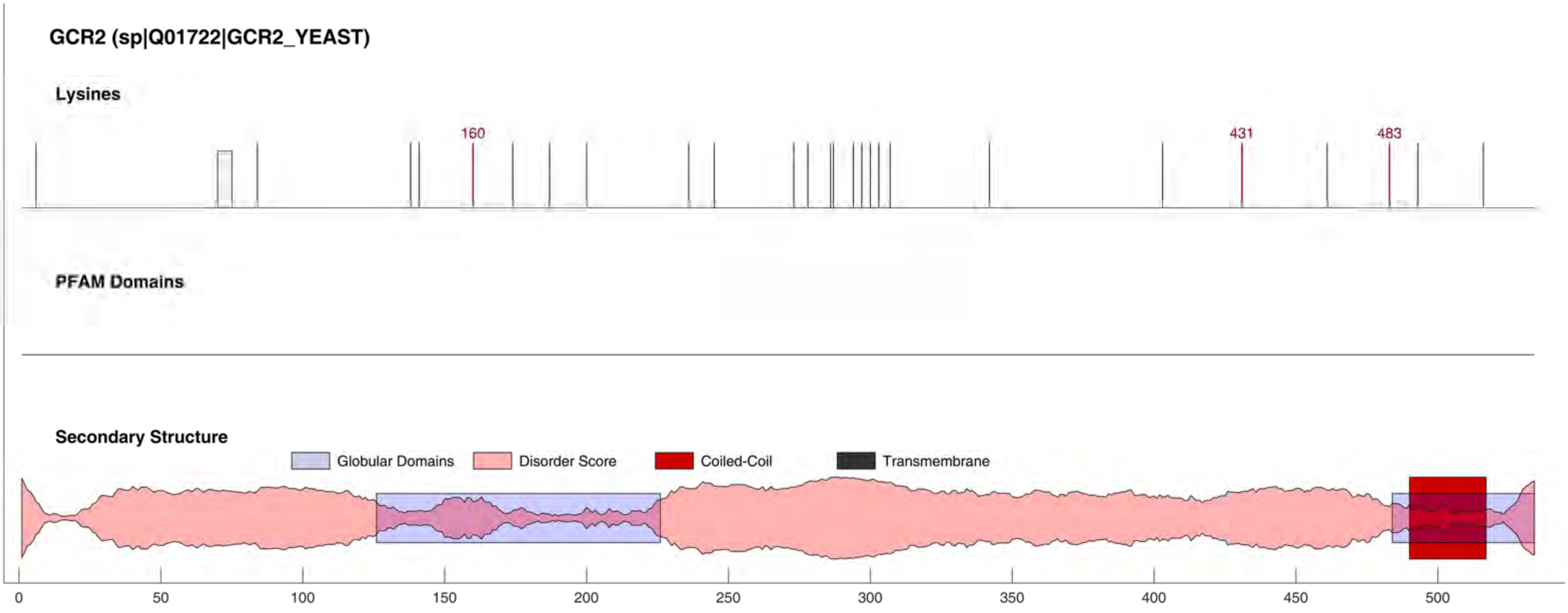
GCR2 (sp|Q01722|GCR2_YEAST)
Lysines
160
431
483
PFAM Domains
Secondary Structure
Globular Domains
Disorder Score
Coiled-Coil
Transmembrane
0
50
100
150
200
250
300
350
400
450
500

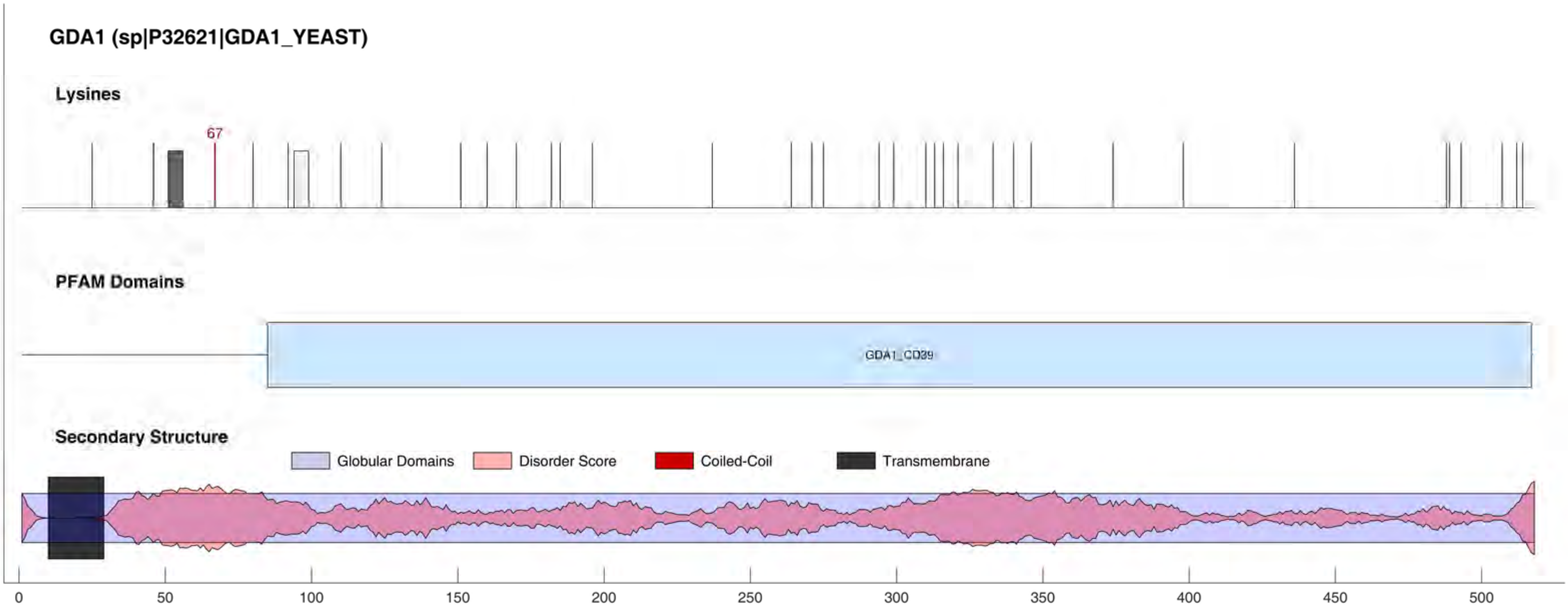
GDA1 (sp|P32621|GDA1_YEAST)
Lysines
67
PFAM Domains
GDA1_CD39
Secondary Structure
Globular Domains
Disorder Score
Coiled-Coil
Transmembrane
0
50
100
150
200
250
300
350
400
450
500

#### GFD1 (sp|Q04839|GFD1_YEAST)

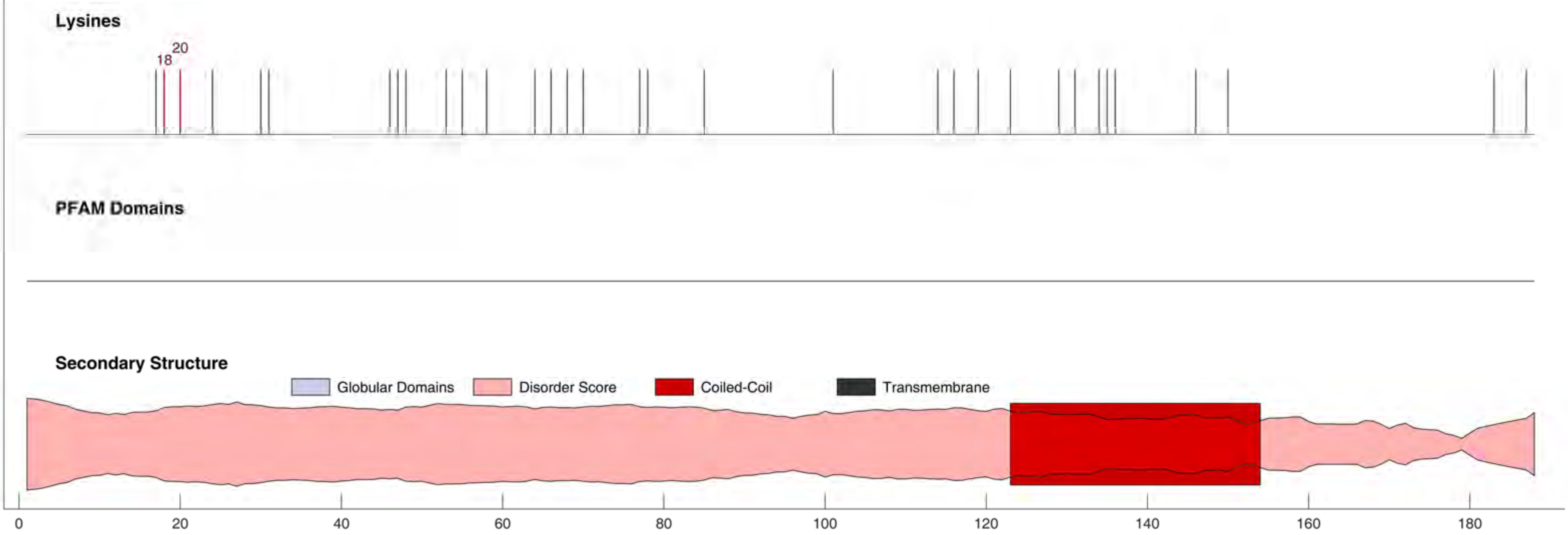

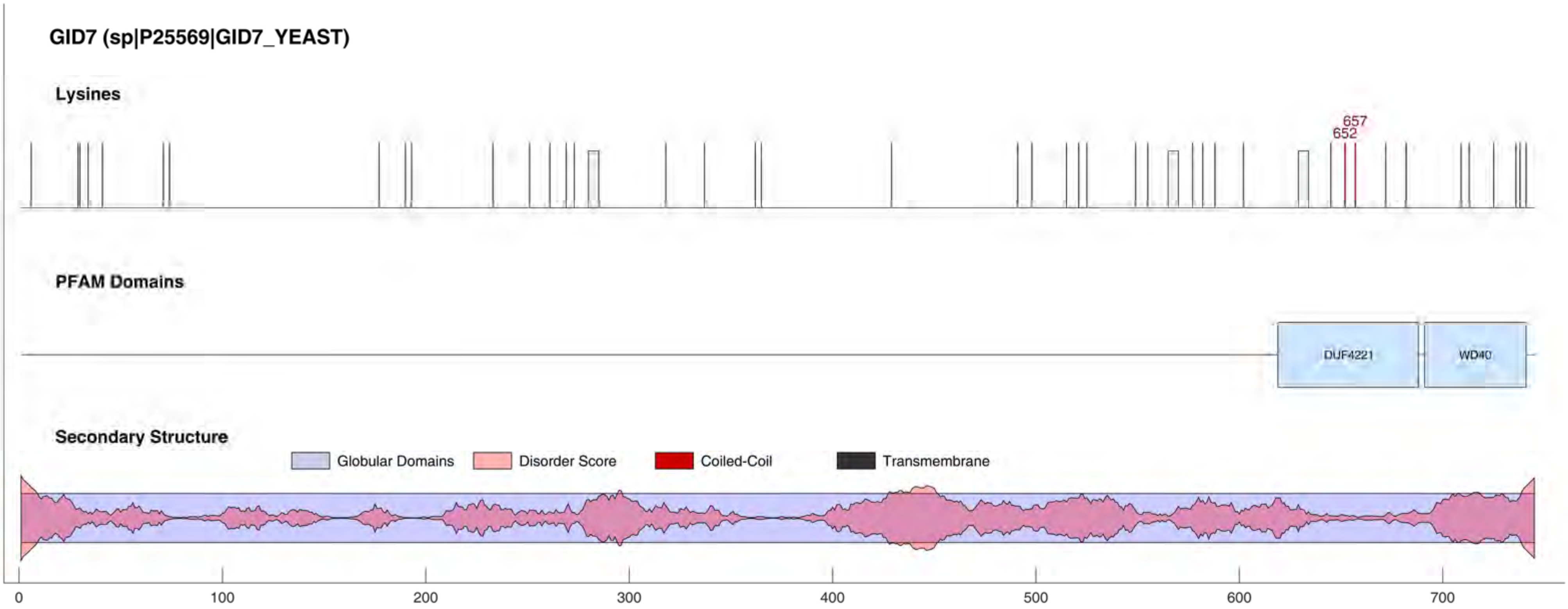
GID7 (sp|P25569|GID7_YEAST)
Lysines
657
652
PFAM Domains
DUF4221
WD40
Secondary Structure
Globular Domains
Disorder Score
Coiled-Coil
Transmembrane
0
100
200
300
400
500
600
700

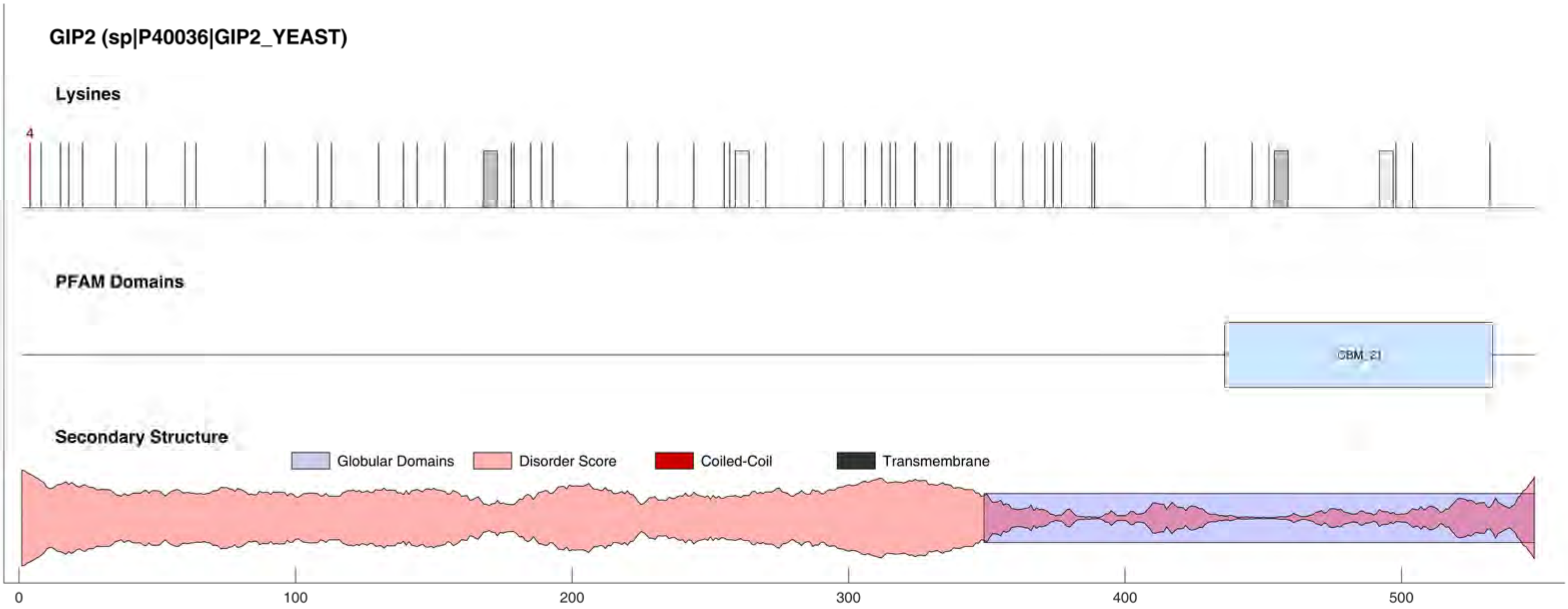
GIP2 (sp|P40036|GIP2_YEAST)
Lysines
4
PFAM Domains
CBM_21
Secondary Structure
Globular Domains
Disorder Score
Coiled-Coil
Transmembrane
0
100
200
300
400
500

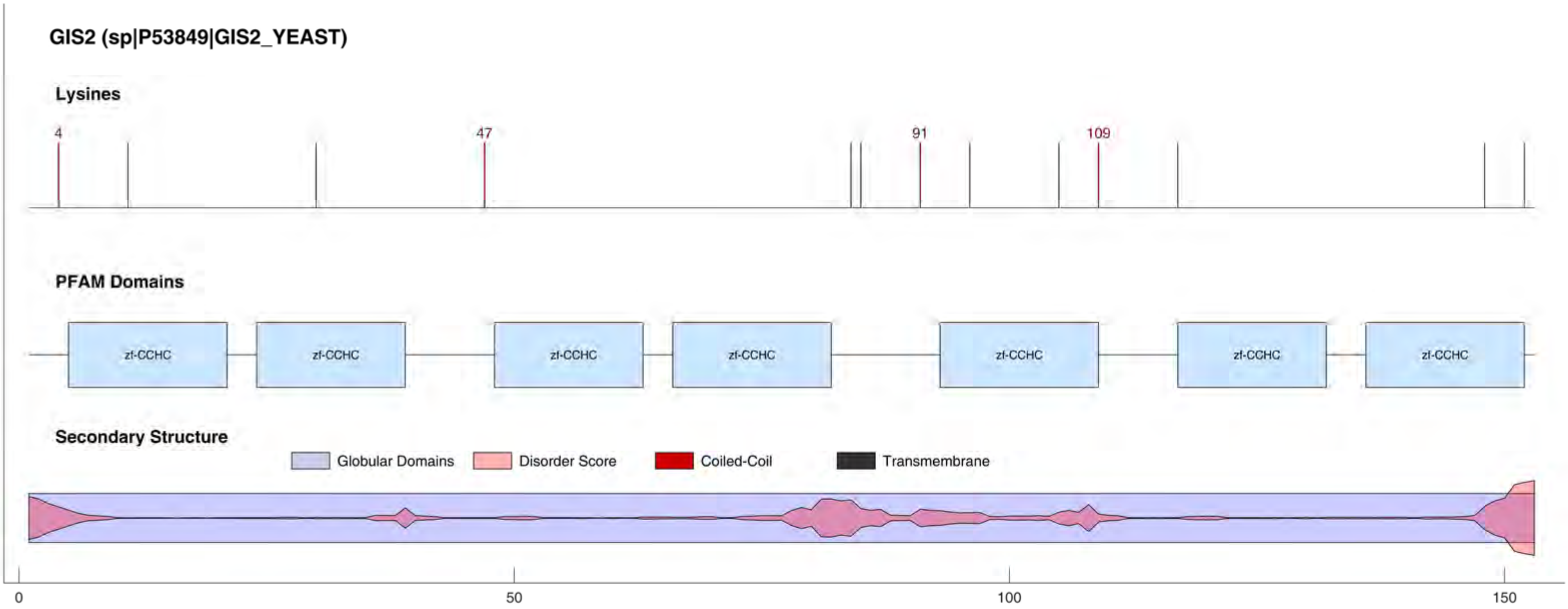

GIS2 (sp|P53849|GIS2_YEAST)
Lysines
4
47
91
109
PFAM Domains
zf-CCHC
zf-CCHC
zf-CCHC
zf-CCHC
zf-CCHC
zf-CCHC
zf-CCHC
Secondary Structure
Globular Domains
Disorder Score
Coiled-Coil
Transmembrane
0
50
100
150

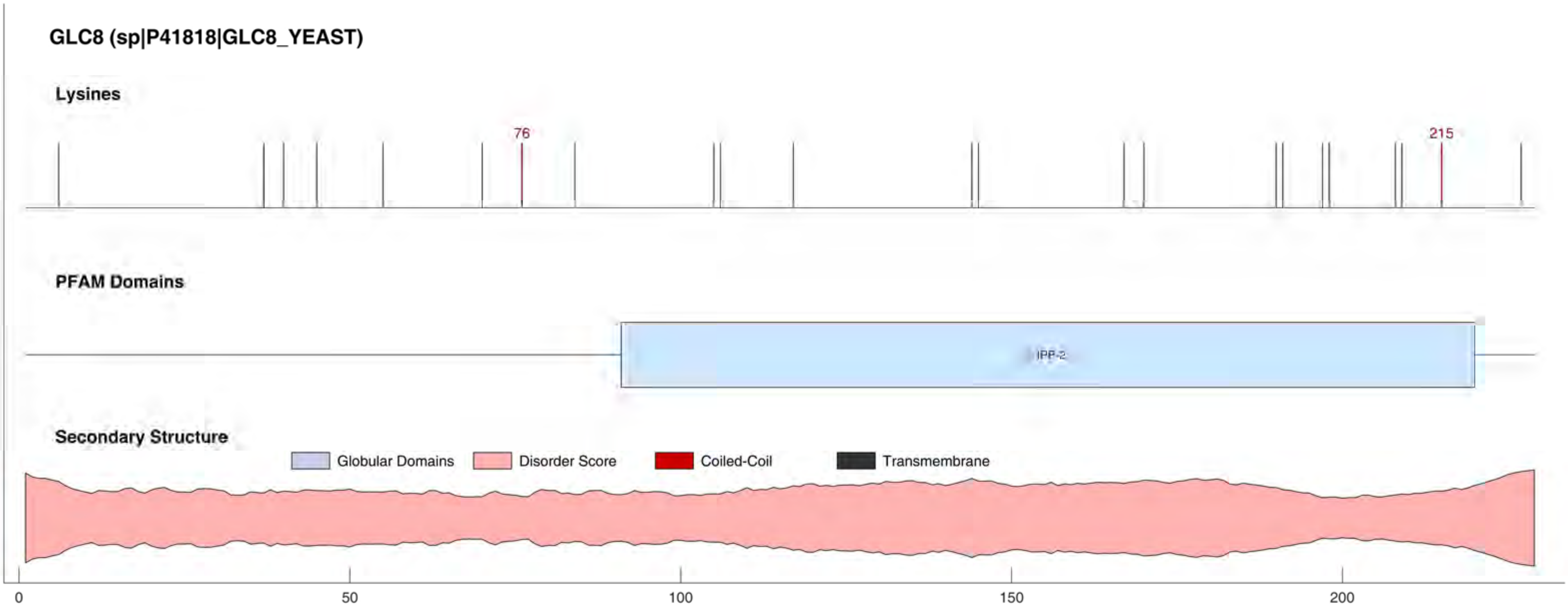

GLC8 (sp|P41818|GLC8_YEAST)
Lysines
76
215
PFAM Domains
IPP-2
Secondary Structure
Globular Domains
Disorder Score
Coiled-Coil
Transmembrane
0
50
100
150
200

**GLPK (sp|P32190|GLPK_YEAST)**

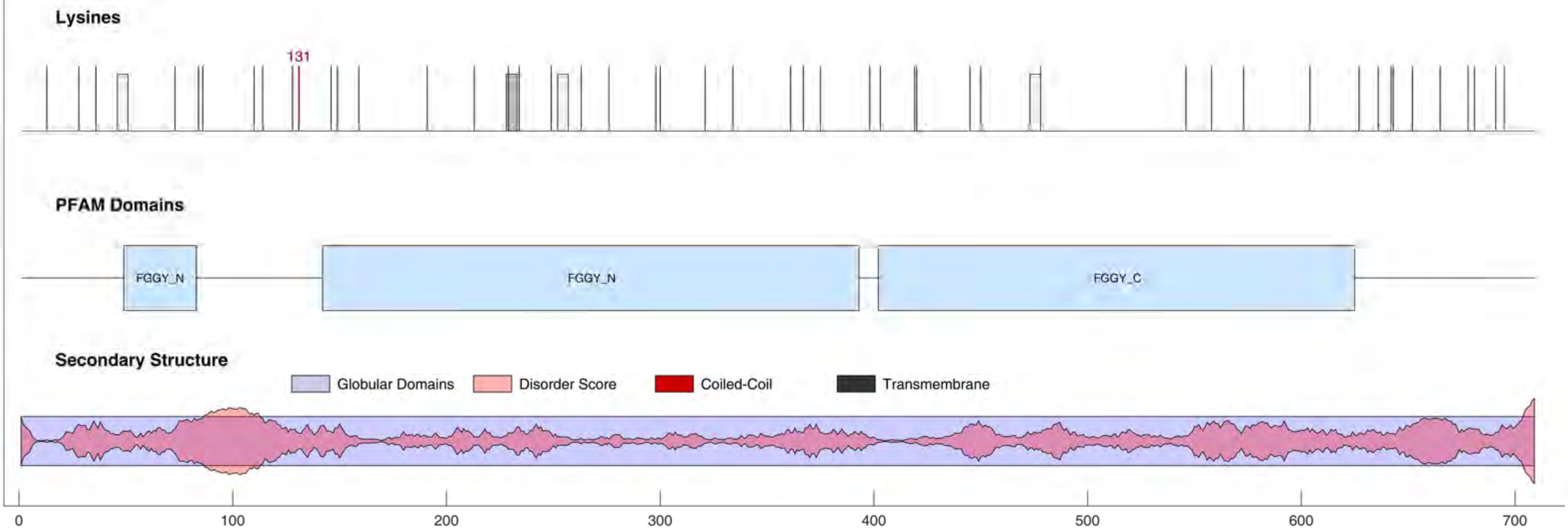

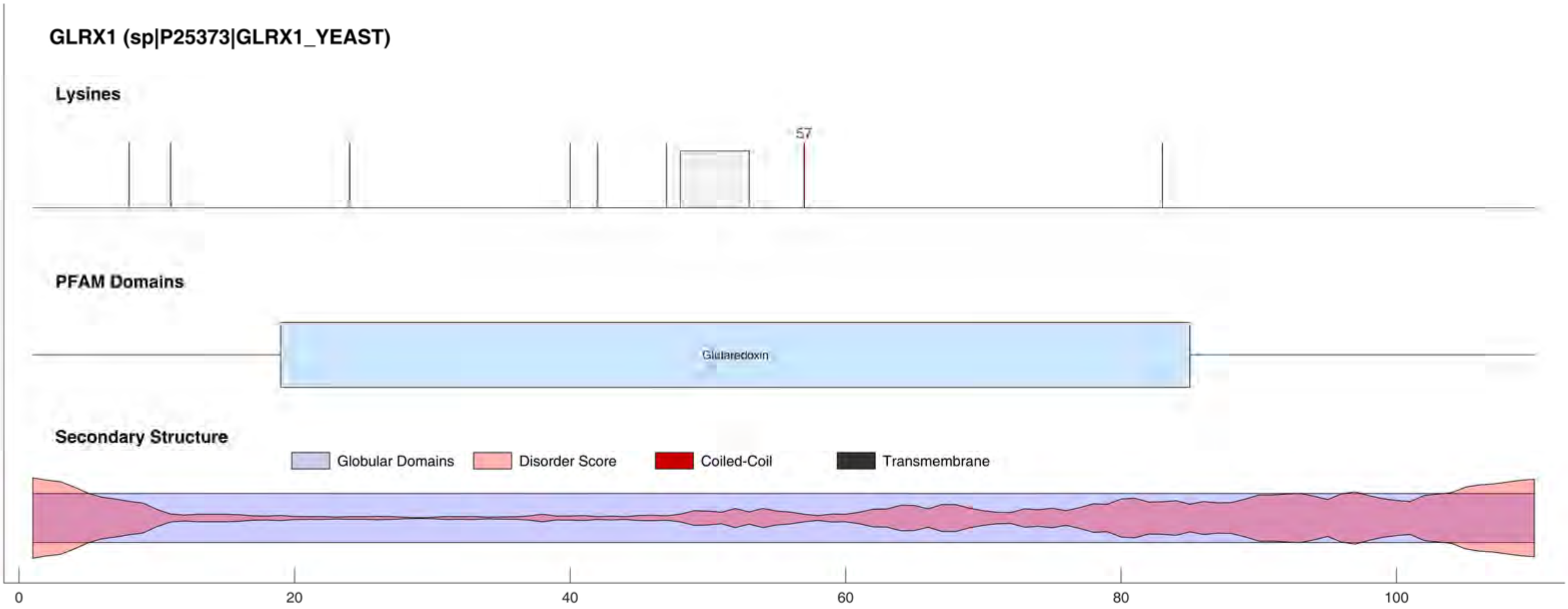
GLRX1 (sp|P25373|GLRX1_YEAST)
Lysines
57
PFAM Domains
Glutaredoxin
Secondary Structure
Globular Domains
Disorder Score
Coiled-Coil
Transmembrane
0
20
40
60
80
100

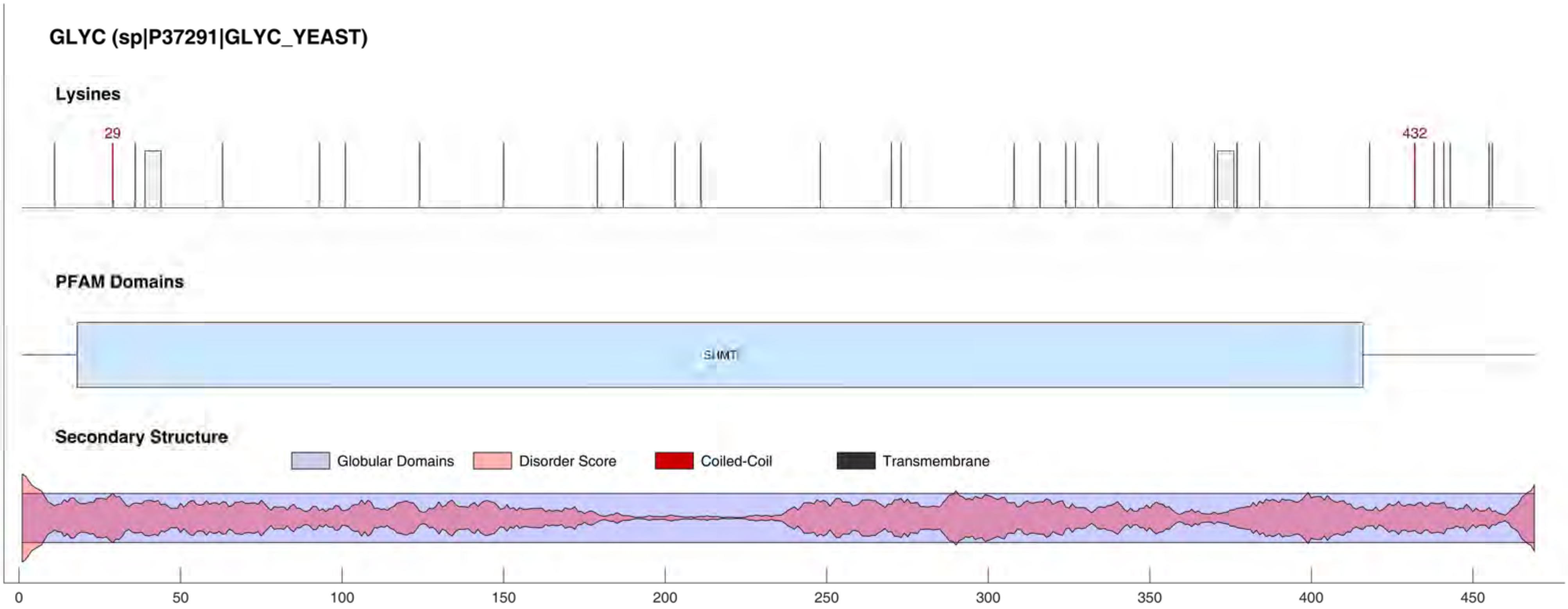
GLYC (sp|P37291|GLYC_YEAST)
Lysines
29
432
PFAM Domains
SHMT
Secondary Structure
Globular Domains
Disorder Score
Coiled-Coil
Transmembrane
0
50
100
150
200
250
300
350
400
450

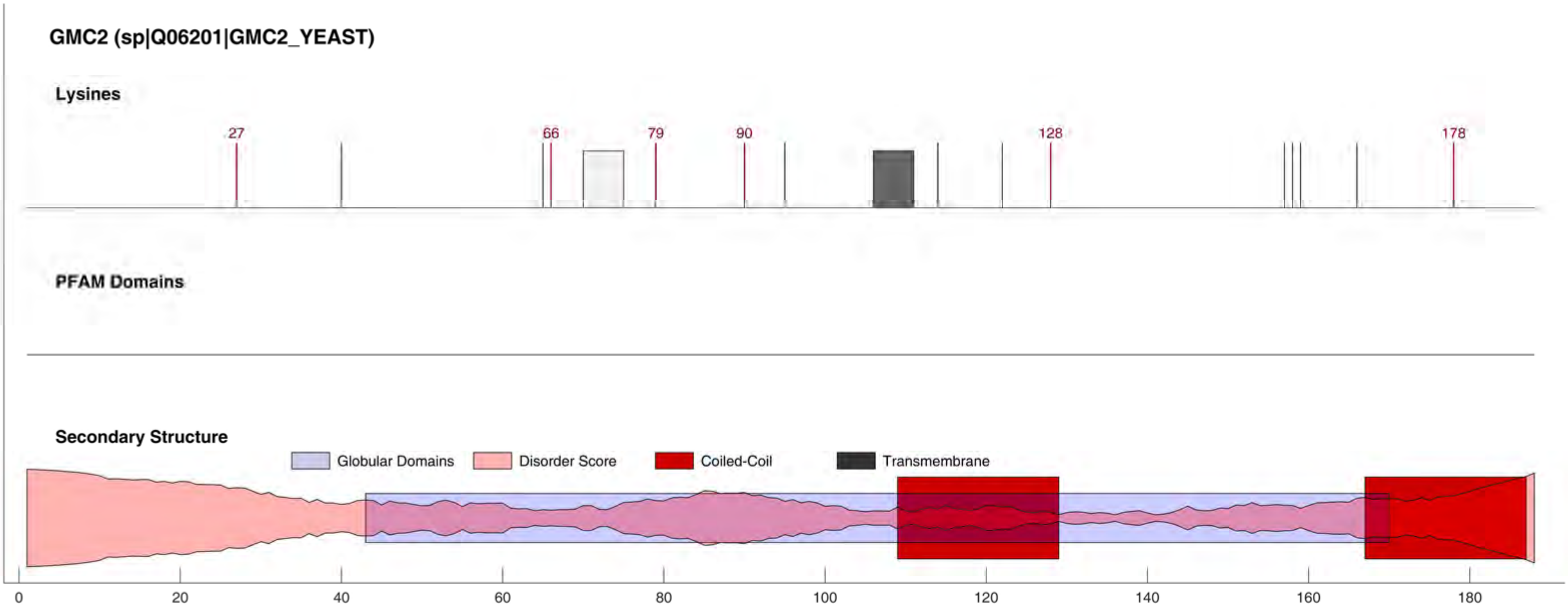

GMC2 (sp|Q06201|GMC2_YEAST)
Lysines
27
66
79
90
128
178
PFAM Domains
Secondary Structure
Globular Domains
Disorder Score
Coiled-Coil
Transmembrane
0
20
40
60
80
100
120
140
160
180

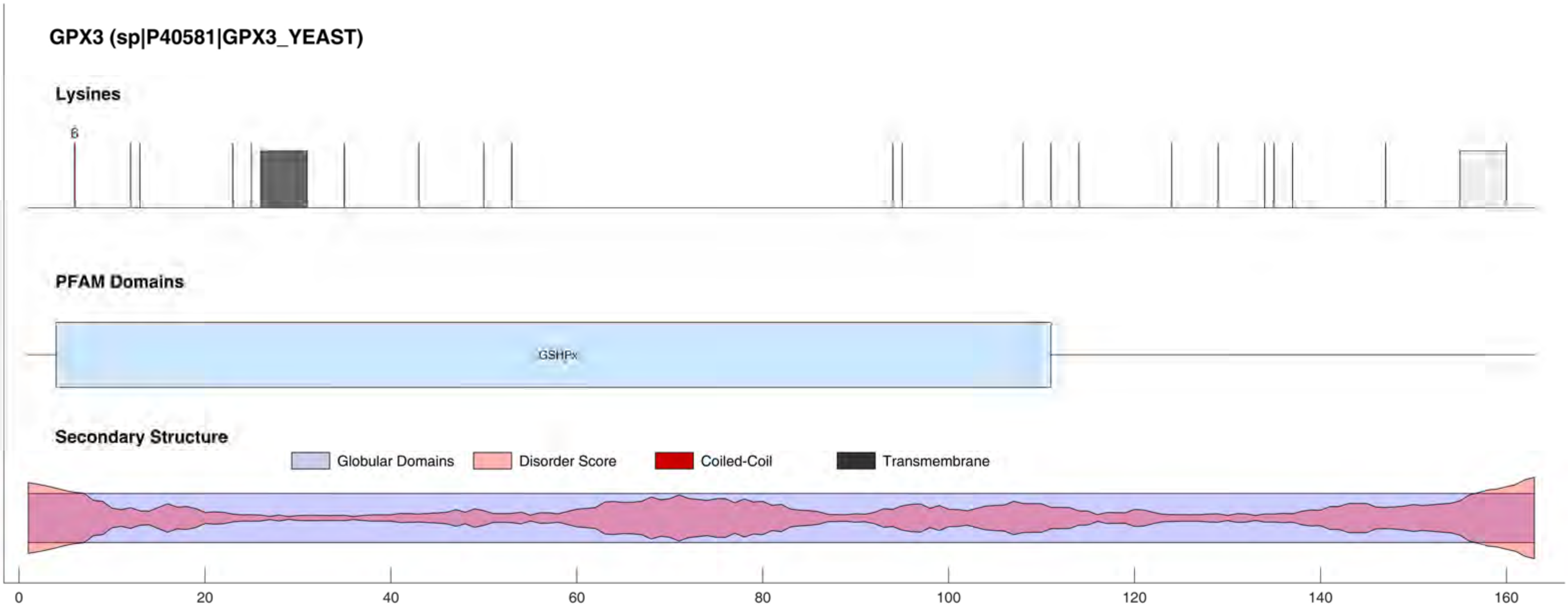

GPX3 (sp|P40581|GPX3_YEAST)
Lysines
6
PFAM Domains
GSHPx
Secondary Structure
Globular Domains
Disorder Score
Coiled-Coil
Transmembrane
0
20
40
60
80
100
120
140
160

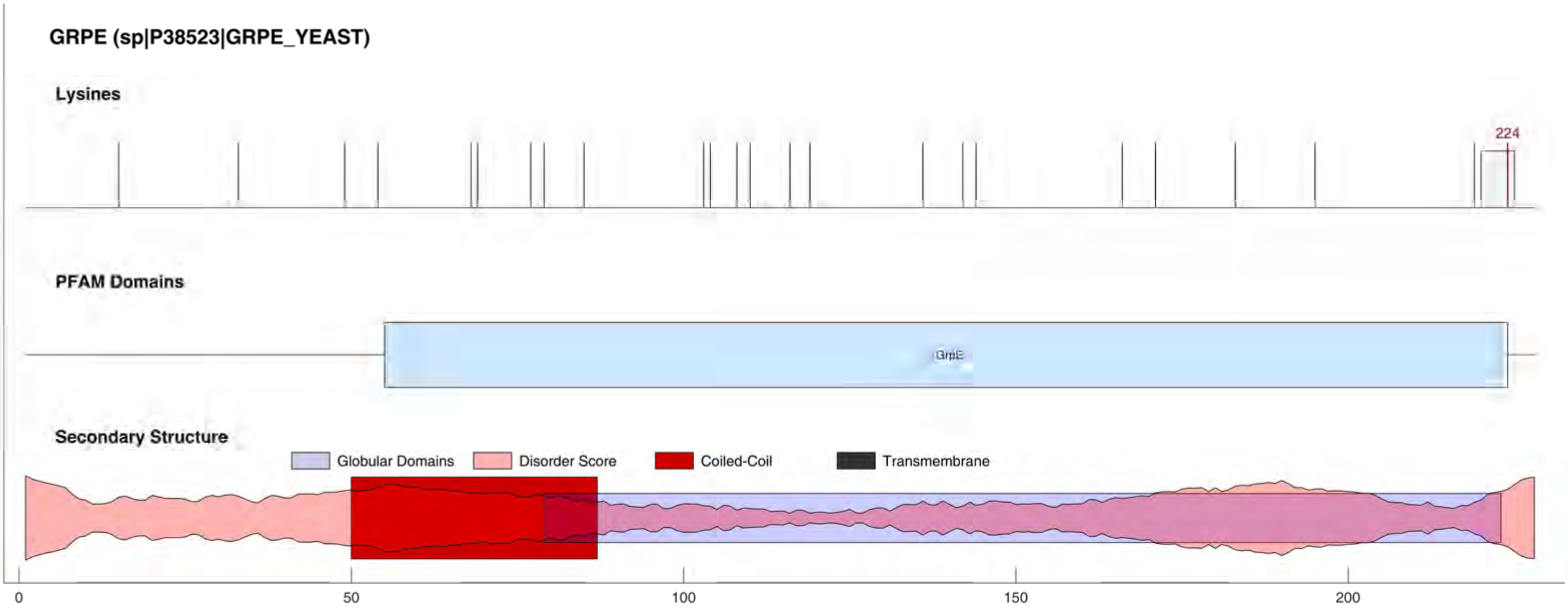

GRPE (sp|P38523|GRPE_YEAST)
Lysines
224
PFAM Domains
GrpE
Secondary Structure
Globular Domains
Disorder Score
Coiled-Coil
Transmembrane
0
50
100
150
200

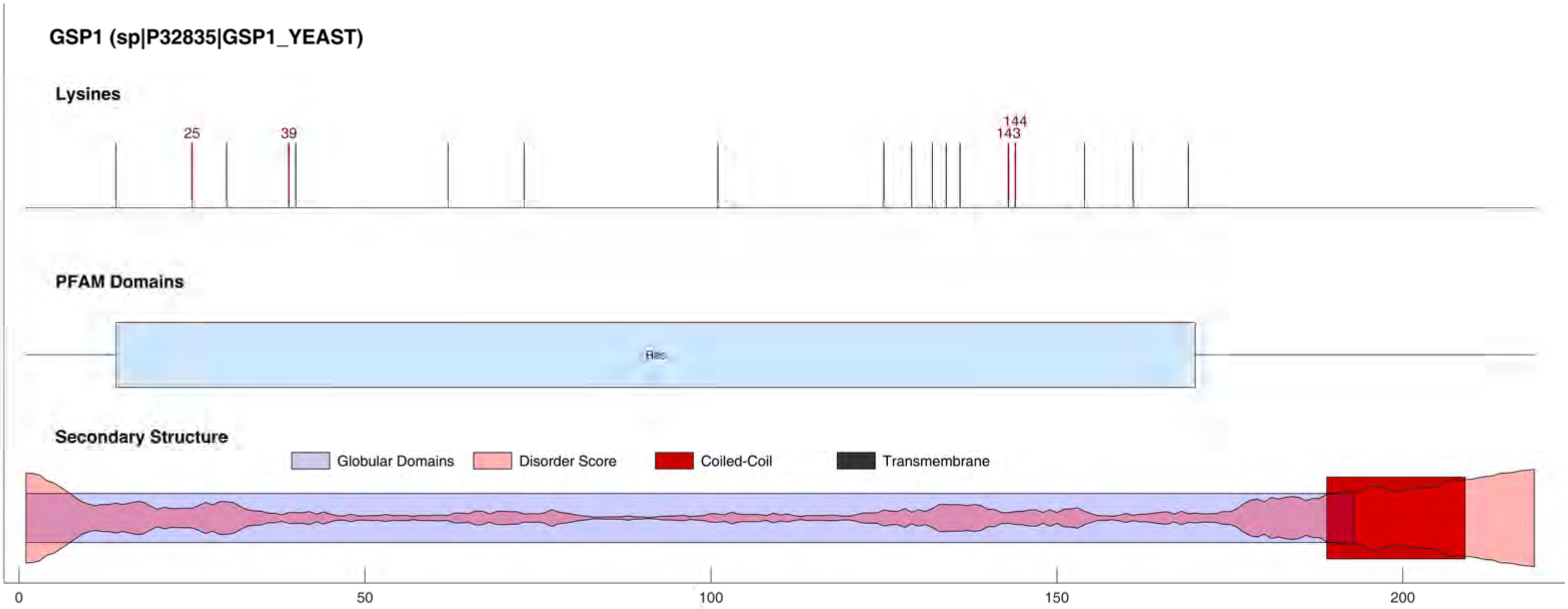
GSP1 (sp|P32835|GSP1_YEAST)
Lysines
25
39
144
143
PFAM Domains
Ras
Secondary Structure
Globular Domains
Disorder Score
Coiled-Coil
Transmembrane
0
50
100
150
200

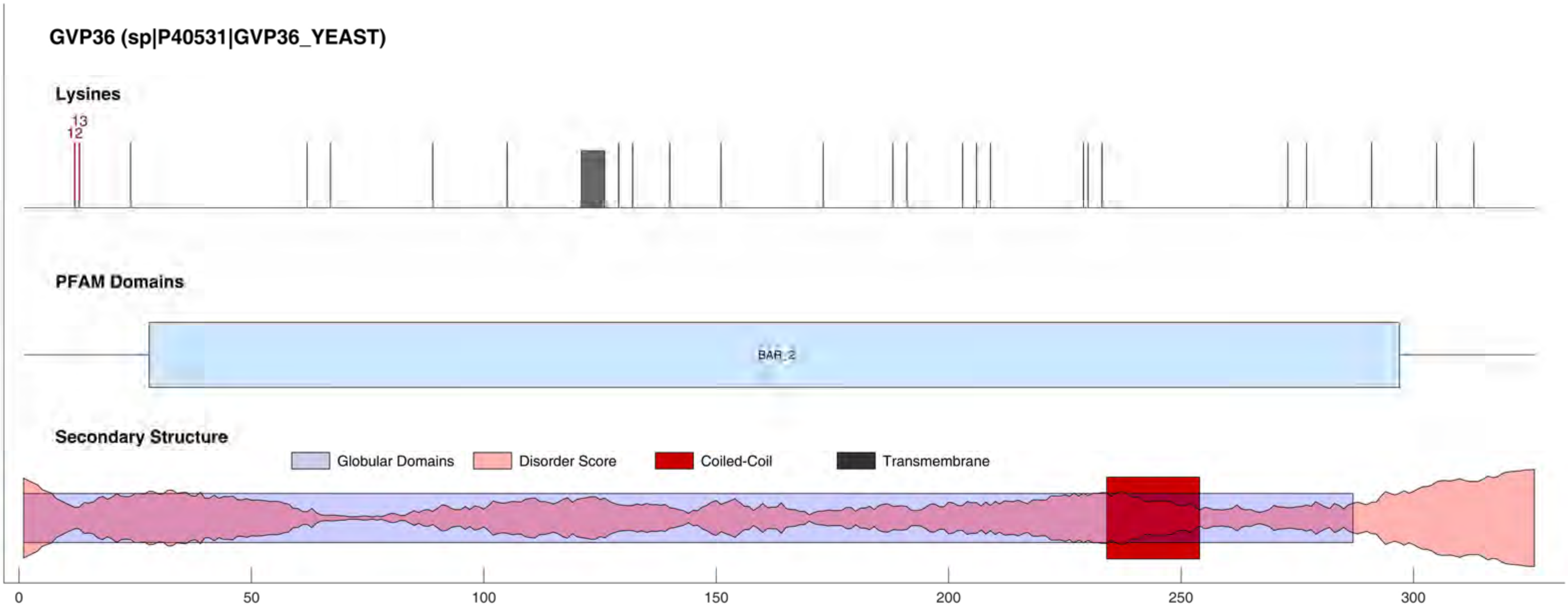

GVP36 (sp|P40531|GVP36_YEAST)
Lysines
13
12
PFAM Domains
BAR_2
Secondary Structure
Globular Domains
Disorder Score
Coiled-Coil
Transmembrane
0
50
100
150
200
250
300

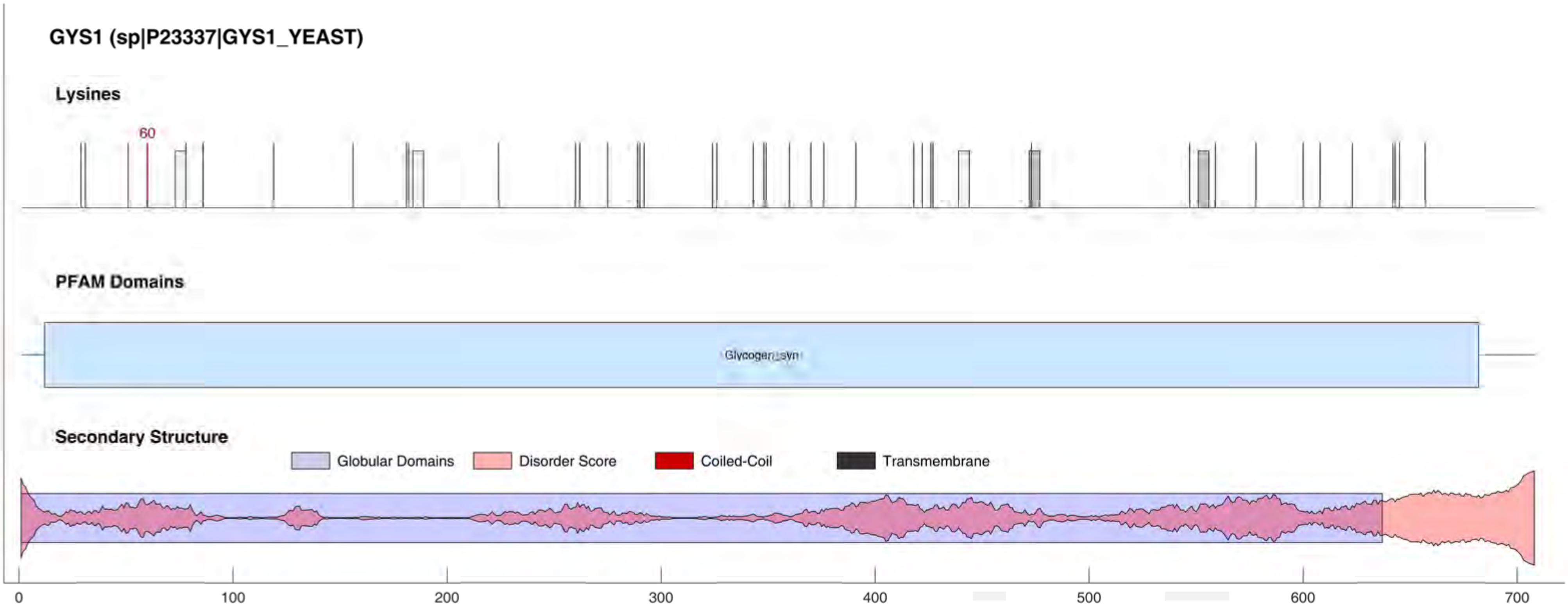
GYS1 (sp|P23337|GYS1_YEAST)
Lysines
60
PFAM Domains
Glycogen_syn
Secondary Structure
Globular Domains
Disorder Score
Coiled-Coil
Transmembrane
0
100
200
300
400
500
600
700

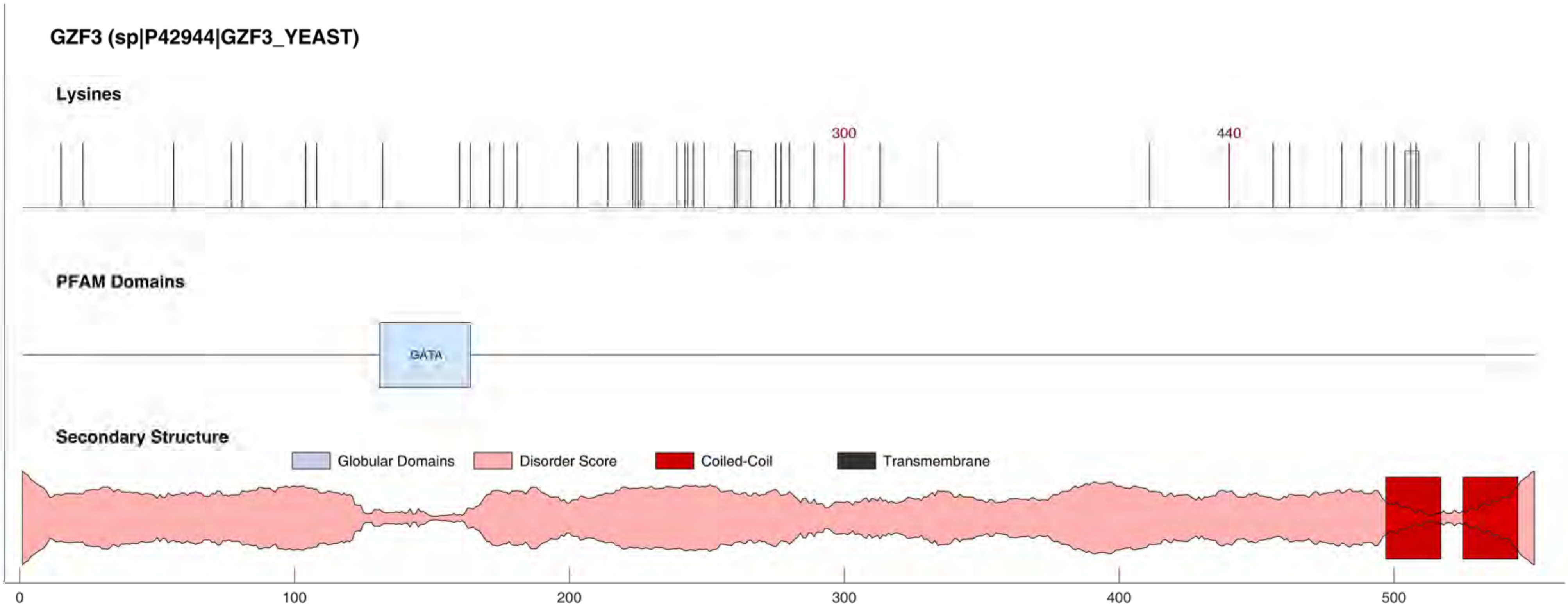

GZF3 (sp|P42944|GZF3_YEAST)
Lysines
300
440
PFAM Domains
GATA
Secondary Structure
Globular Domains
Disorder Score
Coiled-Coil
Transmembrane
0
100
200
300
400
500

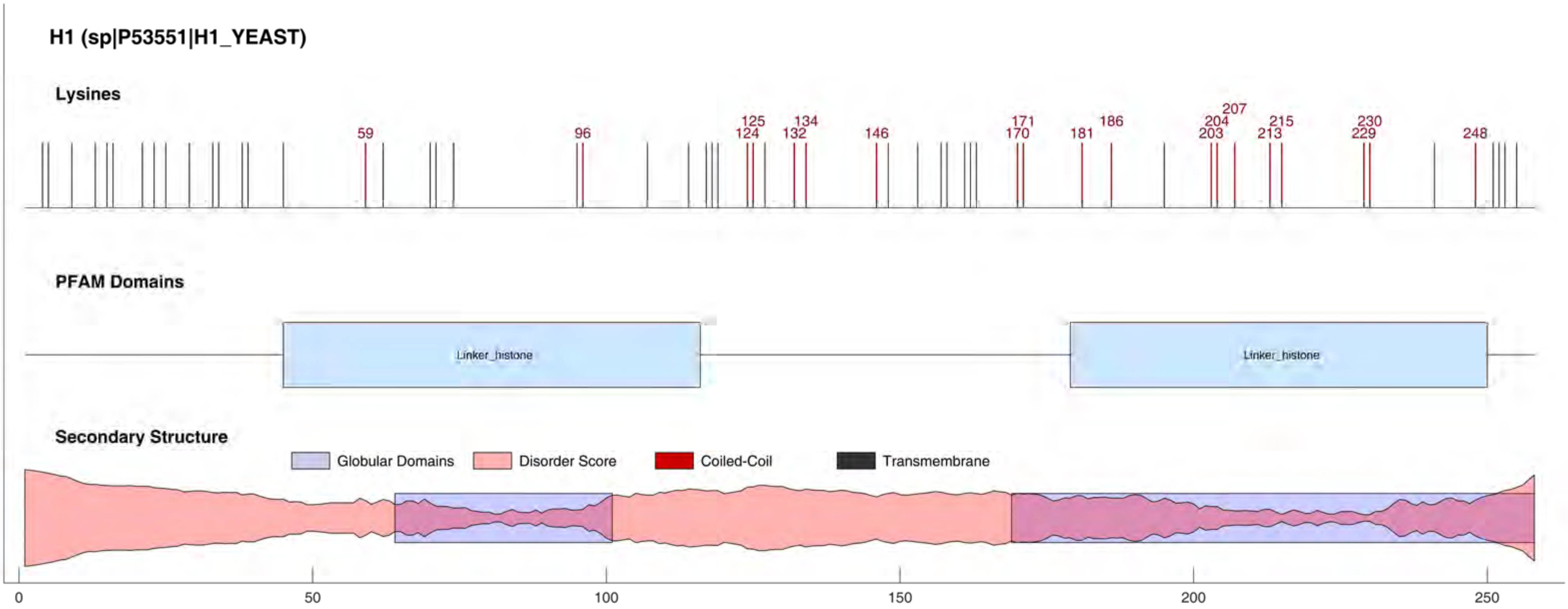
H1 (sp|P53551|H1_YEAST)
Lysines
59
96
125
124
134
132
146
171
170
181
186
207
204
203
215
213
230
229
248
PFAM Domains
Linker_histone
Linker_histone
Secondary Structure
Globular Domains
Disorder Score
Coiled-Coil
Transmembrane
0
50
100
150
200
250

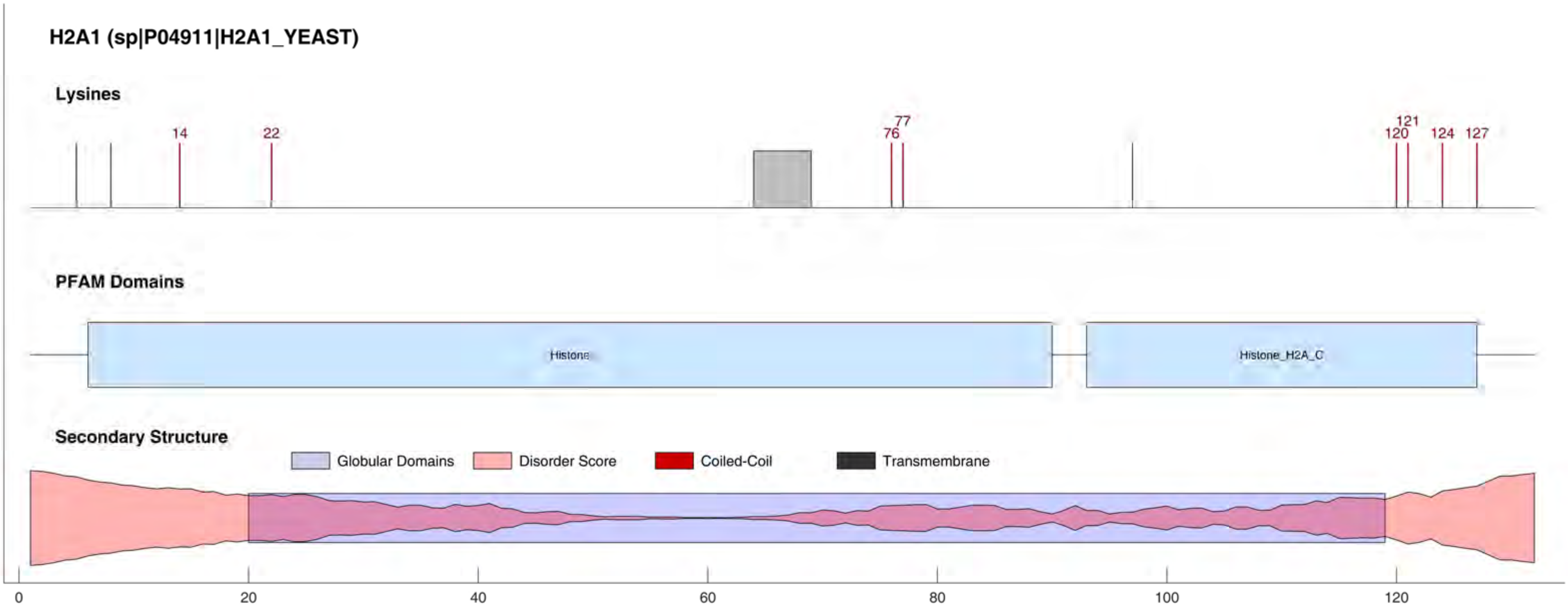
H2A1 (sp|P04911|H2A1_YEAST)
Lysines
14
22
77
76
121
120
124
127
PFAM Domains
Histone
Histone_H2A_C
Secondary Structure
Globular Domains
Disorder Score
Coiled-Coil
Transmembrane
0
20
40
60
80
100
120

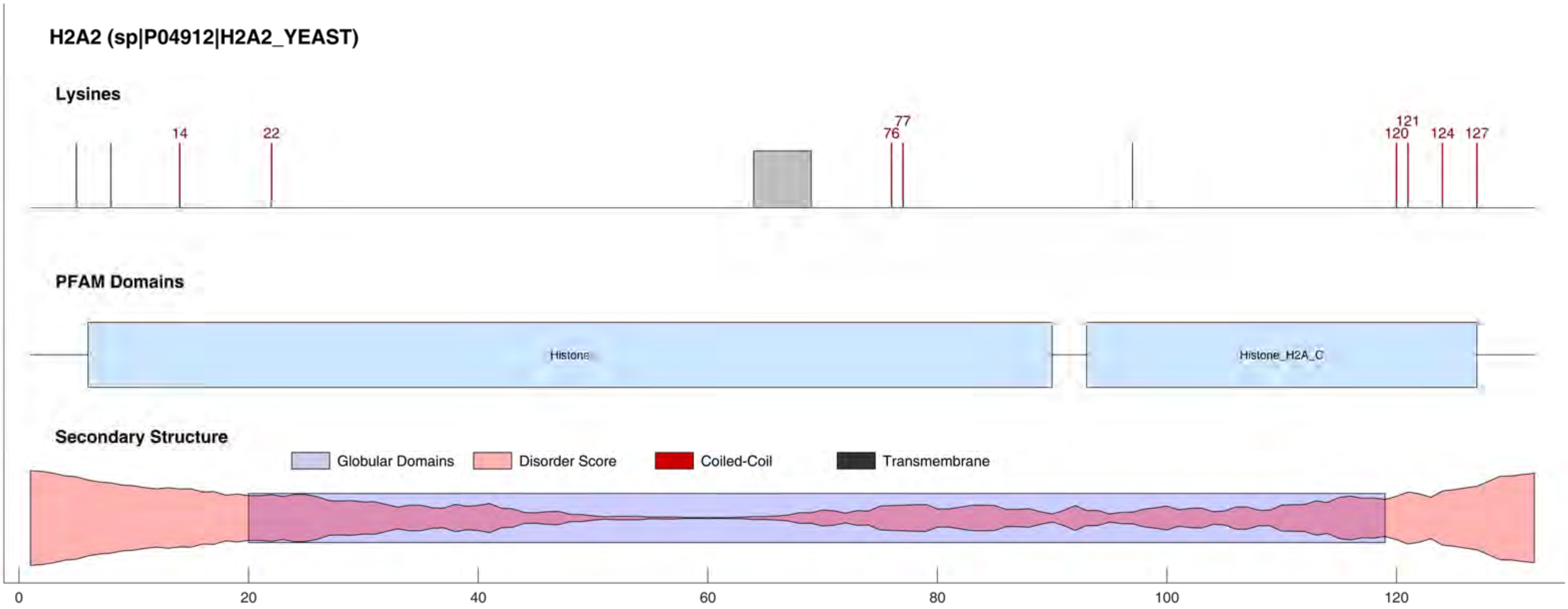
H2A2 (sp|P04912|H2A2_YEAST)
Lysines
14
22
77
76
121
120
124
127
PFAM Domains
Histone
Histone_H2A_C
Secondary Structure
Globular Domains
Disorder Score
Coiled-Coil
Transmembrane
0
20
40
60
80
100
120

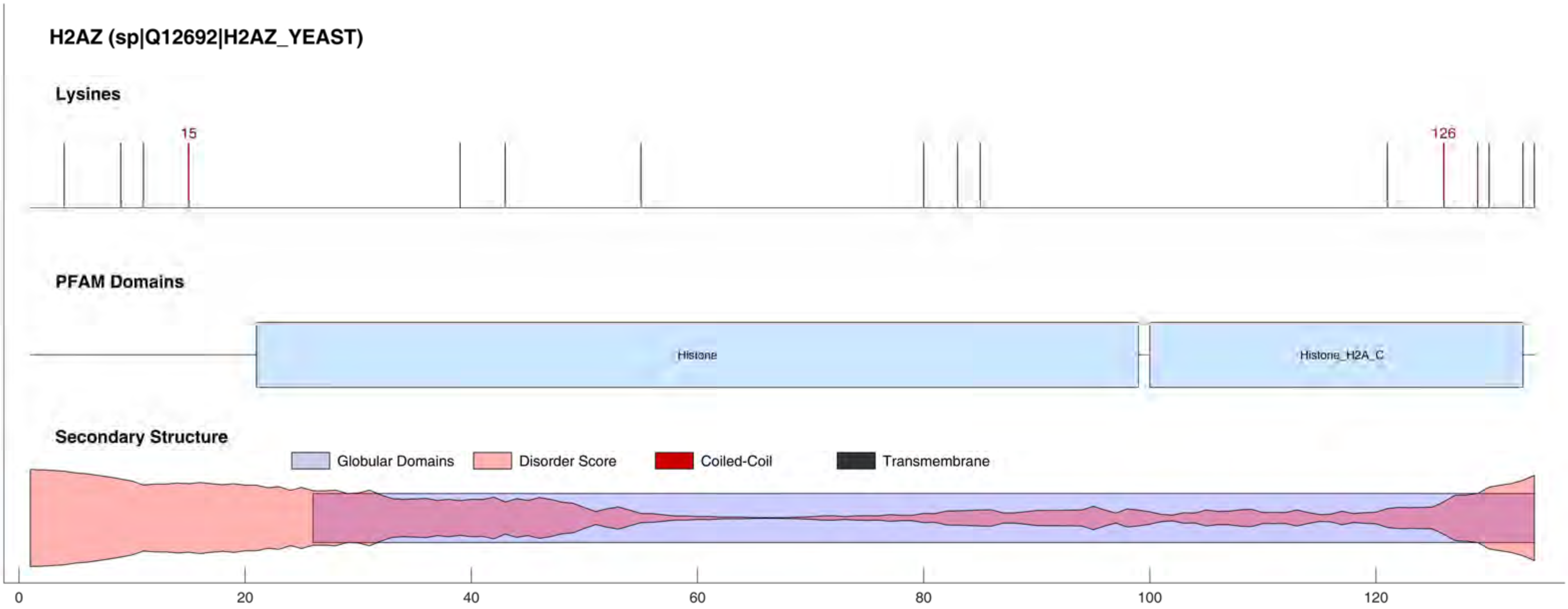

H2AZ (sp|Q12692|H2AZ_YEAST)
Lysines
15
126
PFAM Domains
Histone
Histone_H2A_C
Secondary Structure
Globular Domains
Disorder Score
Coiled-Coil
Transmembrane
0
20
40
60
80
100
120

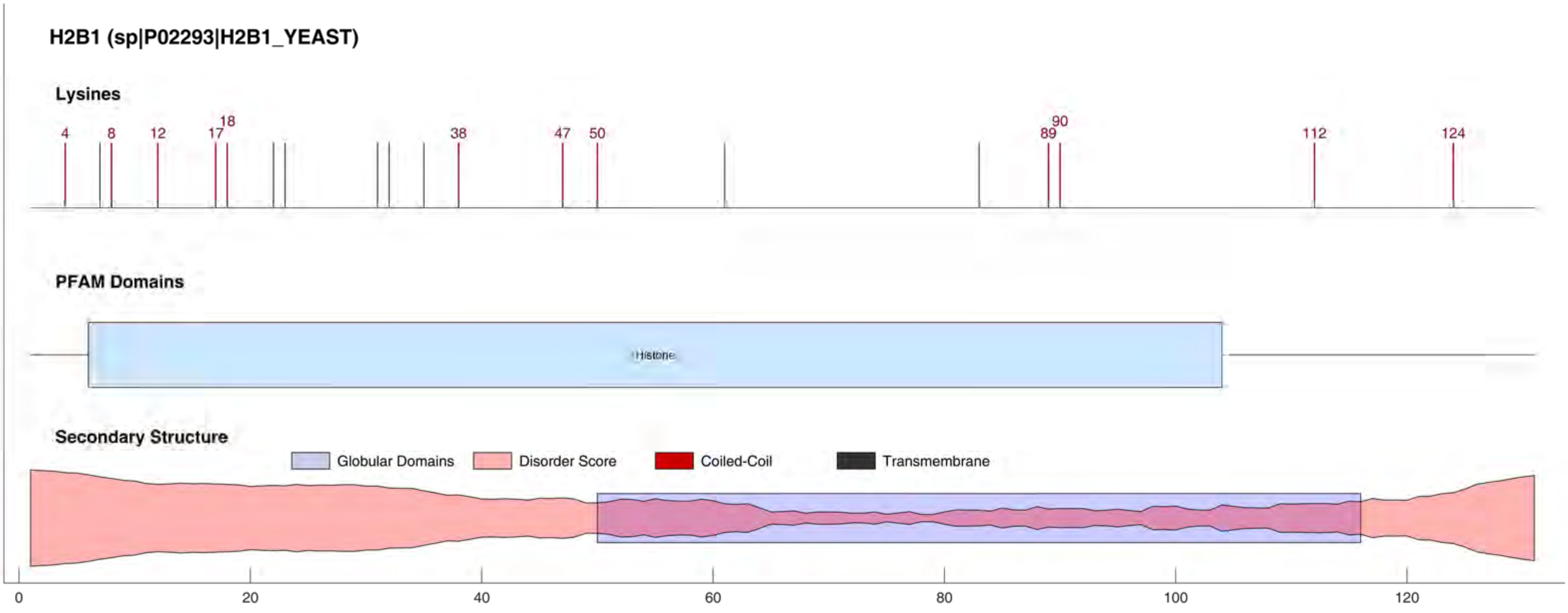
H2B1 (sp|P02293|H2B1_YEAST)
Lysines
4
8
12
17
18
38
47
50
89
90
112
124
PFAM Domains
Histone
Secondary Structure
Globular Domains
Disorder Score
Coiled-Coil
Transmembrane
0
20
40
60
80
100
120

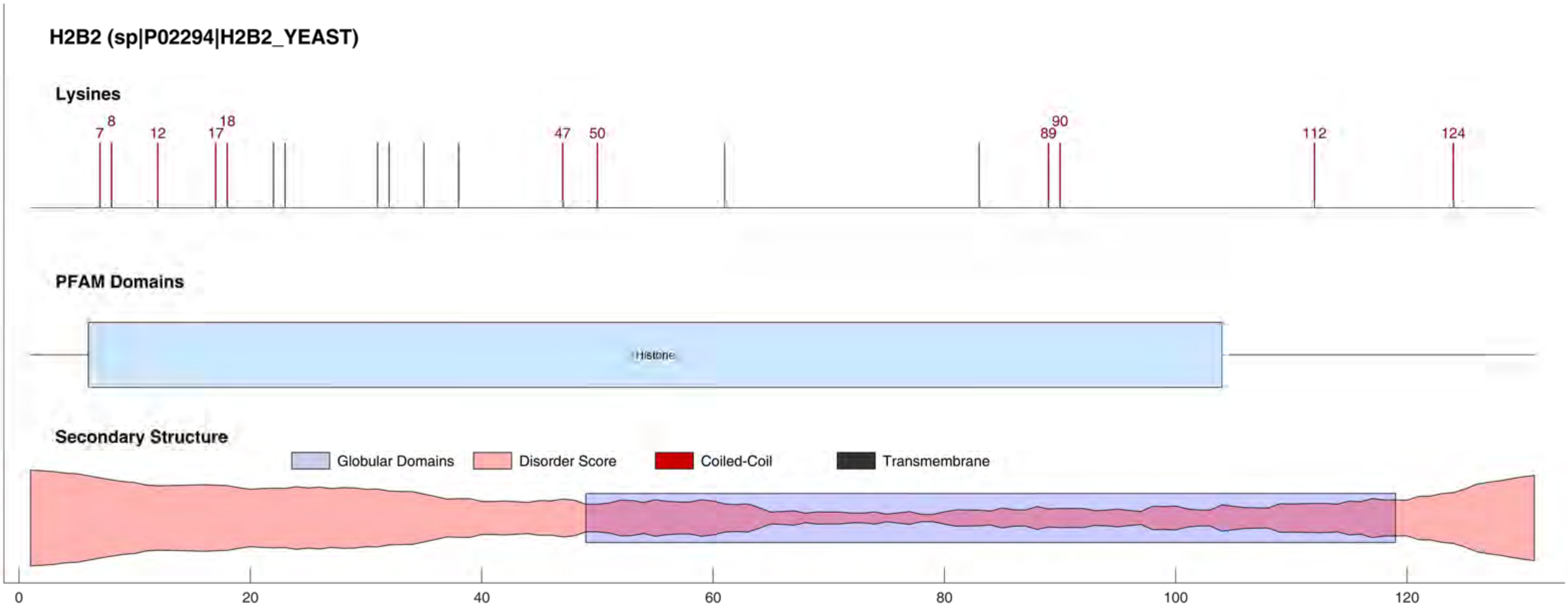
H2B2 (sp|P02294|H2B2_YEAST)
Lysines
7
8
12
17
18
47
50
89
90
112
124
PFAM Domains
Histone
Secondary Structure
Globular Domains
Disorder Score
Coiled-Coil
Transmembrane
0
20
40
60
80
100
120

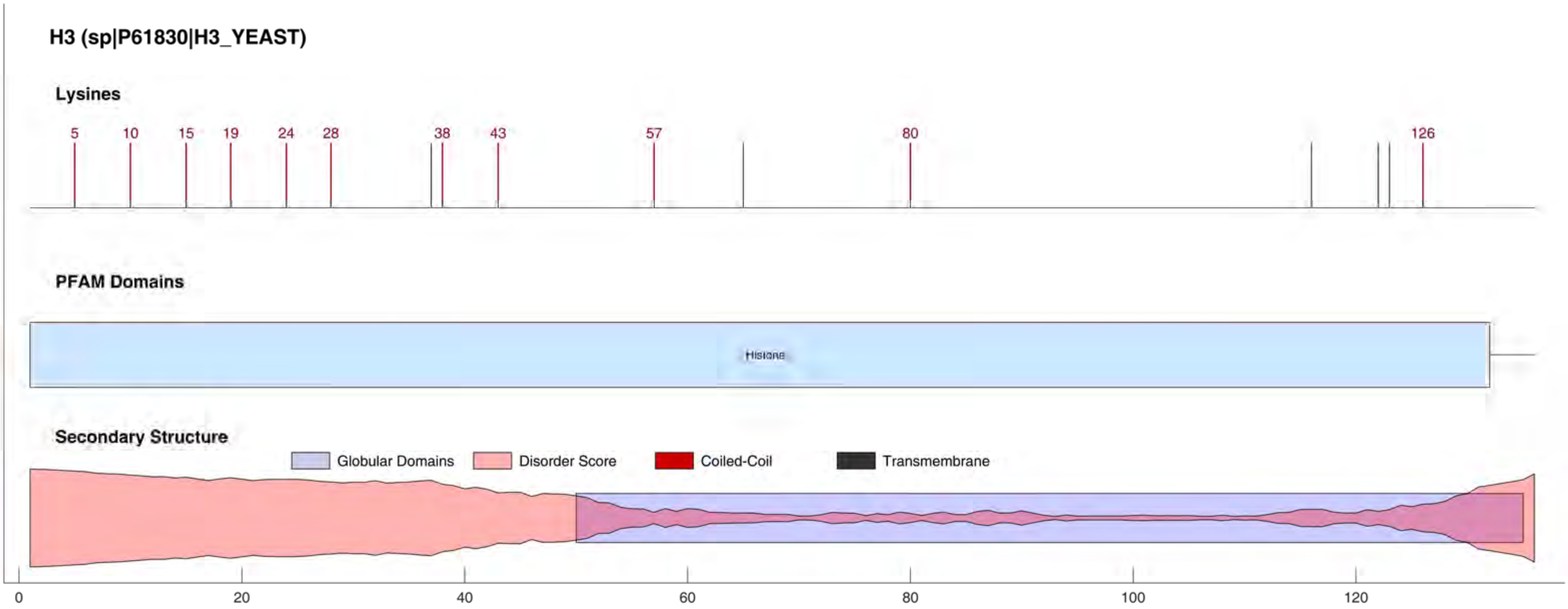
H3 (sp|P61830|H3_YEAST)
Lysines
5
10
15
19
24
28
38
43
57
80
126
PFAM Domains
Histone
Secondary Structure
Globular Domains
Disorder Score
Coiled-Coil
Transmembrane
0
20
40
60
80
100
120

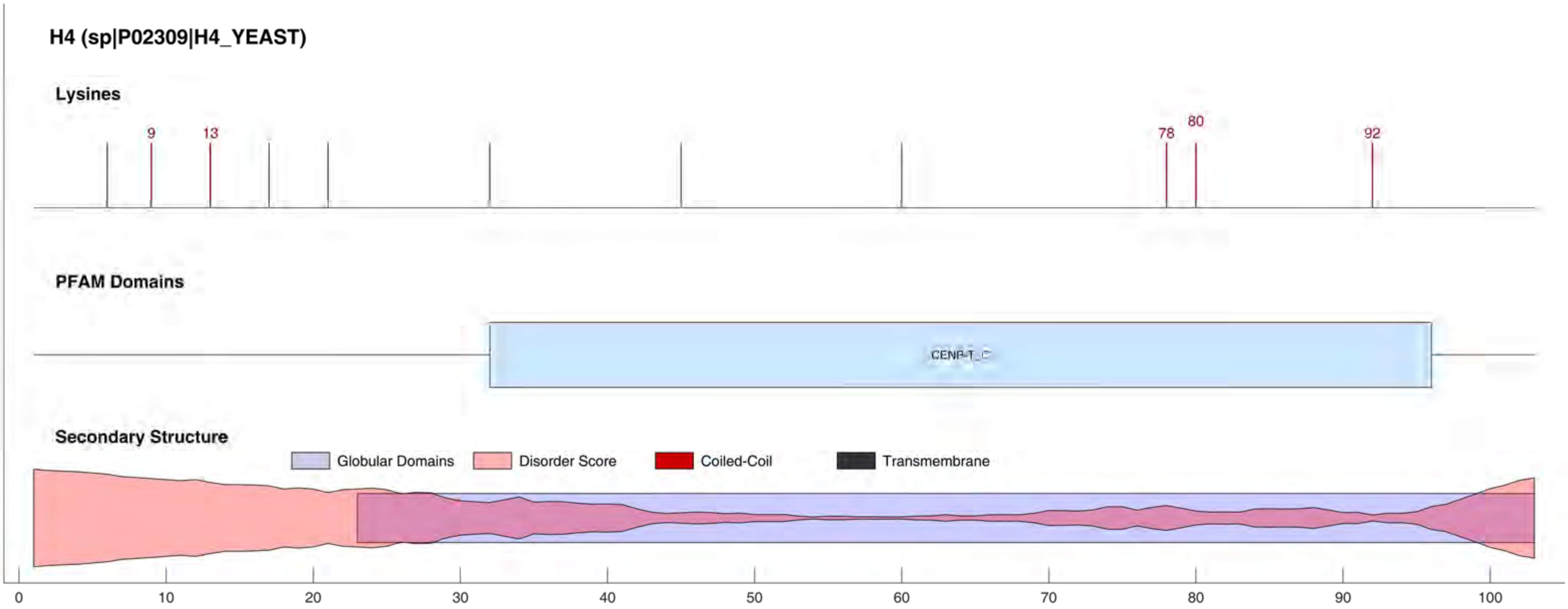

H4 (sp|P02309|H4_YEAST)
Lysines
9
13
78
80
92
PFAM Domains
CENP-T_C
Secondary Structure
Globular Domains
Disorder Score
Coiled-Coil
Transmembrane
0
10
20
30
40
50
60
70
80
90
100

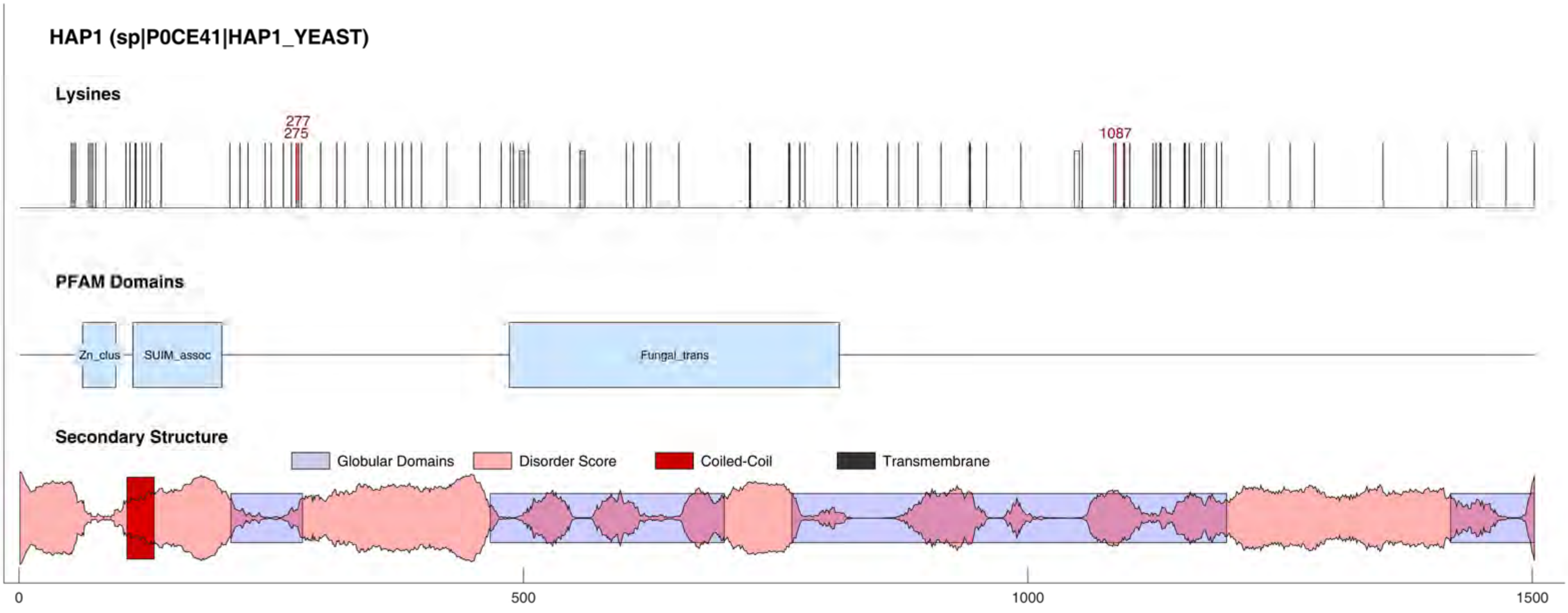
HAP1 (sp|P0CE41|HAP1_YEAST)
Lysines
277
275
1087
PFAM Domains
Zn_clus
SUIM_assoc
Fungal_trans
Secondary Structure
Globular Domains
Disorder Score
Coiled-Coil
Transmembrane
0
500
1000
1500

#### HAP4 (sp|P14064|HAP4_YEAST)

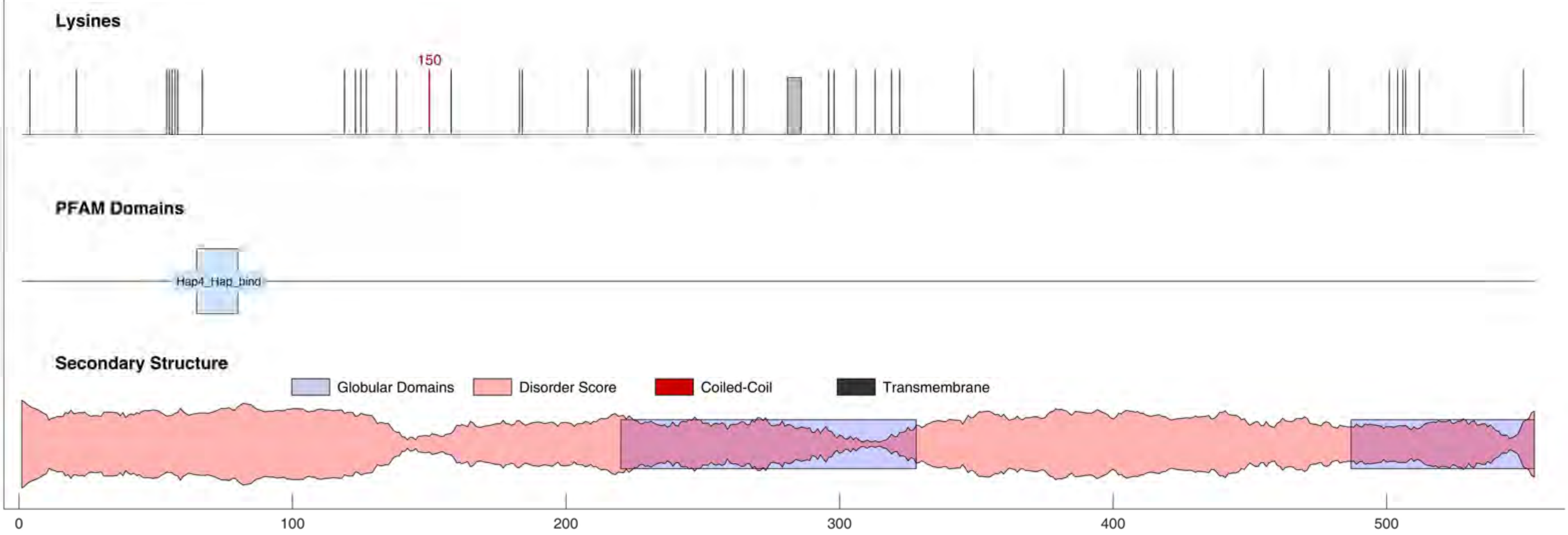

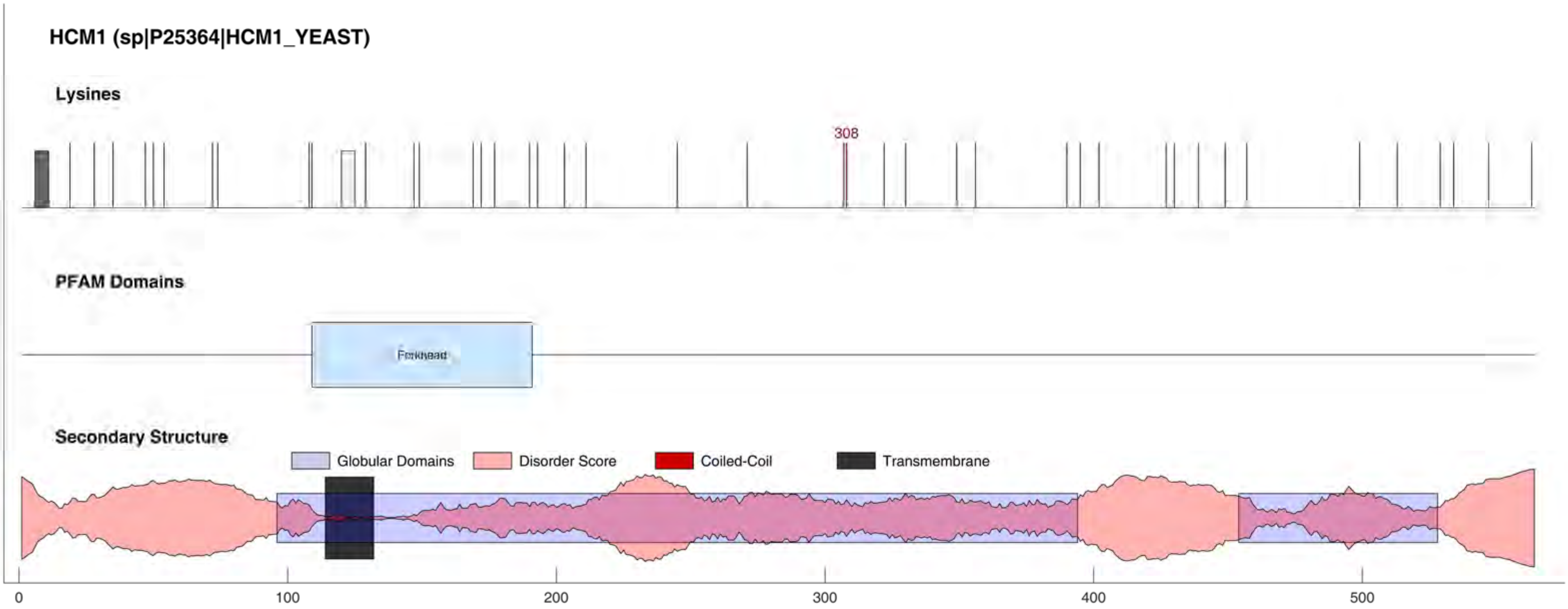

HCM1 (sp|P25364|HCM1_YEAST)
Lysines
308
PFAM Domains
Forkhead
Secondary Structure
Globular Domains
Disorder Score
Coiled-Coil
Transmembrane
0
100
200
300
400
500

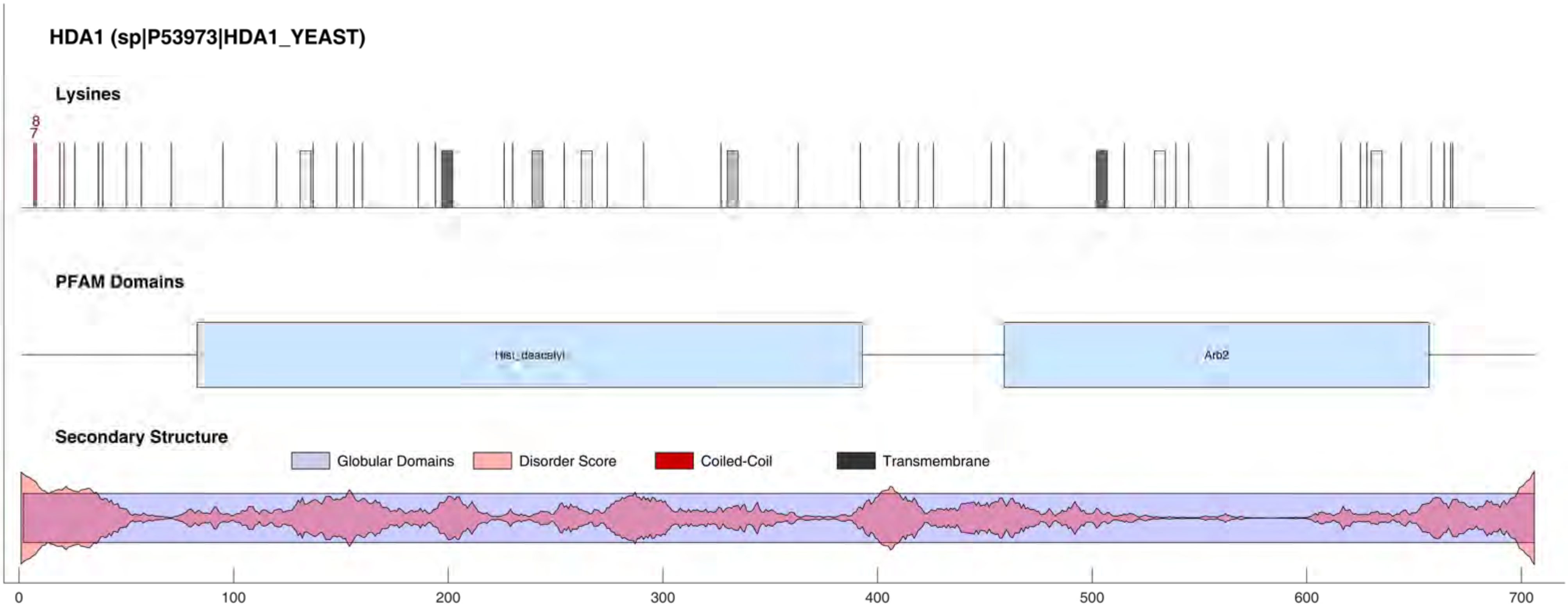
HDA1 (sp|P53973|HDA1_YEAST)
Lysines
8
7
PFAM Domains
Hist_deacetyl
Arb2
Secondary Structure
Globular Domains
Disorder Score
Coiled-Coil
Transmembrane
0
100
200
300
400
500
600
700

HDA2 (sp|Q06629|HDA2_YEAST)

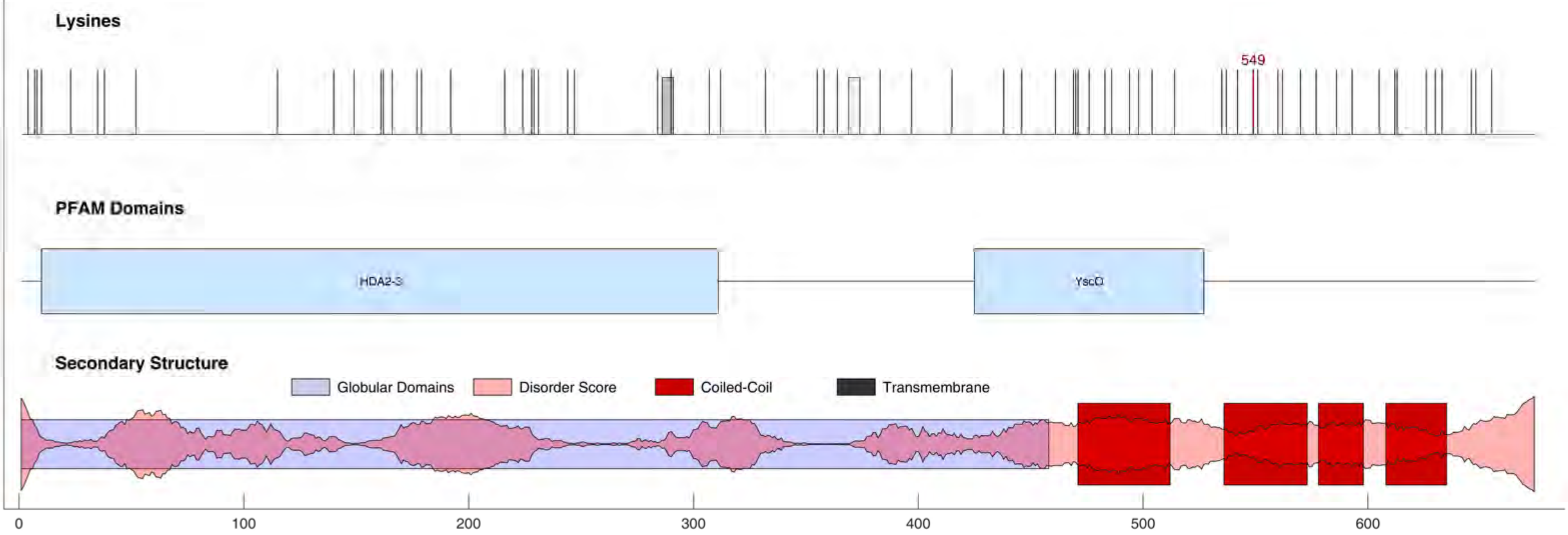

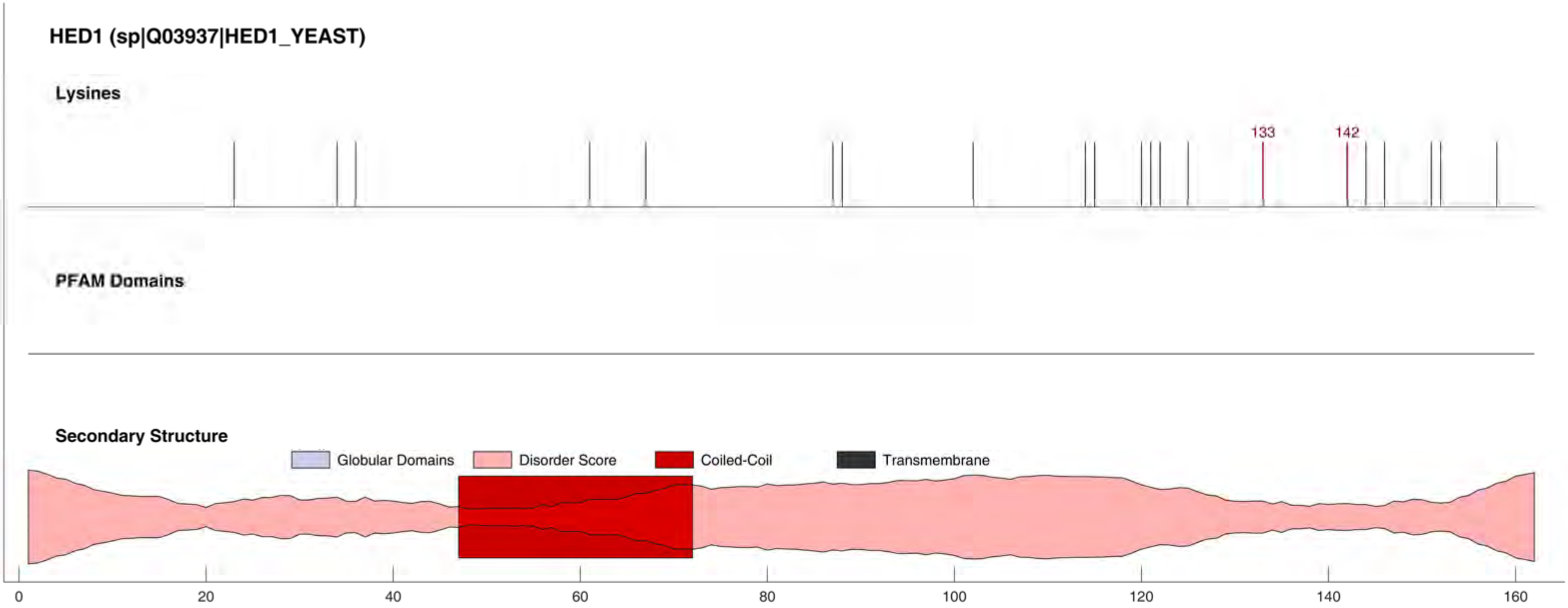
HED1 (sp|Q03937|HED1_YEAST)
Lysines
133
142
PFAM Domains
Secondary Structure
Globular Domains
Disorder Score
Coiled-Coil
Transmembrane
0
20
40
60
80
100
120
140
160

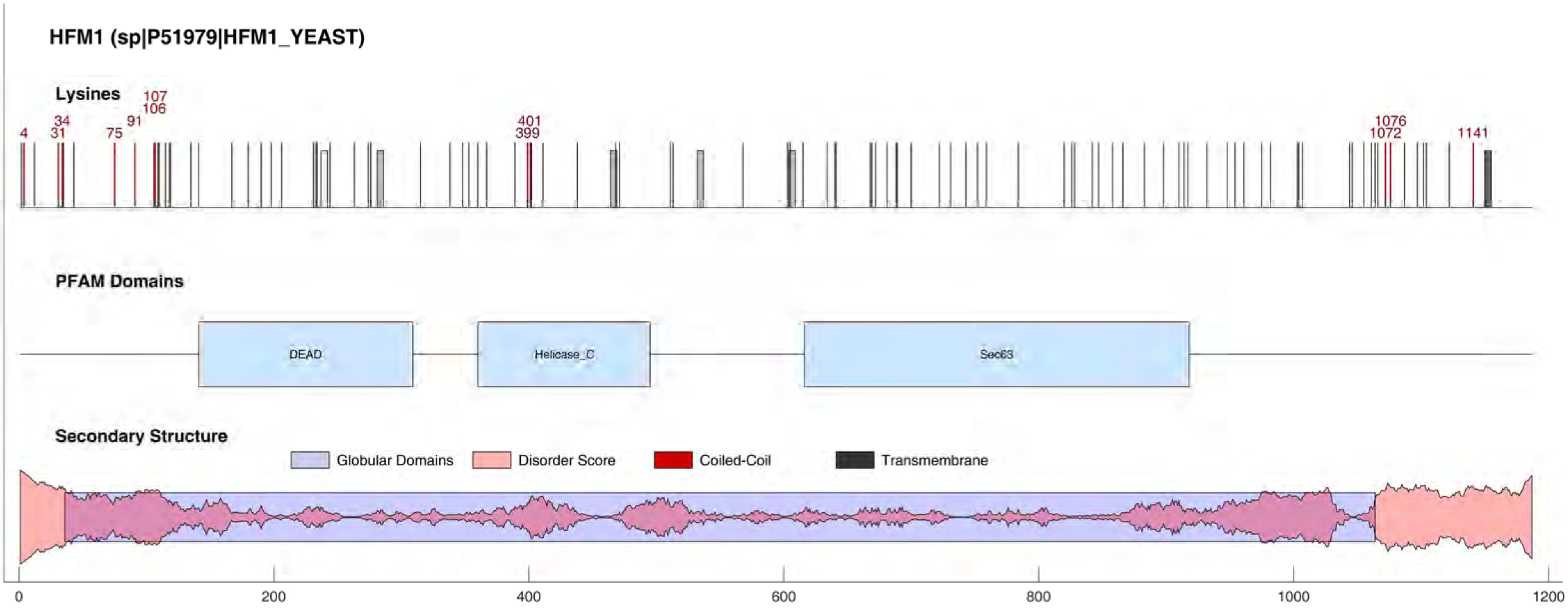
HFM1 (sp|P51979|HFM1_YEAST)
Lysines
4
31
34
75
91
106
107
399
401
1072
1076
1141
PFAM Domains
DEAD
Helicase_C
Sec63
Secondary Structure
Globular Domains
Disorder Score
Coiled-Coil
Transmembrane
0
200
400
600
800
1000
1200

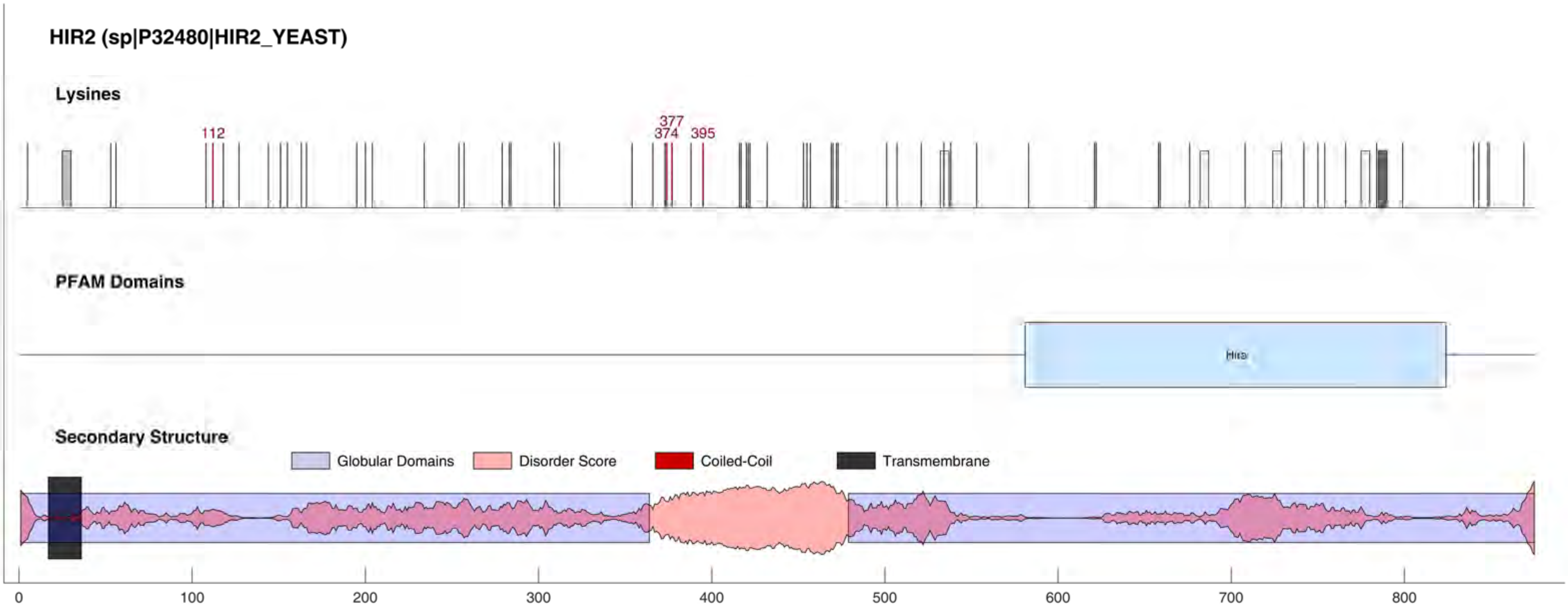
HIR2 (sp|P32480|HIR2_YEAST)
Lysines
112
377
374
395
PFAM Domains
Hira
Secondary Structure
Globular Domains
Disorder Score
Coiled-Coil
Transmembrane
0
100
200
300
400
500
600
700
800

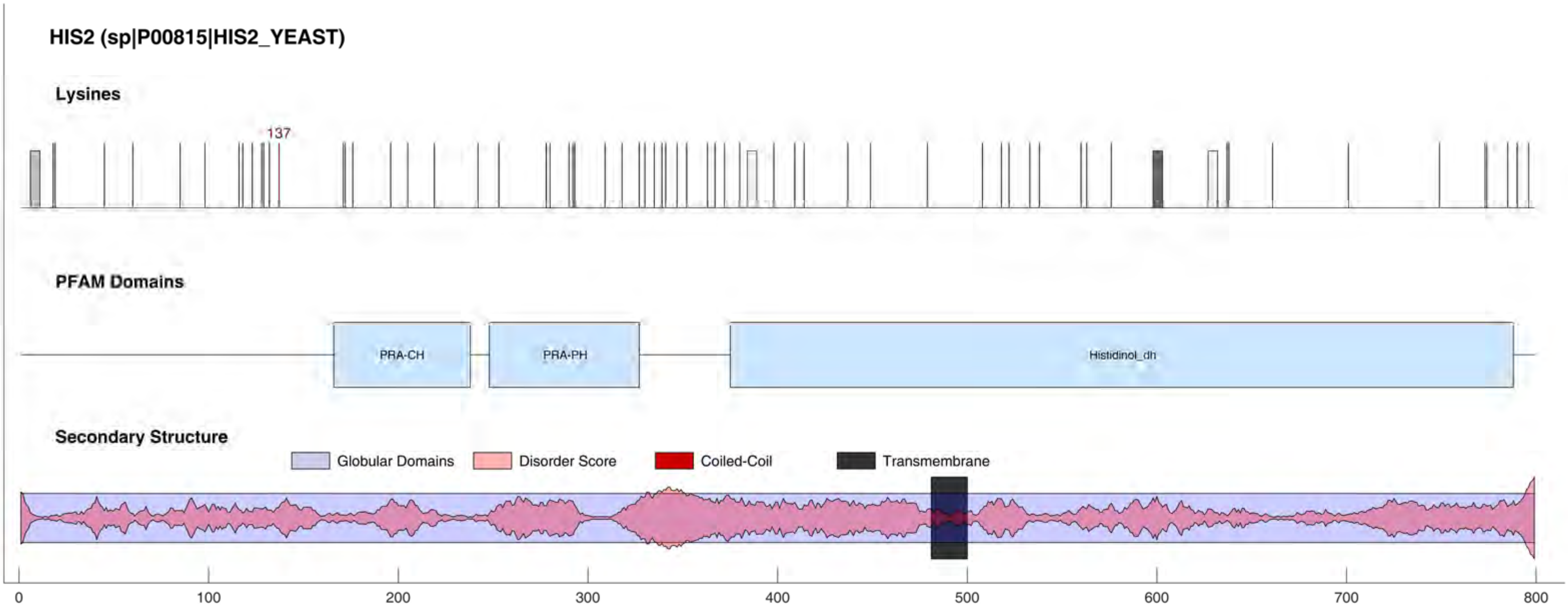
HIS2 (sp|P00815|HIS2_YEAST)
Lysines
137
PFAM Domains
PRA-CH
PRA-PH
Histidinol_dh
Secondary Structure
Globular Domains
Disorder Score
Coiled-Coil
Transmembrane
0
100
200
300
400
500
600
700
800

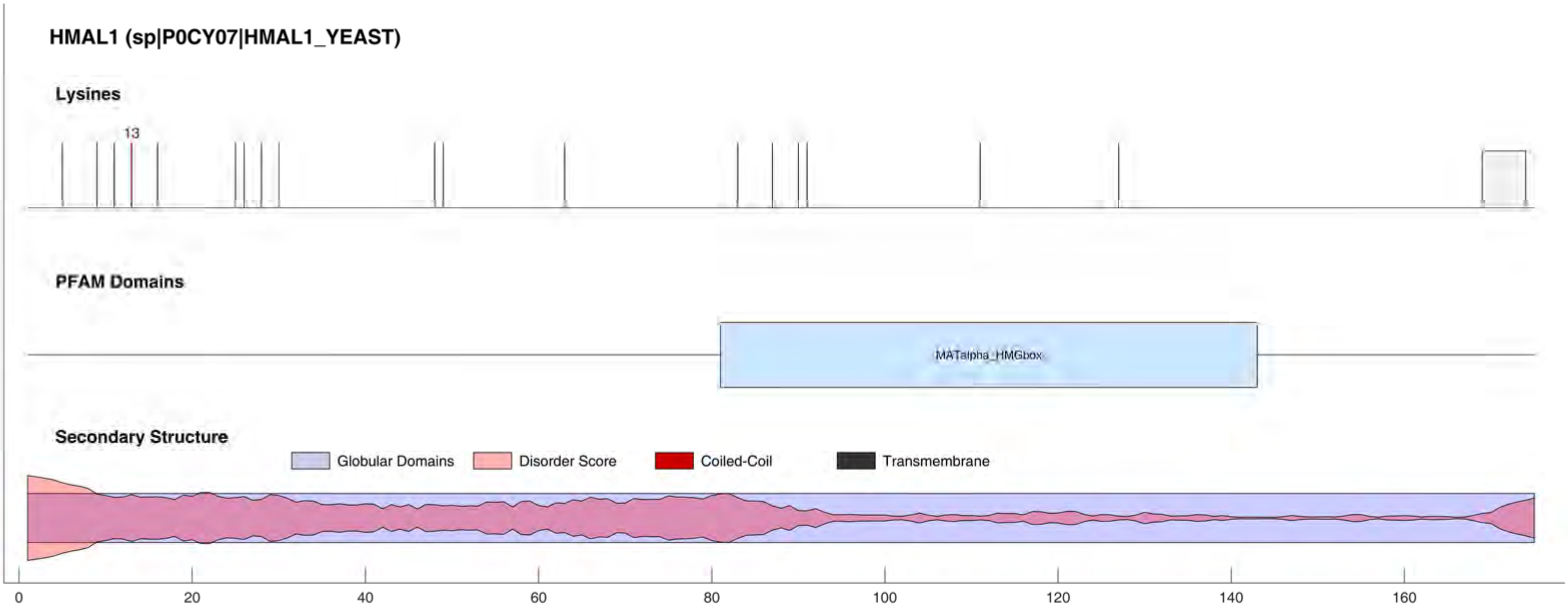

HMAL1 (sp|P0CY07|HMAL1_YEAST)
Lysines
13
PFAM Domains
MATalpha_HMGbox
Secondary Structure
Globular Domains
Disorder Score
Coiled-Coil
Transmembrane
0
20
40
60
80
100
120
140
160

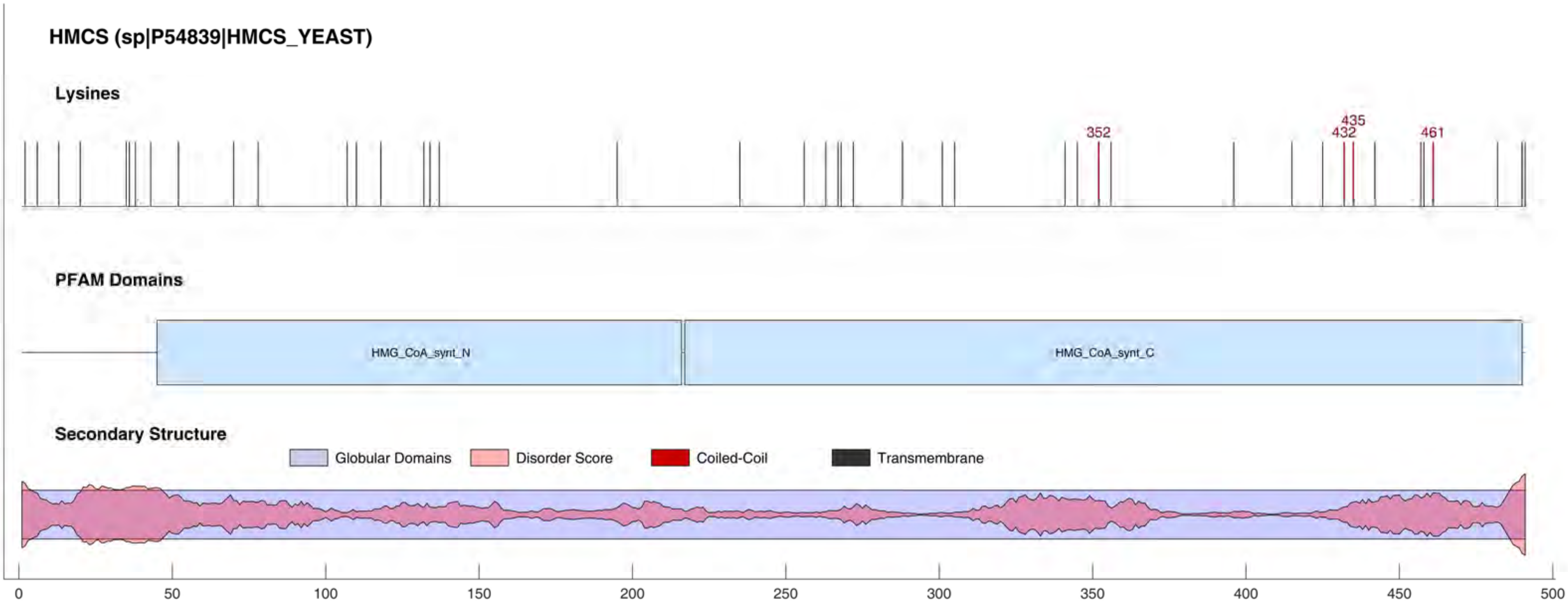

HMCS (sp|P54839|HMCS_YEAST)
Lysines
352
435
432
461
PFAM Domains
HMG_CoA_synt_N
HMG_CoA_synt_C
Secondary Structure
Globular Domains
Disorder Score
Coiled-Coil
Transmembrane
0
50
100
150
200
250
300
350
400
450
500

HMO1 (sp|Q03973|HMO1_YEAST)

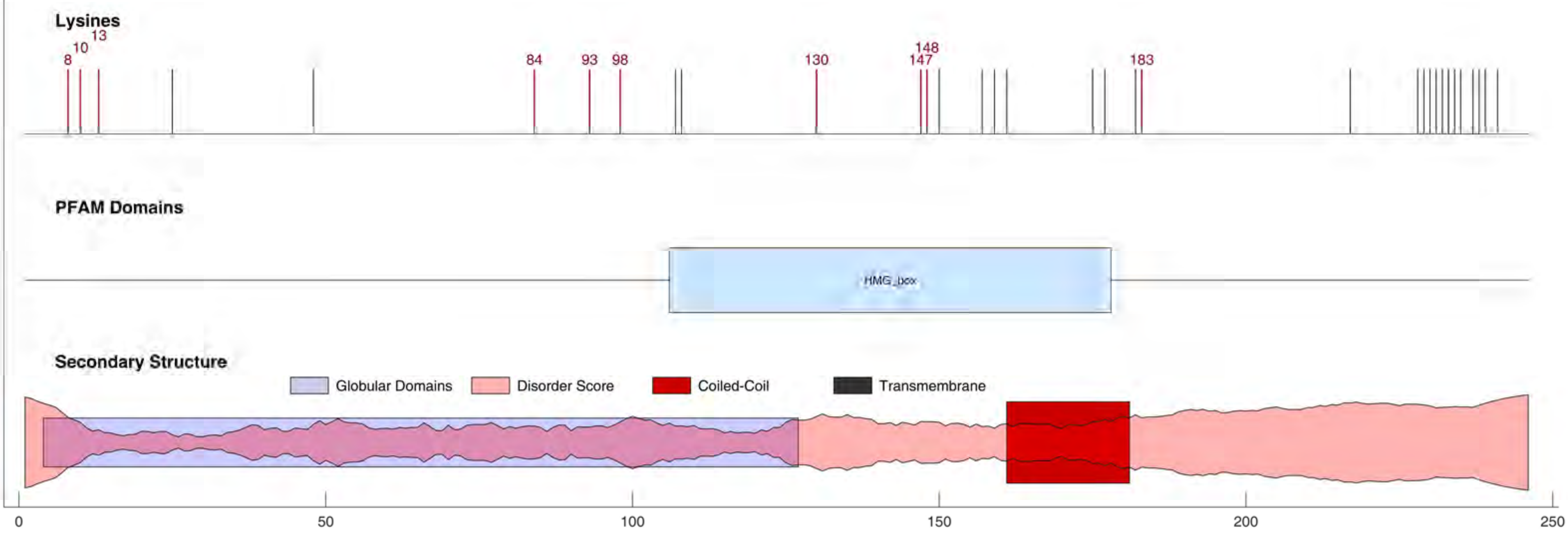

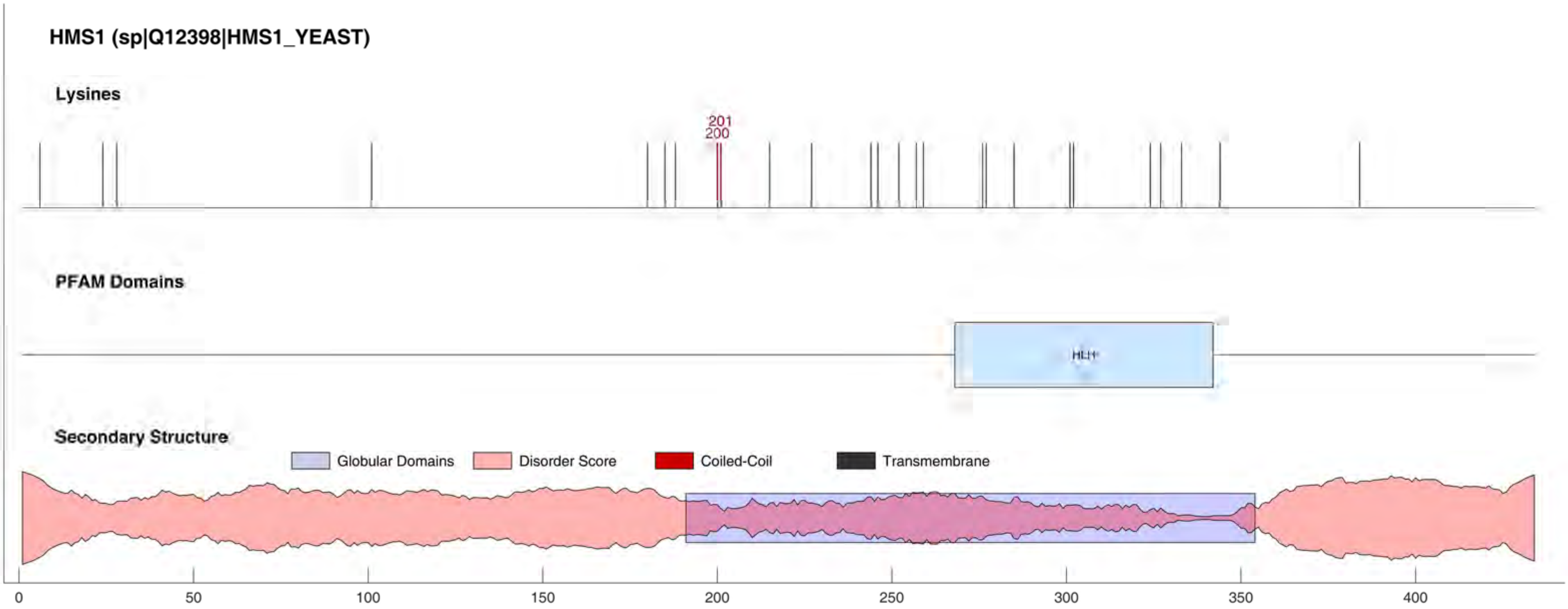
HMS1 (sp|Q12398|HMS1_YEAST)
Lysines
201
200
PFAM Domains
HLH
Secondary Structure
Globular Domains
Disorder Score
Coiled-Coil
Transmembrane
0
50
100
150
200
250
300
350
400

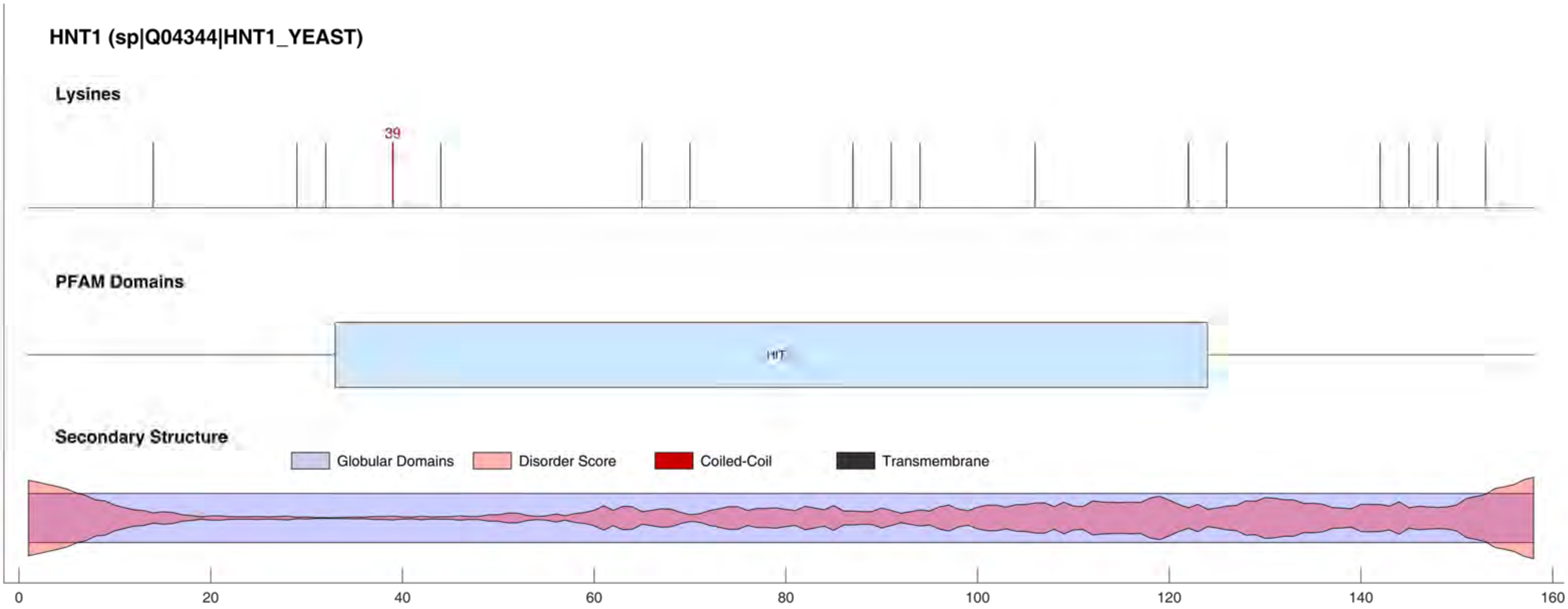
HNT1 (sp|Q04344|HNT1_YEAST)
Lysines
39
PFAM Domains
HIT
Secondary Structure
Globular Domains
Disorder Score
Coiled-Coil
Transmembrane
0
20
40
60
80
100
120
140
160

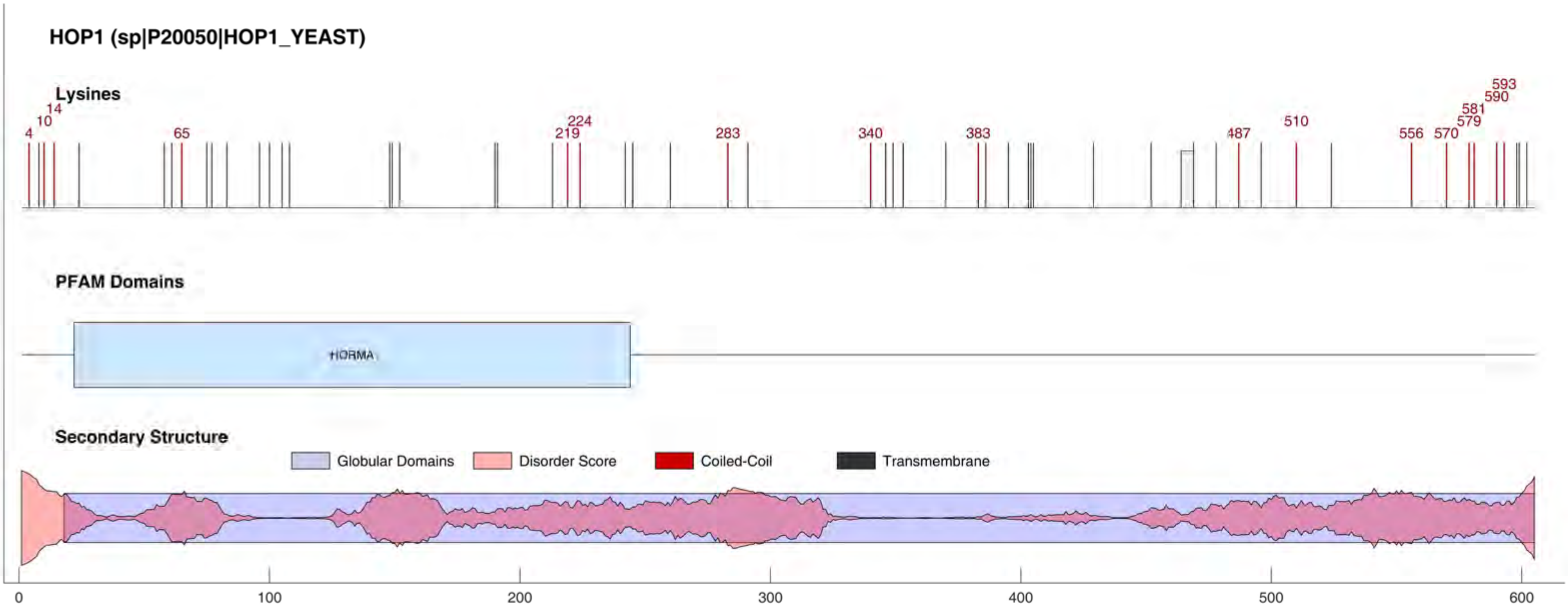

HOP1 (sp|P20050|HOP1_YEAST)
Lysines
4
10
14
65
219
224
283
340
383
487
510
556
570
579
581
590
593
PFAM Domains
HORMA
Secondary Structure
Globular Domains
Disorder Score
Coiled-Coil
Transmembrane
0
100
200
300
400
500
600

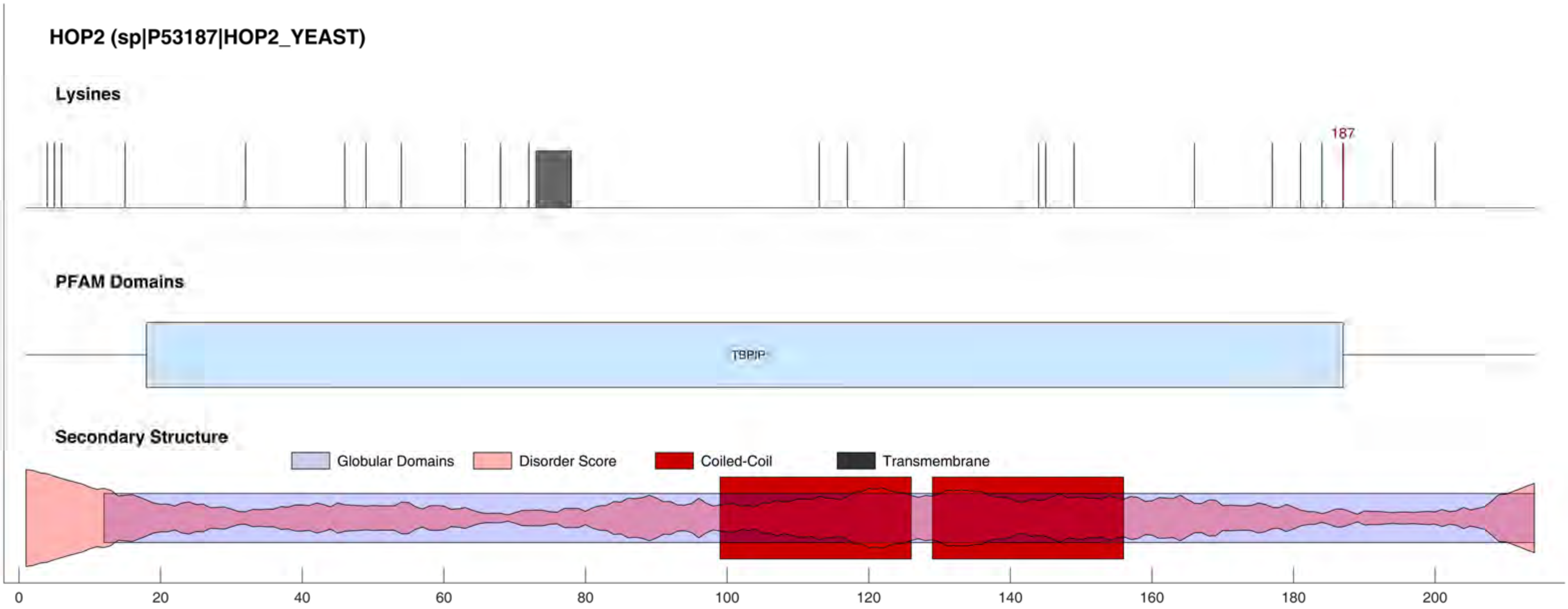
HOP2 (sp|P53187|HOP2_YEAST)
Lysines
187
PFAM Domains
TBPIP
Secondary Structure
Globular Domains
Disorder Score
Coiled-Coil
Transmembrane
0
20
40
60
80
100
120
140
160
180
200

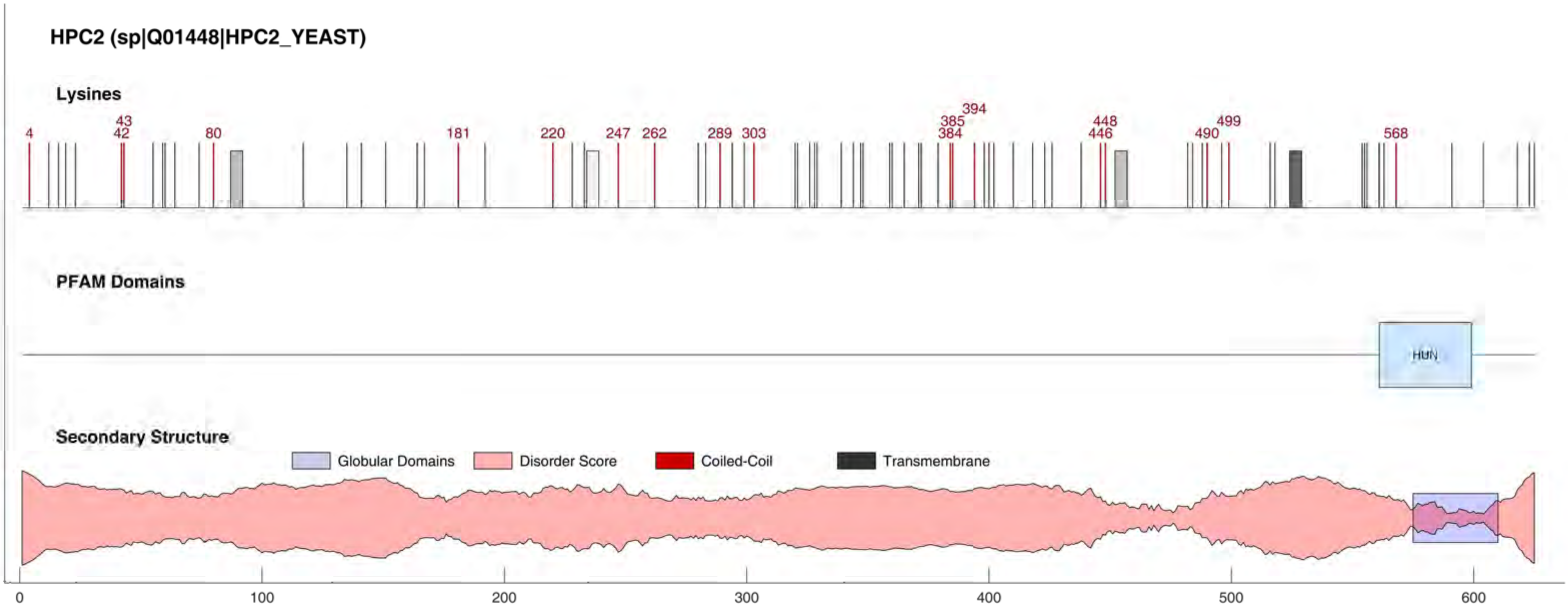

HPC2 (sp|Q01448|HPC2_YEAST)
Lysines
4
43
42
80
181
220
247
262
289
303
394
385
384
448
446
499
490
568
PFAM Domains
HUN
Secondary Structure
Globular Domains
Disorder Score
Coiled-Coil
Transmembrane
0
100
200
300
400
500
600

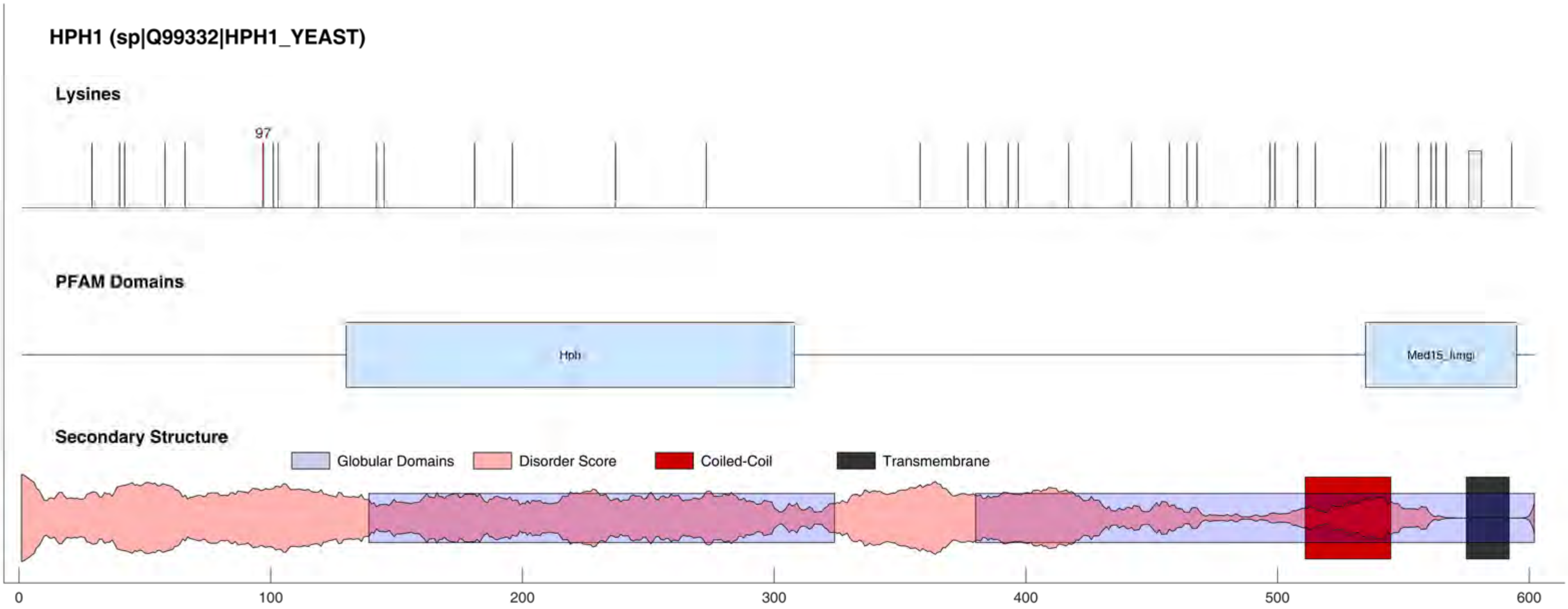

HPH1 (sp|Q99332|HPH1_YEAST)
Lysines
97
PFAM Domains
Hph
Med15_fungi
Secondary Structure
Globular Domains
Disorder Score
Coiled-Coil
Transmembrane
0
100
200
300
400
500
600

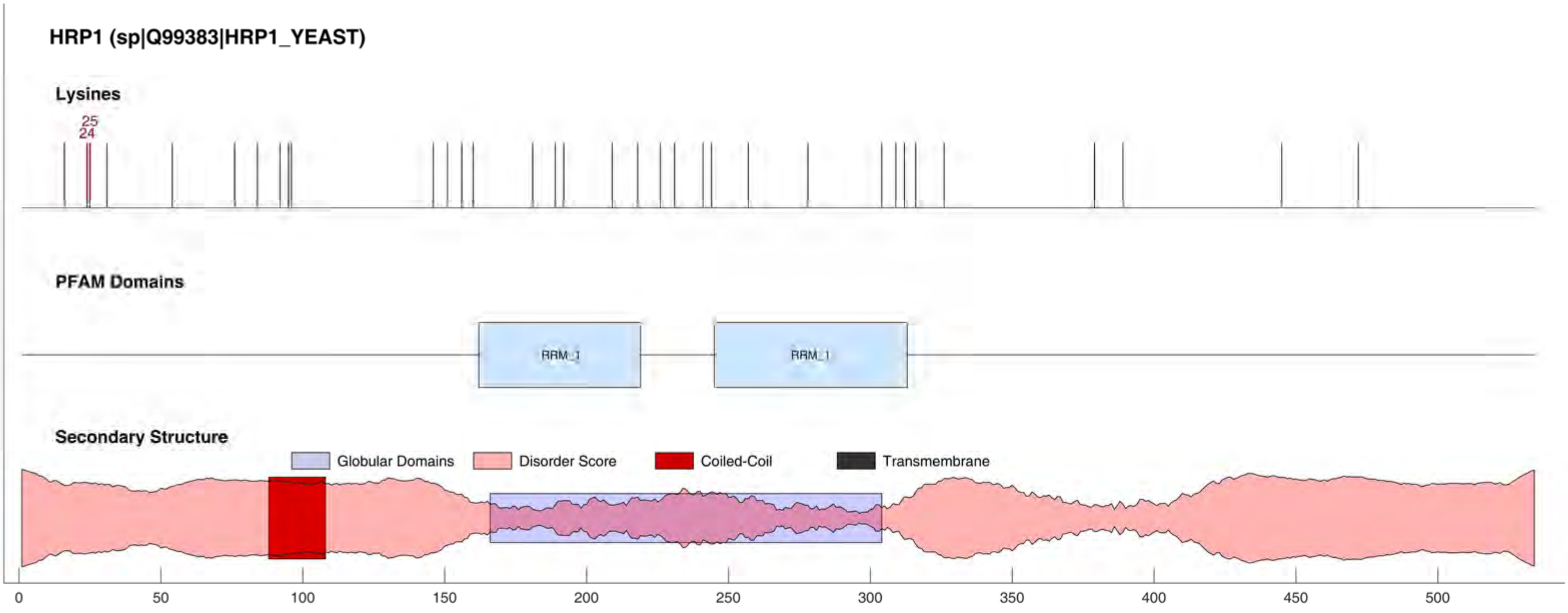
HRP1 (sp|Q99383|HRP1_YEAST)
Lysines
25
24
PFAM Domains
RRM_1
RRM_1
Secondary Structure
Globular Domains
Disorder Score
Coiled-Coil
Transmembrane
0
50
100
150
200
250
300
350
400
450
500

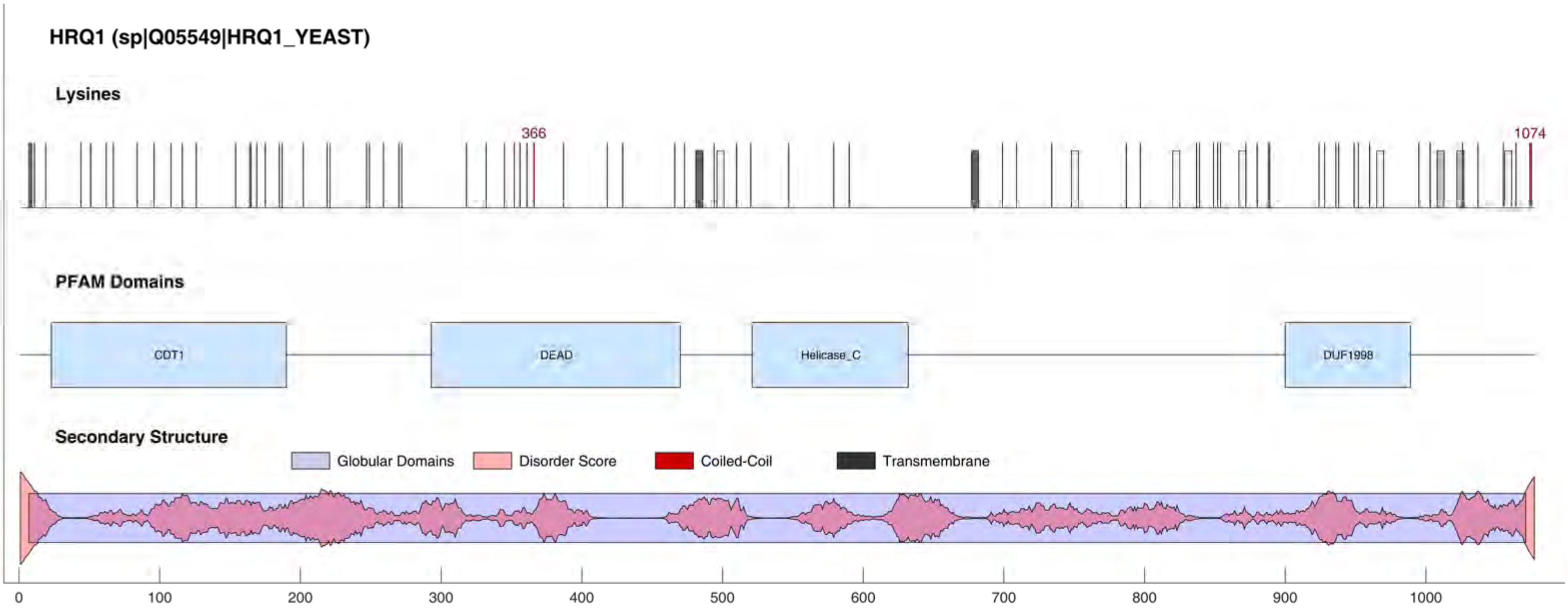
HRQ1 (sp|Q05549|HRQ1_YEAST)
Lysines
366
1074
PFAM Domains
CDT1
DEAD
Helicase_C
DUF1998
Secondary Structure
Globular Domains
Disorder Score
Coiled-Coil
Transmembrane
0
100
200
300
400
500
600
700
800
900
1000

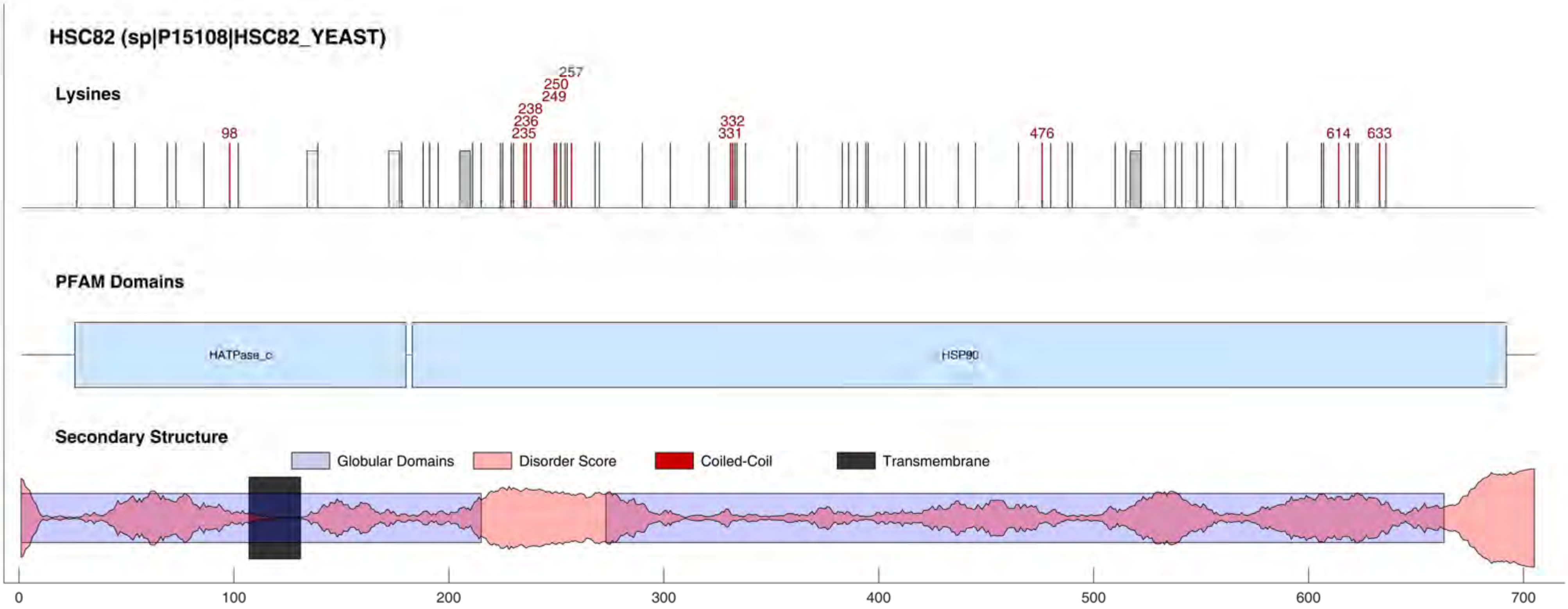

HSC82 (sp|P15108|HSC82_YEAST)
Lysines
98
235
236
238
249
250
257
331
332
476
614
633
PFAM Domains
HATPase_c
HSP90
Secondary Structure
Globular Domains
Disorder Score
Coiled-Coil
Transmembrane
0
100
200
300
400
500
600
700

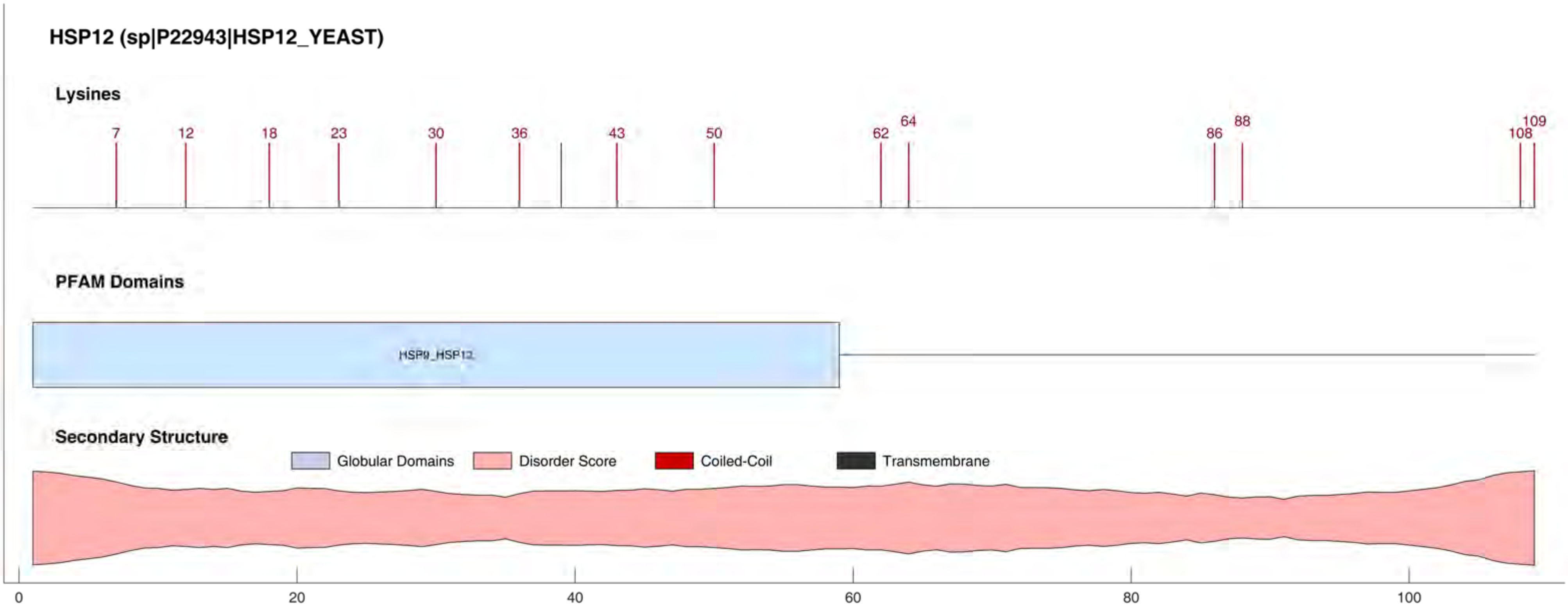

HSP12 (sp|P22943|HSP12_YEAST)
Lysines
7
12
18
23
30
36
43
50
62
64
86
88
108
109
PFAM Domains
HSP9_HSP12
Secondary Structure
Globular Domains
Disorder Score
Coiled-Coil
Transmembrane
0
20
40
60
80
100

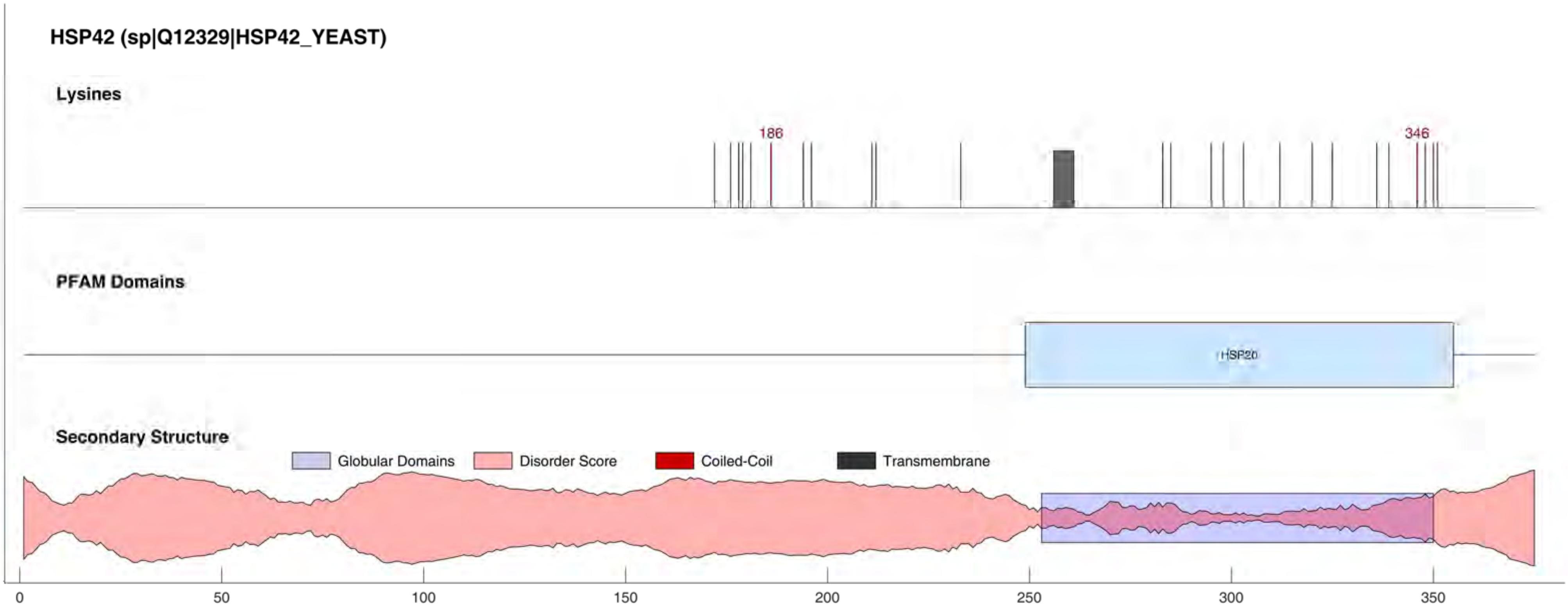

HSP42 (sp|Q12329|HSP42_YEAST)
Lysines
186
346
PFAM Domains
HSP20
Secondary Structure
Globular Domains
Disorder Score
Coiled-Coil
Transmembrane
0
50
100
150
200
250
300
350

#### HSP71 (sp|P10591|HSP71_YEAST)

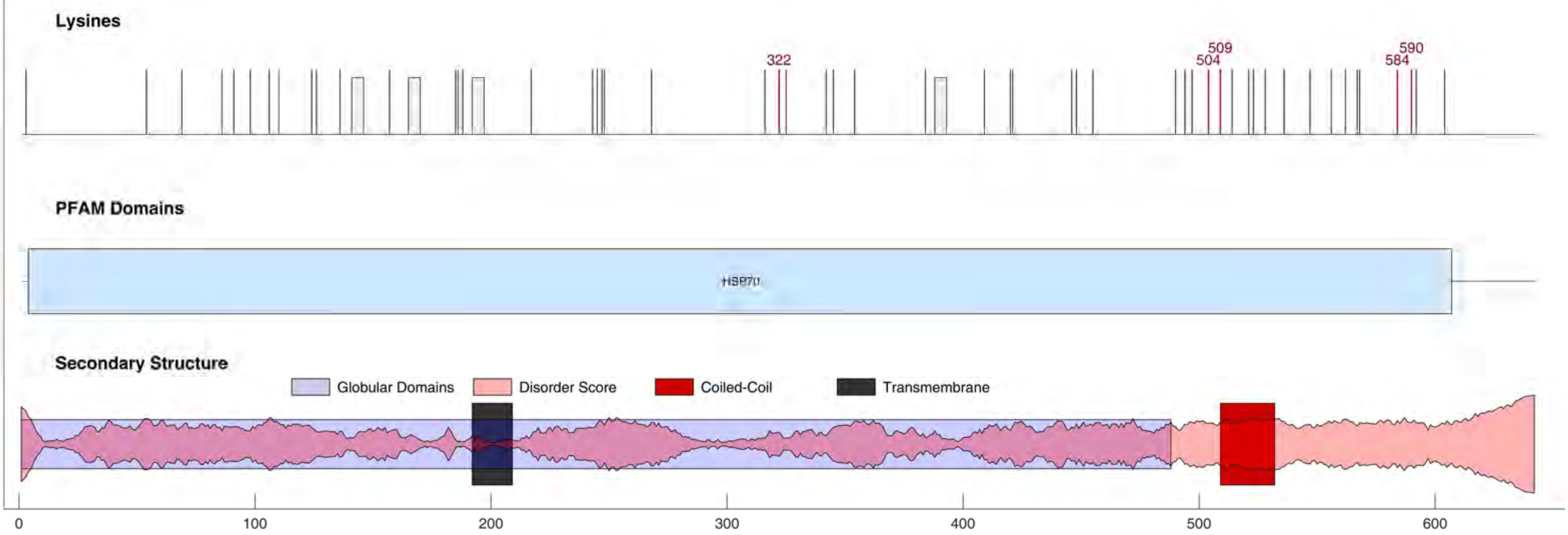

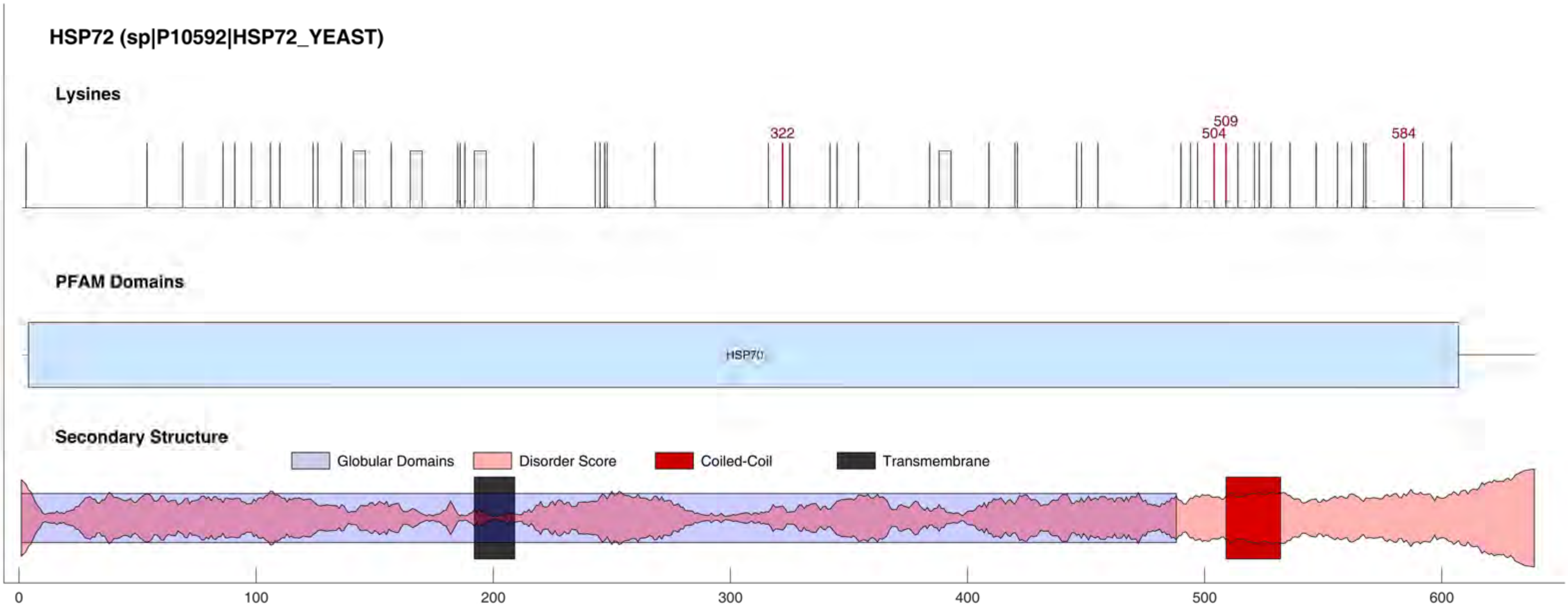
HSP72 (sp|P10592|HSP72_YEAST)
Lysines
322
509
504
584
PFAM Domains
HSP70
Secondary Structure
Globular Domains
Disorder Score
Coiled-Coil
Transmembrane
0
100
200
300
400
500
600

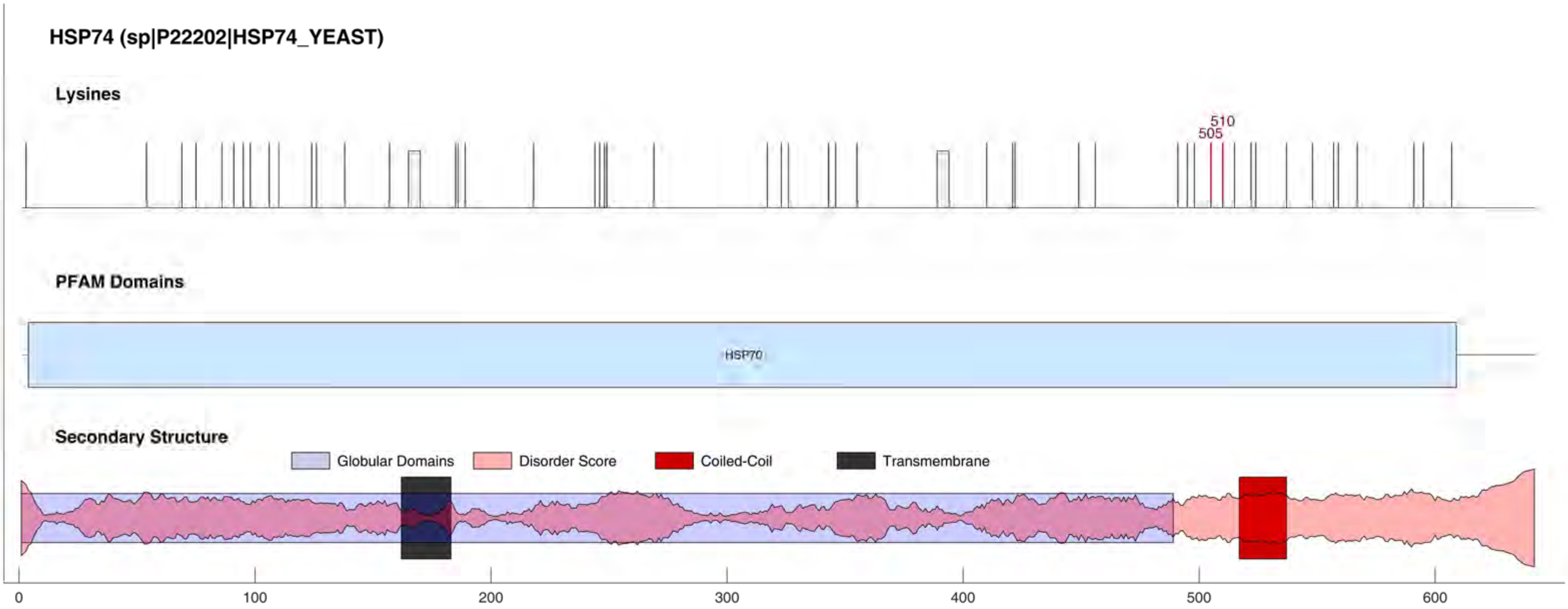
HSP74 (sp|P22202|HSP74_YEAST)
Lysines
510
505
PFAM Domains
HSP70
Secondary Structure
Globular Domains
Disorder Score
Coiled-Coil
Transmembrane
0
100
200
300
400
500
600

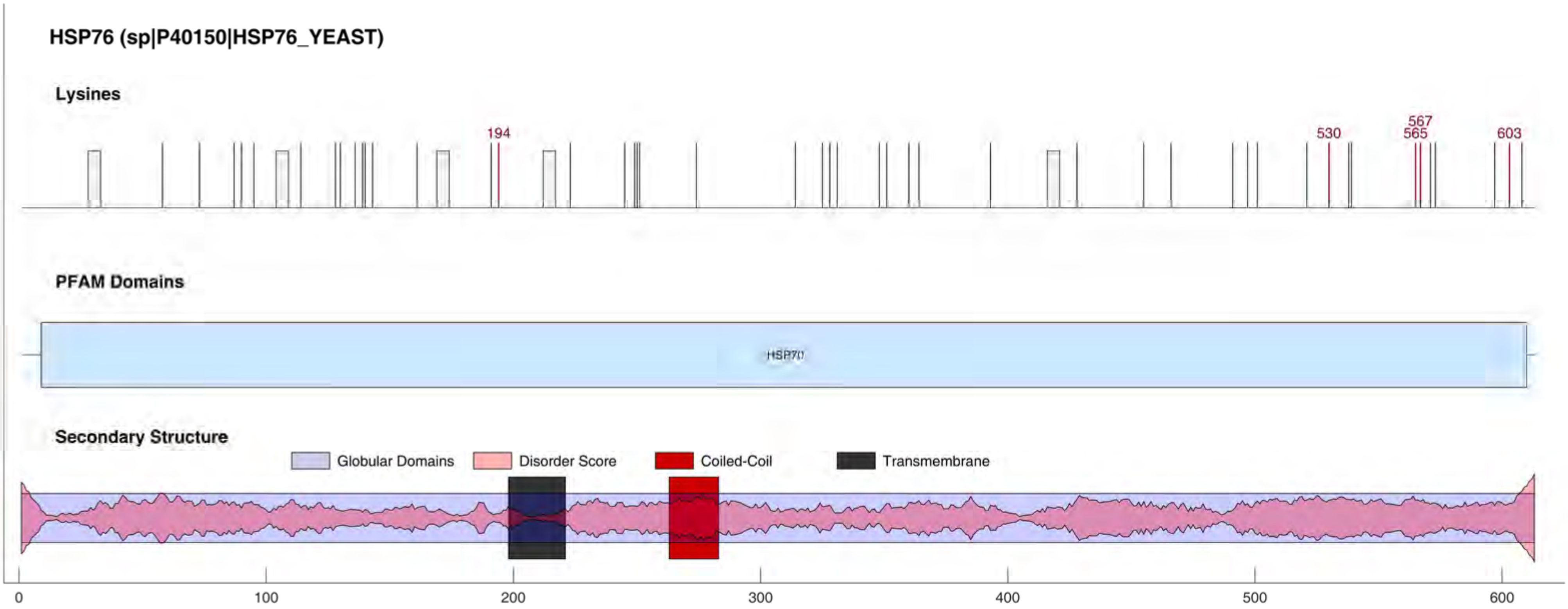
HSP76 (sp|P40150|HSP76_YEAST)
Lysines
194
530
567
565
603
PFAM Domains
HSP70
Secondary Structure
Globular Domains
Disorder Score
Coiled-Coil
Transmembrane
0
100
200
300
400
500
600

#### HSP79 (sp|P32590|HSP79_YEAST)

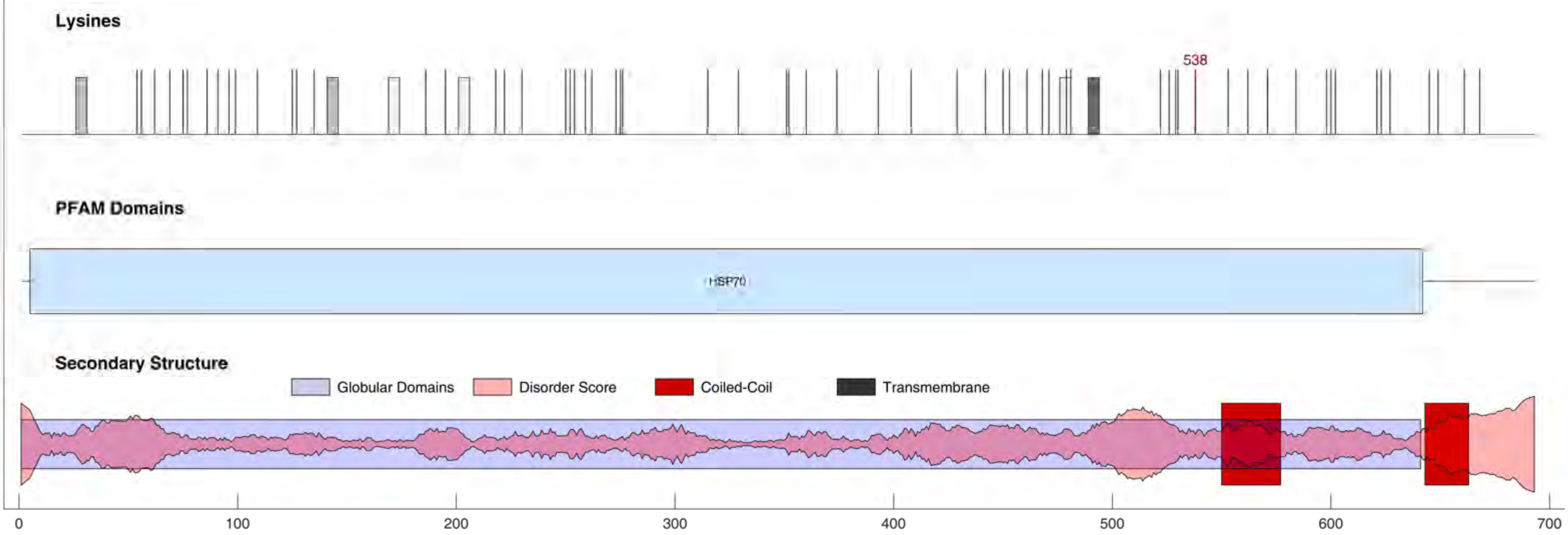

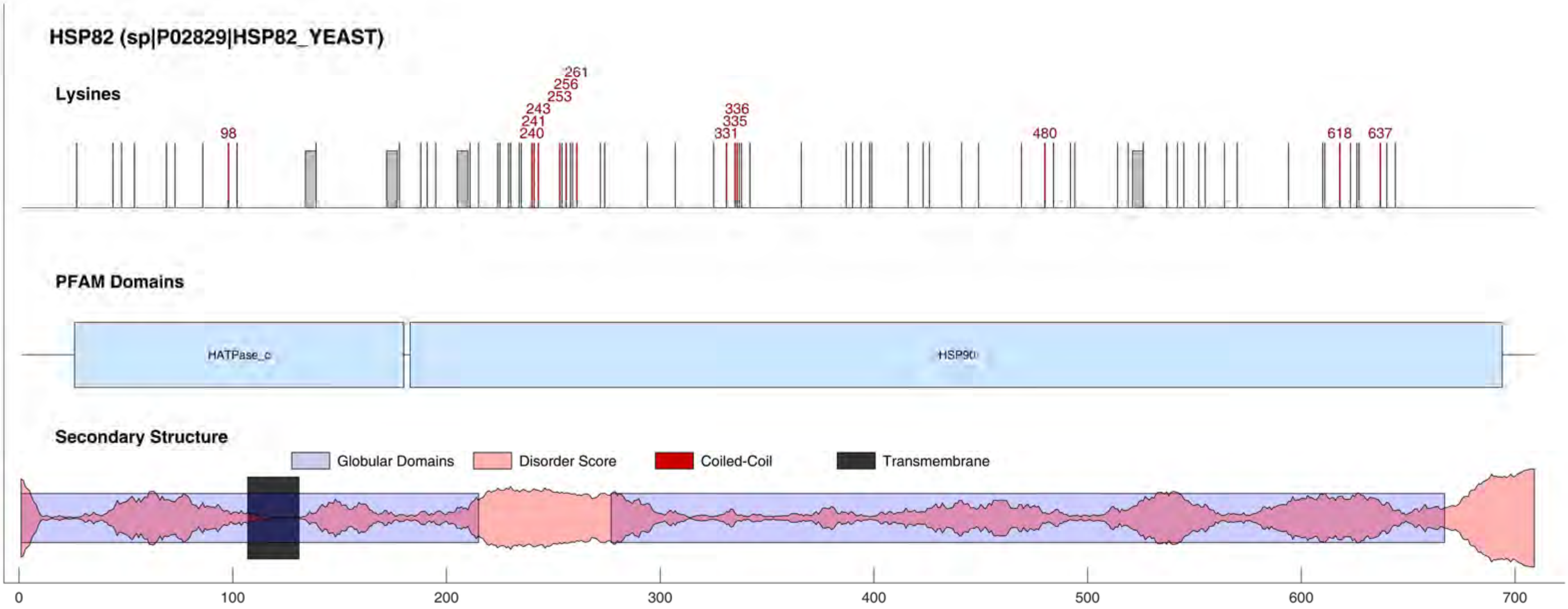
HSP82 (sp|P02829|HSP82_YEAST)
Lysines
98
240
241
243
253
256
261
331
335
336
480
618
637
PFAM Domains
HATPase_c
HSP90
Secondary Structure
Globular Domains
Disorder Score
Coiled-Coil
Transmembrane
0
100
200
300
400
500
600
700

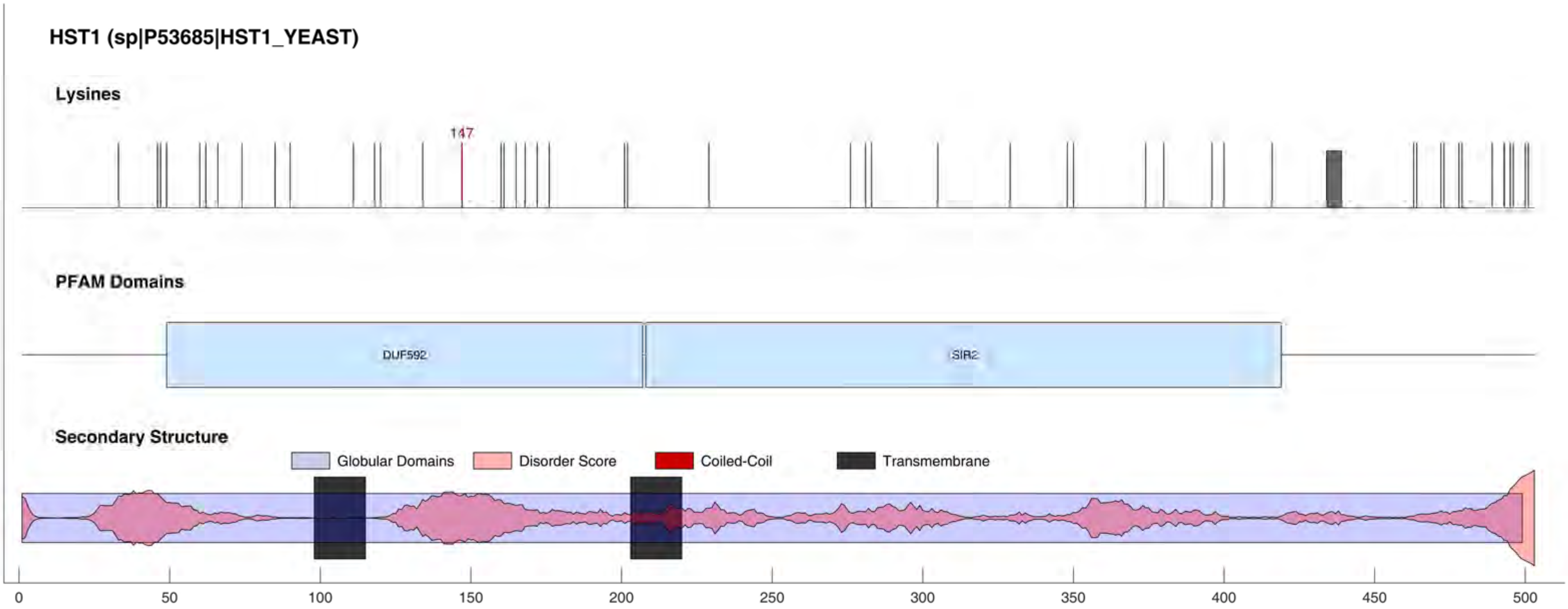

HST1 (sp|P53685|HST1_YEAST)
Lysines
147
PFAM Domains
DUF592
SIR2
Secondary Structure
Globular Domains
Disorder Score
Coiled-Coil
Transmembrane
0
50
100
150
200
250
300
350
400
450
500

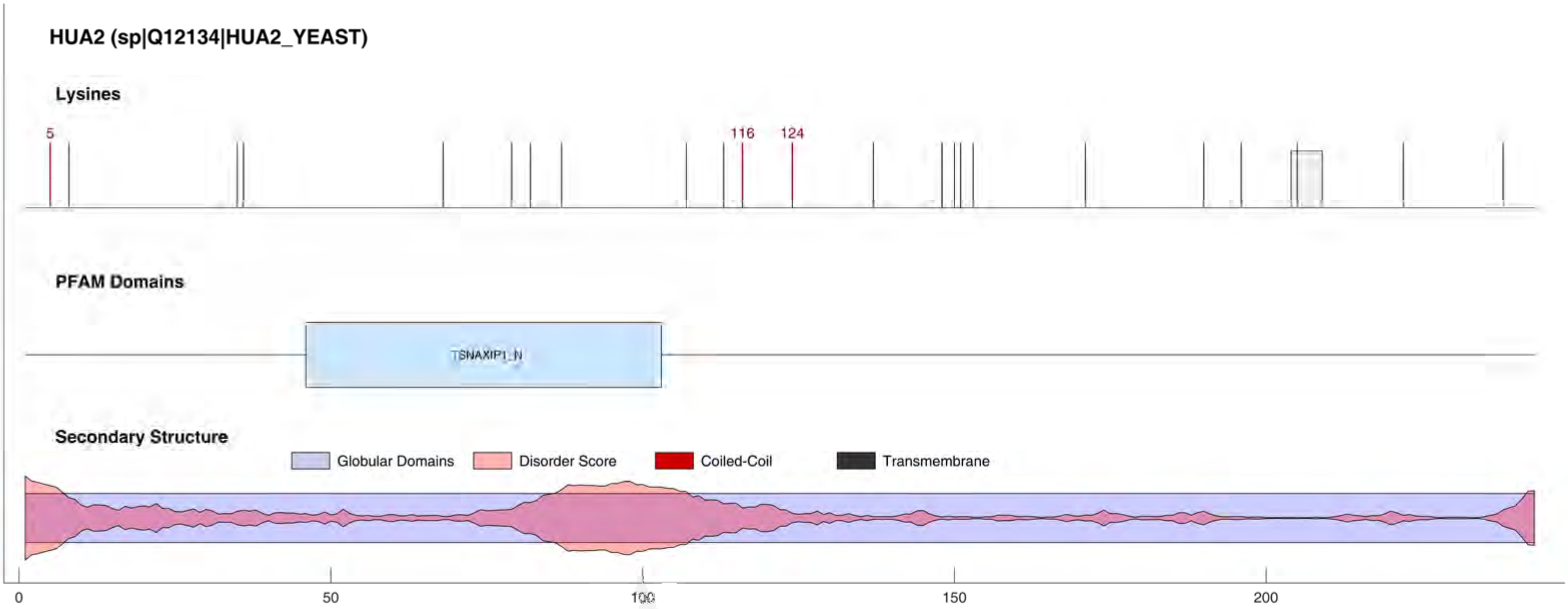
HUA2 (sp|Q12134|HUA2_YEAST)
Lysines
5
116
124
PFAM Domains
TSNAXIP1_N
Secondary Structure
Globular Domains
Disorder Score
Coiled-Coil
Transmembrane
0
50
100
150
200

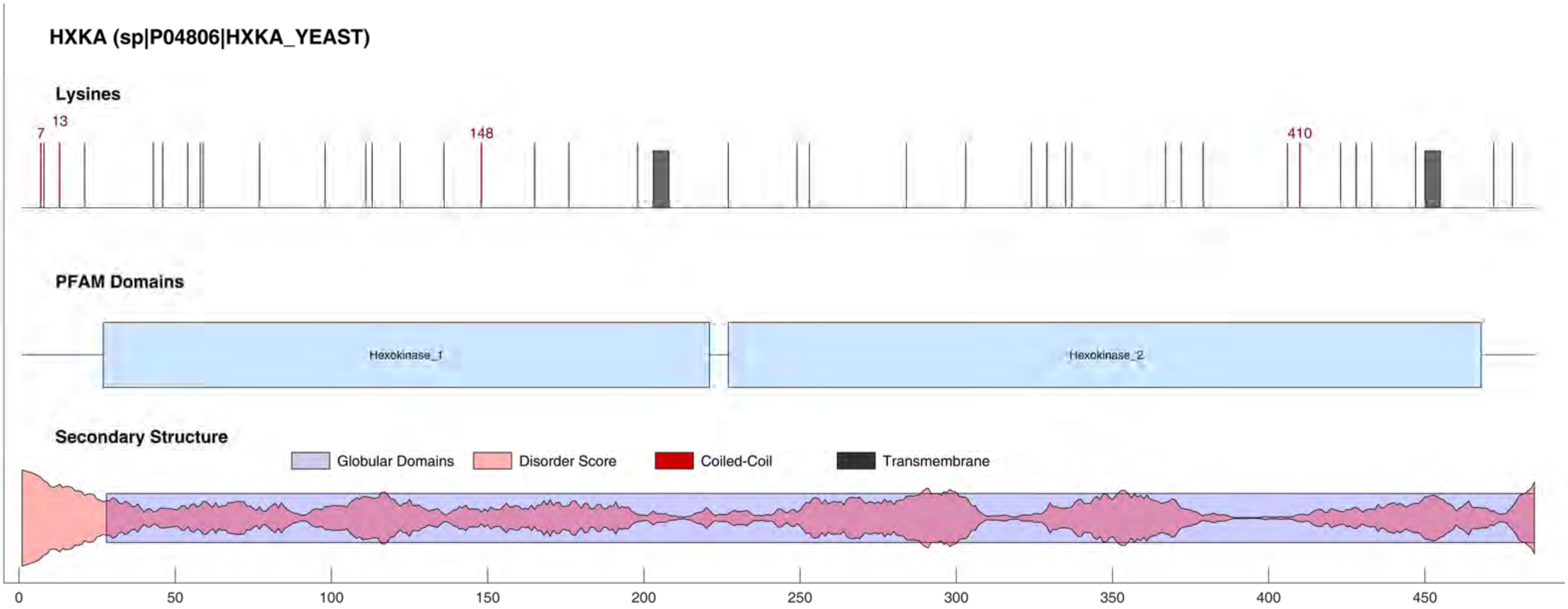
HXKA (sp|P04806|HXKA_YEAST)
Lysines
7
13
148
410
PFAM Domains
Hexokinase_1
Hexokinase_2
Secondary Structure
Globular Domains
Disorder Score
Coiled-Coil
Transmembrane
0
50
100
150
200
250
300
350
400
450

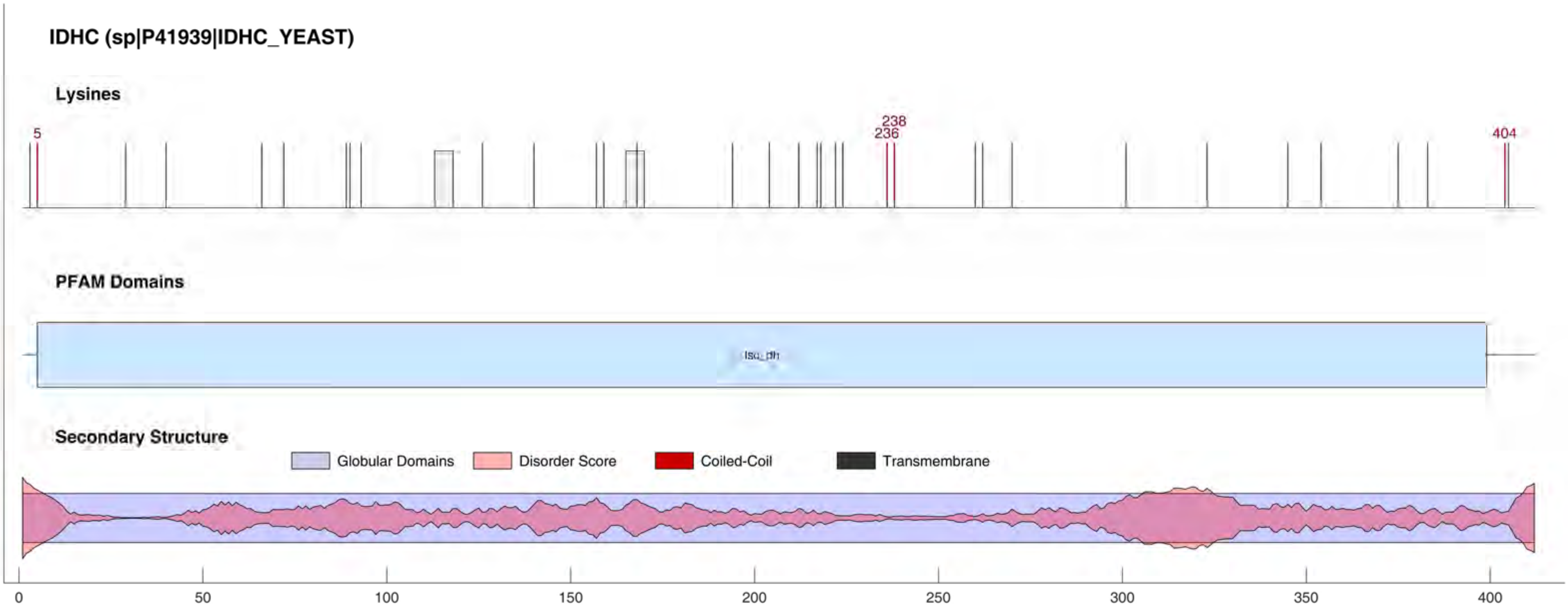
IDHC (sp|P41939|IDHC_YEAST)
Lysines
5
238
236
404
PFAM Domains
Iso_dh
Secondary Structure
Globular Domains
Disorder Score
Coiled-Coil
Transmembrane
0
50
100
150
200
250
300
350
400

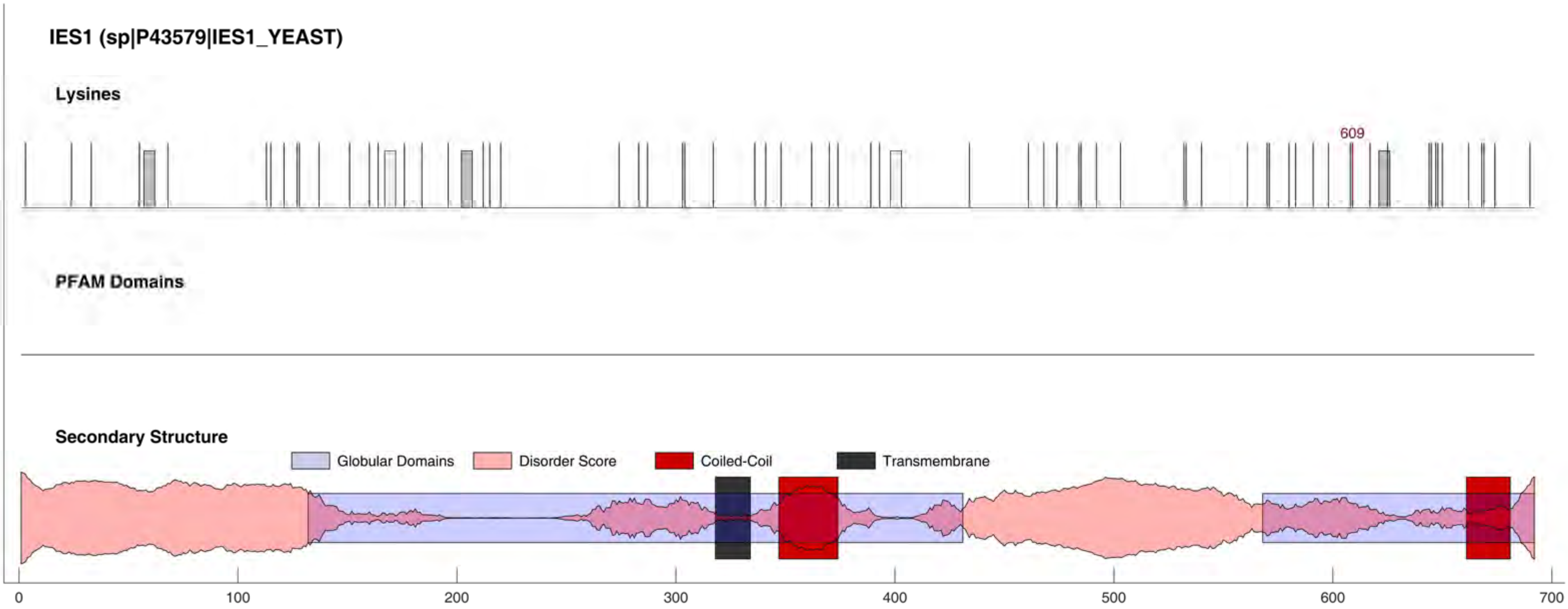
IES1 (sp|P43579|IES1_YEAST)
Lysines
609
PFAM Domains
Secondary Structure
Globular Domains
Disorder Score
Coiled-Coil
Transmembrane
0
100
200
300
400
500
600
700

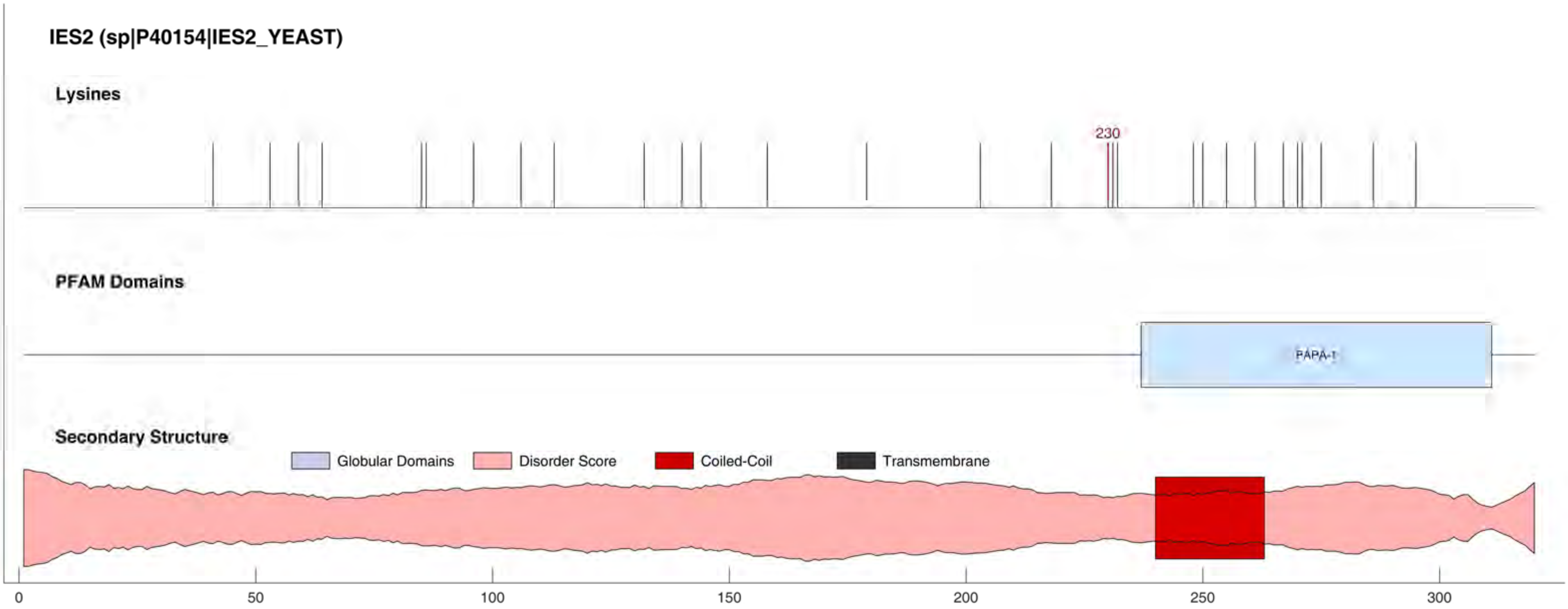
IES2 (sp|P40154|IES2_YEAST)
Lysines
230
PFAM Domains
PAPA-1
Secondary Structure
Globular Domains
Disorder Score
Coiled-Coil
Transmembrane
0
50
100
150
200
250
300

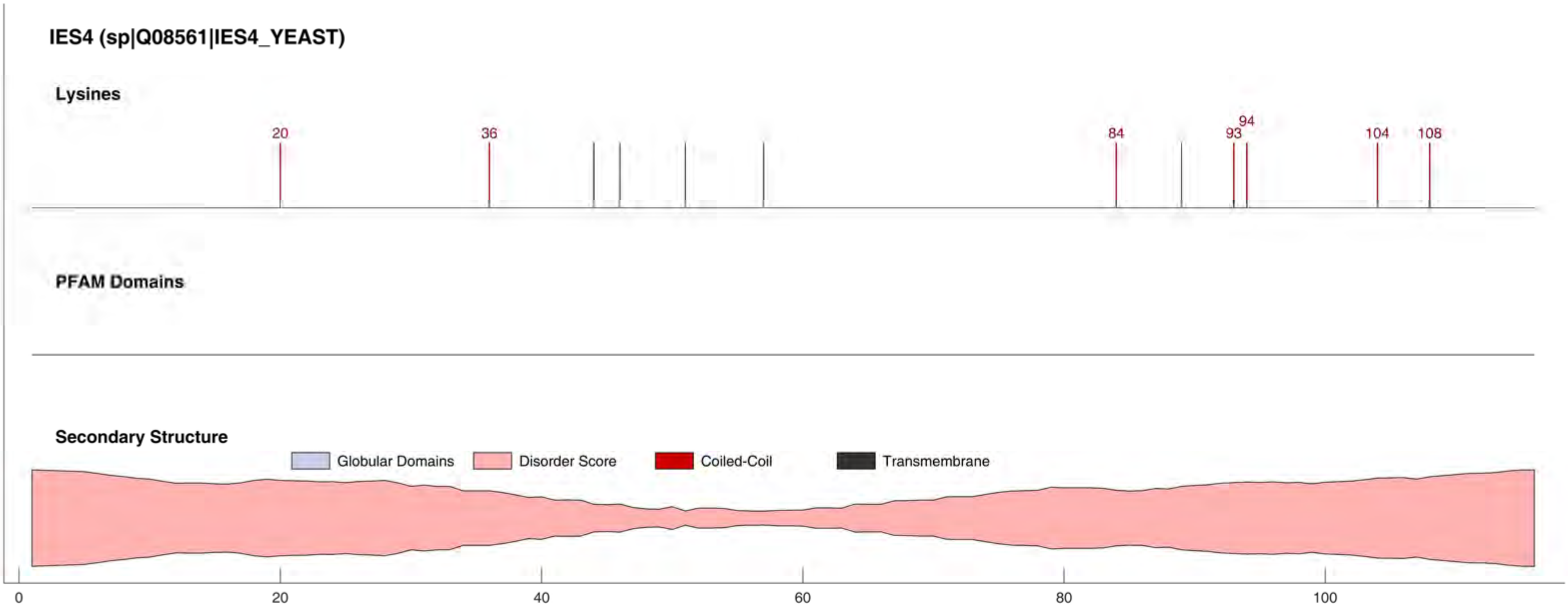
IES4 (sp|Q08561|IES4_YEAST)
Lysines
20
36
84
93
94
104
108
PFAM Domains
Secondary Structure
Globular Domains
Disorder Score
Coiled-Coil
Transmembrane
0
20
40
60
80
100

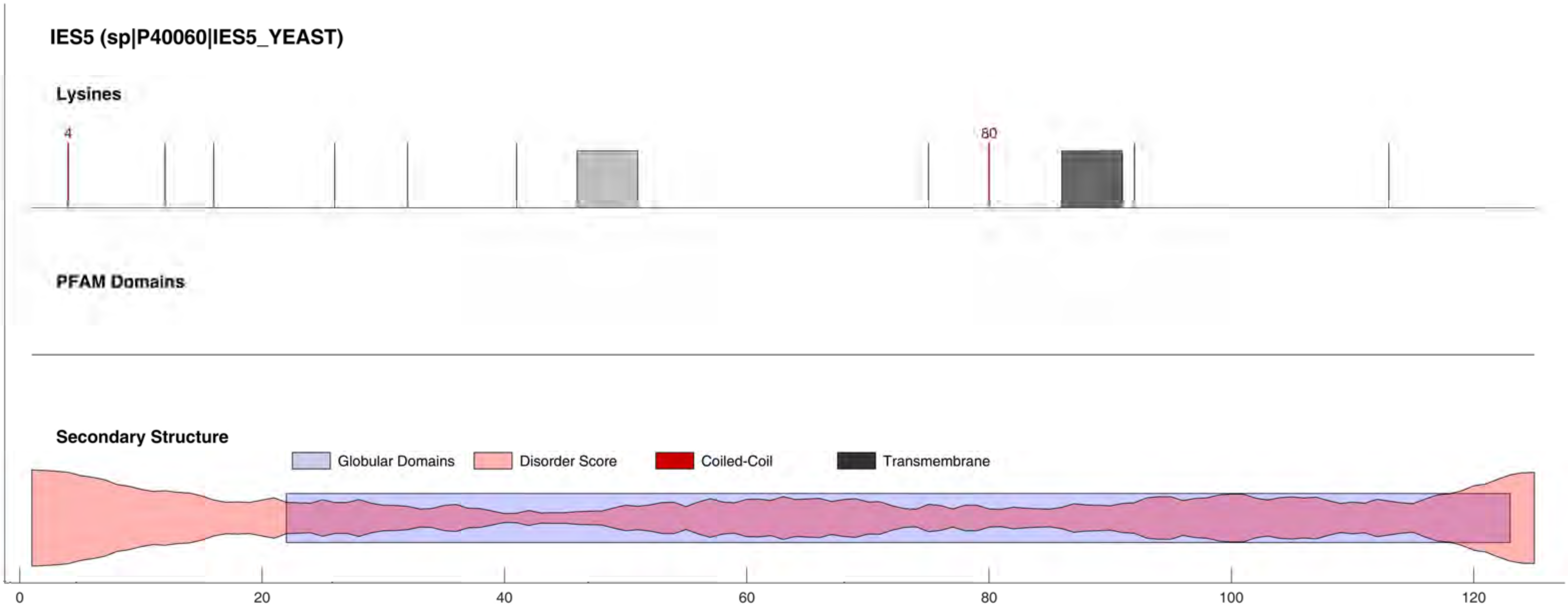
IES5 (sp|P40060|IES5_YEAST)
Lysines
4
80
PFAM Domains
Secondary Structure
Globular Domains
Disorder Score
Coiled-Coil
Transmembrane
0
20
40
60
80
100
120

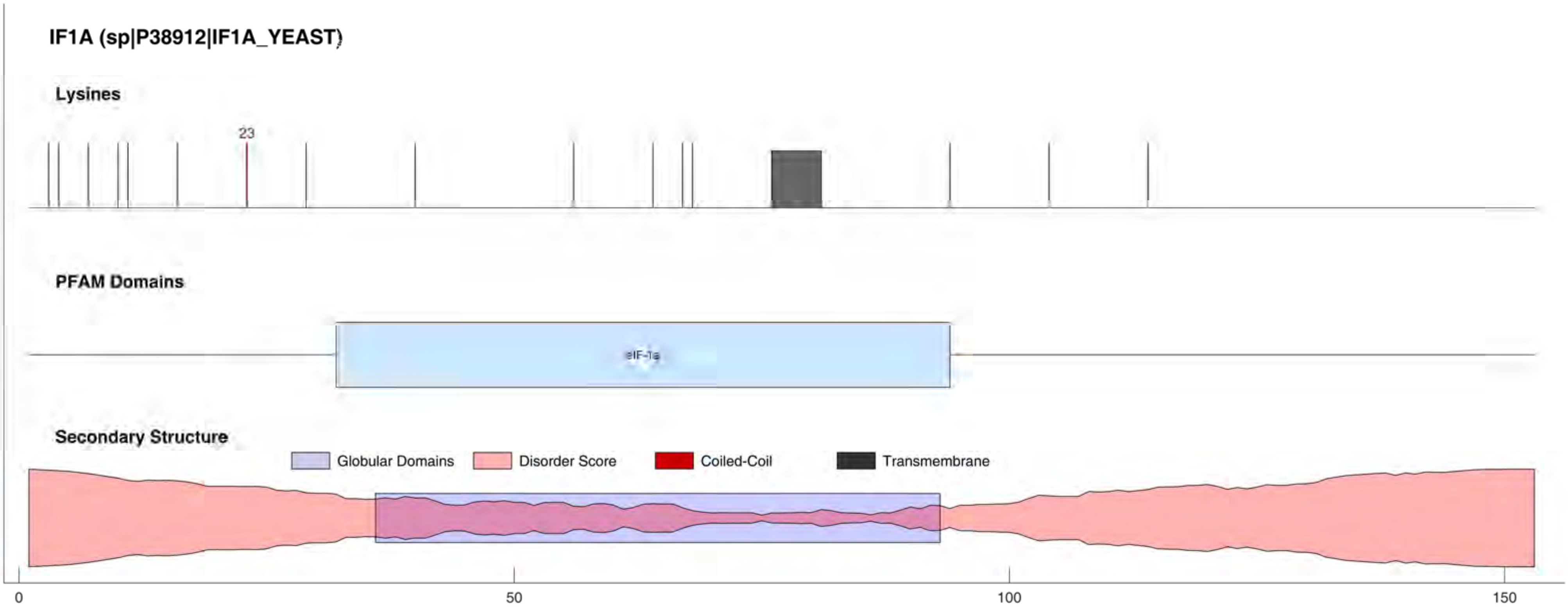
IF1A (sp|P38912|IF1A_YEAST)
Lysines
23
PFAM Domains
eIF-1a
Secondary Structure
Globular Domains
Disorder Score
Coiled-Coil
Transmembrane
0
50
100
150

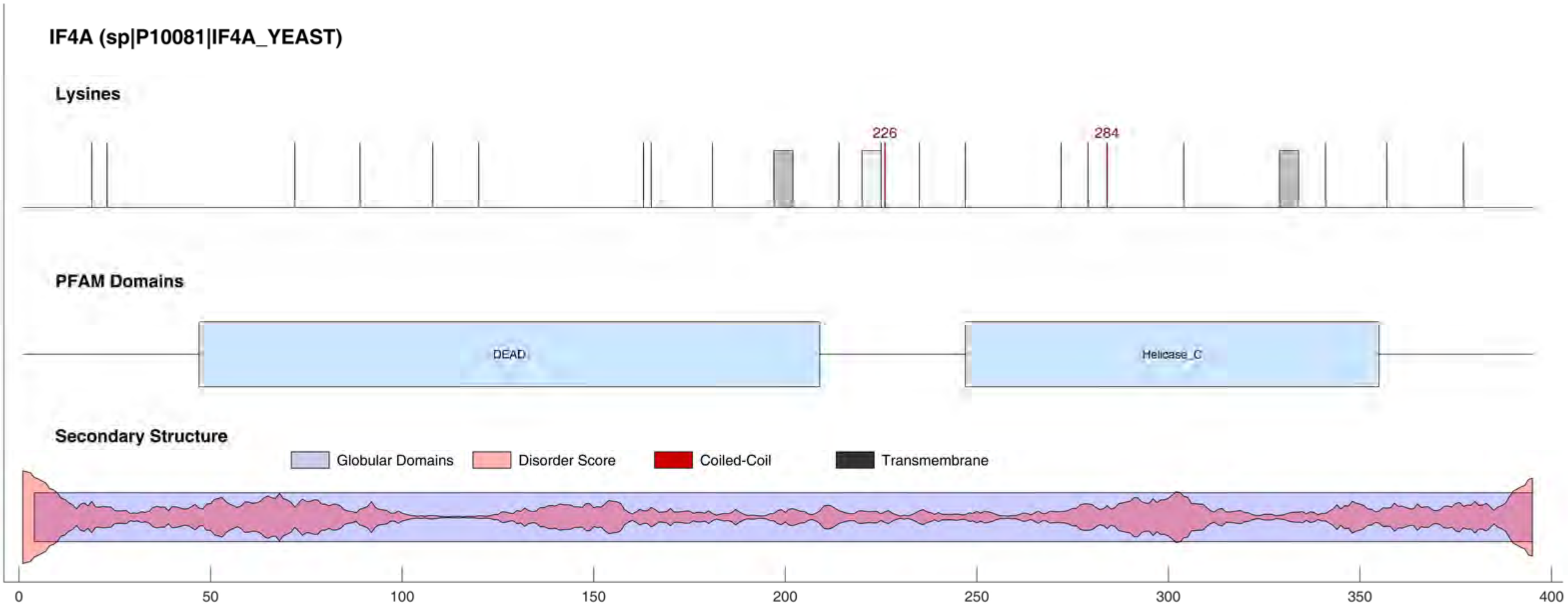
IF4A (sp|P10081|IF4A_YEAST)
Lysines
226
284
PFAM Domains
DEAD
Helicase_C
Secondary Structure
Globular Domains
Disorder Score
Coiled-Coil
Transmembrane
0
50
100
150
200
250
300
350
400

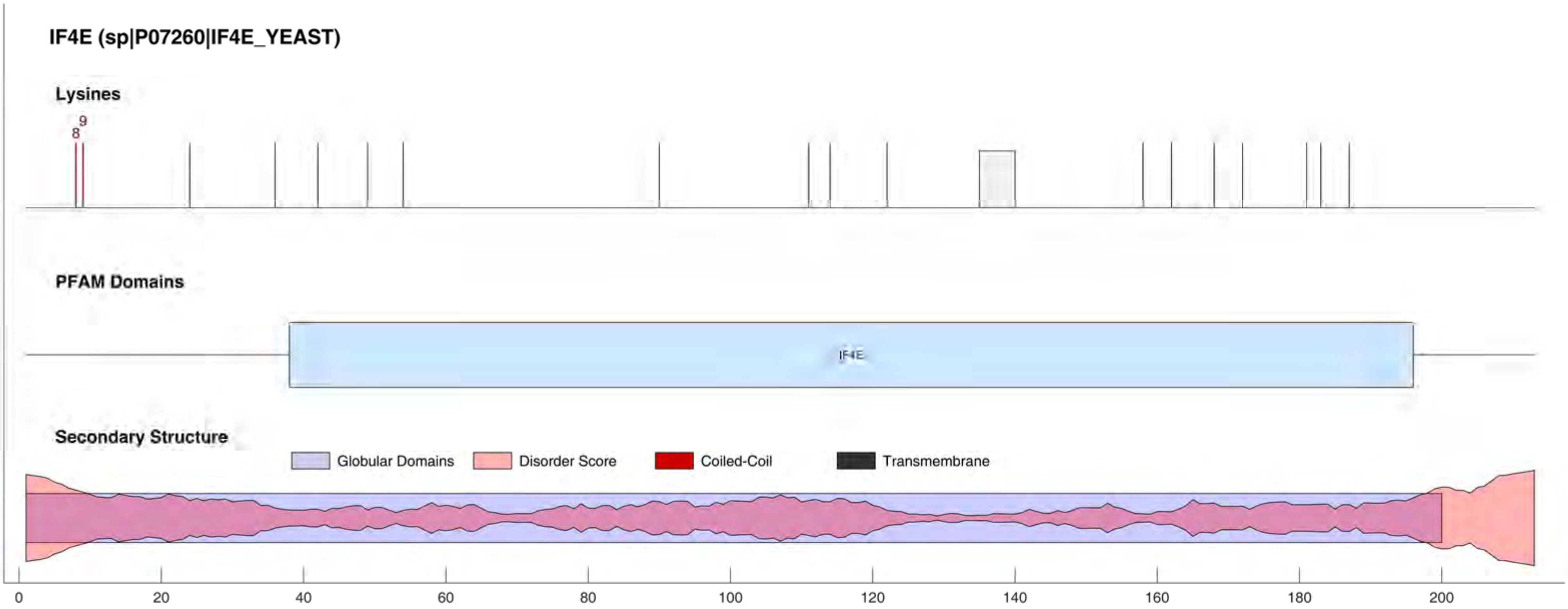

IF4E (sp|P07260|IF4E_YEAST)
Lysines
8
9
PFAM Domains
IF4E
Secondary Structure
Globular Domains
Disorder Score
Coiled-Coil
Transmembrane
0
20
40
60
80
100
120
140
160
180
200

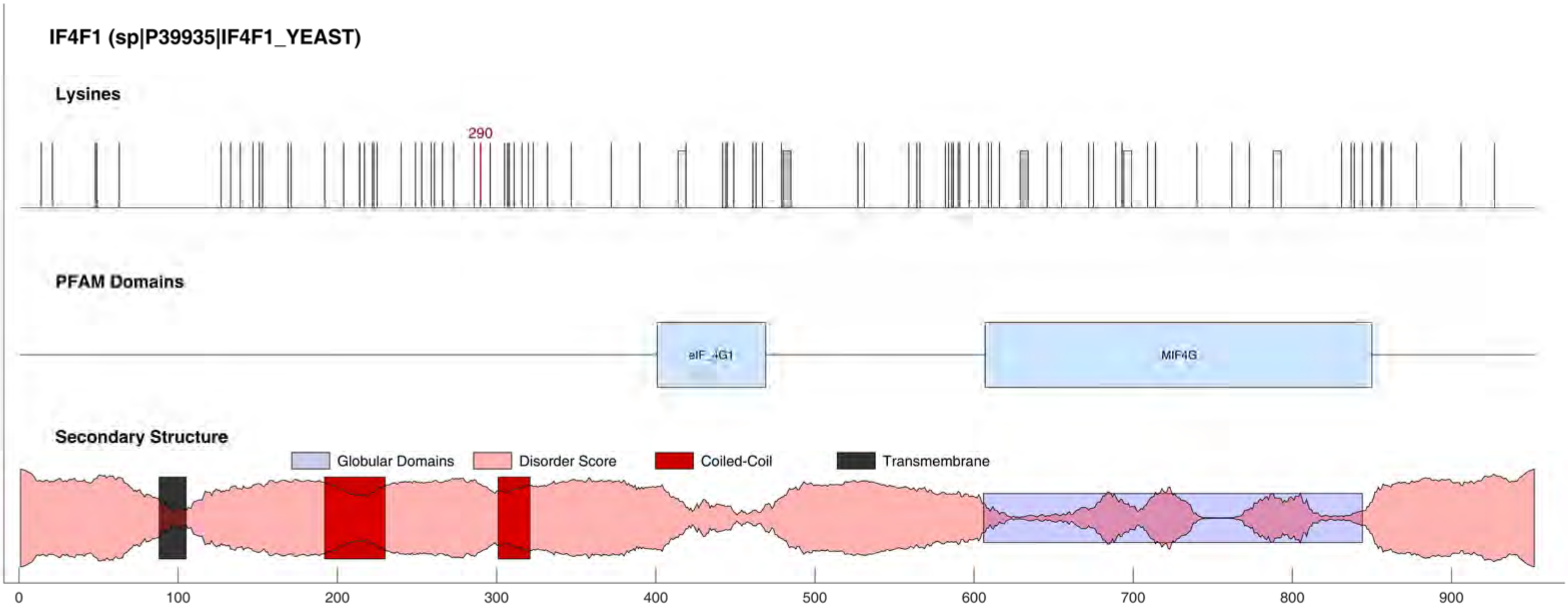
IF4F1 (sp|P39935|IF4F1_YEAST)
Lysines
290
PFAM Domains
eIF_4G1
MIF4G
Secondary Structure
Globular Domains
Disorder Score
Coiled-Coil
Transmembrane
0
100
200
300
400
500
600
700
800
900

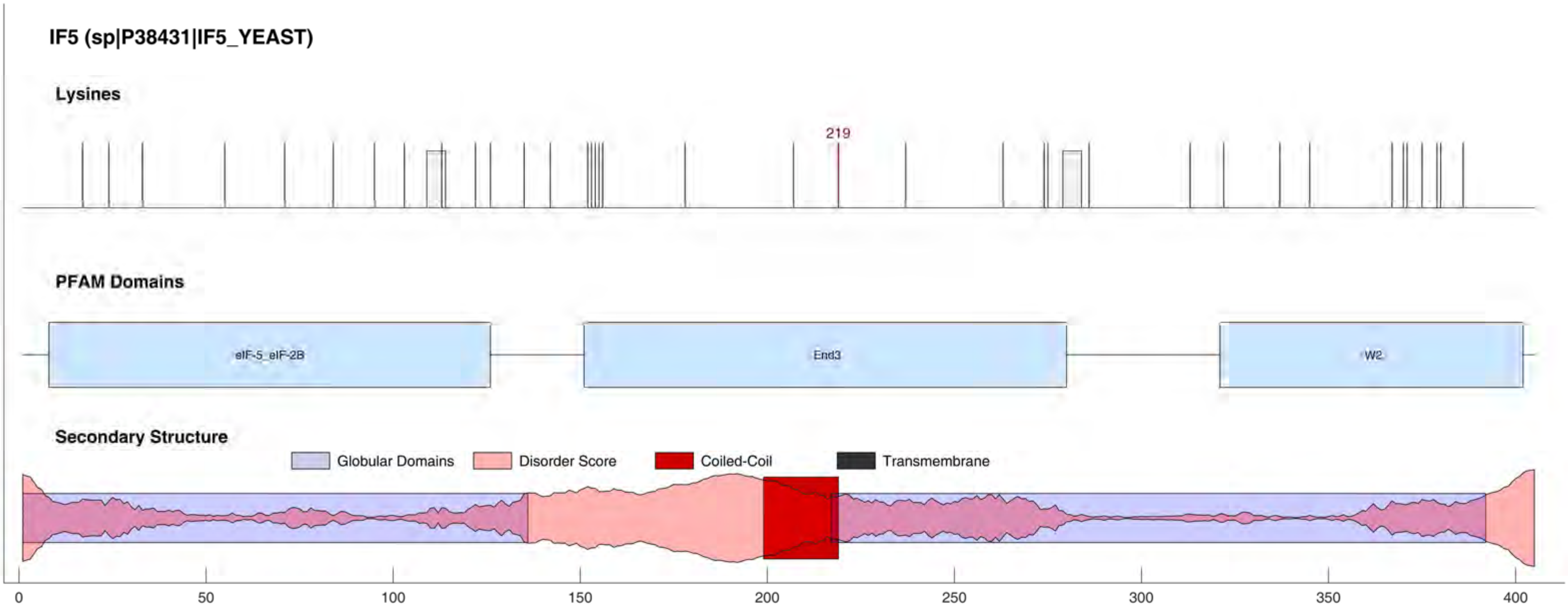
IF5 (sp|P38431|IF5_YEAST)
Lysines
219
PFAM Domains
eIF-5_eIF-2B
End3
W2
Secondary Structure
Globular Domains
Disorder Score
Coiled-Coil
Transmembrane
0
50
100
150
200
250
300
350
400

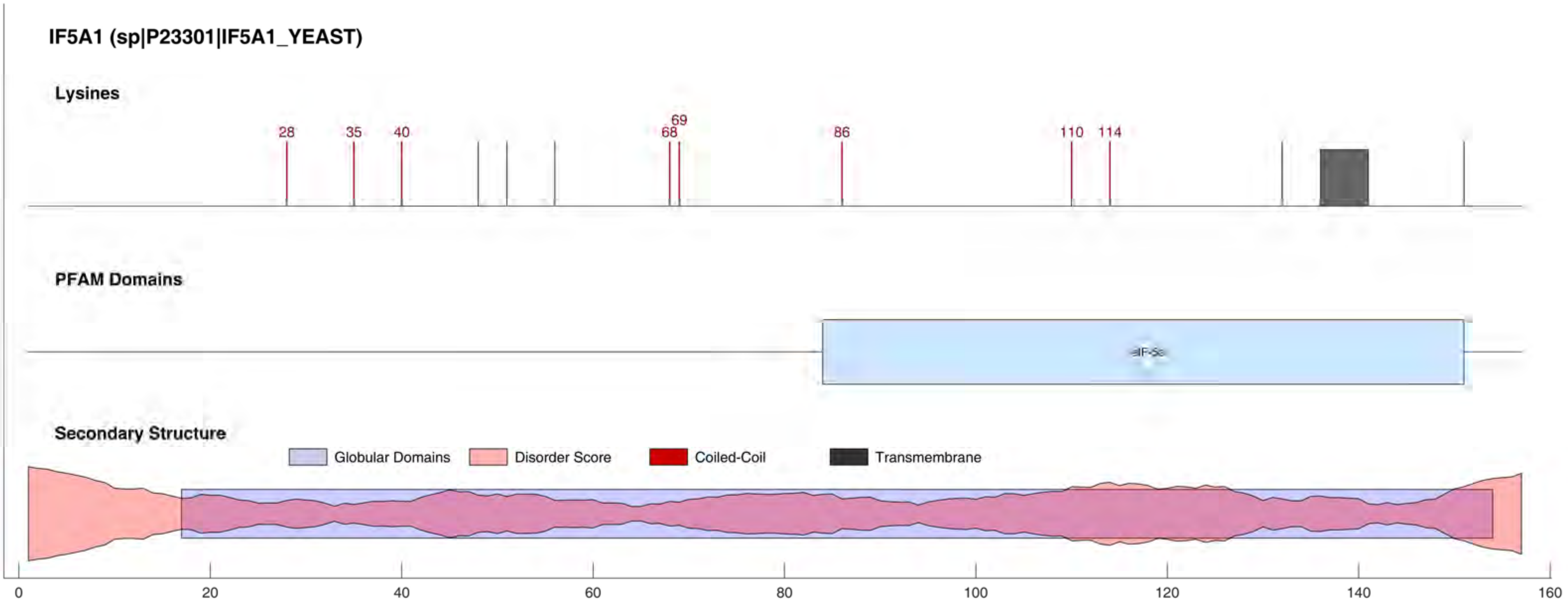
IF5A1 (sp|P23301|IF5A1_YEAST)
Lysines
28
35
40
68
69
86
110
114
PFAM Domains
eIF-5a
Secondary Structure
Globular Domains
Disorder Score
Coiled-Coil
Transmembrane
0
20
40
60
80
100
120
140
160

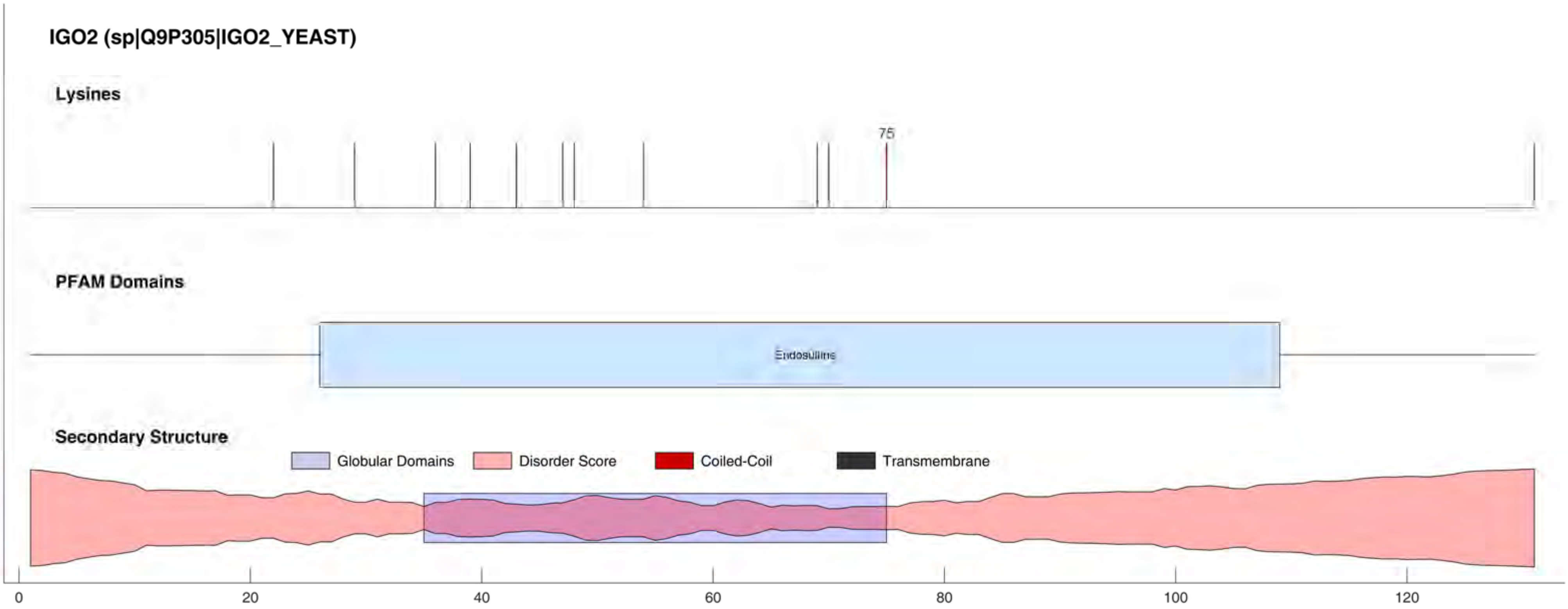
IGO2 (sp|Q9P305|IGO2_YEAST)
Lysines
75
PFAM Domains
Endosulfine
Secondary Structure
Globular Domains
Disorder Score
Coiled-Coil
Transmembrane
0
20
40
60
80
100
120

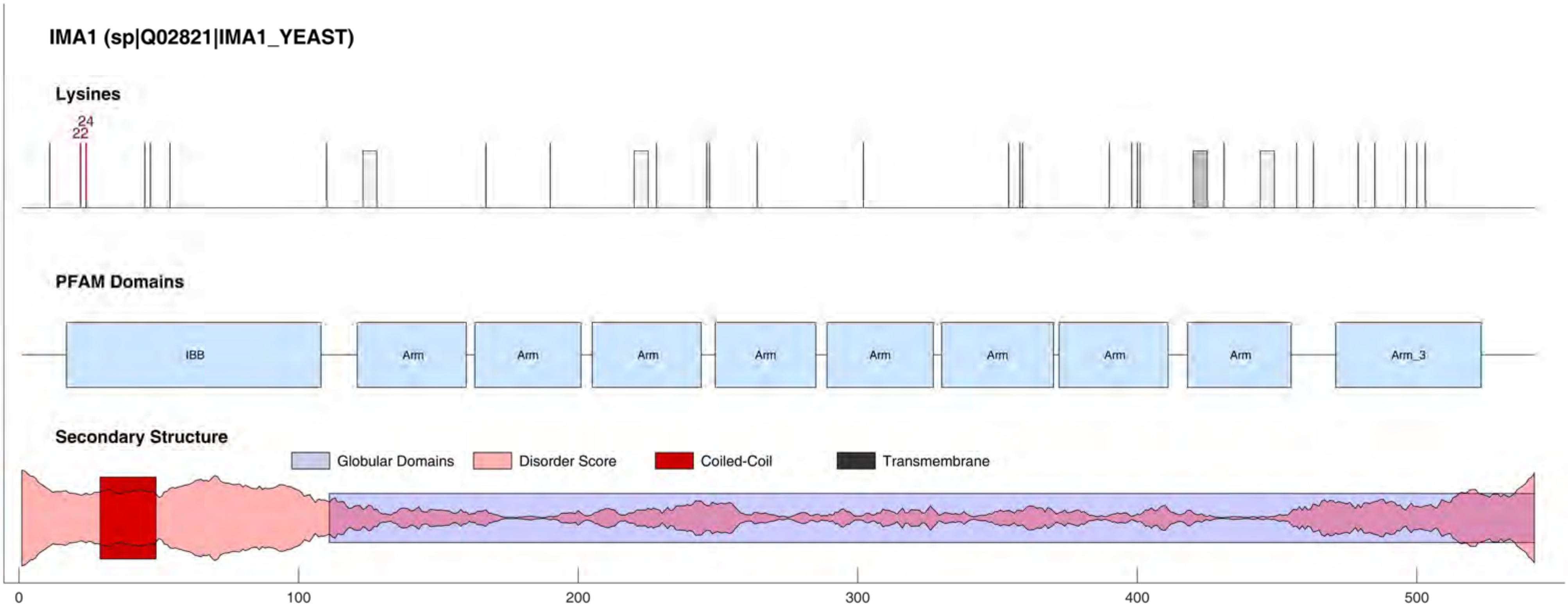
IMA1 (sp|Q02821|IMA1_YEAST)
Lysines
24
22
PFAM Domains
IBB
Arm
Arm
Arm
Arm
Arm
Arm
Arm
Arm
Arm_3
Secondary Structure
Globular Domains
Disorder Score
Coiled-Coil
Transmembrane
0
100
200
300
400
500

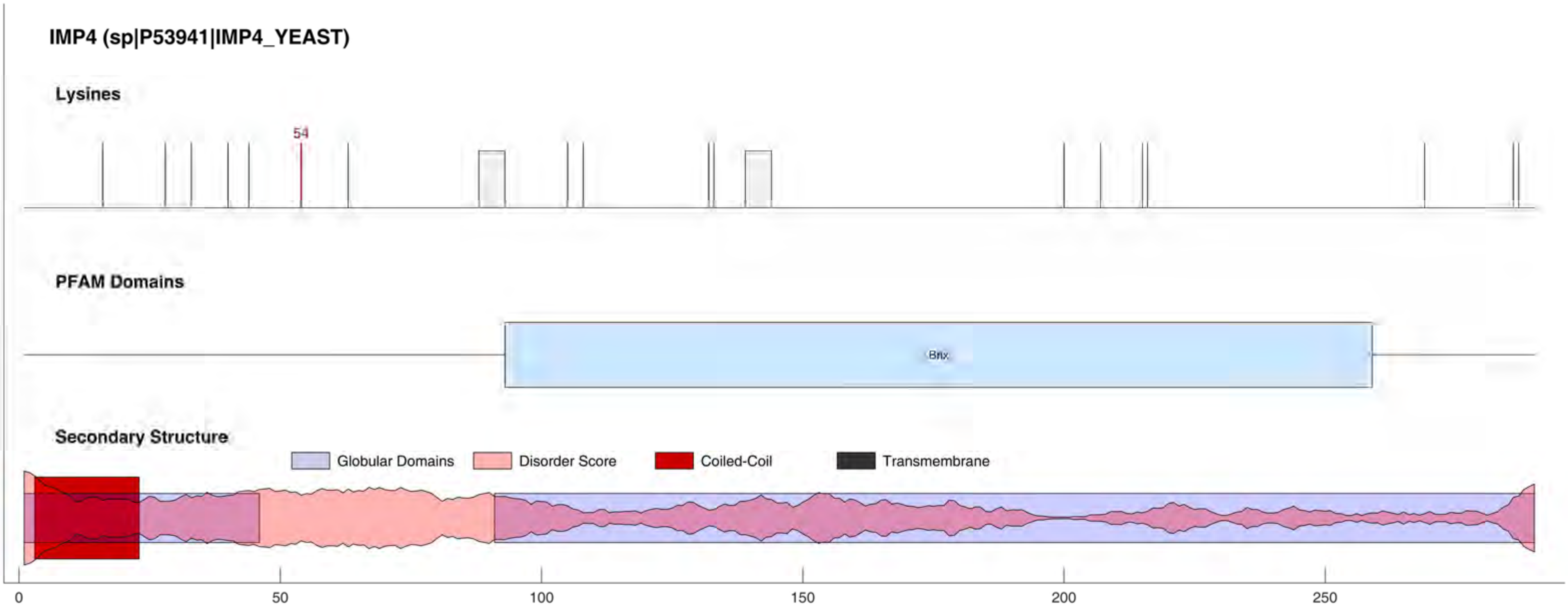
IMP4 (sp|P53941|IMP4_YEAST)
Lysines
54
PFAM Domains
Brix
Secondary Structure
Globular Domains
Disorder Score
Coiled-Coil
Transmembrane
0
50
100
150
200
250

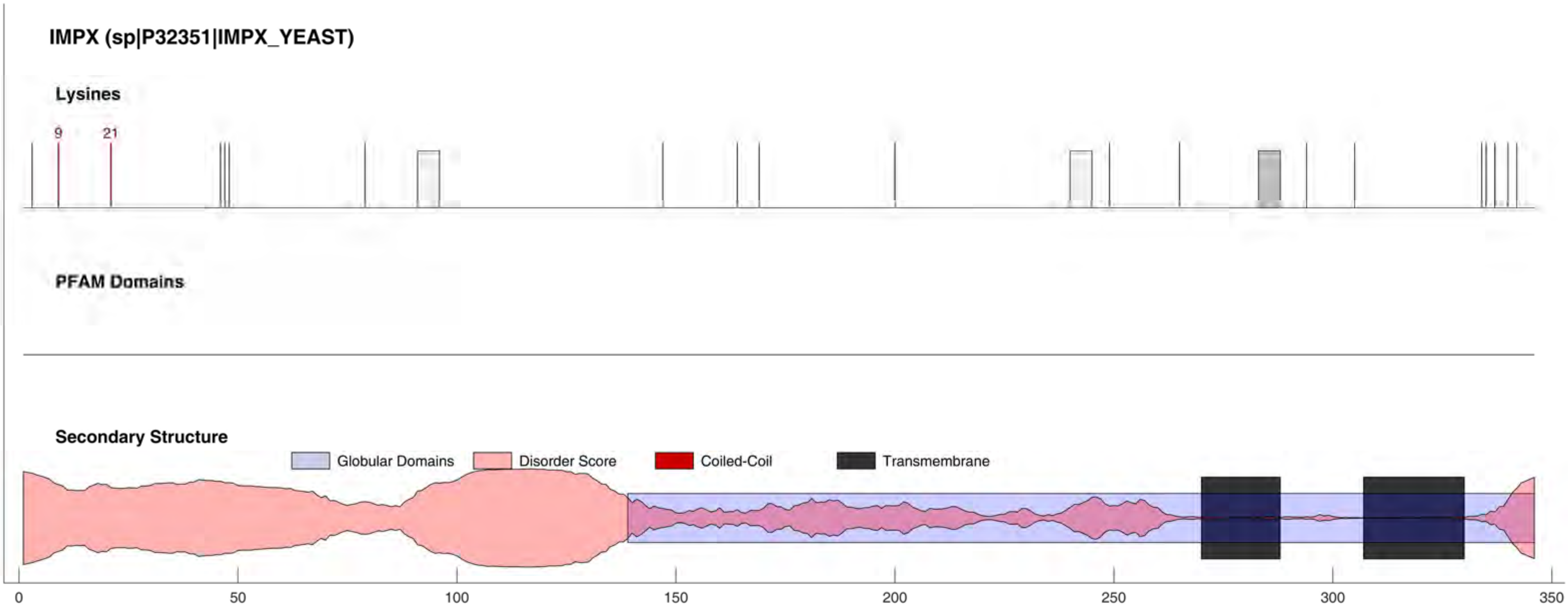
IMPX (sp|P32351|IMPX_YEAST)
Lysines
9
21
PFAM Domains
Secondary Structure
Globular Domains
Disorder Score
Coiled-Coil
Transmembrane
0
50
100
150
200
250
300
350

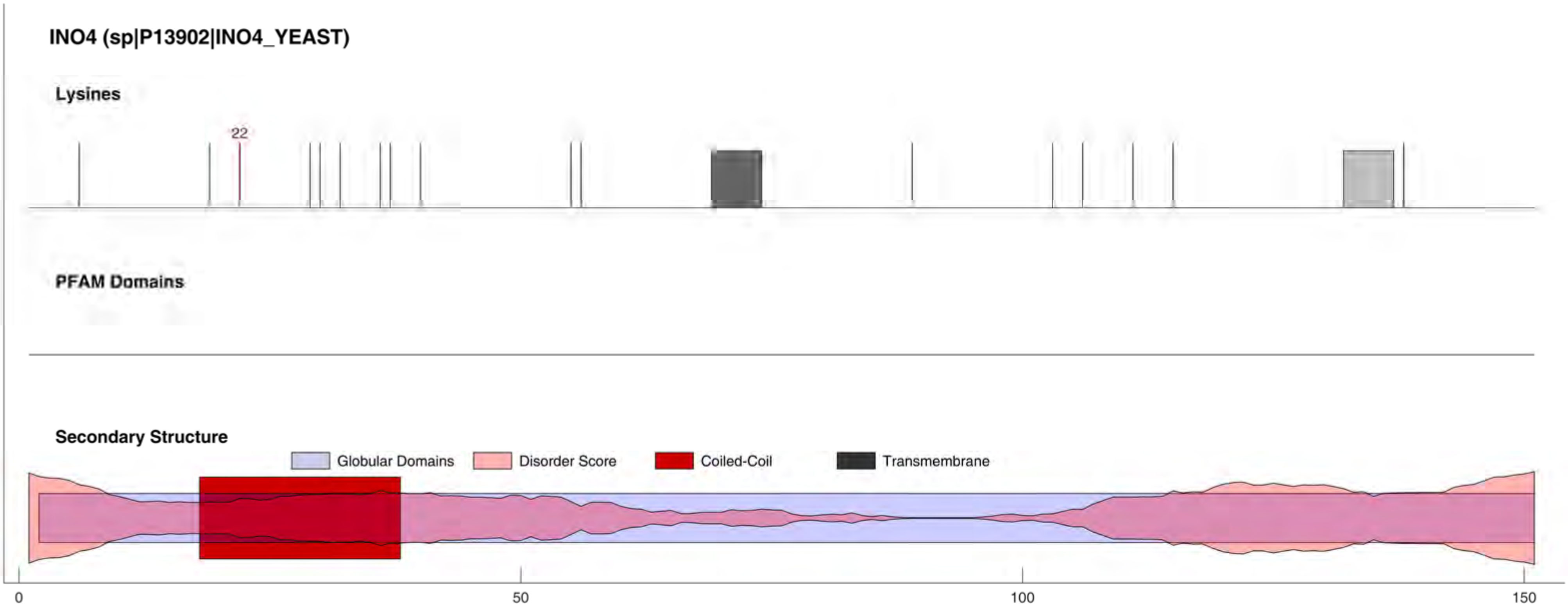
INO4 (sp|P13902|INO4_YEAST)
Lysines
22
PFAM Domains
Secondary Structure
Globular Domains
Disorder Score
Coiled-Coil
Transmembrane
0
50
100
150

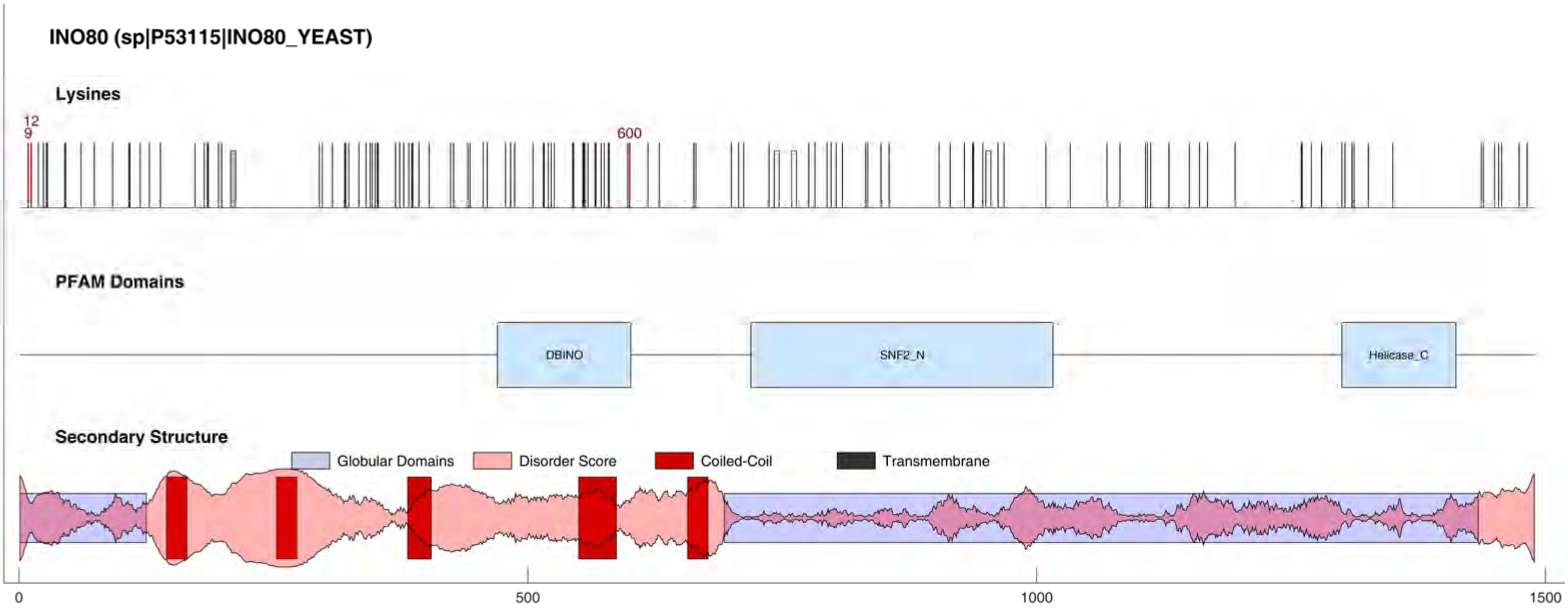

INO80 (sp|P53115|INO80_YEAST)
Lysines
12
9
600
PFAM Domains
DBINO
SNF2_N
Helicase_C
Secondary Structure
Globular Domains
Disorder Score
Coiled-Coil
Transmembrane
0
500
1000
1500

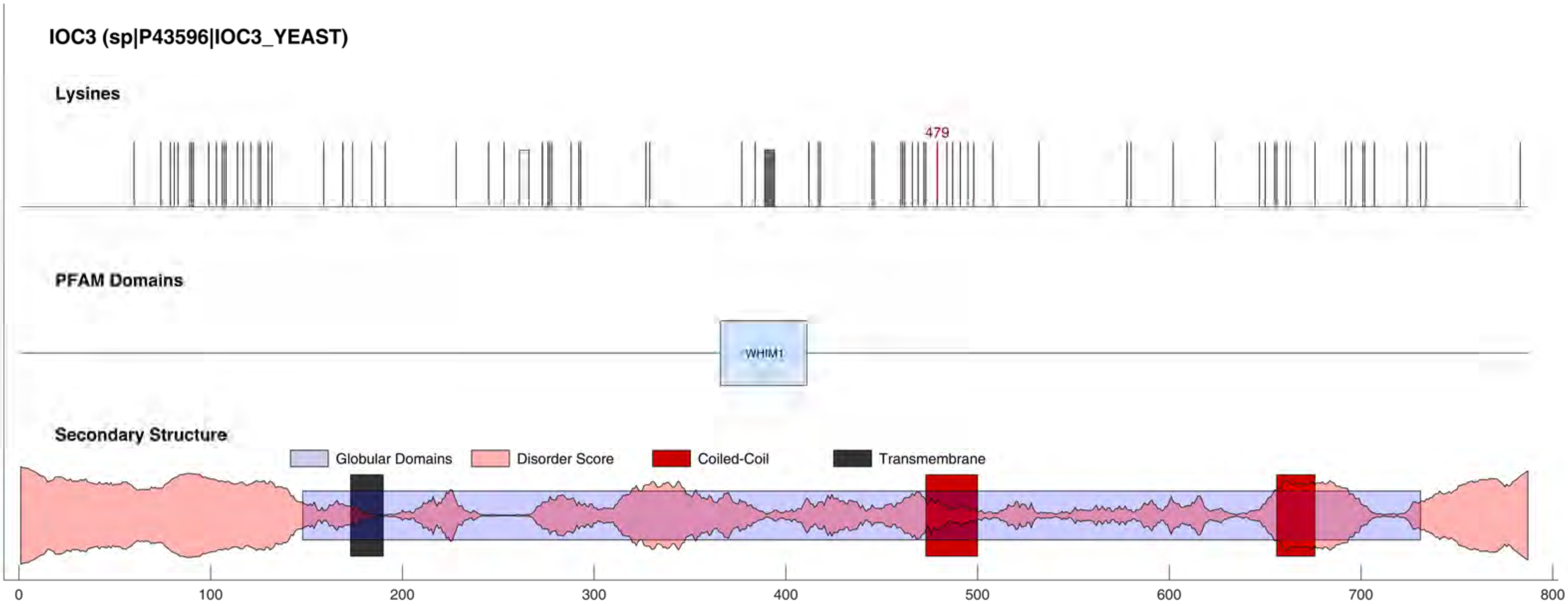
IOC3 (sp|P43596|IOC3_YEAST)
Lysines
479
PFAM Domains
WHIM1
Secondary Structure
Globular Domains
Disorder Score
Coiled-Coil
Transmembrane
0
100
200
300
400
500
600
700
800

#### IPA3 (sp|P01094|IPA3_YEAST)

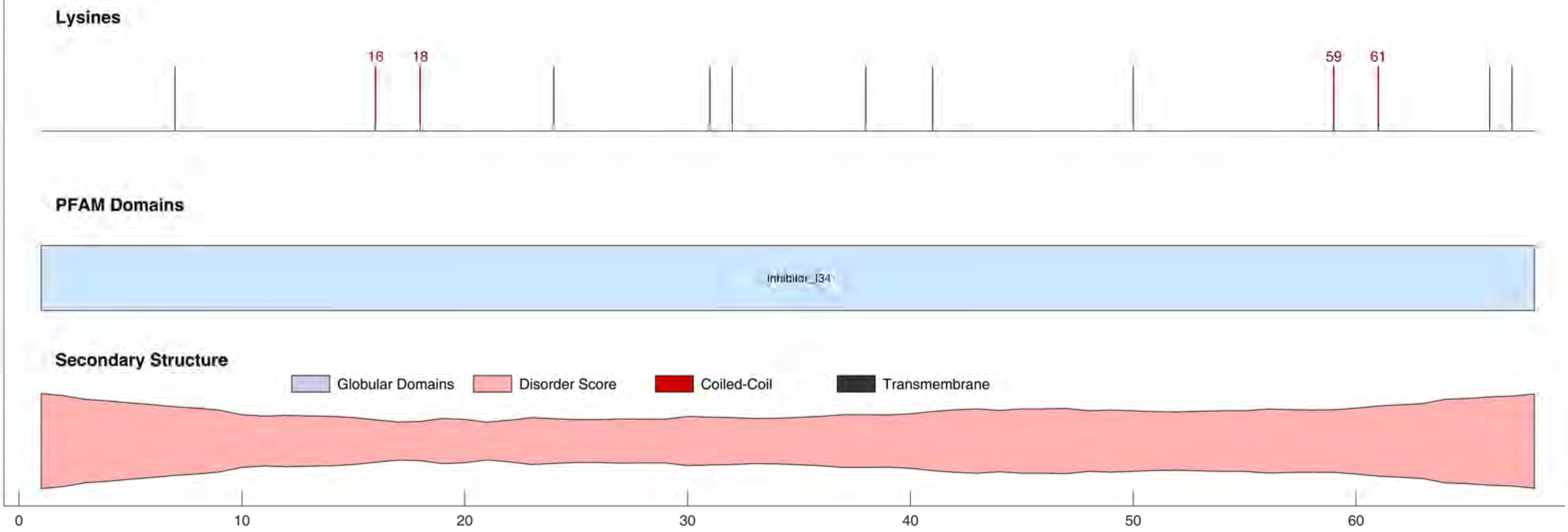

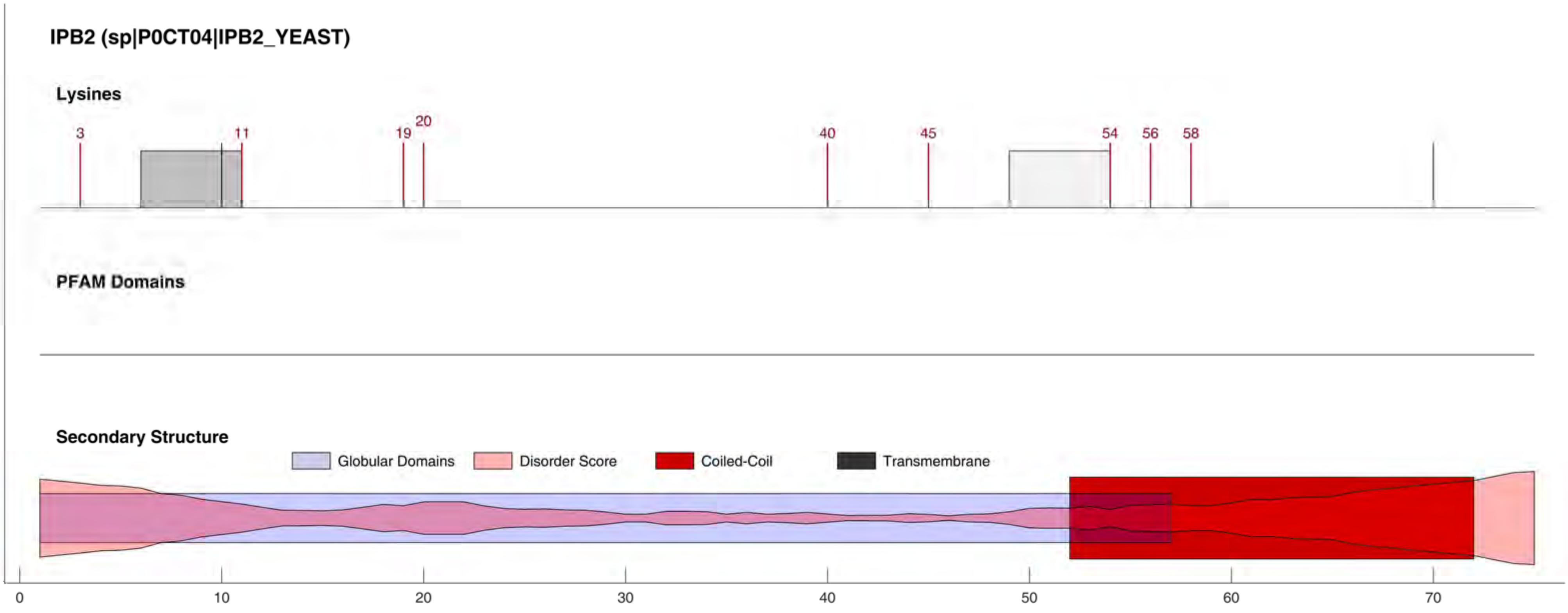

IPB2 (sp|P0CT04|IPB2_YEAST)
Lysines
3
11
19
20
40
45
54
56
58
PFAM Domains
Secondary Structure
Globular Domains
Disorder Score
Coiled-Coil
Transmembrane
0
10
20
30
40
50
60
70

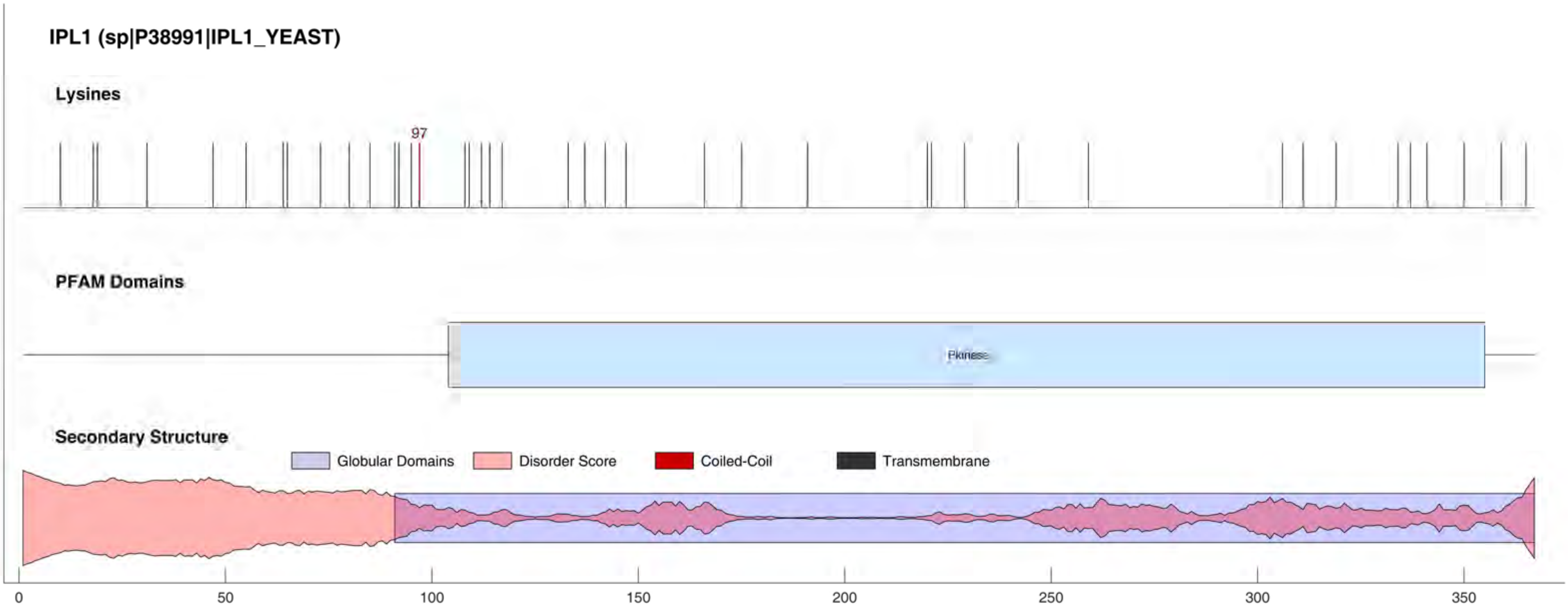
IPL1 (sp|P38991|IPL1_YEAST)
Lysines
97
PFAM Domains
Pkinase
Secondary Structure
Globular Domains
Disorder Score
Coiled-Coil
Transmembrane
0
50
100
150
200
250
300
350

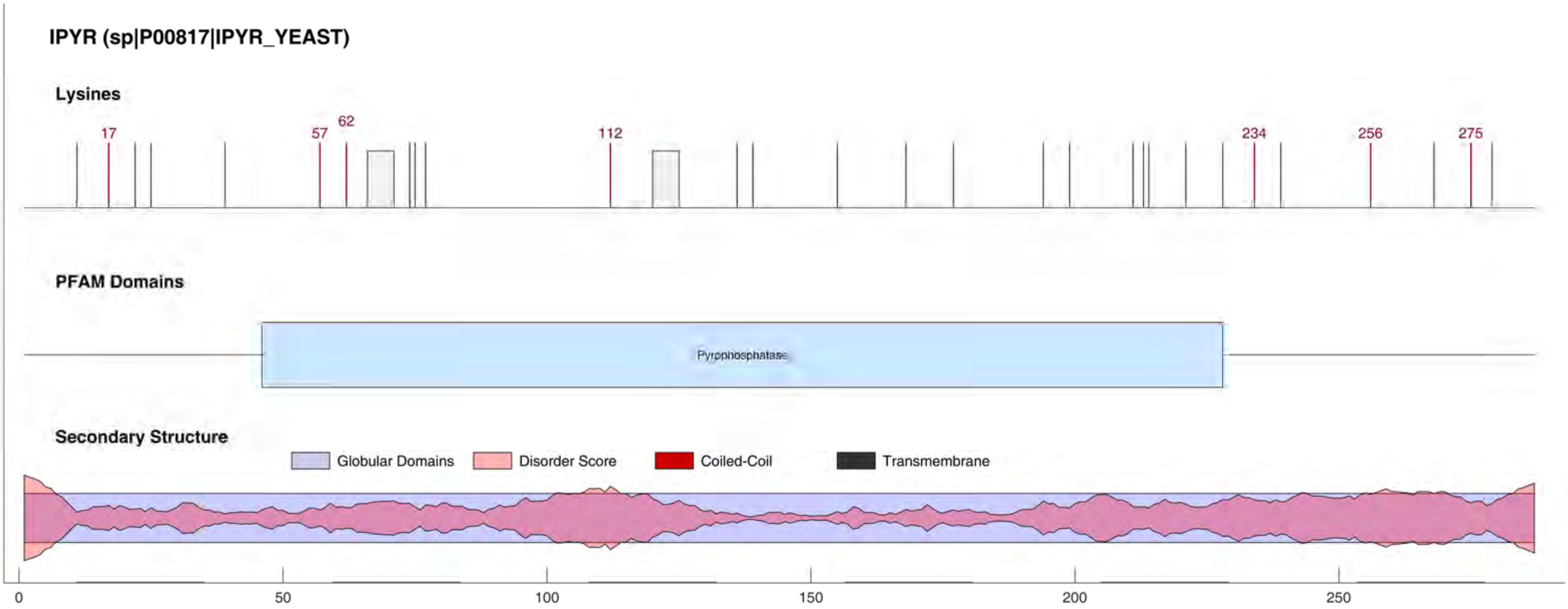

IPYR (sp|P00817|IPYR_YEAST)
Lysines
17
57
62
112
234
256
275
PFAM Domains
Pyrophosphatase
Secondary Structure
Globular Domains
Disorder Score
Coiled-Coil
Transmembrane
0
50
100
150
200
250

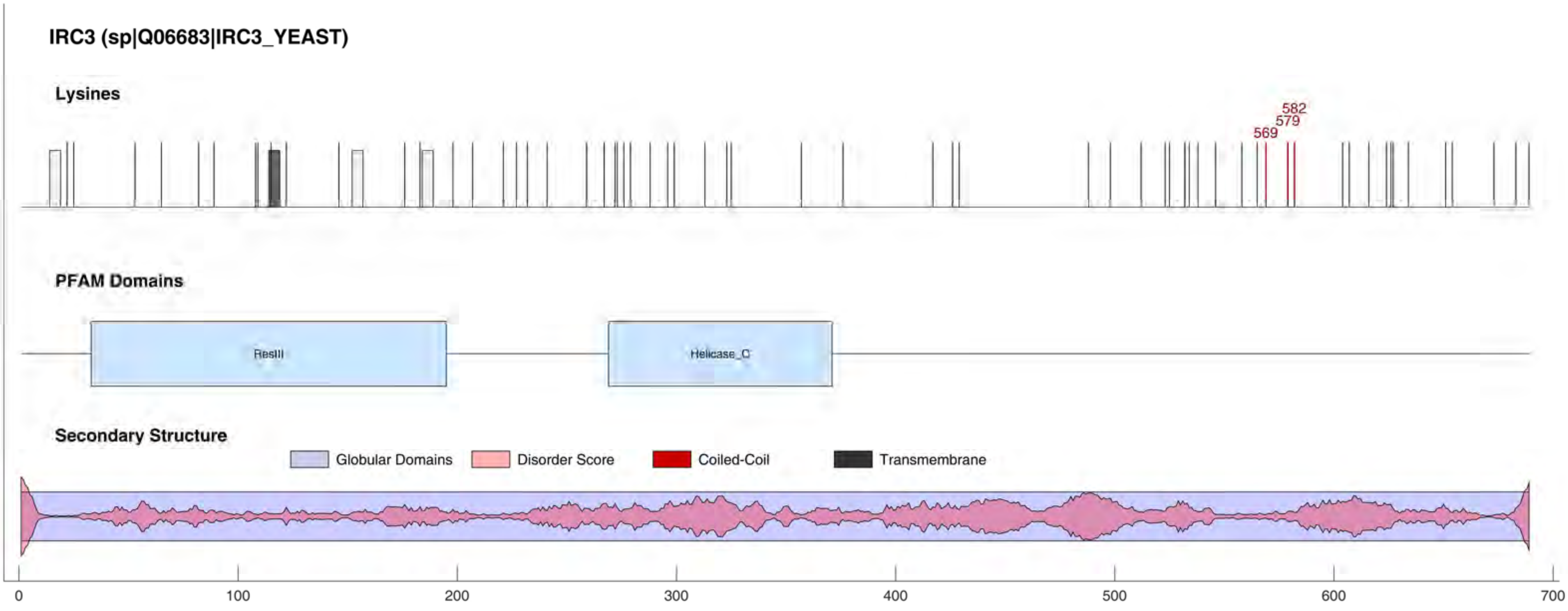

IRC3 (sp|Q06683|IRC3_YEAST)
Lysines
569
579
582
PFAM Domains
ResIII
Helicase_C
Secondary Structure
Globular Domains
Disorder Score
Coiled-Coil
Transmembrane
0
100
200
300
400
500
600
700

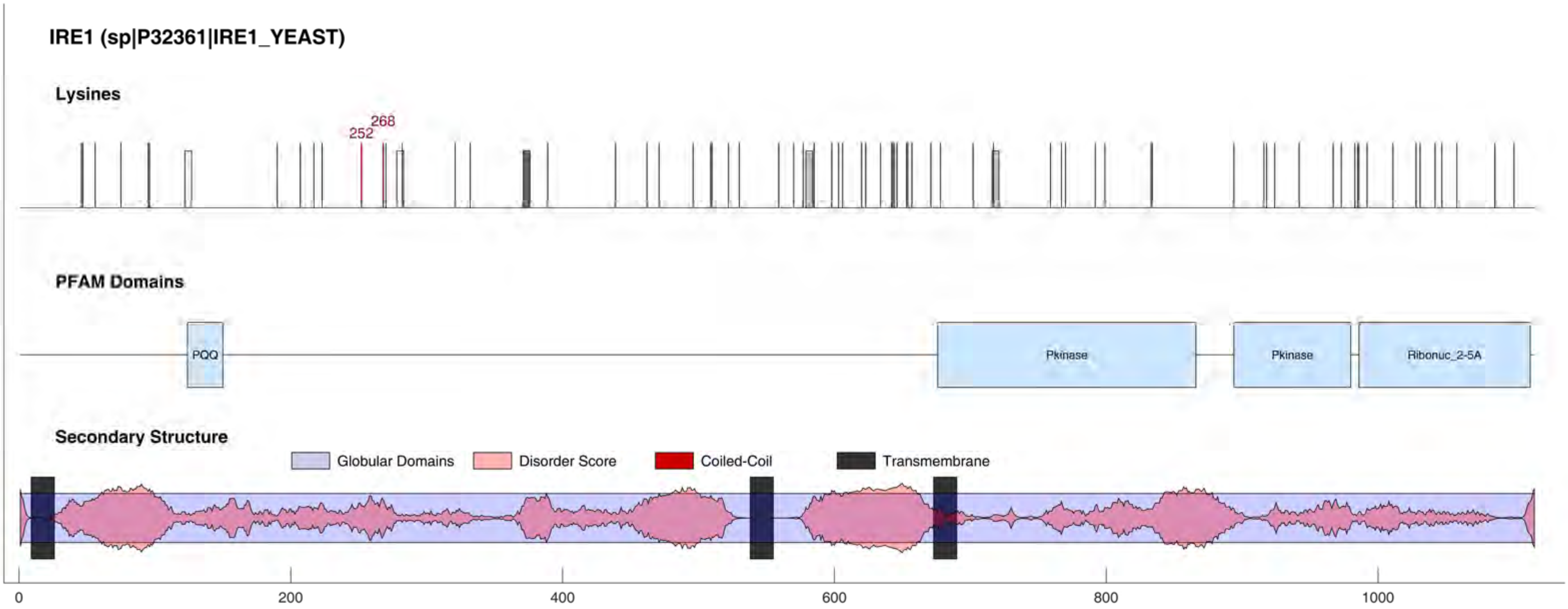
IRE1 (sp|P32361|IRE1_YEAST)
Lysines
252
268
PFAM Domains
PQQ
Pkinase
Pkinase
Ribonuc_2-5A
Secondary Structure
Globular Domains
Disorder Score
Coiled-Coil
Transmembrane
0
200
400
600
800
1000

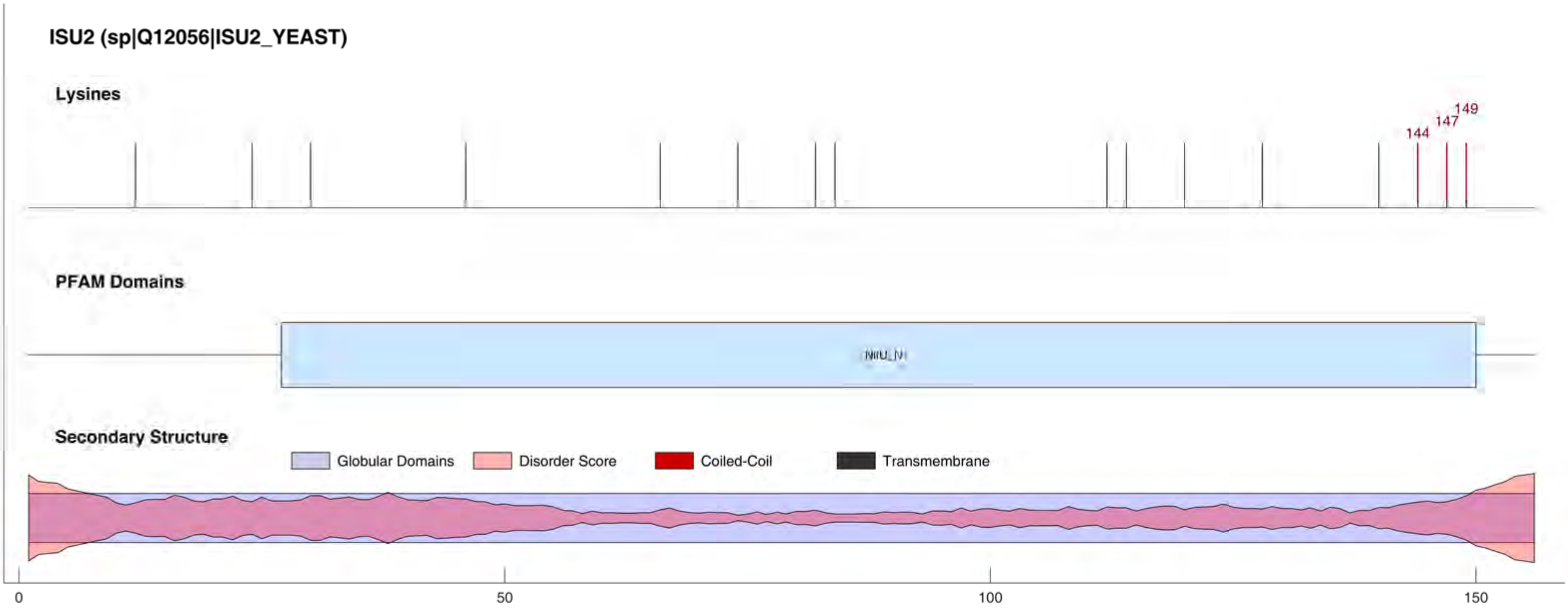

ISU2 (sp|Q12056|ISU2_YEAST)
Lysines
144
147
149
PFAM Domains
NifU_N
Secondary Structure
Globular Domains
Disorder Score
Coiled-Coil
Transmembrane
0
50
100
150

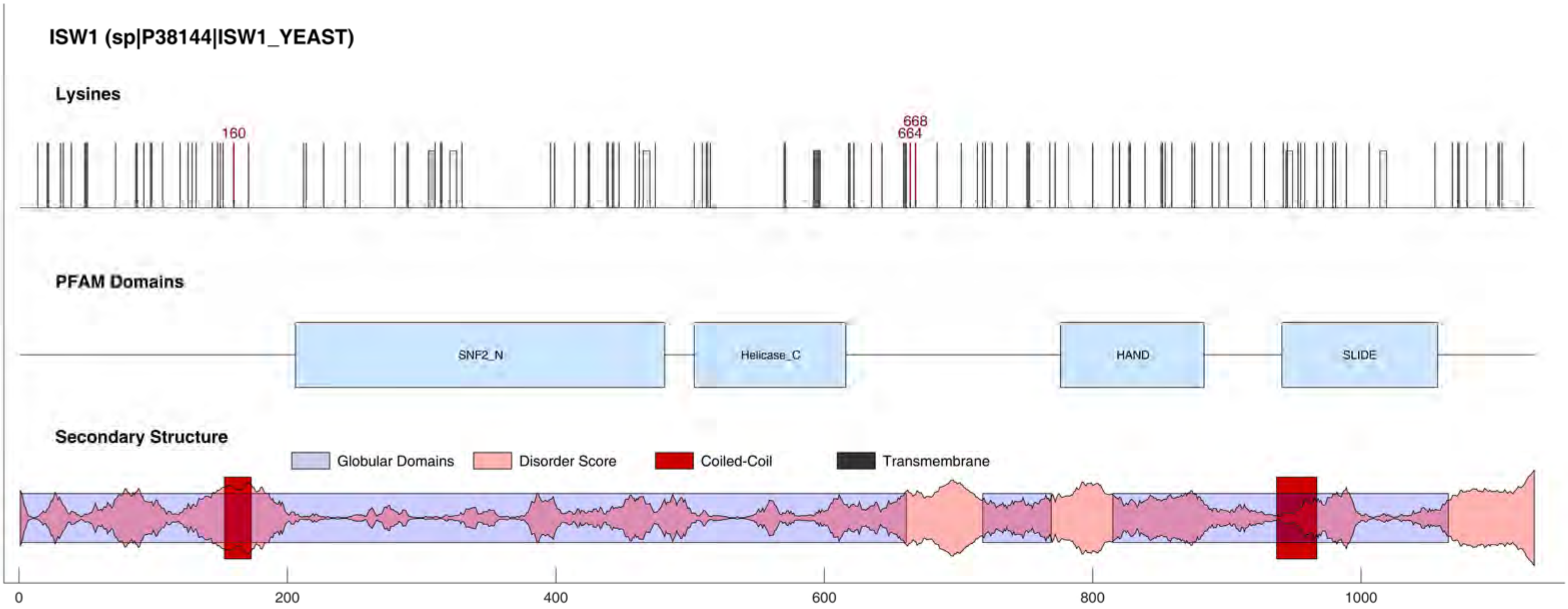

ISW1 (sp|P38144|ISW1_YEAST)
Lysines
160
668
664
PFAM Domains
SNF2_N
Helicase_C
HAND
SLIDE
Secondary Structure
Globular Domains
Disorder Score
Coiled-Coil
Transmembrane
0
200
400
600
800
1000

#### IWR1 (sp|Q07532|IWR1_YEAST)

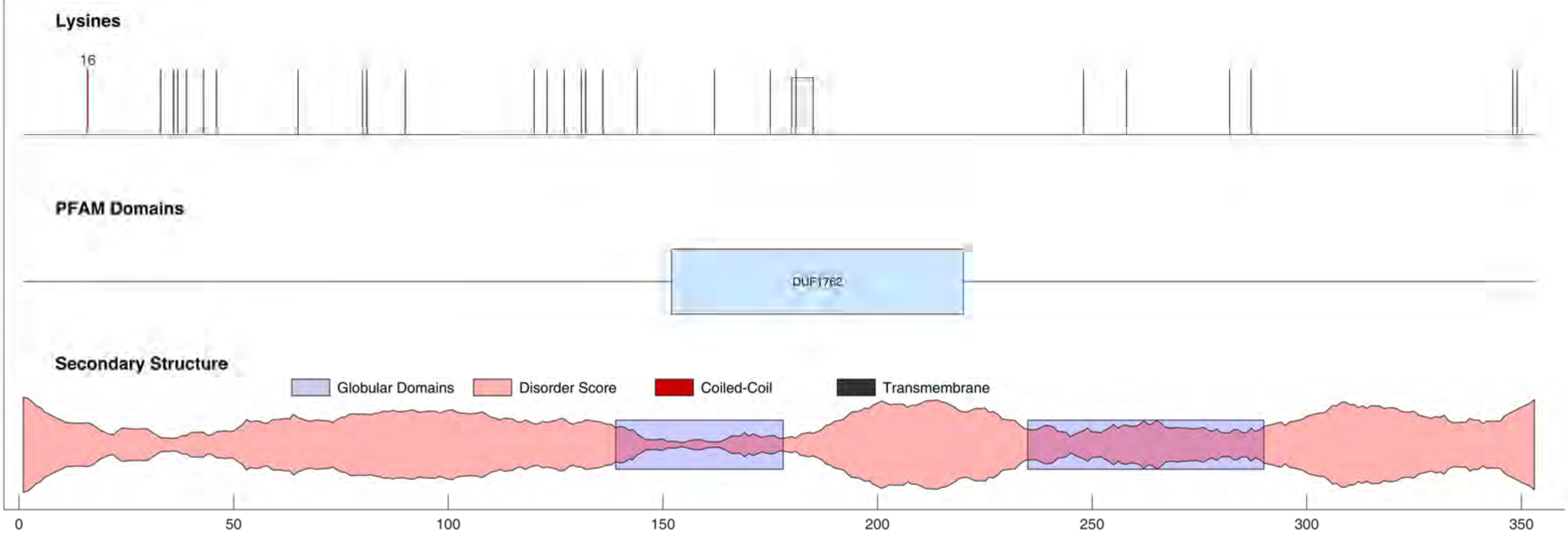

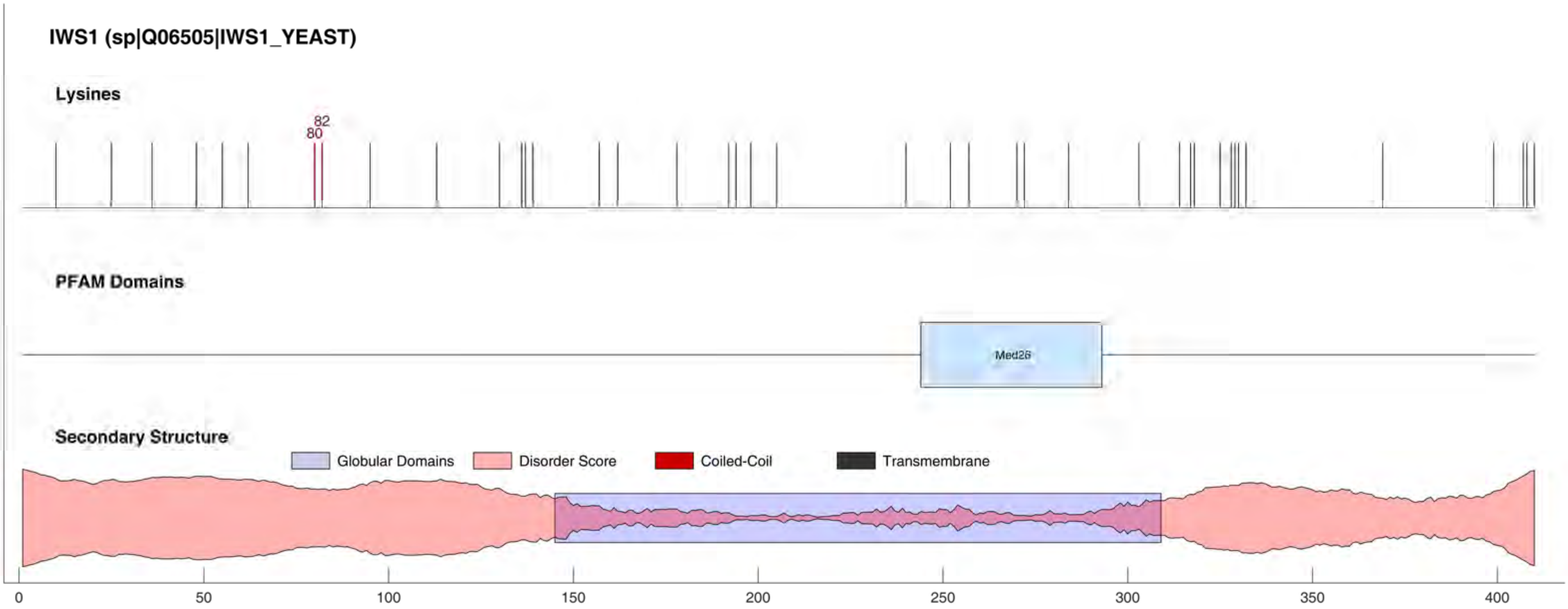

IWS1 (sp|Q06505|IWS1_YEAST)
Lysines
82
80
PFAM Domains
Med26
Secondary Structure
Globular Domains
Disorder Score
Coiled-Coil
Transmembrane
0
50
100
150
200
250
300
350
400

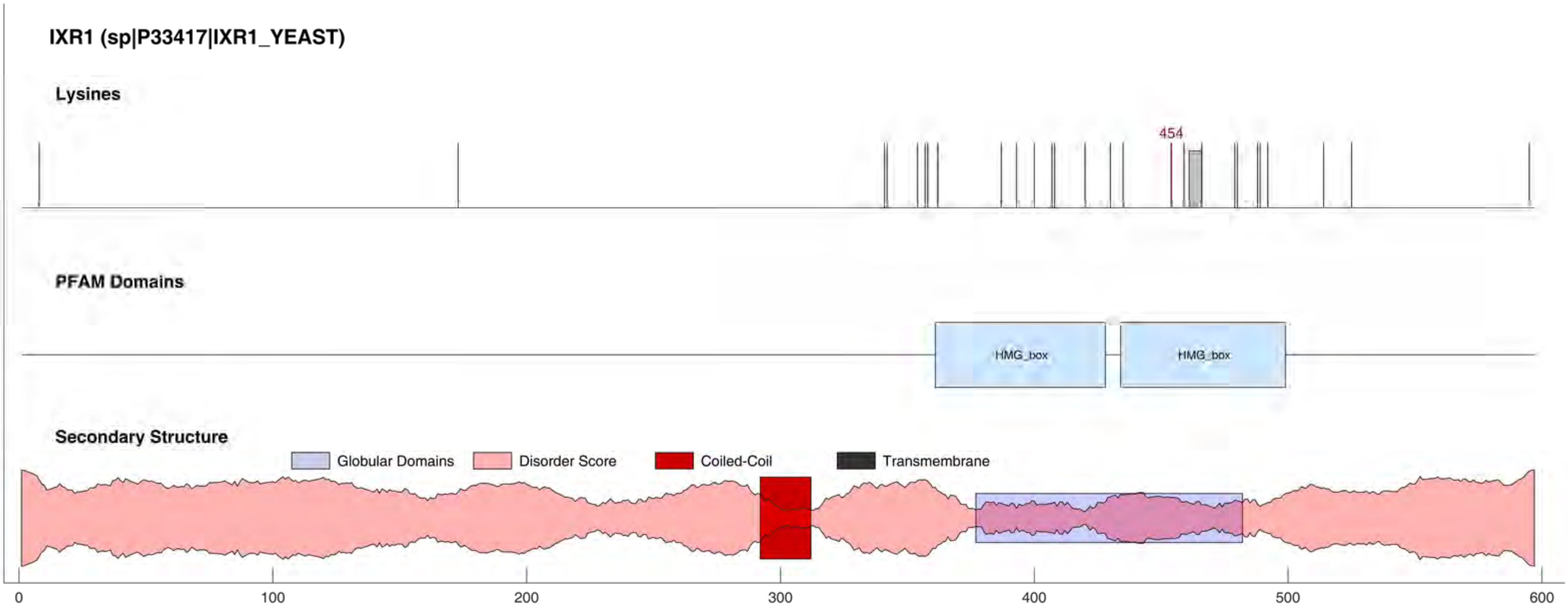

IXR1 (sp|P33417|IXR1_YEAST)
Lysines
454
PFAM Domains
HMG_box
HMG_box
Secondary Structure
Globular Domains
Disorder Score
Coiled-Coil
Transmembrane
0
100
200
300
400
500
600

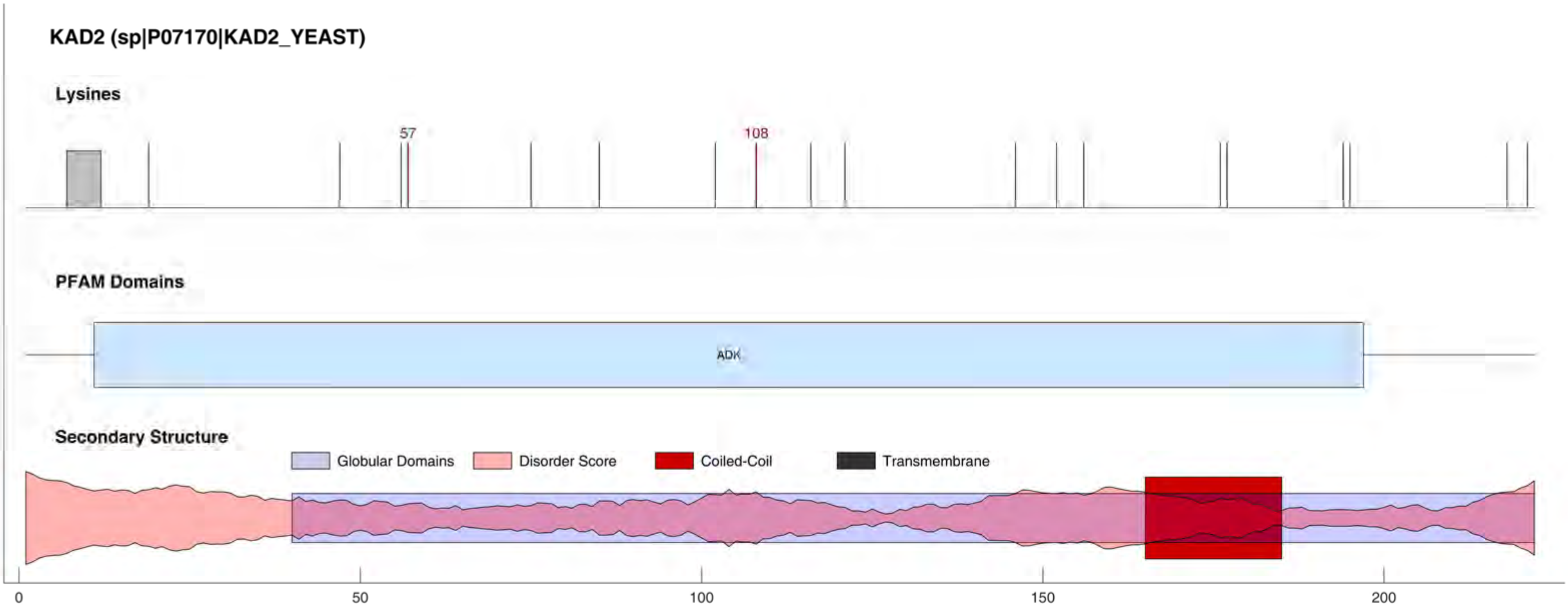
KAD2 (sp|P07170|KAD2_YEAST)
Lysines
57
108
PFAM Domains
ADK
Secondary Structure
Globular Domains
Disorder Score
Coiled-Coil
Transmembrane
0
50
100
150
200

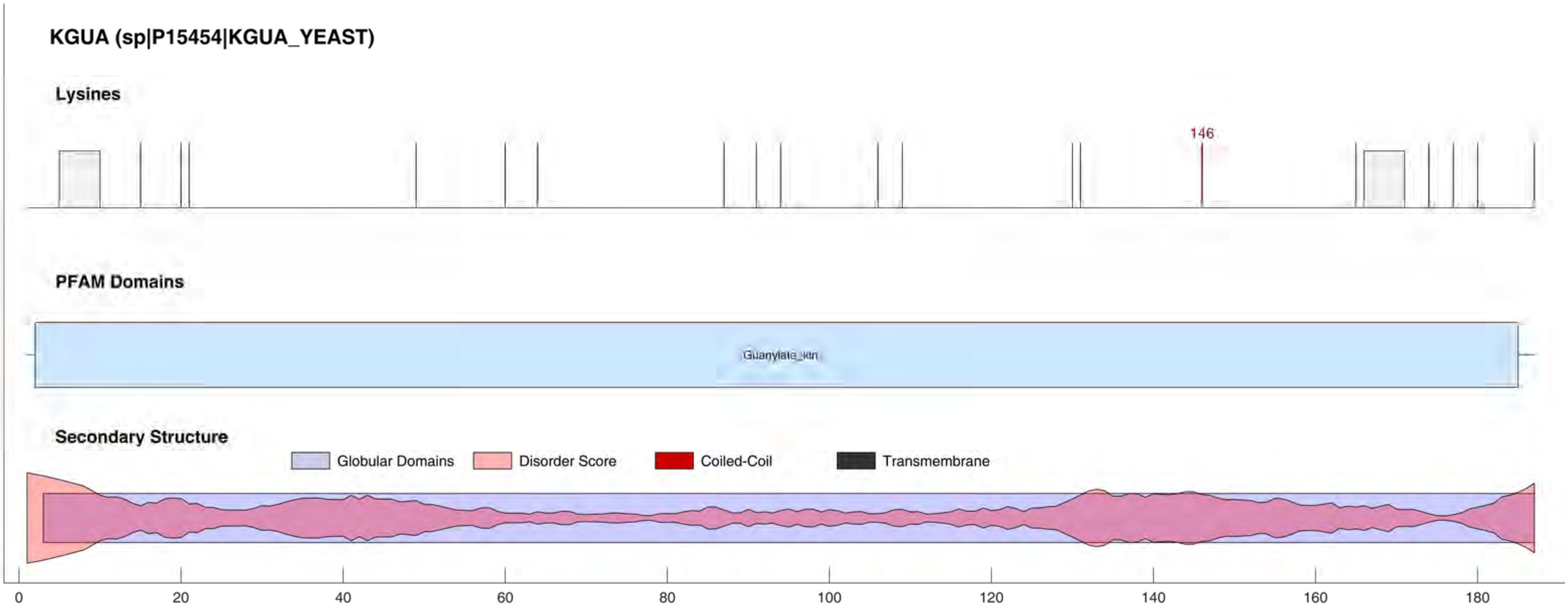

KGUA (sp|P15454|KGUA_YEAST)
Lysines
146
PFAM Domains
Guanylate_kin
Secondary Structure
Globular Domains
Disorder Score
Coiled-Coil
Transmembrane
0
20
40
60
80
100
120
140
160
180

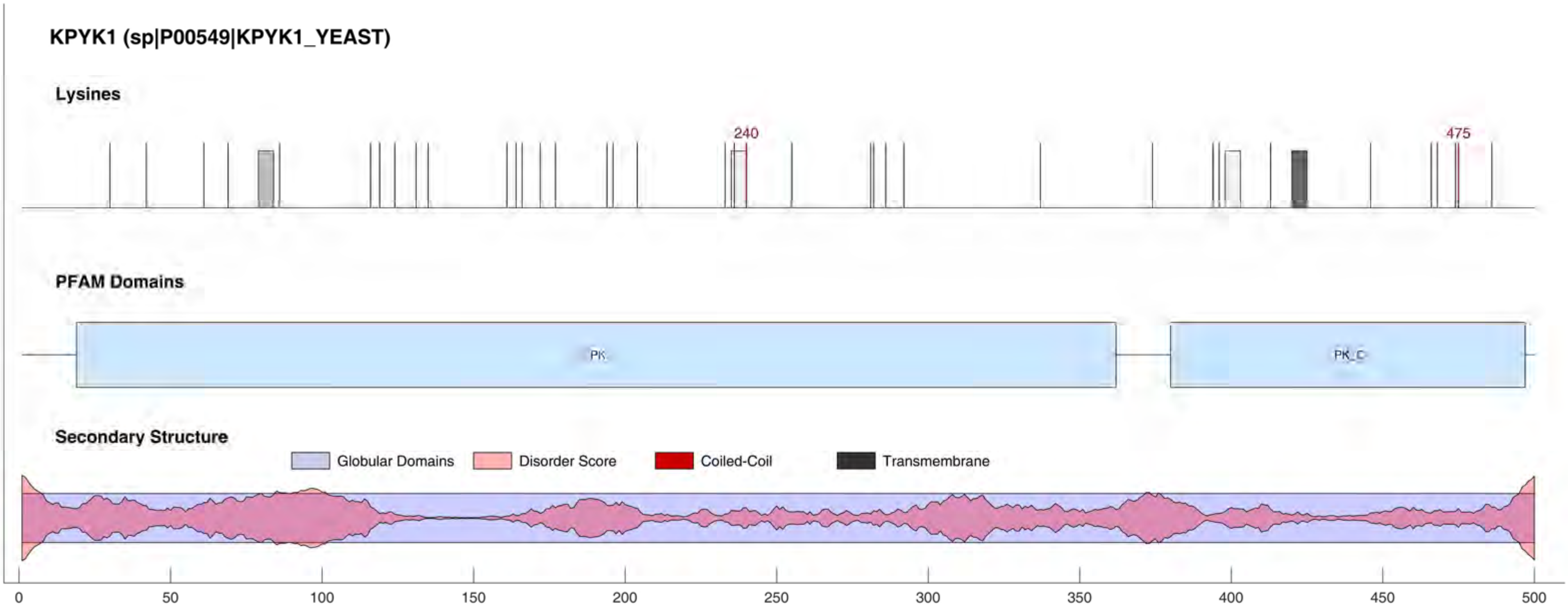

KPYK1 (sp|P00549|KPYK1_YEAST)
Lysines
240
475
PFAM Domains
PK
PK_C
Secondary Structure
Globular Domains
Disorder Score
Coiled-Coil
Transmembrane
0
50
100
150
200
250
300
350
400
450
500

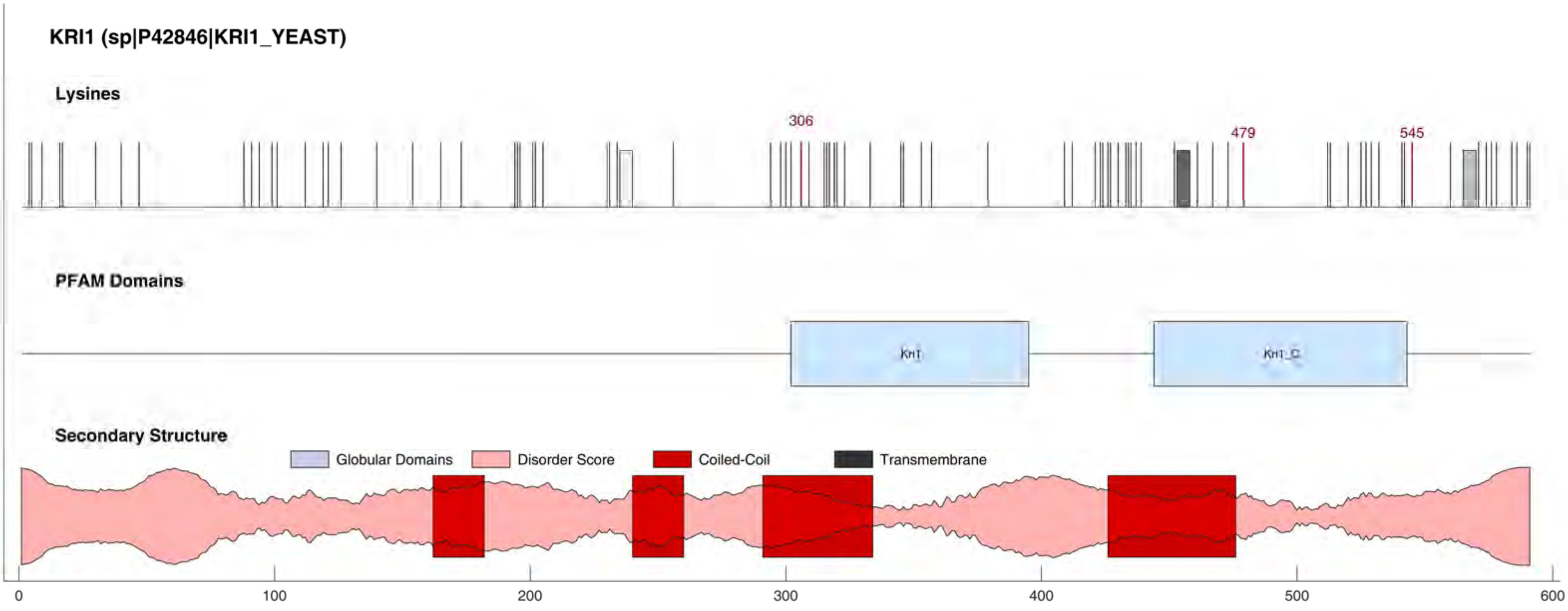
KRI1 (sp|P42846|KRI1_YEAST)
Lysines
306
479
545
PFAM Domains
Kri1
Kri1_C
Secondary Structure
Globular Domains
Disorder Score
Coiled-Coil
Transmembrane
0
100
200
300
400
500
600

#### KRR1 (sp|P25586|KRR1_YEAST)

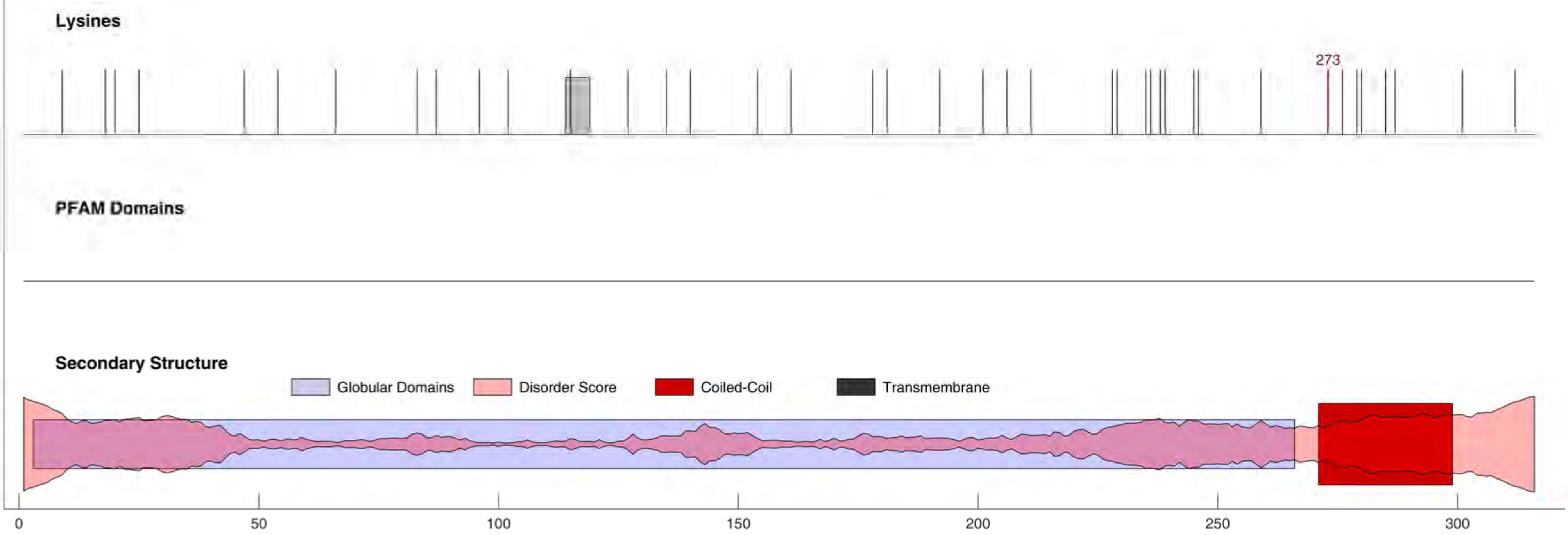

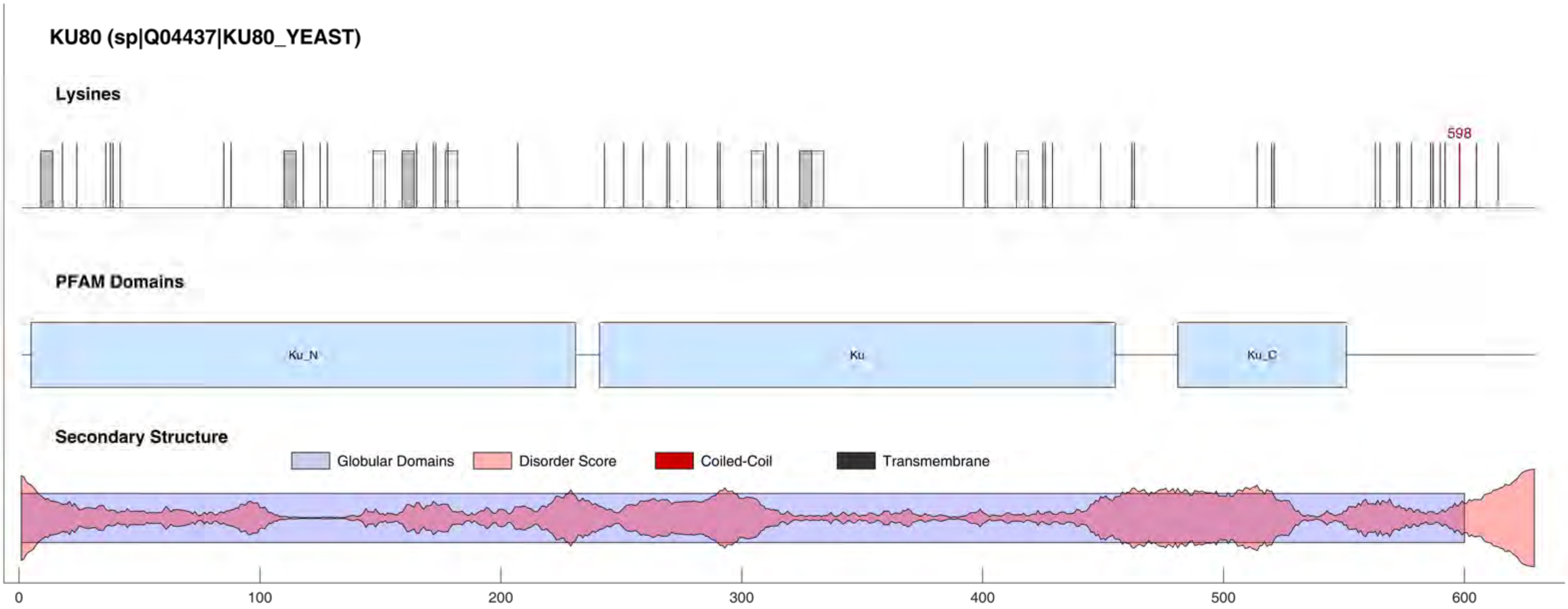
KU80 (sp|Q04437|KU80_YEAST)
Lysines
598
PFAM Domains
Ku_N
Ku
Ku_C
Secondary Structure
Globular Domains
Disorder Score
Coiled-Coil
Transmembrane
0
100
200
300
400
500
600

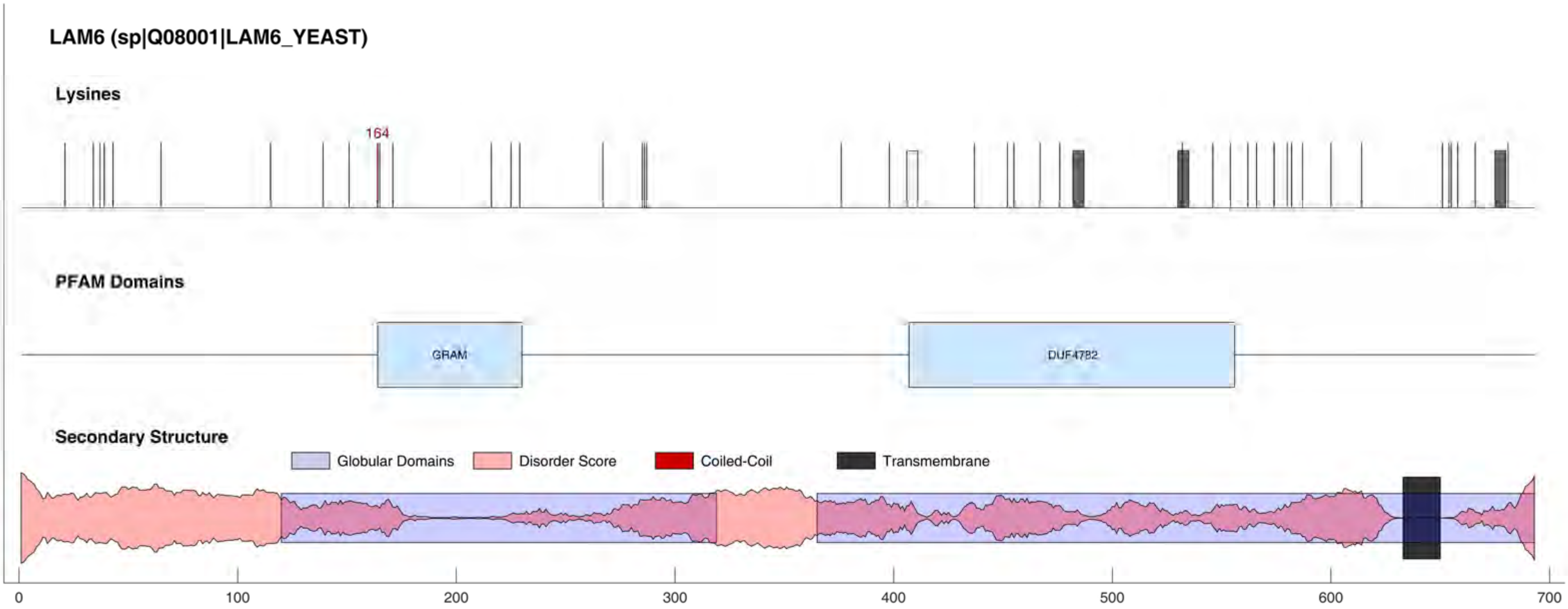
LAM6 (sp|Q08001|LAM6_YEAST)
Lysines
164
PFAM Domains
GRAM
DUF4782
Secondary Structure
Globular Domains
Disorder Score
Coiled-Coil
Transmembrane
0
100
200
300
400
500
600
700

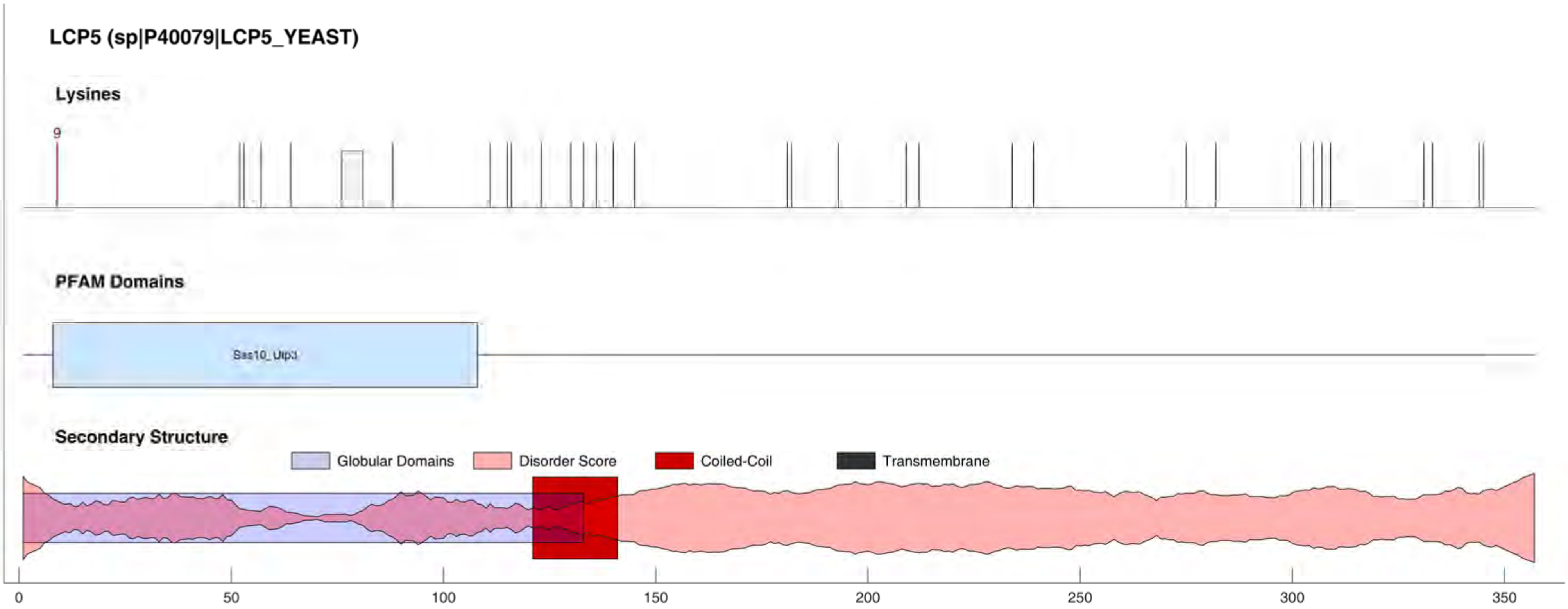

LCP5 (sp|P40079|LCP5_YEAST)
Lysines
9
PFAM Domains
Sas10_Utp3
Secondary Structure
Globular Domains
Disorder Score
Coiled-Coil
Transmembrane
0
50
100
150
200
250
300
350

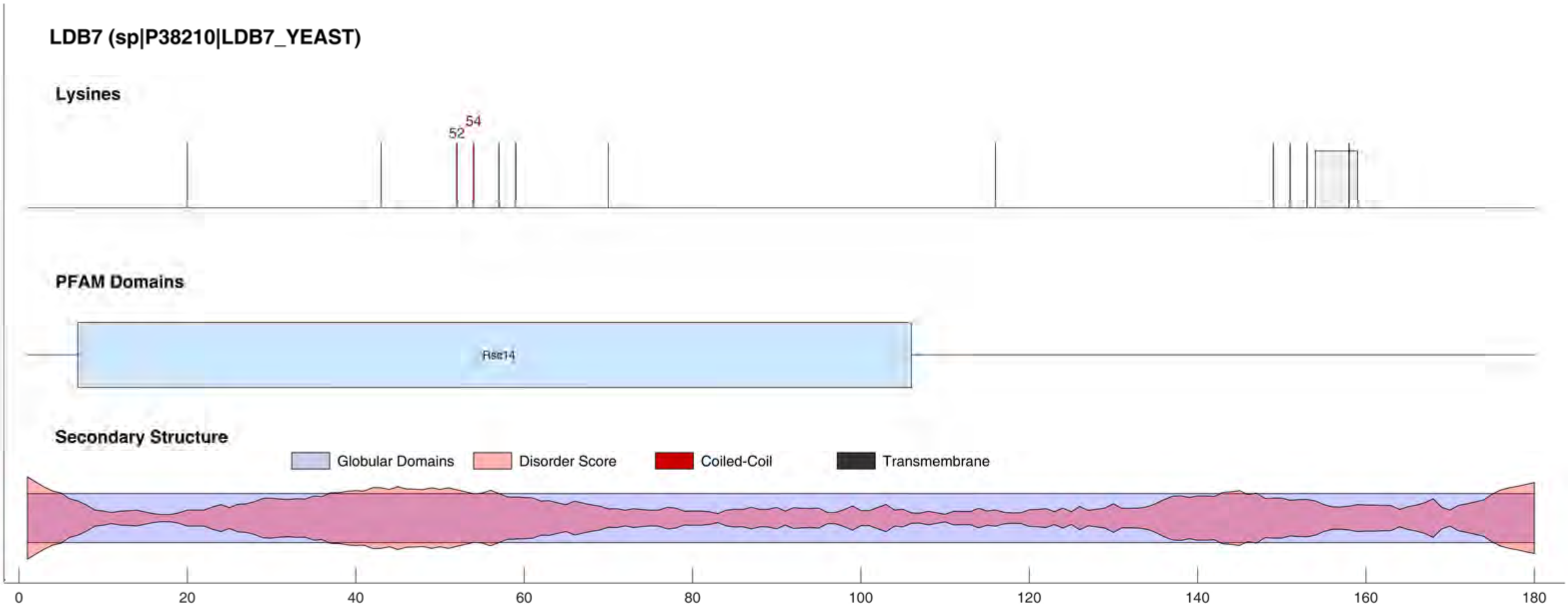

LDB7 (sp|P38210|LDB7_YEAST)
Lysines
52
54
PFAM Domains
Rsc14
Secondary Structure
Globular Domains
Disorder Score
Coiled-Coil
Transmembrane
0
20
40
60
80
100
120
140
160
180

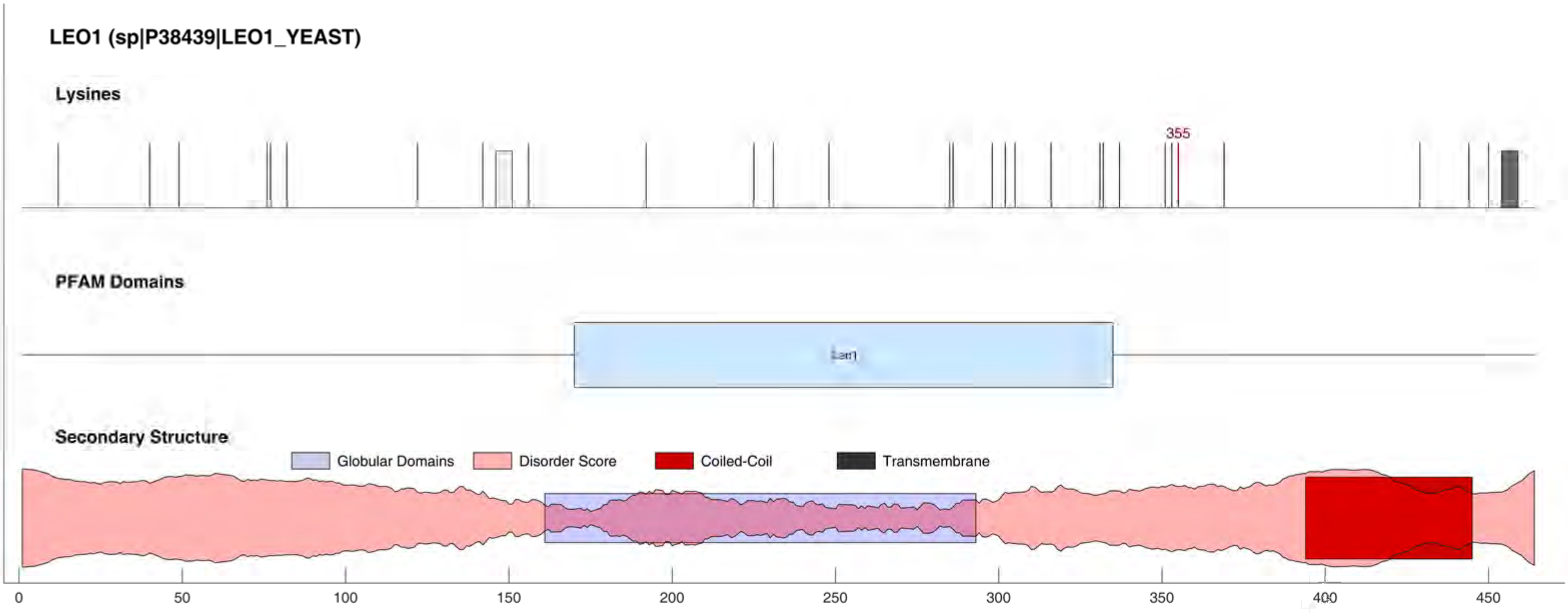
LEO1 (sp|P38439|LEO1_YEAST)
Lysines
355
PFAM Domains
Secondary Structure
Globular Domains
Disorder Score
Coiled-Coil
Transmembrane
0
50
100
150
200
250
300
350
400
450

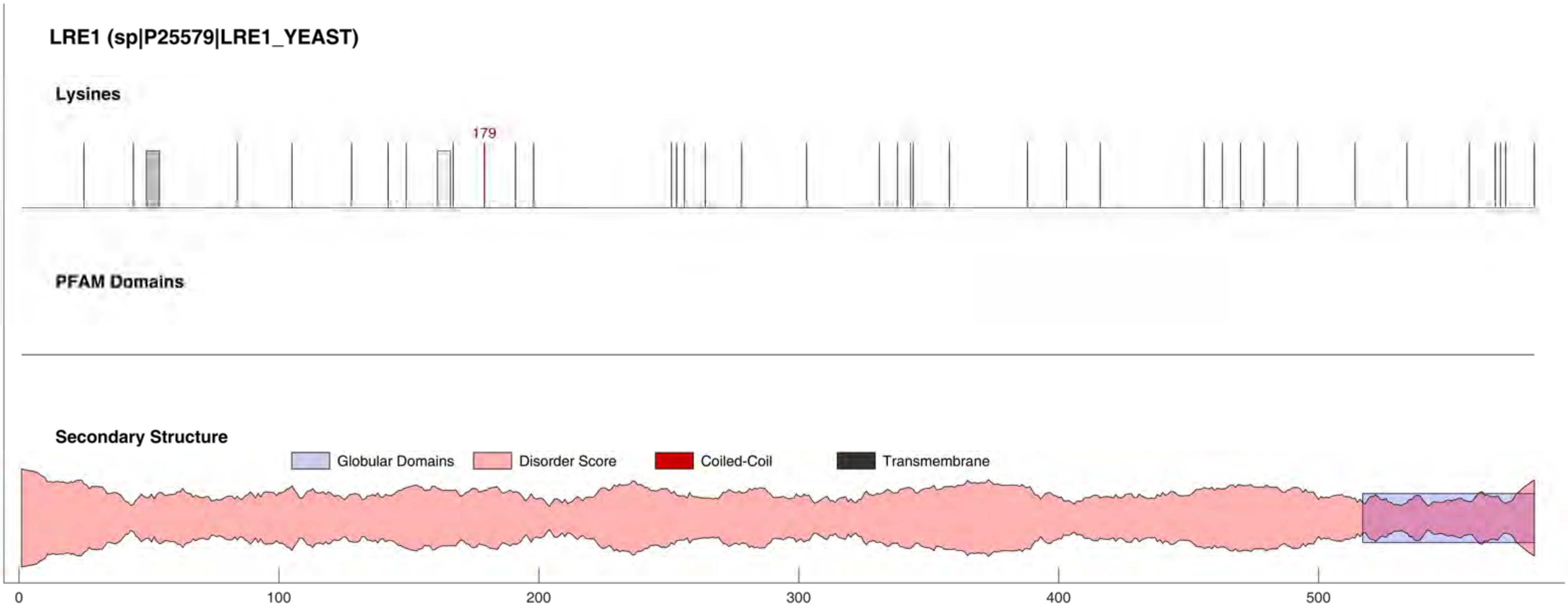
LRE1 (sp|P25579|LRE1_YEAST)
Lysines
179
PFAM Domains
Secondary Structure
Globular Domains
Disorder Score
Coiled-Coil
Transmembrane
0
100
200
300
400
500

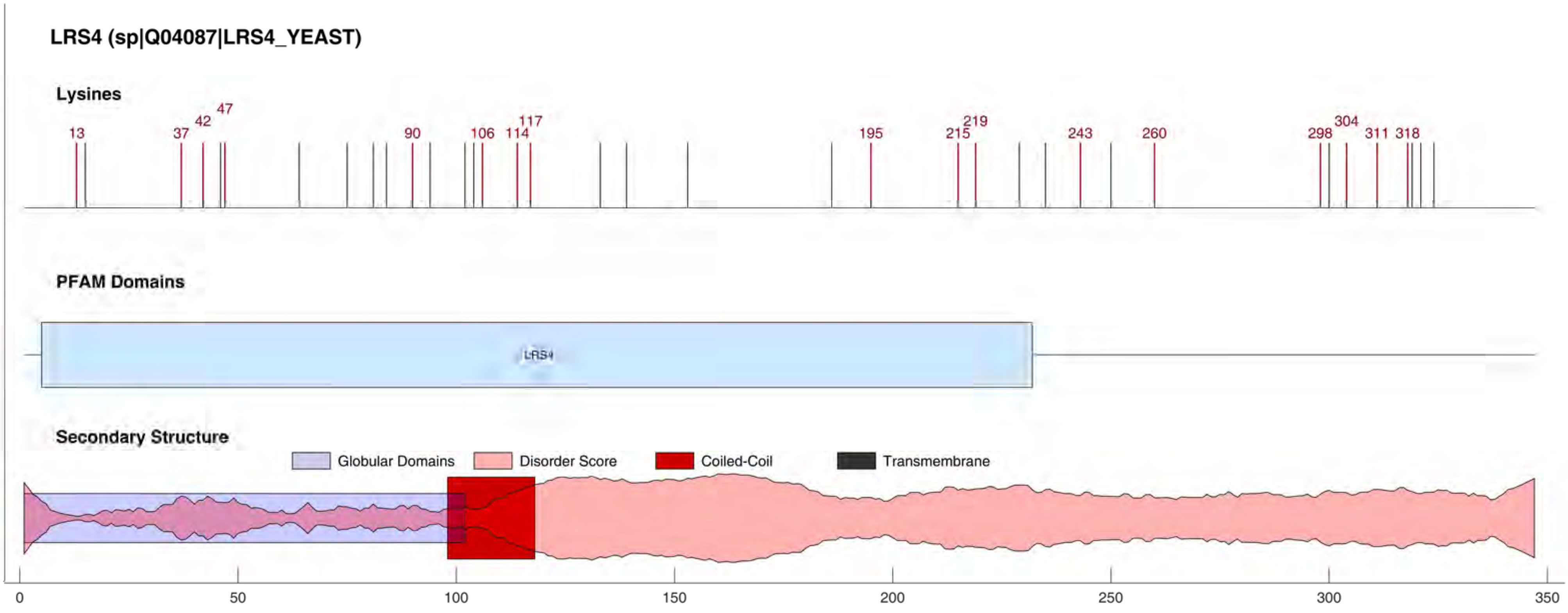

LRS4 (sp|Q04087|LRS4_YEAST)
Lysines
13
37
42
47
90
106
114
117
195
215
219
243
260
298
304
311
318
PFAM Domains
LRS4
Secondary Structure
Globular Domains
Disorder Score
Coiled-Coil
Transmembrane
0
50
100
150
200
250
300
350

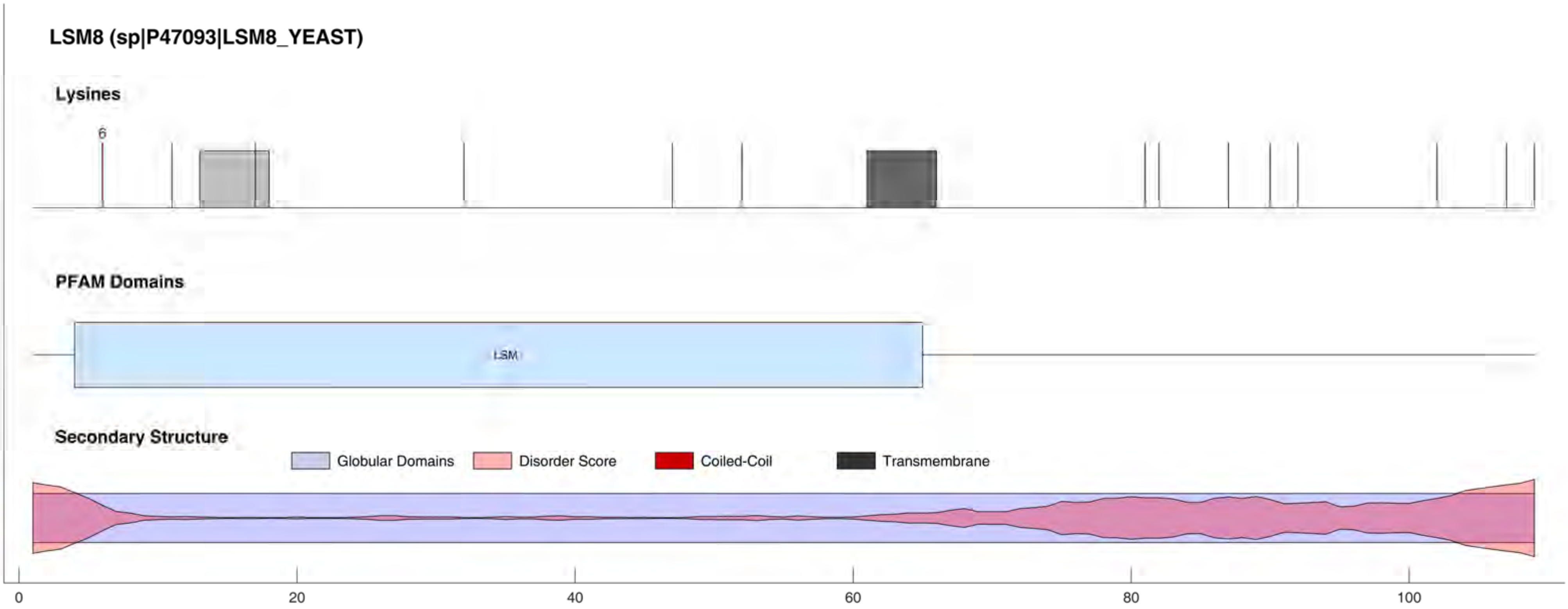
LSM8 (sp|P47093|LSM8_YEAST)
Lysines
6
PFAM Domains
LSM
Secondary Structure
Globular Domains
Disorder Score
Coiled-Coil
Transmembrane
0
20
40
60
80
100

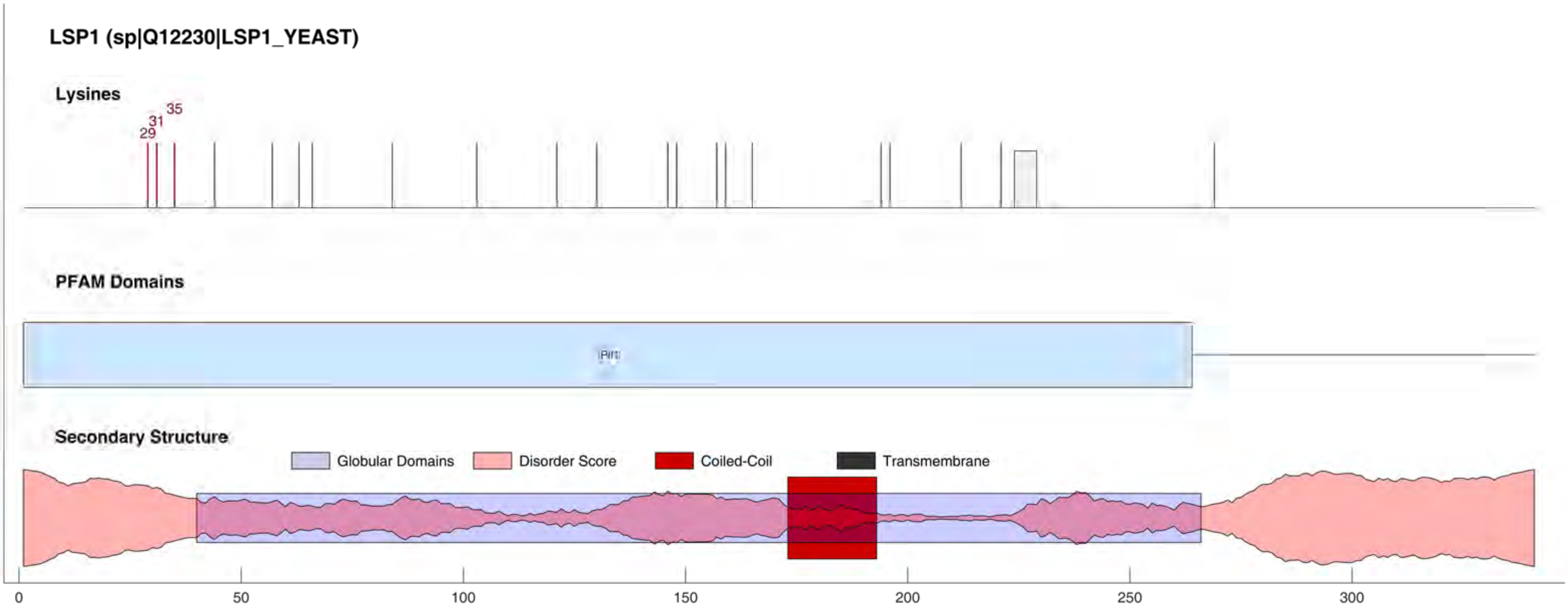
LSP1 (sp|Q12230|LSP1_YEAST)
Lysines
29
31
35
PFAM Domains
Pil1
Secondary Structure
Globular Domains
Disorder Score
Coiled-Coil
Transmembrane
0
50
100
150
200
250
300

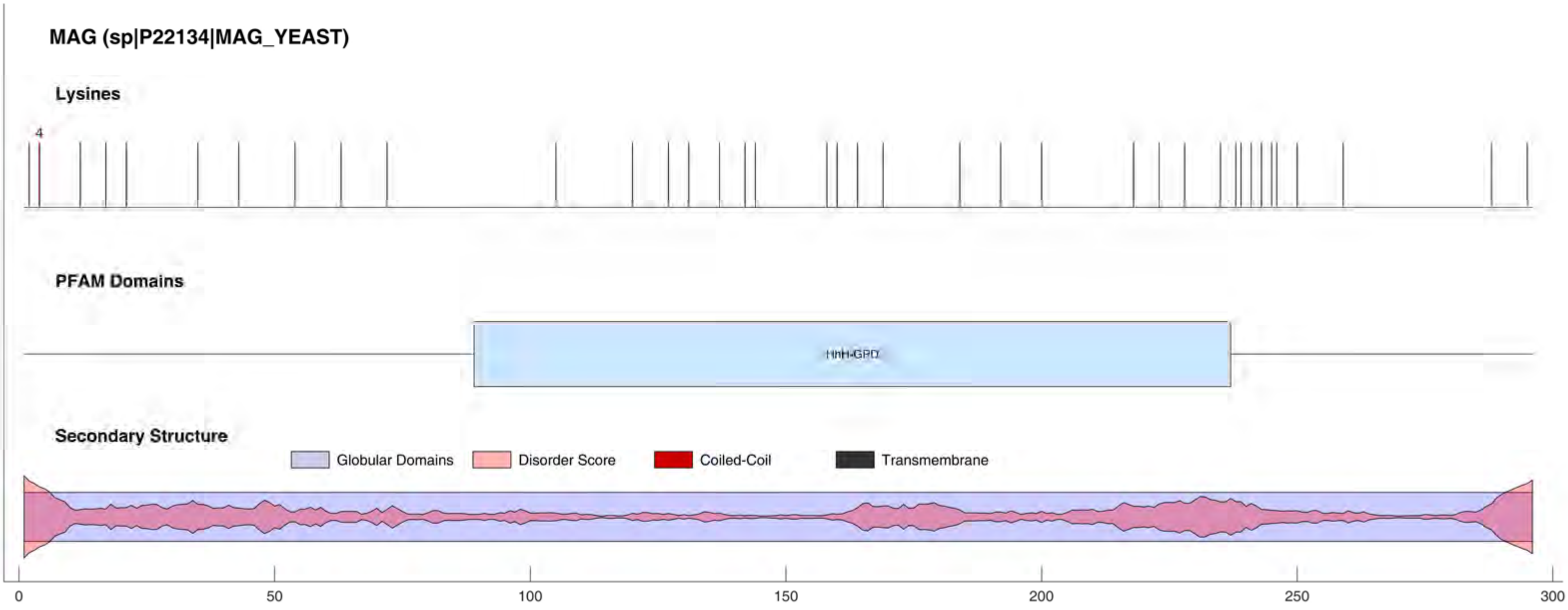
MAG (sp|P22134|MAG_YEAST)
Lysines
4
PFAM Domains
HhH-GPD
Secondary Structure
Globular Domains
Disorder Score
Coiled-Coil
Transmembrane
0
50
100
150
200
250
300

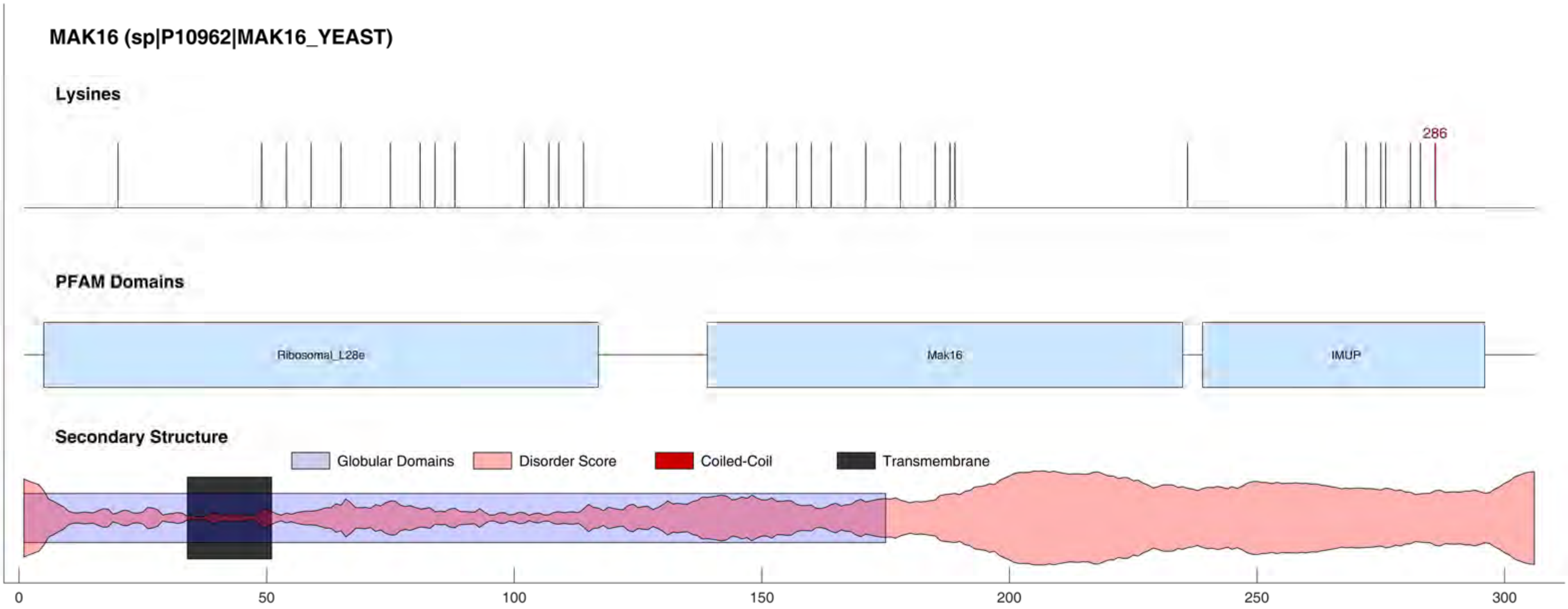
MAK16 (sp|P10962|MAK16_YEAST)
Lysines
286
PFAM Domains
Ribosomal_L28e
Mak16
IMUP
Secondary Structure
Globular Domains
Disorder Score
Coiled-Coil
Transmembrane
0
50
100
150
200
250
300

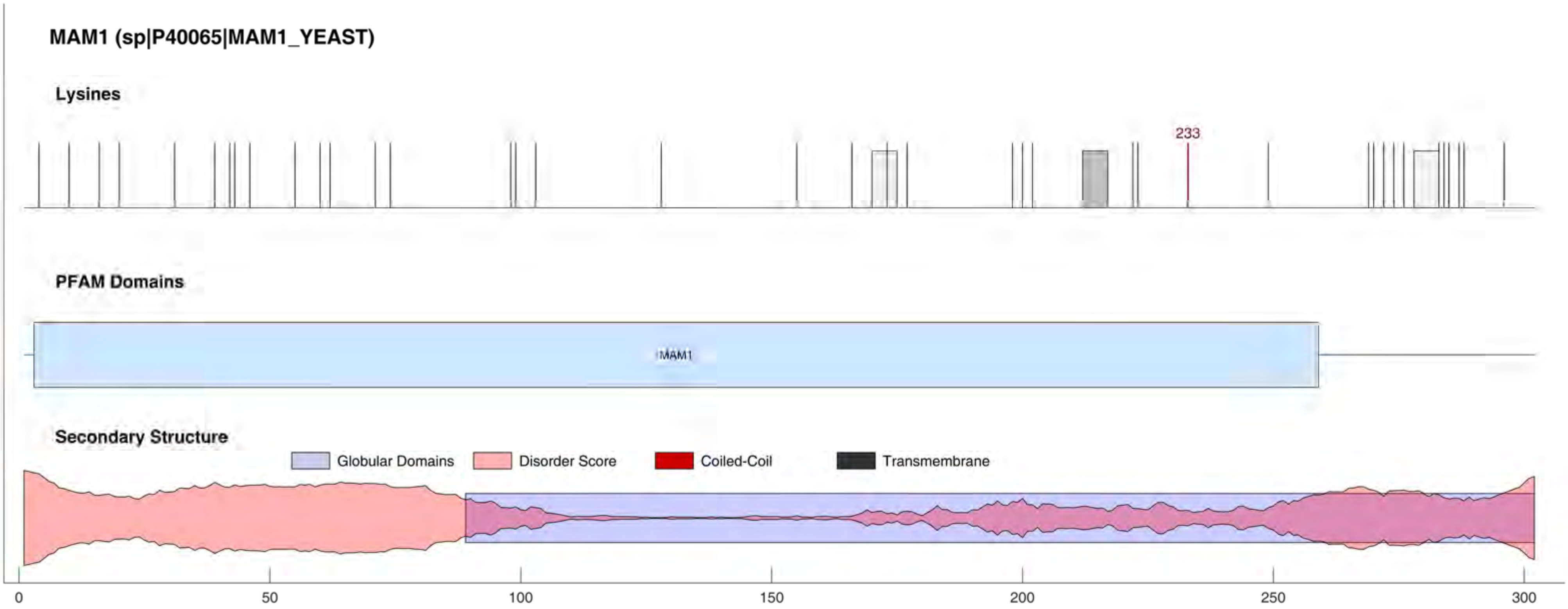
MAM1 (sp|P40065|MAM1_YEAST)
Lysines
233
PFAM Domains
MAM1
Secondary Structure
Globular Domains
Disorder Score
Coiled-Coil
Transmembrane
0
50
100
150
200
250
300

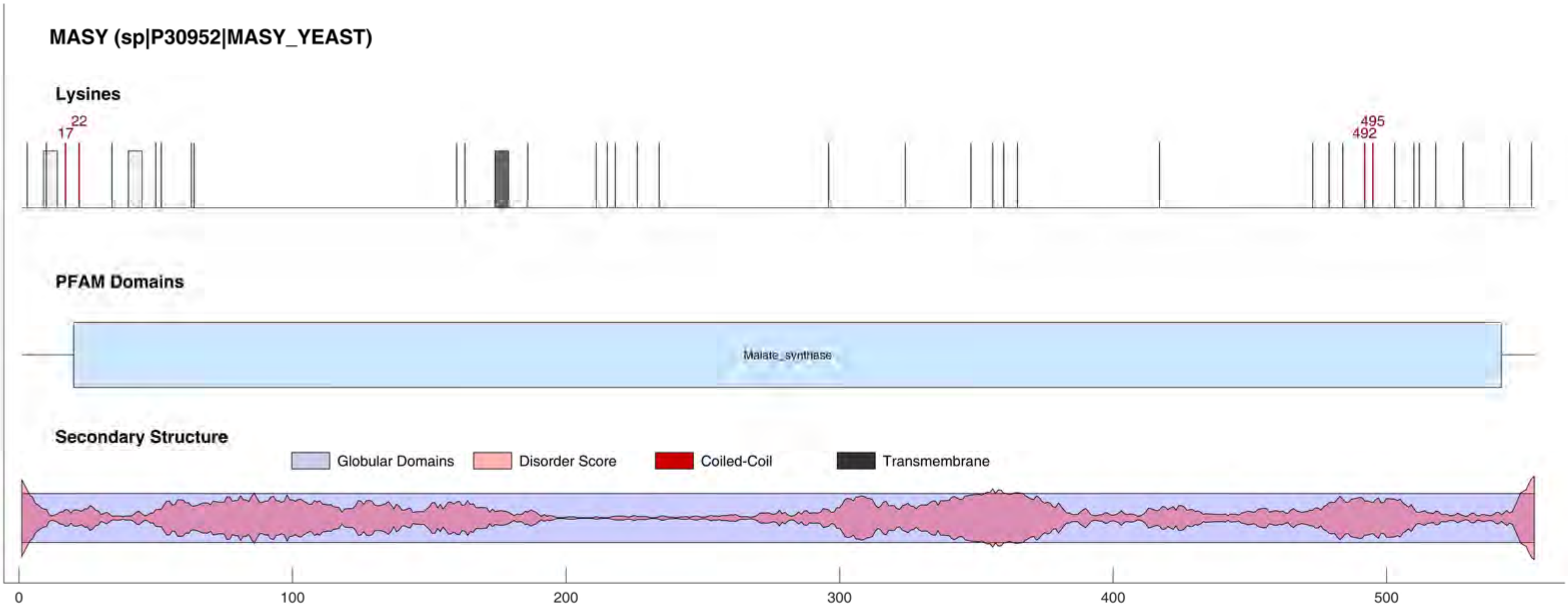
MASY (sp|P30952|MASY_YEAST)
Lysines
17
22
492
495
PFAM Domains
Malate_synthase
Secondary Structure
Globular Domains
Disorder Score
Coiled-Coil
Transmembrane
0
100
200
300
400
500

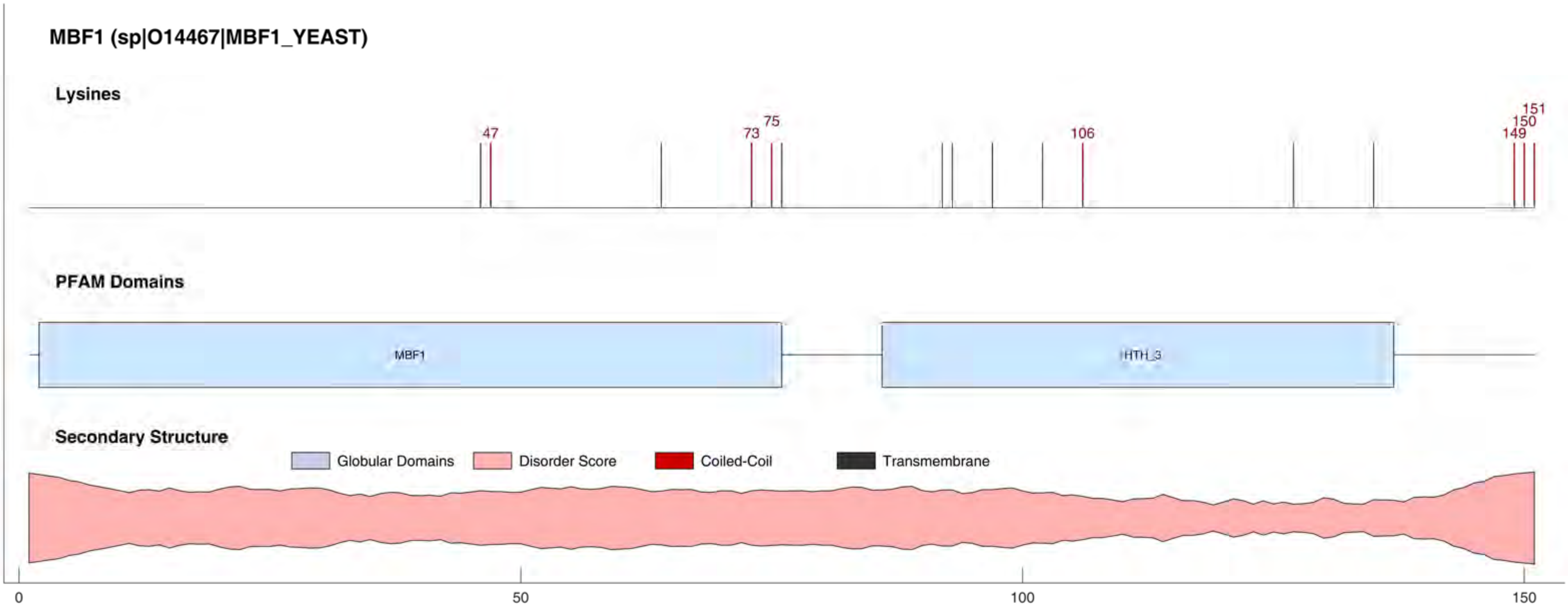
MBF1 (sp|O14467|MBF1_YEAST)
Lysines
47
73
75
106
149
150
151
PFAM Domains
MBF1
HTH_3
Secondary Structure
Globular Domains
Disorder Score
Coiled-Coil
Transmembrane
0
50
100
150

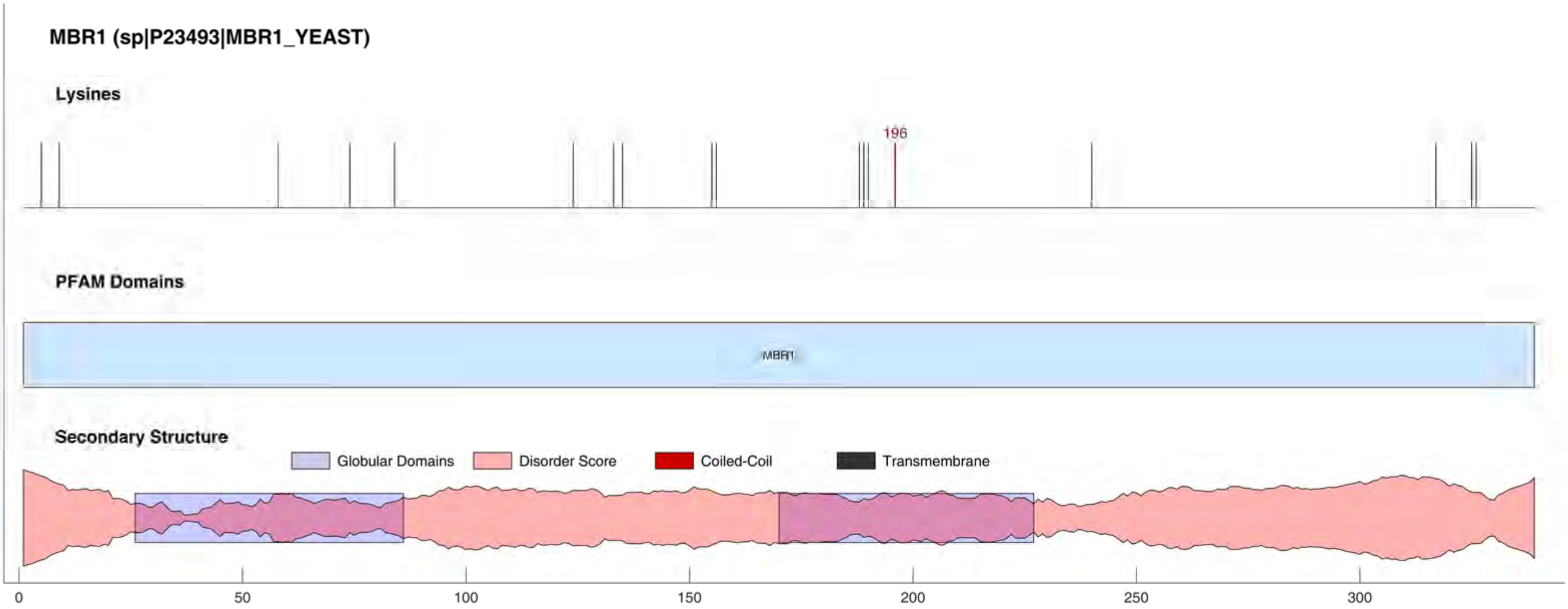
MBR1 (sp|P23493|MBR1_YEAST)
Lysines
196
PFAM Domains
MBR1
Secondary Structure
Globular Domains
Disorder Score
Coiled-Coil
Transmembrane
0
50
100
150
200
250
300

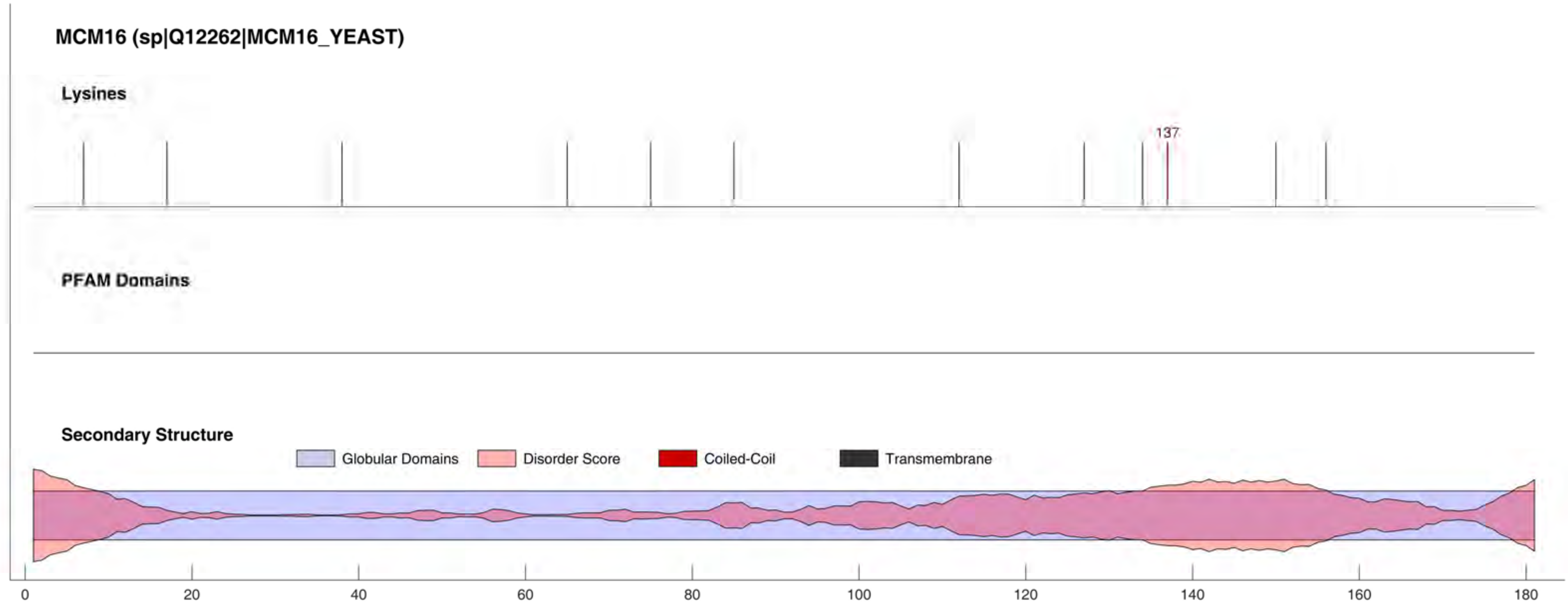
MCM16 (sp|Q12262|MCM16_YEAST)
Lysines
137
PFAM Domains
Secondary Structure
Globular Domains
Disorder Score
Coiled-Coil
Transmembrane
0
20
40
60
80
100
120
140
160
180

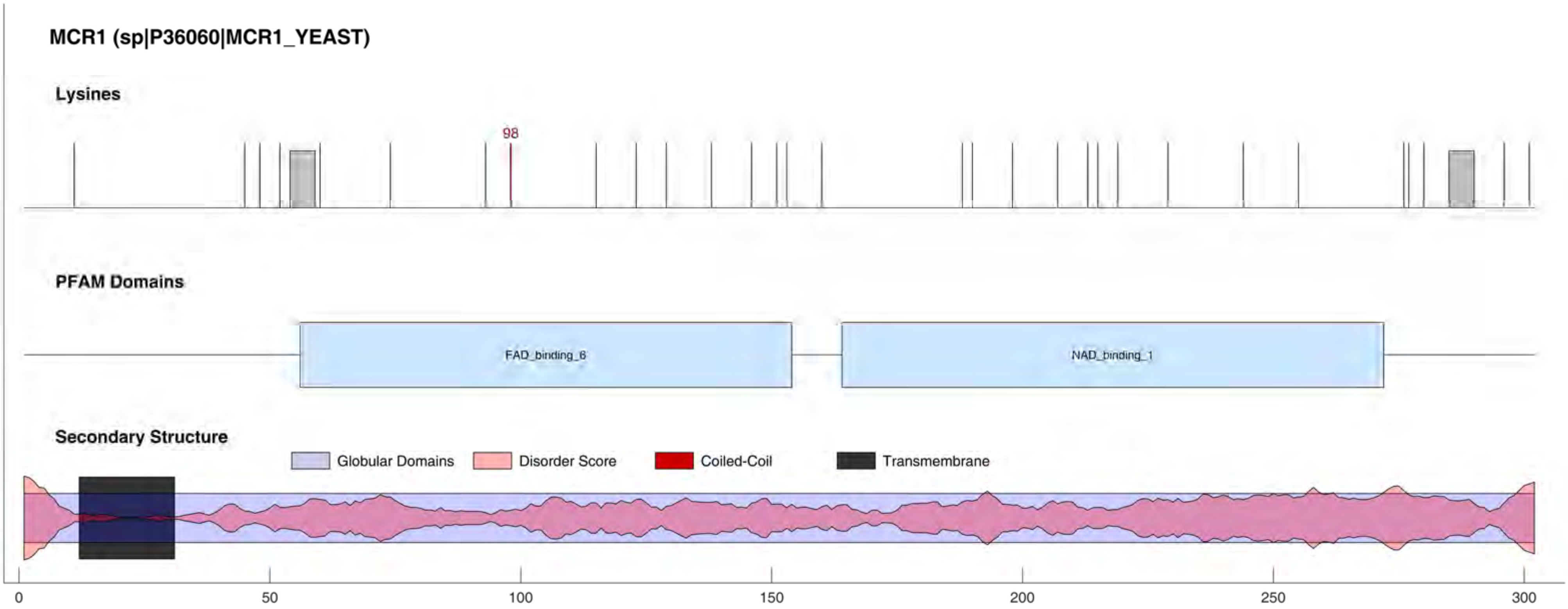

MCR1 (sp|P36060|MCR1_YEAST)
Lysines
98
PFAM Domains
FAD_binding_6
NAD_binding_1
Secondary Structure
Globular Domains
Disorder Score
Coiled-Coil
Transmembrane
0
50
100
150
200
250
300

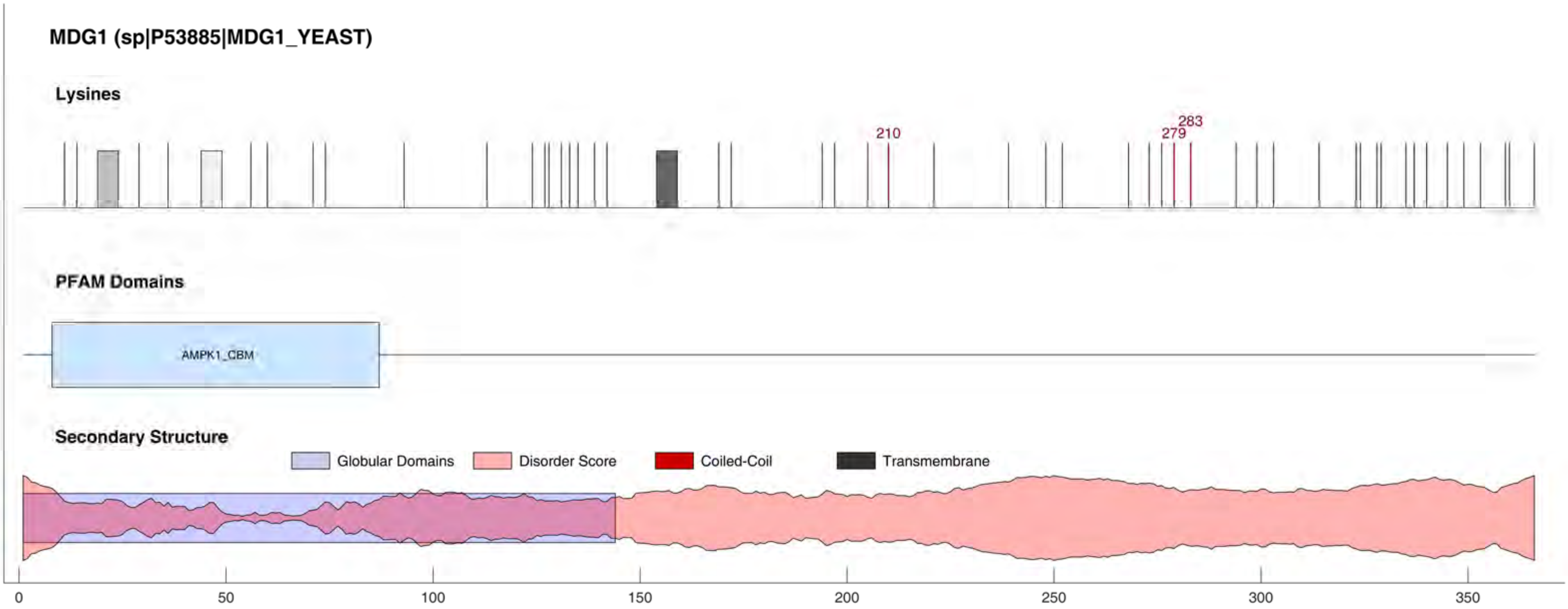
MDG1 (sp|P53885|MDG1_YEAST)
Lysines
210
279
283
PFAM Domains
AMPK1_CBM
Secondary Structure
Globular Domains
Disorder Score
Coiled-Coil
Transmembrane
0
50
100
150
200
250
300
350

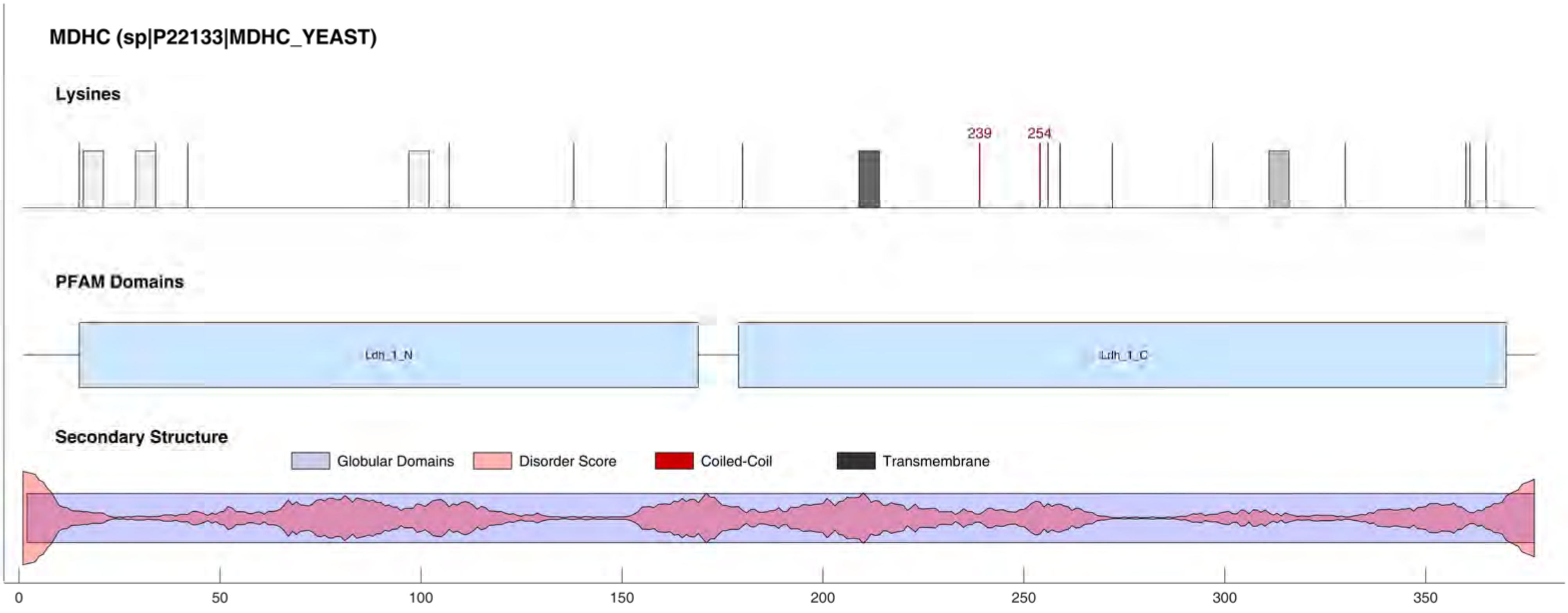
MDHC (sp|P22133|MDHC_YEAST)
Lysines
239
254
PFAM Domains
Ldh_1_N
Ldh_1_C
Secondary Structure
Globular Domains
Disorder Score
Coiled-Coil
Transmembrane
0
50
100
150
200
250
300
350

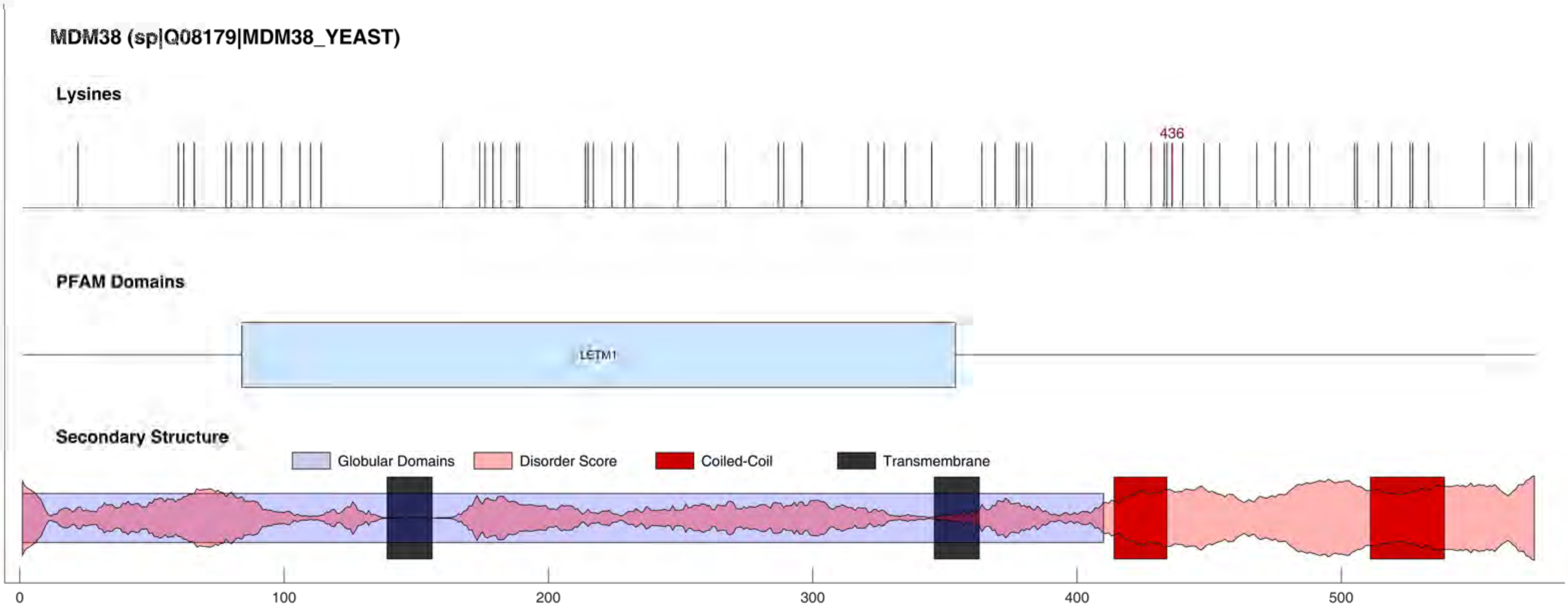
MDM38 (sp|Q08179|MDM38_YEAST)
Lysines
436
PFAM Domains
LETM1
Secondary Structure
Globular Domains
Disorder Score
Coiled-Coil
Transmembrane
0
100
200
300
400
500

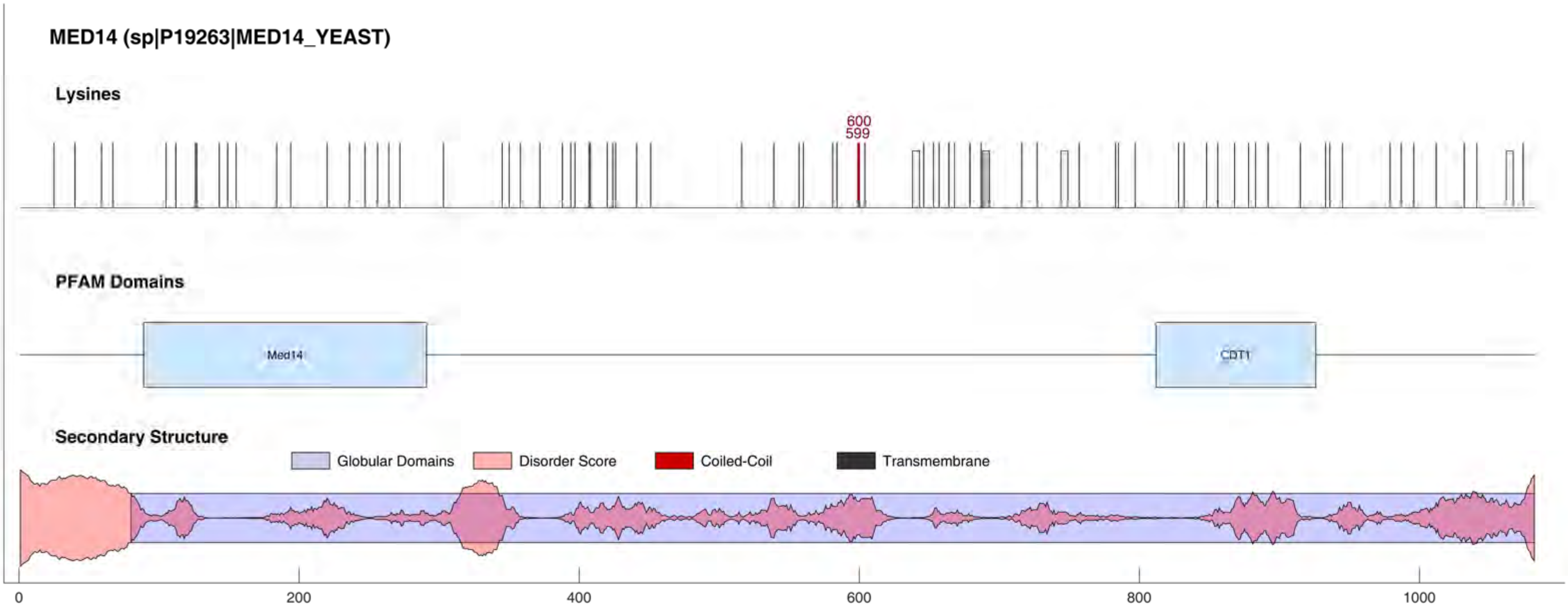

MED14 (sp|P19263|MED14_YEAST)
Lysines
600
599
PFAM Domains
Med14
CDT1
Secondary Structure
Globular Domains
Disorder Score
Coiled-Coil
Transmembrane
0
200
400
600
800
1000

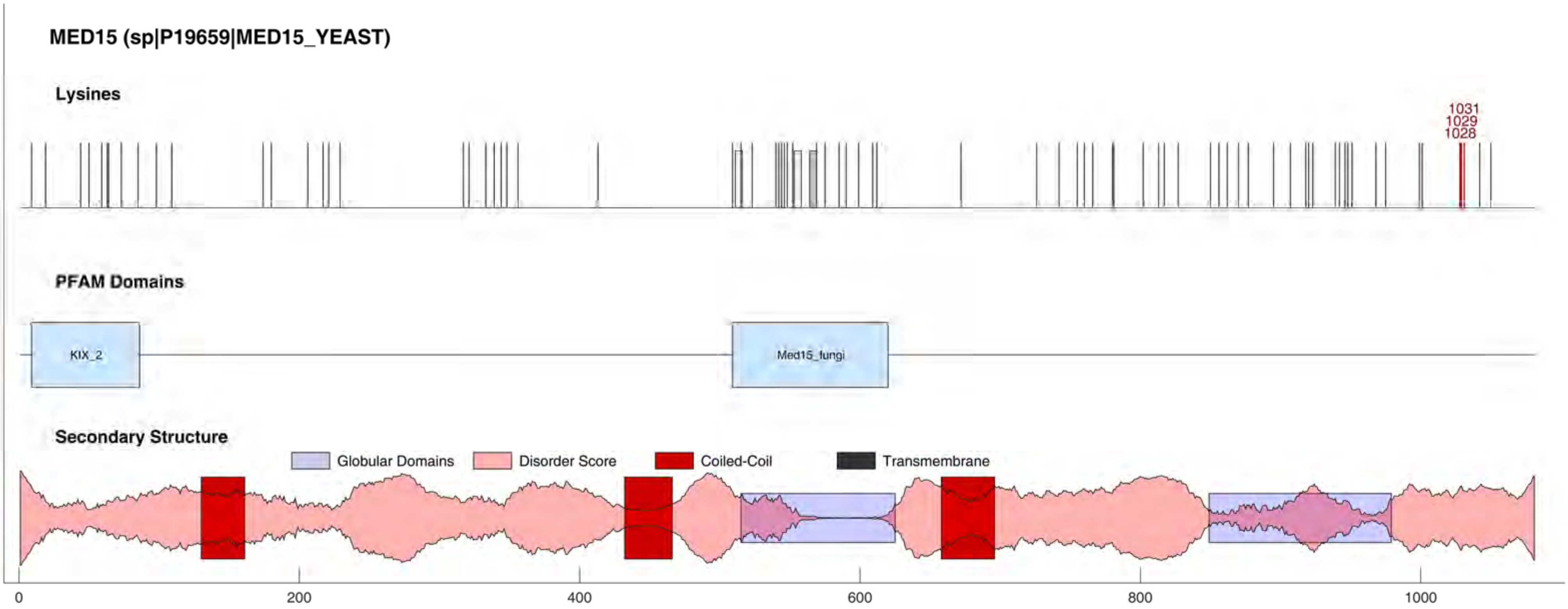
MED15 (sp|P19659|MED15_YEAST)
Lysines
1031
1029
1028
PFAM Domains
KIX_2
Med15_fungi
Secondary Structure
Globular Domains
Disorder Score
Coiled-Coil
Transmembrane
0
200
400
600
800
1000

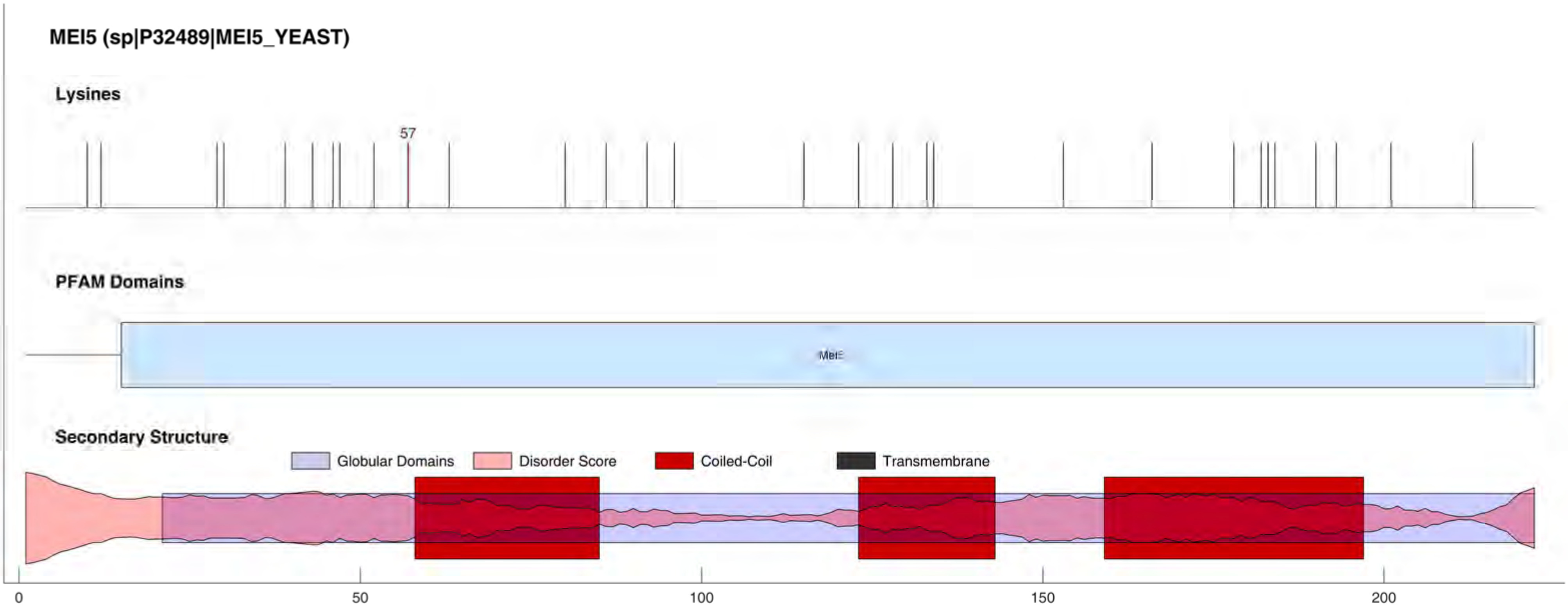
MEI5 (sp|P32489|MEI5_YEAST)
Lysines
57
PFAM Domains
Secondary Structure
Globular Domains
Disorder Score
Coiled-Coil
Transmembrane
0
50
100
150
200

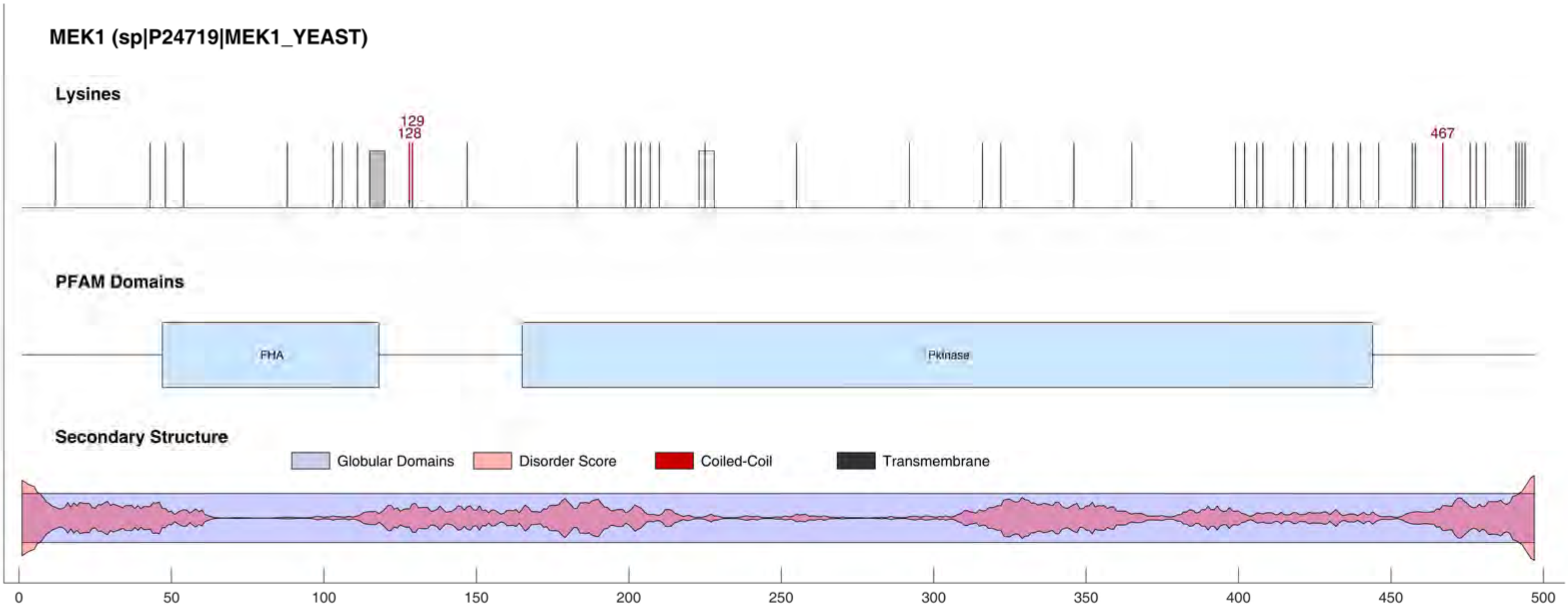
MEK1 (sp|P24719|MEK1_YEAST)
Lysines
129
128
467
PFAM Domains
FHA
Pkinase
Secondary Structure
Globular Domains
Disorder Score
Coiled-Coil
Transmembrane
0
50
100
150
200
250
300
350
400
450
500

#### MET4 (sp|P32389|MET4_YEAST)

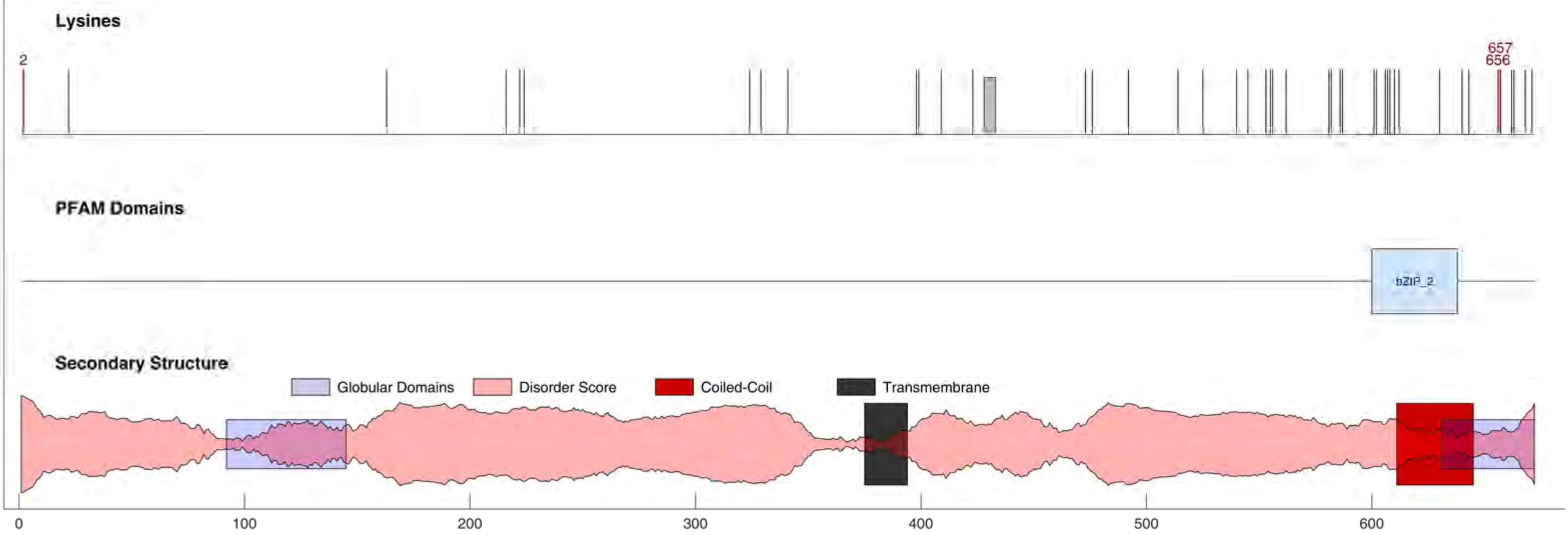

#### MET28 (sp|P40573|MET28_YEAST)

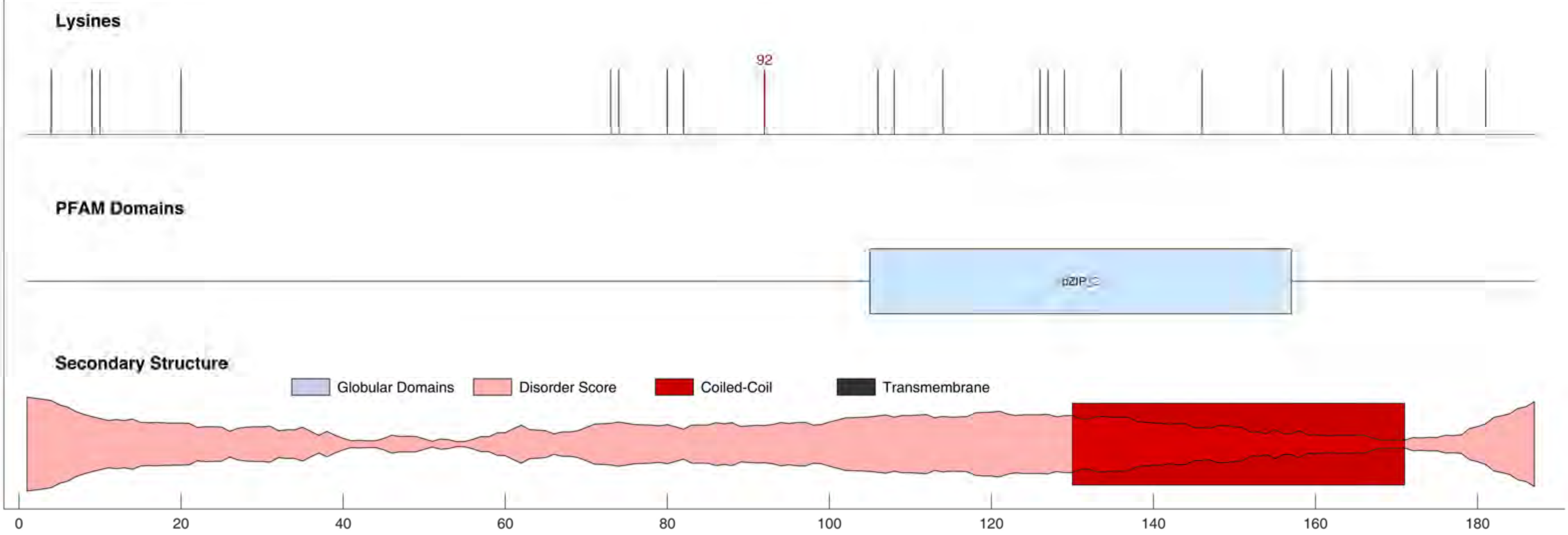

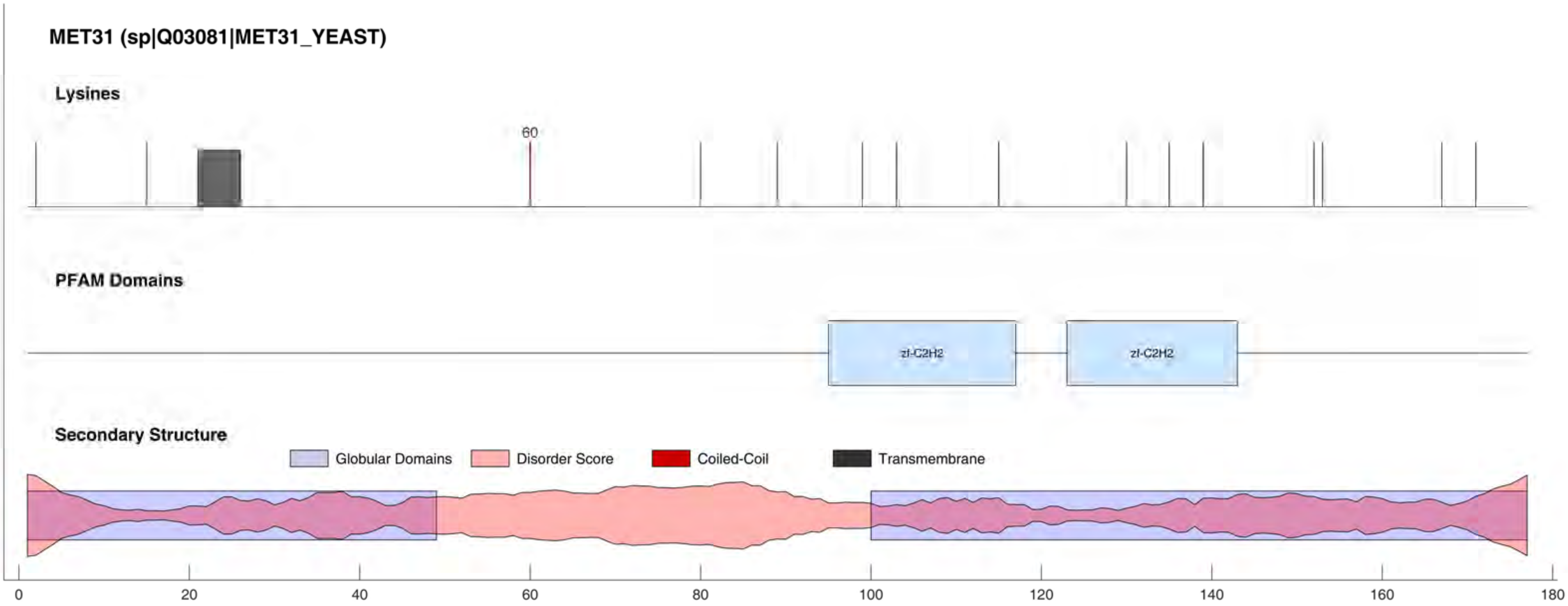

MET31 (sp|Q03081|MET31_YEAST)
Lysines
60
PFAM Domains
zf-C2H2
zf-C2H2
Secondary Structure
Globular Domains
Disorder Score
Coiled-Coil
Transmembrane
0
20
40
60
80
100
120
140
160
180

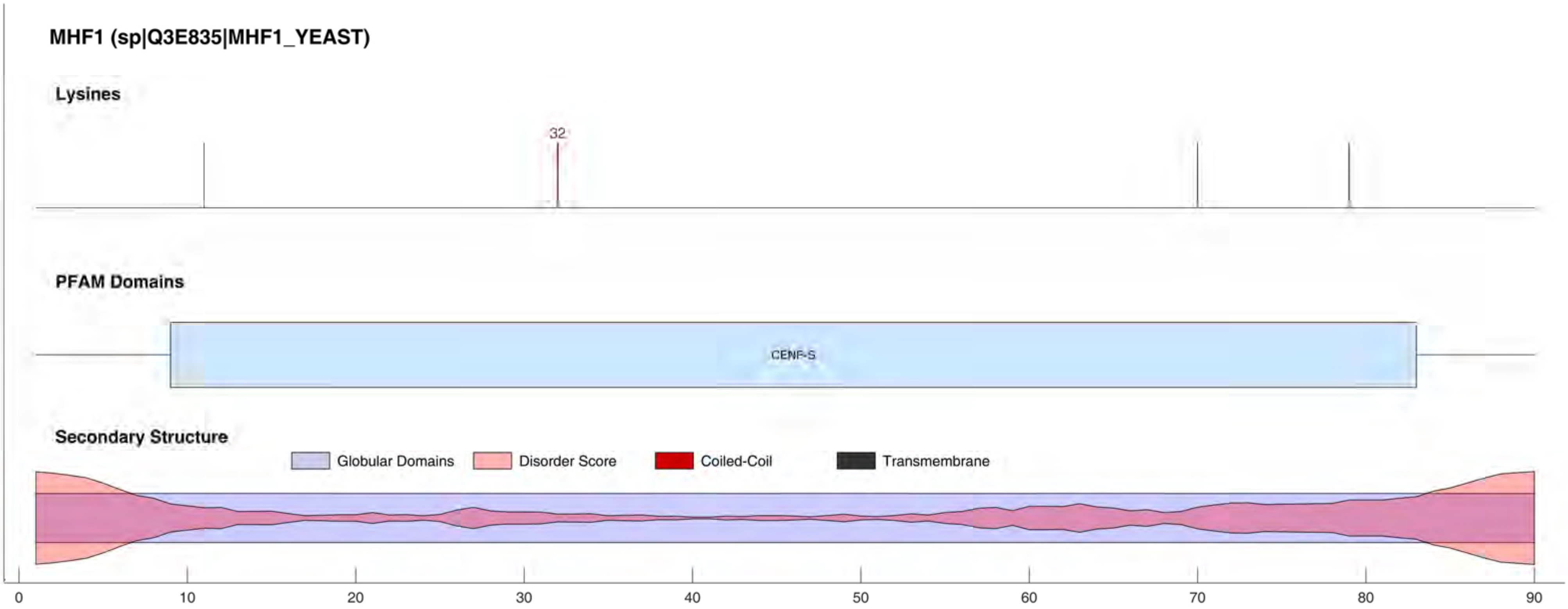
MHF1 (sp|Q3E835|MHF1_YEAST)
Lysines
32
PFAM Domains
CENP-S
Secondary Structure
Globular Domains
Disorder Score
Coiled-Coil
Transmembrane
0
10
20
30
40
50
60
70
80
90

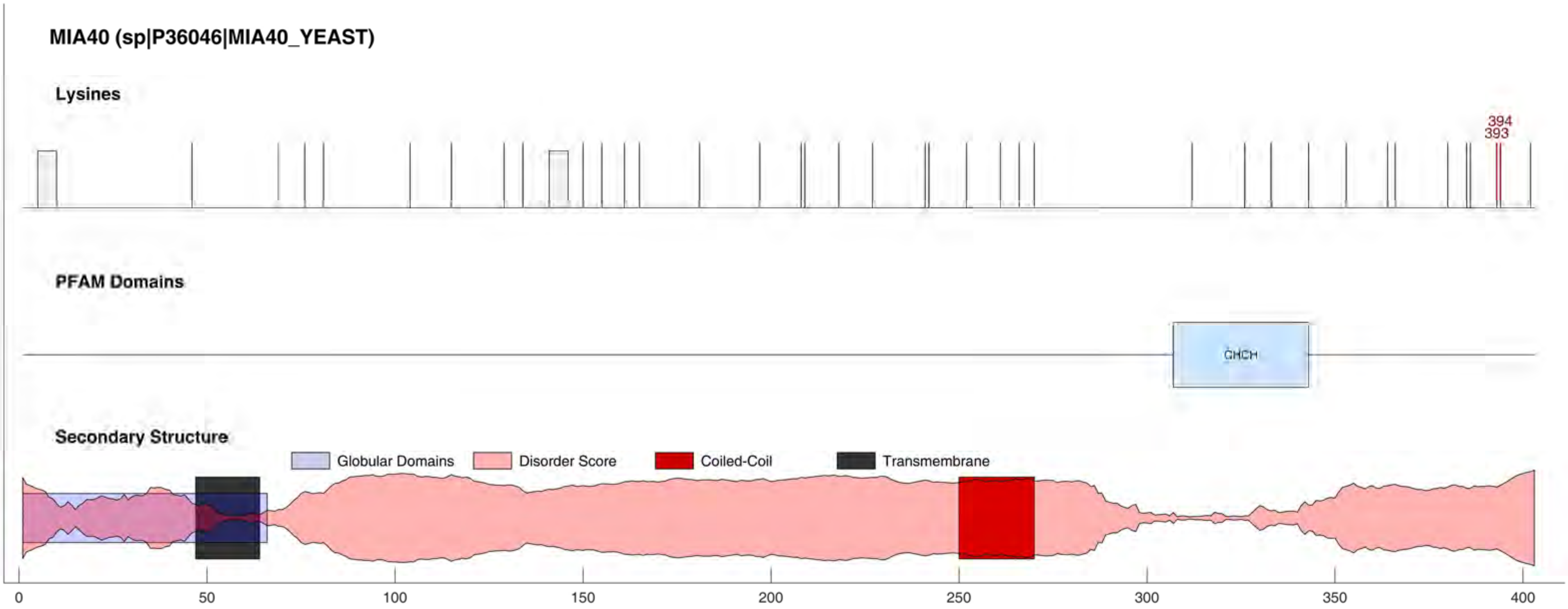
MIA40 (sp|P36046|MIA40_YEAST)
Lysines
394
393
PFAM Domains
CHCH
Secondary Structure
Globular Domains
Disorder Score
Coiled-Coil
Transmembrane
0
50
100
150
200
250
300
350
400

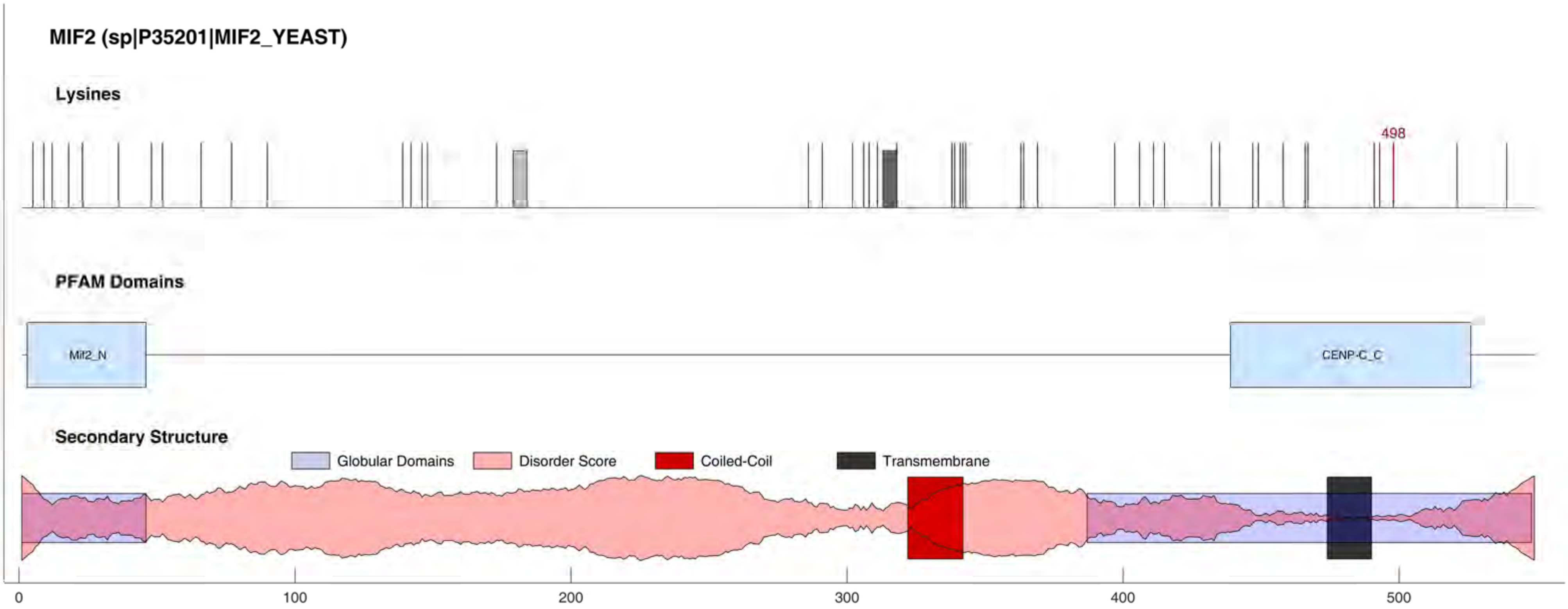
MIF2 (sp|P35201|MIF2_YEAST)
Lysines
498
PFAM Domains
Mif2_N
CENP-C_C
Secondary Structure
Globular Domains
Disorder Score
Coiled-Coil
Transmembrane
0
100
200
300
400
500

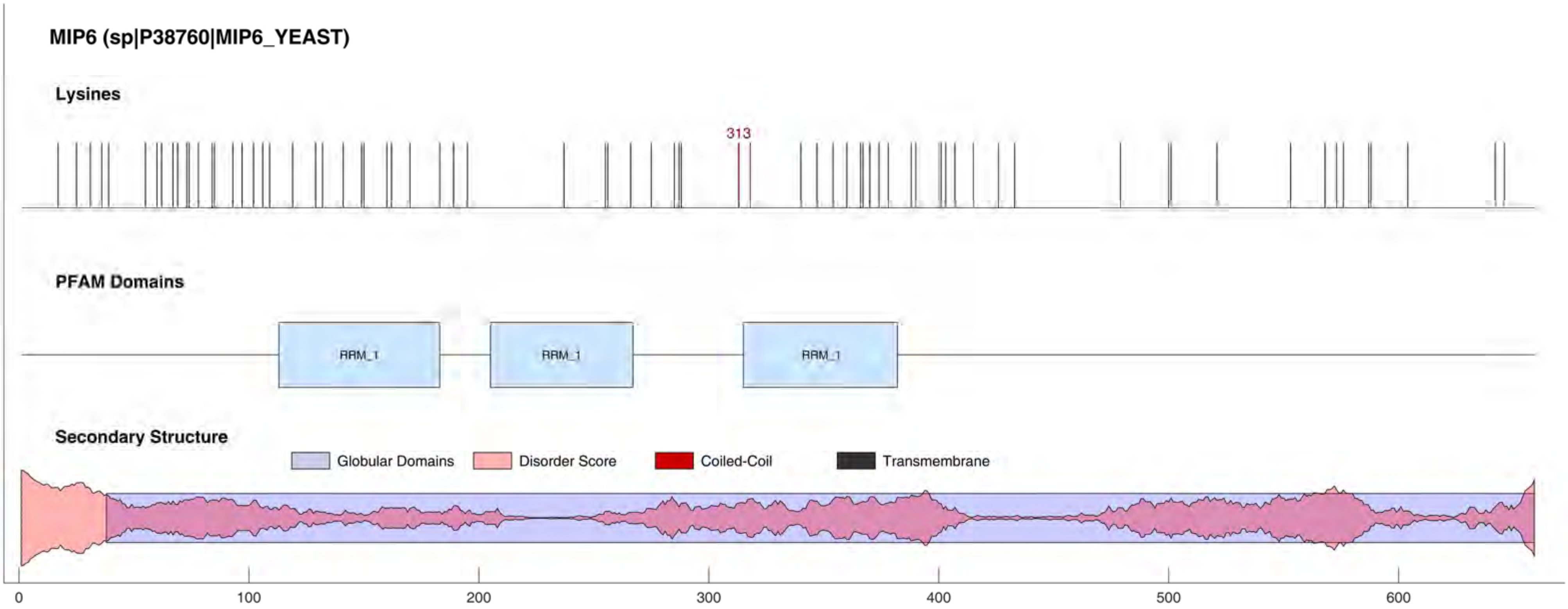
MIP6 (sp|P38760|MIP6_YEAST)
Lysines
313
PFAM Domains
RRM_1
RRM_1
RRM_1
Secondary Structure
Globular Domains
Disorder Score
Coiled-Coil
Transmembrane
0
100
200
300
400
500
600

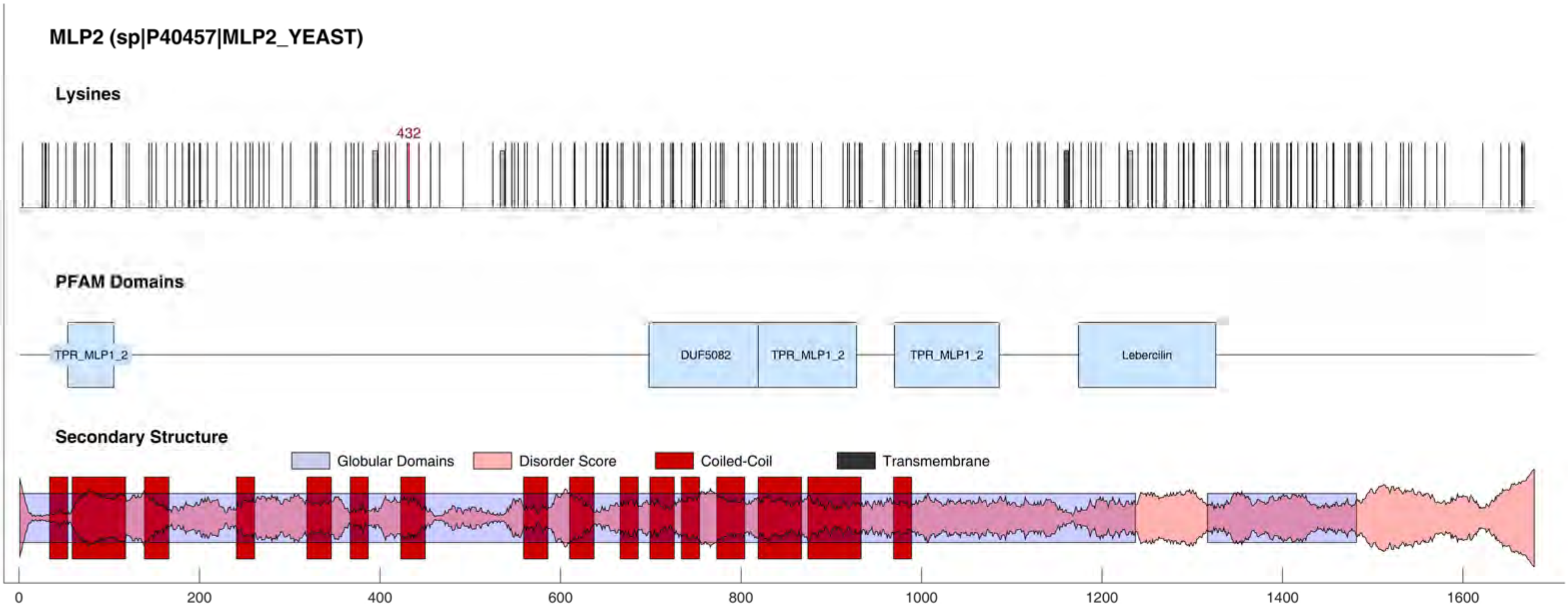
MLP2 (sp|P40457|MLP2_YEAST)
Lysines
432
PFAM Domains
TPR_MLP1_2
DUF5082
TPR_MLP1_2
TPR_MLP1_2
Lebercilin
Secondary Structure
Globular Domains
Disorder Score
Coiled-Coil
Transmembrane
0
200
400
600
800
1000
1200
1400
1600

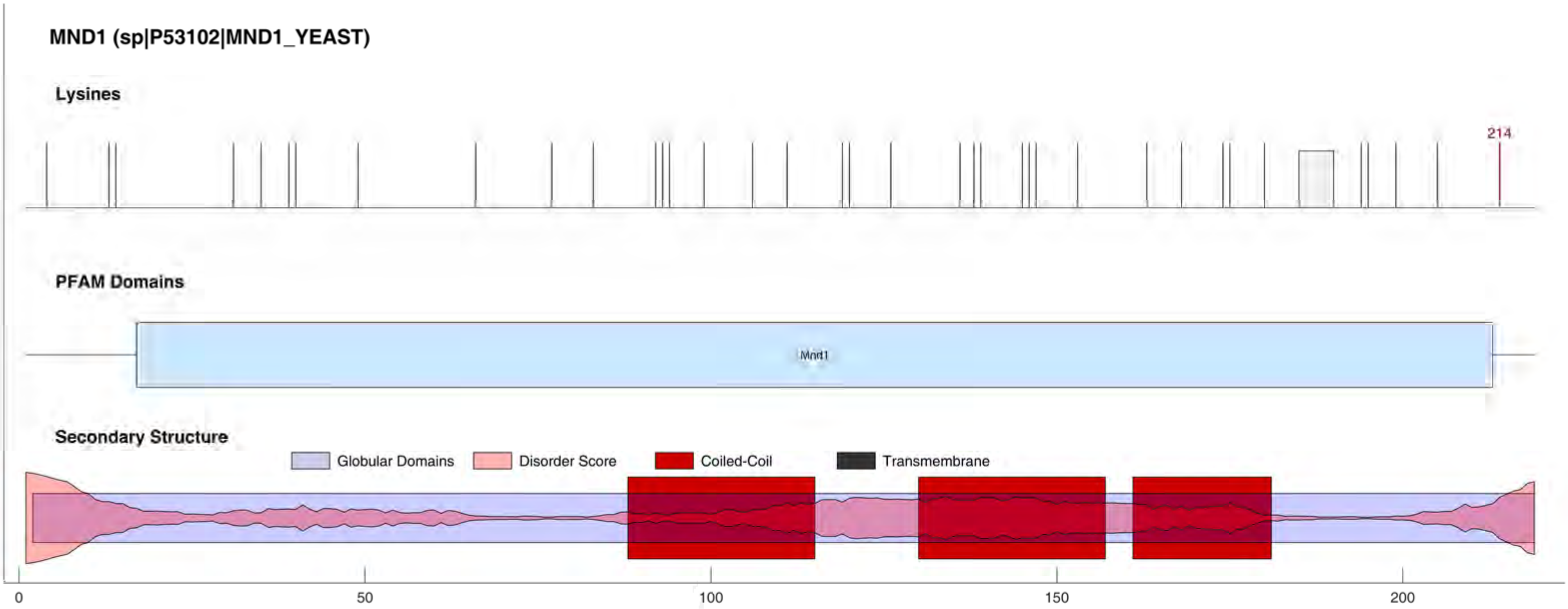
MND1 (sp|P53102|MND1_YEAST)
Lysines
214
PFAM Domains
Mnd1
Secondary Structure
Globular Domains
Disorder Score
Coiled-Coil
Transmembrane
0
50
100
150
200

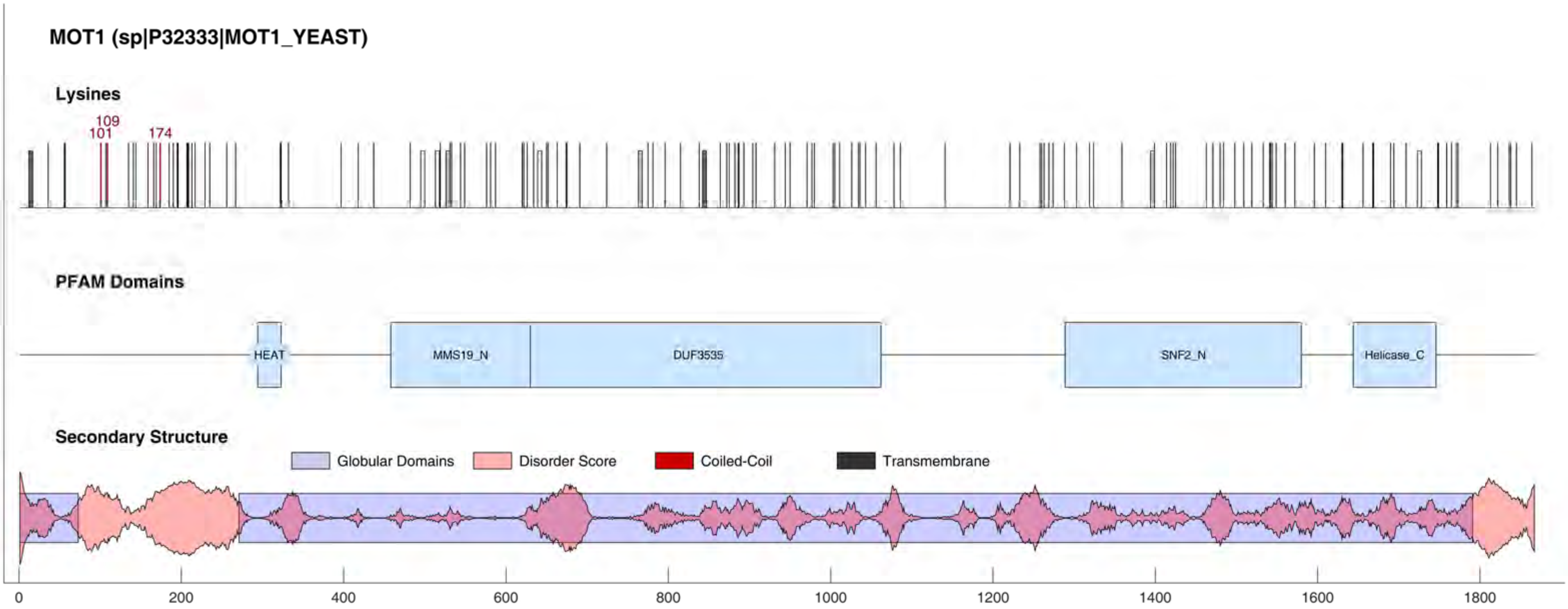
MOT1 (sp|P32333|MOT1_YEAST)
Lysines
109
101
174
PFAM Domains
HEAT
MMS19_N
DUF3535
SNF2_N
Helicase_C
Secondary Structure
Globular Domains
Disorder Score
Coiled-Coil
Transmembrane
0
200
400
600
800
1000
1200
1400
1600
1800

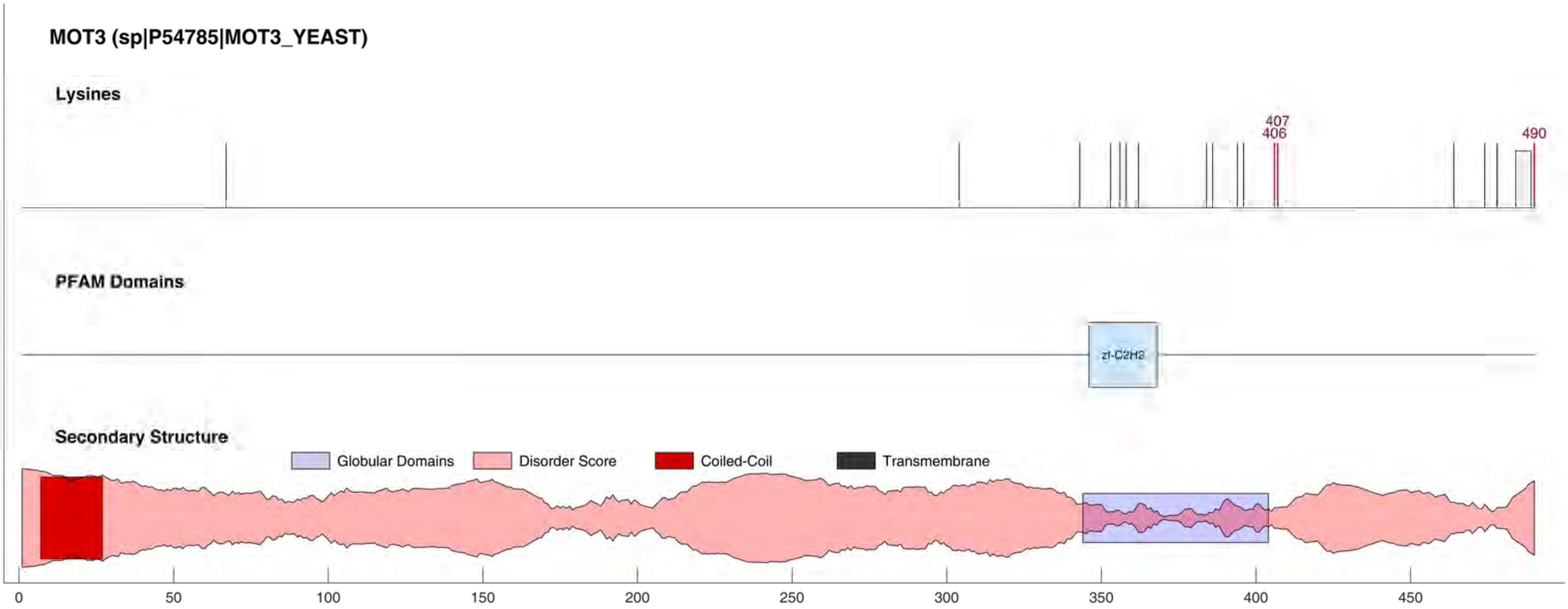
MOT3 (sp|P54785|MOT3_YEAST)
Lysines
407
406
490
PFAM Domains
zf-C2H2
Secondary Structure
Globular Domains
Disorder Score
Coiled-Coil
Transmembrane
0
50
100
150
200
250
300
350
400
450

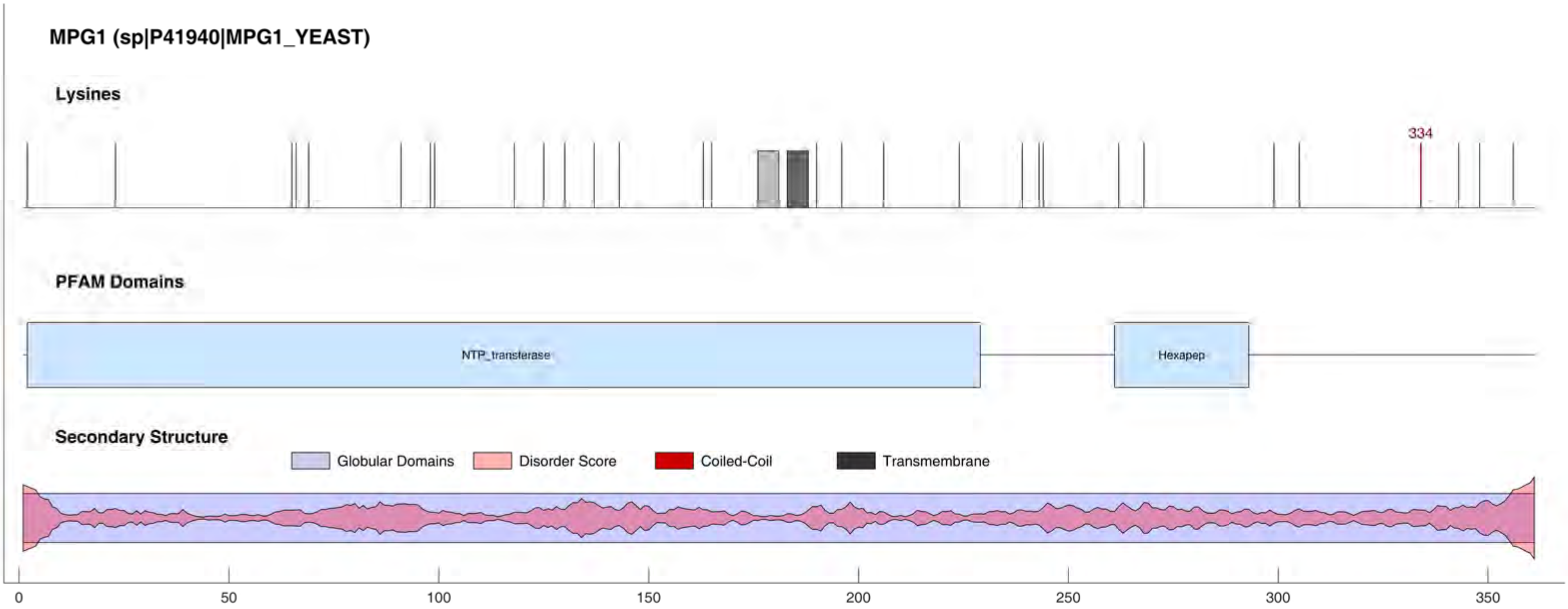
MPG1 (sp|P41940|MPG1_YEAST)
Lysines
334
PFAM Domains
NTP_transferase
Hexapep
Secondary Structure
Globular Domains
Disorder Score
Coiled-Coil
Transmembrane
0
50
100
150
200
250
300
350

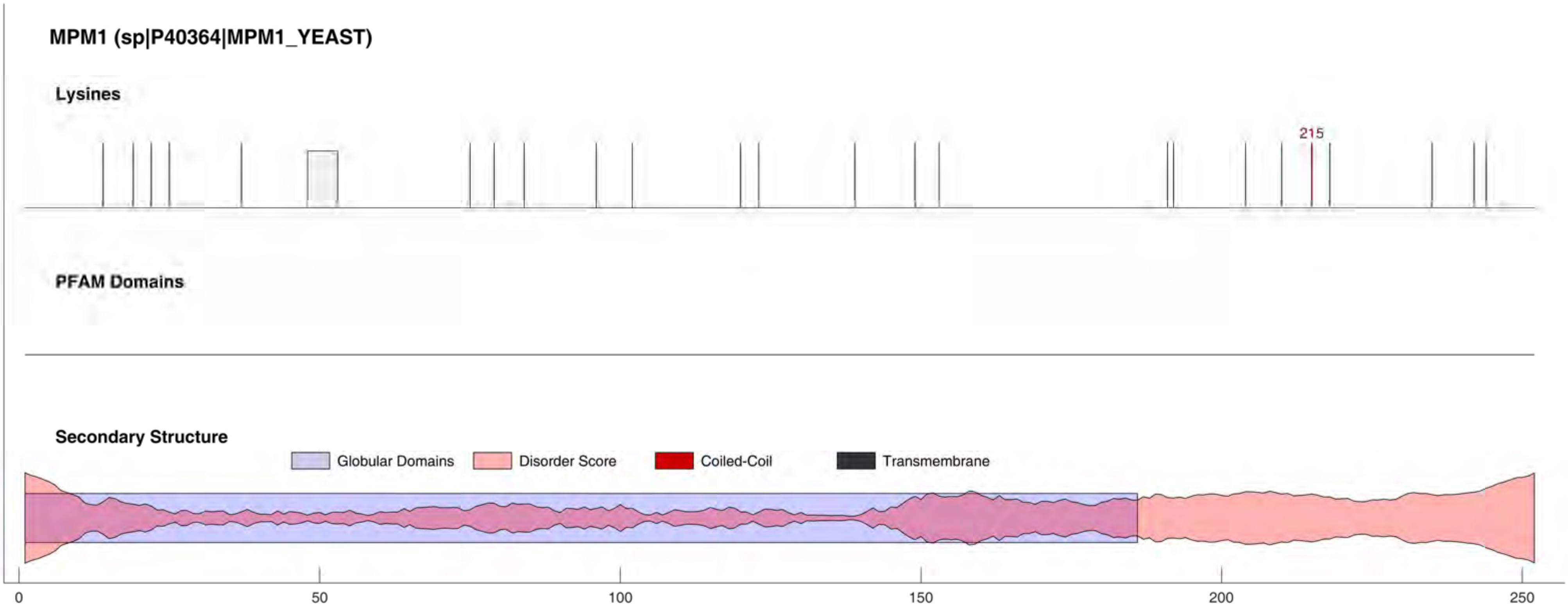
MPM1 (sp|P40364|MPM1_YEAST)
Lysines
215
PFAM Domains
Secondary Structure
Globular Domains
Disorder Score
Coiled-Coil
Transmembrane
0
50
100
150
200
250

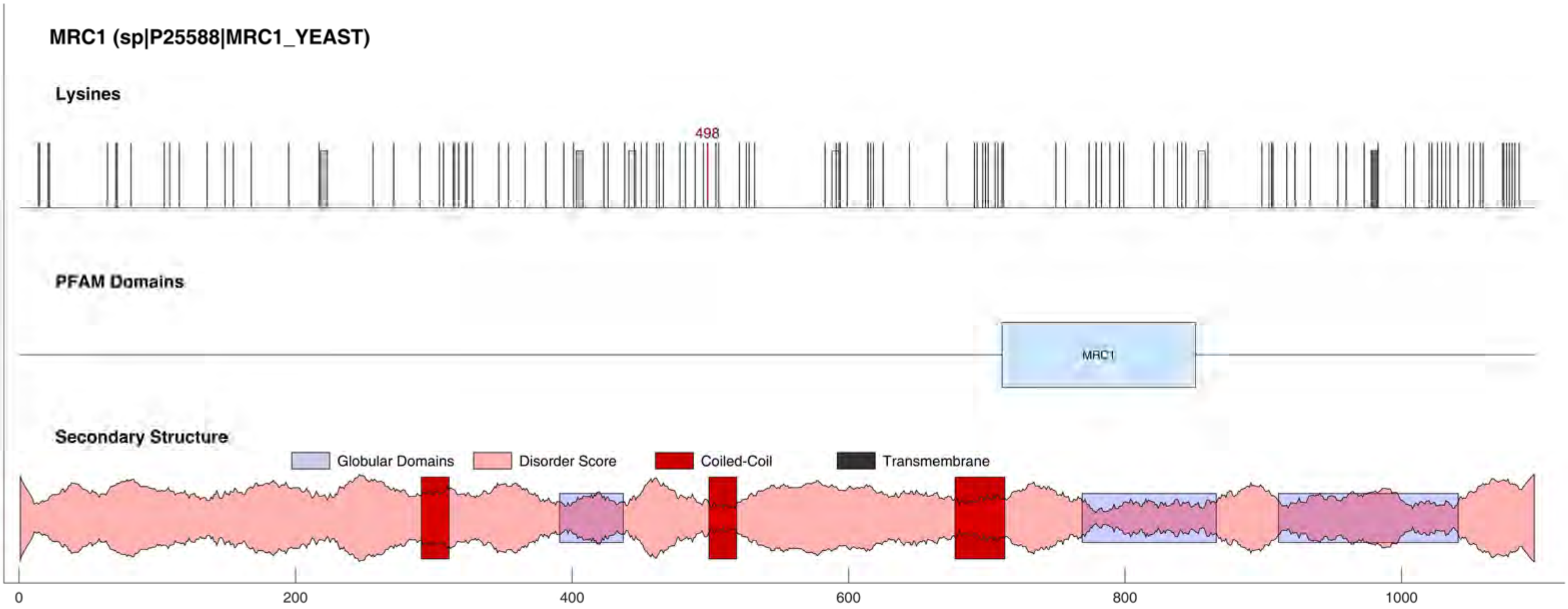

MRC1 (sp|P25588|MRC1_YEAST)
Lysines
498
PFAM Domains
MRC1
Secondary Structure
Globular Domains
Disorder Score
Coiled-Coil
Transmembrane
0
200
400
600
800
1000

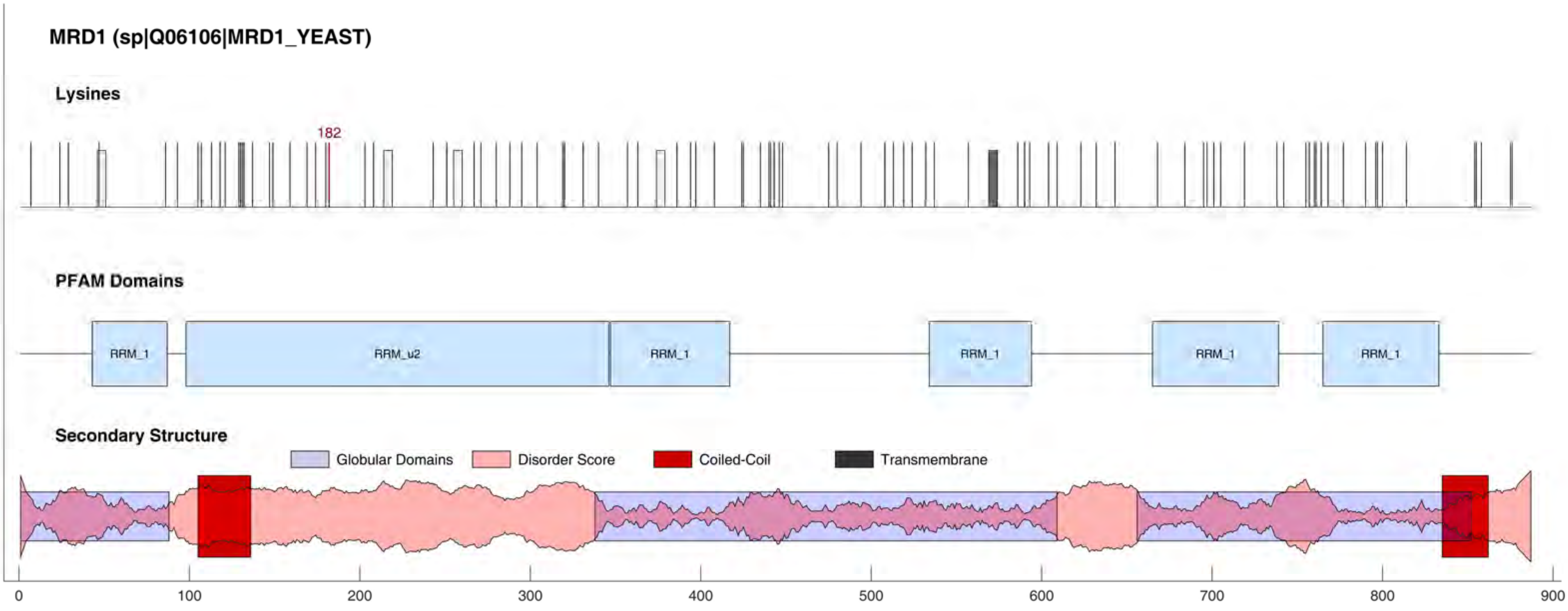

MRD1 (sp|Q06106|MRD1_YEAST)
Lysines
182
PFAM Domains
RRM_1
RRM_u2
RRM_1
RRM_1
RRM_1
RRM_1
Secondary Structure
Globular Domains
Disorder Score
Coiled-Coil
Transmembrane
0
100
200
300
400
500
600
700
800
900

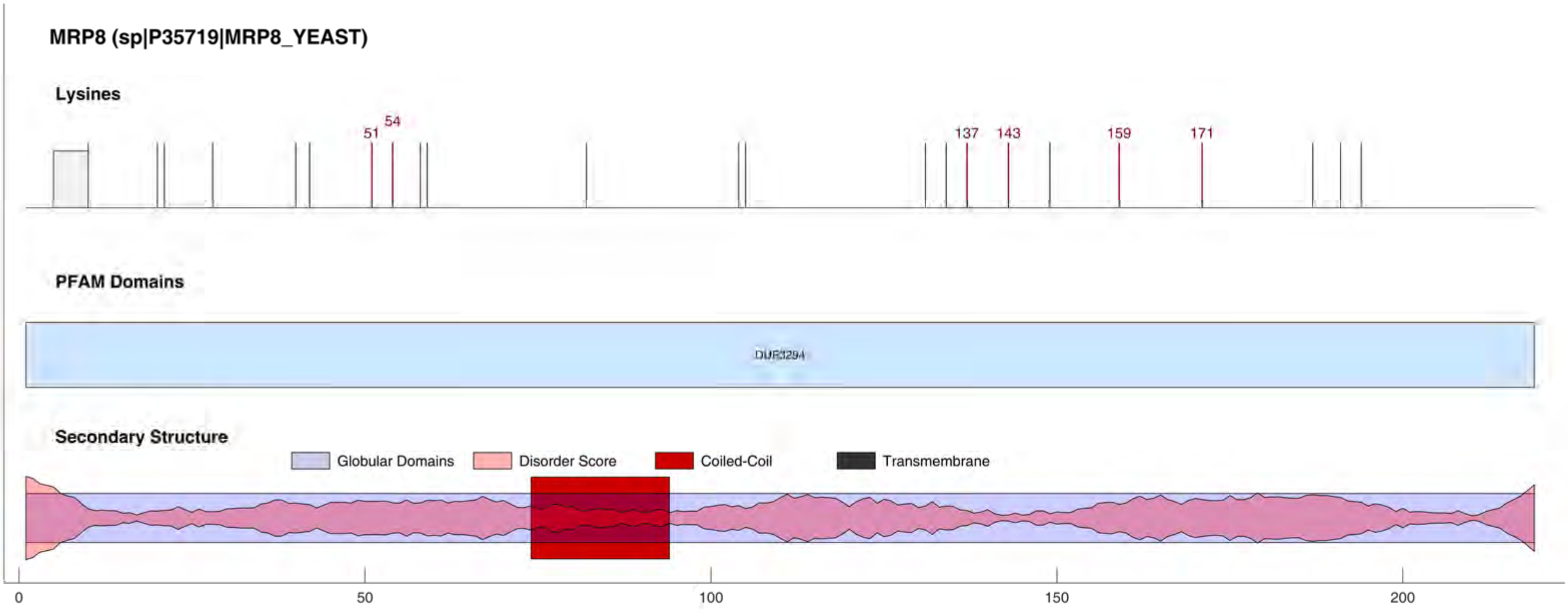

MRP8 (sp|P35719|MRP8_YEAST)
Lysines
51
54
137
143
159
171
PFAM Domains
DUF3294
Secondary Structure
Globular Domains
Disorder Score
Coiled-Coil
Transmembrane
0
50
100
150
200

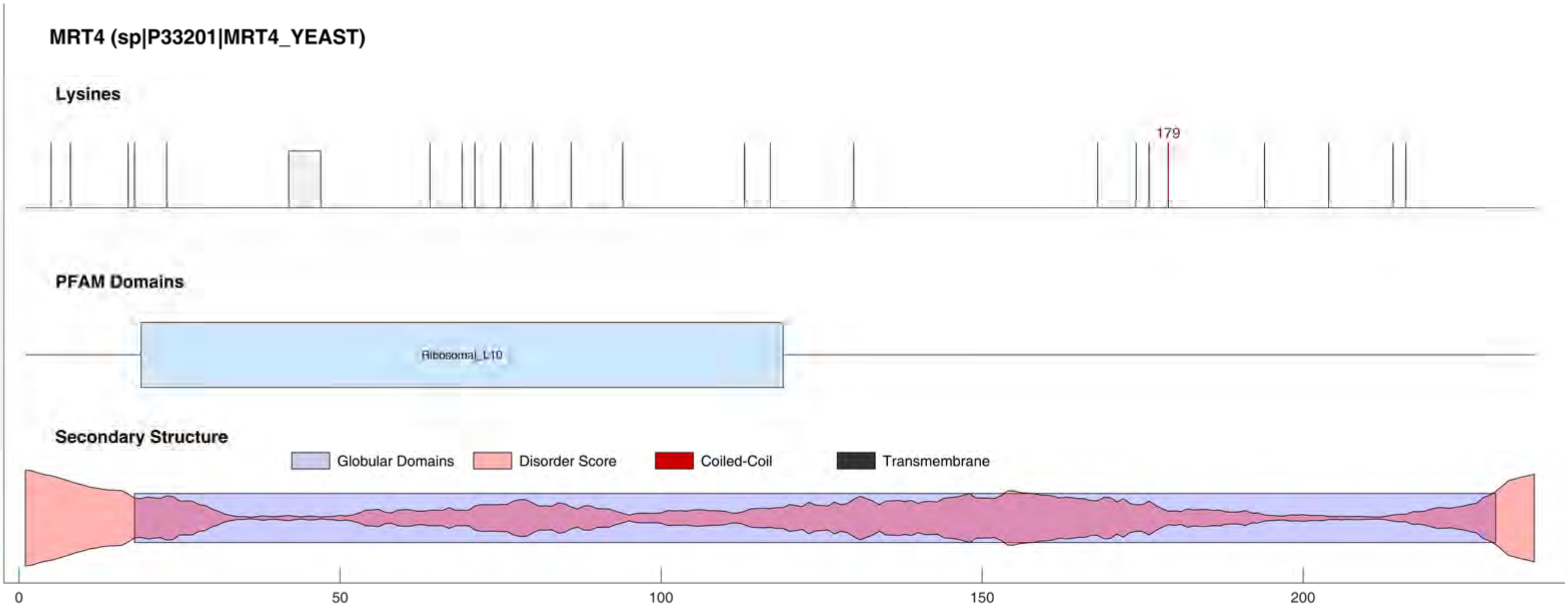

MRT4 (sp|P33201|MRT4_YEAST)
Lysines
179
PFAM Domains
Ribosomal_L10
Secondary Structure
Globular Domains
Disorder Score
Coiled-Coil
Transmembrane
0
50
100
150
200

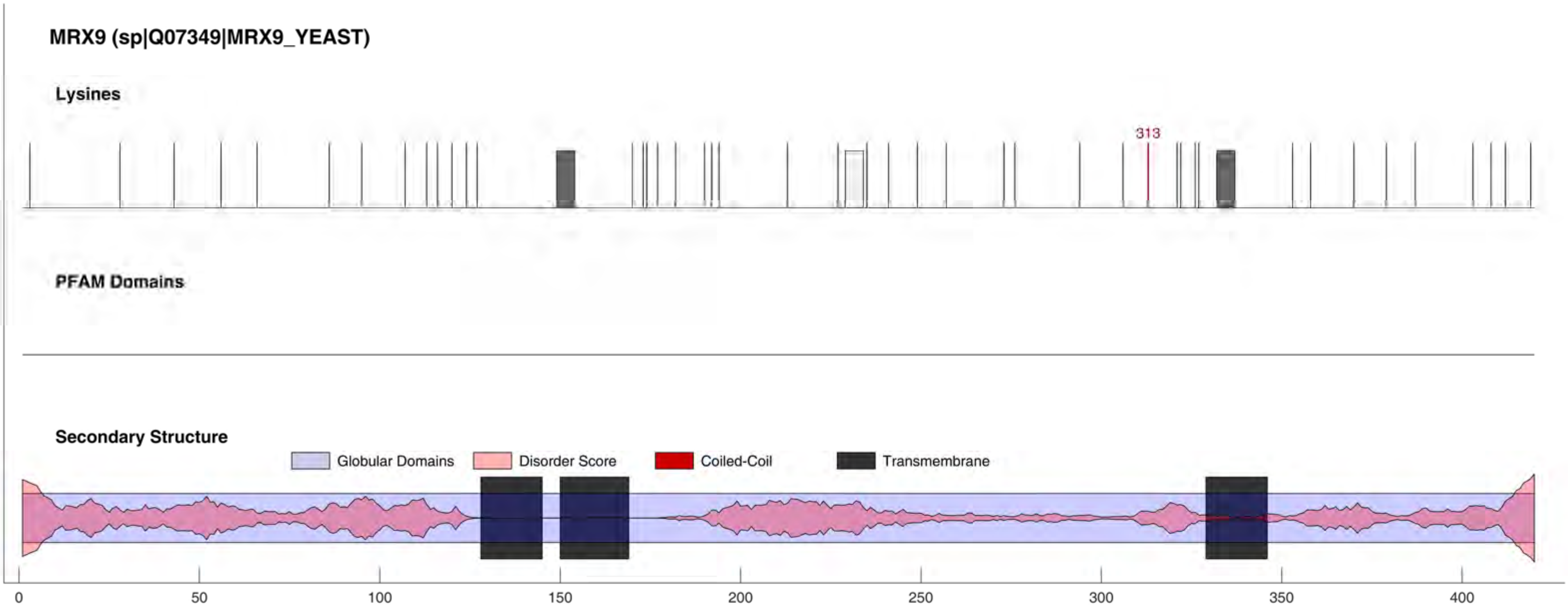

MRX9 (sp|Q07349|MRX9_YEAST)
Lysines
313
PFAM Domains
Secondary Structure
Globular Domains
Disorder Score
Coiled-Coil
Transmembrane
0
50
100
150
200
250
300
350
400

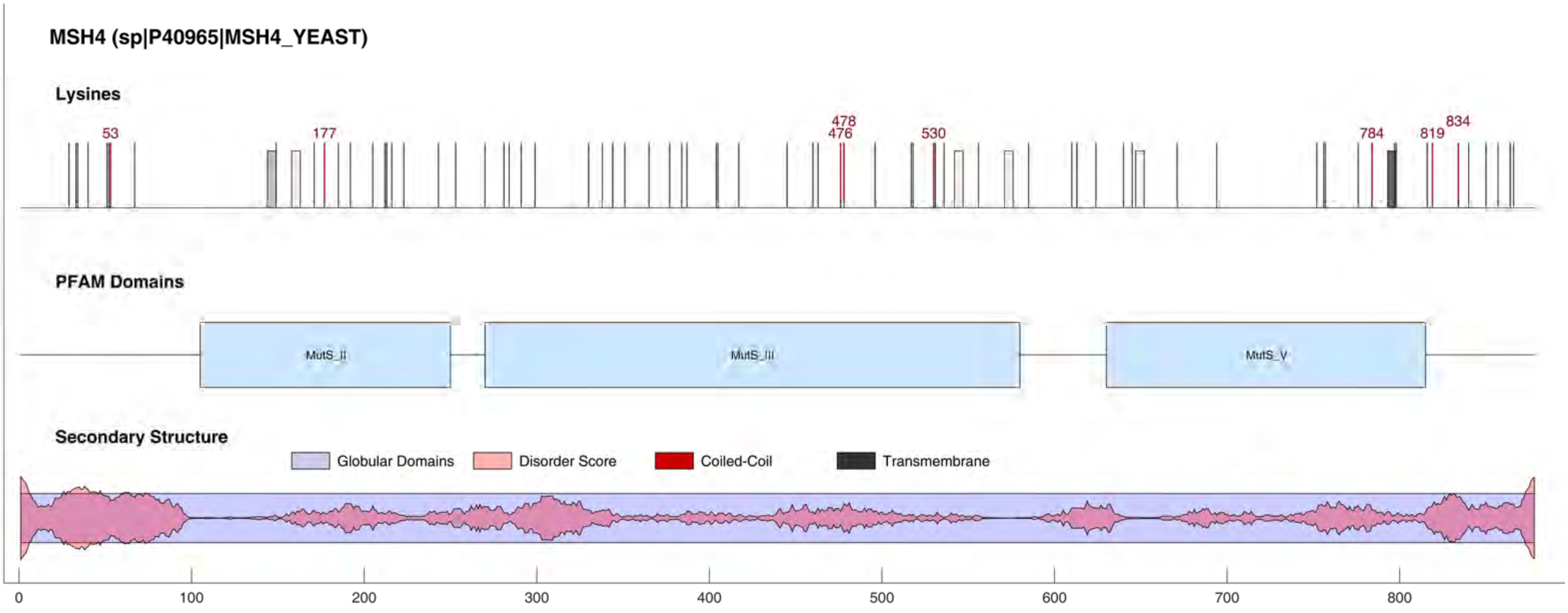

MSH4 (sp|P40965|MSH4_YEAST)
Lysines
53
177
478
476
530
784
819
834
PFAM Domains
MutS_II
MutS_III
MutS_V
Secondary Structure
Globular Domains
Disorder Score
Coiled-Coil
Transmembrane
0
100
200
300
400
500
600
700
800

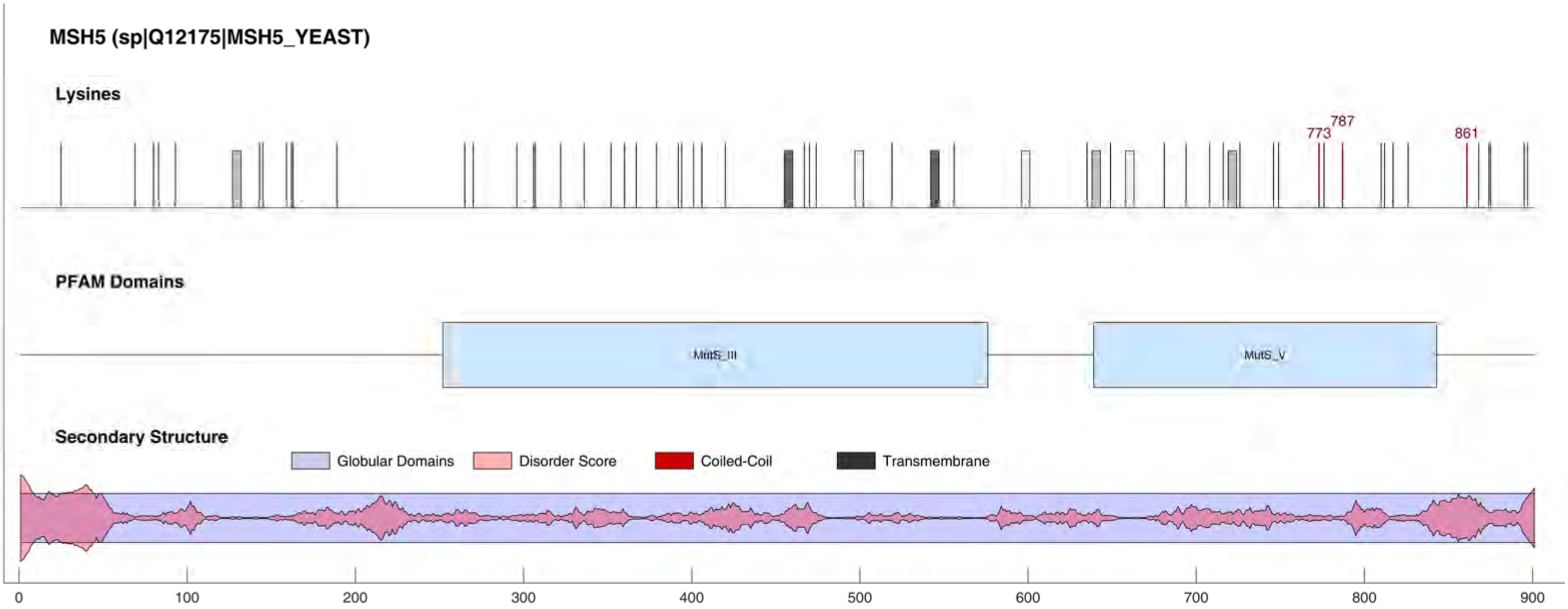

MSH5 (sp|Q12175|MSH5_YEAST)
Lysines
773
787
861
PFAM Domains
MutS_III
MutS_V
Secondary Structure
Globular Domains
Disorder Score
Coiled-Coil
Transmembrane
0
100
200
300
400
500
600
700
800
900

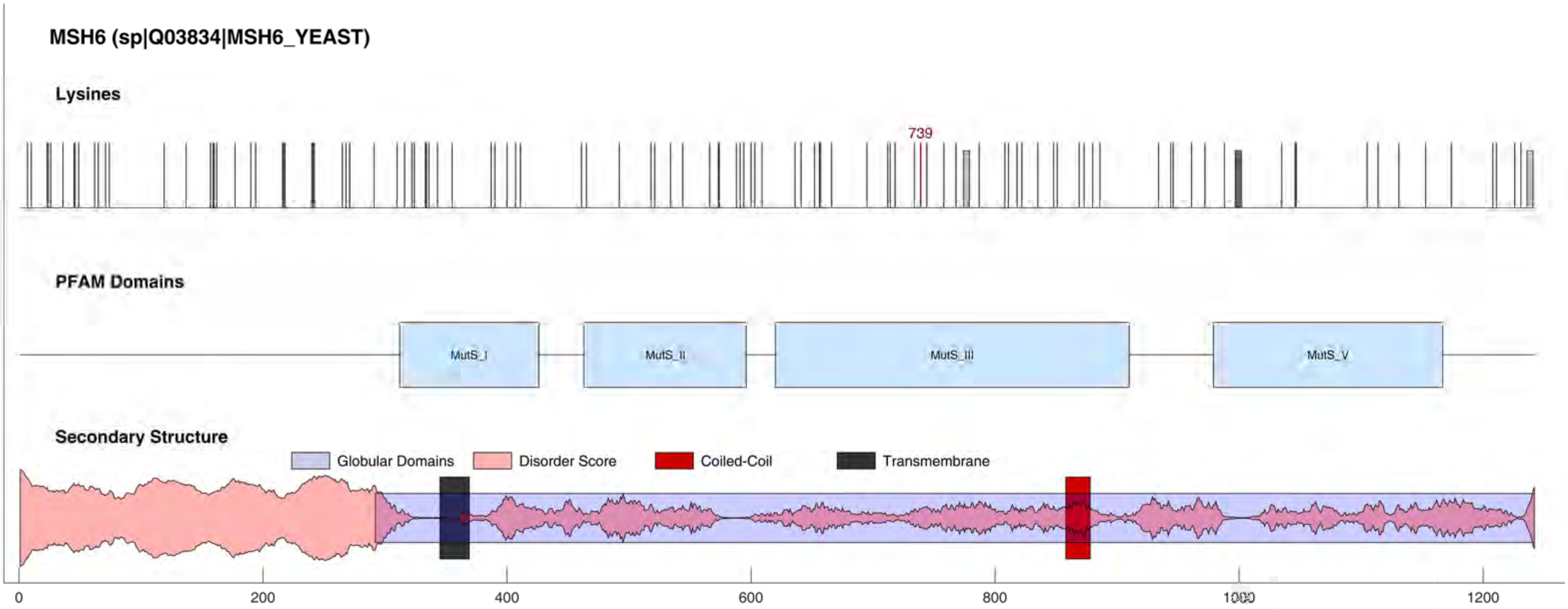
MSH6 (sp|Q03834|MSH6_YEAST)
Lysines
739
PFAM Domains
MutS_I
MutS_II
MutS_III
MutS_V
Secondary Structure
Globular Domains
Disorder Score
Coiled-Coil
Transmembrane
0
200
400
600
800
1000
1200

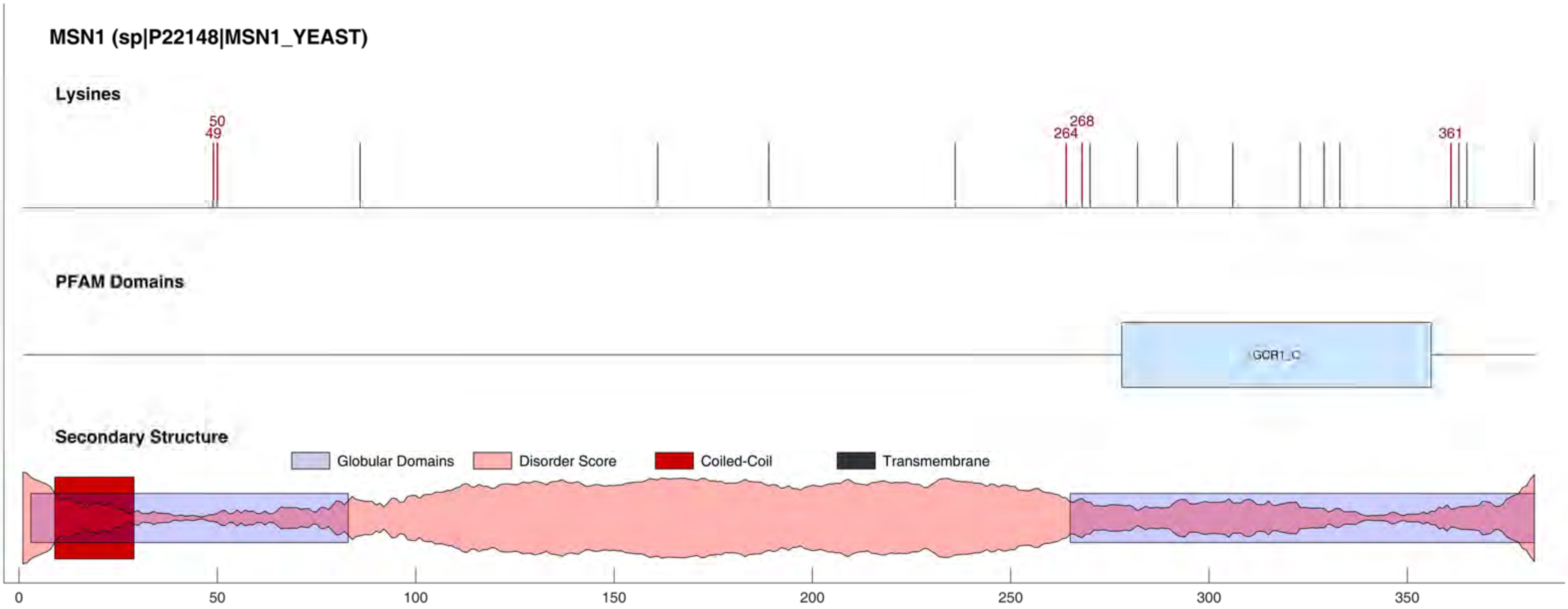

MSN1 (sp|P22148|MSN1_YEAST)
Lysines
50
49
268
264
361
PFAM Domains
GCR1_C
Secondary Structure
Globular Domains
Disorder Score
Coiled-Coil
Transmembrane
0
50
100
150
200
250
300
350

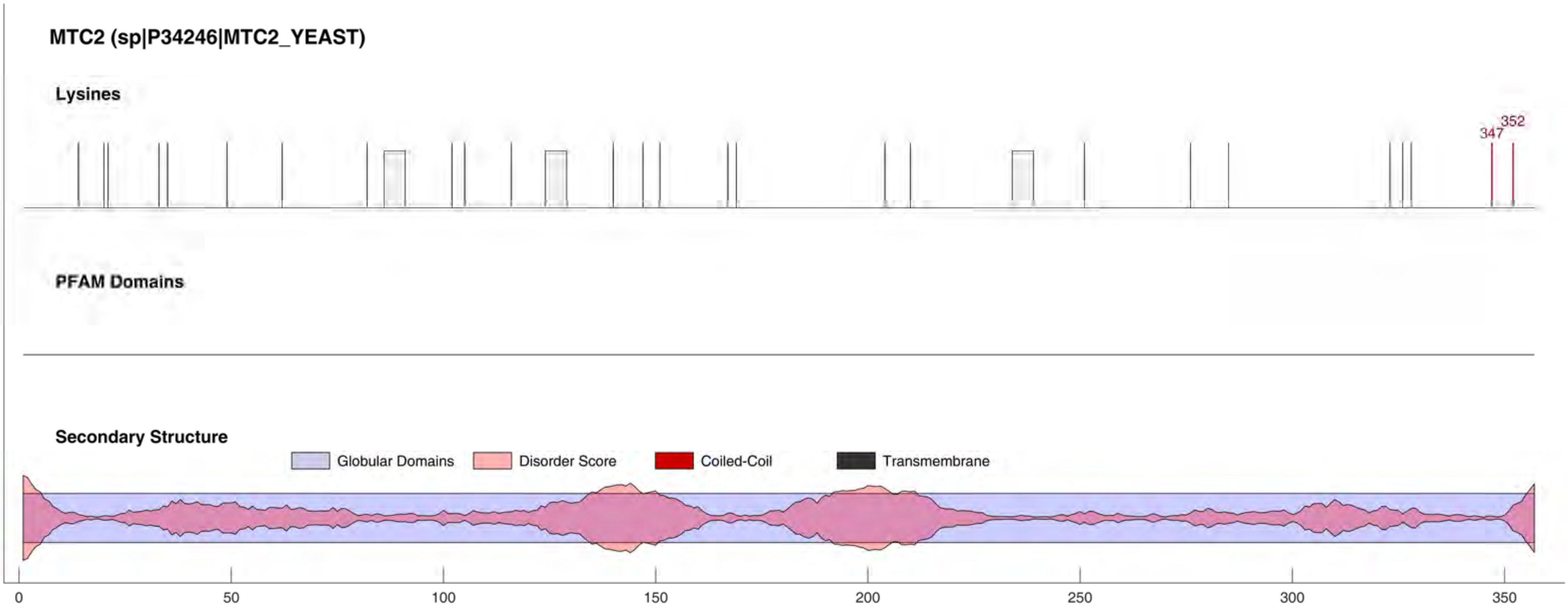
MTC2 (sp|P34246|MTC2_YEAST)
Lysines
352
347
PFAM Domains
Secondary Structure
Globular Domains
Disorder Score
Coiled-Coil
Transmembrane
0
50
100
150
200
250
300
350

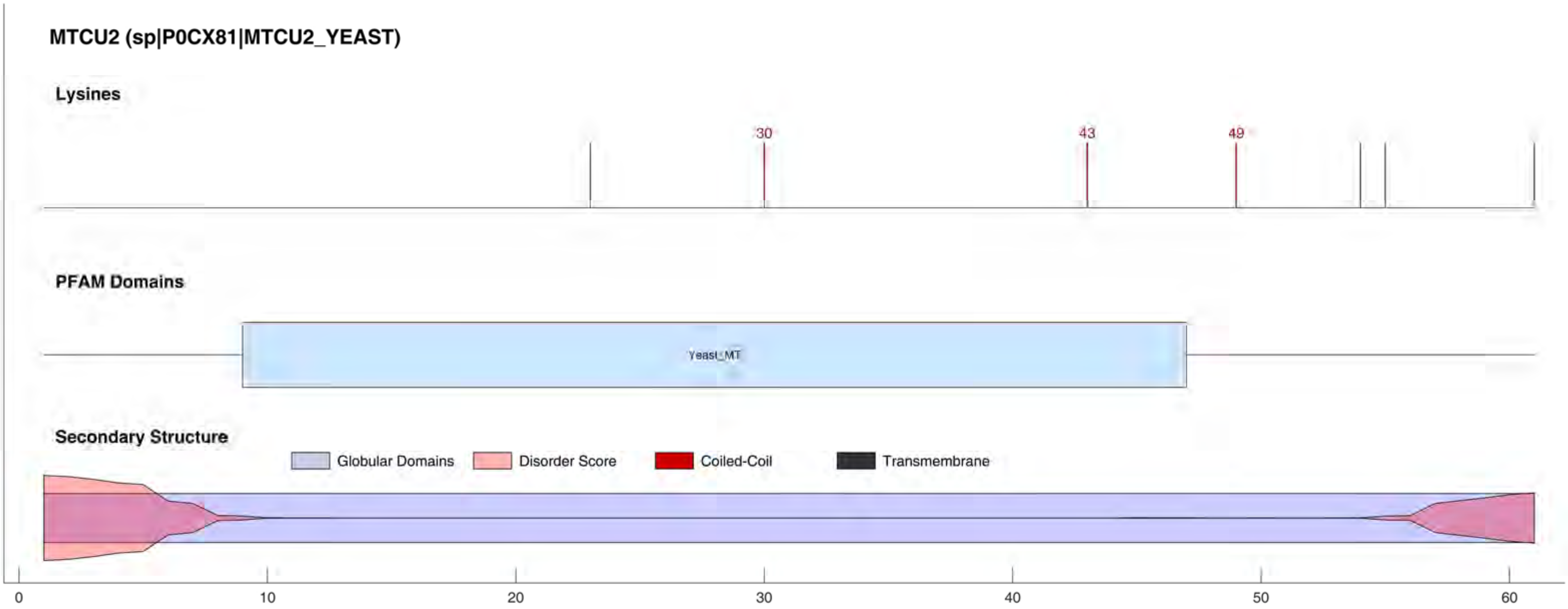
MTCU2 (sp|P0CX81|MTCU2_YEAST)
Lysines
30
43
49
PFAM Domains
Yeast_MT
Secondary Structure
Globular Domains
Disorder Score
Coiled-Coil
Transmembrane
0
10
20
30
40
50
60

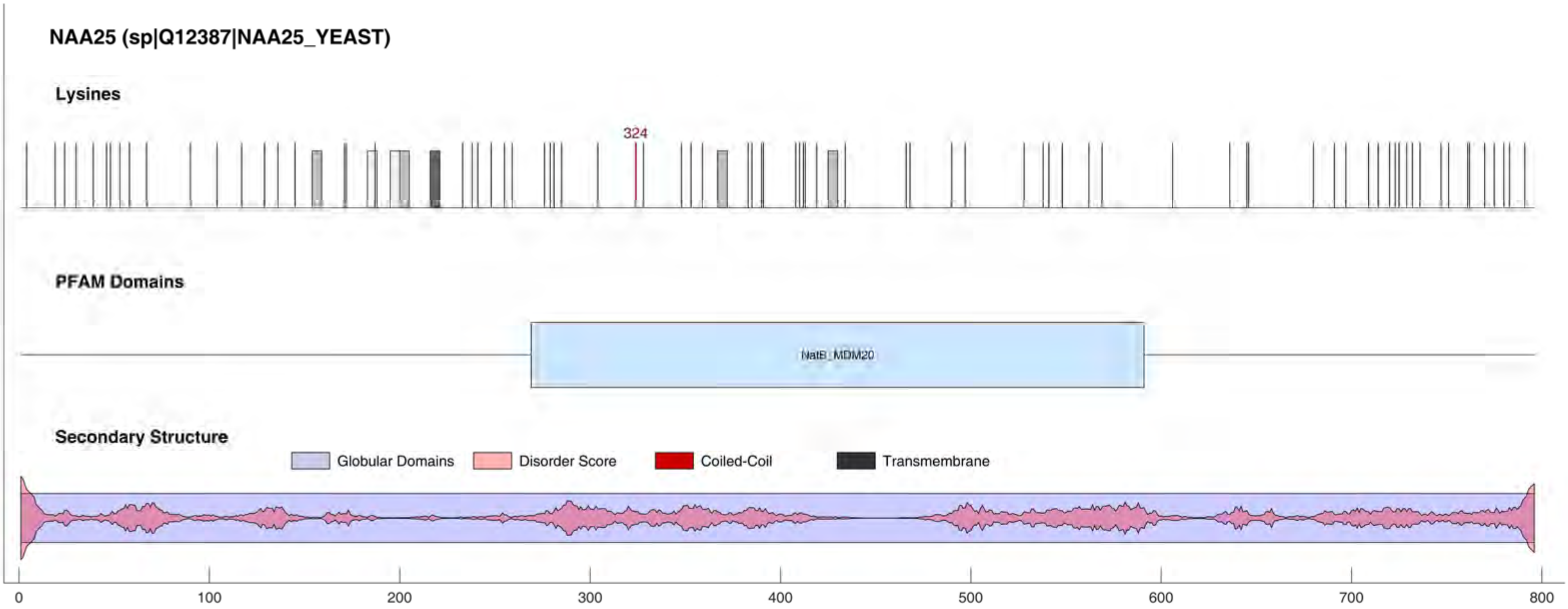

NAA25 (sp|Q12387|NAA25_YEAST)
Lysines
324
PFAM Domains
NatB_MDM20
Secondary Structure
Globular Domains
Disorder Score
Coiled-Coil
Transmembrane
0
100
200
300
400
500
600
700
800

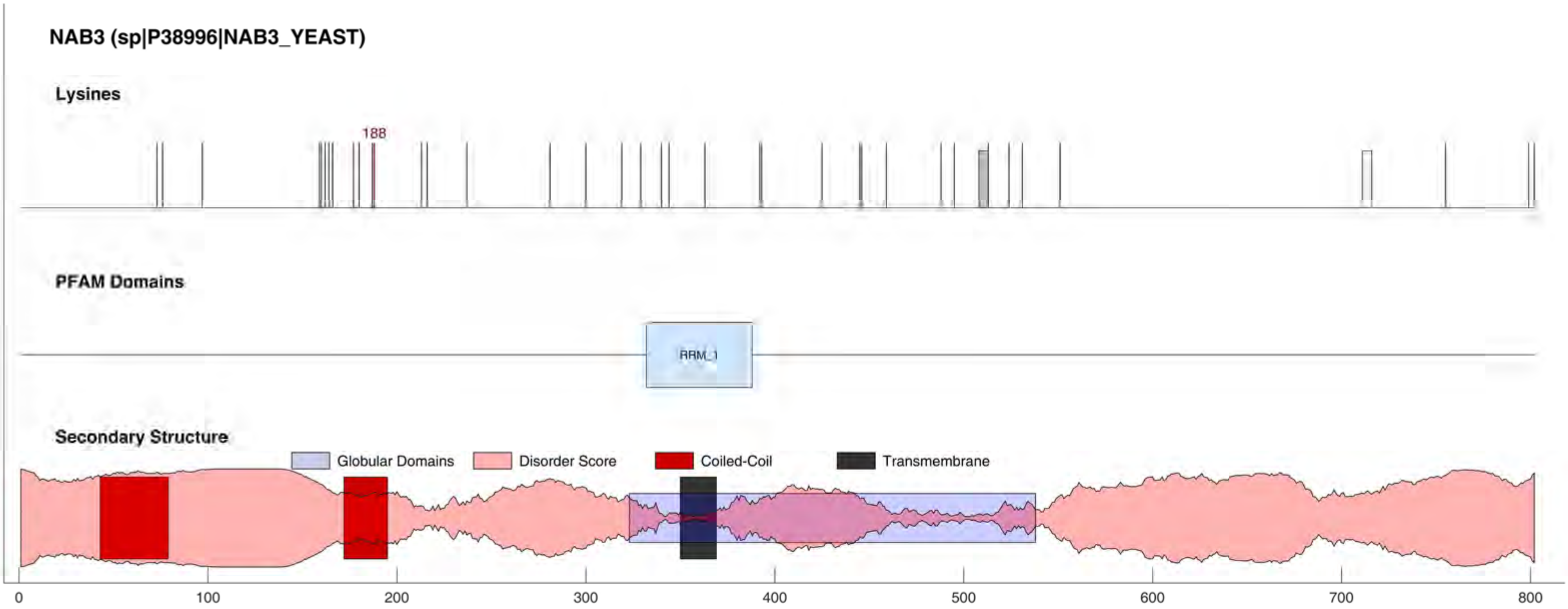
NAB3 (sp|P38996|NAB3_YEAST)
Lysines
188
PFAM Domains
RRM_1
Secondary Structure
Globular Domains
Disorder Score
Coiled-Coil
Transmembrane
0
100
200
300
400
500
600
700
800

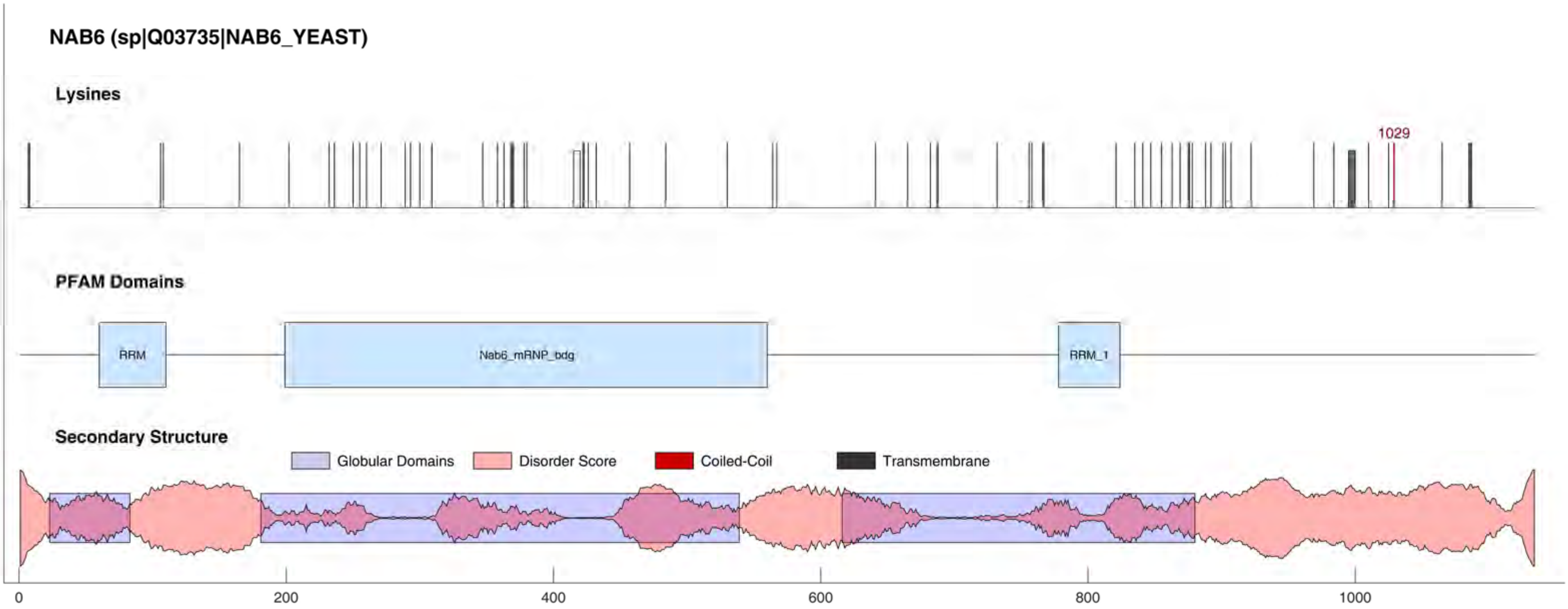
NAB6 (sp|Q03735|NAB6_YEAST)
Lysines
1029
PFAM Domains
RRM
Nab6_mRNP_bdg
RRM_1
Secondary Structure
Globular Domains
Disorder Score
Coiled-Coil
Transmembrane
0
200
400
600
800
1000

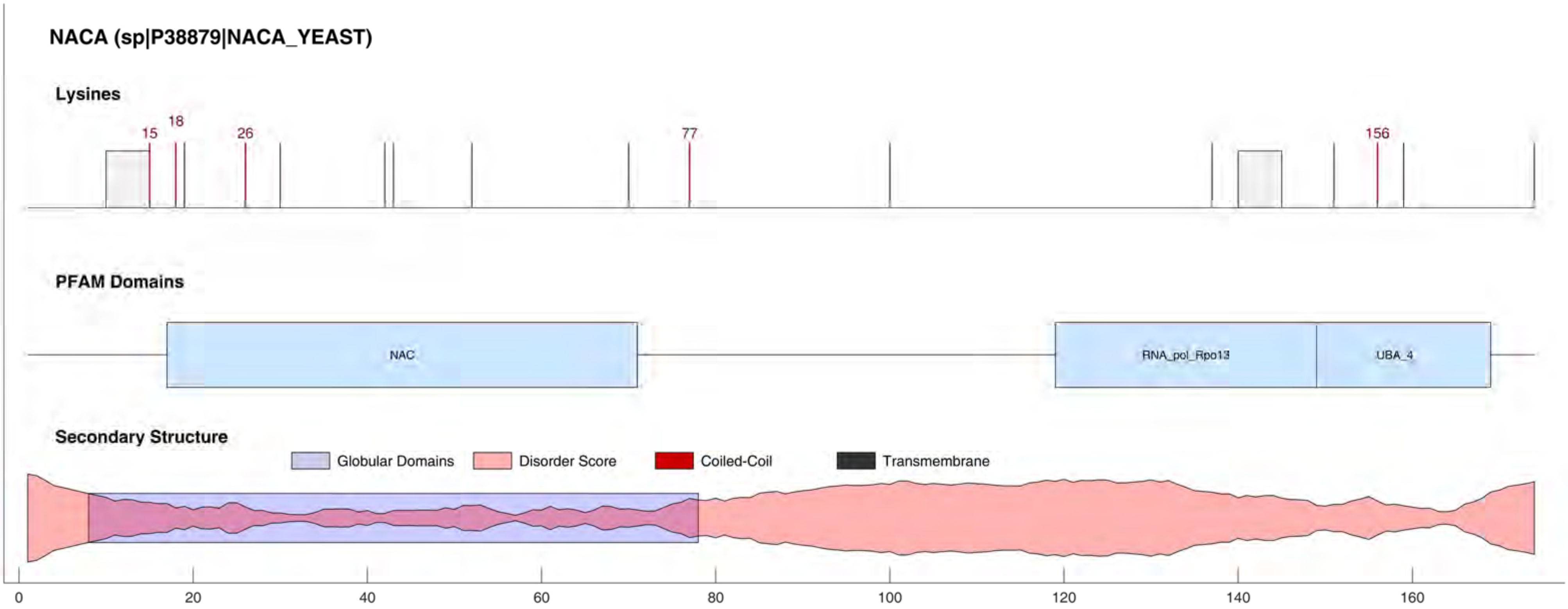
NACA (sp|P38879|NACA_YEAST)
Lysines
15
18
26
77
156
PFAM Domains
NAC
RNA_pol_Rpo13
UBA_4
Secondary Structure
Globular Domains
Disorder Score
Coiled-Coil
Transmembrane
0
20
40
60
80
100
120
140
160

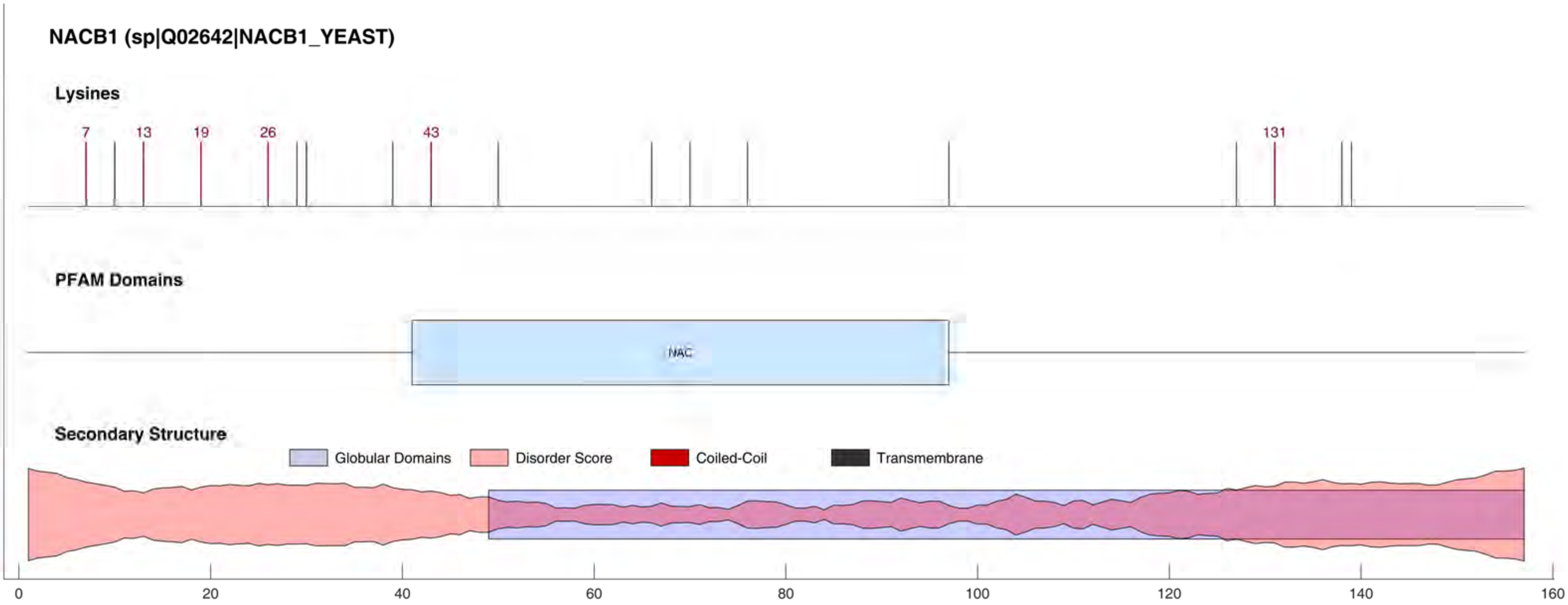

NACB1 (sp|Q02642|NACB1_YEAST)
Lysines
7
13
19
26
43
131
PFAM Domains
NAC
Secondary Structure
Globular Domains
Disorder Score
Coiled-Coil
Transmembrane
0
20
40
60
80
100
120
140
160

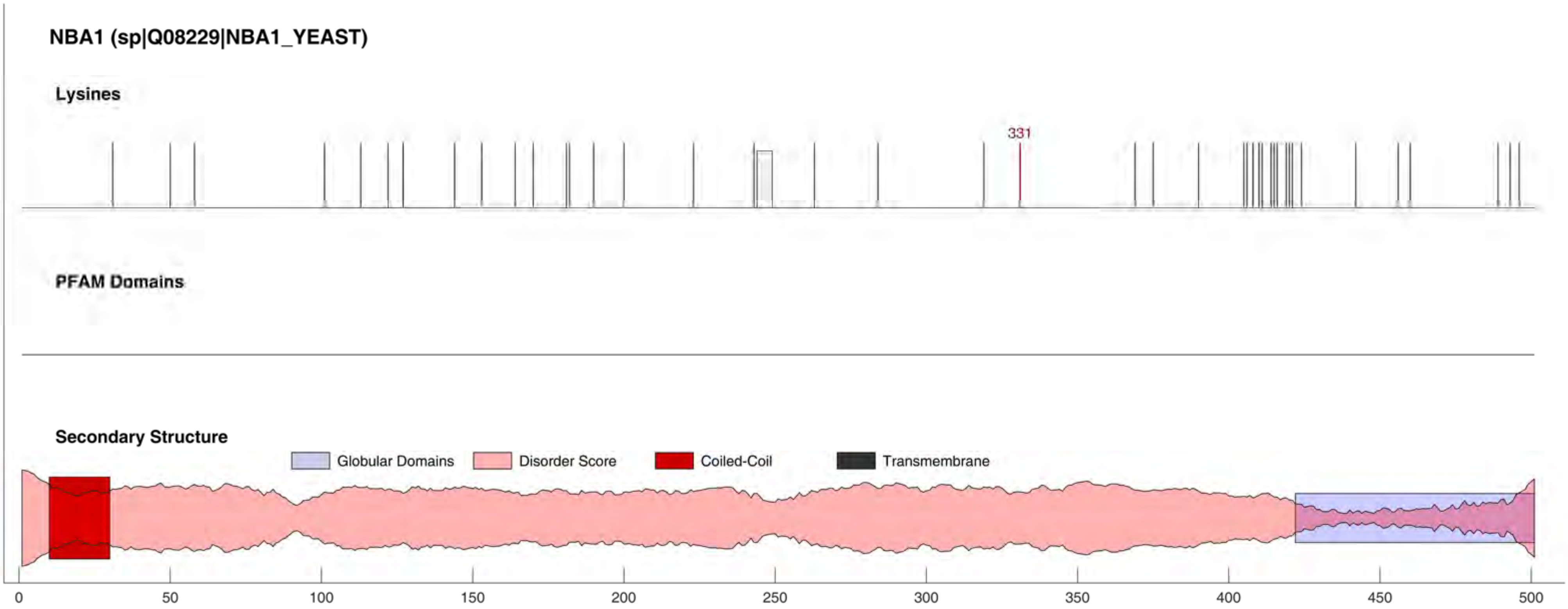
NBA1 (sp|Q08229|NBA1_YEAST)
Lysines
331
PFAM Domains
Secondary Structure
Globular Domains
Disorder Score
Coiled-Coil
Transmembrane
0
50
100
150
200
250
300
350
400
450
500

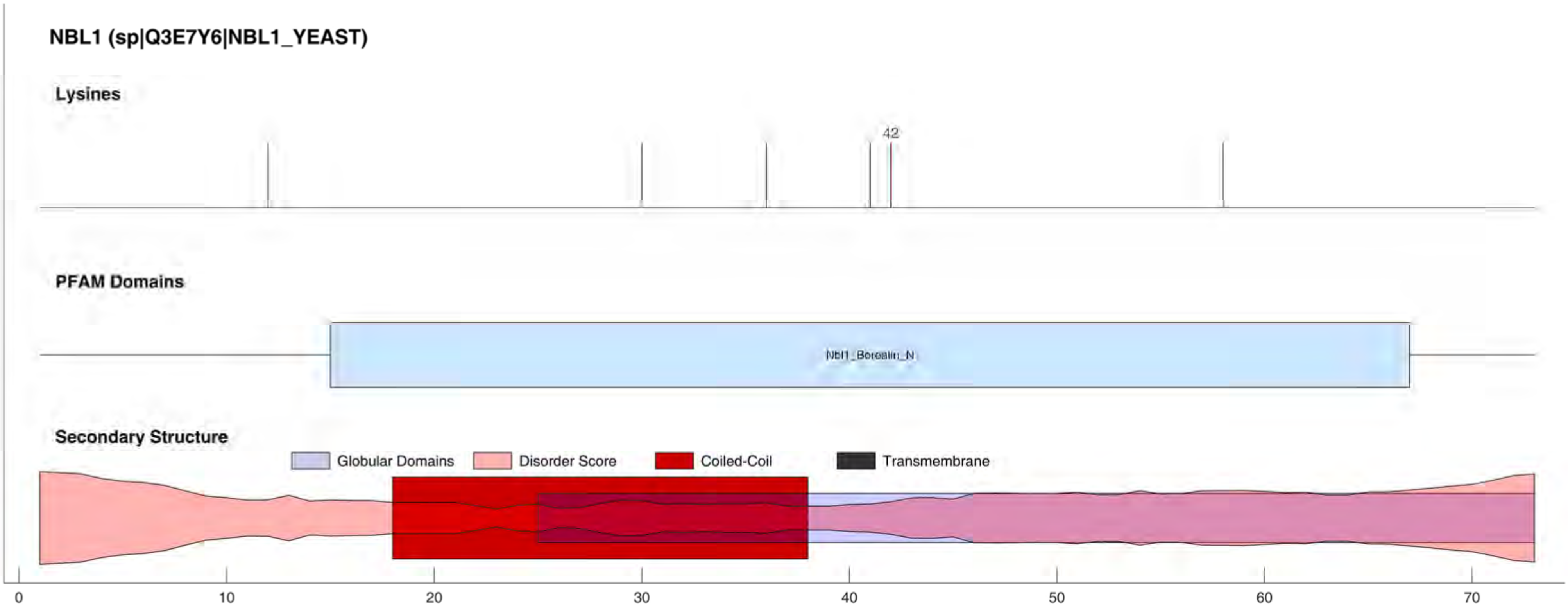

NBL1 (sp|Q3E7Y6|NBL1_YEAST)
Lysines
42
PFAM Domains
Nbl1_Borealin_N
Secondary Structure
Globular Domains
Disorder Score
Coiled-Coil
Transmembrane
0
10
20
30
40
50
60
70

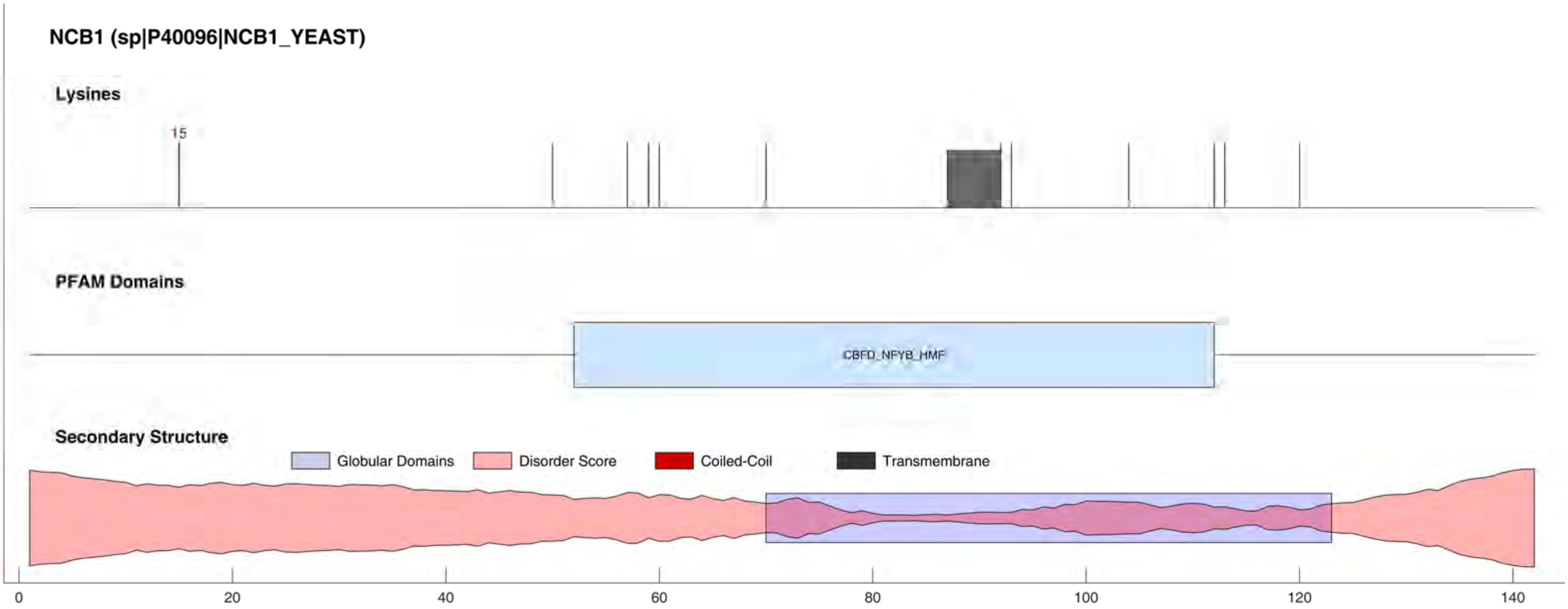
NCB1 (sp|P40096|NCB1_YEAST)
Lysines
15
PFAM Domains
CBFD_NFYB_HMF
Secondary Structure
Globular Domains
Disorder Score
Coiled-Coil
Transmembrane
0
20
40
60
80
100
120
140

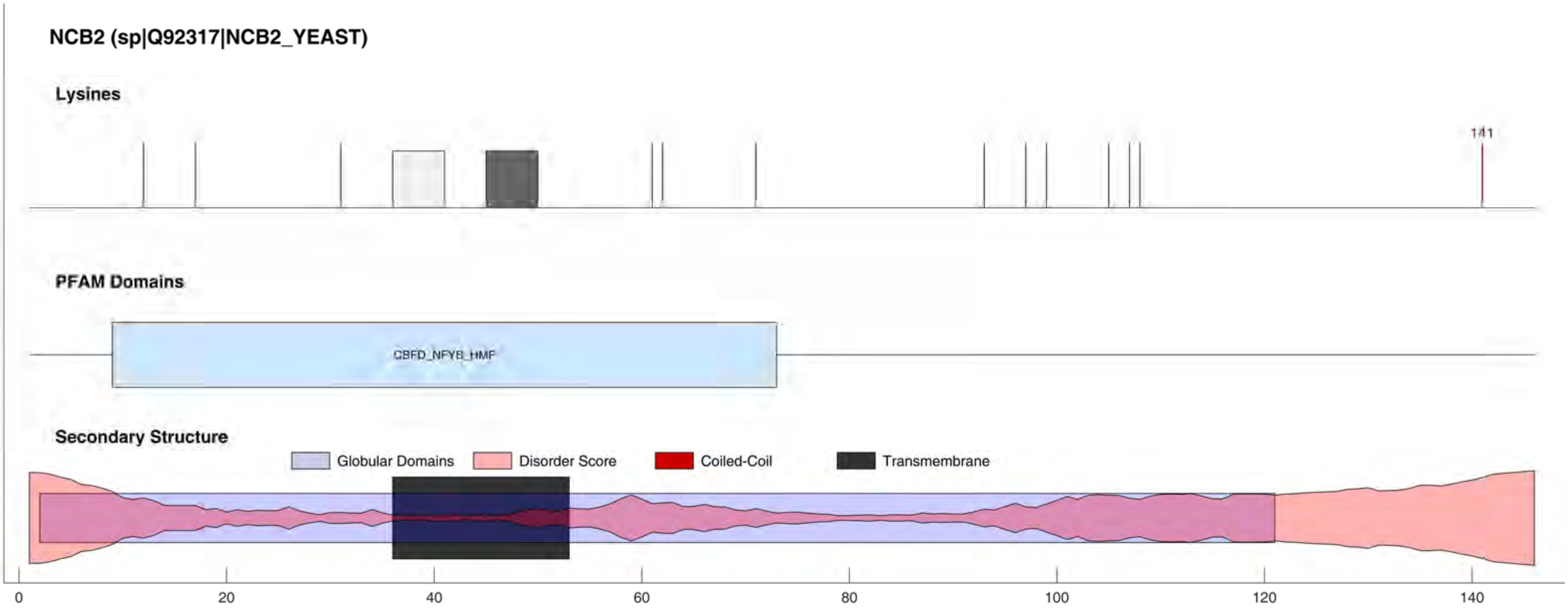
NCB2 (sp|Q92317|NCB2_YEAST)
Lysines
141
PFAM Domains
CBFD_NFYB_HMF
Secondary Structure
Globular Domains
Disorder Score
Coiled-Coil
Transmembrane
0
20
40
60
80
100
120
140

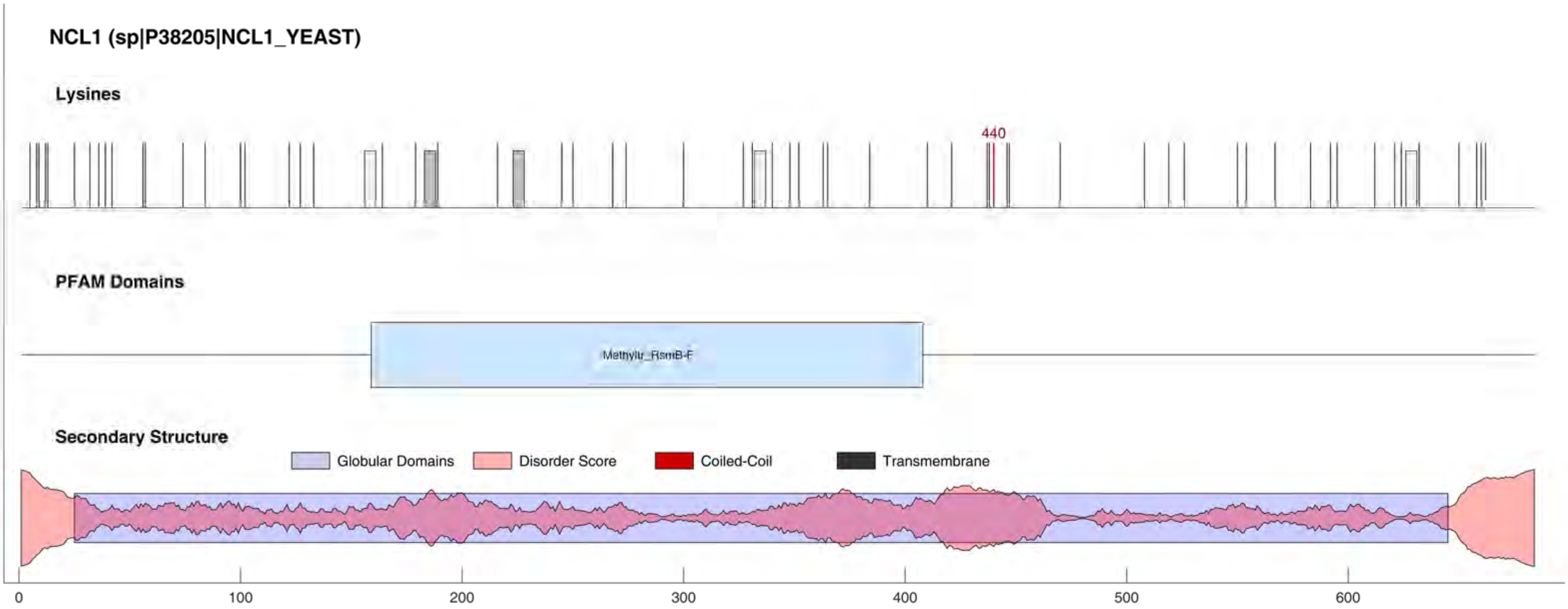
NCL1 (sp|P38205|NCL1_YEAST)
Lysines
440
PFAM Domains
Methyltr_RsmB-F
Secondary Structure
Globular Domains
Disorder Score
Coiled-Coil
Transmembrane
0
100
200
300
400
500
600

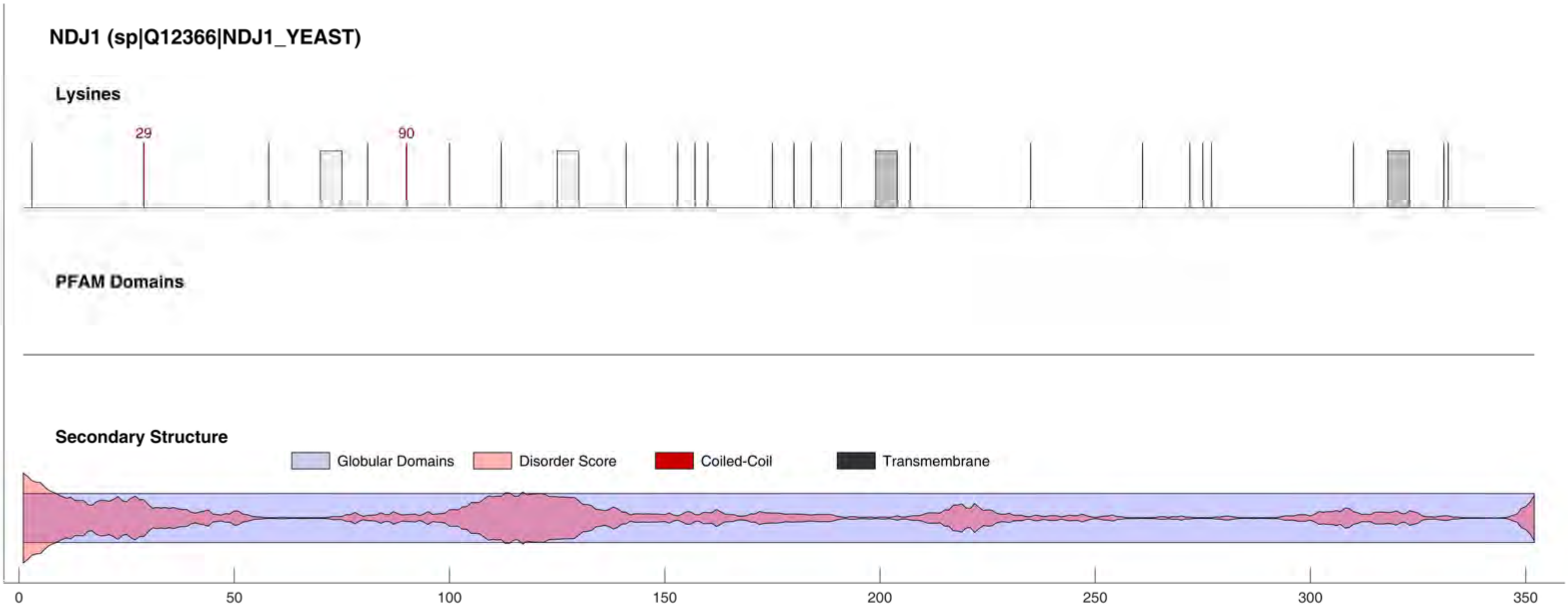
NDJ1 (sp|Q12366|NDJ1_YEAST)
Lysines
29
90
PFAM Domains
Secondary Structure
Globular Domains
Disorder Score
Coiled-Coil
Transmembrane
0
50
100
150
200
250
300
350

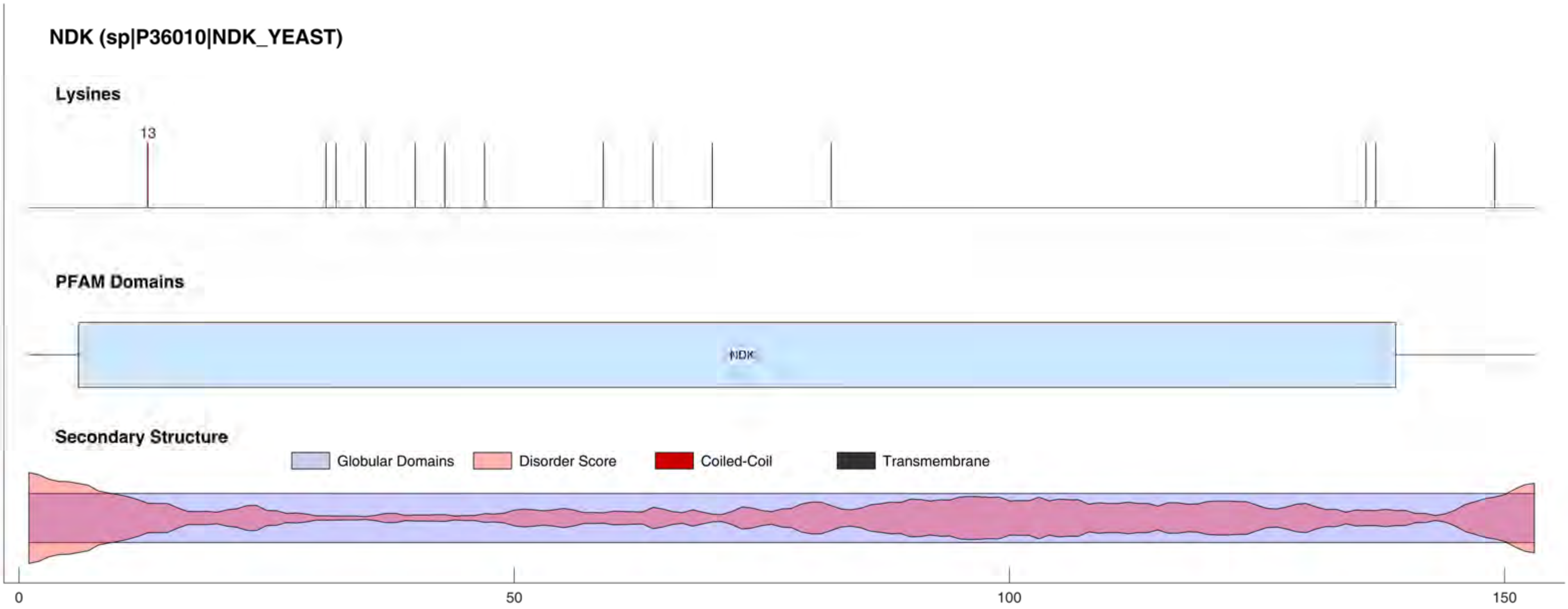
NDK (sp|P36010|NDK_YEAST)
Lysines
13
PFAM Domains
NDK
Secondary Structure
Globular Domains
Disorder Score
Coiled-Coil
Transmembrane
0
50
100
150

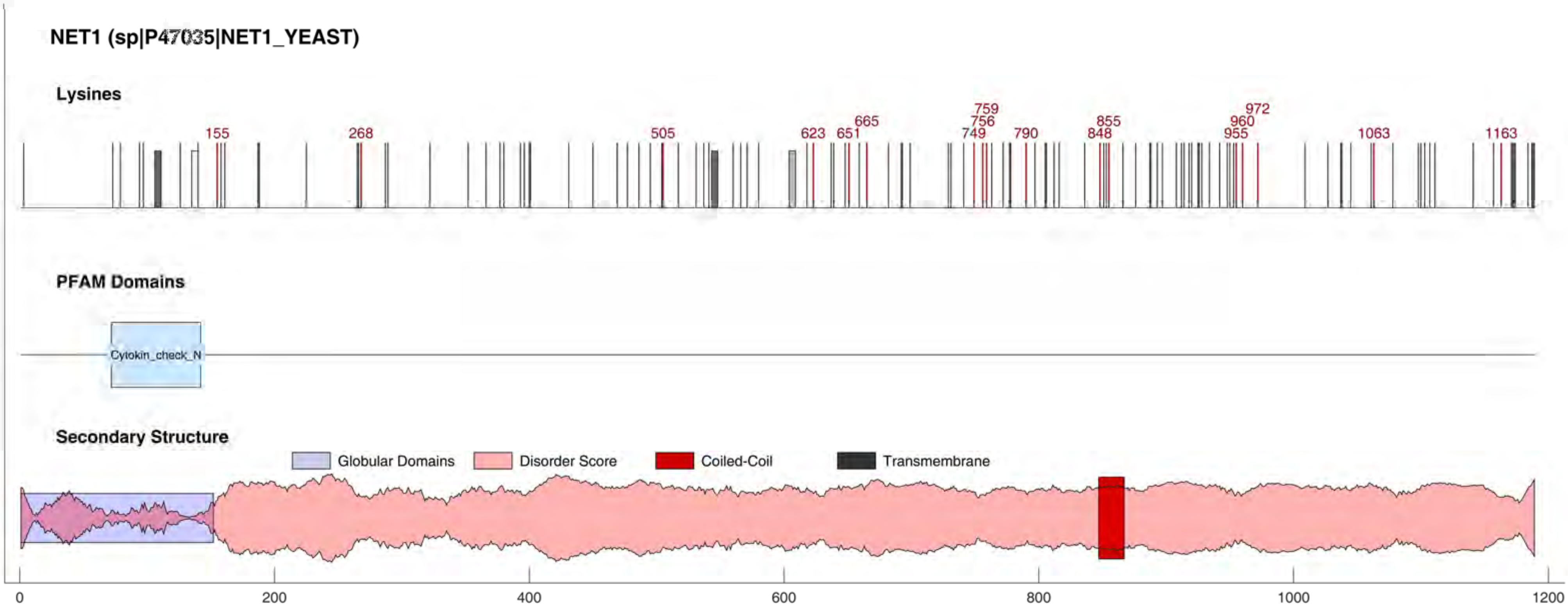
NET1 (sp|P47035|NET1_YEAST)
Lysines
155
268
505
623
651
665
749
756
759
790
848
855
955
960
972
1063
1163
PFAM Domains
Cytokin_check_N
Secondary Structure
Globular Domains
Disorder Score
Coiled-Coil
Transmembrane
0
200
400
600
800
1000
1200

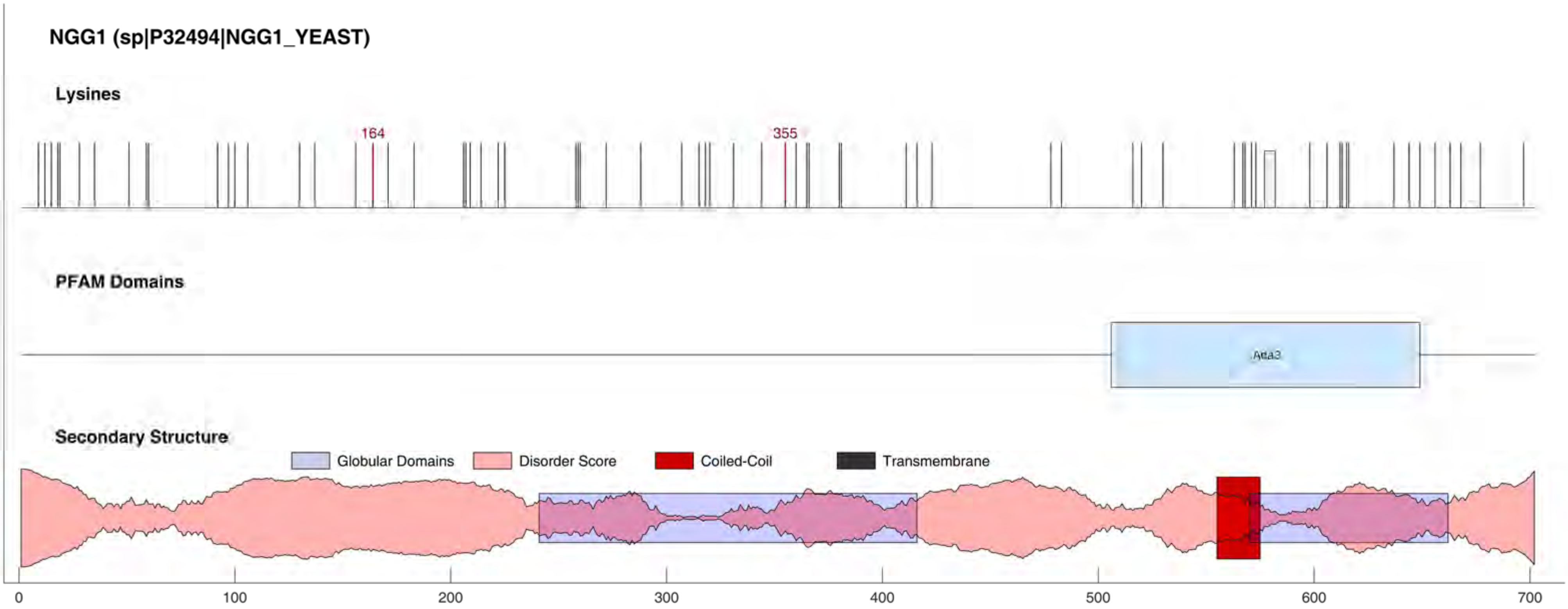
NGG1 (sp|P32494|NGG1_YEAST)
Lysines
164
355
PFAM Domains
Ada3
Secondary Structure
Globular Domains
Disorder Score
Coiled-Coil
Transmembrane
0
100
200
300
400
500
600
700

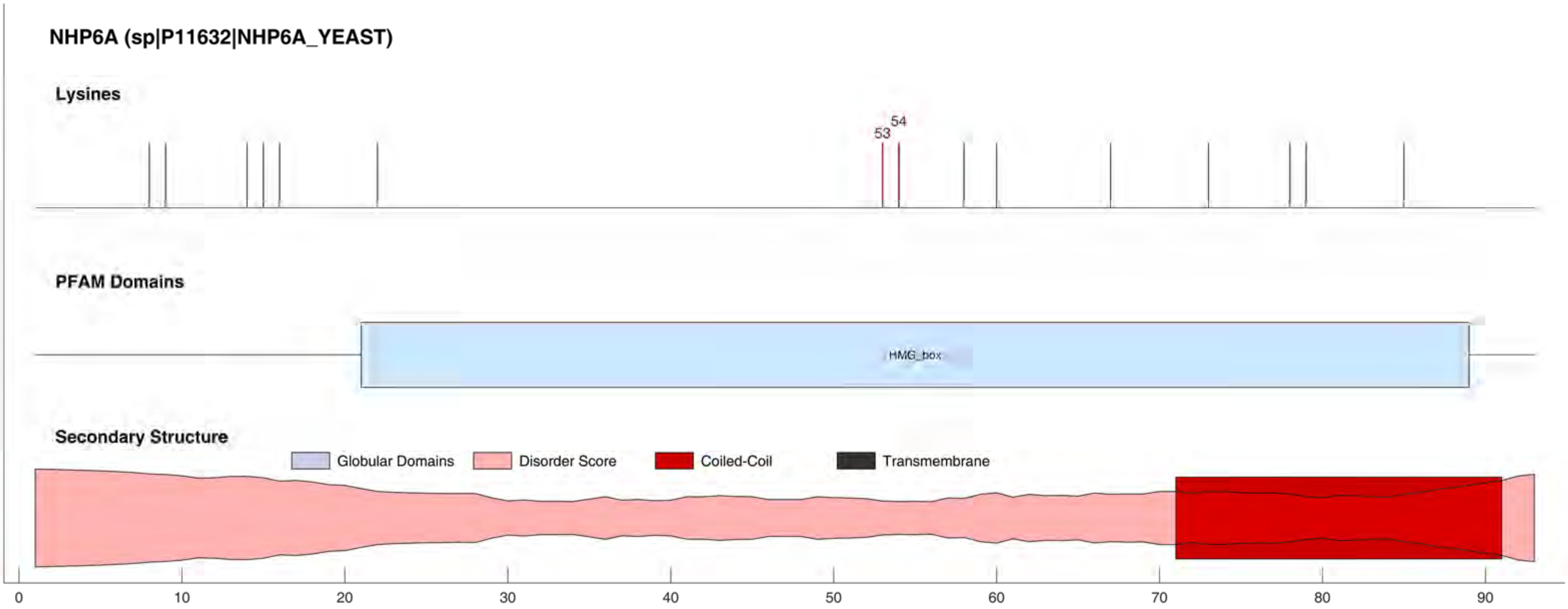
NHP6A (sp|P11632|NHP6A_YEAST)
Lysines
53
54
PFAM Domains
HMG_box
Secondary Structure
Globular Domains
Disorder Score
Coiled-Coil
Transmembrane
0
10
20
30
40
50
60
70
80
90

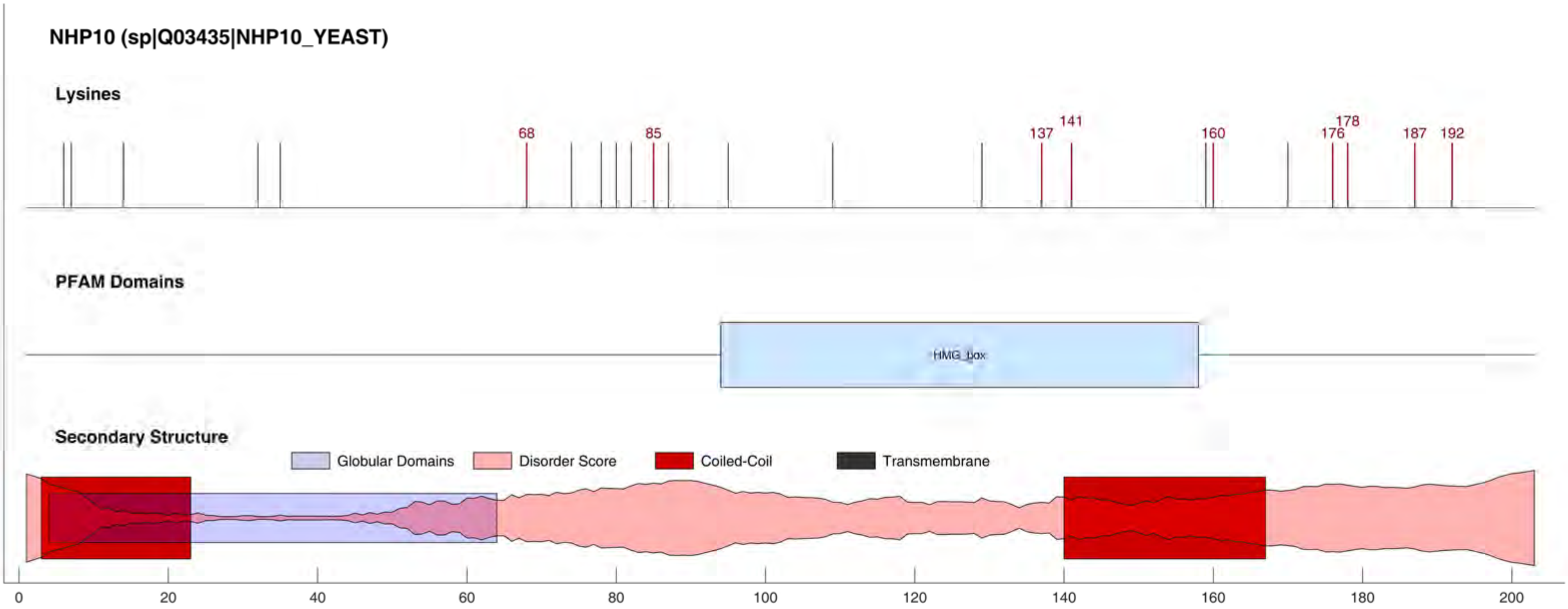
NHP10 (sp|Q03435|NHP10_YEAST)
Lysines
68
85
137
141
160
176
178
187
192
PFAM Domains
HMG_box
Secondary Structure
Globular Domains
Disorder Score
Coiled-Coil
Transmembrane
0
20
40
60
80
100
120
140
160
180
200

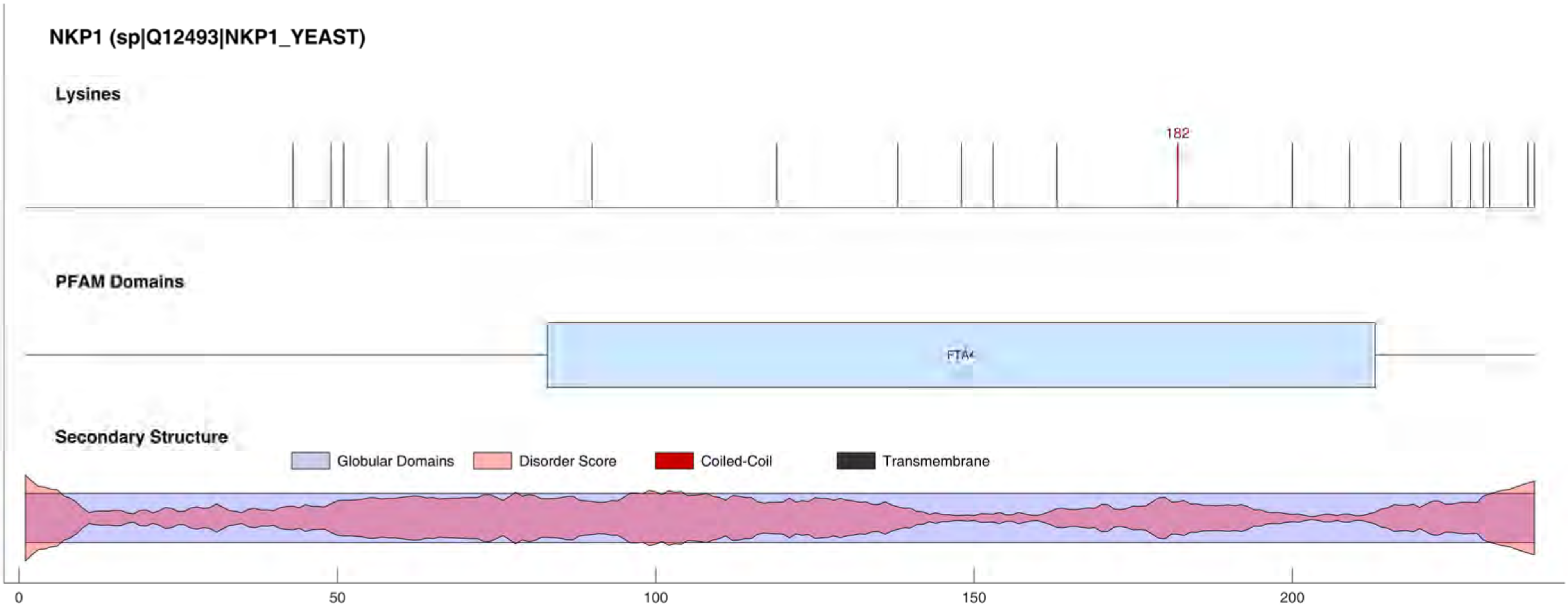
NKP1 (sp|Q12493|NKP1_YEAST)
Lysines
182
PFAM Domains
FTA4
Secondary Structure
Globular Domains
Disorder Score
Coiled-Coil
Transmembrane
0
50
100
150
200

NMA1 (sp|Q06178|NMA1_YEAST)

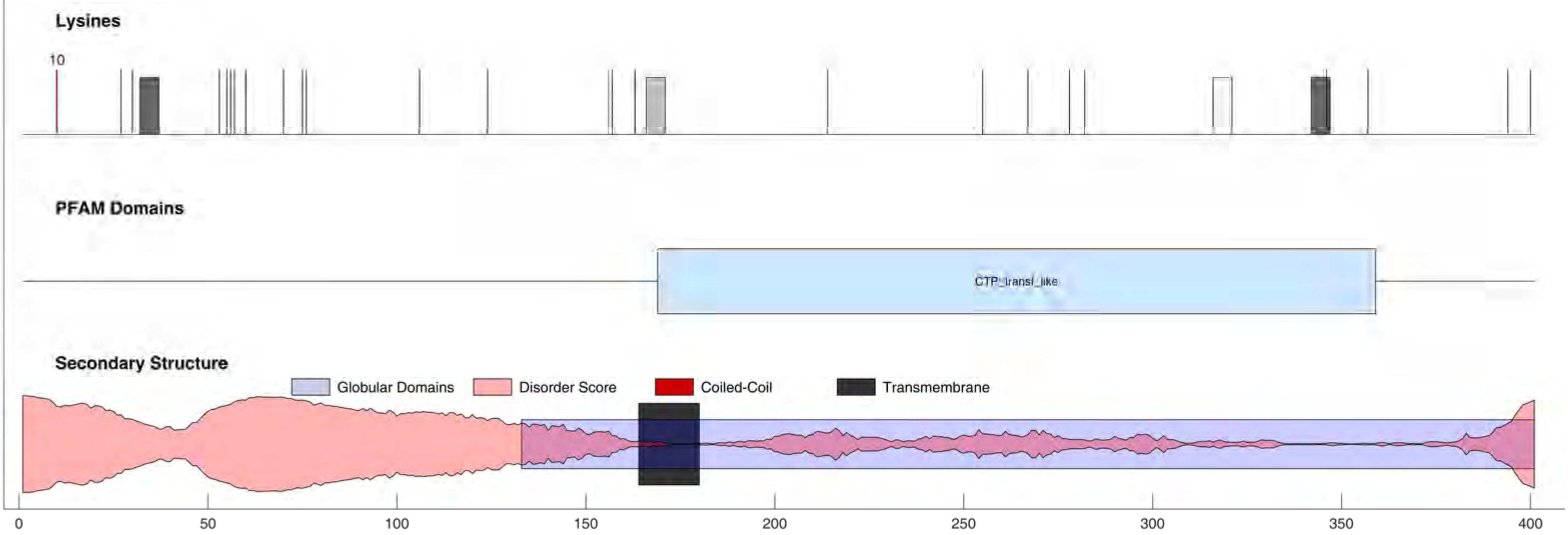

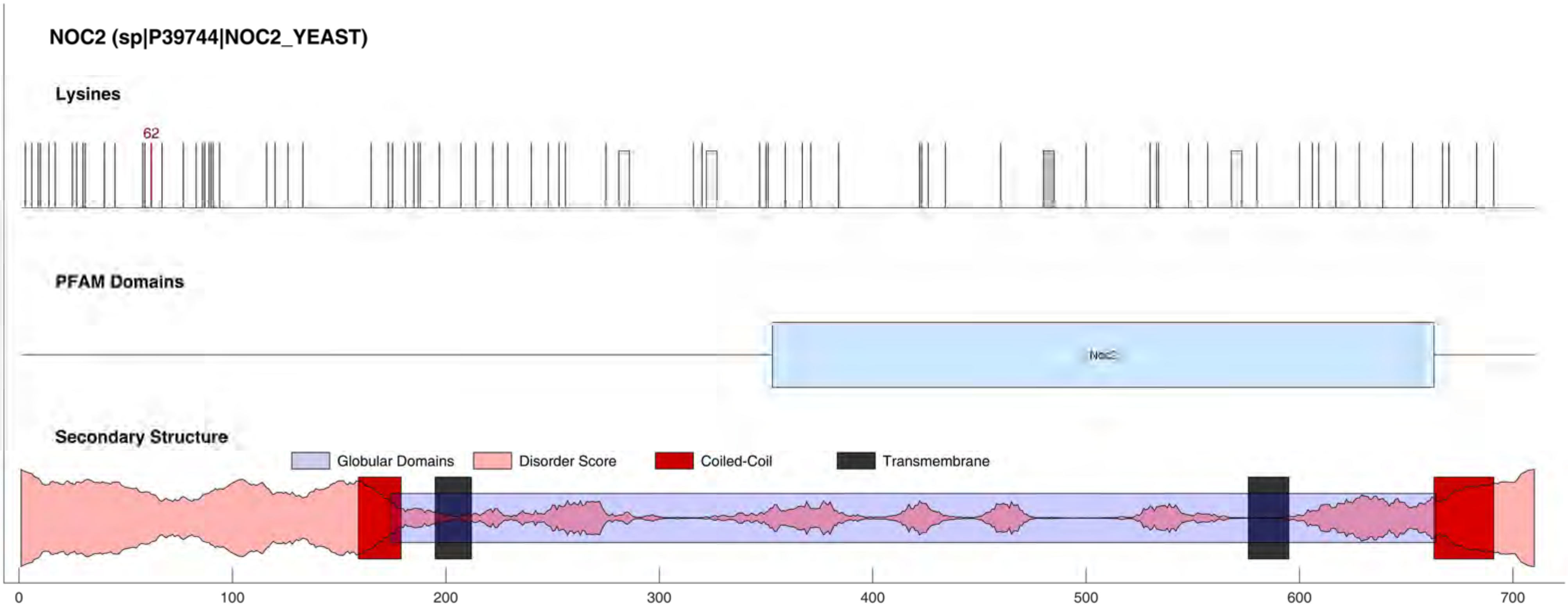
NOC2 (sp|P39744|NOC2_YEAST)
Lysines
62
PFAM Domains
Noc2
Secondary Structure
Globular Domains
Disorder Score
Coiled-Coil
Transmembrane
0
100
200
300
400
500
600
700

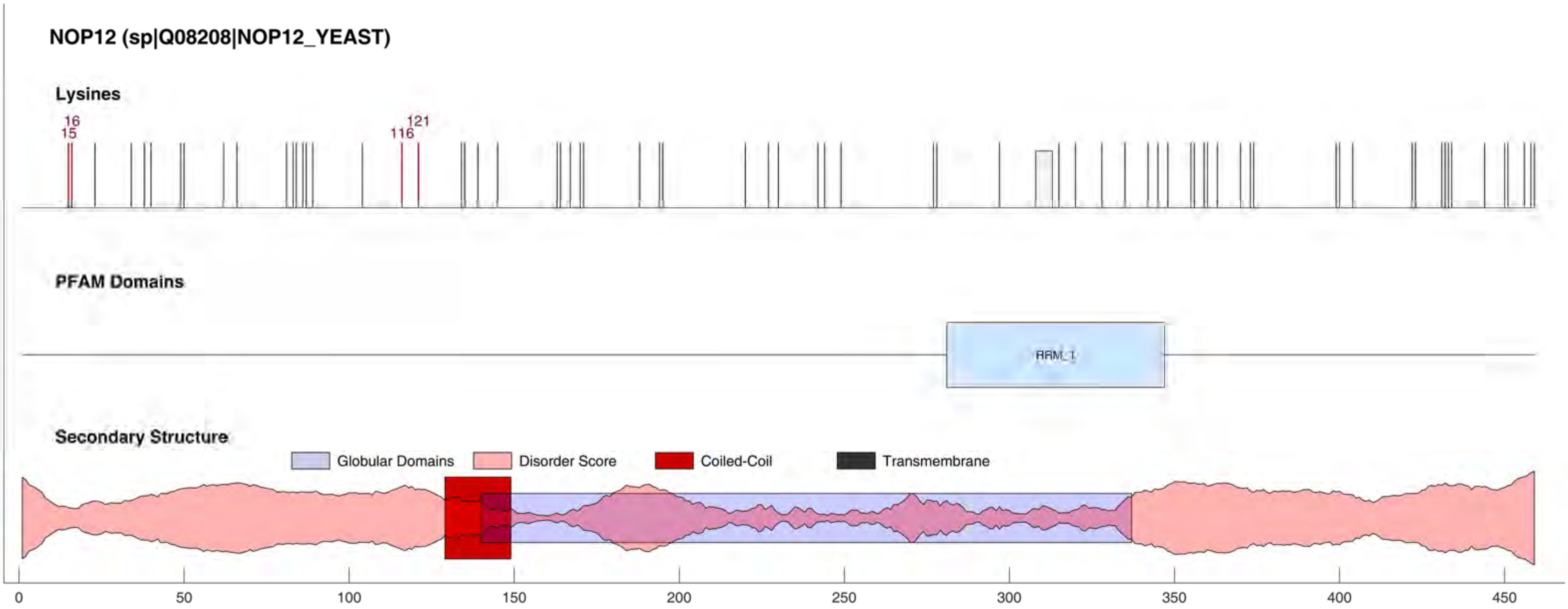
NOP12 (sp|Q08208|NOP12_YEAST)
Lysines
16
15
121
116
PFAM Domains
RRM_1
Secondary Structure
Globular Domains
Disorder Score
Coiled-Coil
Transmembrane
0
50
100
150
200
250
300
350
400
450

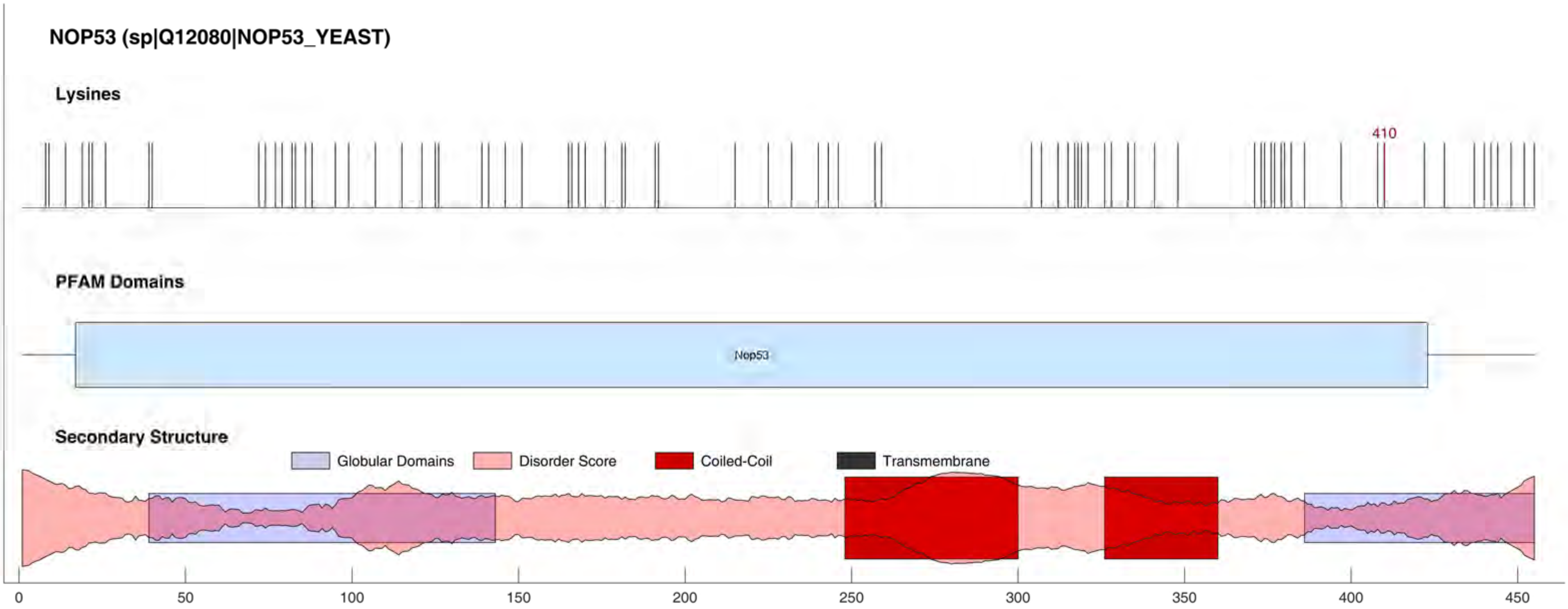
NOP53 (sp|Q12080|NOP53_YEAST)
Lysines
410
PFAM Domains
Nop53
Secondary Structure
Globular Domains
Disorder Score
Coiled-Coil
Transmembrane
0
50
100
150
200
250
300
350
400
450

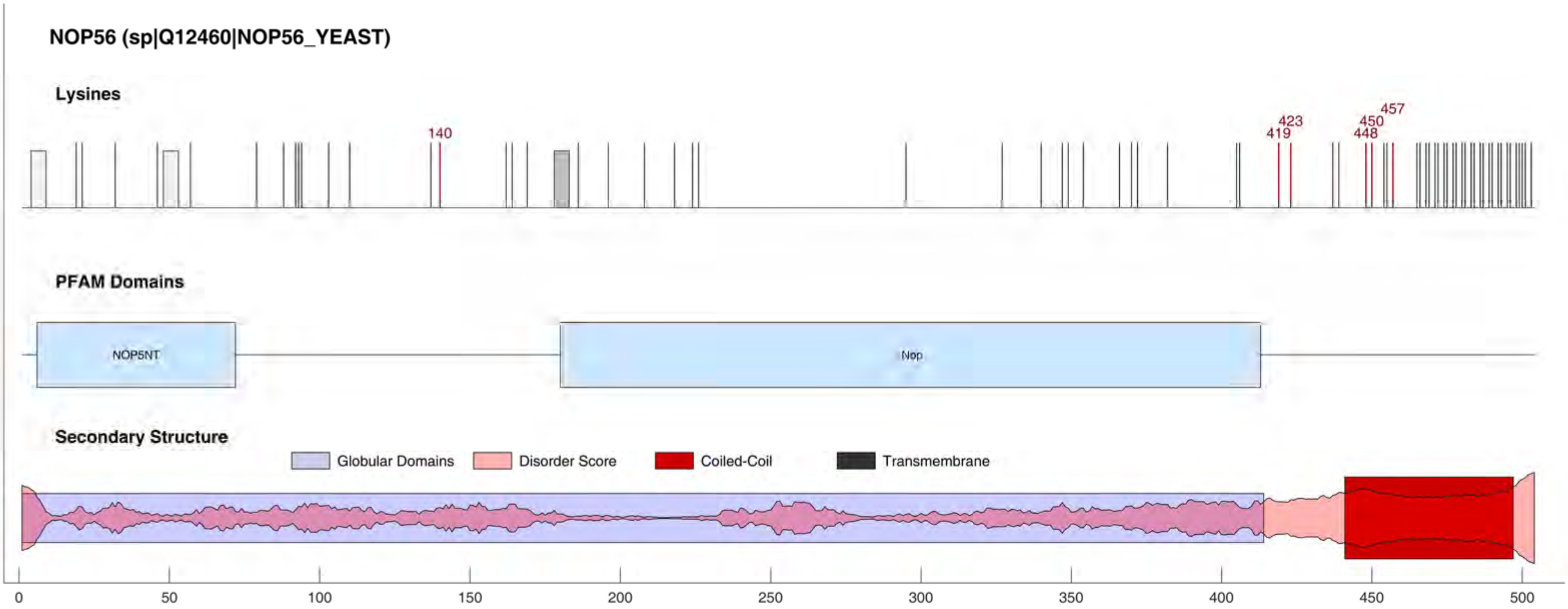
NOP56 (sp|Q12460|NOP56_YEAST)
Lysines
140
419
423
448
450
457
PFAM Domains
NOP5NT
Nop
Secondary Structure
Globular Domains
Disorder Score
Coiled-Coil
Transmembrane
0
50
100
150
200
250
300
350
400
450
500

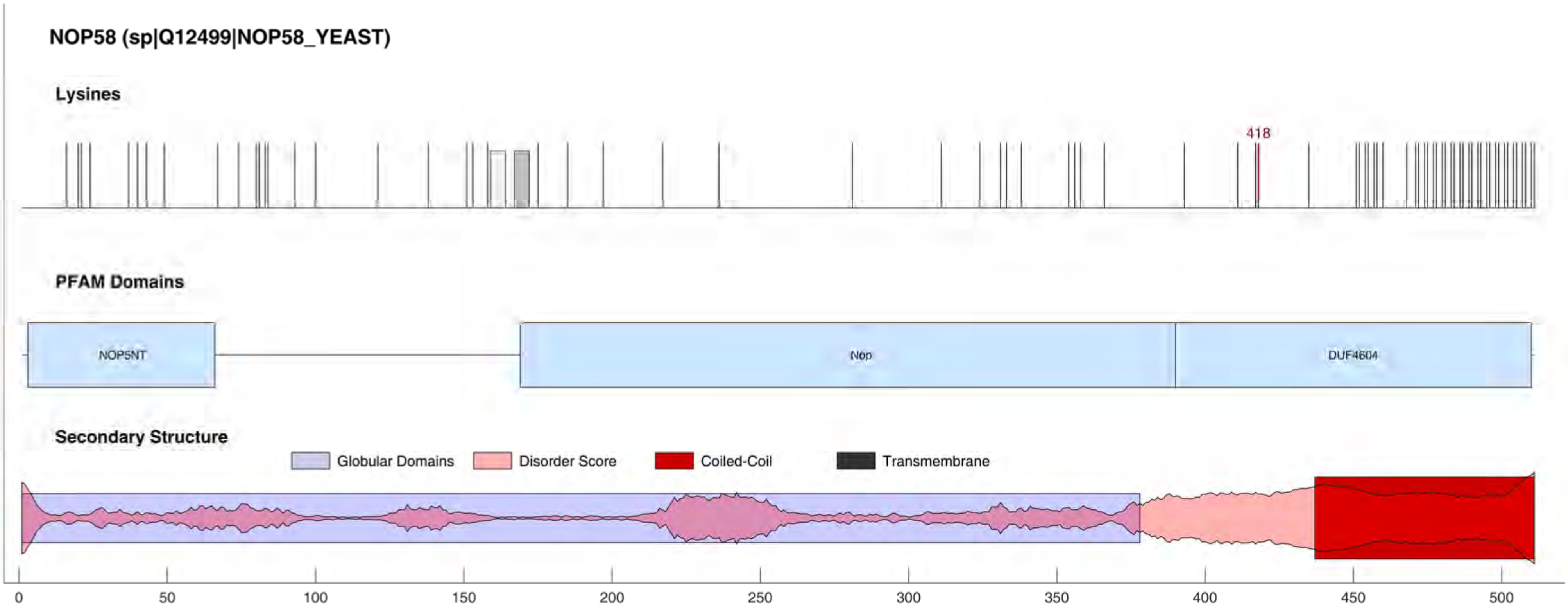
NOP58 (sp|Q12499|NOP58_YEAST)
Lysines
418
PFAM Domains
NOP5NT
Nop
DUF4604
Secondary Structure
Globular Domains
Disorder Score
Coiled-Coil
Transmembrane
0
50
100
150
200
250
300
350
400
450
500

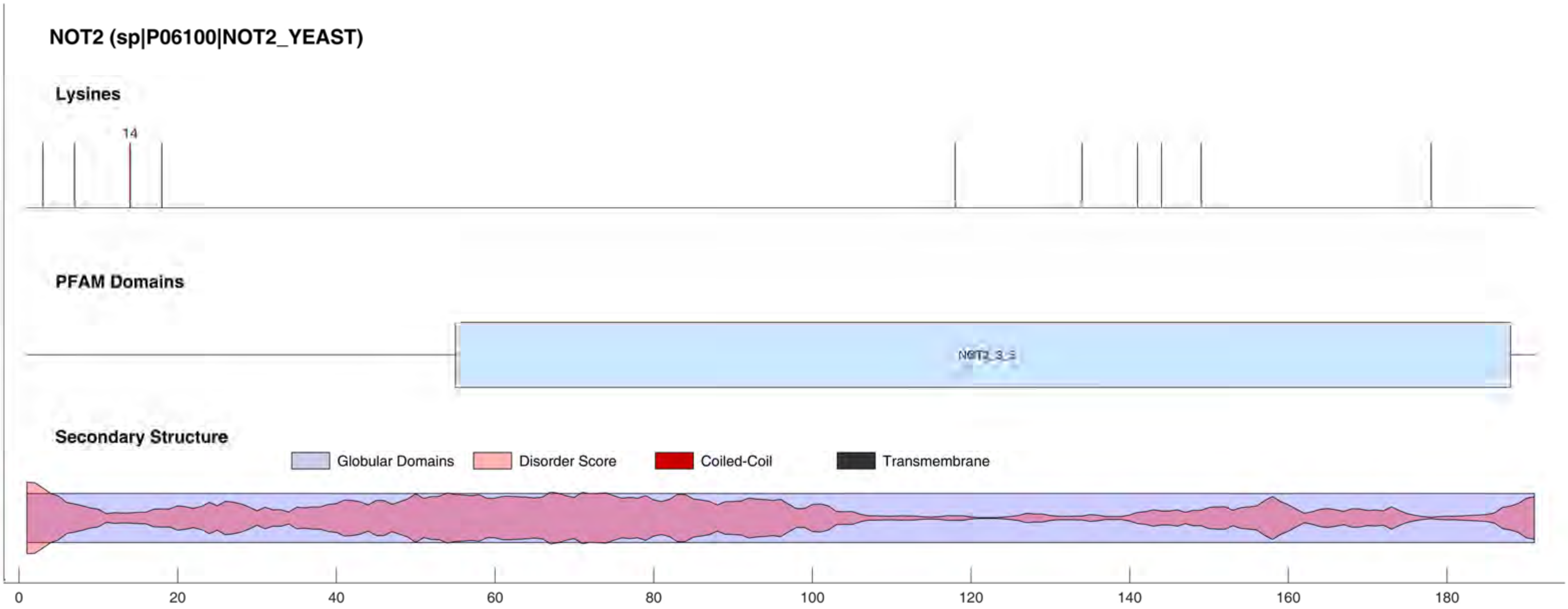
NOT2 (sp|P06100|NOT2_YEAST)
Lysines
14
PFAM Domains
Secondary Structure
Globular Domains
Disorder Score
Coiled-Coil
Transmembrane
0
20
40
60
80
100
120
140
160
180

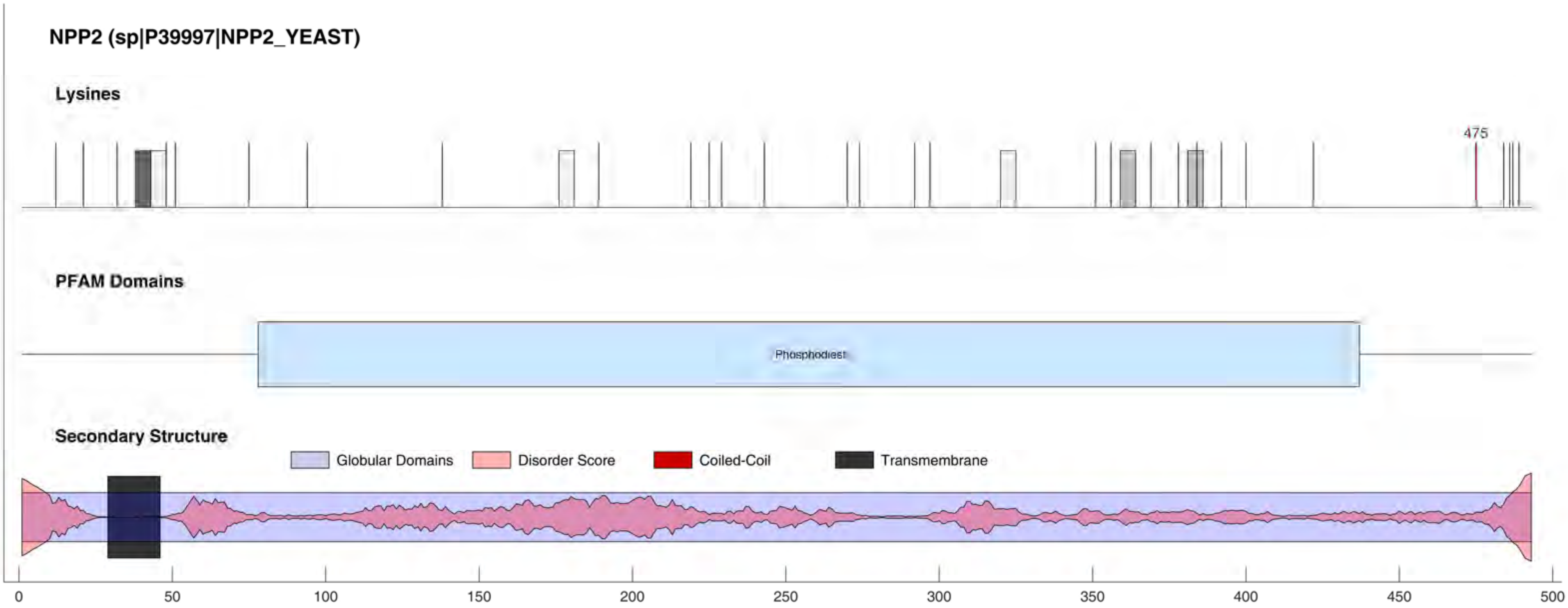
NPP2 (sp|P39997|NPP2_YEAST)
Lysines
475
PFAM Domains
Phosphodiest
Secondary Structure
Globular Domains
Disorder Score
Coiled-Coil
Transmembrane
0
50
100
150
200
250
300
350
400
450
500

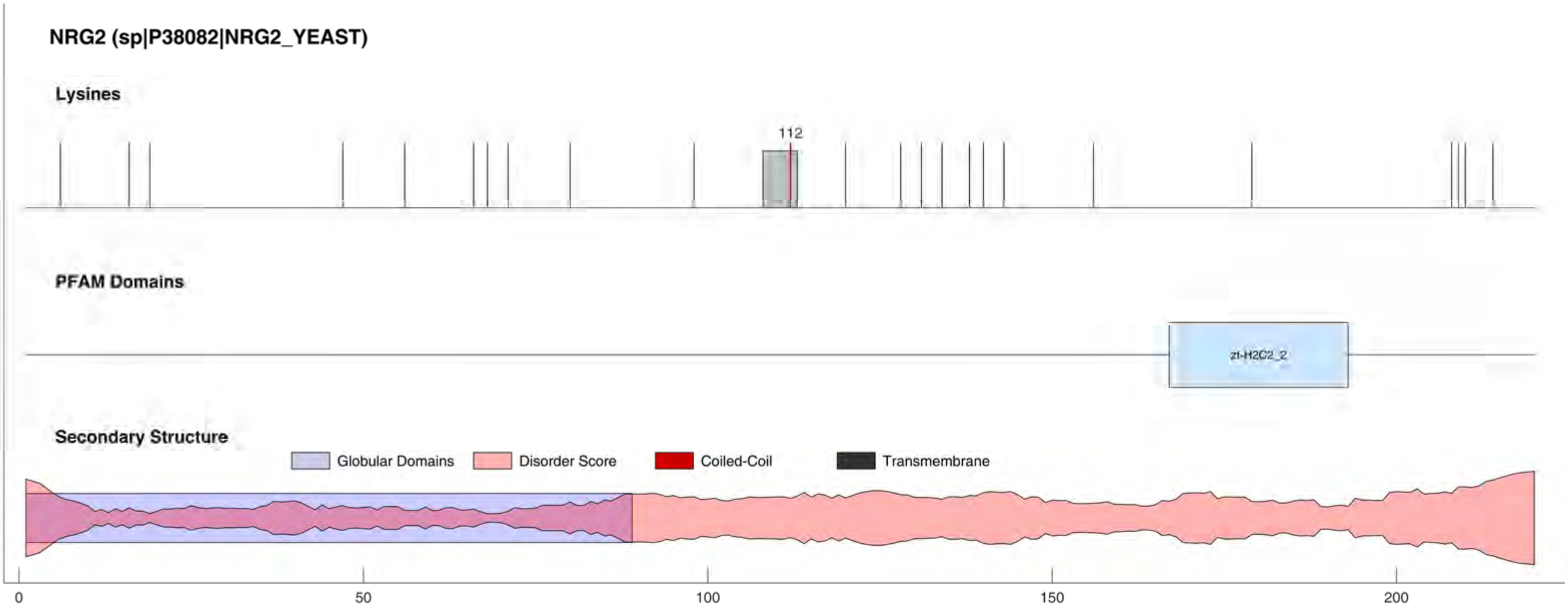
NRG2 (sp|P38082|NRG2_YEAST)
Lysines
112
PFAM Domains
zf-H2C2_2
Secondary Structure
Globular Domains
Disorder Score
Coiled-Coil
Transmembrane
0
50
100
150
200

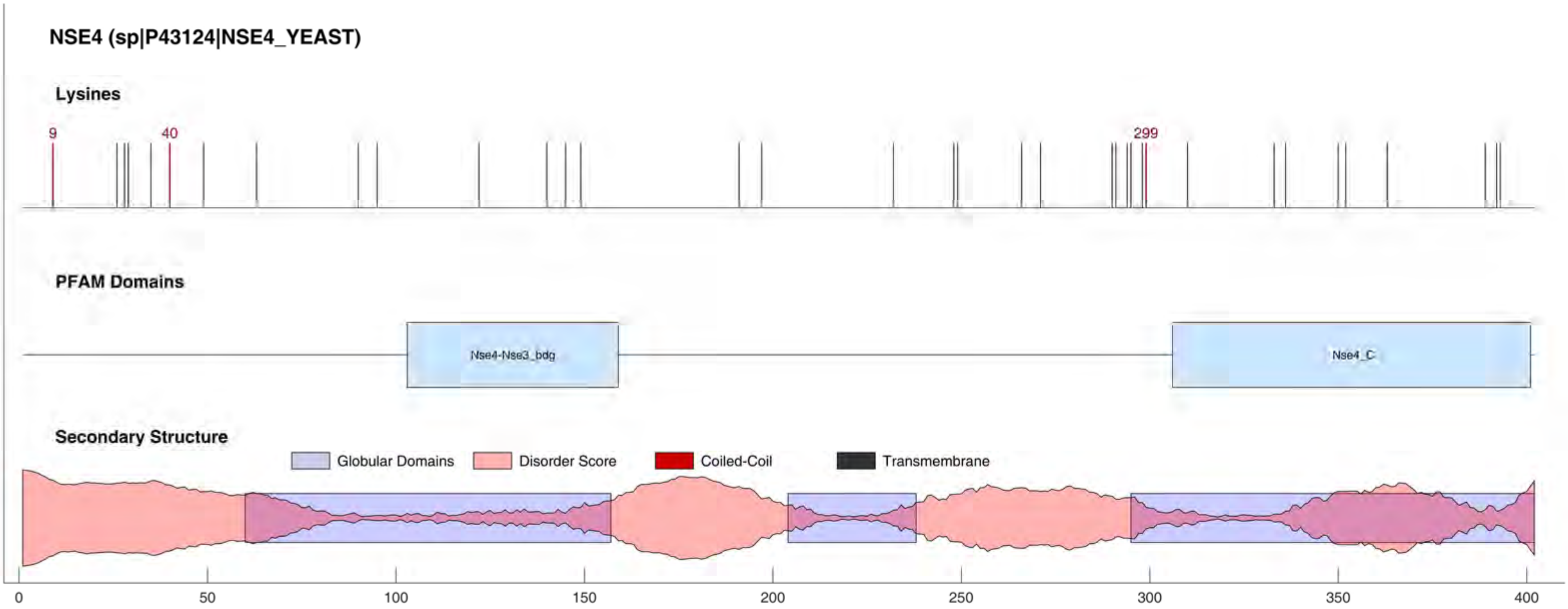
NSE4 (sp|P43124|NSE4_YEAST)
Lysines
9
40
299
PFAM Domains
Nse4-Nse3_bdg
Nse4_C
Secondary Structure
Globular Domains
Disorder Score
Coiled-Coil
Transmembrane
0
50
100
150
200
250
300
350
400

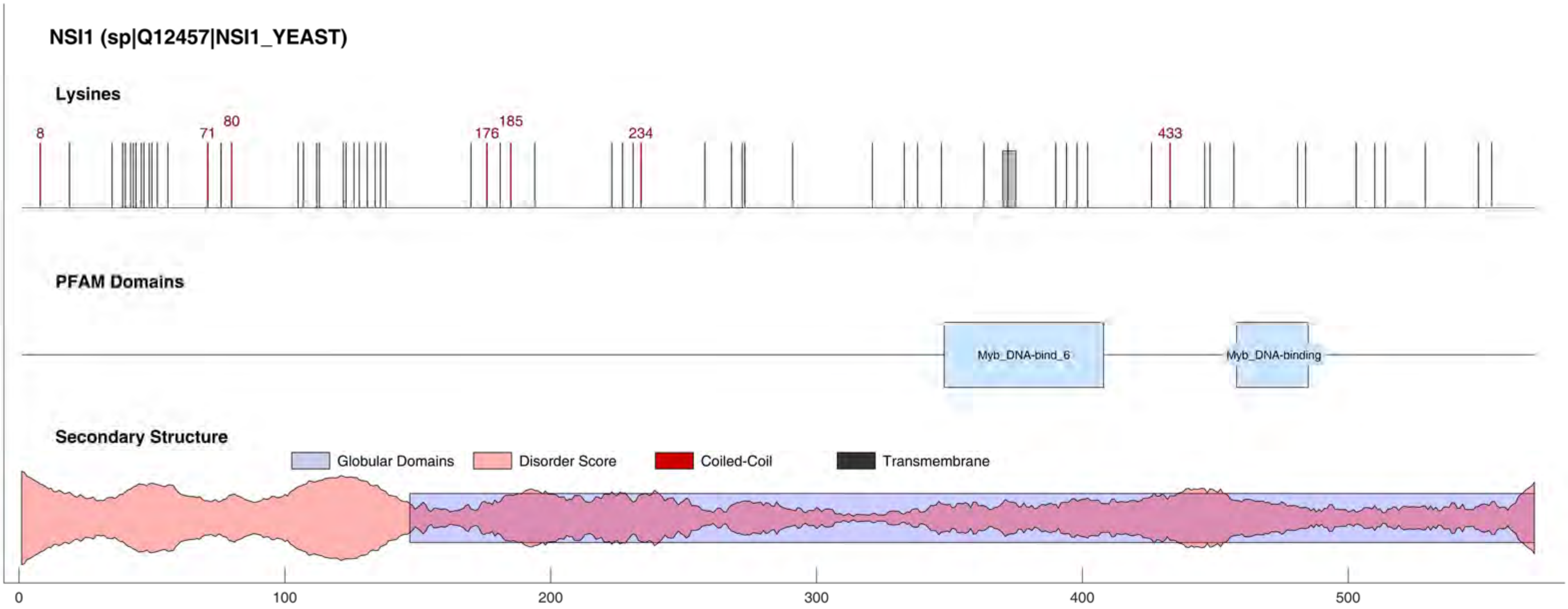
NSI1 (sp|Q12457|NSI1_YEAST)
Lysines
8
71
80
176
185
234
433
PFAM Domains
Myb_DNA-bind_6
Myb_DNA-binding
Secondary Structure
Globular Domains
Disorder Score
Coiled-Coil
Transmembrane
0
100
200
300
400
500

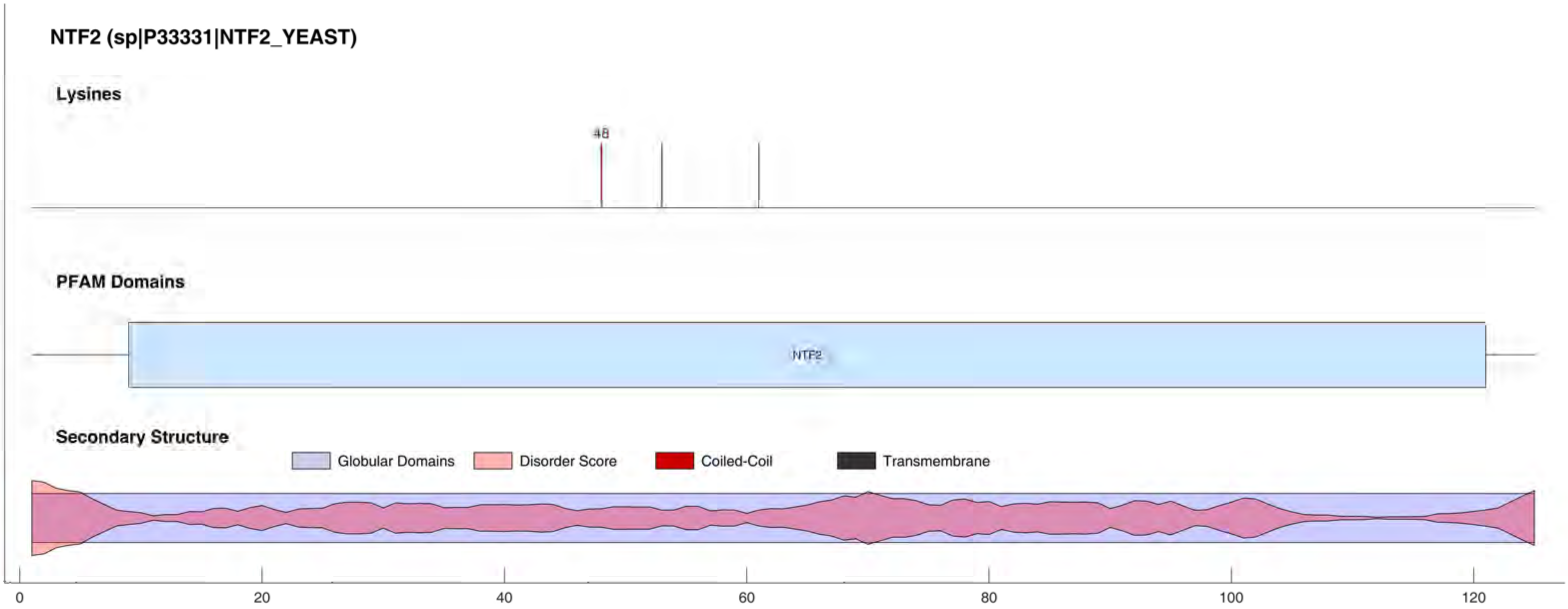
NTF2 (sp|P33331|NTF2_YEAST)
Lysines
48
PFAM Domains
NTF2
Secondary Structure
Globular Domains
Disorder Score
Coiled-Coil
Transmembrane
0
20
40
60
80
100
120

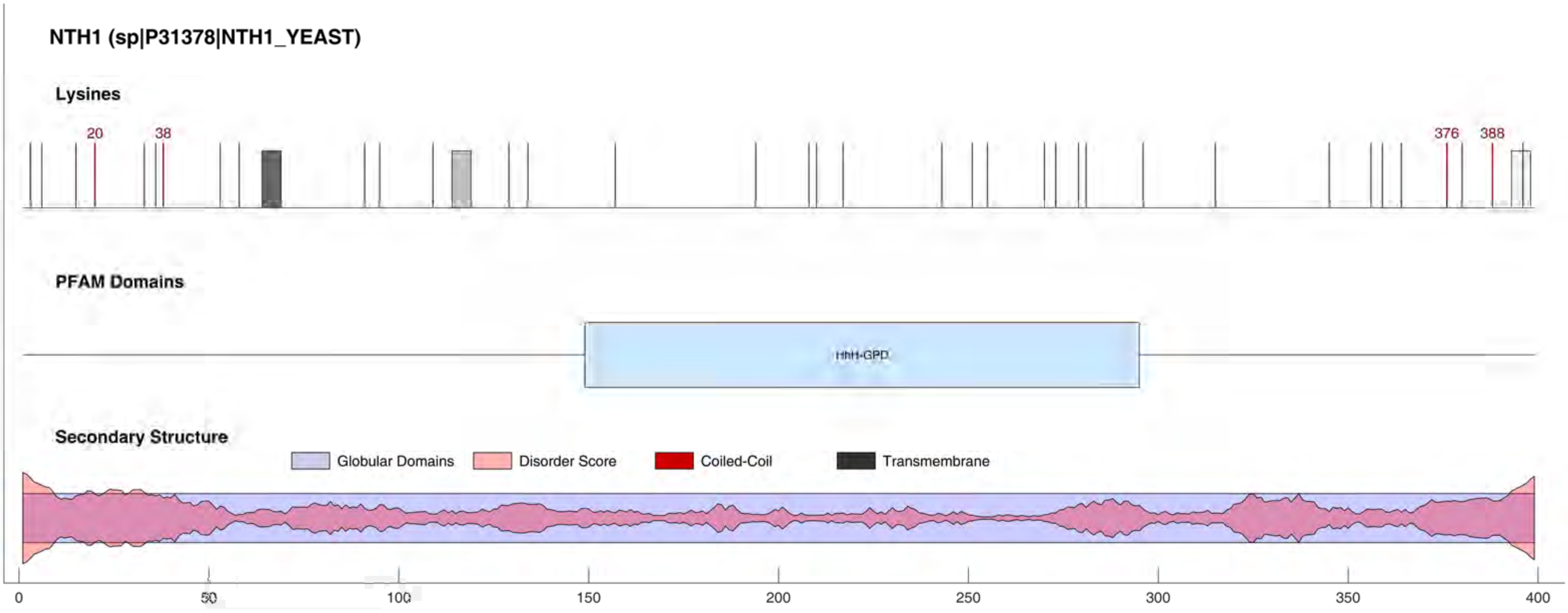
NTH1 (sp|P31378|NTH1_YEAST)
Lysines
20
38
376
388
PFAM Domains
HhH-GPD
Secondary Structure
Globular Domains
Disorder Score
Coiled-Coil
Transmembrane
0
50
100
150
200
250
300
350
400

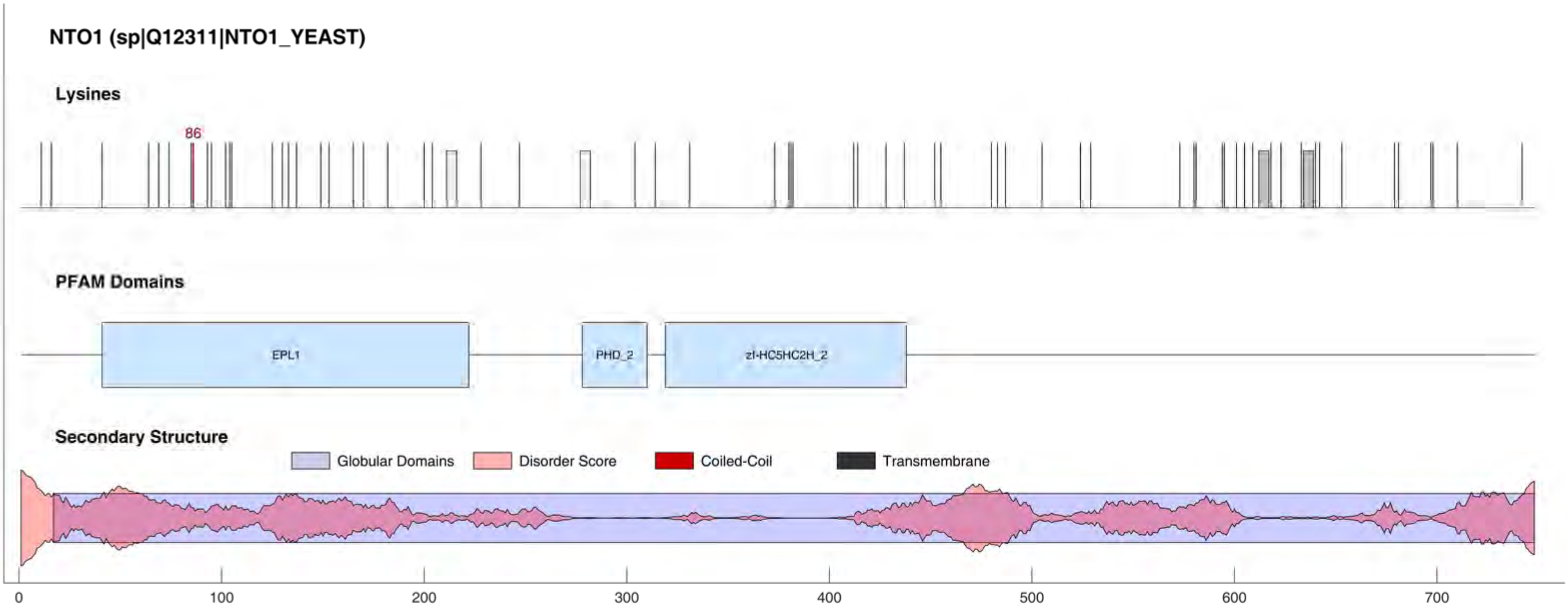

NTO1 (sp|Q12311|NTO1_YEAST)
Lysines
86
PFAM Domains
EPL1
PHD_2
zf-HC5HC2H_2
Secondary Structure
Globular Domains
Disorder Score
Coiled-Coil
Transmembrane
0
100
200
300
400
500
600
700

#### NU159 (sp|P40477|NU159_YEAST)

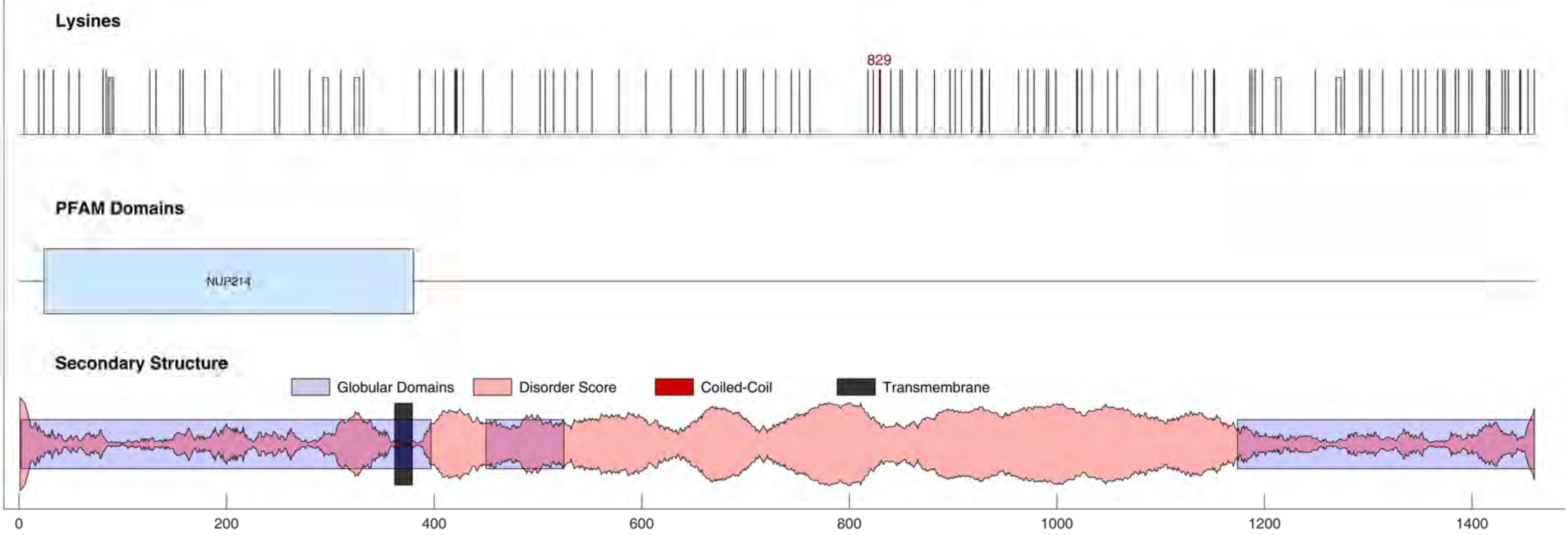

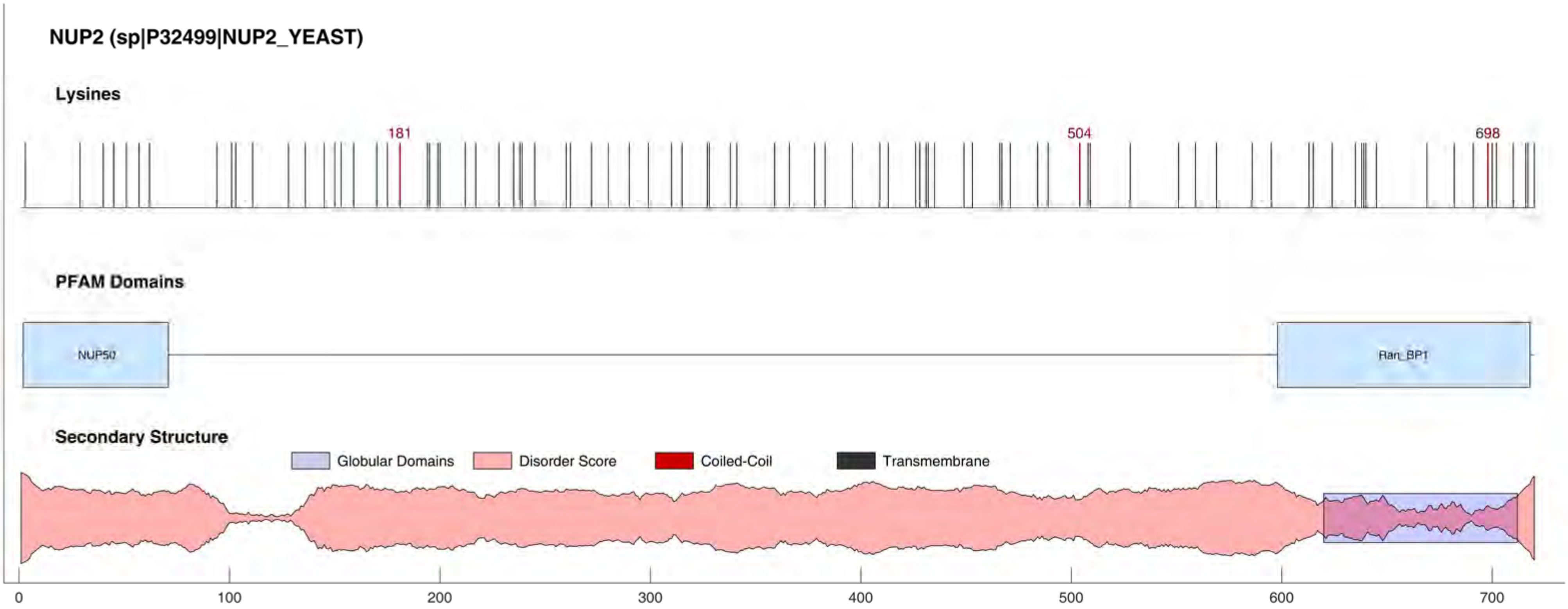
NUP2 (sp|P32499|NUP2_YEAST)
Lysines
181
504
698
PFAM Domains
NUP50
Ran_BP1
Secondary Structure
Globular Domains
Disorder Score
Coiled-Coil
Transmembrane
0
100
200
300
400
500
600
700

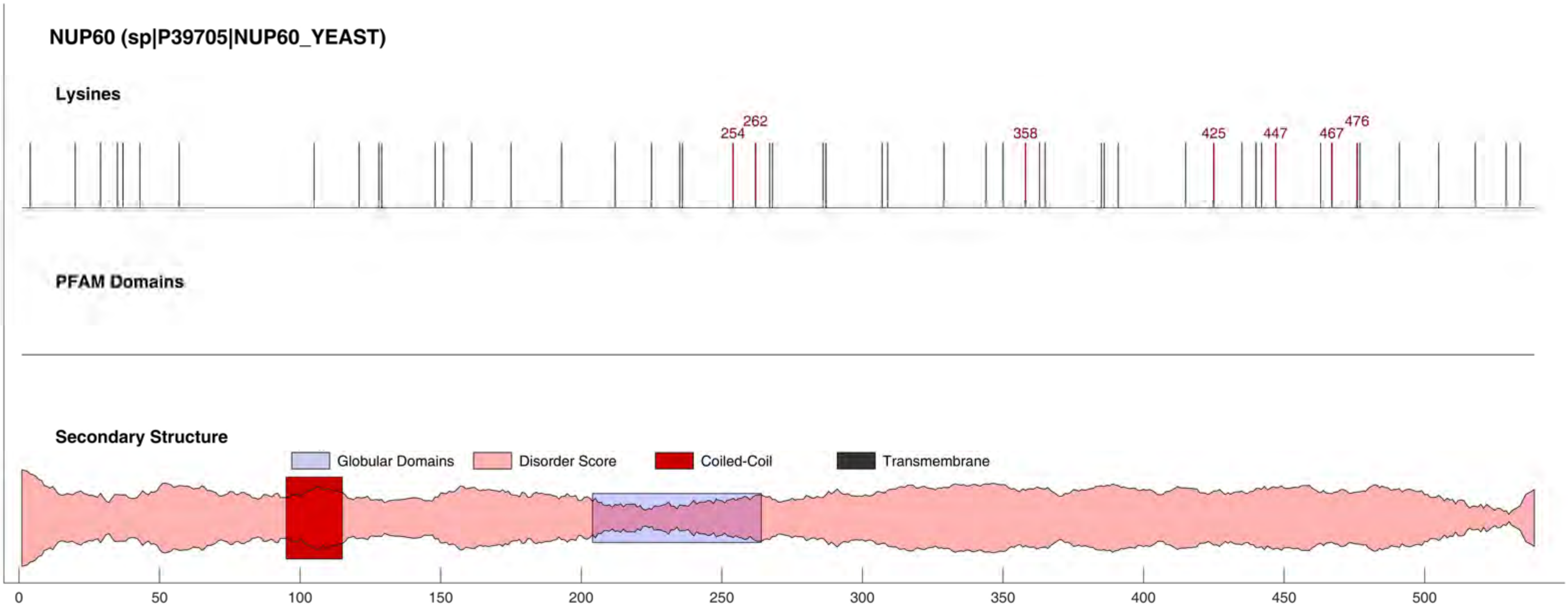

NUP60 (sp|P39705|NUP60_YEAST)
Lysines
254
262
358
425
447
467
476
PFAM Domains
Secondary Structure
Globular Domains
Disorder Score
Coiled-Coil
Transmembrane
0
50
100
150
200
250
300
350
400
450
500

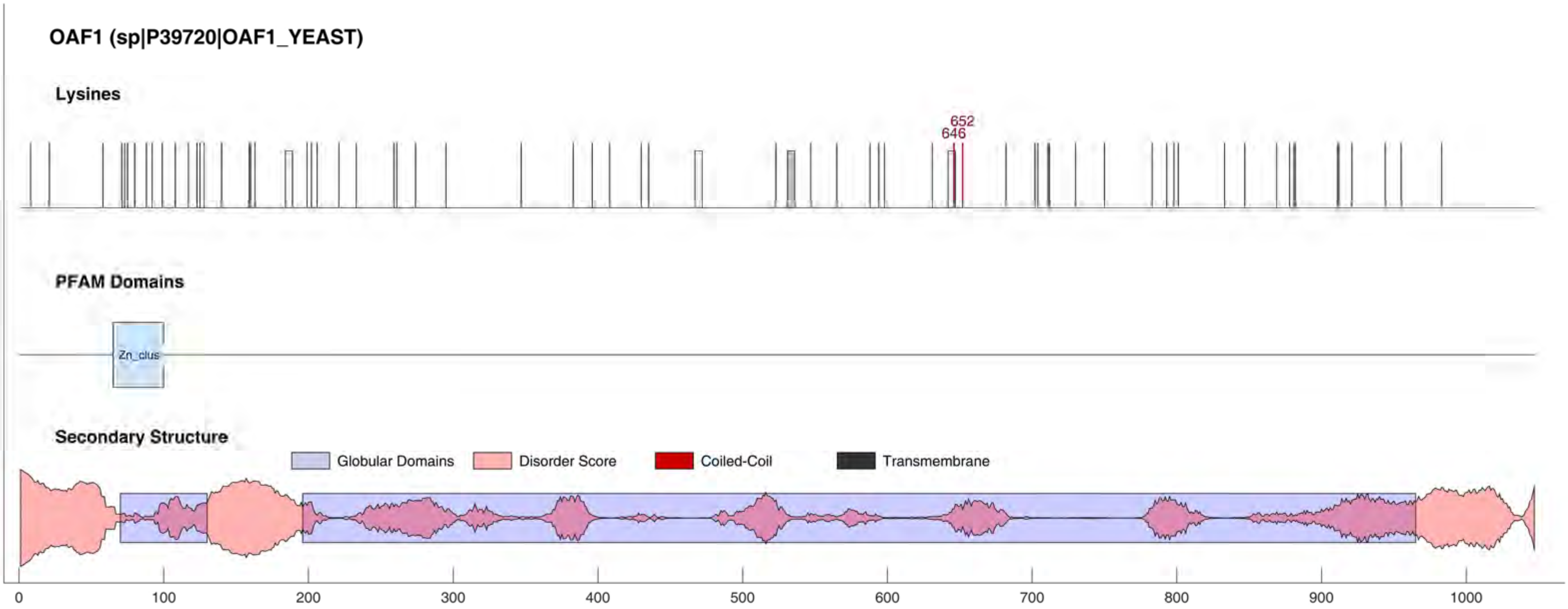

OAF1 (sp|P39720|OAF1_YEAST)
Lysines
652
646
PFAM Domains
Zn_clus
Secondary Structure
Globular Domains
Disorder Score
Coiled-Coil
Transmembrane
0
100
200
300
400
500
600
700
800
900
1000

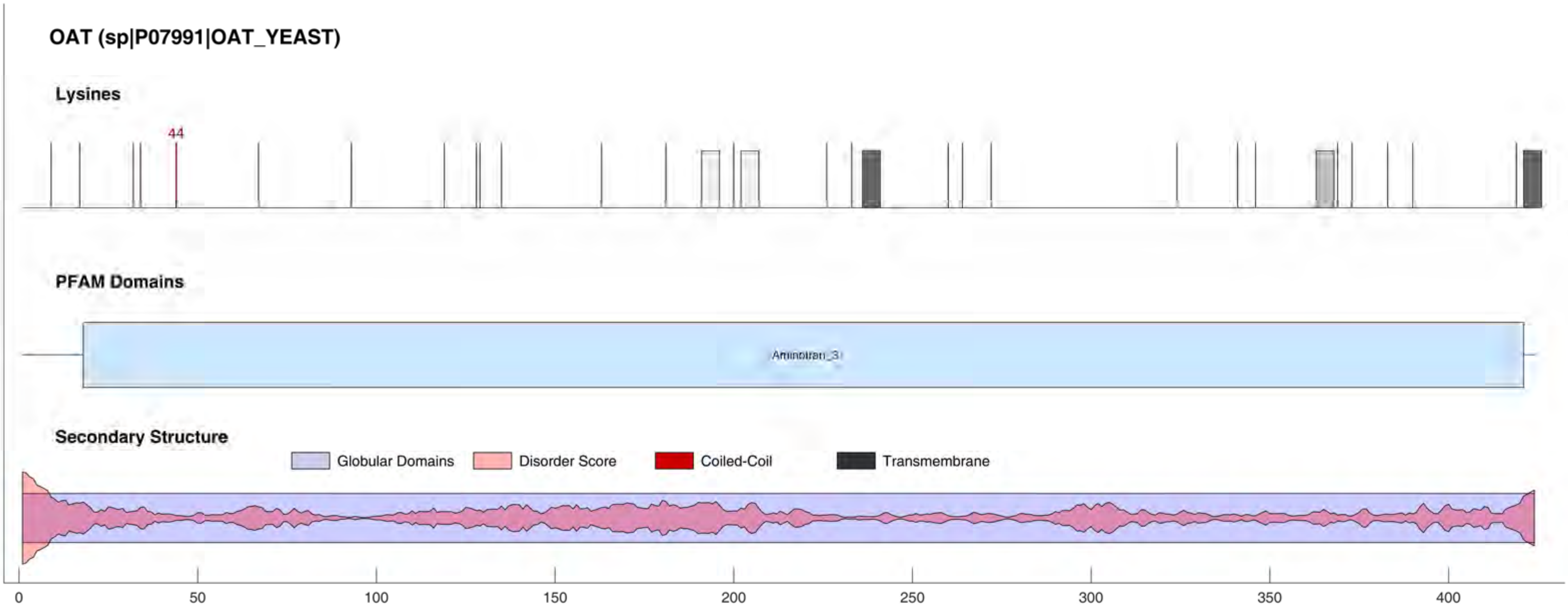
OAT (sp|P07991|OAT_YEAST)
Lysines
44
PFAM Domains
Aminotran_3
Secondary Structure
Globular Domains
Disorder Score
Coiled-Coil
Transmembrane
0
50
100
150
200
250
300
350
400

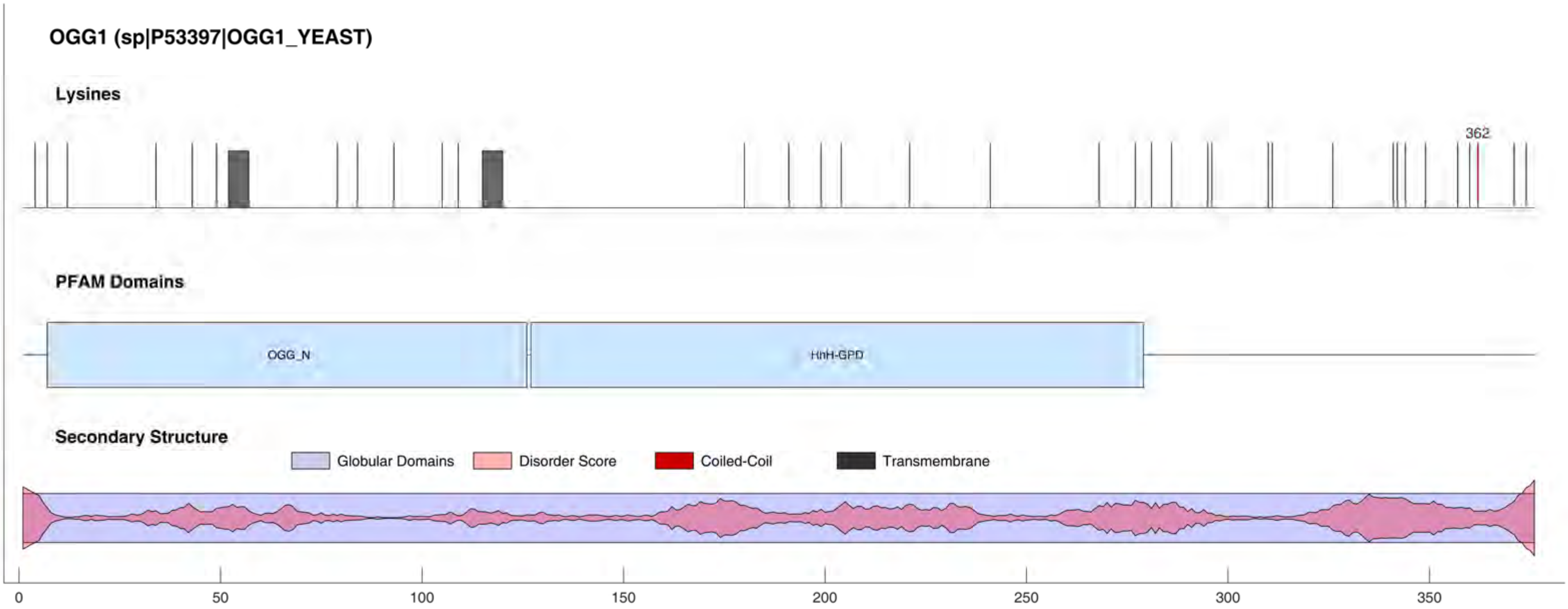

OGG1 (sp|P53397|OGG1_YEAST)
Lysines
362
PFAM Domains
OGG_N
HhH-GPD
Secondary Structure
Globular Domains
Disorder Score
Coiled-Coil
Transmembrane
0
50
100
150
200
250
300
350

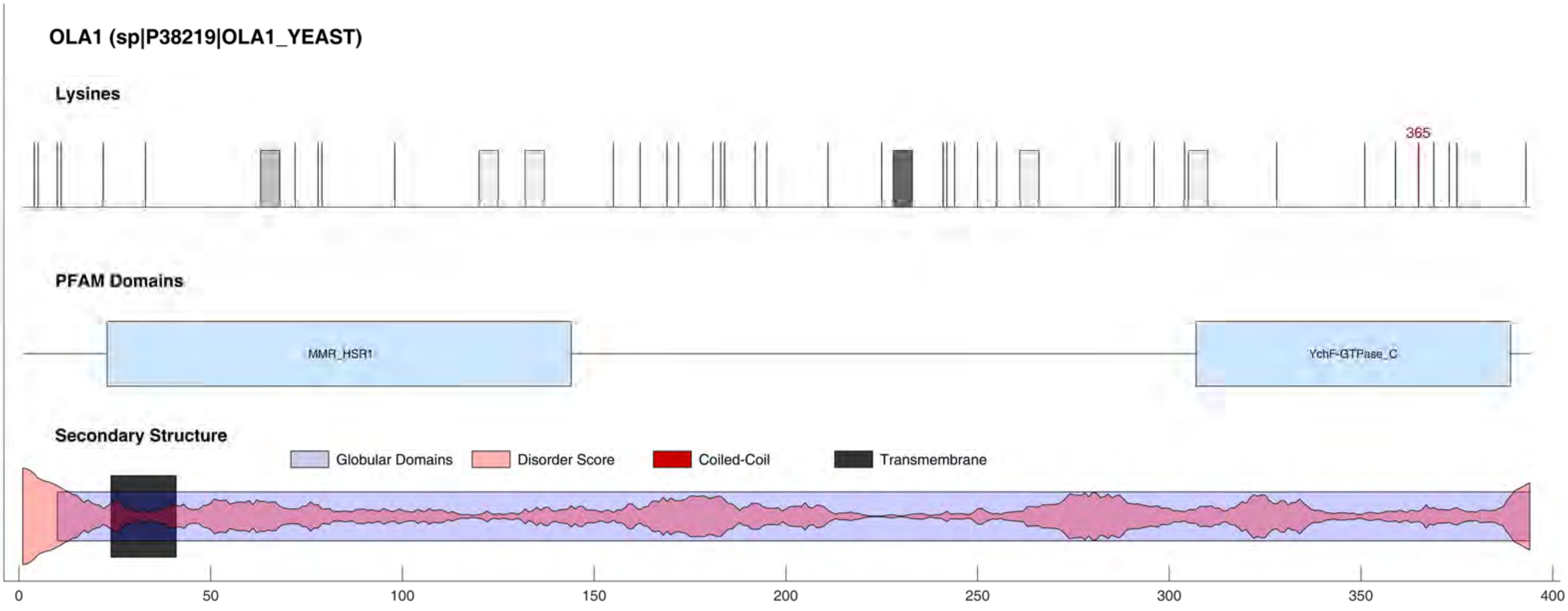

OLA1 (sp|P38219|OLA1_YEAST)
Lysines
365
PFAM Domains
MMR_HSR1
YchF-GTPase_C
Secondary Structure
Globular Domains
Disorder Score
Coiled-Coil
Transmembrane
0
50
100
150
200
250
300
350
400

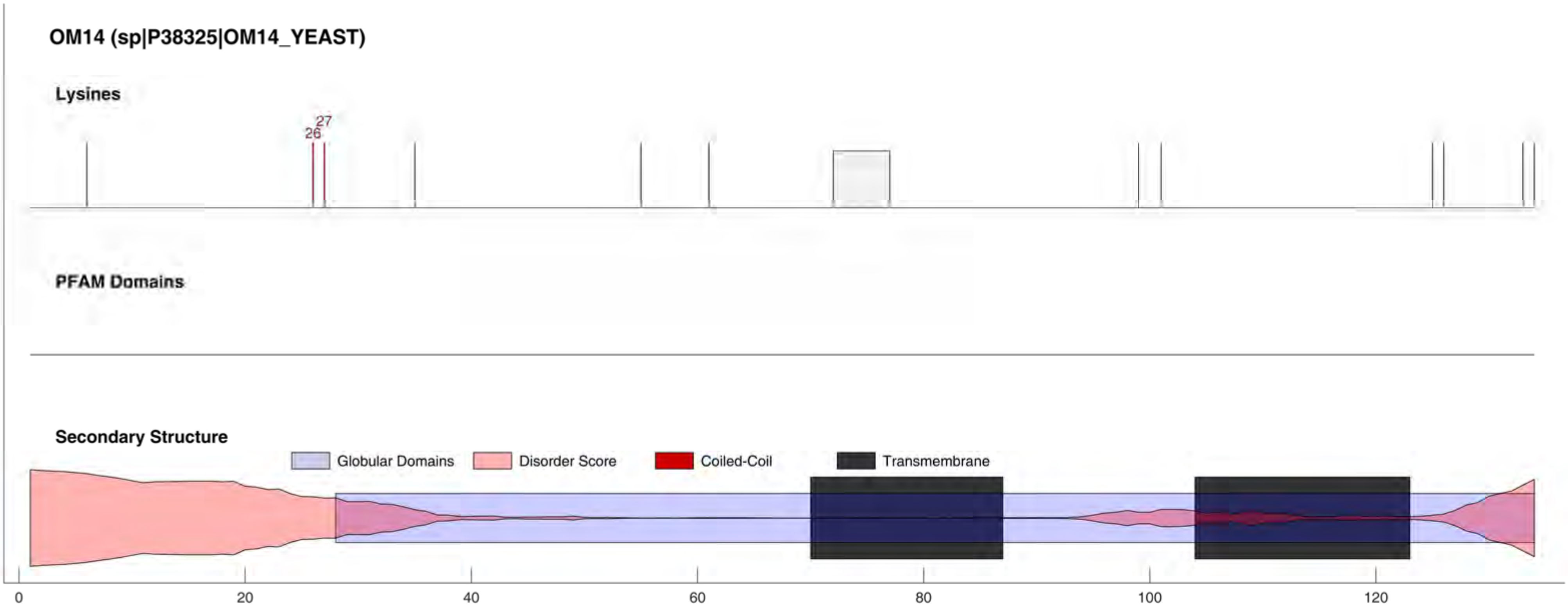

OM14 (sp|P38325|OM14_YEAST)
Lysines
26
27
PFAM Domains
Secondary Structure
Globular Domains
Disorder Score
Coiled-Coil
Transmembrane
0
20
40
60
80
100
120

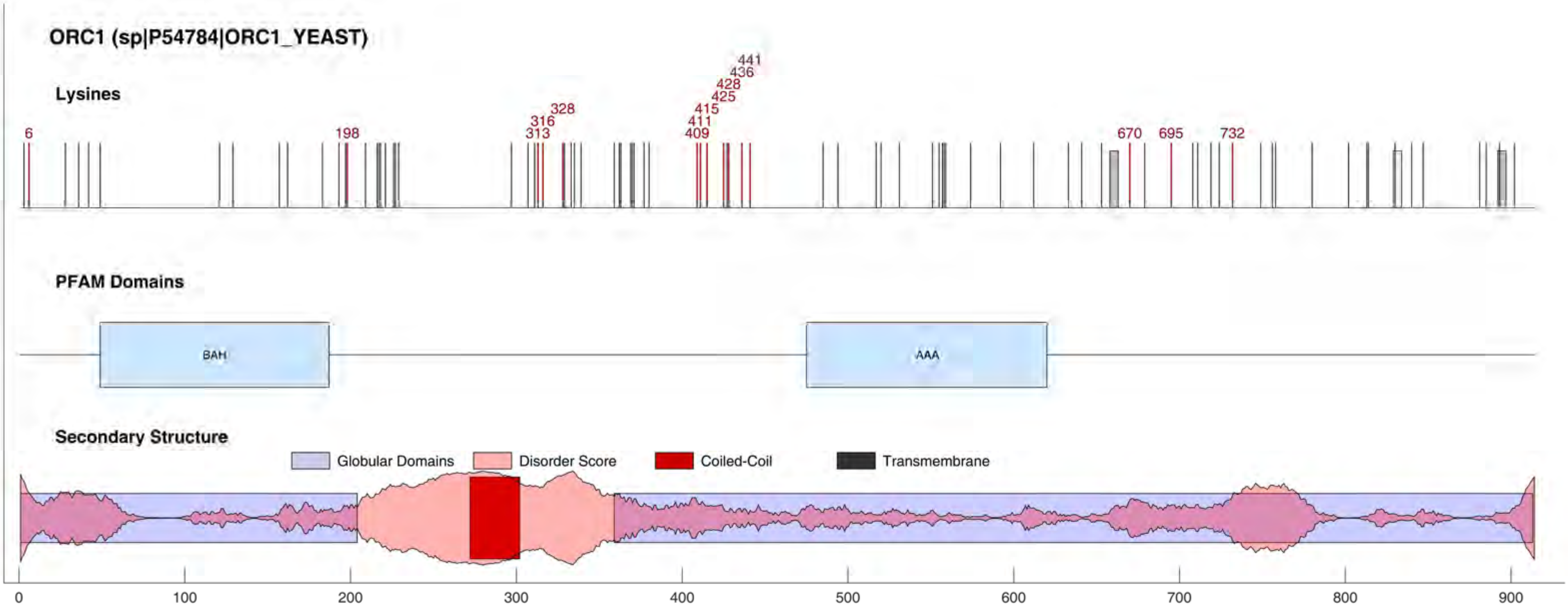
ORC1 (sp|P54784|ORC1_YEAST)
Lysines
6
198
313
316
328
409
411
415
425
428
436
441
670
695
732
PFAM Domains
BAH
AAA
Secondary Structure
Globular Domains
Disorder Score
Coiled-Coil
Transmembrane
0
100
200
300
400
500
600
700
800
900

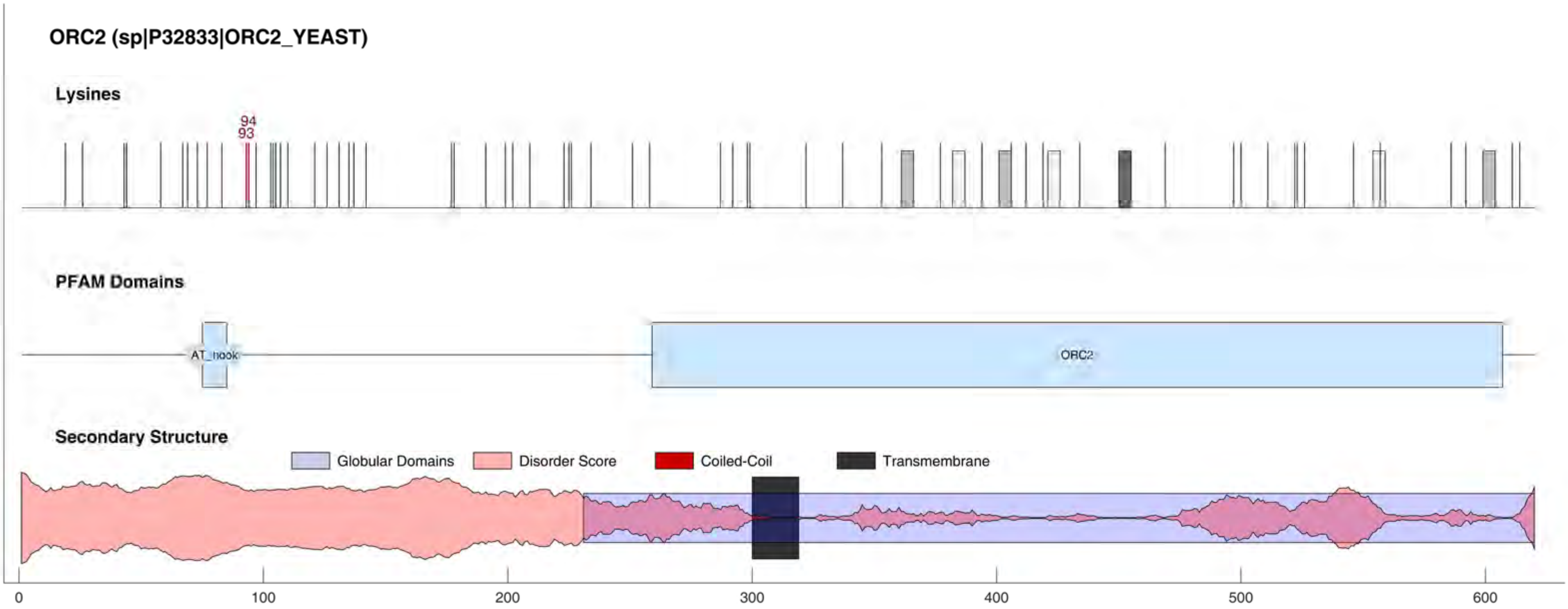

ORC2 (sp|P32833|ORC2_YEAST)
Lysines
94
93
PFAM Domains
AT_hook
ORC2
Secondary Structure
Globular Domains
Disorder Score
Coiled-Coil
Transmembrane
0
100
200
300
400
500
600

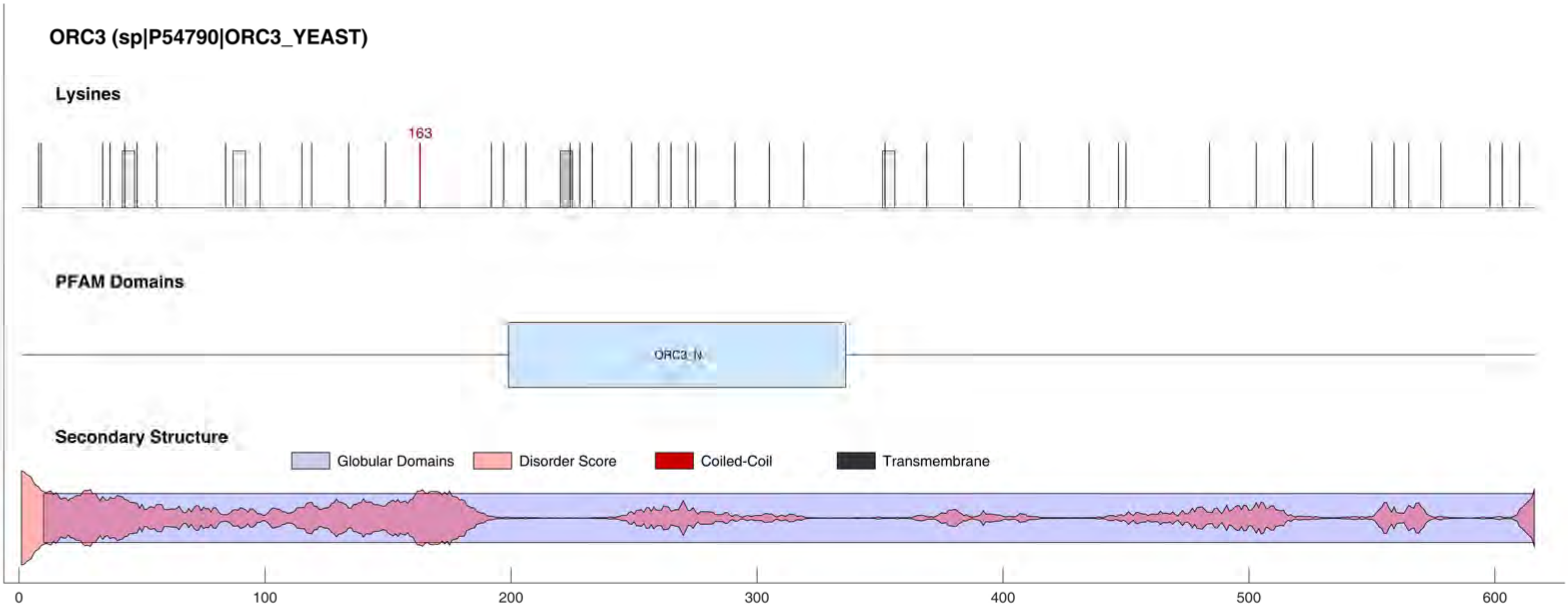
ORC3 (sp|P54790|ORC3_YEAST)
Lysines
163
PFAM Domains
ORC3_N
Secondary Structure
Globular Domains
Disorder Score
Coiled-Coil
Transmembrane
0
100
200
300
400
500
600

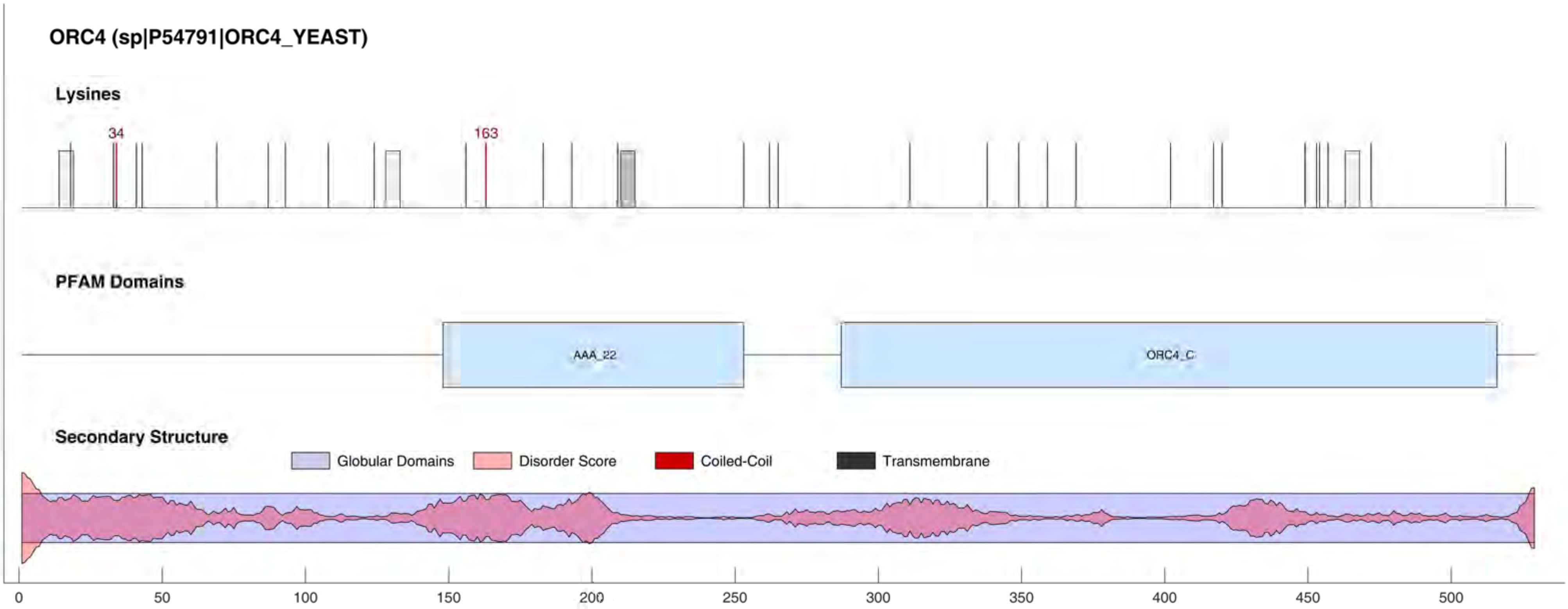
ORC4 (sp|P54791|ORC4_YEAST)
Lysines
34
163
PFAM Domains
AAA_22
ORC4_C
Secondary Structure
Globular Domains
Disorder Score
Coiled-Coil
Transmembrane
0
50
100
150
200
250
300
350
400
450
500

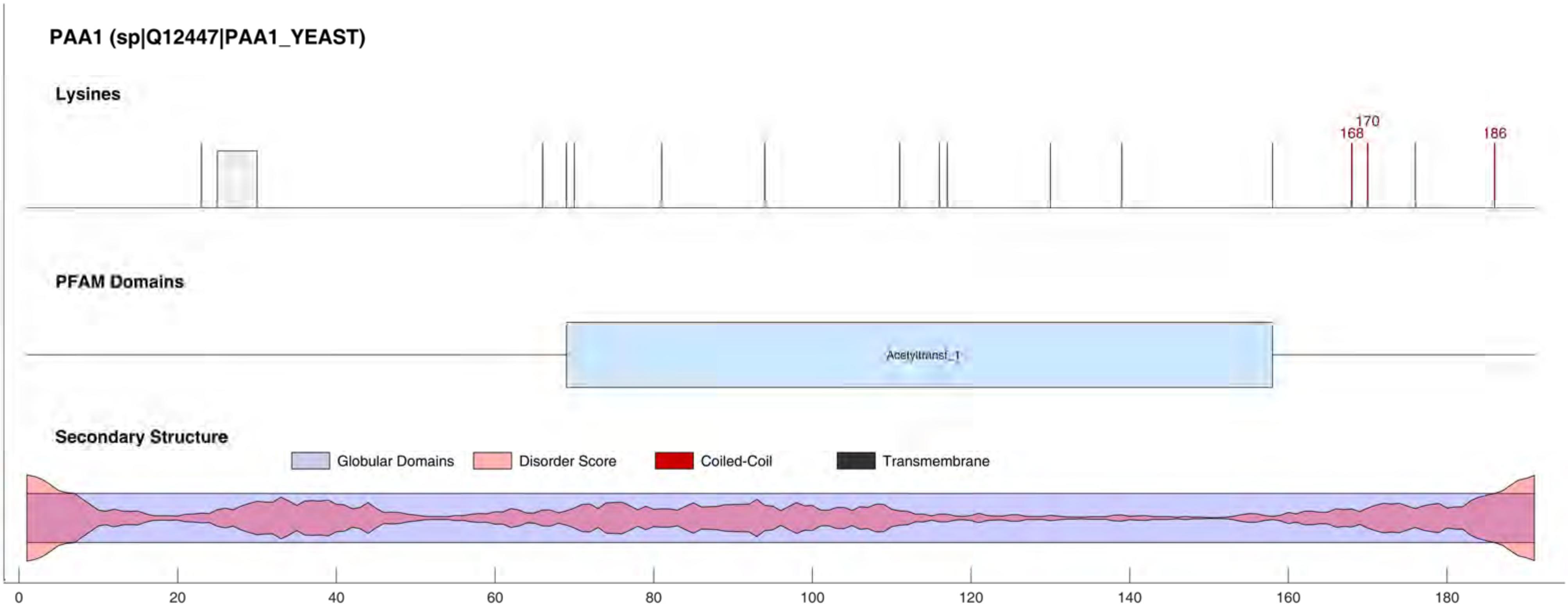
PAA1 (sp|Q12447|PAA1_YEAST)
Lysines
170
168
186
PFAM Domains
Acetyltransf_1
Secondary Structure
Globular Domains
Disorder Score
Coiled-Coil
Transmembrane
0
20
40
60
80
100
120
140
160
180

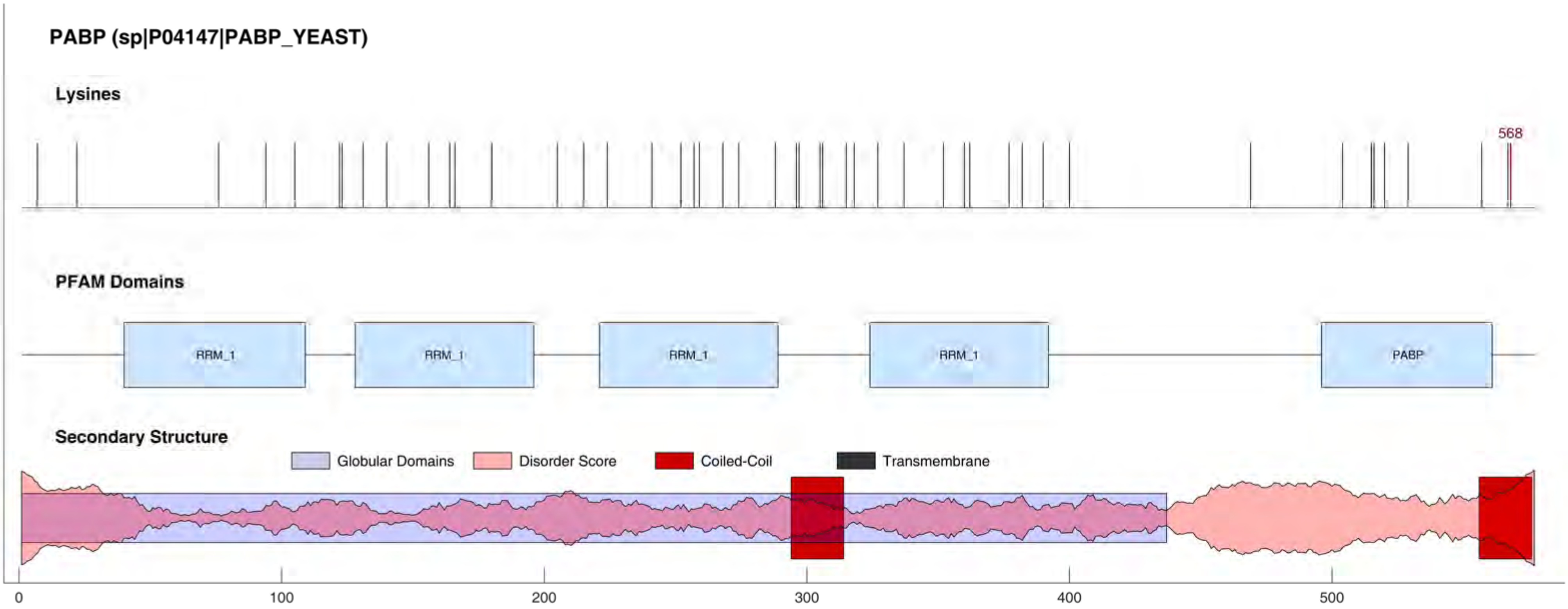
PABP (sp|P04147|PABP_YEAST)
Lysines
568
PFAM Domains
RRM_1
RRM_1
RRM_1
RRM_1
PABP
Secondary Structure
Globular Domains
Disorder Score
Coiled-Coil
Transmembrane
0
100
200
300
400
500

PAF1 (sp|P38351|PAF1_YEAST)

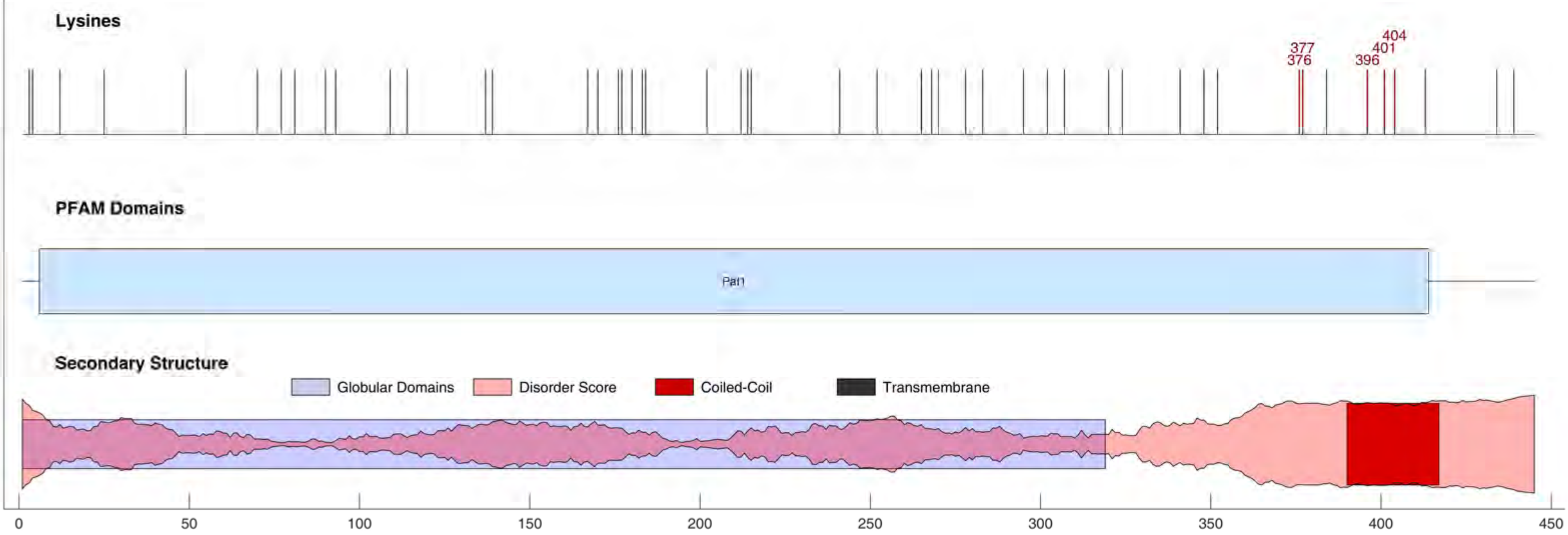

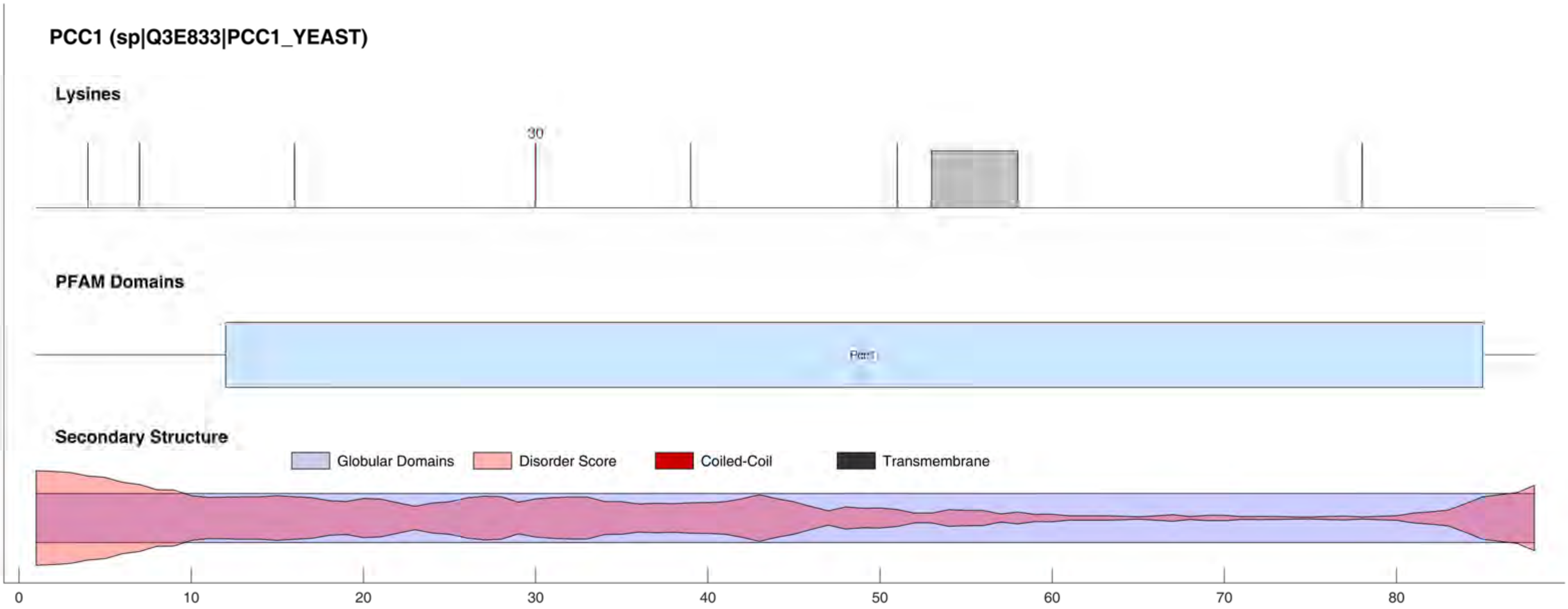
PCC1 (sp|Q3E833|PCC1_YEAST)
Lysines
30
PFAM Domains
Pcc1
Secondary Structure
Globular Domains
Disorder Score
Coiled-Coil
Transmembrane
0
10
20
30
40
50
60
70
80

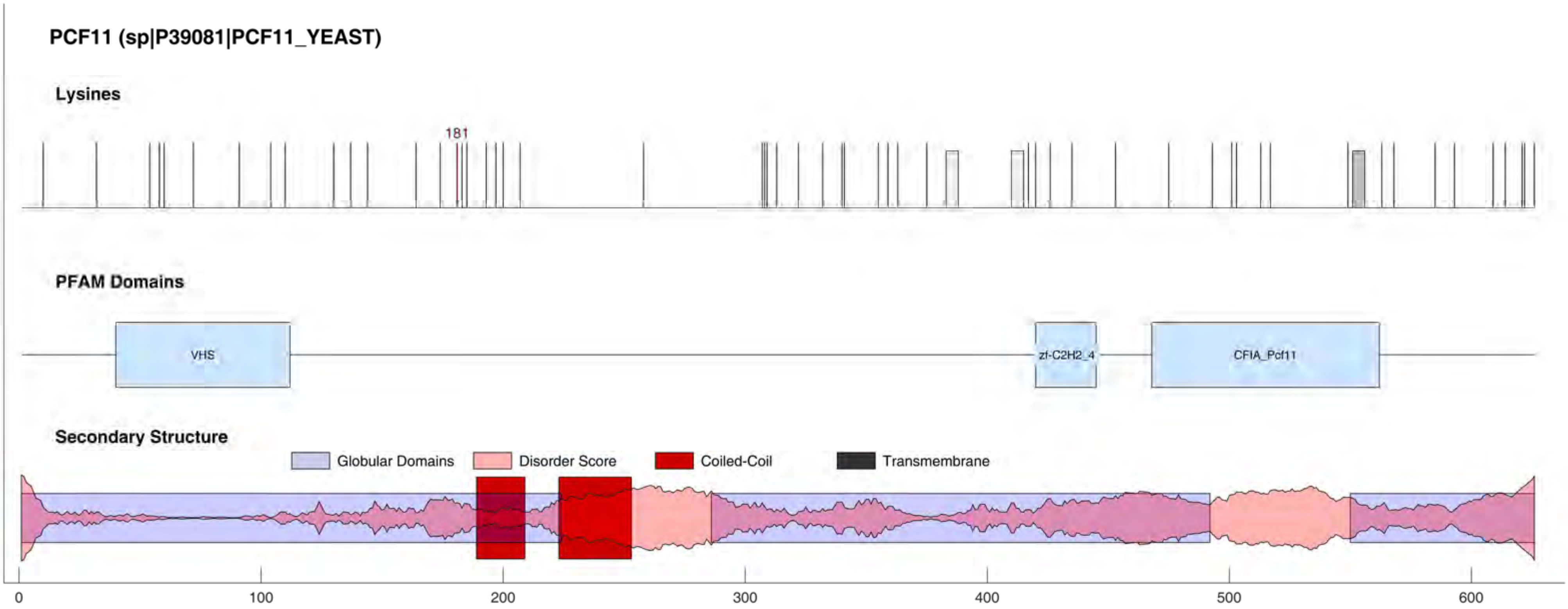

PCF11 (sp|P39081|PCF11_YEAST)
Lysines
181
PFAM Domains
VHS
zf-C2H2_4
CFIA_Pcf11
Secondary Structure
Globular Domains
Disorder Score
Coiled-Coil
Transmembrane
0
100
200
300
400
500
600

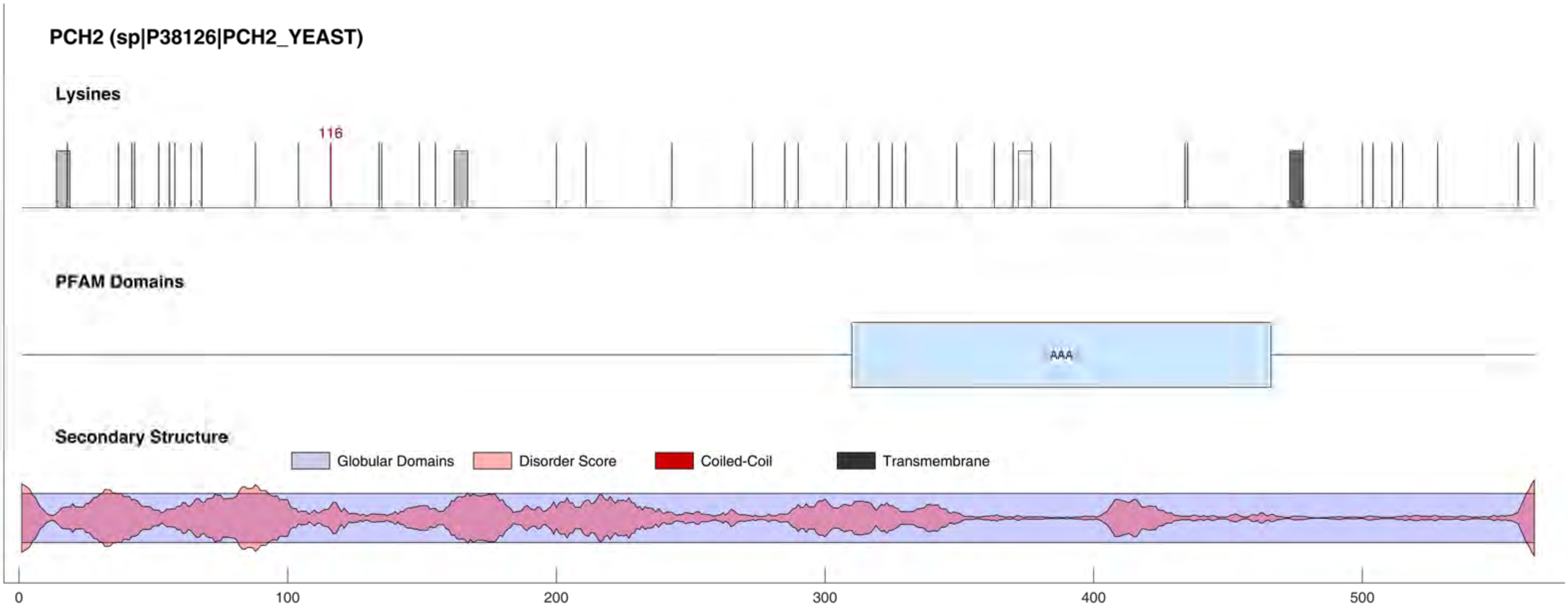
PCH2 (sp|P38126|PCH2_YEAST)
Lysines
116
PFAM Domains
AAA
Secondary Structure
Globular Domains
Disorder Score
Coiled-Coil
Transmembrane
0
100
200
300
400
500

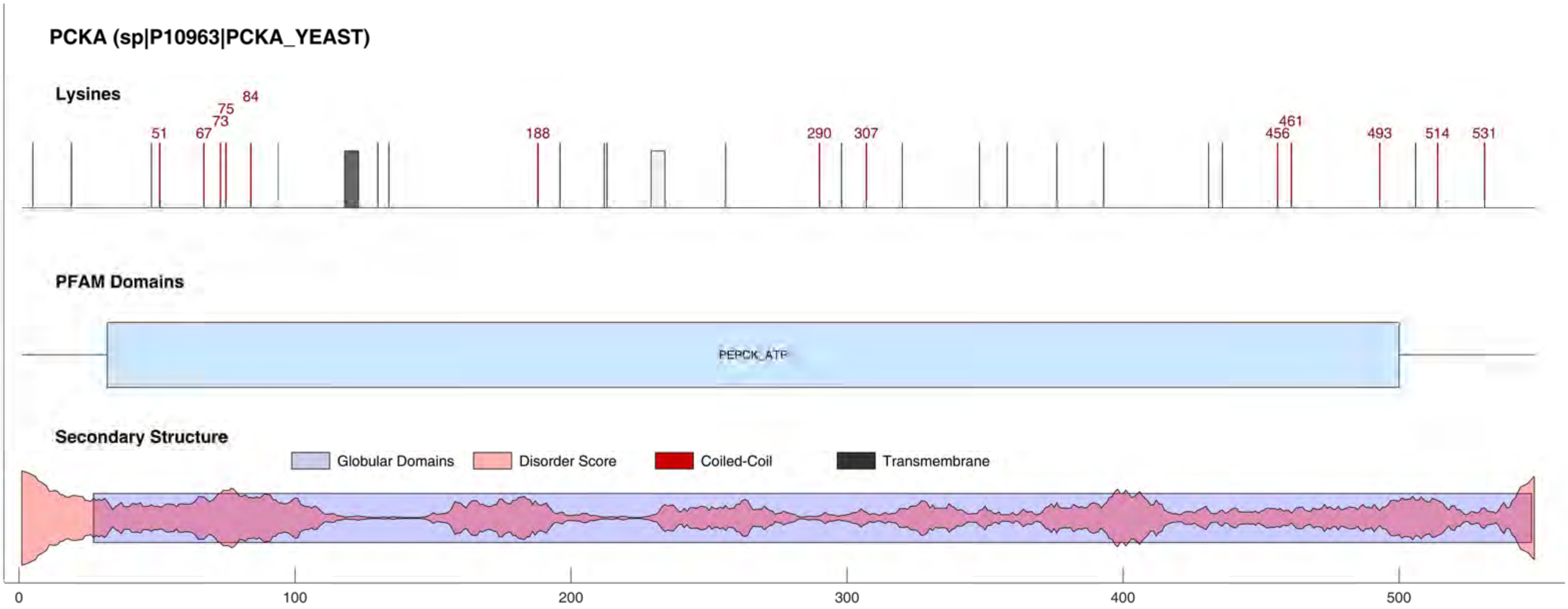
PCKA (sp|P10963|PCKA_YEAST)
Lysines
51
67
73
75
84
188
290
307
456
461
493
514
531
PFAM Domains
PEPCK_ATP
Secondary Structure
Globular Domains
Disorder Score
Coiled-Coil
Transmembrane
0
100
200
300
400
500

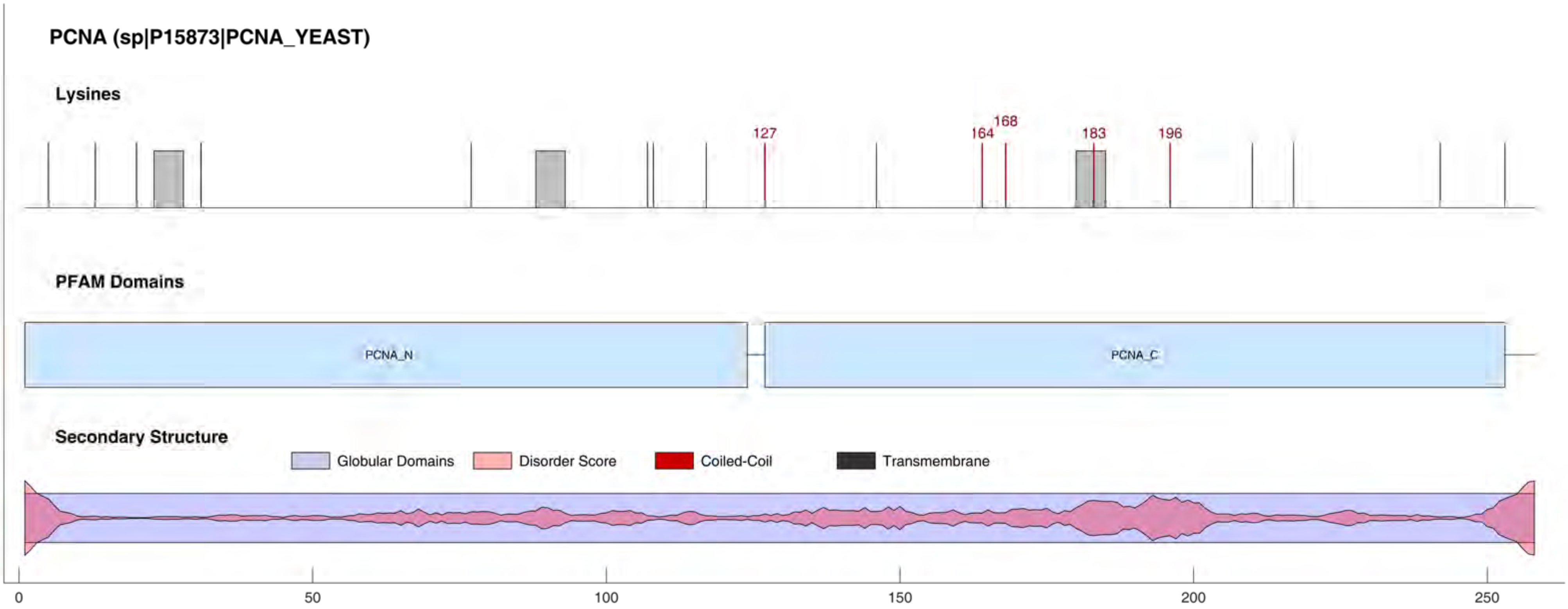
PCNA (sp|P15873|PCNA_YEAST)
Lysines
127
164
168
183
196
PFAM Domains
PCNA_N
PCNA_C
Secondary Structure
Globular Domains
Disorder Score
Coiled-Coil
Transmembrane
0
50
100
150
200
250

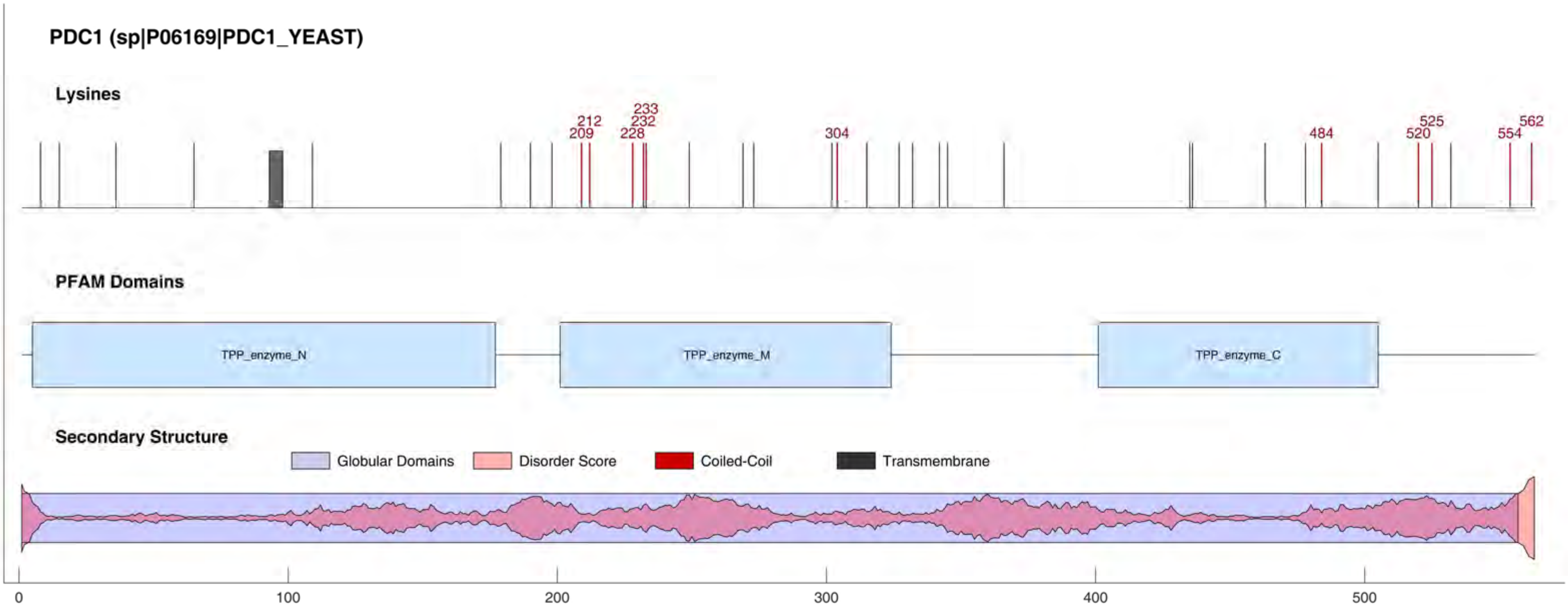

PDC1 (sp|P06169|PDC1_YEAST)
Lysines
209
212
228
232
233
304
484
520
525
554
562
PFAM Domains
TPP_enzyme_N
TPP_enzyme_M
TPP_enzyme_C
Secondary Structure
Globular Domains
Disorder Score
Coiled-Coil
Transmembrane
0
100
200
300
400
500

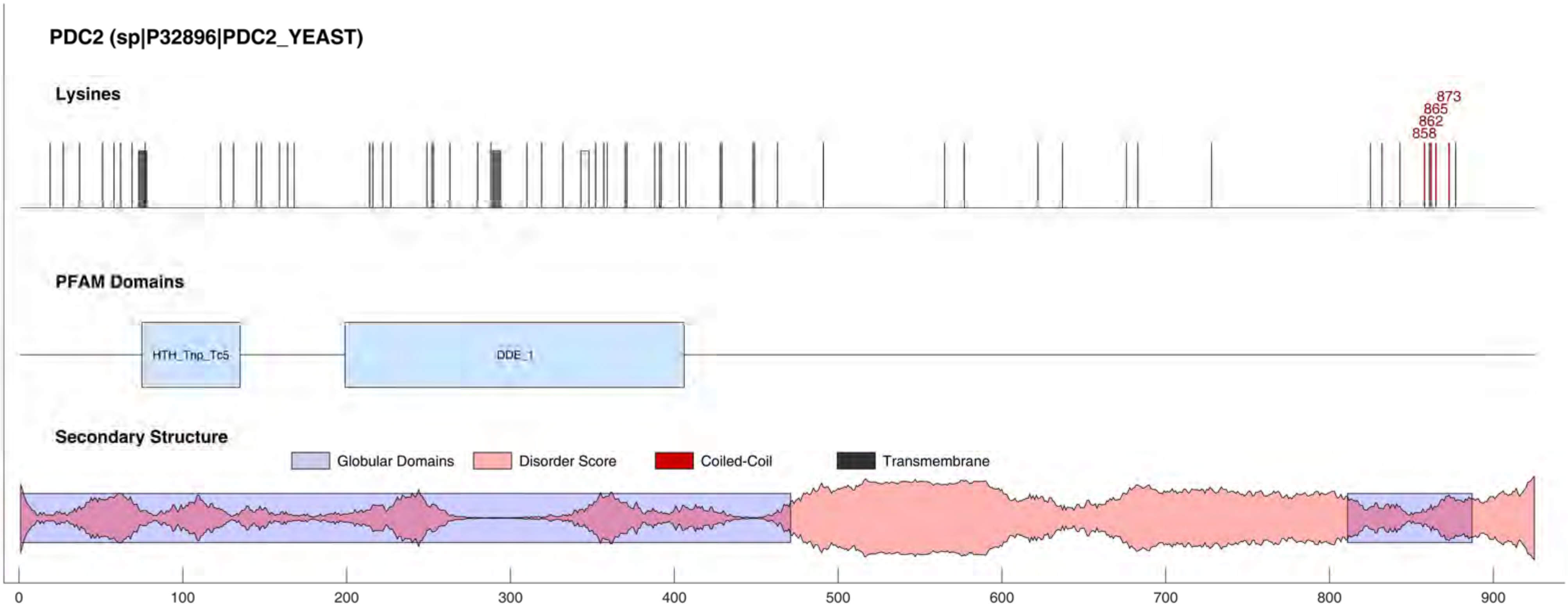
PDC2 (sp|P32896|PDC2_YEAST)
Lysines
873
865
862
858
PFAM Domains
HTH_Tnp_Tc5
DDE_1
Secondary Structure
Globular Domains
Disorder Score
Coiled-Coil
Transmembrane
0
100
200
300
400
500
600
700
800
900

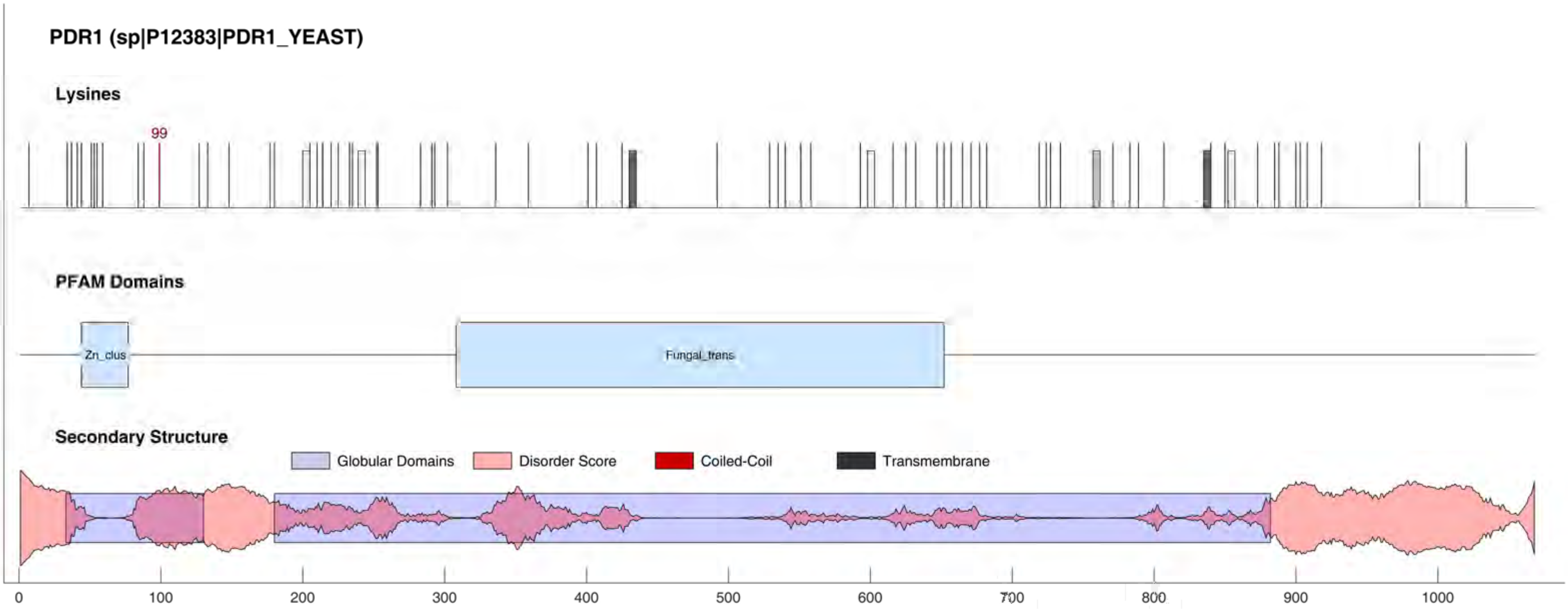

PDR1 (sp|P12383|PDR1_YEAST)
Lysines
99
PFAM Domains
Zn_clus
Fungal_trans
Secondary Structure
Globular Domains
Disorder Score
Coiled-Coil
Transmembrane
0
100
200
300
400
500
600
700
800
900
1000

#### PDS5 (sp|Q04264|PDS5_YEAST)

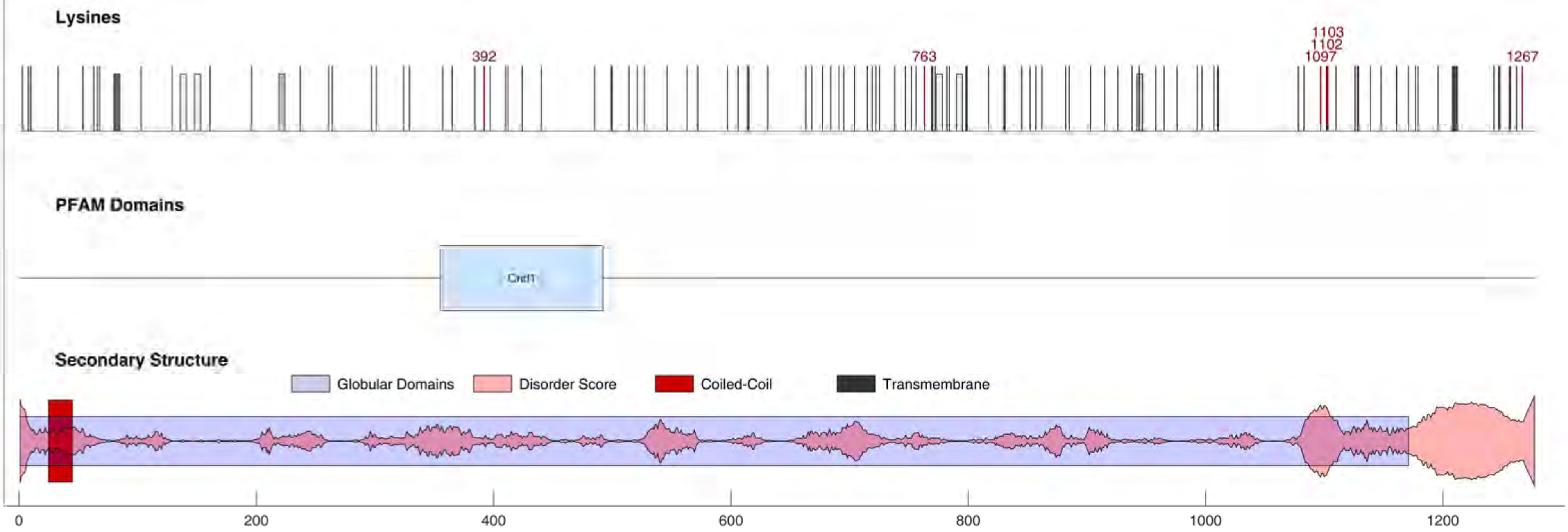

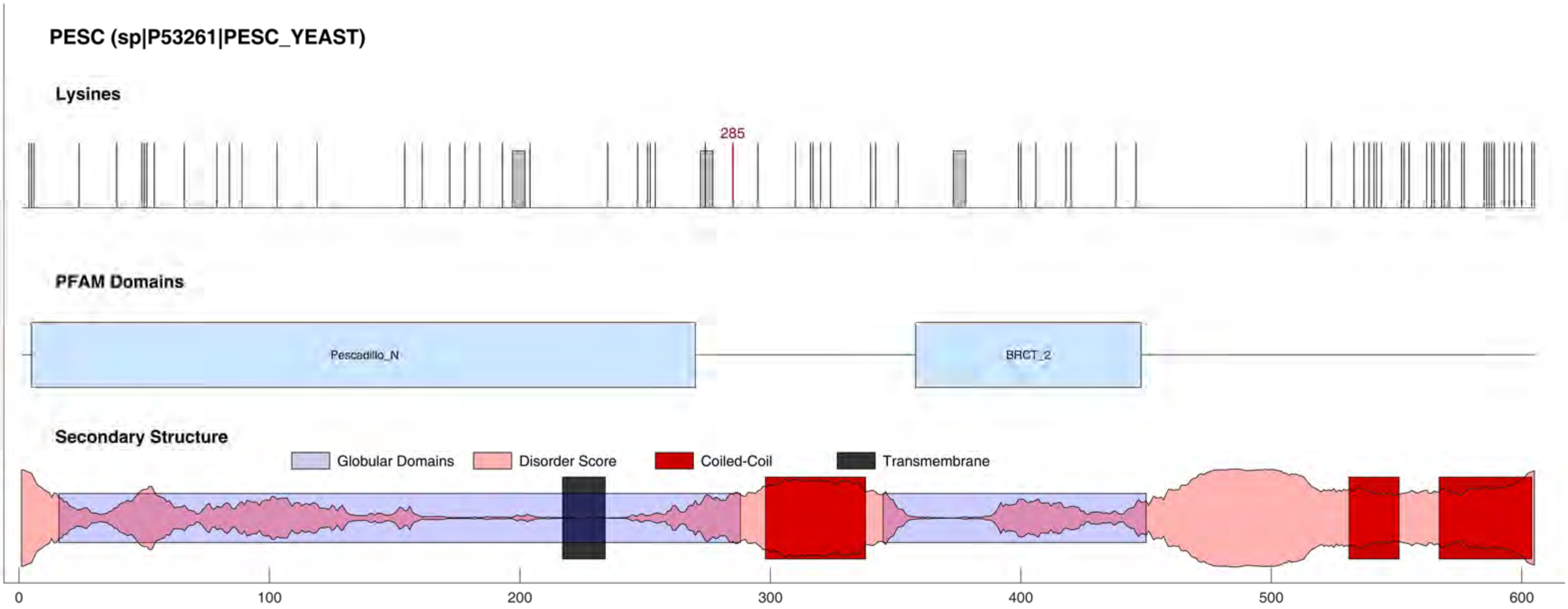
PESC (sp|P53261|PESC_YEAST)
Lysines
285
PFAM Domains
Pescadillo_N
BRCT_2
Secondary Structure
Globular Domains
Disorder Score
Coiled-Coil
Transmembrane
0
100
200
300
400
500
600

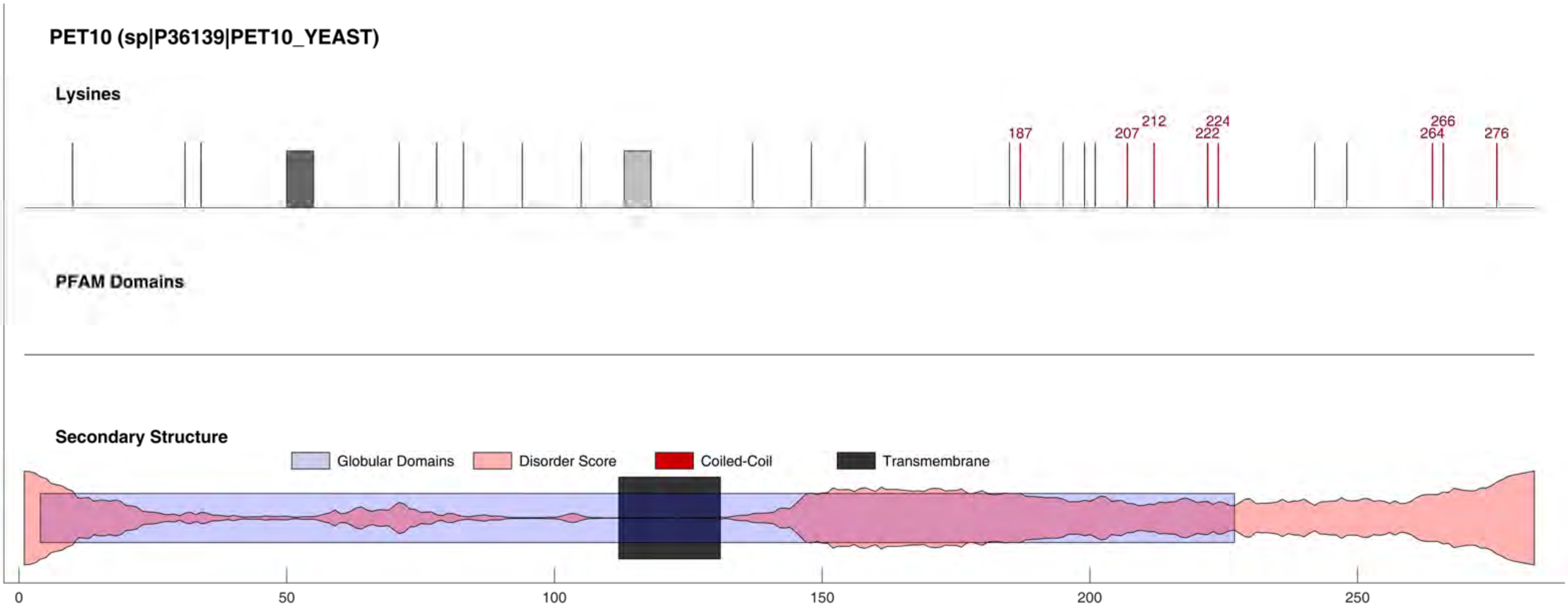
PET10 (sp|P36139|PET10_YEAST)
Lysines
187
207
212
222
224
264
266
276
PFAM Domains
Secondary Structure
Globular Domains
Disorder Score
Coiled-Coil
Transmembrane
0
50
100
150
200
250

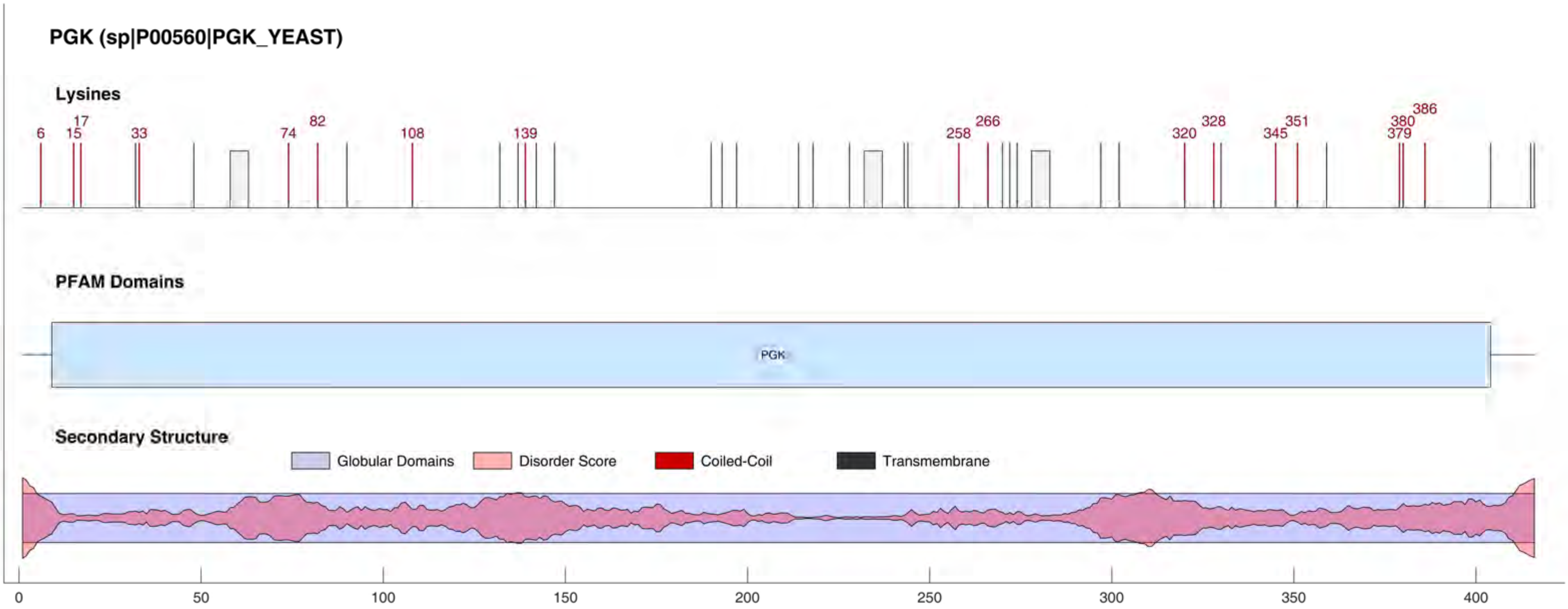

PGK (sp|P00560|PGK_YEAST)
Lysines
6
15
17
33
74
82
108
139
258
266
320
328
345
351
379
380
386
PFAM Domains
PGK
Secondary Structure
Globular Domains
Disorder Score
Coiled-Coil
Transmembrane
0
50
100
150
200
250
300
350
400

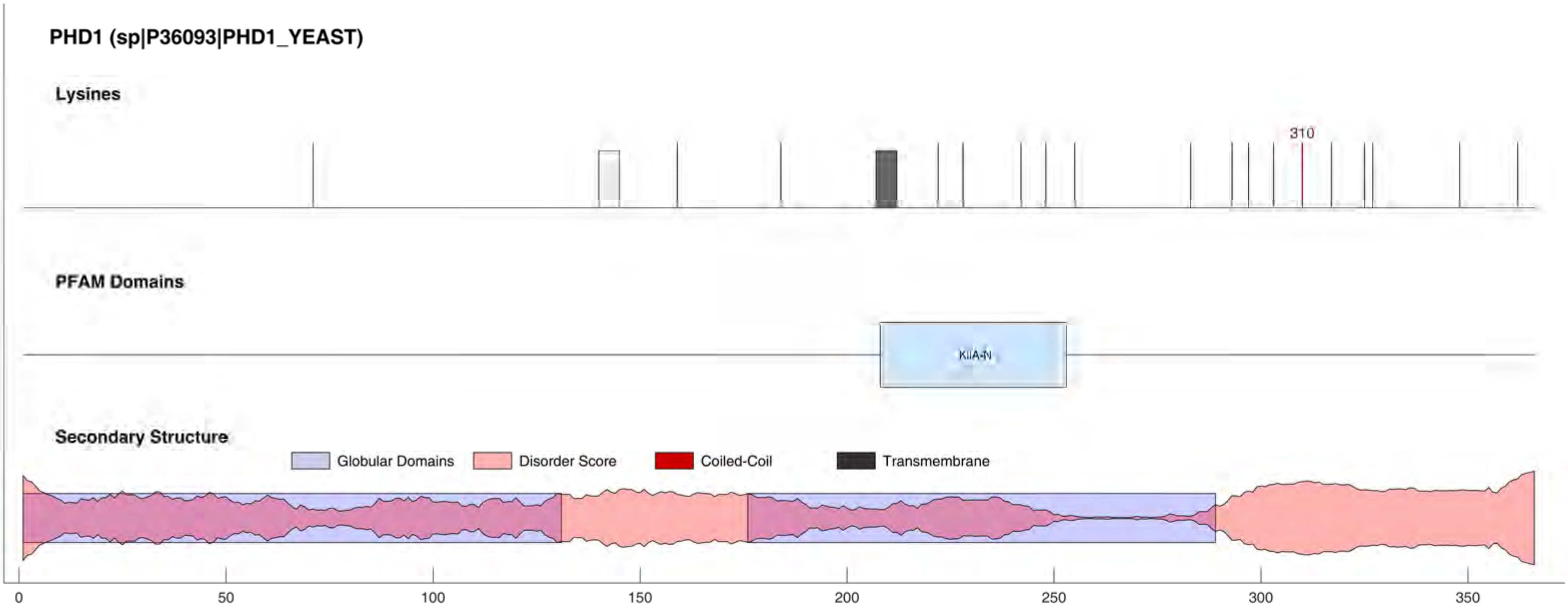

PHD1 (sp|P36093|PHD1_YEAST)
Lysines
310
PFAM Domains
KilA-N
Secondary Structure
Globular Domains
Disorder Score
Coiled-Coil
Transmembrane
0
50
100
150
200
250
300
350

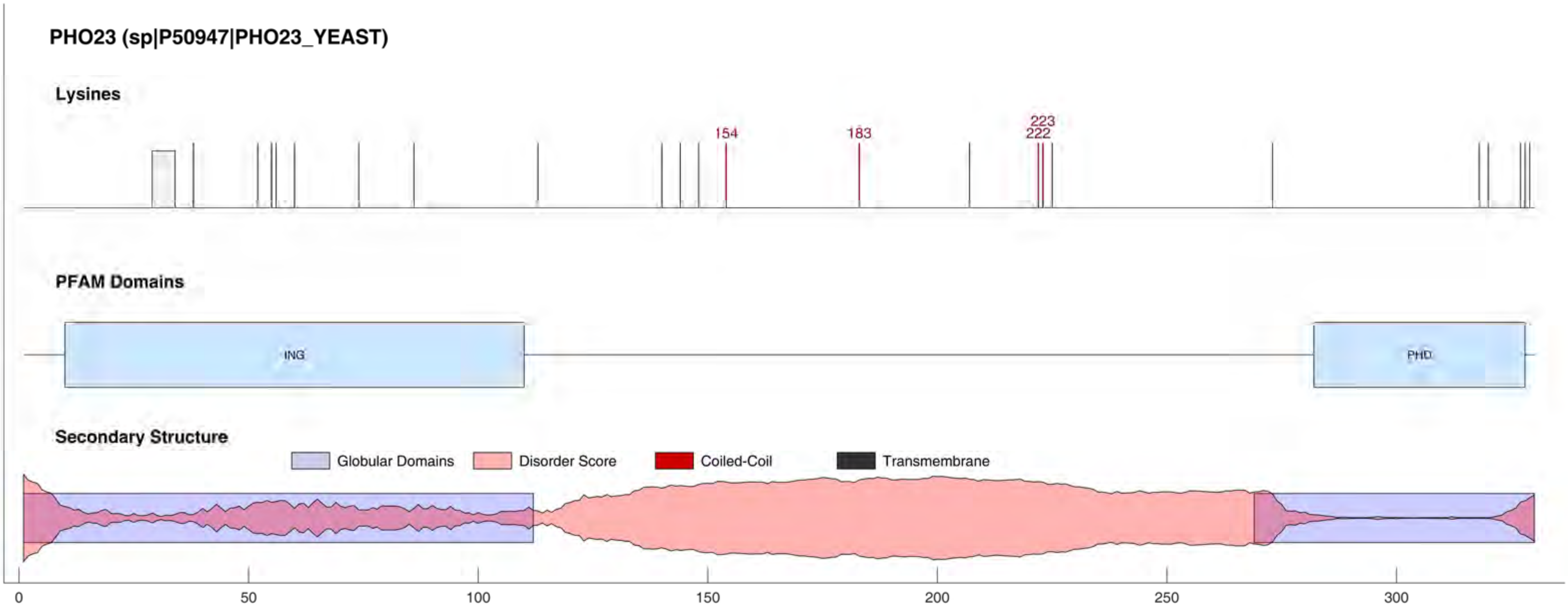
PHO23 (sp|P50947|PHO23_YEAST)
Lysines
154
183
223
222
PFAM Domains
ING
PHD
Secondary Structure
Globular Domains
Disorder Score
Coiled-Coil
Transmembrane
0
50
100
150
200
250
300

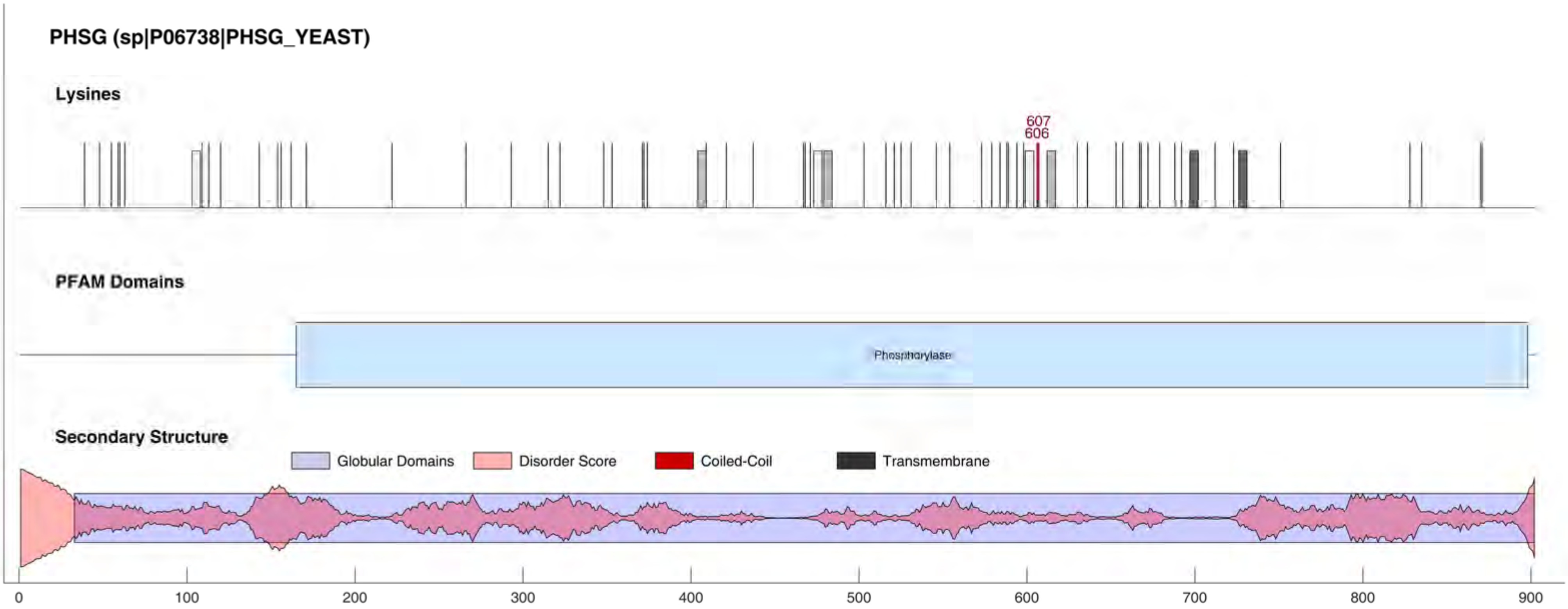
PHSG (sp|P06738|PHSG_YEAST)
Lysines
607
606
PFAM Domains
Phosphorylase
Secondary Structure
Globular Domains
Disorder Score
Coiled-Coil
Transmembrane
0
100
200
300
400
500
600
700
800
900

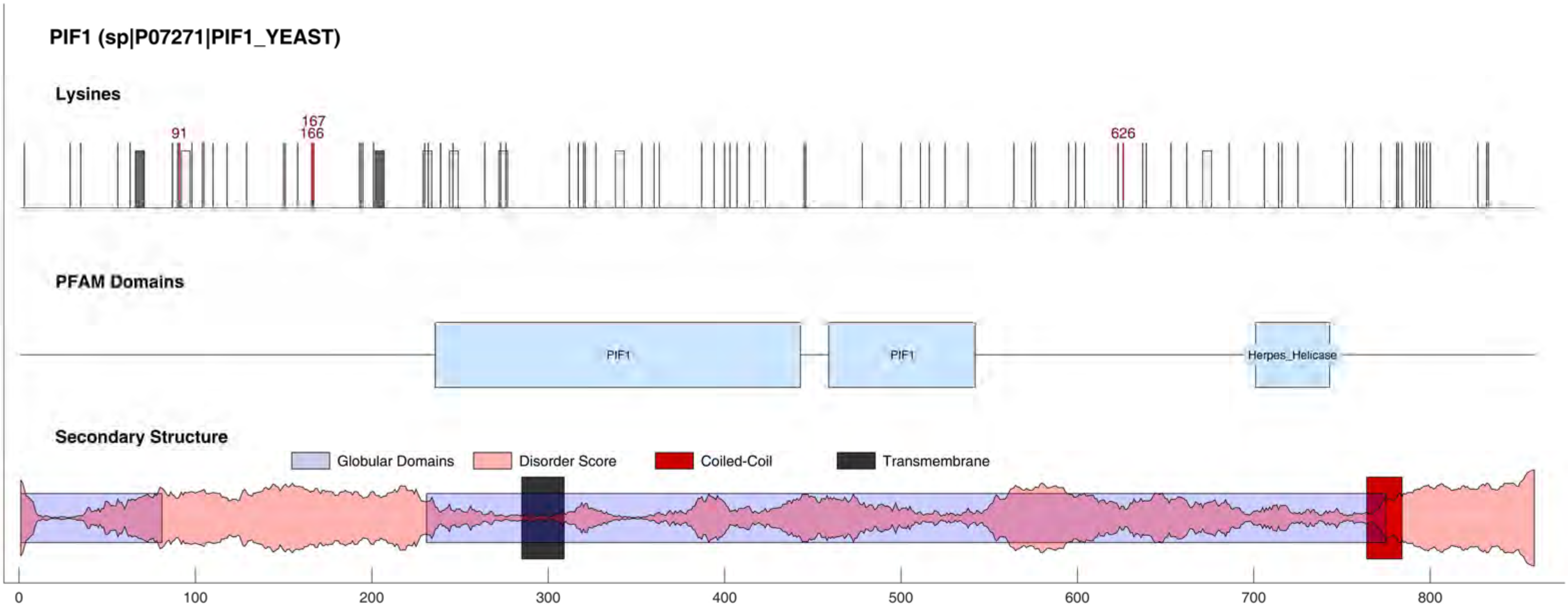
PIF1 (sp|P07271|PIF1_YEAST)
Lysines
91
167
166
626
PFAM Domains
PIF1
PIF1
Herpes_Helicase
Secondary Structure
Globular Domains
Disorder Score
Coiled-Coil
Transmembrane
0
100
200
300
400
500
600
700
800

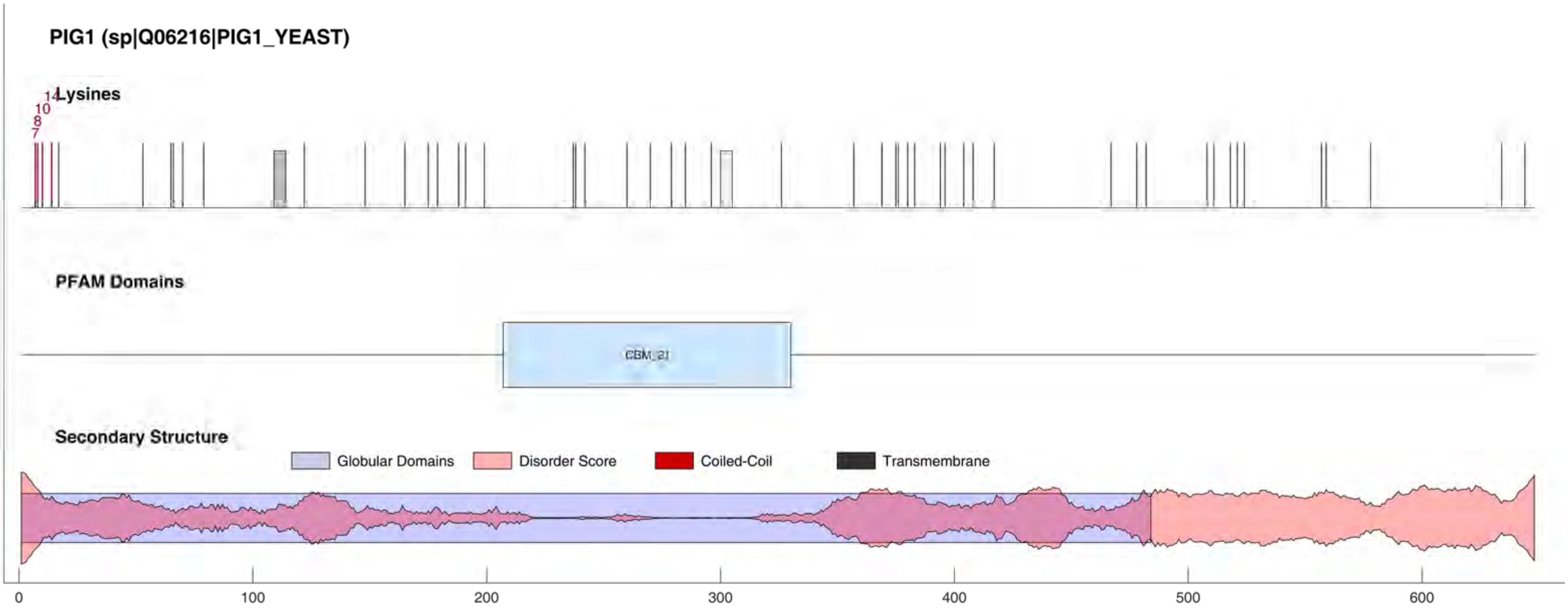

PIG1 (sp|Q06216|PIG1_YEAST)
Lysines
7
8
10
14
PFAM Domains
CBM_21
Secondary Structure
Globular Domains
Disorder Score
Coiled-Coil
Transmembrane
0
100
200
300
400
500
600

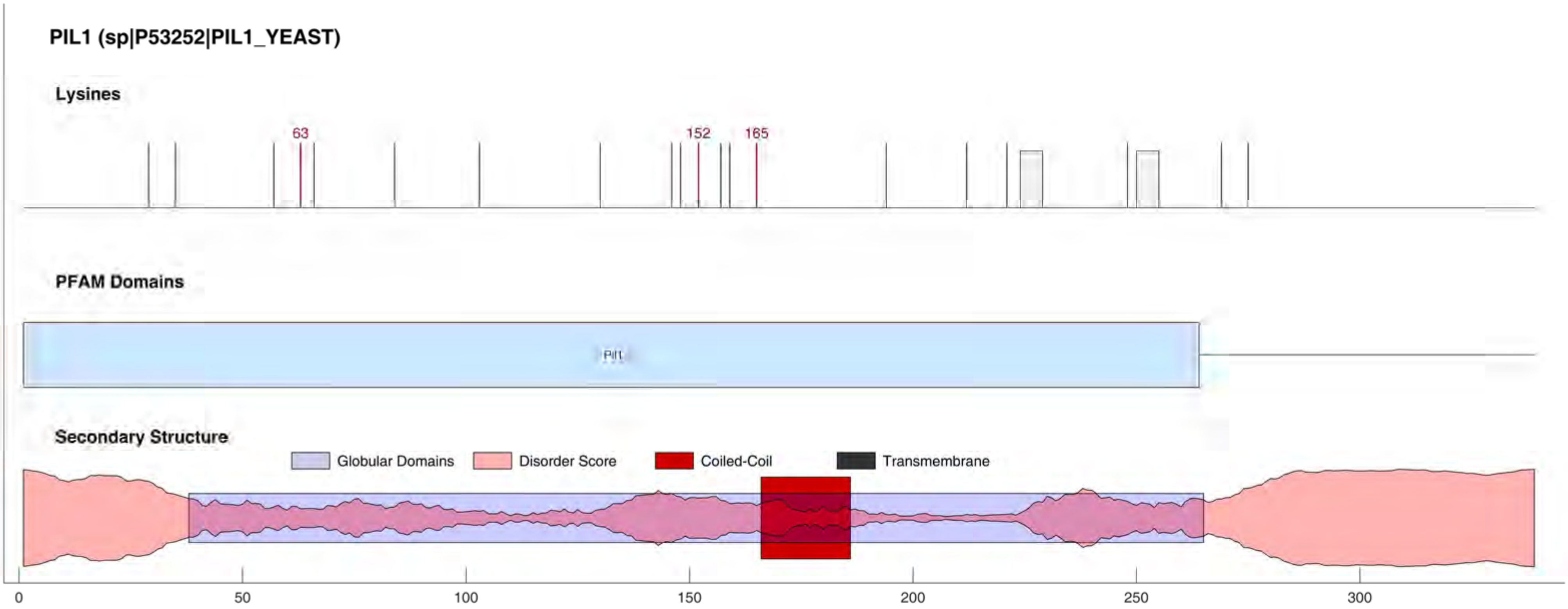

PIL1 (sp|P53252|PIL1_YEAST)
Lysines
63
152
165
PFAM Domains
Pil1
Secondary Structure
Globular Domains
Disorder Score
Coiled-Coil
Transmembrane
0
50
100
150
200
250
300

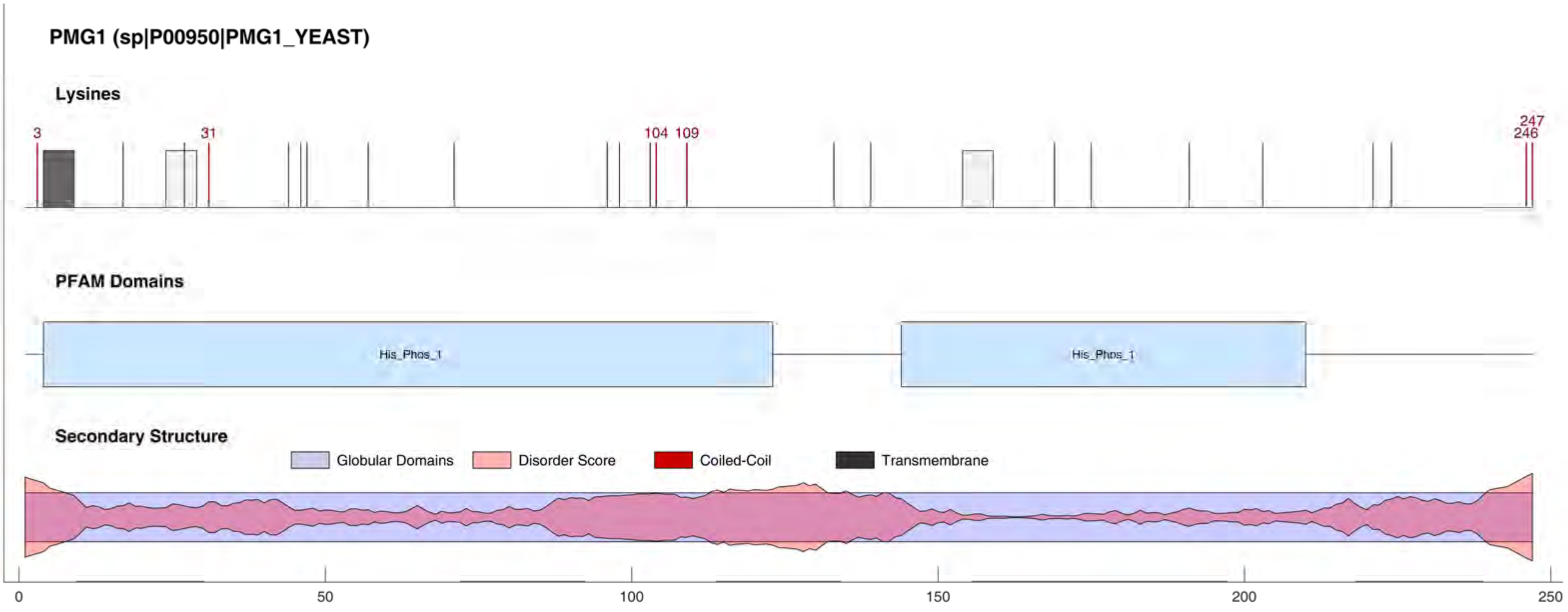

PMG1 (sp|P00950|PMG1_YEAST)
Lysines
3
31
104
109
247
246
PFAM Domains
His_Phos_1
His_Phos_1
Secondary Structure
Globular Domains
Disorder Score
Coiled-Coil
Transmembrane
0
50
100
150
200
250

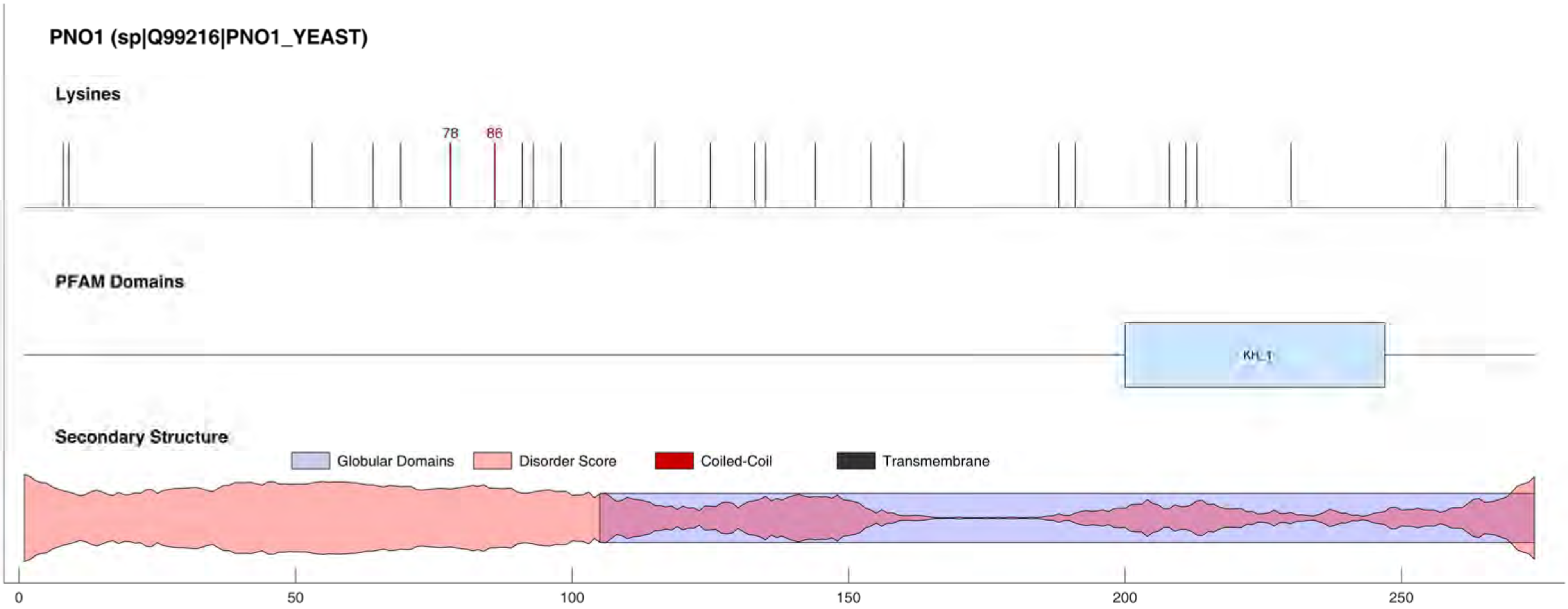
PNO1 (sp|Q99216|PNO1_YEAST)
Lysines
78
86
PFAM Domains
KH_1
Secondary Structure
Globular Domains
Disorder Score
Coiled-Coil
Transmembrane
0
50
100
150
200
250

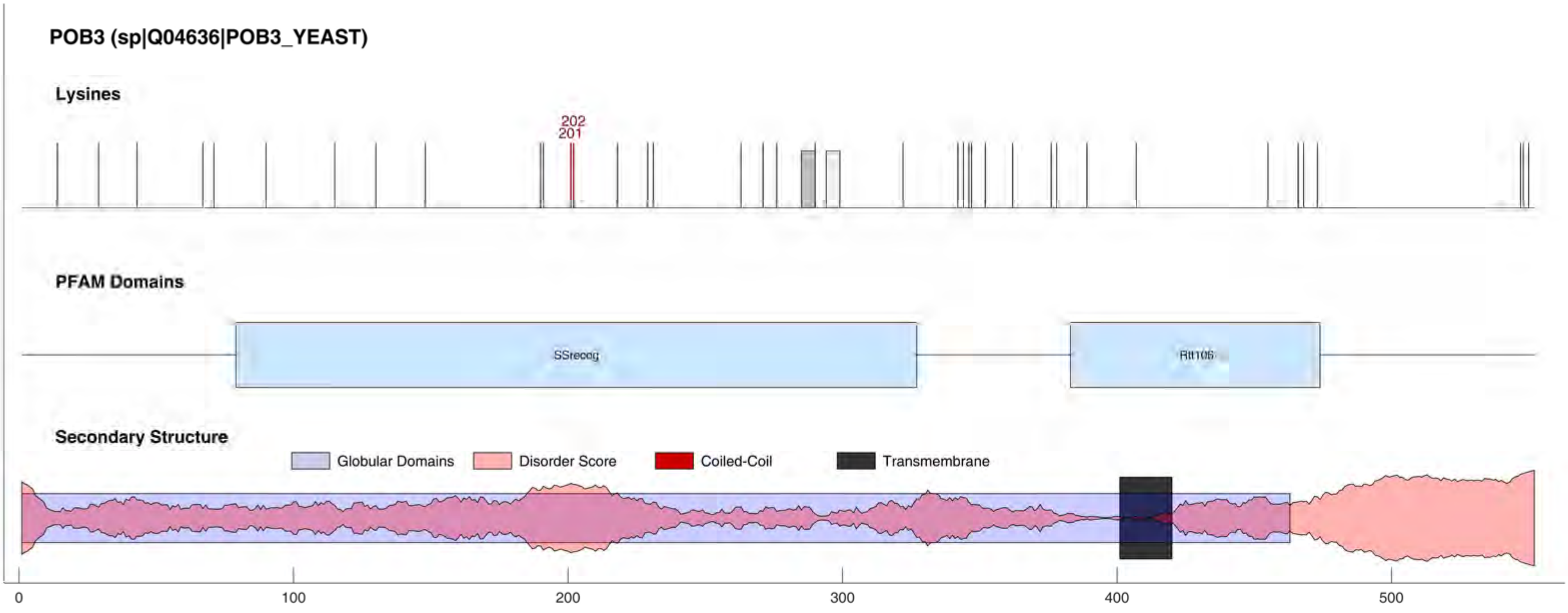

POB3 (sp|Q04636|POB3_YEAST)
Lysines
202
201
PFAM Domains
SSrecog
Rtt106
Secondary Structure
Globular Domains
Disorder Score
Coiled-Coil
Transmembrane
0
100
200
300
400
500

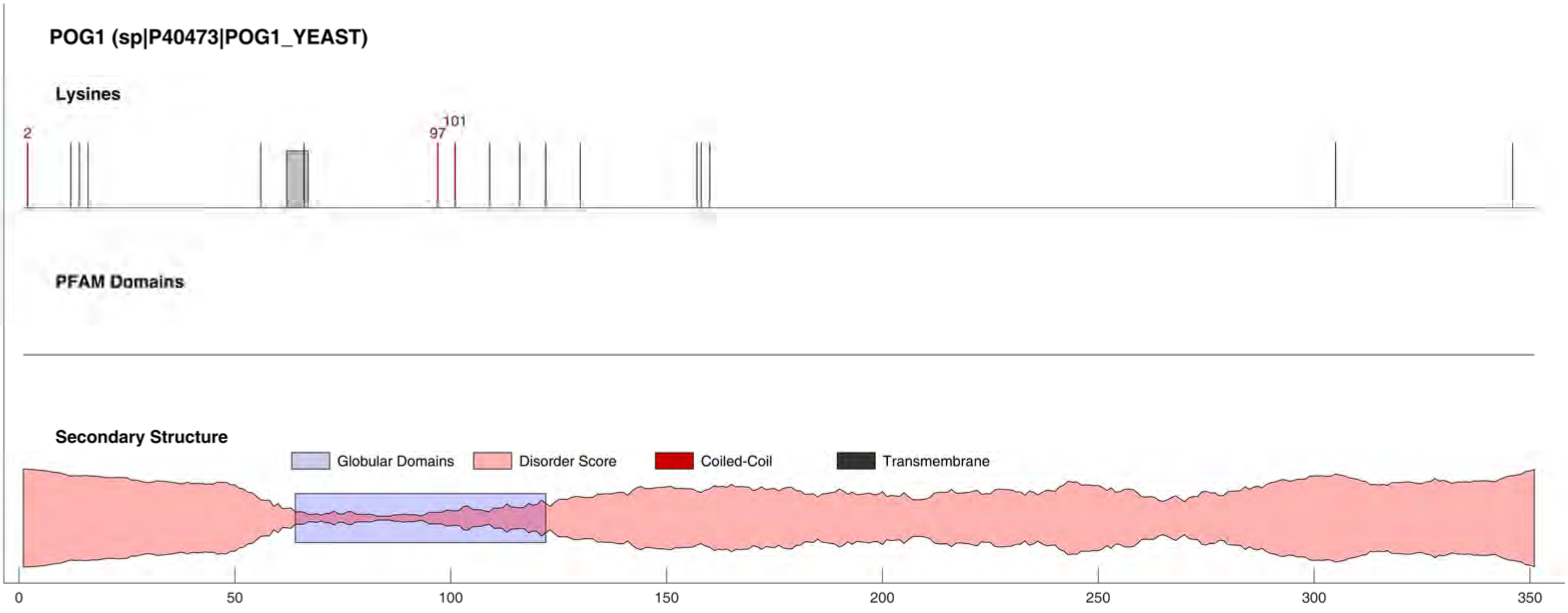
POG1 (sp|P40473|POG1_YEAST)
Lysines
2
97
101
PFAM Domains
Secondary Structure
Globular Domains
Disorder Score
Coiled-Coil
Transmembrane
0
50
100
150
200
250
300
350

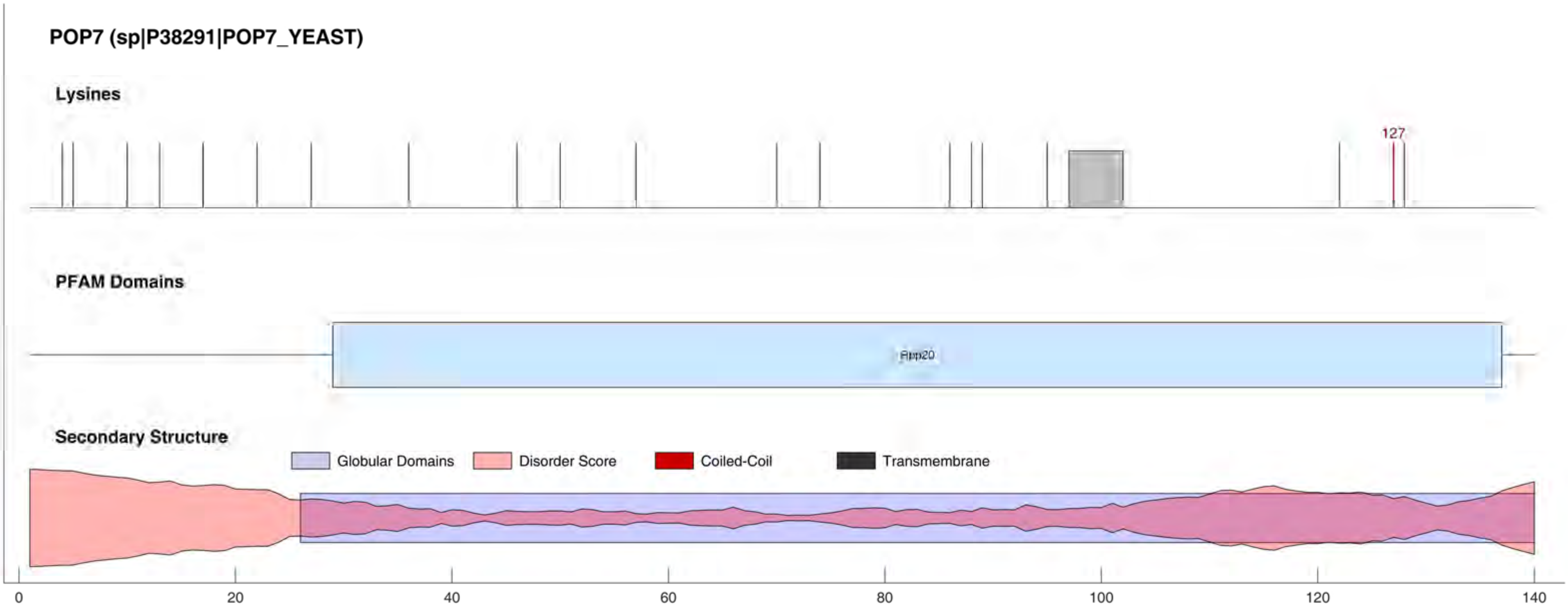

POP7 (sp|P38291|POP7_YEAST)
Lysines
127
PFAM Domains
Rpp20
Secondary Structure
Globular Domains
Disorder Score
Coiled-Coil
Transmembrane
0
20
40
60
80
100
120
140

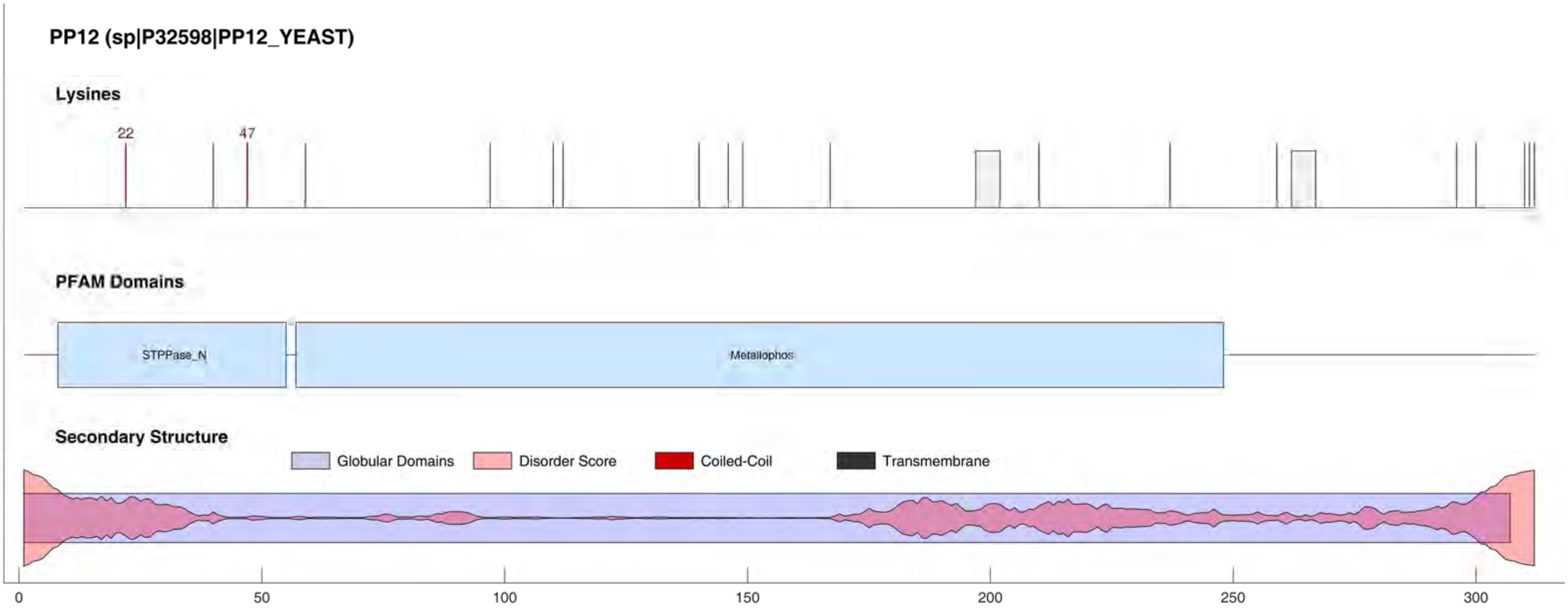
PP12 (sp|P32598|PP12_YEAST)
Lysines
22
47
PFAM Domains
STPPase_N
Metallophos
Secondary Structure
Globular Domains
Disorder Score
Coiled-Coil
Transmembrane
0
50
100
150
200
250
300

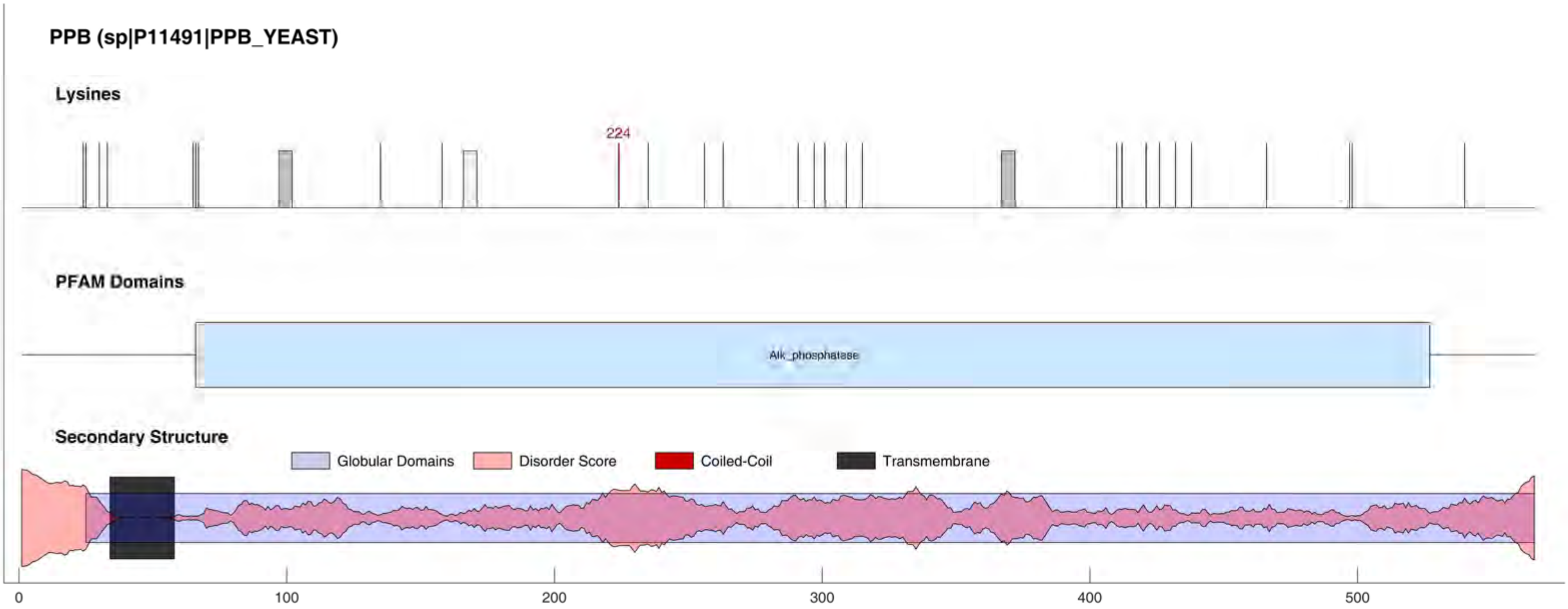
PPB (sp|P11491|PPB_YEAST)
Lysines
224
PFAM Domains
Alk_phosphatase
Secondary Structure
Globular Domains
Disorder Score
Coiled-Coil
Transmembrane
0
100
200
300
400
500

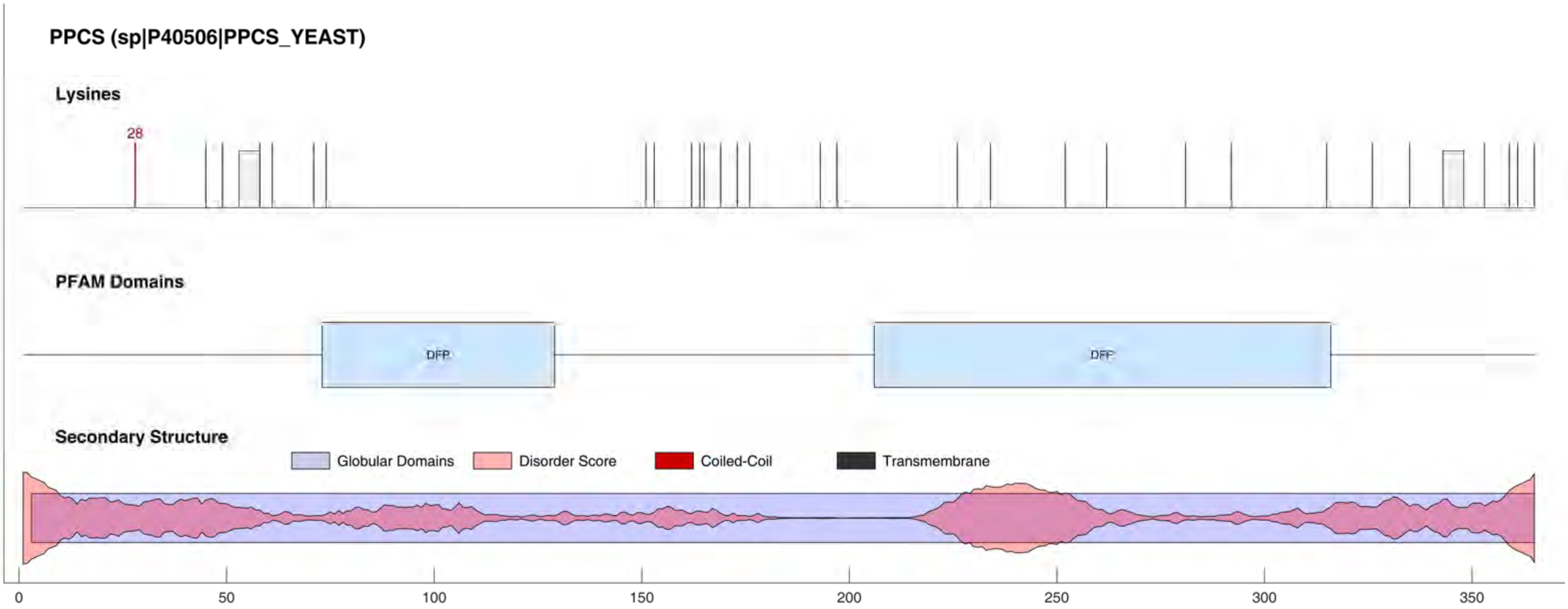

PPCS (sp|P40506|PPCS_YEAST)
Lysines
28
PFAM Domains
DFP
DFP
Secondary Structure
Globular Domains
Disorder Score
Coiled-Coil
Transmembrane
0
50
100
150
200
250
300
350

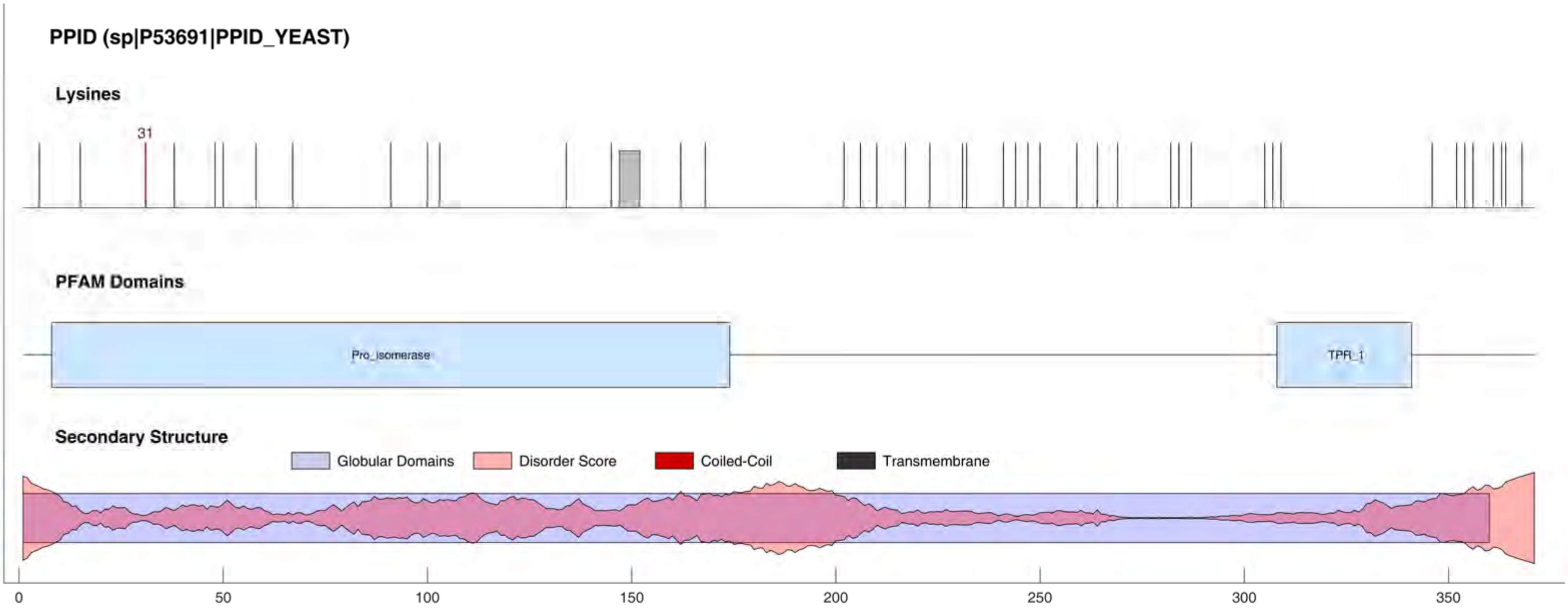

PPID (sp|P53691|PPID_YEAST)
Lysines
31
PFAM Domains
Pro_isomerase
TPR_1
Secondary Structure
Globular Domains
Disorder Score
Coiled-Coil
Transmembrane
0
50
100
150
200
250
300
350

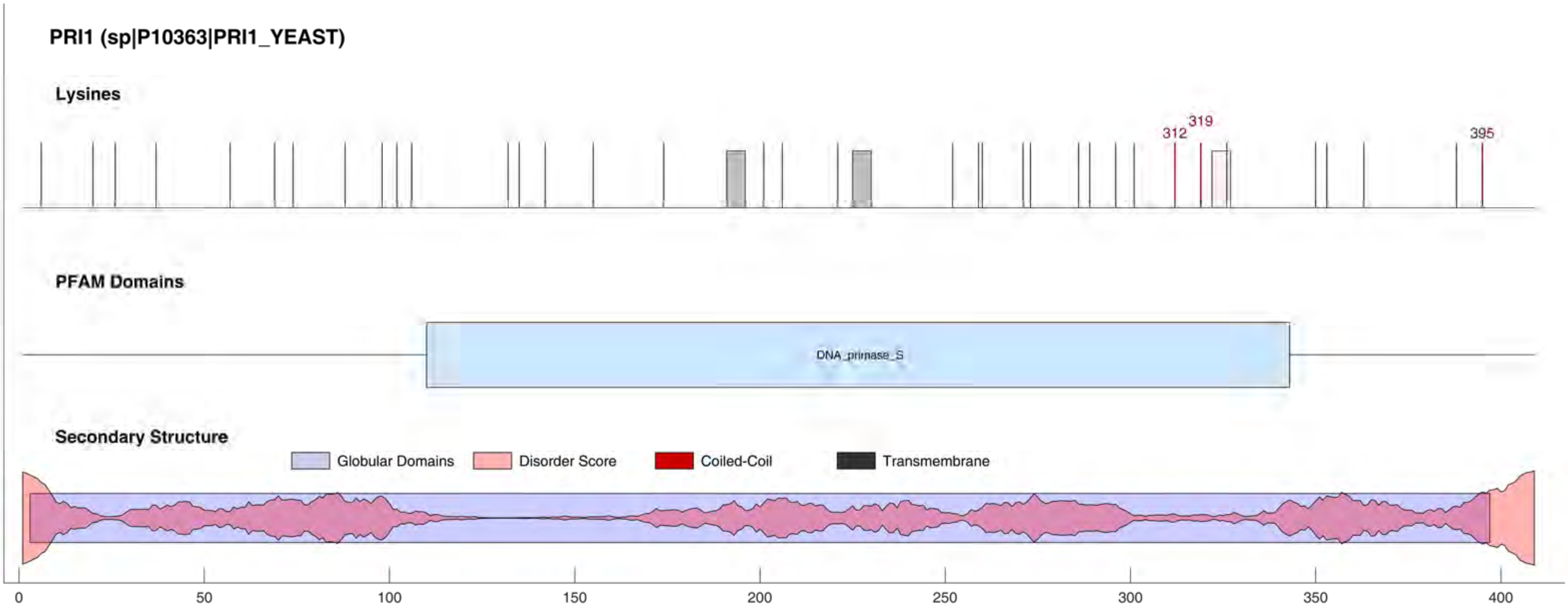
PRI1 (sp|P10363|PRI1_YEAST)
Lysines
312
319
395
PFAM Domains
DNA_primase_S
Secondary Structure
Globular Domains
Disorder Score
Coiled-Coil
Transmembrane
0
50
100
150
200
250
300
350
400

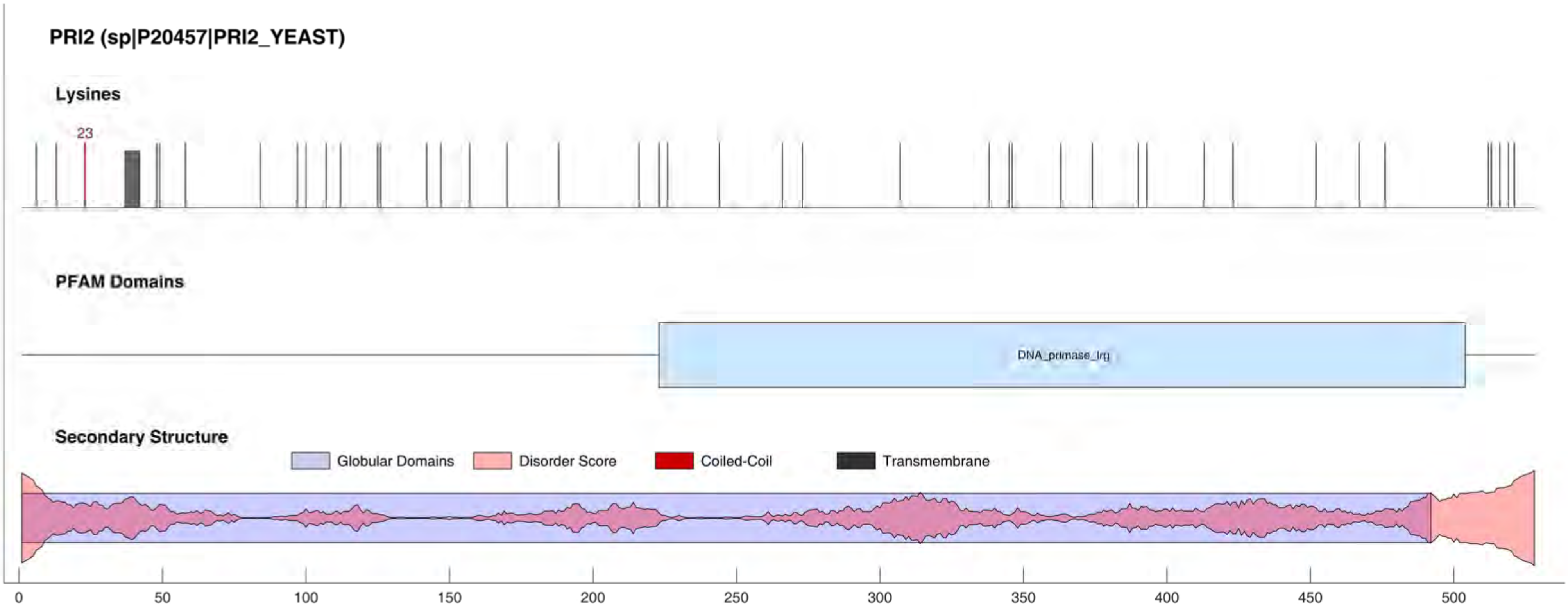
PRI2 (sp|P20457|PRI2_YEAST)
Lysines
23
PFAM Domains
DNA_primase_lrg
Secondary Structure
Globular Domains
Disorder Score
Coiled-Coil
Transmembrane
0
50
100
150
200
250
300
350
400
450
500

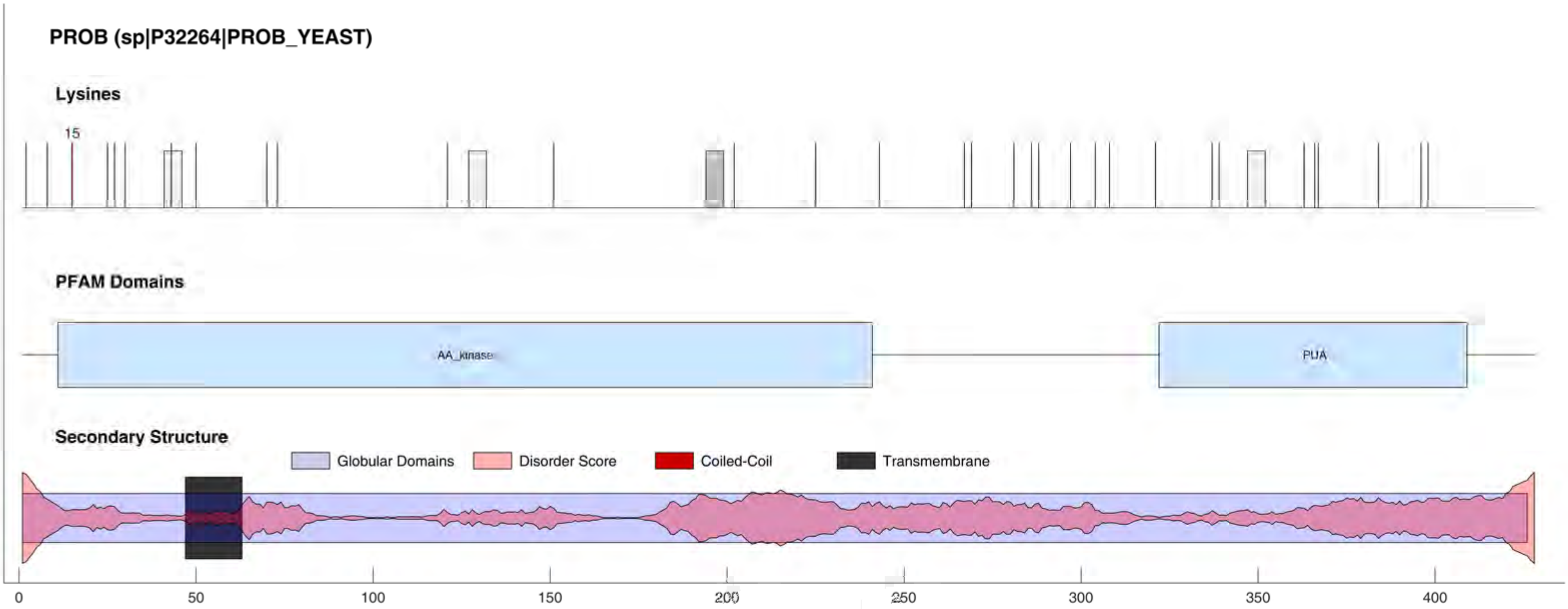
PROB (sp|P32264|PROB_YEAST)
Lysines
15
PFAM Domains
AA_kinase
PUA
Secondary Structure
Globular Domains
Disorder Score
Coiled-Coil
Transmembrane
0
50
100
150
200
250
300
350
400

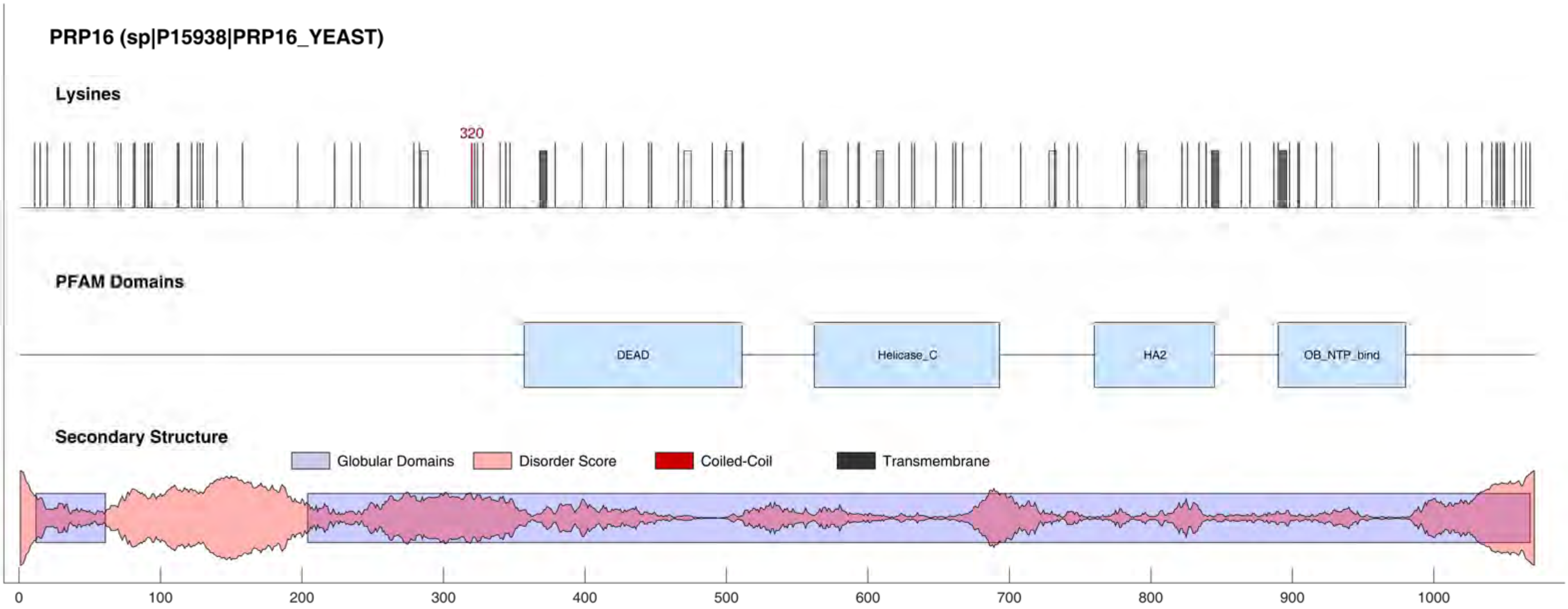

PRP16 (sp|P15938|PRP16_YEAST)
Lysines
320
PFAM Domains
DEAD
Helicase_C
HA2
OB_NTP_bind
Secondary Structure
Globular Domains
Disorder Score
Coiled-Coil
Transmembrane
0
100
200
300
400
500
600
700
800
900
1000

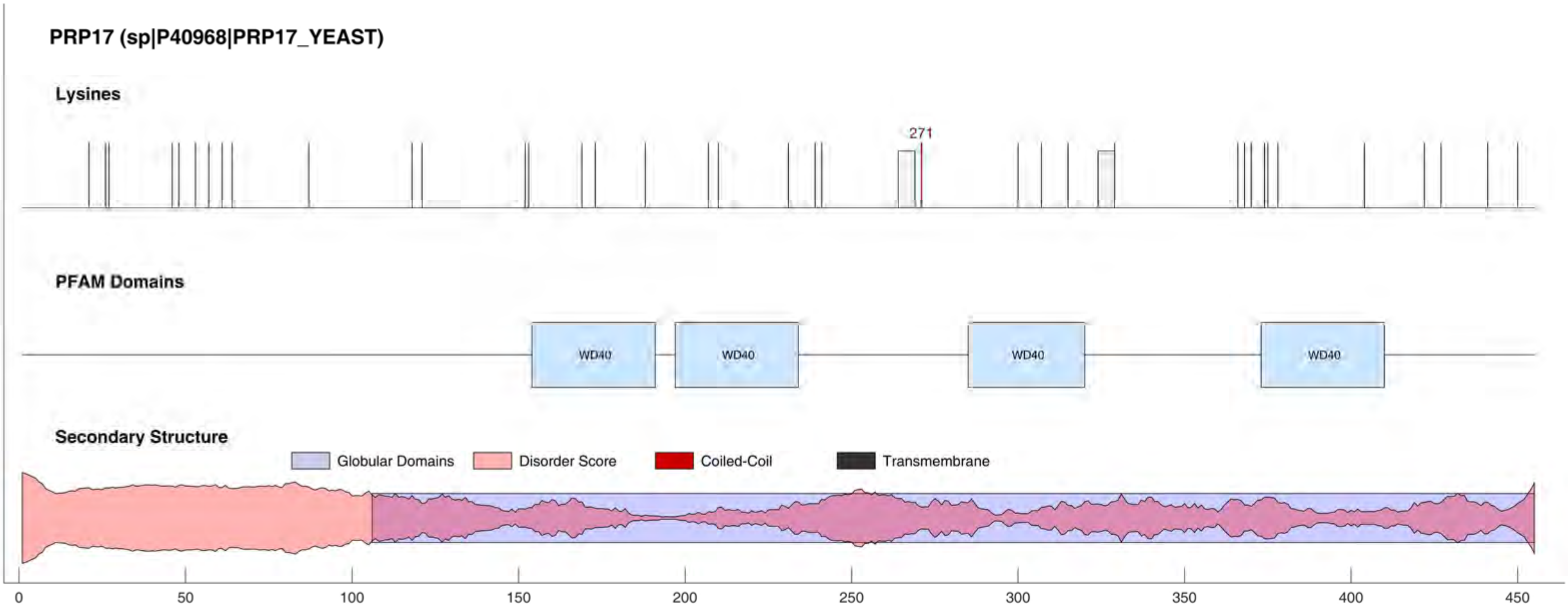
PRP17 (sp|P40968|PRP17_YEAST)
Lysines
271
PFAM Domains
WD40
WD40
WD40
WD40
Secondary Structure
Globular Domains
Disorder Score
Coiled-Coil
Transmembrane
0
50
100
150
200
250
300
350
400
450

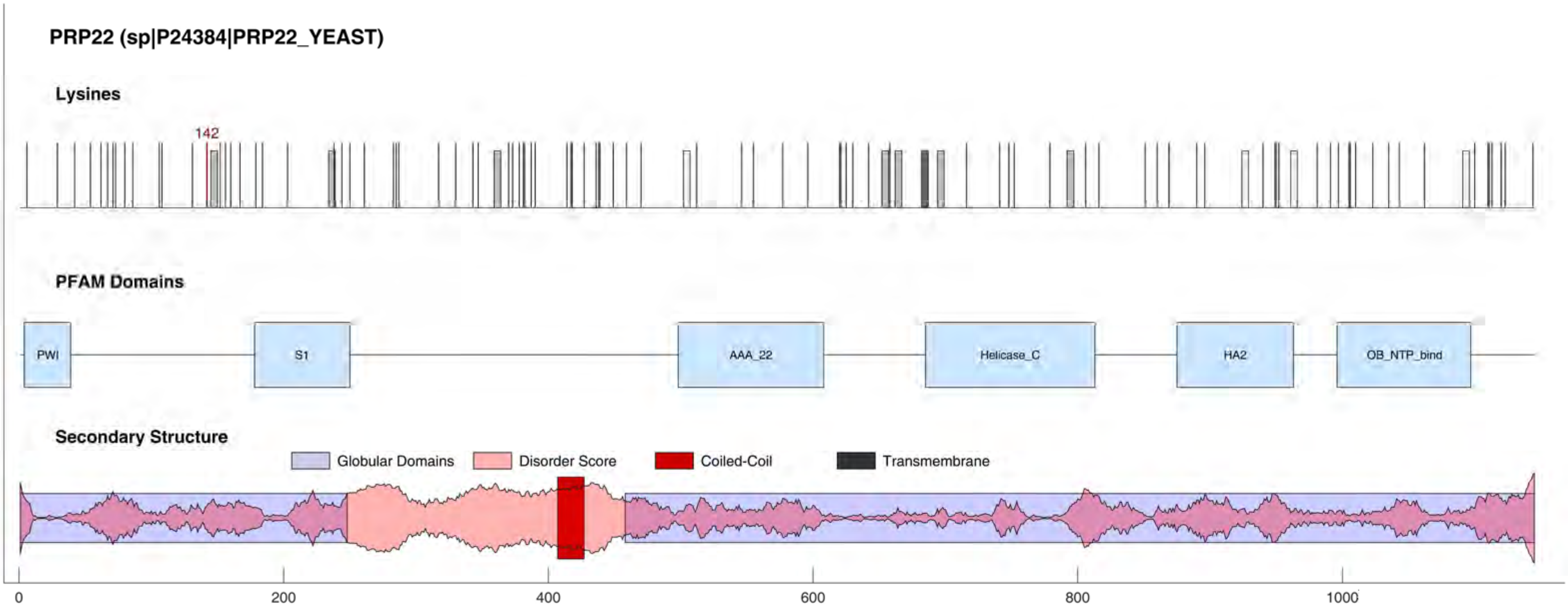

PRP22 (sp|P24384|PRP22_YEAST)
Lysines
142
PFAM Domains
PWI
S1
AAA_22
Helicase_C
HA2
OB_NTP_bind
Secondary Structure
Globular Domains
Disorder Score
Coiled-Coil
Transmembrane
0
200
400
600
800
1000

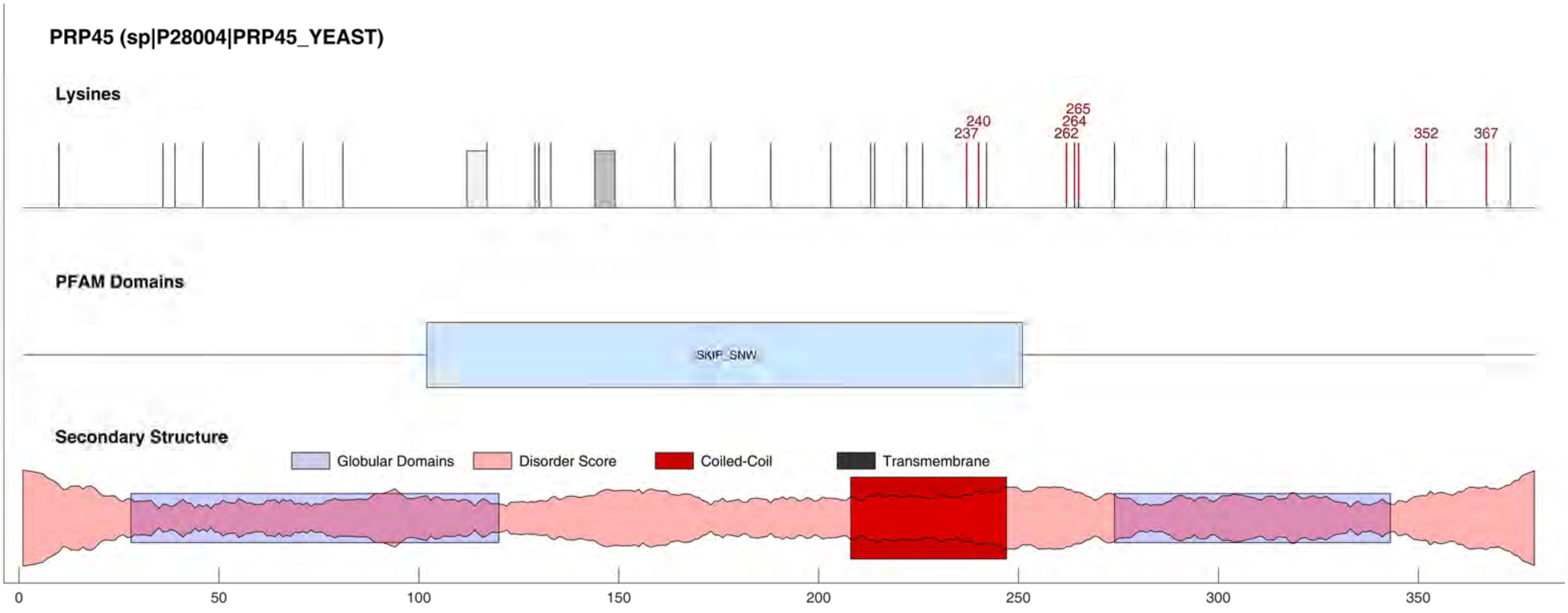
PRP45 (sp|P28004|PRP45_YEAST)
Lysines
237
240
262
264
265
352
367
PFAM Domains
SKIP_SNW
Secondary Structure
Globular Domains
Disorder Score
Coiled-Coil
Transmembrane
0
50
100
150
200
250
300
350

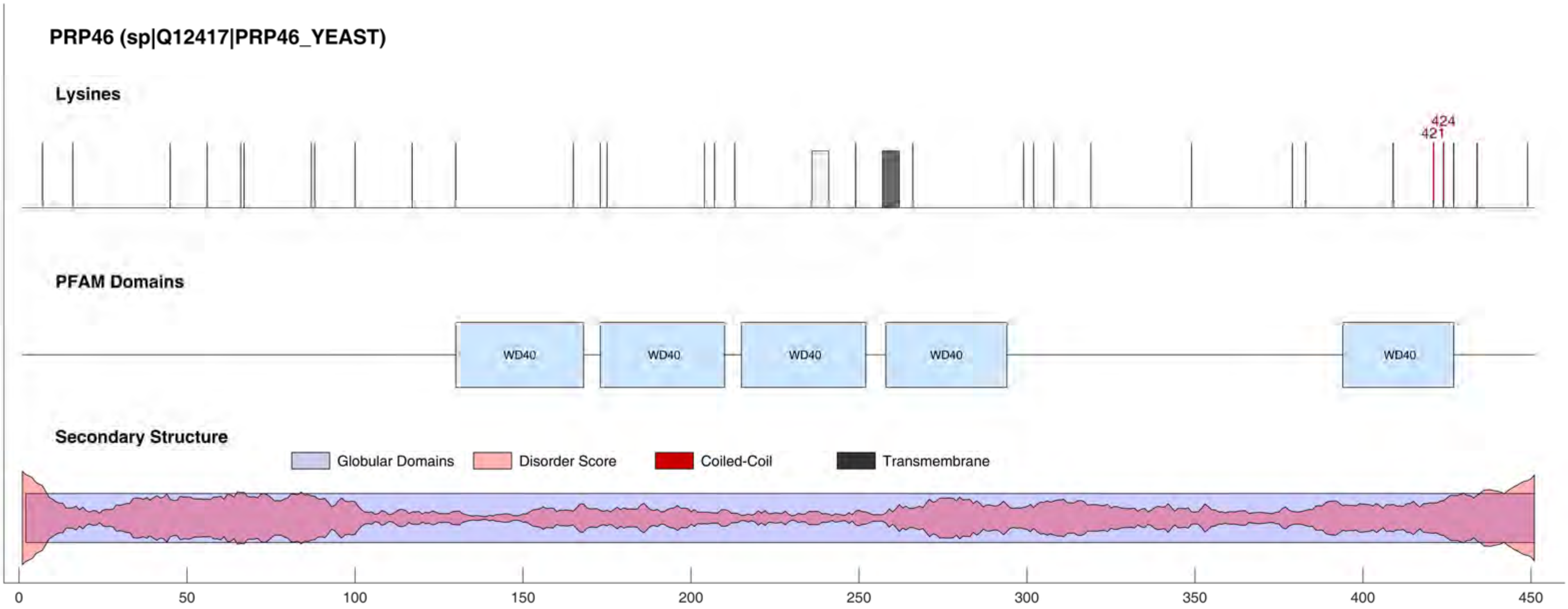
PRP46 (sp|Q12417|PRP46_YEAST)
Lysines
424
421
PFAM Domains
WD40
WD40
WD40
WD40
WD40
Secondary Structure
Globular Domains
Disorder Score
Coiled-Coil
Transmembrane
0
50
100
150
200
250
300
350
400
450

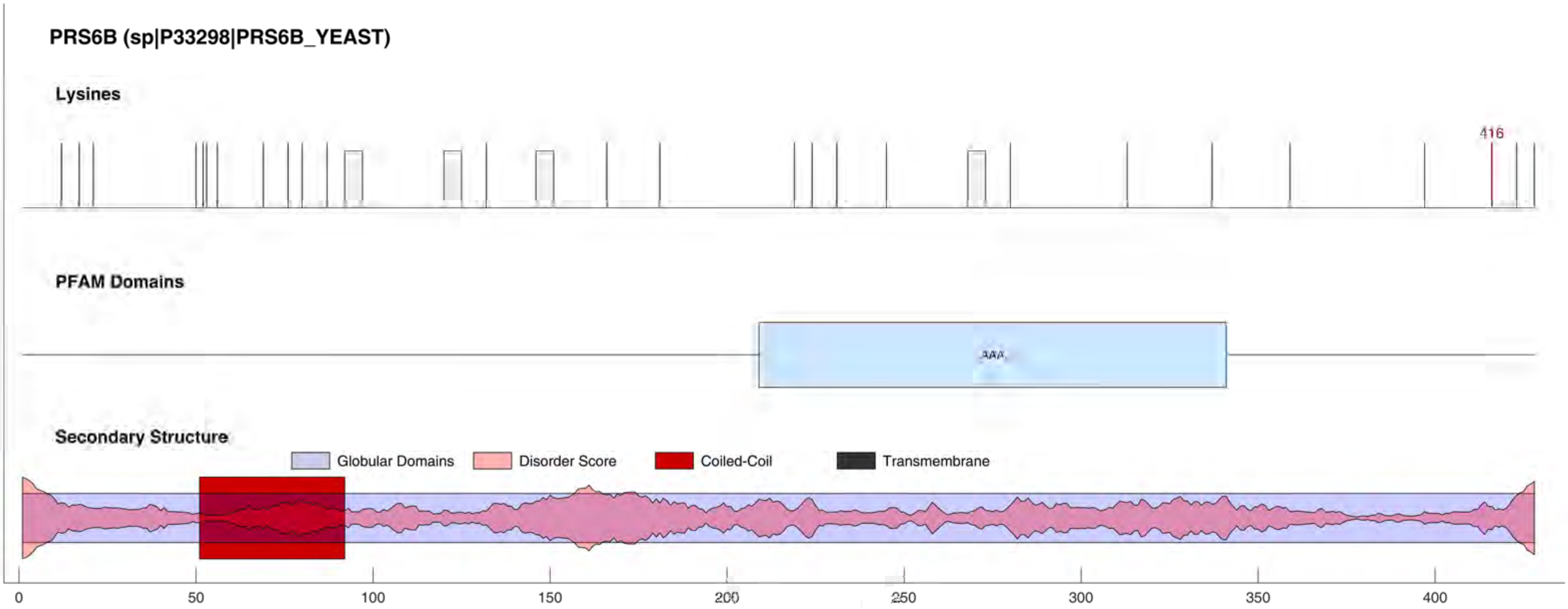
PRS6B (sp|P33298|PRS6B_YEAST)
Lysines
416
PFAM Domains
AAA
Secondary Structure
Globular Domains
Disorder Score
Coiled-Coil
Transmembrane
0
50
100
150
200
250
300
350
400

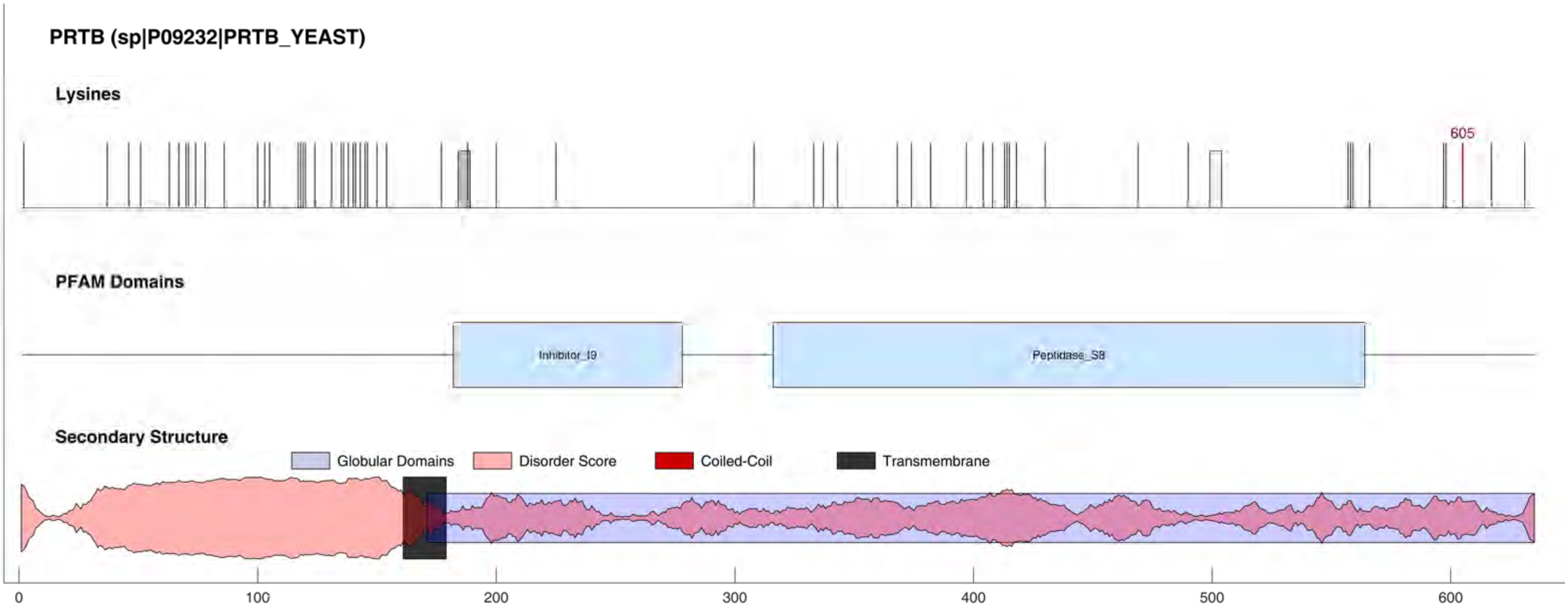
PRTB (sp|P09232|PRTB_YEAST)
Lysines
605
PFAM Domains
Inhibitor_I9
Peptidase_S8
Secondary Structure
Globular Domains
Disorder Score
Coiled-Coil
Transmembrane
0
100
200
300
400
500
600

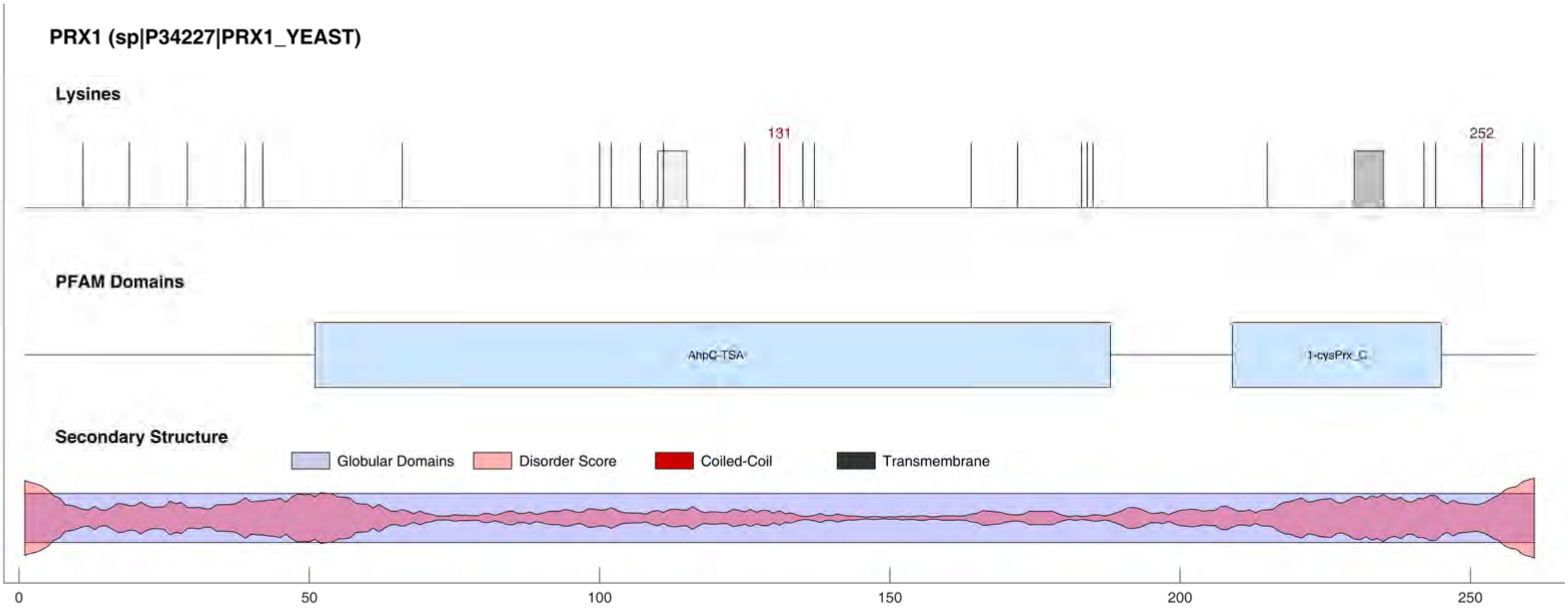
PRX1 (sp|P34227|PRX1_YEAST)
Lysines
131
252
PFAM Domains
AhpC-TSA
1-cysPrx_C
Secondary Structure
Globular Domains
Disorder Score
Coiled-Coil
Transmembrane
0
50
100
150
200
250

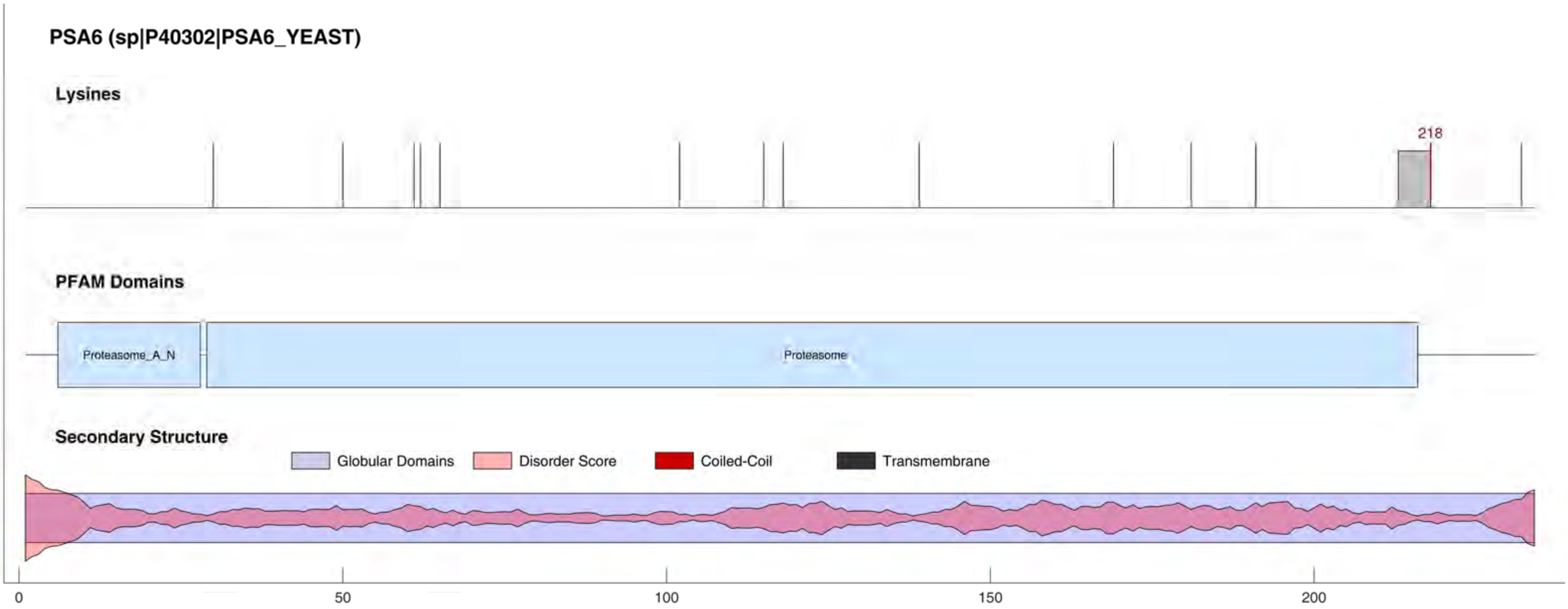
PSA6 (sp|P40302|PSA6_YEAST)
Lysines
218
PFAM Domains
Proteasome_A_N
Proteasome
Secondary Structure
Globular Domains
Disorder Score
Coiled-Coil
Transmembrane
0
50
100
150
200

#### PSB5 (sp|P30656|PSB5_YEAST)

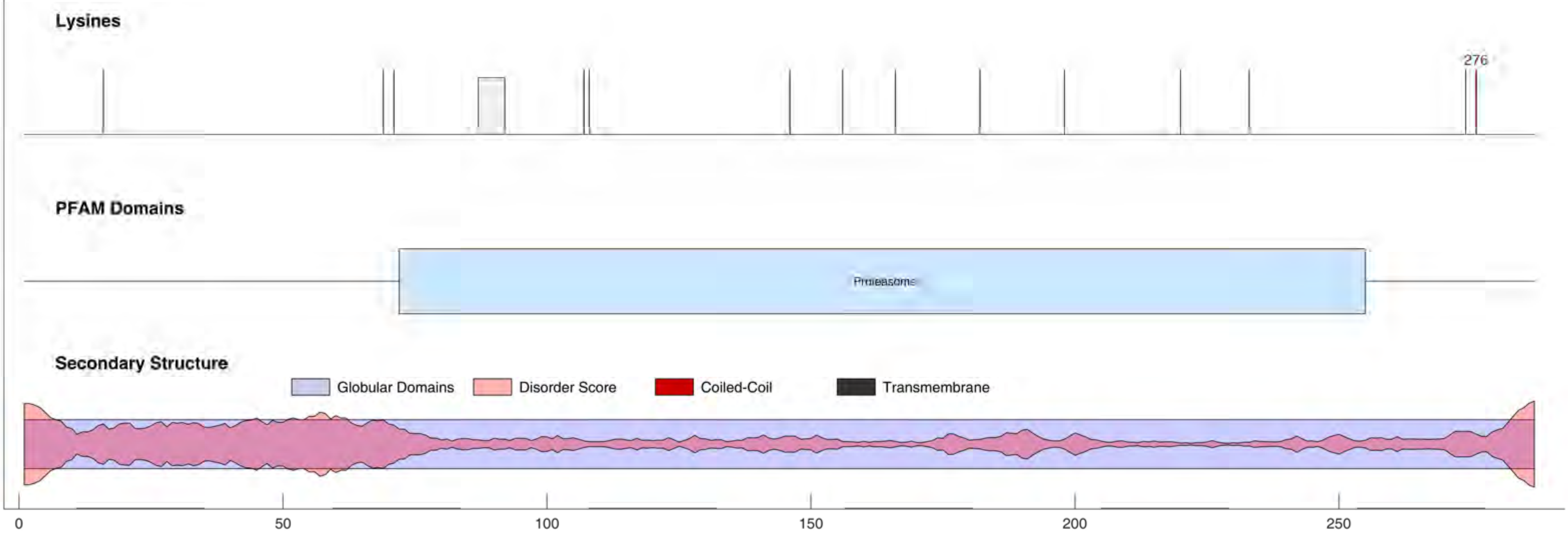

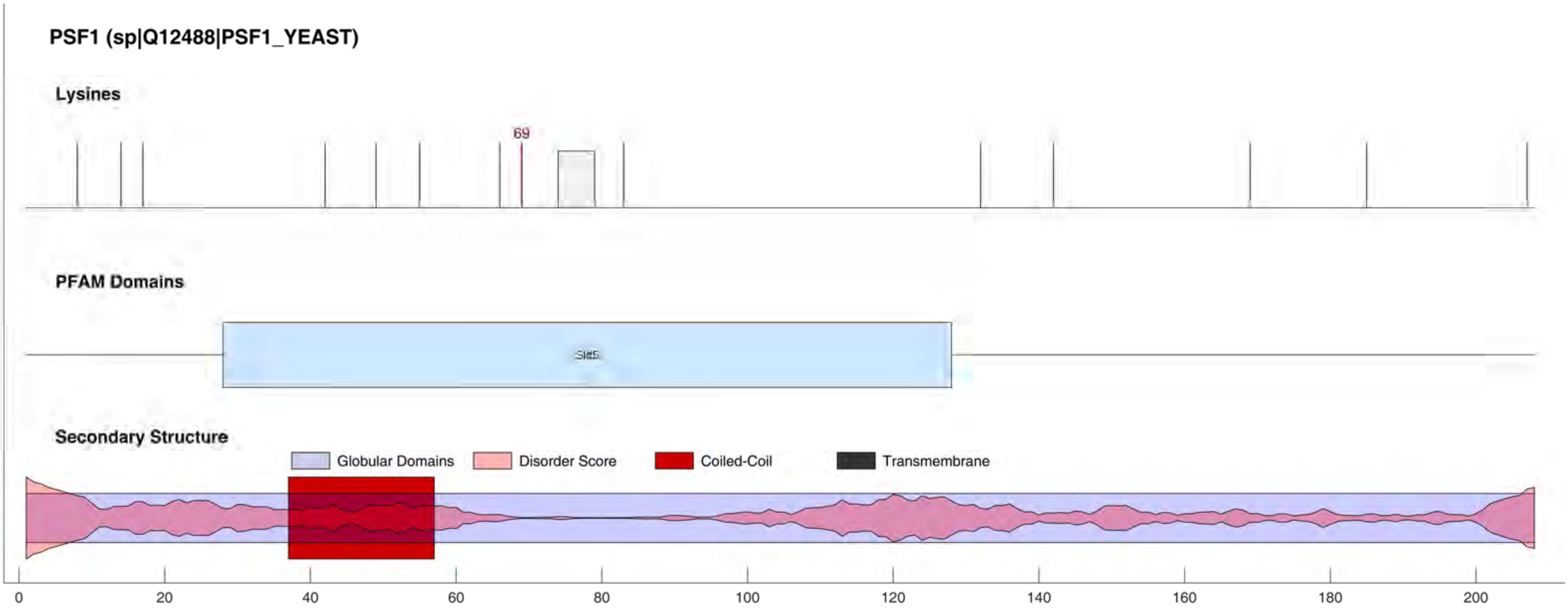
PSF1 (sp|Q12488|PSF1_YEAST)
Lysines
69
PFAM Domains
Sld5
Secondary Structure
Globular Domains
Disorder Score
Coiled-Coil
Transmembrane
0
20
40
60
80
100
120
140
160
180
200

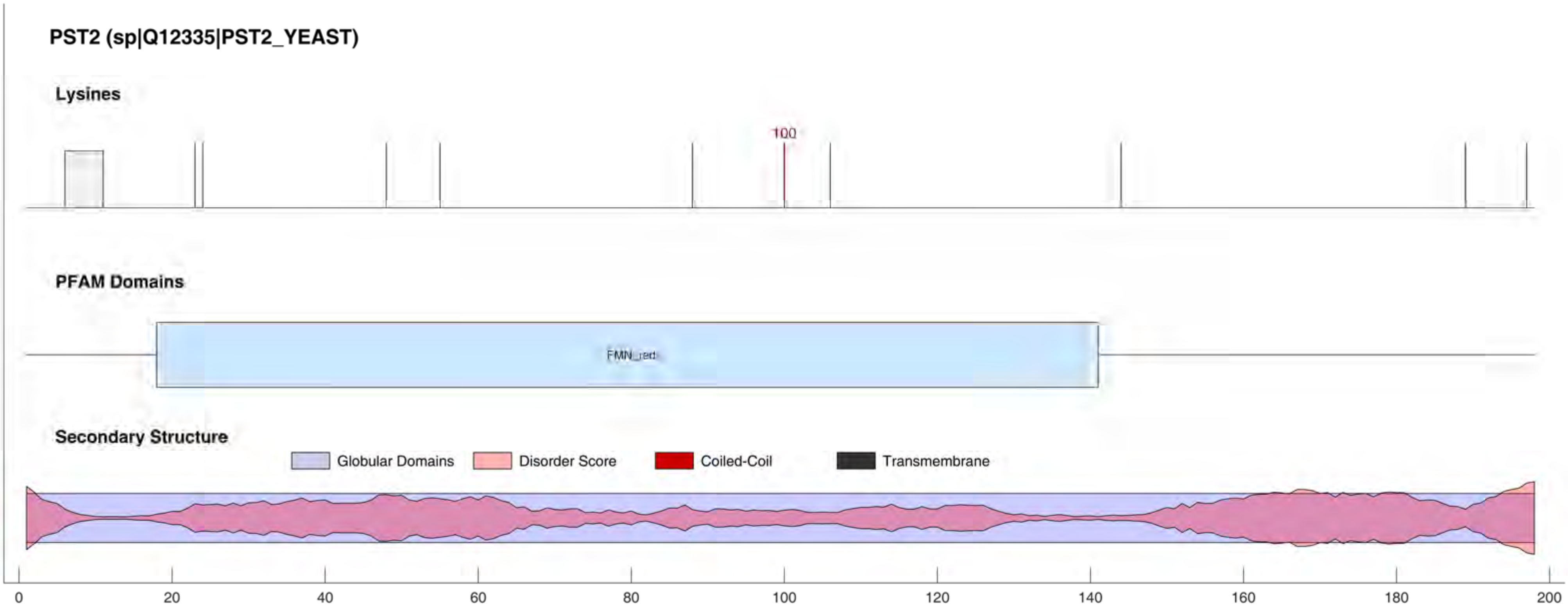

PST2 (sp|Q12335|PST2_YEAST)
Lysines
100
PFAM Domains
FMN_red
Secondary Structure
Globular Domains
Disorder Score
Coiled-Coil
Transmembrane
0
20
40
60
80
100
120
140
160
180
200

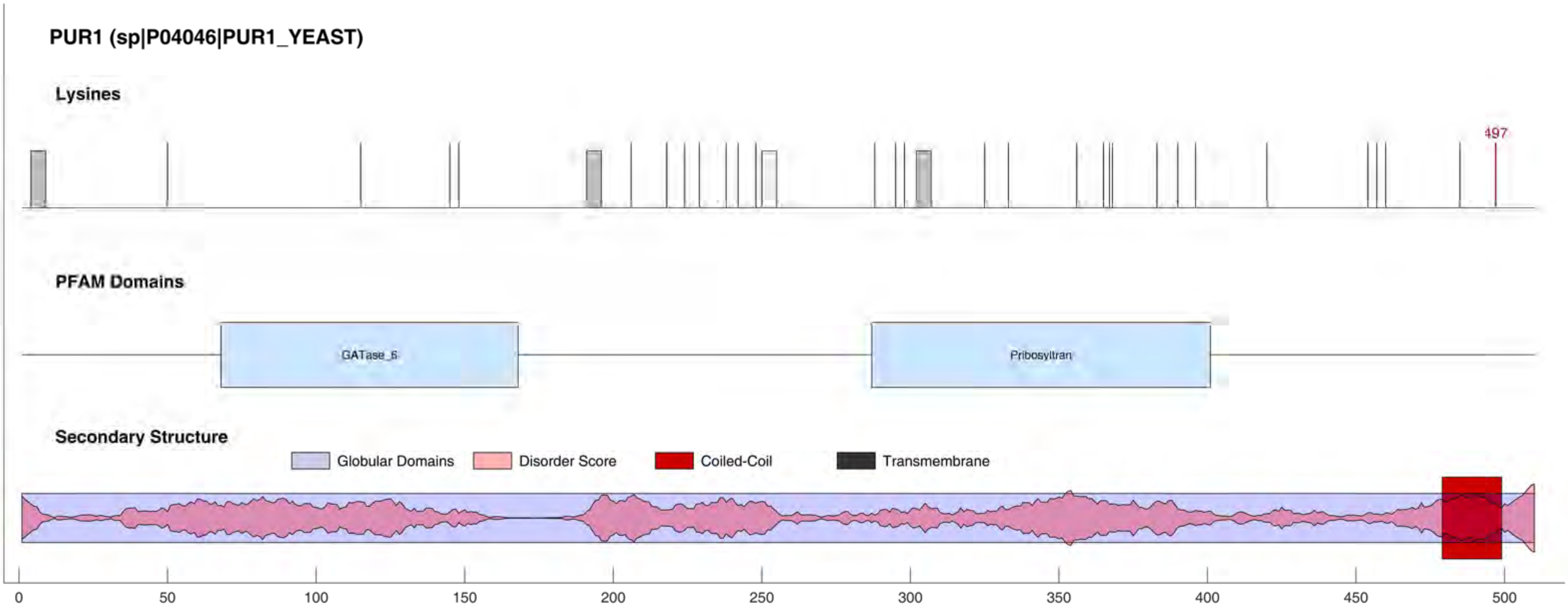
PUR1 (sp|P04046|PUR1_YEAST)
Lysines
497
PFAM Domains
GATase_6
Pribosyltran
Secondary Structure
Globular Domains
Disorder Score
Coiled-Coil
Transmembrane
0
50
100
150
200
250
300
350
400
450
500

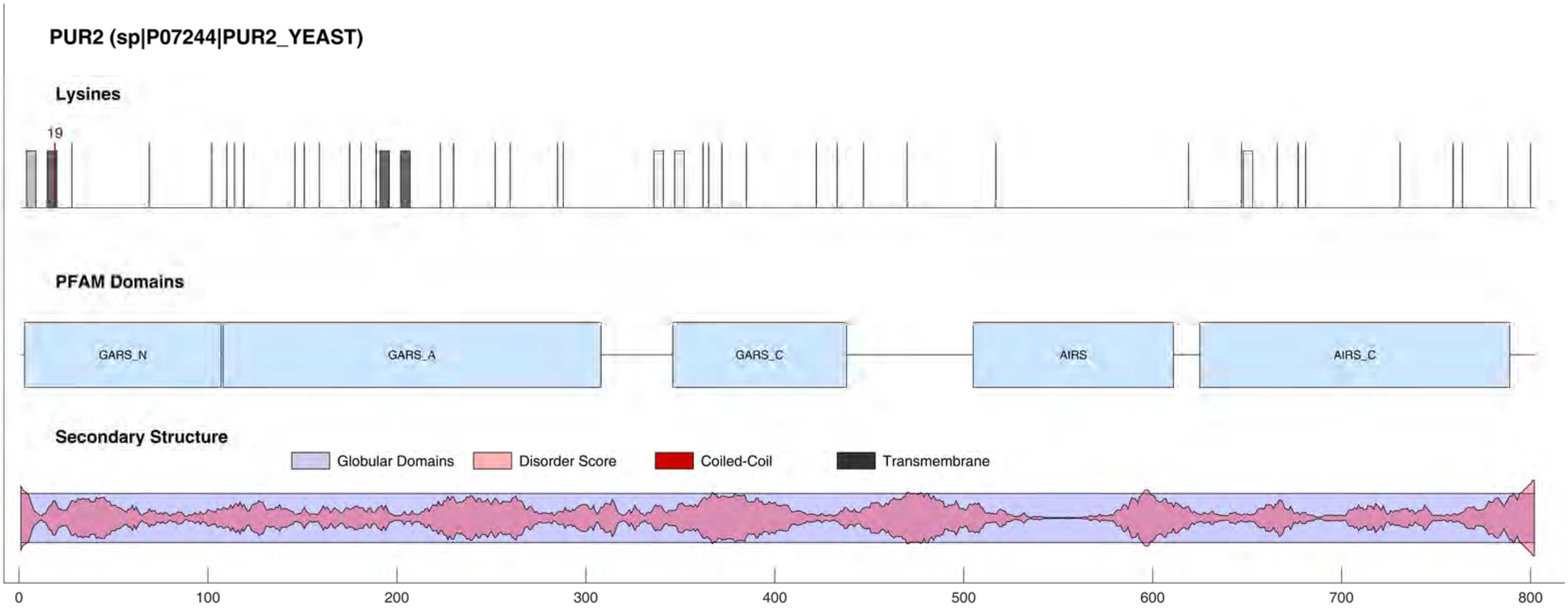

PUR2 (sp|P07244|PUR2_YEAST)
Lysines
19
PFAM Domains
GARS_N
GARS_A
GARS_C
AIRS
AIRS_C
Secondary Structure
Globular Domains
Disorder Score
Coiled-Coil
Transmembrane
0
100
200
300
400
500
600
700
800

#### PUR7 (sp|P27616|PUR7_YEAST)

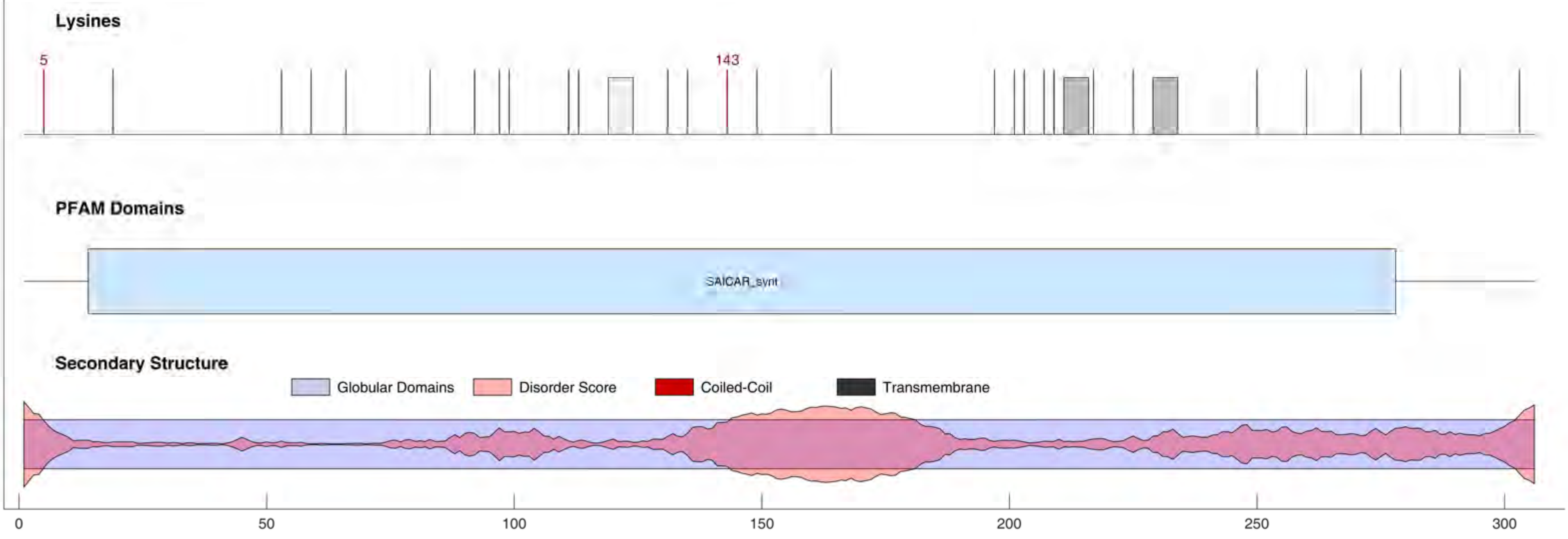

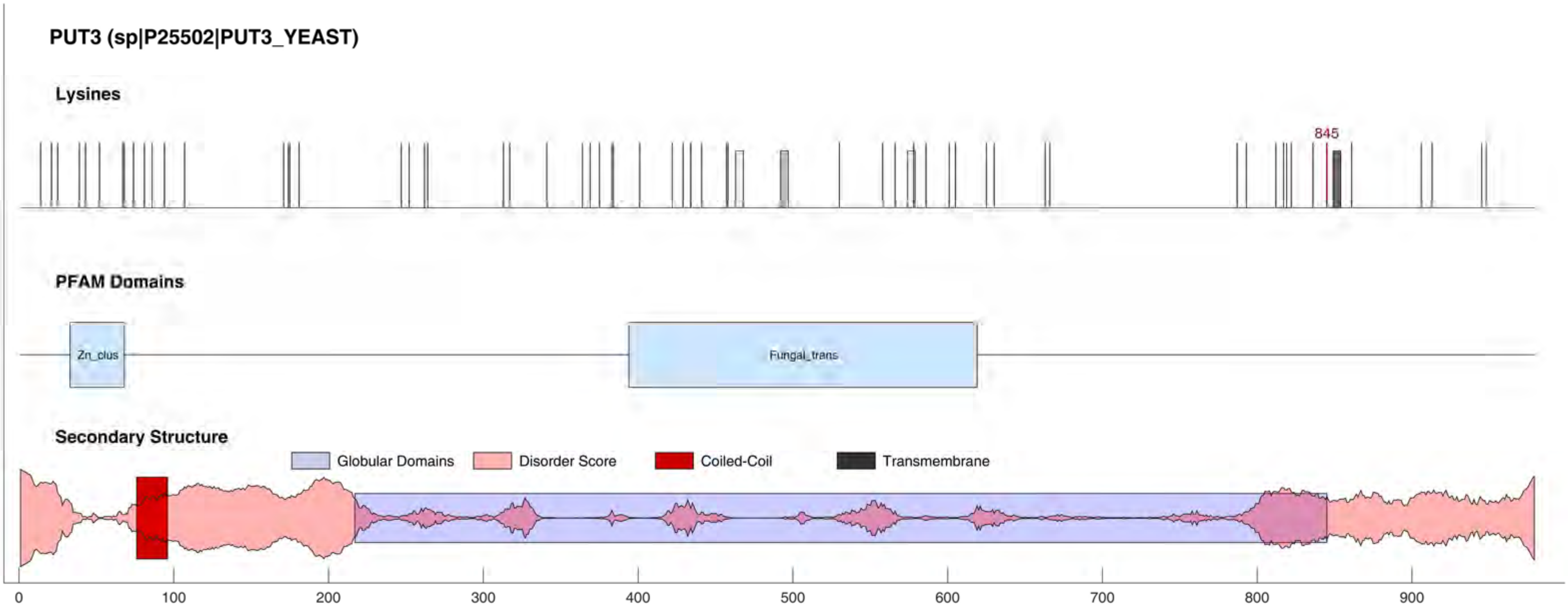

PUT3 (sp|P25502|PUT3_YEAST)
Lysines
845
PFAM Domains
Zn_clus
Fungal_trans
Secondary Structure
Globular Domains
Disorder Score
Coiled-Coil
Transmembrane
0
100
200
300
400
500
600
700
800
900

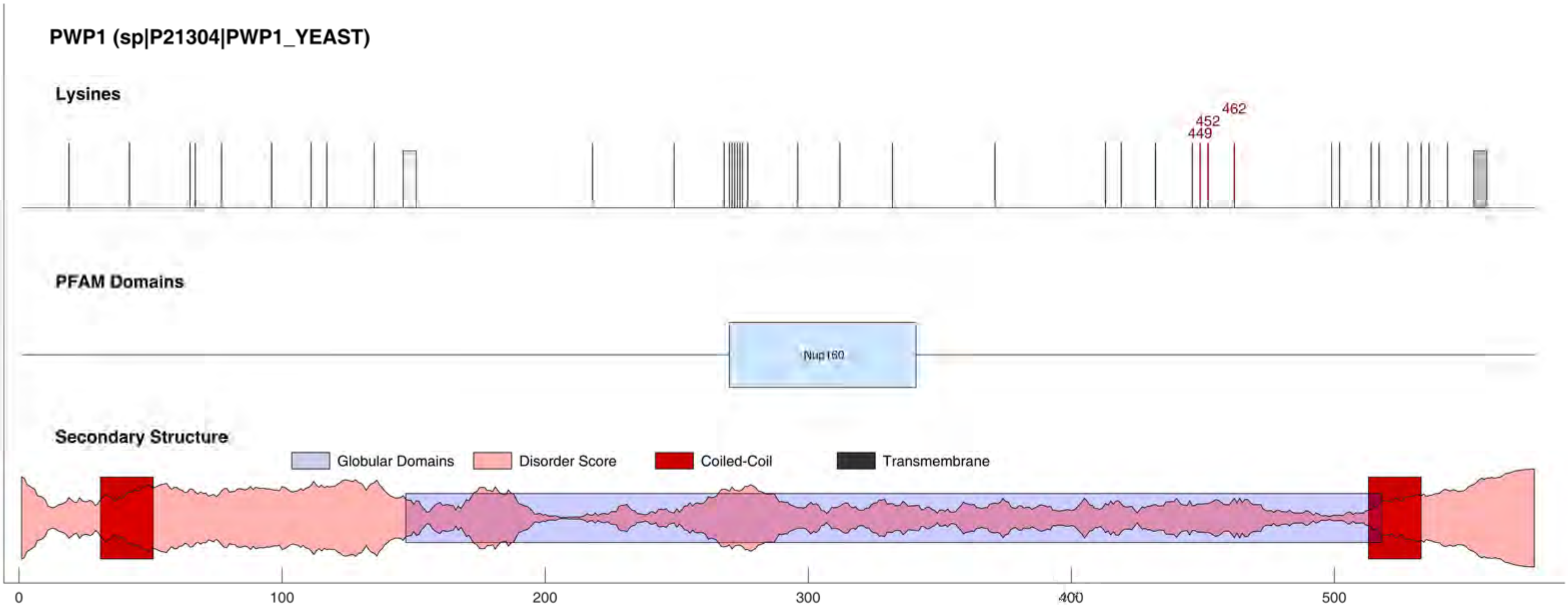
PWP1 (sp|P21304|PWP1_YEAST)
Lysines
462
452
449
PFAM Domains
Nup160
Secondary Structure
Globular Domains
Disorder Score
Coiled-Coil
Transmembrane
0
100
200
300
400
500

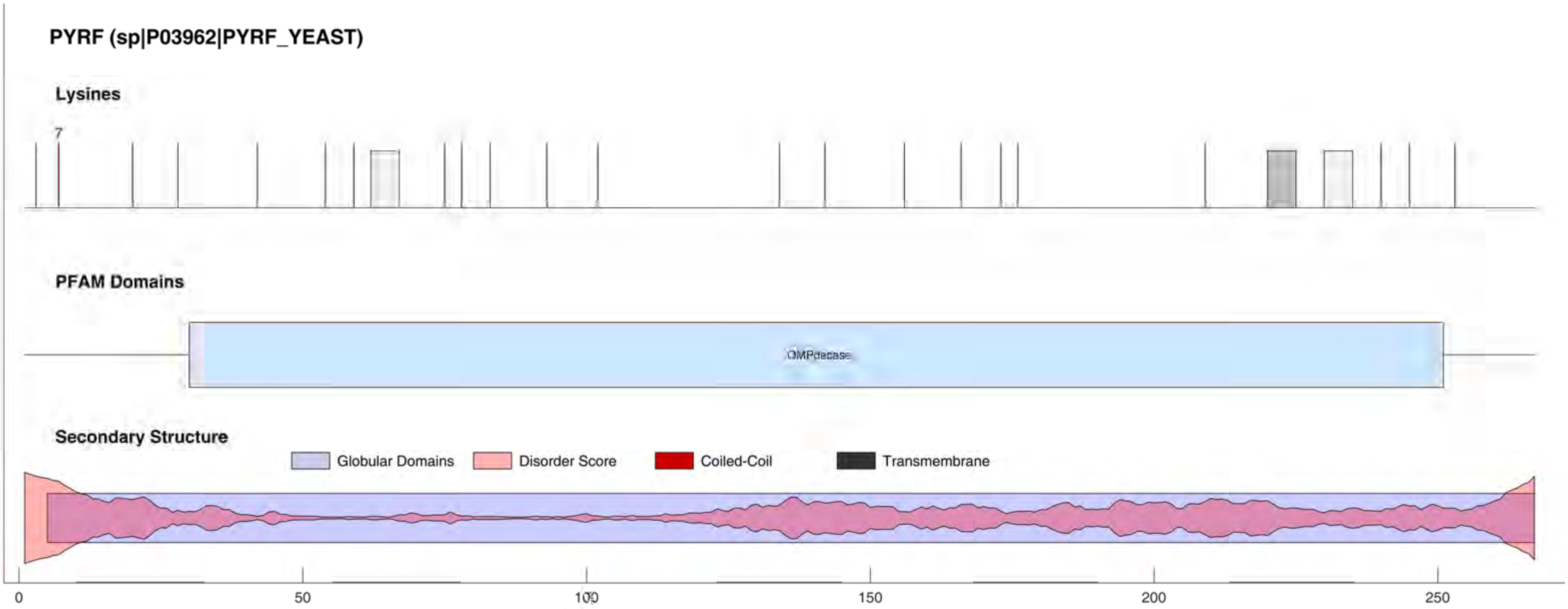
PYRF (sp|P03962|PYRF_YEAST)
Lysines
7
PFAM Domains
OMPdecase
Secondary Structure
Globular Domains
Disorder Score
Coiled-Coil
Transmembrane
0
50
100
150
200
250

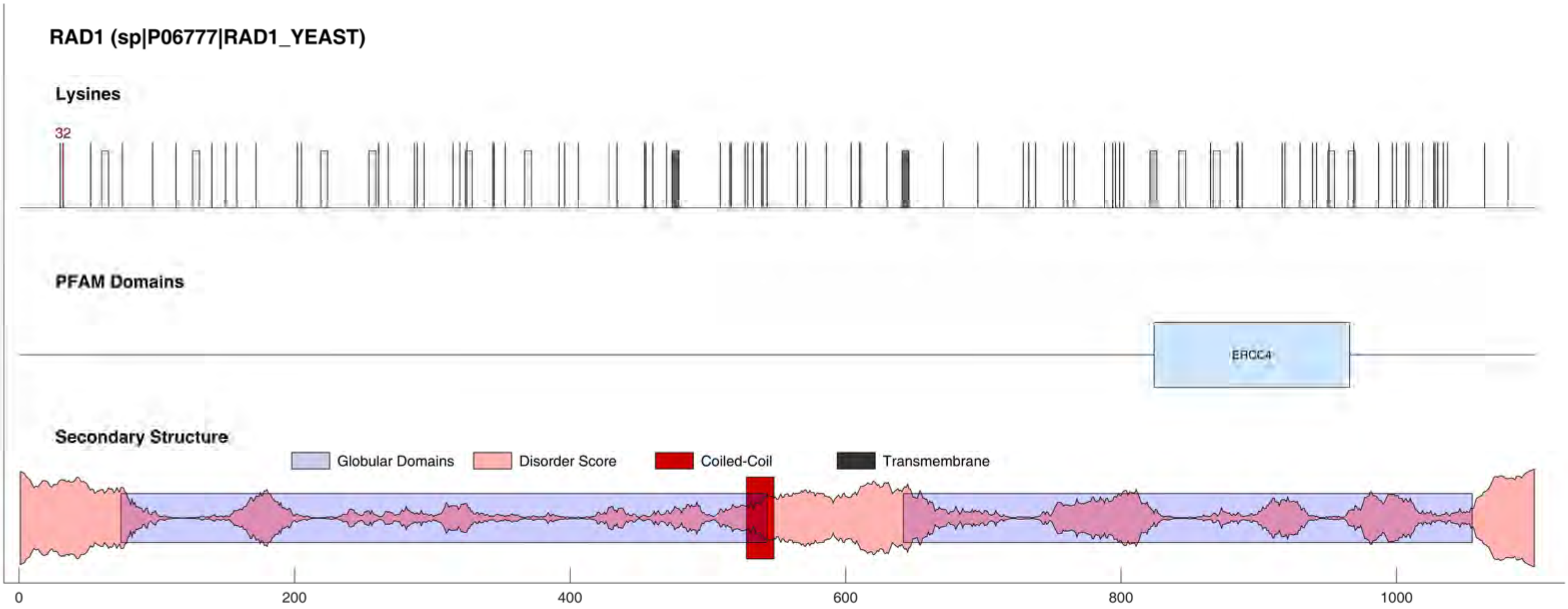
RAD1 (sp|P06777|RAD1_YEAST)
Lysines
32
PFAM Domains
ERCC4
Secondary Structure
Globular Domains
Disorder Score
Coiled-Coil
Transmembrane
0
200
400
600
800
1000

#### RAD5 (sp|P32849|RAD5_YEAST)

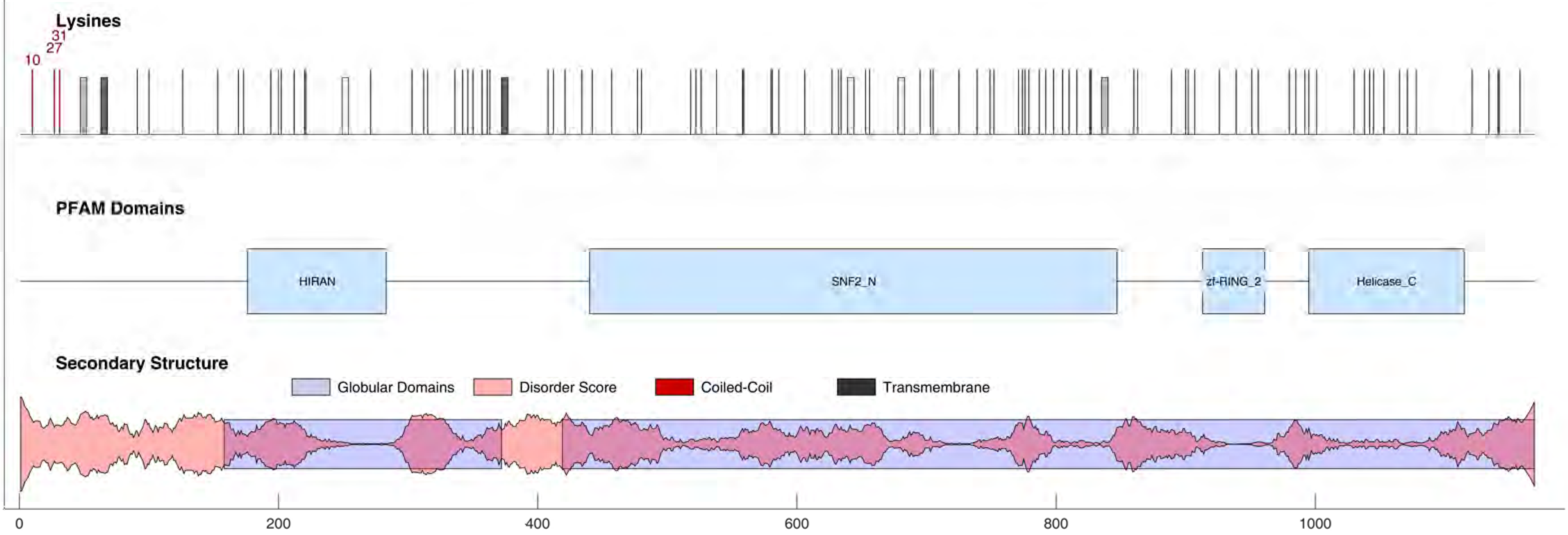

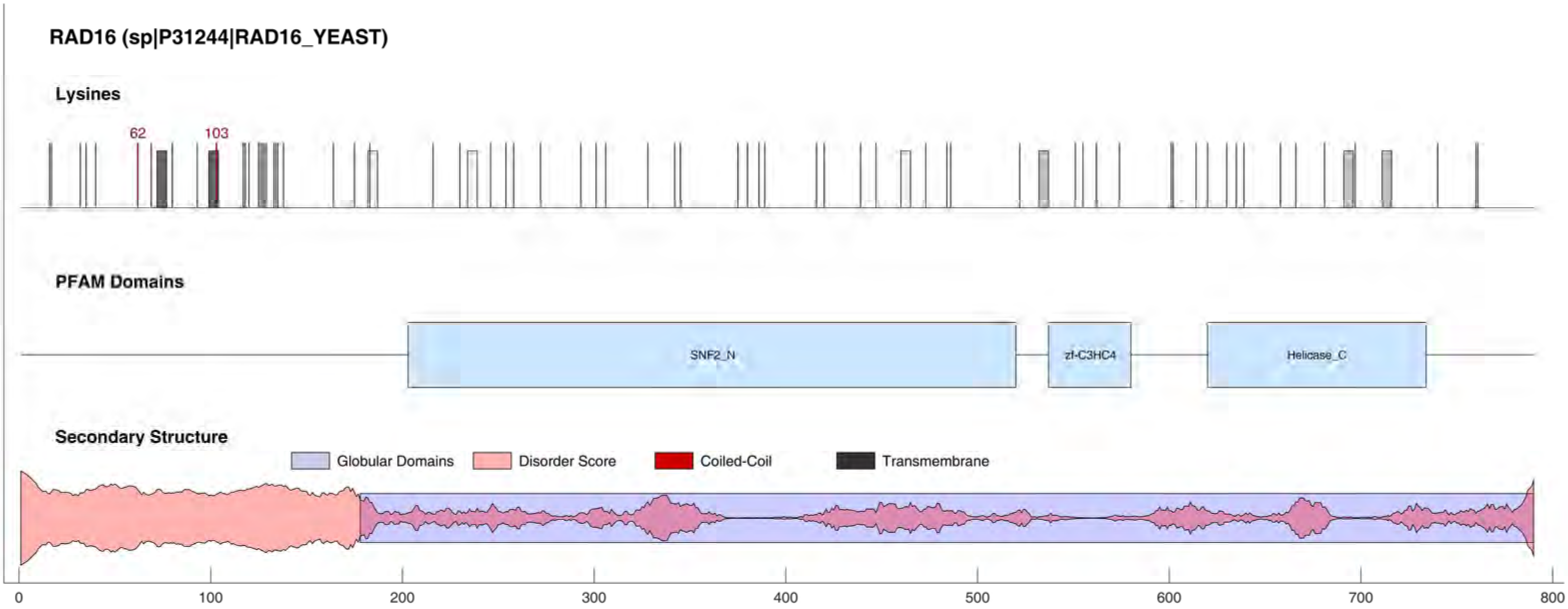

RAD16 (sp|P31244|RAD16_YEAST)
Lysines
62
103
PFAM Domains
SNF2_N
zf-C3HC4
Helicase_C
Secondary Structure
Globular Domains
Disorder Score
Coiled-Coil
Transmembrane
0
100
200
300
400
500
600
700
800

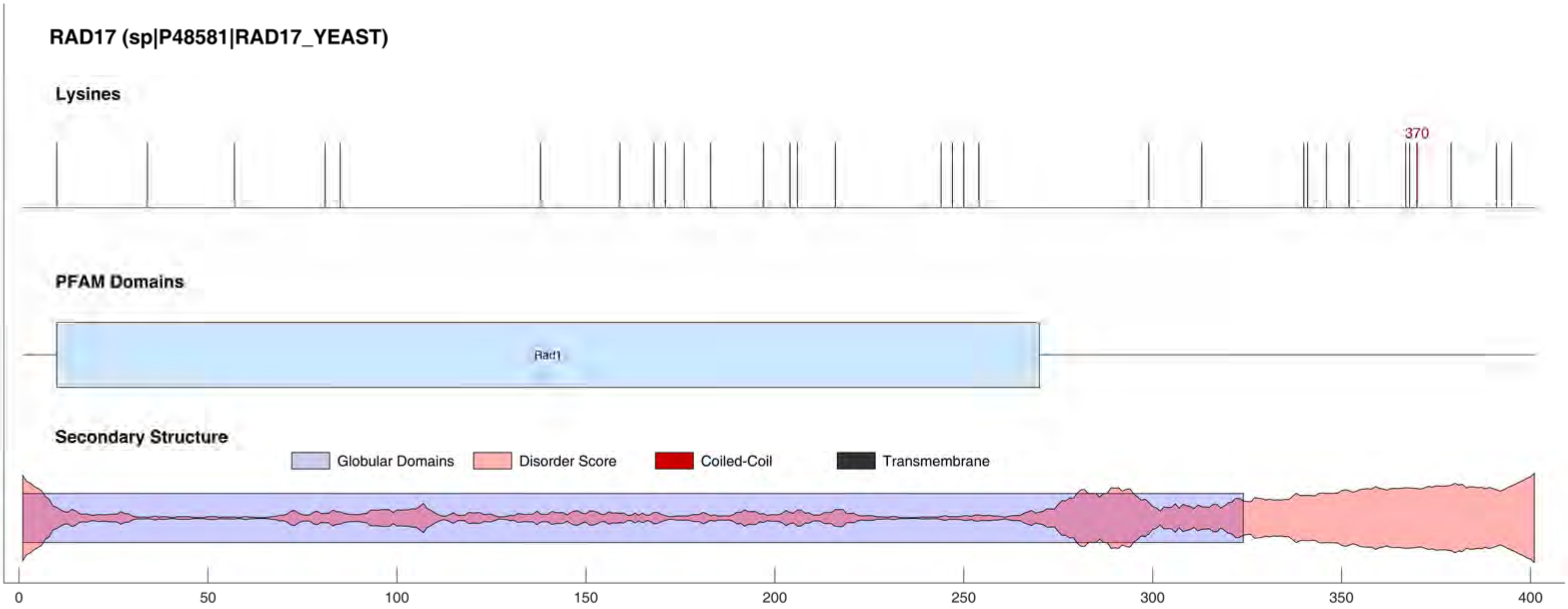
RAD17 (sp|P48581|RAD17_YEAST)
Lysines
370
PFAM Domains
Rad1
Secondary Structure
Globular Domains
Disorder Score
Coiled-Coil
Transmembrane
0
50
100
150
200
250
300
350
400

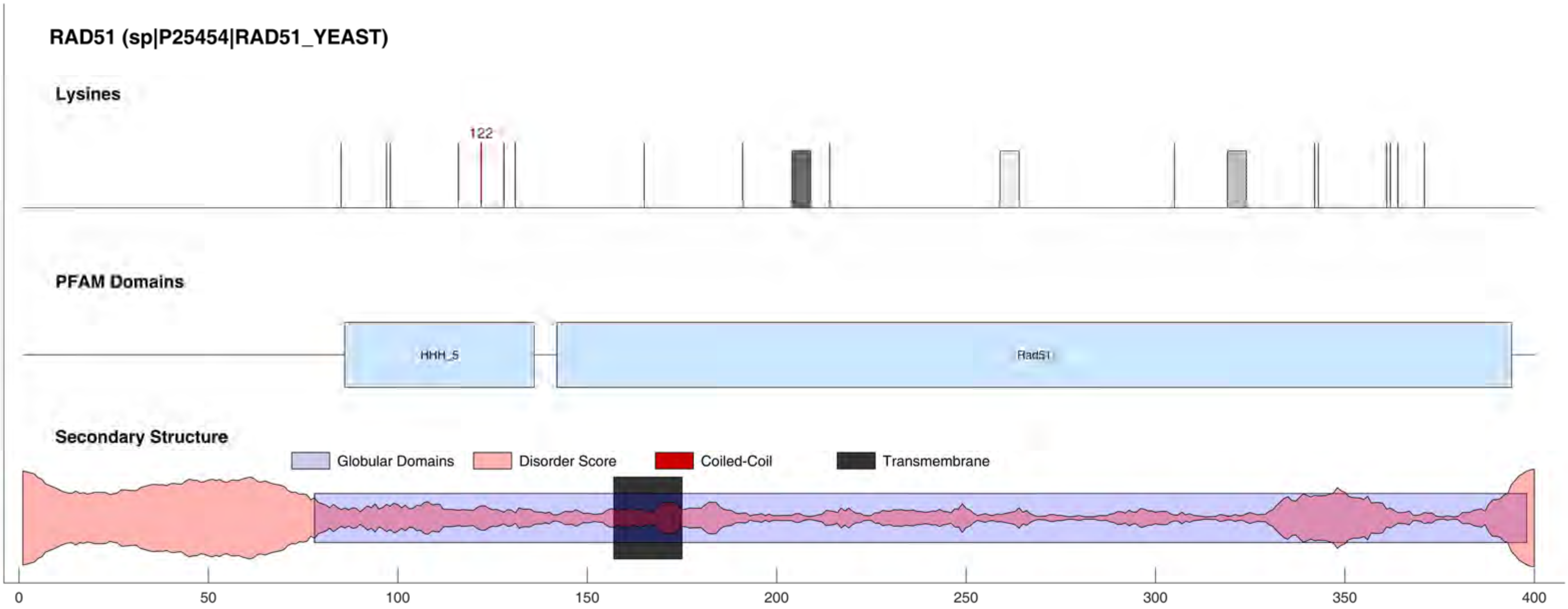
RAD51 (sp|P25454|RAD51_YEAST)
Lysines
122
PFAM Domains
HHH_5
Rad51
Secondary Structure
Globular Domains
Disorder Score
Coiled-Coil
Transmembrane
0
50
100
150
200
250
300
350
400

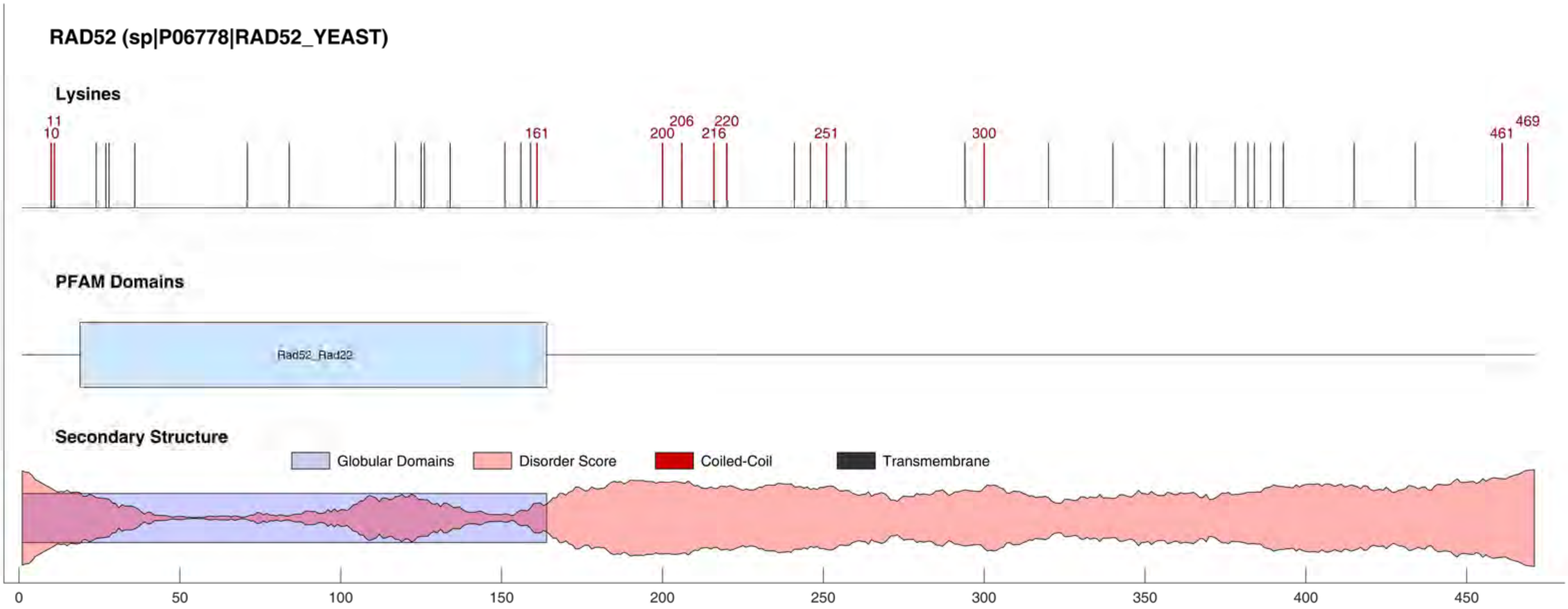
RAD52 (sp|P06778|RAD52_YEAST)
Lysines
11
10
161
200
206
216
220
251
300
461
469
PFAM Domains
Rad52_Rad22
Secondary Structure
Globular Domains
Disorder Score
Coiled-Coil
Transmembrane
0
50
100
150
200
250
300
350
400
450

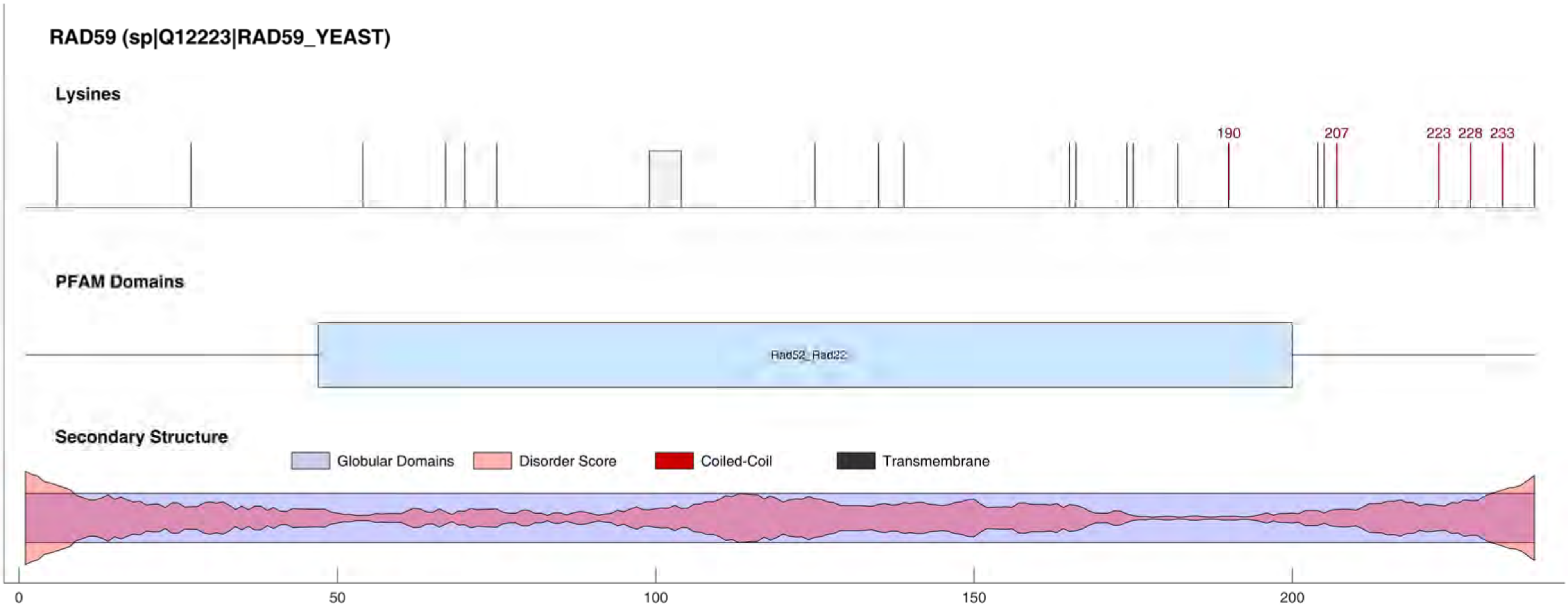

RAD59 (sp|Q12223|RAD59_YEAST)
Lysines
190
207
223
228
233
PFAM Domains
Rad52_Rad22
Secondary Structure
Globular Domains
Disorder Score
Coiled-Coil
Transmembrane
0
50
100
150
200

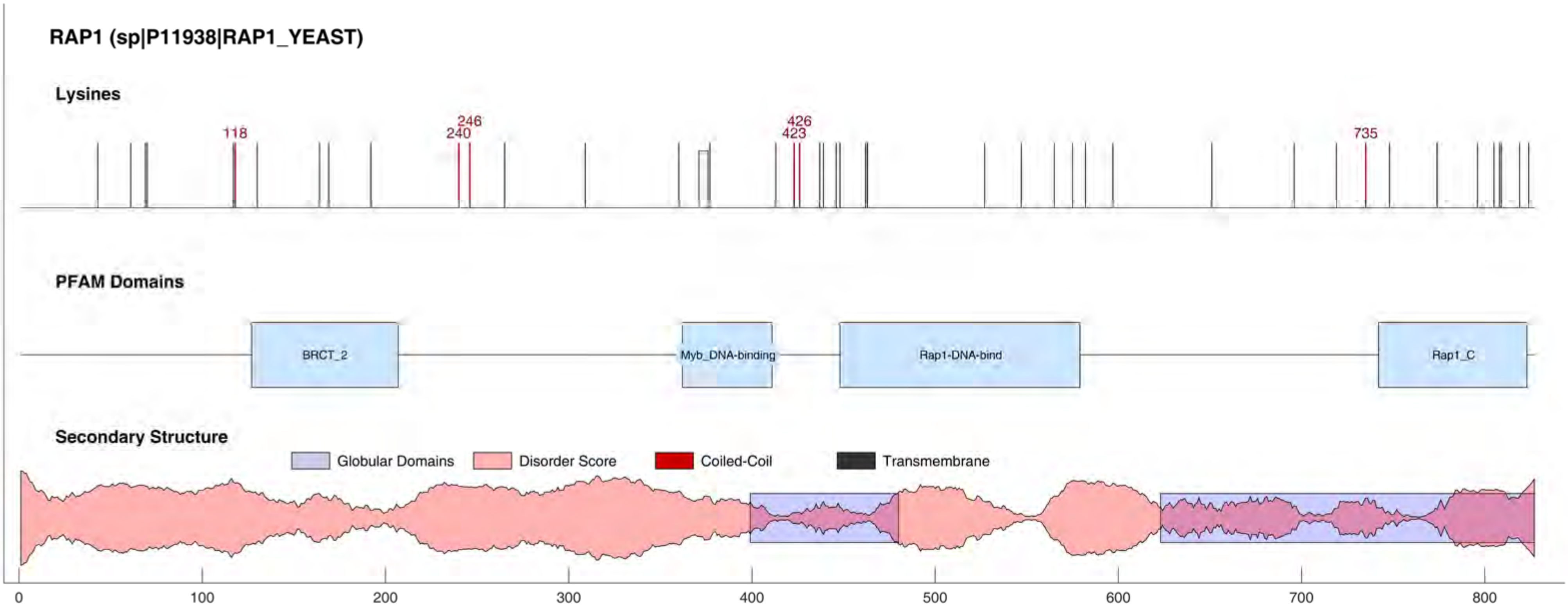
RAP1 (sp|P11938|RAP1_YEAST)
Lysines
118
240
246
423
426
735
PFAM Domains
BRCT_2
Myb_DNA-binding
Rap1-DNA-bind
Rap1_C
Secondary Structure
Globular Domains
Disorder Score
Coiled-Coil
Transmembrane
0
100
200
300
400
500
600
700
800

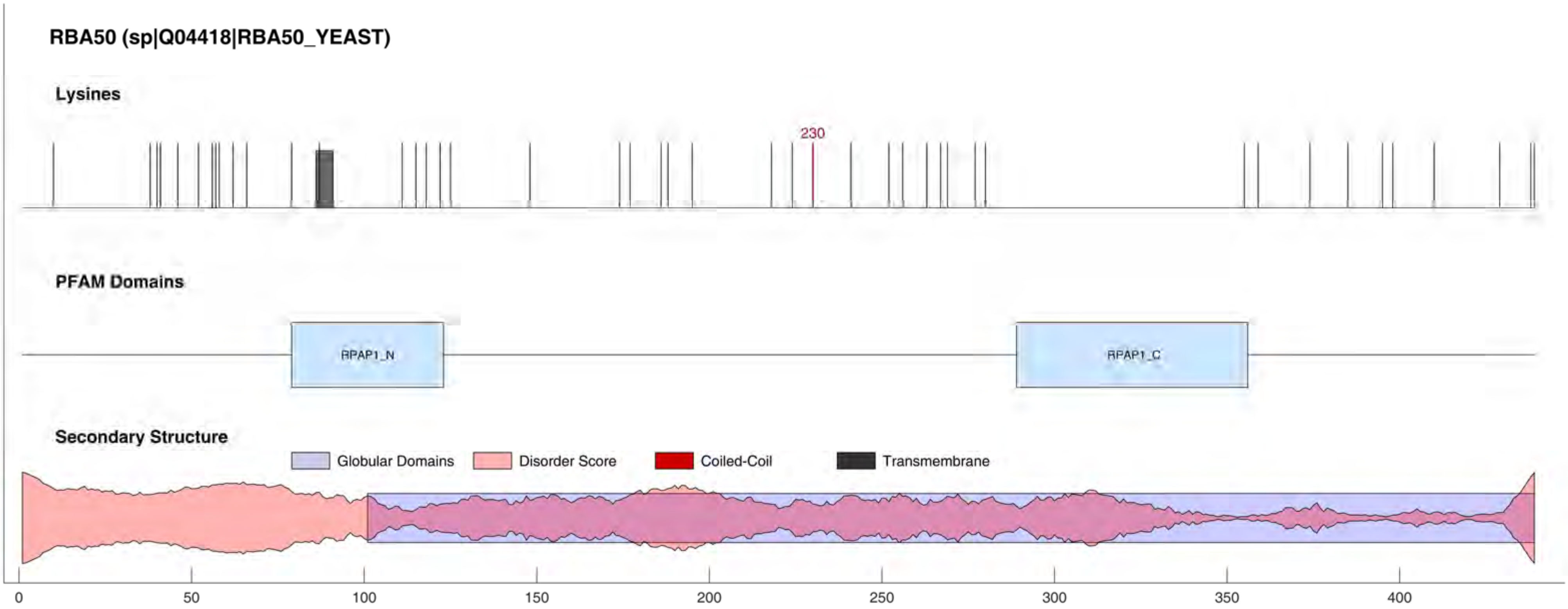
RBA50 (sp|Q04418|RBA50_YEAST)
Lysines
230
PFAM Domains
RPAP1_N
RPAP1_C
Secondary Structure
Globular Domains
Disorder Score
Coiled-Coil
Transmembrane
0
50
100
150
200
250
300
350
400

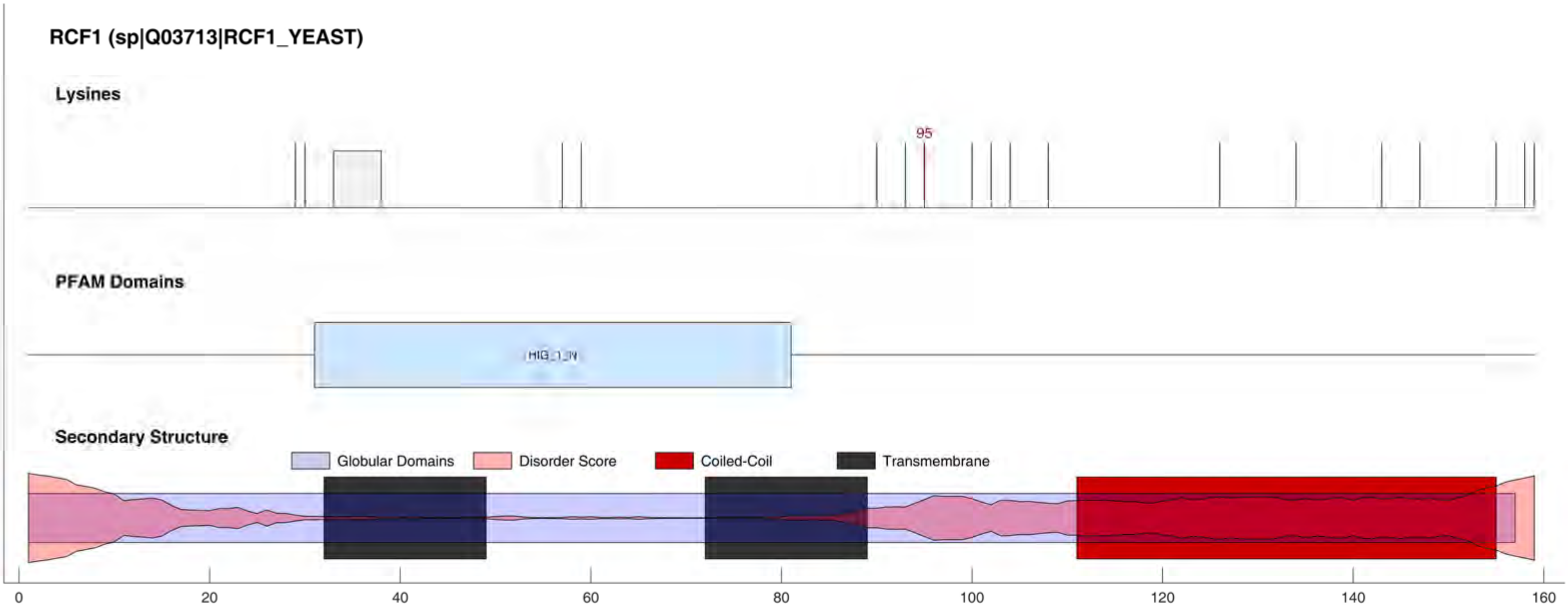

RCF1 (sp|Q03713|RCF1_YEAST)
Lysines
95
PFAM Domains
HIG_1_N
Secondary Structure
Globular Domains
Disorder Score
Coiled-Coil
Transmembrane
0
20
40
60
80
100
120
140
160

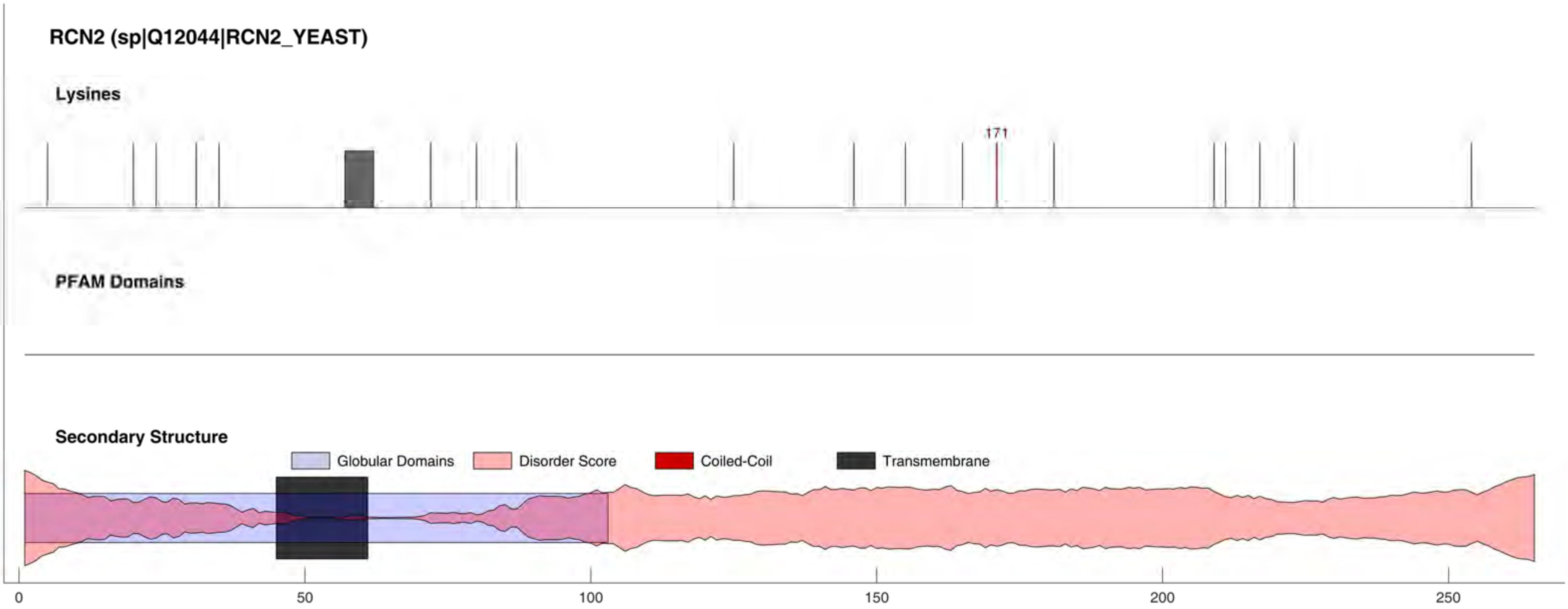

RCN2 (sp|Q12044|RCN2_YEAST)
Lysines
171
PFAM Domains
Secondary Structure
Globular Domains
Disorder Score
Coiled-Coil
Transmembrane
0
50
100
150
200
250

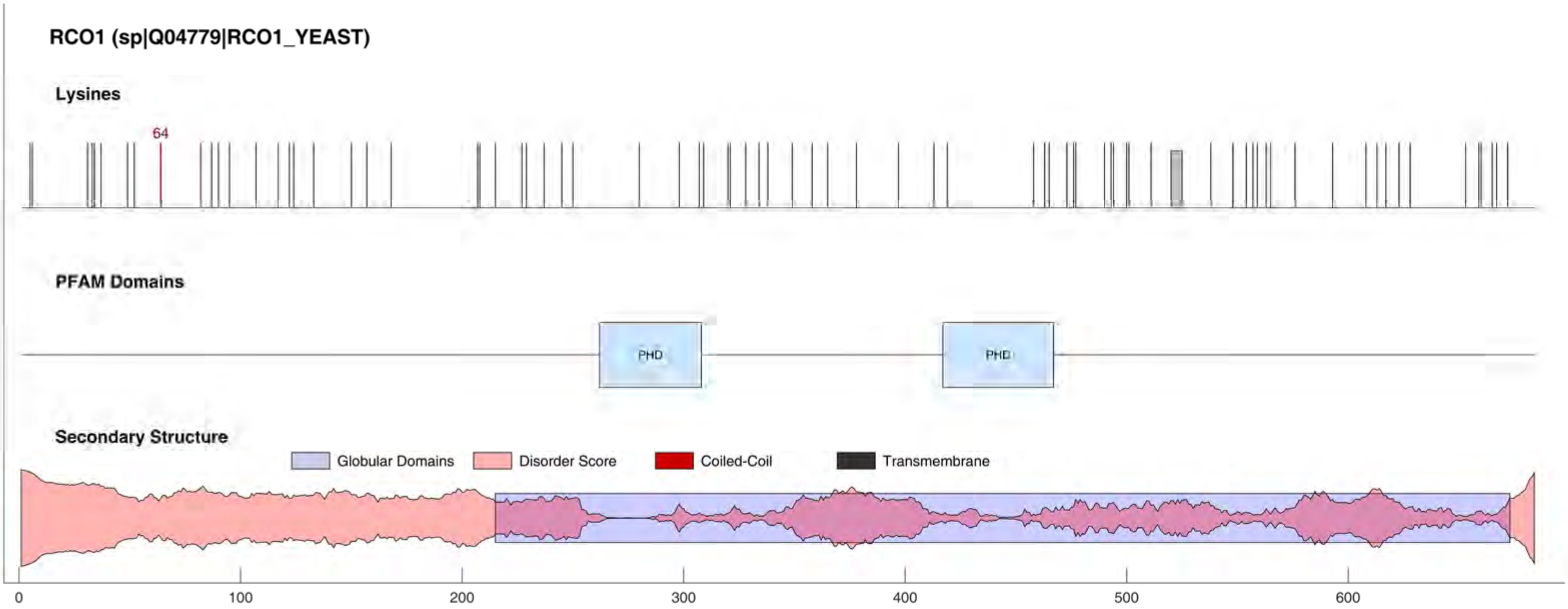

RCO1 (sp|Q04779|RCO1_YEAST)
Lysines
64
PFAM Domains
PHD
PHD
Secondary Structure
Globular Domains
Disorder Score
Coiled-Coil
Transmembrane
0
100
200
300
400
500
600

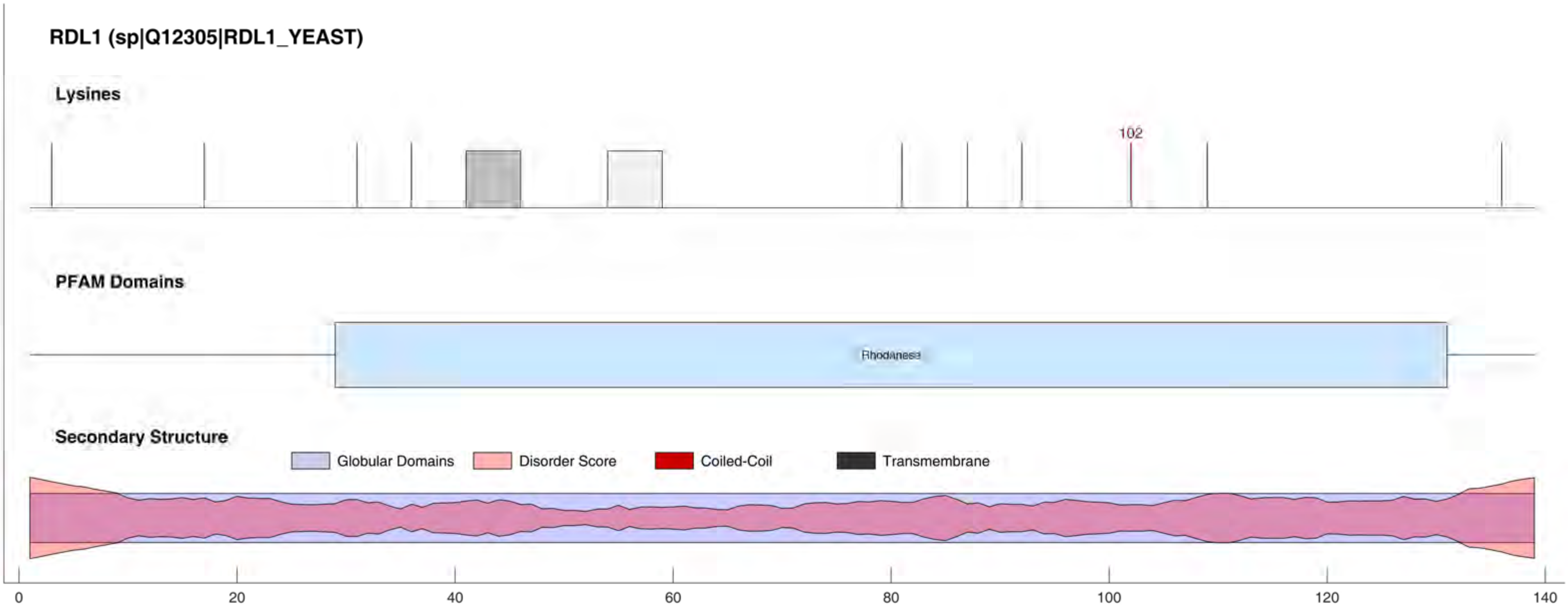
RDL1 (sp|Q12305|RDL1_YEAST)
Lysines
102
PFAM Domains
Rhodanese
Secondary Structure
Globular Domains
Disorder Score
Coiled-Coil
Transmembrane
0
20
40
60
80
100
120
140

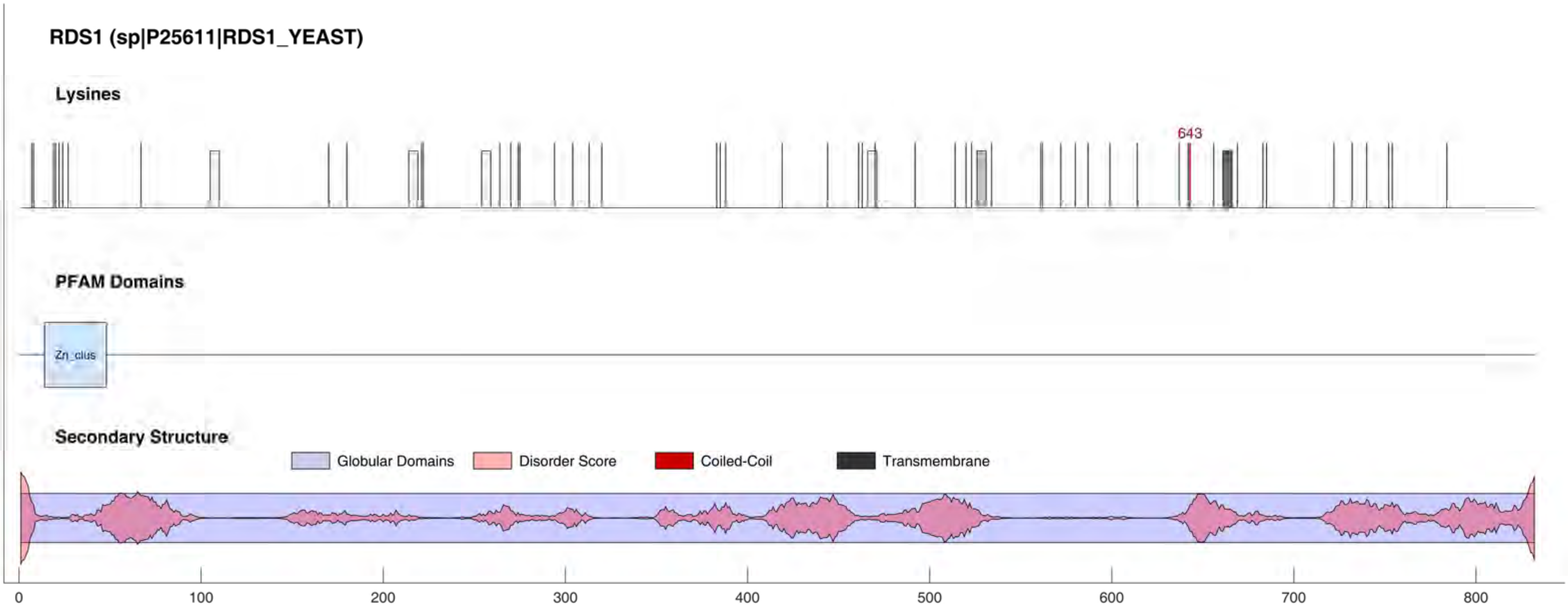
RDS1 (sp|P25611|RDS1_YEAST)
Lysines
643
PFAM Domains
Zn_clus
Secondary Structure
Globular Domains
Disorder Score
Coiled-Coil
Transmembrane
0
100
200
300
400
500
600
700
800

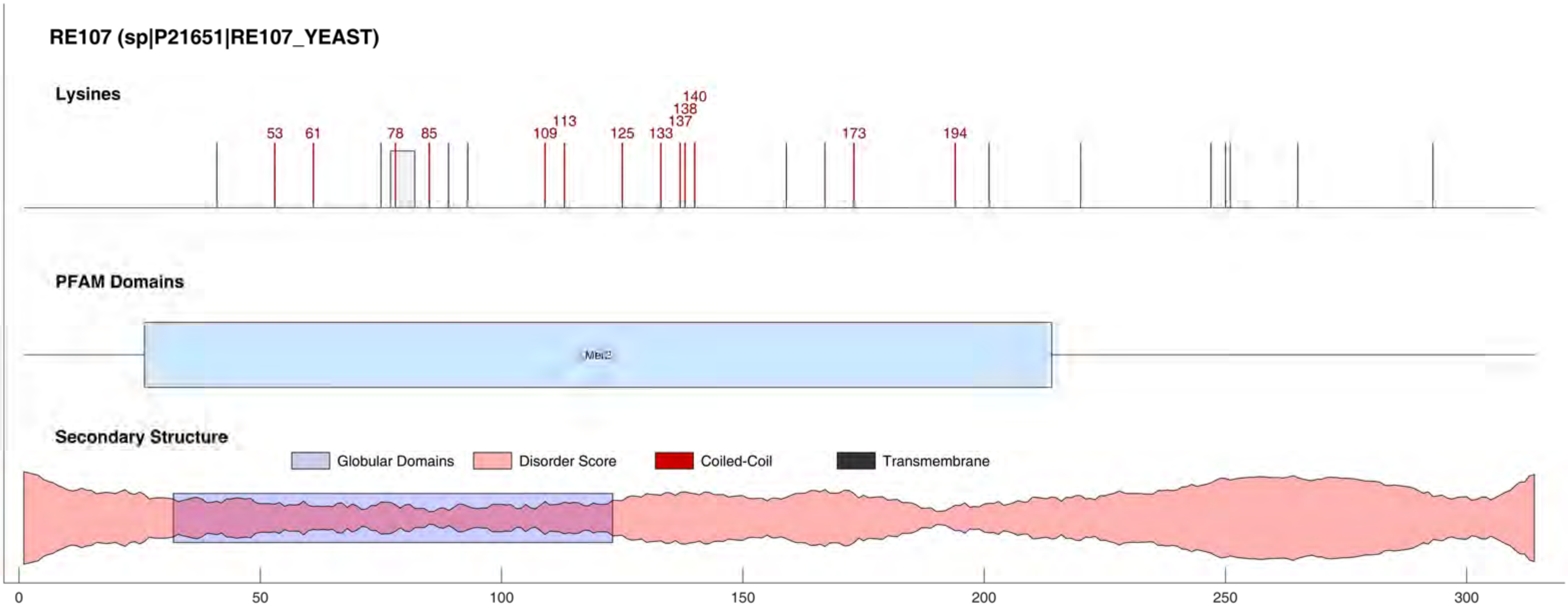
RE107 (sp|P21651|RE107_YEAST)
Lysines
53
61
78
85
109
113
125
133
137
138
140
173
194
PFAM Domains
Secondary Structure
Globular Domains
Disorder Score
Coiled-Coil
Transmembrane
0
50
100
150
200
250
300

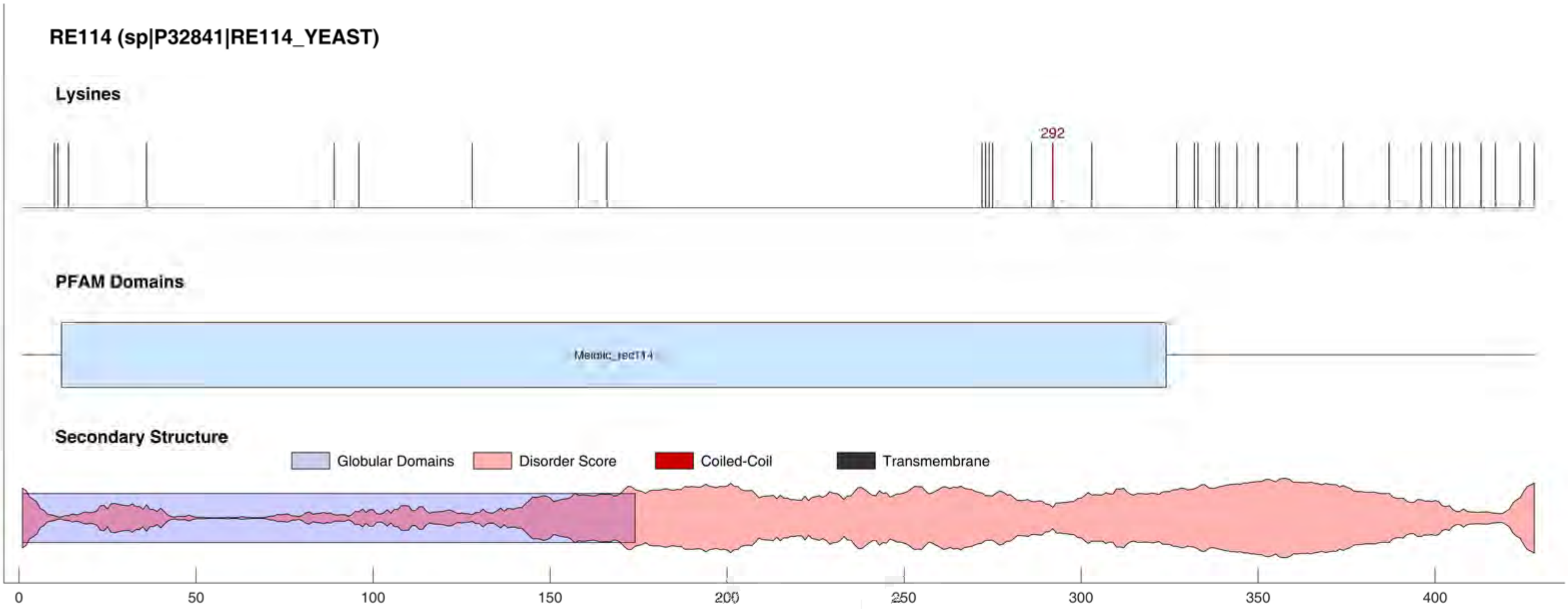

RE114 (sp|P32841|RE114_YEAST)
Lysines
292
PFAM Domains
Secondary Structure
Globular Domains
Disorder Score
Coiled-Coil
Transmembrane
0
50
100
150
200
250
300
350
400

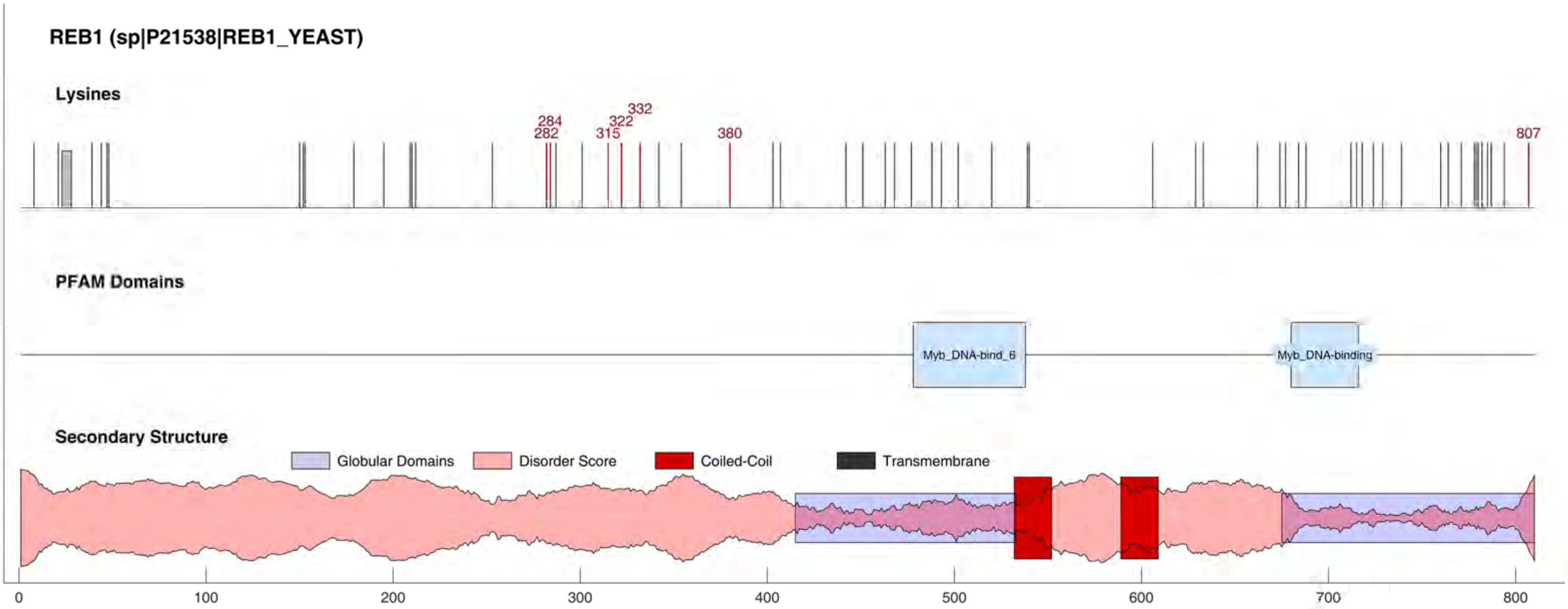
REB1 (sp|P21538|REB1_YEAST)
Lysines
284
282
332
322
315
380
807
PFAM Domains
Myb_DNA-bind_6
Myb_DNA-binding
Secondary Structure
Globular Domains
Disorder Score
Coiled-Coil
Transmembrane
0
100
200
300
400
500
600
700
800

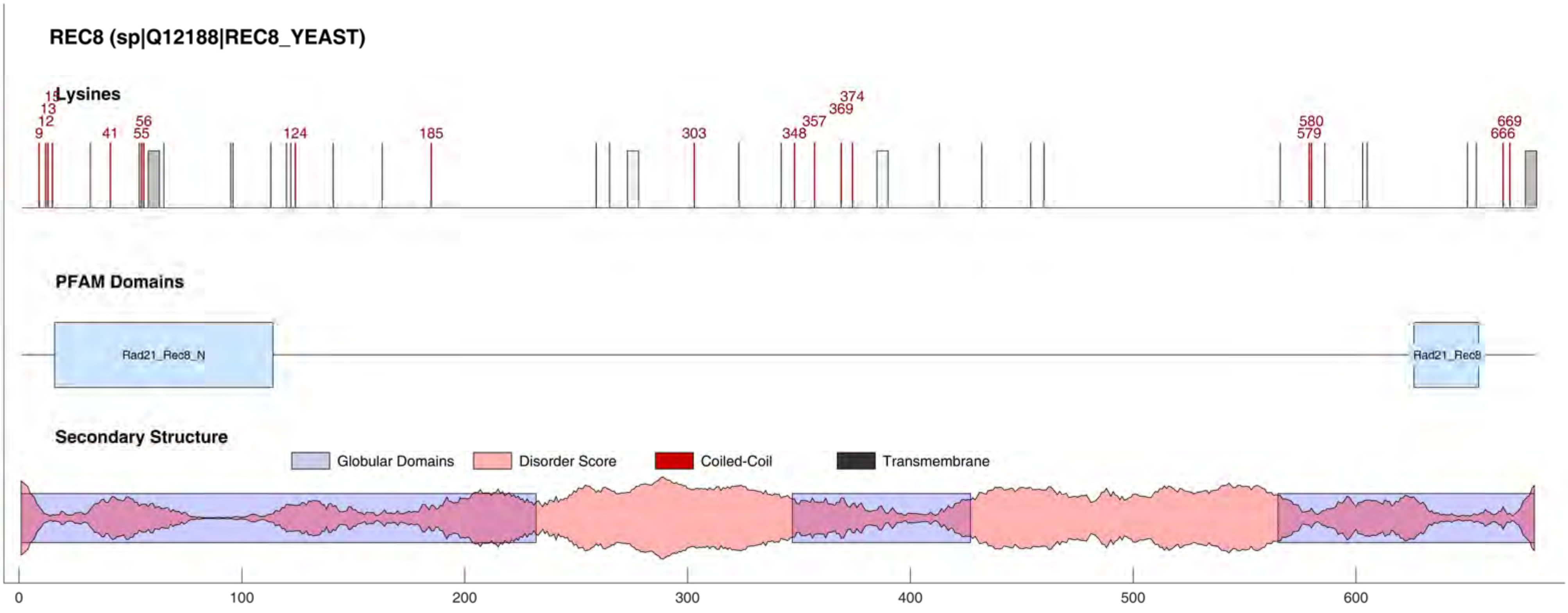
REC8 (sp|Q12188|REC8_YEAST)
Lysines
9
12
13
15
41
55
56
124
185
303
348
357
369
374
579
580
666
669
PFAM Domains
Rad21_Rec8_N
Rad21_Rec8
Secondary Structure
Globular Domains
Disorder Score
Coiled-Coil
Transmembrane
0
100
200
300
400
500
600

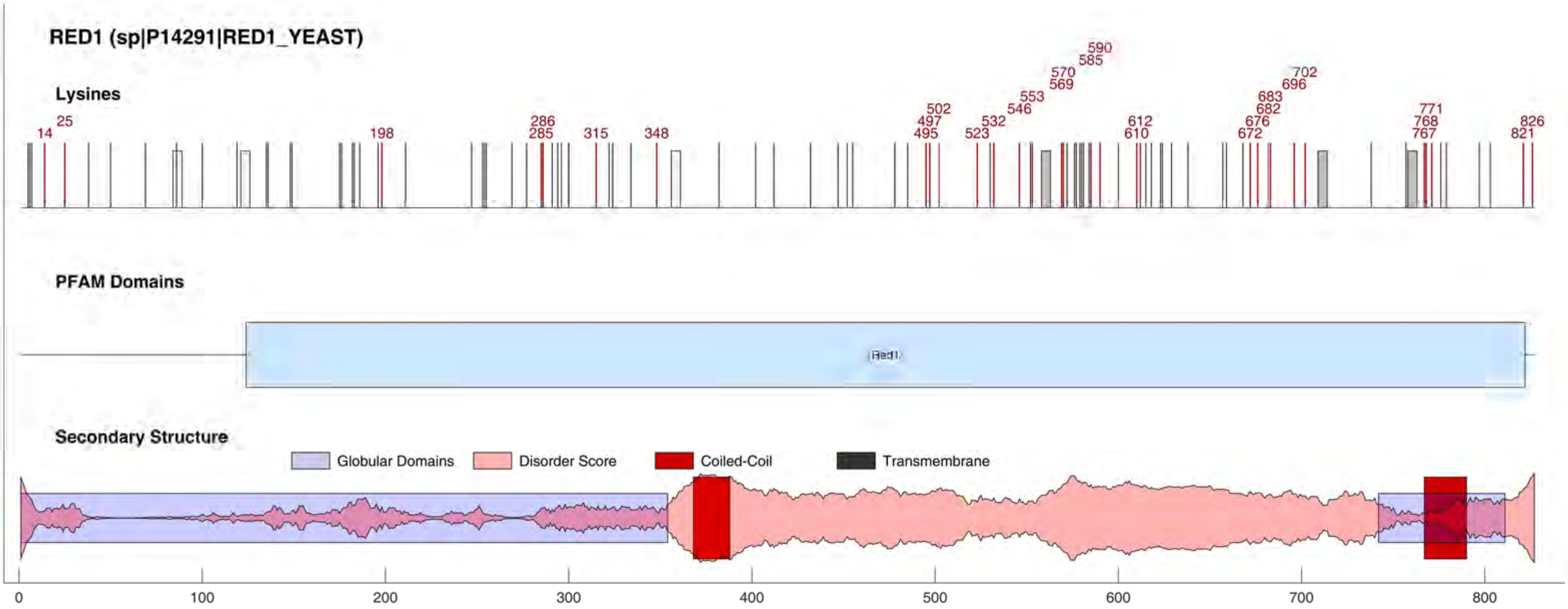
RED1 (sp|P14291|RED1_YEAST)
Lysines
14
25
198
285
286
315
348
495
497
502
523
532
546
553
569
570
585
590
610
612
672
676
682
683
696
702
767
768
771
821
826
PFAM Domains
Red1
Secondary Structure
Globular Domains
Disorder Score
Coiled-Coil
Transmembrane
0
100
200
300
400
500
600
700
800

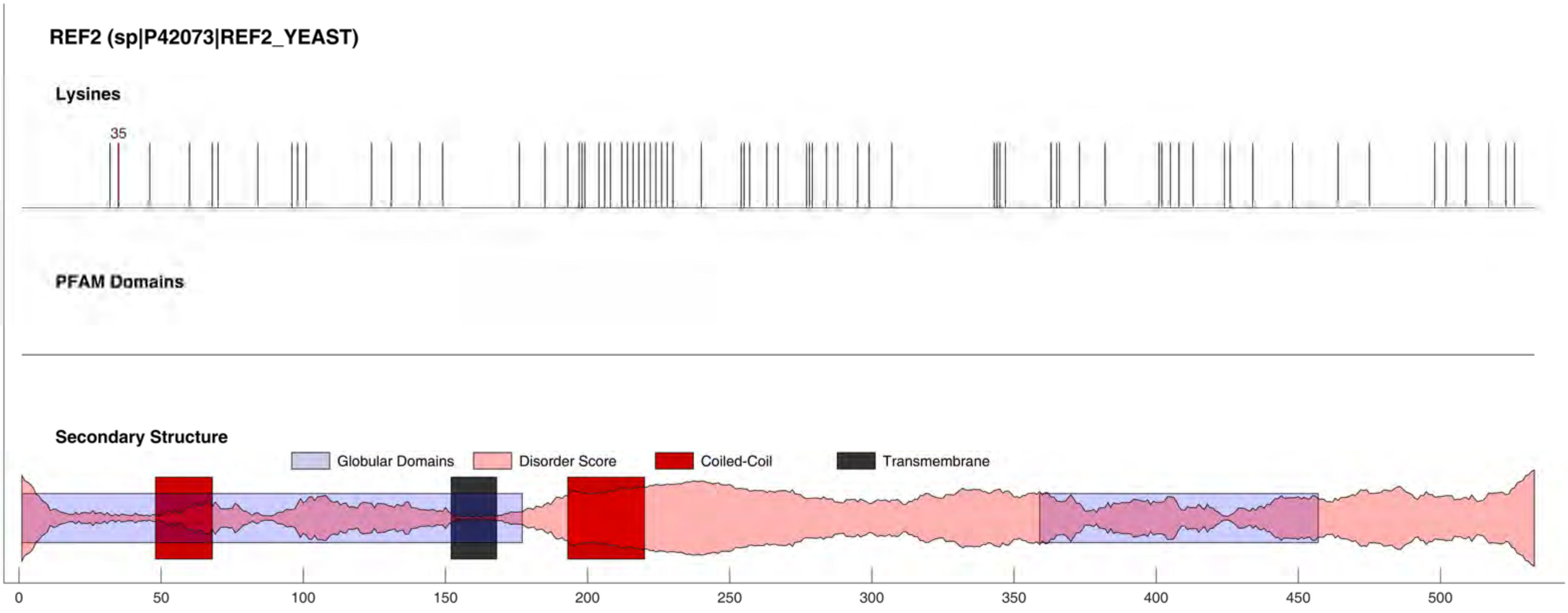

REF2 (sp|P42073|REF2_YEAST)
Lysines
35
PFAM Domains
Secondary Structure
Globular Domains
Disorder Score
Coiled-Coil
Transmembrane
0
50
100
150
200
250
300
350
400
450
500

#### REP1 (sp|P03871|REP1_YEAST)

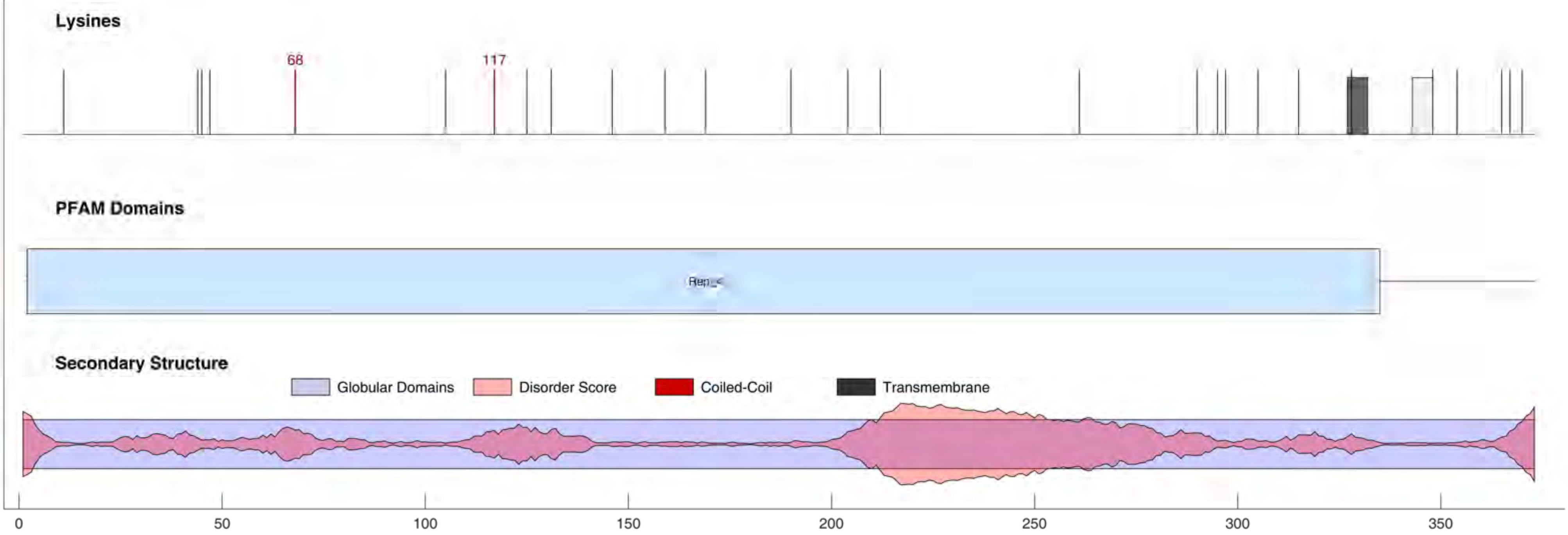

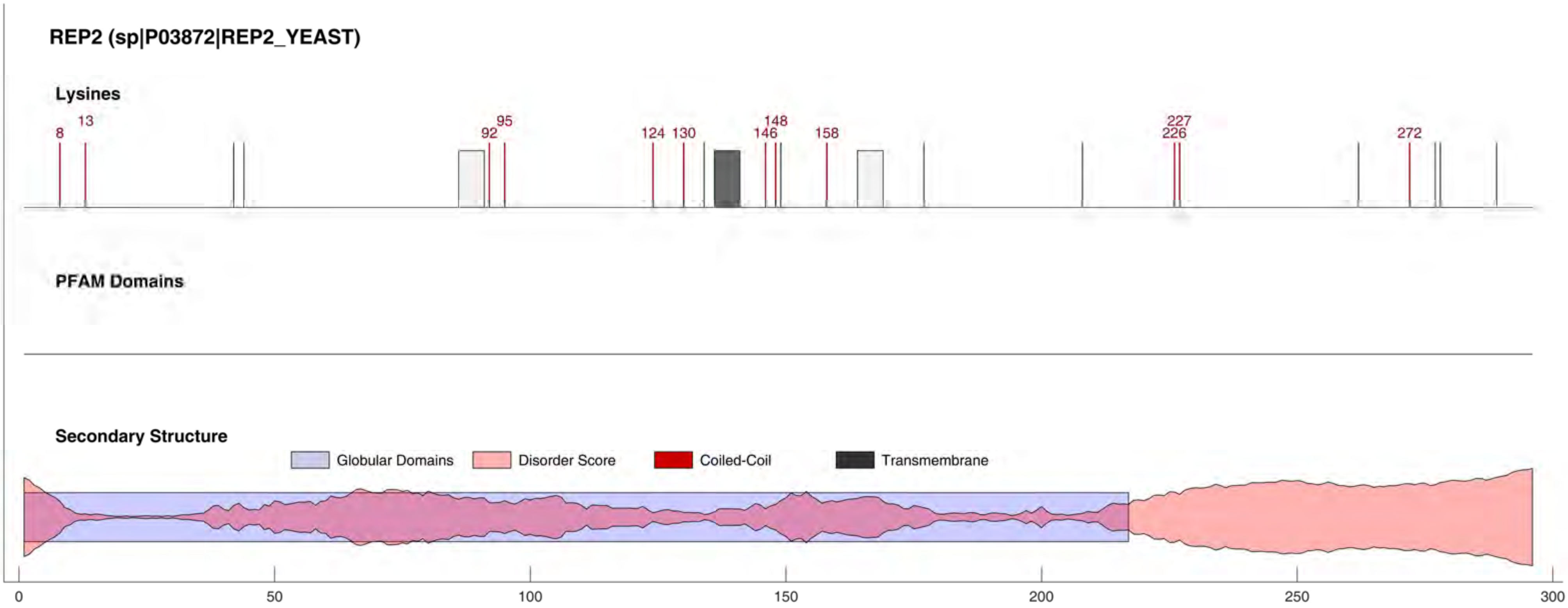
REP2 (sp|P03872|REP2_YEAST)
Lysines
8
13
92
95
124
130
146
148
158
226
227
272
PFAM Domains
Secondary Structure
Globular Domains
Disorder Score
Coiled-Coil
Transmembrane
0
50
100
150
200
250
300

#### RFA1 (sp|P22336|RFA1_YEAST)

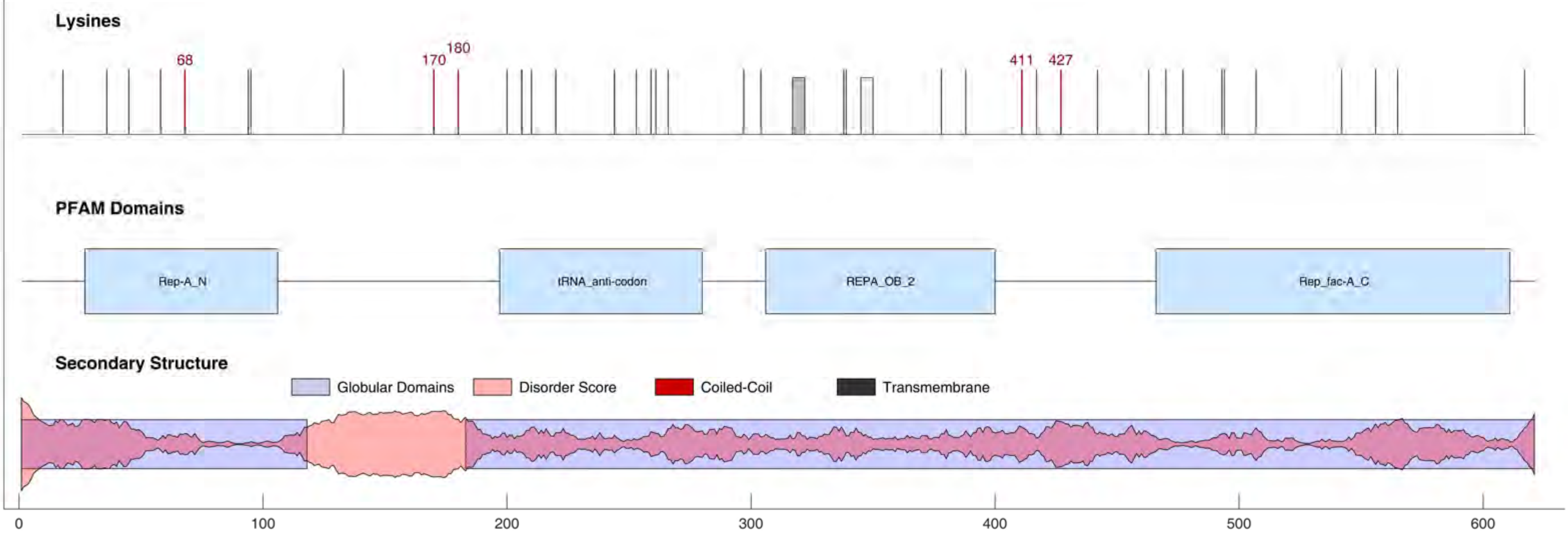

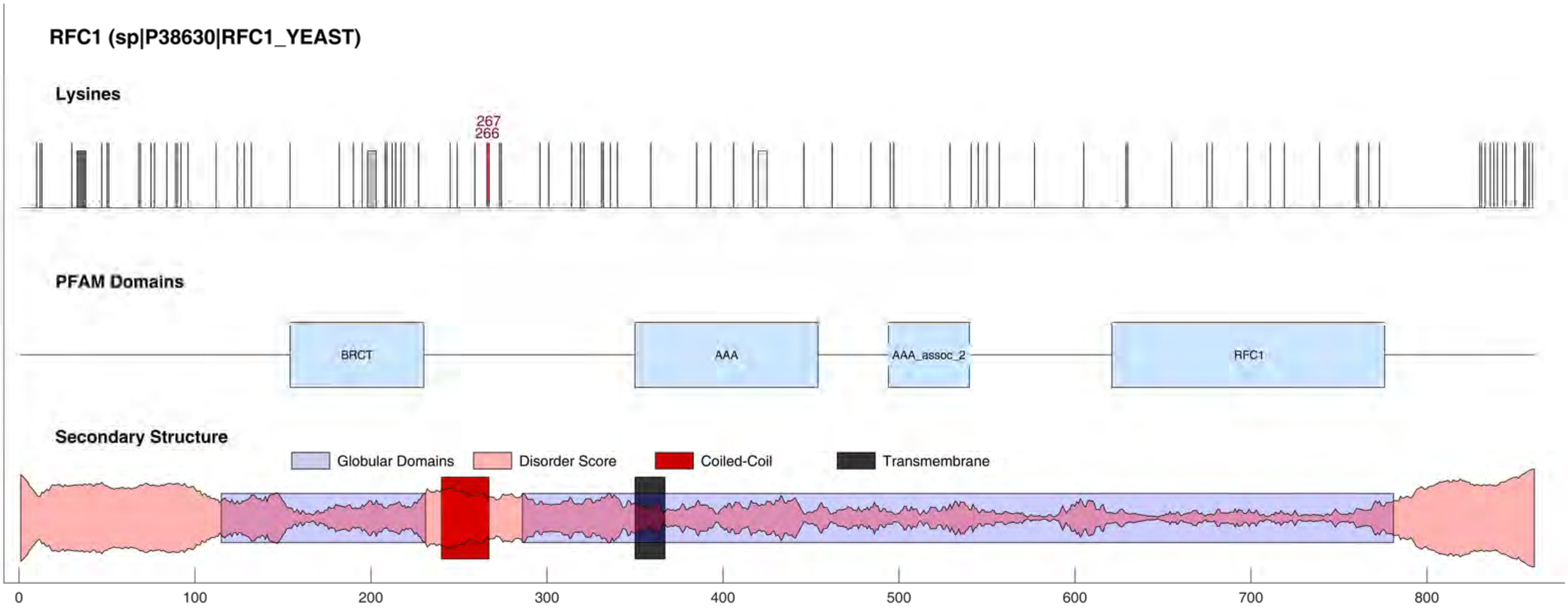
RFC1 (sp|P38630|RFC1_YEAST)
Lysines
267
266
PFAM Domains
BRCT
AAA
AAA_assoc_2
RFC1
Secondary Structure
Globular Domains
Disorder Score
Coiled-Coil
Transmembrane
0
100
200
300
400
500
600
700
800

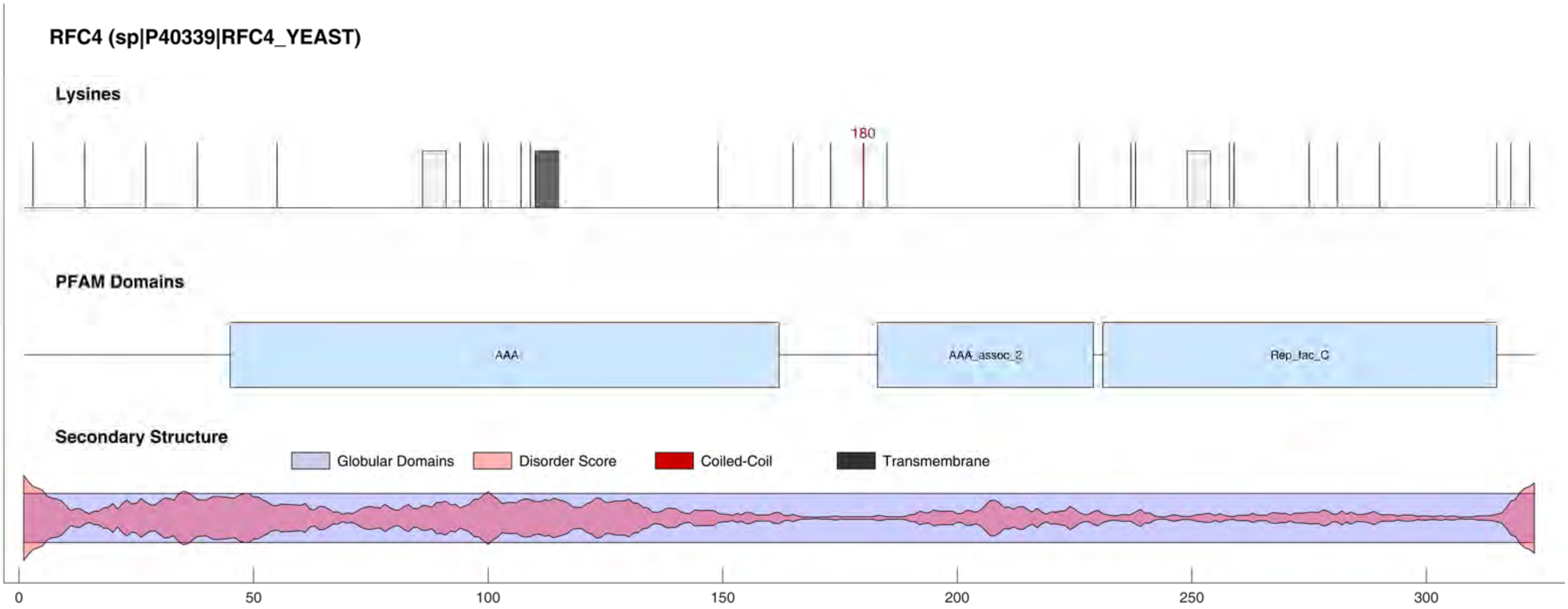
RFC4 (sp|P40339|RFC4_YEAST)
Lysines
180
PFAM Domains
AAA
AAA_assoc_2
Rep_fac_C
Secondary Structure
Globular Domains
Disorder Score
Coiled-Coil
Transmembrane
0
50
100
150
200
250
300

#### RGA1 (sp|P39083|RGA1_YEAST)

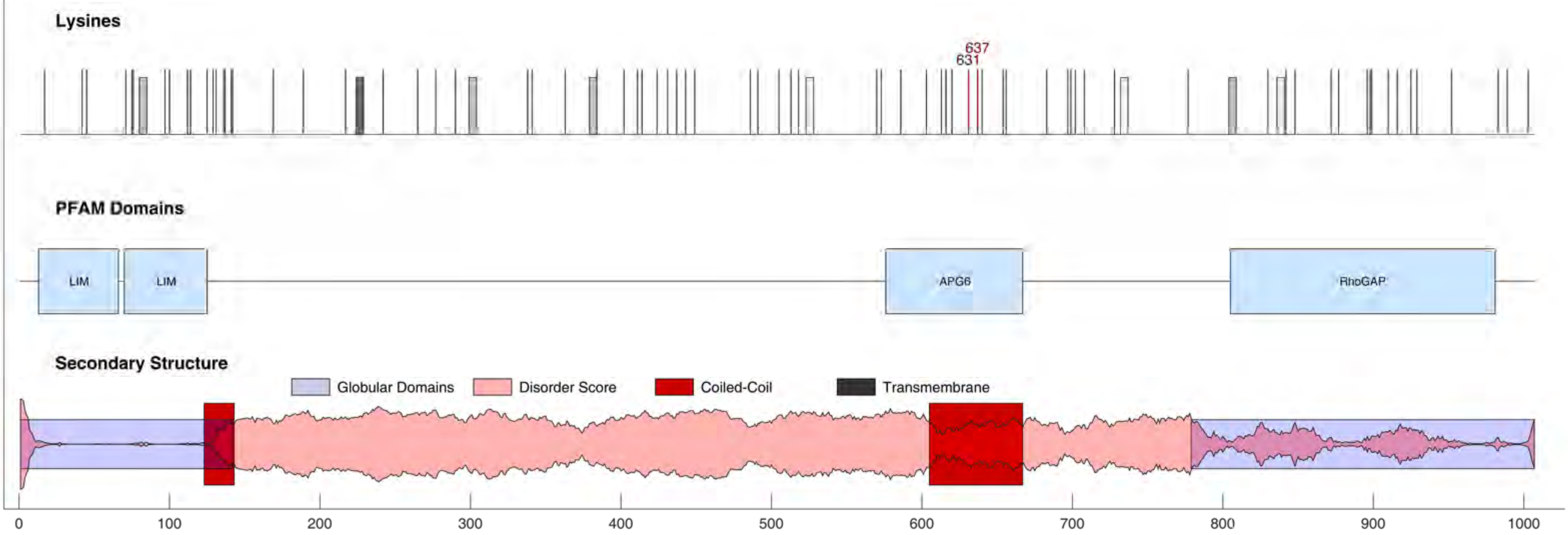

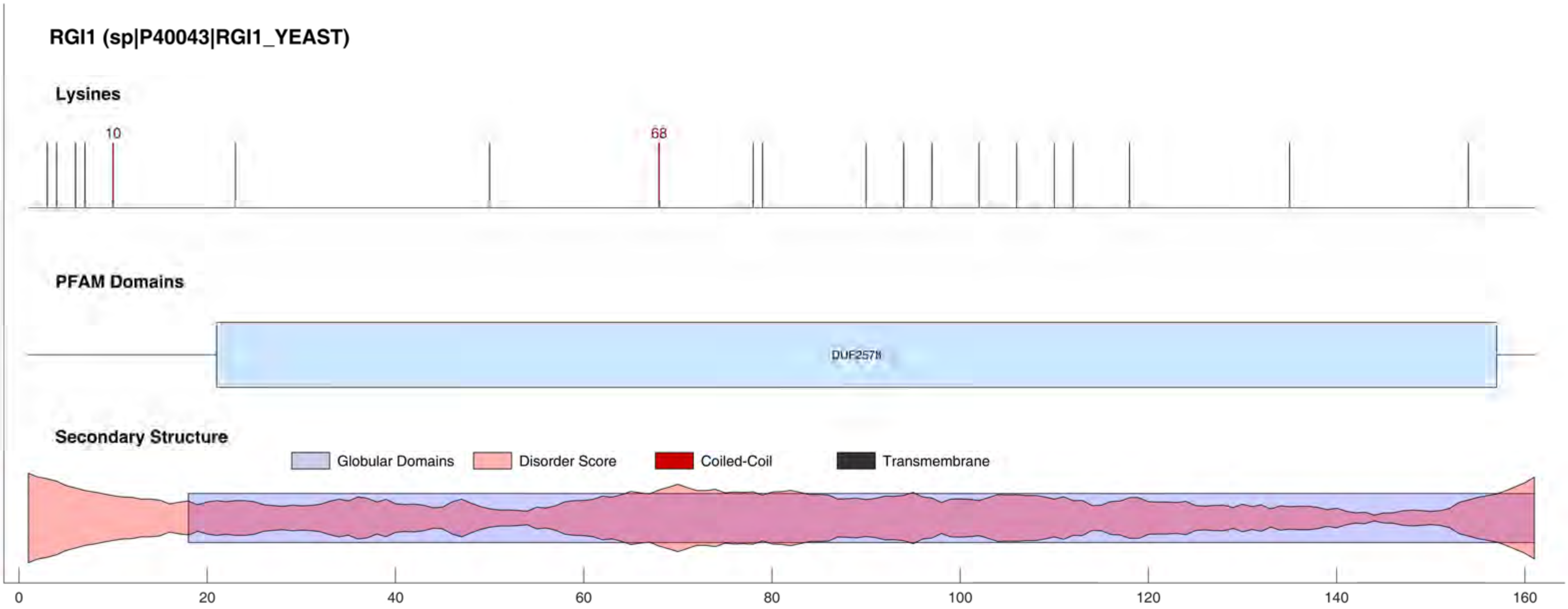
RGI1 (sp|P40043|RGI1_YEAST)
Lysines
10
68
PFAM Domains
DUF2578
Secondary Structure
Globular Domains
Disorder Score
Coiled-Coil
Transmembrane
0
20
40
60
80
100
120
140
160

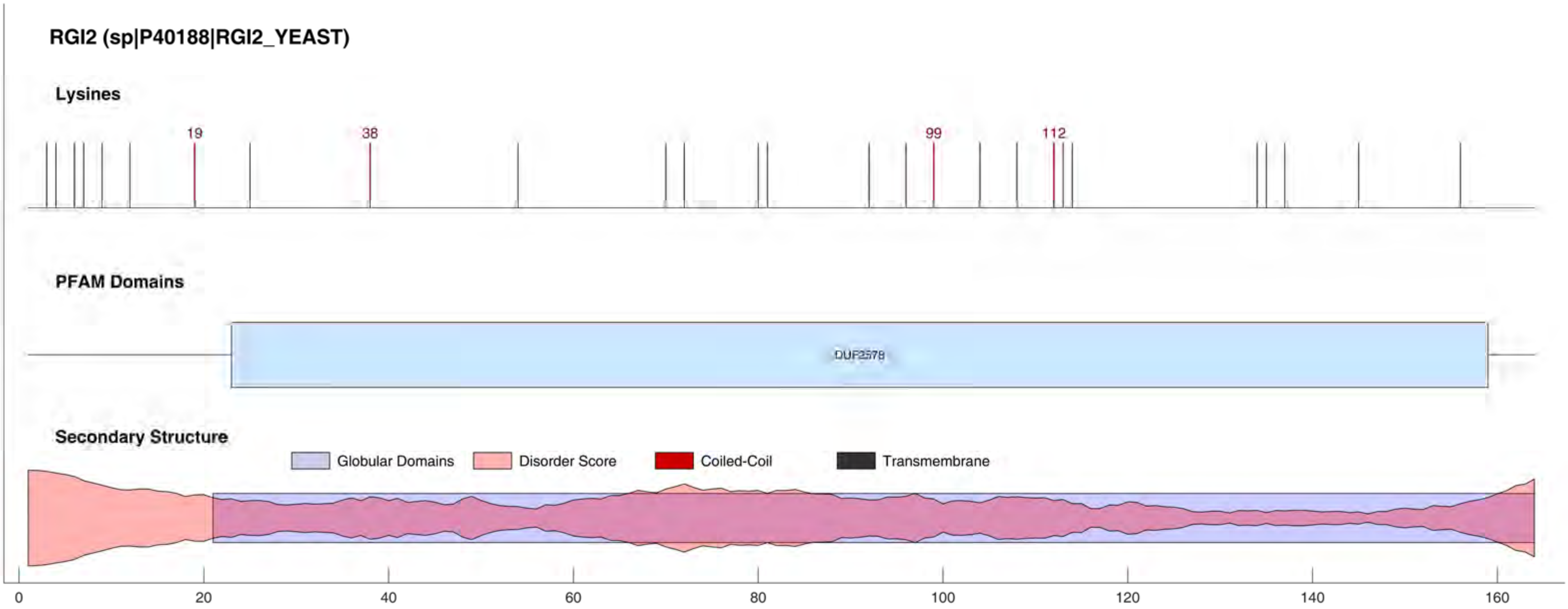
RGI2 (sp|P40188|RGI2_YEAST)
Lysines
19
38
99
112
PFAM Domains
DUF2578
Secondary Structure
Globular Domains
Disorder Score
Coiled-Coil
Transmembrane
0
20
40
60
80
100
120
140
160

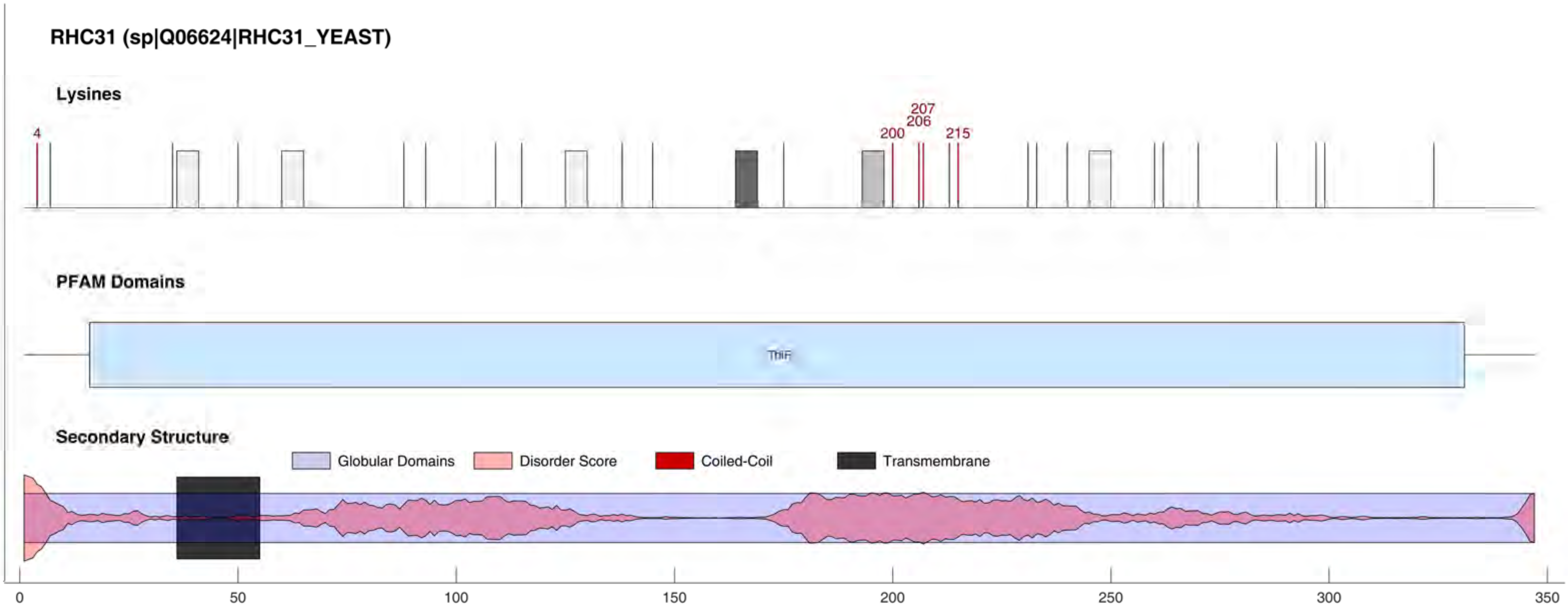
RHC31 (sp|Q06624|RHC31_YEAST)
Lysines
4
200
206
207
215
PFAM Domains
ThiF
Secondary Structure
Globular Domains
Disorder Score
Coiled-Coil
Transmembrane
0
50
100
150
200
250
300
350

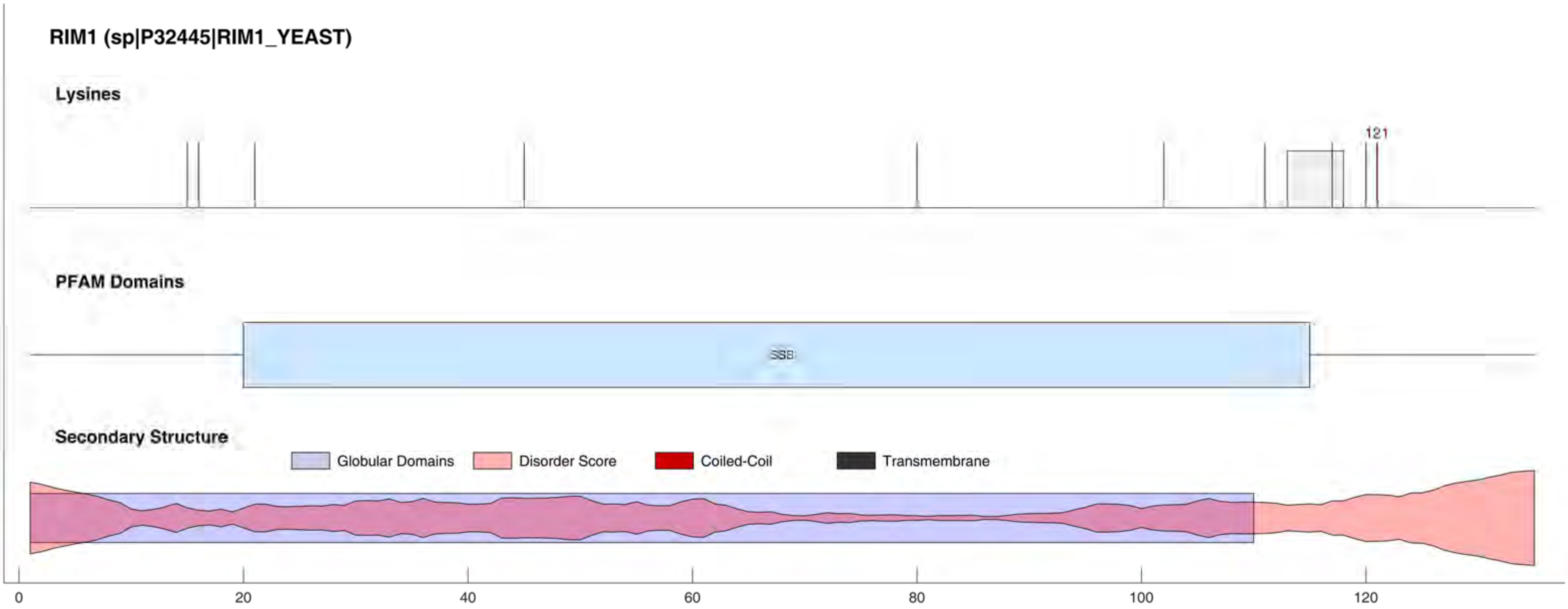

RIM1 (sp|P32445|RIM1_YEAST)
Lysines
121
PFAM Domains
SSB
Secondary Structure
Globular Domains
Disorder Score
Coiled-Coil
Transmembrane
0
20
40
60
80
100
120

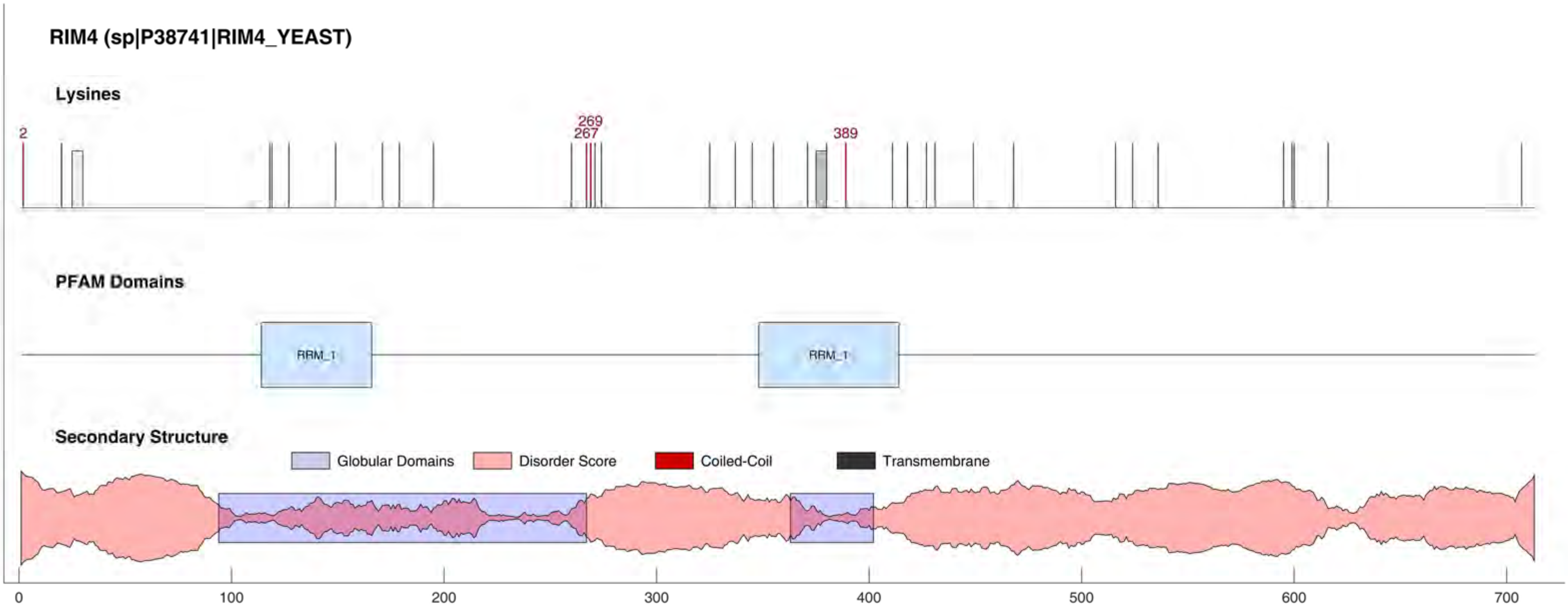
RIM4 (sp|P38741|RIM4_YEAST)
Lysines
2
269
267
389
PFAM Domains
RRM_1
RRM_1
Secondary Structure
Globular Domains
Disorder Score
Coiled-Coil
Transmembrane
0
100
200
300
400
500
600
700

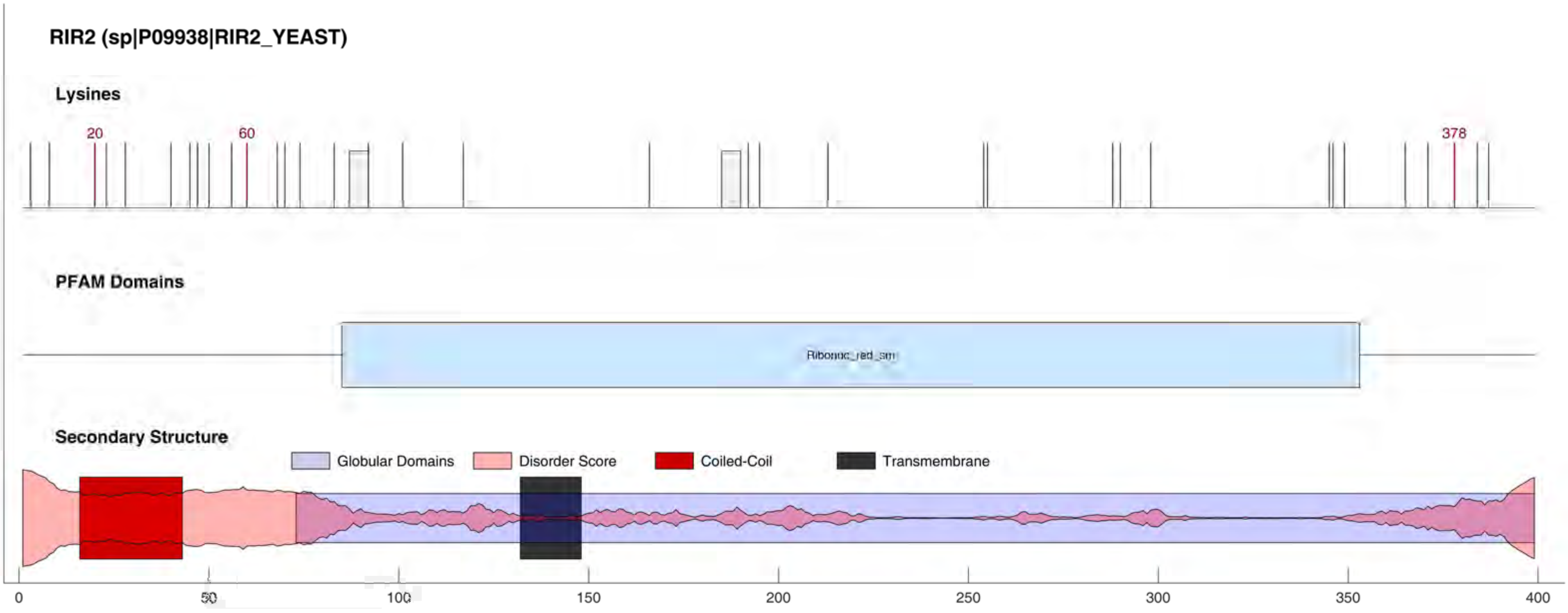
RIR2 (sp|P09938|RIR2_YEAST)
Lysines
20
60
378
PFAM Domains
Ribonuc_red_sm
Secondary Structure
Globular Domains
Disorder Score
Coiled-Coil
Transmembrane
0
50
100
150
200
250
300
350
400

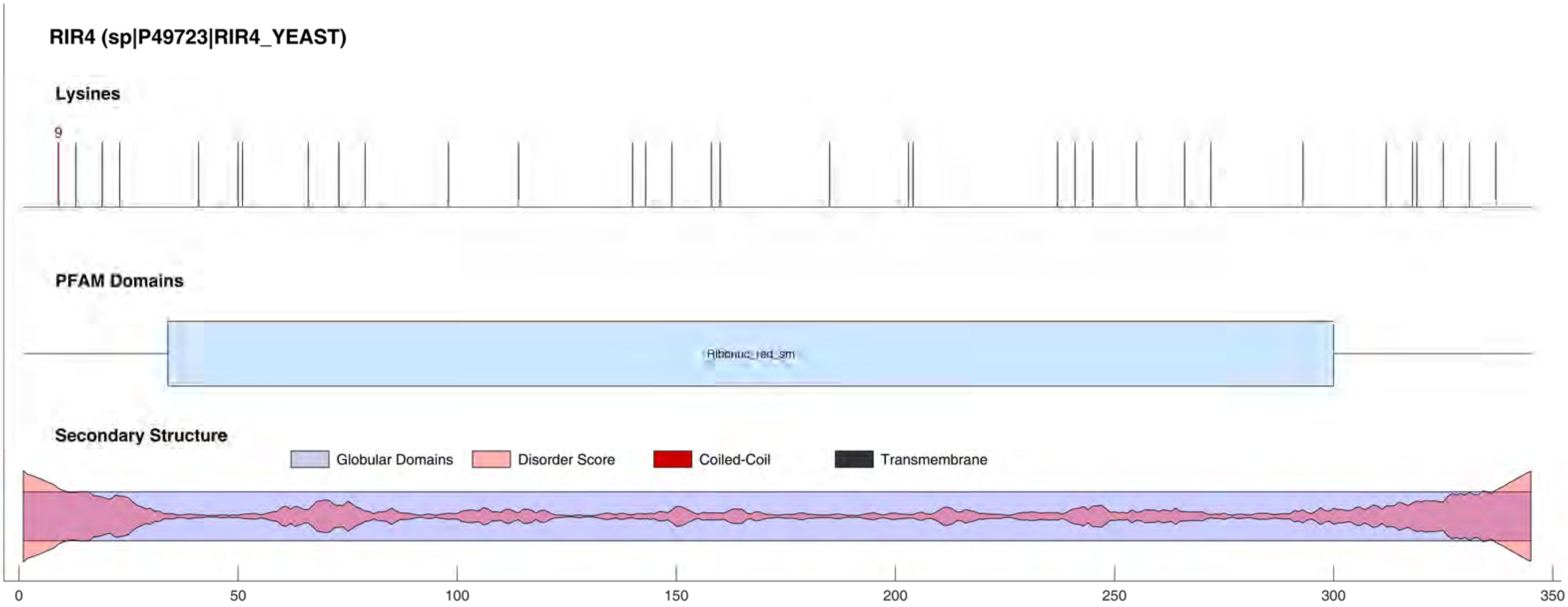

RIR4 (sp|P49723|RIR4_YEAST)
Lysines
9
PFAM Domains
Ribonuc_red_sm
Secondary Structure
Globular Domains
Disorder Score
Coiled-Coil
Transmembrane
0
50
100
150
200
250
300
350

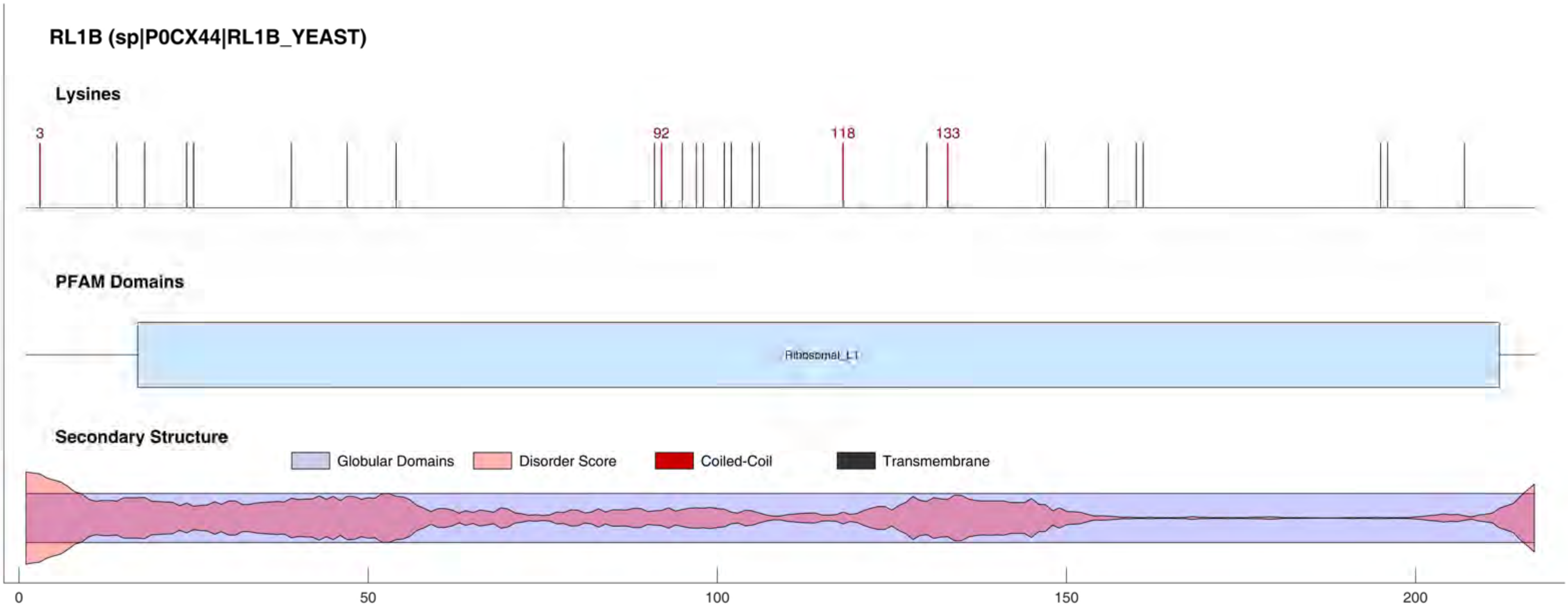

RL1B (sp|P0CX44|RL1B_YEAST)
Lysines
3
92
118
133
PFAM Domains
Ribosomal_L1
Secondary Structure
Globular Domains
Disorder Score
Coiled-Coil
Transmembrane
0
50
100
150
200

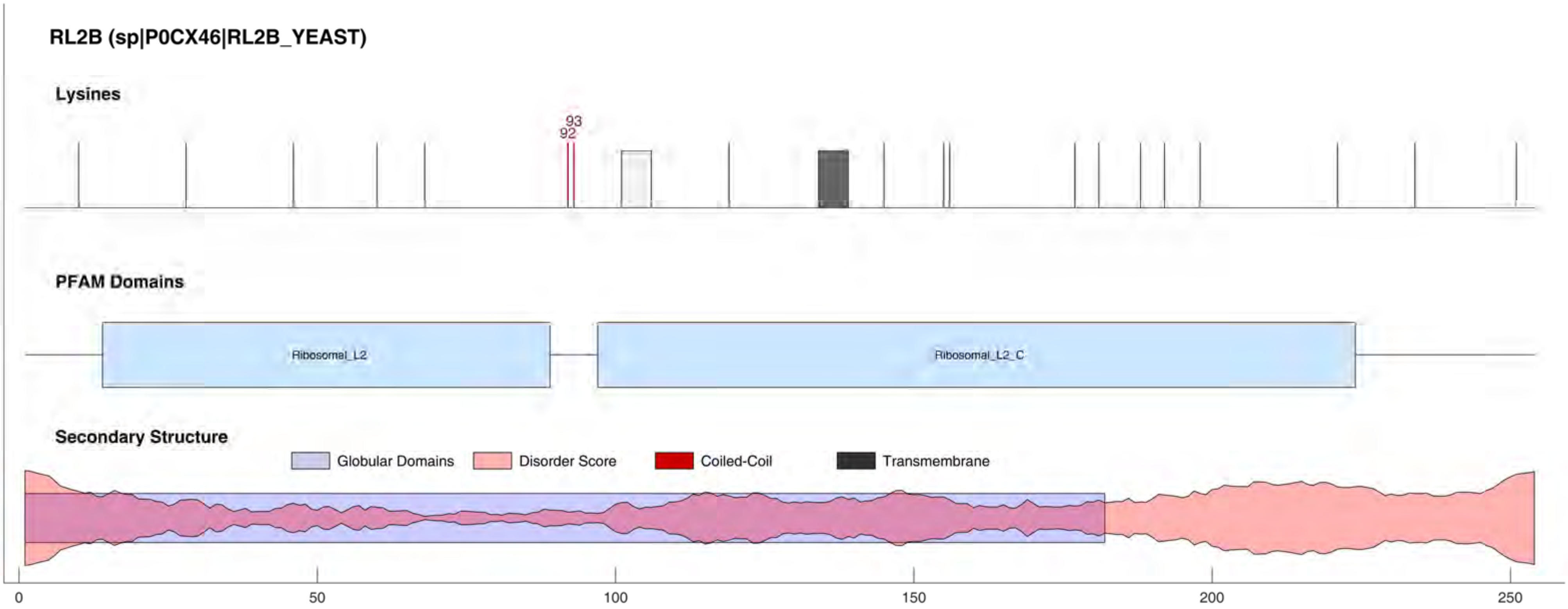

RL2B (sp|P0CX46|RL2B_YEAST)
Lysines
93
92
PFAM Domains
Ribosomal_L2
Ribosomal_L2_C
Secondary Structure
Globular Domains
Disorder Score
Coiled-Coil
Transmembrane
0
50
100
150
200
250

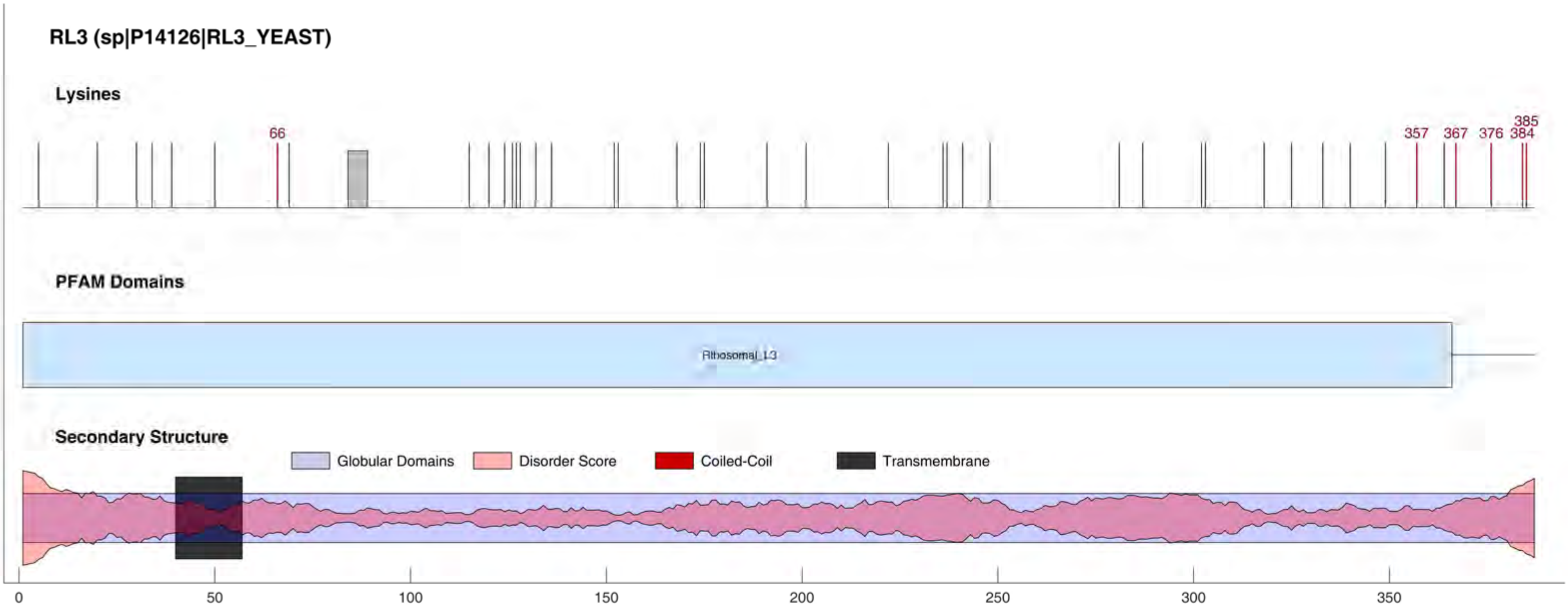
RL3 (sp|P14126|RL3_YEAST)
Lysines
66
357
367
376
384
385
PFAM Domains
Ribosomal_L3
Secondary Structure
Globular Domains
Disorder Score
Coiled-Coil
Transmembrane
0
50
100
150
200
250
300
350

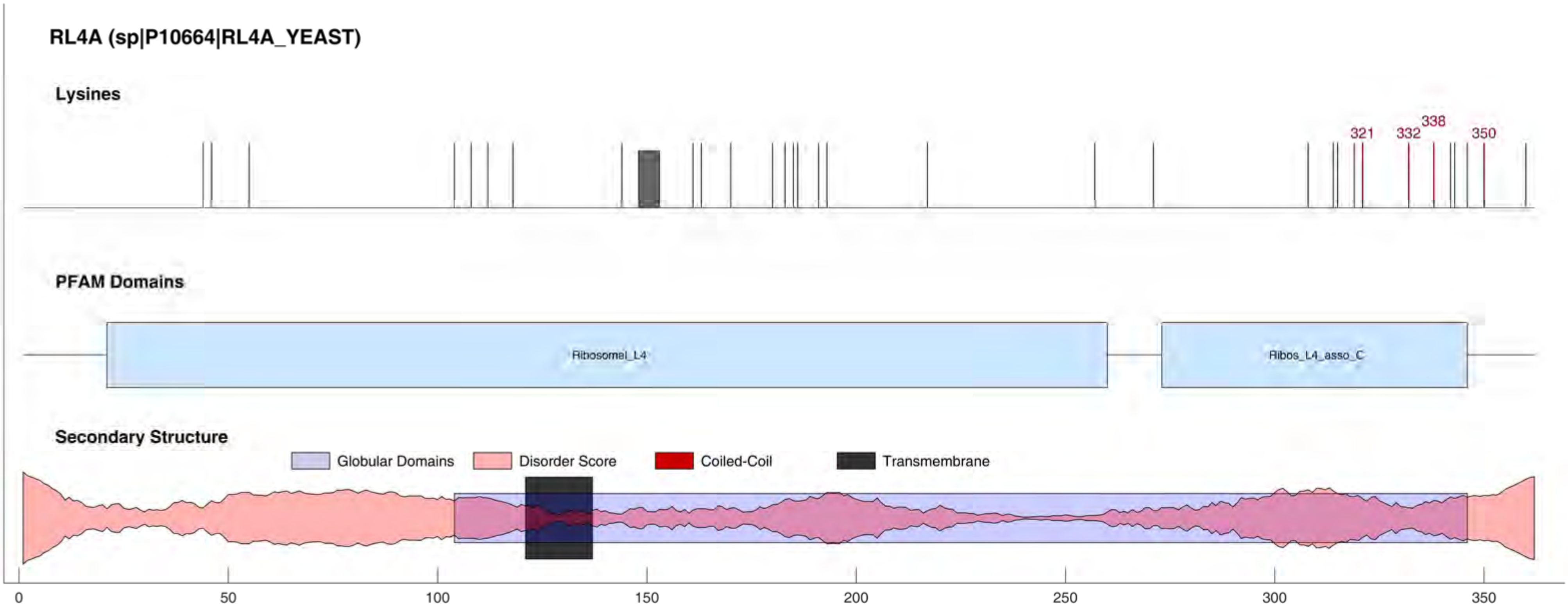
RL4A (sp|P10664|RL4A_YEAST)
Lysines
321
332
338
350
PFAM Domains
Ribosomal_L4
Ribos_L4_asso_C
Secondary Structure
Globular Domains
Disorder Score
Coiled-Coil
Transmembrane
0
50
100
150
200
250
300
350

#### RL5 (sp|P26321|RL5_YEAST)

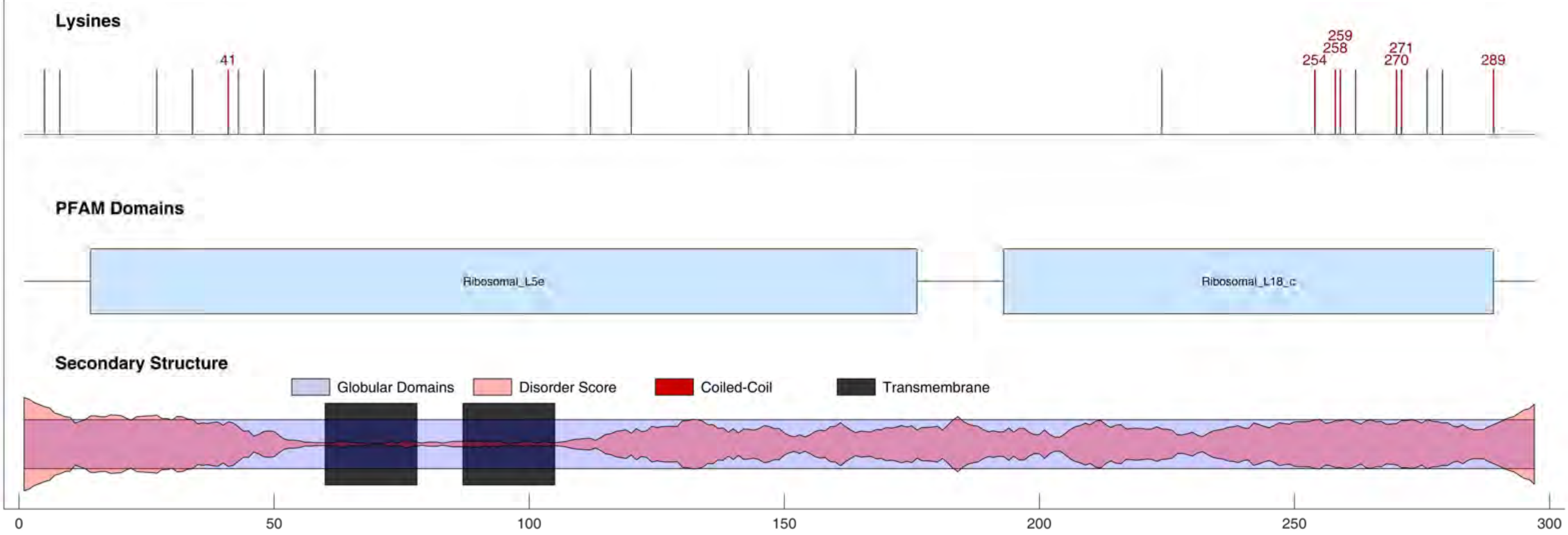

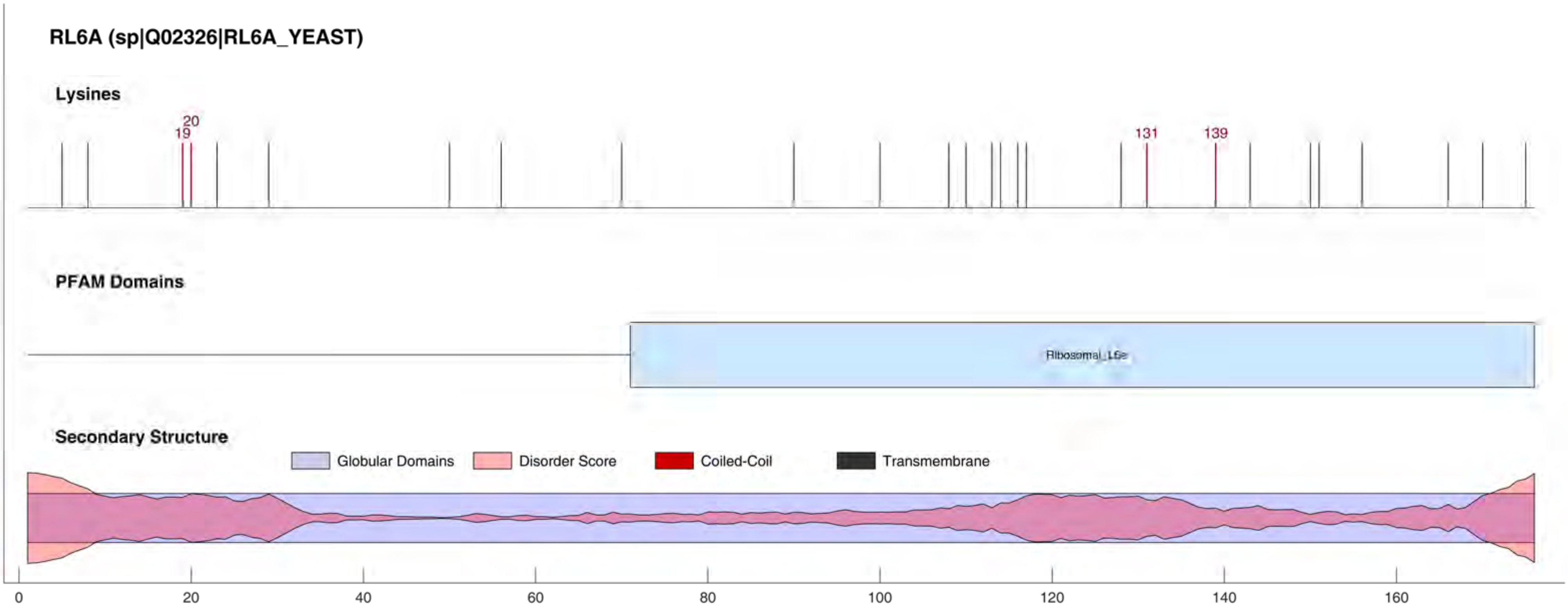
RL6A (sp|Q02326|RL6A_YEAST)
Lysines
19
20
131
139
PFAM Domains
Ribosomal_L6e
Secondary Structure
Globular Domains
Disorder Score
Coiled-Coil
Transmembrane
0
20
40
60
80
100
120
140
160

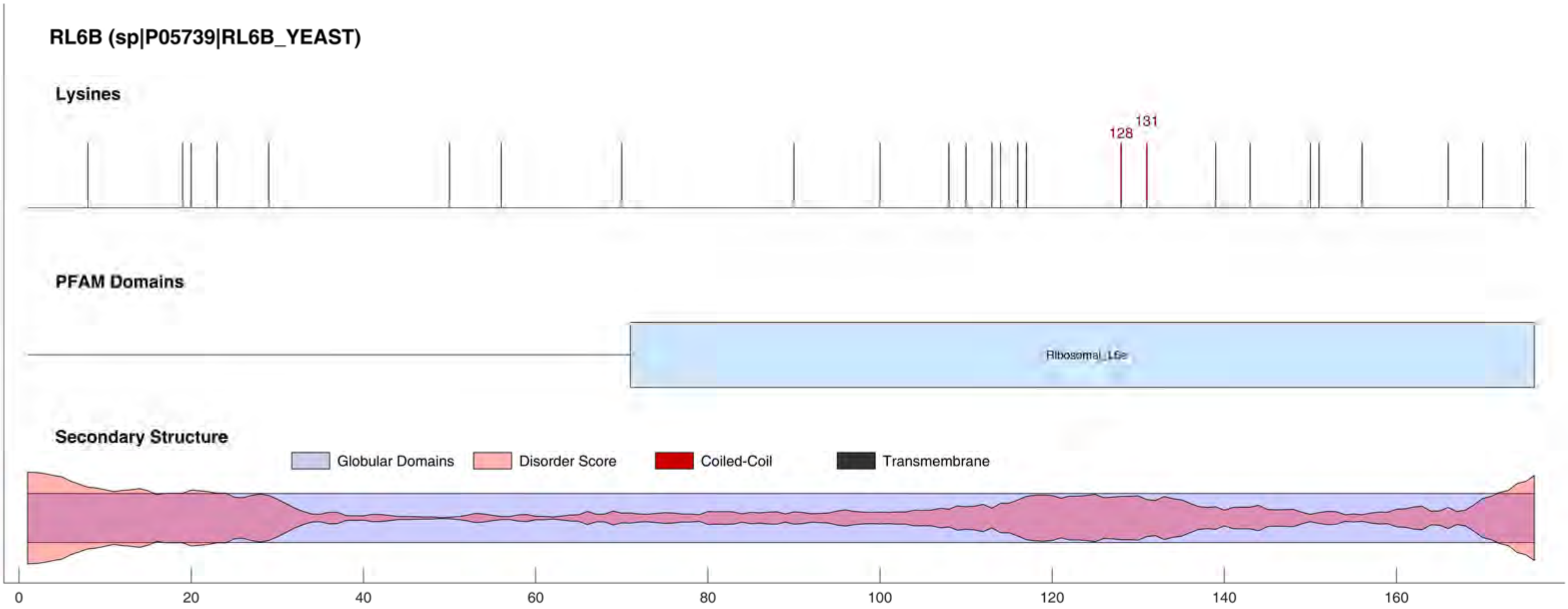

RL6B (sp|P05739|RL6B_YEAST)
Lysines
128
131
PFAM Domains
Ribosomal_L6e
Secondary Structure
Globular Domains
Disorder Score
Coiled-Coil
Transmembrane
0
20
40
60
80
100
120
140
160

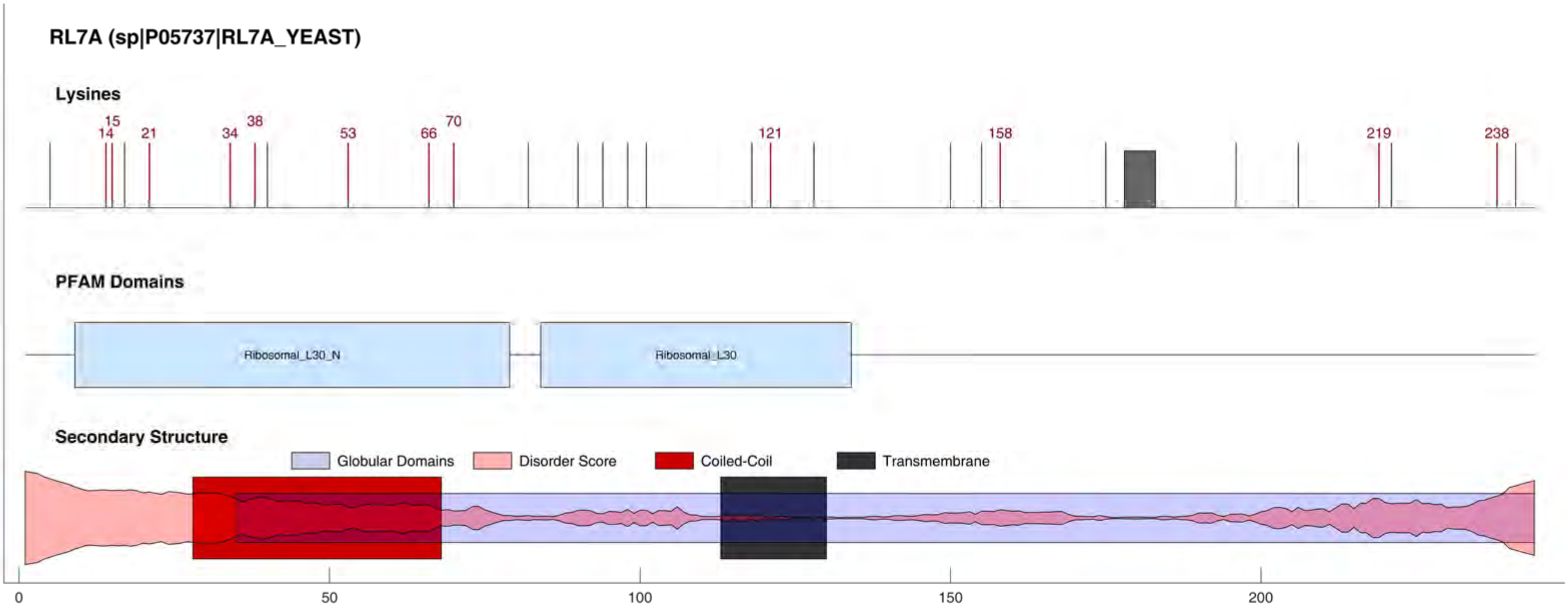
RL7A (sp|P05737|RL7A_YEAST)
Lysines
14
15
21
34
38
53
66
70
121
158
219
238
PFAM Domains
Ribosomal_L30_N
Ribosomal_L30
Secondary Structure
Globular Domains
Disorder Score
Coiled-Coil
Transmembrane
0
50
100
150
200

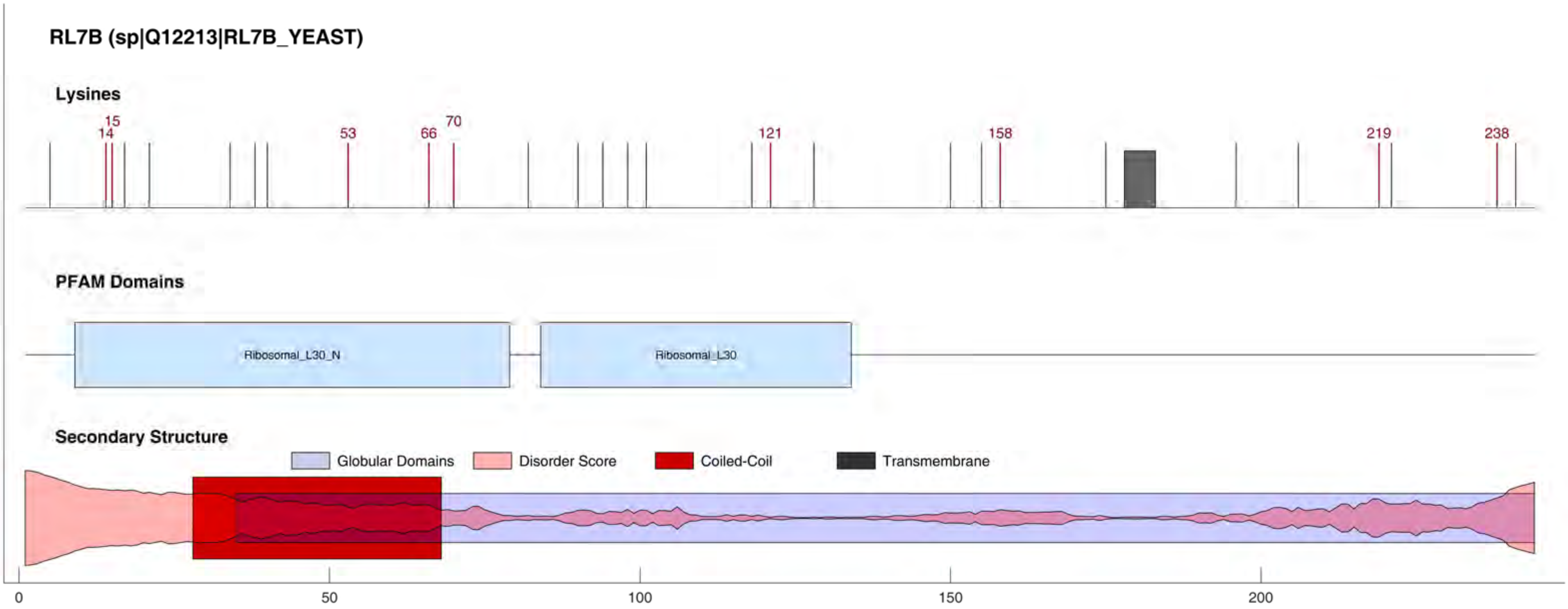
RL7B (sp|Q12213|RL7B_YEAST)
Lysines
14
15
53
66
70
121
158
219
238
PFAM Domains
Ribosomal_L30_N
Ribosomal_L30
Secondary Structure
Globular Domains
Disorder Score
Coiled-Coil
Transmembrane
0
50
100
150
200

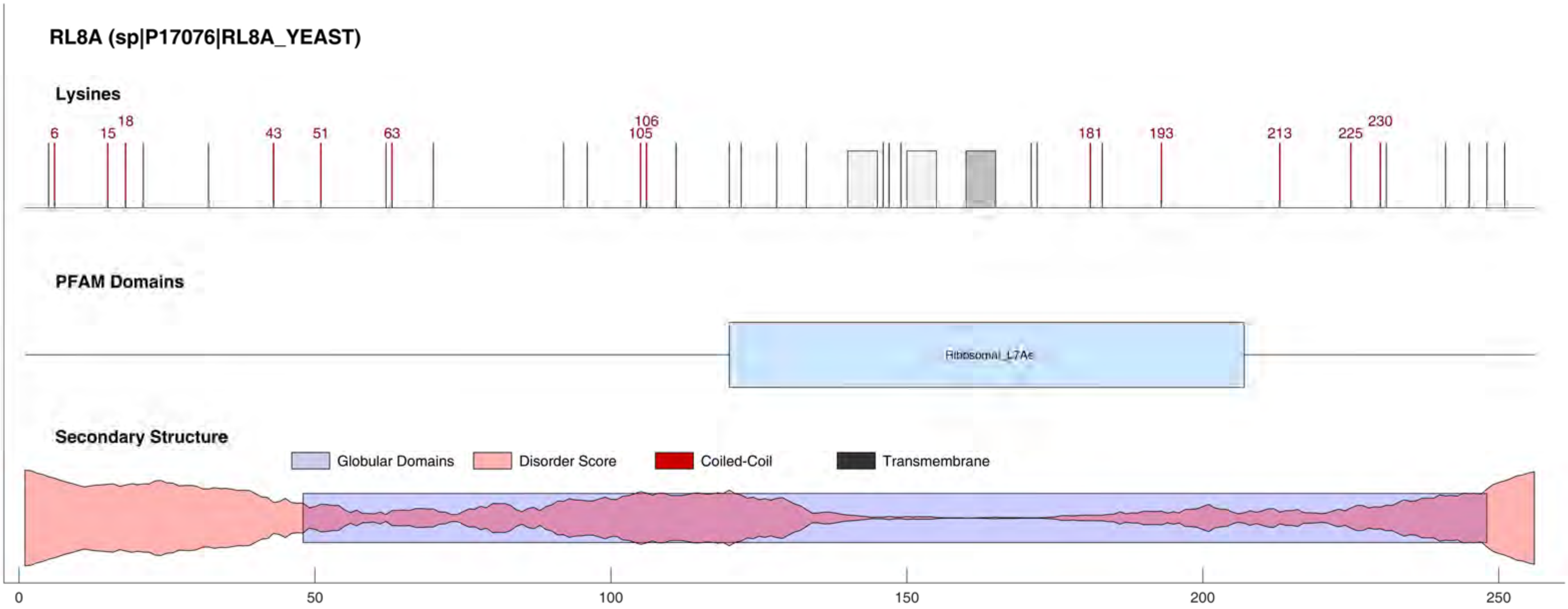
RL8A (sp|P17076|RL8A_YEAST)
Lysines
6
15
18
43
51
63
106
105
181
193
213
225
230
PFAM Domains
Ribosomal_L7Ae
Secondary Structure
Globular Domains
Disorder Score
Coiled-Coil
Transmembrane
0
50
100
150
200
250

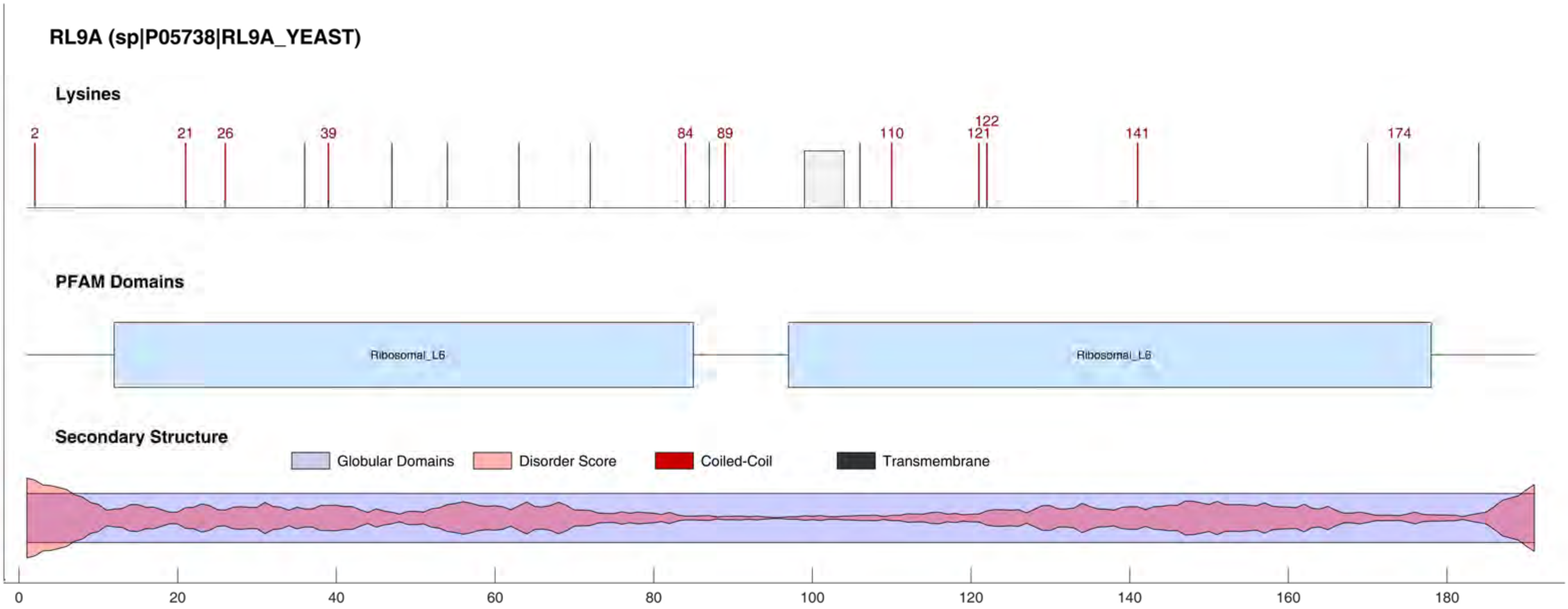

RL9A (sp|P05738|RL9A_YEAST)
Lysines
2
21
26
39
84
89
110
121
122
141
174
PFAM Domains
Ribosomal_L6
Ribosomal_L6
Secondary Structure
Globular Domains
Disorder Score
Coiled-Coil
Transmembrane
0
20
40
60
80
100
120
140
160
180

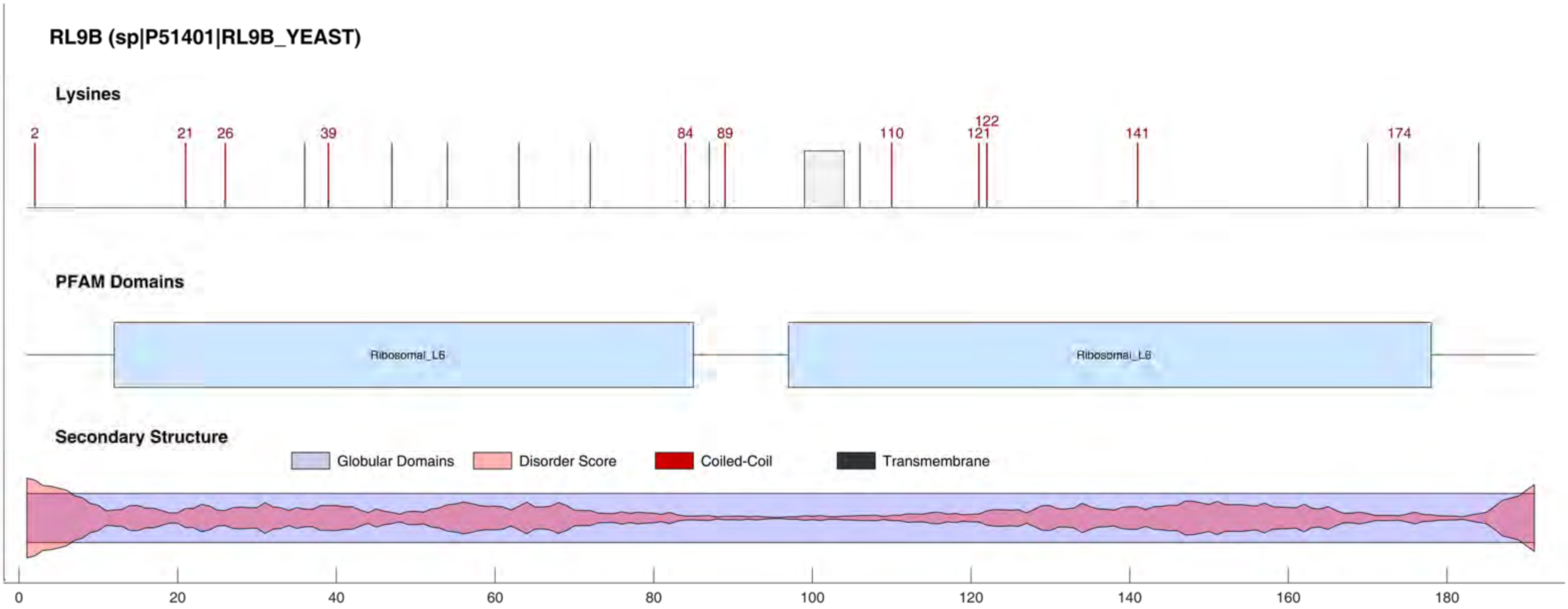
RL9B (sp|P51401|RL9B_YEAST)
Lysines
2
21
26
39
84
89
110
121
122
141
174
PFAM Domains
Ribosomal_L6
Ribosomal_L6
Secondary Structure
Globular Domains
Disorder Score
Coiled-Coil
Transmembrane
0
20
40
60
80
100
120
140
160
180

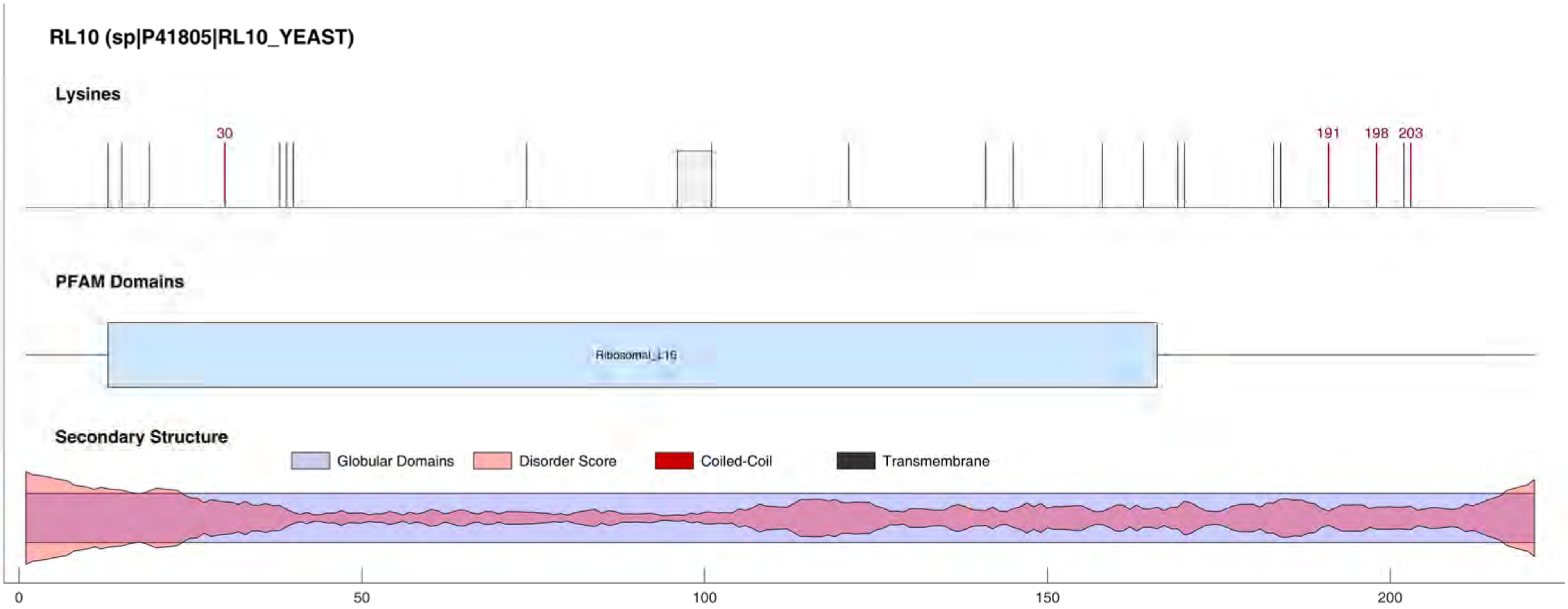
RL10 (sp|P41805|RL10_YEAST)
Lysines
30
191
198
203
PFAM Domains
Ribosomal_L16
Secondary Structure
Globular Domains
Disorder Score
Coiled-Coil
Transmembrane
0
50
100
150
200

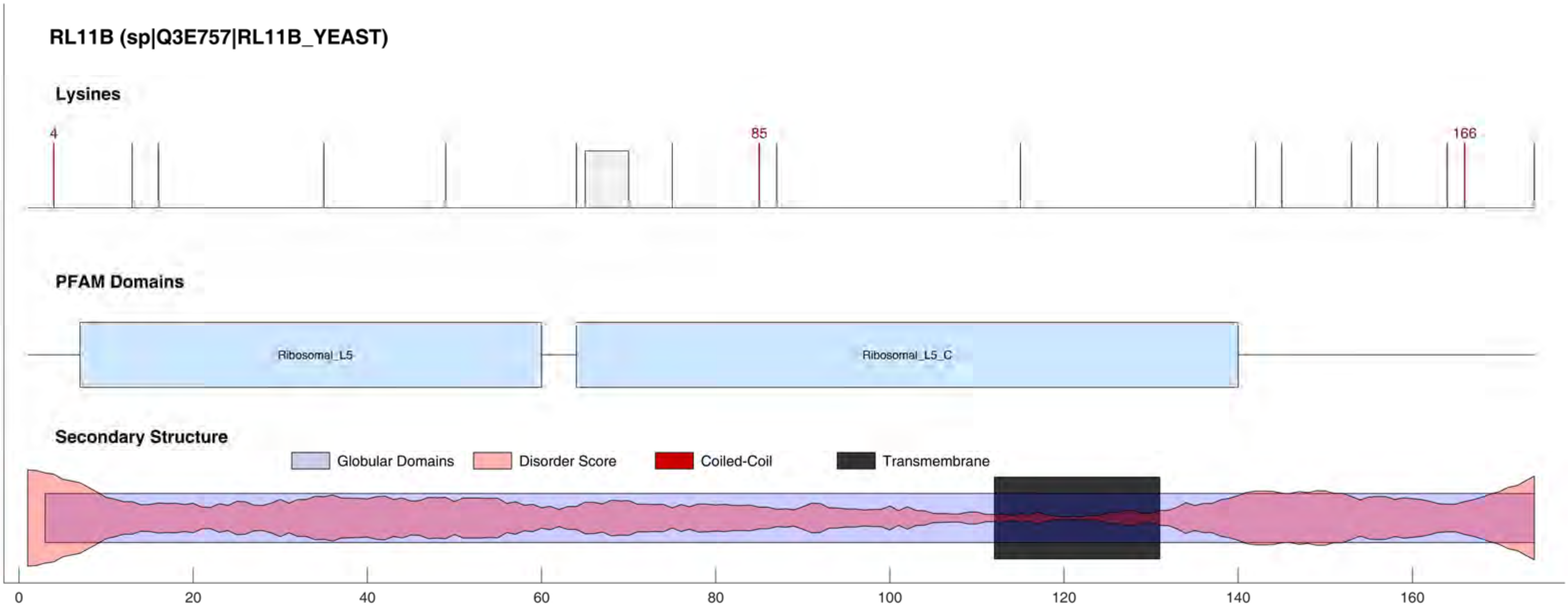
RL11B (sp|Q3E757|RL11B_YEAST)
Lysines
4
85
166
PFAM Domains
Ribosomal_L5
Ribosomal_L5_C
Secondary Structure
Globular Domains
Disorder Score
Coiled-Coil
Transmembrane
0
20
40
60
80
100
120
140
160

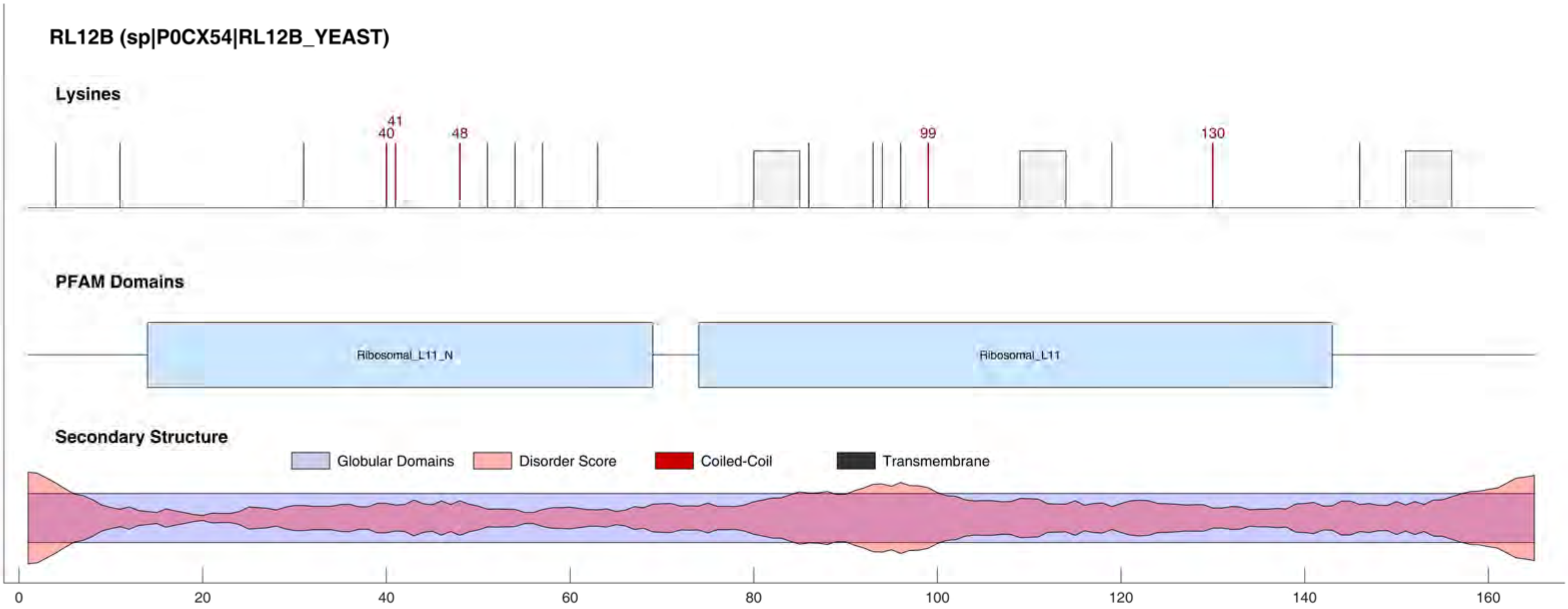
RL12B (sp|P0CX54|RL12B_YEAST)
Lysines
41
40
48
99
130
PFAM Domains
Ribosomal_L11_N
Ribosomal_L11
Secondary Structure
Globular Domains
Disorder Score
Coiled-Coil
Transmembrane
0
20
40
60
80
100
120
140
160

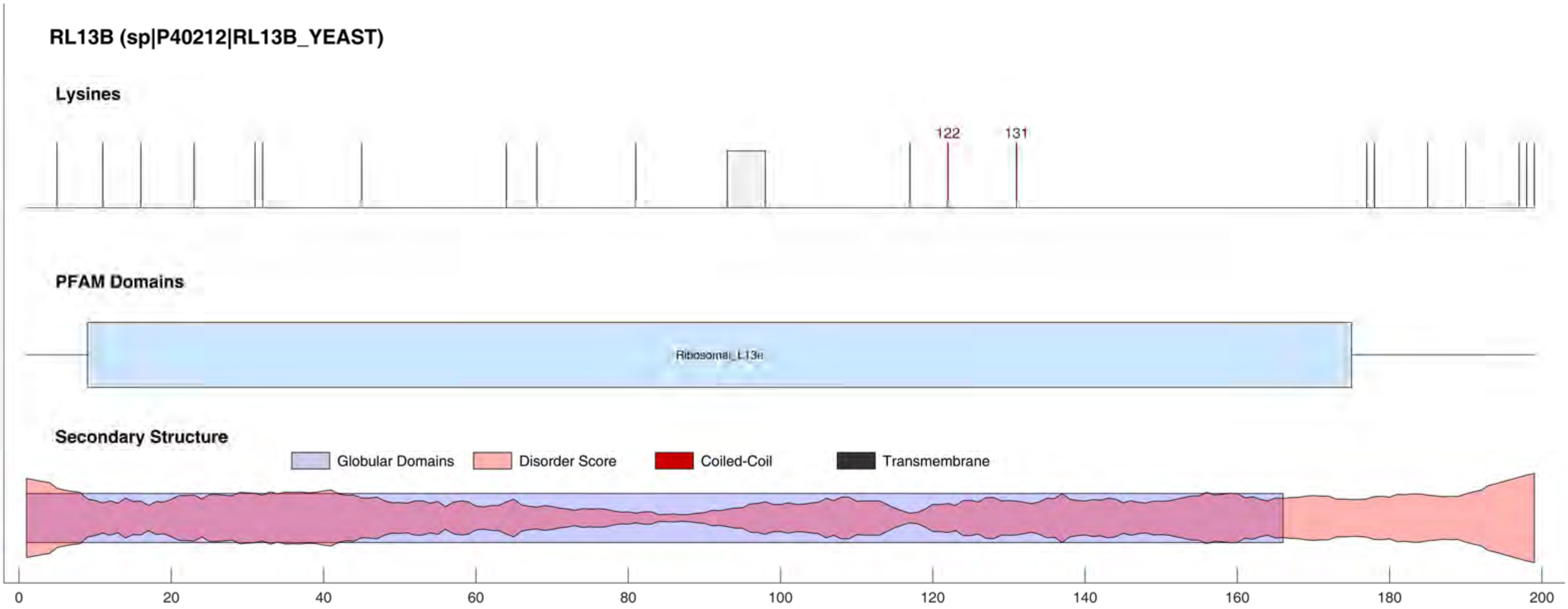
RL13B (sp|P40212|RL13B_YEAST)
Lysines
122
131
PFAM Domains
Ribosomal_L13e
Secondary Structure
Globular Domains
Disorder Score
Coiled-Coil
Transmembrane
0
20
40
60
80
100
120
140
160
180
200

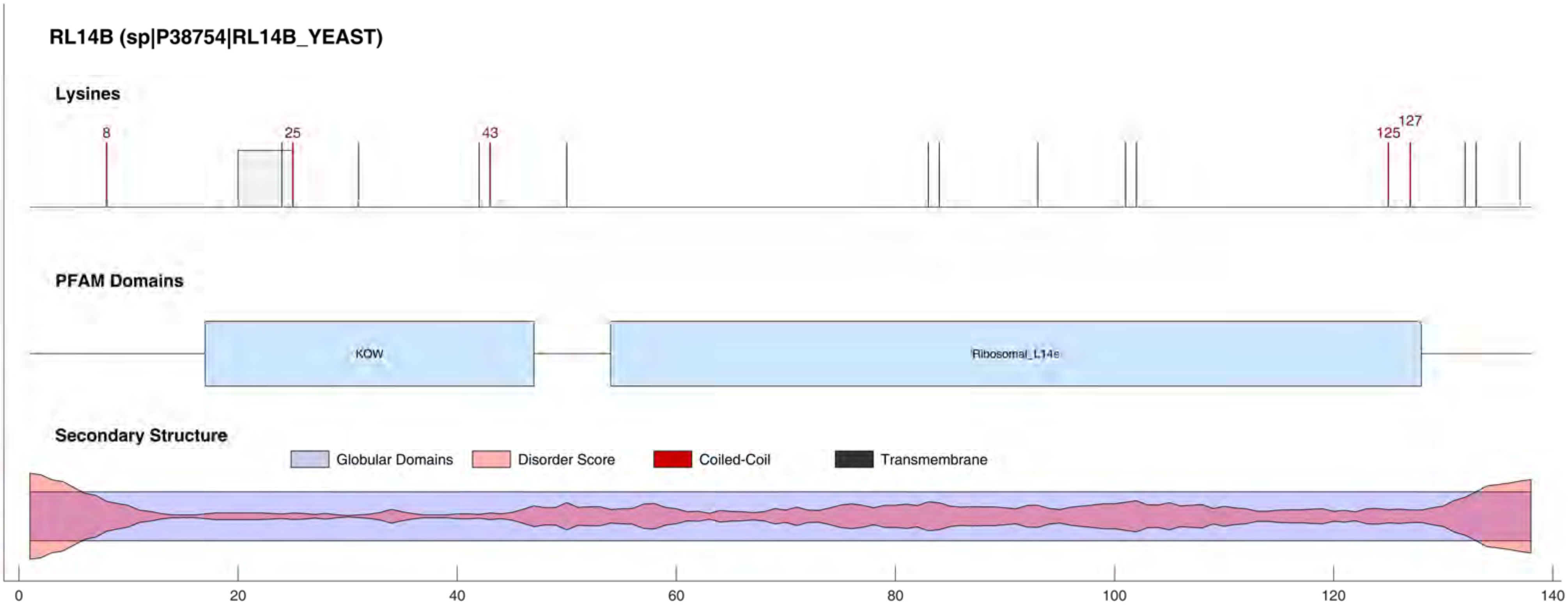
RL14B (sp|P38754|RL14B_YEAST)
Lysines
8
25
43
125
127
PFAM Domains
KOW
Ribosomal_L14e
Secondary Structure
Globular Domains
Disorder Score
Coiled-Coil
Transmembrane
0
20
40
60
80
100
120
140

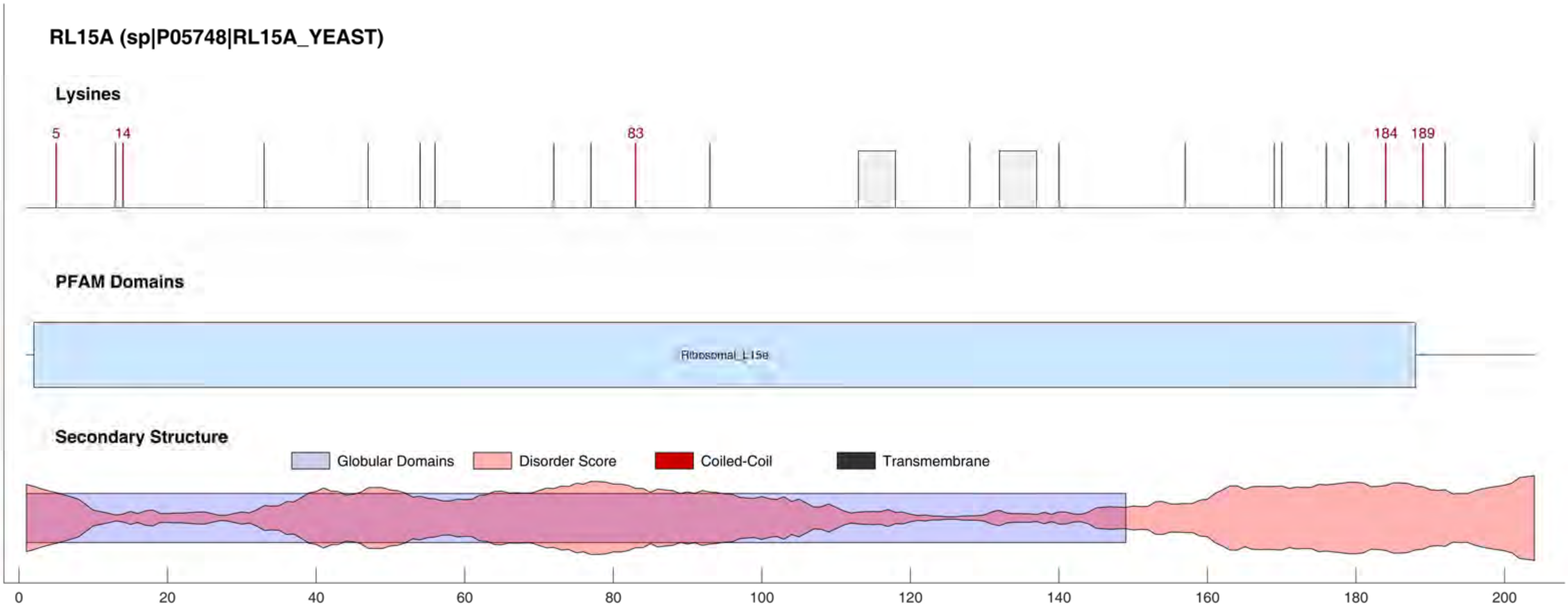
RL15A (sp|P05748|RL15A_YEAST)
Lysines
5
14
83
184
189
PFAM Domains
Ribosomal_L15e
Secondary Structure
Globular Domains
Disorder Score
Coiled-Coil
Transmembrane
0
20
40
60
80
100
120
140
160
180
200

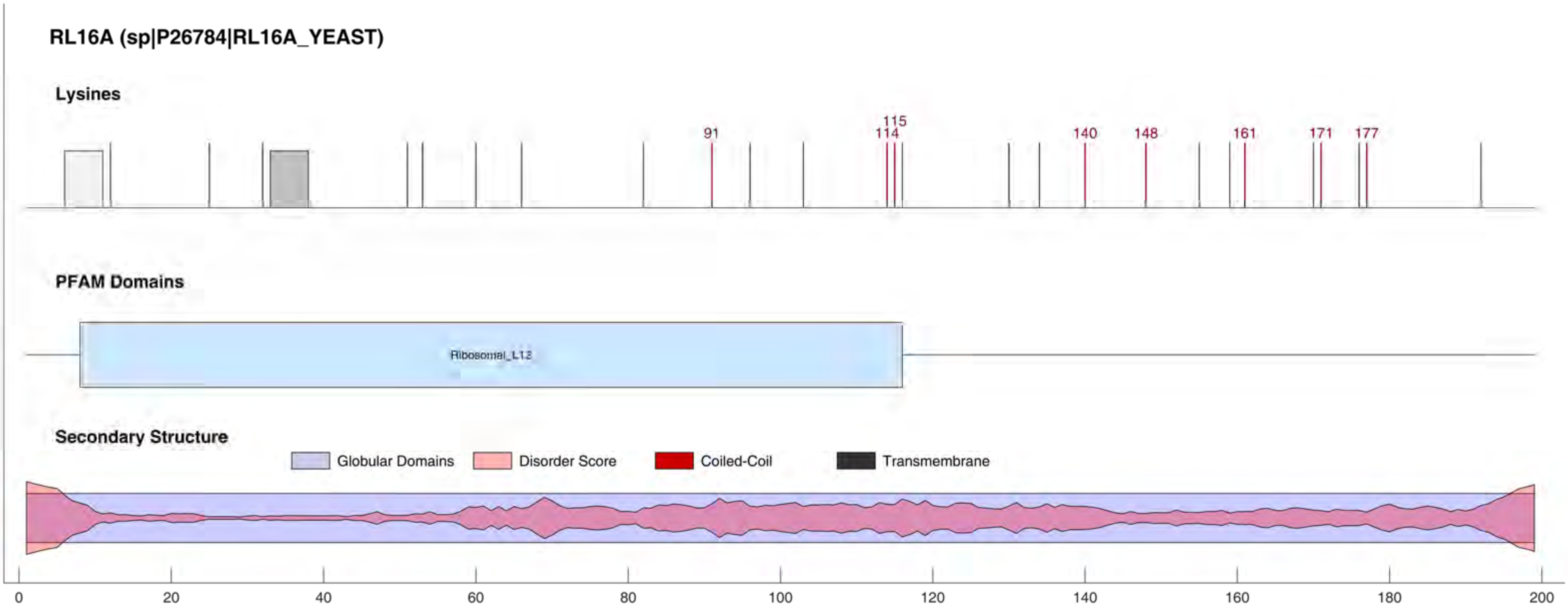

RL16A (sp|P26784|RL16A_YEAST)
Lysines
91
115
114
140
148
161
171
177
PFAM Domains
Ribosomal_L13
Secondary Structure
Globular Domains
Disorder Score
Coiled-Coil
Transmembrane
0
20
40
60
80
100
120
140
160
180
200

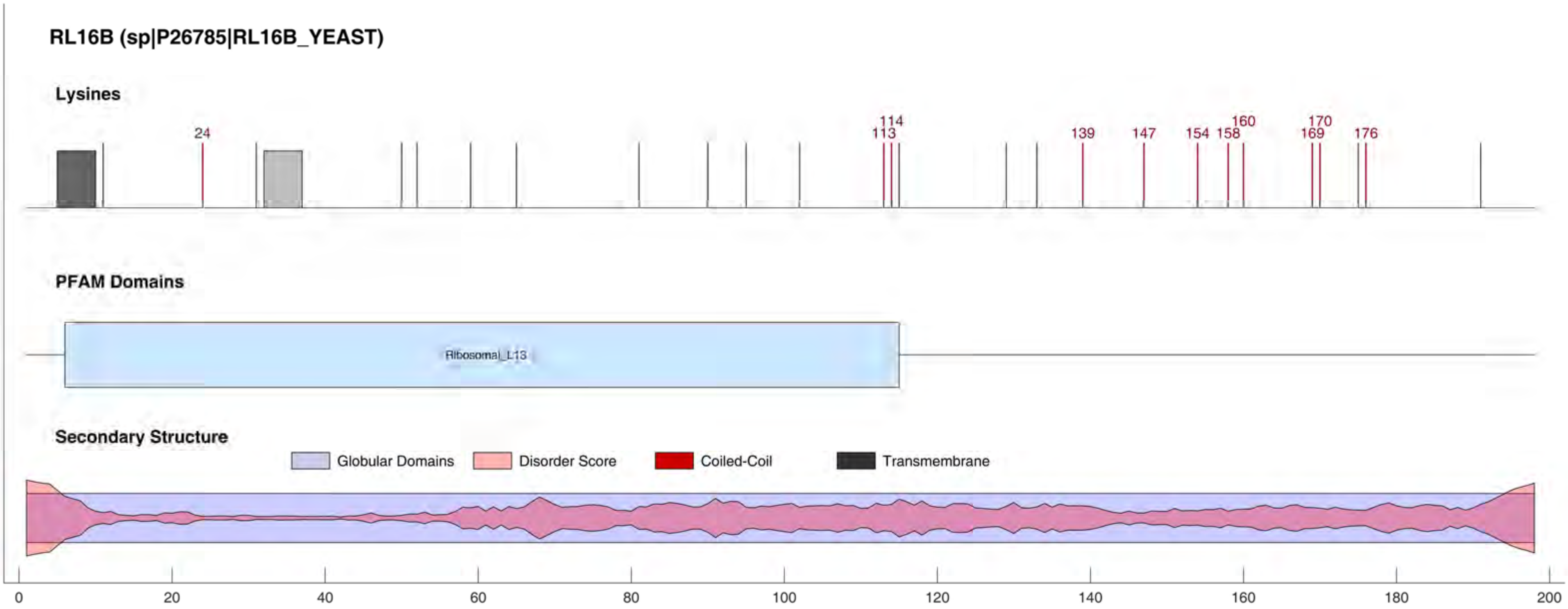

RL16B (sp|P26785|RL16B_YEAST)
Lysines
24
113
114
139
147
154
158
160
169
170
176
PFAM Domains
Ribosomal_L13
Secondary Structure
Globular Domains
Disorder Score
Coiled-Coil
Transmembrane
0
20
40
60
80
100
120
140
160
180
200

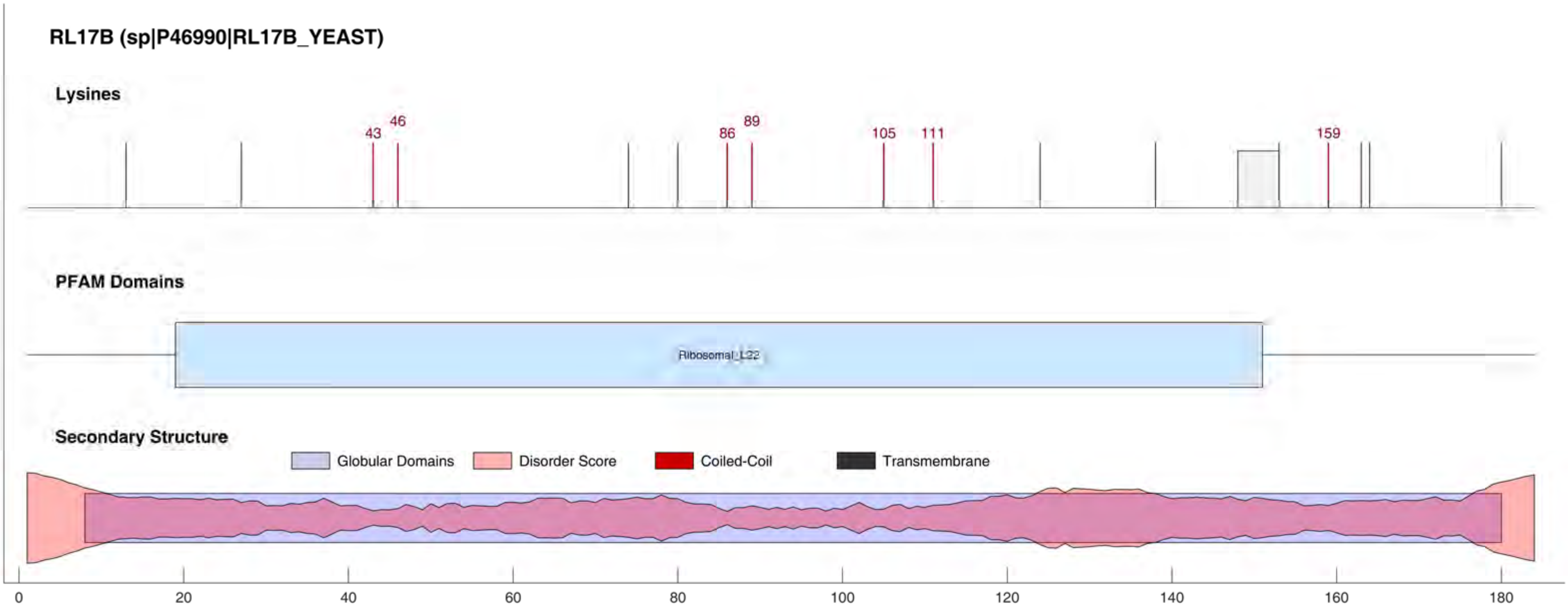
RL17B (sp|P46990|RL17B_YEAST)
Lysines
43
46
86
89
105
111
159
PFAM Domains
Ribosomal_L22
Secondary Structure
Globular Domains
Disorder Score
Coiled-Coil
Transmembrane
0
20
40
60
80
100
120
140
160
180

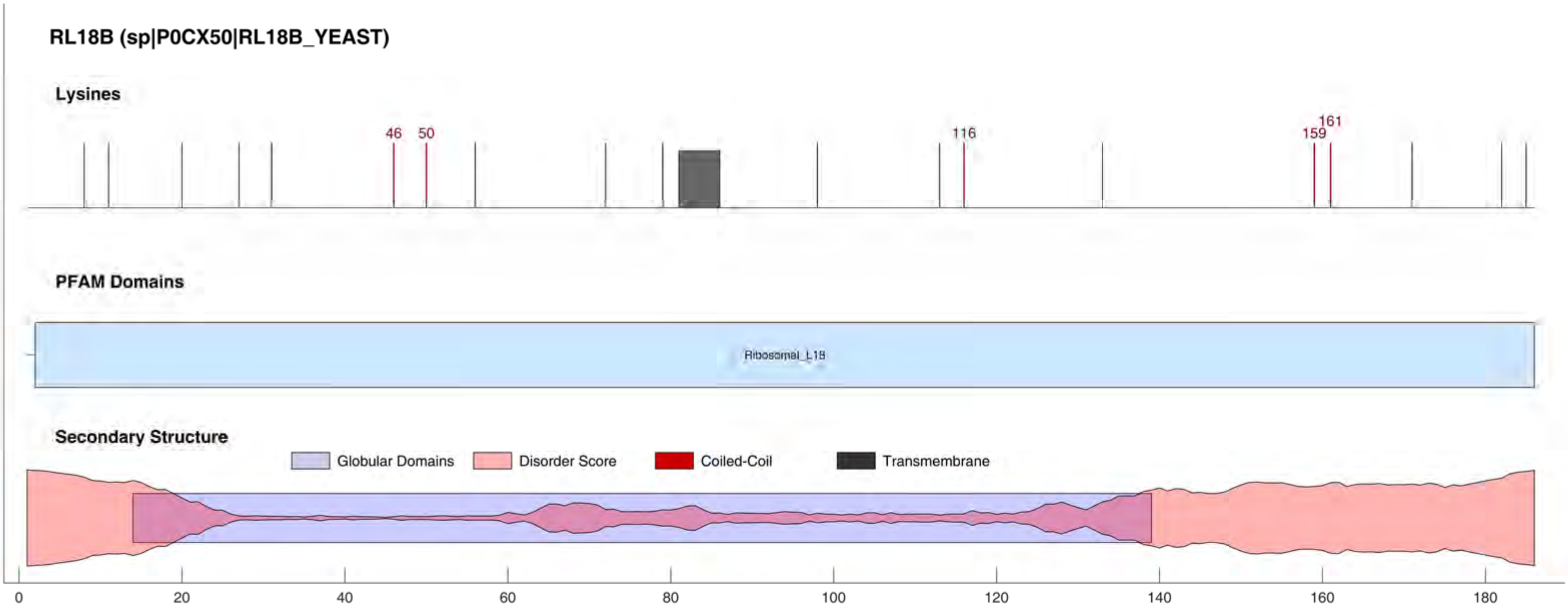
RL18B (sp|P0CX50|RL18B_YEAST)
Lysines
46
50
116
159
161
PFAM Domains
Ribosomal_L18
Secondary Structure
Globular Domains
Disorder Score
Coiled-Coil
Transmembrane
0
20
40
60
80
100
120
140
160
180

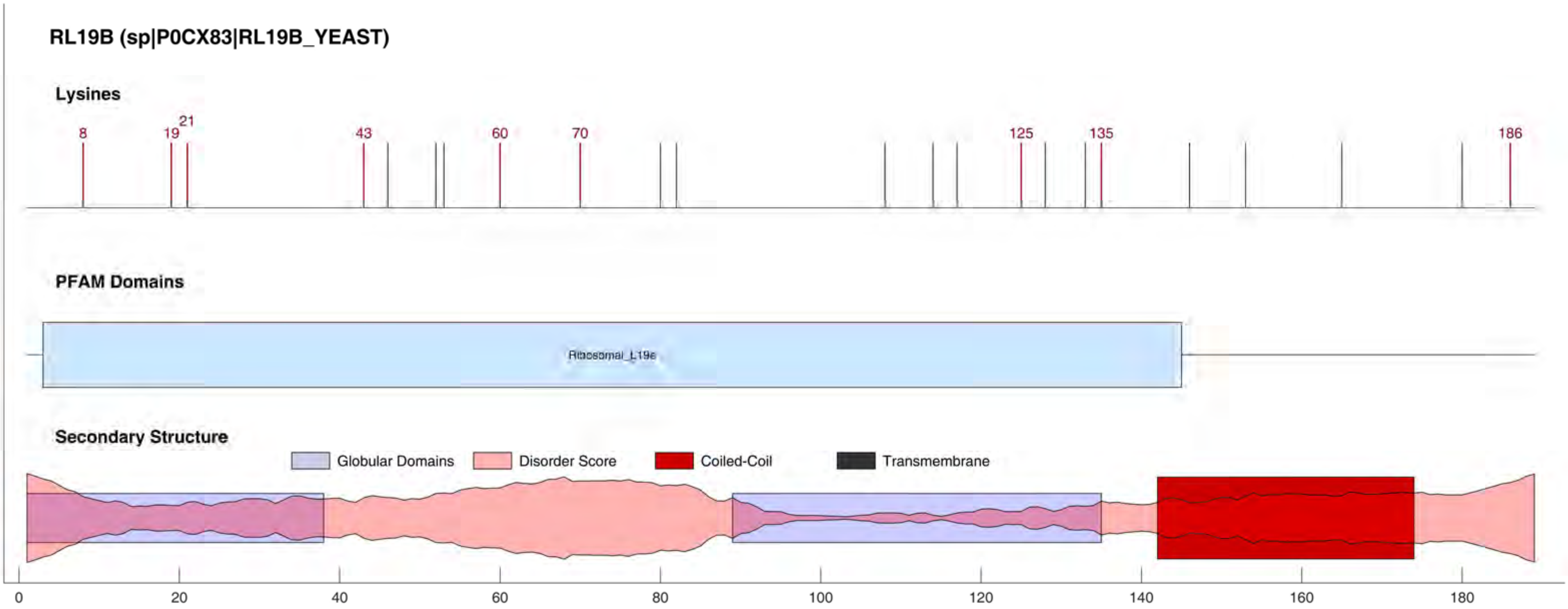

RL19B (sp|P0CX83|RL19B_YEAST)
Lysines
8
19
21
43
60
70
125
135
186
PFAM Domains
Ribosomal_L19e
Secondary Structure
Globular Domains
Disorder Score
Coiled-Coil
Transmembrane
0
20
40
60
80
100
120
140
160
180

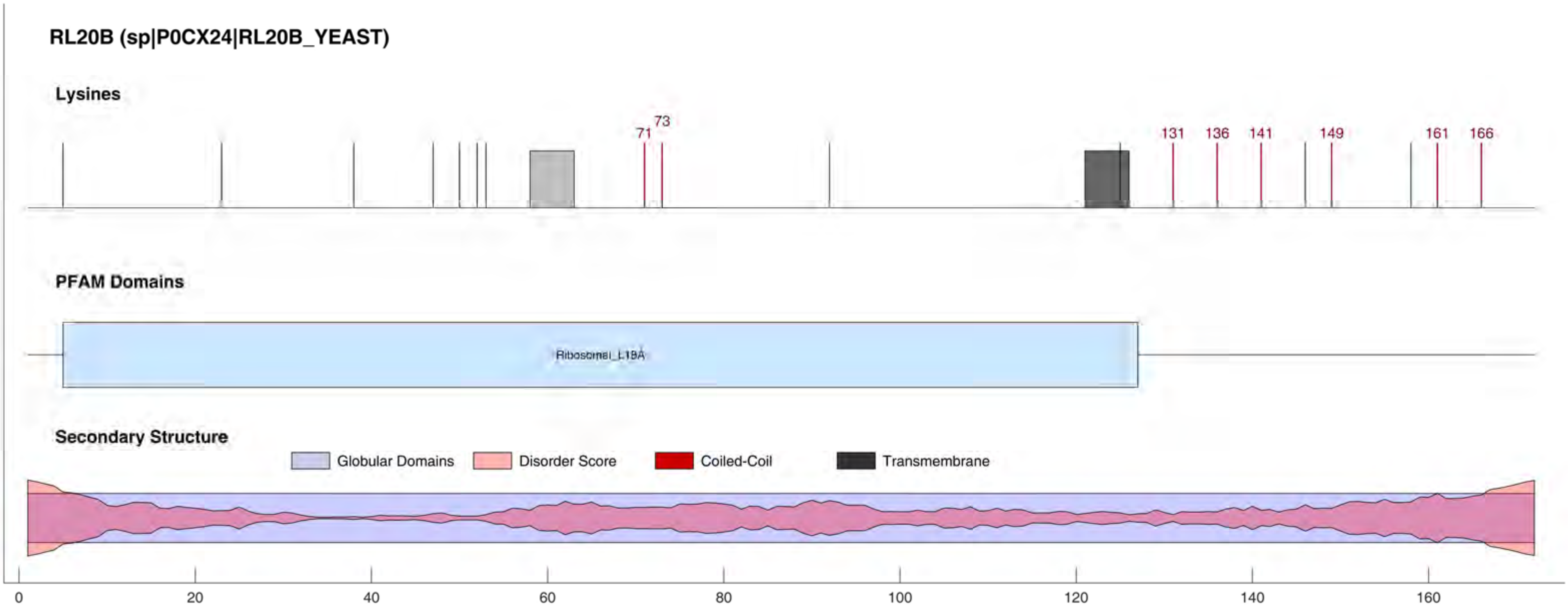

RL20B (sp|P0CX24|RL20B_YEAST)
Lysines
71
73
131
136
141
149
161
166
PFAM Domains
Ribosomal_L18A
Secondary Structure
Globular Domains
Disorder Score
Coiled-Coil
Transmembrane
0
20
40
60
80
100
120
140
160

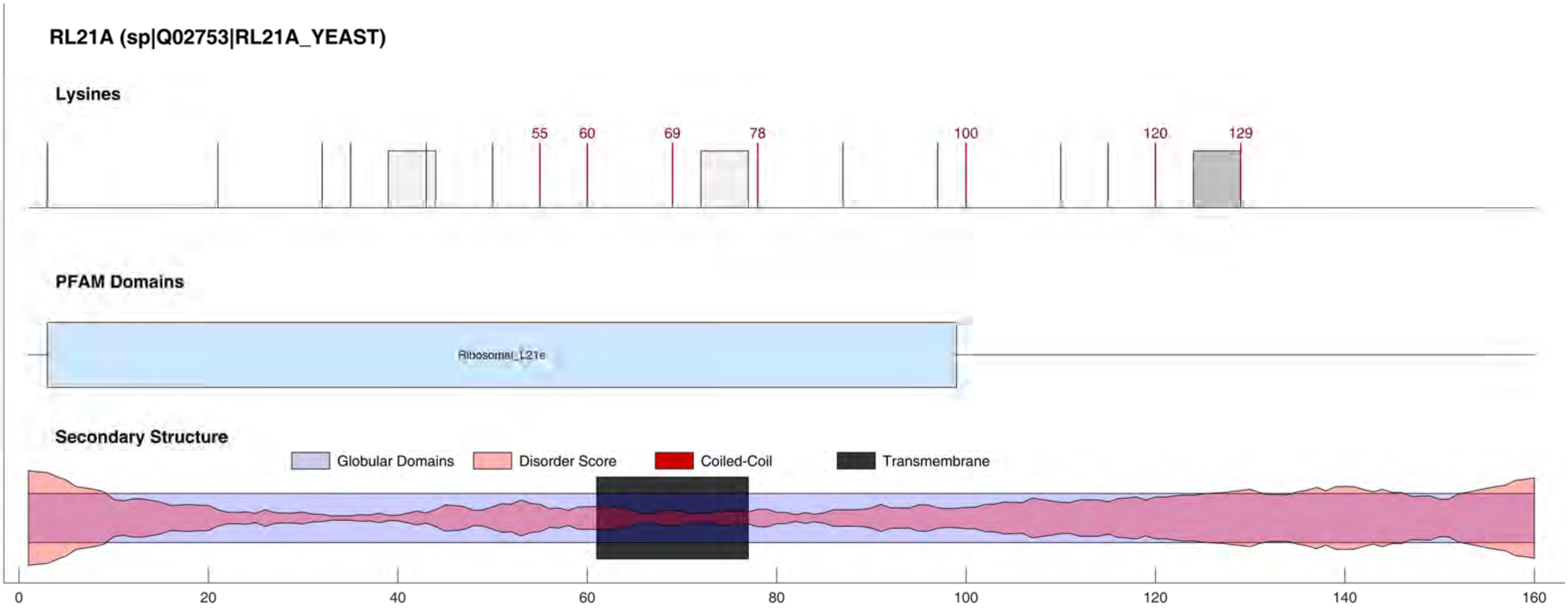
RL21A (sp|Q02753|RL21A_YEAST)
Lysines
55
60
69
78
100
120
129
PFAM Domains
Ribosomal_L21e
Secondary Structure
Globular Domains
Disorder Score
Coiled-Coil
Transmembrane
0
20
40
60
80
100
120
140
160

RL21B (sp|Q12672|RL21B_YEAST)

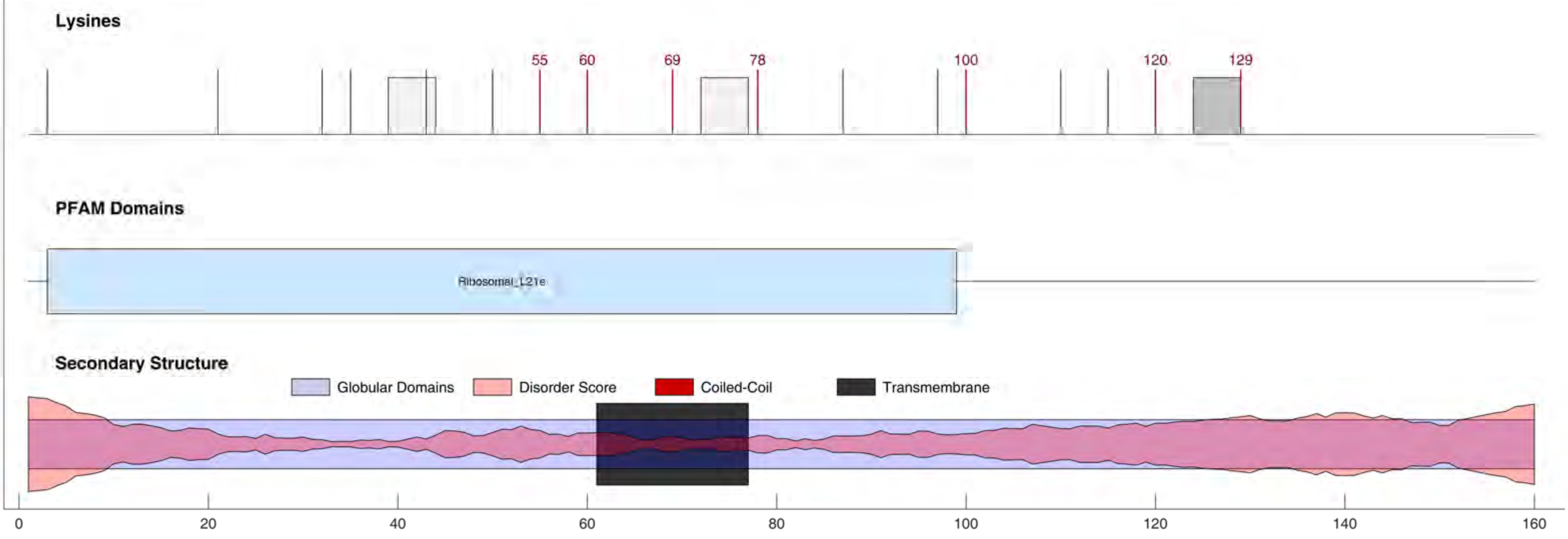

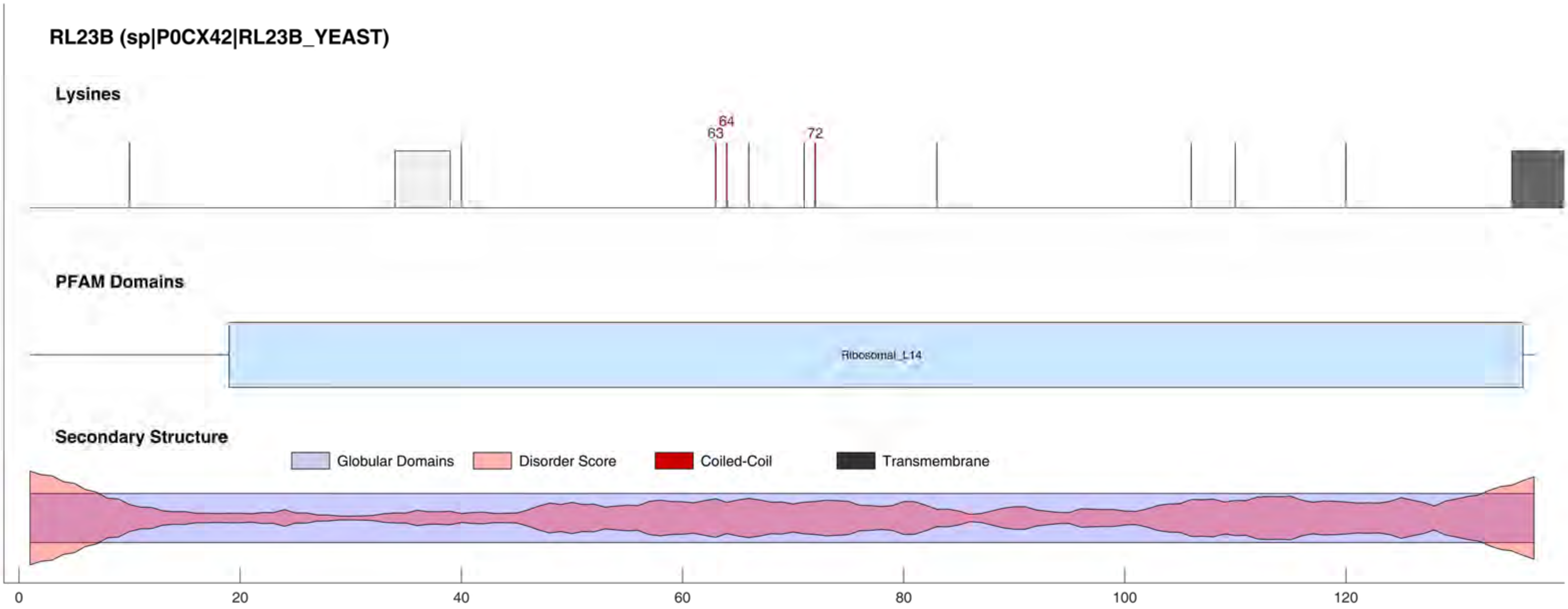
RL23B (sp|P0CX42|RL23B_YEAST)
Lysines
63
64
72
PFAM Domains
Ribosomal_L14
Secondary Structure
Globular Domains
Disorder Score
Coiled-Coil
Transmembrane
0
20
40
60
80
100
120

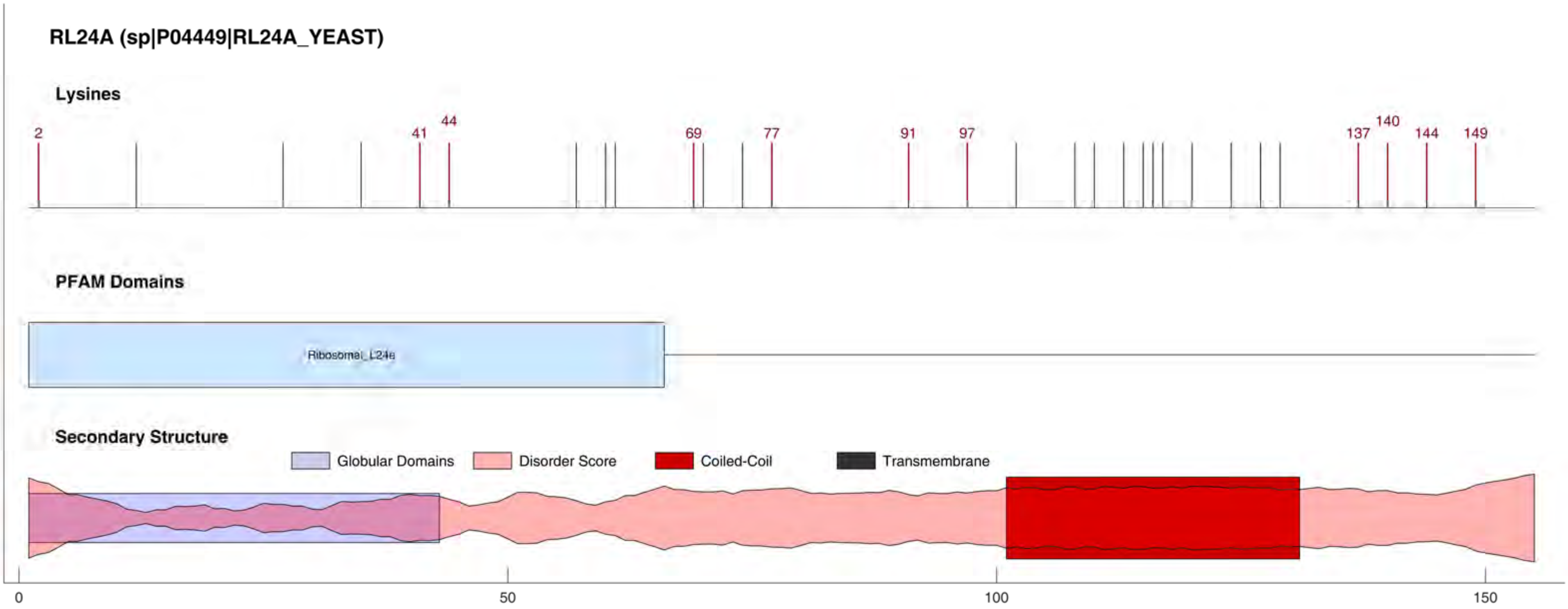

RL24A (sp|P04449|RL24A_YEAST)
Lysines
2
41
44
69
77
91
97
137
140
144
149
PFAM Domains
Ribosomal_L24e
Secondary Structure
Globular Domains
Disorder Score
Coiled-Coil
Transmembrane
0
50
100
150

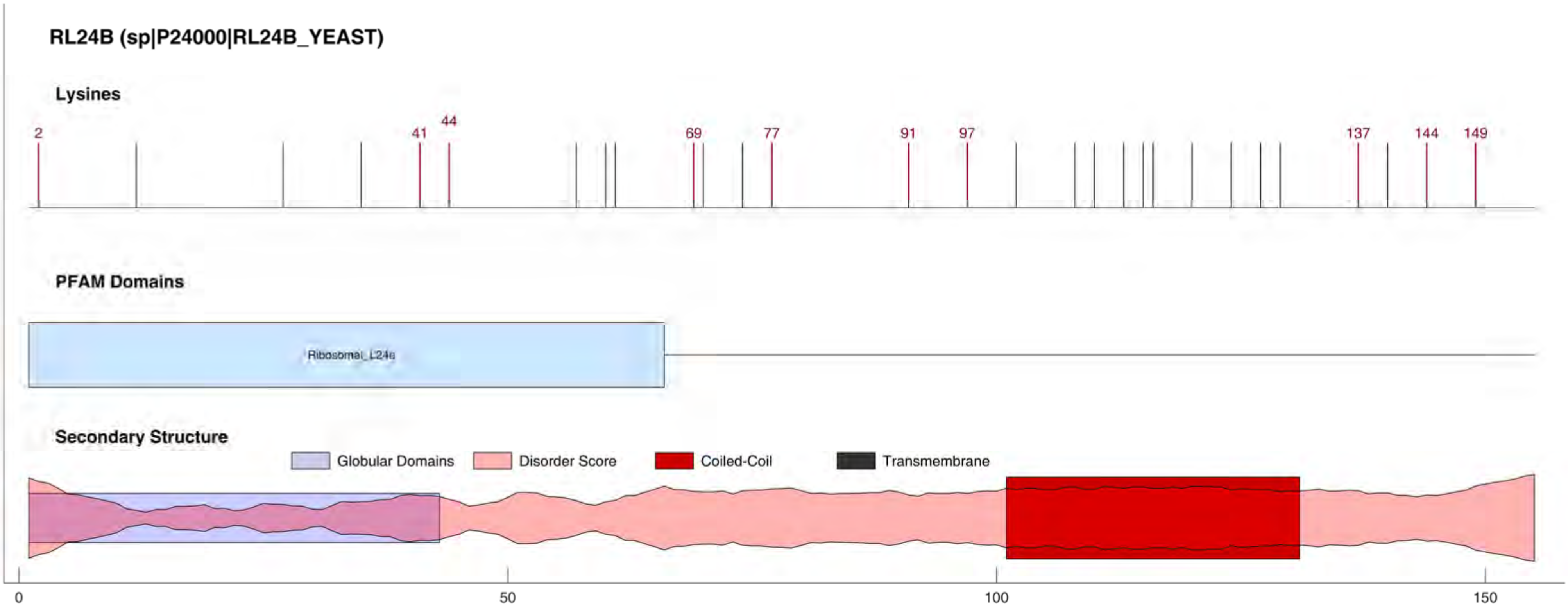
RL24B (sp|P24000|RL24B_YEAST)
Lysines
2
41
44
69
77
91
97
137
144
149
PFAM Domains
Ribosomal_L24e
Secondary Structure
Globular Domains
Disorder Score
Coiled-Coil
Transmembrane
0
50
100
150

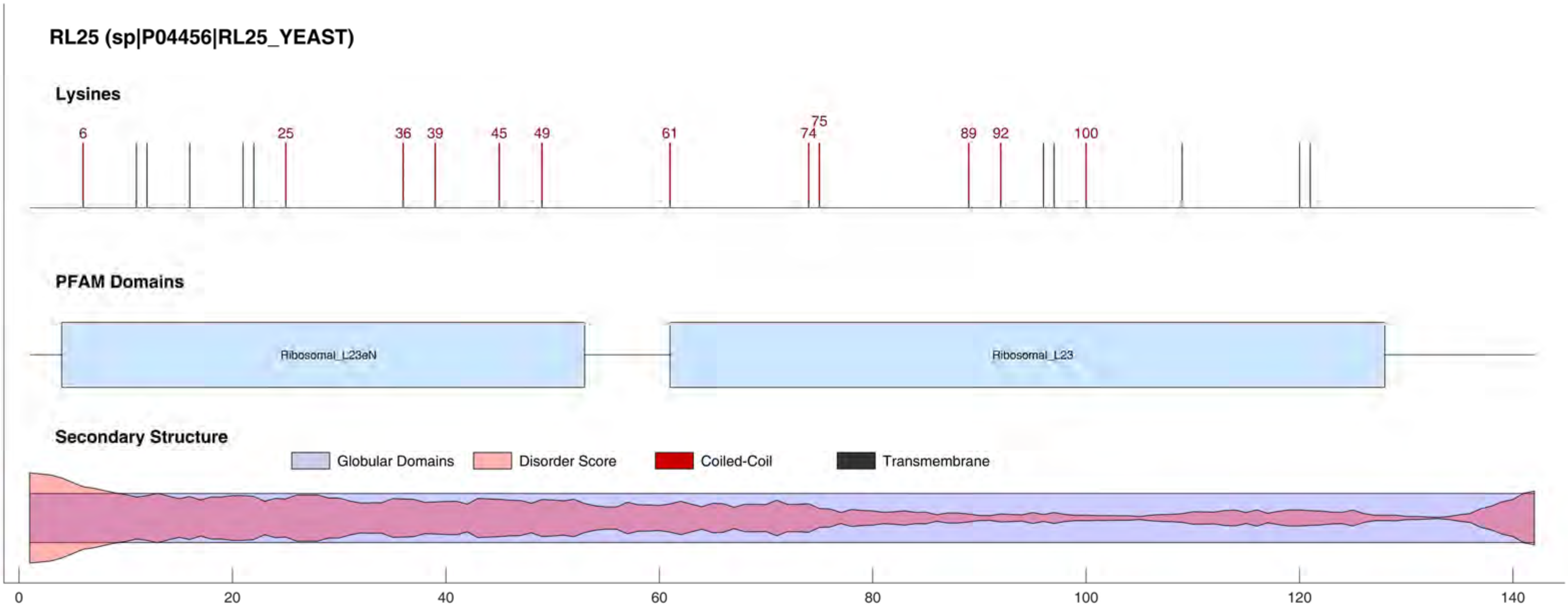
RL25 (sp|P04456|RL25_YEAST)
Lysines
6
25
36
39
45
49
61
74
75
89
92
100
PFAM Domains
Ribosomal_L23eN
Ribosomal_L23
Secondary Structure
Globular Domains
Disorder Score
Coiled-Coil
Transmembrane
0
20
40
60
80
100
120
140

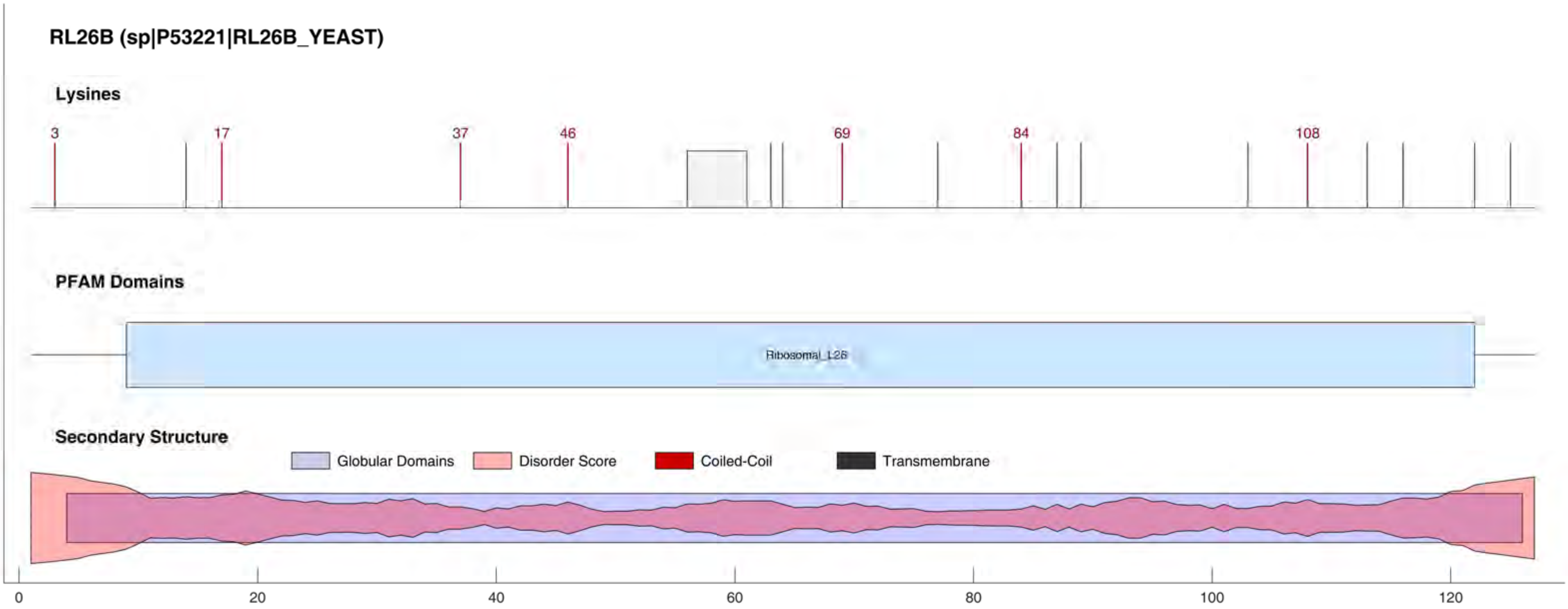
RL26B (sp|P53221|RL26B_YEAST)
Lysines
3
17
37
46
69
84
108
PFAM Domains
Ribosomal_L26
Secondary Structure
Globular Domains
Disorder Score
Coiled-Coil
Transmembrane
0
20
40
60
80
100
120

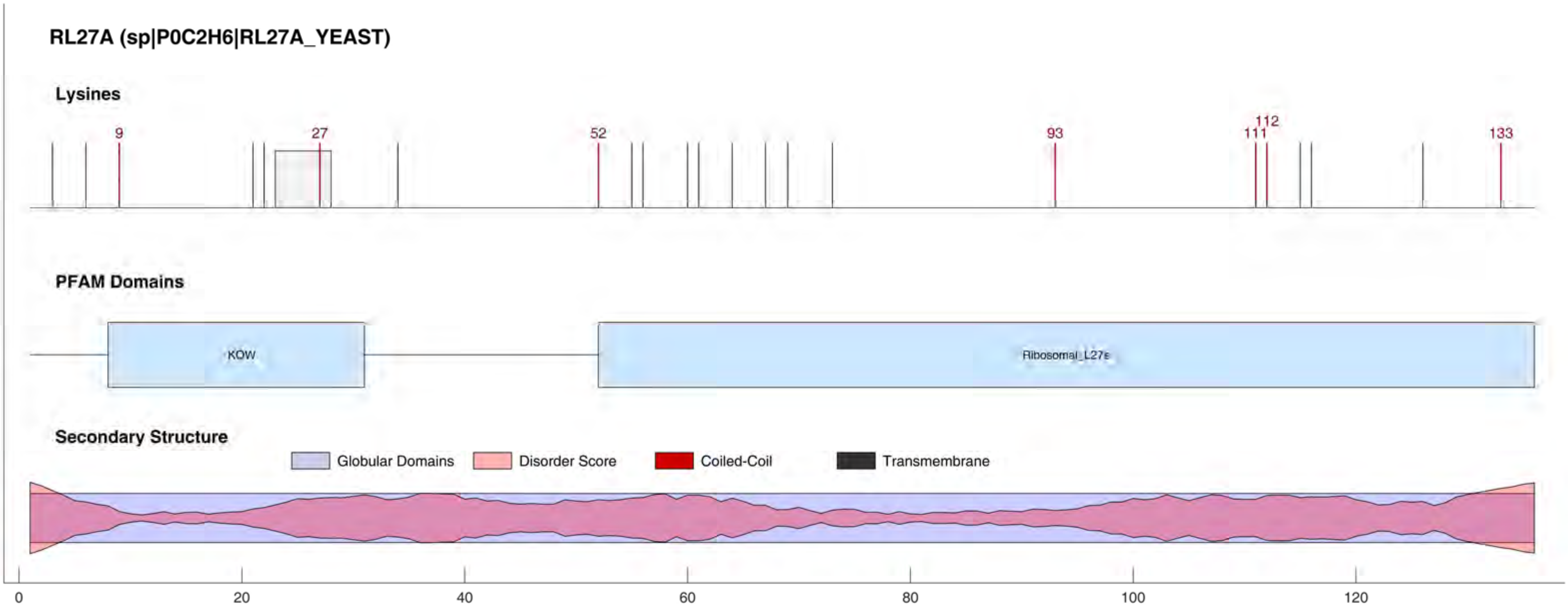
RL27A (sp|P0C2H6|RL27A_YEAST)
Lysines
9
27
52
93
111
112
133
PFAM Domains
KOW
Ribosomal_L27e
Secondary Structure
Globular Domains
Disorder Score
Coiled-Coil
Transmembrane
0
20
40
60
80
100
120

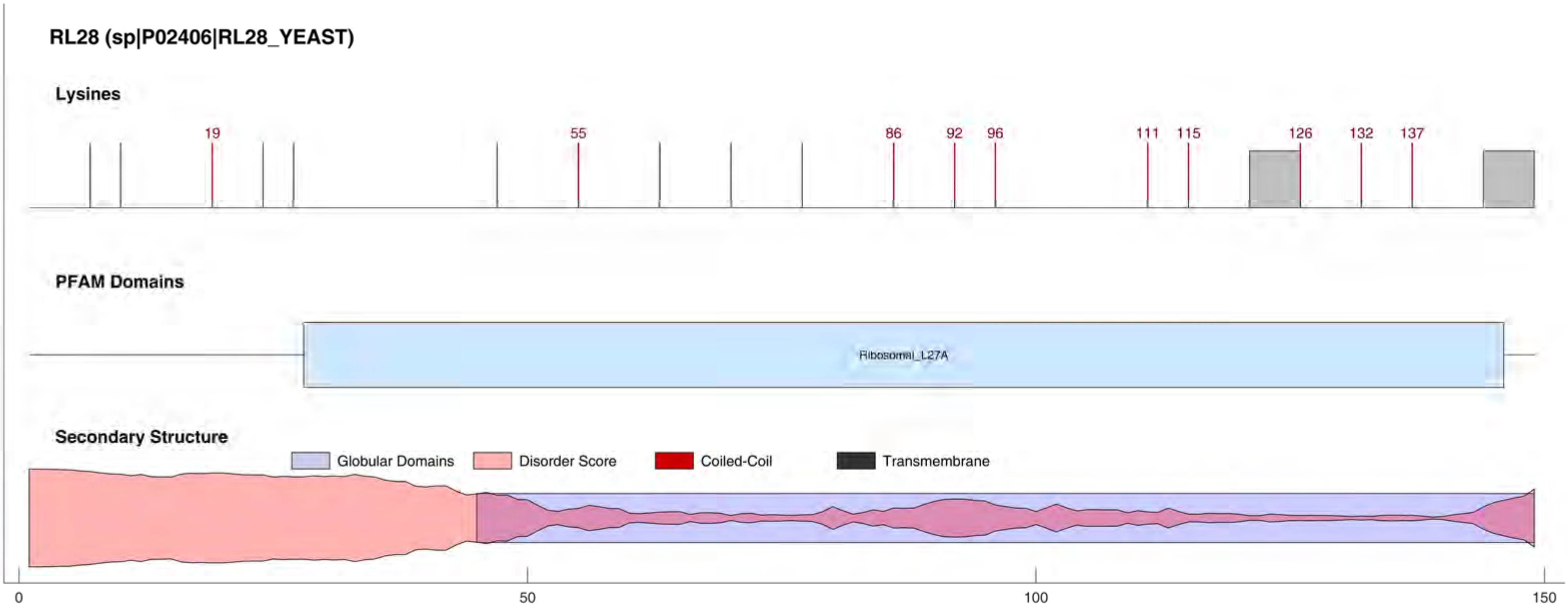
RL28 (sp|P02406|RL28_YEAST)
Lysines
19
55
86
92
96
111
115
126
132
137
PFAM Domains
Ribosomal_L27A
Secondary Structure
Globular Domains
Disorder Score
Coiled-Coil
Transmembrane
0
50
100
150

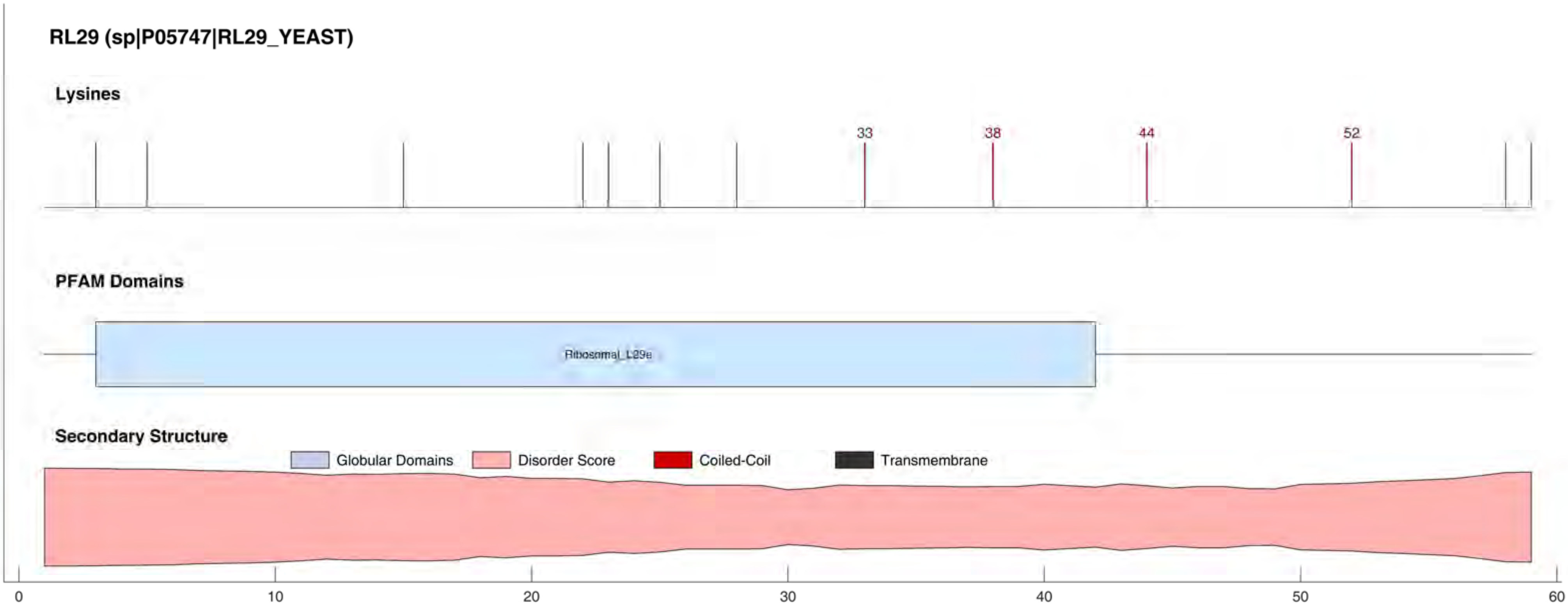
RL29 (sp|P05747|RL29_YEAST)
Lysines
33
38
44
52
PFAM Domains
Ribosomal_L29e
Secondary Structure
Globular Domains
Disorder Score
Coiled-Coil
Transmembrane
0
10
20
30
40
50
60

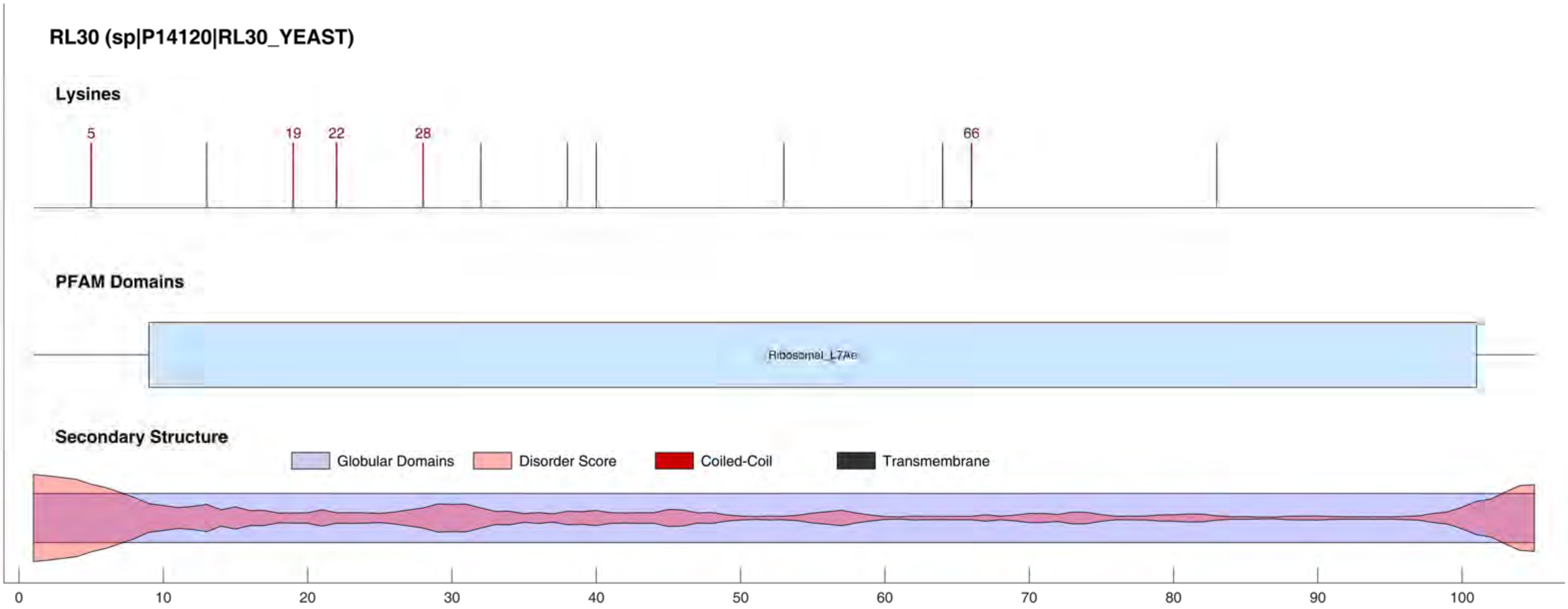
RL30 (sp|P14120|RL30_YEAST)
Lysines
5
19
22
28
66
PFAM Domains
Ribosomal_L7Ae
Secondary Structure
Globular Domains
Disorder Score
Coiled-Coil
Transmembrane
0
10
20
30
40
50
60
70
80
90
100

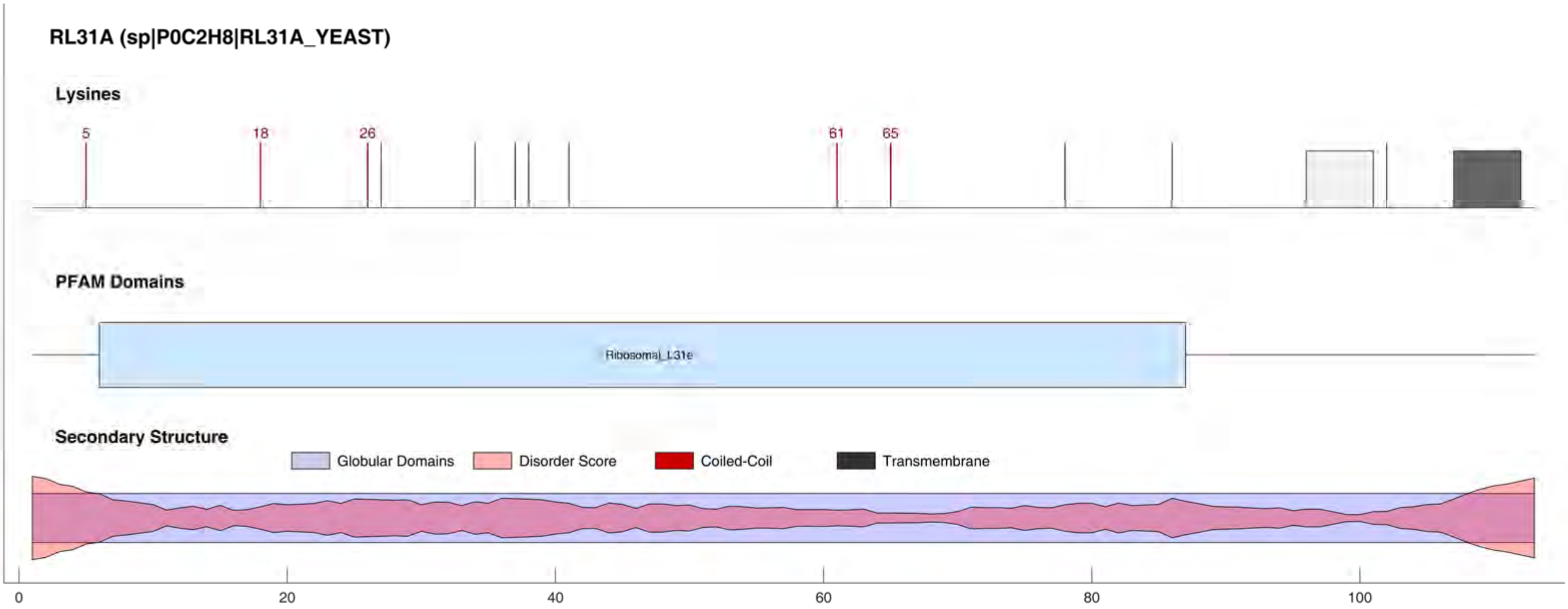
RL31A (sp|P0C2H8|RL31A_YEAST)
Lysines
5
18
26
61
65
PFAM Domains
Ribosomal_L31e
Secondary Structure
Globular Domains
Disorder Score
Coiled-Coil
Transmembrane
0
20
40
60
80
100

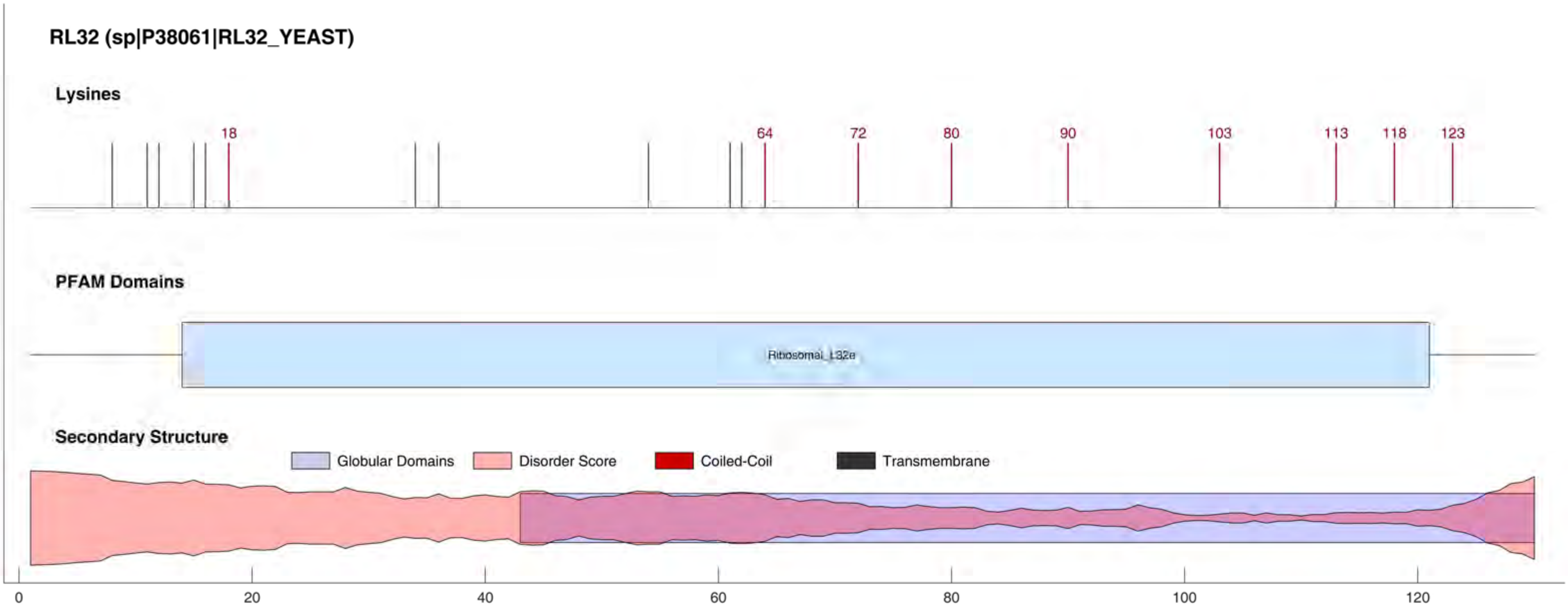
RL32 (sp|P38061|RL32_YEAST)
Lysines
18
64
72
80
90
103
113
118
123
PFAM Domains
Ribosomal_L32e
Secondary Structure
Globular Domains
Disorder Score
Coiled-Coil
Transmembrane
0
20
40
60
80
100
120

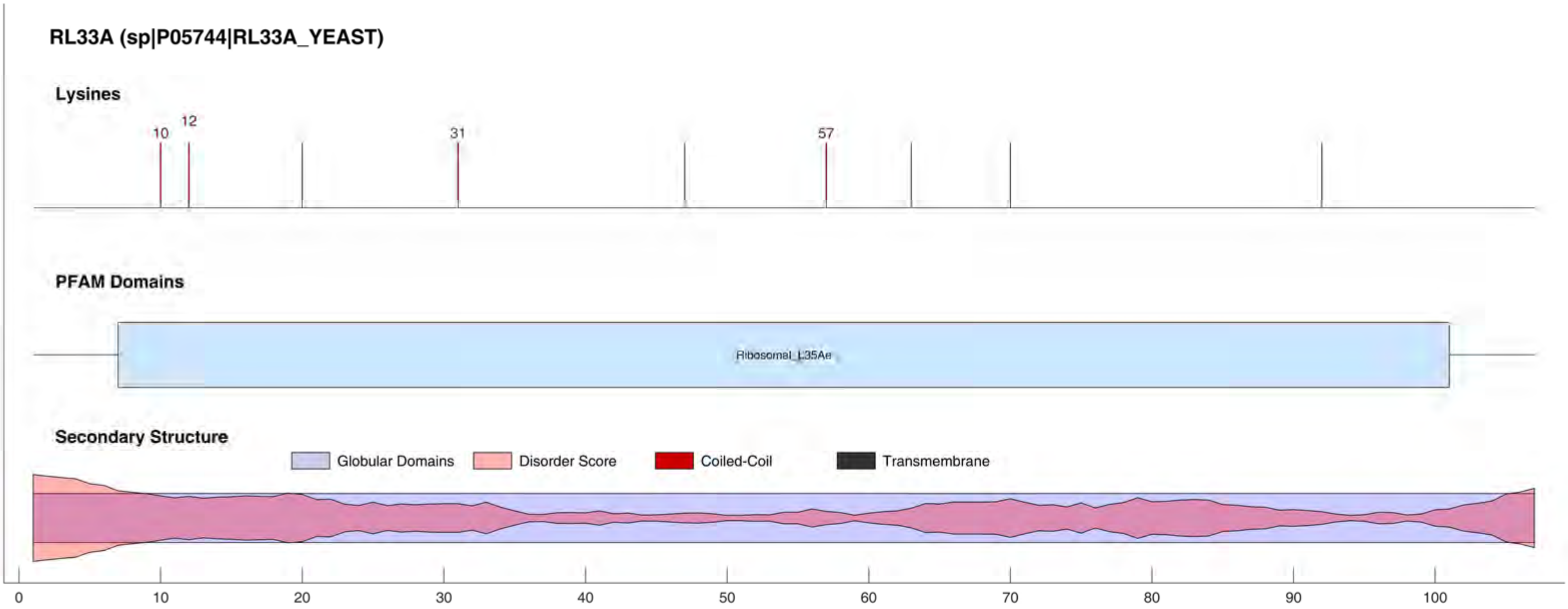
RL33A (sp|P05744|RL33A_YEAST)
Lysines
10
12
31
57
PFAM Domains
Ribosomal_L35Ae
Secondary Structure
Globular Domains
Disorder Score
Coiled-Coil
Transmembrane
0
10
20
30
40
50
60
70
80
90
100

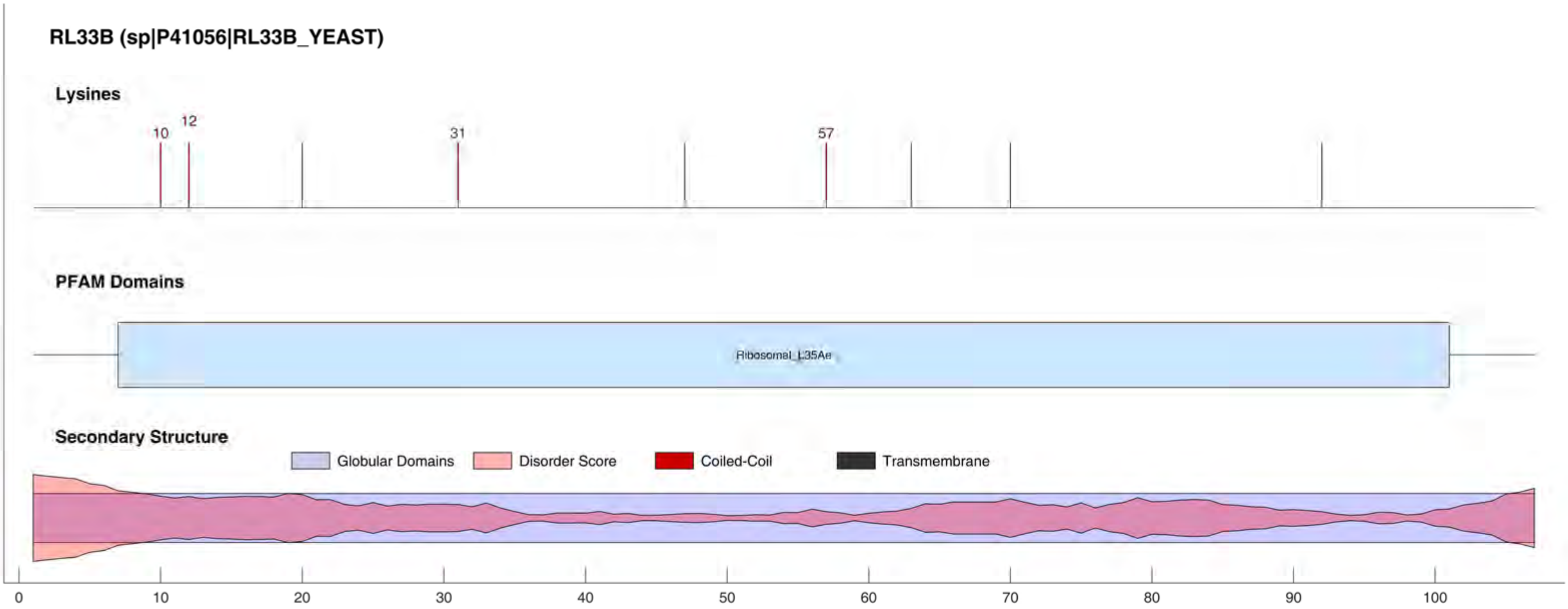

RL33B (sp|P41056|RL33B_YEAST)
Lysines
10
12
31
57
PFAM Domains
Ribosomal_L35Ae
Secondary Structure
Globular Domains
Disorder Score
Coiled-Coil
Transmembrane
0
10
20
30
40
50
60
70
80
90
100

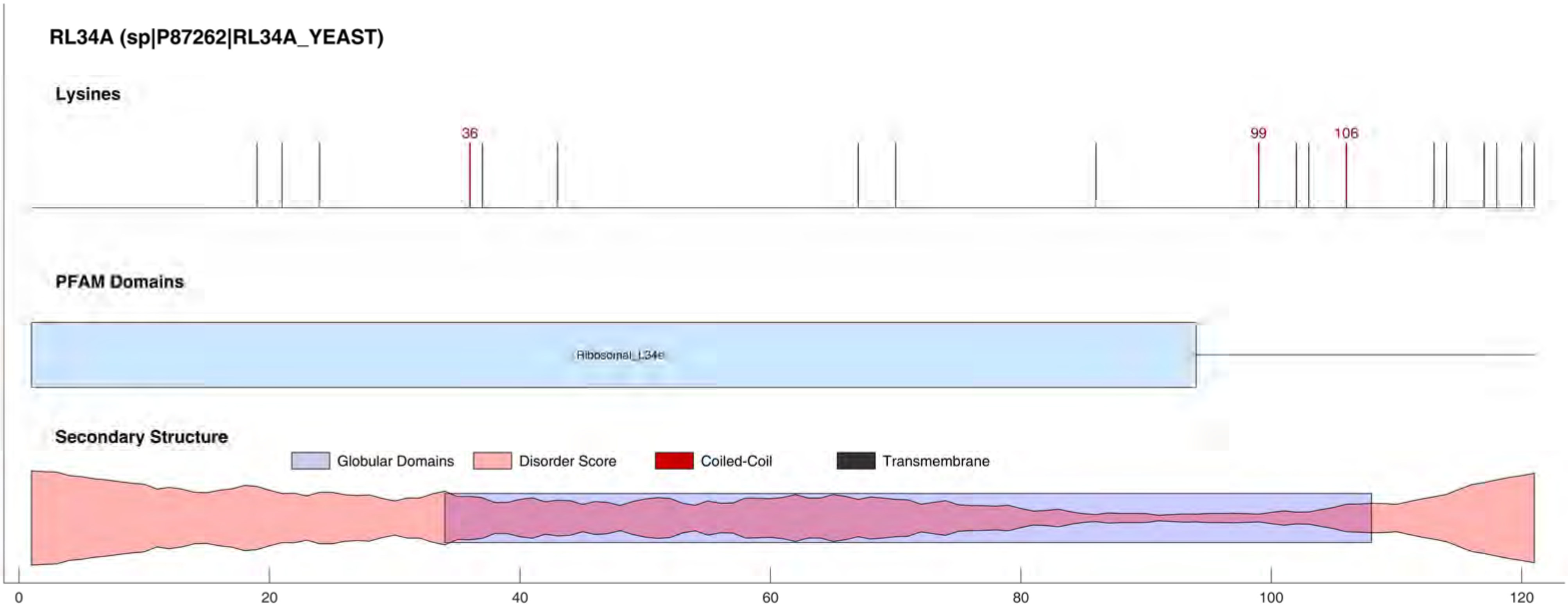
RL34A (sp|P87262|RL34A_YEAST)
Lysines
36
99
106
PFAM Domains
Ribosomal_L34e
Secondary Structure
Globular Domains
Disorder Score
Coiled-Coil
Transmembrane
0
20
40
60
80
100
120

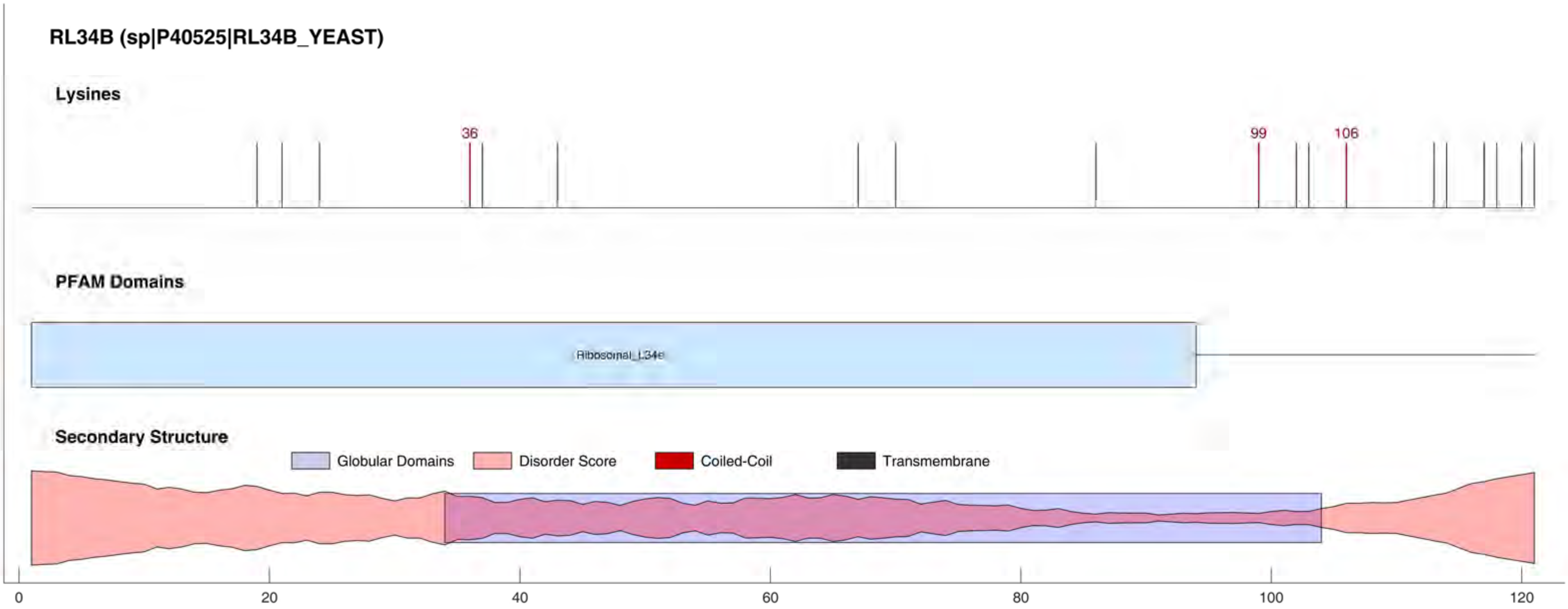
RL34B (sp|P40525|RL34B_YEAST)
Lysines
36
99
106
PFAM Domains
Ribosomal_L34e
Secondary Structure
Globular Domains
Disorder Score
Coiled-Coil
Transmembrane
0
20
40
60
80
100
120

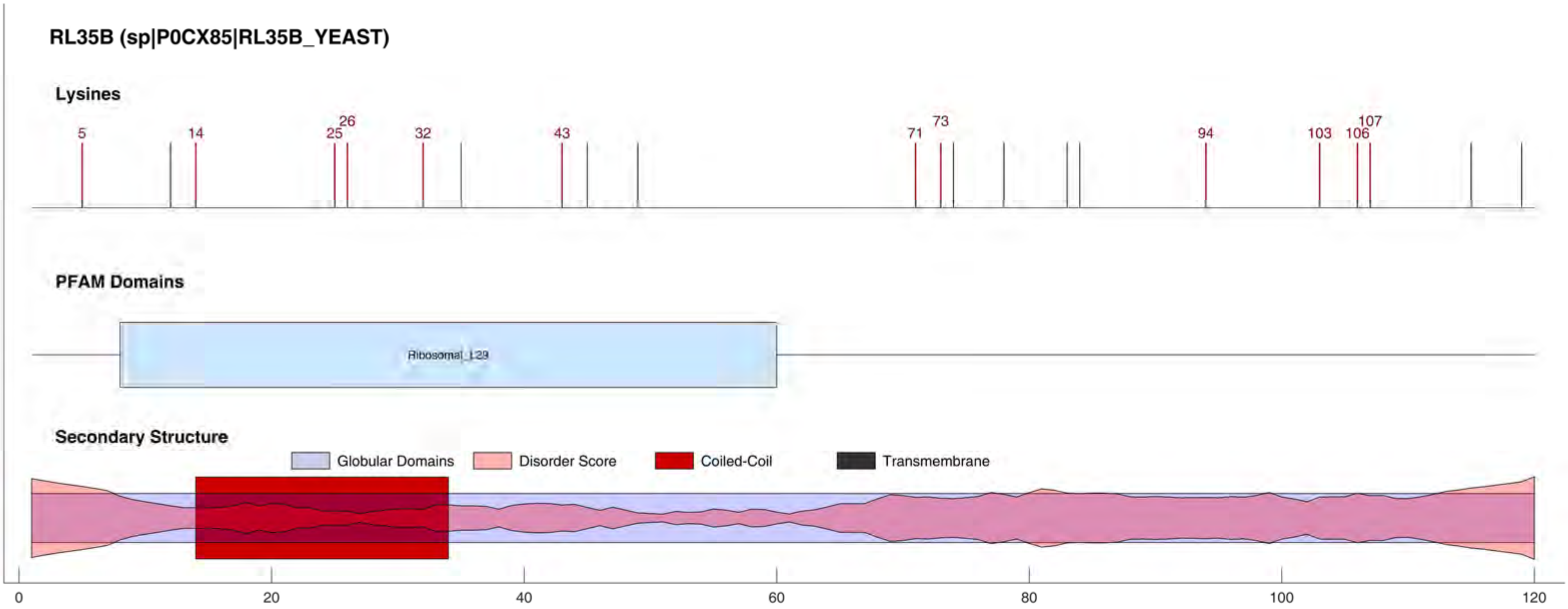

RL35B (sp|P0CX85|RL35B_YEAST)
Lysines
5
14
25
26
32
43
71
73
94
103
106
107
PFAM Domains
Ribosomal_L29
Secondary Structure
Globular Domains
Disorder Score
Coiled-Coil
Transmembrane
0
20
40
60
80
100
120

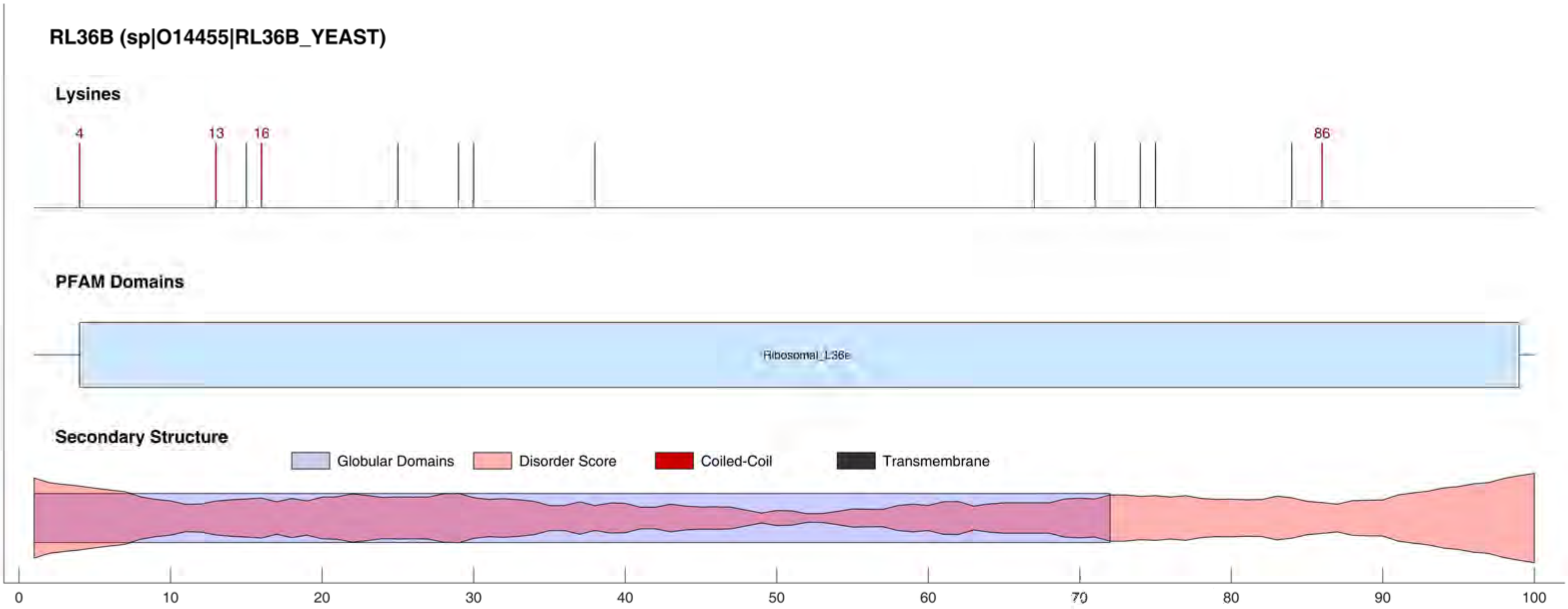

RL36B (sp|O14455|RL36B_YEAST)
Lysines
4
13
16
86
PFAM Domains
Ribosomal_L36e
Secondary Structure
Globular Domains
Disorder Score
Coiled-Coil
Transmembrane
0
10
20
30
40
50
60
70
80
90
100

#### RL37A (sp|P49166|RL37A_YEAST)

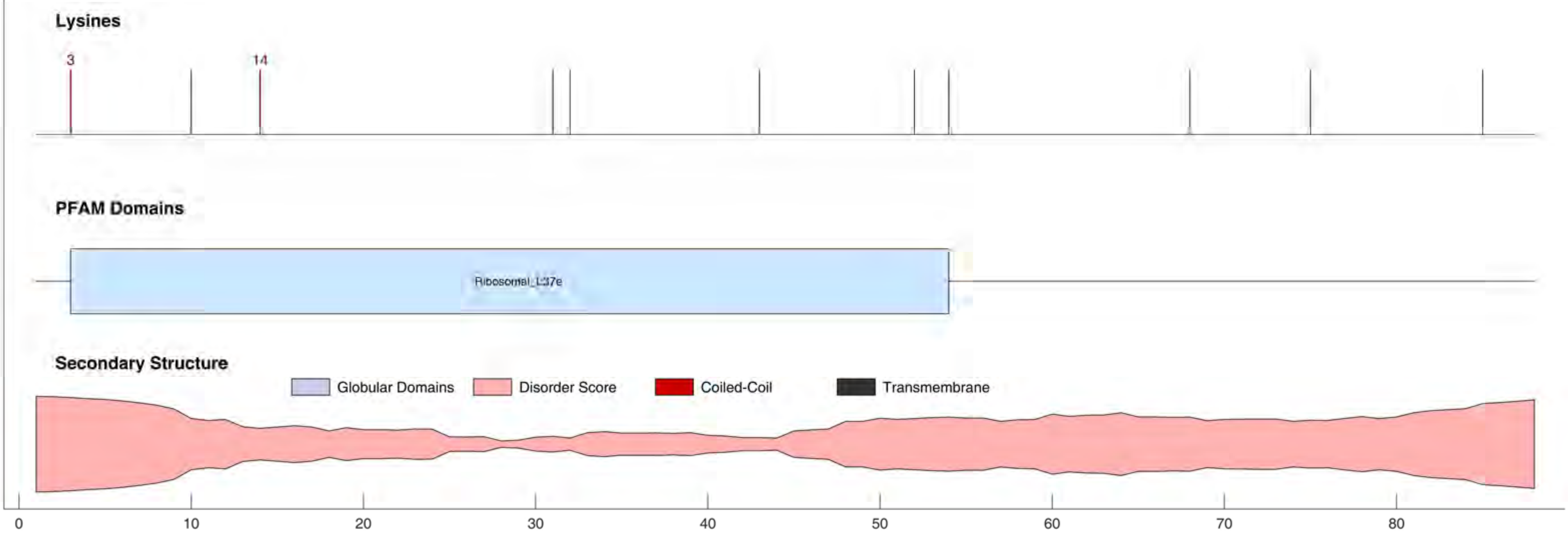

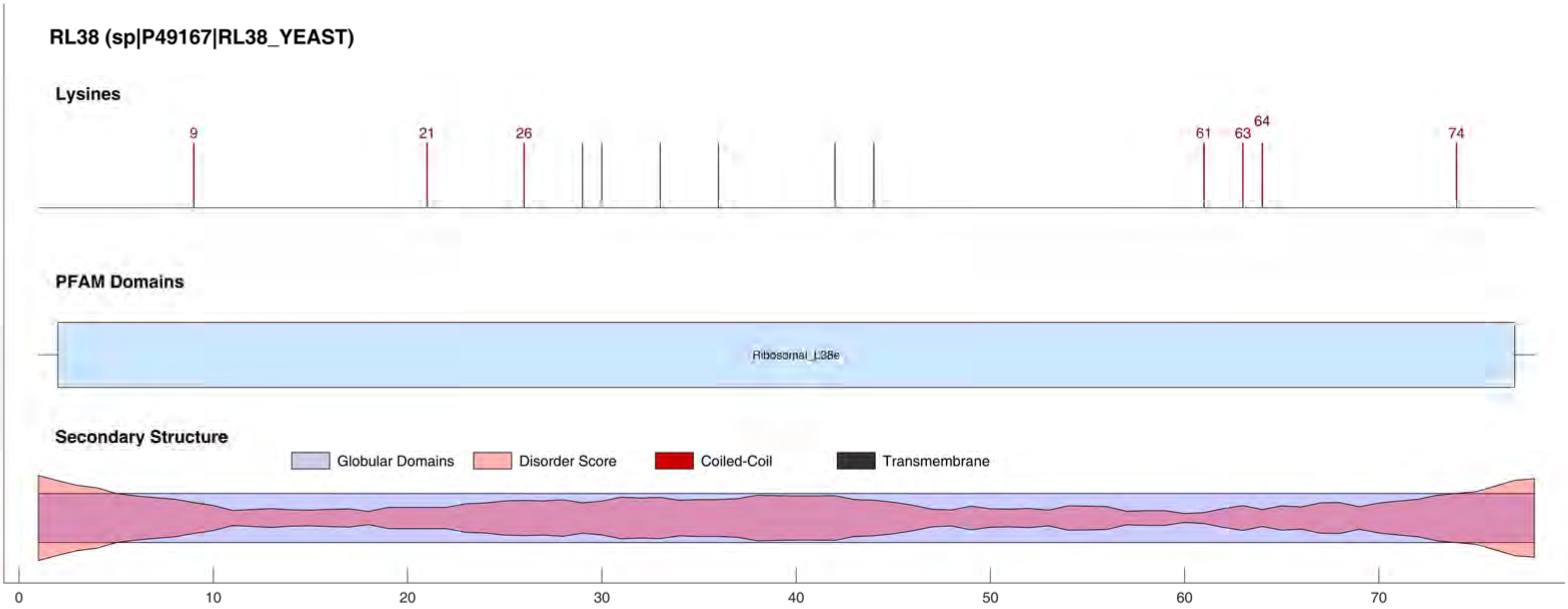

RL38 (sp|P49167|RL38_YEAST)
Lysines
9
21
26
61
63
64
74
PFAM Domains
Ribosomal_L38e
Secondary Structure
Globular Domains
Disorder Score
Coiled-Coil
Transmembrane
0
10
20
30
40
50
60
70

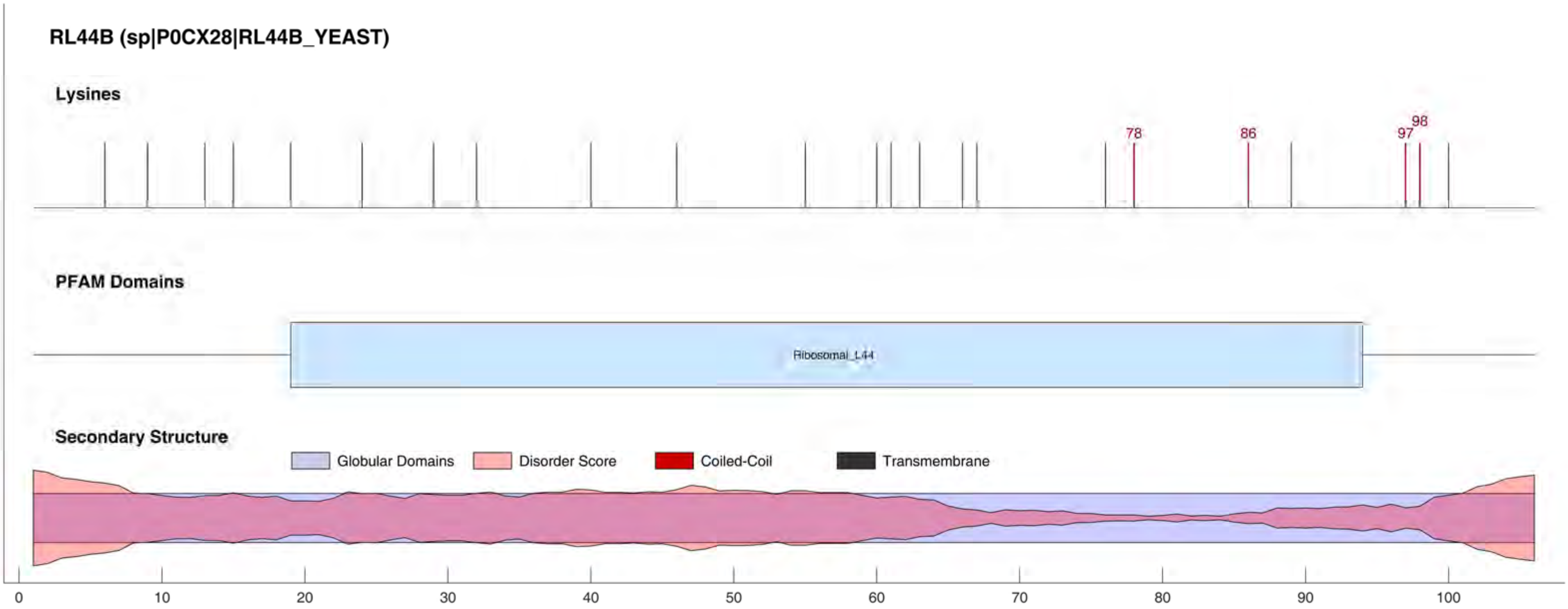
RL44B (sp|P0CX28|RL44B_YEAST)
Lysines
78
86
97
98
PFAM Domains
Ribosomal_L44
Secondary Structure
Globular Domains
Disorder Score
Coiled-Coil
Transmembrane
0
10
20
30
40
50
60
70
80
90
100

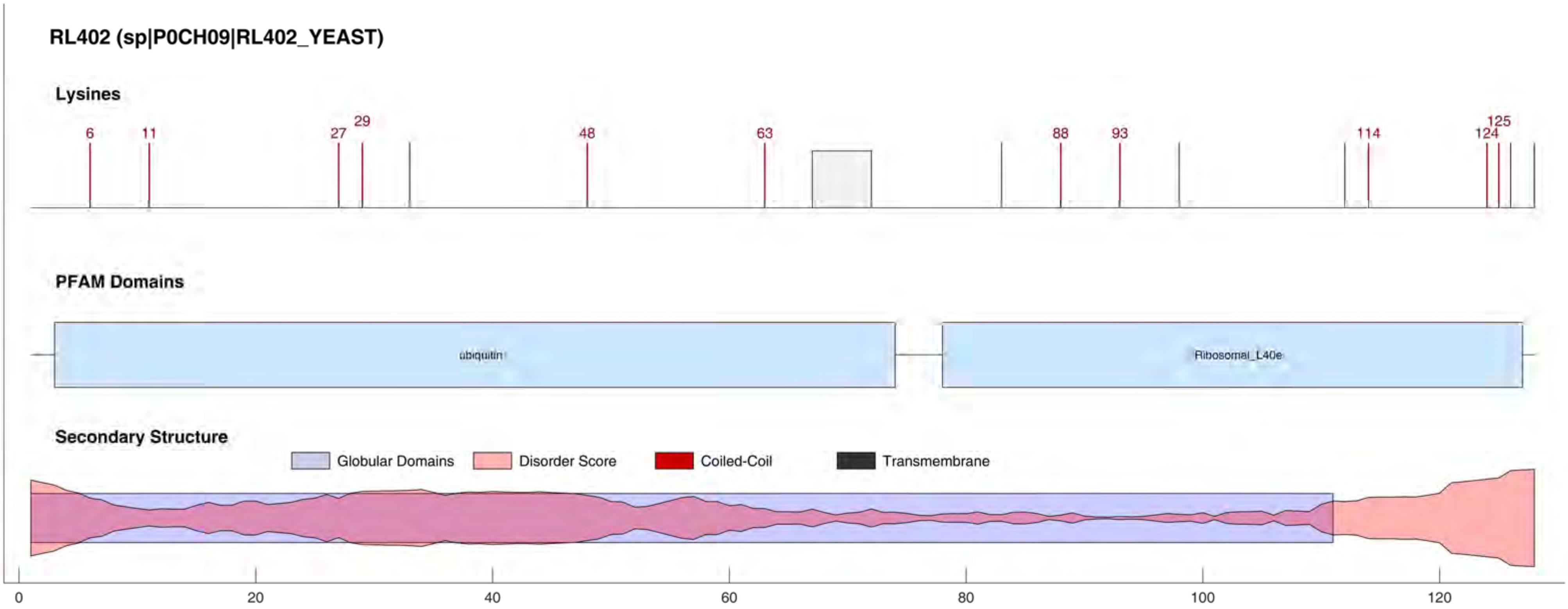

RL402 (sp|P0CH09|RL402_YEAST)
Lysines
6
11
27
29
48
63
88
93
114
124
125
PFAM Domains
ubiquitin
Ribosomal_L40e
Secondary Structure
Globular Domains
Disorder Score
Coiled-Coil
Transmembrane
0
20
40
60
80
100
120

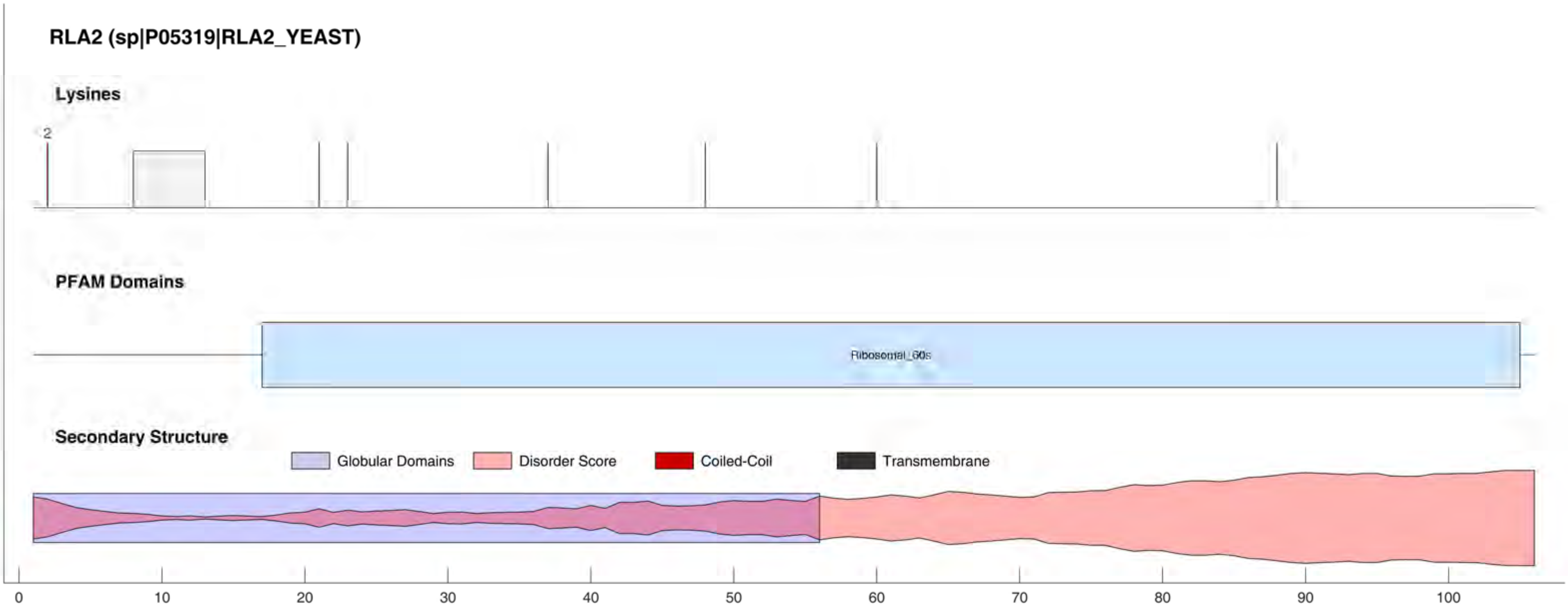
RLA2 (sp|P05319|RLA2_YEAST)
Lysines
2
PFAM Domains
Ribosomal_60s
Secondary Structure
Globular Domains
Disorder Score
Coiled-Coil
Transmembrane
0
10
20
30
40
50
60
70
80
90
100

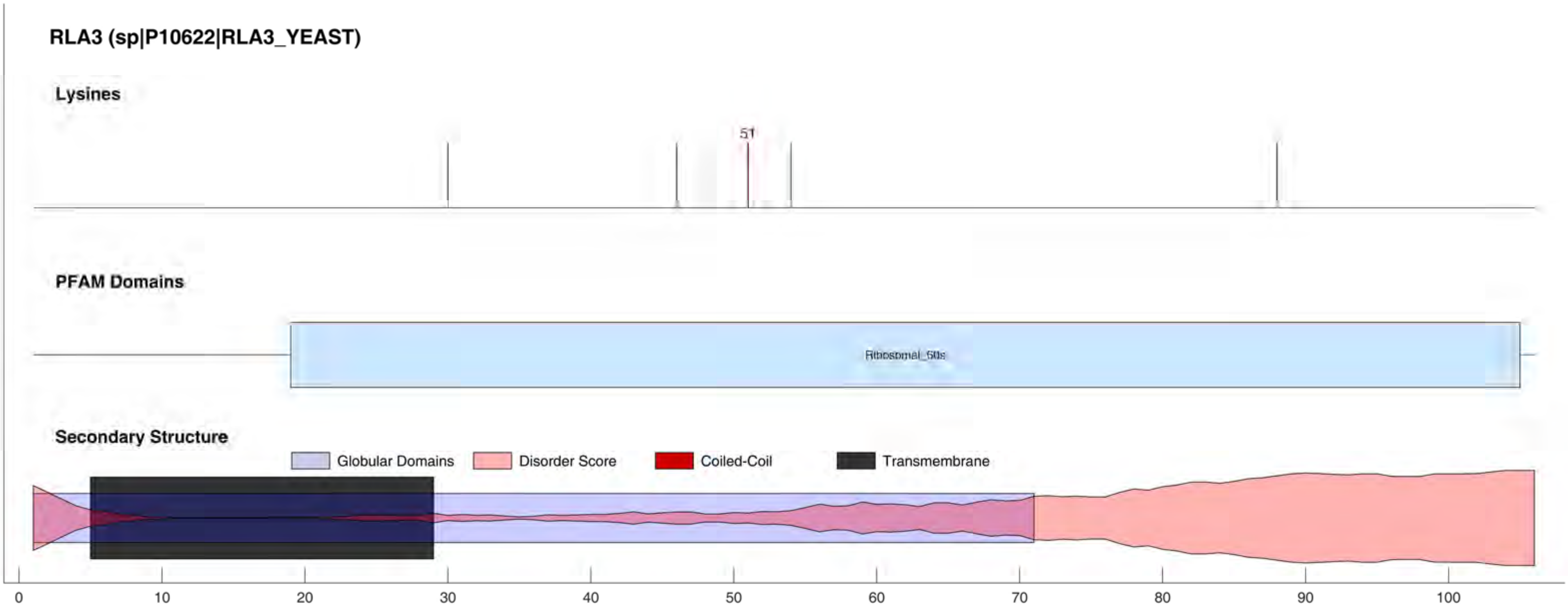
RLA3 (sp|P10622|RLA3_YEAST)
Lysines
51
PFAM Domains
Ribosomal_60s
Secondary Structure
Globular Domains
Disorder Score
Coiled-Coil
Transmembrane
0
10
20
30
40
50
60
70
80
90
100

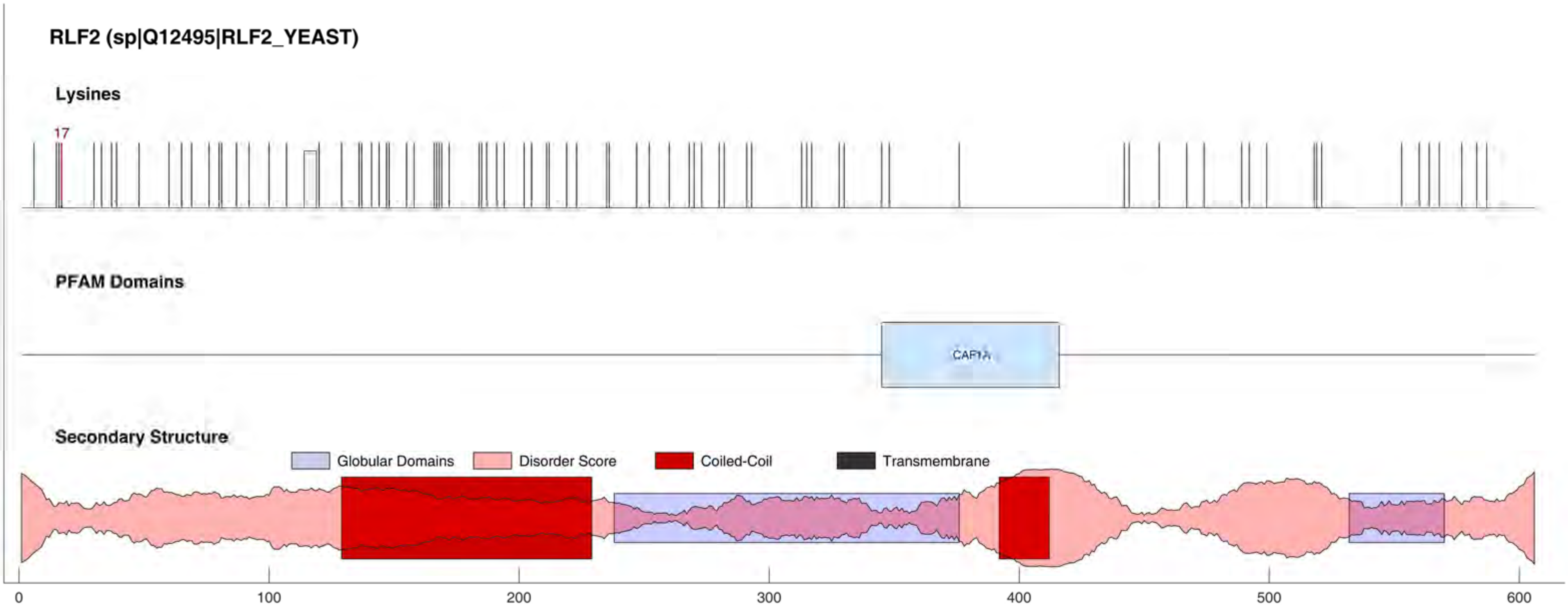
RLF2 (sp|Q12495|RLF2_YEAST)
Lysines
17
PFAM Domains
CAF1A
Secondary Structure
Globular Domains
Disorder Score
Coiled-Coil
Transmembrane
0
100
200
300
400
500
600

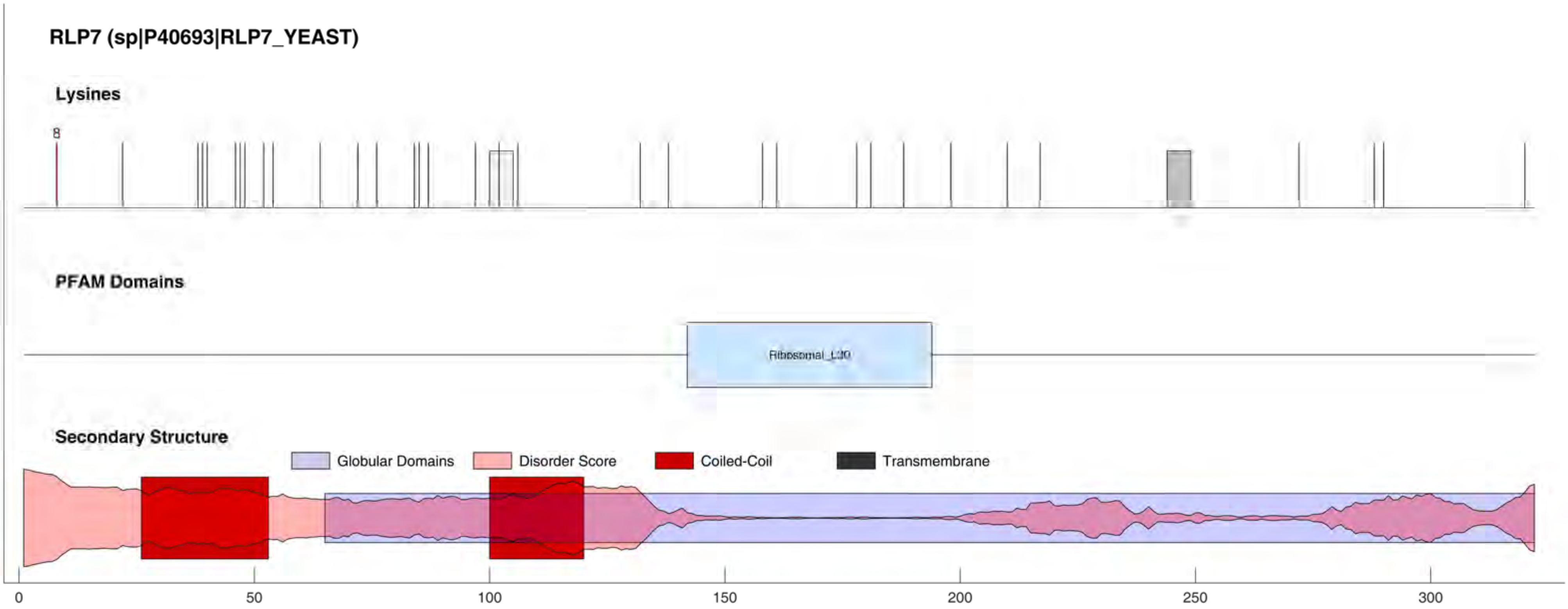
RLP7 (sp|P40693|RLP7_YEAST)
Lysines
8
PFAM Domains
Ribosomal_L30
Secondary Structure
Globular Domains
Disorder Score
Coiled-Coil
Transmembrane
0
50
100
150
200
250
300

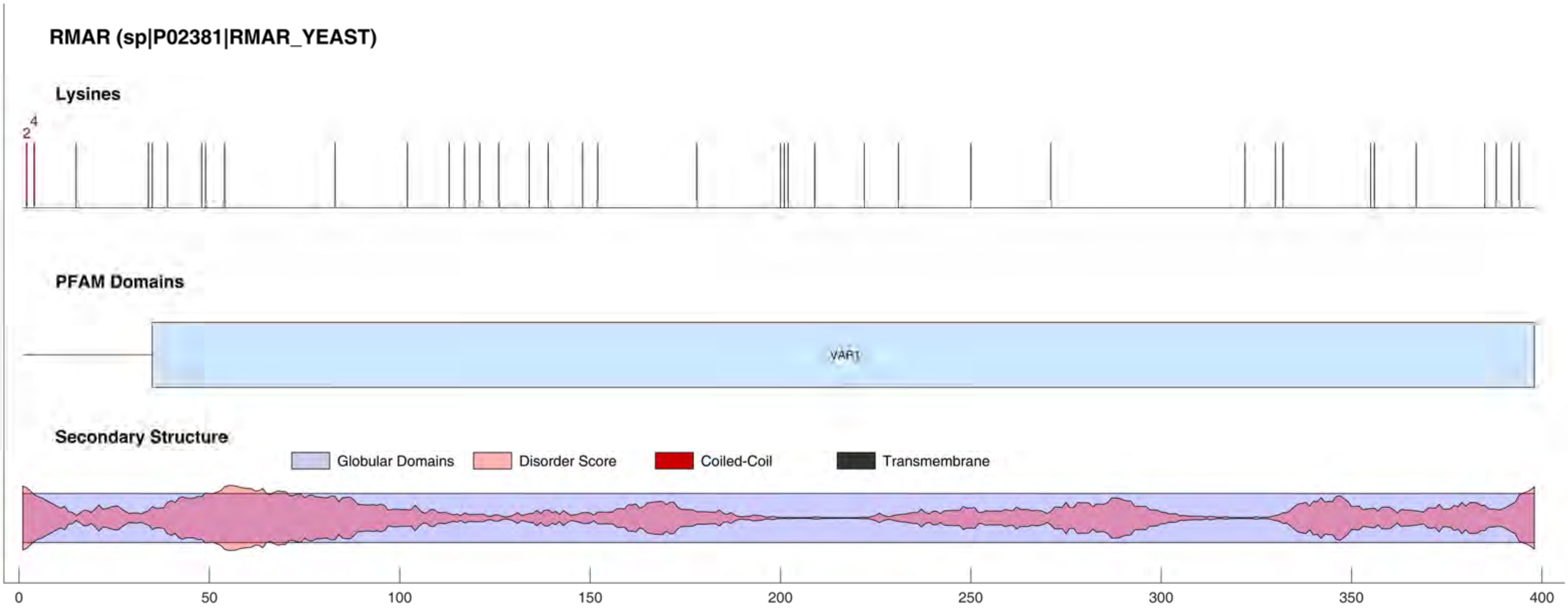
RMAR (sp|P02381|RMAR_YEAST)
Lysines
2
4
PFAM Domains
VAR1
Secondary Structure
Globular Domains
Disorder Score
Coiled-Coil
Transmembrane
0
50
100
150
200
250
300
350
400

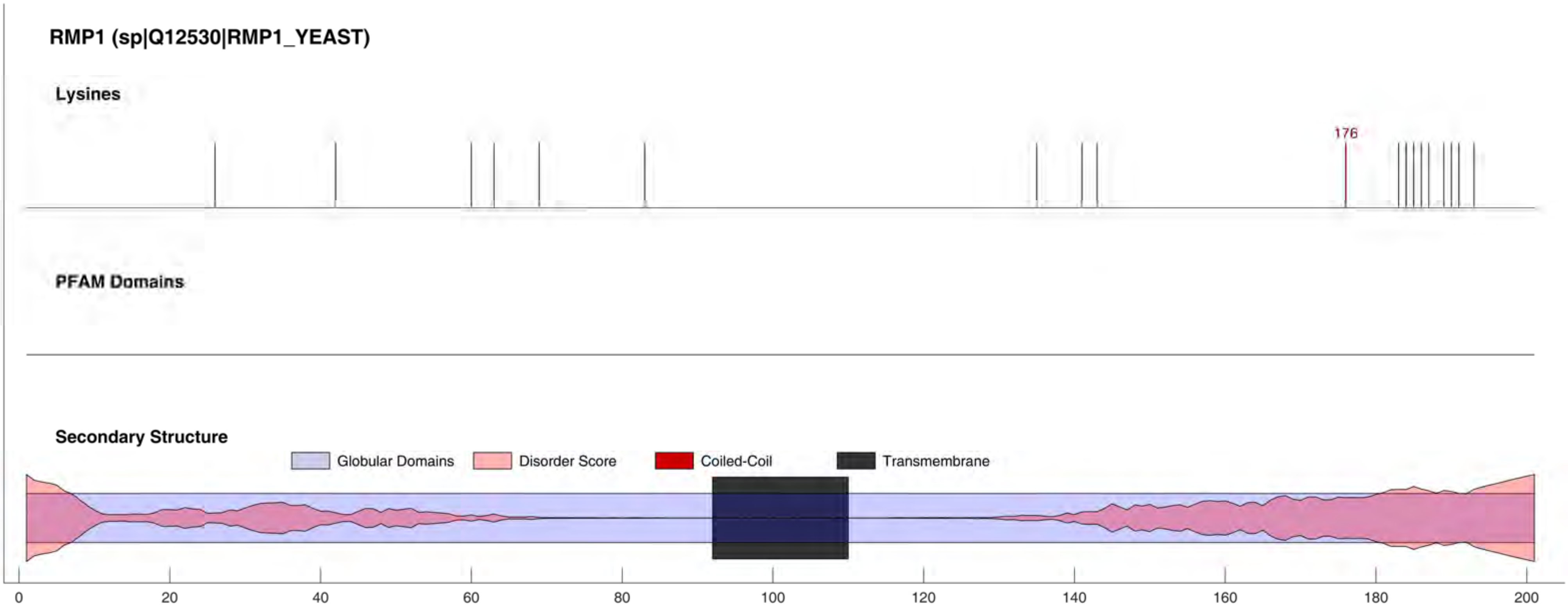
RMP1 (sp|Q12530|RMP1_YEAST)
Lysines
176
PFAM Domains
Secondary Structure
Globular Domains
Disorder Score
Coiled-Coil
Transmembrane
0
20
40
60
80
100
120
140
160
180
200

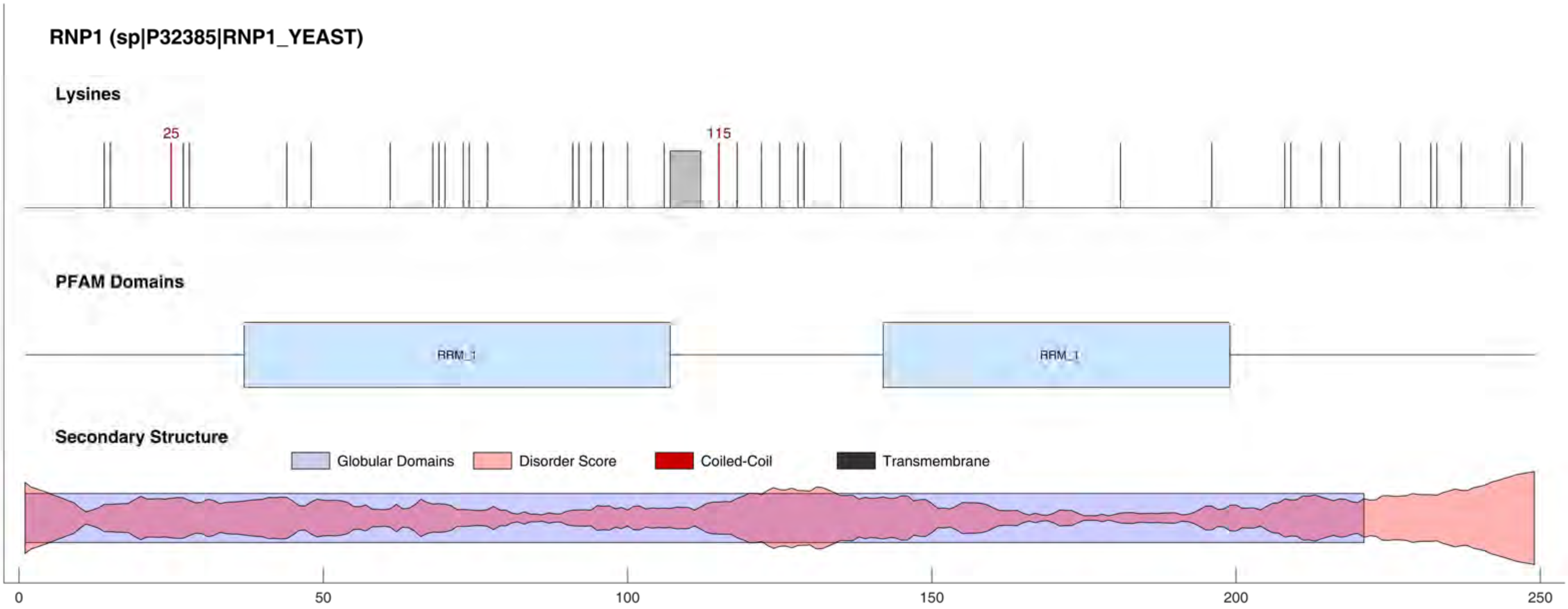
RNP1 (sp|P32385|RNP1_YEAST)
Lysines
25
115
PFAM Domains
RRM_1
RRM_1
Secondary Structure
Globular Domains
Disorder Score
Coiled-Coil
Transmembrane
0
50
100
150
200
250

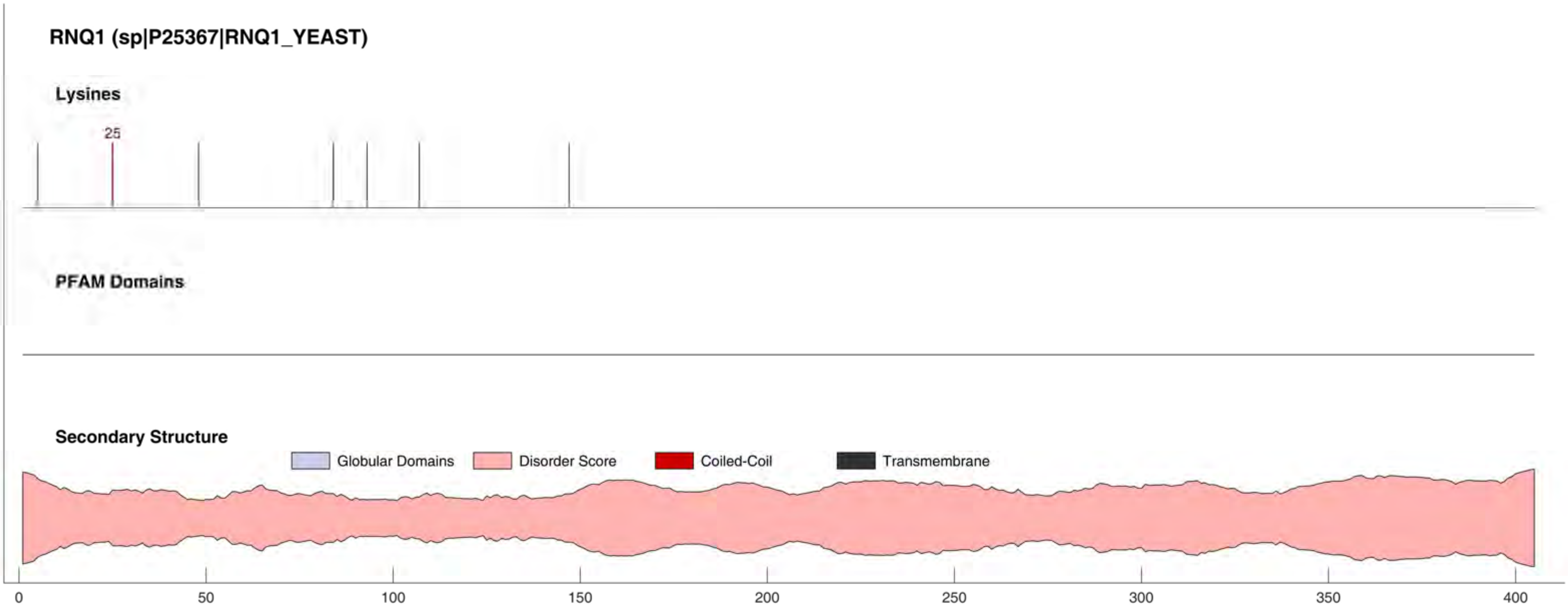
RNQ1 (sp|P25367|RNQ1_YEAST)
Lysines
25
PFAM Domains
Secondary Structure
Globular Domains
Disorder Score
Coiled-Coil
Transmembrane
0
50
100
150
200
250
300
350
400

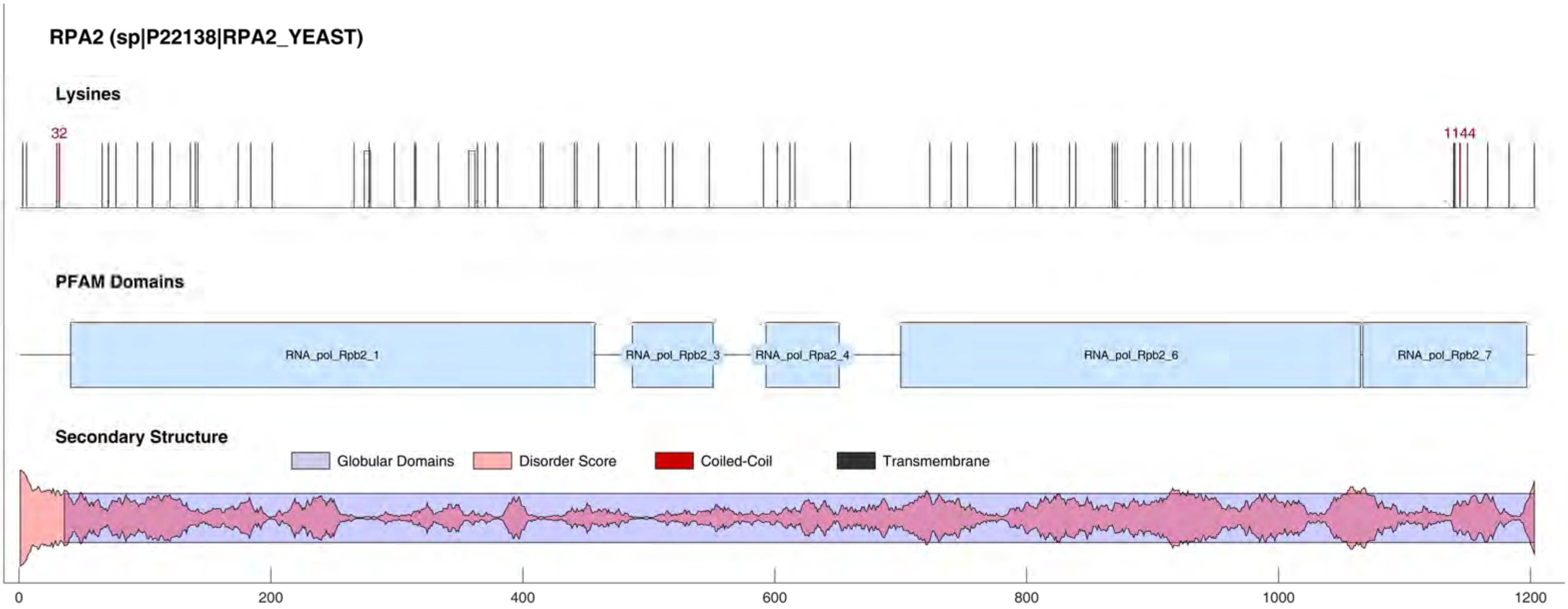
RPA2 (sp|P22138|RPA2_YEAST)
Lysines
32
1144
PFAM Domains
RNA_pol_Rpb2_1
RNA_pol_Rpb2_3
RNA_pol_Rpa2_4
RNA_pol_Rpb2_6
RNA_pol_Rpb2_7
Secondary Structure
Globular Domains
Disorder Score
Coiled-Coil
Transmembrane
0
200
400
600
800
1000
1200

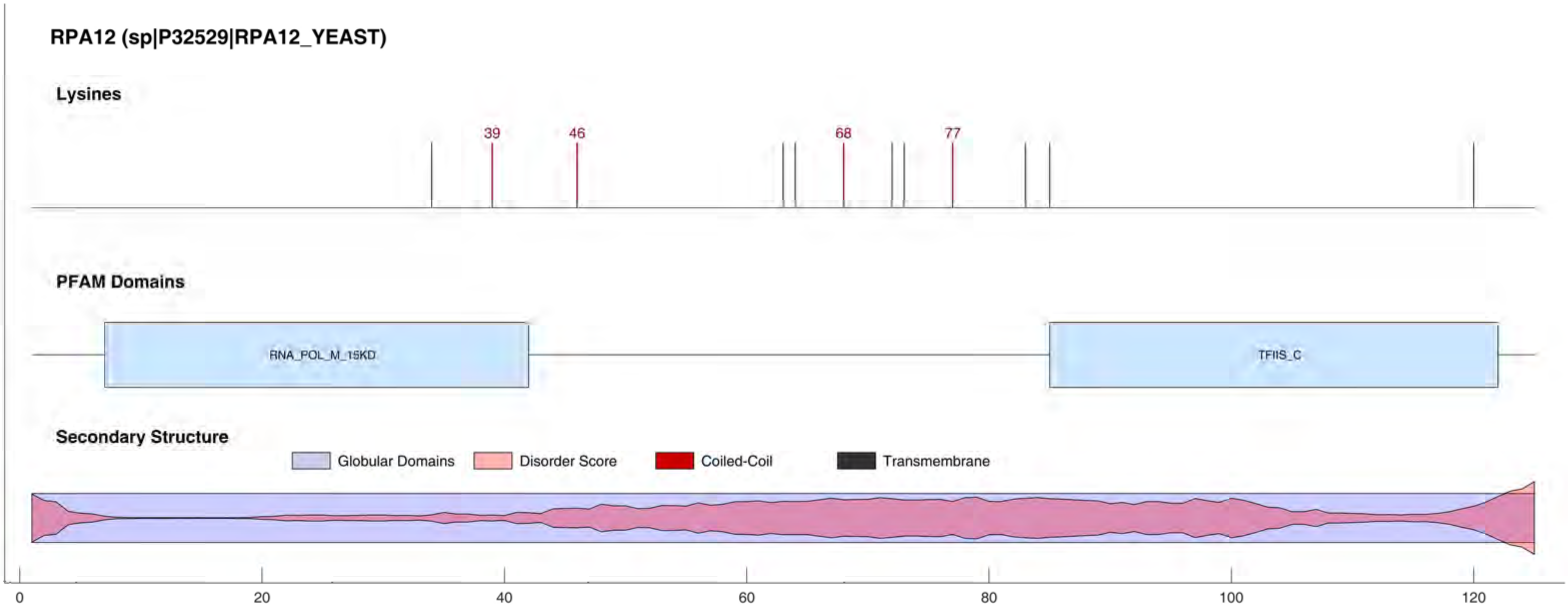
RPA12 (sp|P32529|RPA12_YEAST)
Lysines
39
46
68
77
PFAM Domains
RNA_POL_M_15KD
TFIIS_C
Secondary Structure
Globular Domains
Disorder Score
Coiled-Coil
Transmembrane
0
20
40
60
80
100
120

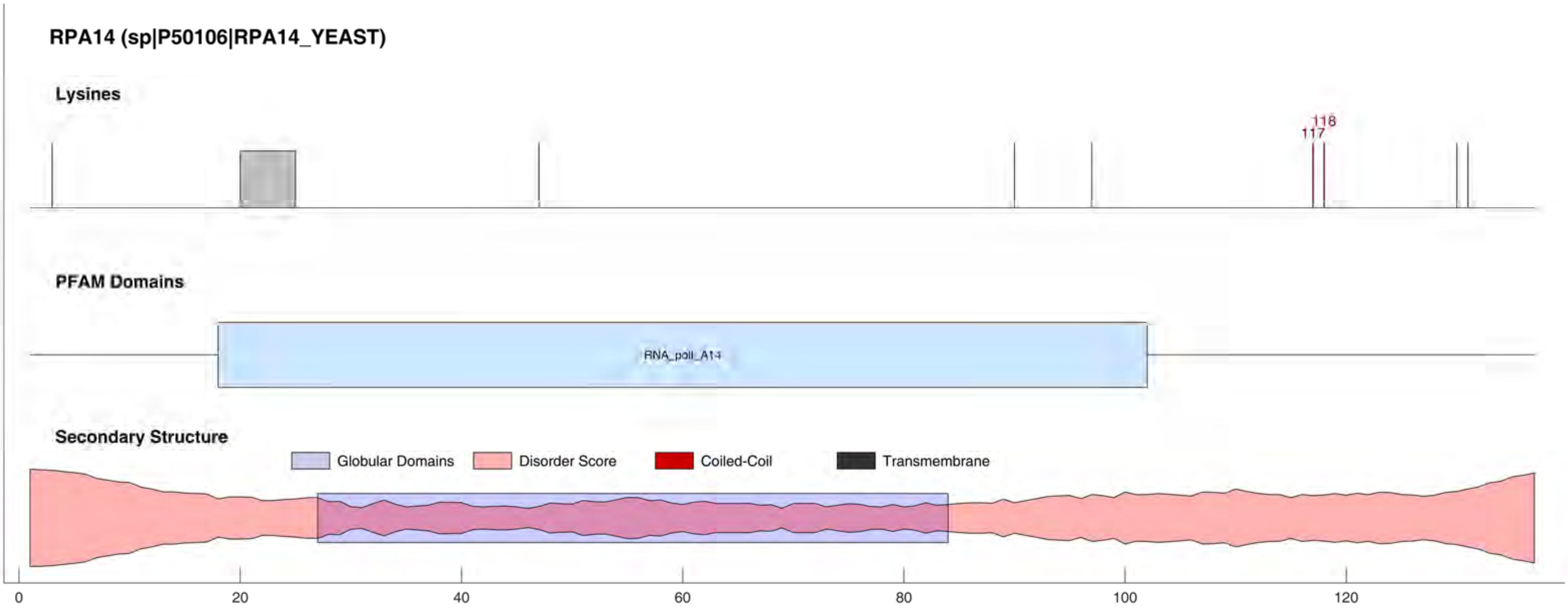
RPA14 (sp|P50106|RPA14_YEAST)
Lysines
118
117
PFAM Domains
RNA_pol_A14
Secondary Structure
Globular Domains
Disorder Score
Coiled-Coil
Transmembrane
0
20
40
60
80
100
120

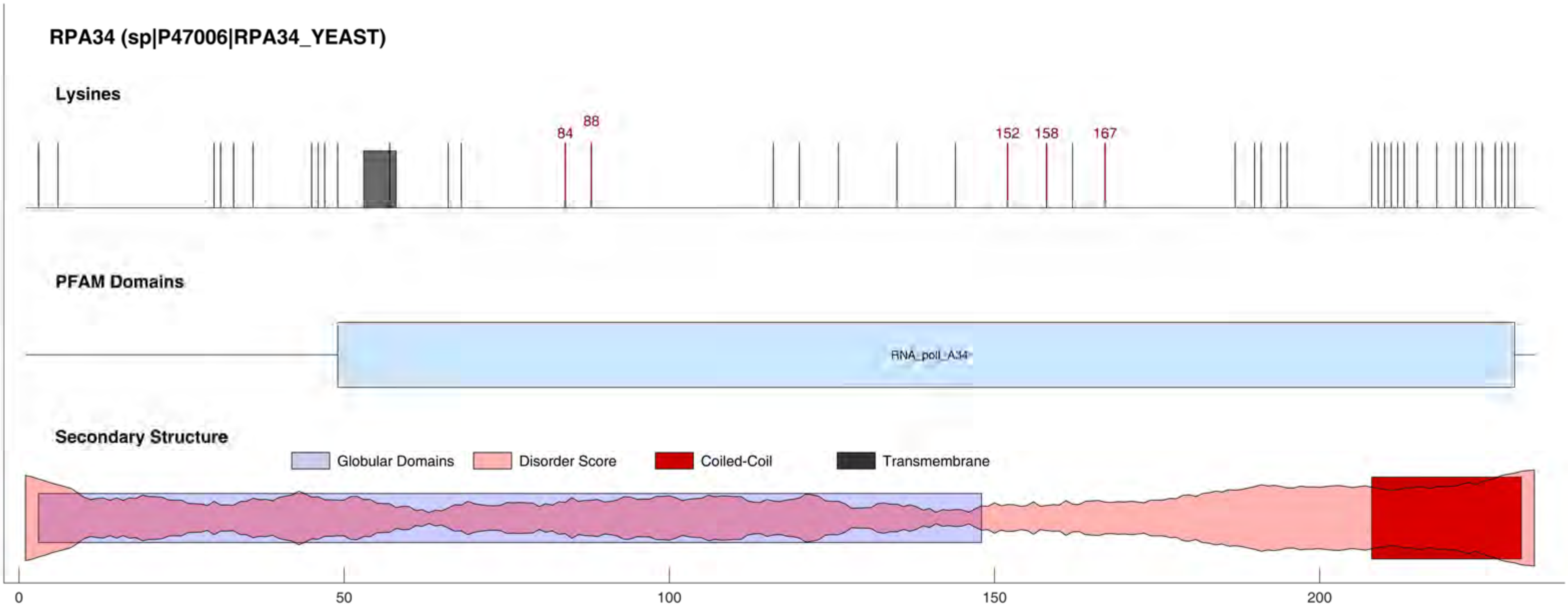
RPA34 (sp|P47006|RPA34_YEAST)
Lysines
84
88
152
158
167
PFAM Domains
RNA_pol_I_A34
Secondary Structure
Globular Domains
Disorder Score
Coiled-Coil
Transmembrane
0
50
100
150
200

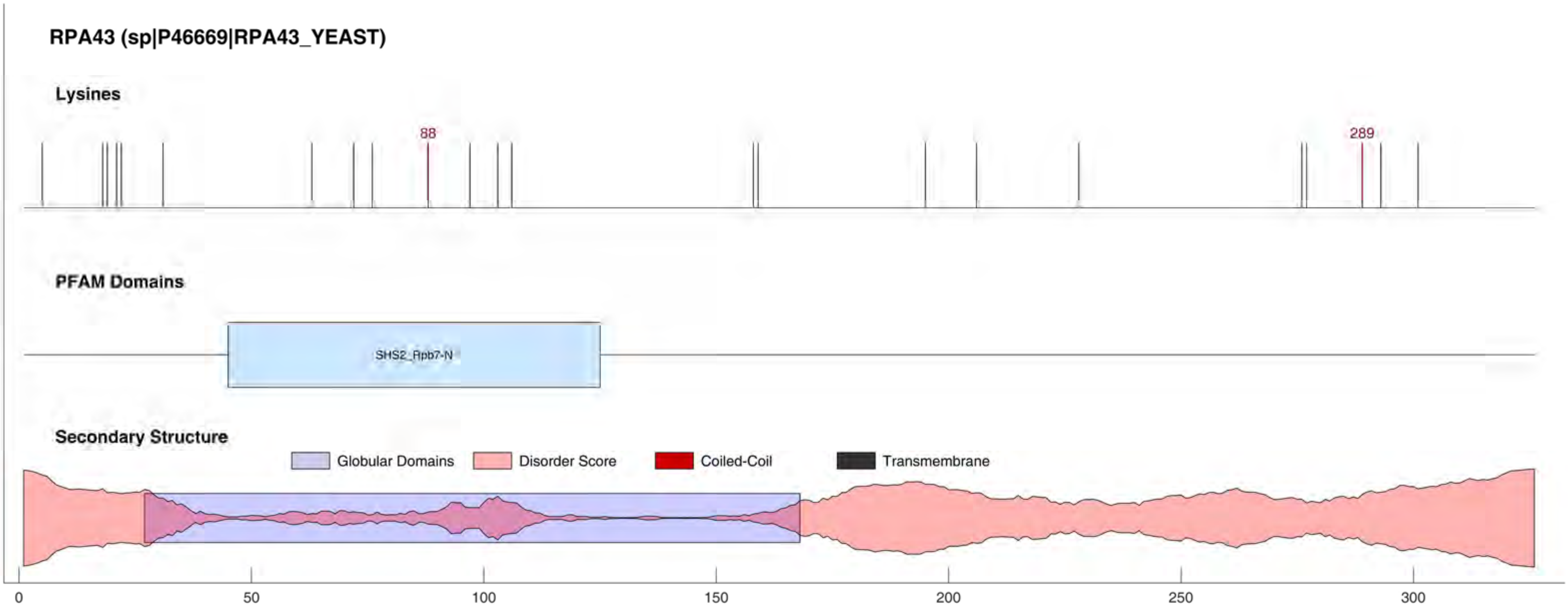
RPA43 (sp|P46669|RPA43_YEAST)
Lysines
88
289
PFAM Domains
SHS2_Rpb7-N
Secondary Structure
Globular Domains
Disorder Score
Coiled-Coil
Transmembrane
0
50
100
150
200
250
300

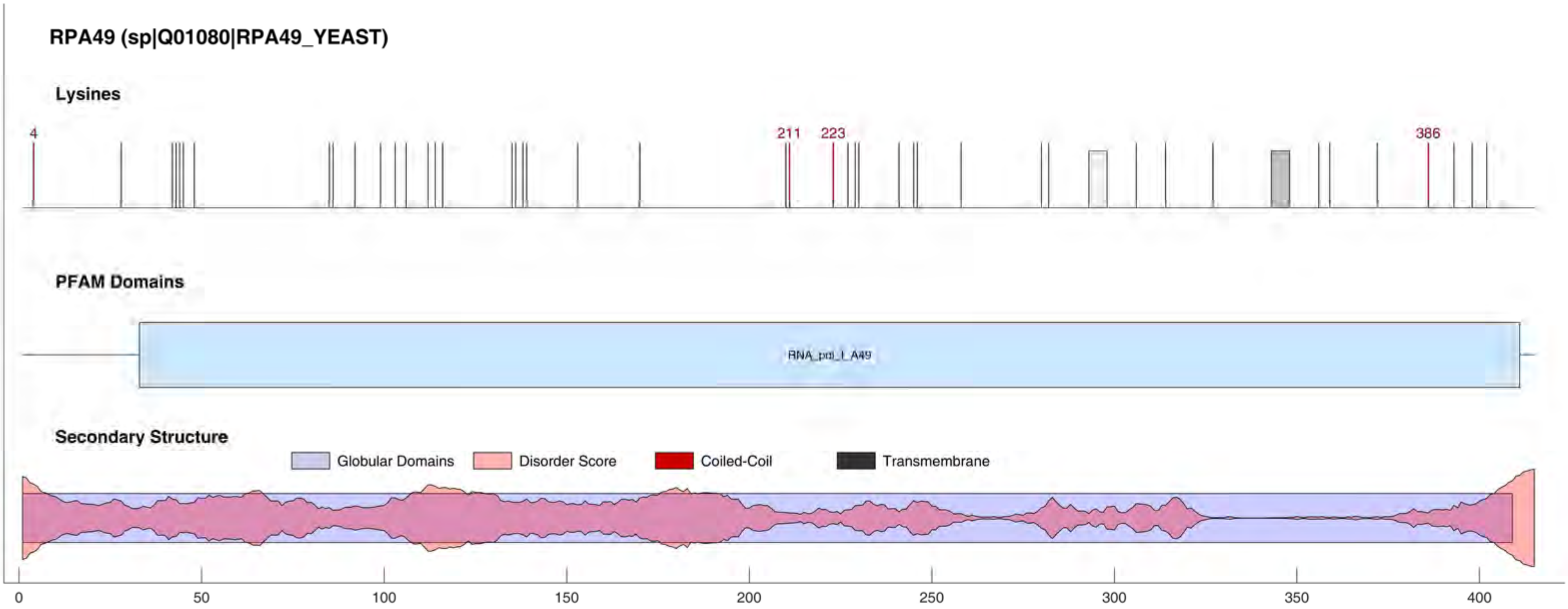
RPA49 (sp|Q01080|RPA49_YEAST)
Lysines
4
211
223
386
PFAM Domains
RNA_pol_I_A49
Secondary Structure
Globular Domains
Disorder Score
Coiled-Coil
Transmembrane
0
50
100
150
200
250
300
350
400

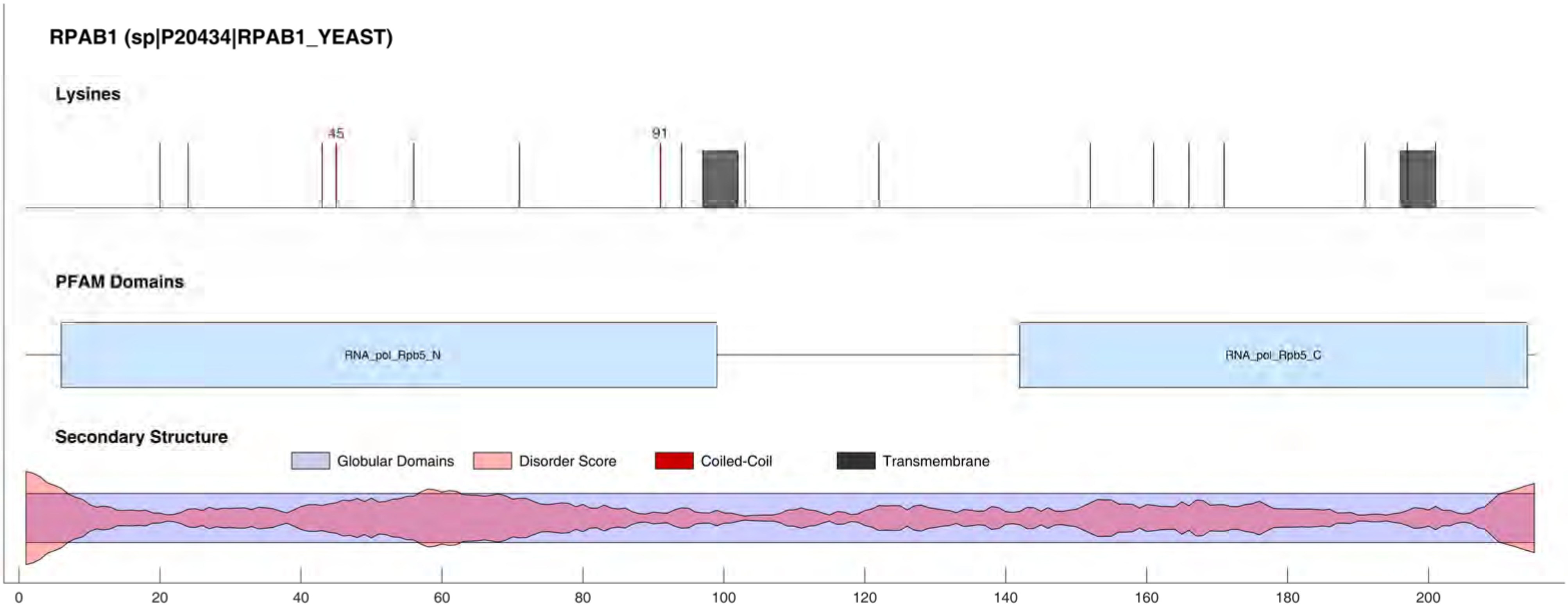

RPAB1 (sp|P20434|RPAB1_YEAST)
Lysines
45
91
PFAM Domains
RNA_pol_Rpb5_N
RNA_pol_Rpb5_C
Secondary Structure
Globular Domains
Disorder Score
Coiled-Coil
Transmembrane
0
20
40
60
80
100
120
140
160
180
200

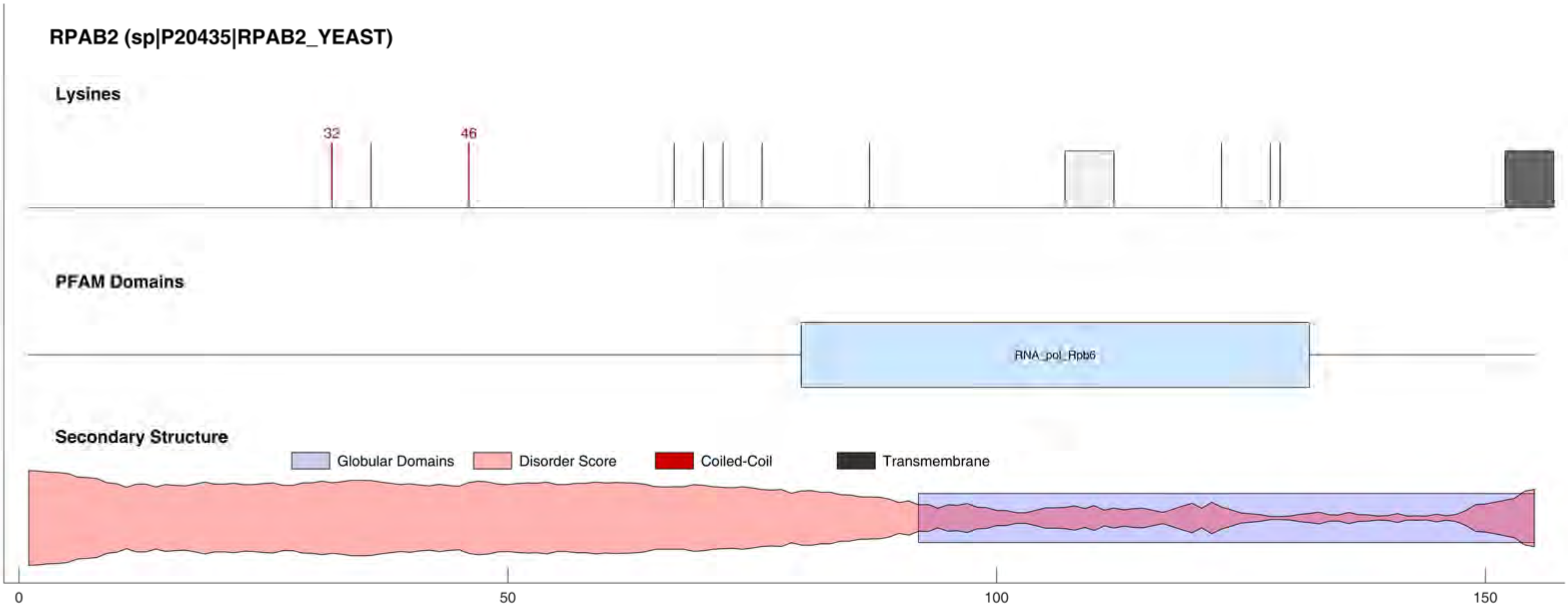

RPAB2 (sp|P20435|RPAB2_YEAST)
Lysines
32
46
PFAM Domains
RNA_pol_Rpb6
Secondary Structure
Globular Domains
Disorder Score
Coiled-Coil
Transmembrane
0
50
100
150

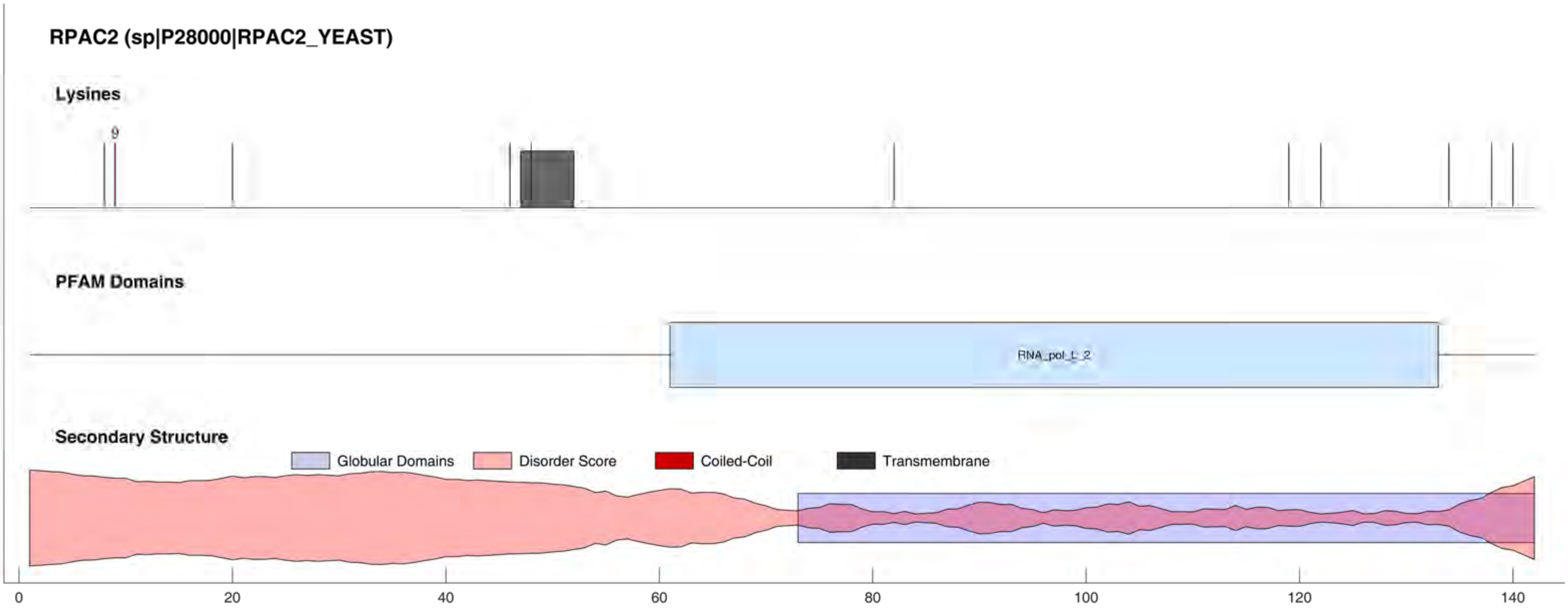

RPAC2 (sp|P28000|RPAC2_YEAST)
Lysines
9
PFAM Domains
RNA_pol_L_2
Secondary Structure
Globular Domains
Disorder Score
Coiled-Coil
Transmembrane
0
20
40
60
80
100
120
140

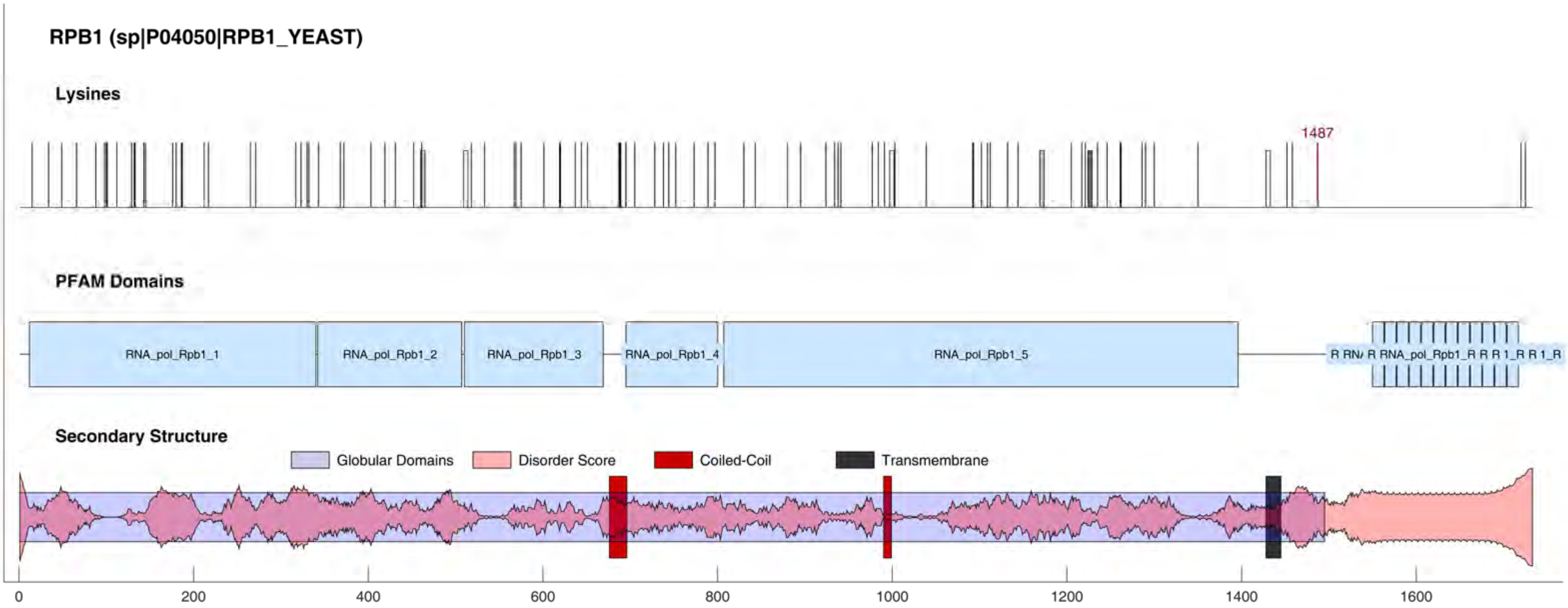
RPB1 (sp|P04050|RPB1_YEAST)
Lysines
1487
PFAM Domains
RNA_pol_Rpb1_1
RNA_pol_Rpb1_2
RNA_pol_Rpb1_3
RNA_pol_Rpb1_4
RNA_pol_Rpb1_5
Secondary Structure
Globular Domains
Disorder Score
Coiled-Coil
Transmembrane
0
200
400
600
800
1000
1200
1400
1600

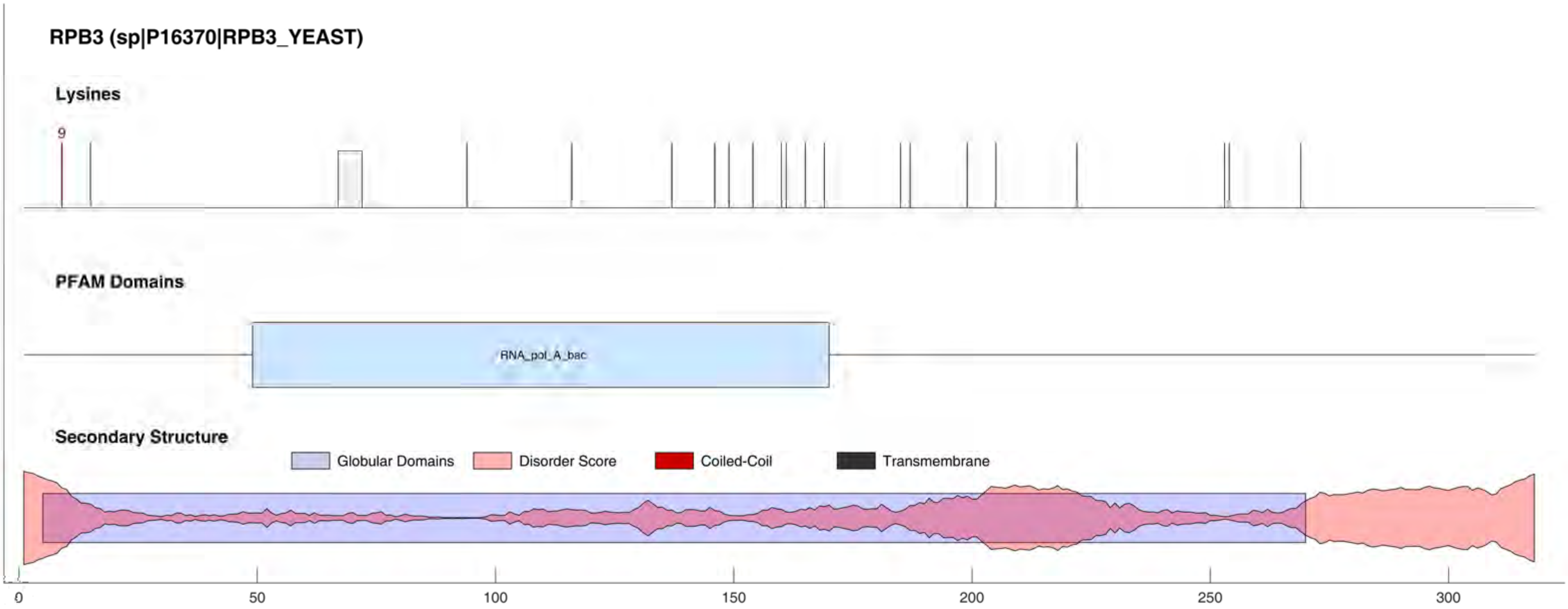
RPB3 (sp|P16370|RPB3_YEAST)
Lysines
9
PFAM Domains
RNA_pol_A_bac
Secondary Structure
Globular Domains
Disorder Score
Coiled-Coil
Transmembrane
0
50
100
150
200
250
300

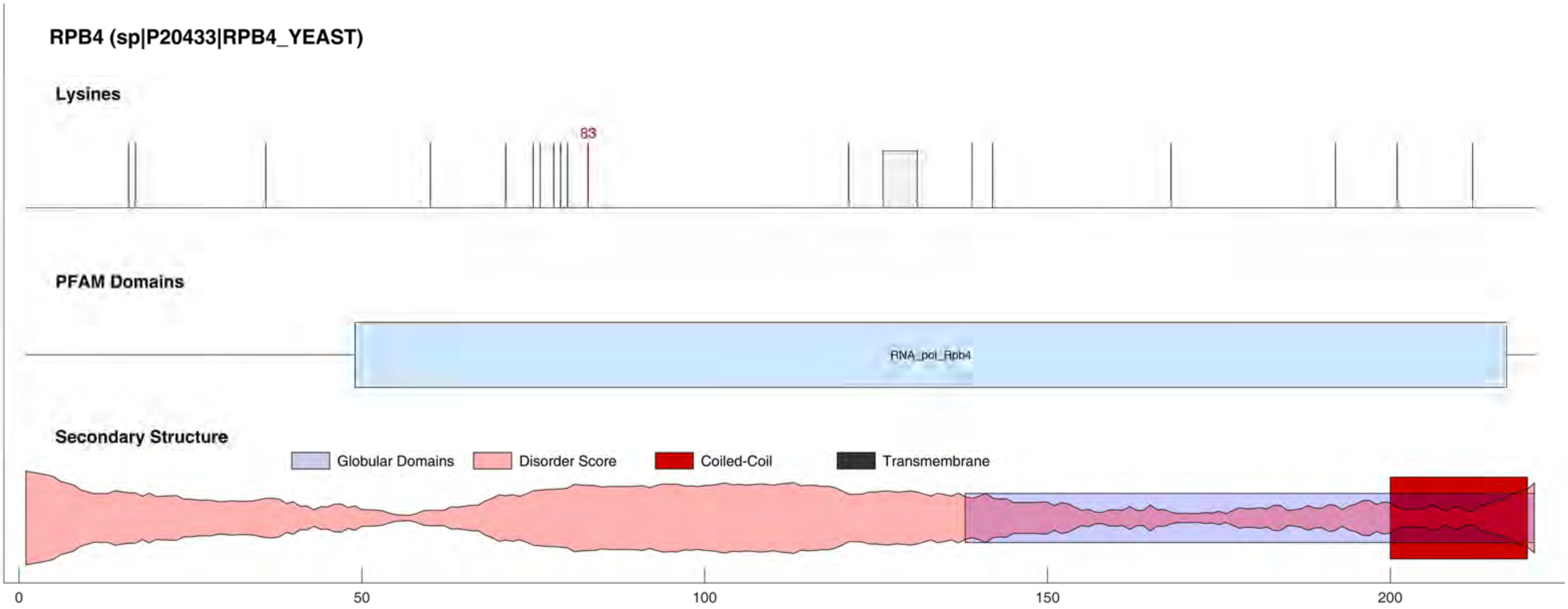

RPB4 (sp|P20433|RPB4_YEAST)
Lysines
83
PFAM Domains
RNA_pol_Rpb4
Secondary Structure
Globular Domains
Disorder Score
Coiled-Coil
Transmembrane
0
50
100
150
200

#### RPC2 (sp|P22276|RPC2_YEAST)

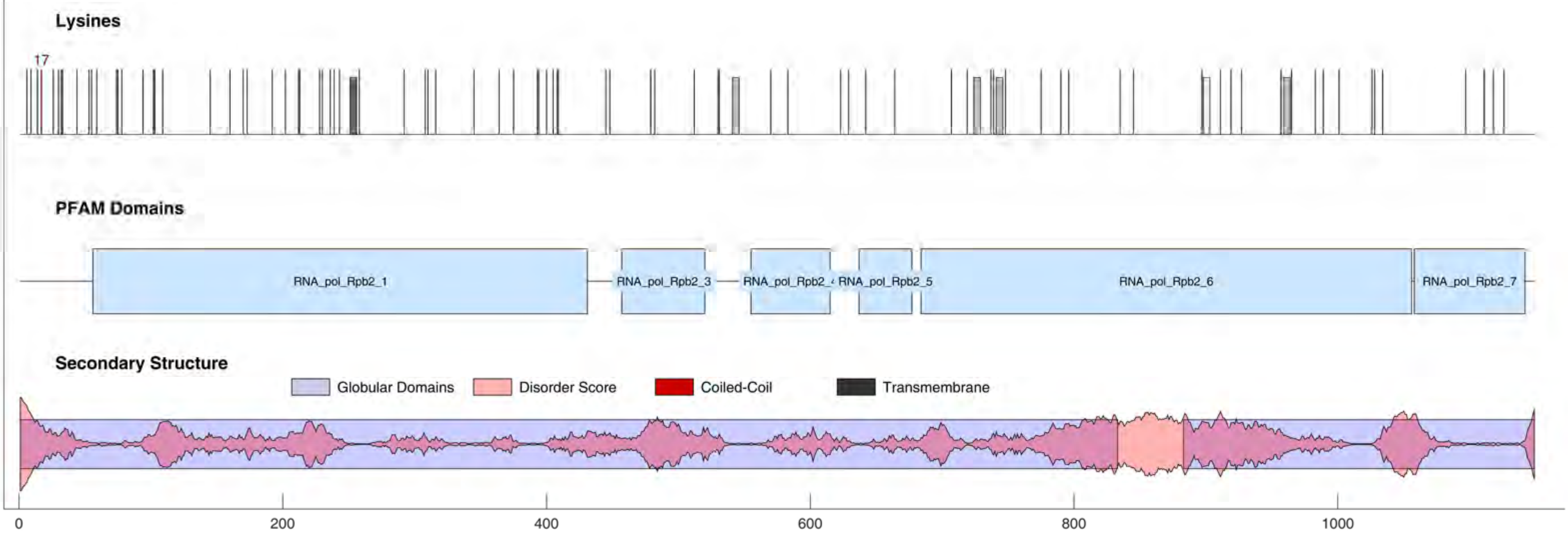

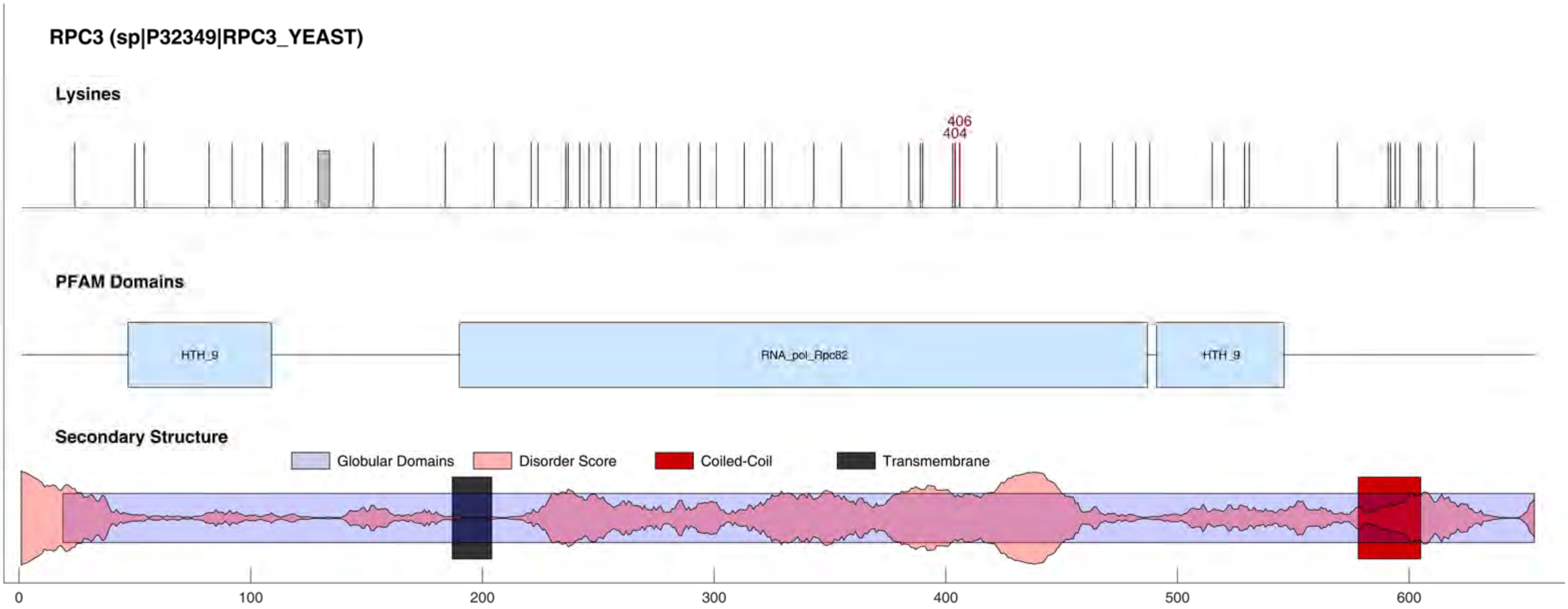
RPC3 (sp|P32349|RPC3_YEAST)
Lysines
406
404
PFAM Domains
HTH_9
RNA_pol_Rpc82
HTH_9
Secondary Structure
Globular Domains
Disorder Score
Coiled-Coil
Transmembrane
0
100
200
300
400
500
600

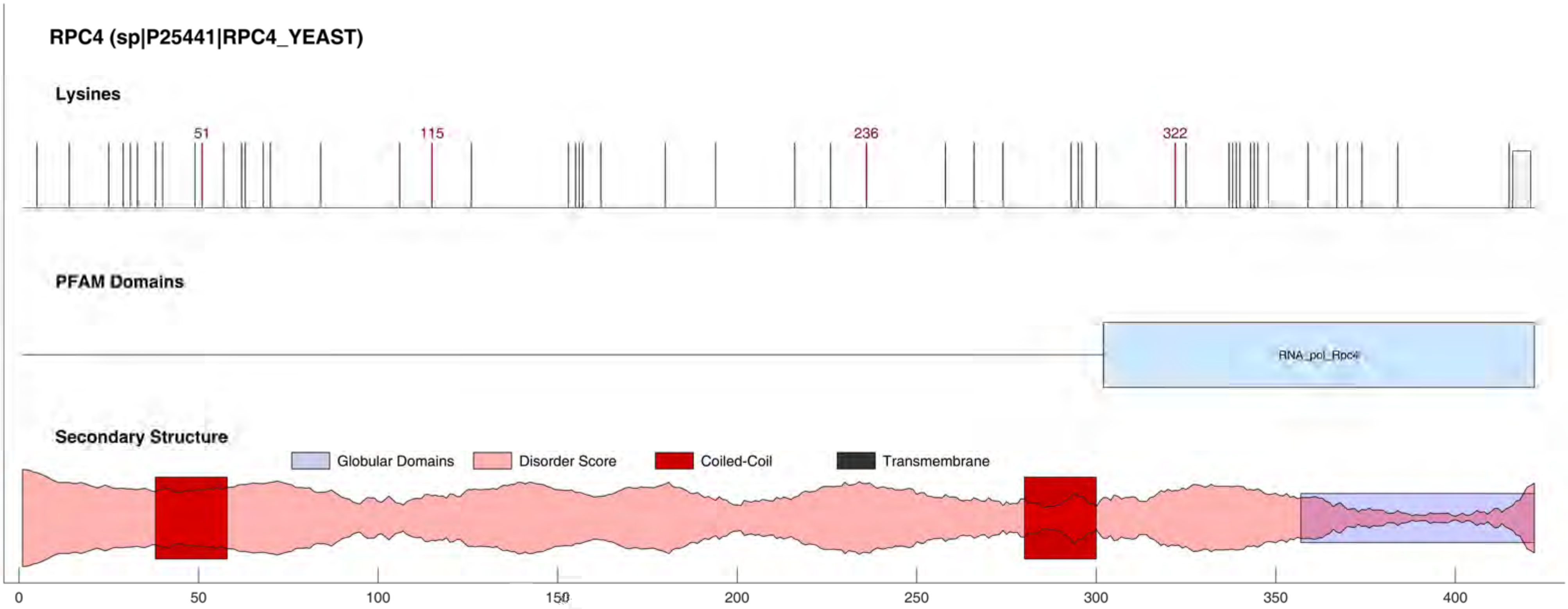

RPC4 (sp|P25441|RPC4_YEAST)
Lysines
51
115
236
322
PFAM Domains
RNA_pol_Rpc4
Secondary Structure
Globular Domains
Disorder Score
Coiled-Coil
Transmembrane
0
50
100
150
200
250
300
350
400

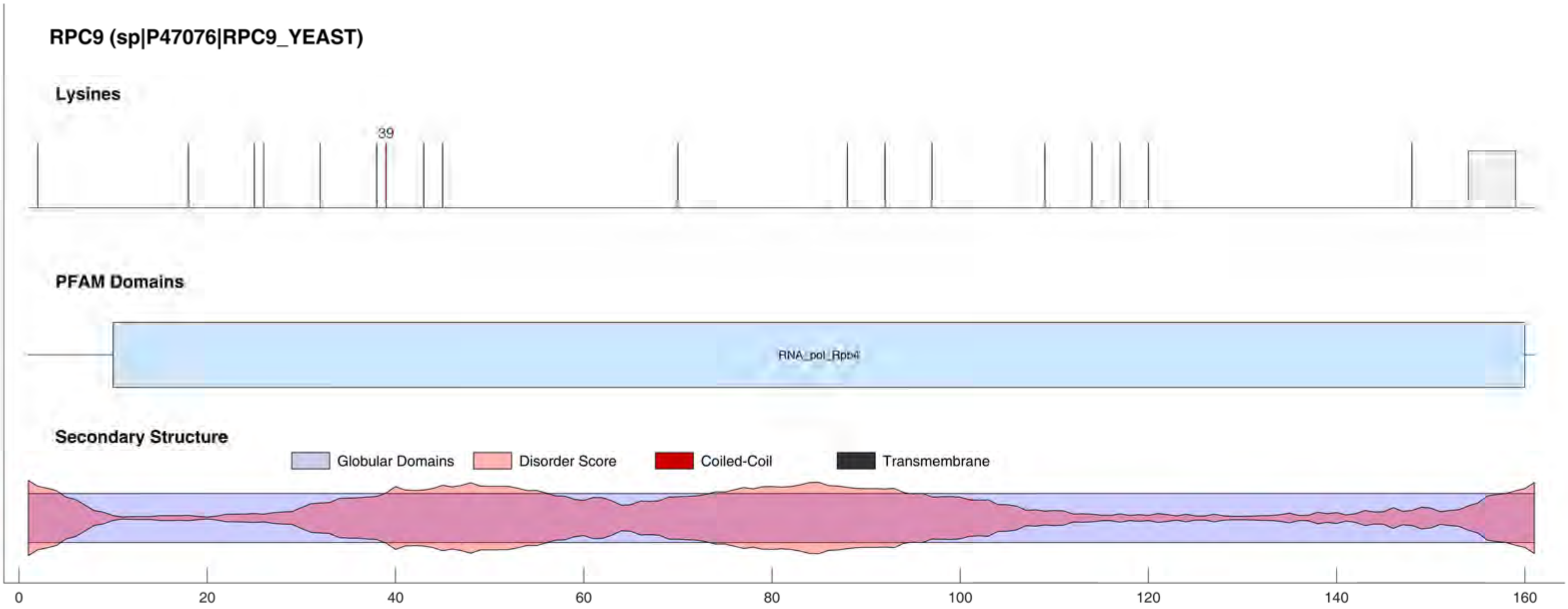

RPC9 (sp|P47076|RPC9_YEAST)
Lysines
39
PFAM Domains
RNA_pol_Rpb4
Secondary Structure
Globular Domains
Disorder Score
Coiled-Coil
Transmembrane
0
20
40
60
80
100
120
140
160

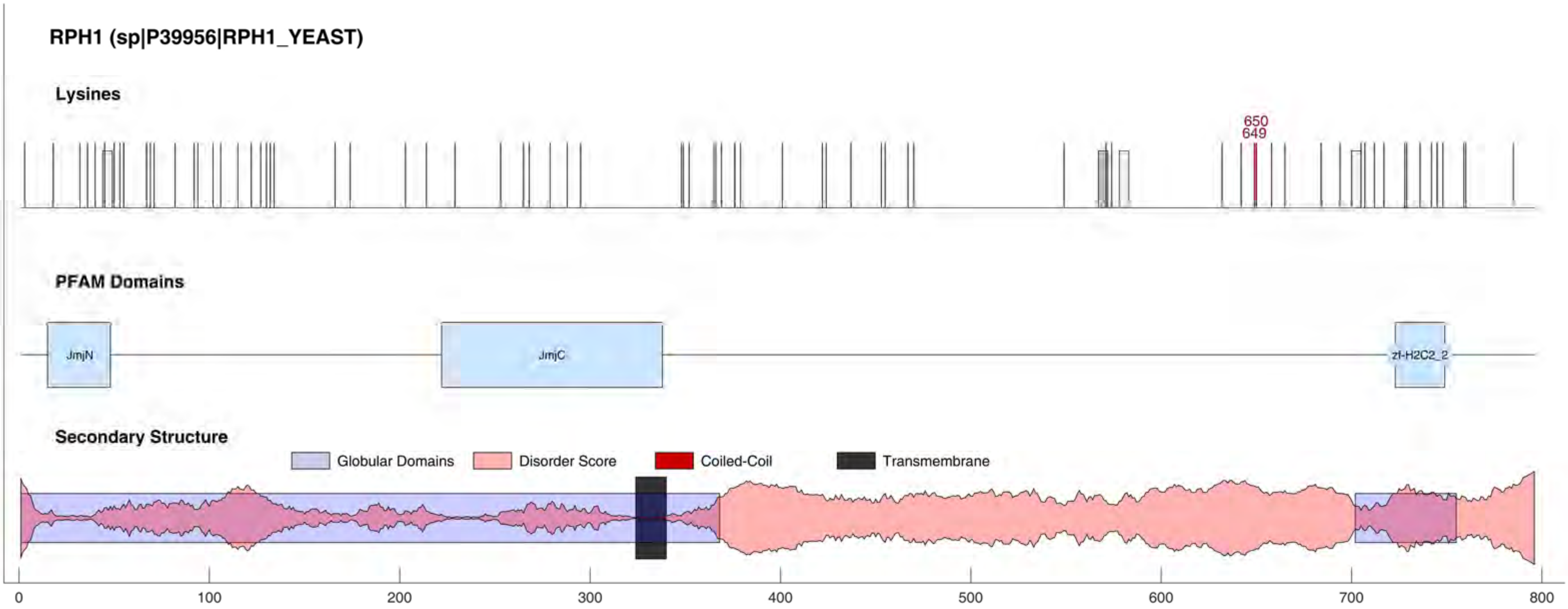

RPH1 (sp|P39956|RPH1_YEAST)
Lysines
650
649
PFAM Domains
JmjN
JmjC
zf-H2C2_2
Secondary Structure
Globular Domains
Disorder Score
Coiled-Coil
Transmembrane
0
100
200
300
400
500
600
700
800

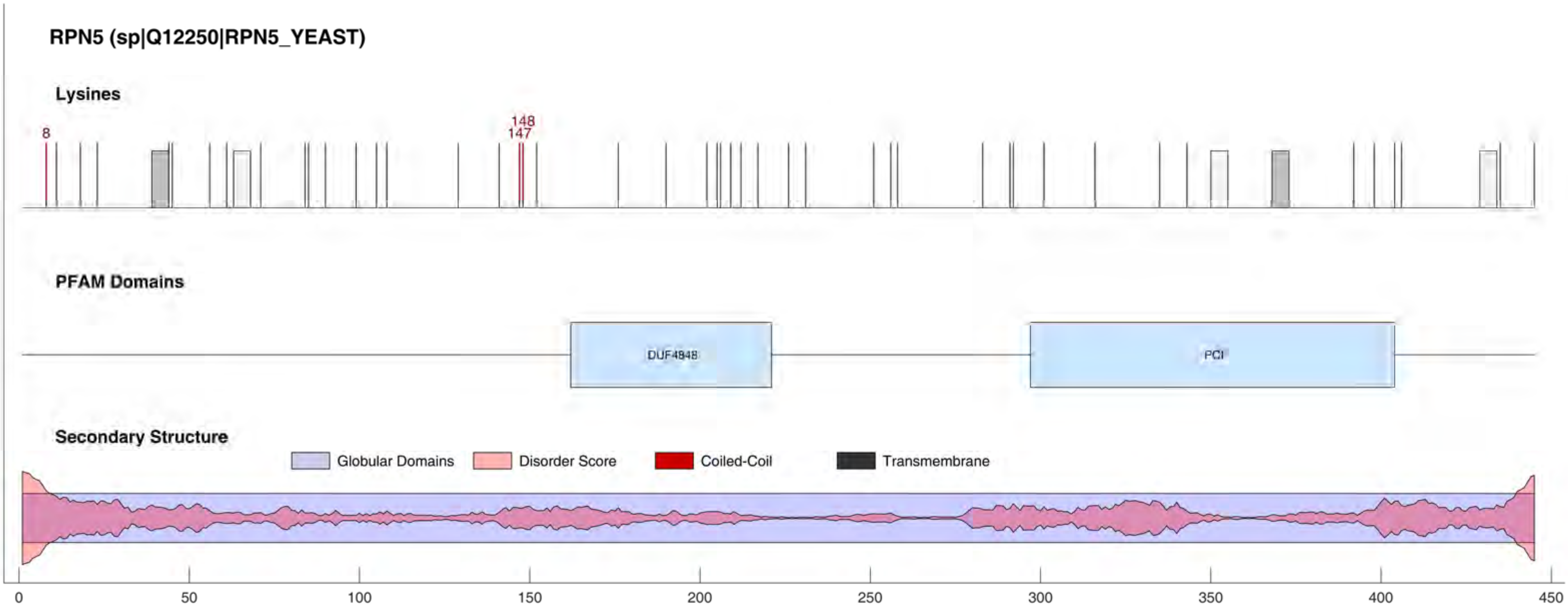

RPN5 (sp|Q12250|RPN5_YEAST)
Lysines
8
148
147
PFAM Domains
DUF4848
PCI
Secondary Structure
Globular Domains
Disorder Score
Coiled-Coil
Transmembrane
0
50
100
150
200
250
300
350
400
450

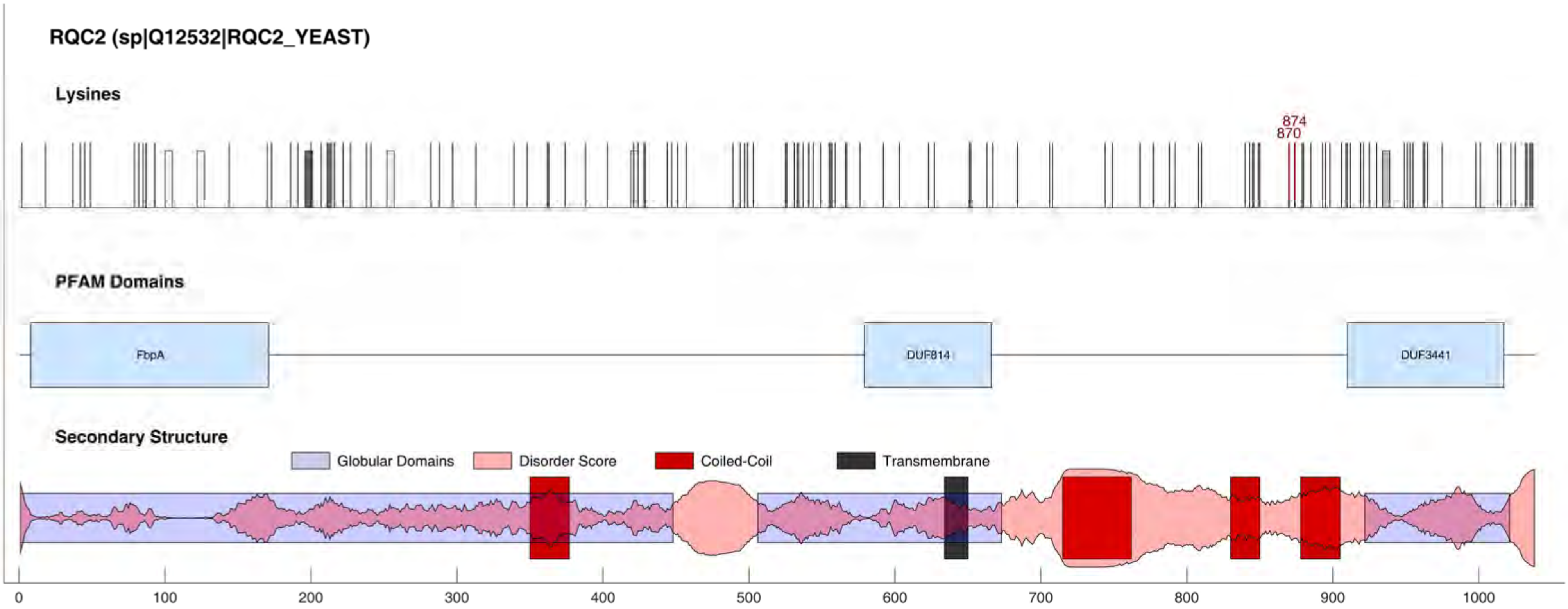
RQC2 (sp|Q12532|RQC2_YEAST)
Lysines
874
870
PFAM Domains
FbpA
DUF814
DUF3441
Secondary Structure
Globular Domains
Disorder Score
Coiled-Coil
Transmembrane
0
100
200
300
400
500
600
700
800
900
1000

RRN10 (sp|P38204|RRN10_YEAST)

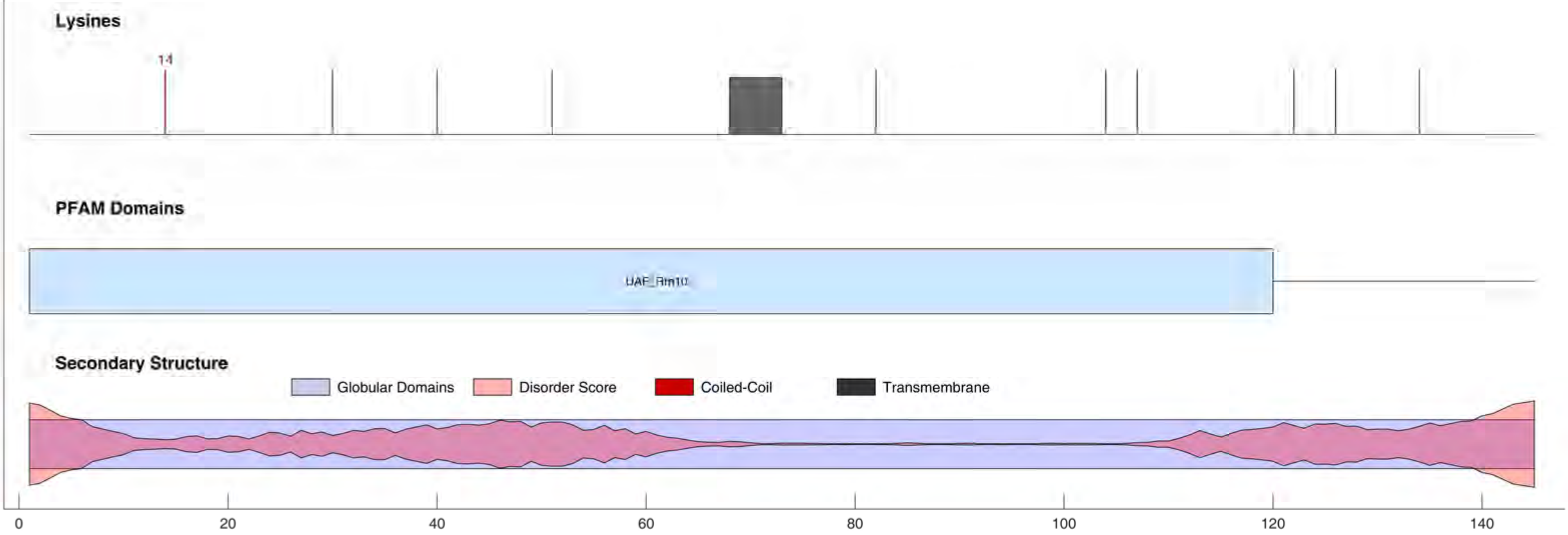

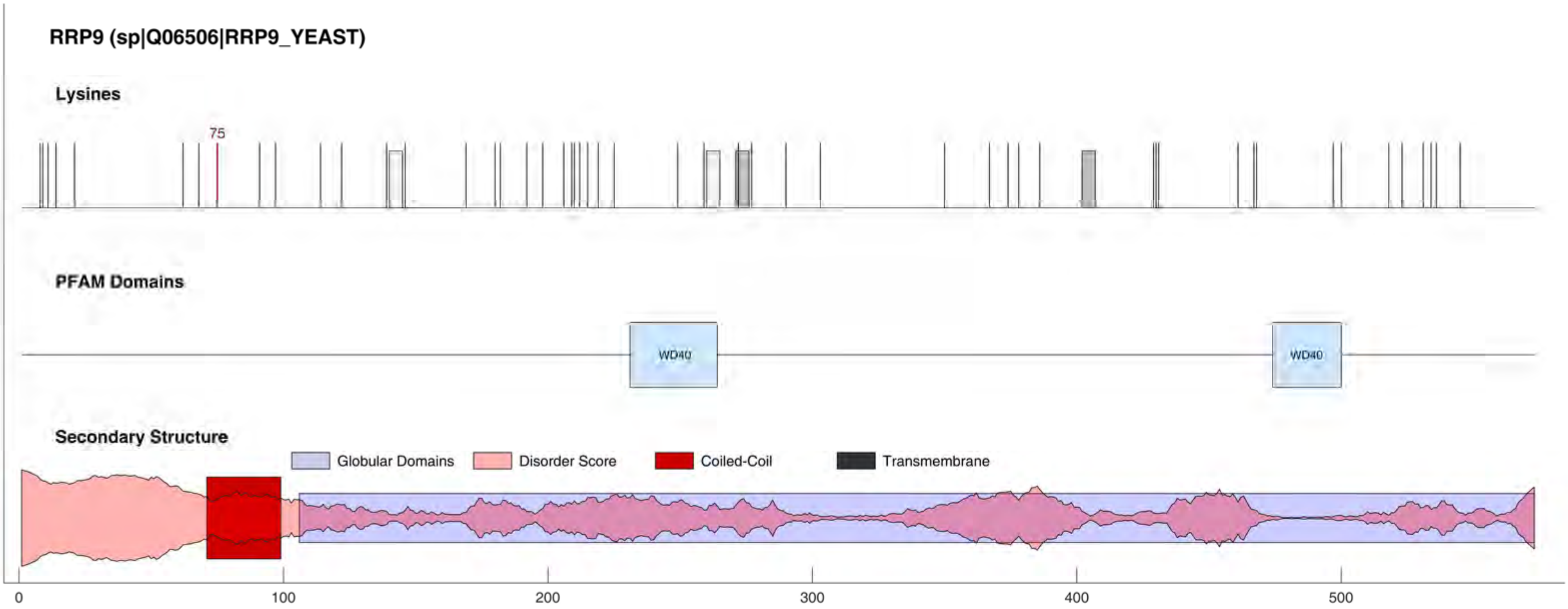
RRP9 (sp|Q06506|RRP9_YEAST)
Lysines
75
PFAM Domains
WD40
WD40
Secondary Structure
Globular Domains
Disorder Score
Coiled-Coil
Transmembrane
0
100
200
300
400
500

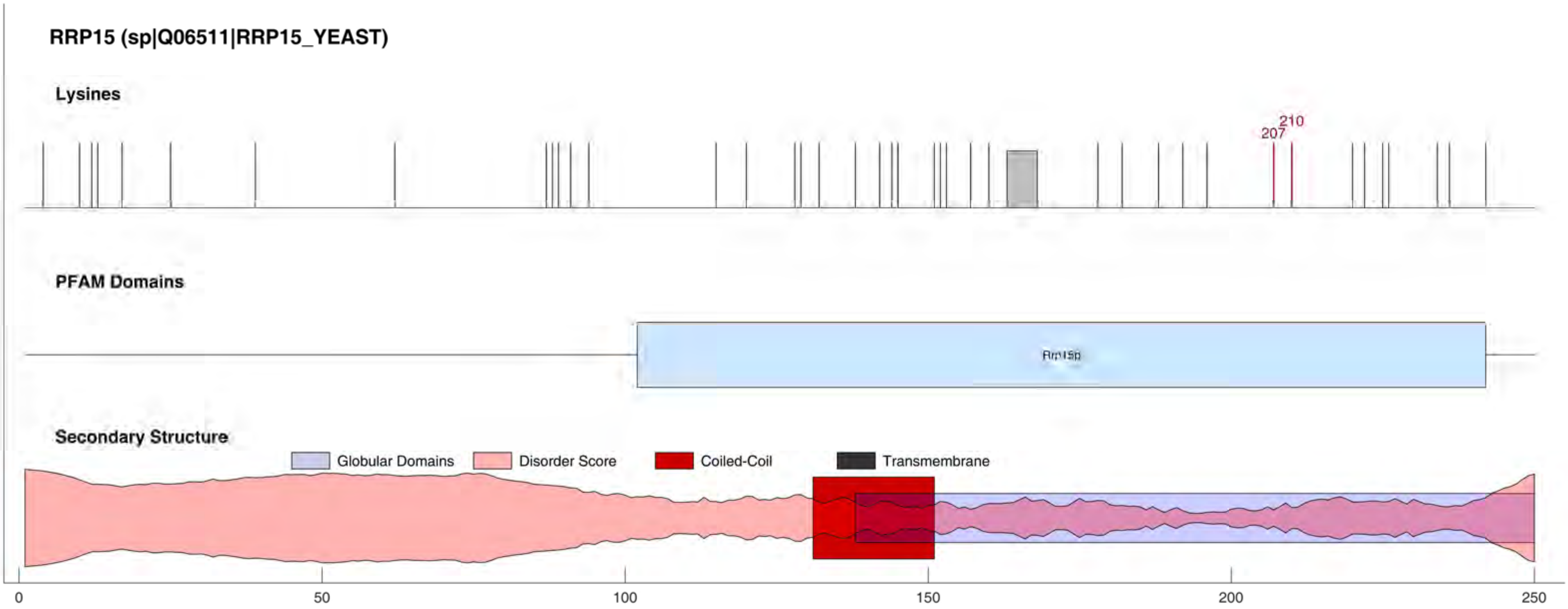
RRP15 (sp|Q06511|RRP15_YEAST)
Lysines
210
207
PFAM Domains
Rrp15p
Secondary Structure
Globular Domains
Disorder Score
Coiled-Coil
Transmembrane
0
50
100
150
200
250

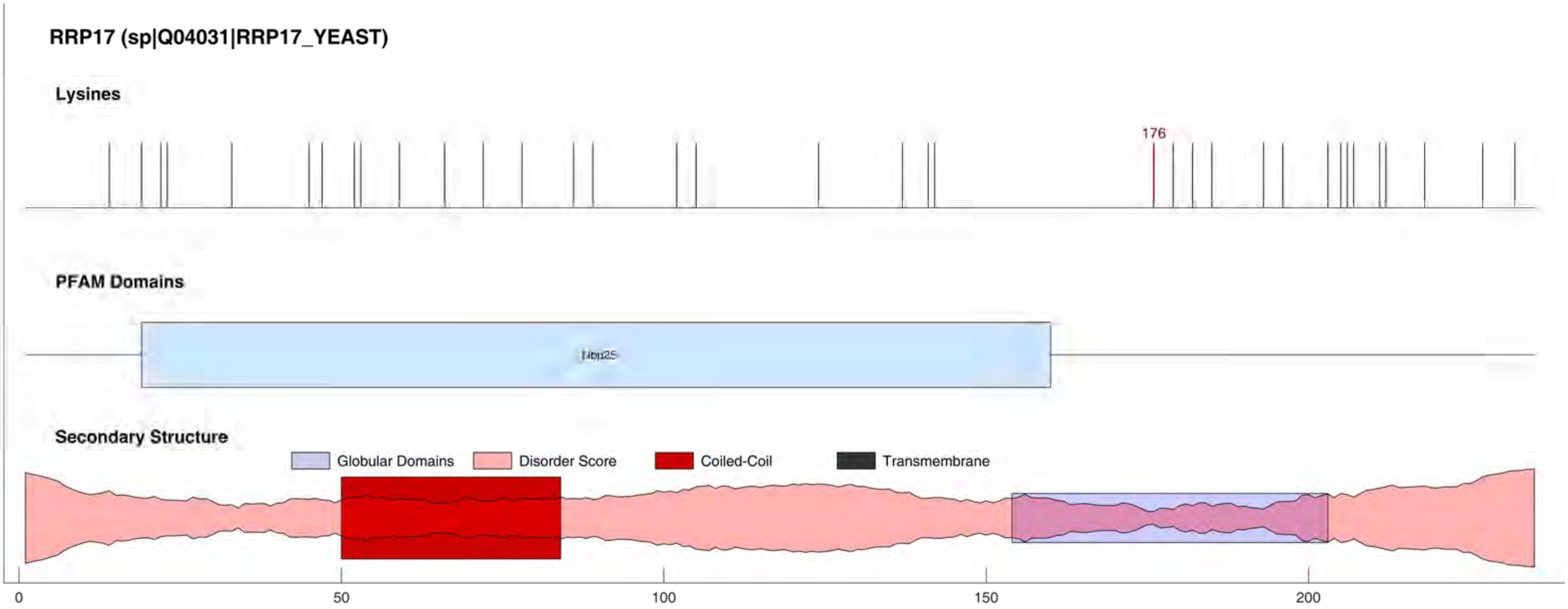
RRP17 (sp|Q04031|RRP17_YEAST)
Lysines
176
PFAM Domains
Nop25
Secondary Structure
Globular Domains
Disorder Score
Coiled-Coil
Transmembrane
0
50
100
150
200

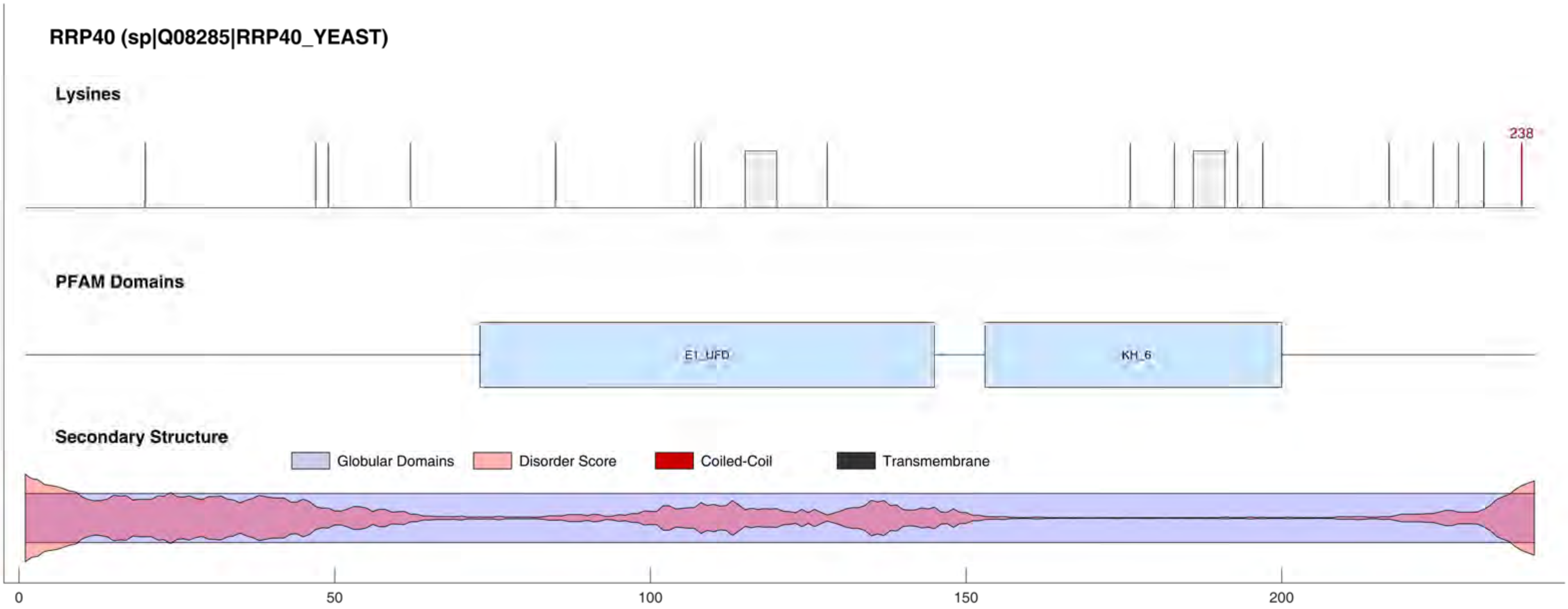
RRP40 (sp|Q08285|RRP40_YEAST)
Lysines
238
PFAM Domains
E1_UFD
KH_6
Secondary Structure
Globular Domains
Disorder Score
Coiled-Coil
Transmembrane
0
50
100
150
200

RS2 (sp|P25443|RS2_YEAST)

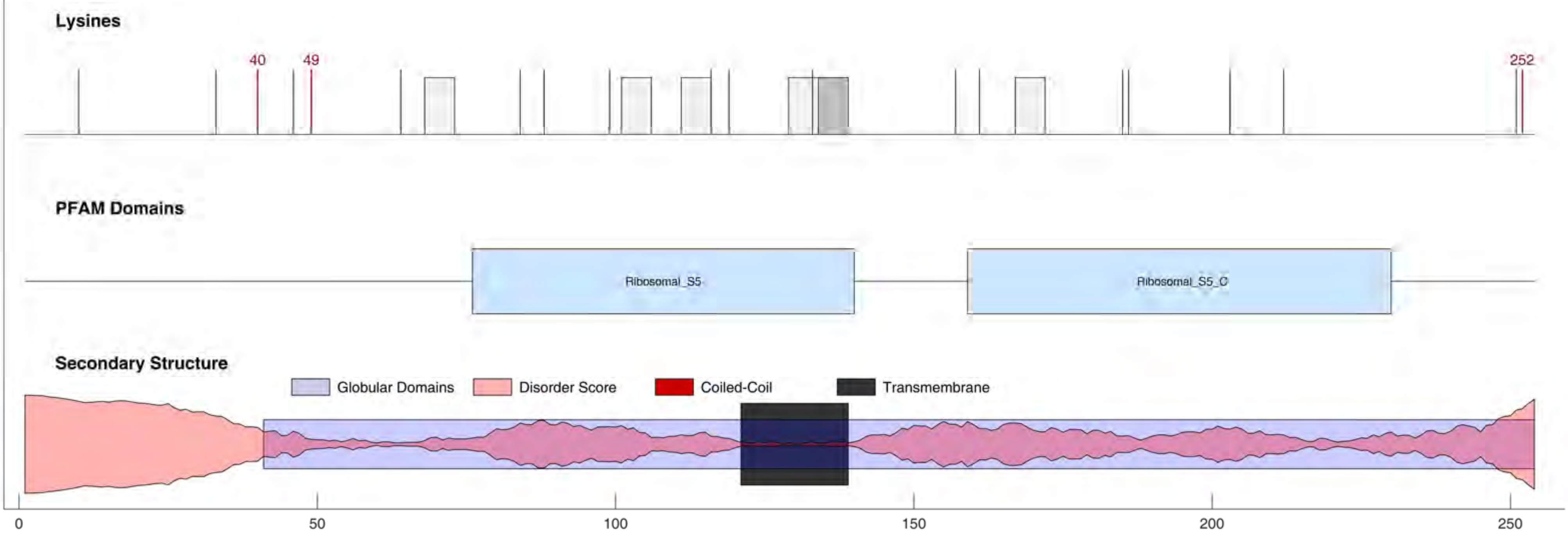

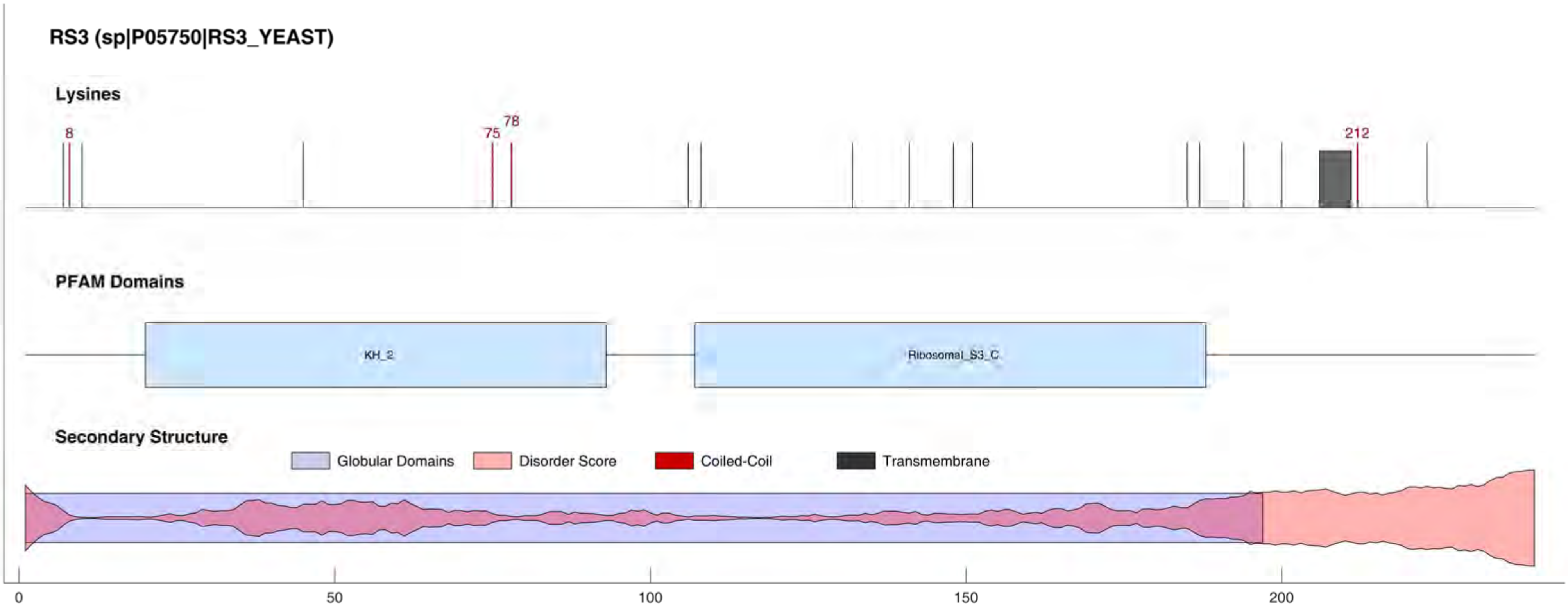

RS3 (sp|P05750|RS3_YEAST)
Lysines
8
75
78
212
PFAM Domains
KH_2
Ribosomal_S3_C
Secondary Structure
Globular Domains
Disorder Score
Coiled-Coil
Transmembrane
0
50
100
150
200

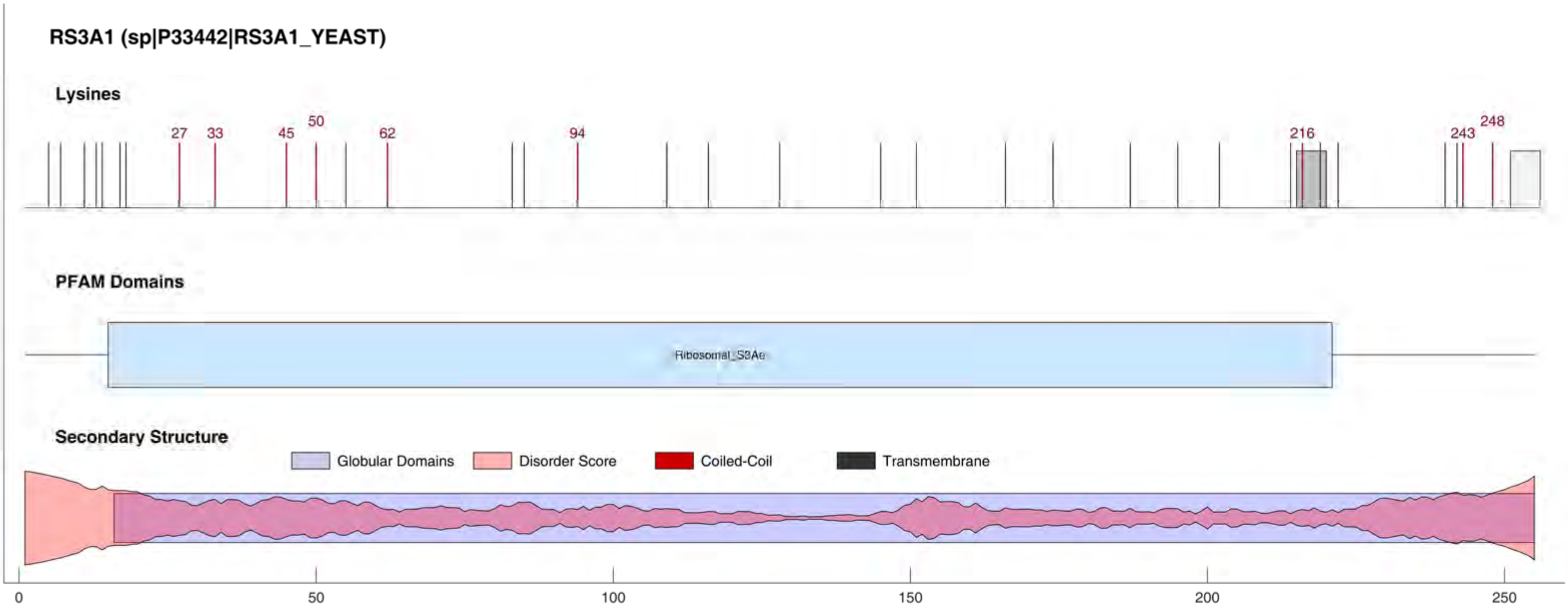
RS3A1 (sp|P33442|RS3A1_YEAST)
Lysines
27
33
45
50
62
94
216
243
248
PFAM Domains
Ribosomal_S3Ae
Secondary Structure
Globular Domains
Disorder Score
Coiled-Coil
Transmembrane
0
50
100
150
200
250

RS3A2 (sp|P23248|RS3A2_YEAST)

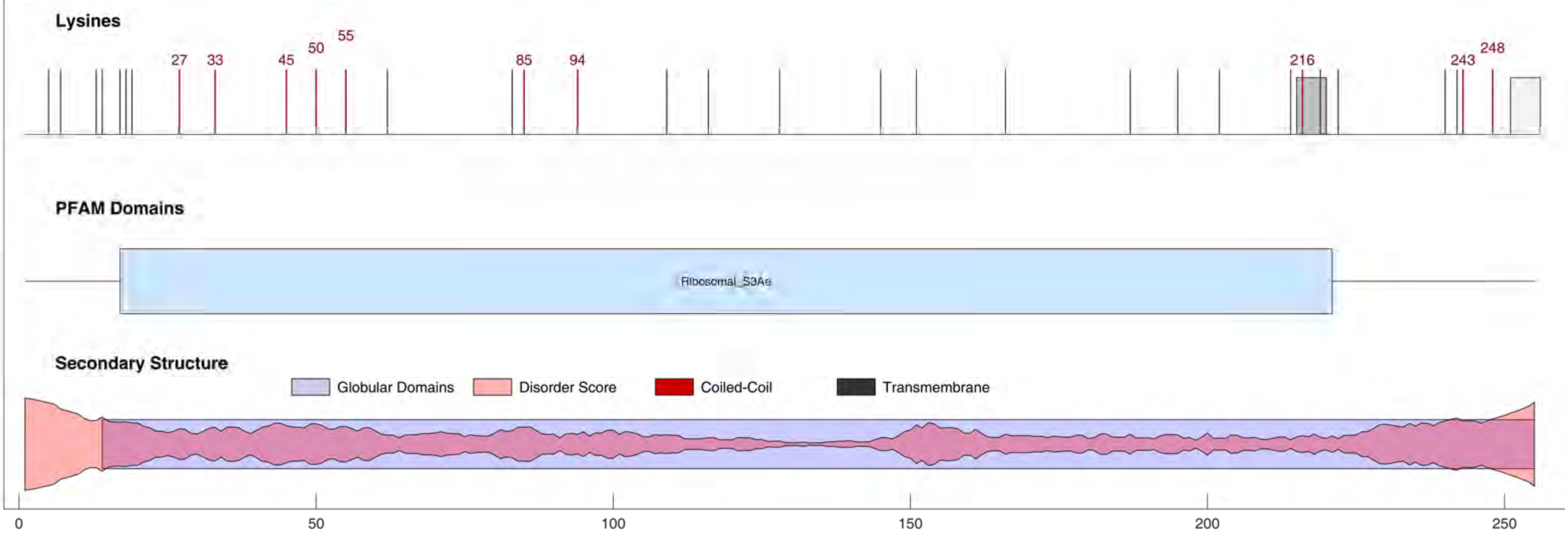

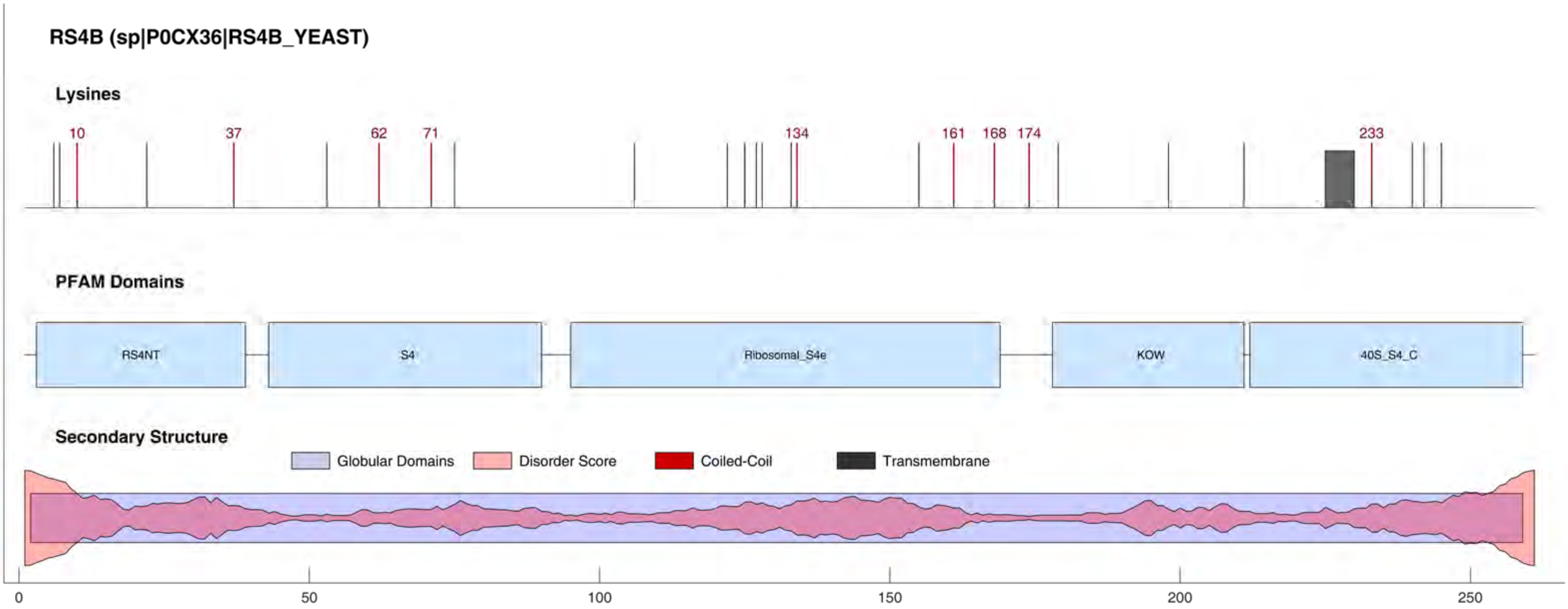

RS4B (sp|P0CX36|RS4B_YEAST)
Lysines
10
37
62
71
134
161
168
174
233
PFAM Domains
RS4NT
S4
Ribosomal_S4e
KOW
40S_S4_C
Secondary Structure
Globular Domains
Disorder Score
Coiled-Coil
Transmembrane
0
50
100
150
200
250

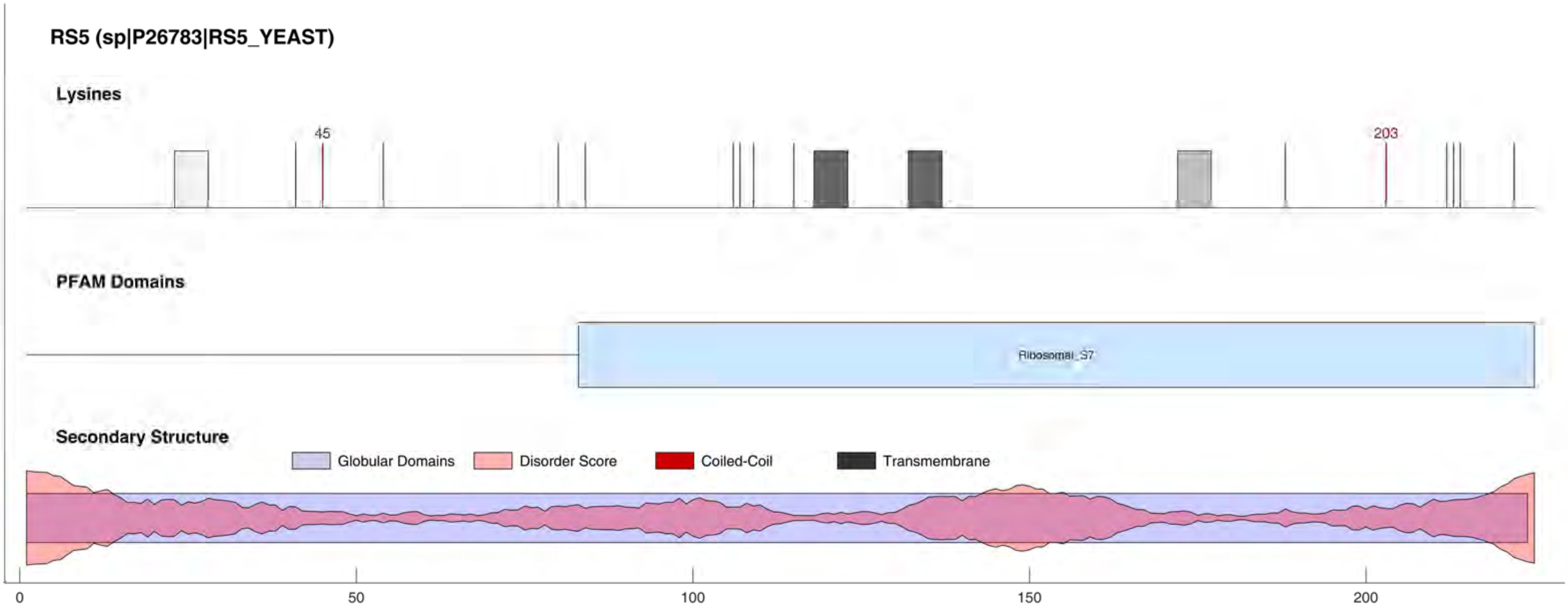
RS5 (sp|P26783|RS5_YEAST)
Lysines
45
203
PFAM Domains
Ribosomal_S7
Secondary Structure
Globular Domains
Disorder Score
Coiled-Coil
Transmembrane
0
50
100
150
200

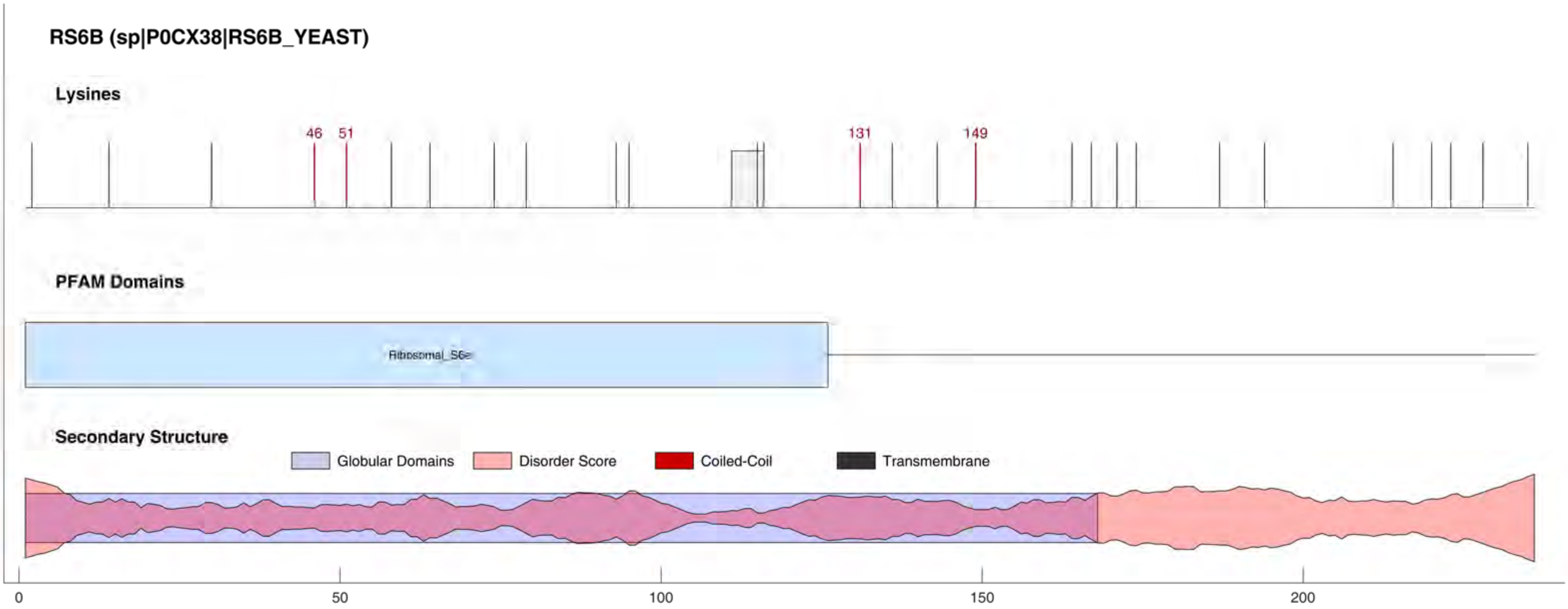
RS6B (sp|P0CX38|RS6B_YEAST)
Lysines
46
51
131
149
PFAM Domains
Ribosomal_S6e
Secondary Structure
Globular Domains
Disorder Score
Coiled-Coil
Transmembrane
0
50
100
150
200

### RS7A (sp|P26786|RS7A_YEAST)

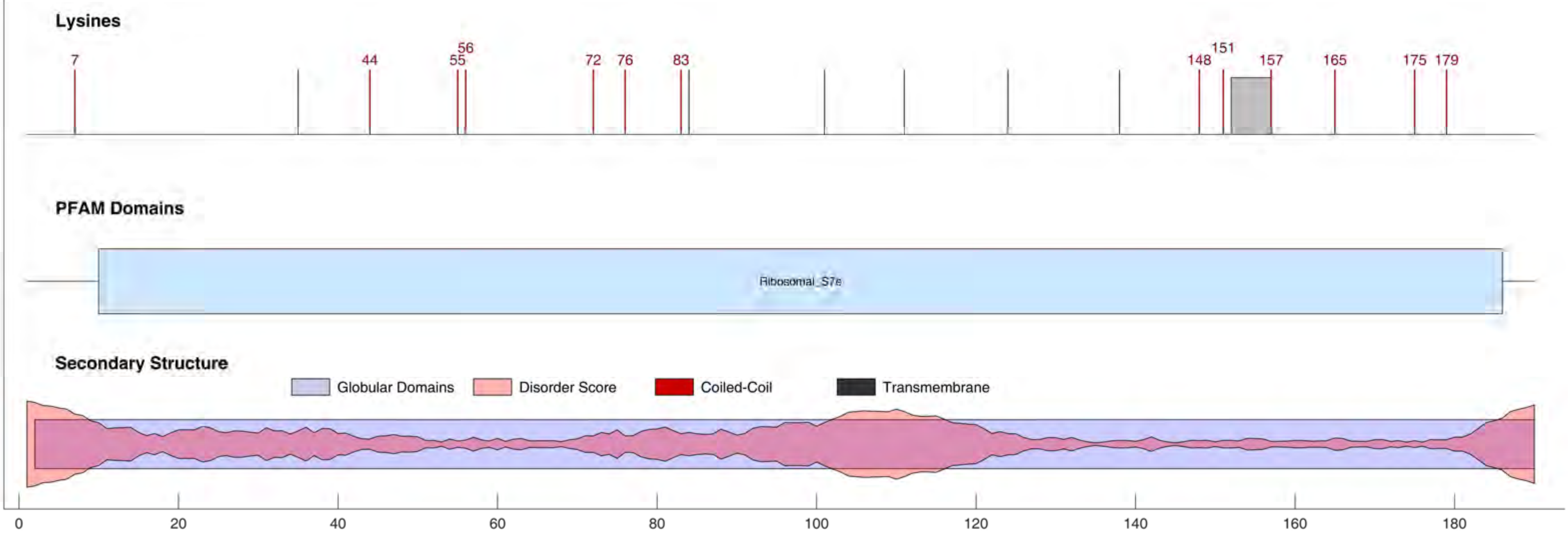

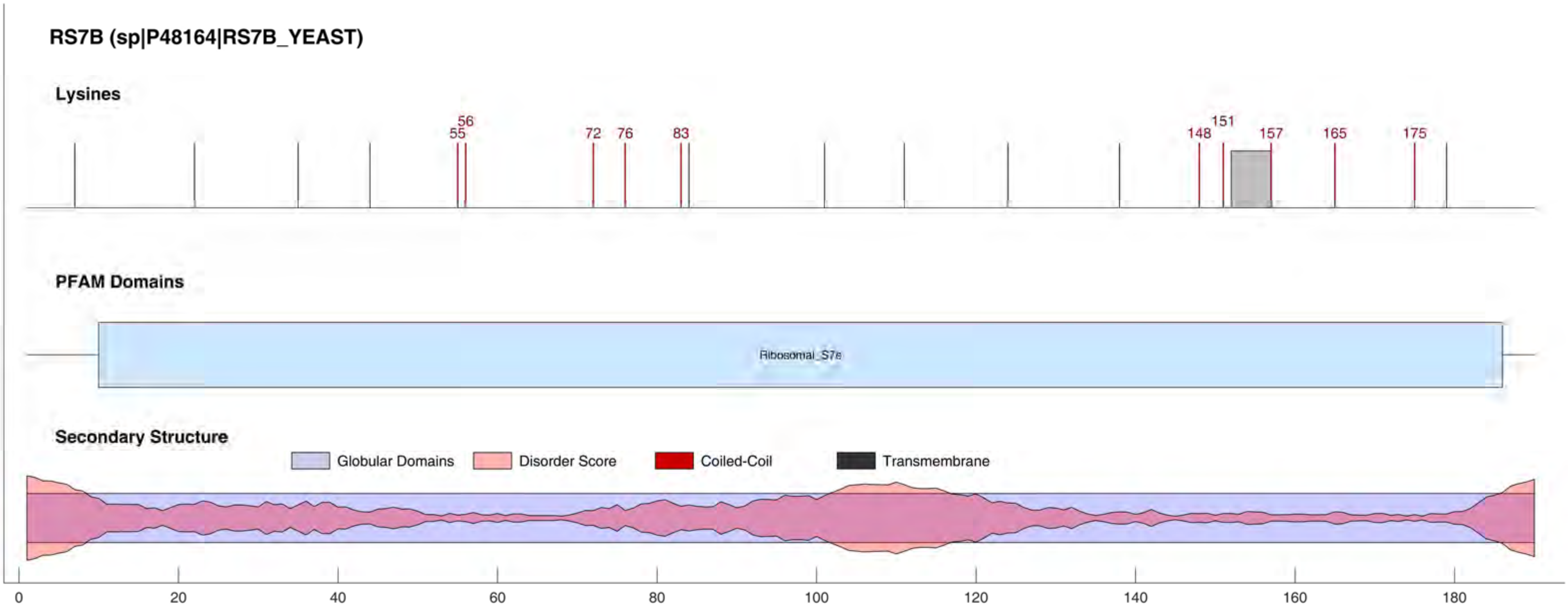
RS7B (sp|P48164|RS7B_YEAST)
Lysines
55
56
72
76
83
148
151
157
165
175
PFAM Domains
Ribosomal_S7e
Secondary Structure
Globular Domains
Disorder Score
Coiled-Coil
Transmembrane
0
20
40
60
80
100
120
140
160
180

RS8B (sp|P0CX40|RS8B_YEAST)

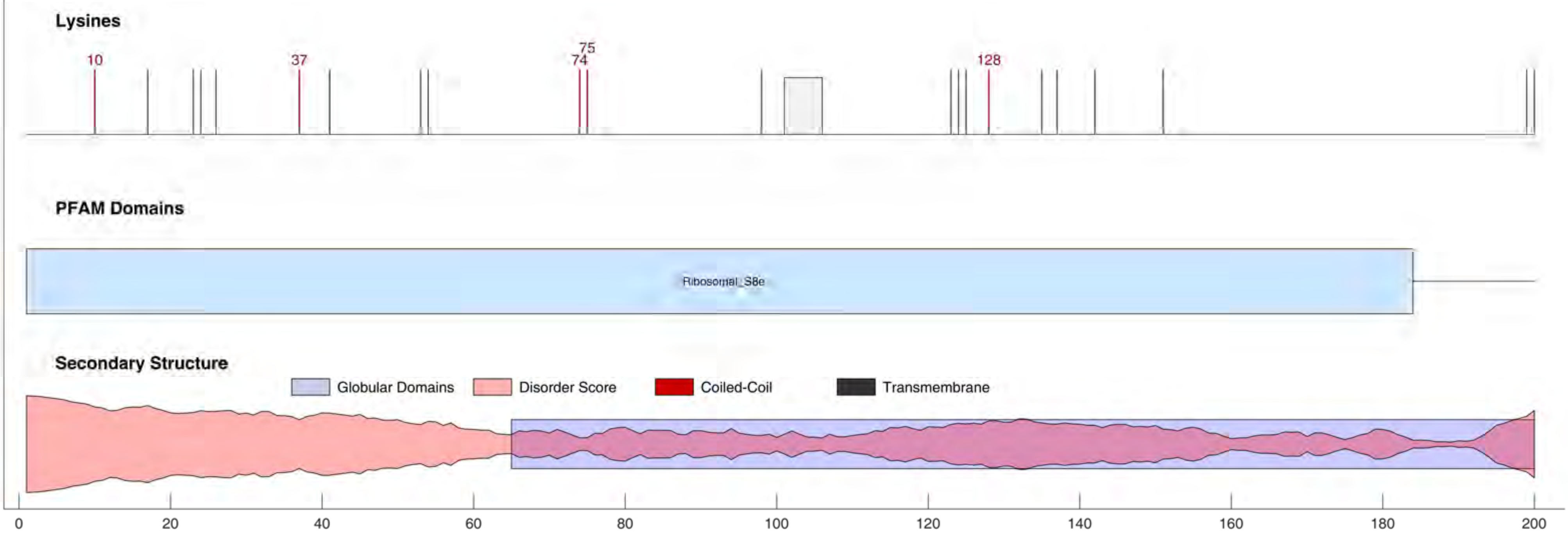

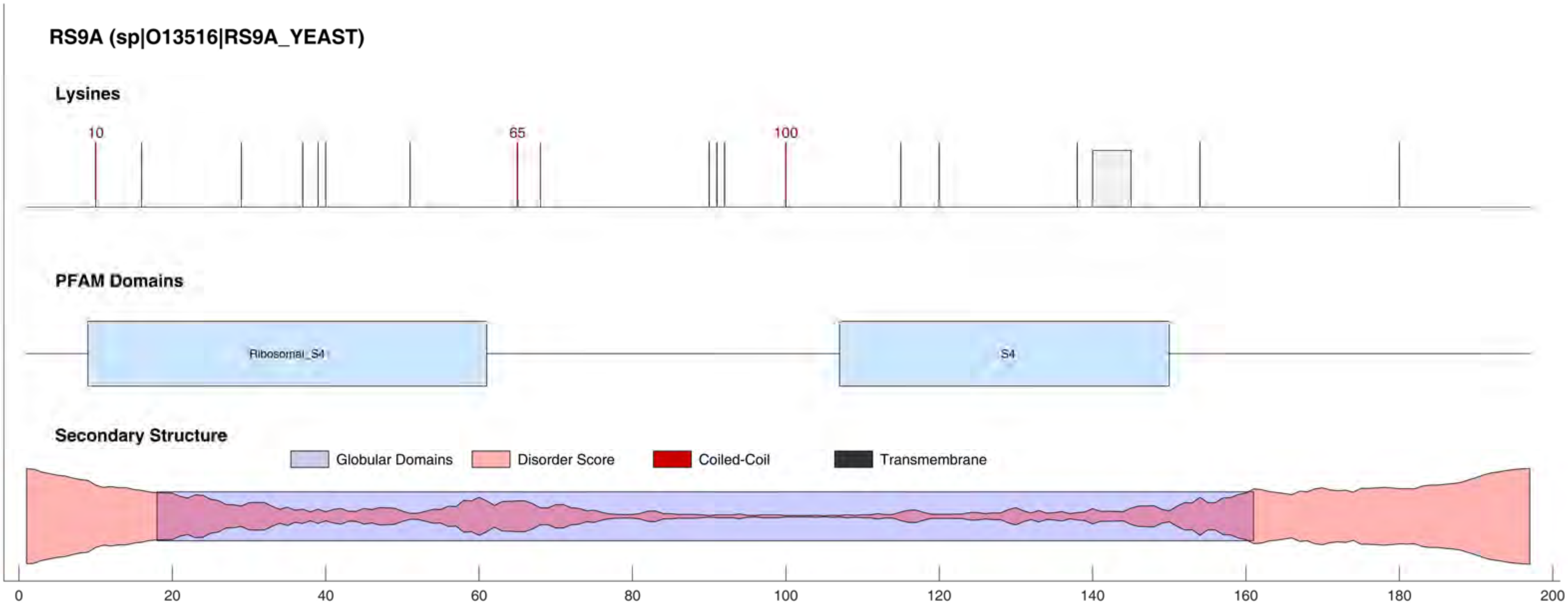
RS9A (sp|O13516|RS9A_YEAST)
Lysines
10
65
100
PFAM Domains
Ribosomal_S4
S4
Secondary Structure
Globular Domains
Disorder Score
Coiled-Coil
Transmembrane
0
20
40
60
80
100
120
140
160
180
200

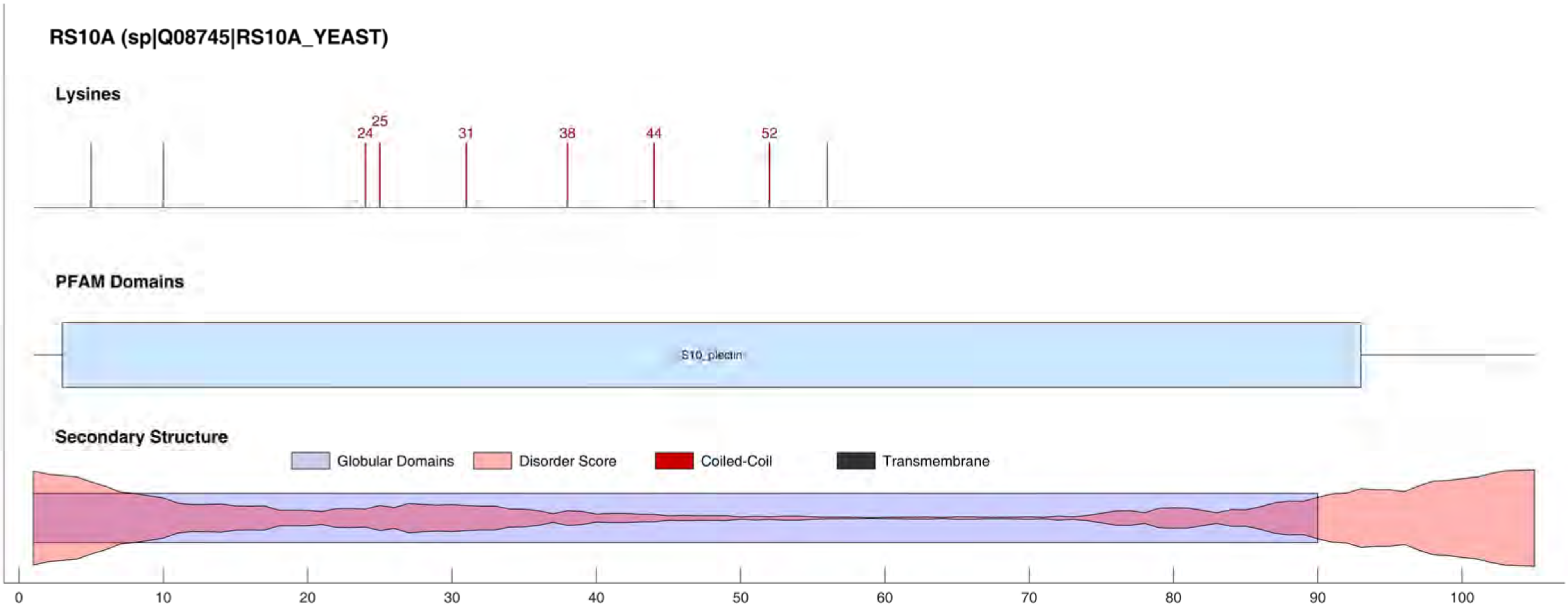

RS10A (sp|Q08745|RS10A_YEAST)
Lysines
24
25
31
38
44
52
PFAM Domains
S10_plectin
Secondary Structure
Globular Domains
Disorder Score
Coiled-Coil
Transmembrane
0
10
20
30
40
50
60
70
80
90
100

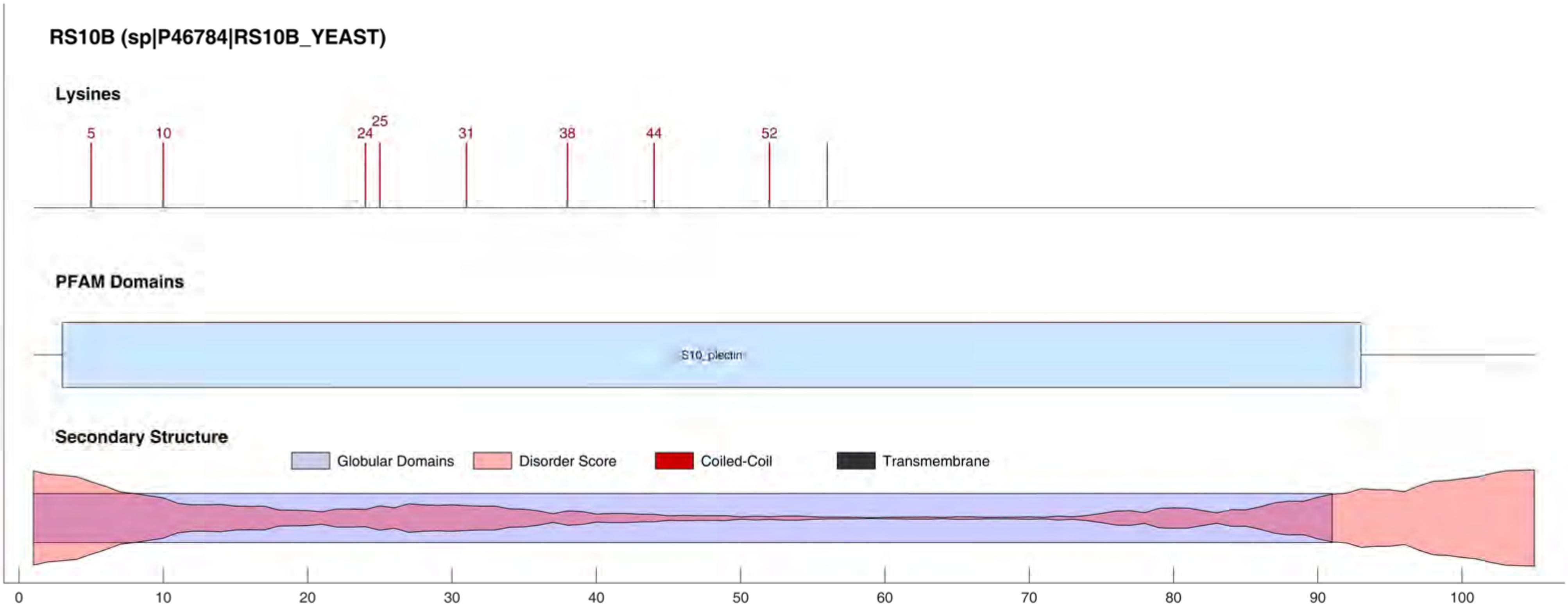

RS10B (sp|P46784|RS10B_YEAST)
Lysines
5
10
24
25
31
38
44
52
PFAM Domains
S10_plectin
Secondary Structure
Globular Domains
Disorder Score
Coiled-Coil
Transmembrane
0
10
20
30
40
50
60
70
80
90
100

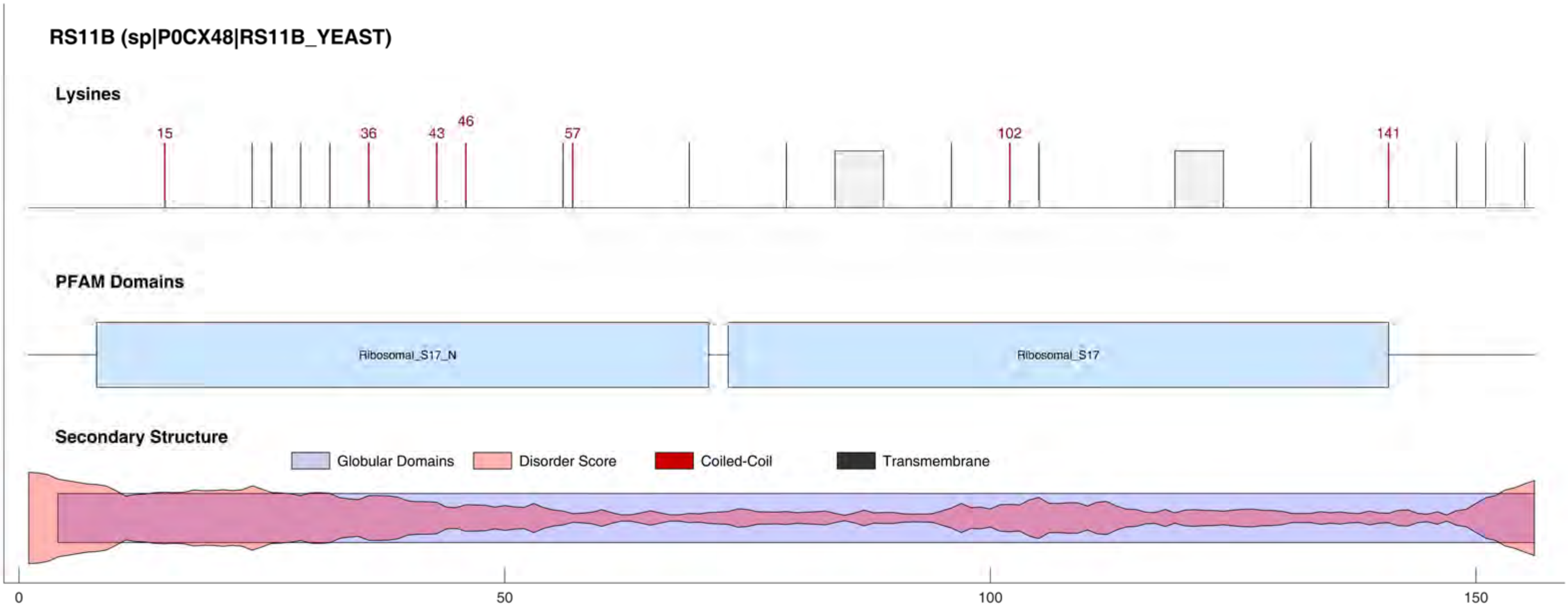
RS11B (sp|P0CX48|RS11B_YEAST)
Lysines
15
36
43
46
57
102
141
PFAM Domains
Ribosomal_S17_N
Ribosomal_S17
Secondary Structure
Globular Domains
Disorder Score
Coiled-Coil
Transmembrane
0
50
100
150

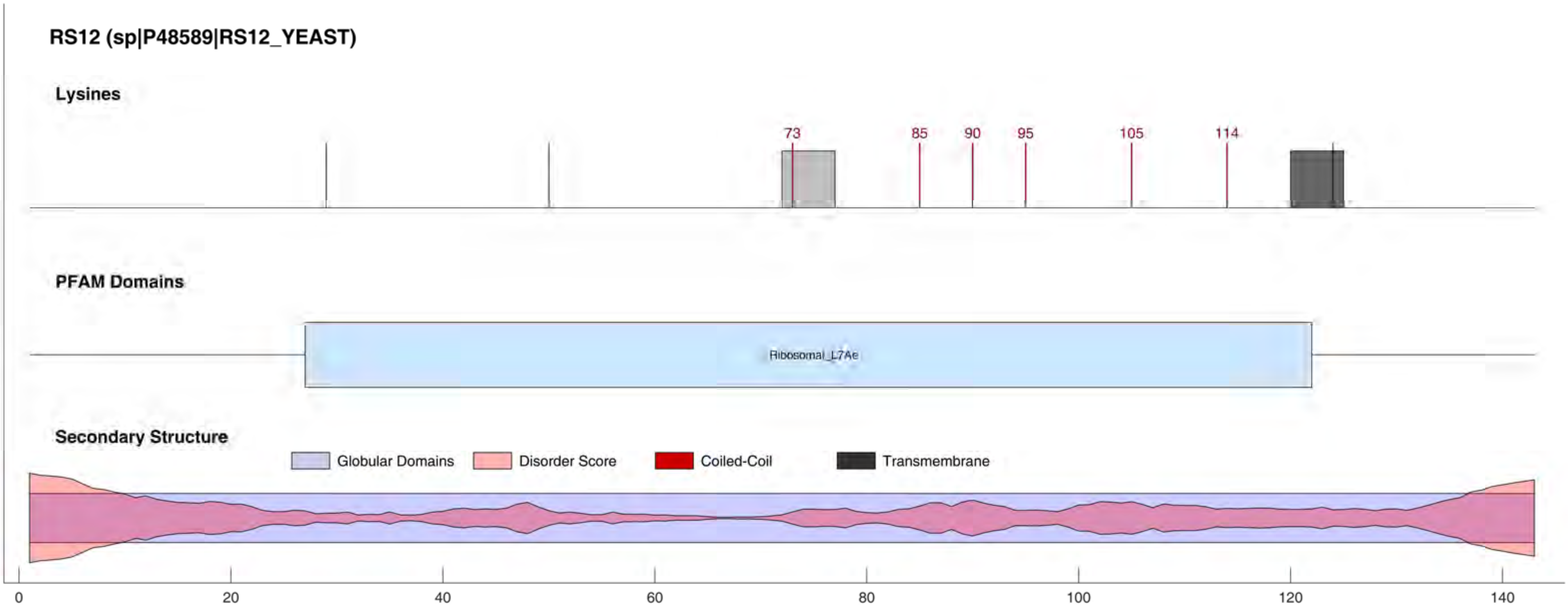
RS12 (sp|P48589|RS12_YEAST)
Lysines
73
85
90
95
105
114
PFAM Domains
Ribosomal_L7Ae
Secondary Structure
Globular Domains
Disorder Score
Coiled-Coil
Transmembrane
0
20
40
60
80
100
120
140

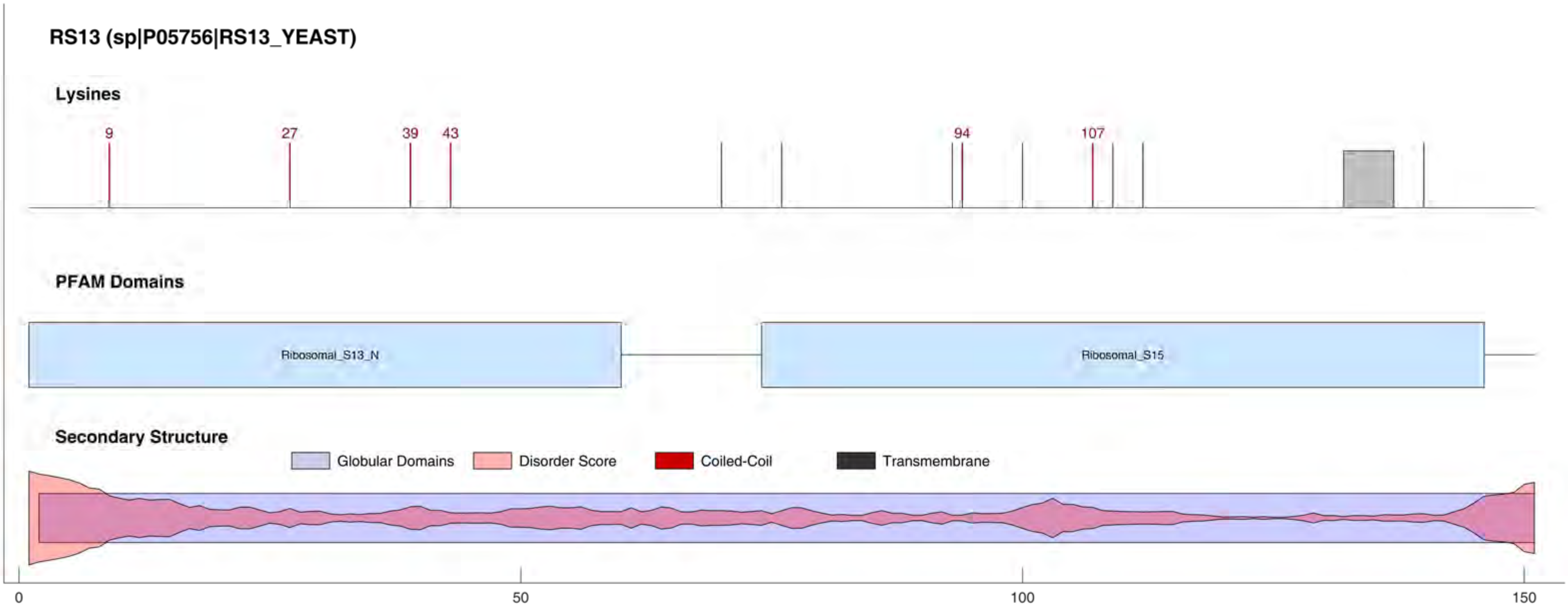
RS13 (sp|P05756|RS13_YEAST)
Lysines
9
27
39
43
94
107
PFAM Domains
Ribosomal_S13_N
Ribosomal_S15
Secondary Structure
Globular Domains
Disorder Score
Coiled-Coil
Transmembrane
0
50
100
150

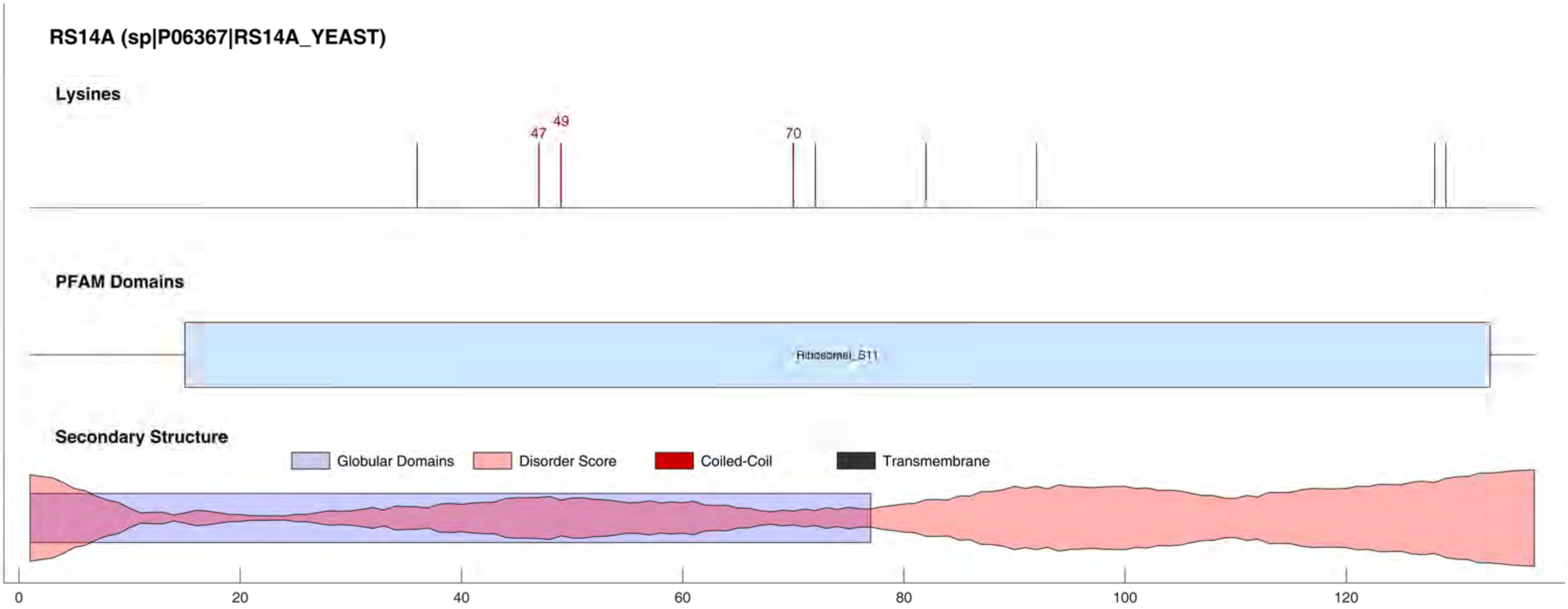

RS14A (sp|P06367|RS14A_YEAST)
Lysines
47
49
70
PFAM Domains
Ribosomal_S11
Secondary Structure
Globular Domains
Disorder Score
Coiled-Coil
Transmembrane
0
20
40
60
80
100
120

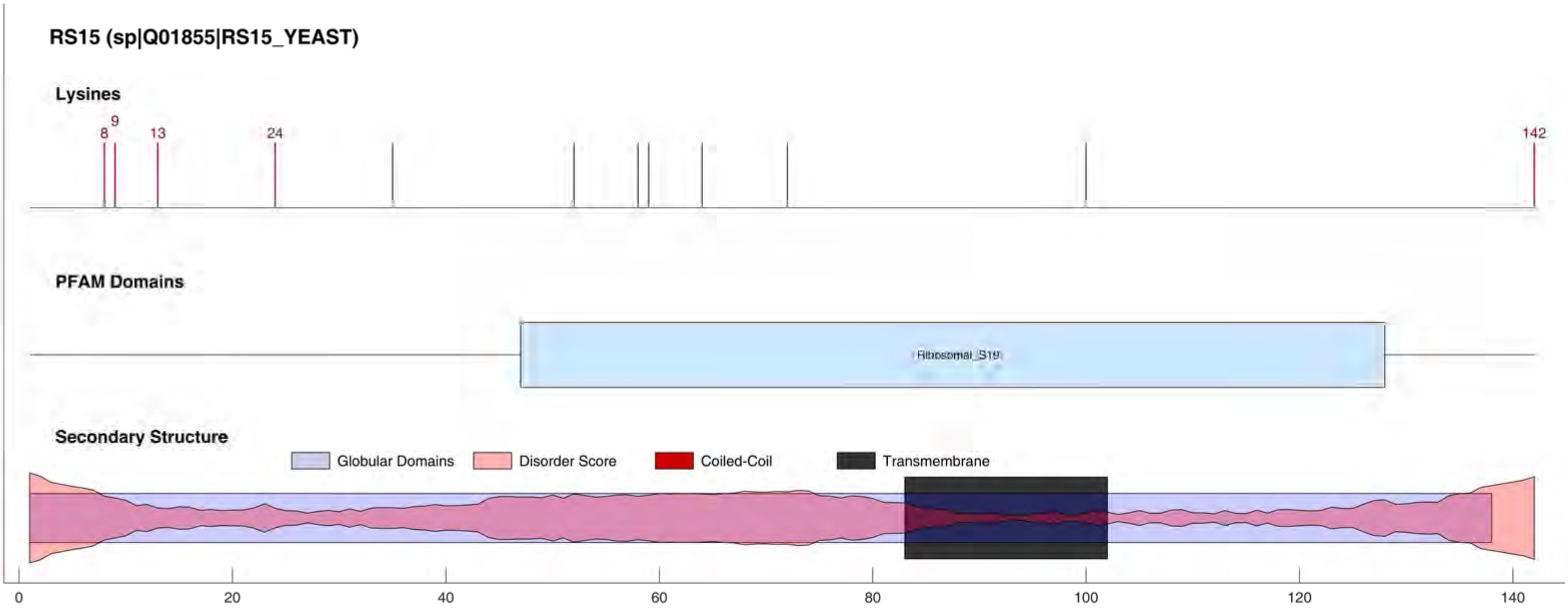
RS15 (sp|Q01855|RS15_YEAST)
Lysines
8
9
13
24
142
PFAM Domains
Ribosomal_S19
Secondary Structure
Globular Domains
Disorder Score
Coiled-Coil
Transmembrane
0
20
40
60
80
100
120
140

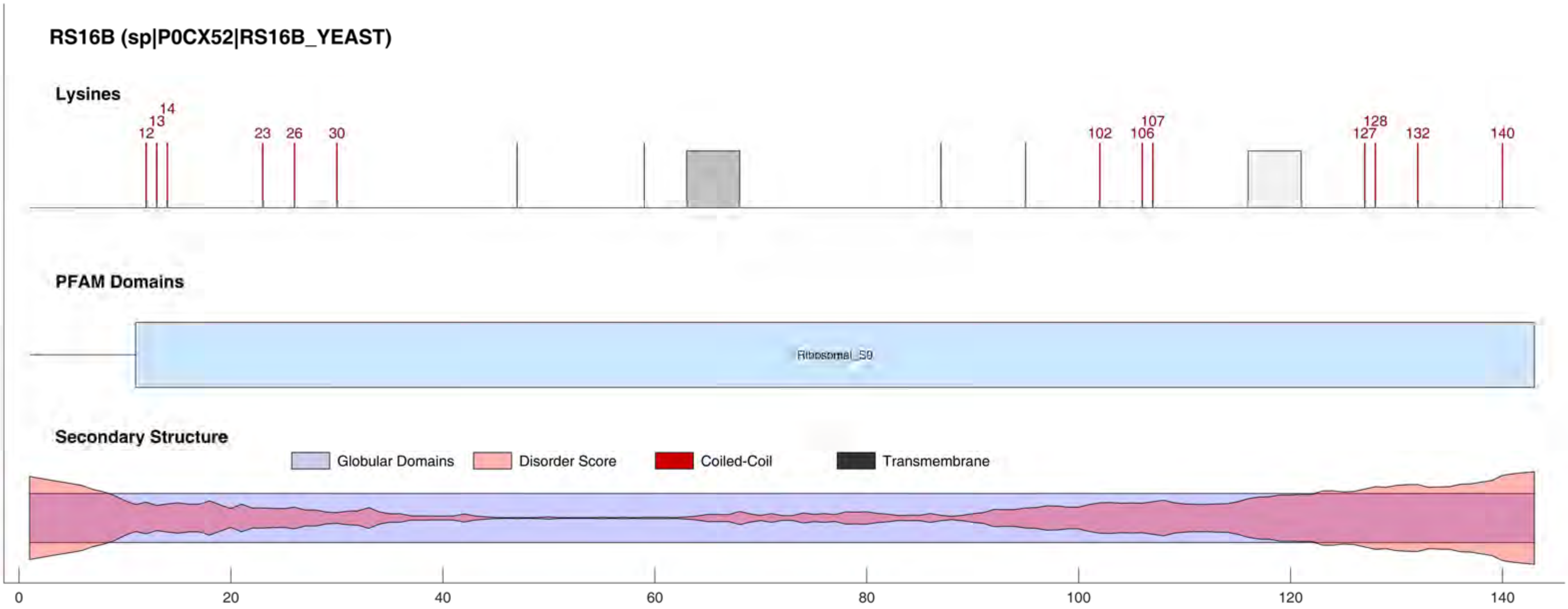

RS16B (sp|P0CX52|RS16B_YEAST)
Lysines
12
13
14
23
26
30
102
106
107
127
128
132
140
PFAM Domains
Ribosomal_S9
Secondary Structure
Globular Domains
Disorder Score
Coiled-Coil
Transmembrane
0
20
40
60
80
100
120
140

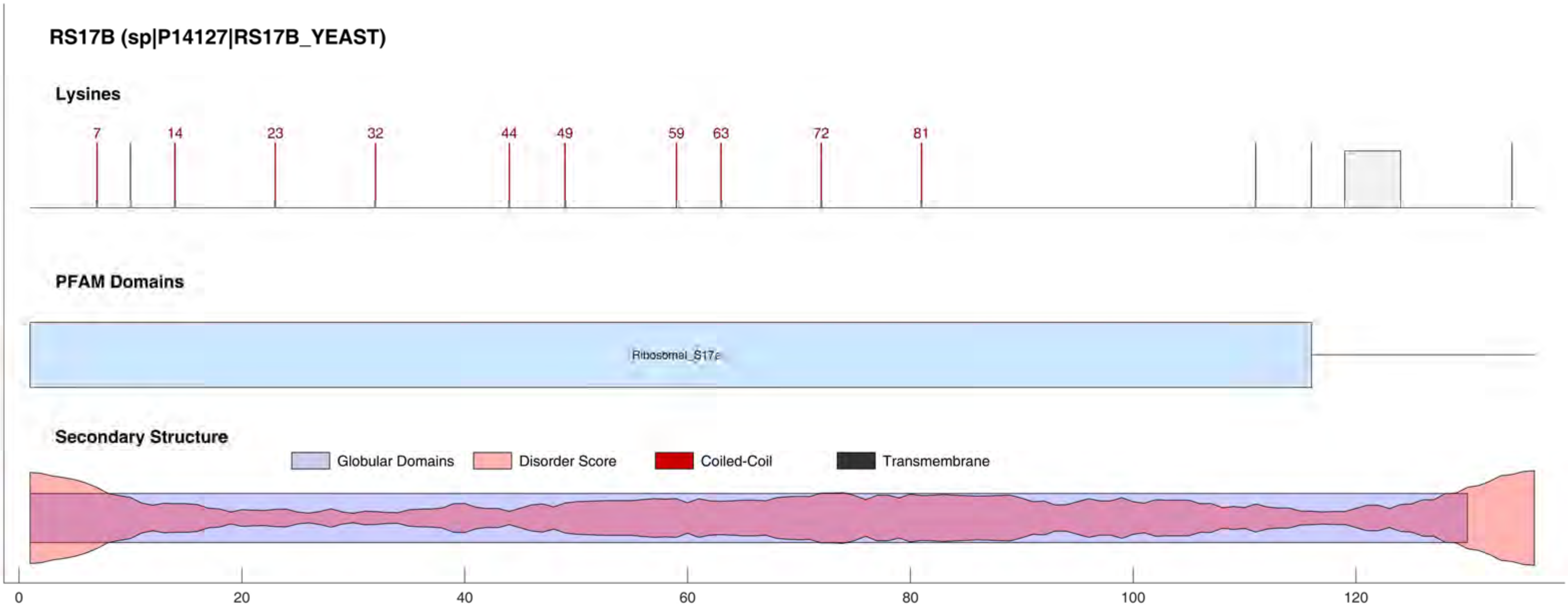

RS17B (sp|P14127|RS17B_YEAST)
Lysines
7
14
23
32
44
49
59
63
72
81
PFAM Domains
Ribosomal_S17e
Secondary Structure
Globular Domains
Disorder Score
Coiled-Coil
Transmembrane
0
20
40
60
80
100
120

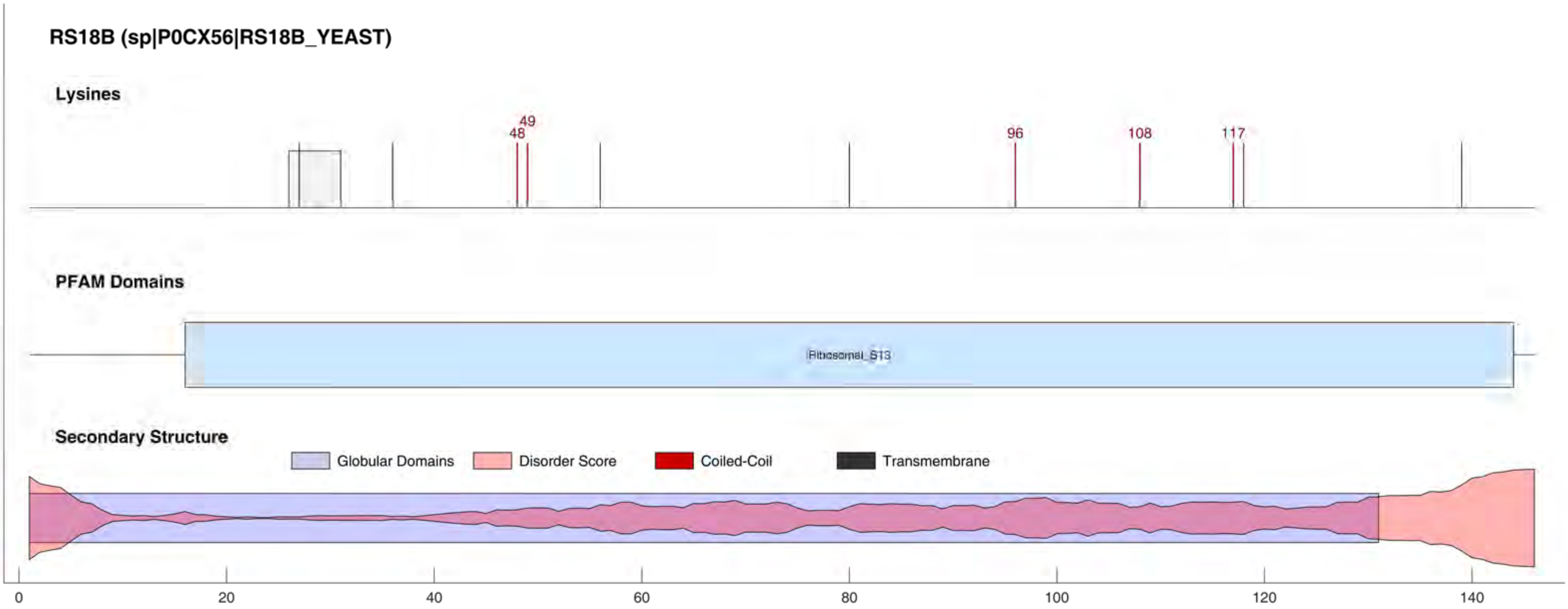

RS18B (sp|P0CX56|RS18B_YEAST)
Lysines
48
49
96
108
117
PFAM Domains
Ribosomal_S13
Secondary Structure
Globular Domains
Disorder Score
Coiled-Coil
Transmembrane
0
20
40
60
80
100
120
140

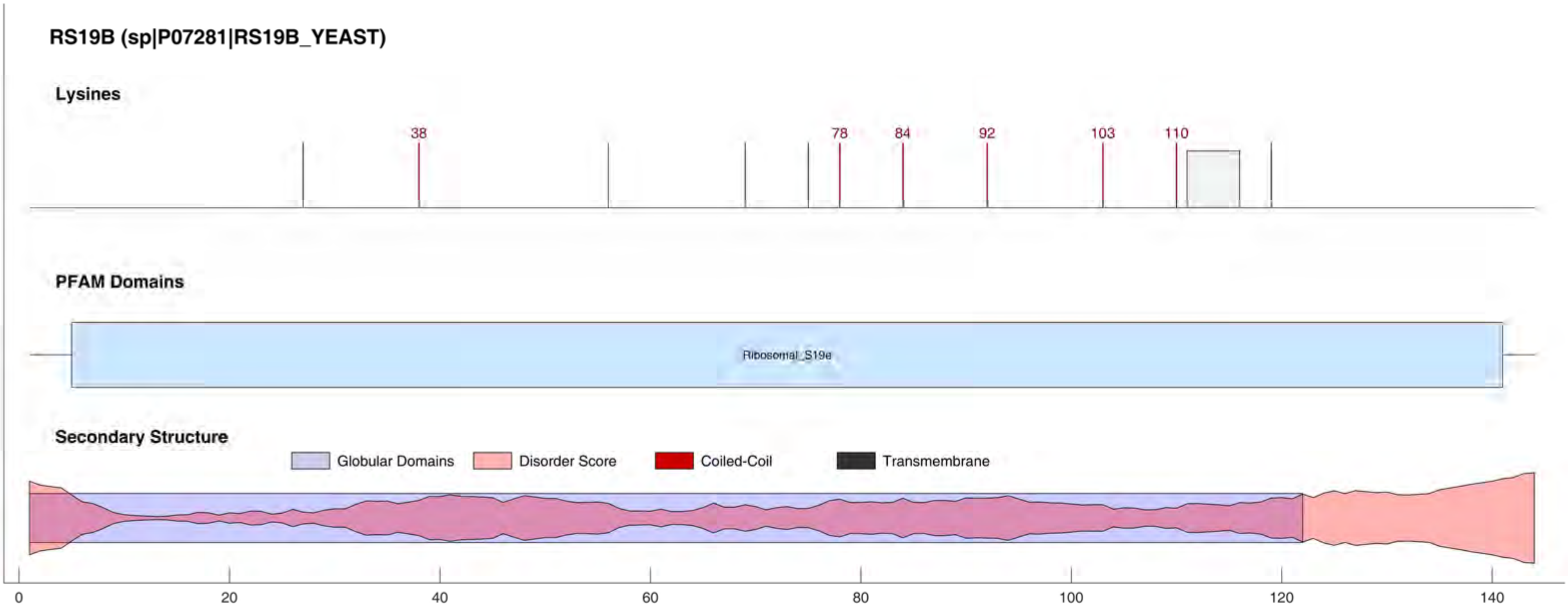
RS19B (sp|P07281|RS19B_YEAST)
Lysines
38
78
84
92
103
110
PFAM Domains
Ribosomal_S19e
Secondary Structure
Globular Domains
Disorder Score
Coiled-Coil
Transmembrane
0
20
40
60
80
100
120
140

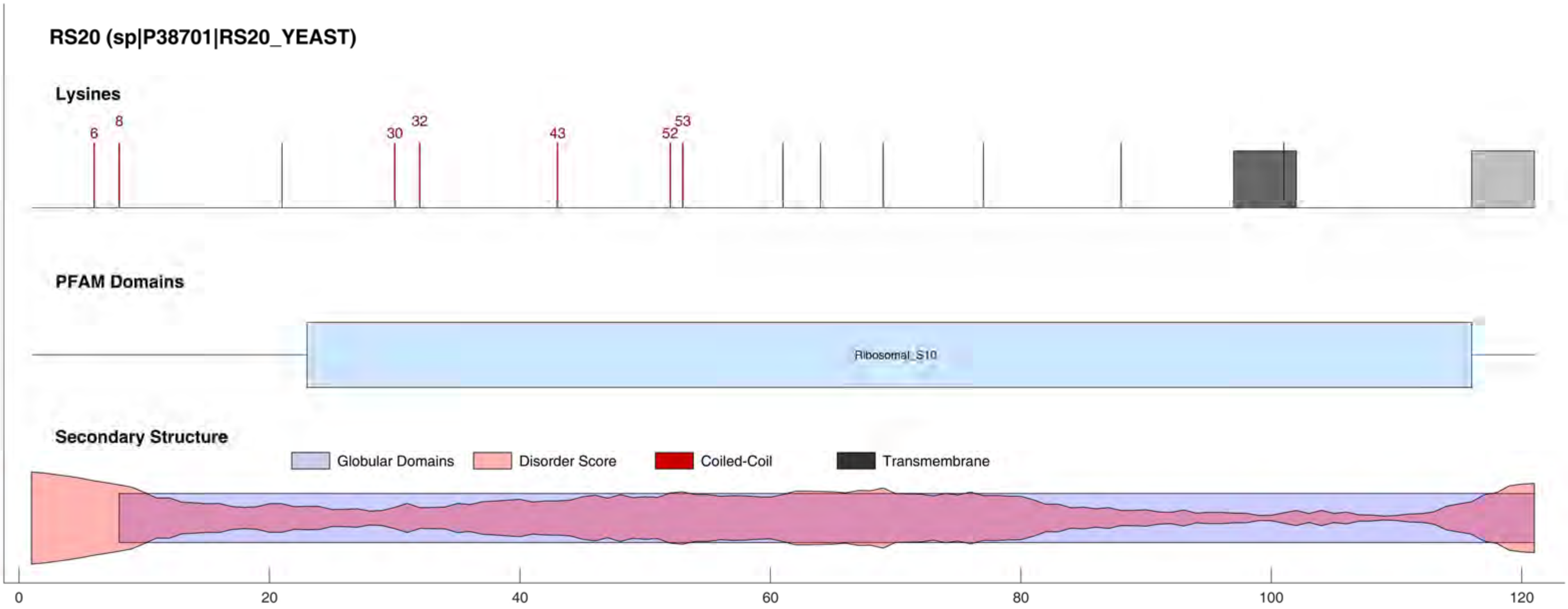
RS20 (sp|P38701|RS20_YEAST)
Lysines
6
8
30
32
43
52
53
PFAM Domains
Ribosomal_S10
Secondary Structure
Globular Domains
Disorder Score
Coiled-Coil
Transmembrane
0
20
40
60
80
100
120

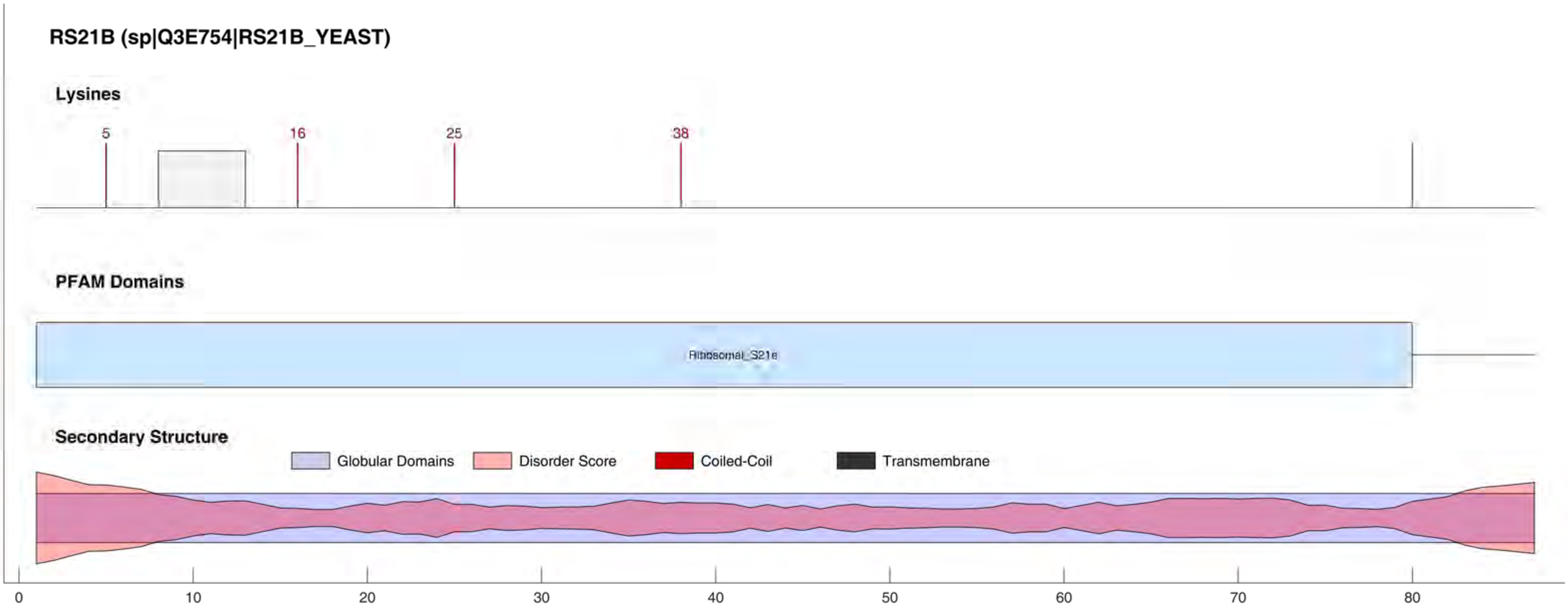
RS21B (sp|Q3E754|RS21B_YEAST)
Lysines
5
16
25
38
PFAM Domains
Ribosomal_S21e
Secondary Structure
Globular Domains
Disorder Score
Coiled-Coil
Transmembrane
0
10
20
30
40
50
60
70
80

#### RS22B (sp|Q3E7Y3|RS22B_YEAST)

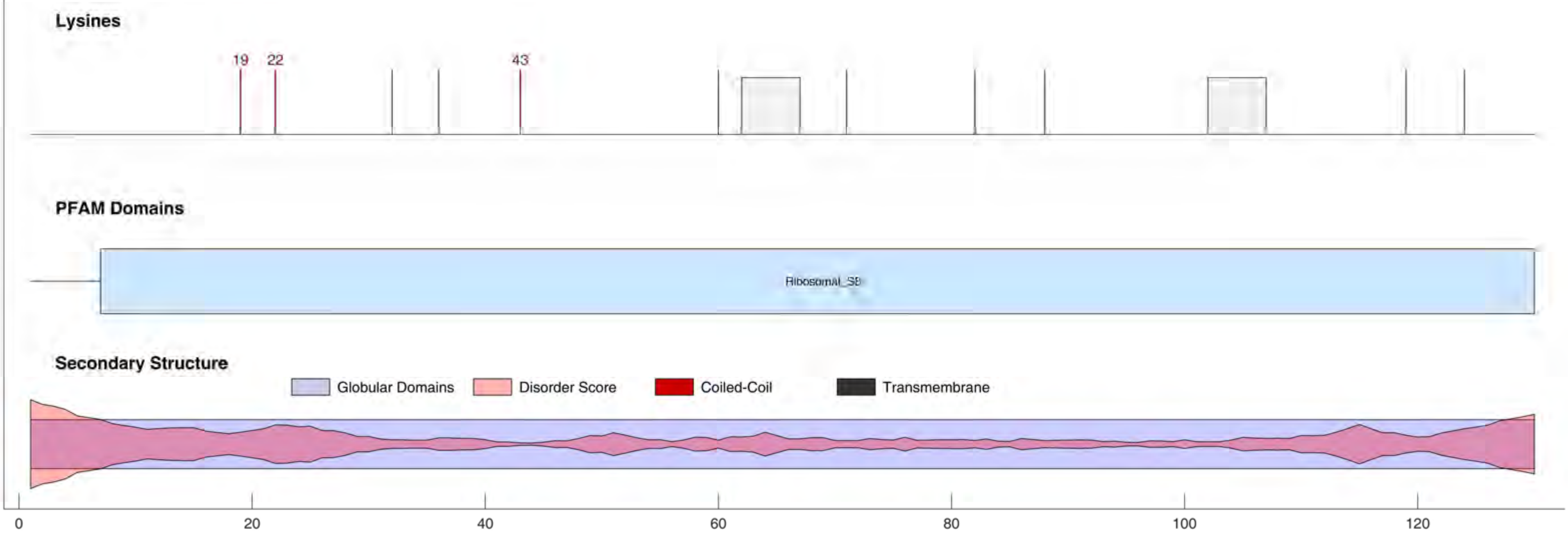

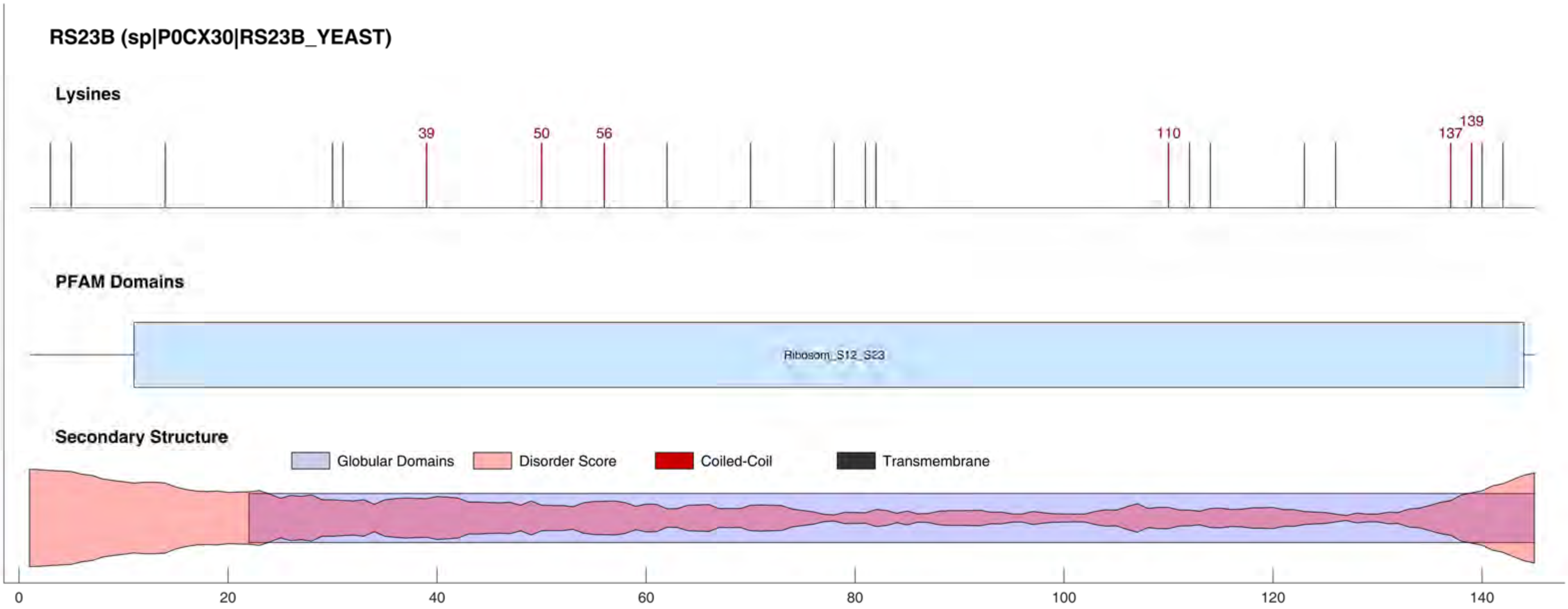

RS23B (sp|P0CX30|RS23B_YEAST)
Lysines
39
50
56
110
137
139
PFAM Domains
Ribosom_S12_S23
Secondary Structure
Globular Domains
Disorder Score
Coiled-Coil
Transmembrane
0
20
40
60
80
100
120
140

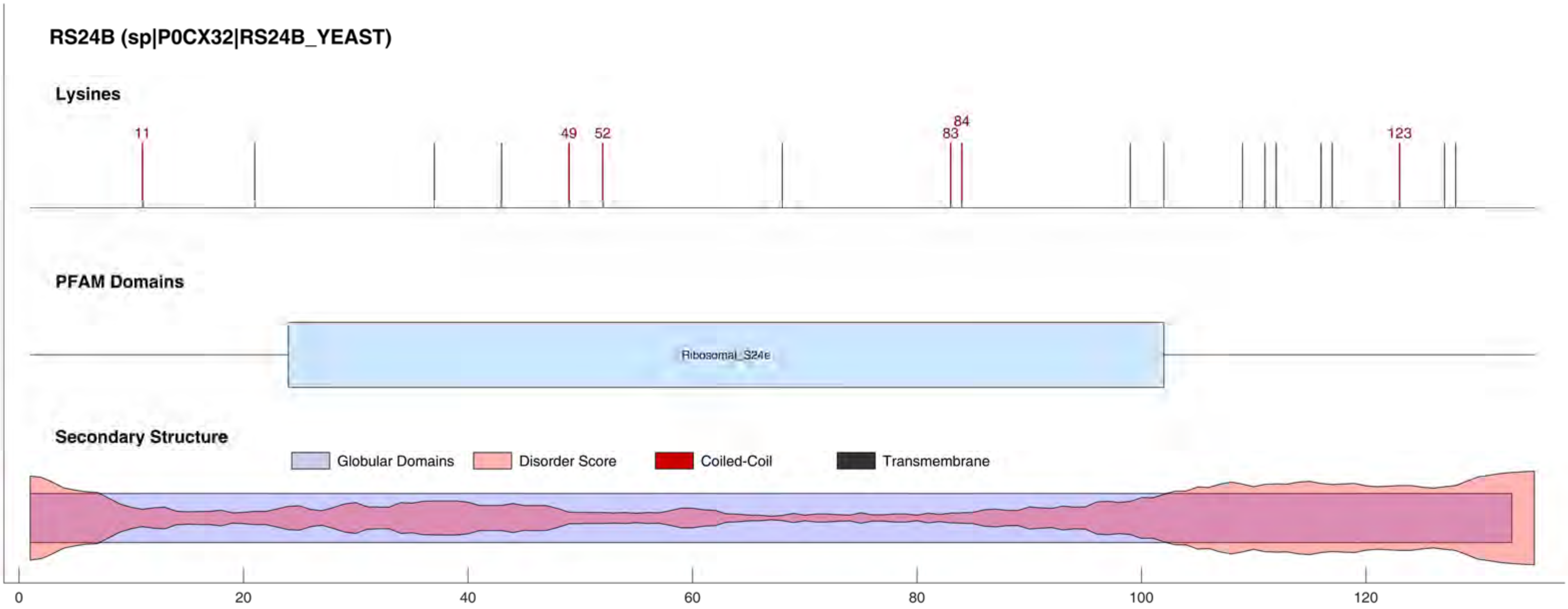
RS24B (sp|P0CX32|RS24B_YEAST)
Lysines
11
49
52
83
84
123
PFAM Domains
Ribosomal_S24e
Secondary Structure
Globular Domains
Disorder Score
Coiled-Coil
Transmembrane
0
20
40
60
80
100
120

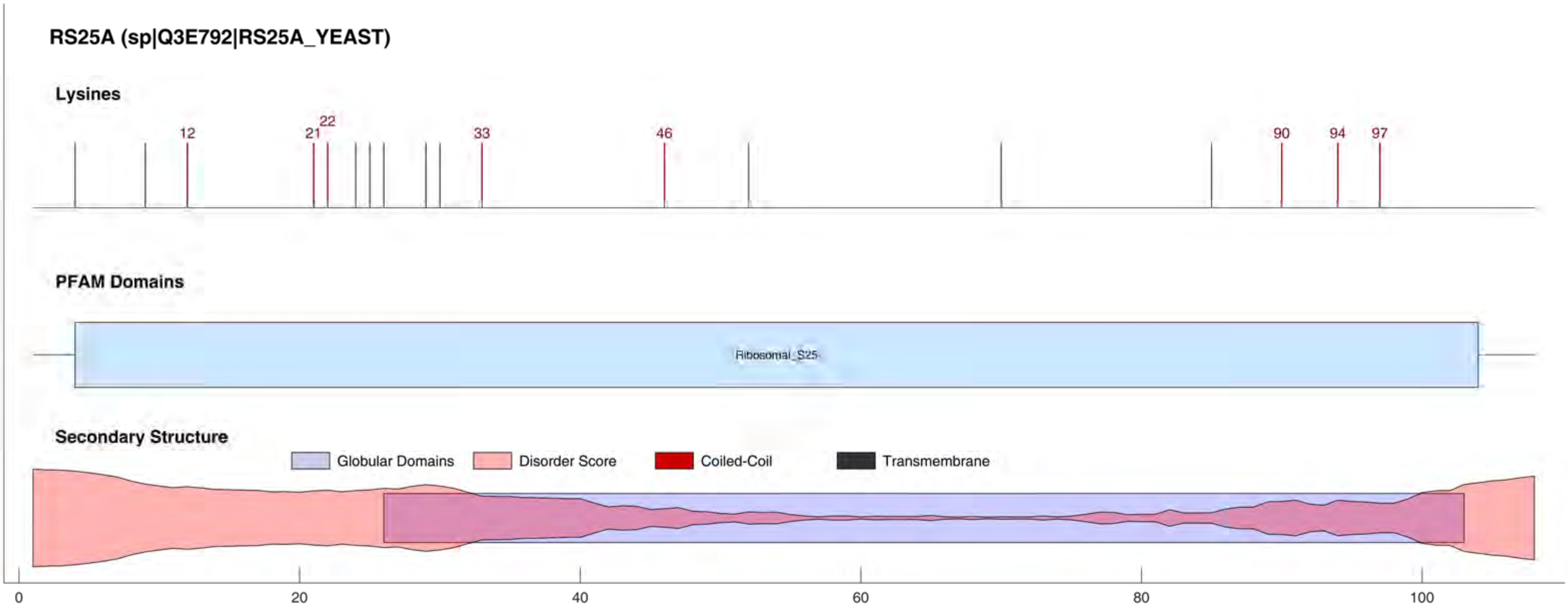

RS25A (sp|Q3E792|RS25A_YEAST)
Lysines
12
21
22
33
46
90
94
97
PFAM Domains
Ribosomal_S25
Secondary Structure
Globular Domains
Disorder Score
Coiled-Coil
Transmembrane
0
20
40
60
80
100

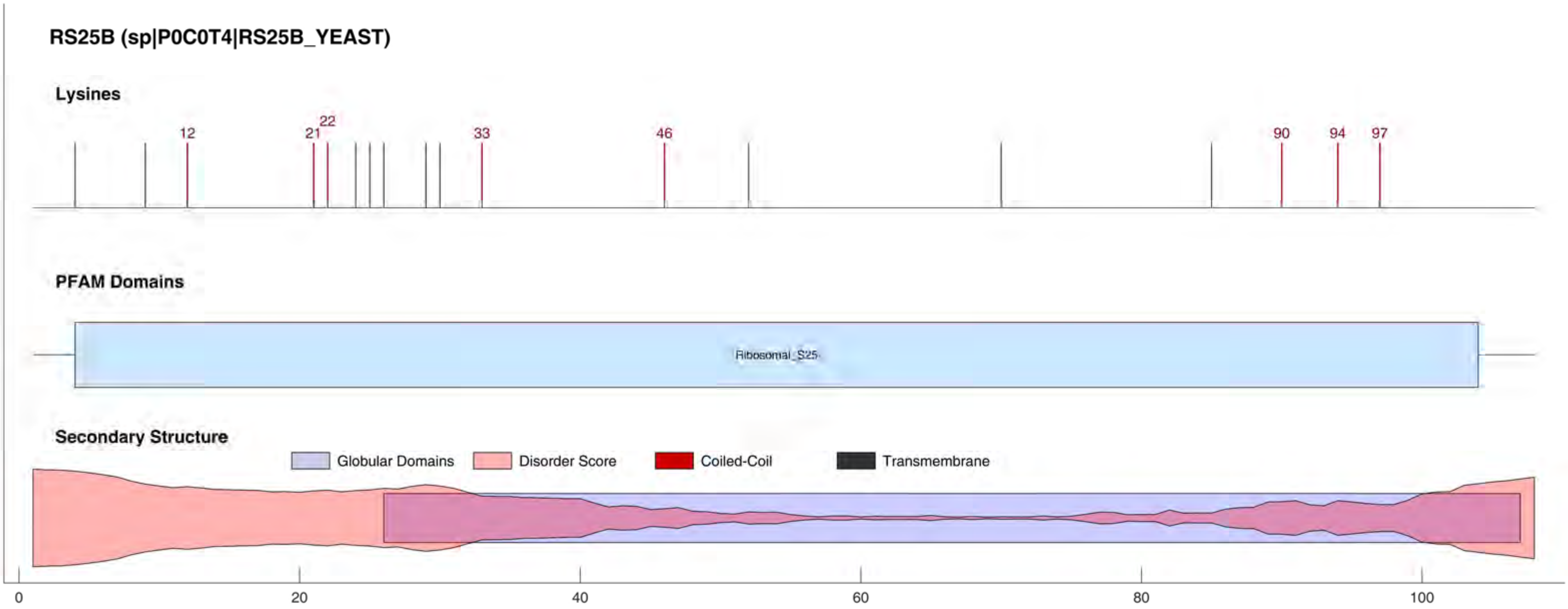
RS25B (sp|P0C0T4|RS25B_YEAST)
Lysines
12
21
22
33
46
90
94
97
PFAM Domains
Ribosomal_S25
Secondary Structure
Globular Domains
Disorder Score
Coiled-Coil
Transmembrane
0
20
40
60
80
100

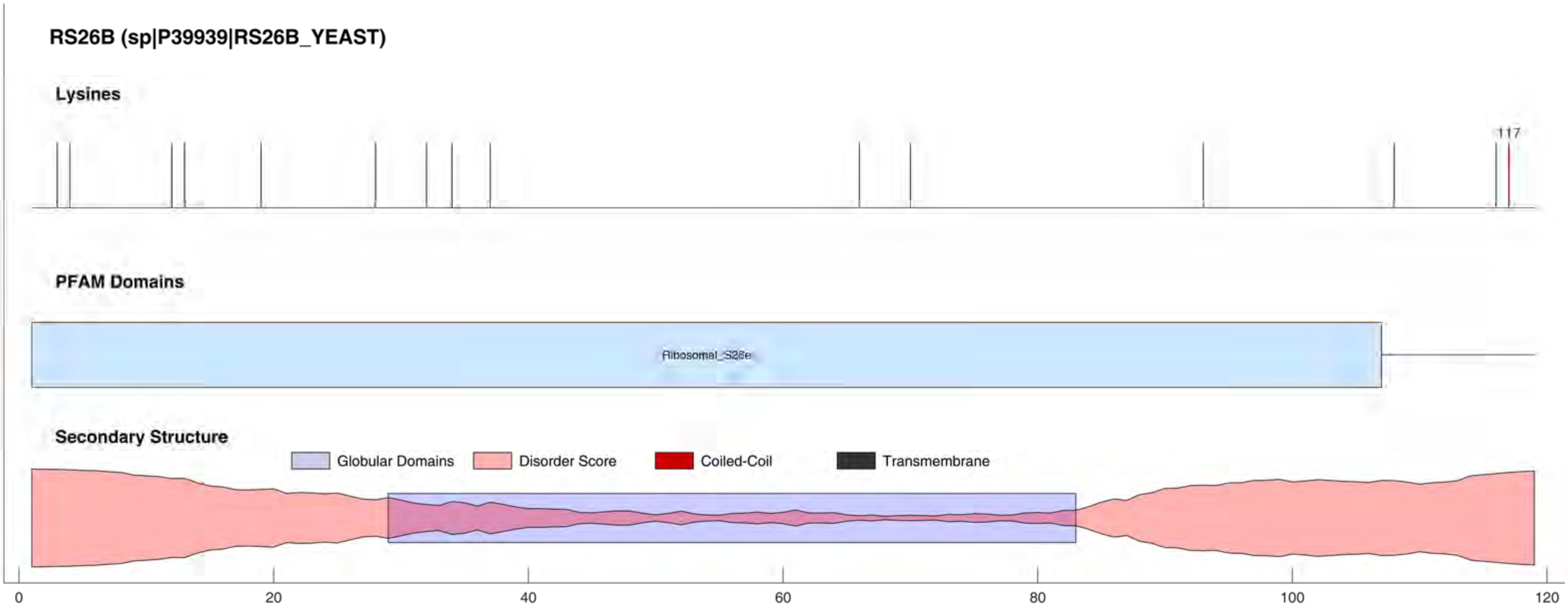

RS26B (sp|P39939|RS26B_YEAST)
Lysines
117
PFAM Domains
Ribosomal_S28e
Secondary Structure
Globular Domains
Disorder Score
Coiled-Coil
Transmembrane
0
20
40
60
80
100
120

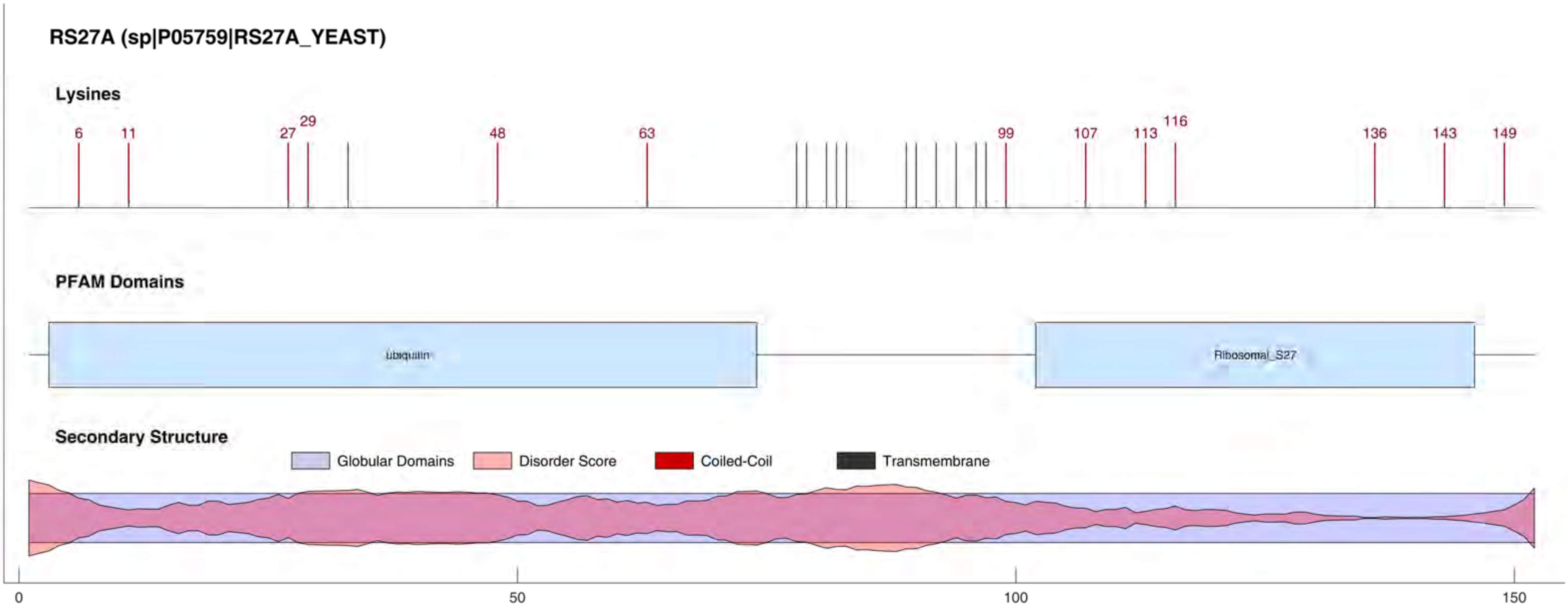

RS27A (sp|P05759|RS27A_YEAST)
Lysines
6
11
27
29
48
63
99
107
113
116
136
143
149
PFAM Domains
ubiquitin
Ribosomal_S27
Secondary Structure
Globular Domains
Disorder Score
Coiled-Coil
Transmembrane
0
50
100
150

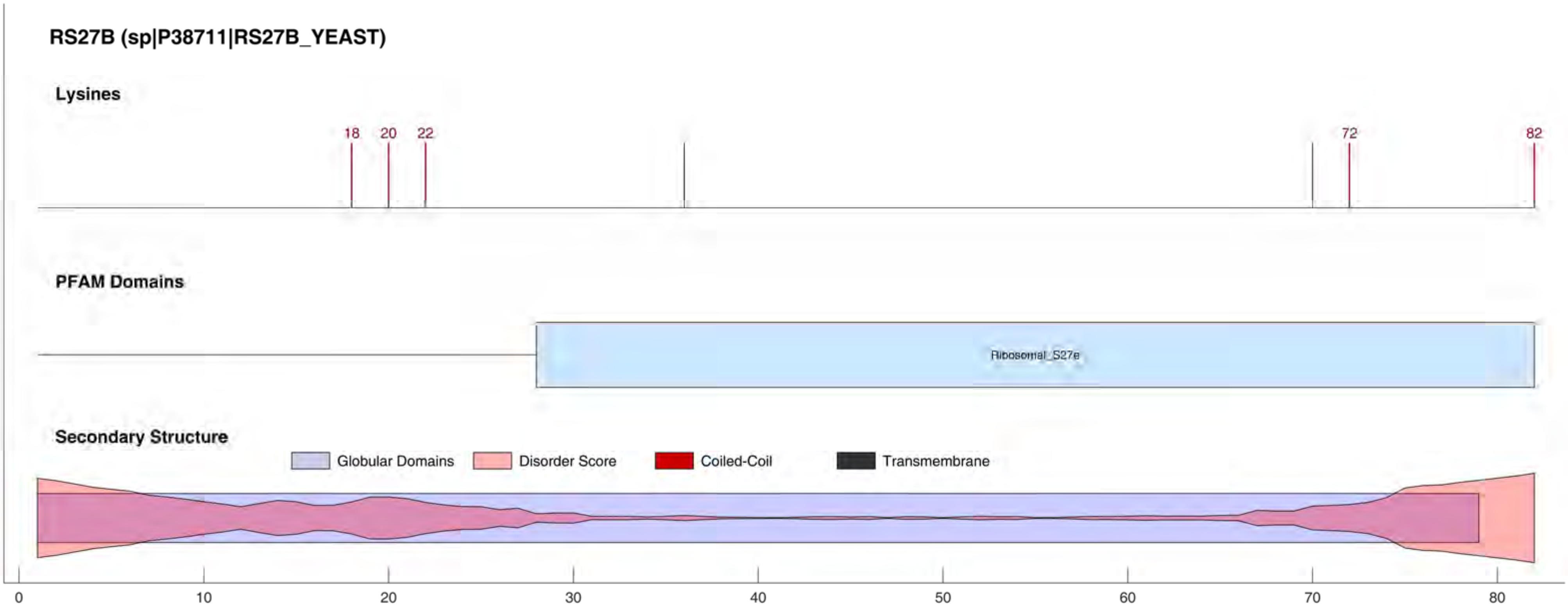
RS27B (sp|P38711|RS27B_YEAST)
Lysines
18
20
22
72
82
PFAM Domains
Ribosomal_S27e
Secondary Structure
Globular Domains
Disorder Score
Coiled-Coil
Transmembrane
0
10
20
30
40
50
60
70
80

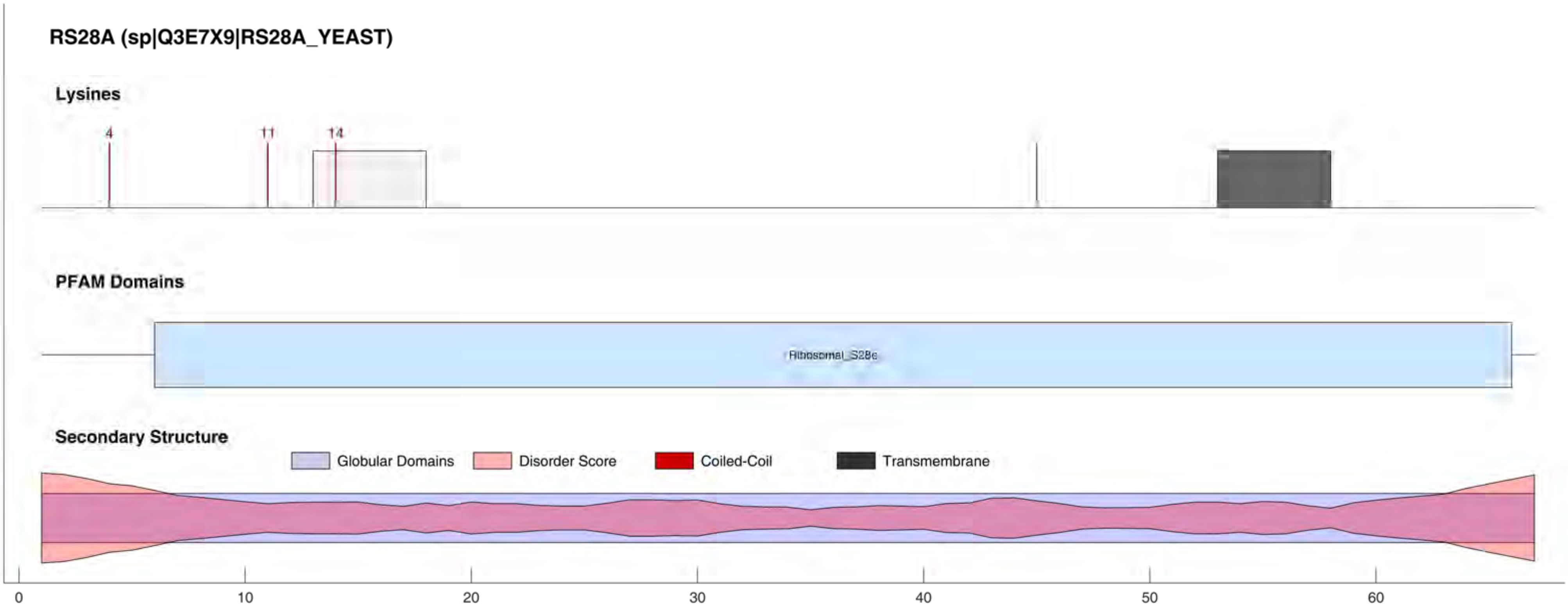

RS28A (sp|Q3E7X9|RS28A_YEAST)
Lysines
4
11
14
PFAM Domains
Ribosomal_S28e
Secondary Structure
Globular Domains
Disorder Score
Coiled-Coil
Transmembrane
0
10
20
30
40
50
60

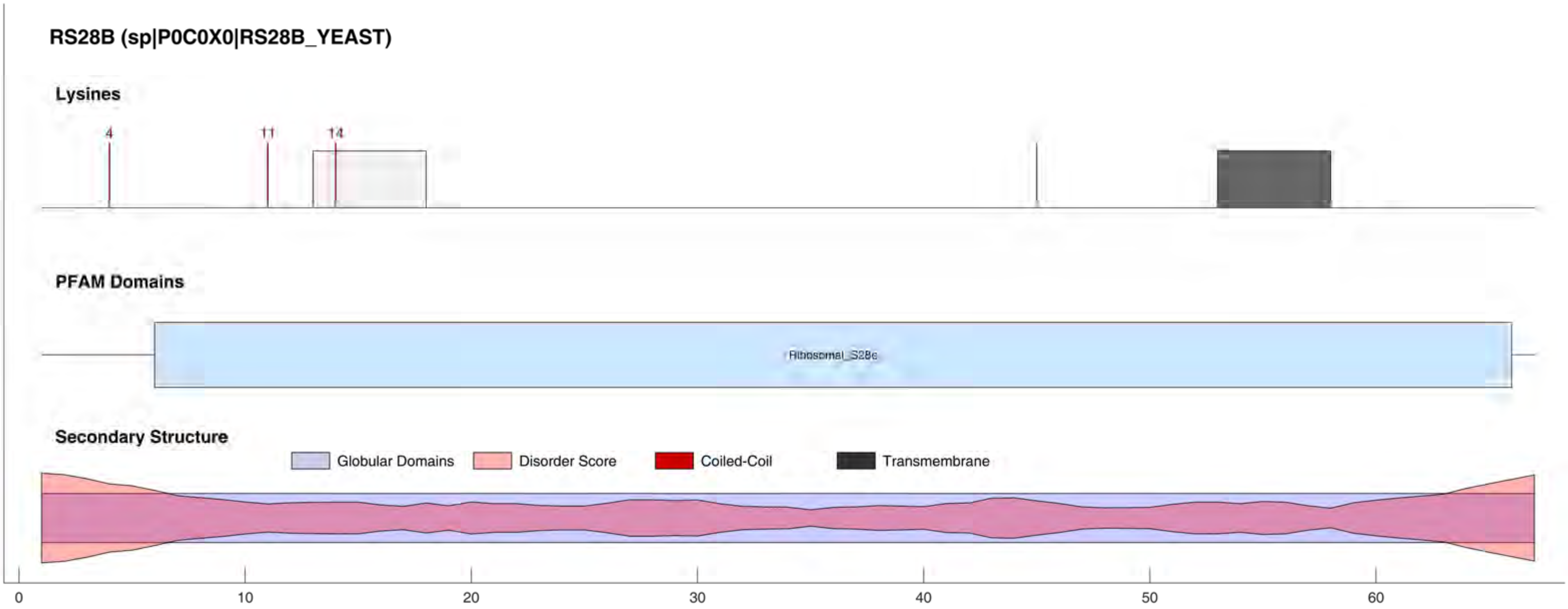

RS28B (sp|P0C0X0|RS28B_YEAST)
Lysines
4
11
14
PFAM Domains
Ribosomal_S28e
Secondary Structure
Globular Domains
Disorder Score
Coiled-Coil
Transmembrane
0
10
20
30
40
50
60

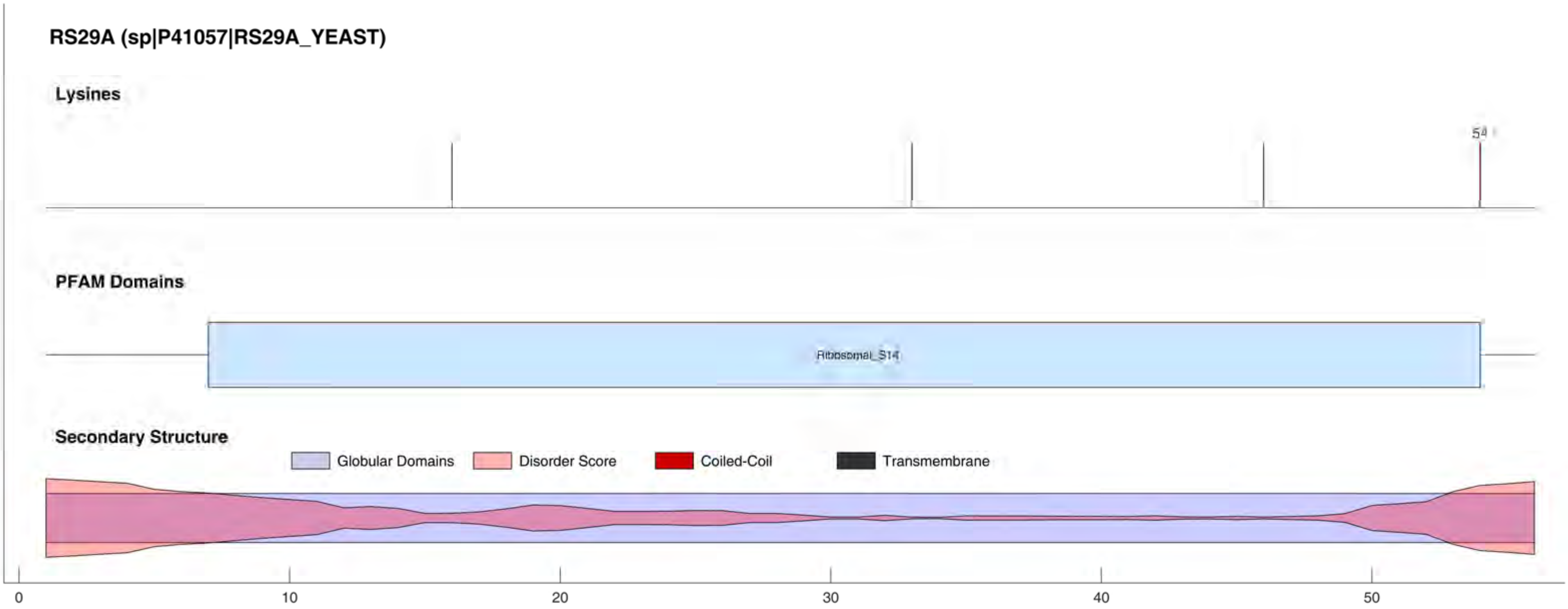
RS29A (sp|P41057|RS29A_YEAST)
Lysines
54
PFAM Domains
Ribosomal_S14
Secondary Structure
Globular Domains
Disorder Score
Coiled-Coil
Transmembrane
0
10
20
30
40
50

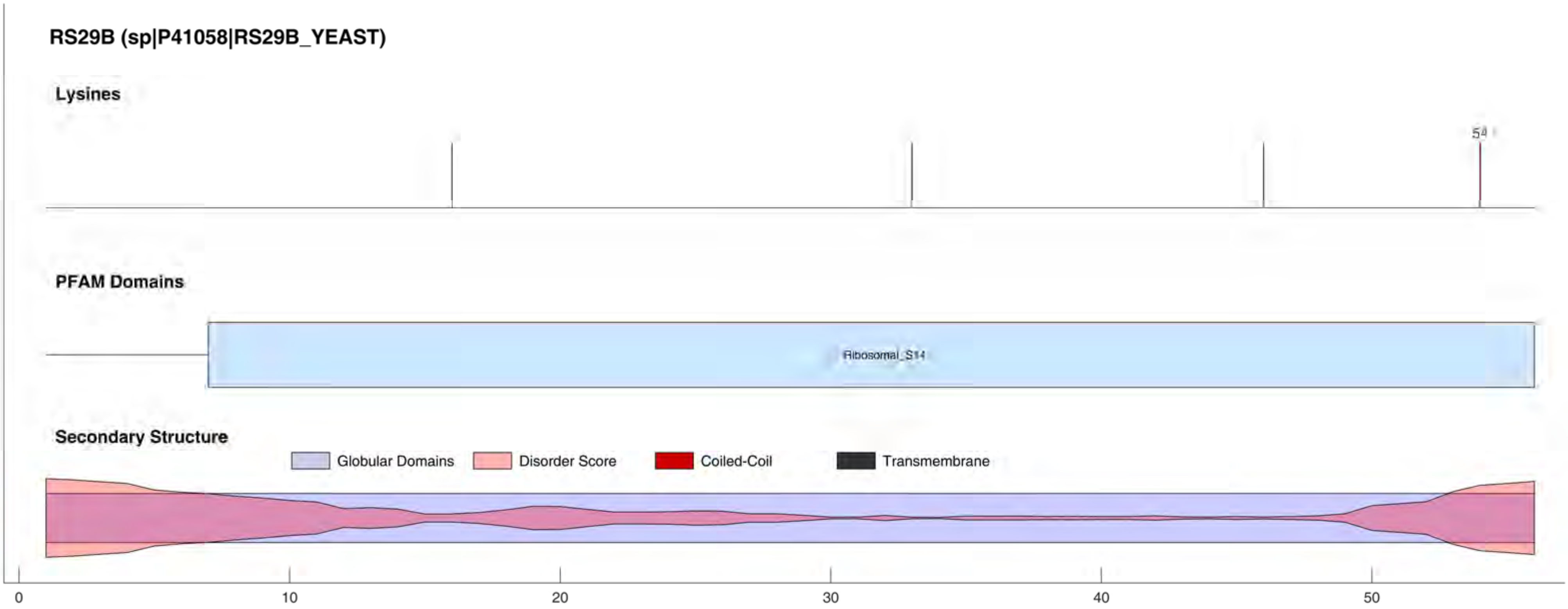
RS29B (sp|P41058|RS29B_YEAST)
Lysines
54
PFAM Domains
Ribosomal_S14
Secondary Structure
Globular Domains
Disorder Score
Coiled-Coil
Transmembrane
0
10
20
30
40
50

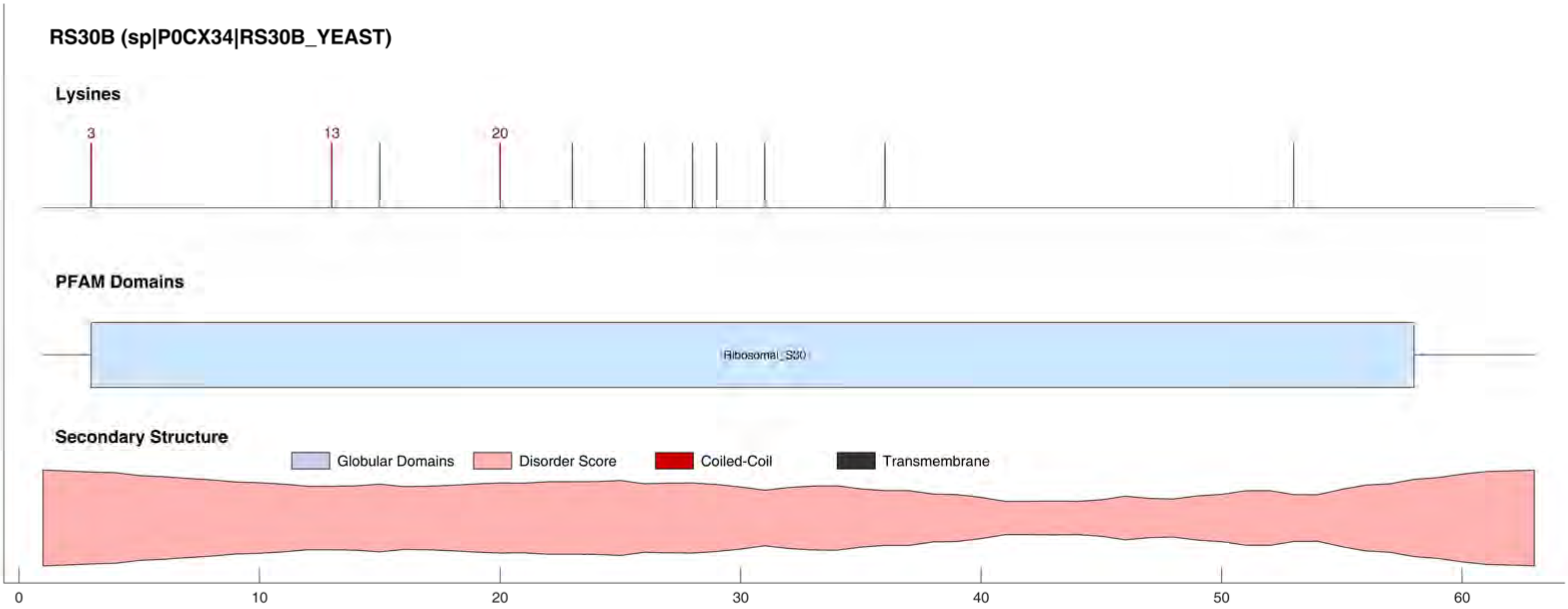
RS30B (sp|P0CX34|RS30B_YEAST)
Lysines
3
13
20
PFAM Domains
Ribosomal_S30
Secondary Structure
Globular Domains
Disorder Score
Coiled-Coil
Transmembrane
0
10
20
30
40
50
60

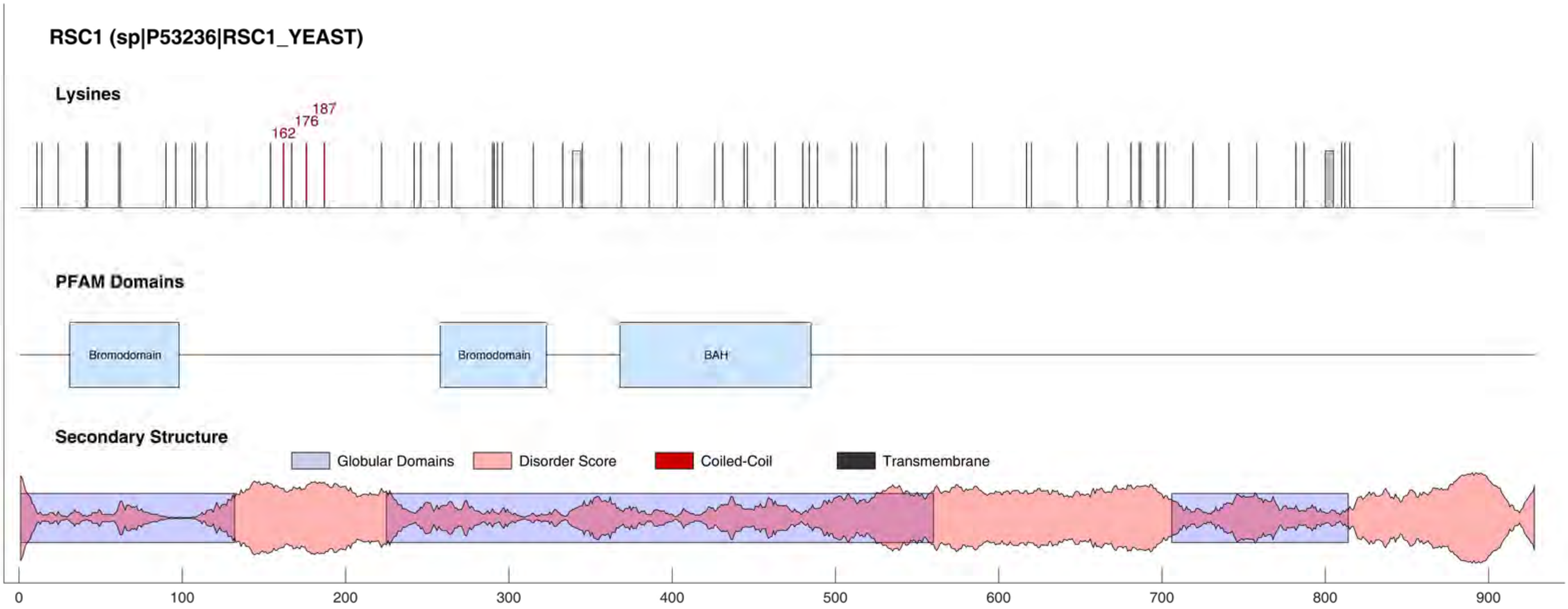

RSC1 (sp|P53236|RSC1_YEAST)
Lysines
162
176
187
PFAM Domains
Bromodomain
Bromodomain
BAH
Secondary Structure
Globular Domains
Disorder Score
Coiled-Coil
Transmembrane
0
100
200
300
400
500
600
700
800
900

RSC2 (sp|Q06488|RSC2_YEAST)

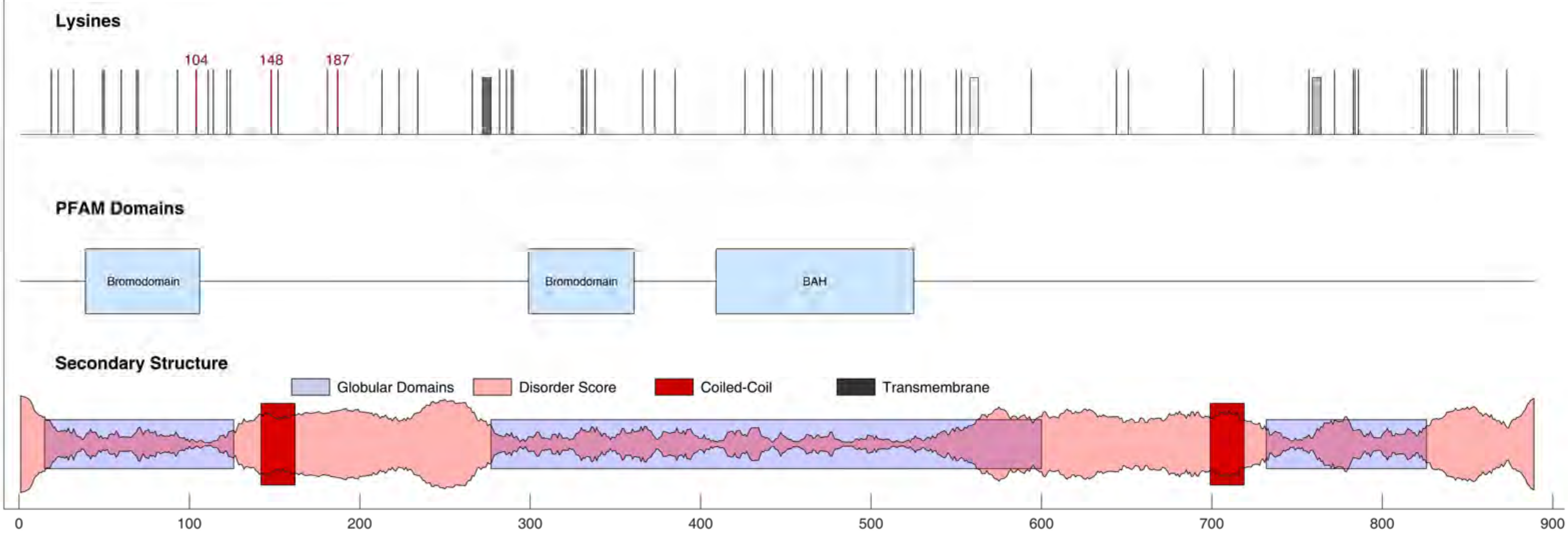

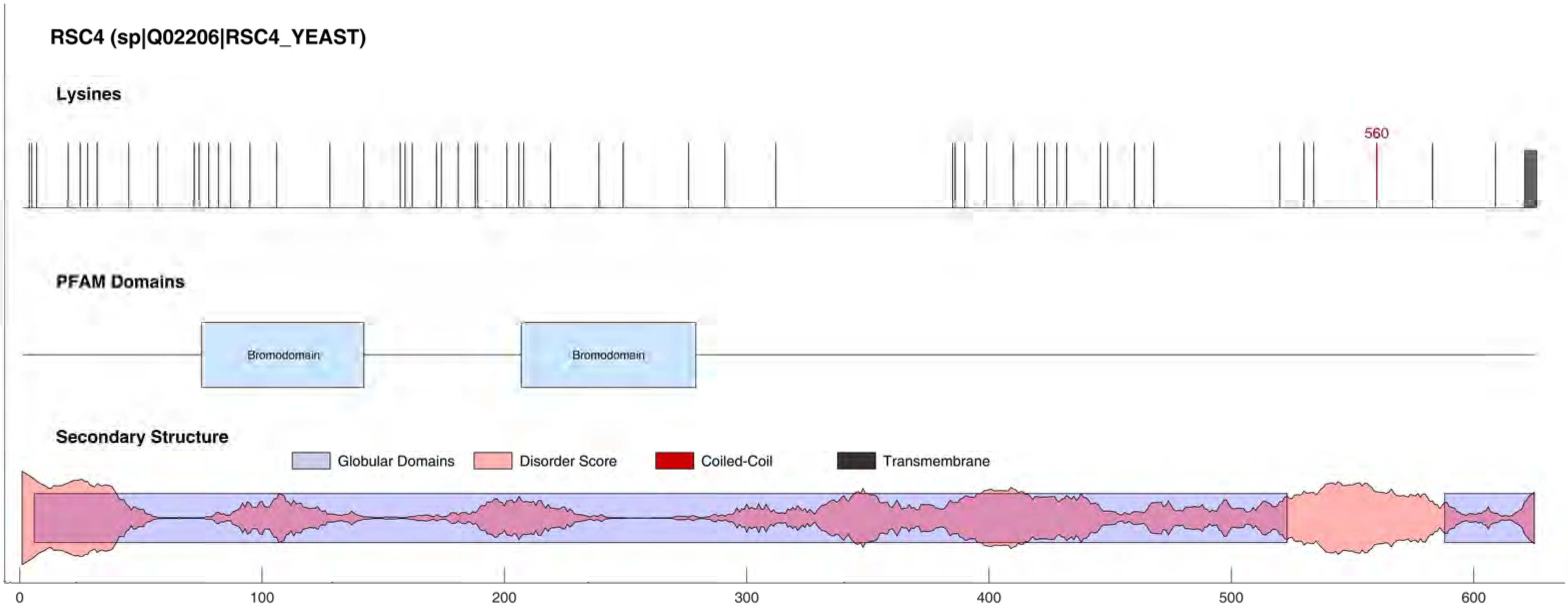
RSC4 (sp|Q02206|RSC4_YEAST)
Lysines
560
PFAM Domains
Bromodomain
Bromodomain
Secondary Structure
Globular Domains
Disorder Score
Coiled-Coil
Transmembrane
0
100
200
300
400
500
600

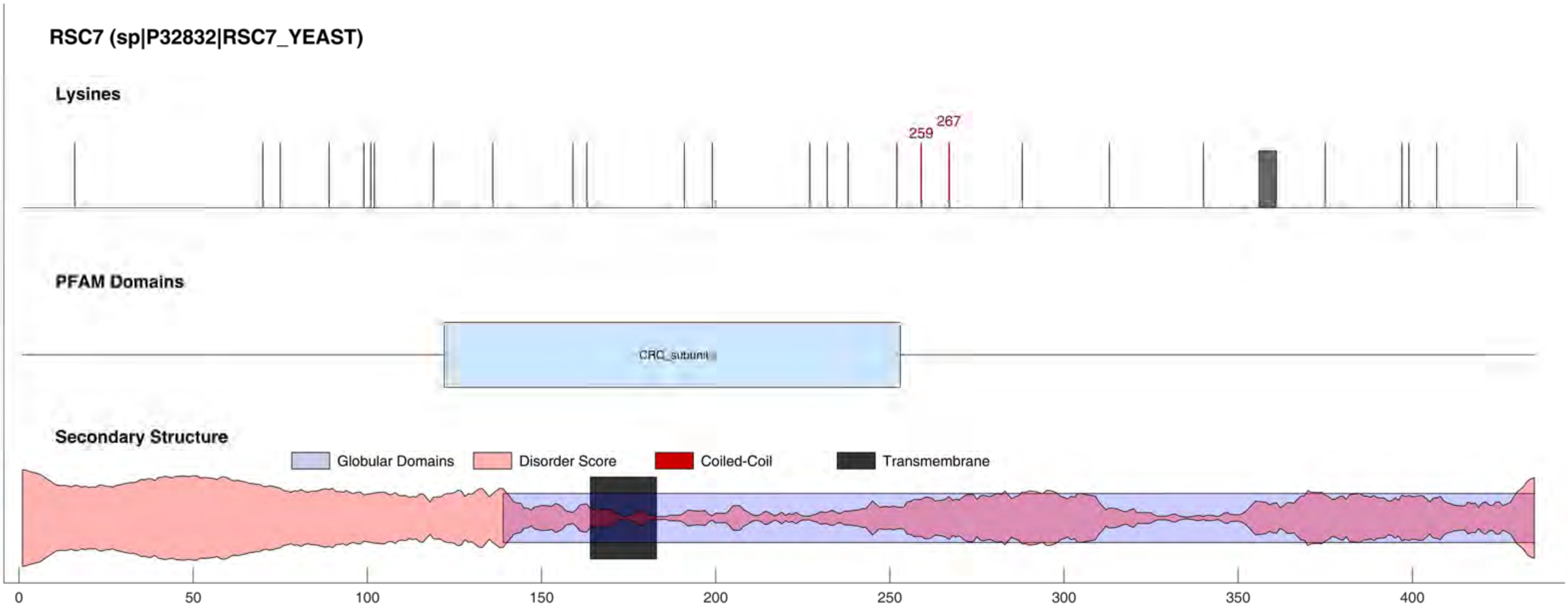
RSC7 (sp|P32832|RSC7_YEAST)
Lysines
259
267
PFAM Domains
CRC_subunit
Secondary Structure
Globular Domains
Disorder Score
Coiled-Coil
Transmembrane
0
50
100
150
200
250
300
350
400

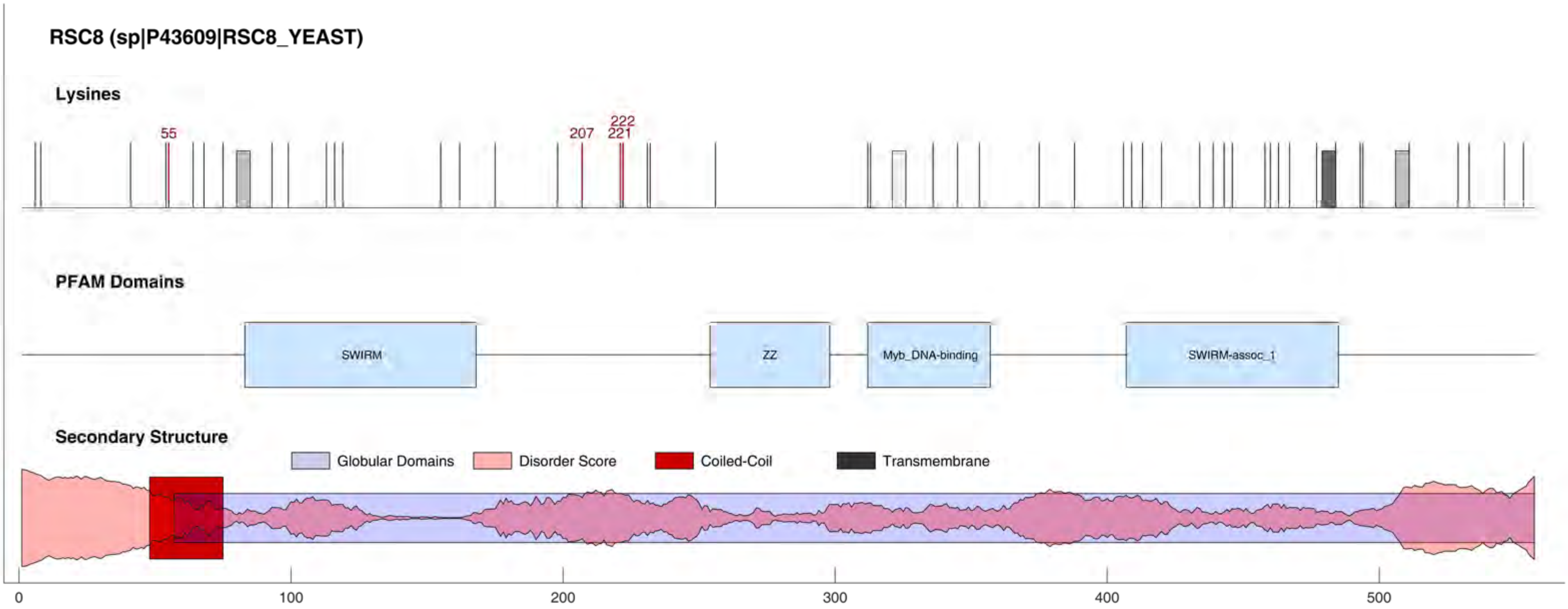

RSC8 (sp|P43609|RSC8_YEAST)
Lysines
55
207
221
222
PFAM Domains
SWIRM
ZZ
Myb_DNA-binding
SWIRM-assoc_1
Secondary Structure
Globular Domains
Disorder Score
Coiled-Coil
Transmembrane
0
100
200
300
400
500

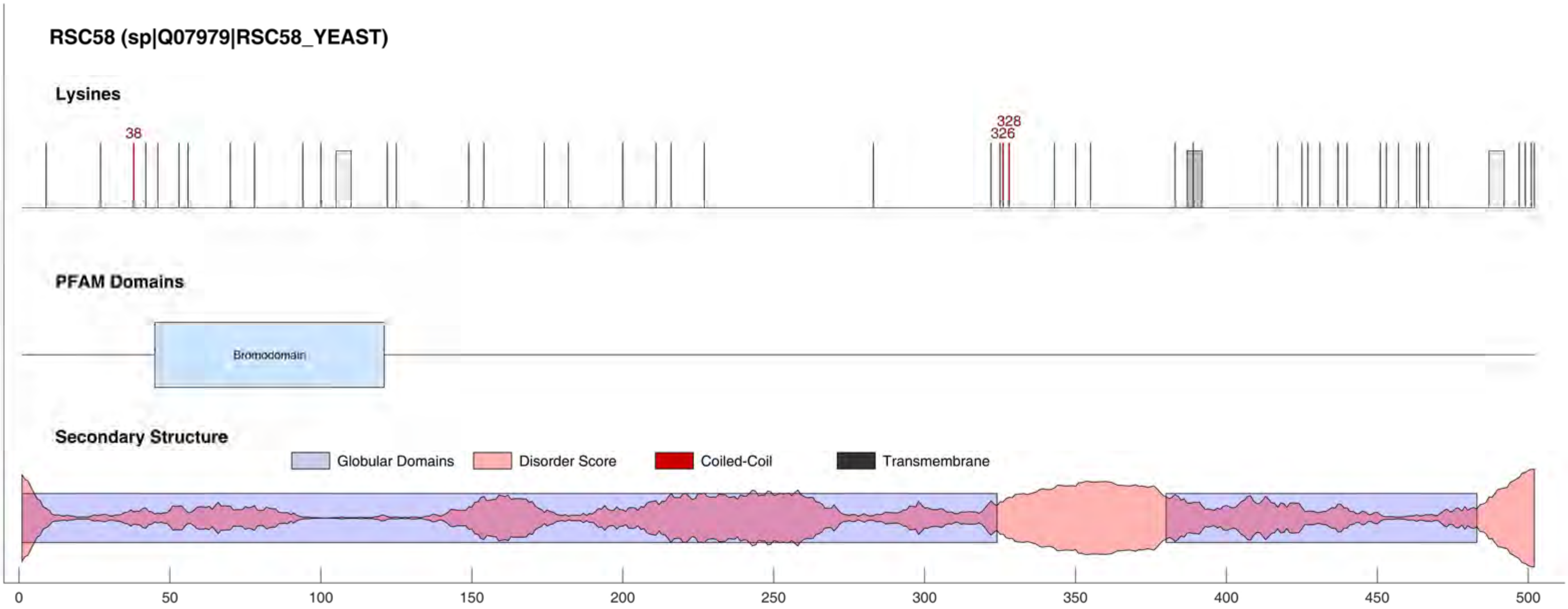

RSC58 (sp|Q07979|RSC58_YEAST)
Lysines
38
328
326
PFAM Domains
Bromodomain
Secondary Structure
Globular Domains
Disorder Score
Coiled-Coil
Transmembrane
0
50
100
150
200
250
300
350
400
450
500

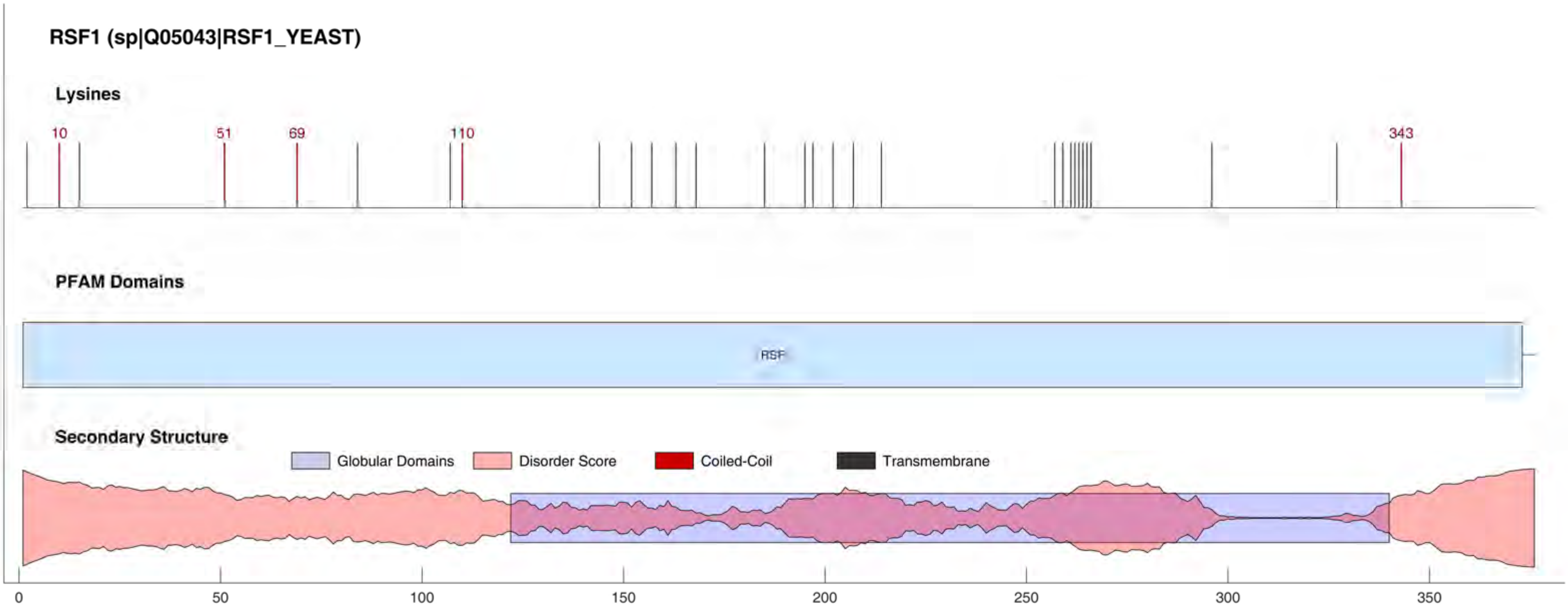
RSF1 (sp|Q05043|RSF1_YEAST)
Lysines
10
51
69
110
343
PFAM Domains
RSF
Secondary Structure
Globular Domains
Disorder Score
Coiled-Coil
Transmembrane
0
50
100
150
200
250
300
350

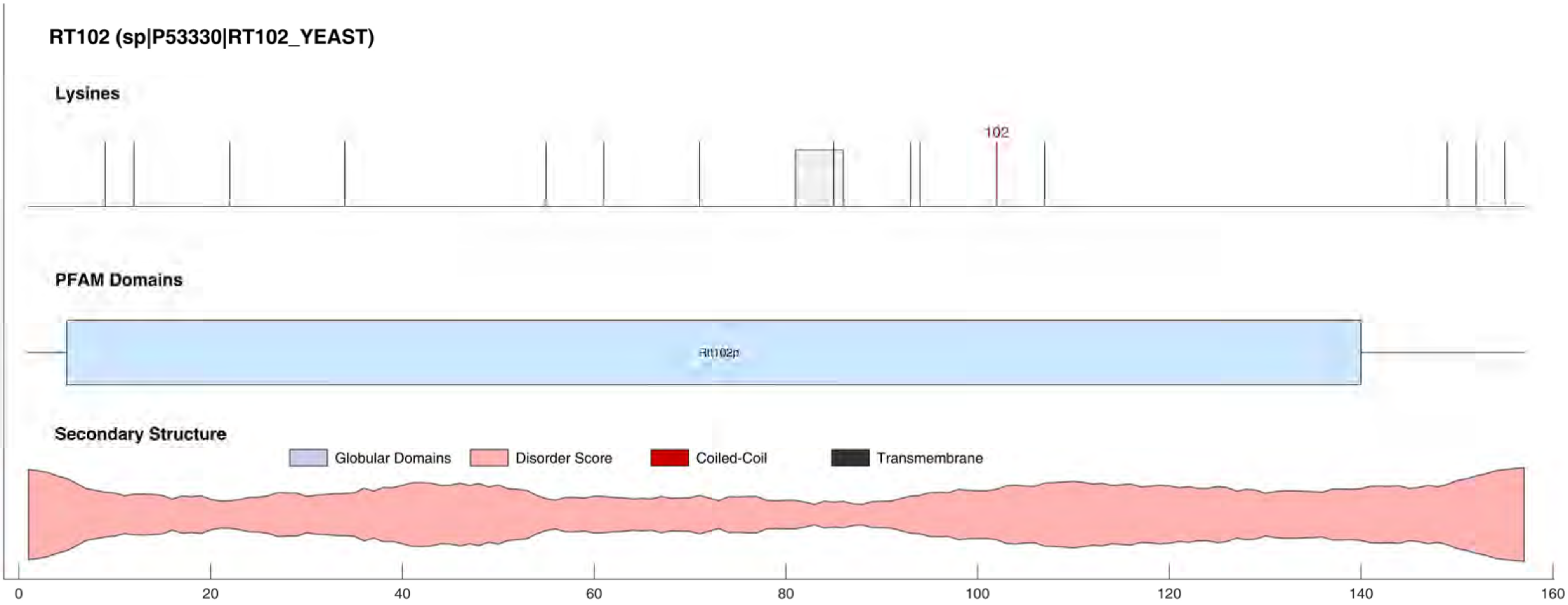
RT102 (sp|P53330|RT102_YEAST)
Lysines
102
PFAM Domains
Rtt102p
Secondary Structure
Globular Domains
Disorder Score
Coiled-Coil
Transmembrane
0
20
40
60
80
100
120
140
160

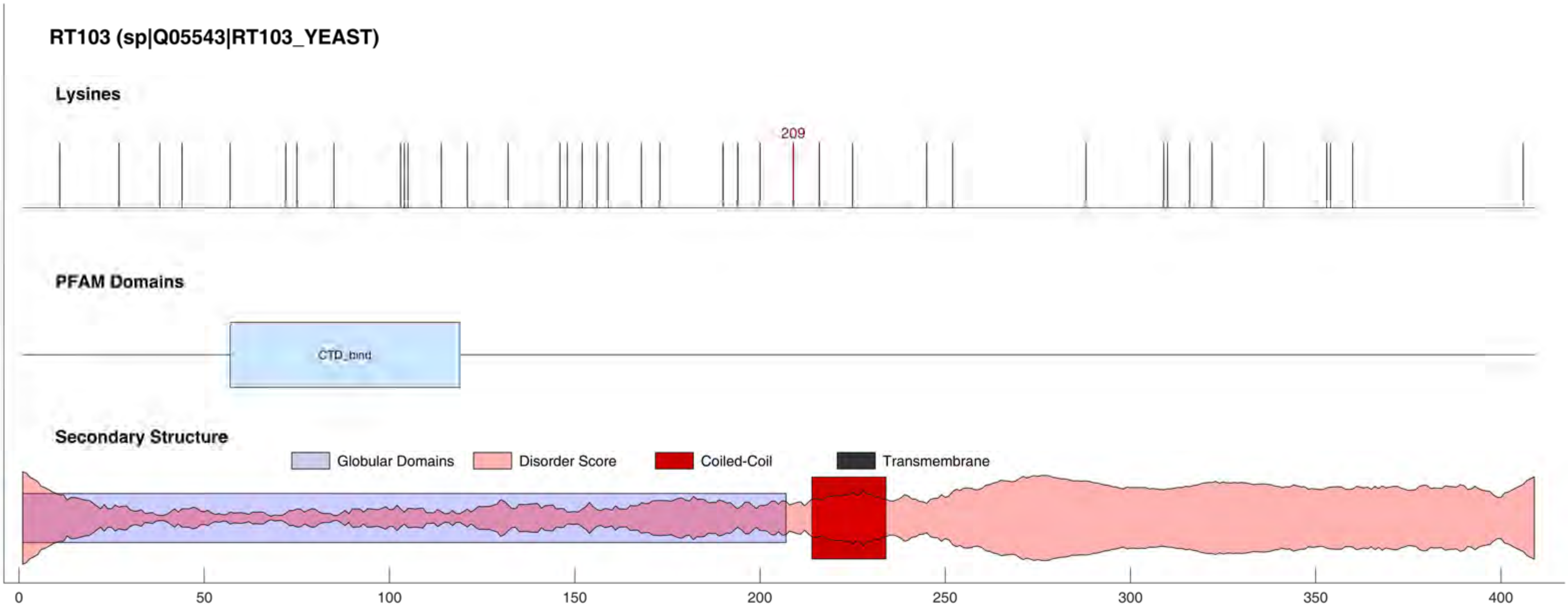
RT103 (sp|Q05543|RT103_YEAST)
Lysines
209
PFAM Domains
CTD_bind
Secondary Structure
Globular Domains
Disorder Score
Coiled-Coil
Transmembrane
0
50
100
150
200
250
300
350
400

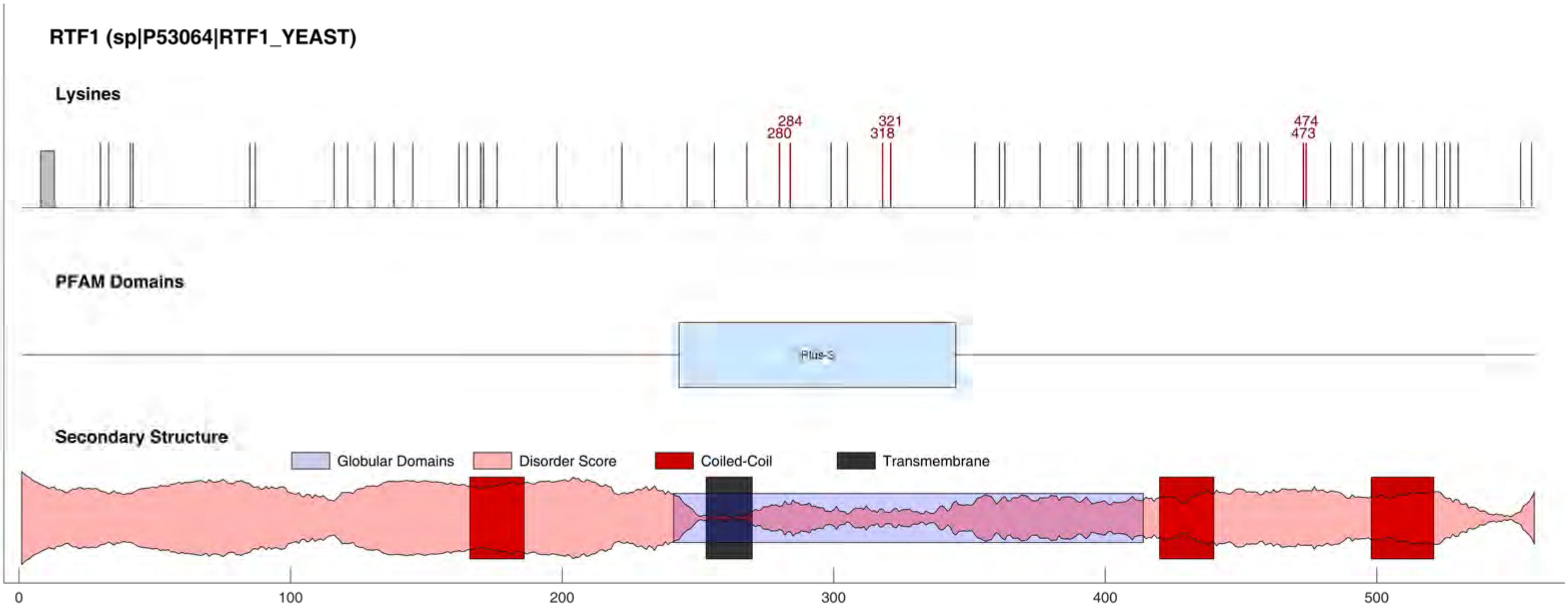

RTF1 (sp|P53064|RTF1_YEAST)
Lysines
284
280
321
318
474
473
PFAM Domains
Plus-3
Secondary Structure
Globular Domains
Disorder Score
Coiled-Coil
Transmembrane
0
100
200
300
400
500

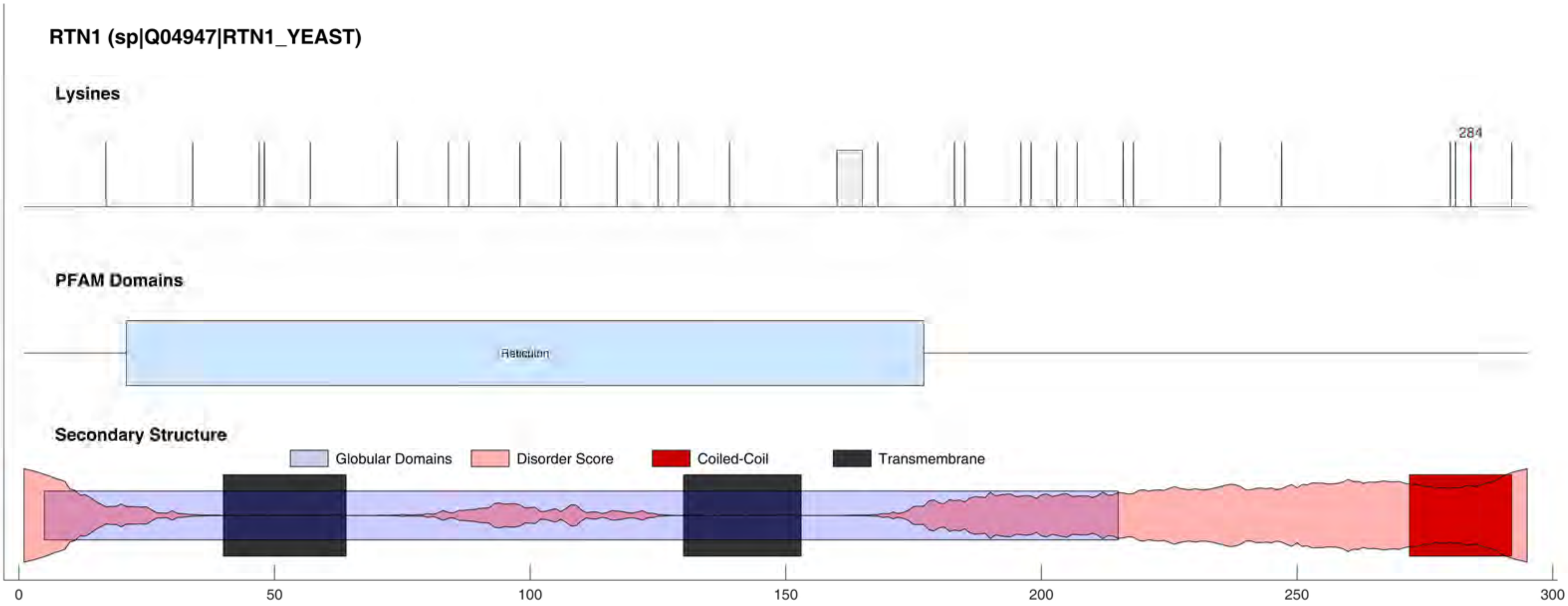

RTN1 (sp|Q04947|RTN1_YEAST)
Lysines
284
PFAM Domains
Reticulon
Secondary Structure
Globular Domains
Disorder Score
Coiled-Coil
Transmembrane
0
50
100
150
200
250
300

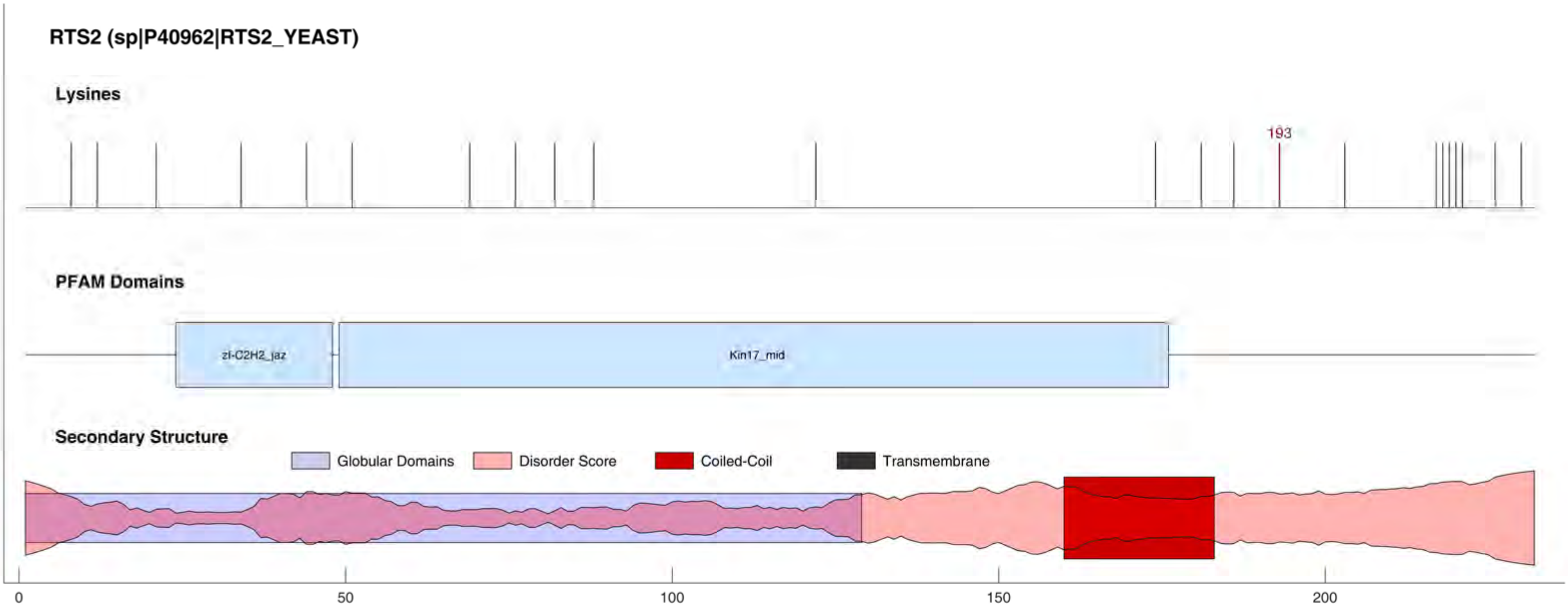
RTS2 (sp|P40962|RTS2_YEAST)
Lysines
193
PFAM Domains
zf-C2H2_jaz
Kin17_mid
Secondary Structure
Globular Domains
Disorder Score
Coiled-Coil
Transmembrane
0
50
100
150
200

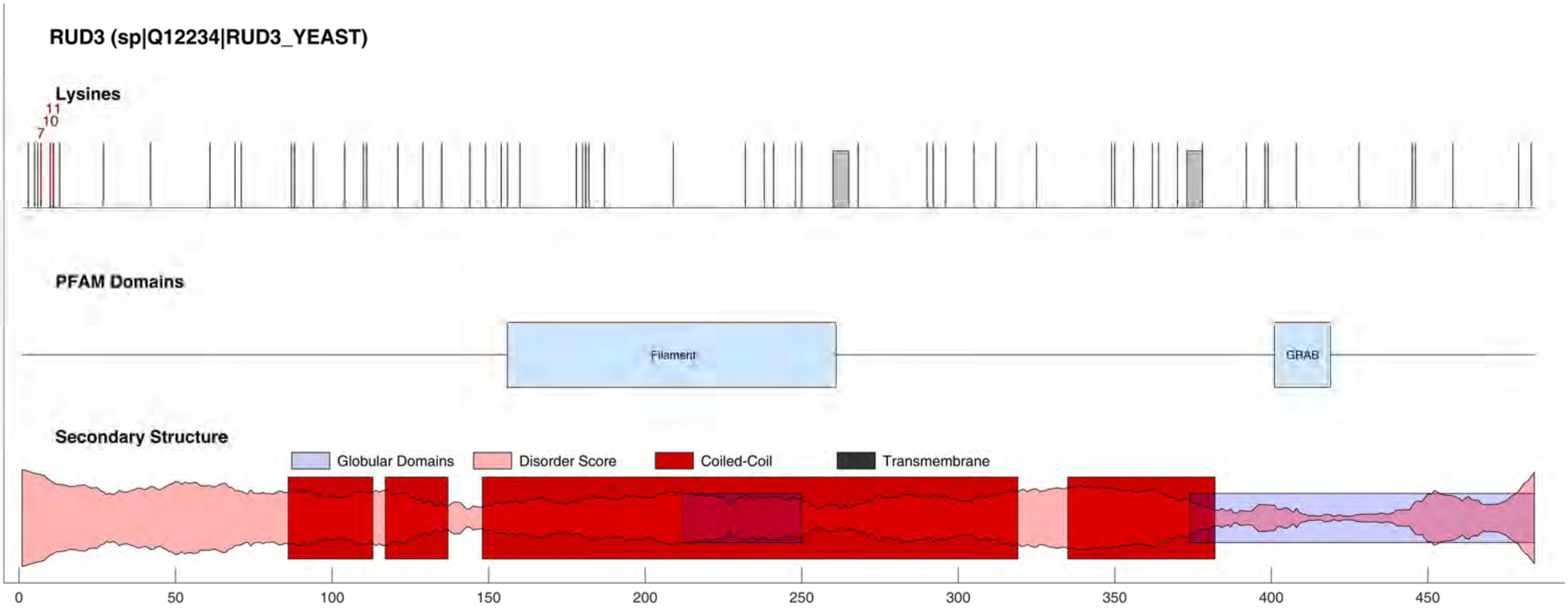

RUD3 (sp|Q12234|RUD3_YEAST)
Lysines
11
10
7
PFAM Domains
Filament
GRAB
Secondary Structure
Globular Domains
Disorder Score
Coiled-Coil
Transmembrane
0
50
100
150
200
250
300
350
400
450

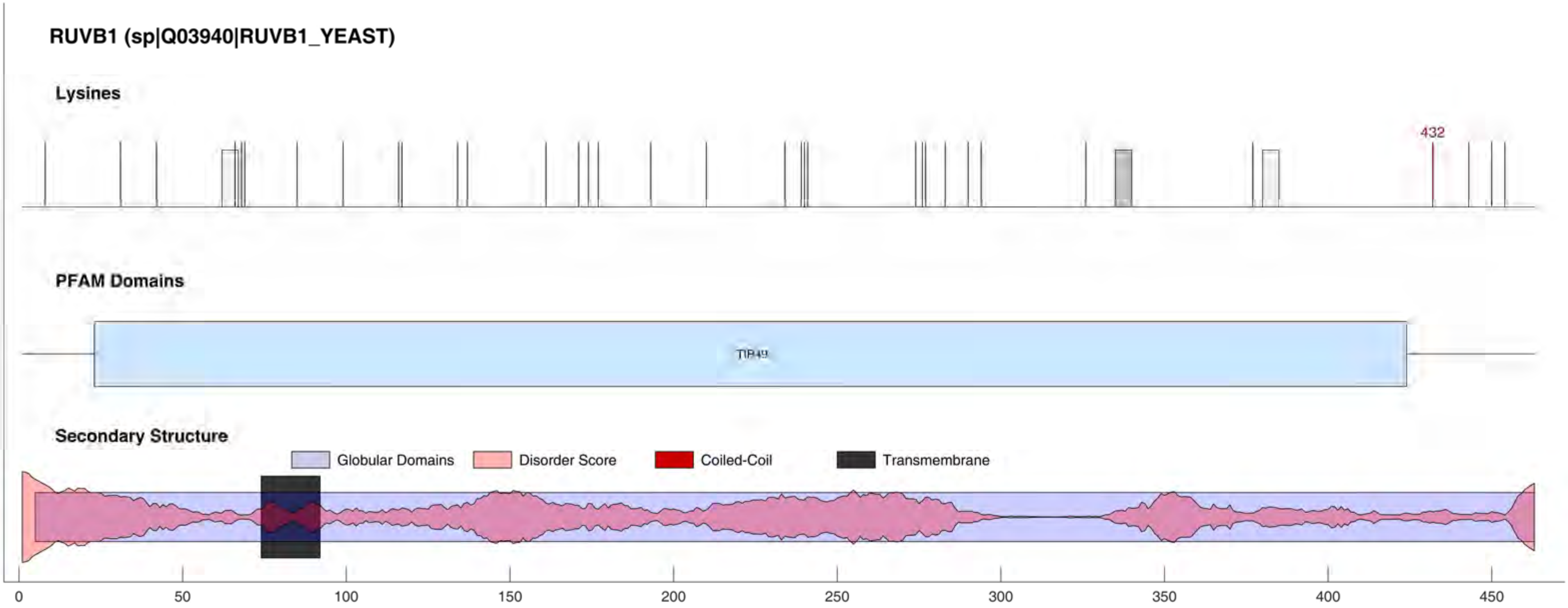

RUVB1 (sp|Q03940|RUVB1_YEAST)
Lysines
432
PFAM Domains
TIP49
Secondary Structure
Globular Domains
Disorder Score
Coiled-Coil
Transmembrane
0
50
100
150
200
250
300
350
400
450

#### RUVB2 (sp|Q12464|RUVB2_YEAST)

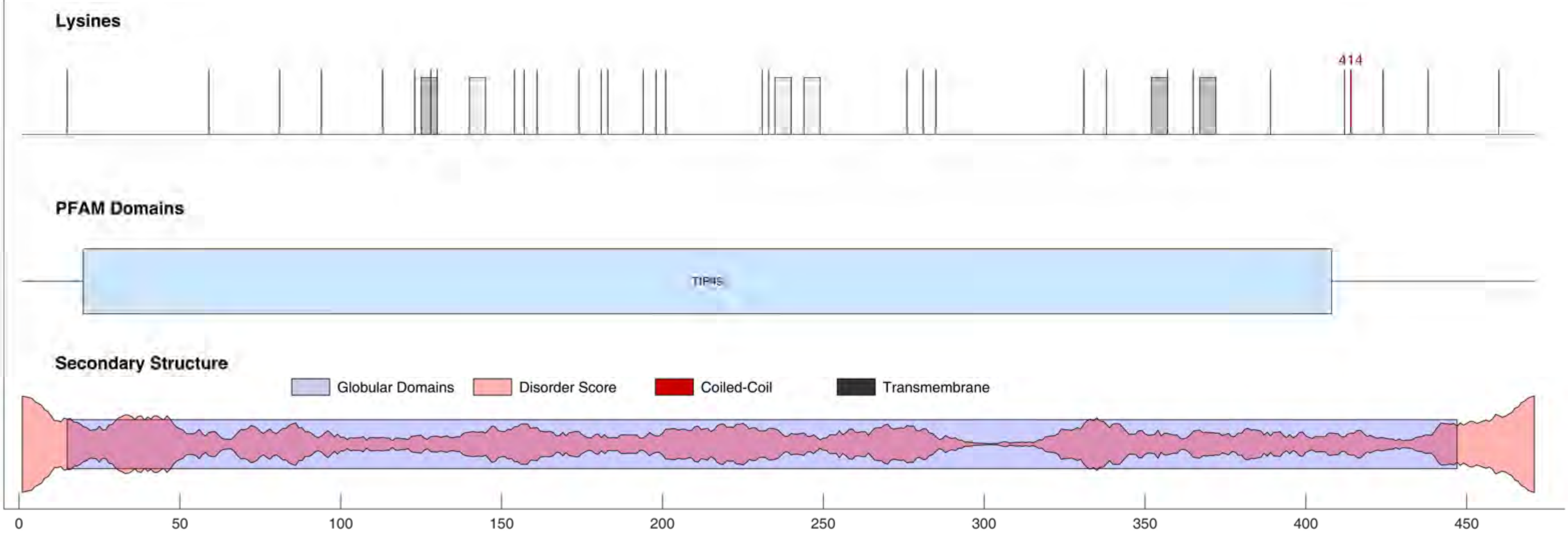

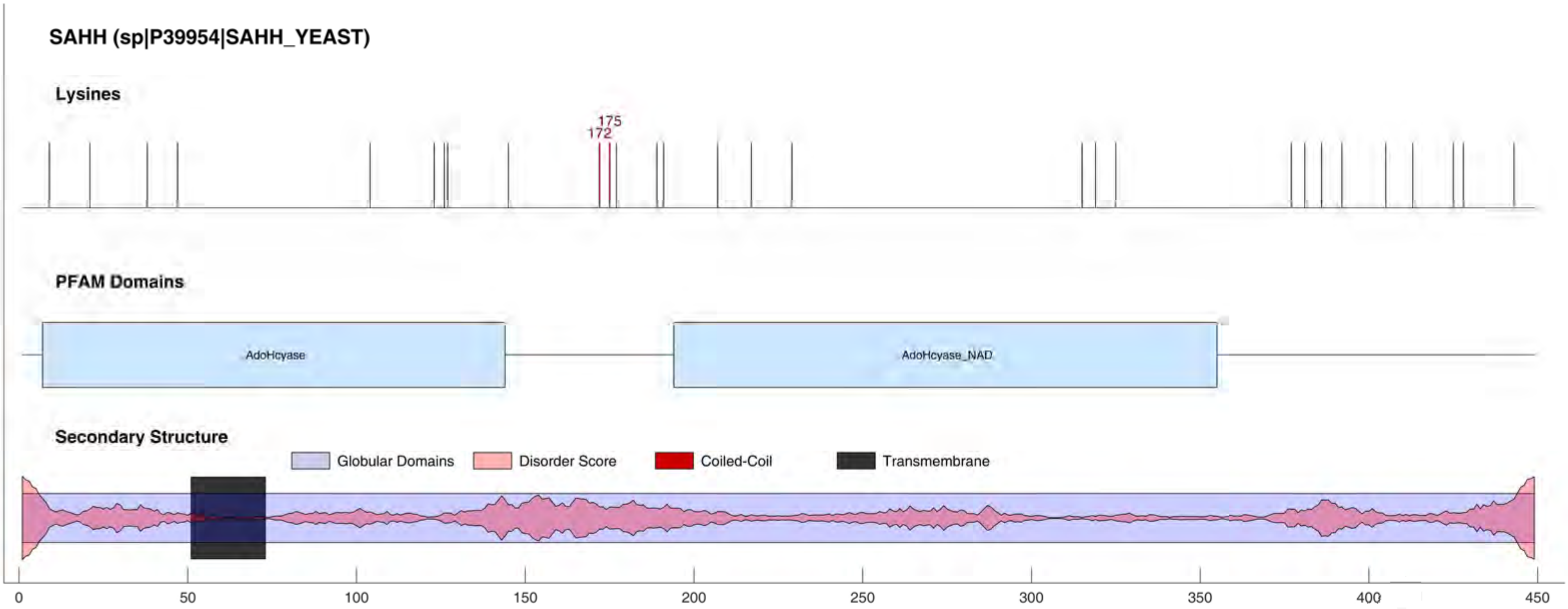
SAHH (sp|P39954|SAHH_YEAST)
Lysines
175
172
PFAM Domains
AdoHcyase
AdoHcyase_NAD
Secondary Structure
Globular Domains
Disorder Score
Coiled-Coil
Transmembrane
0
50
100
150
200
250
300
350
400
450

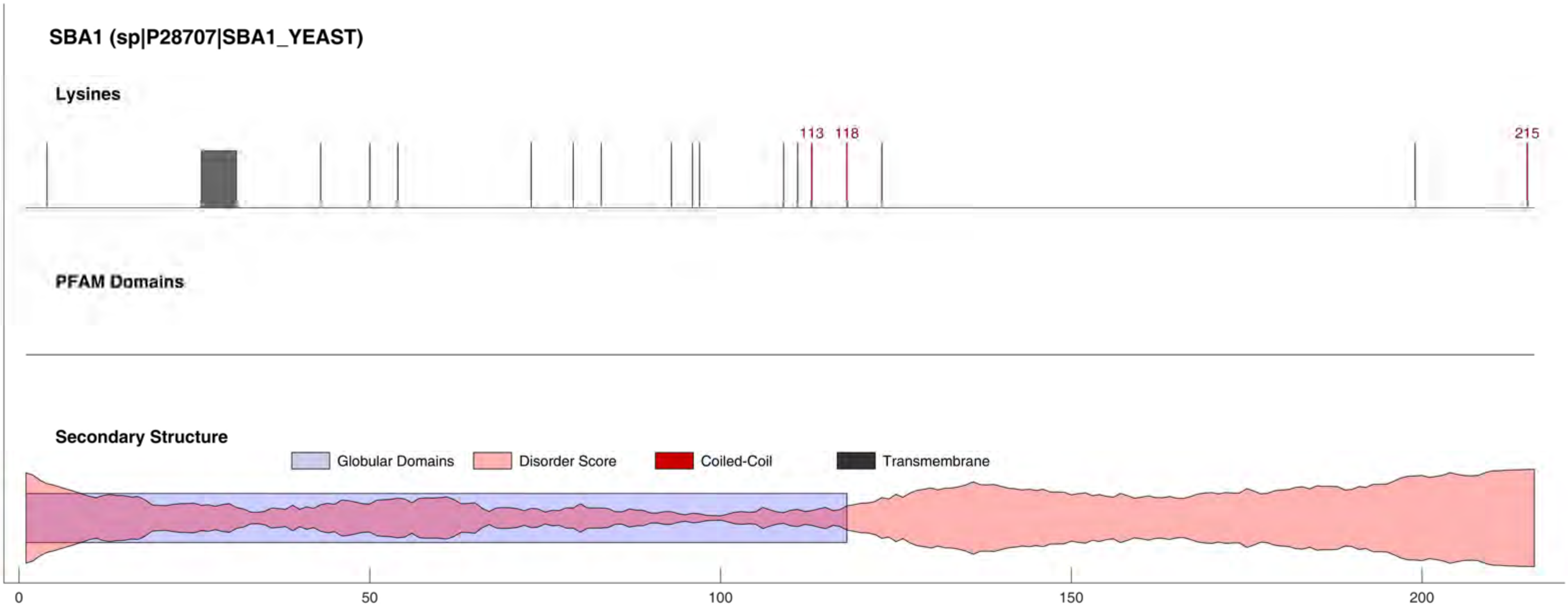

SBA1 (sp|P28707|SBA1_YEAST)
Lysines
113
118
215
PFAM Domains
Secondary Structure
Globular Domains
Disorder Score
Coiled-Coil
Transmembrane
0
50
100
150
200

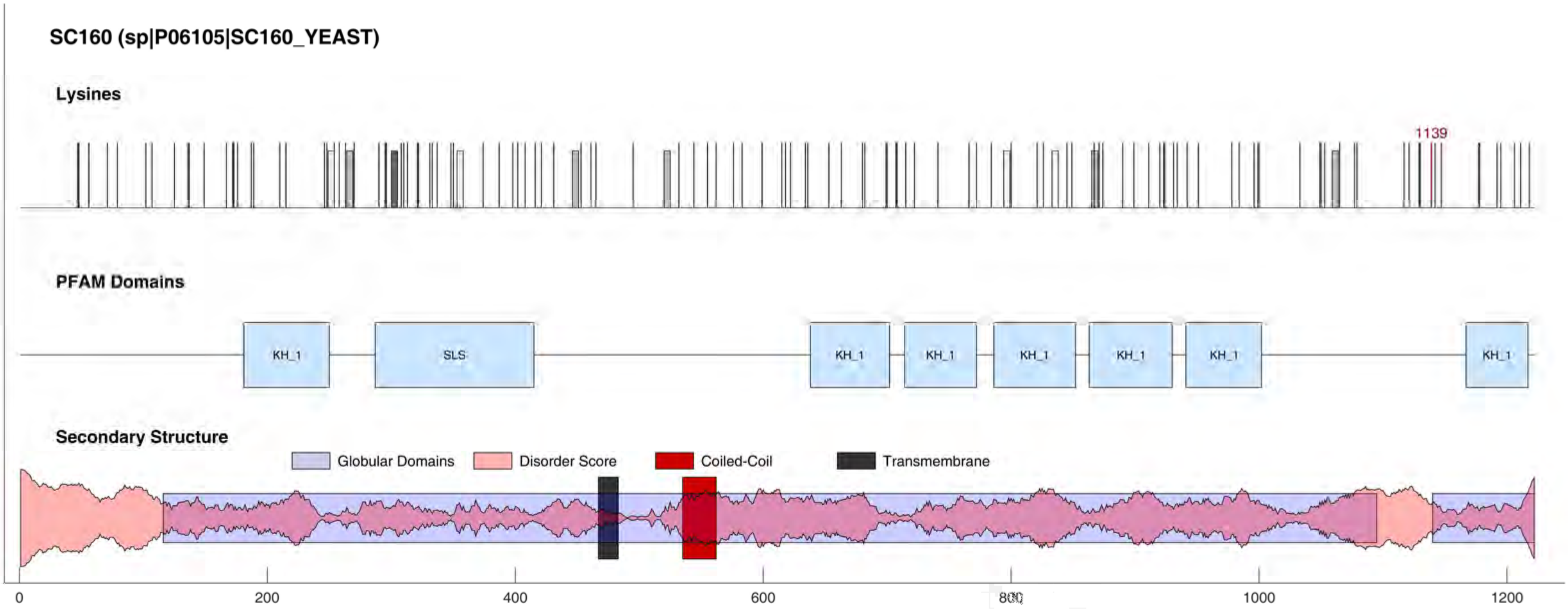
SC160 (sp|P06105|SC160_YEAST)
Lysines
1139
PFAM Domains
KH_1
SLS
KH_1
KH_1
KH_1
KH_1
KH_1
KH_1
Secondary Structure
Globular Domains
Disorder Score
Coiled-Coil
Transmembrane
0
200
400
600
800
1000
1200

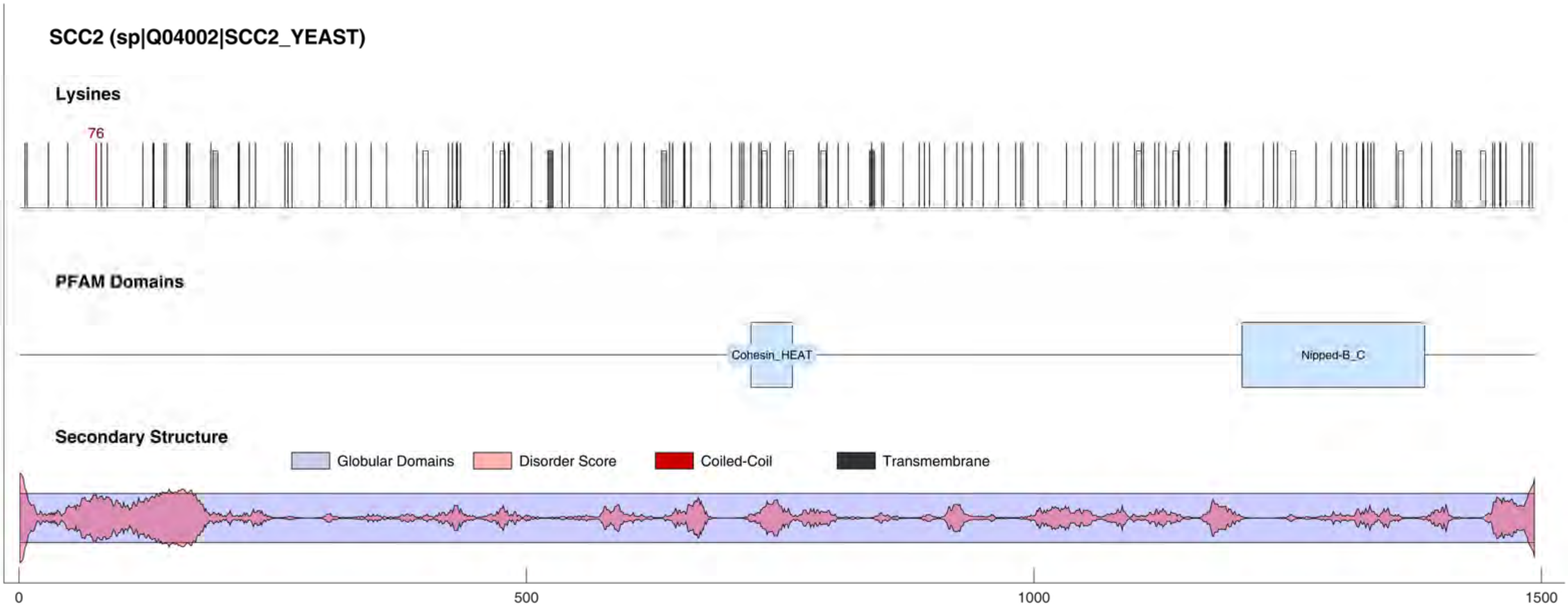

SCC2 (sp|Q04002|SCC2_YEAST)
Lysines
76
PFAM Domains
Cohesin_HEAT
Nipped-B_C
Secondary Structure
Globular Domains
Disorder Score
Coiled-Coil
Transmembrane
0
500
1000
1500

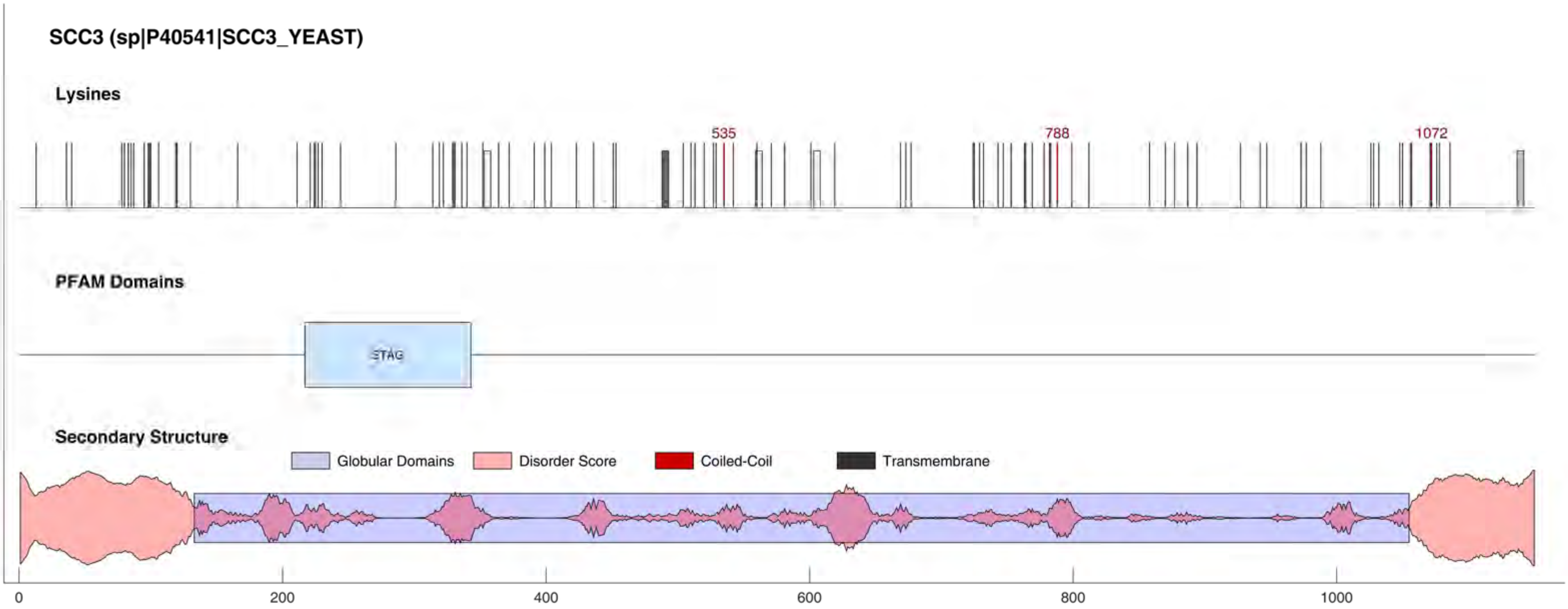
SCC3 (sp|P40541|SCC3_YEAST)
Lysines
535
788
1072
PFAM Domains
STAG
Secondary Structure
Globular Domains
Disorder Score
Coiled-Coil
Transmembrane
0
200
400
600
800
1000

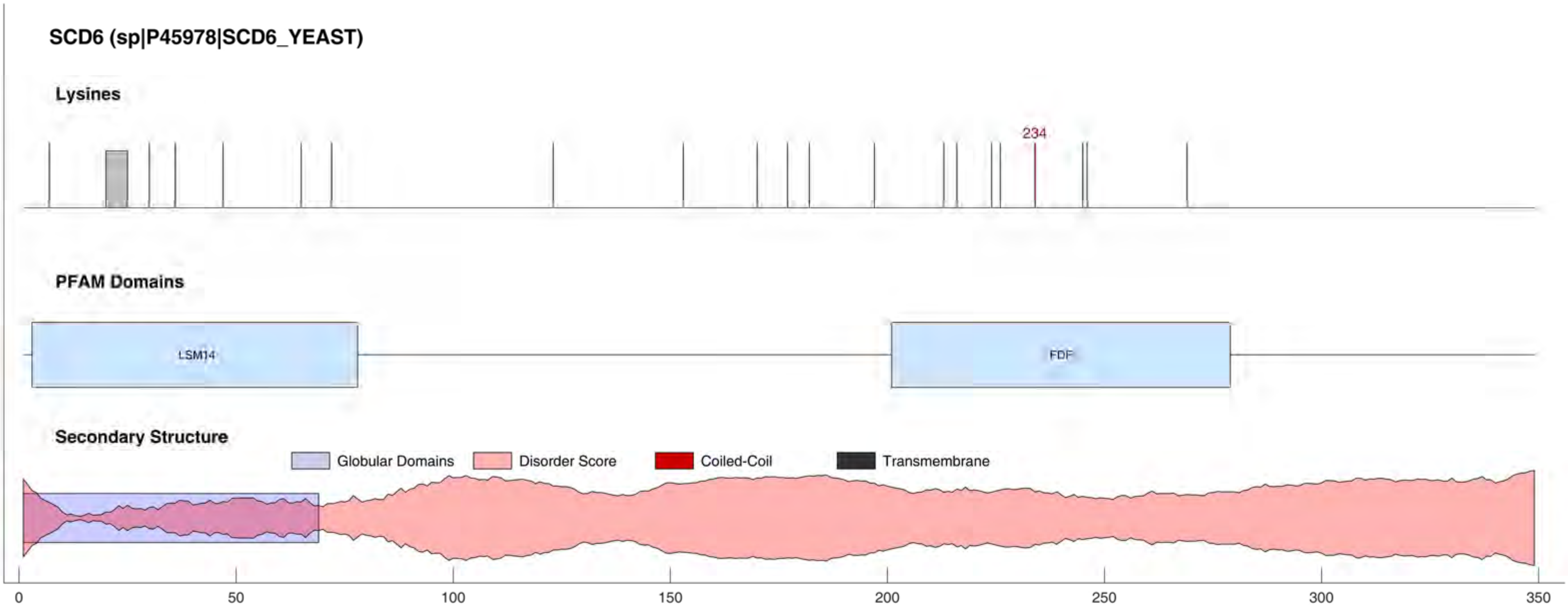
SCD6 (sp|P45978|SCD6_YEAST)
Lysines
234
PFAM Domains
LSM14
FDF
Secondary Structure
Globular Domains
Disorder Score
Coiled-Coil
Transmembrane
0
50
100
150
200
250
300
350

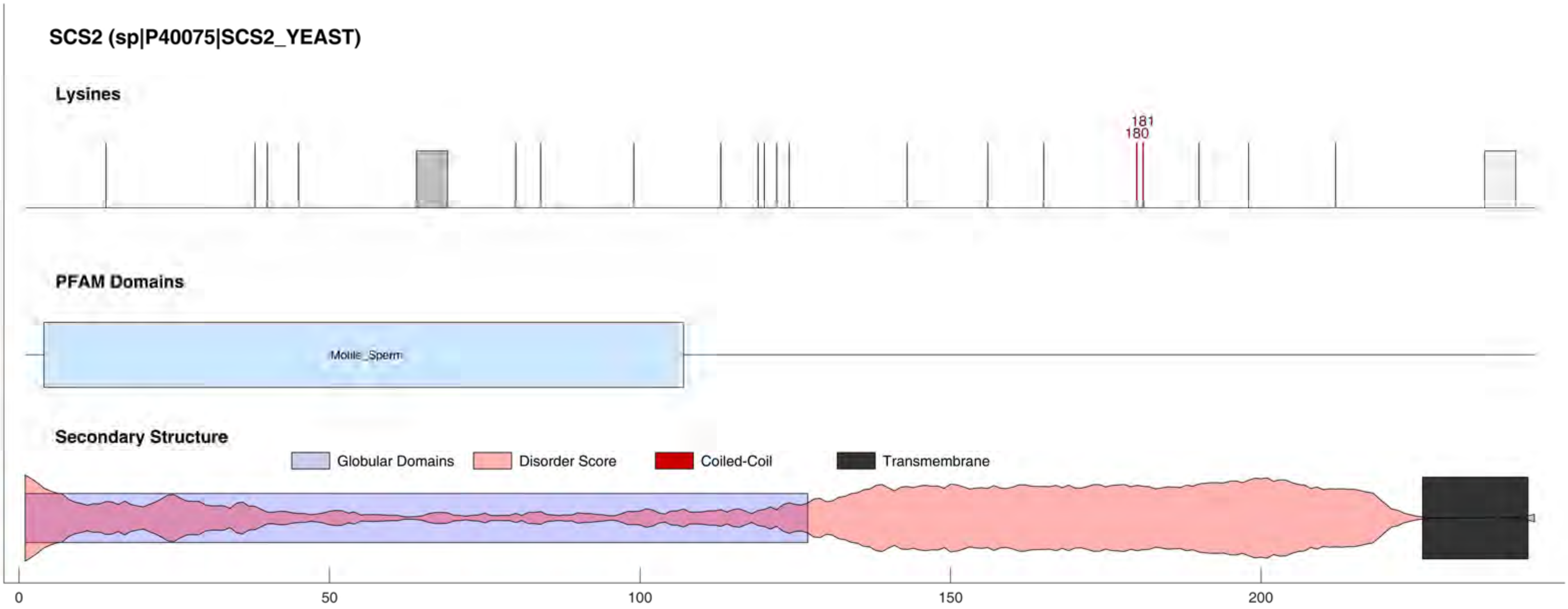
SCS2 (sp|P40075|SCS2_YEAST)
Lysines
181
180
PFAM Domains
Motile_Sperm
Secondary Structure
Globular Domains
Disorder Score
Coiled-Coil
Transmembrane
0
50
100
150
200

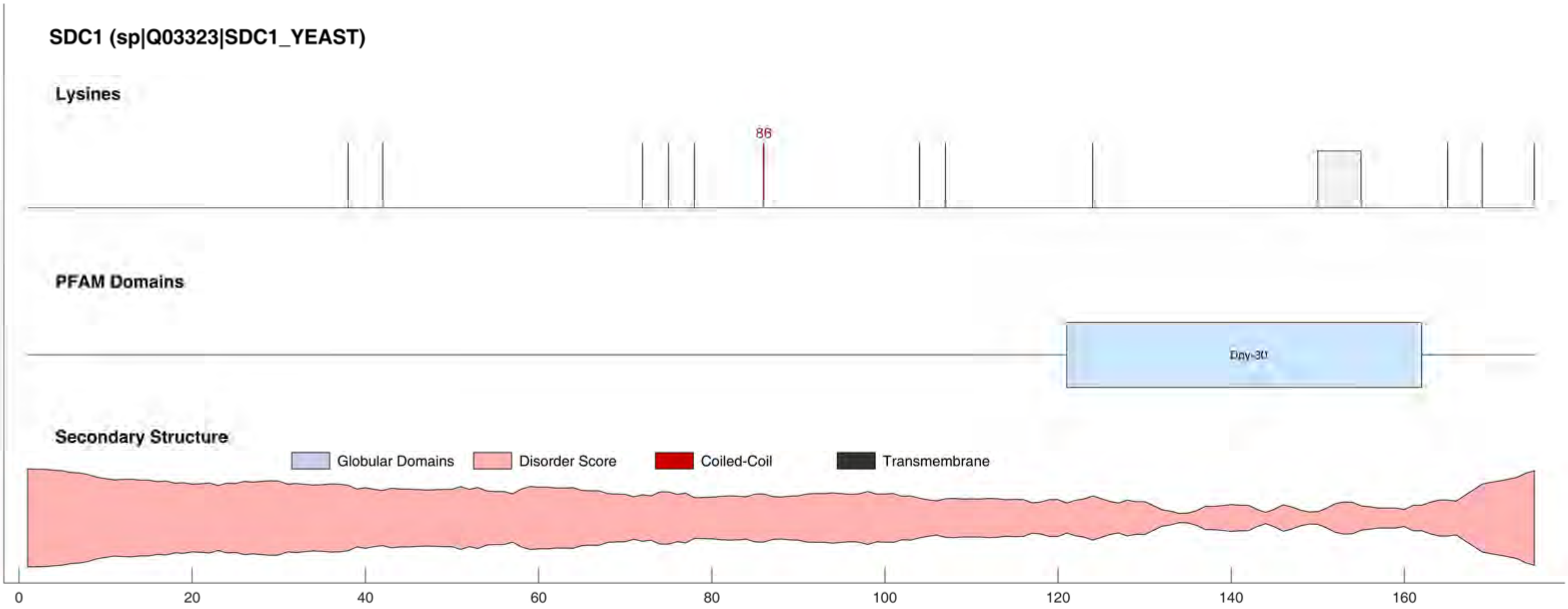
SDC1 (sp|Q03323|SDC1_YEAST)
Lysines
86
PFAM Domains
Dpy-30
Secondary Structure
Globular Domains
Disorder Score
Coiled-Coil
Transmembrane
0
20
40
60
80
100
120
140
160

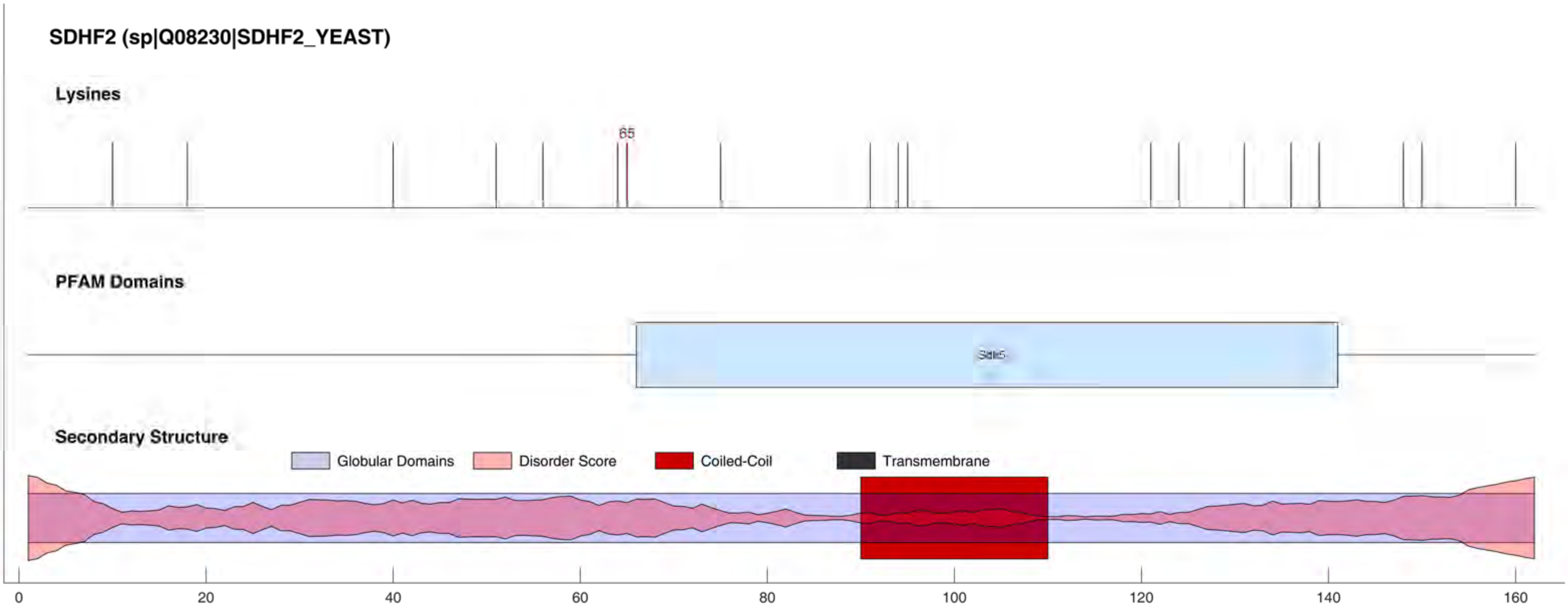
SDHF2 (sp|Q08230|SDHF2_YEAST)
Lysines
65
PFAM Domains
Sdh5
Secondary Structure
Globular Domains
Disorder Score
Coiled-Coil
Transmembrane
0
20
40
60
80
100
120
140
160

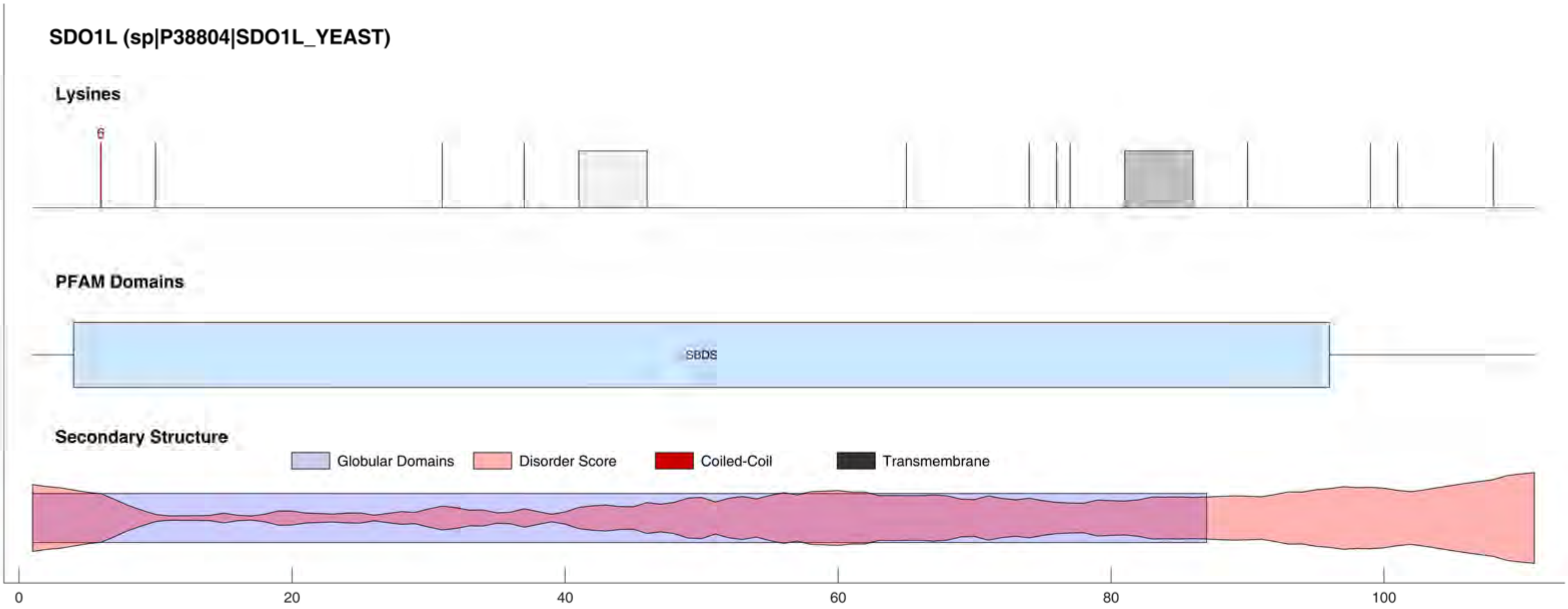
SDO1L (sp|P38804|SDO1L_YEAST)
Lysines
6
PFAM Domains
SBDS
Secondary Structure
Globular Domains
Disorder Score
Coiled-Coil
Transmembrane
0
20
40
60
80
100

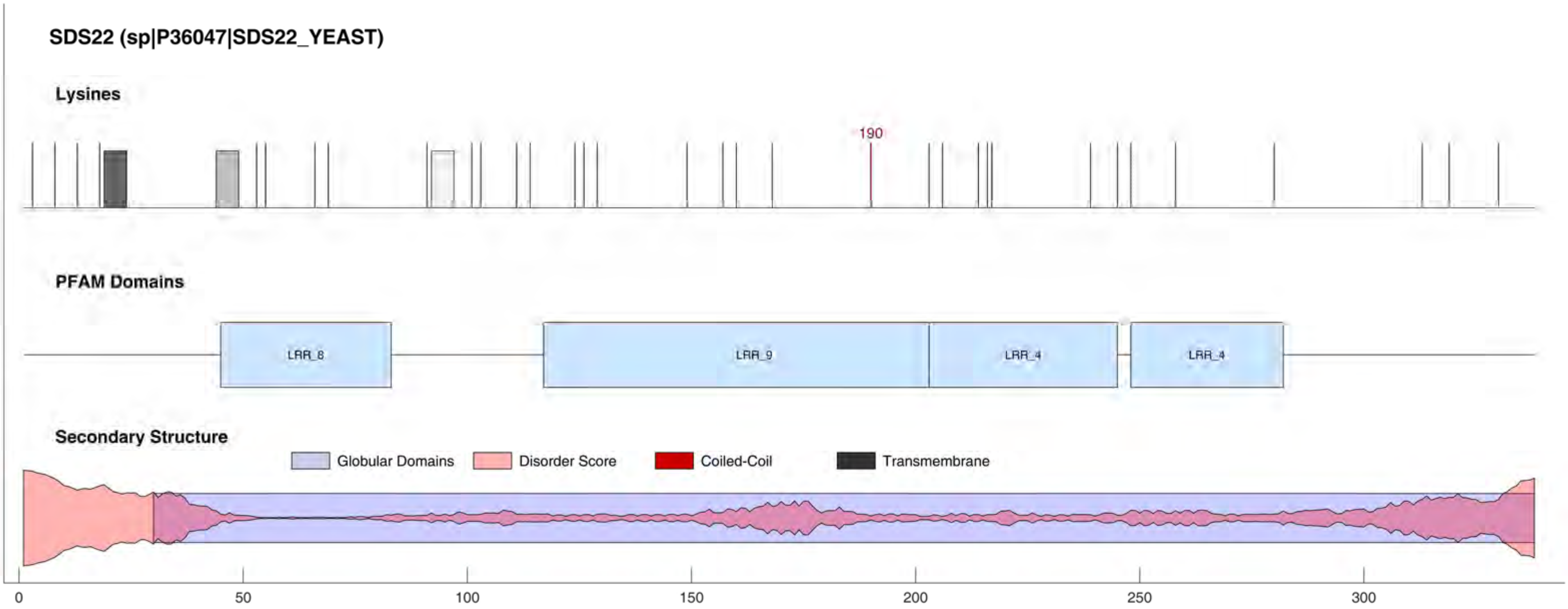
SDS22 (sp|P36047|SDS22_YEAST)
Lysines
190
PFAM Domains
LRR_8
LRR_9
LRR_4
LRR_4
Secondary Structure
Globular Domains
Disorder Score
Coiled-Coil
Transmembrane
0
50
100
150
200
250
300

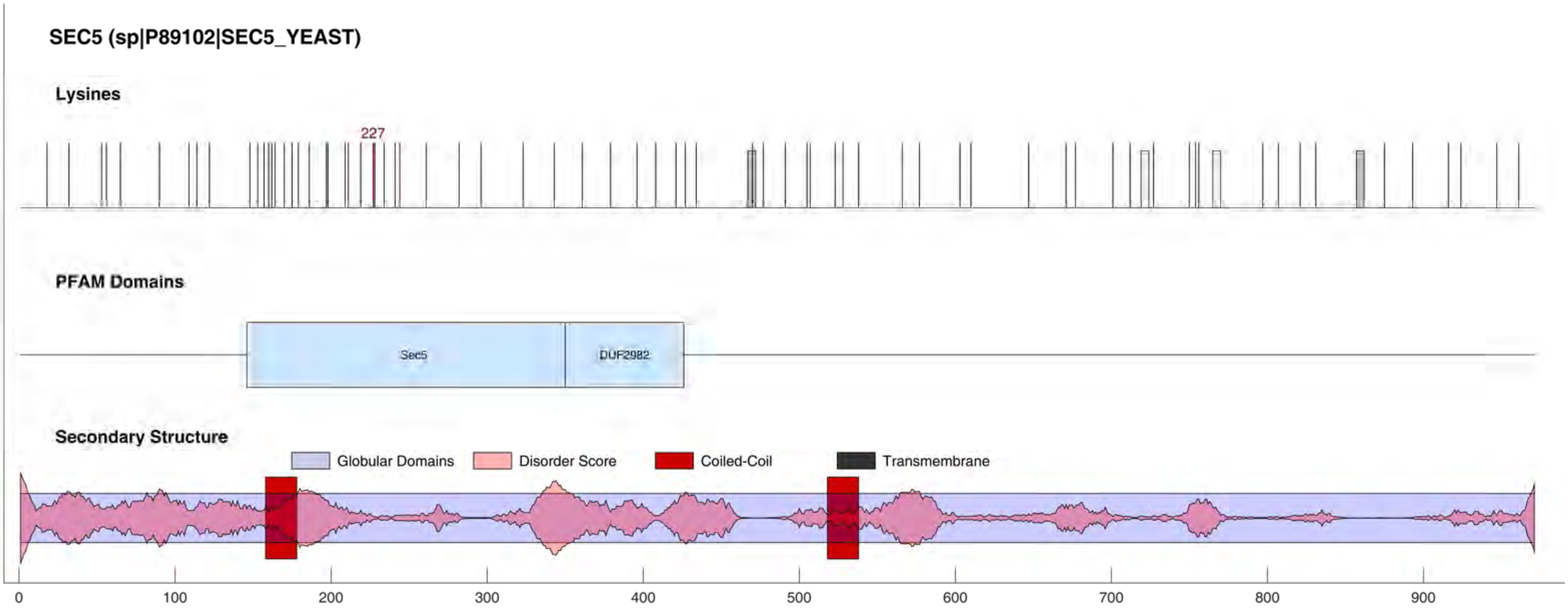
SEC5 (sp|P89102|SEC5_YEAST)
Lysines
227
PFAM Domains
Sec5
DUF2982
Secondary Structure
Globular Domains
Disorder Score
Coiled-Coil
Transmembrane
0
100
200
300
400
500
600
700
800
900

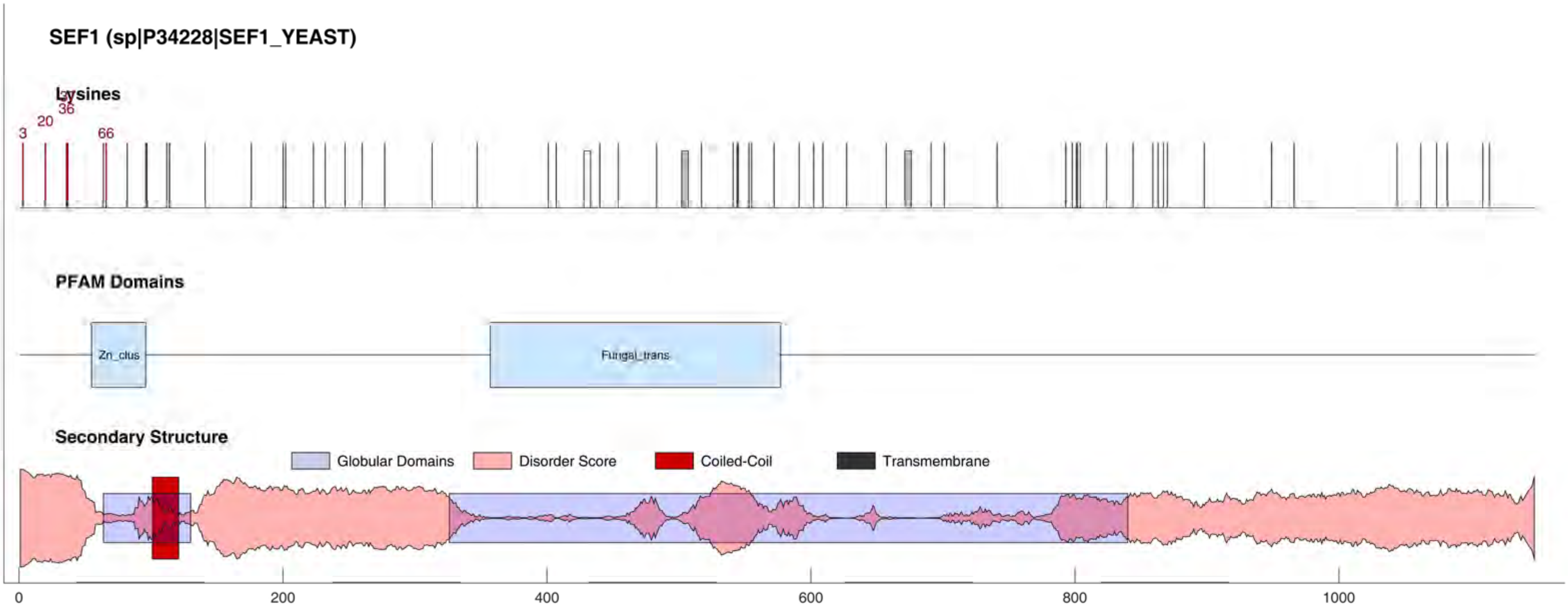
SEF1 (sp|P34228|SEF1_YEAST)
Lysines
3
20
36
66
PFAM Domains
Zn_clus
Fungal_trans
Secondary Structure
Globular Domains
Disorder Score
Coiled-Coil
Transmembrane
0
200
400
600
800
1000

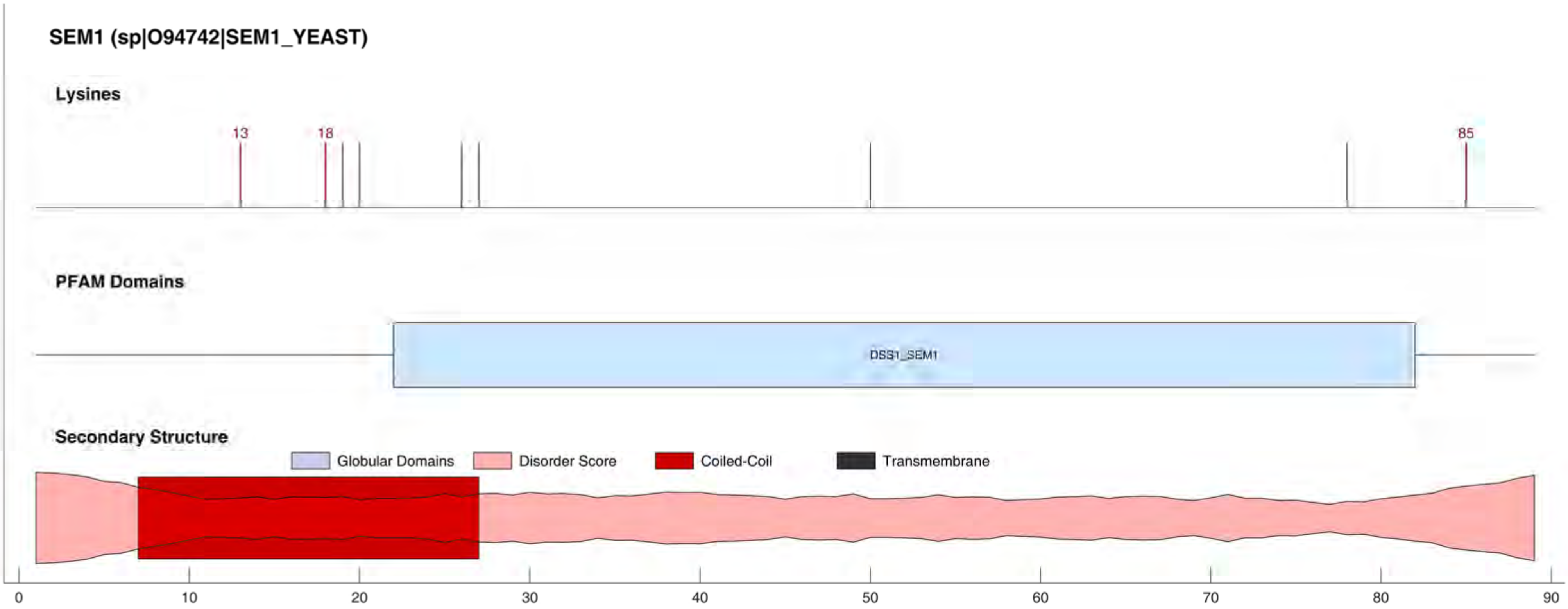
SEM1 (sp|O94742|SEM1_YEAST)
Lysines
13
18
85
PFAM Domains
DSS1_SEM1
Secondary Structure
Globular Domains
Disorder Score
Coiled-Coil
Transmembrane
0
10
20
30
40
50
60
70
80
90

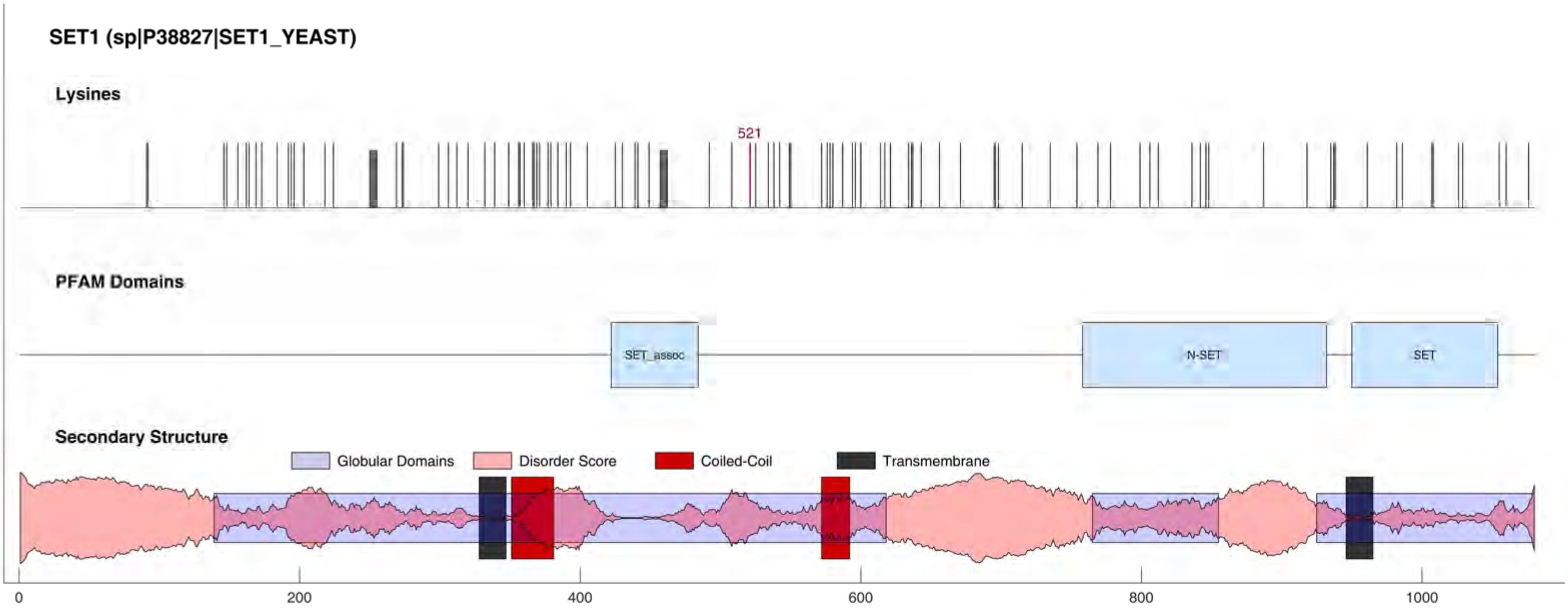
SET1 (sp|P38827|SET1_YEAST)
Lysines
521
PFAM Domains
SET_assoc
N-SET
SET
Secondary Structure
Globular Domains
Disorder Score
Coiled-Coil
Transmembrane
0
200
400
600
800
1000

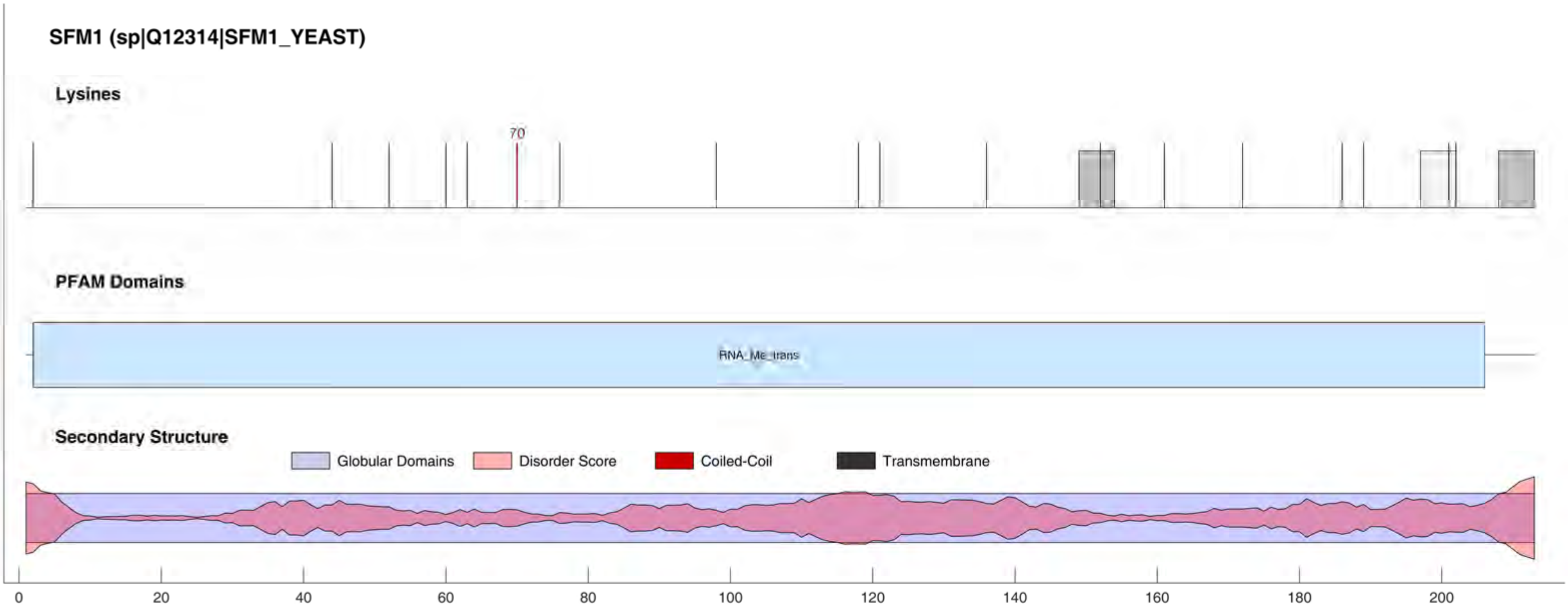
SFM1 (sp|Q12314|SFM1_YEAST)
Lysines
70
PFAM Domains
RNA_Me_trans
Secondary Structure
Globular Domains
Disorder Score
Coiled-Coil
Transmembrane
0
20
40
60
80
100
120
140
160
180
200

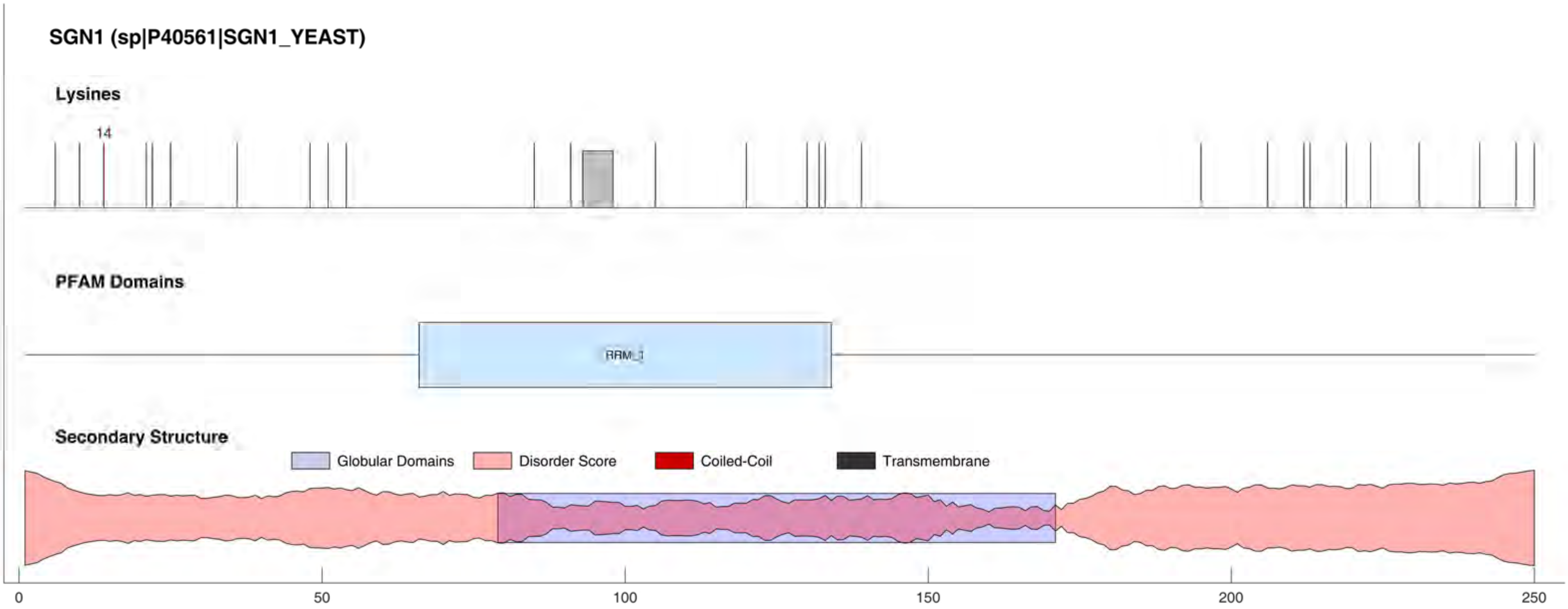

SGN1 (sp|P40561|SGN1_YEAST)
Lysines
14
PFAM Domains
RRM_1
Secondary Structure
Globular Domains
Disorder Score
Coiled-Coil
Transmembrane
0
50
100
150
200
250

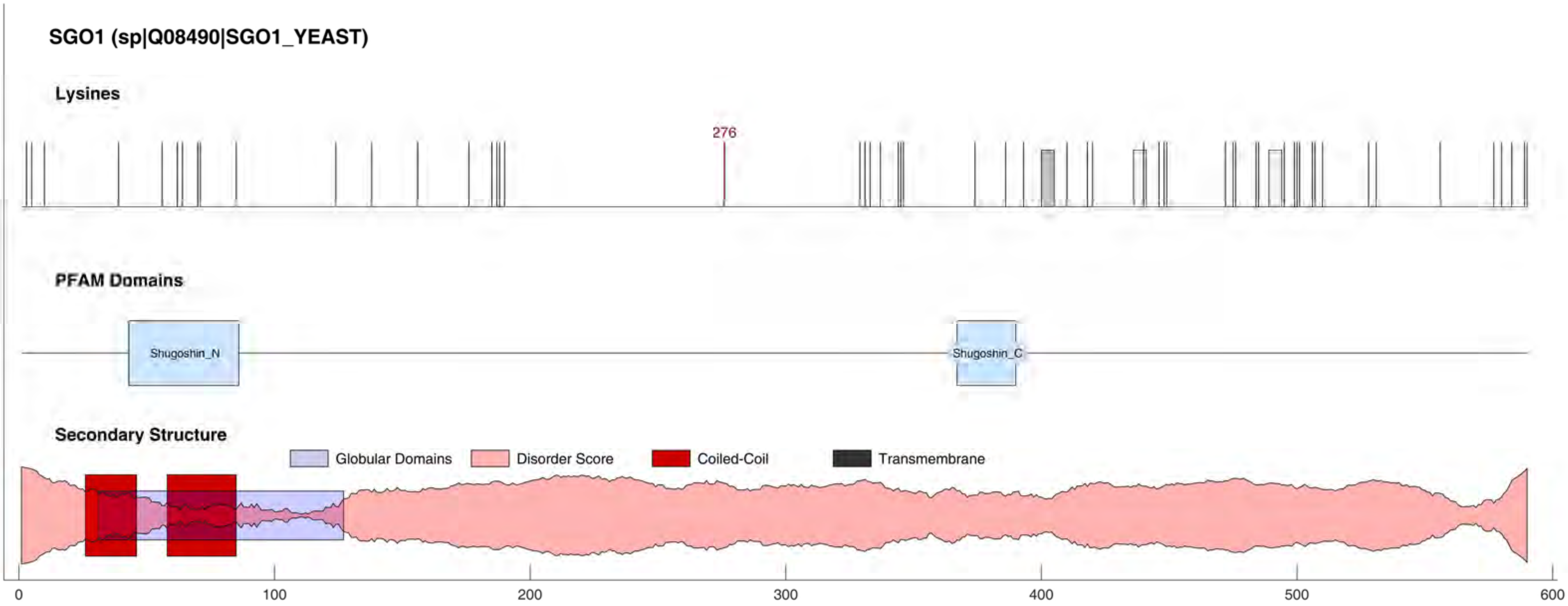
SGO1 (sp|Q08490|SGO1_YEAST)
Lysines
276
PFAM Domains
Shugoshin_N
Shugoshin_C
Secondary Structure
Globular Domains
Disorder Score
Coiled-Coil
Transmembrane
0
100
200
300
400
500
600

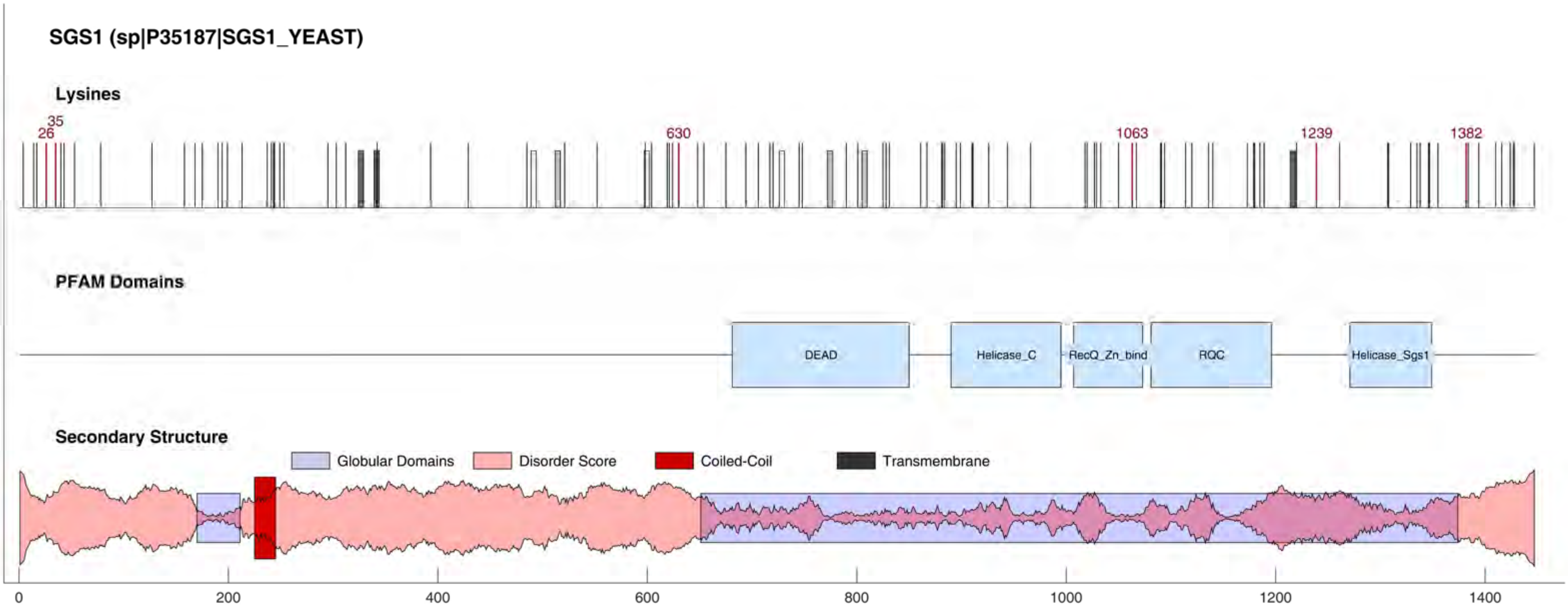

SGS1 (sp|P35187|SGS1_YEAST)
Lysines
26
35
630
1063
1239
1382
PFAM Domains
DEAD
Helicase_C
RecQ_Zn_bind
RQC
Helicase_Sgs1
Secondary Structure
Globular Domains
Disorder Score
Coiled-Coil
Transmembrane
0
200
400
600
800
1000
1200
1400

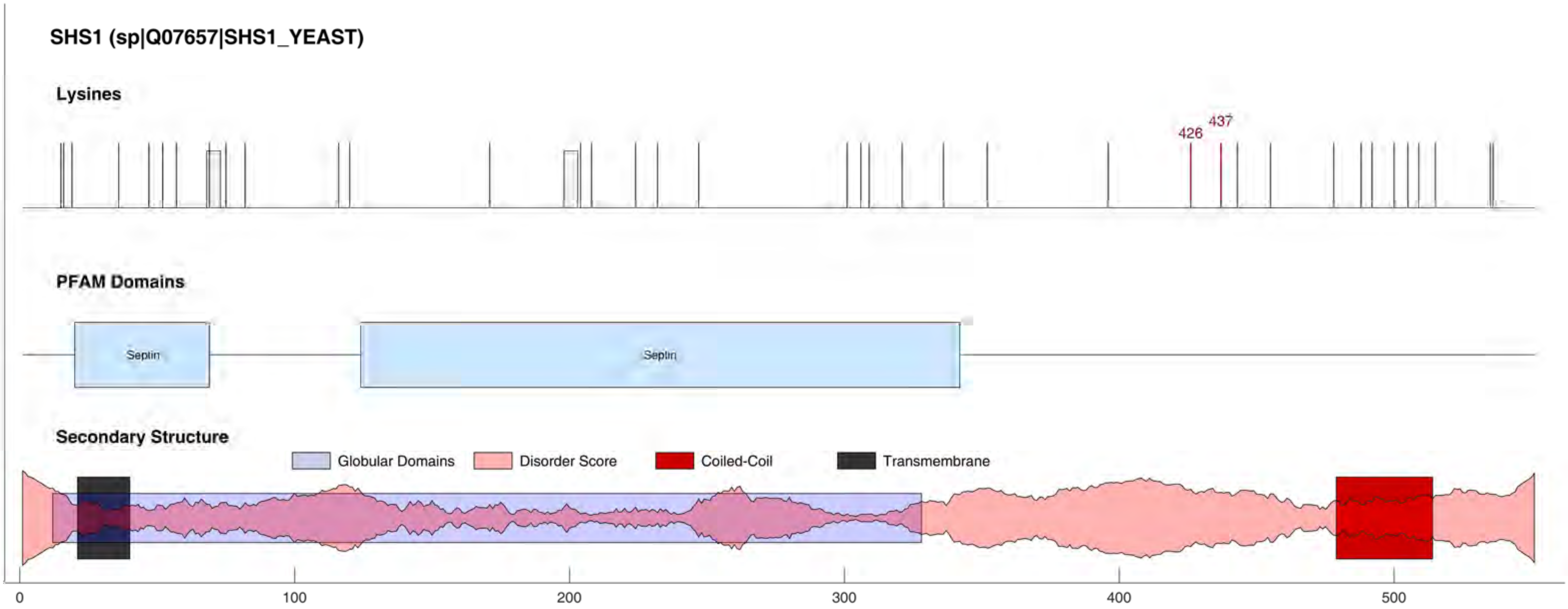
SHS1 (sp|Q07657|SHS1_YEAST)
Lysines
426
437
PFAM Domains
Septin
Septin
Secondary Structure
Globular Domains
Disorder Score
Coiled-Coil
Transmembrane
0
100
200
300
400
500

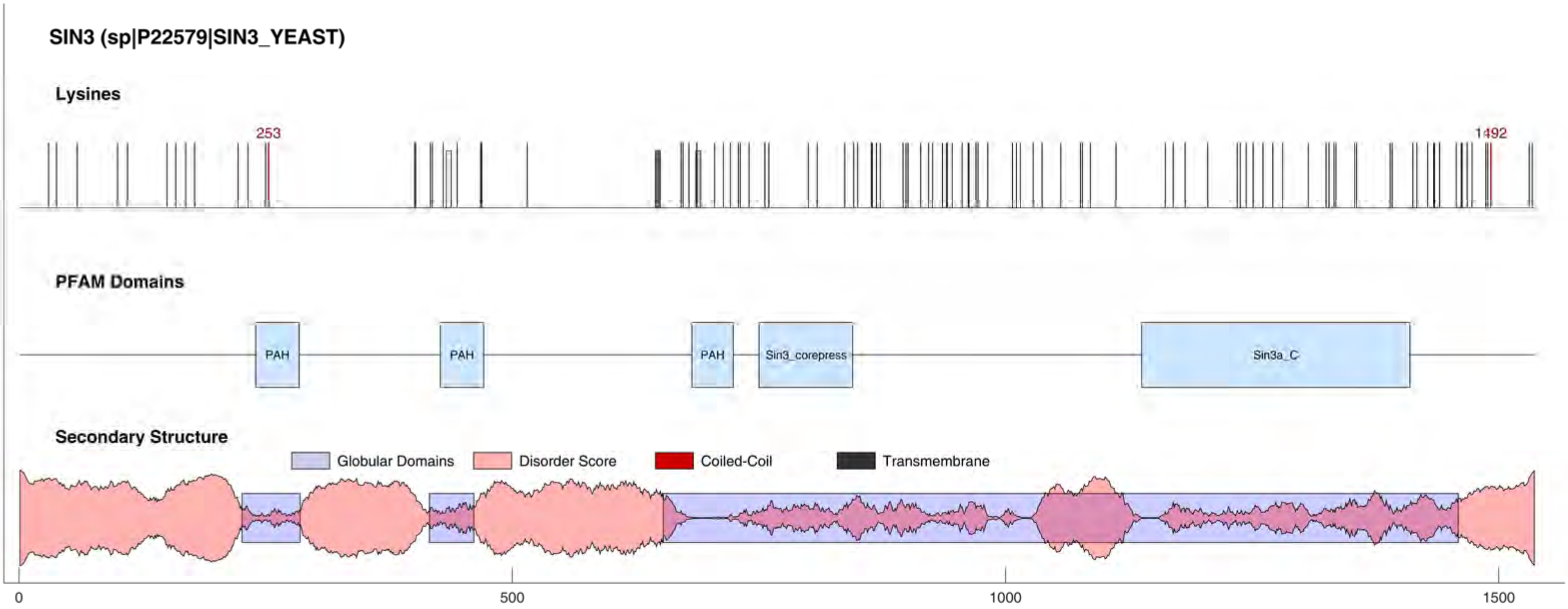
SIN3 (sp|P22579|SIN3_YEAST)
Lysines
253
1492
PFAM Domains
PAH
PAH
PAH
Sin3_corepress
Sin3a_C
Secondary Structure
Globular Domains
Disorder Score
Coiled-Coil
Transmembrane
0
500
1000
1500

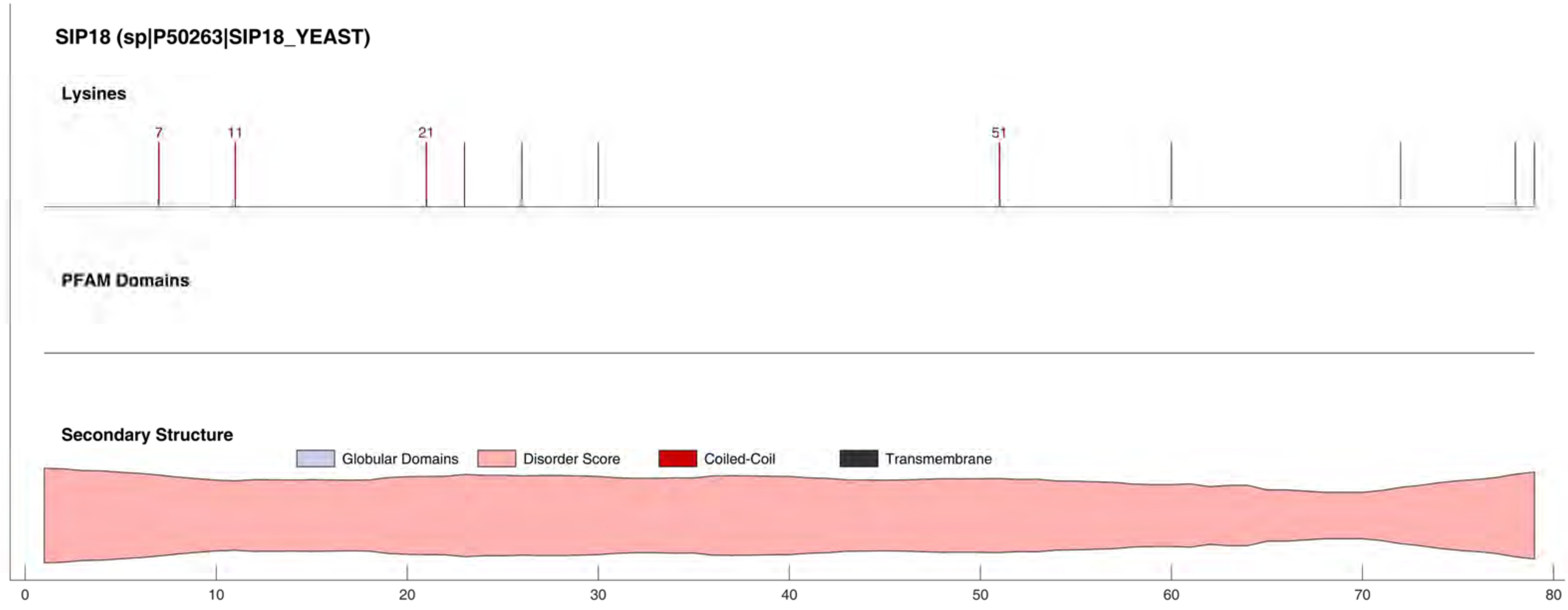
SIP18 (sp|P50263|SIP18_YEAST)
Lysines
7
11
21
51
PFAM Domains
Secondary Structure
Globular Domains
Disorder Score
Coiled-Coil
Transmembrane
0
10
20
30
40
50
60
70
80

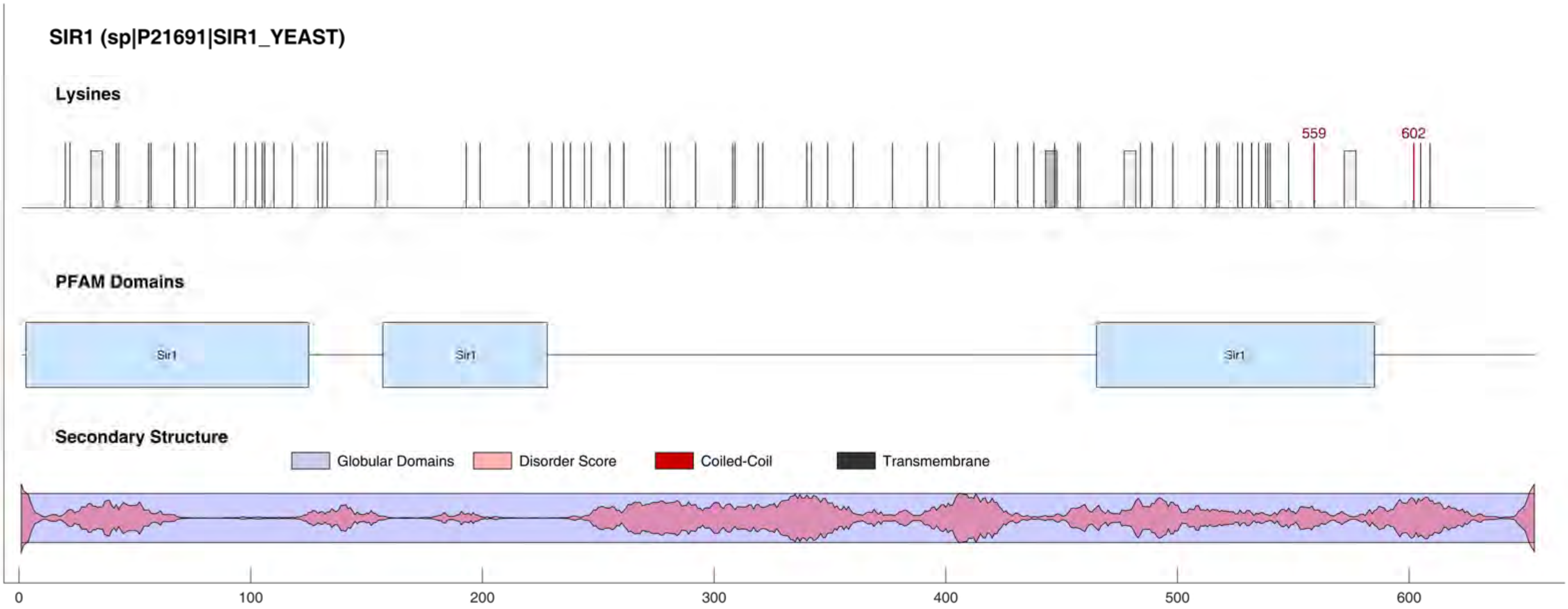
SIR1 (sp|P21691|SIR1_YEAST)
Lysines
559
602
PFAM Domains
Sir1
Sir1
Sir1
Secondary Structure
Globular Domains
Disorder Score
Coiled-Coil
Transmembrane
0
100
200
300
400
500
600

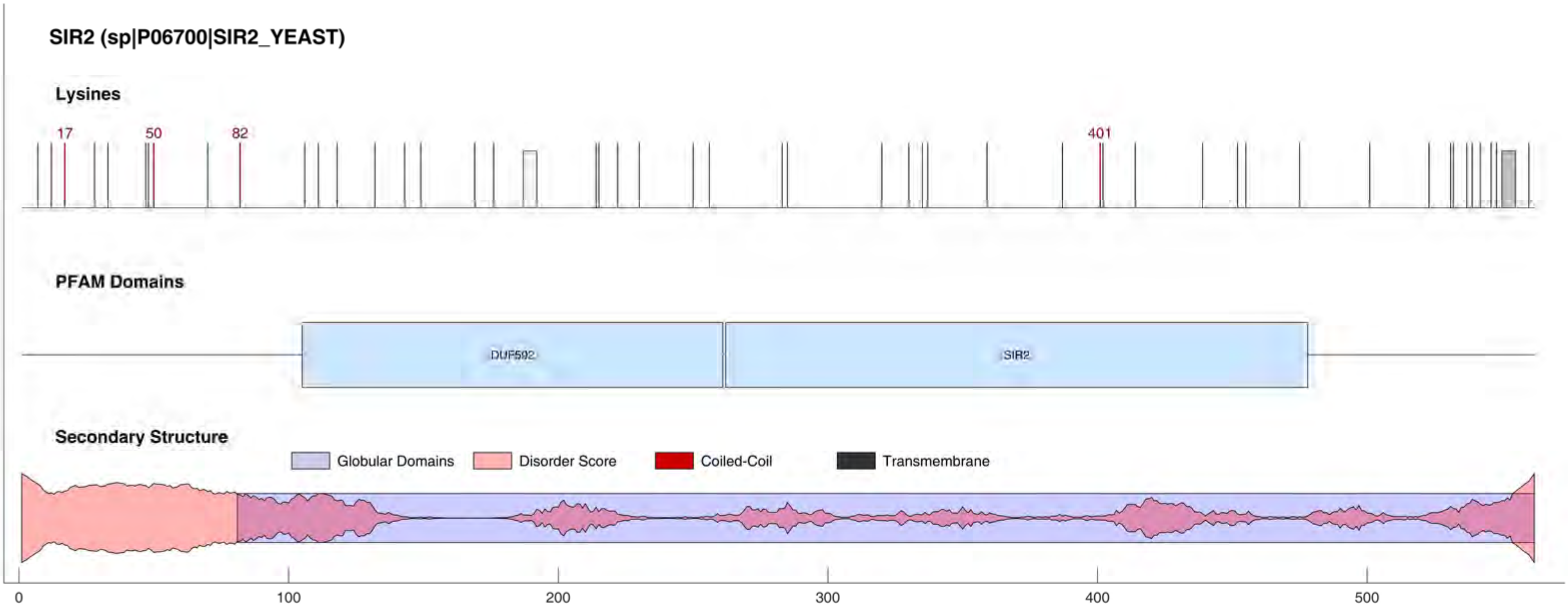
SIR2 (sp|P06700|SIR2_YEAST)
Lysines
17
50
82
401
PFAM Domains
DUF592
SIR2
Secondary Structure
Globular Domains
Disorder Score
Coiled-Coil
Transmembrane
0
100
200
300
400
500

#### SIR3 (sp|P06701|SIR3_YEAST)

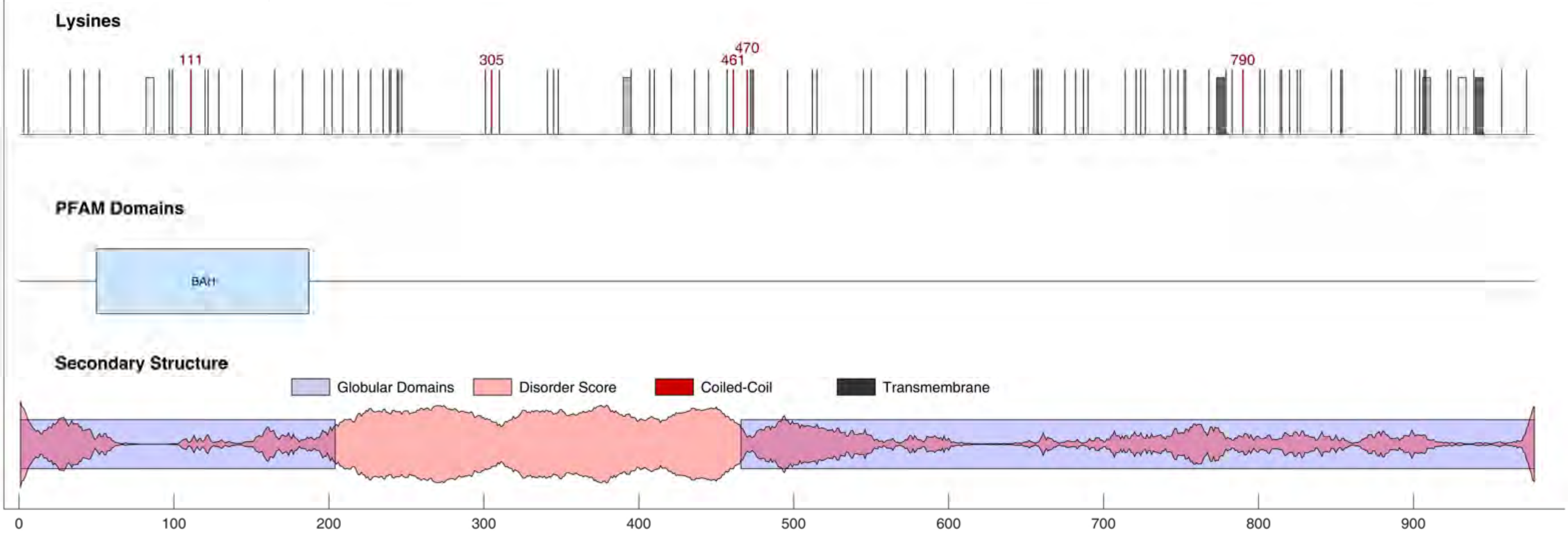

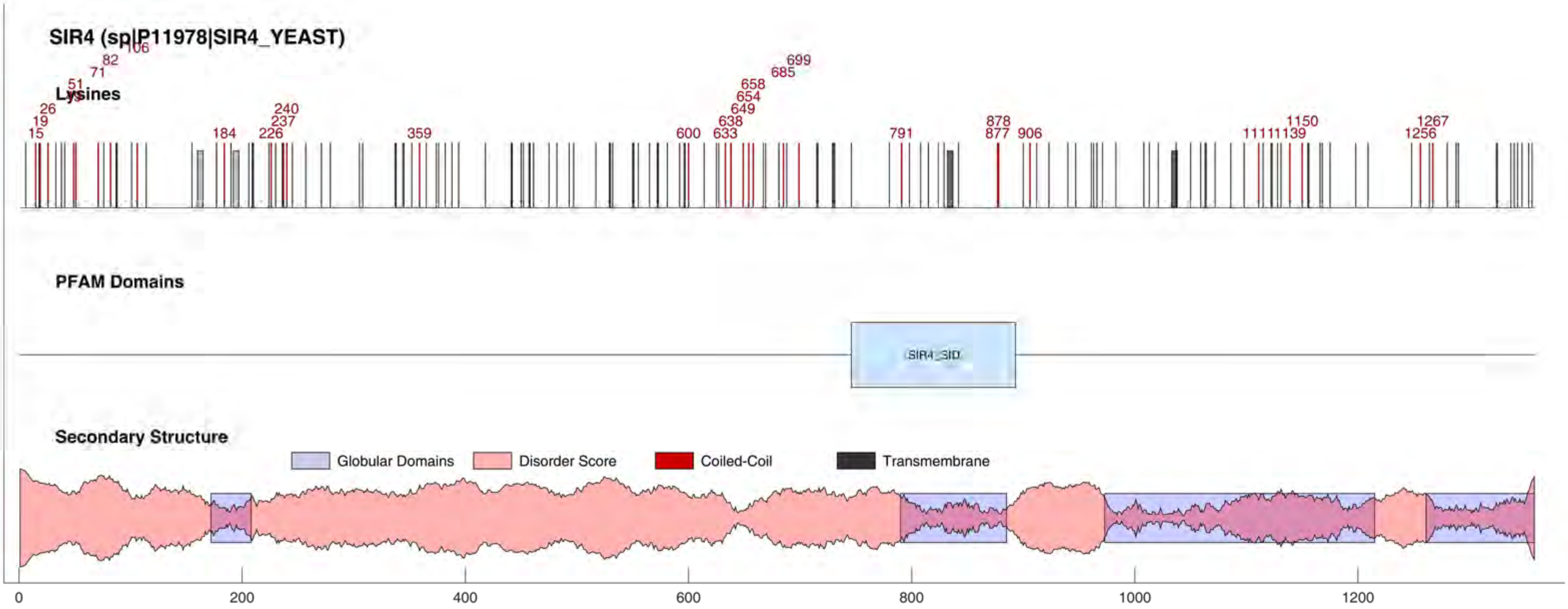

SIR4 (sp|P11978|SIR4_YEAST)
Lysines
15
19
26
51
71
82
106
184
226
237
240
359
600
633
638
649
654
658
685
699
791
877
878
906
1150
1256
1267
PFAM Domains
SIR4_SID
Secondary Structure
Globular Domains
Disorder Score
Coiled-Coil
Transmembrane
0
200
400
600
800
1000
1200

#### SIZ1 (sp|Q04195|SIZ1_YEAST)

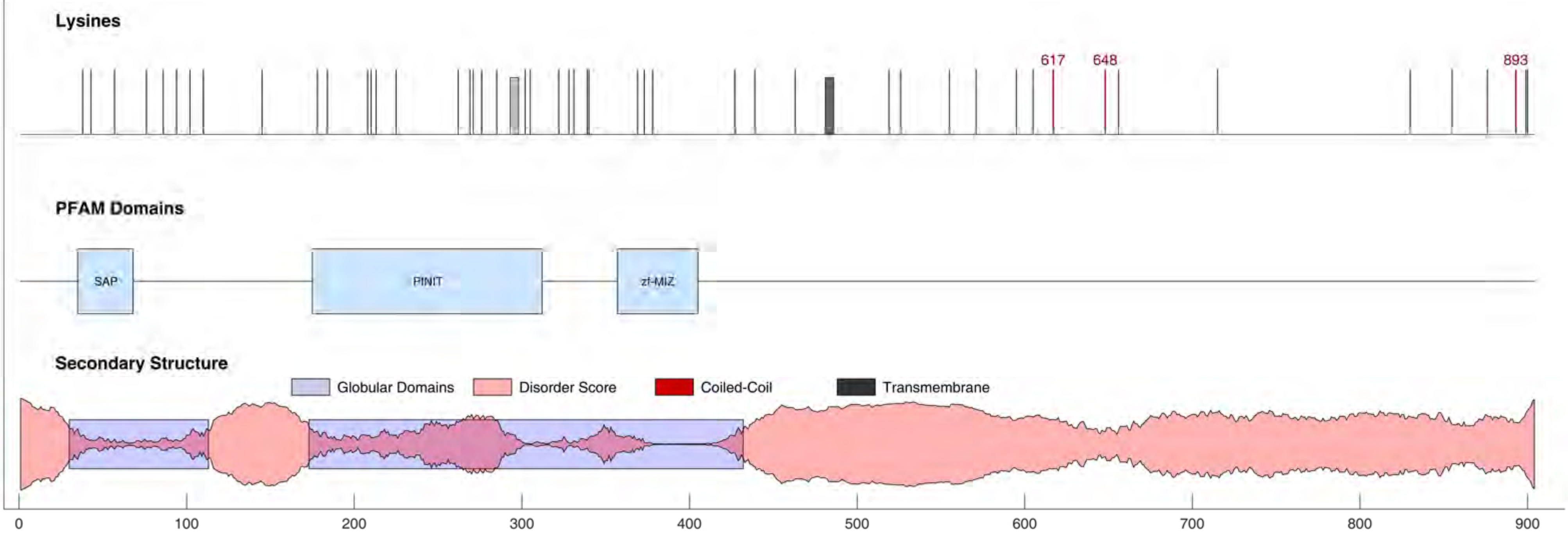

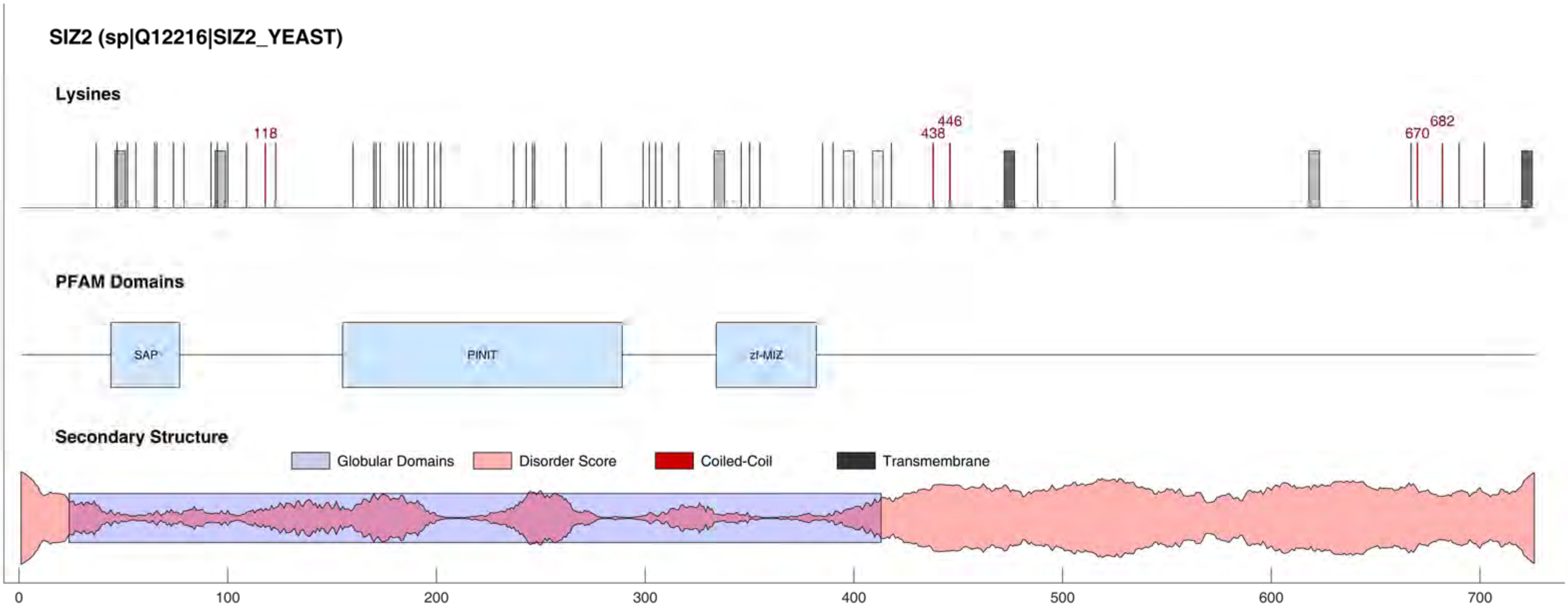

SIZ2 (sp|Q12216|SIZ2_YEAST)
Lysines
118
438
446
670
682
PFAM Domains
SAP
PINIT
zf-MIZ
Secondary Structure
Globular Domains
Disorder Score
Coiled-Coil
Transmembrane
0
100
200
300
400
500
600
700

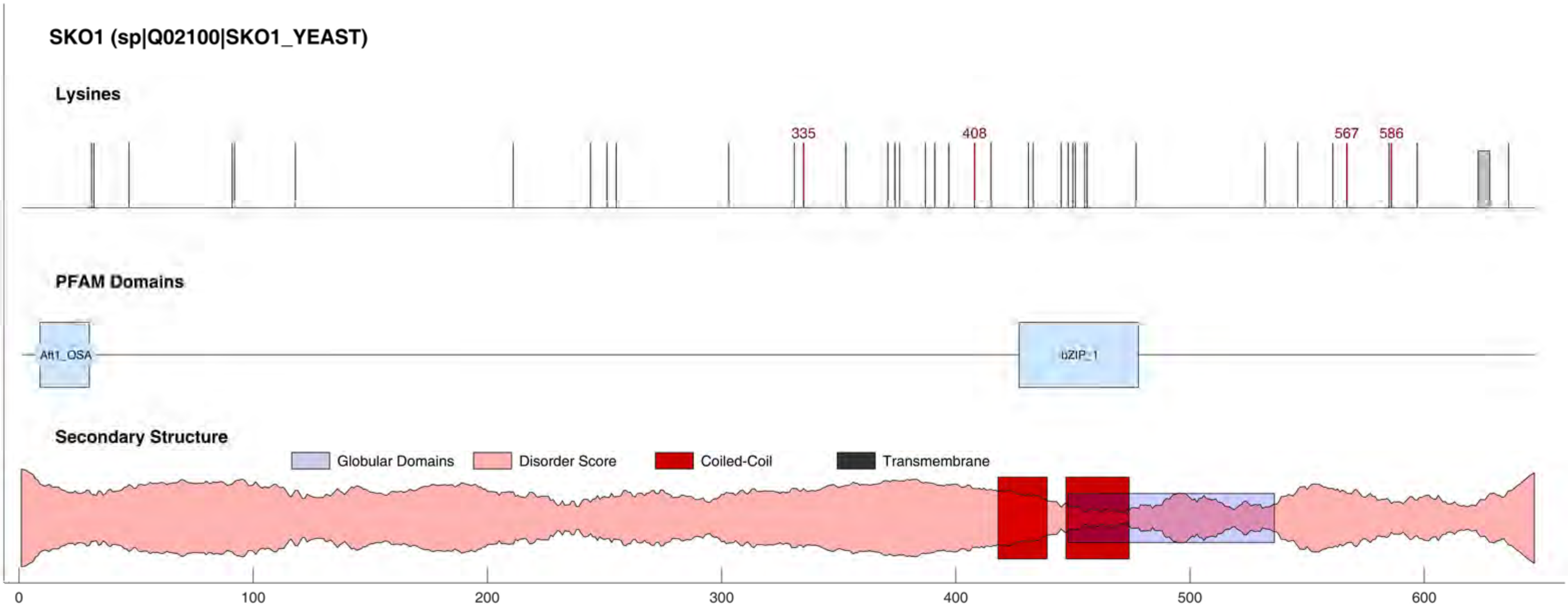
SKO1 (sp|Q02100|SKO1_YEAST)
Lysines
335
408
567
586
PFAM Domains
Aft1_OSA
bZIP_1
Secondary Structure
Globular Domains
Disorder Score
Coiled-Coil
Transmembrane
0
100
200
300
400
500
600

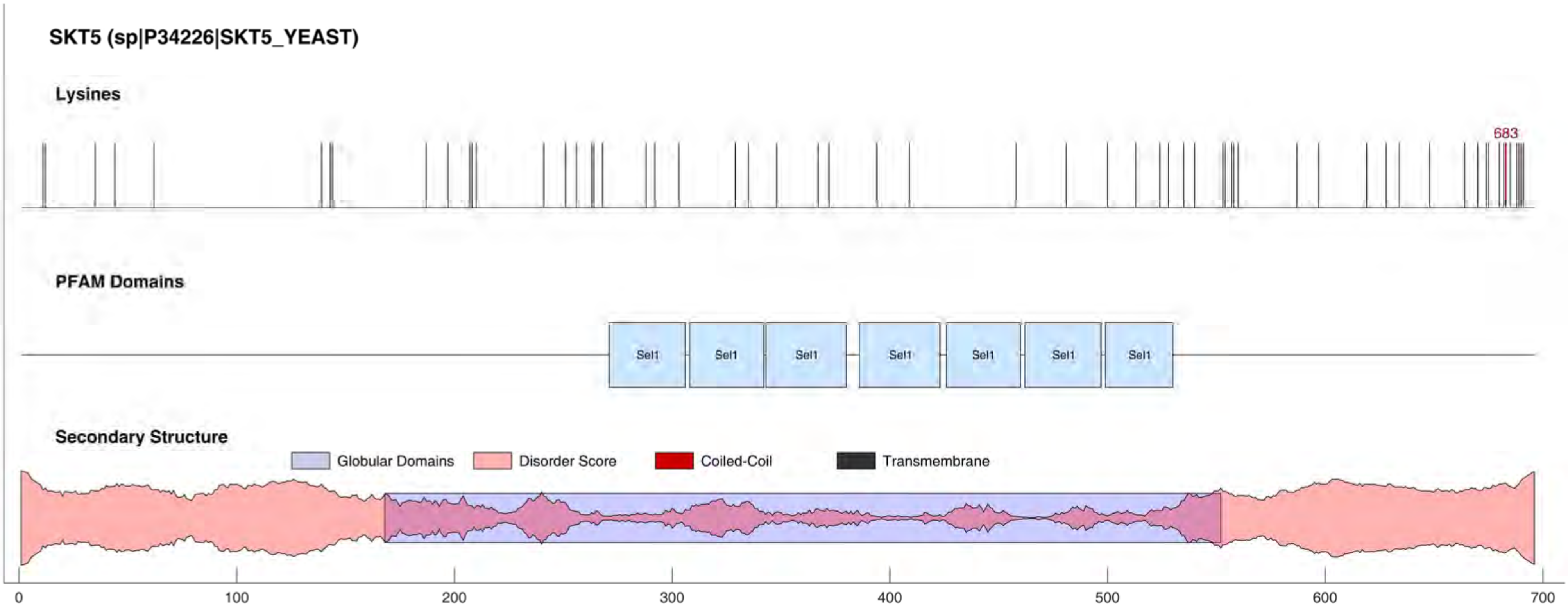
SKT5 (sp|P34226|SKT5_YEAST)
Lysines
683
PFAM Domains
Sel1
Sel1
Sel1
Sel1
Sel1
Sel1
Sel1
Secondary Structure
Globular Domains
Disorder Score
Coiled-Coil
Transmembrane
0
100
200
300
400
500
600
700

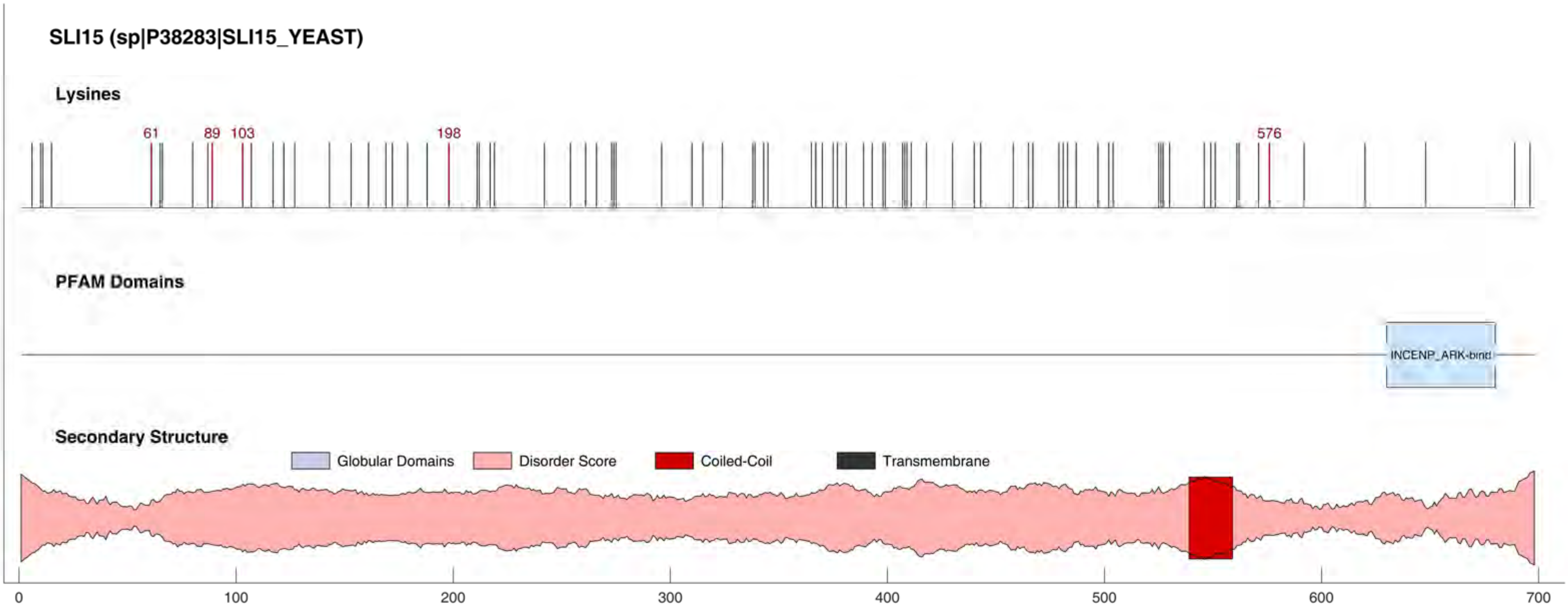
SLI15 (sp|P38283|SLI15_YEAST)
Lysines
61
89
103
198
576
PFAM Domains
INCENP_ARK-bind
Secondary Structure
Globular Domains
Disorder Score
Coiled-Coil
Transmembrane
0
100
200
300
400
500
600
700

#### SLK19 (sp|Q08581|SLK19_YEAST)

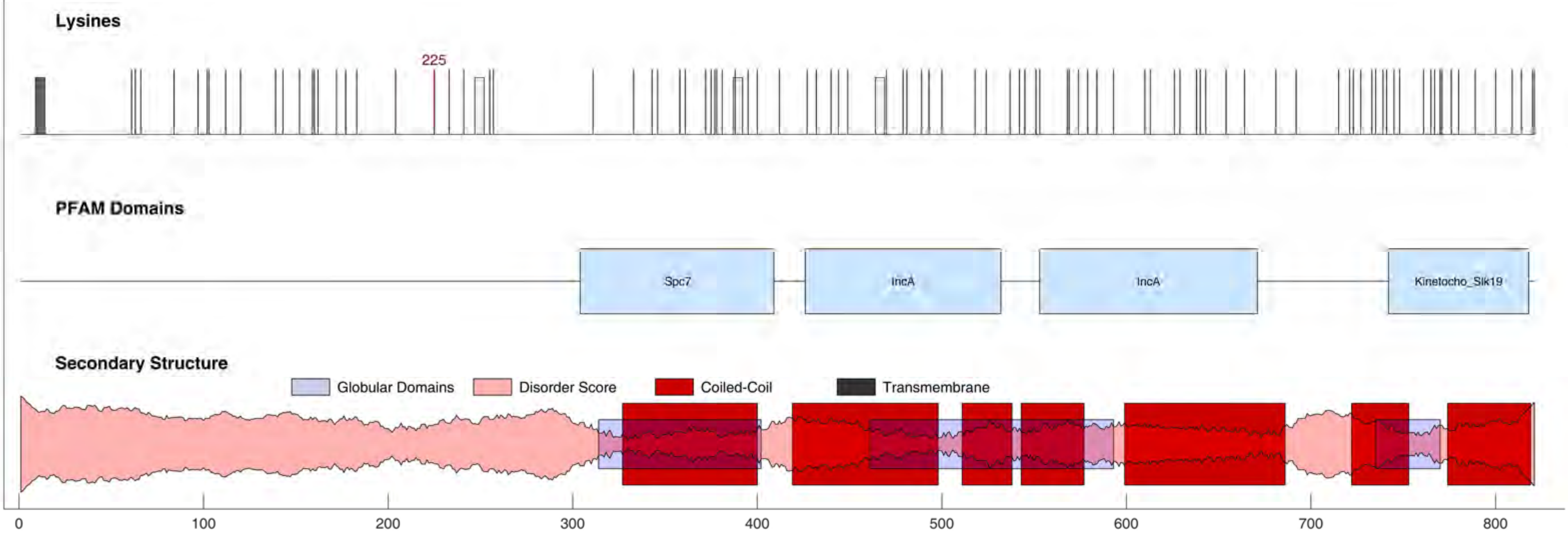

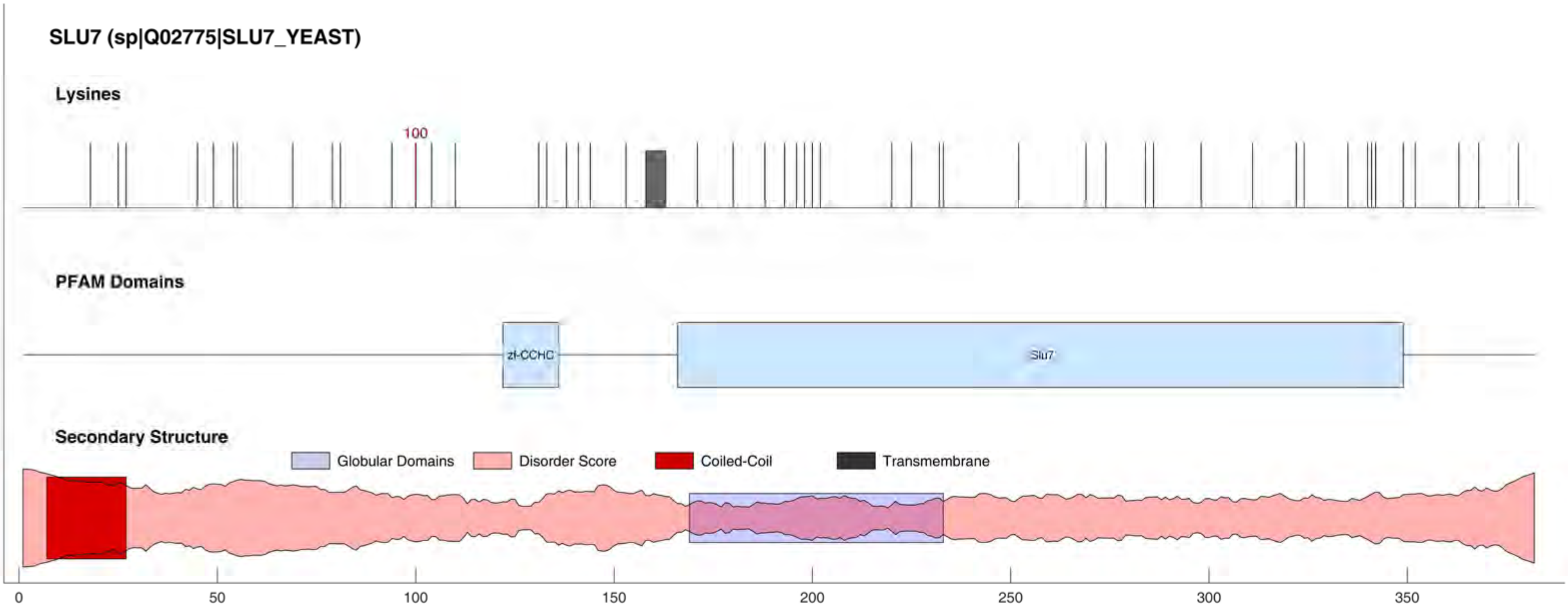

SLU7 (sp|Q02775|SLU7_YEAST)
Lysines
100
PFAM Domains
zf-CCHC
Slu7
Secondary Structure
Globular Domains
Disorder Score
Coiled-Coil
Transmembrane
0
50
100
150
200
250
300
350

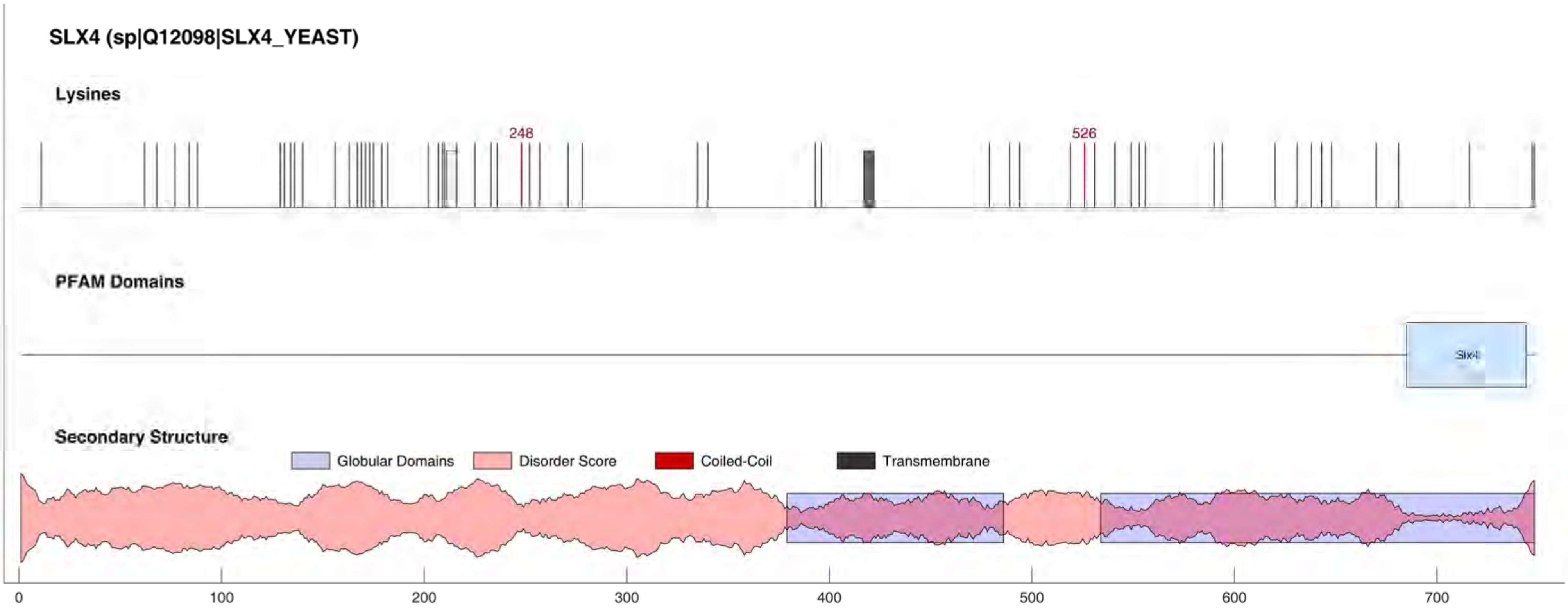

SLX4 (sp|Q12098|SLX4_YEAST)
Lysines
248
526
PFAM Domains
Slx4
Secondary Structure
Globular Domains
Disorder Score
Coiled-Coil
Transmembrane
0
100
200
300
400
500
600
700

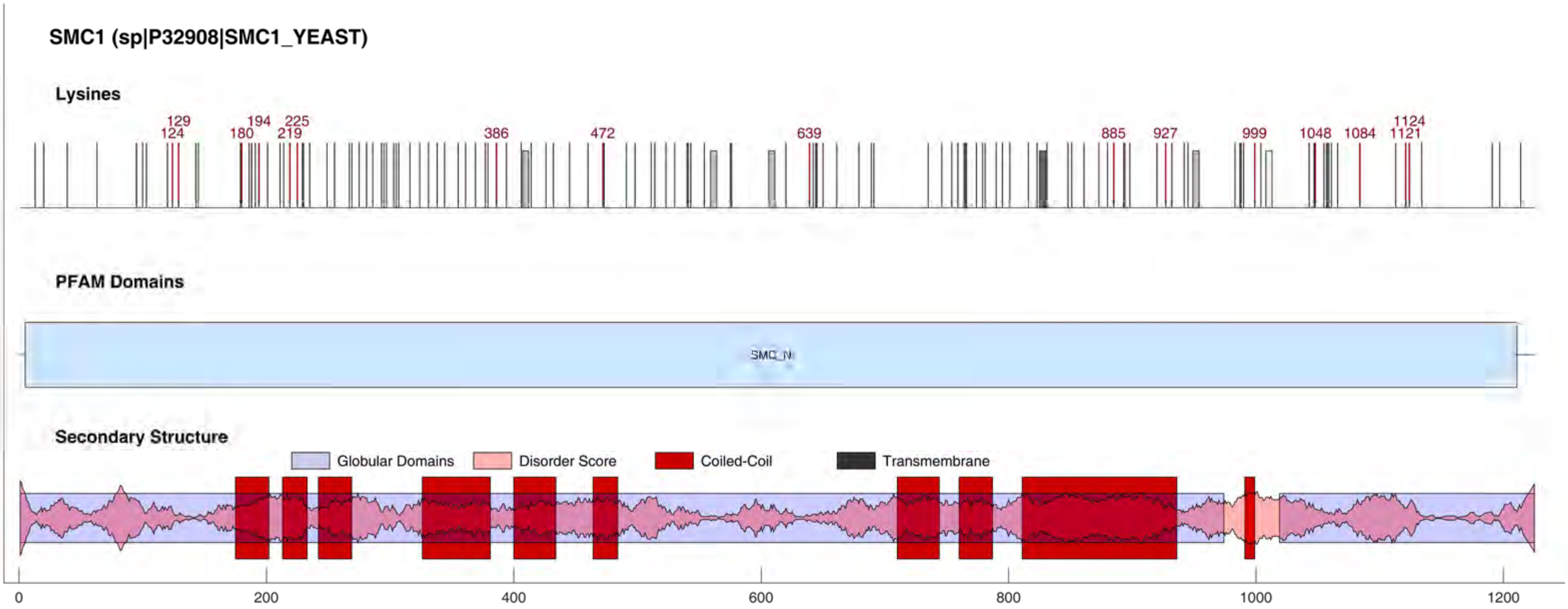

SMC1 (sp|P32908|SMC1_YEAST)
Lysines
129
124
180
194
225
219
386
472
639
885
927
999
1048
1084
1124
1121
PFAM Domains
SMC_N
Secondary Structure
Globular Domains
Disorder Score
Coiled-Coil
Transmembrane
0
200
400
600
800
1000
1200

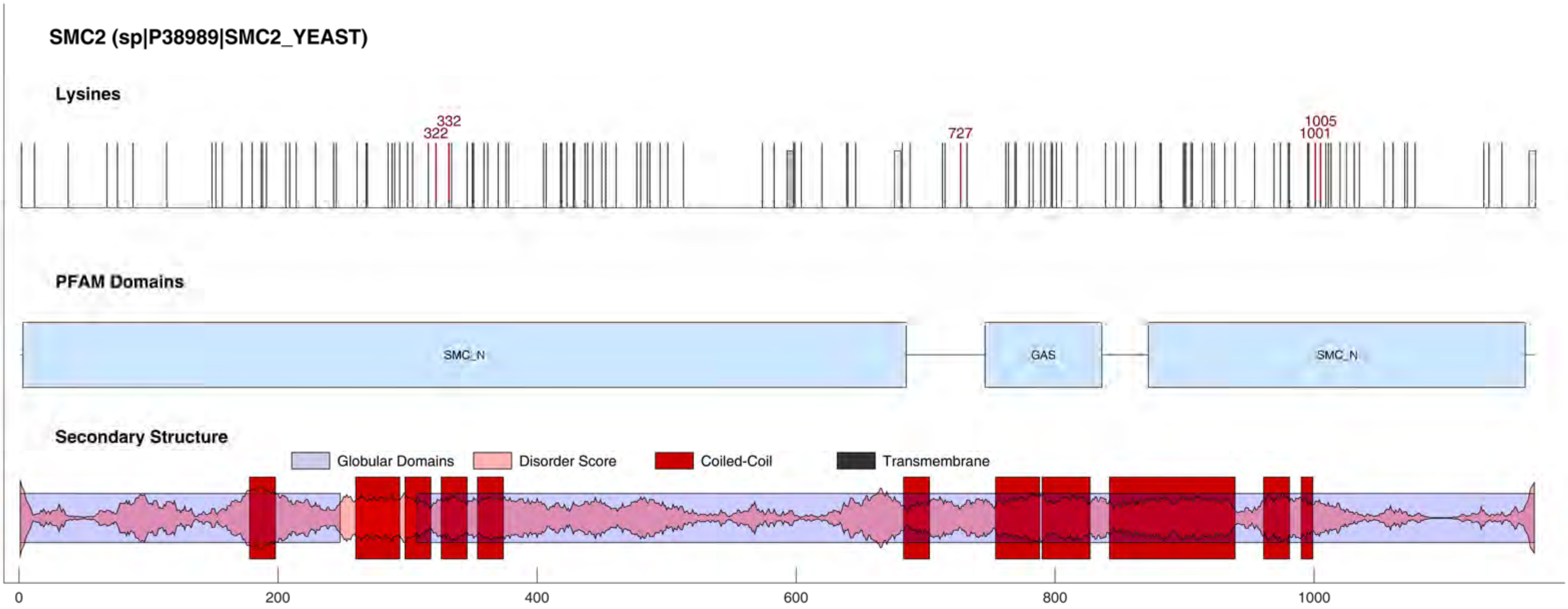

SMC2 (sp|P38989|SMC2_YEAST)
Lysines
332
322
727
1005
1001
PFAM Domains
SMC_N
GAS
SMC_N
Secondary Structure
Globular Domains
Disorder Score
Coiled-Coil
Transmembrane
0
200
400
600
800
1000

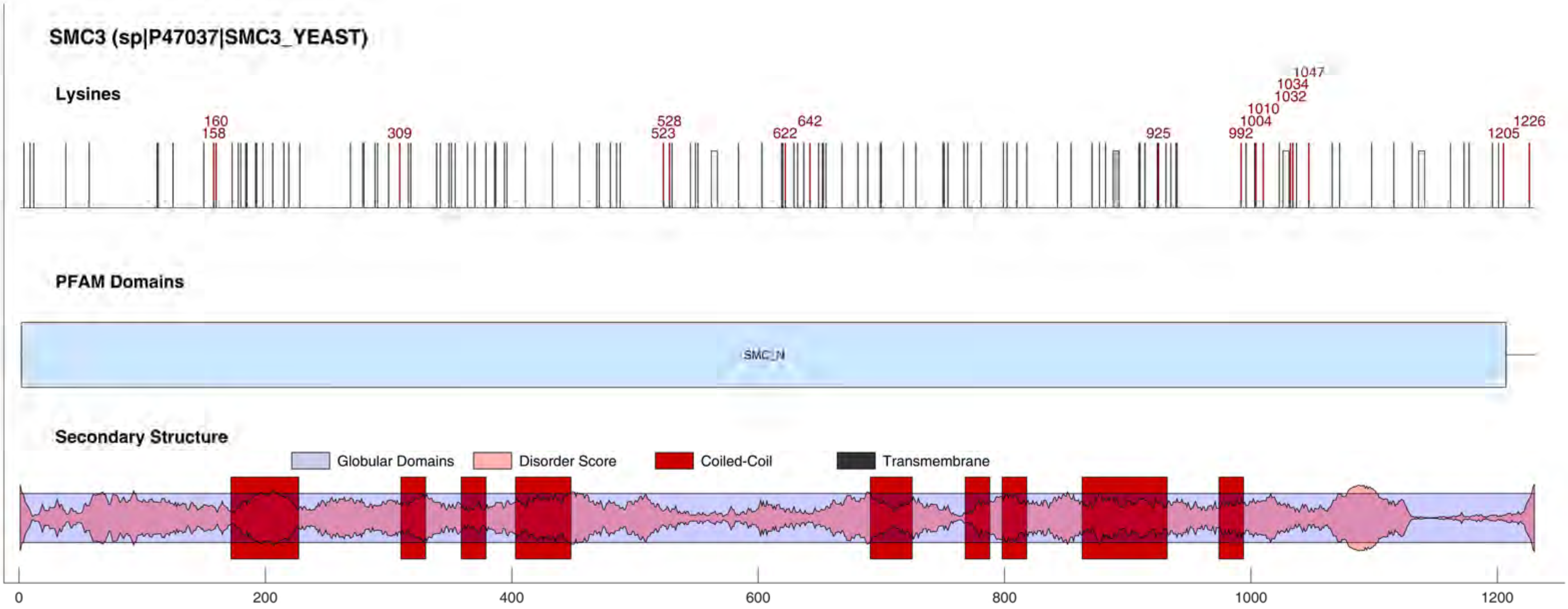

SMC3 (sp|P47037|SMC3_YEAST)
Lysines
160
158
309
528
523
642
622
925
1047
1034
1032
1010
1004
992
1226
1205
PFAM Domains
SMC_N
Secondary Structure
Globular Domains
Disorder Score
Coiled-Coil
Transmembrane
0
200
400
600
800
1000
1200

#### SMC4 (sp|Q12267|SMC4_YEAST)

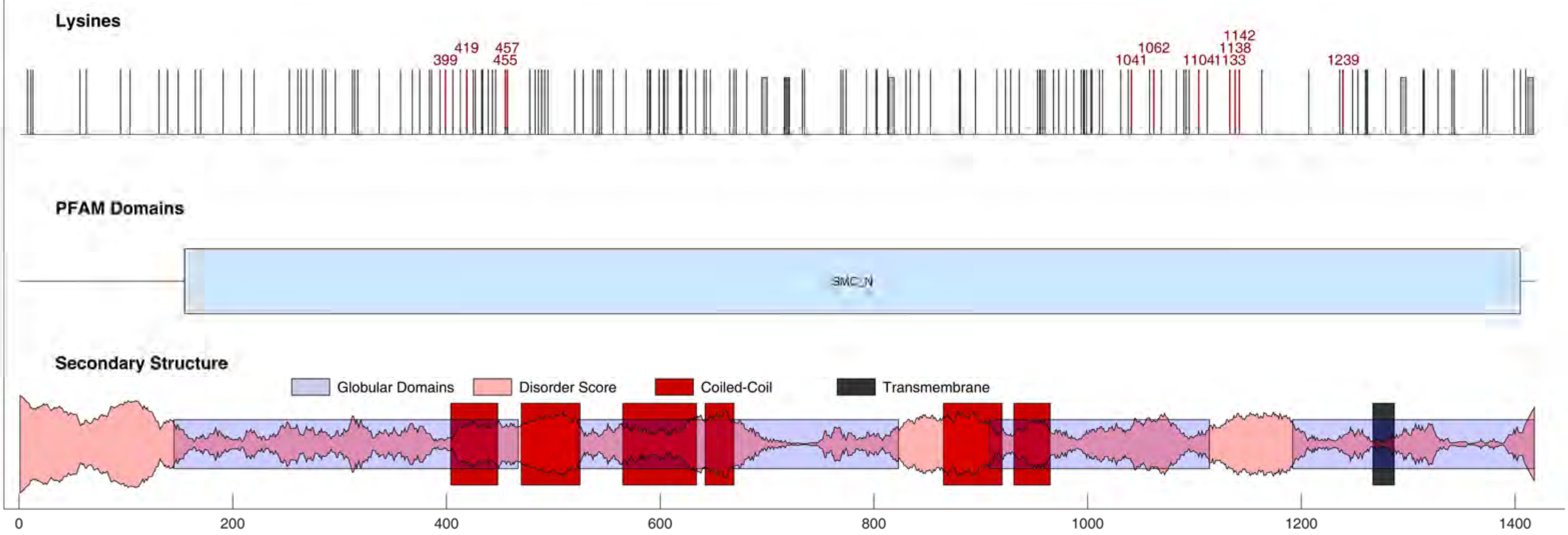

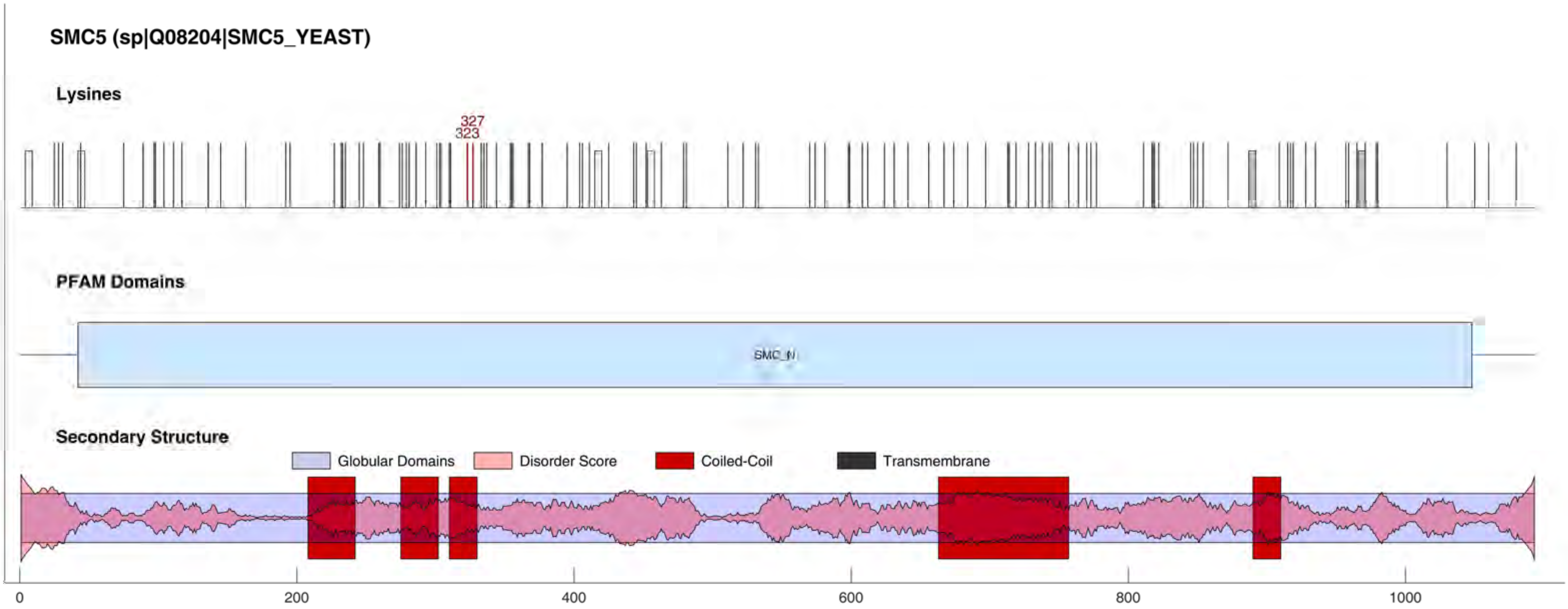
SMC5 (sp|Q08204|SMC5_YEAST)
Lysines
327
323
PFAM Domains
SMC_N
Secondary Structure
Globular Domains
Disorder Score
Coiled-Coil
Transmembrane
0
200
400
600
800
1000

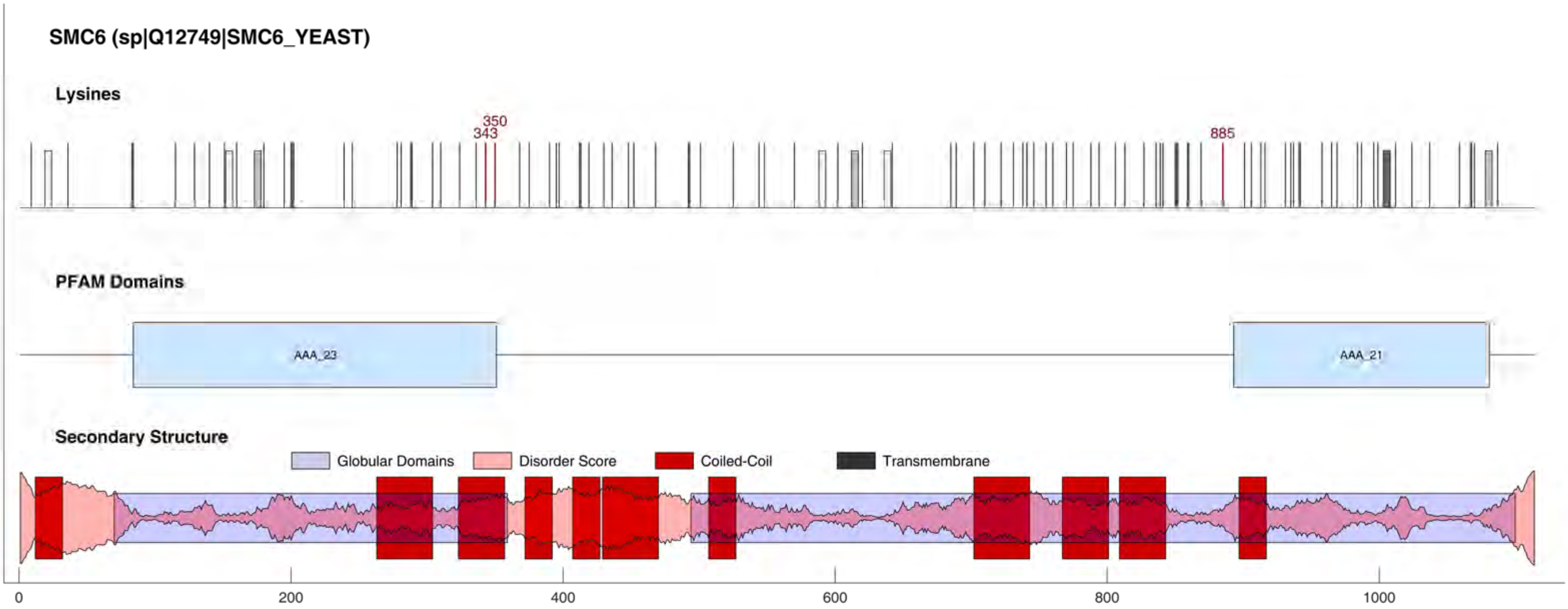

SMC6 (sp|Q12749|SMC6_YEAST)
Lysines
350
343
885
PFAM Domains
AAA_23
AAA_21
Secondary Structure
Globular Domains
Disorder Score
Coiled-Coil
Transmembrane
0
200
400
600
800
1000

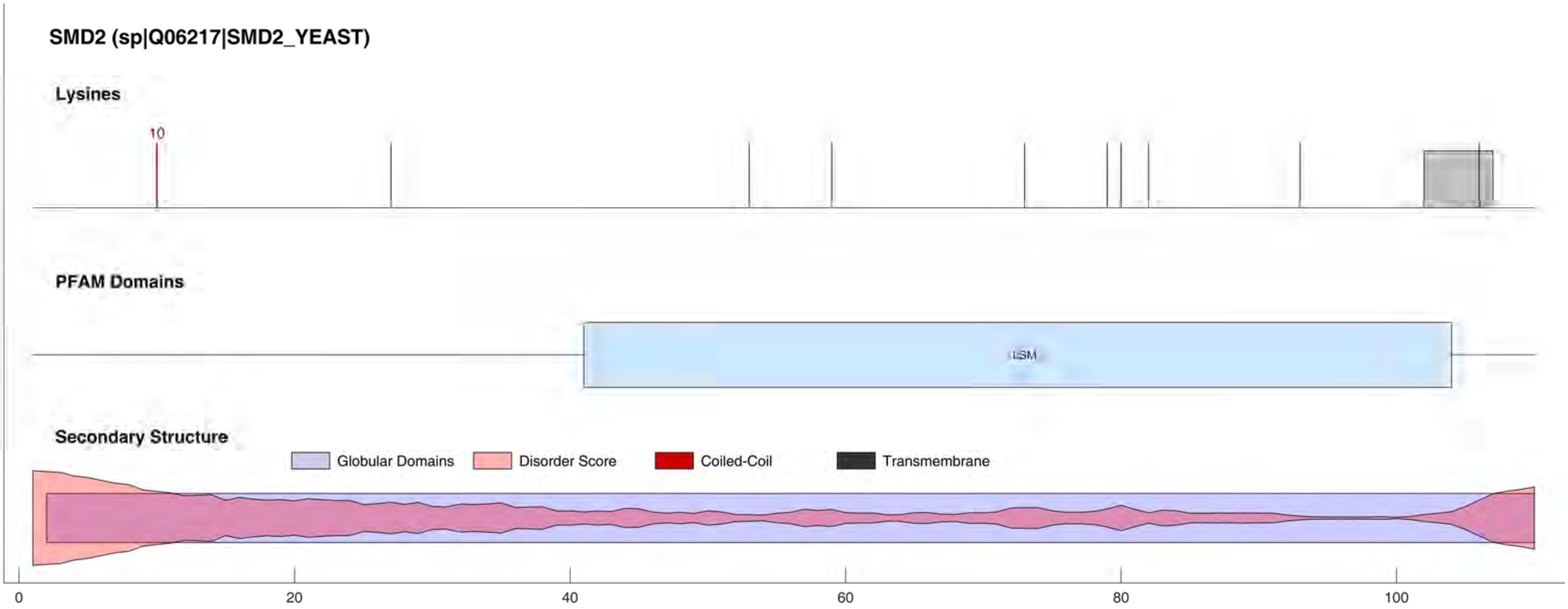
SMD2 (sp|Q06217|SMD2_YEAST)
Lysines
10
PFAM Domains
LSM
Secondary Structure
Globular Domains
Disorder Score
Coiled-Coil
Transmembrane
0
20
40
60
80
100

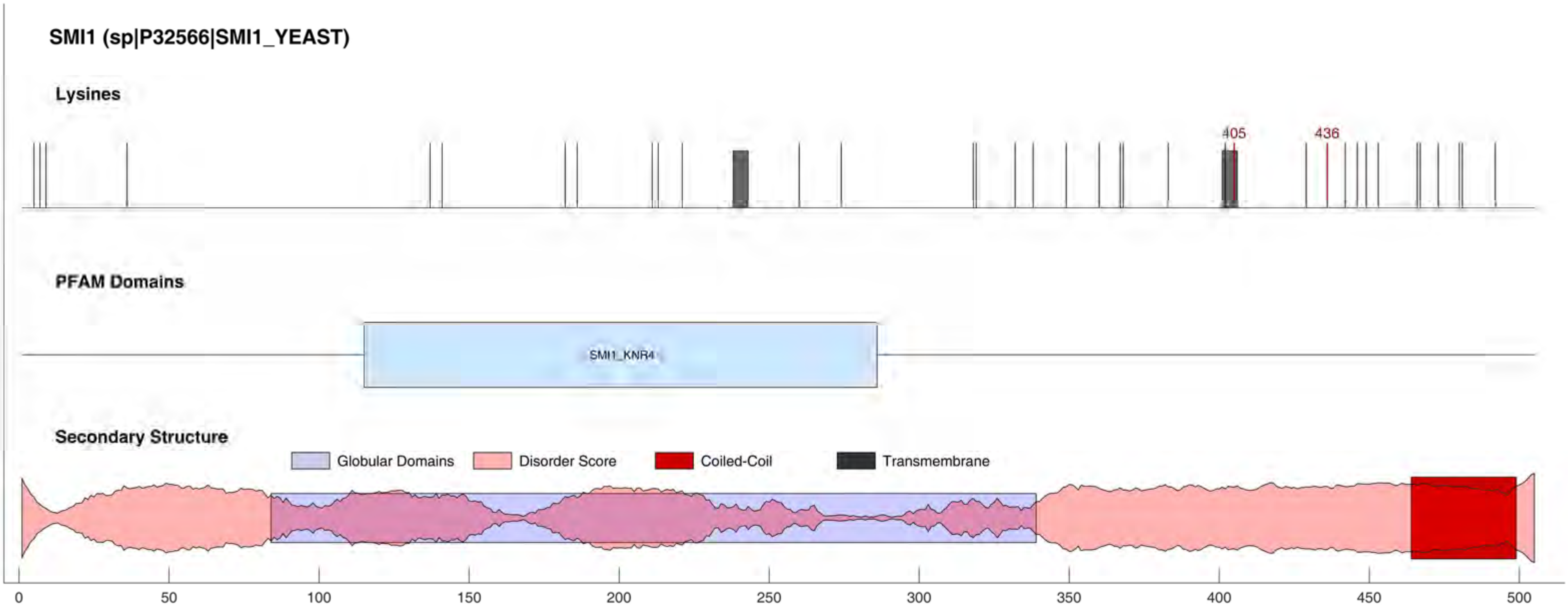
SMI1 (sp|P32566|SMI1_YEAST)
Lysines
405
436
PFAM Domains
SMI1_KNR4
Secondary Structure
Globular Domains
Disorder Score
Coiled-Coil
Transmembrane
0
50
100
150
200
250
300
350
400
450
500

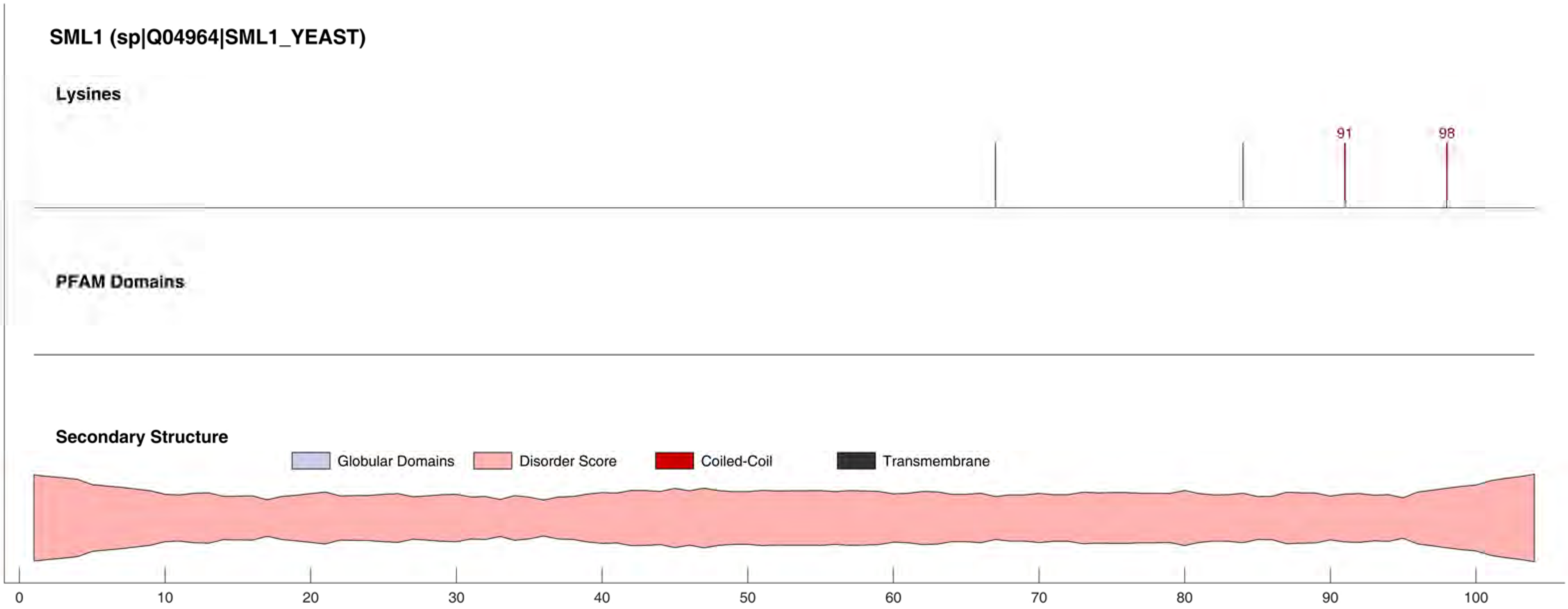
SML1 (sp|Q04964|SML1_YEAST)
Lysines
91
98
PFAM Domains
Secondary Structure
Globular Domains
Disorder Score
Coiled-Coil
Transmembrane
0
10
20
30
40
50
60
70
80
90
100

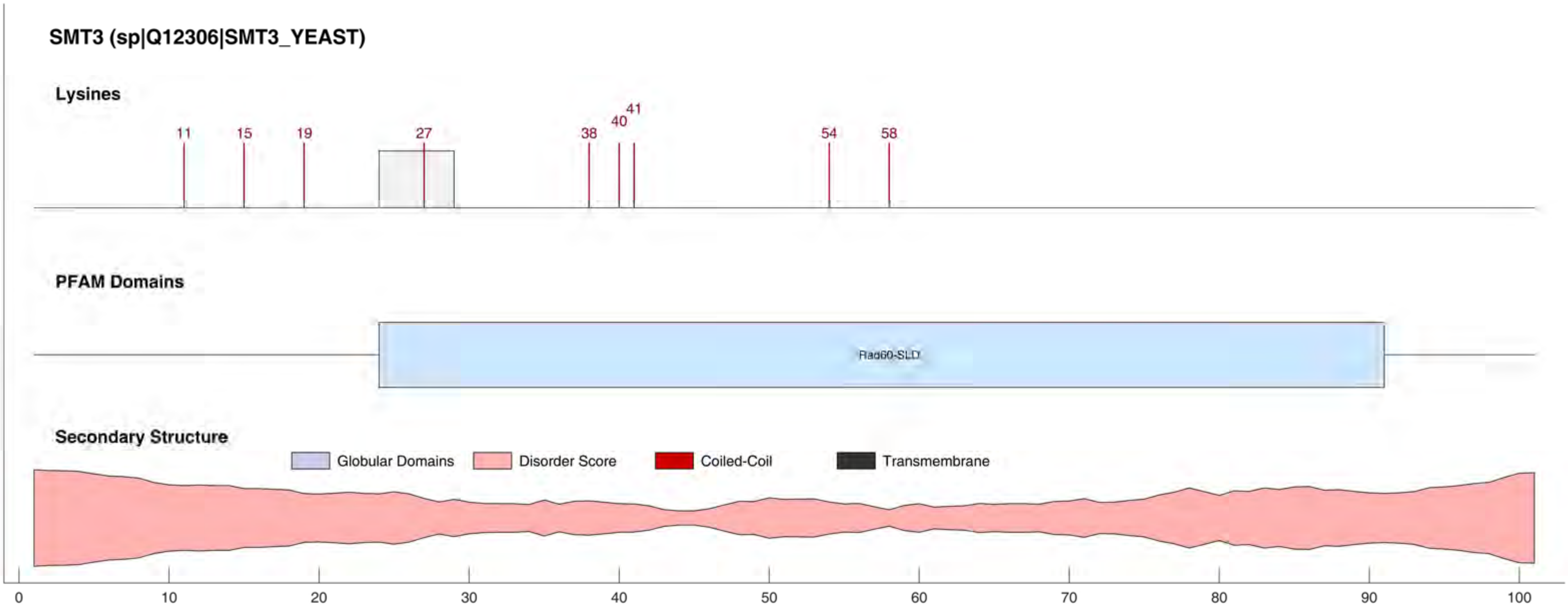

SMT3 (sp|Q12306|SMT3_YEAST)
Lysines
11
15
19
27
38
40
41
54
58
PFAM Domains
Rad60-SLD
Secondary Structure
Globular Domains
Disorder Score
Coiled-Coil
Transmembrane
0
10
20
30
40
50
60
70
80
90
100

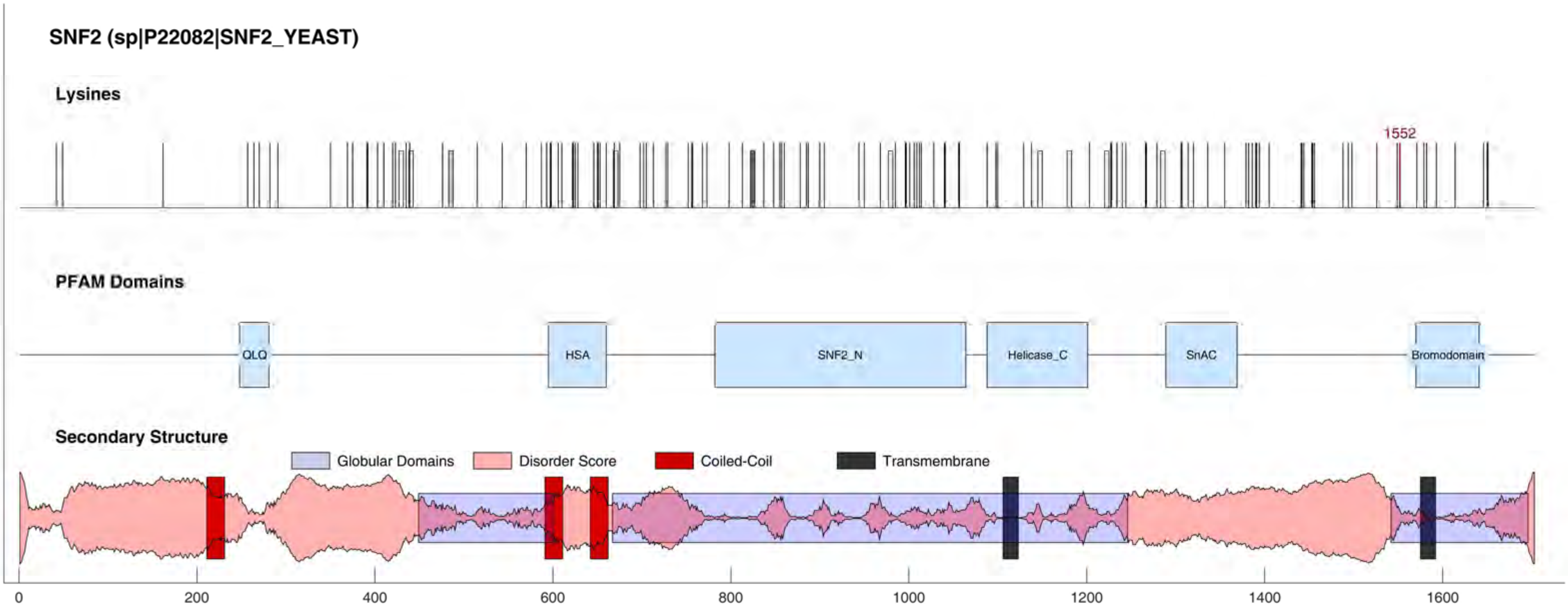
SNF2 (sp|P22082|SNF2_YEAST)
Lysines
1552
PFAM Domains
QLQ
HSA
SNF2_N
Helicase_C
SnAC
Bromodomain
Secondary Structure
Globular Domains
Disorder Score
Coiled-Coil
Transmembrane
0
200
400
600
800
1000
1200
1400
1600

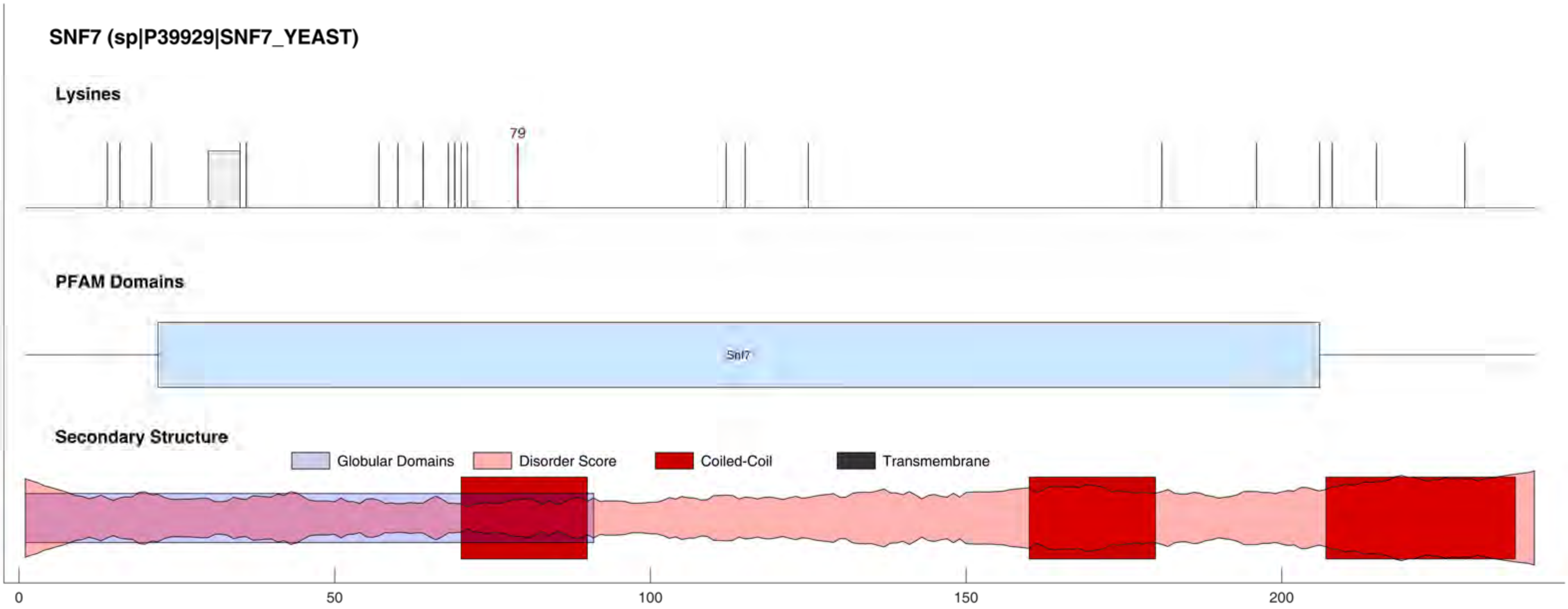

SNF7 (sp|P39929|SNF7_YEAST)
Lysines
79
PFAM Domains
Snf7
Secondary Structure
Globular Domains
Disorder Score
Coiled-Coil
Transmembrane
0
50
100
150
200

#### SNF12 (sp|P53628|SNF12_YEAST)

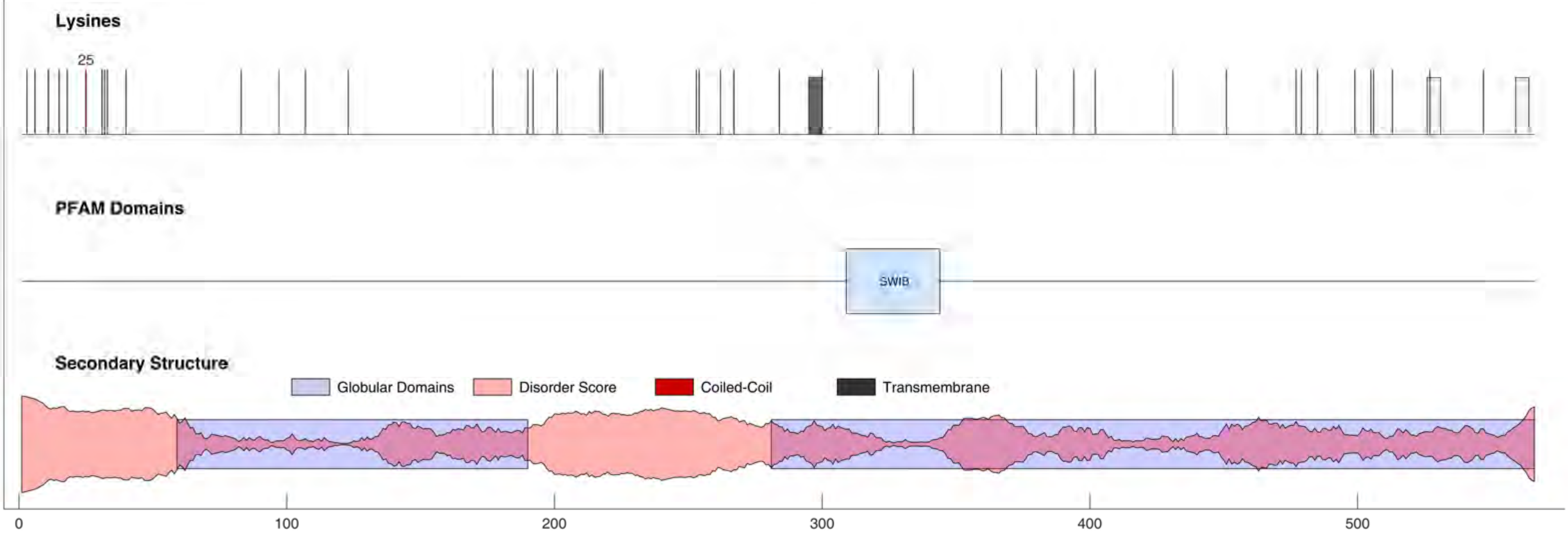

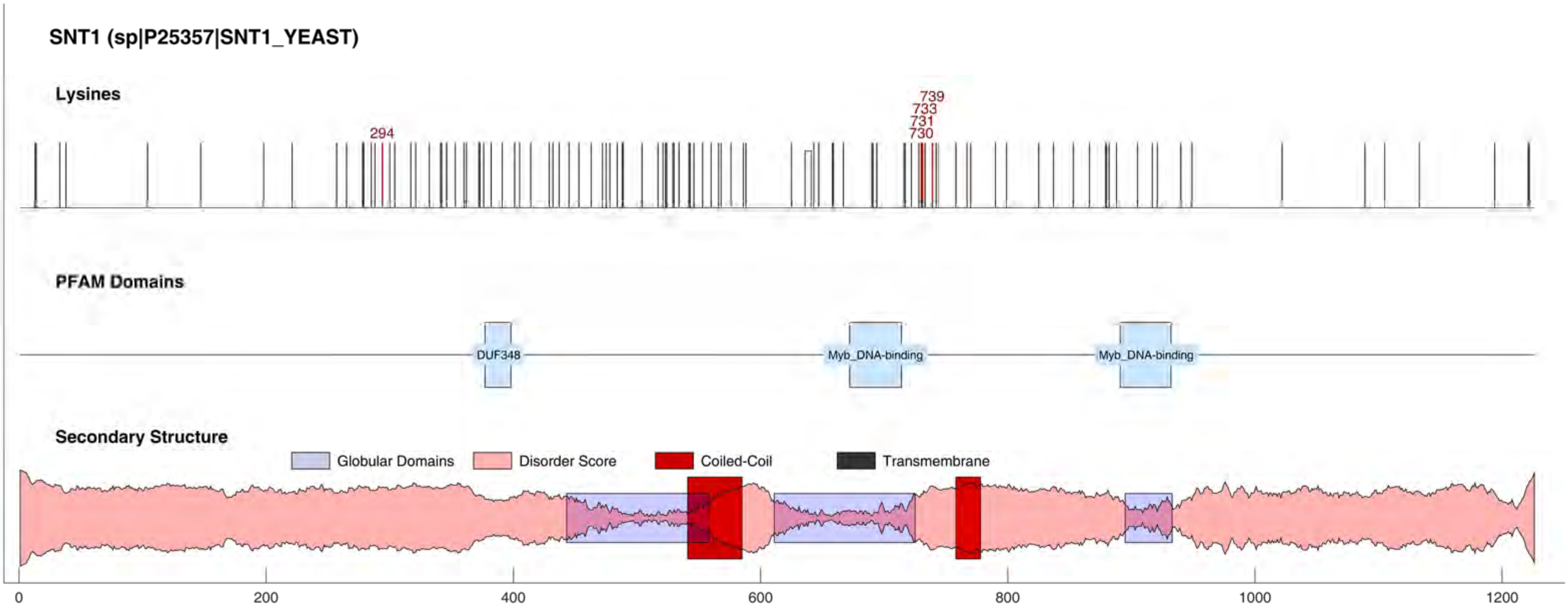
SNT1 (sp|P25357|SNT1_YEAST)
Lysines
294
739
733
731
730
PFAM Domains
DUF348
Myb_DNA-binding
Myb_DNA-binding
Secondary Structure
Globular Domains
Disorder Score
Coiled-Coil
Transmembrane
0
200
400
600
800
1000
1200

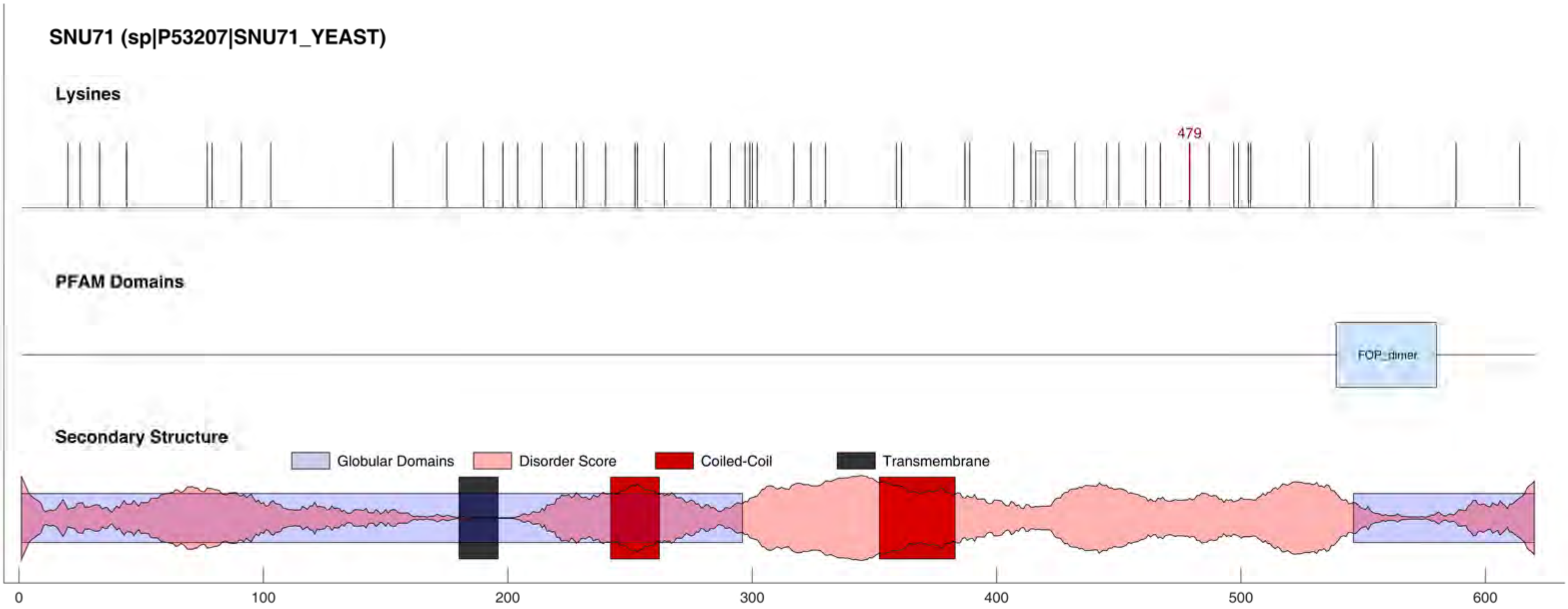
SNU71 (sp|P53207|SNU71_YEAST)
Lysines
479
PFAM Domains
FOP_dimer
Secondary Structure
Globular Domains
Disorder Score
Coiled-Coil
Transmembrane
0
100
200
300
400
500
600

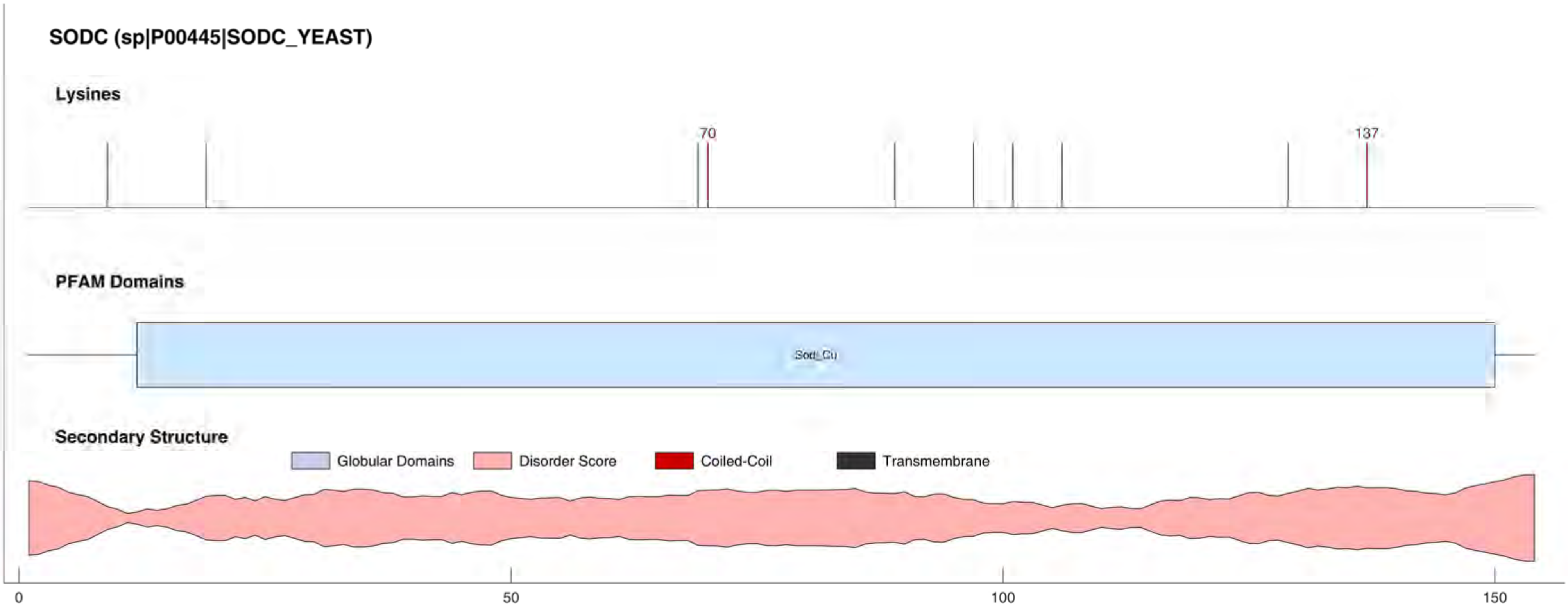

SODC (sp|P00445|SODC_YEAST)
Lysines
70
137
PFAM Domains
Sod_Cu
Secondary Structure
Globular Domains
Disorder Score
Coiled-Coil
Transmembrane
0
50
100
150

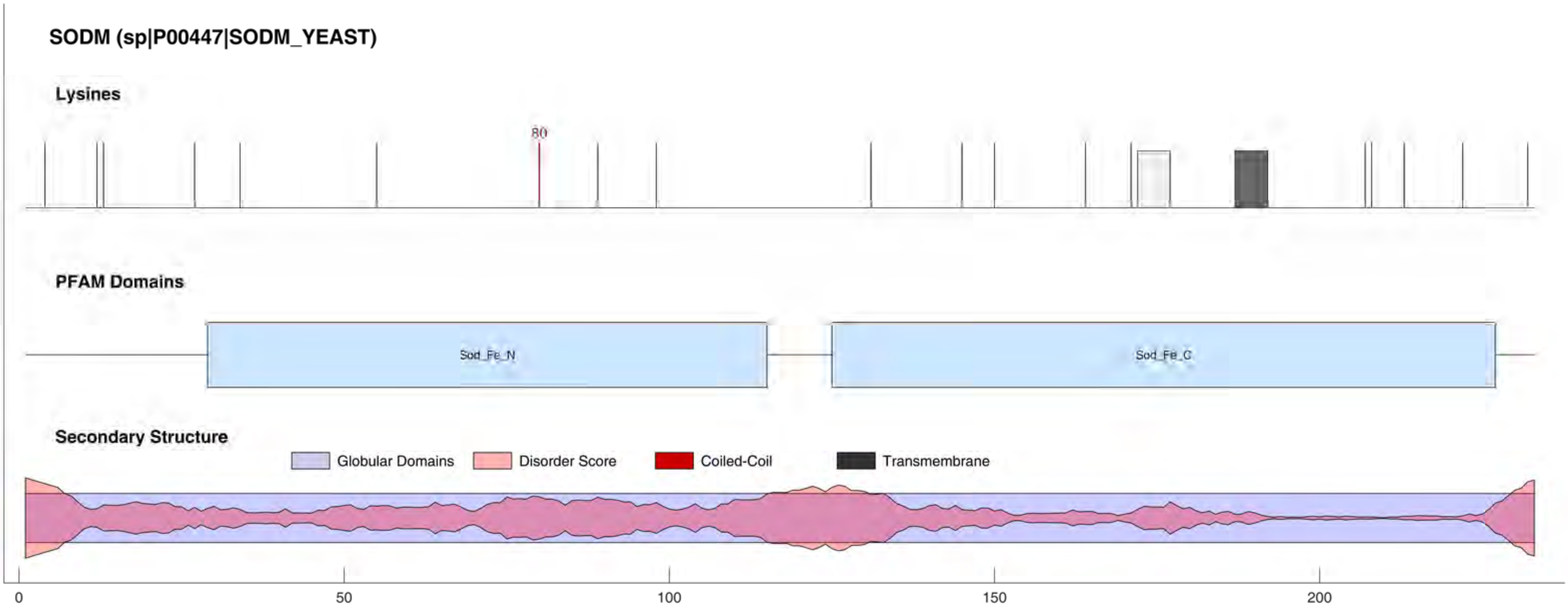
SODM (sp|P00447|SODM_YEAST)
Lysines
80
PFAM Domains
Sod_Fe_N
Sod_Fe_C
Secondary Structure
Globular Domains
Disorder Score
Coiled-Coil
Transmembrane
0
50
100
150
200

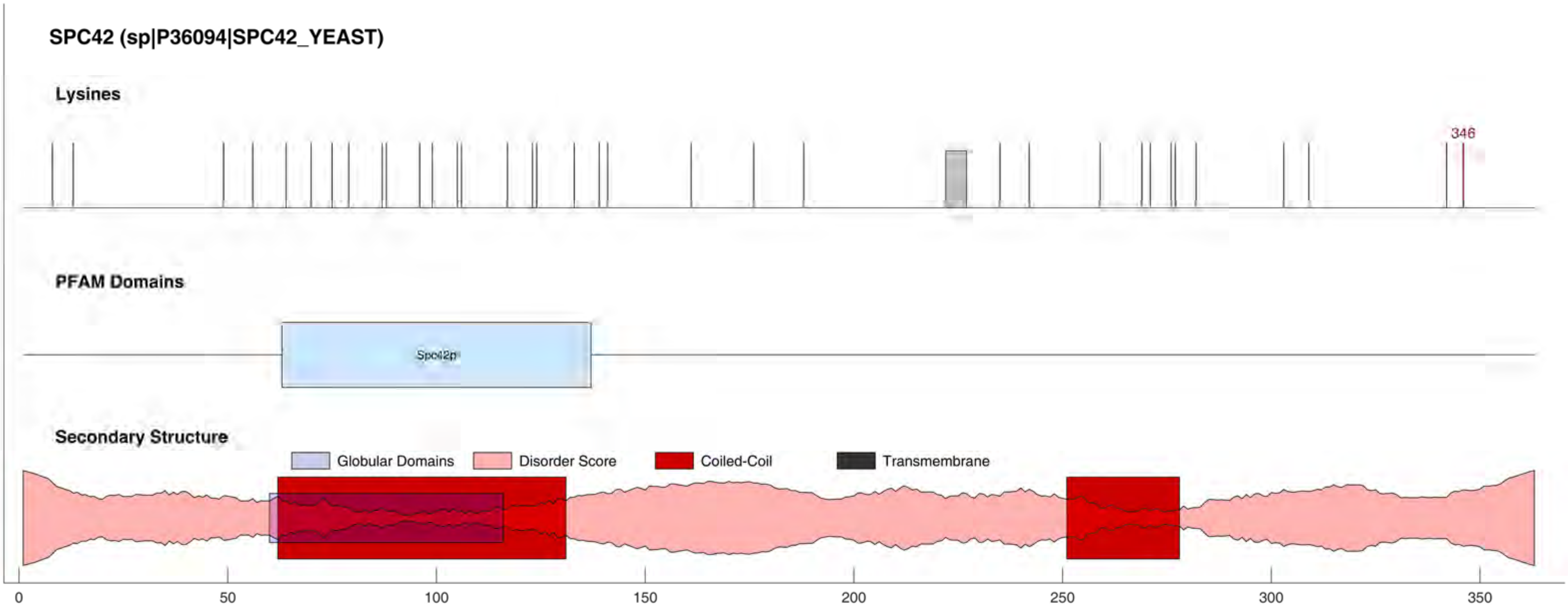
SPC42 (sp|P36094|SPC42_YEAST)
Lysines
346
PFAM Domains
Spc42p
Secondary Structure
Globular Domains
Disorder Score
Coiled-Coil
Transmembrane
0
50
100
150
200
250
300
350

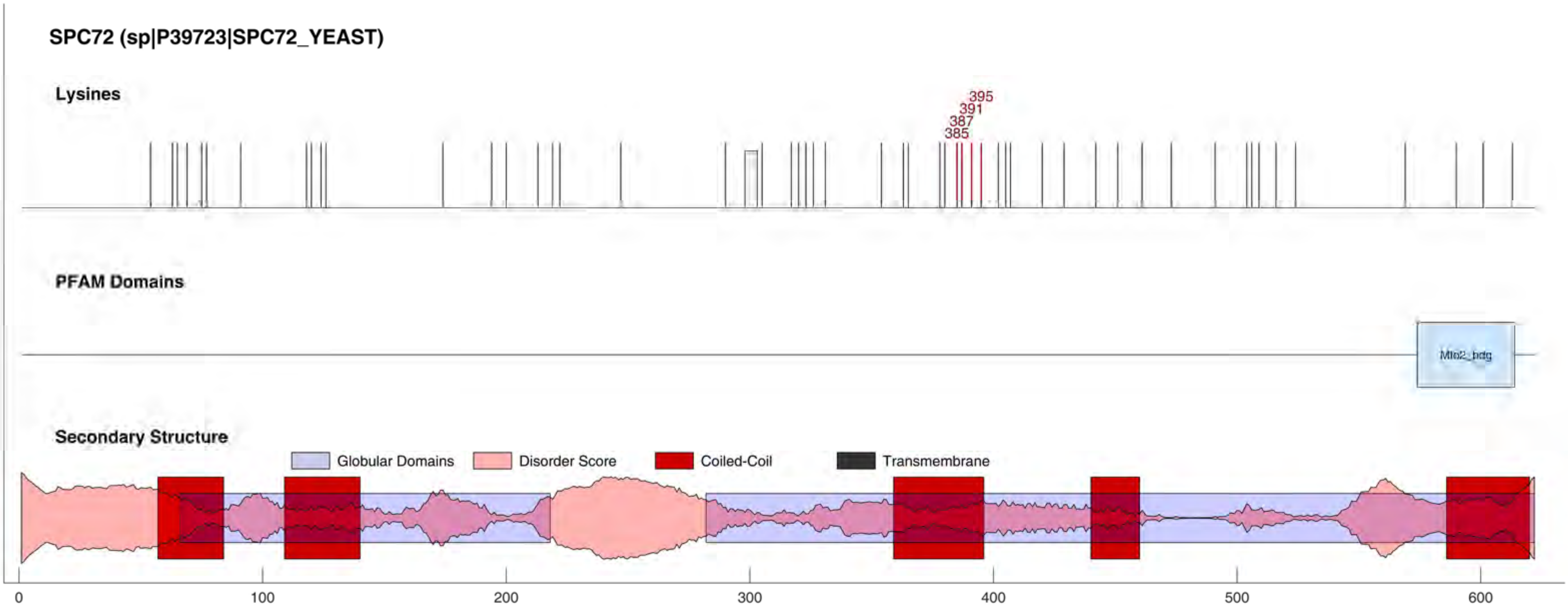
SPC72 (sp|P39723|SPC72_YEAST)
Lysines
395
391
387
385
PFAM Domains
Mlo2_bdg
Secondary Structure
Globular Domains
Disorder Score
Coiled-Coil
Transmembrane
0
100
200
300
400
500
600

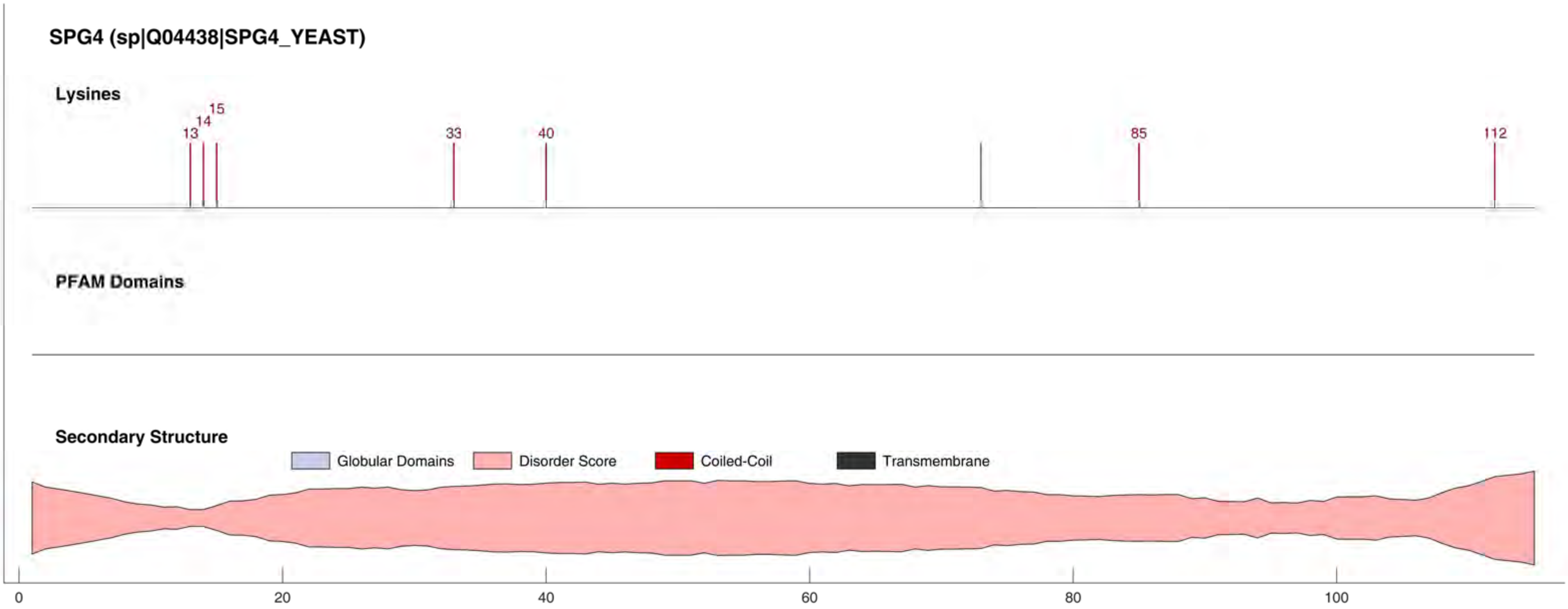
SPG4 (sp|Q04438|SPG4_YEAST)
Lysines
13
14
15
33
40
85
112
PFAM Domains
Secondary Structure
Globular Domains
Disorder Score
Coiled-Coil
Transmembrane
0
20
40
60
80
100

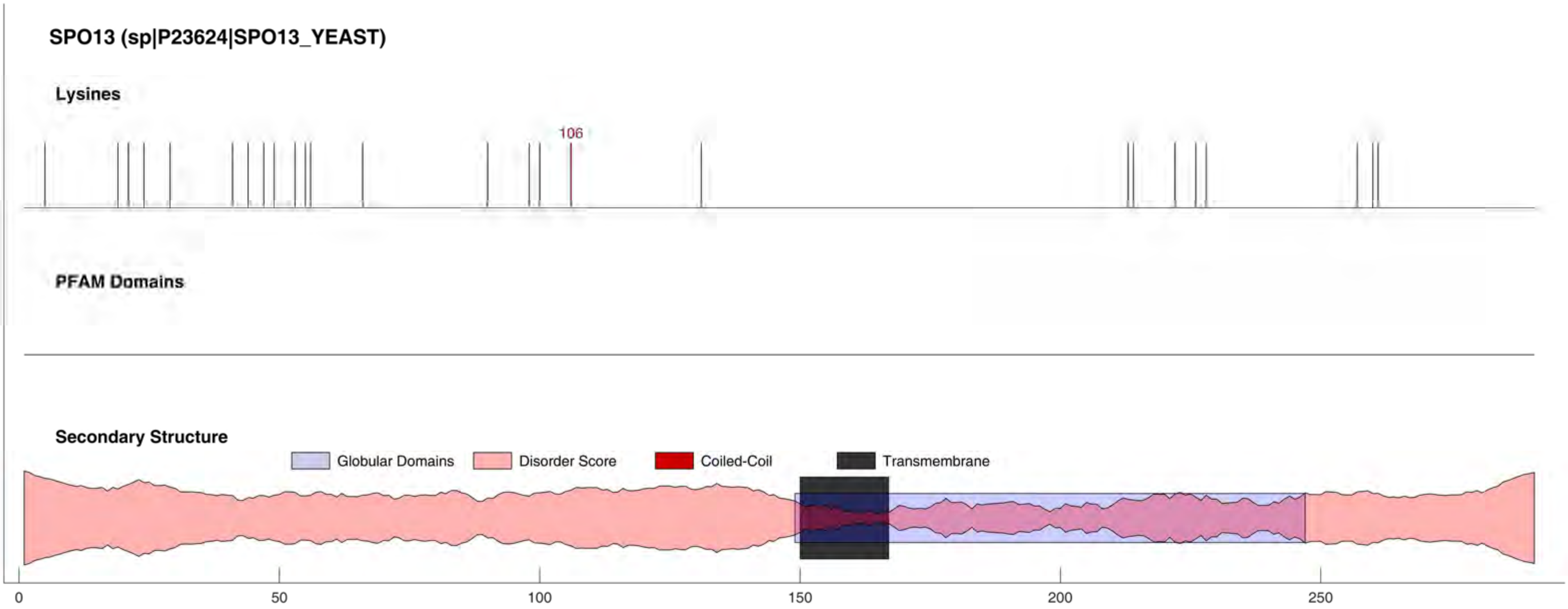

SPO13 (sp|P23624|SPO13_YEAST)
Lysines
106
PFAM Domains
Secondary Structure
Globular Domains
Disorder Score
Coiled-Coil
Transmembrane
0
50
100
150
200
250

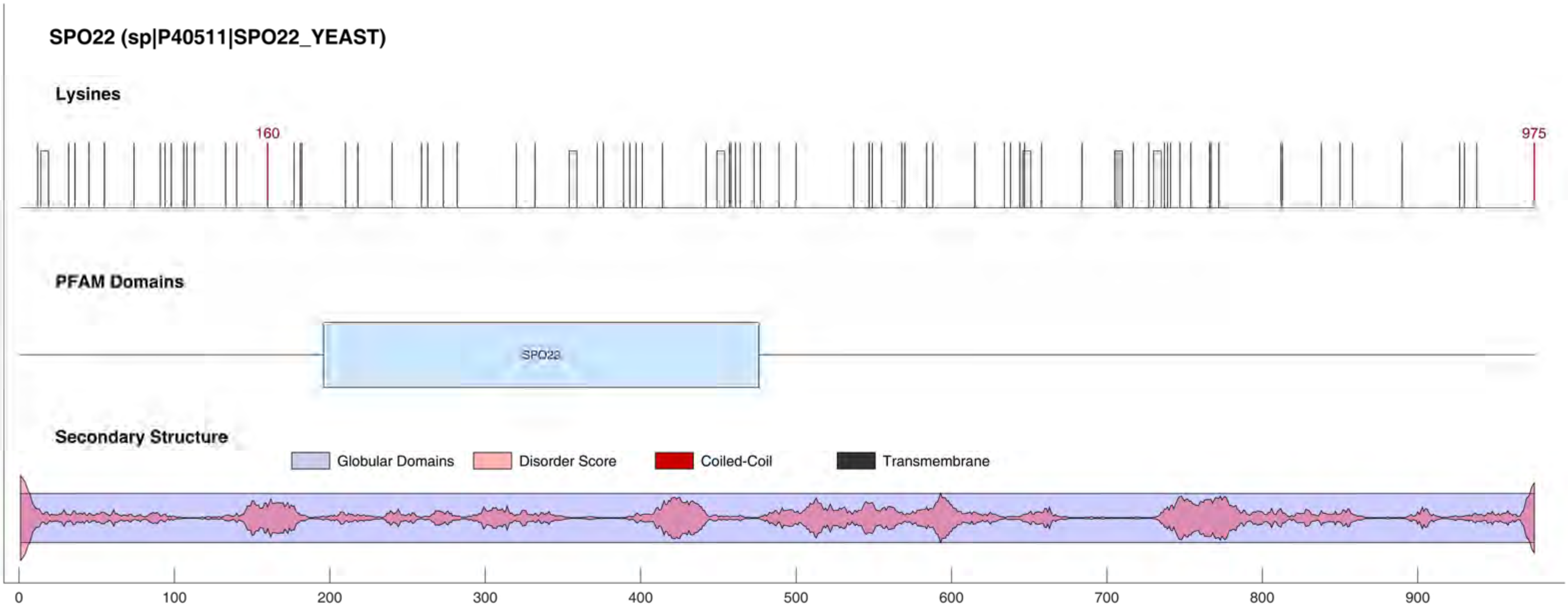
SPO22 (sp|P40511|SPO22_YEAST)
Lysines
160
975
PFAM Domains
SPO22
Secondary Structure
Globular Domains
Disorder Score
Coiled-Coil
Transmembrane
0
100
200
300
400
500
600
700
800
900

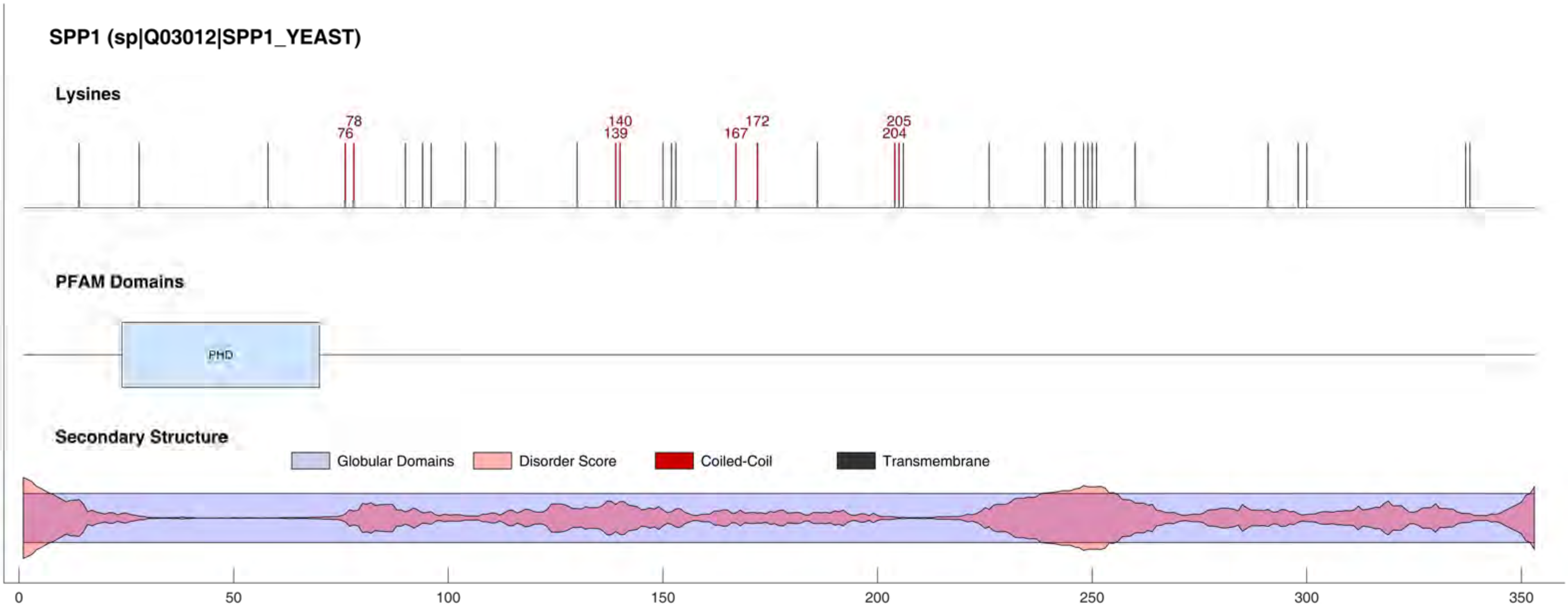
SPP1 (sp|Q03012|SPP1_YEAST)
Lysines
78
76
140
139
172
167
205
204
PFAM Domains
PHD
Secondary Structure
Globular Domains
Disorder Score
Coiled-Coil
Transmembrane
0
50
100
150
200
250
300
350

SPP41 (sp|P38904|SPP41_YEAST)

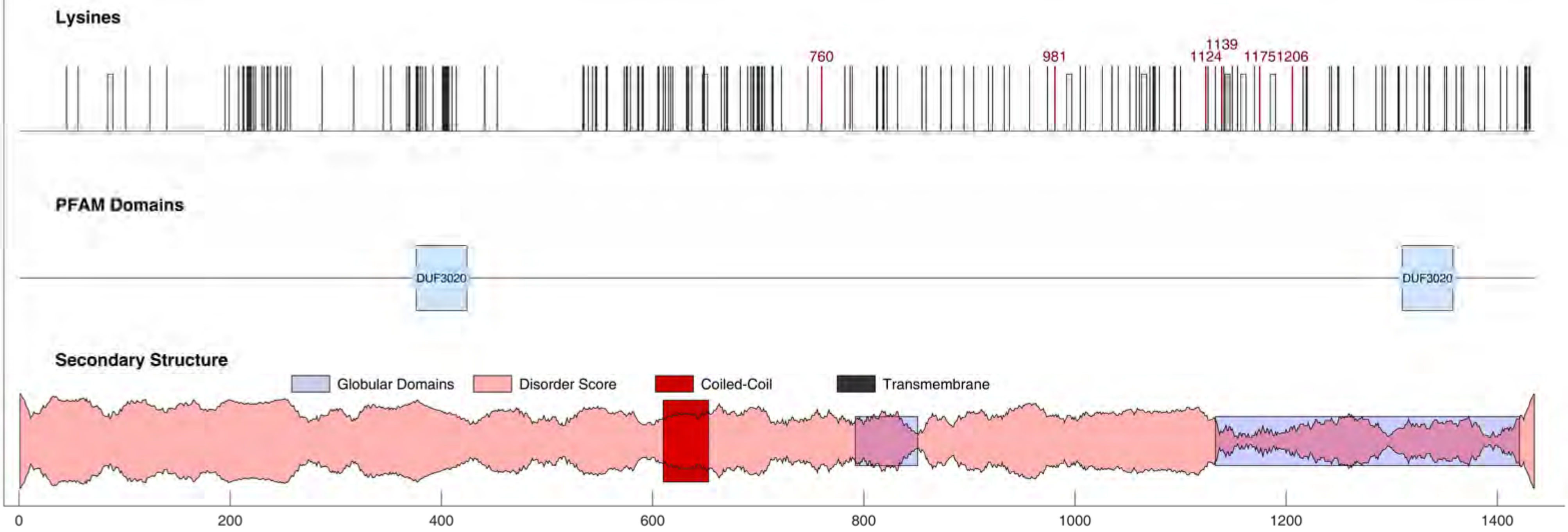

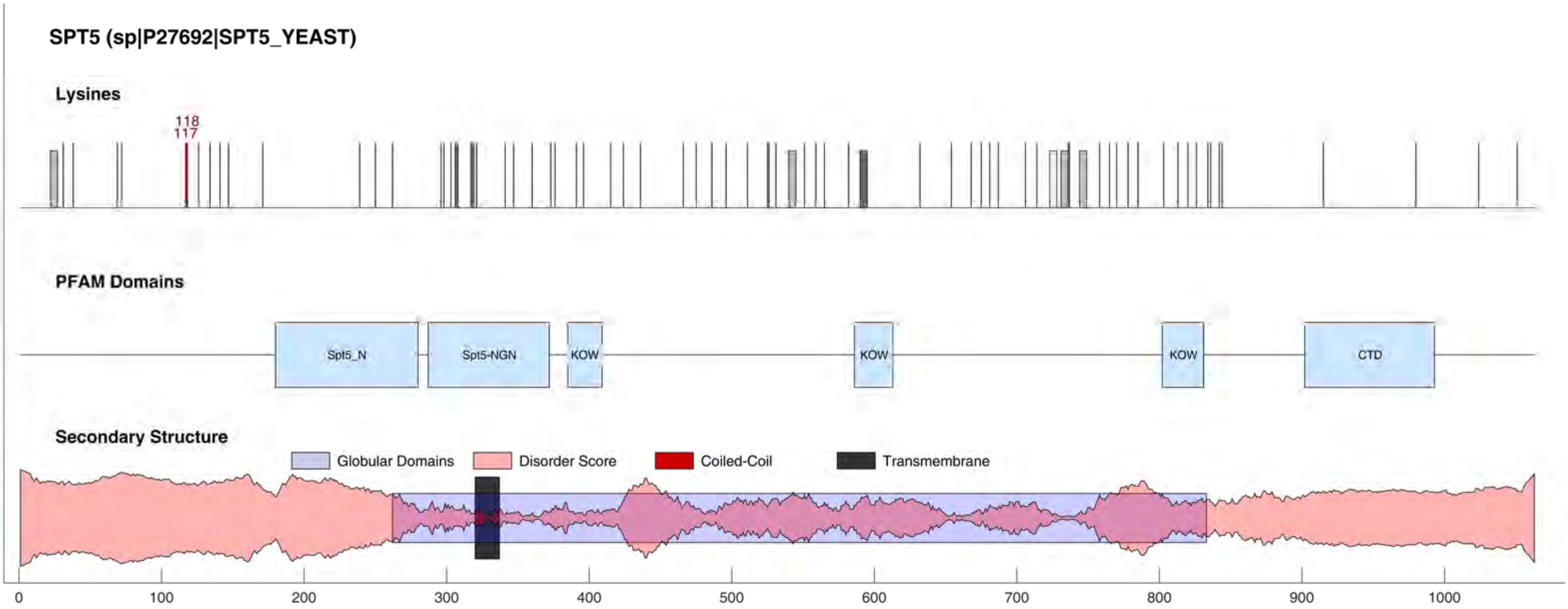
SPT5 (sp|P27692|SPT5_YEAST)
Lysines
118
117
PFAM Domains
Spt5_N
Spt5-NGN
KOW
KOW
KOW
CTD
Secondary Structure
Globular Domains
Disorder Score
Coiled-Coil
Transmembrane
0
100
200
300
400
500
600
700
800
900
1000

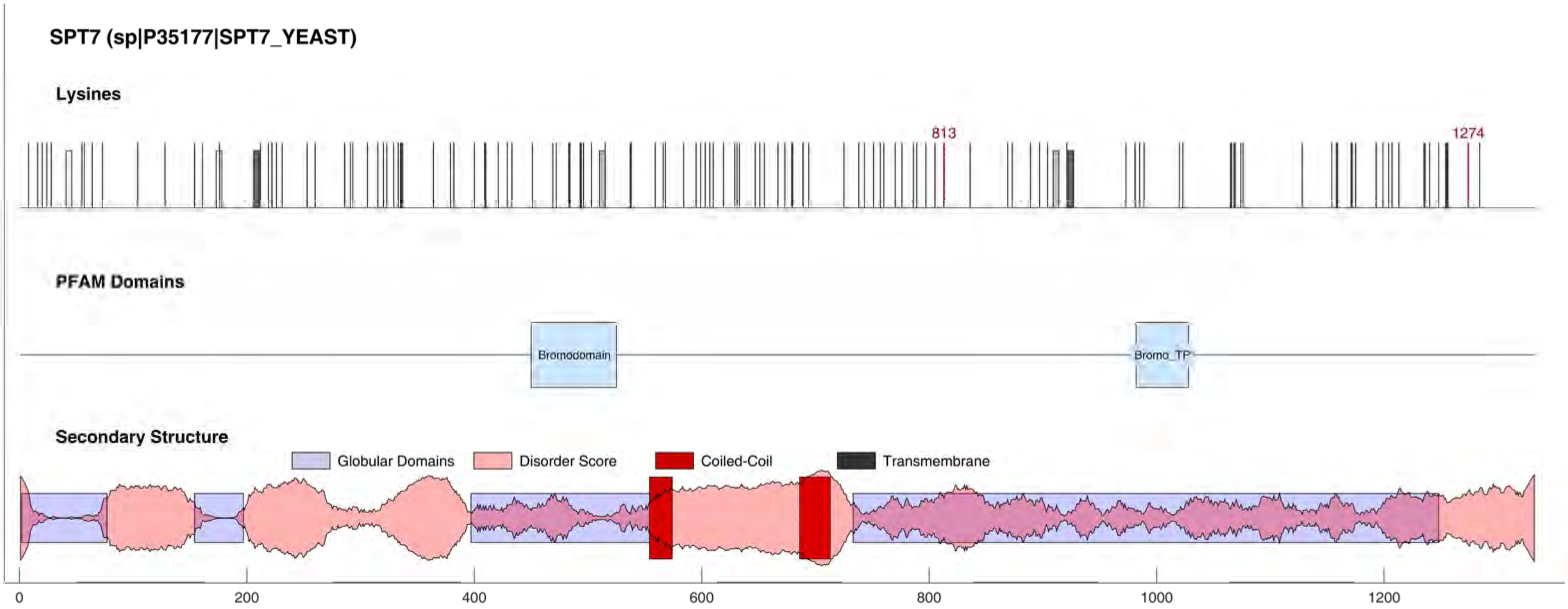

SPT7 (sp|P35177|SPT7_YEAST)
Lysines
813
1274
PFAM Domains
Bromodomain
Bromo_TP
Secondary Structure
Globular Domains
Disorder Score
Coiled-Coil
Transmembrane
0
200
400
600
800
1000
1200

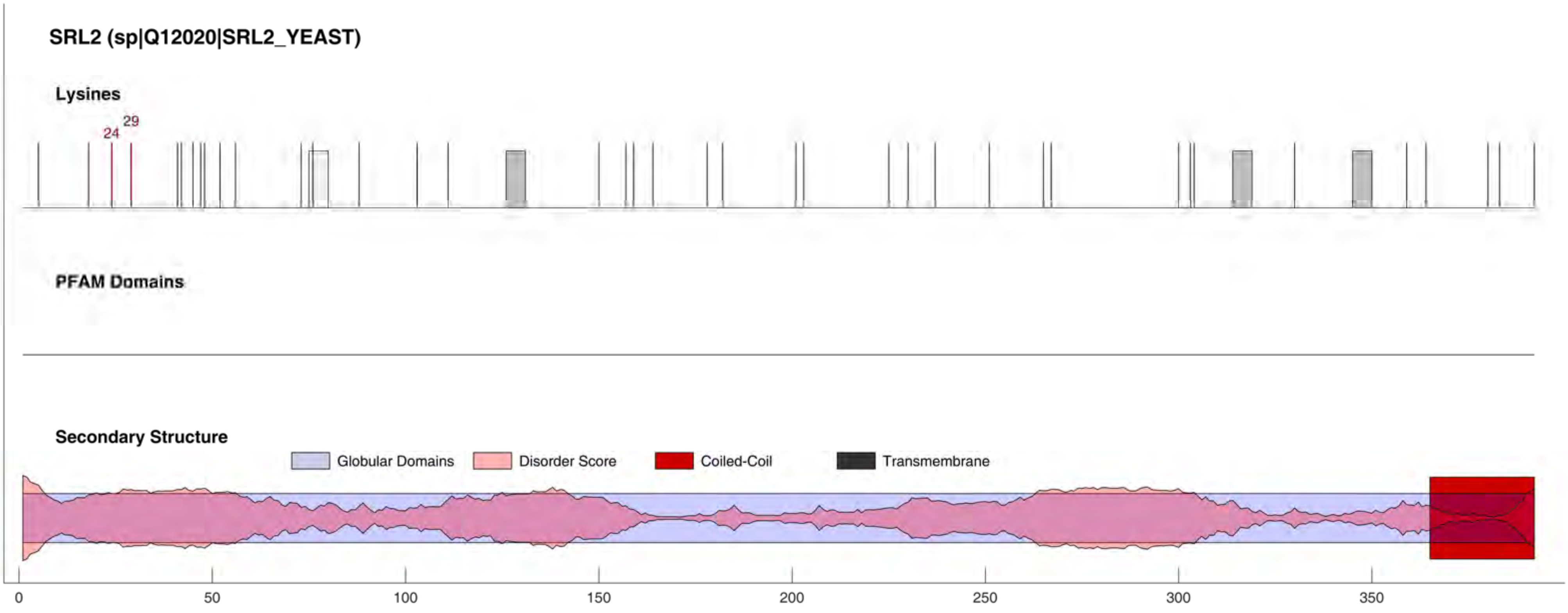
SRL2 (sp|Q12020|SRL2_YEAST)
Lysines
24
29
PFAM Domains
Secondary Structure
Globular Domains
Disorder Score
Coiled-Coil
Transmembrane
0
50
100
150
200
250
300
350

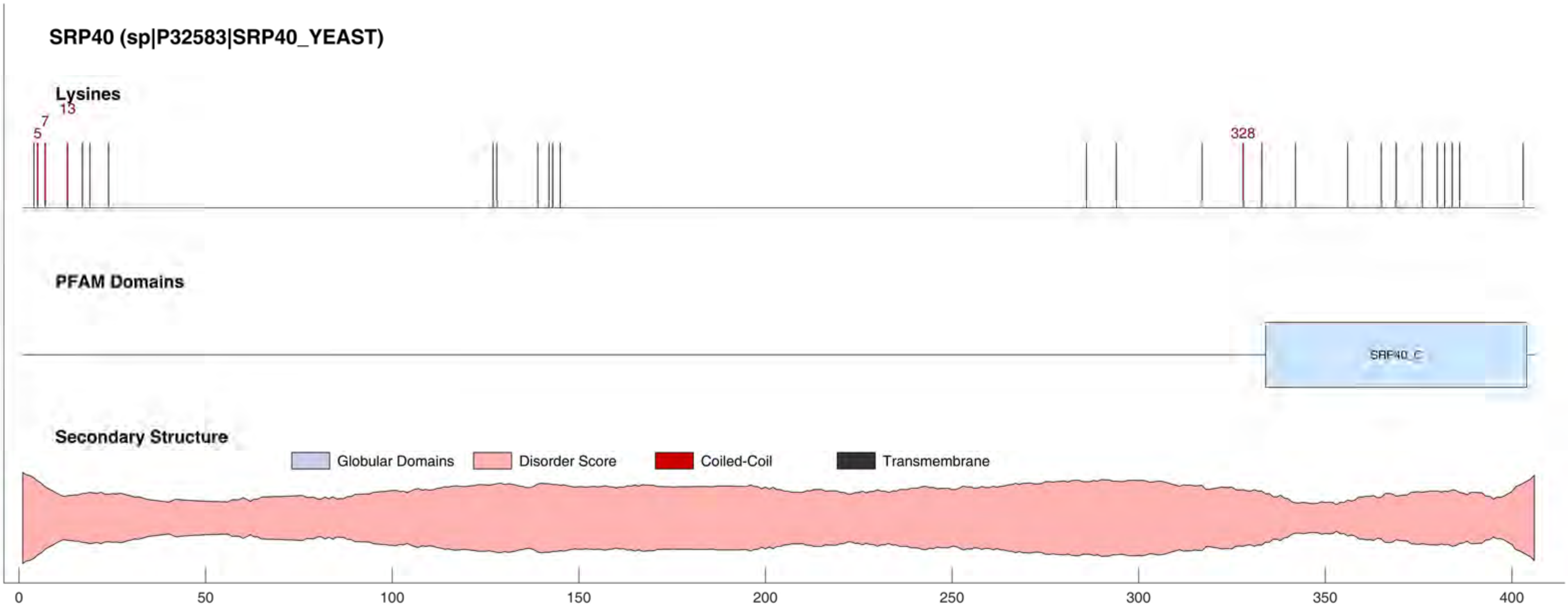

SRP40 (sp|P32583|SRP40_YEAST)
Lysines
5
7
13
328
PFAM Domains
SRP40_C
Secondary Structure
Globular Domains
Disorder Score
Coiled-Coil
Transmembrane
0
50
100
150
200
250
300
350
400

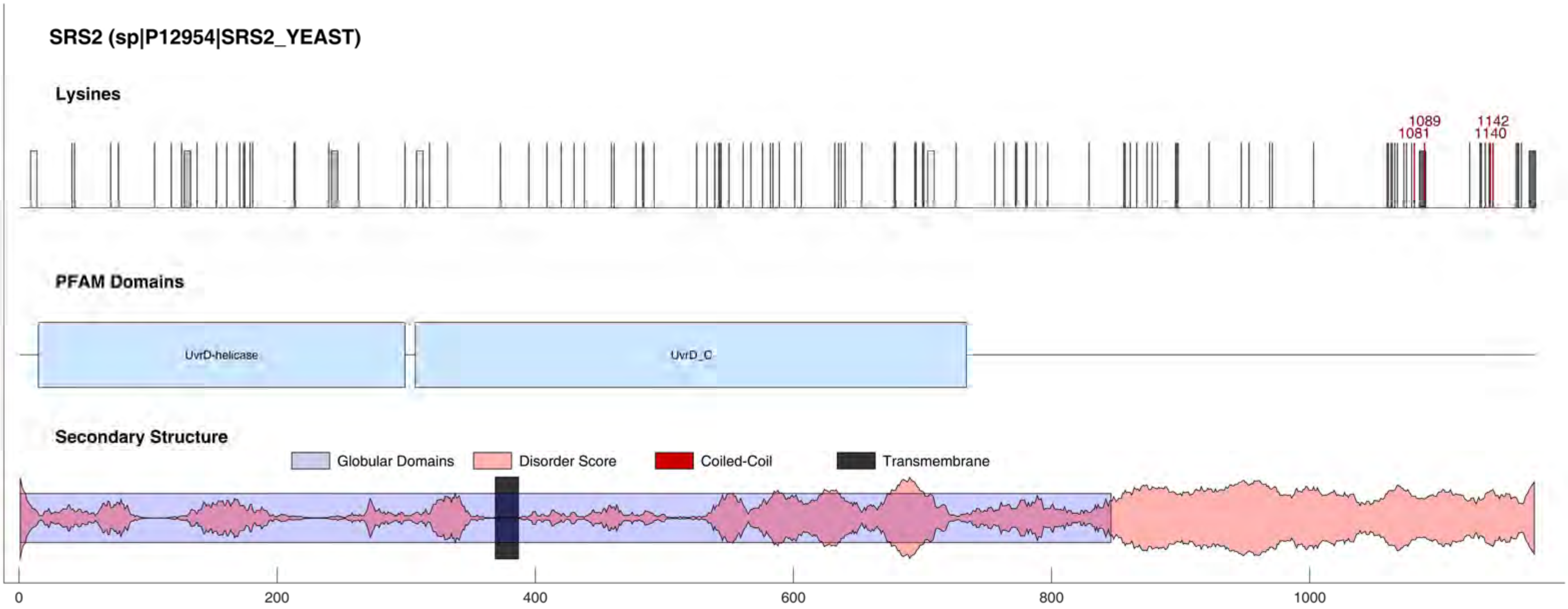
SRS2 (sp|P12954|SRS2_YEAST)
Lysines
1089
1081
1142
1140
PFAM Domains
UvrD-helicase
UvrD_C
Secondary Structure
Globular Domains
Disorder Score
Coiled-Coil
Transmembrane
0
200
400
600
800
1000

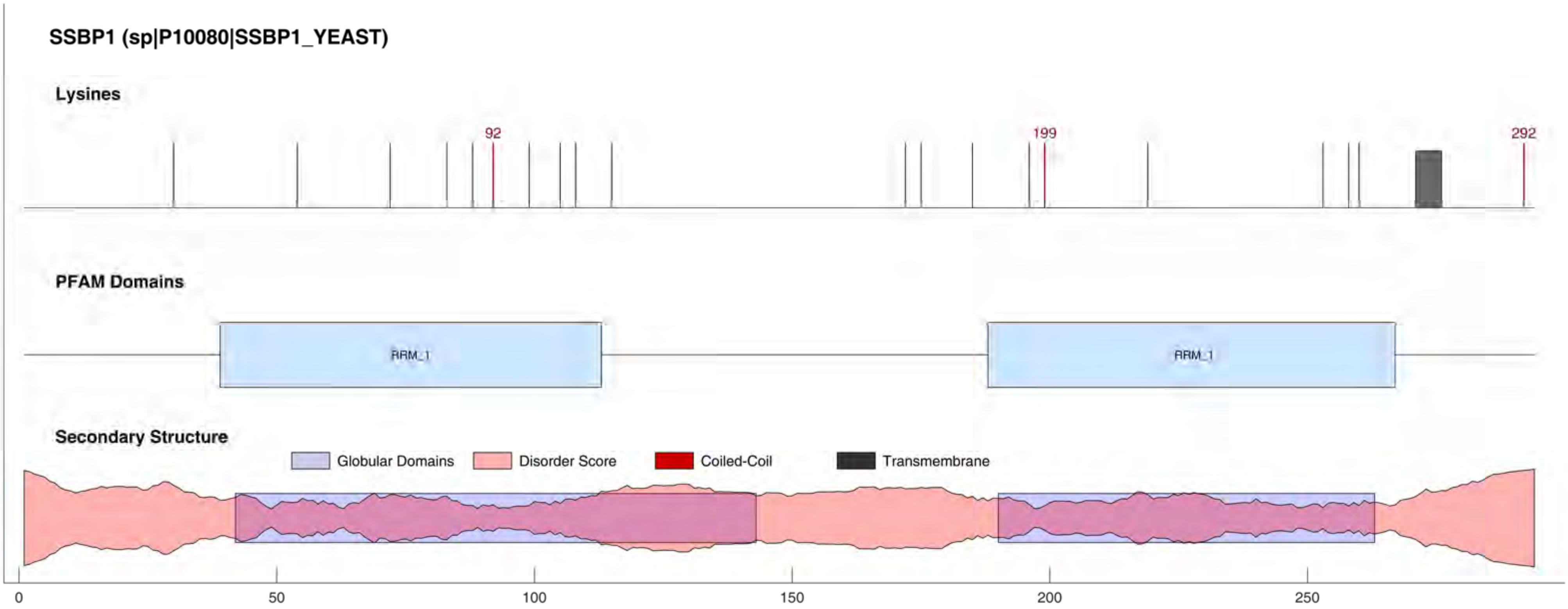

SSBP1 (sp|P10080|SSBP1_YEAST)
Lysines
92
199
292
PFAM Domains
RRM_1
RRM_1
Secondary Structure
Globular Domains
Disorder Score
Coiled-Coil
Transmembrane
0
50
100
150
200
250

#### SSP1 (sp|P38871|SSP1_YEAST)

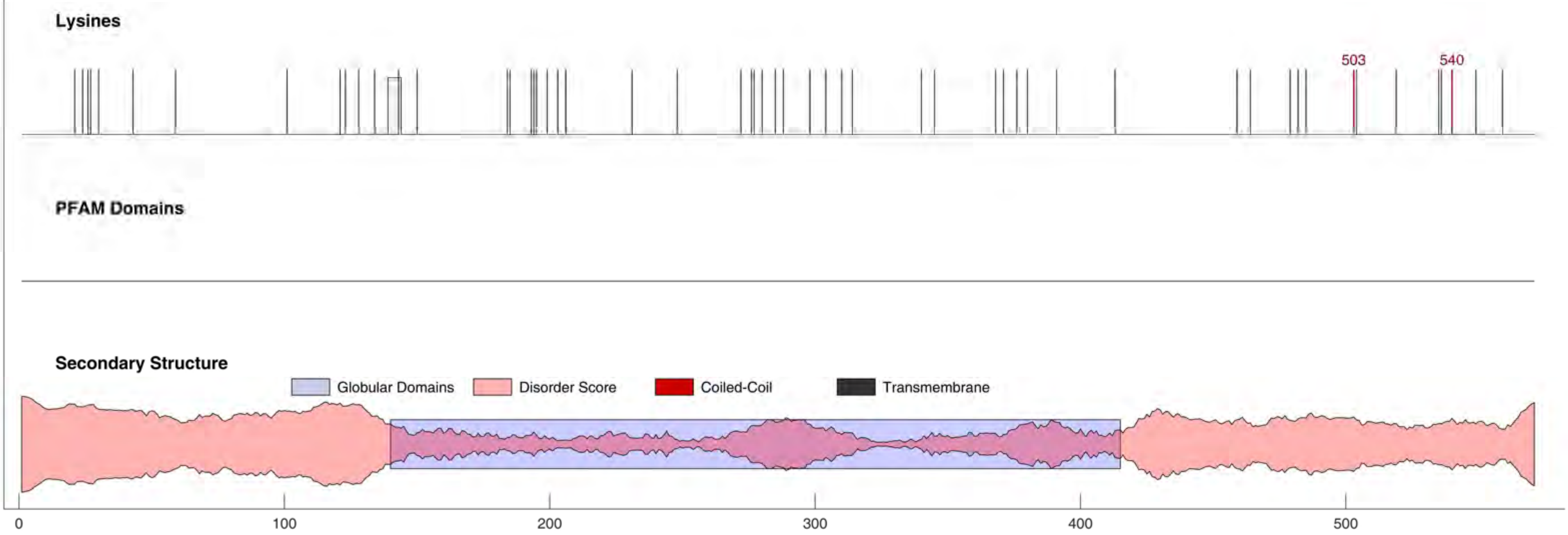

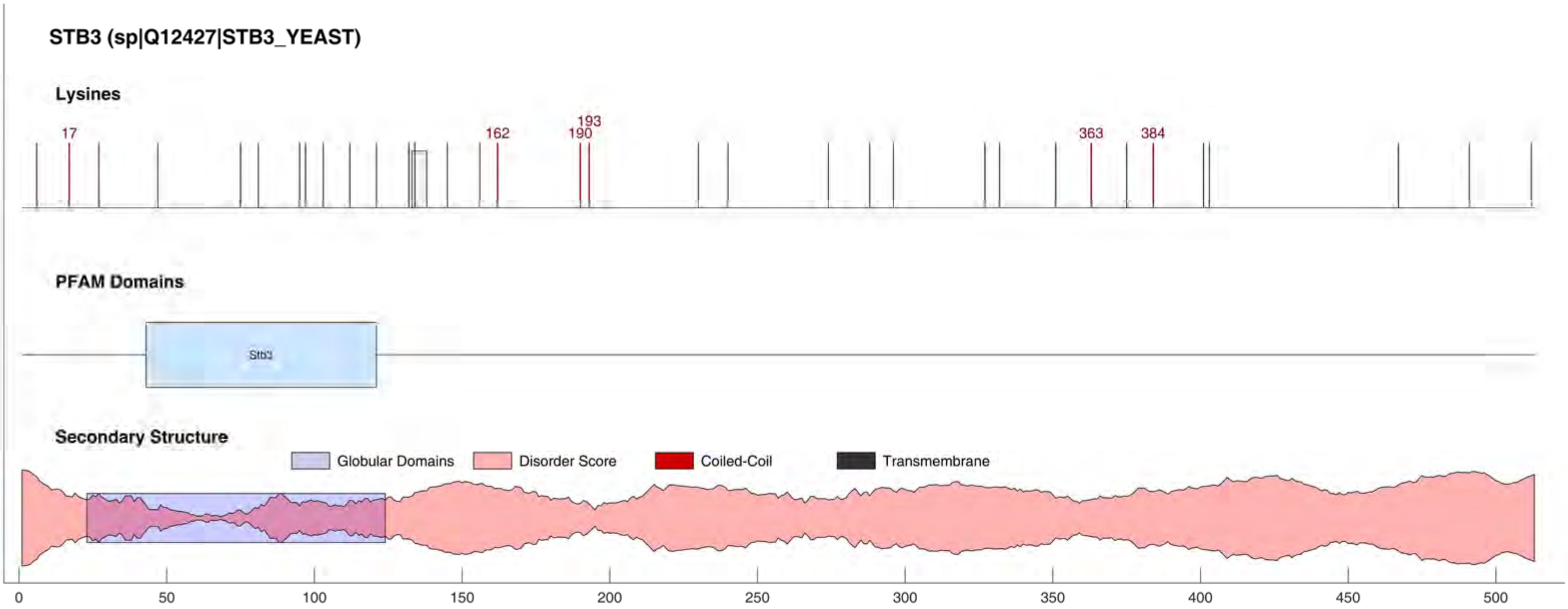
STB3 (sp|Q12427|STB3_YEAST)
Lysines
17
162
193
190
363
384
PFAM Domains
Stb3
Secondary Structure
Globular Domains
Disorder Score
Coiled-Coil
Transmembrane
0
50
100
150
200
250
300
350
400
450
500

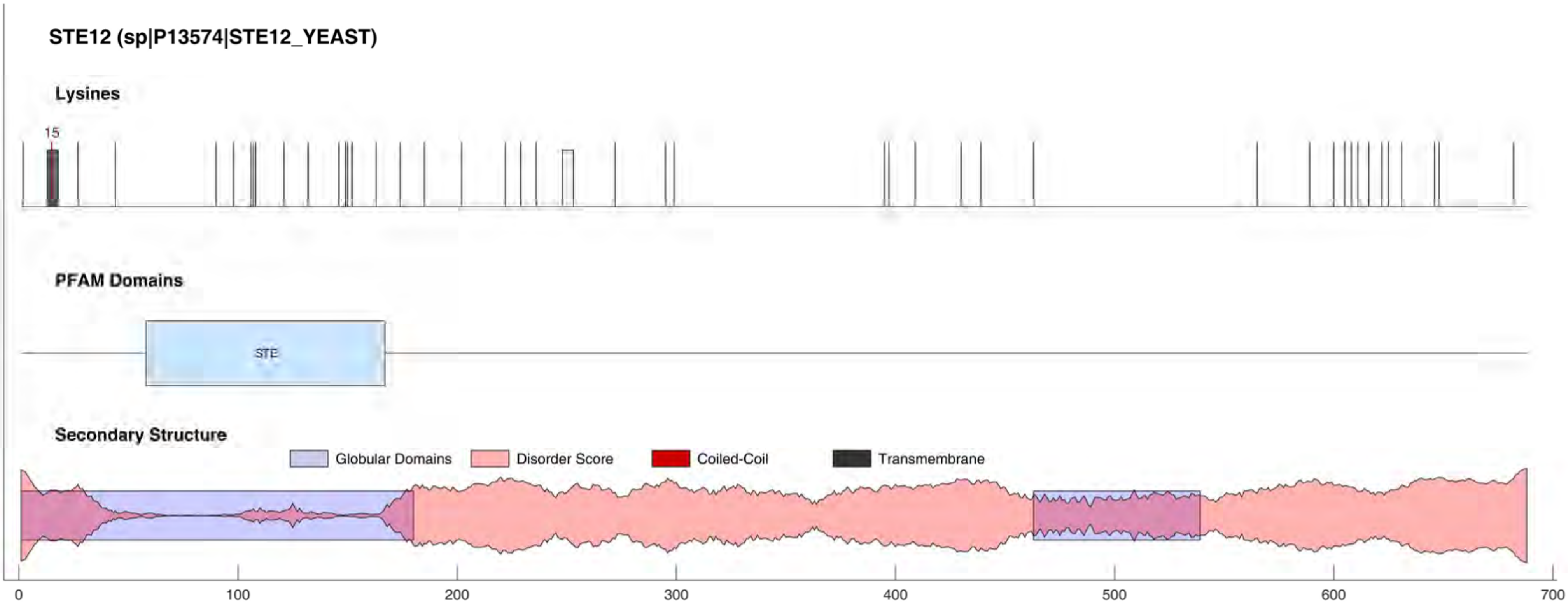

STE12 (sp|P13574|STE12_YEAST)
Lysines
15
PFAM Domains
STE
Secondary Structure
Globular Domains
Disorder Score
Coiled-Coil
Transmembrane
0
100
200
300
400
500
600
700

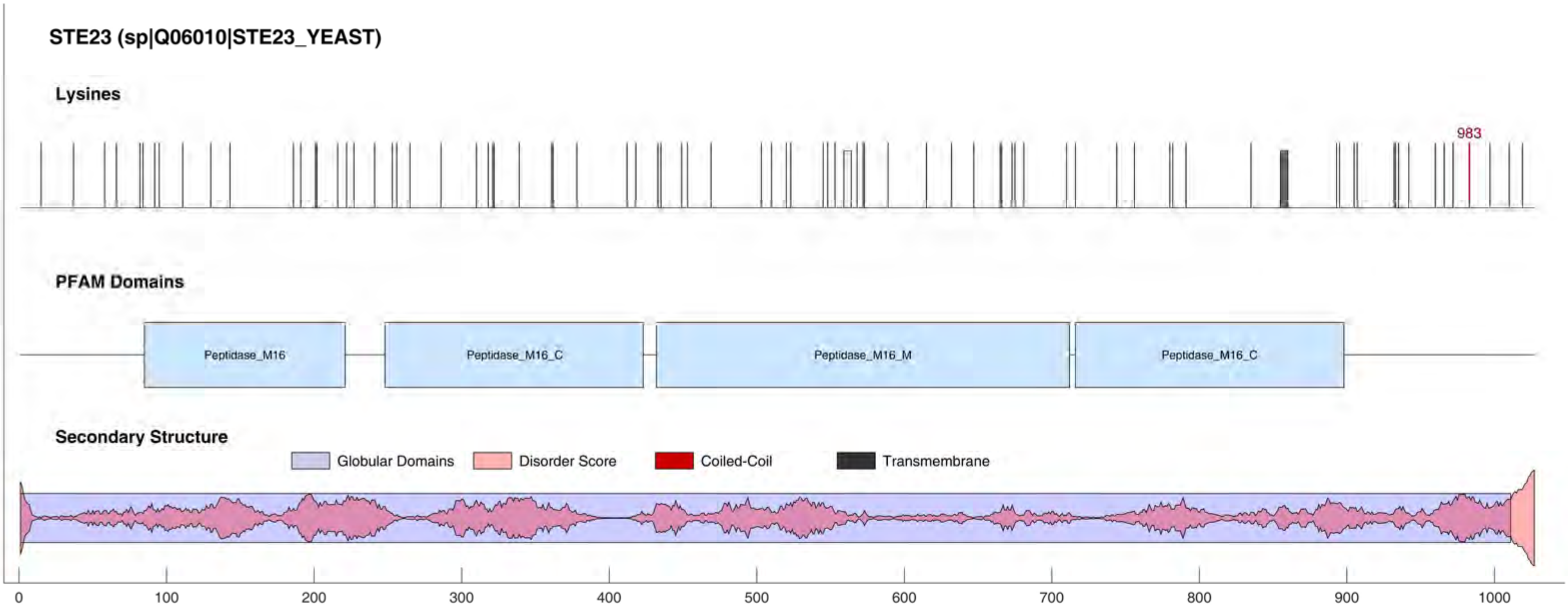
STE23 (sp|Q06010|STE23_YEAST)
Lysines
983
PFAM Domains
Peptidase_M16
Peptidase_M16_C
Peptidase_M16_M
Peptidase_M16_C
Secondary Structure
Globular Domains
Disorder Score
Coiled-Coil
Transmembrane
0
100
200
300
400
500
600
700
800
900
1000

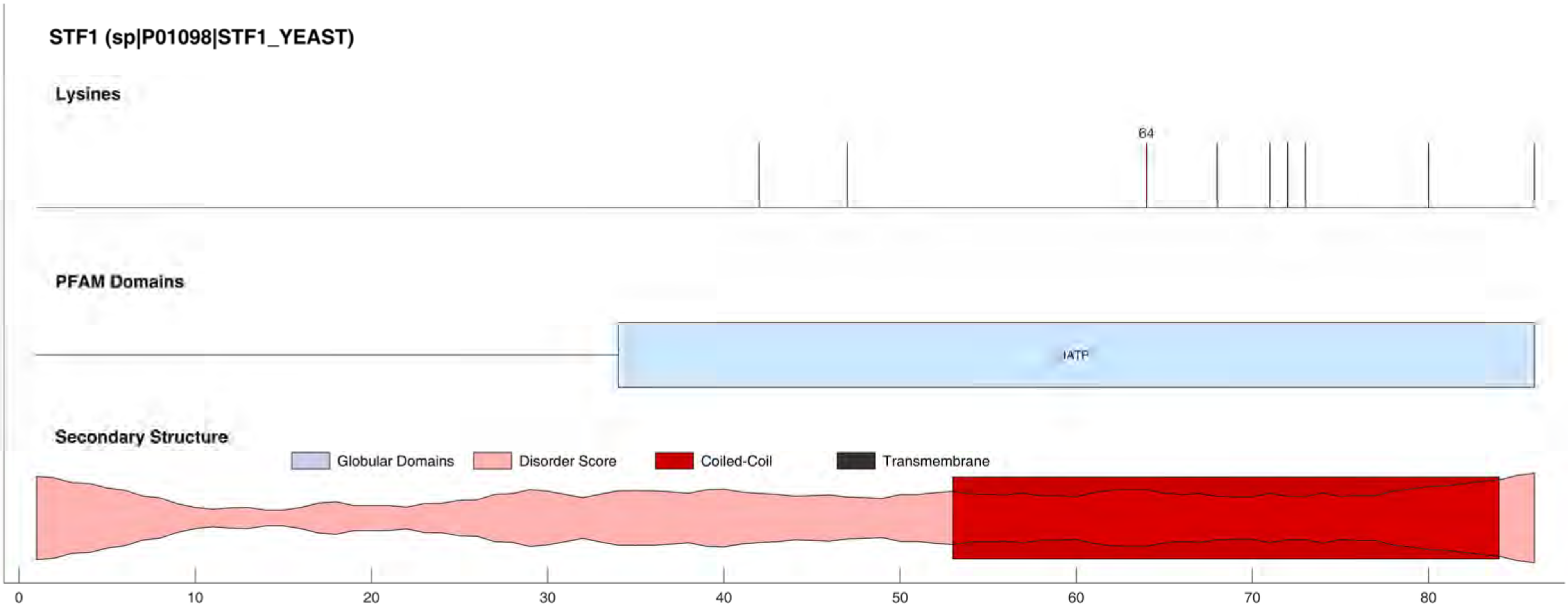
STF1 (sp|P01098|STF1_YEAST)
Lysines
64
PFAM Domains
IATP
Secondary Structure
Globular Domains
Disorder Score
Coiled-Coil
Transmembrane
0
10
20
30
40
50
60
70
80

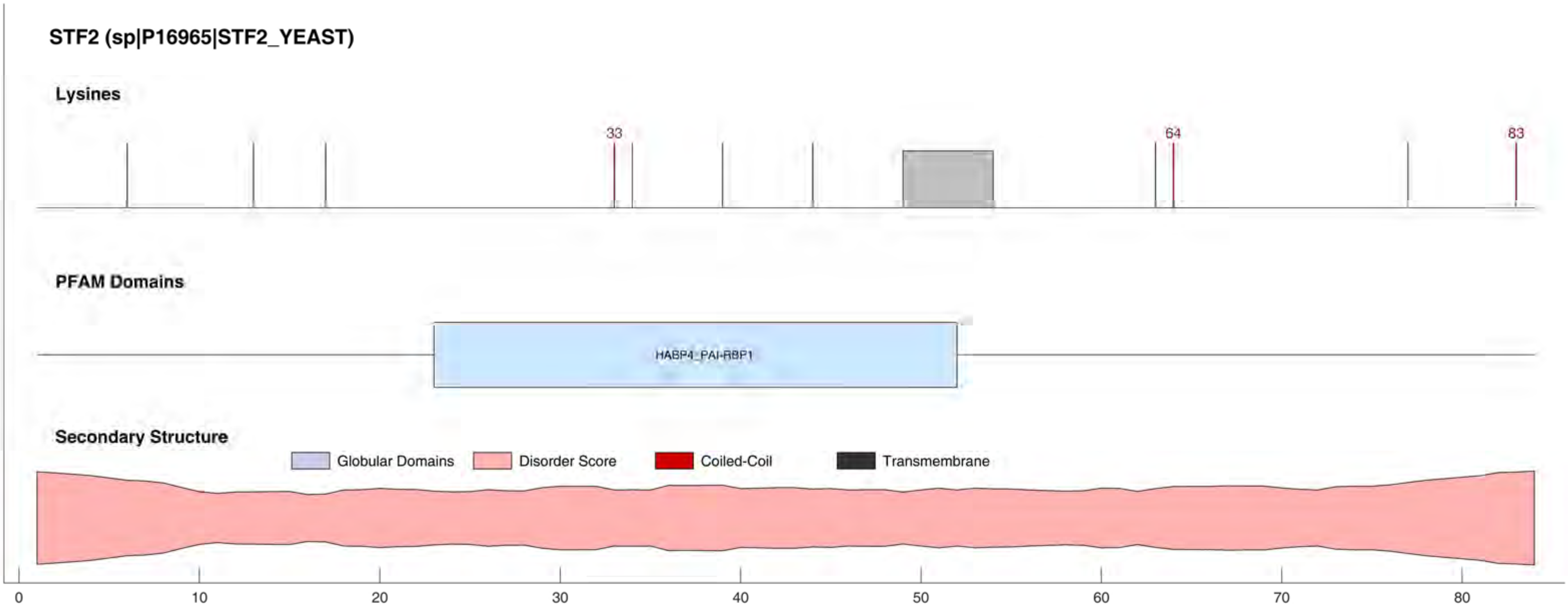
STF2 (sp|P16965|STF2_YEAST)
Lysines
33
64
83
PFAM Domains
HABP4_PAI-RBP1
Secondary Structure
Globular Domains
Disorder Score
Coiled-Coil
Transmembrane
0
10
20
30
40
50
60
70
80

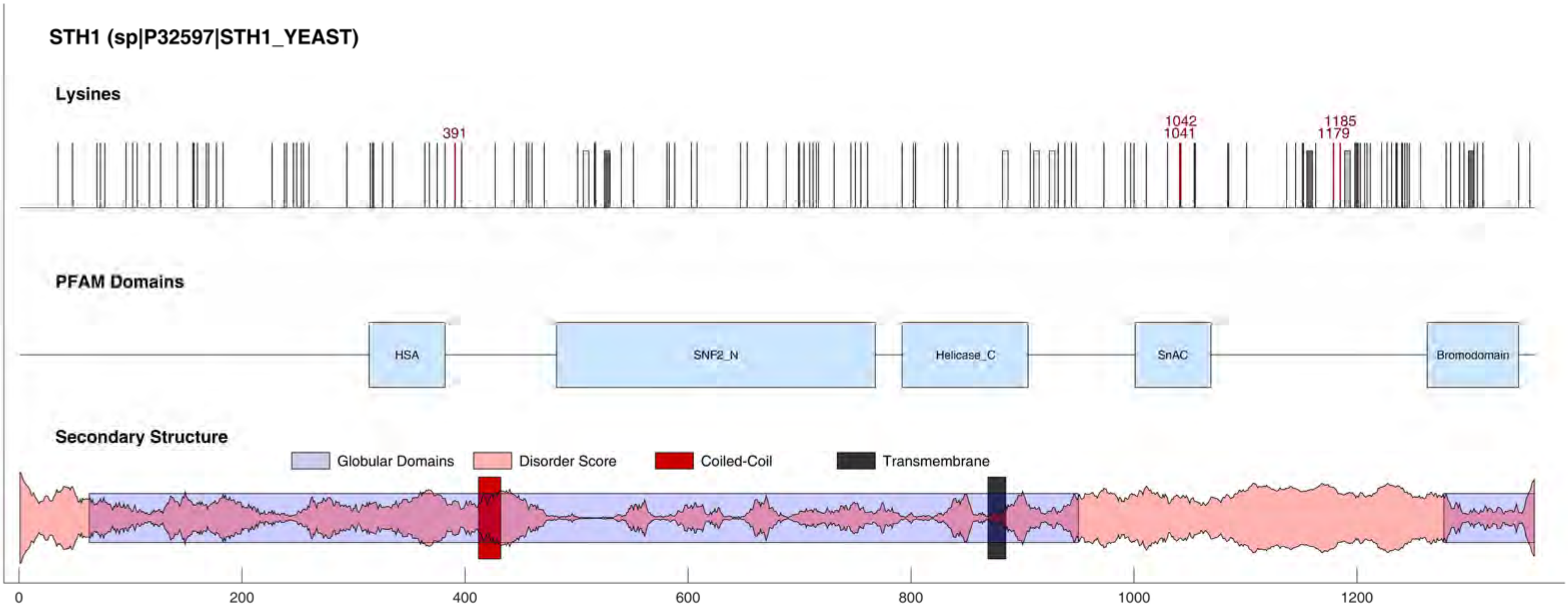

STH1 (sp|P32597|STH1_YEAST)
Lysines
391
1042
1041
1185
1179
PFAM Domains
HSA
SNF2_N
Helicase_C
SnAC
Bromodomain
Secondary Structure
Globular Domains
Disorder Score
Coiled-Coil
Transmembrane
0
200
400
600
800
1000
1200

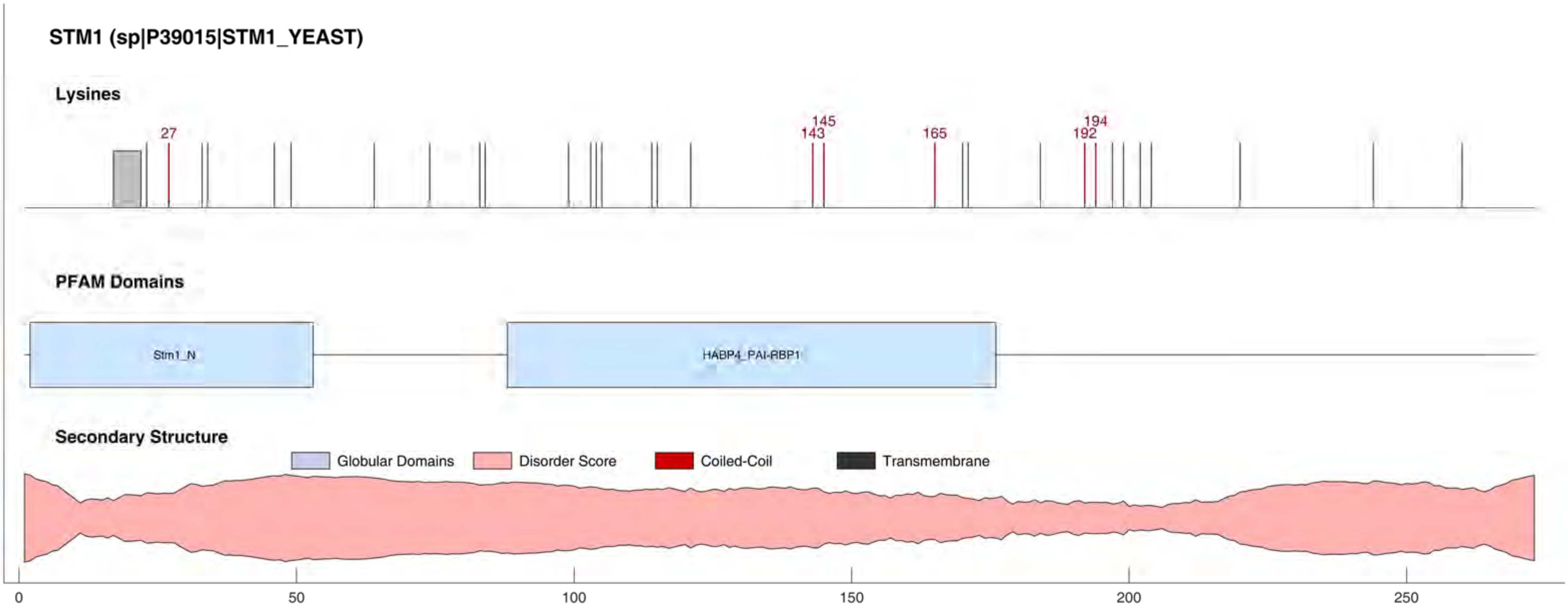

STM1 (sp|P39015|STM1_YEAST)
Lysines
27
143
145
165
192
194
PFAM Domains
Stm1_N
HABP4_PAI-RBP1
Secondary Structure
Globular Domains
Disorder Score
Coiled-Coil
Transmembrane
0
50
100
150
200
250

#### STP3 (sp|Q05937|STP3_YEAST)

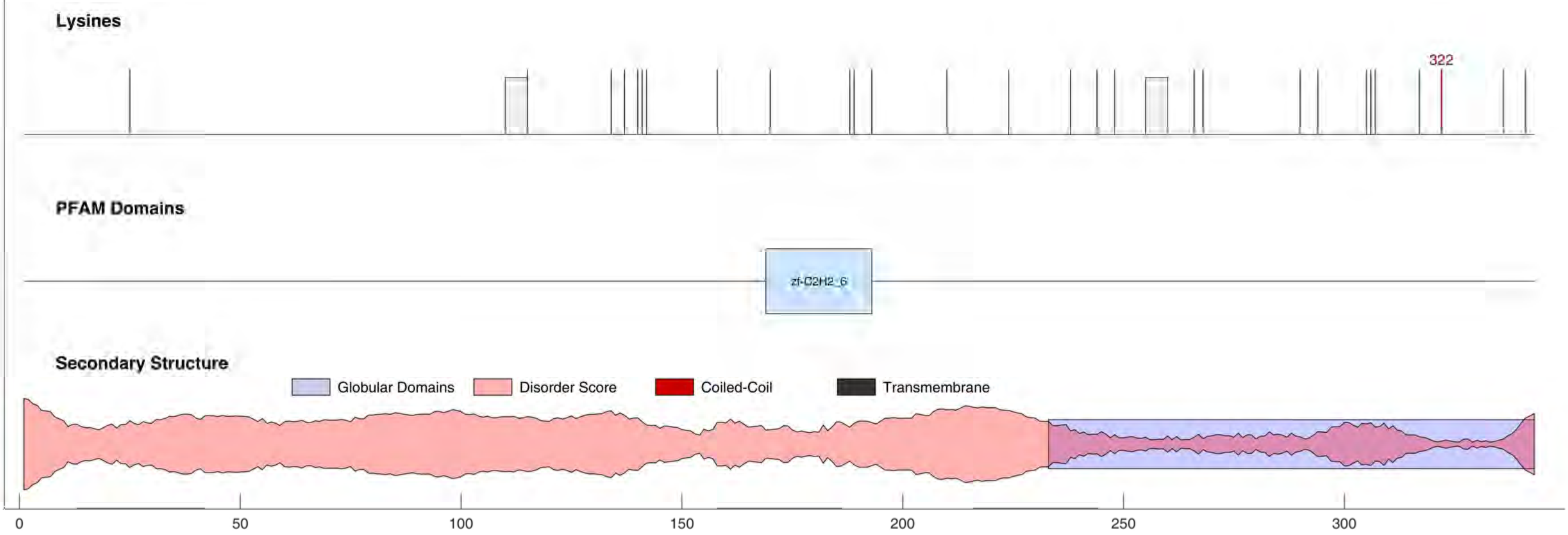

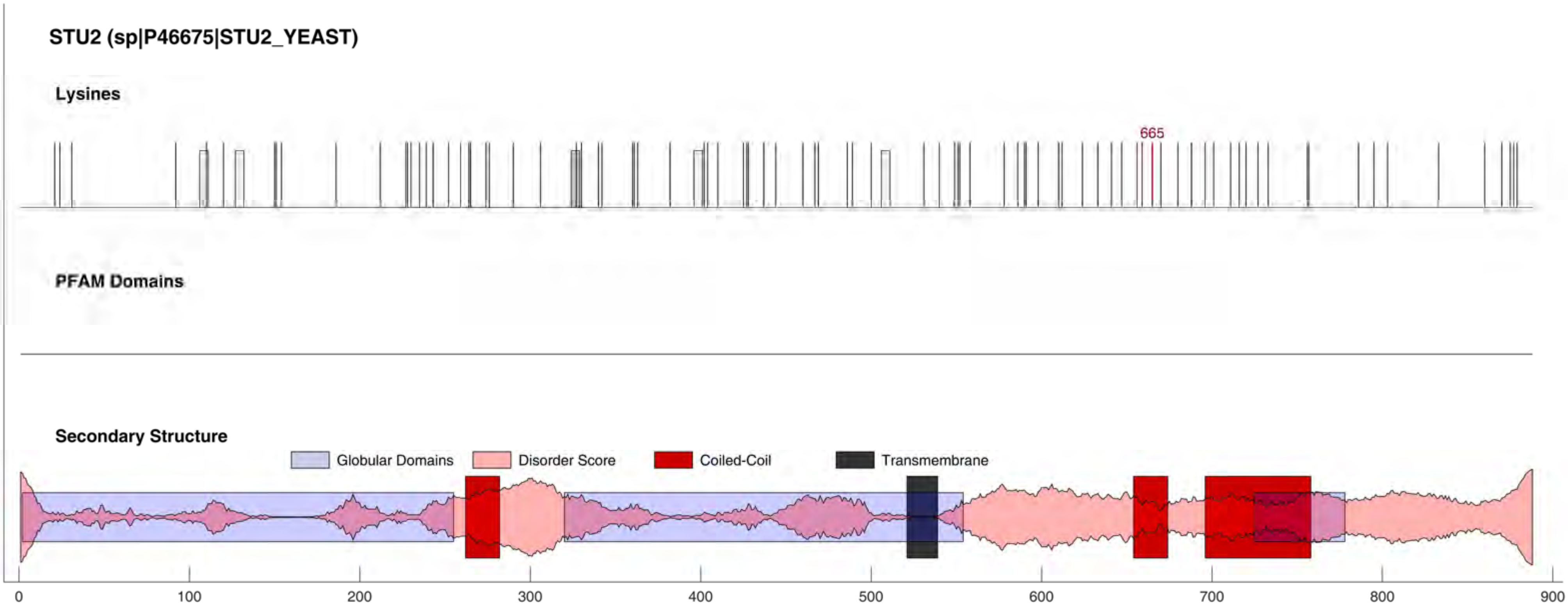
STU2 (sp|P46675|STU2_YEAST)
Lysines
665
PFAM Domains
Secondary Structure
Globular Domains
Disorder Score
Coiled-Coil
Transmembrane
0
100
200
300
400
500
600
700
800
900

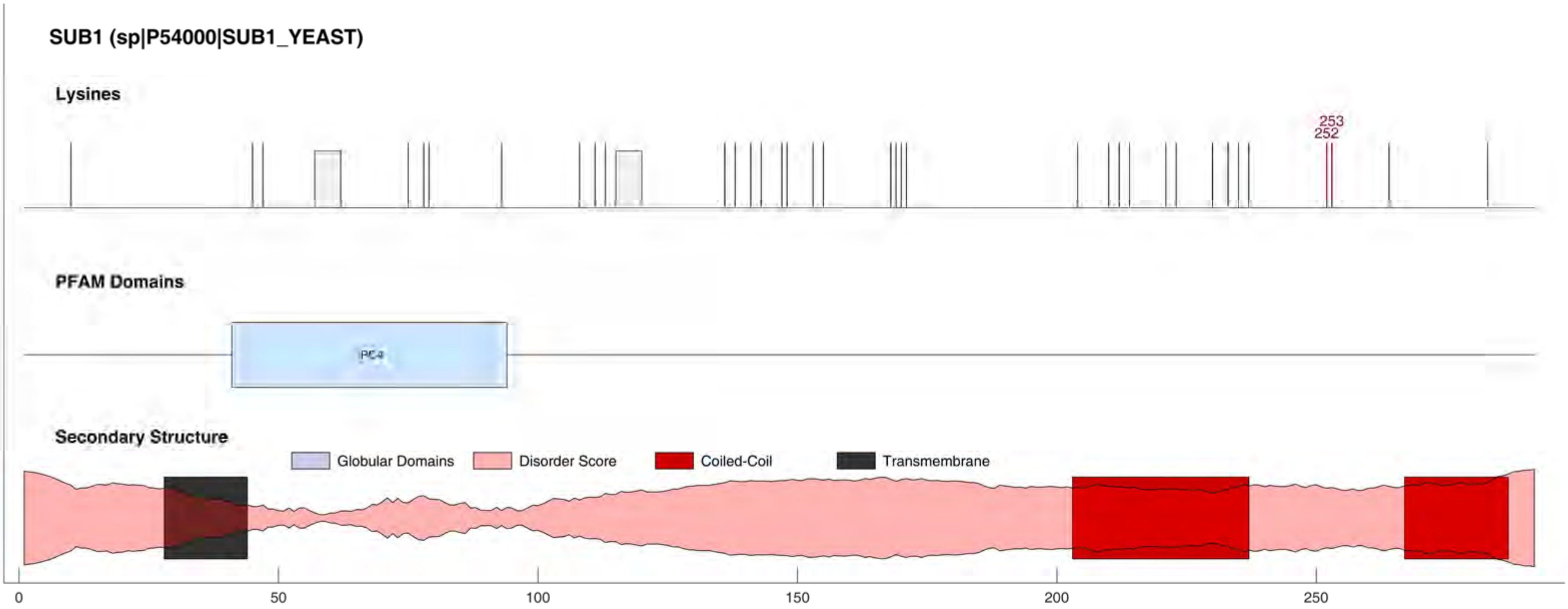
SUB1 (sp|P54000|SUB1_YEAST)
Lysines
253
252
PFAM Domains
PC4
Secondary Structure
Globular Domains
Disorder Score
Coiled-Coil
Transmembrane
0
50
100
150
200
250

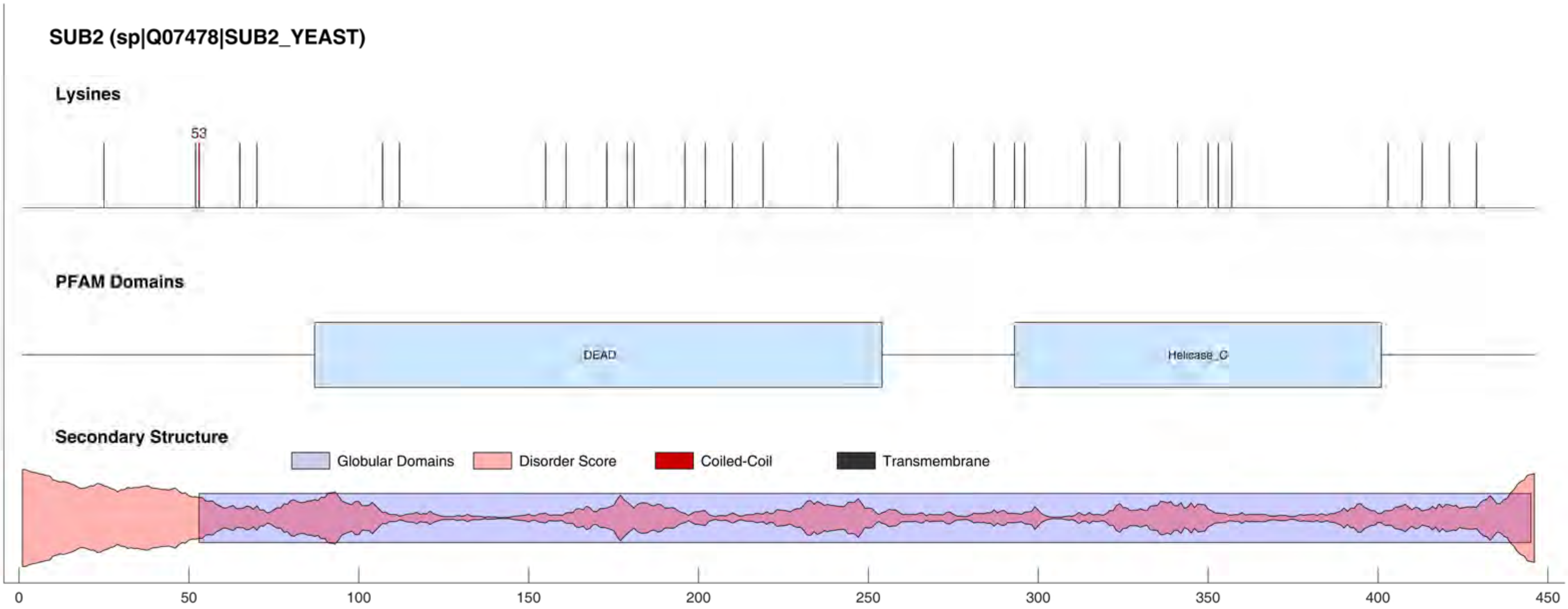
SUB2 (sp|Q07478|SUB2_YEAST)
Lysines
53
PFAM Domains
DEAD
Helicase_C
Secondary Structure
Globular Domains
Disorder Score
Coiled-Coil
Transmembrane
0
50
100
150
200
250
300
350
400
450

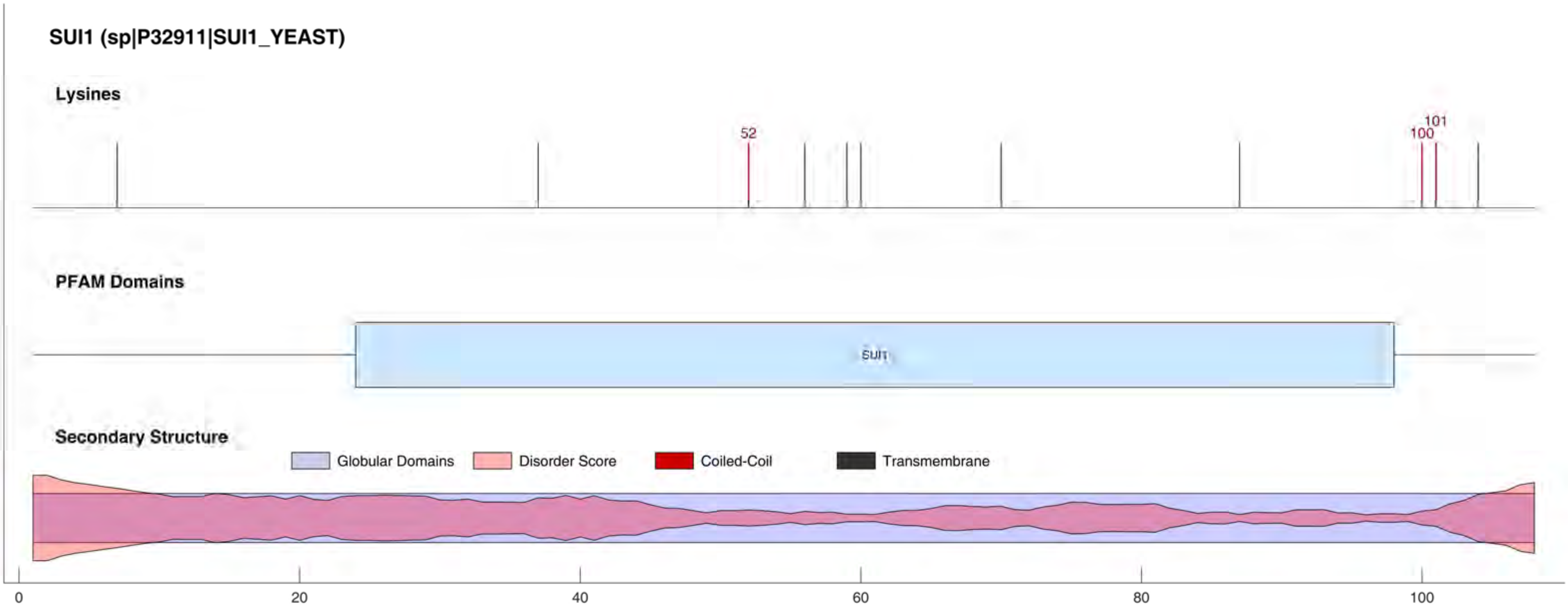
SUI1 (sp|P32911|SUI1_YEAST)
Lysines
52
100
101
PFAM Domains
SUI1
Secondary Structure
Globular Domains
Disorder Score
Coiled-Coil
Transmembrane
0
20
40
60
80
100

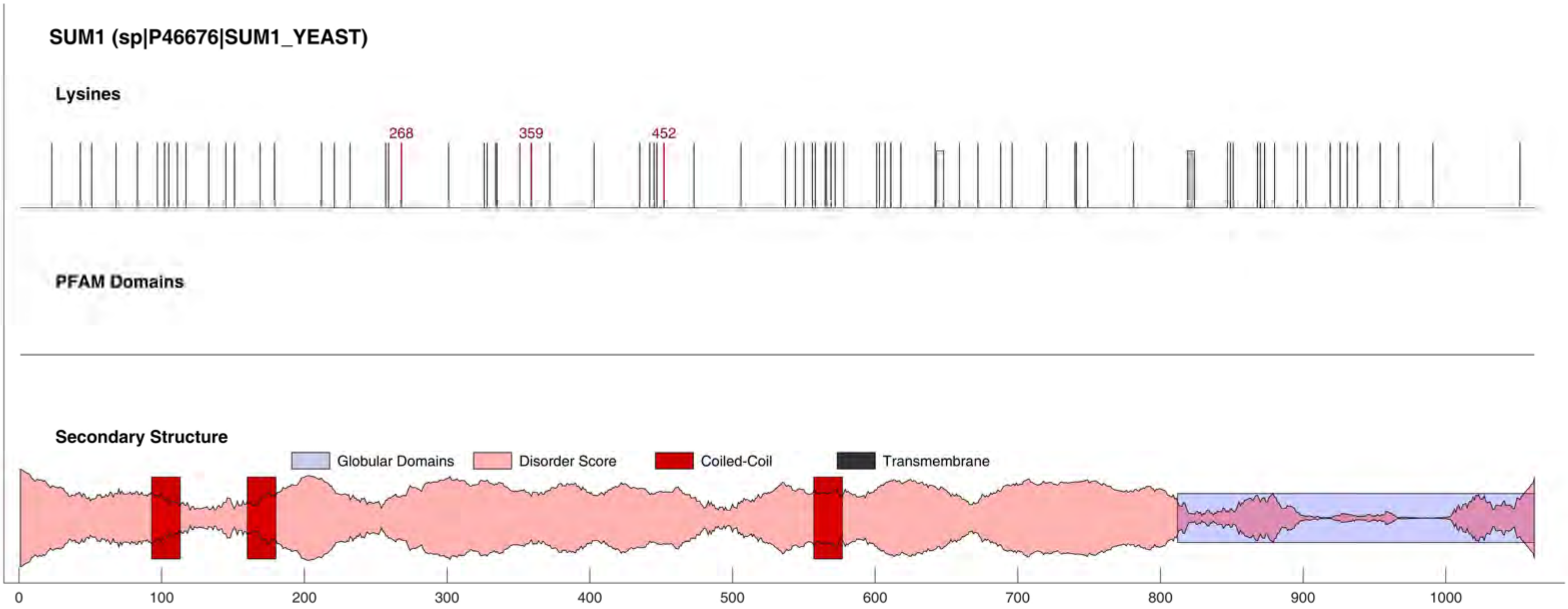

SUM1 (sp|P46676|SUM1_YEAST)
Lysines
268
359
452
PFAM Domains
Secondary Structure
Globular Domains
Disorder Score
Coiled-Coil
Transmembrane
0
100
200
300
400
500
600
700
800
900
1000

#### SWC3 (sp|P31376|SWC3_YEAST)

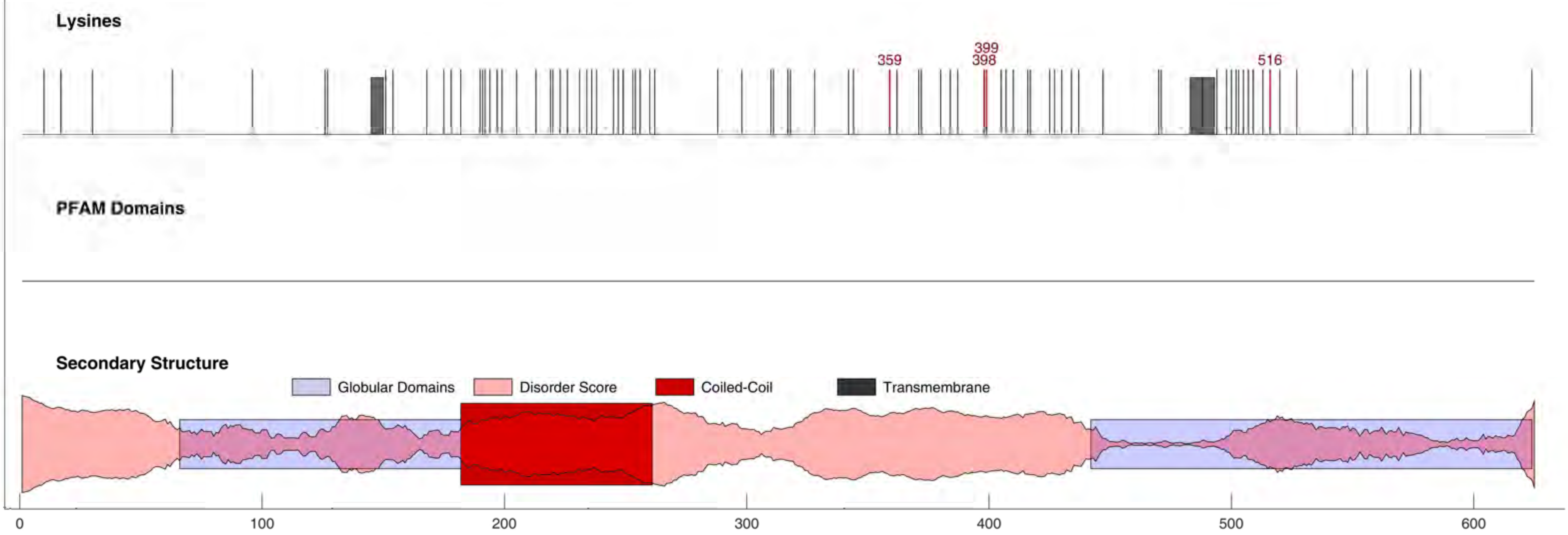

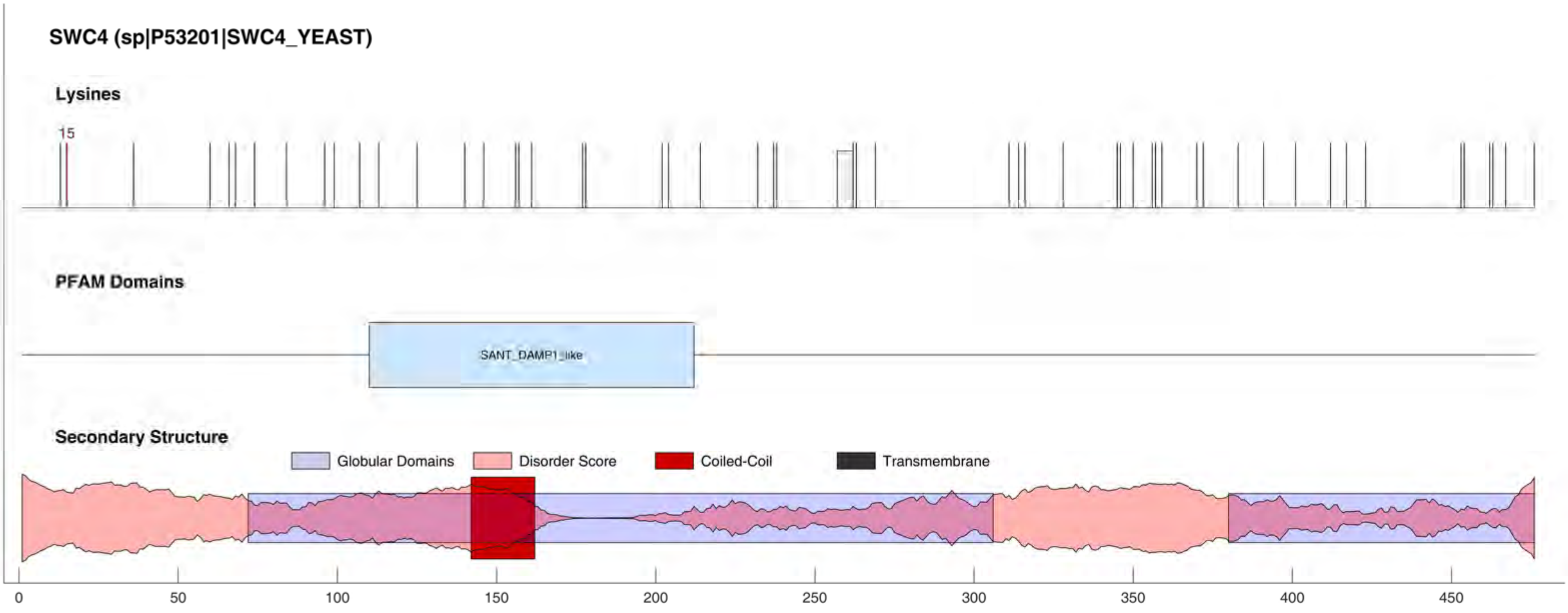
SWC4 (sp|P53201|SWC4_YEAST)
Lysines
15
PFAM Domains
SANT_DAMP1_like
Secondary Structure
Globular Domains
Disorder Score
Coiled-Coil
Transmembrane
0
50
100
150
200
250
300
350
400
450

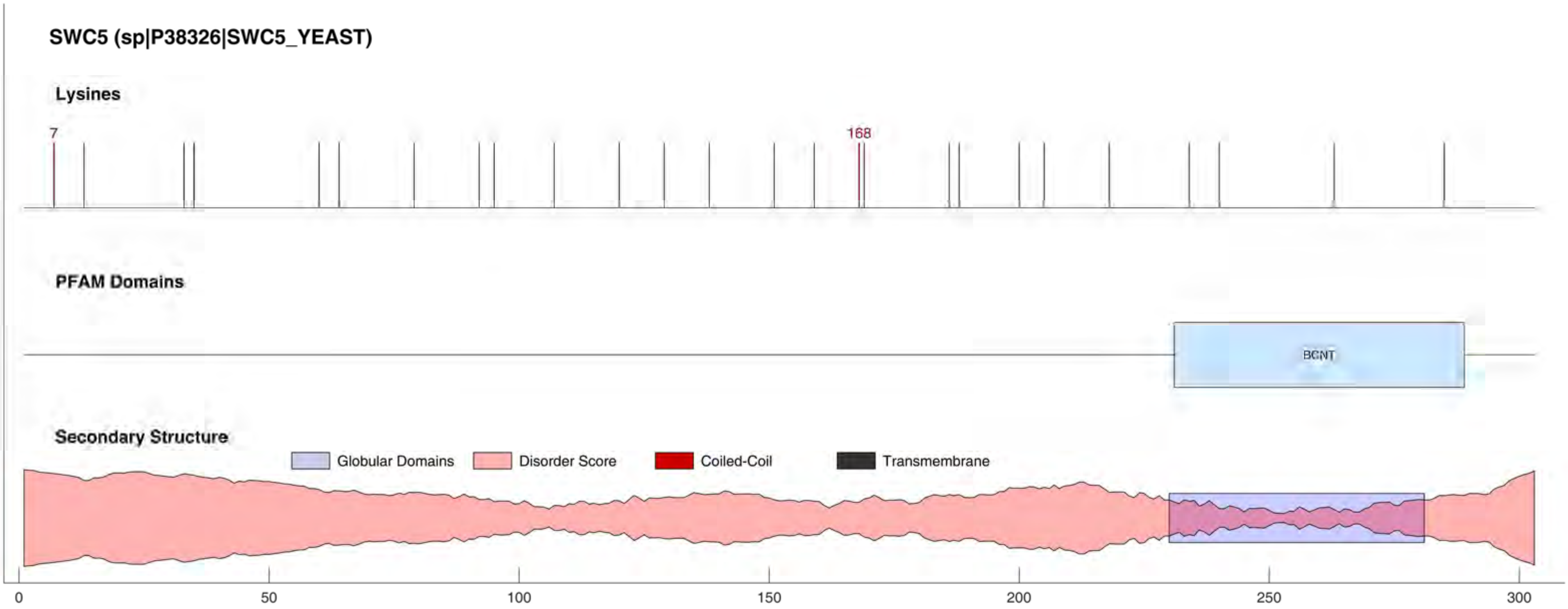

SWC5 (sp|P38326|SWC5_YEAST)
Lysines
7
168
PFAM Domains
BCNT
Secondary Structure
Globular Domains
Disorder Score
Coiled-Coil
Transmembrane
0
50
100
150
200
250
300

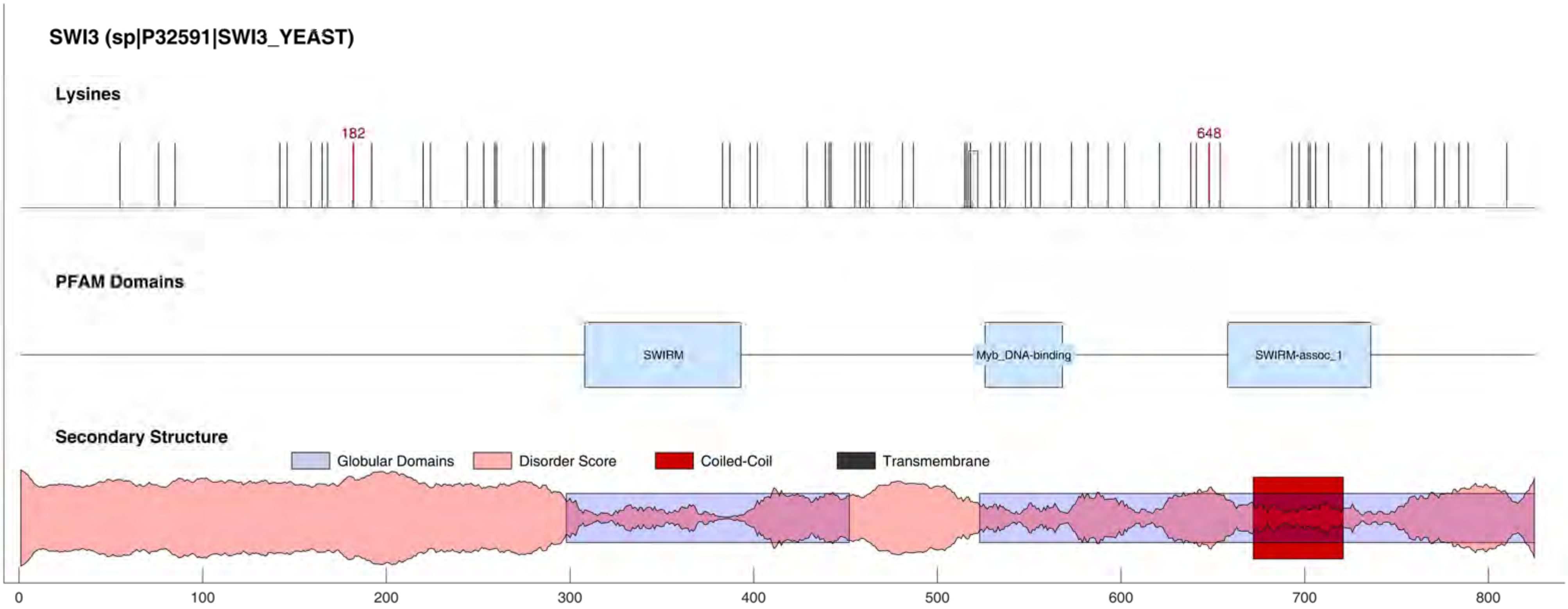

SWI3 (sp|P32591|SWI3_YEAST)
Lysines
182
648
PFAM Domains
SWIRM
Myb_DNA-binding
SWIRM-assoc_1
Secondary Structure
Globular Domains
Disorder Score
Coiled-Coil
Transmembrane
0
100
200
300
400
500
600
700
800

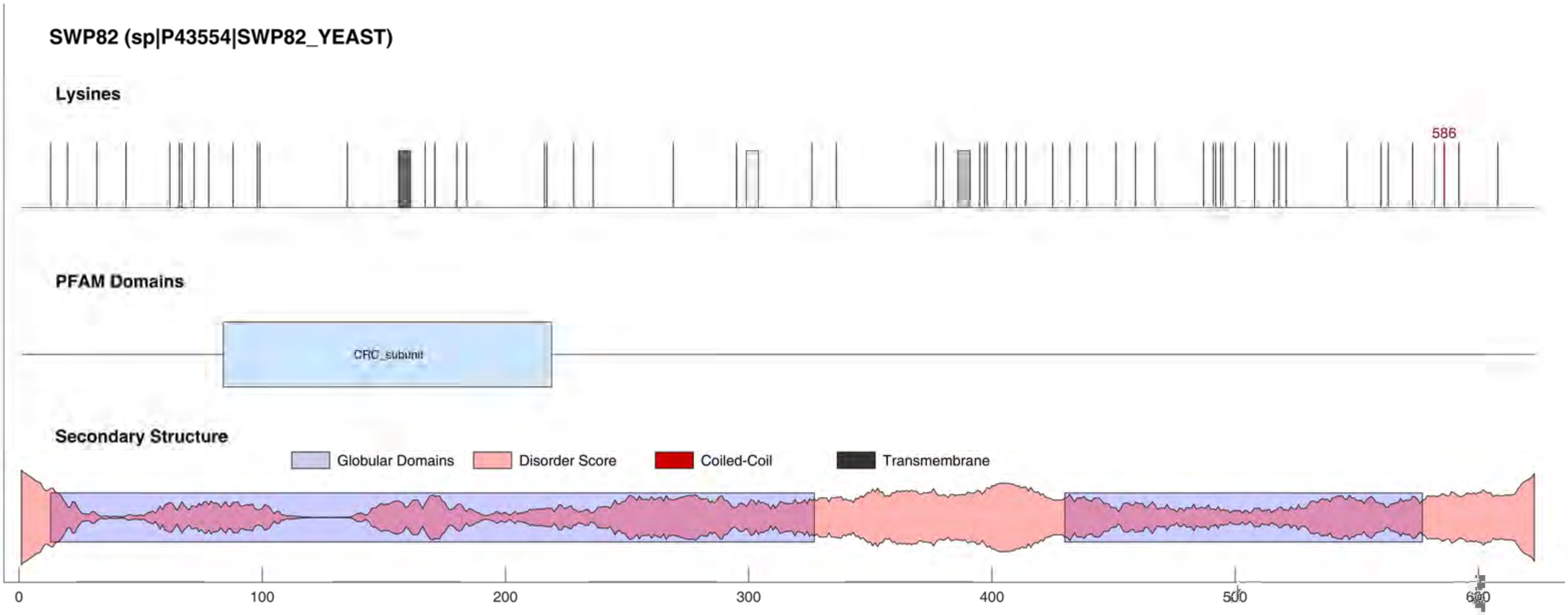

SWP82 (sp|P43554|SWP82_YEAST)
Lysines
586
PFAM Domains
CRC_subunit
Secondary Structure
Globular Domains
Disorder Score
Coiled-Coil
Transmembrane
0
100
200
300
400
500
600

#### SWR1 (sp|Q05471|SWR1_YEAST)

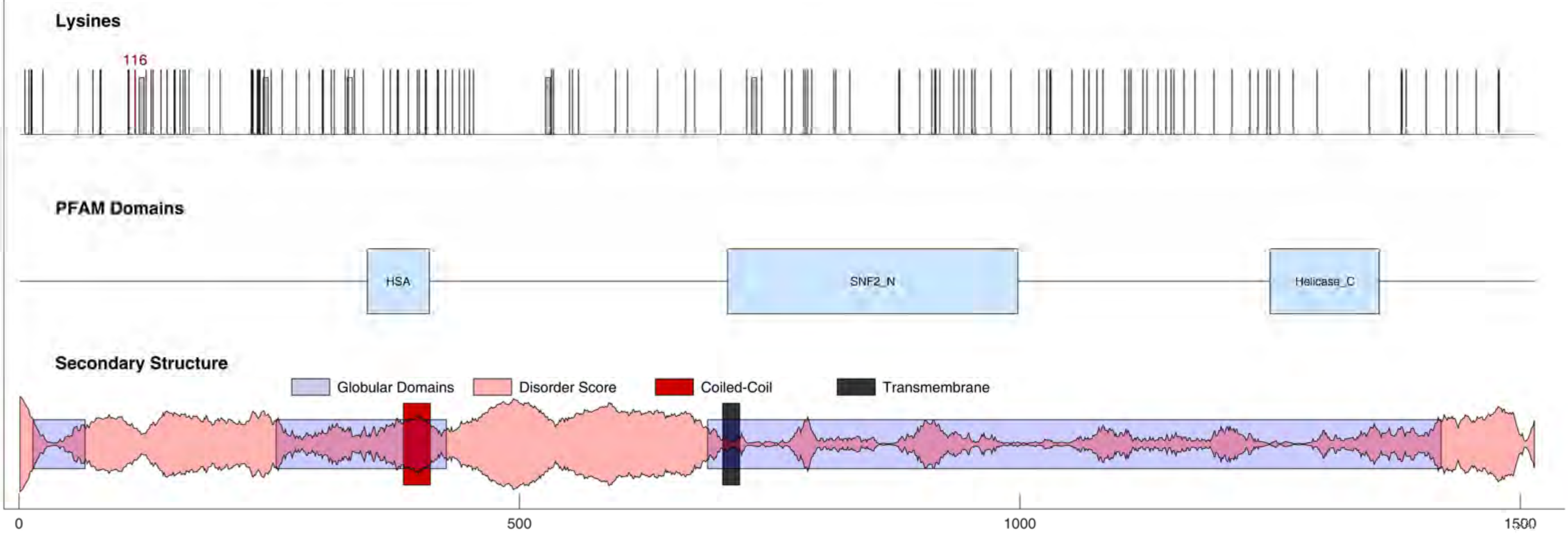

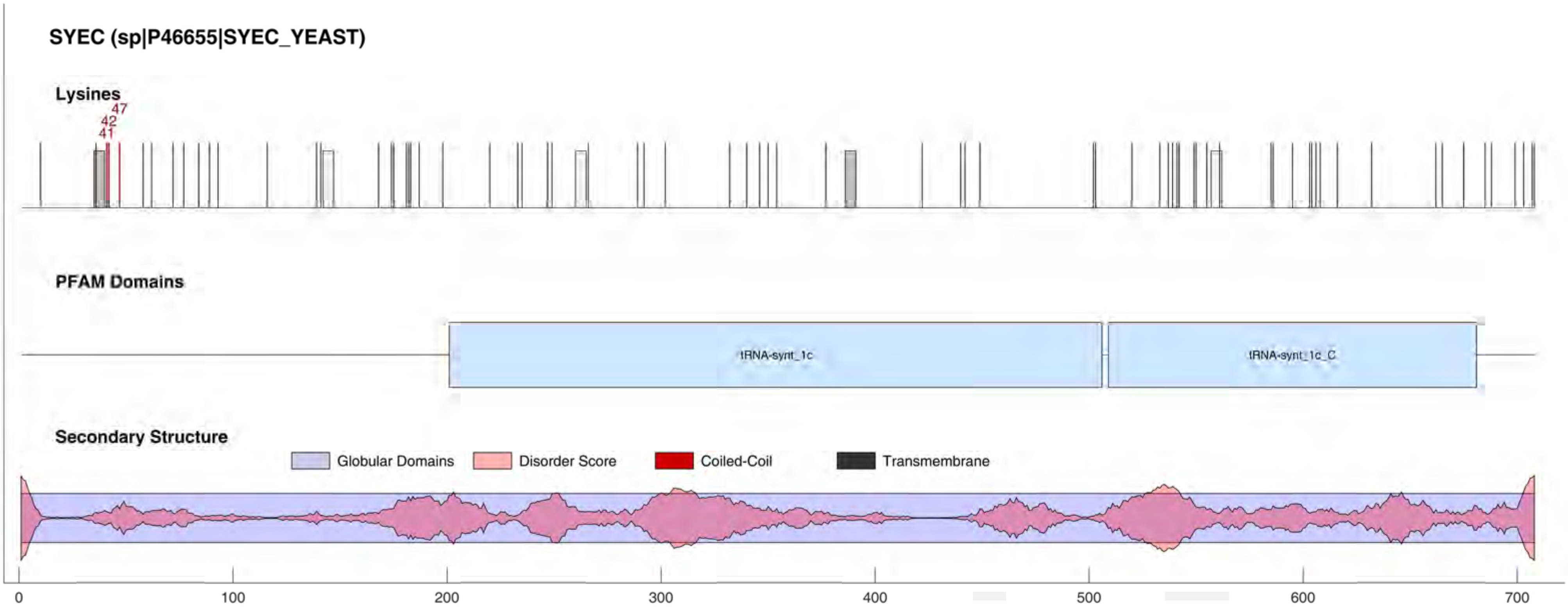
SYEC (sp|P46655|SYEC_YEAST)
Lysines
41
42
47
PFAM Domains
tRNA-synt_1c
tRNA-synt_1c_C
Secondary Structure
Globular Domains
Disorder Score
Coiled-Coil
Transmembrane
0
100
200
300
400
500
600
700

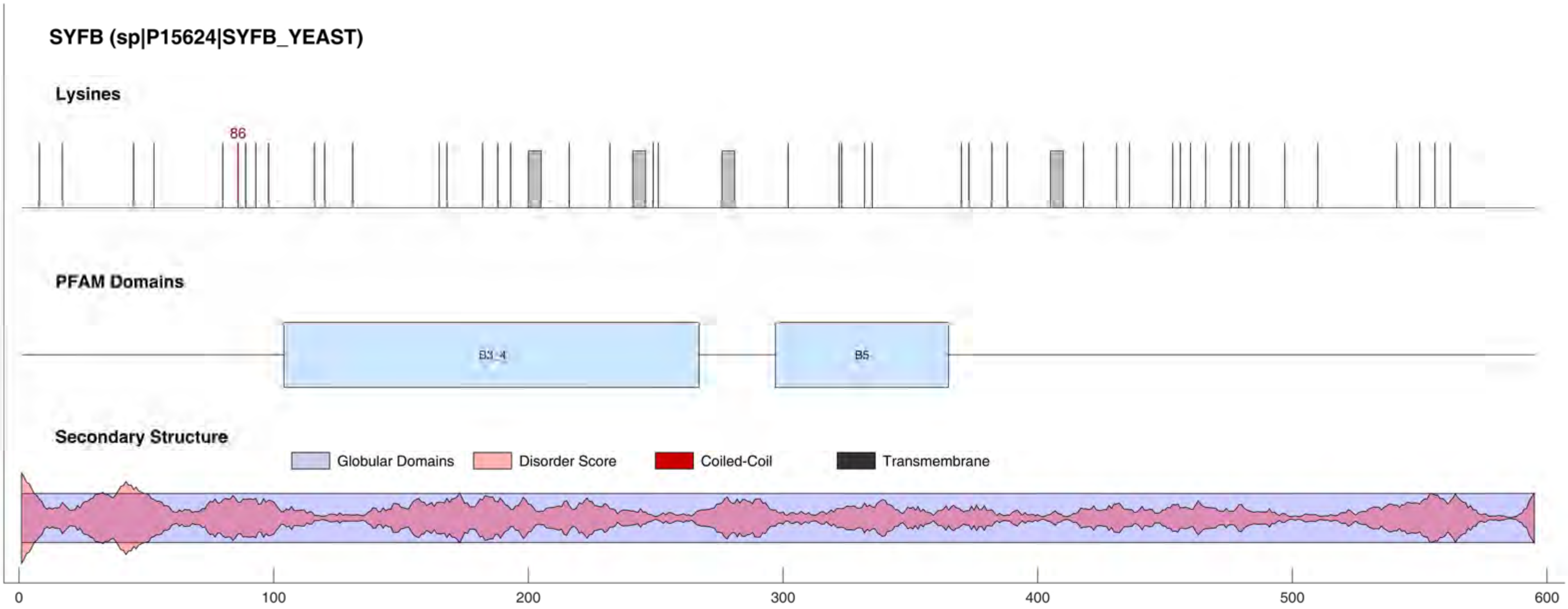
SYFB (sp|P15624|SYFB_YEAST)
Lysines
86
PFAM Domains
B3_4
B5
Secondary Structure
Globular Domains
Disorder Score
Coiled-Coil
Transmembrane
0
100
200
300
400
500
600

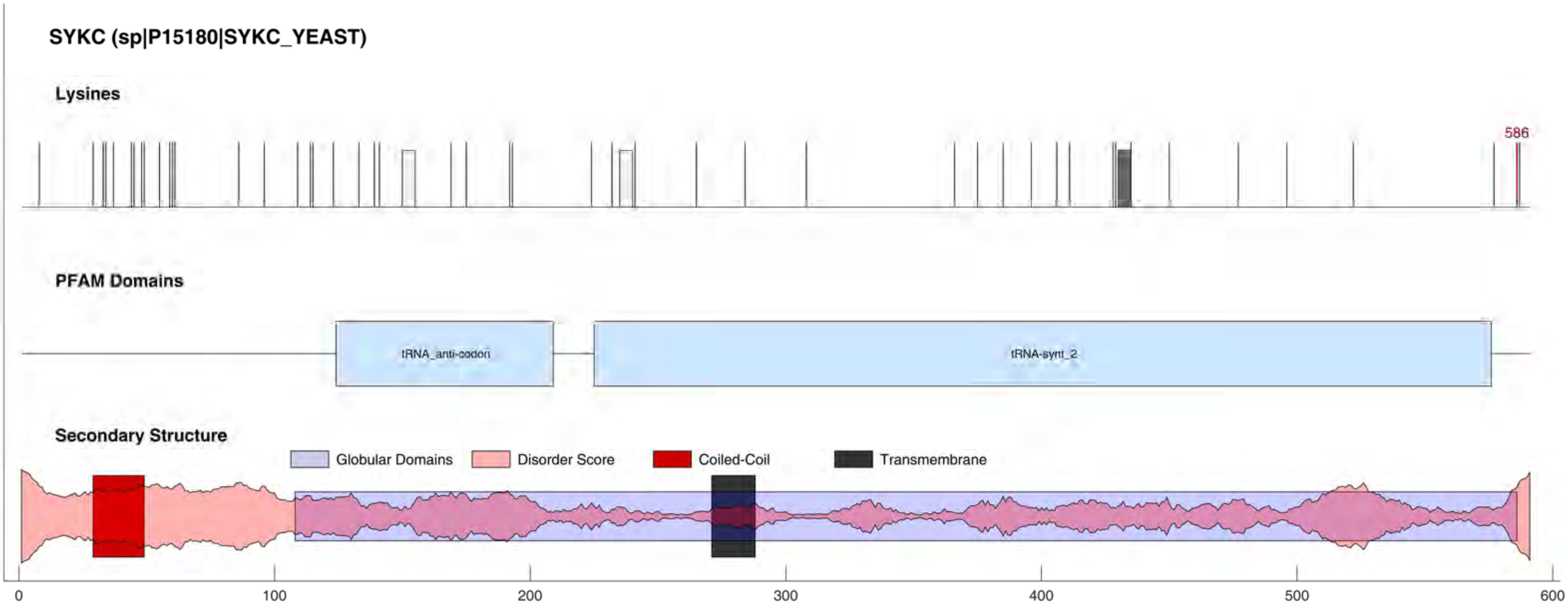
SYKC (sp|P15180|SYKC_YEAST)
Lysines
586
PFAM Domains
tRNA_anti-codon
tRNA-synt_2
Secondary Structure
Globular Domains
Disorder Score
Coiled-Coil
Transmembrane
0
100
200
300
400
500
600

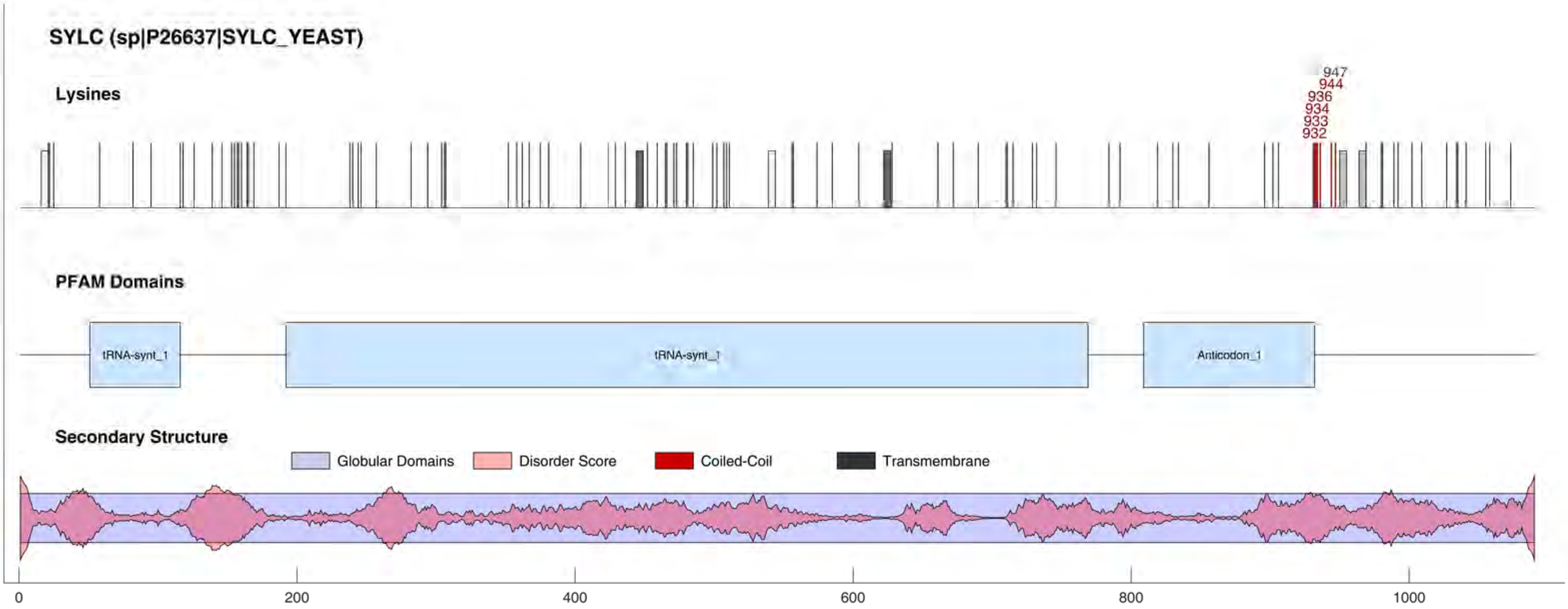
SYLC (sp|P26637|SYLC_YEAST)
Lysines
947
944
936
934
933
932
PFAM Domains
tRNA-synt_1
tRNA-synt_1
Anticodon_1
Secondary Structure
Globular Domains
Disorder Score
Coiled-Coil
Transmembrane
0
200
400
600
800
1000

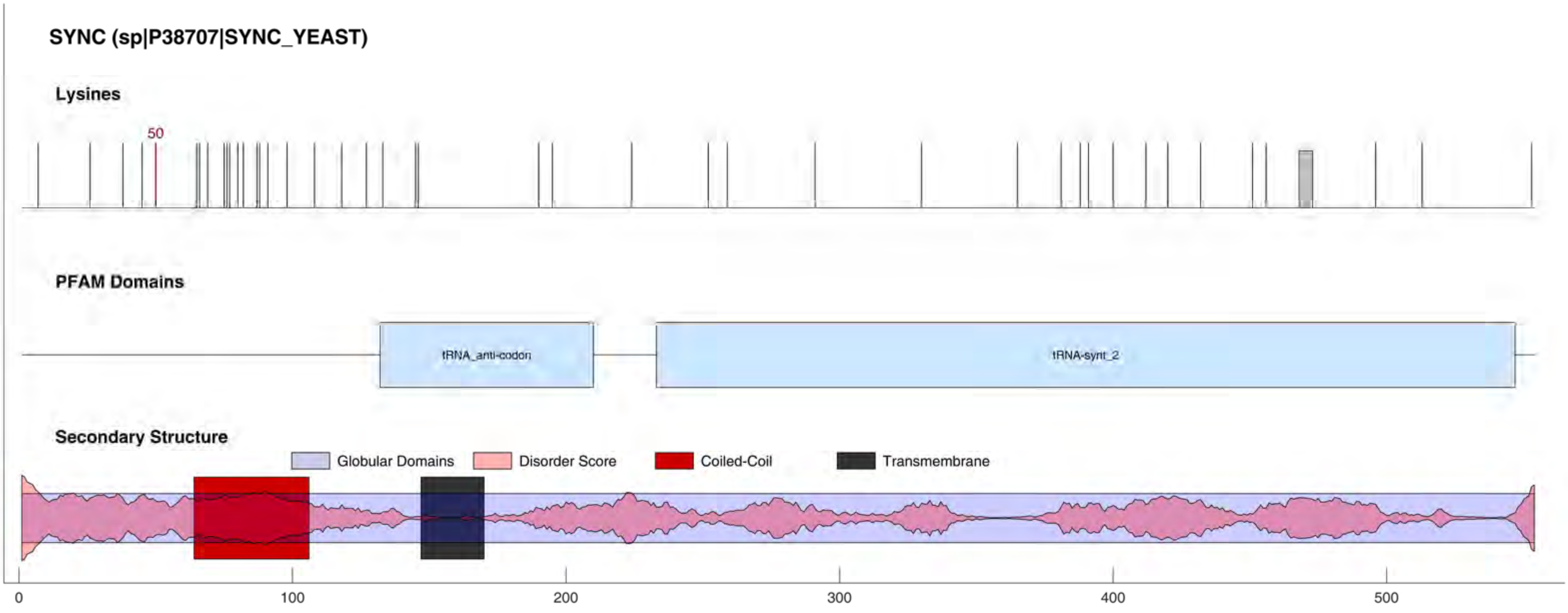

SYNC (sp|P38707|SYNC_YEAST)
Lysines
50
PFAM Domains
tRNA_anti-codon
tRNA-synt_2
Secondary Structure
Globular Domains
Disorder Score
Coiled-Coil
Transmembrane
0
100
200
300
400
500

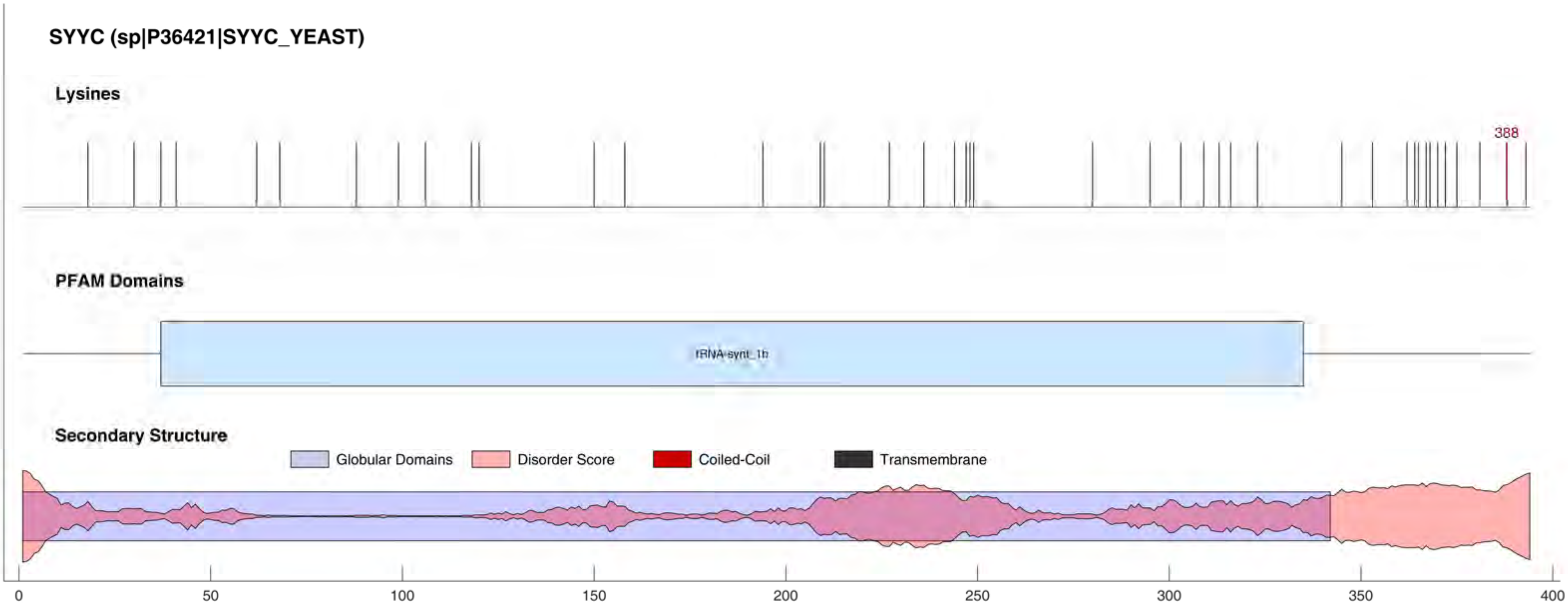

SYYC (sp|P36421|SYYC_YEAST)
Lysines
388
PFAM Domains
tRNA-synt_1b
Secondary Structure
Globular Domains
Disorder Score
Coiled-Coil
Transmembrane
0
50
100
150
200
250
300
350
400

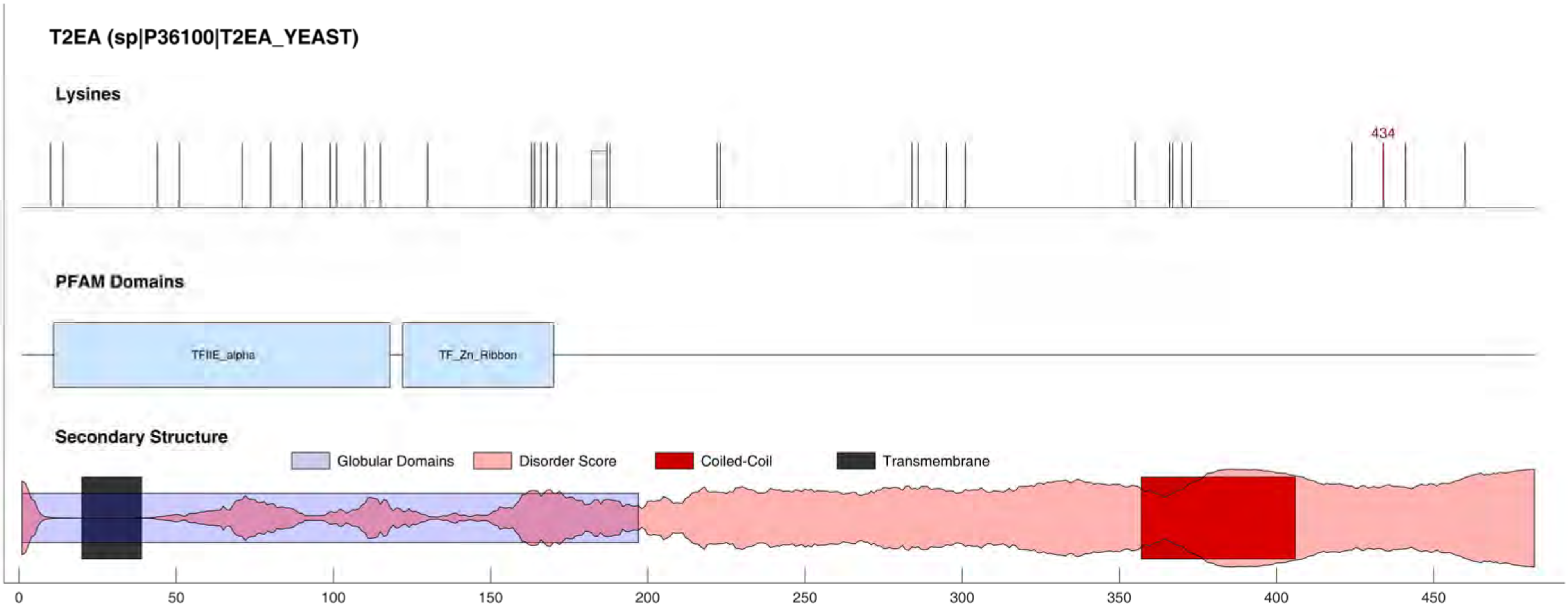
T2EA (sp|P36100|T2EA_YEAST)
Lysines
434
PFAM Domains
TFIIE_alpha
TF_Zn_Ribbon
Secondary Structure
Globular Domains
Disorder Score
Coiled-Coil
Transmembrane
0
50
100
150
200
250
300
350
400
450

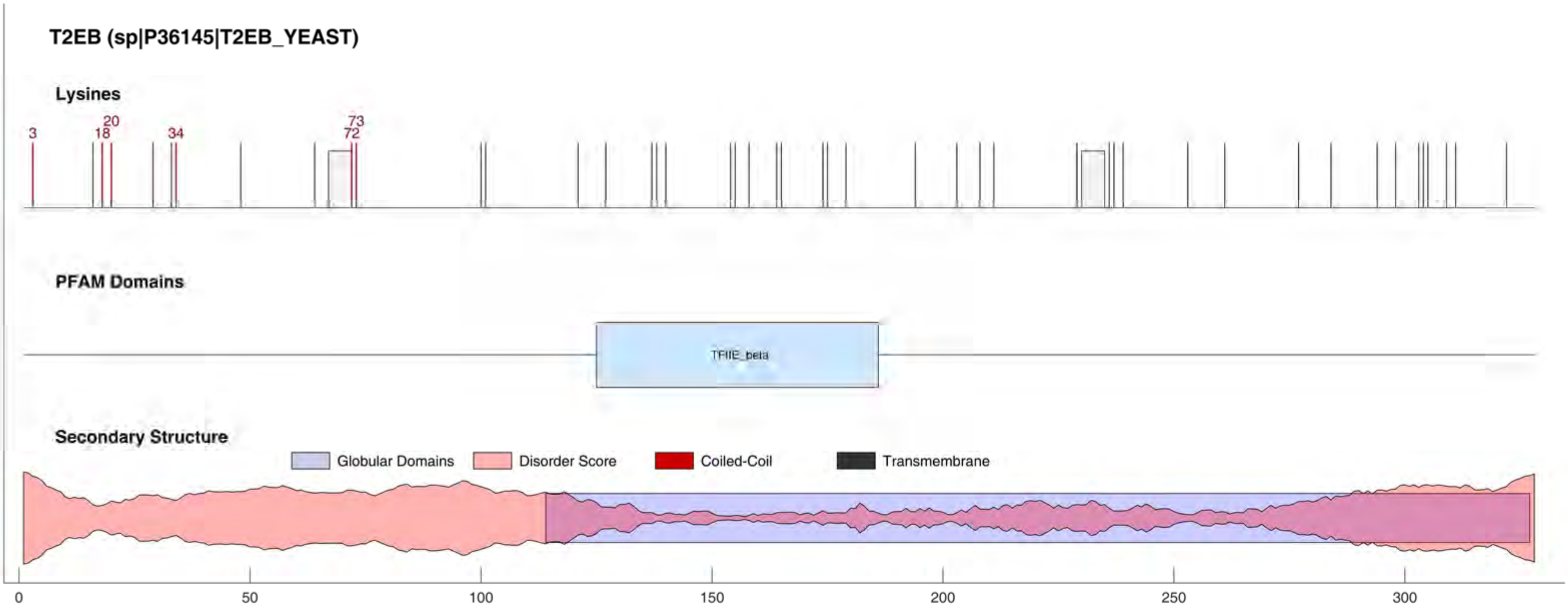
T2EB (sp|P36145|T2EB_YEAST)
Lysines
3
18
20
34
72
73
PFAM Domains
TFIIE_beta
Secondary Structure
Globular Domains
Disorder Score
Coiled-Coil
Transmembrane
0
50
100
150
200
250
300

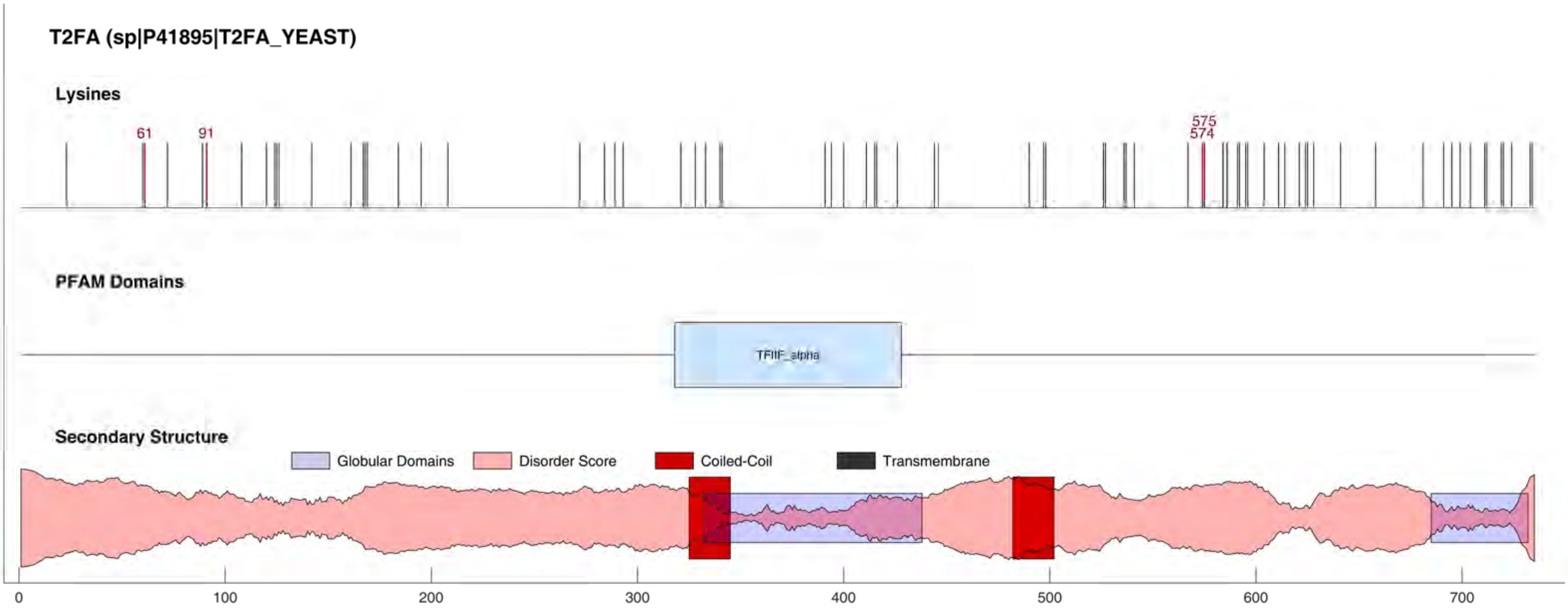
T2FA (sp|P41895|T2FA_YEAST)
Lysines
61
91
575
574
PFAM Domains
TFIIF_alpha
Secondary Structure
Globular Domains
Disorder Score
Coiled-Coil
Transmembrane
0
100
200
300
400
500
600
700

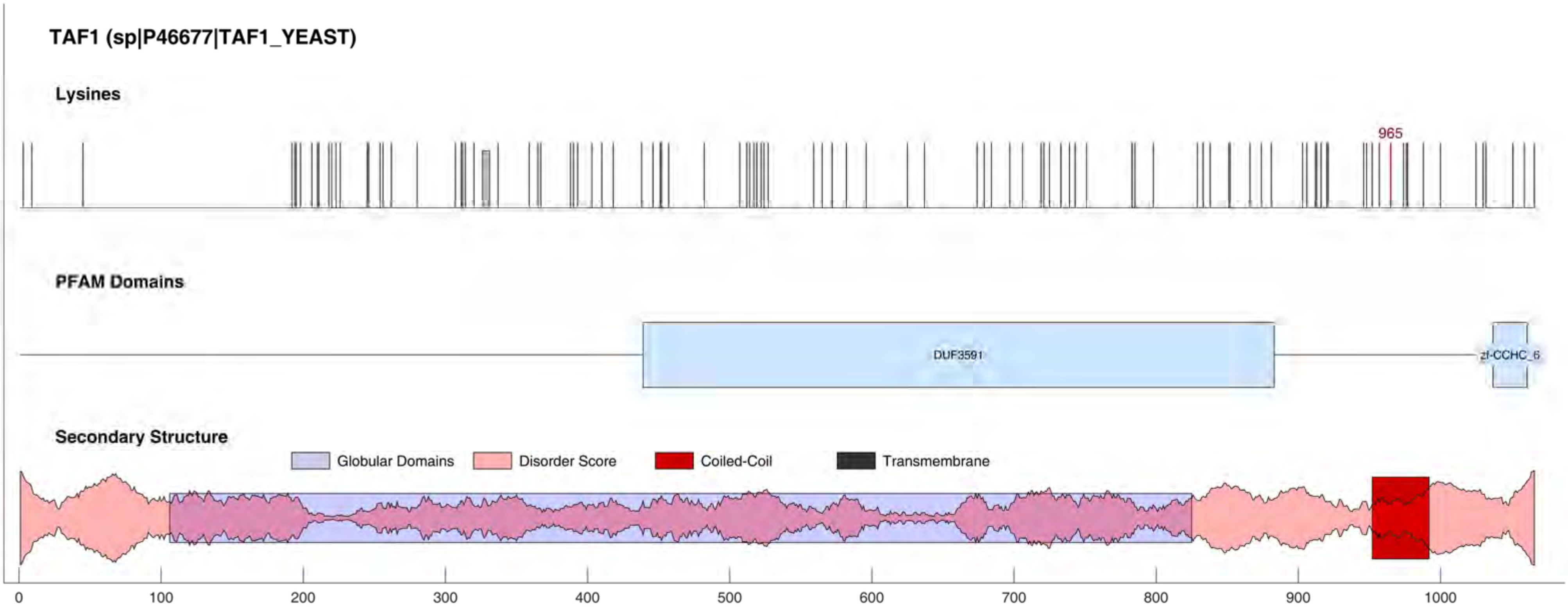
TAF1 (sp|P46677|TAF1_YEAST)
Lysines
965
PFAM Domains
DUF3591
zf-CCHC_6
Secondary Structure
Globular Domains
Disorder Score
Coiled-Coil
Transmembrane
0
100
200
300
400
500
600
700
800
900
1000

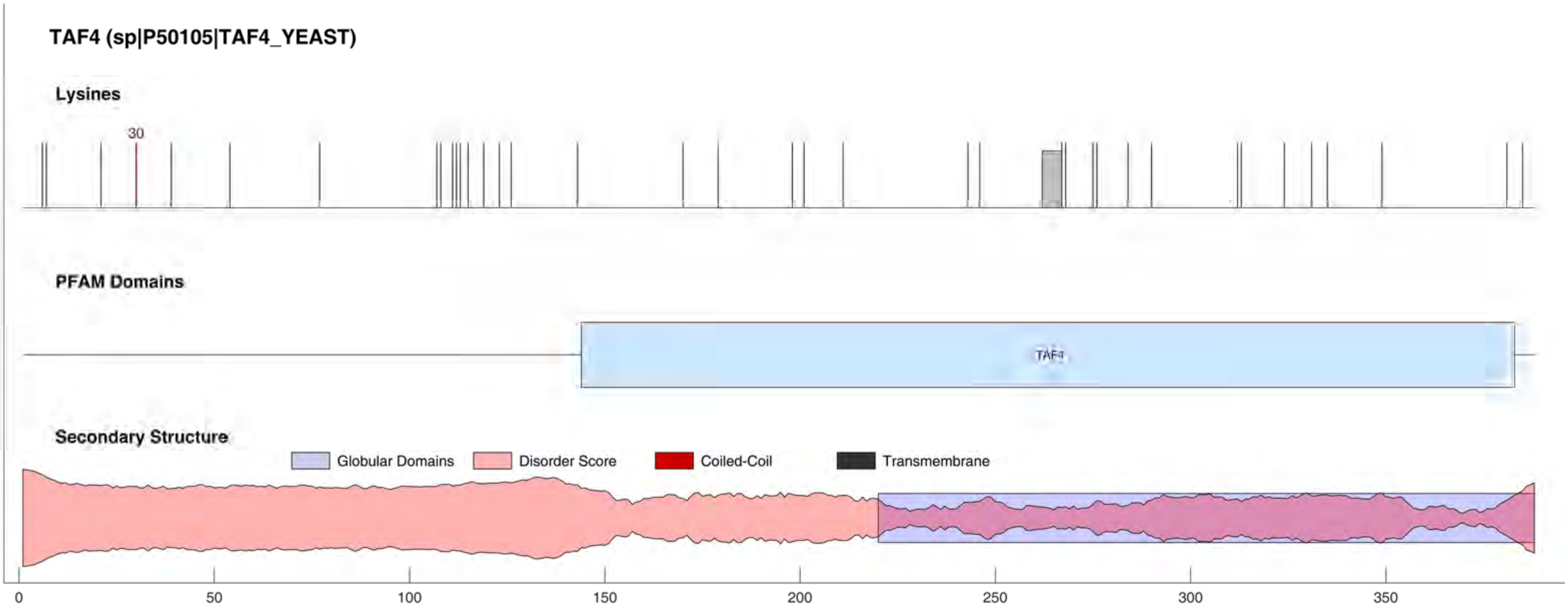
TAF4 (sp|P50105|TAF4_YEAST)
Lysines
30
PFAM Domains
TAF4
Secondary Structure
Globular Domains
Disorder Score
Coiled-Coil
Transmembrane
0
50
100
150
200
250
300
350

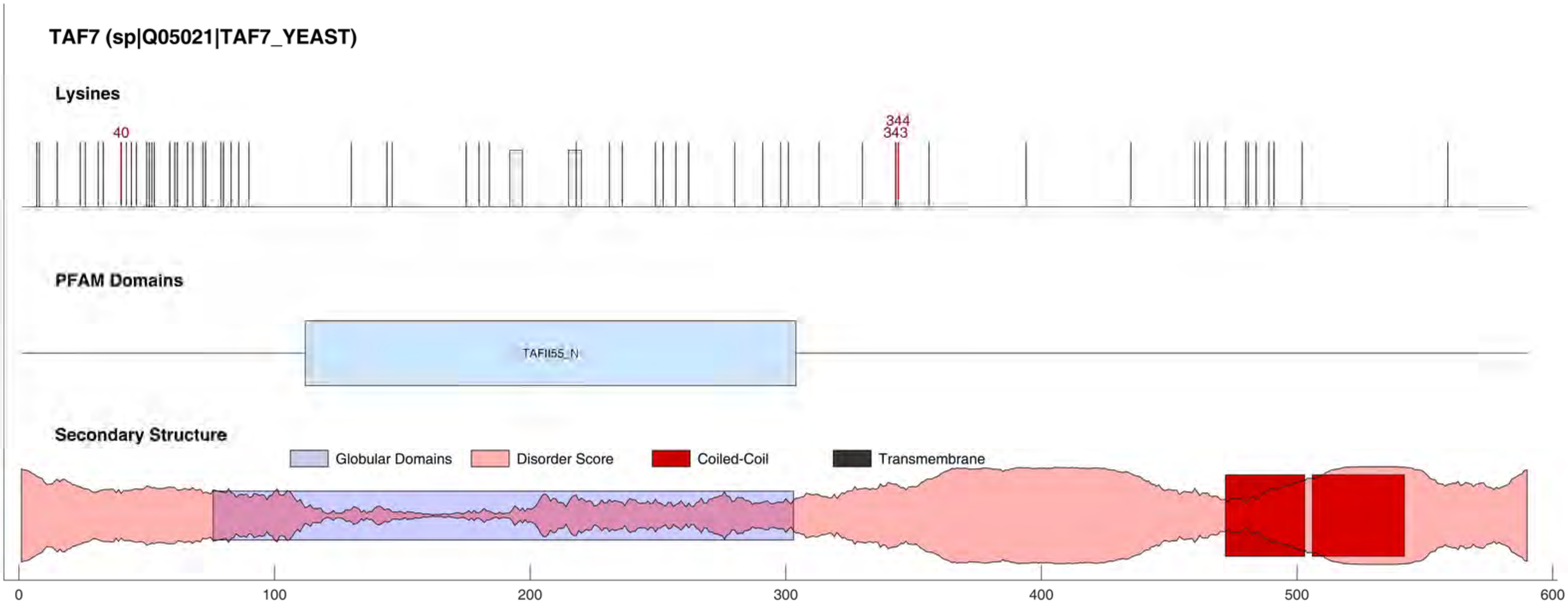
TAF7 (sp|Q05021|TAF7_YEAST)
Lysines
40
344
343
PFAM Domains
TAFII55_N
Secondary Structure
Globular Domains
Disorder Score
Coiled-Coil
Transmembrane
0
100
200
300
400
500
600

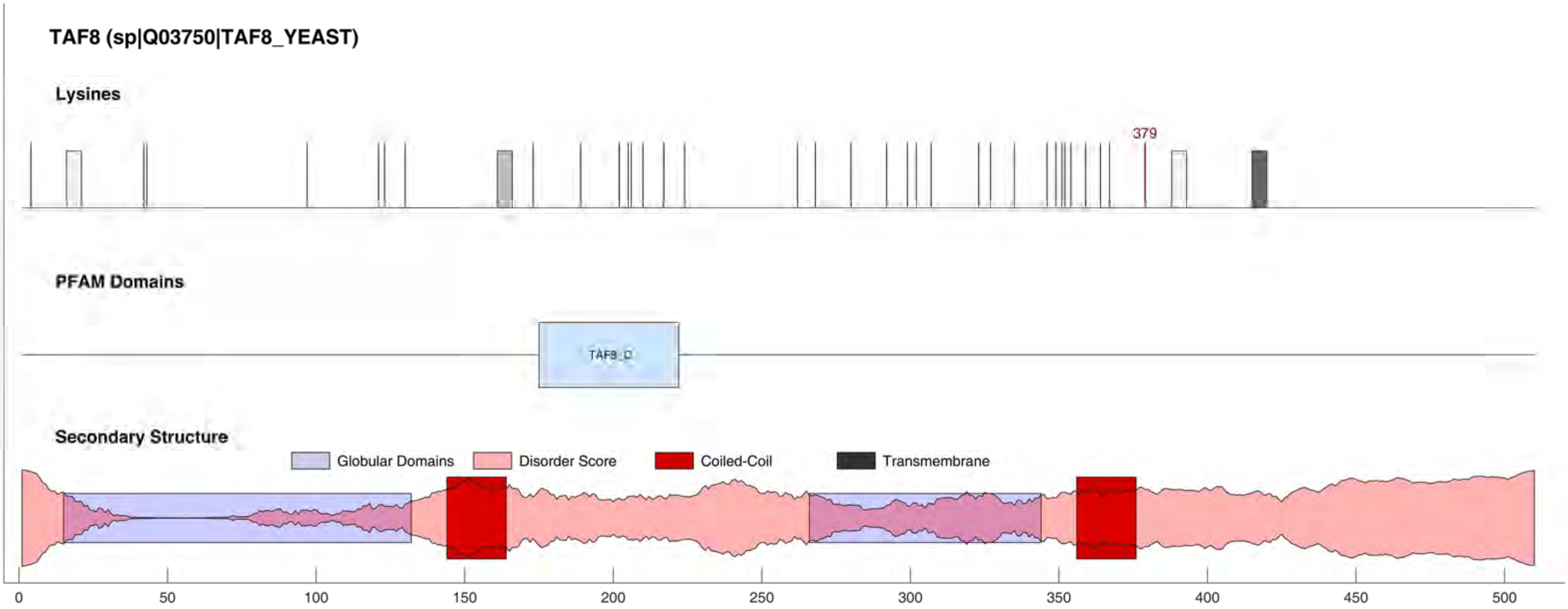
TAF8 (sp|Q03750|TAF8_YEAST)
Lysines
379
PFAM Domains
TAF8_C
Secondary Structure
Globular Domains
Disorder Score
Coiled-Coil
Transmembrane
0
50
100
150
200
250
300
350
400
450
500

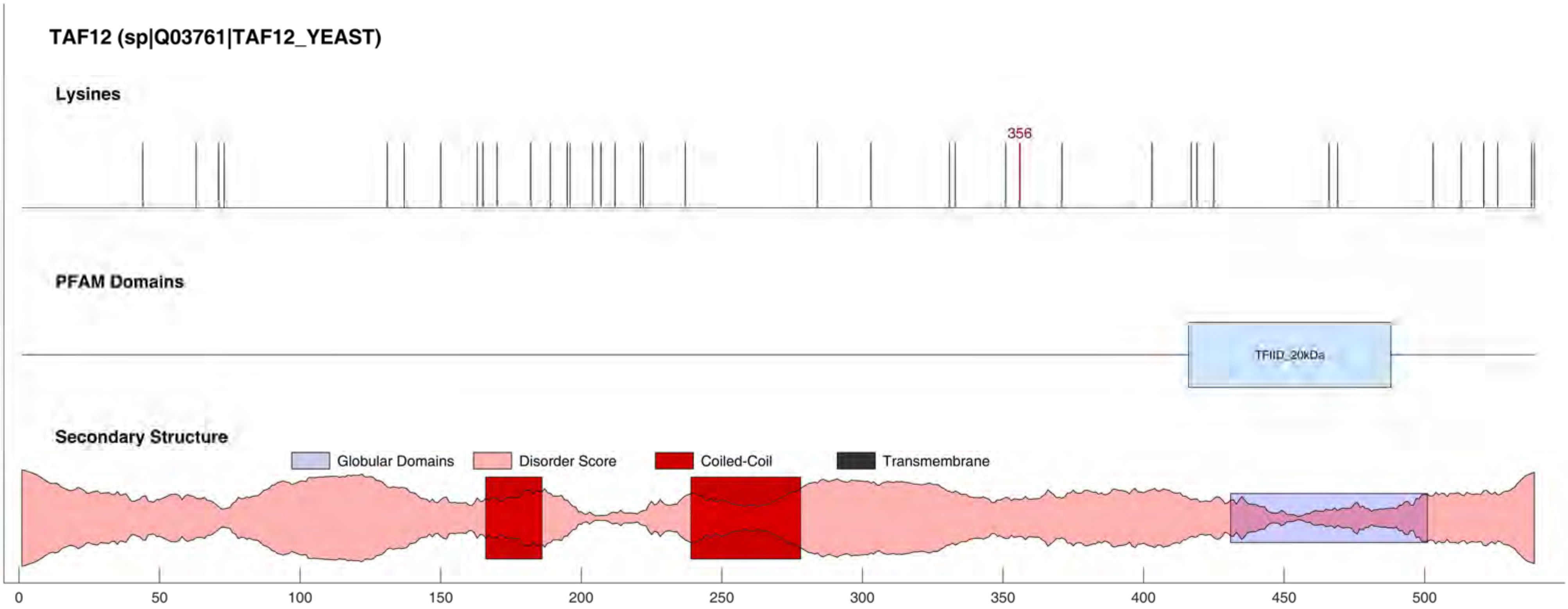

TAF12 (sp|Q03761|TAF12_YEAST)
Lysines
356
PFAM Domains
TFIID_20kDa
Secondary Structure
Globular Domains
Disorder Score
Coiled-Coil
Transmembrane
0
50
100
150
200
250
300
350
400
450
500

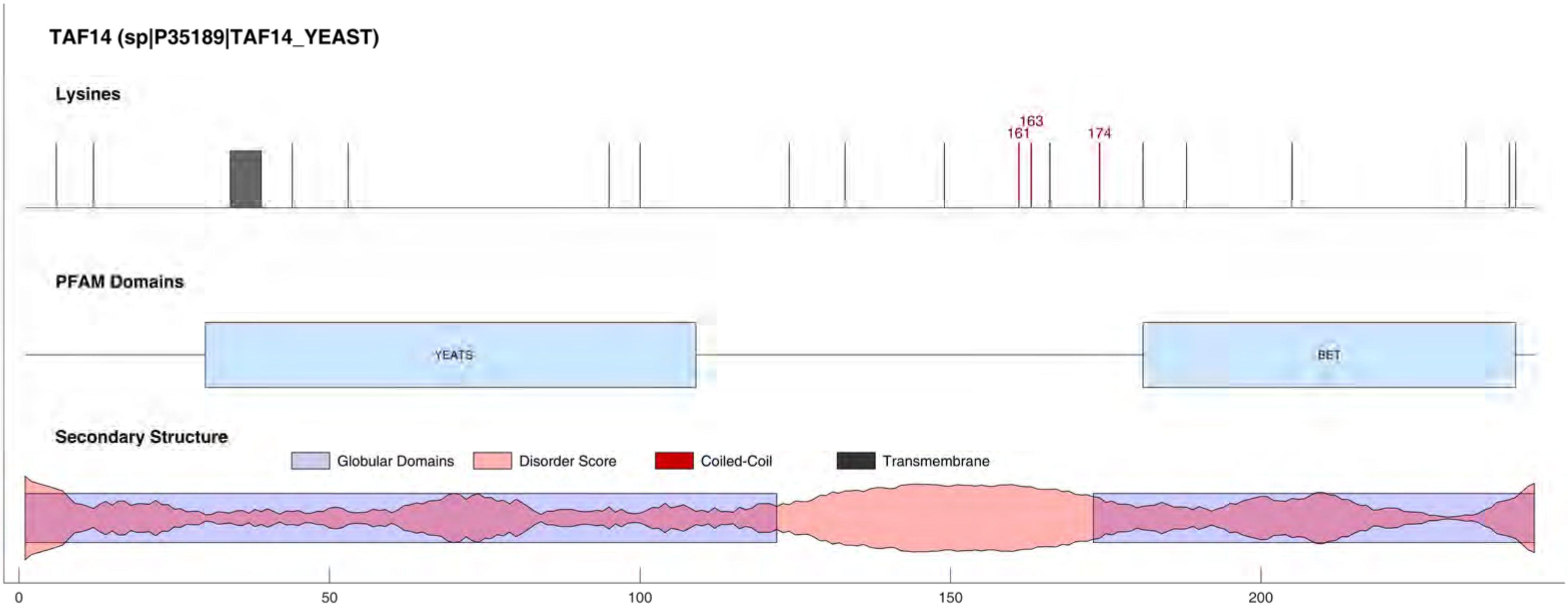
TAF14 (sp|P35189|TAF14_YEAST)
Lysines
161
163
174
PFAM Domains
YEATS
BET
Secondary Structure
Globular Domains
Disorder Score
Coiled-Coil
Transmembrane
0
50
100
150
200

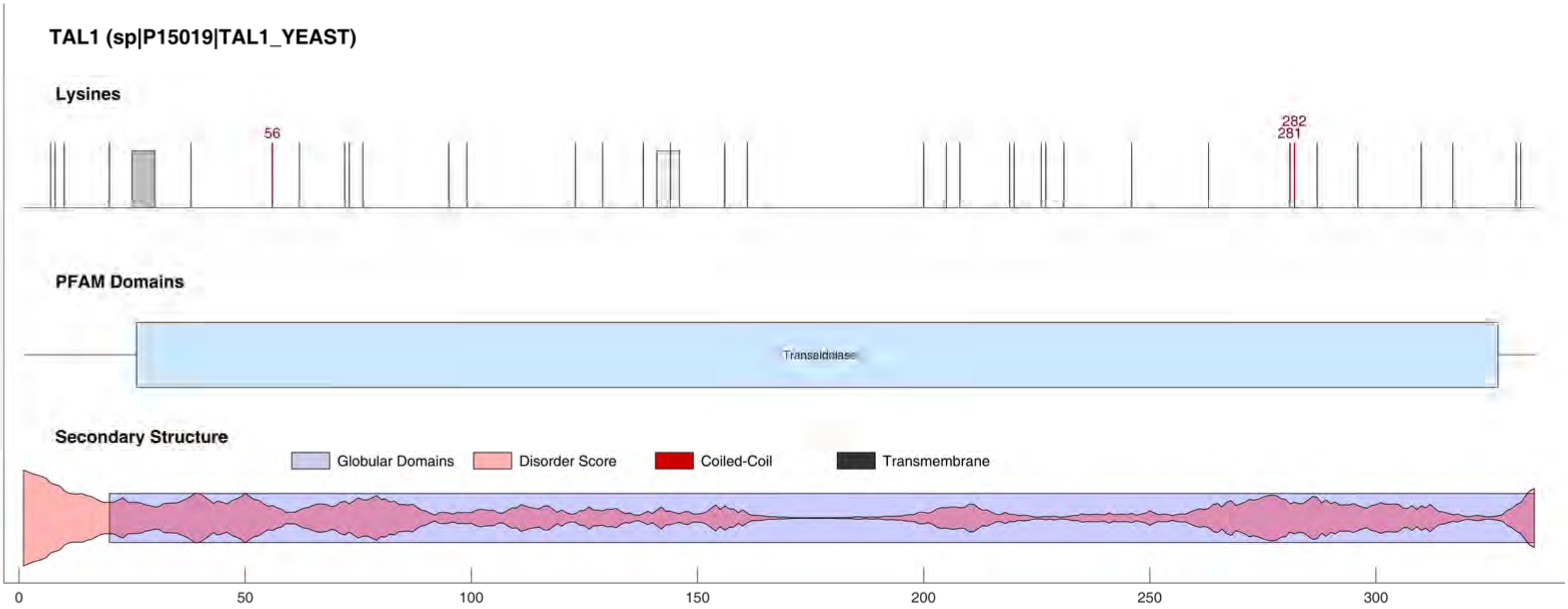

TAL1 (sp|P15019|TAL1_YEAST)
Lysines
56
282
281
PFAM Domains
Transaldolase
Secondary Structure
Globular Domains
Disorder Score
Coiled-Coil
Transmembrane
0
50
100
150
200
250
300

#### TAL2 (sp|P53228|TAL2_YEAST)

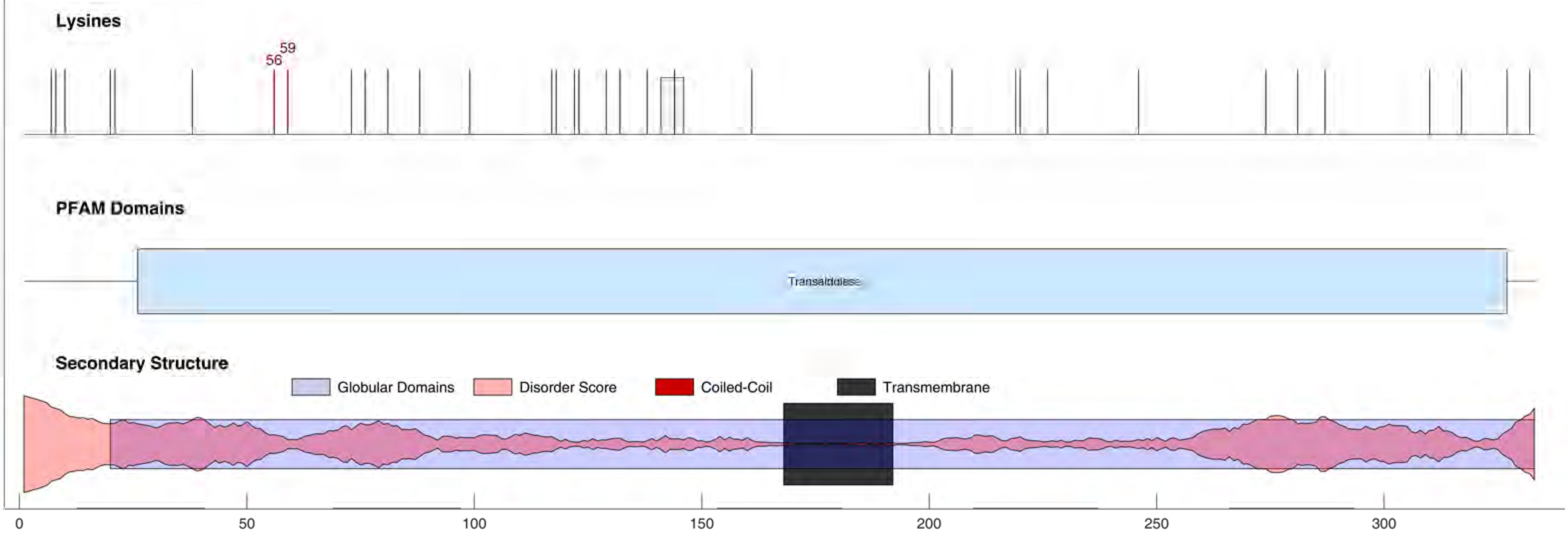

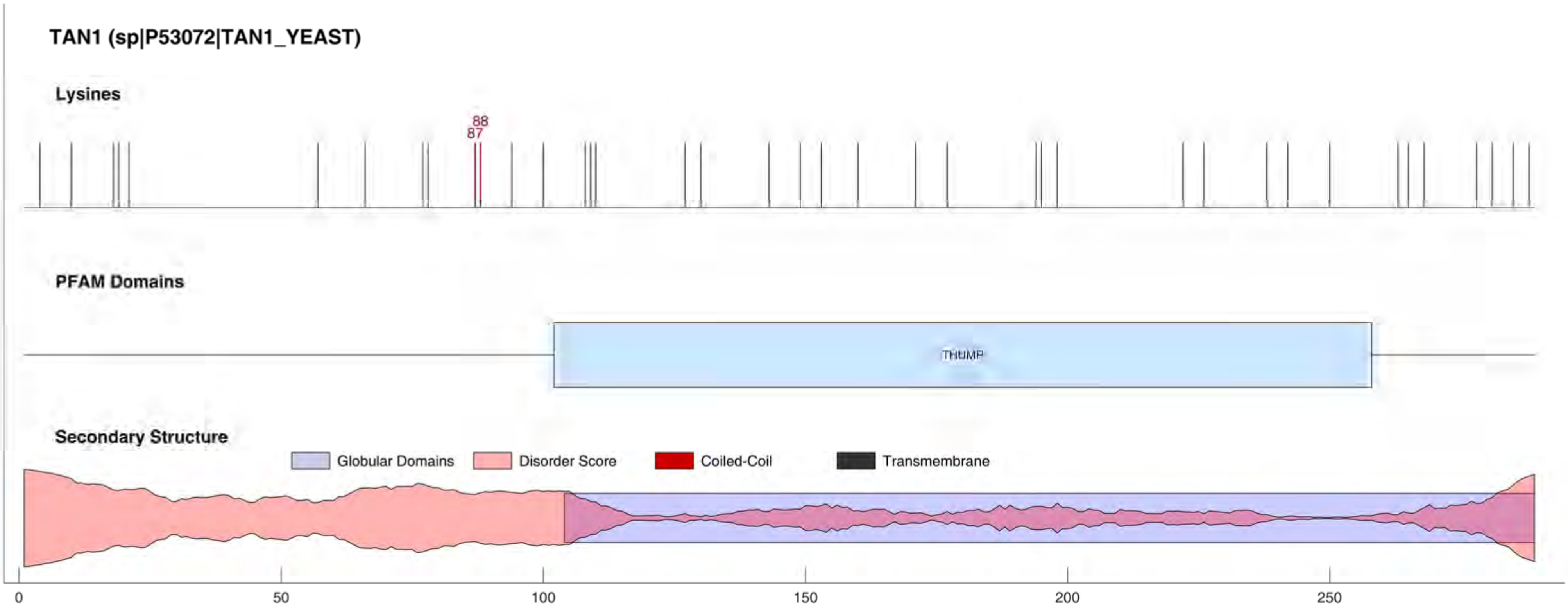
TAN1 (sp|P53072|TAN1_YEAST)
Lysines
88
87
PFAM Domains
THUMP
Secondary Structure
Globular Domains
Disorder Score
Coiled-Coil
Transmembrane
0
50
100
150
200
250

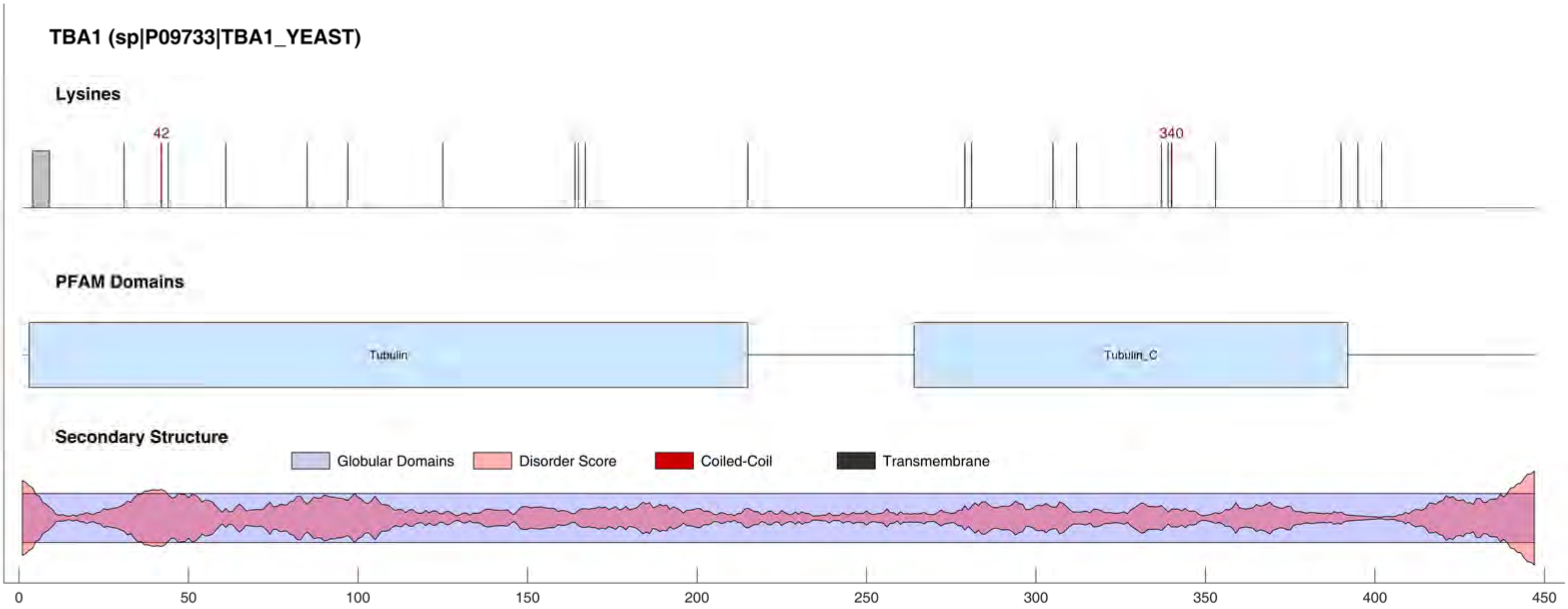

TBA1 (sp|P09733|TBA1_YEAST)
Lysines
42
340
PFAM Domains
Tubulin
Tubulin_C
Secondary Structure
Globular Domains
Disorder Score
Coiled-Coil
Transmembrane
0
50
100
150
200
250
300
350
400
450

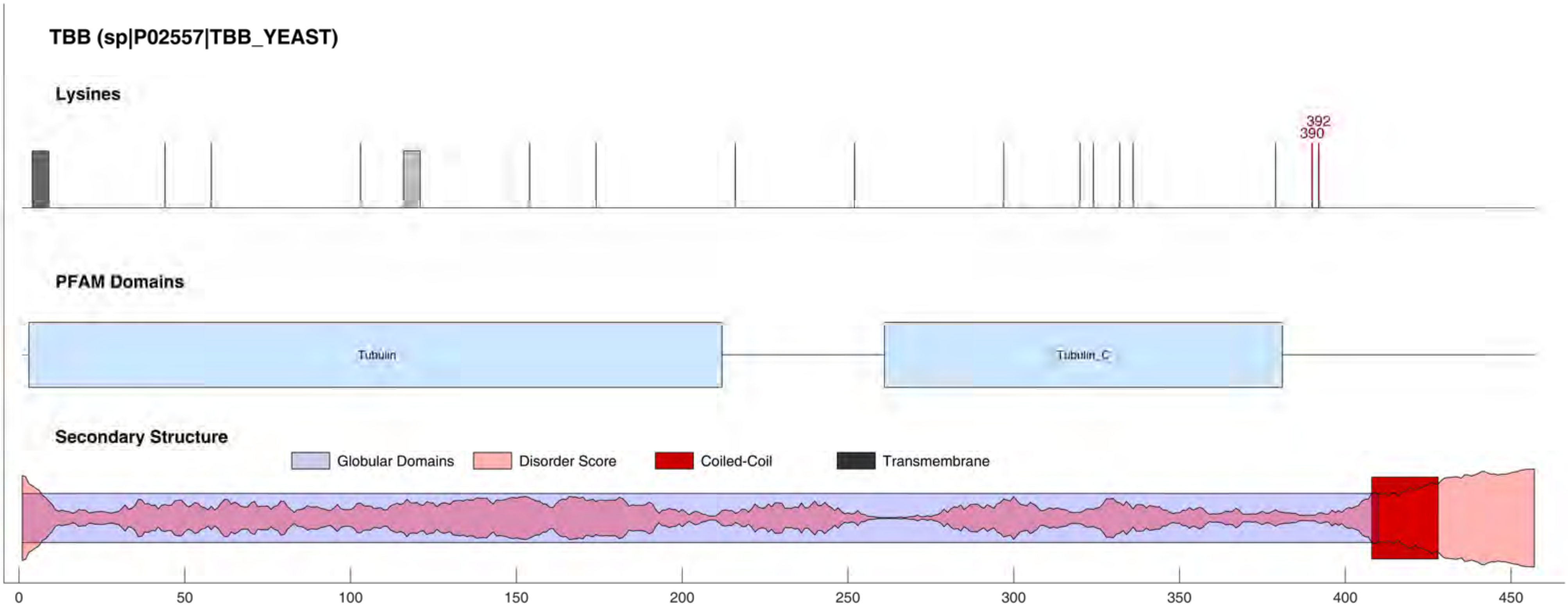
TBB (sp|P02557|TBB_YEAST)
Lysines
392
390
PFAM Domains
Tubulin
Tubulin_C
Secondary Structure
Globular Domains
Disorder Score
Coiled-Coil
Transmembrane
0
50
100
150
200
250
300
350
400
450

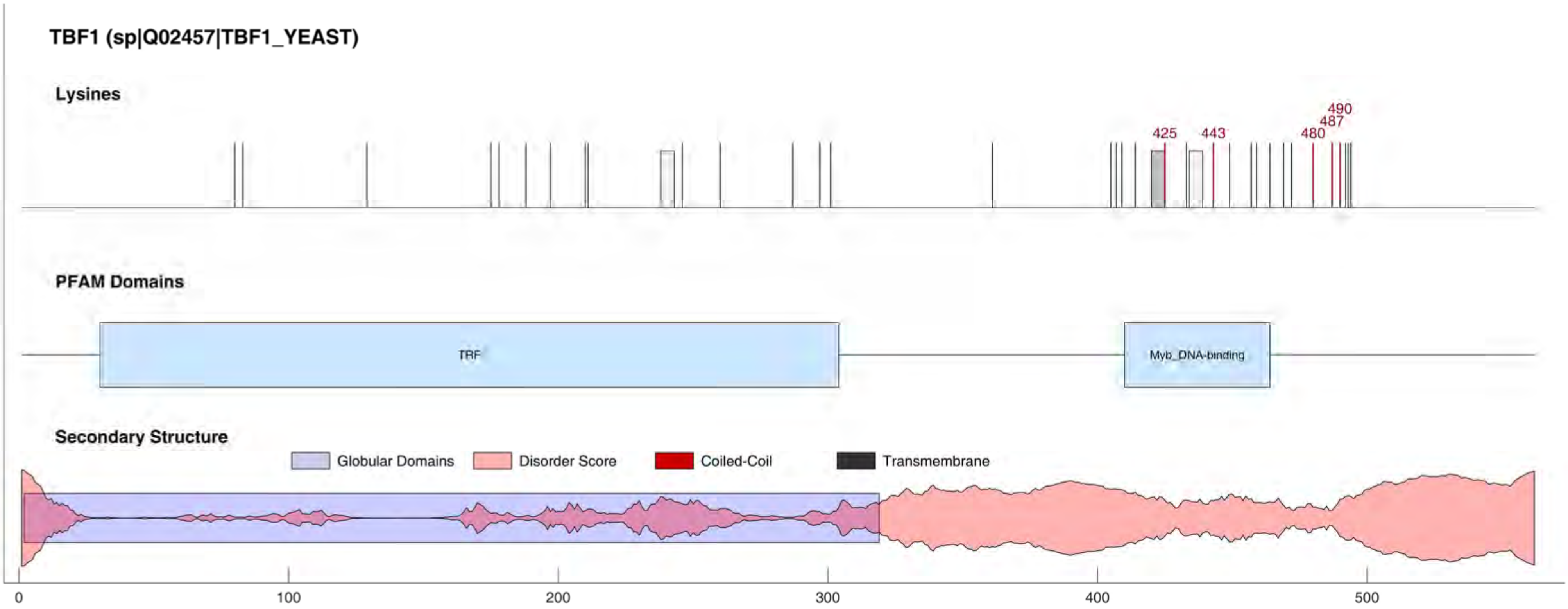

TBF1 (sp|Q02457|TBF1_YEAST)
Lysines
425
443
480
487
490
PFAM Domains
TRF
Myb_DNA-binding
Secondary Structure
Globular Domains
Disorder Score
Coiled-Coil
Transmembrane
0
100
200
300
400
500

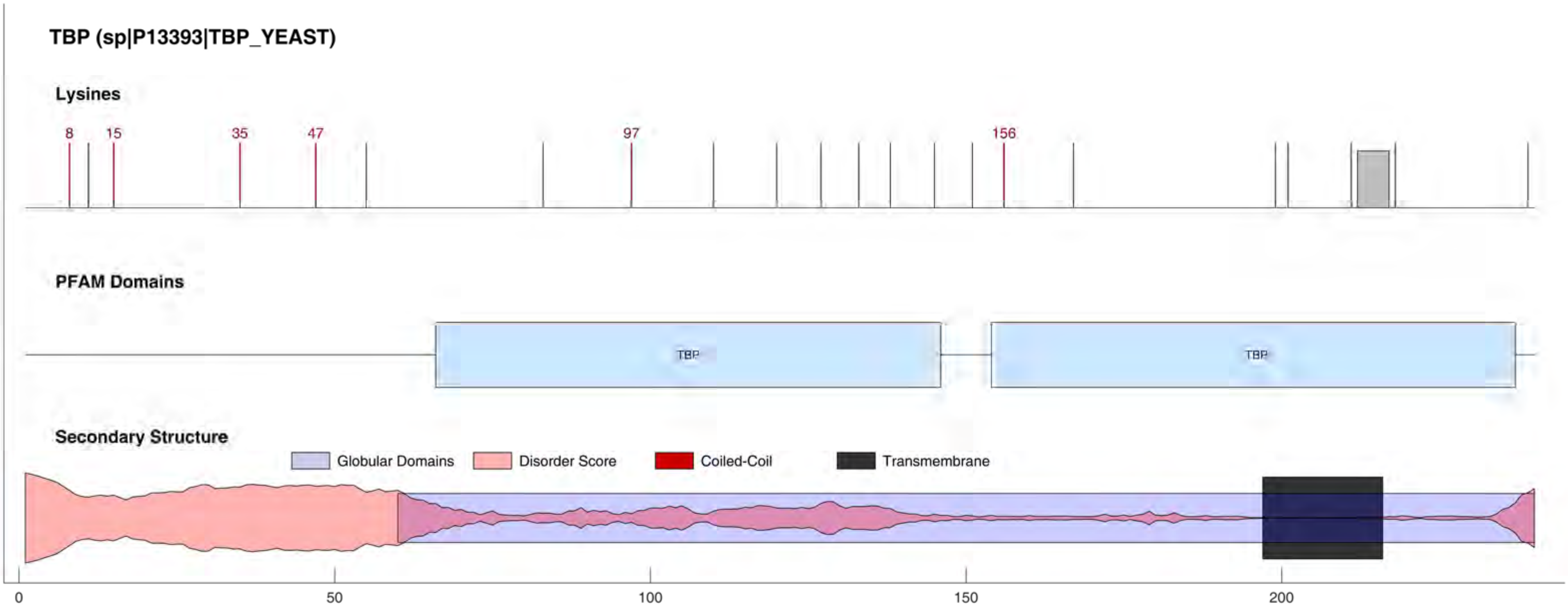

TBP (sp|P13393|TBP_YEAST)
Lysines
8
15
35
47
97
156
PFAM Domains
TBP
TBP
Secondary Structure
Globular Domains
Disorder Score
Coiled-Coil
Transmembrane
0
50
100
150
200

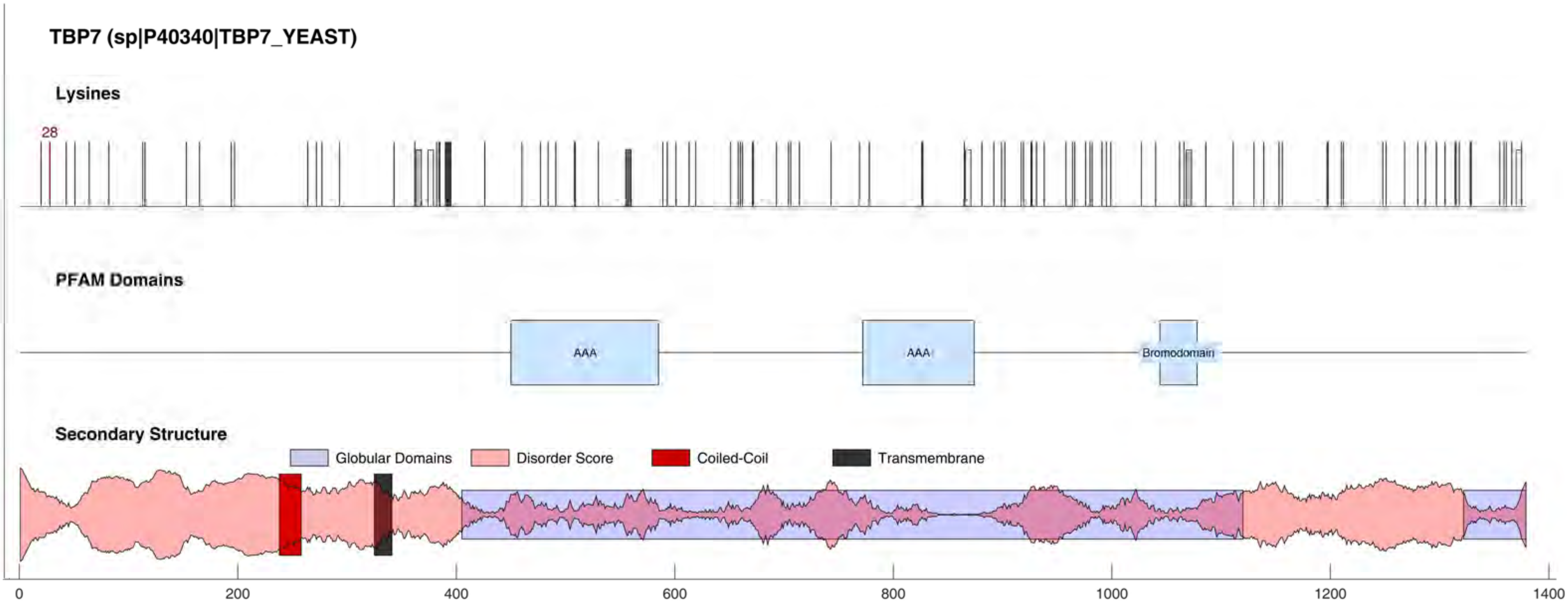

TBP7 (sp|P40340|TBP7_YEAST)
Lysines
28
PFAM Domains
AAA
AAA
Bromodomain
Secondary Structure
Globular Domains
Disorder Score
Coiled-Coil
Transmembrane
0
200
400
600
800
1000
1200
1400

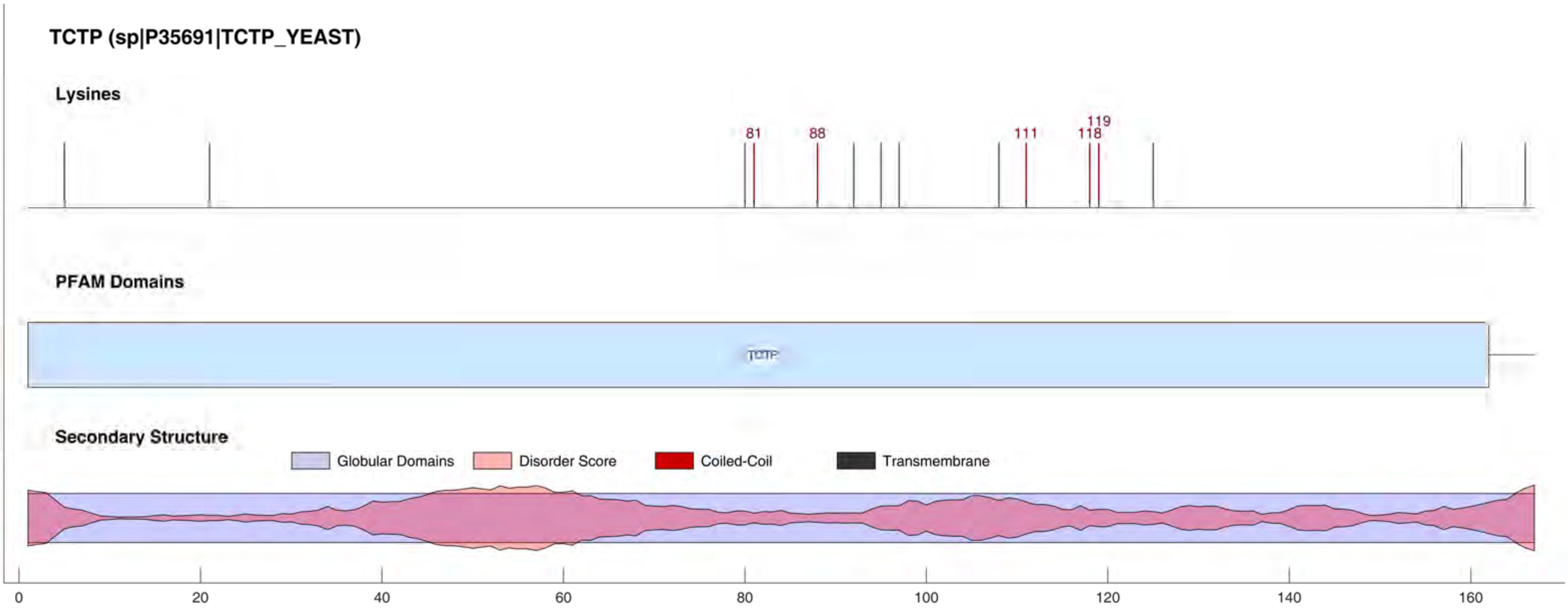

TCTP (sp|P35691|TCTP_YEAST)
Lysines
81
88
111
118
119
PFAM Domains
TCTP
Secondary Structure
Globular Domains
Disorder Score
Coiled-Coil
Transmembrane
0
20
40
60
80
100
120
140
160

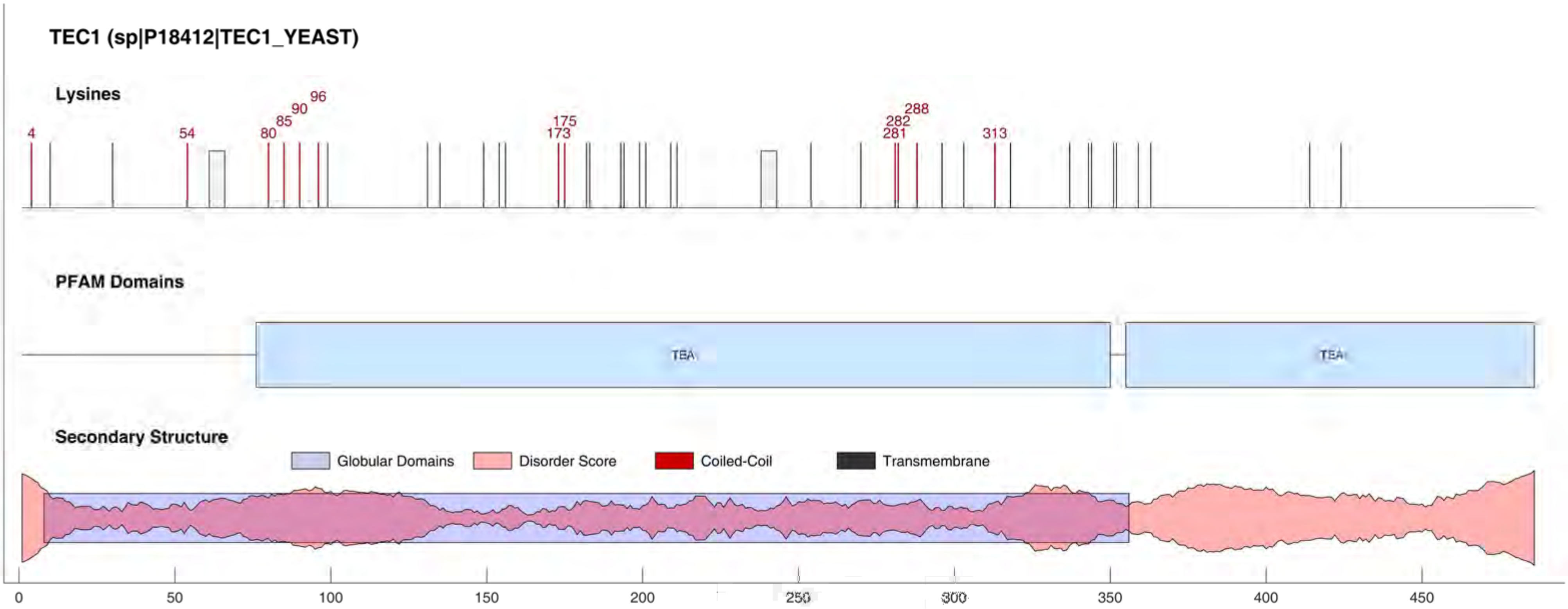
TEC1 (sp|P18412|TEC1_YEAST)
Lysines
4
54
80
85
90
96
173
175
281
282
288
313
PFAM Domains
TEA
TEA
Secondary Structure
Globular Domains
Disorder Score
Coiled-Coil
Transmembrane
0
50
100
150
200
250
300
350
400
450

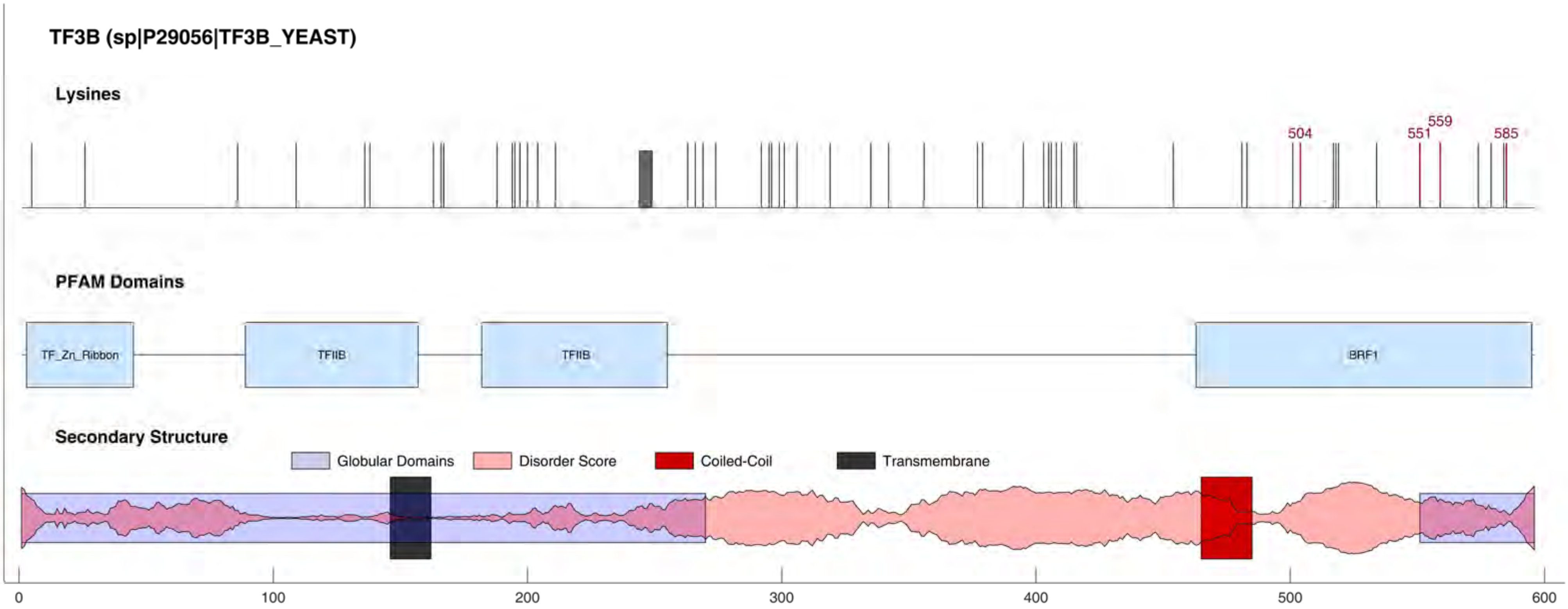

TF3B (sp|P29056|TF3B_YEAST)
Lysines
504
551
559
585
PFAM Domains
TF_Zn_Ribbon
TFIIB
TFIIB
BRF1
Secondary Structure
Globular Domains
Disorder Score
Coiled-Coil
Transmembrane
0
100
200
300
400
500
600

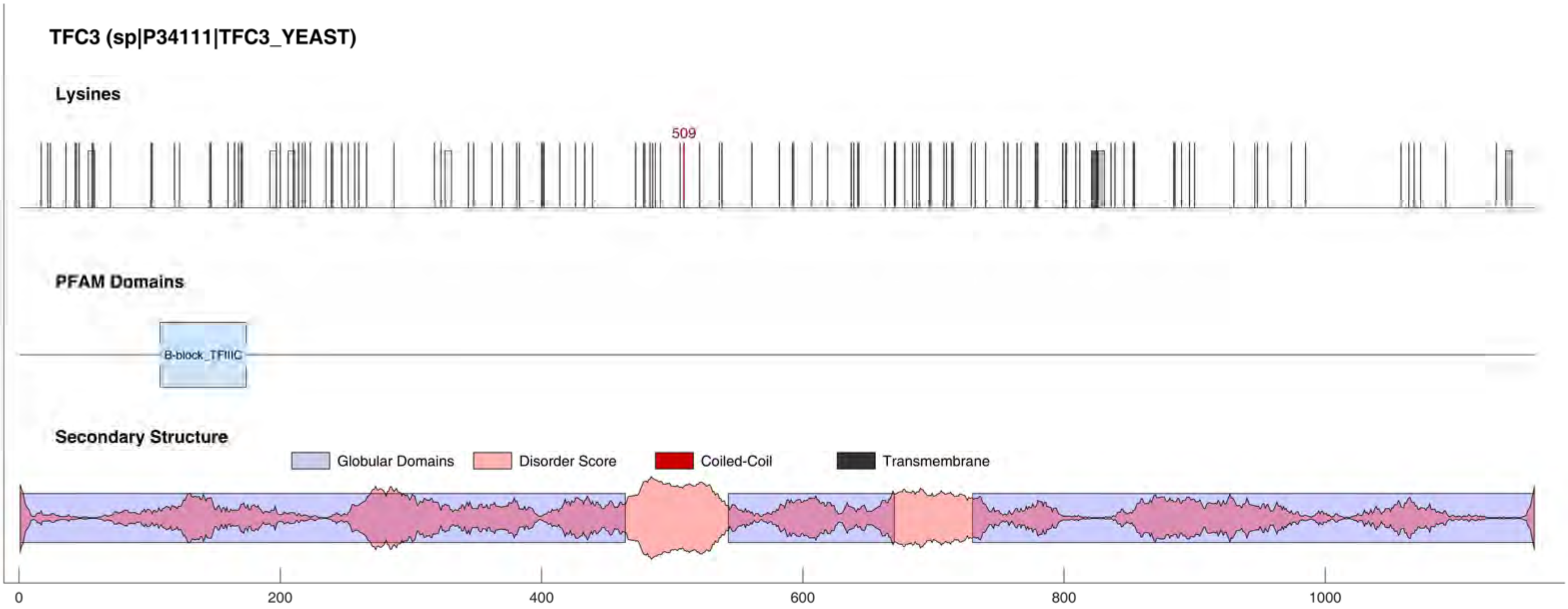
TFC3 (sp|P34111|TFC3_YEAST)
Lysines
509
PFAM Domains
B-block_TFIIIC
Secondary Structure
Globular Domains
Disorder Score
Coiled-Coil
Transmembrane
0
200
400
600
800
1000

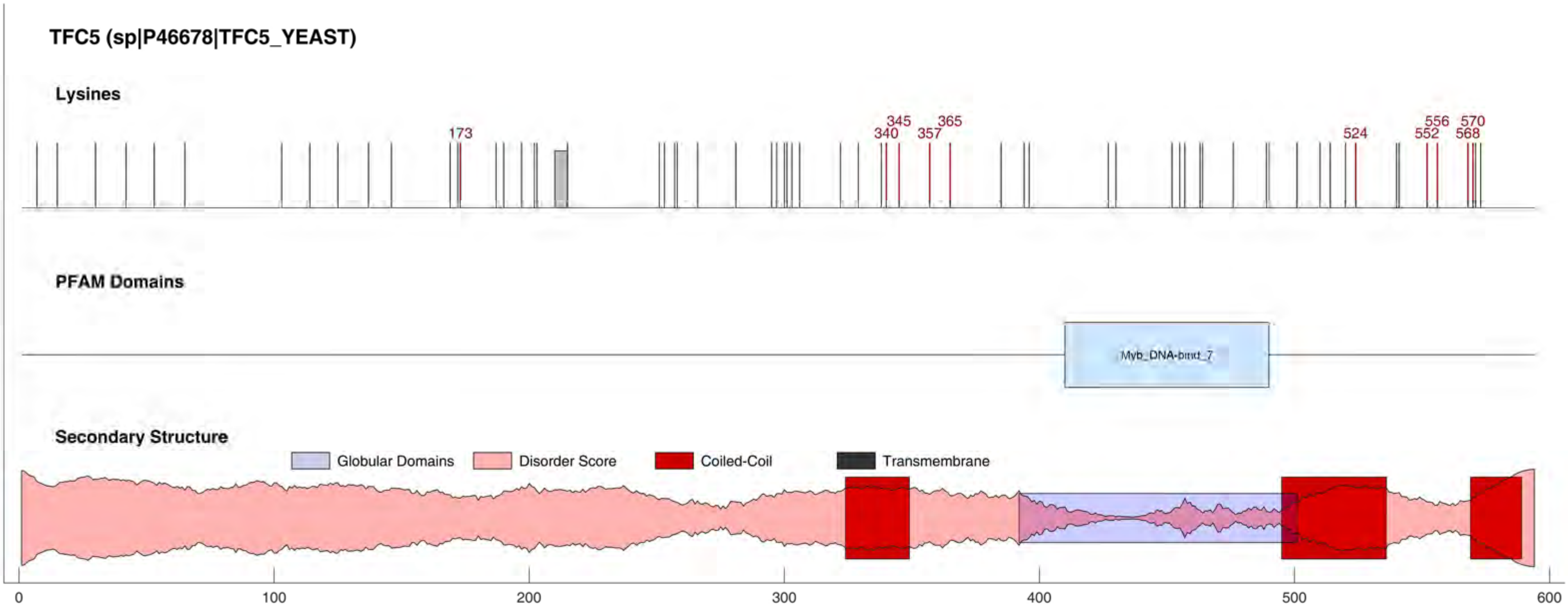
TFC5 (sp|P46678|TFC5_YEAST)
Lysines
173
340
345
357
365
524
552
556
568
570
PFAM Domains
Myb_DNA-bind_7
Secondary Structure
Globular Domains
Disorder Score
Coiled-Coil
Transmembrane
0
100
200
300
400
500
600

#### TFC7 (sp|Q12415|TFC7_YEAST)

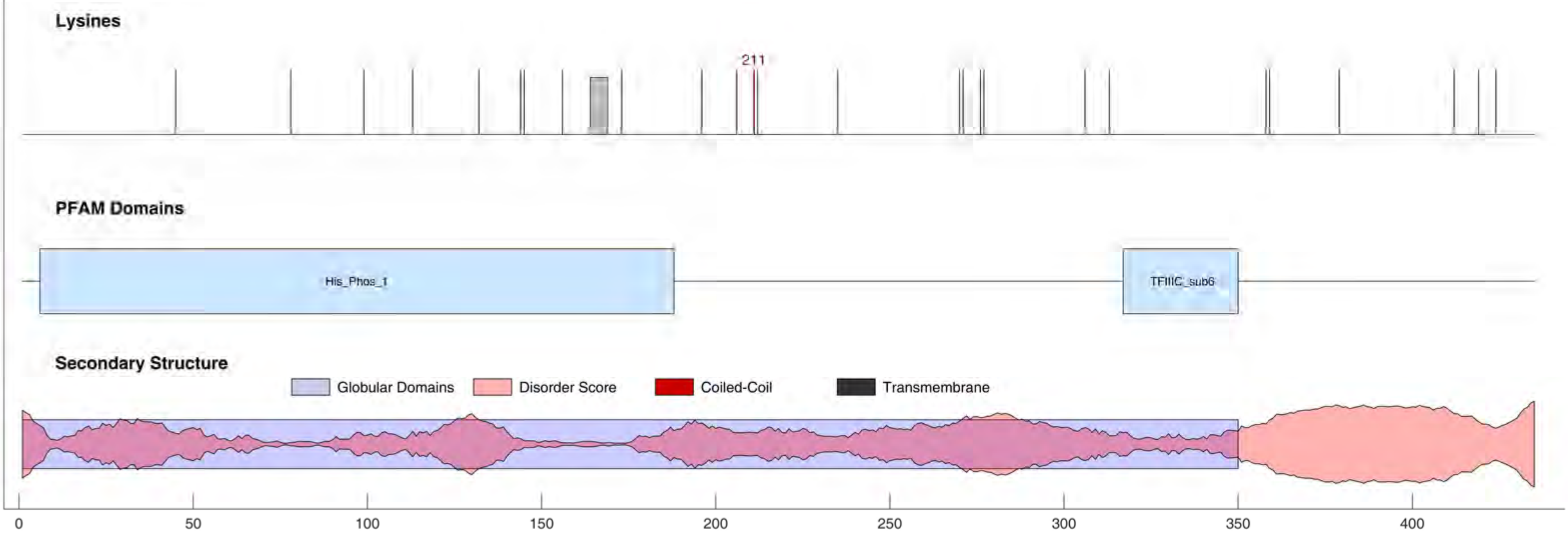

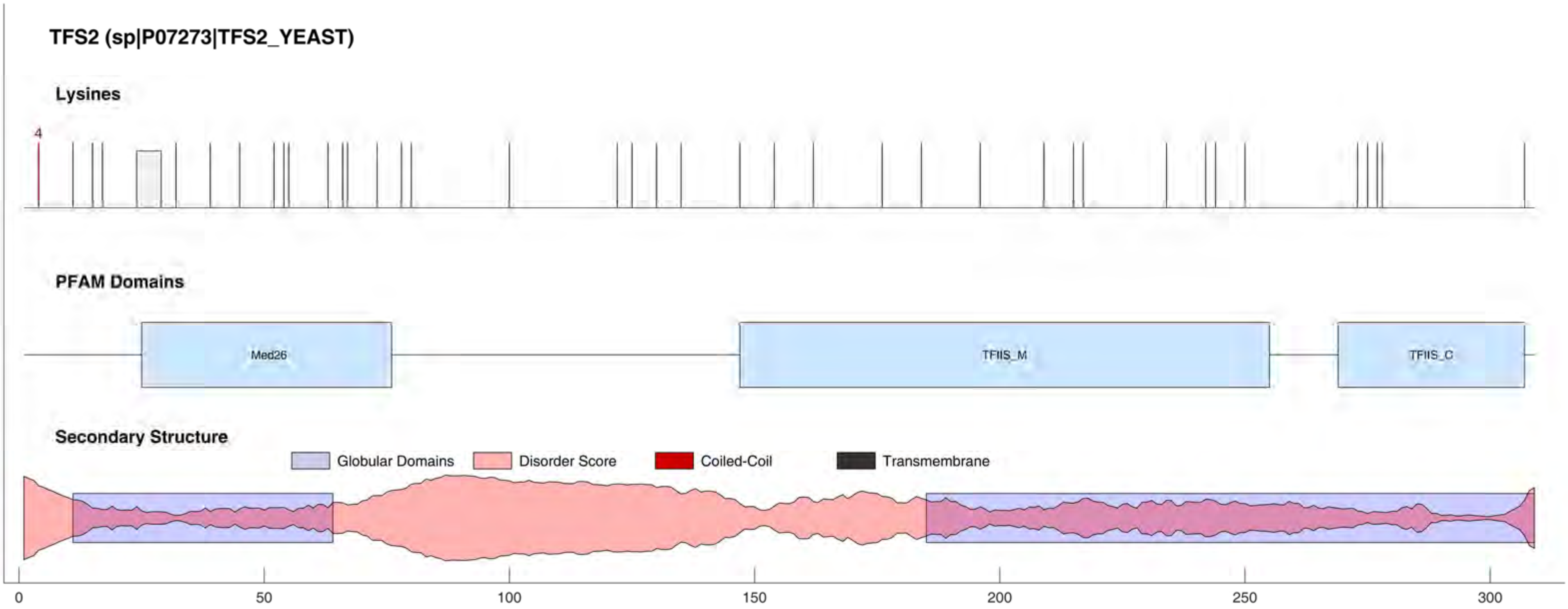
TFS2 (sp|P07273|TFS2_YEAST)
Lysines
4
PFAM Domains
Med26
TFIIS_M
TFIIS_C
Secondary Structure
Globular Domains
Disorder Score
Coiled-Coil
Transmembrane
0
50
100
150
200
250
300

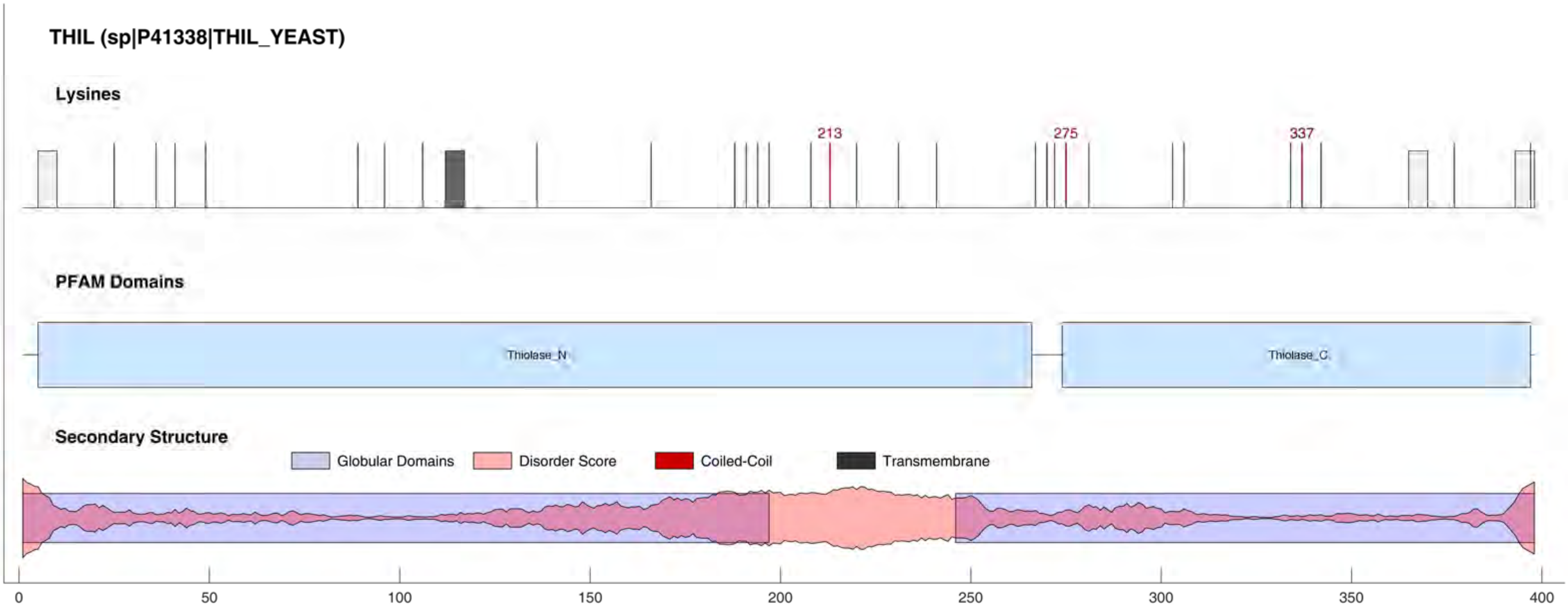

THIL (sp|P41338|THIL_YEAST)
Lysines
213
275
337
PFAM Domains
Thiolase_N
Thiolase_C
Secondary Structure
Globular Domains
Disorder Score
Coiled-Coil
Transmembrane
0
50
100
150
200
250
300
350
400

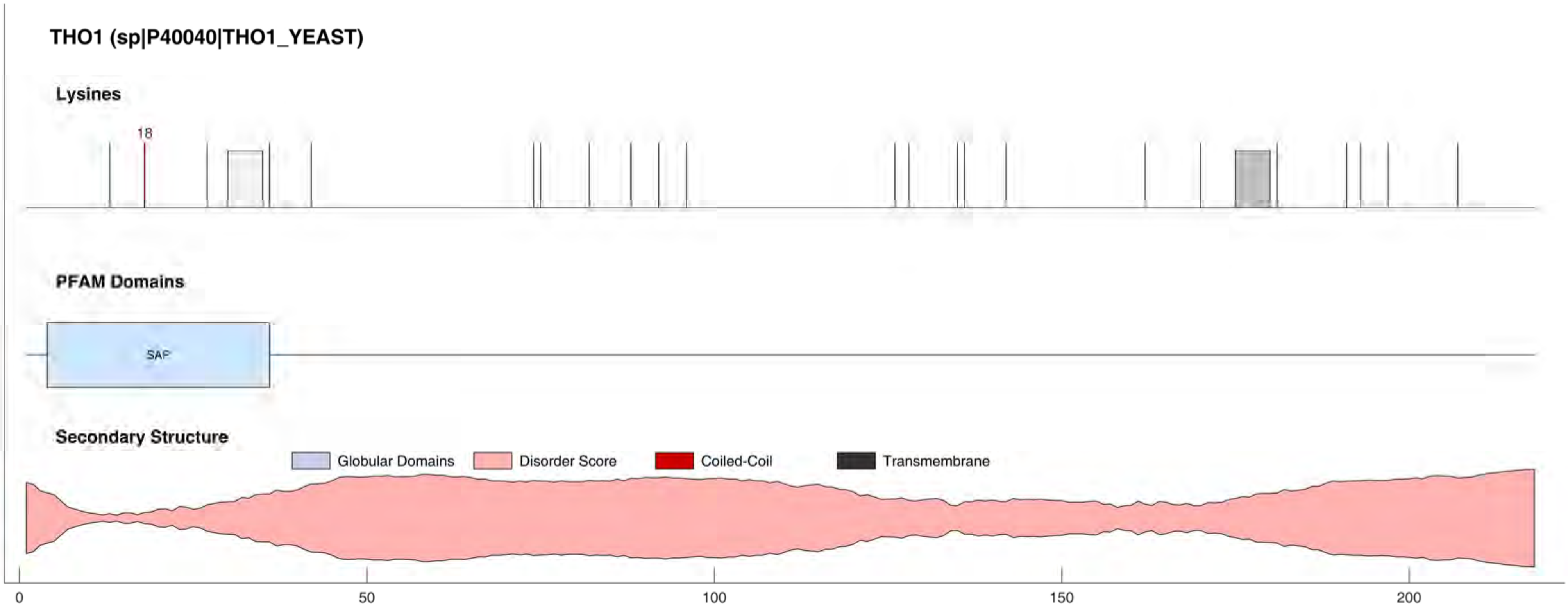

THO1 (sp|P40040|THO1_YEAST)
Lysines
18
PFAM Domains
SAP
Secondary Structure
Globular Domains
Disorder Score
Coiled-Coil
Transmembrane
0
50
100
150
200

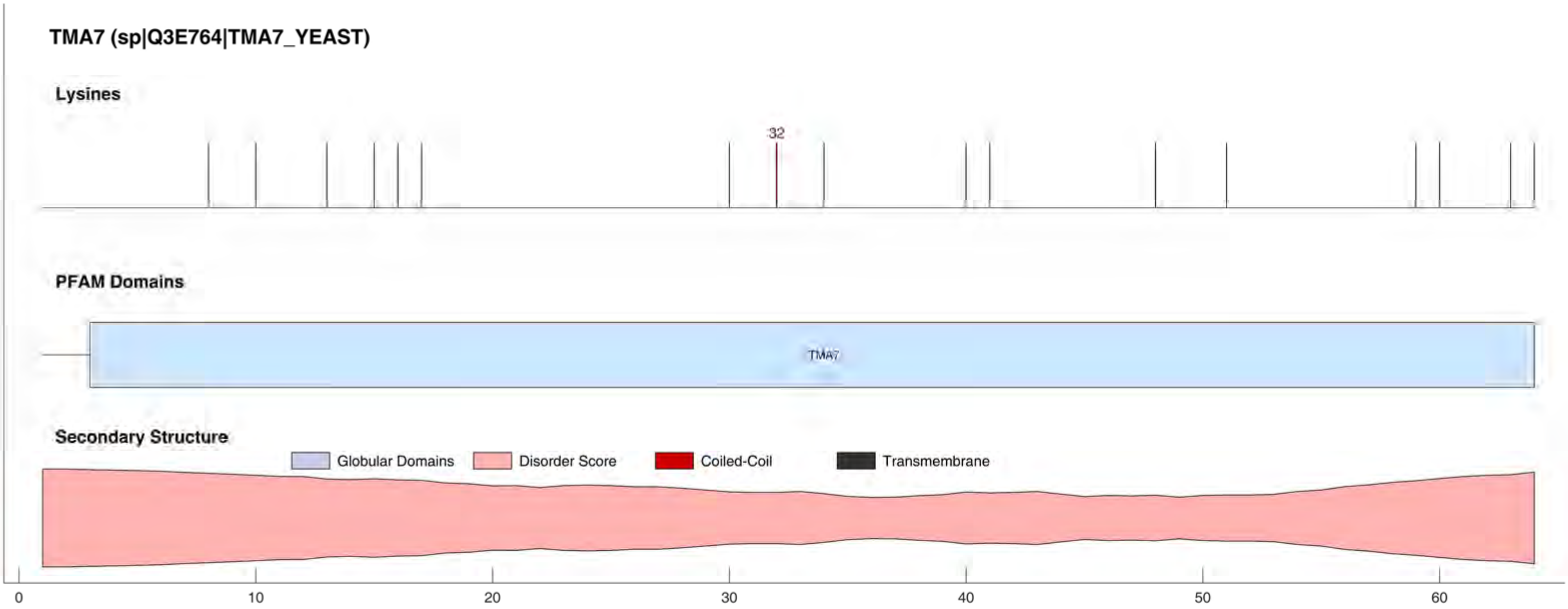
TMA7 (sp|Q3E764|TMA7_YEAST)
Lysines
32
PFAM Domains
TMA7
Secondary Structure
Globular Domains
Disorder Score
Coiled-Coil
Transmembrane
0
10
20
30
40
50
60

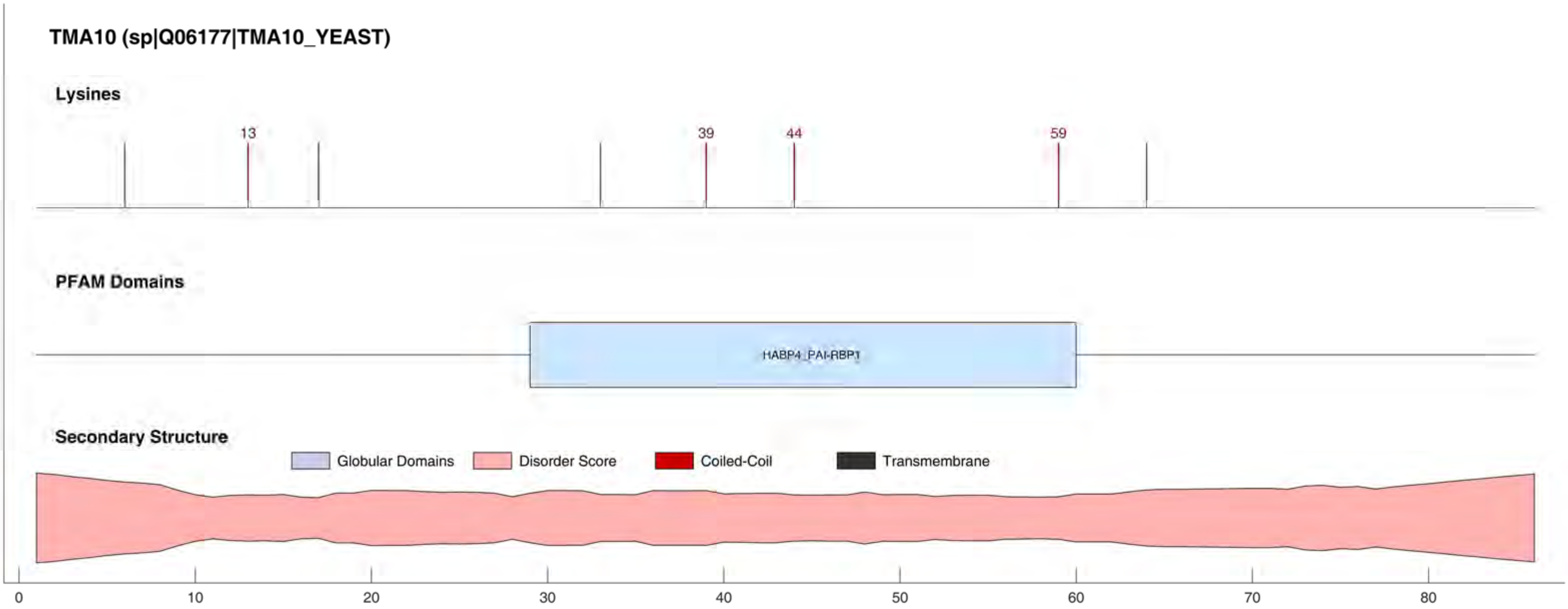
TMA10 (sp|Q06177|TMA10_YEAST)
Lysines
13
39
44
59
PFAM Domains
HABP4_PAI-RBP1
Secondary Structure
Globular Domains
Disorder Score
Coiled-Coil
Transmembrane
0
10
20
30
40
50
60
70
80

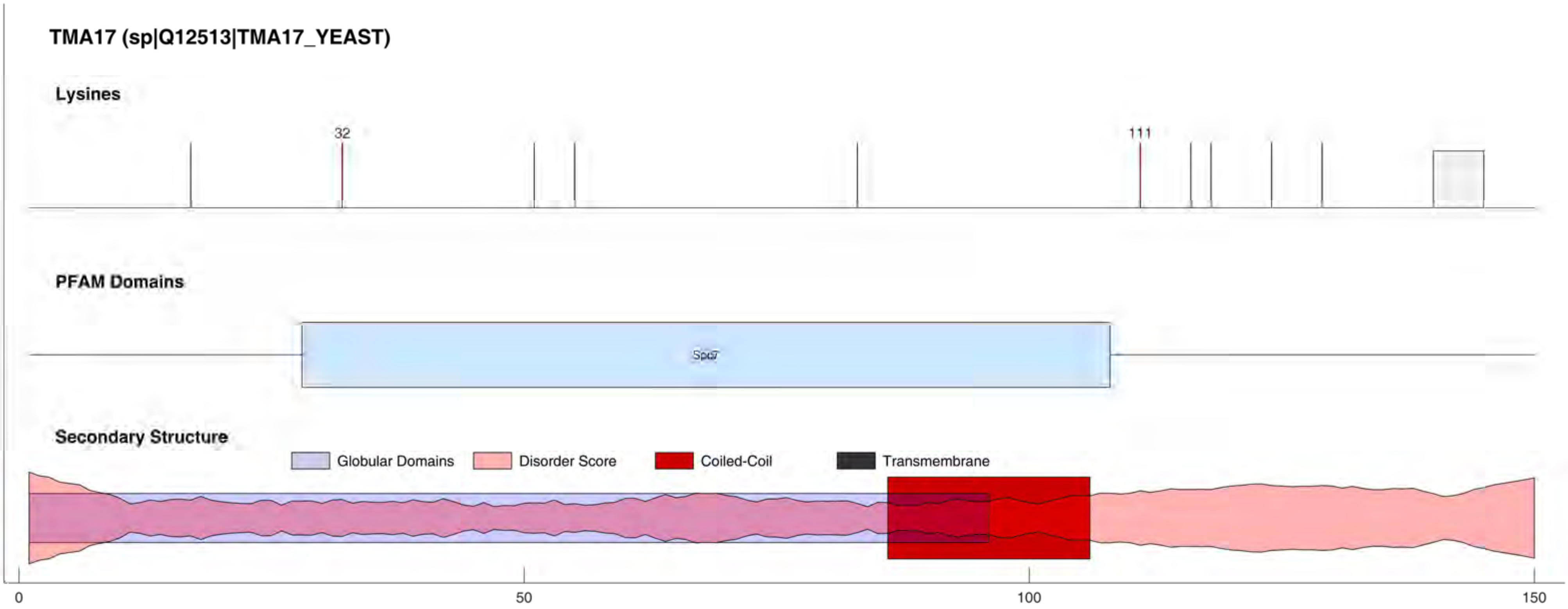

TMA17 (sp|Q12513|TMA17_YEAST)
Lysines
32
111
PFAM Domains
Secondary Structure
Globular Domains
Disorder Score
Coiled-Coil
Transmembrane
0
50
100
150

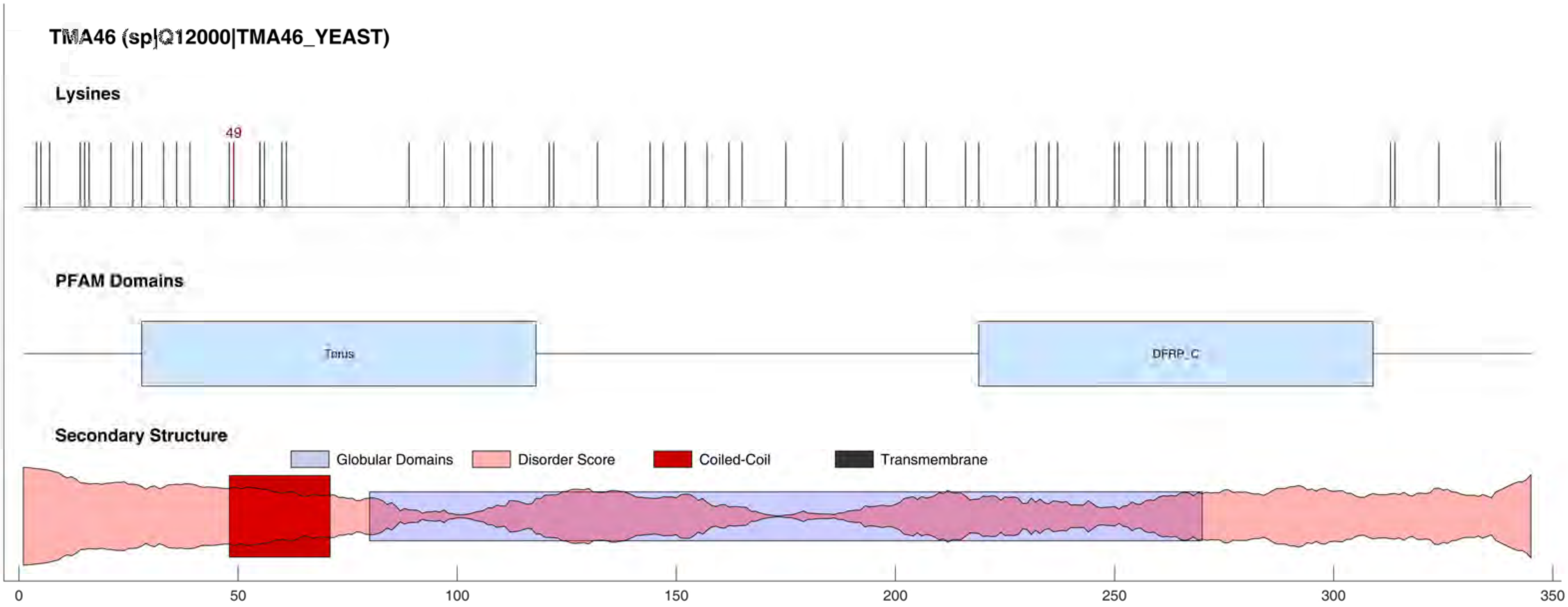

TMA46 (sp|Q12000|TMA46_YEAST)
Lysines
49
PFAM Domains
Torus
DFRP_C
Secondary Structure
Globular Domains
Disorder Score
Coiled-Coil
Transmembrane
0
50
100
150
200
250
300
350

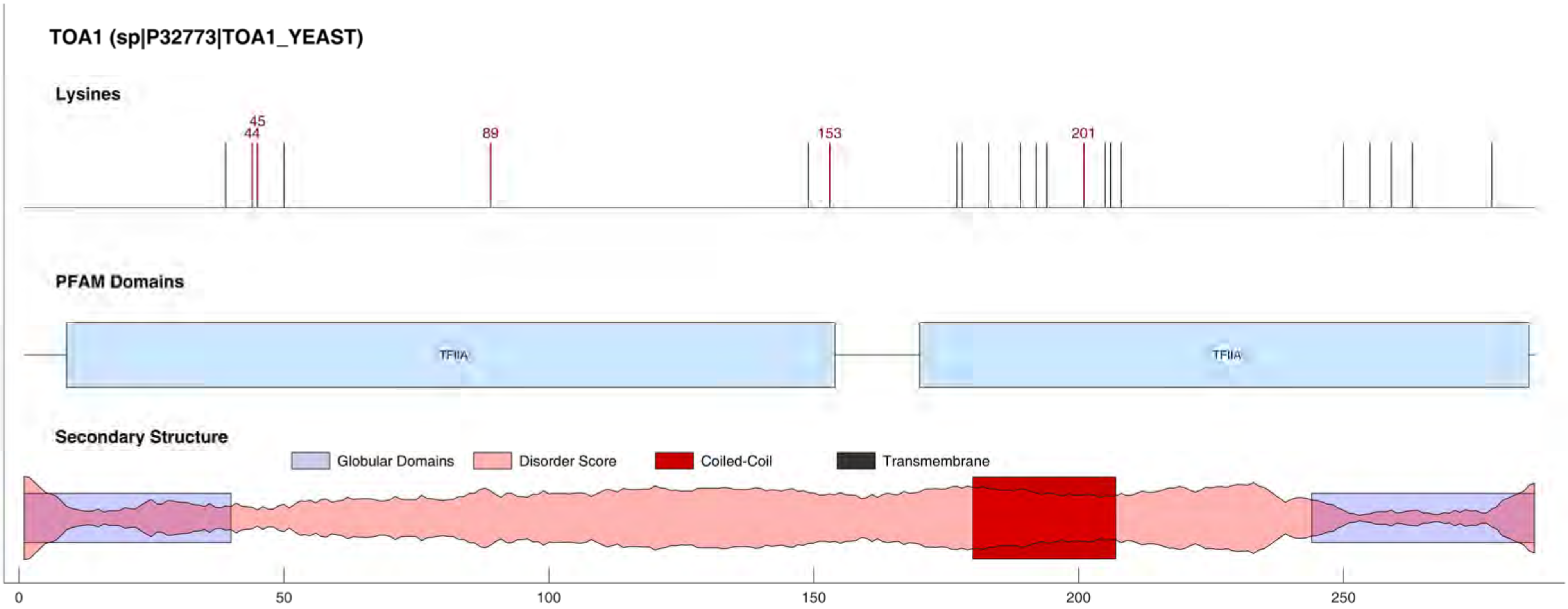

TOA1 (sp|P32773|TOA1_YEAST)
Lysines
44
45
89
153
201
PFAM Domains
TFIIA
TFIIA
Secondary Structure
Globular Domains
Disorder Score
Coiled-Coil
Transmembrane
0
50
100
150
200
250

#### TOF2 (sp|Q02208|TOF2_YEAST)

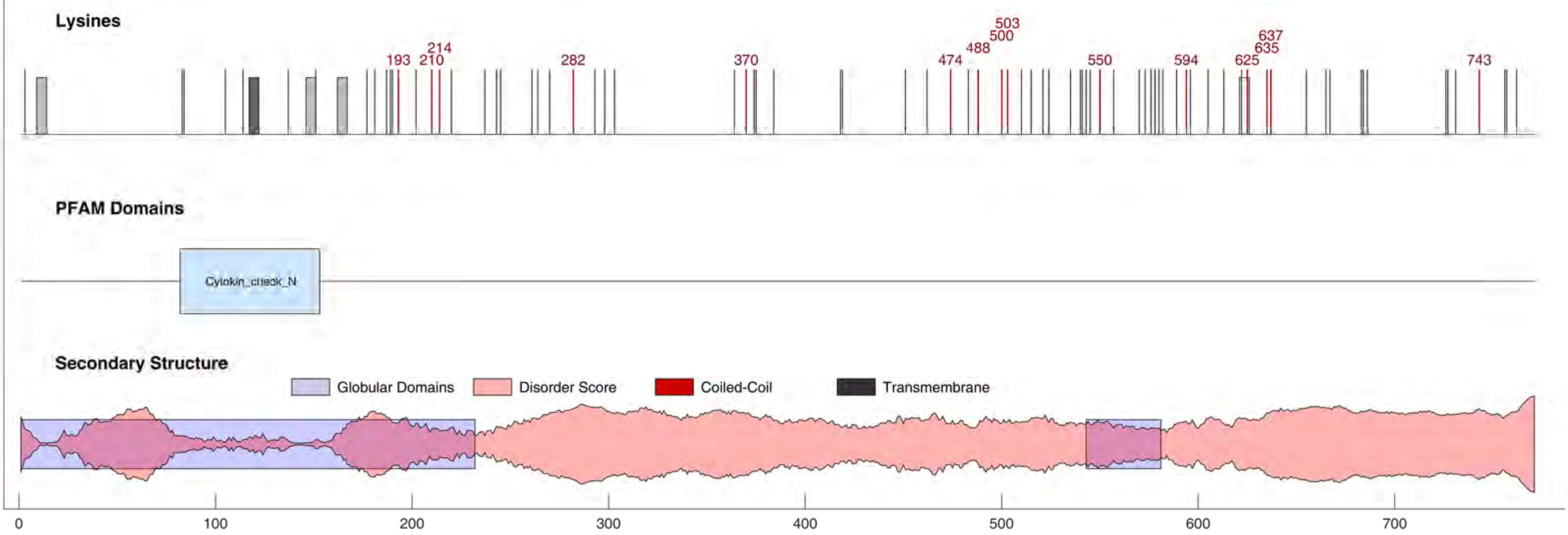

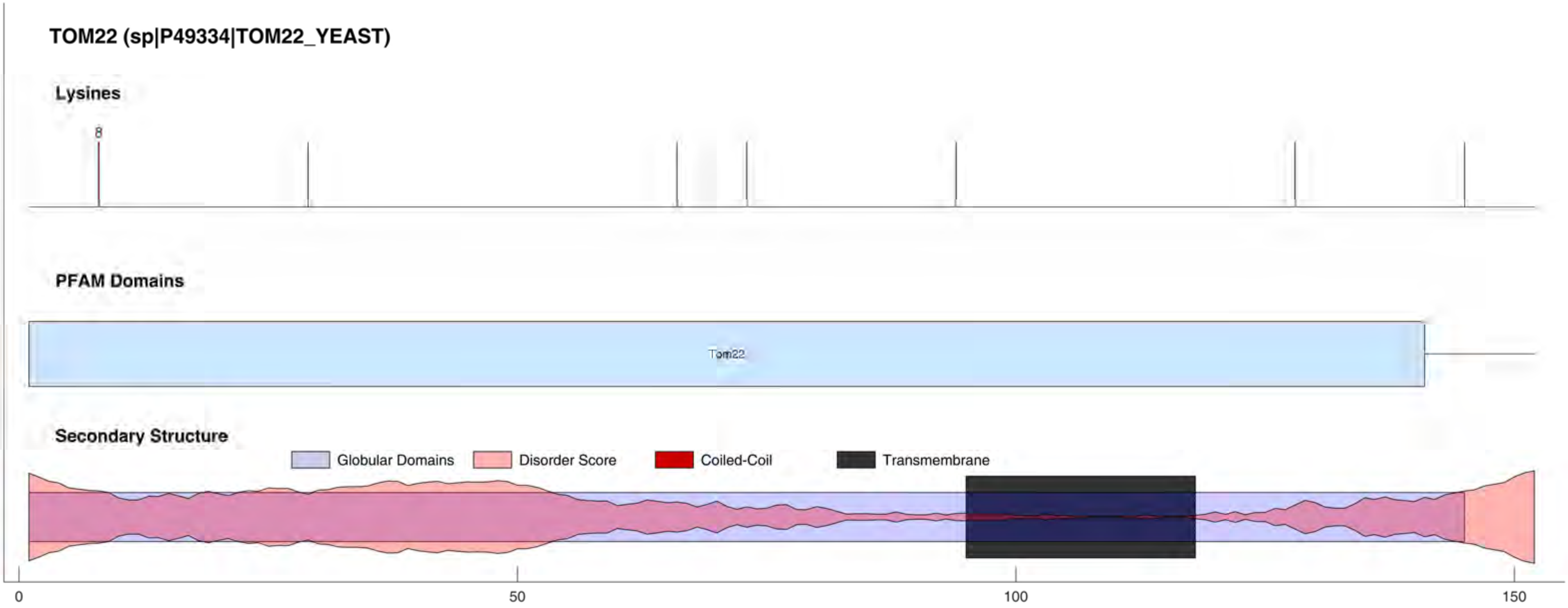
TOM22 (sp|P49334|TOM22_YEAST)
Lysines
8
PFAM Domains
Tom22
Secondary Structure
Globular Domains
Disorder Score
Coiled-Coil
Transmembrane
0
50
100
150

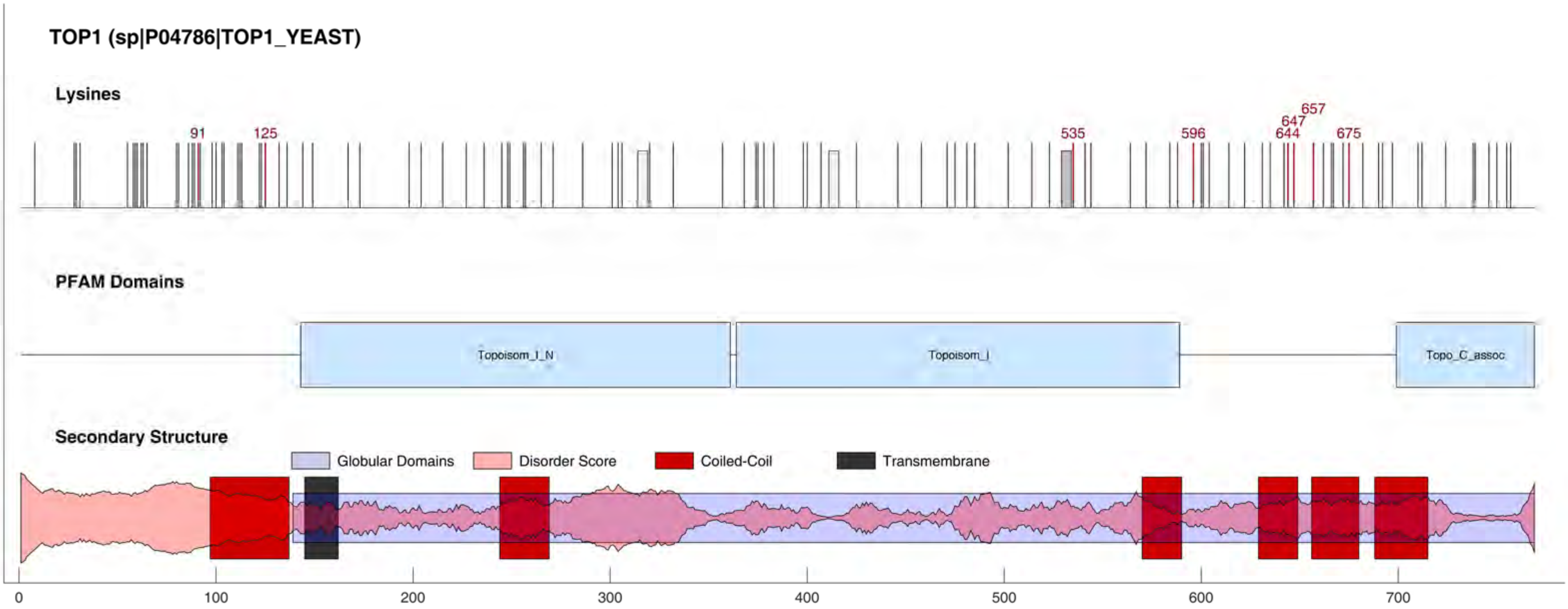

TOP1 (sp|P04786|TOP1_YEAST)
Lysines
91
125
535
596
657
647
644
675
PFAM Domains
Topoisom_I_N
Topoisom_I
Topo_C_assoc
Secondary Structure
Globular Domains
Disorder Score
Coiled-Coil
Transmembrane
0
100
200
300
400
500
600
700

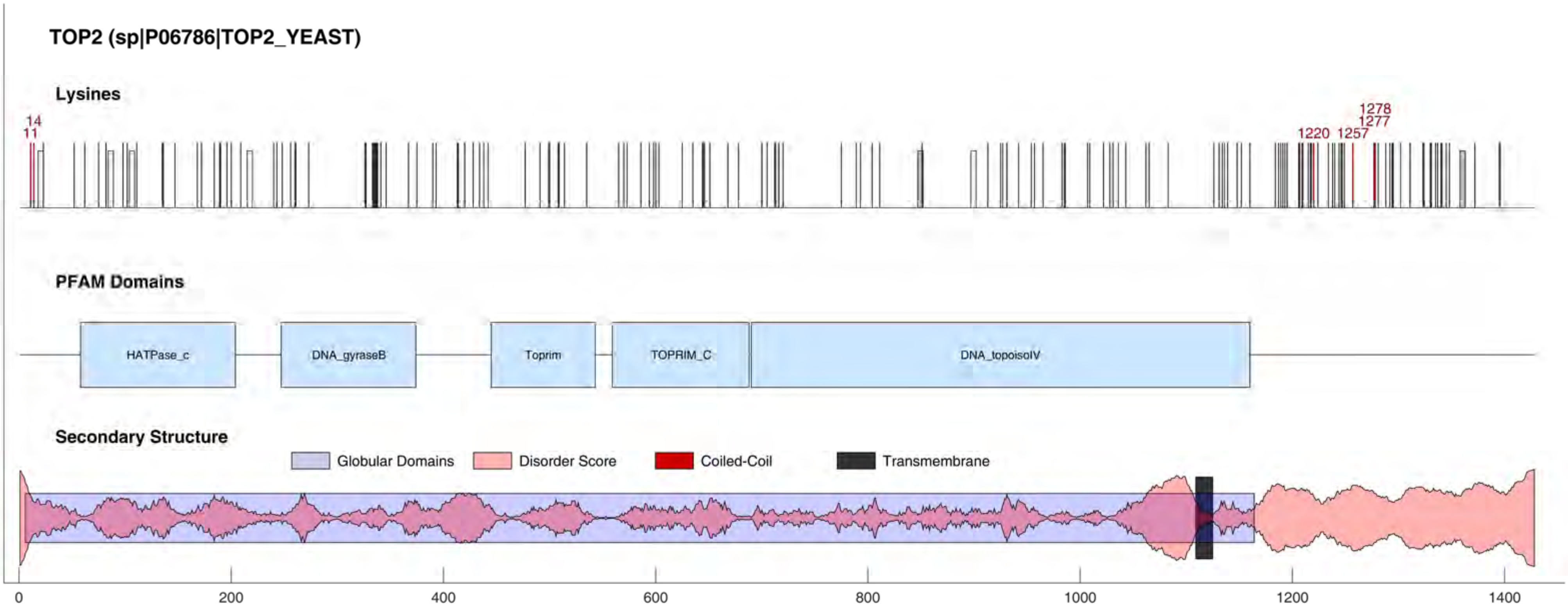

TOP2 (sp|P06786|TOP2_YEAST)
Lysines
14
11
1220
1257
1277
1278
PFAM Domains
HATPase_c
DNA_gyraseB
Toprim
TOPRIM_C
DNA_topoisoIV
Secondary Structure
Globular Domains
Disorder Score
Coiled-Coil
Transmembrane
0
200
400
600
800
1000
1200
1400

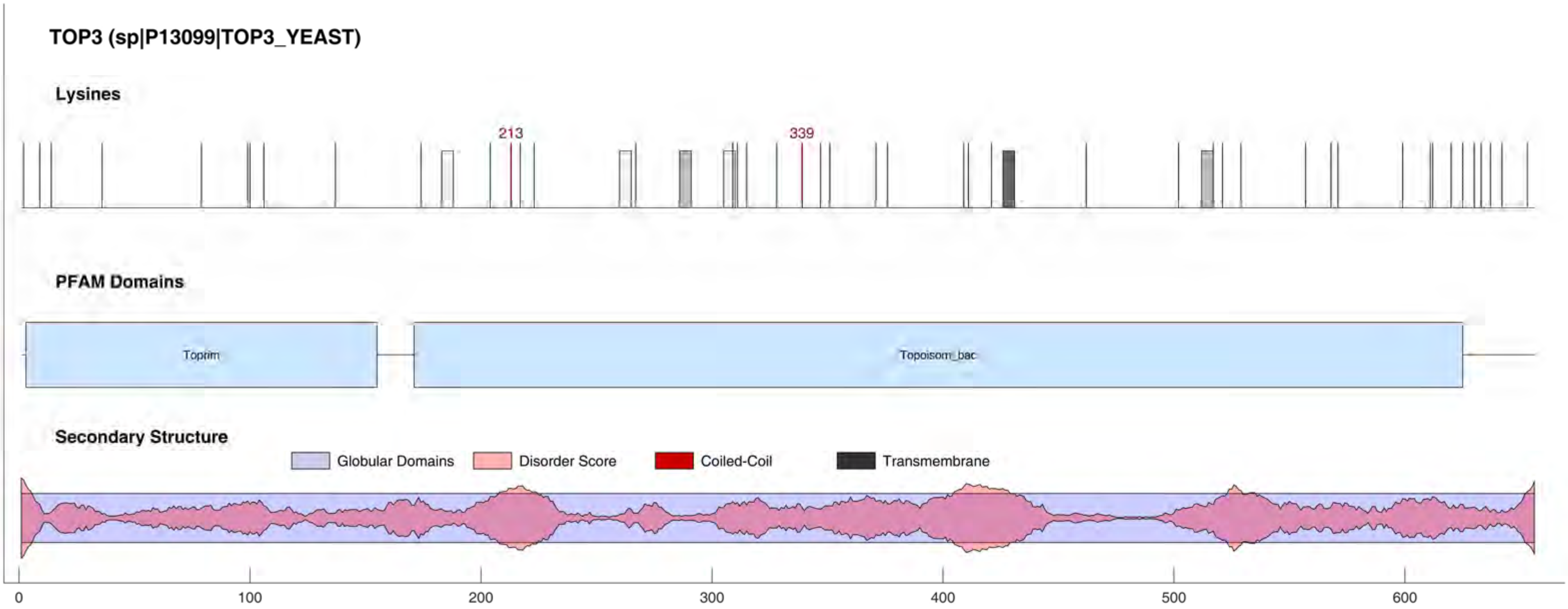

TOP3 (sp|P13099|TOP3_YEAST)
Lysines
213
339
PFAM Domains
Toprim
Topoisom_bac
Secondary Structure
Globular Domains
Disorder Score
Coiled-Coil
Transmembrane
0
100
200
300
400
500
600

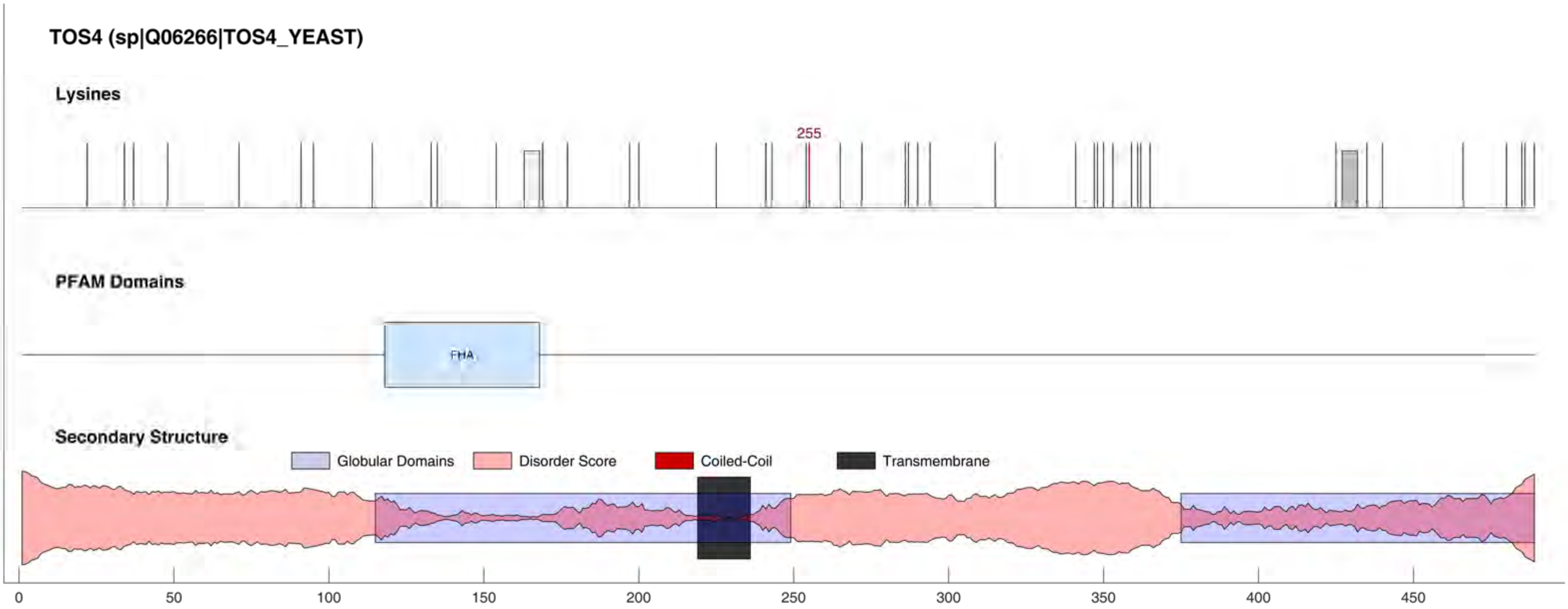

TOS4 (sp|Q06266|TOS4_YEAST)
Lysines
255
PFAM Domains
FHA
Secondary Structure
Globular Domains
Disorder Score
Coiled-Coil
Transmembrane
0
50
100
150
200
250
300
350
400
450

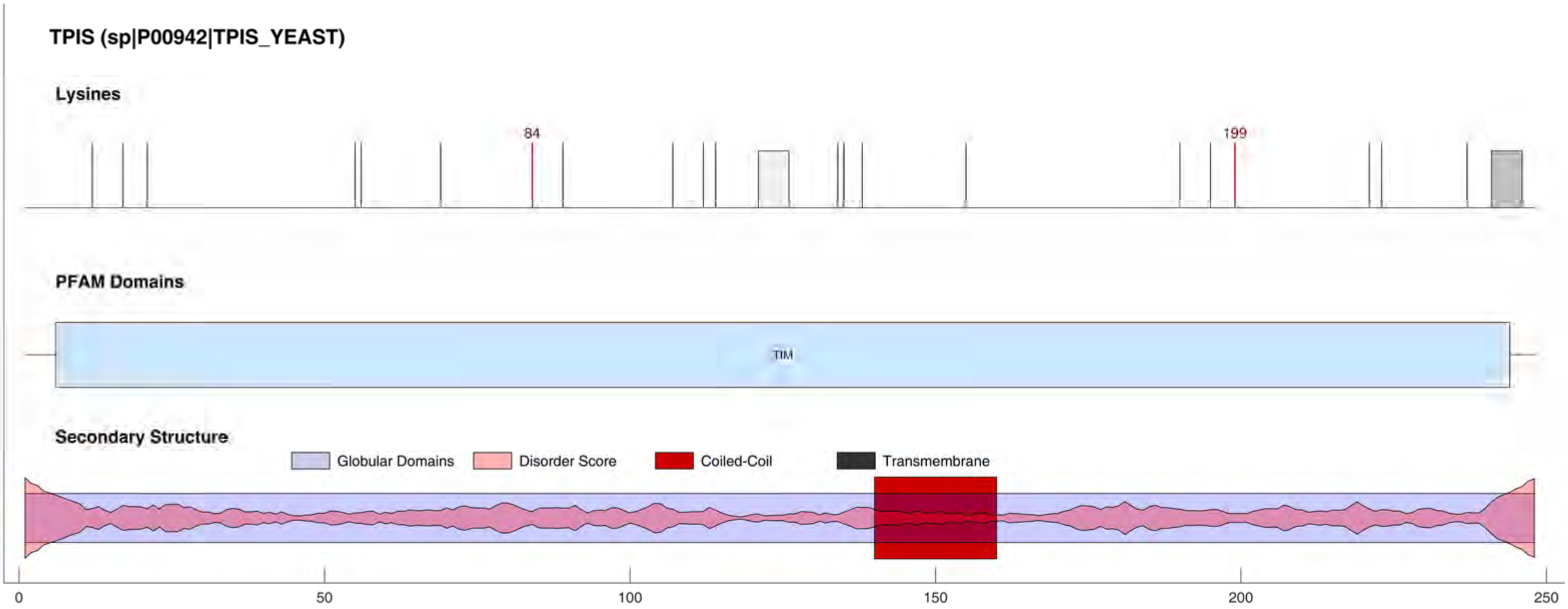
TPIS (sp|P00942|TPIS_YEAST)
Lysines
84
199
PFAM Domains
TIM
Secondary Structure
Globular Domains
Disorder Score
Coiled-Coil
Transmembrane
0
50
100
150
200
250

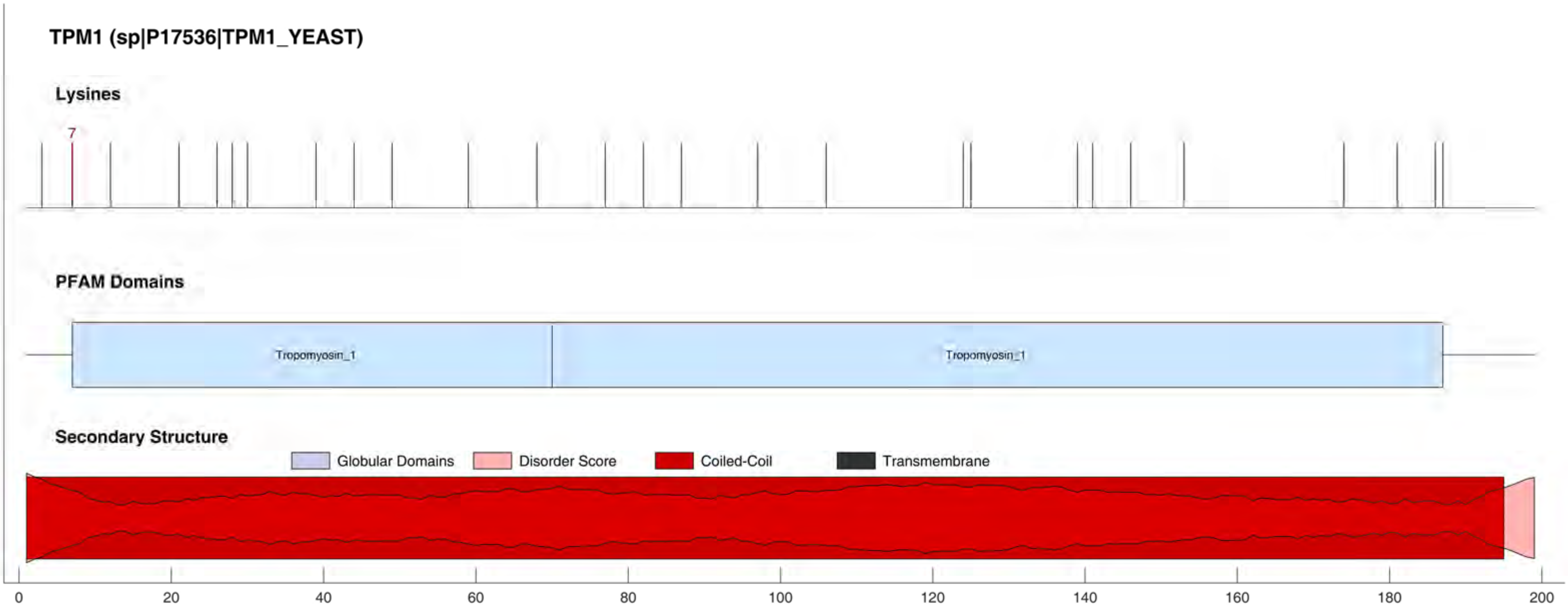

TPM1 (sp|P17536|TPM1_YEAST)
Lysines
7
PFAM Domains
Tropomyosin_1
Tropomyosin_1
Secondary Structure
Globular Domains
Disorder Score
Coiled-Coil
Transmembrane
0
20
40
60
80
100
120
140
160
180
200

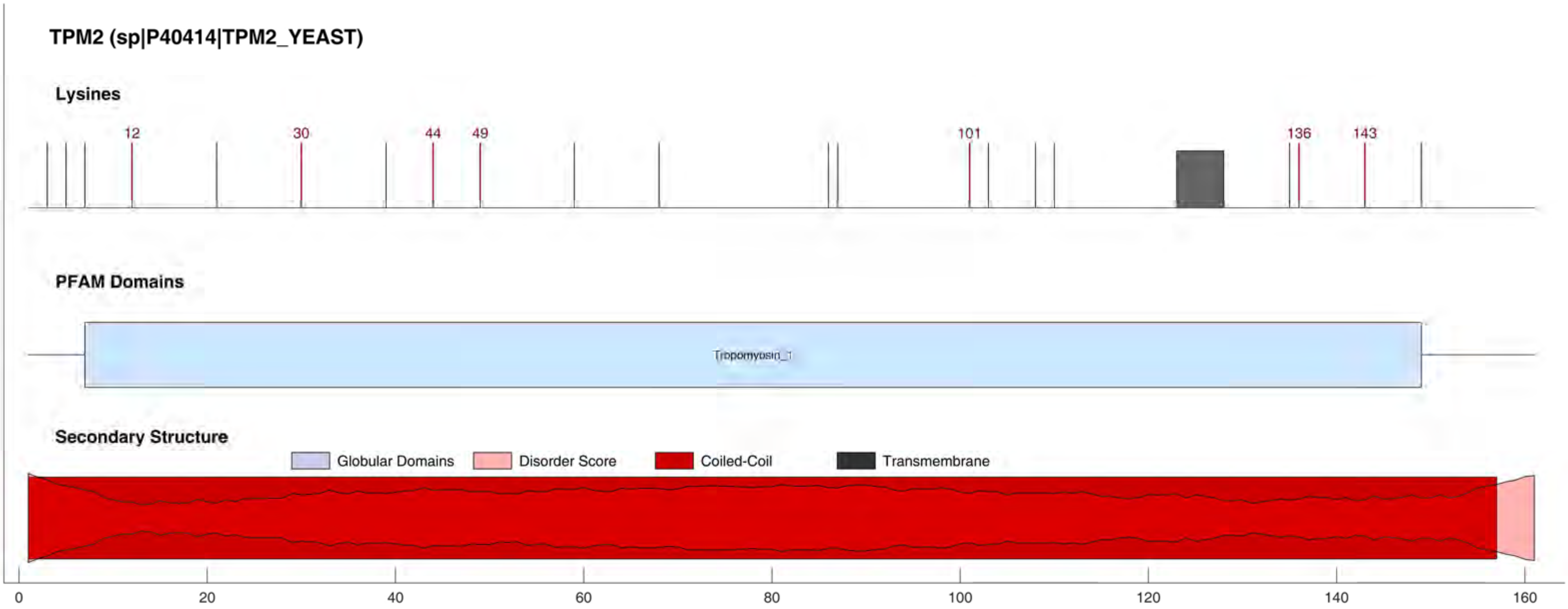

TPM2 (sp|P40414|TPM2_YEAST)
Lysines
12
30
44
49
101
136
143
PFAM Domains
Tropomyosin_1
Secondary Structure
Globular Domains
Disorder Score
Coiled-Coil
Transmembrane
0
20
40
60
80
100
120
140
160

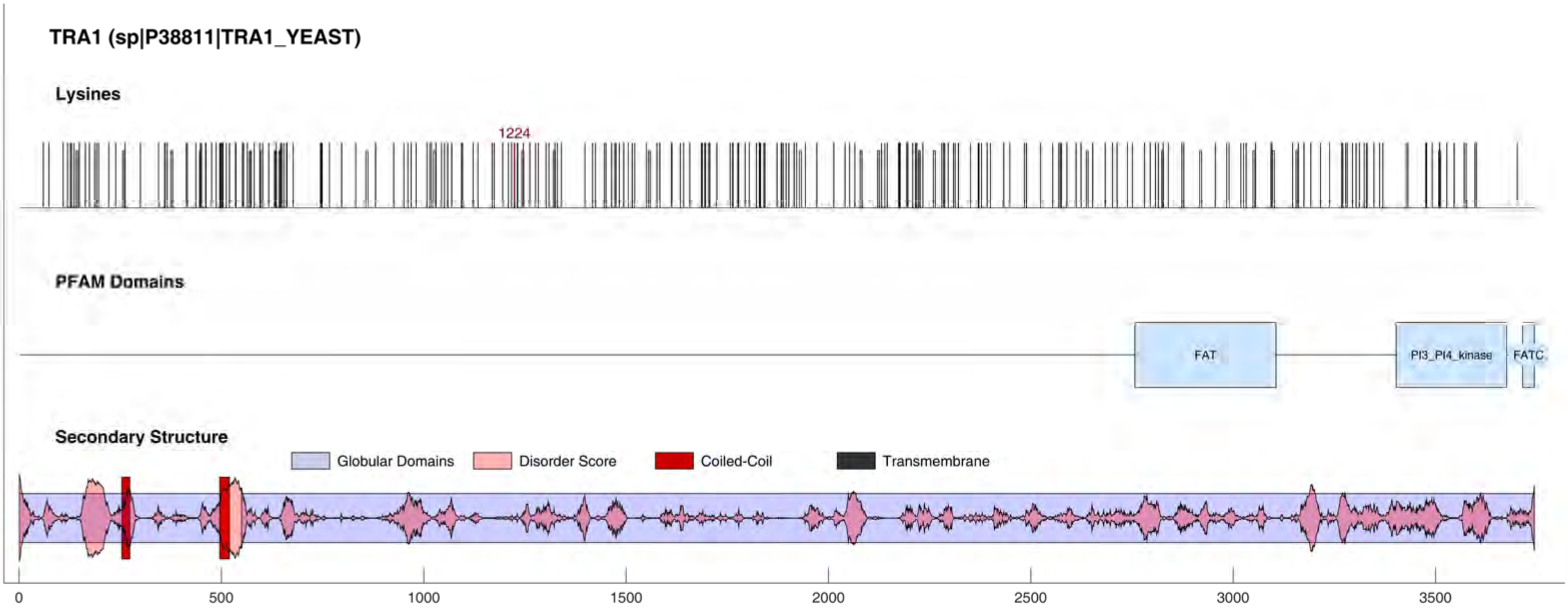
TRA1 (sp|P38811|TRA1_YEAST)
Lysines
1224
PFAM Domains
FAT
PI3_PI4_kinase
FATC
Secondary Structure
Globular Domains
Disorder Score
Coiled-Coil
Transmembrane
0
500
1000
1500
2000
2500
3000
3500

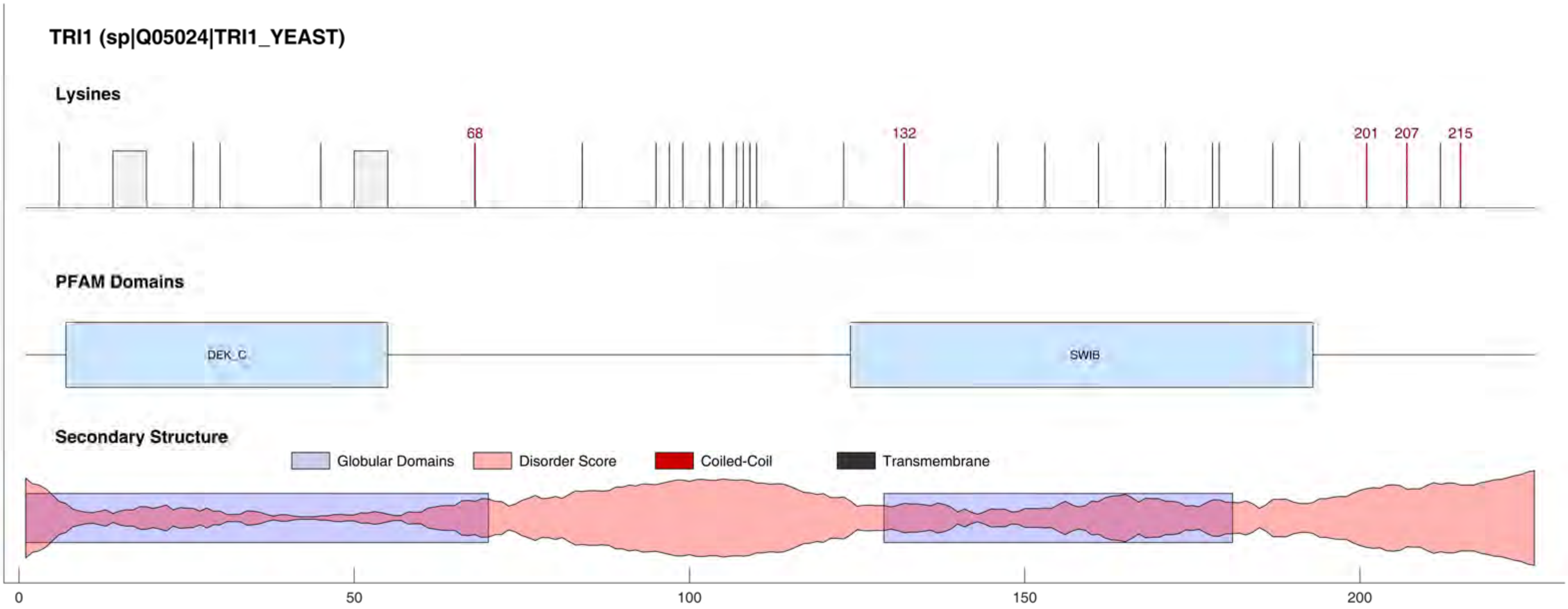
TRI1 (sp|Q05024|TRI1_YEAST)
Lysines
68
132
201
207
215
PFAM Domains
DEK_C
SWIB
Secondary Structure
Globular Domains
Disorder Score
Coiled-Coil
Transmembrane
0
50
100
150
200

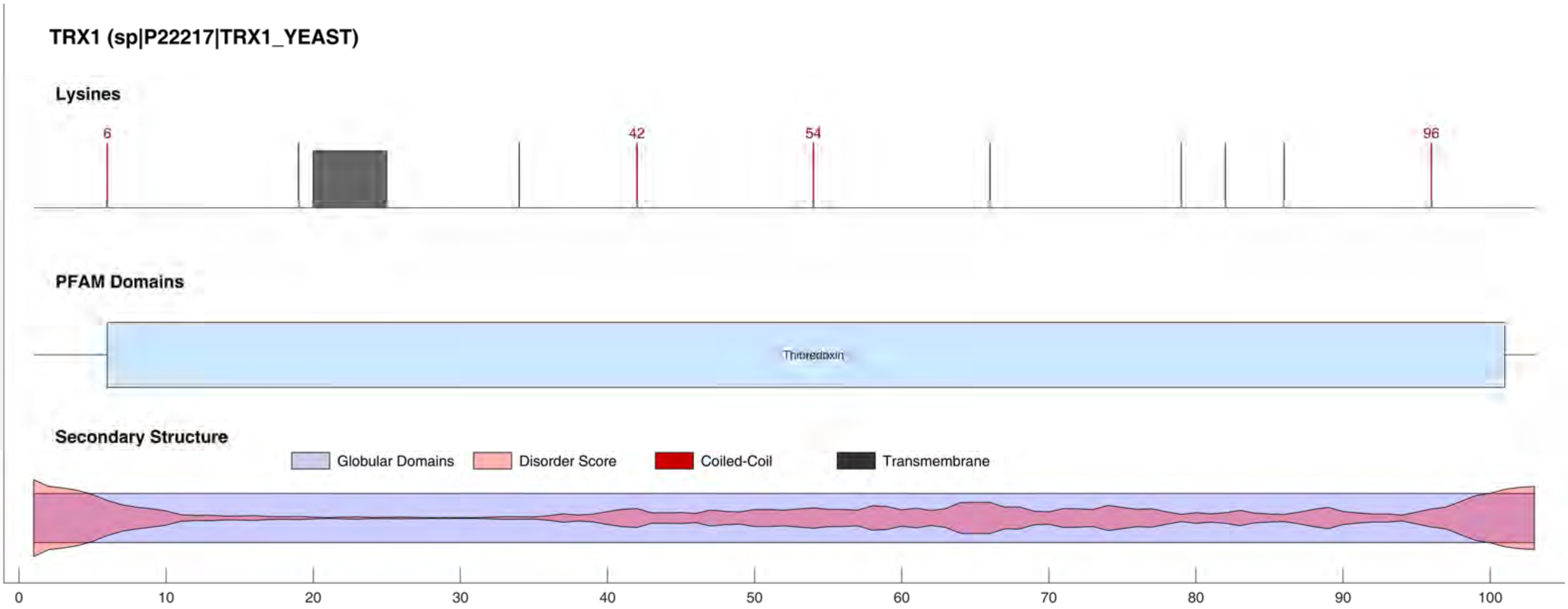

TRX1 (sp|P22217|TRX1_YEAST)
Lysines
6
42
54
96
PFAM Domains
Thioredoxin
Secondary Structure
Globular Domains
Disorder Score
Coiled-Coil
Transmembrane
0
10
20
30
40
50
60
70
80
90
100

#### TRX2 (sp|P22803|TRX2_YEAST)

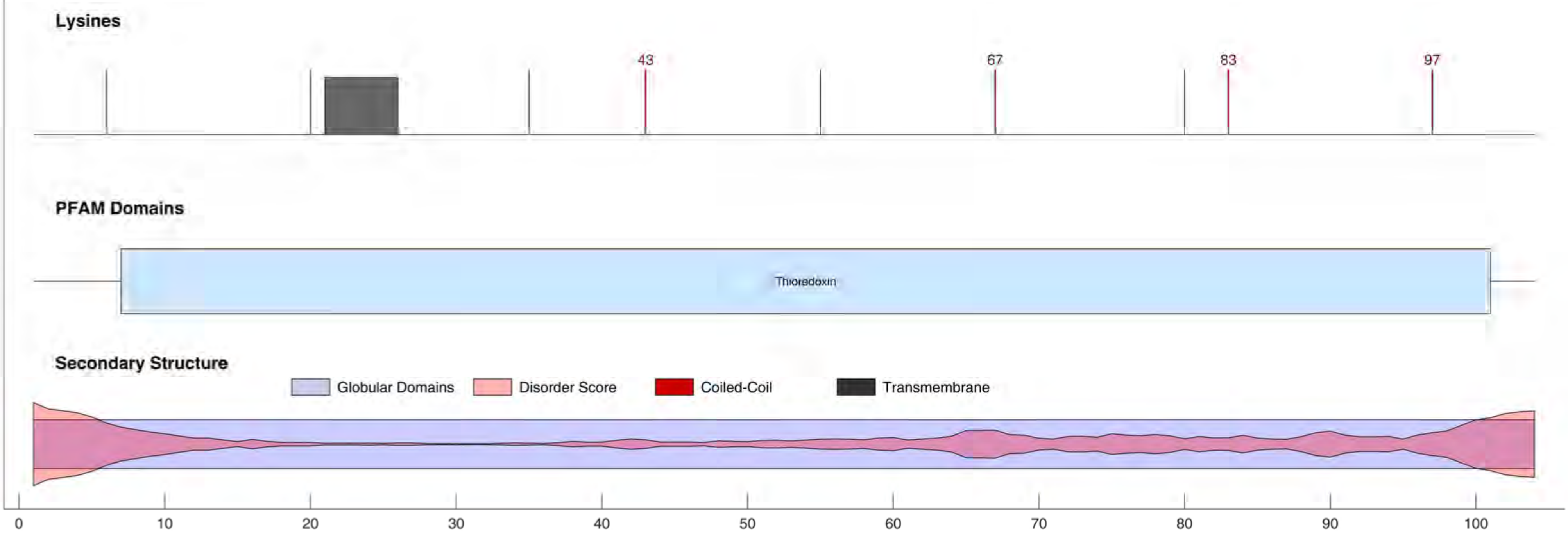

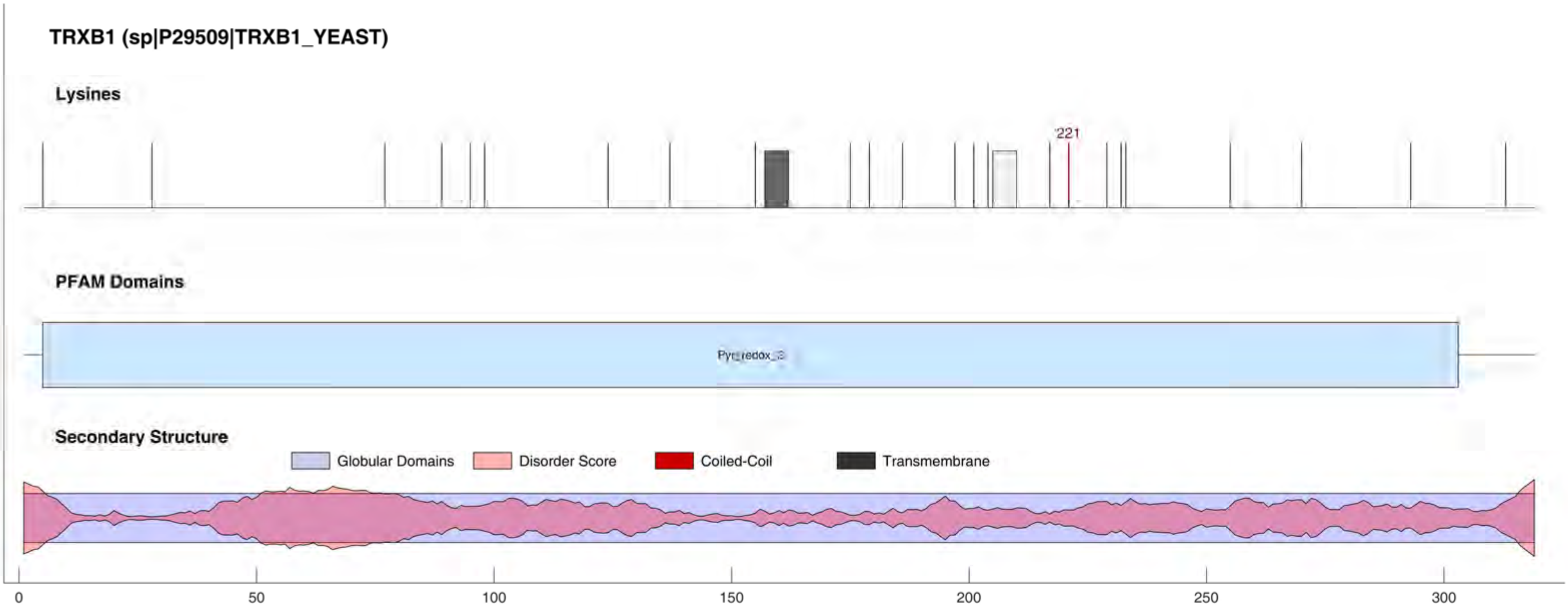
TRXB1 (sp|P29509|TRXB1_YEAST)
Lysines
221
PFAM Domains
Pyr_redox_2
Secondary Structure
Globular Domains
Disorder Score
Coiled-Coil
Transmembrane
0
50
100
150
200
250
300

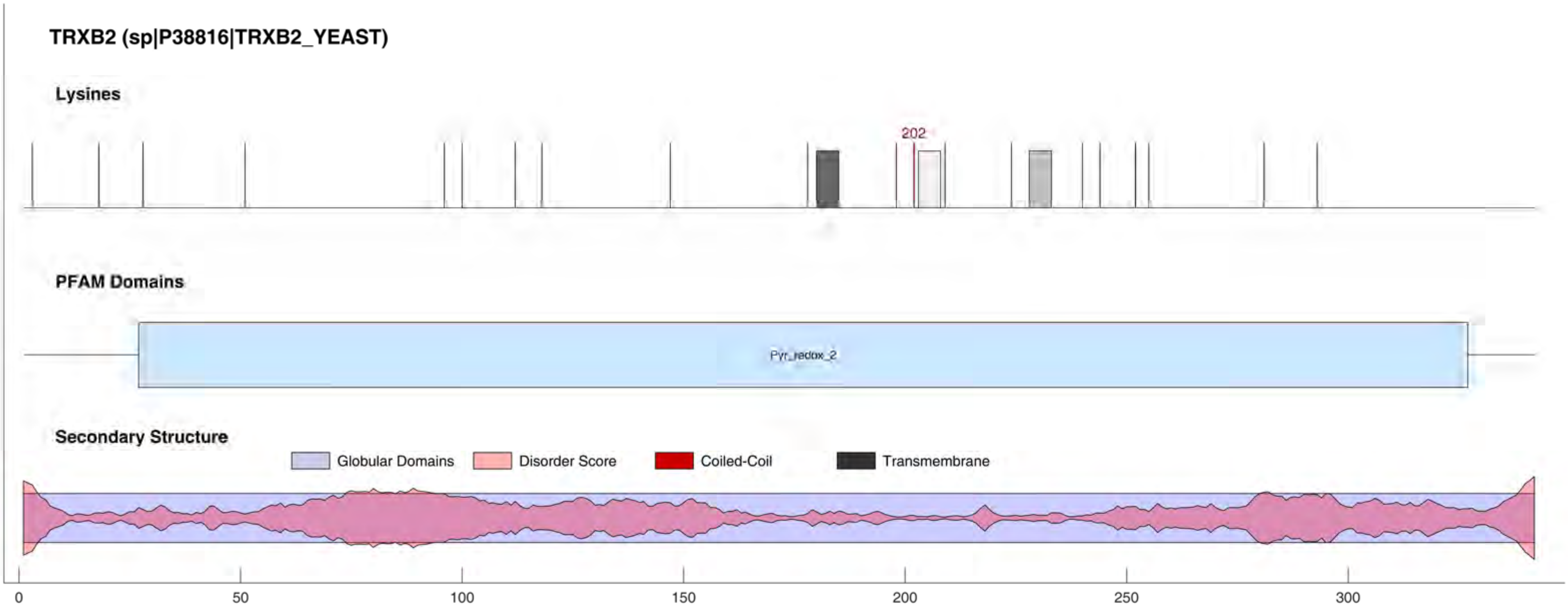
TRXB2 (sp|P38816|TRXB2_YEAST)
Lysines
202
PFAM Domains
Pyr_redox_2
Secondary Structure
Globular Domains
Disorder Score
Coiled-Coil
Transmembrane
0
50
100
150
200
250
300

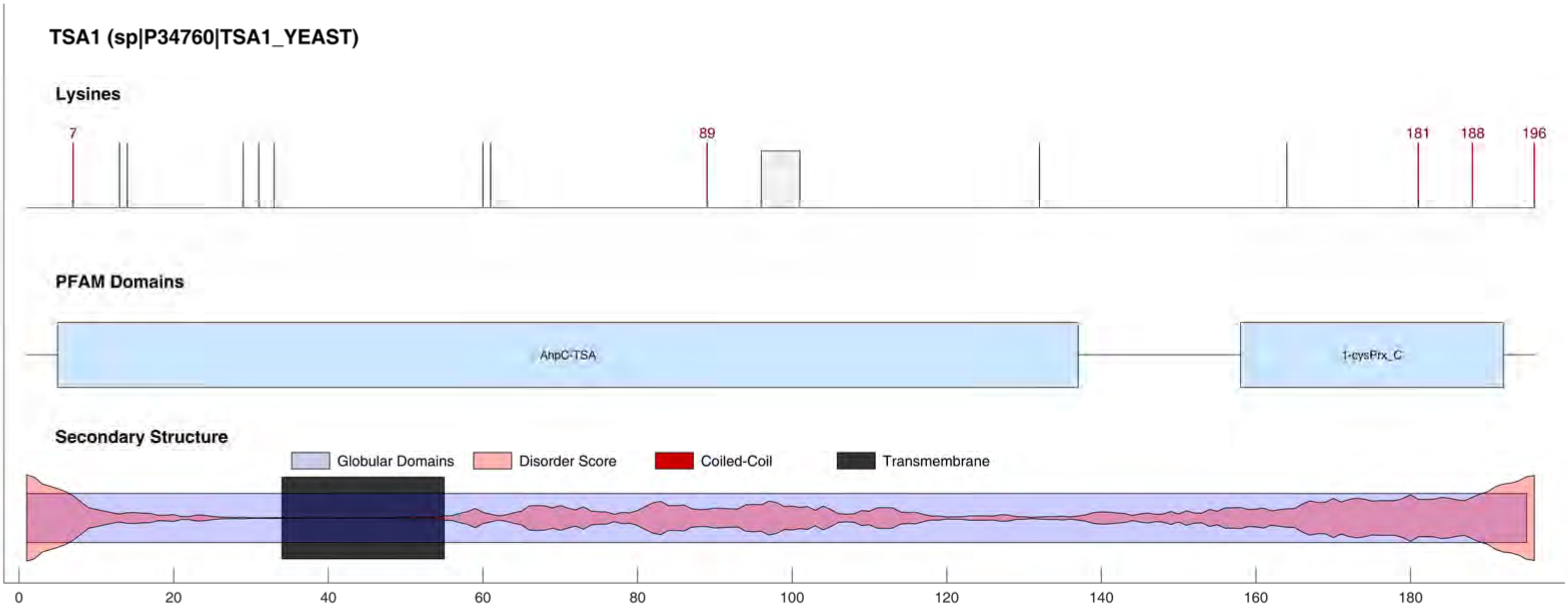
TSA1 (sp|P34760|TSA1_YEAST)
Lysines
7
89
181
188
196
PFAM Domains
AhpC-TSA
1-cysPrx_C
Secondary Structure
Globular Domains
Disorder Score
Coiled-Coil
Transmembrane
0
20
40
60
80
100
120
140
160
180

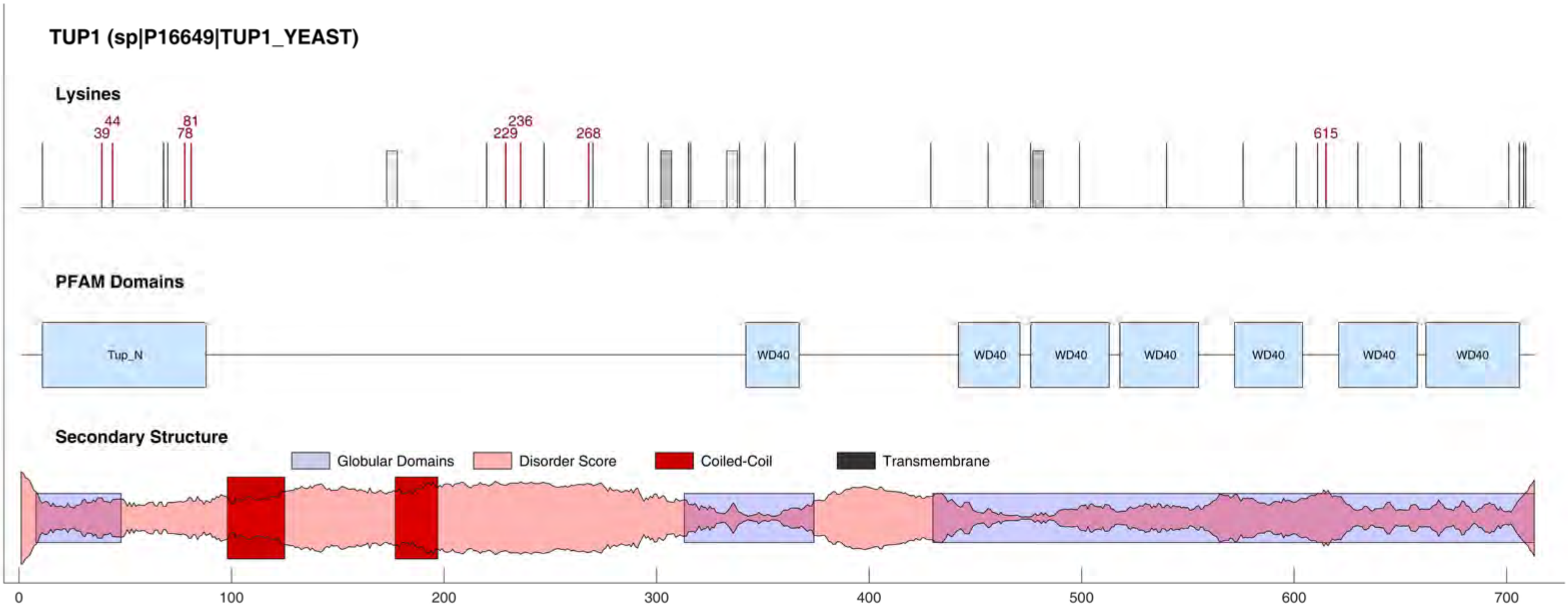
TUP1 (sp|P16649|TUP1_YEAST)
Lysines
39
44
78
81
229
236
268
615
PFAM Domains
Tup_N
WD40
WD40
WD40
WD40
WD40
WD40
WD40
Secondary Structure
Globular Domains
Disorder Score
Coiled-Coil
Transmembrane
0
100
200
300
400
500
600
700

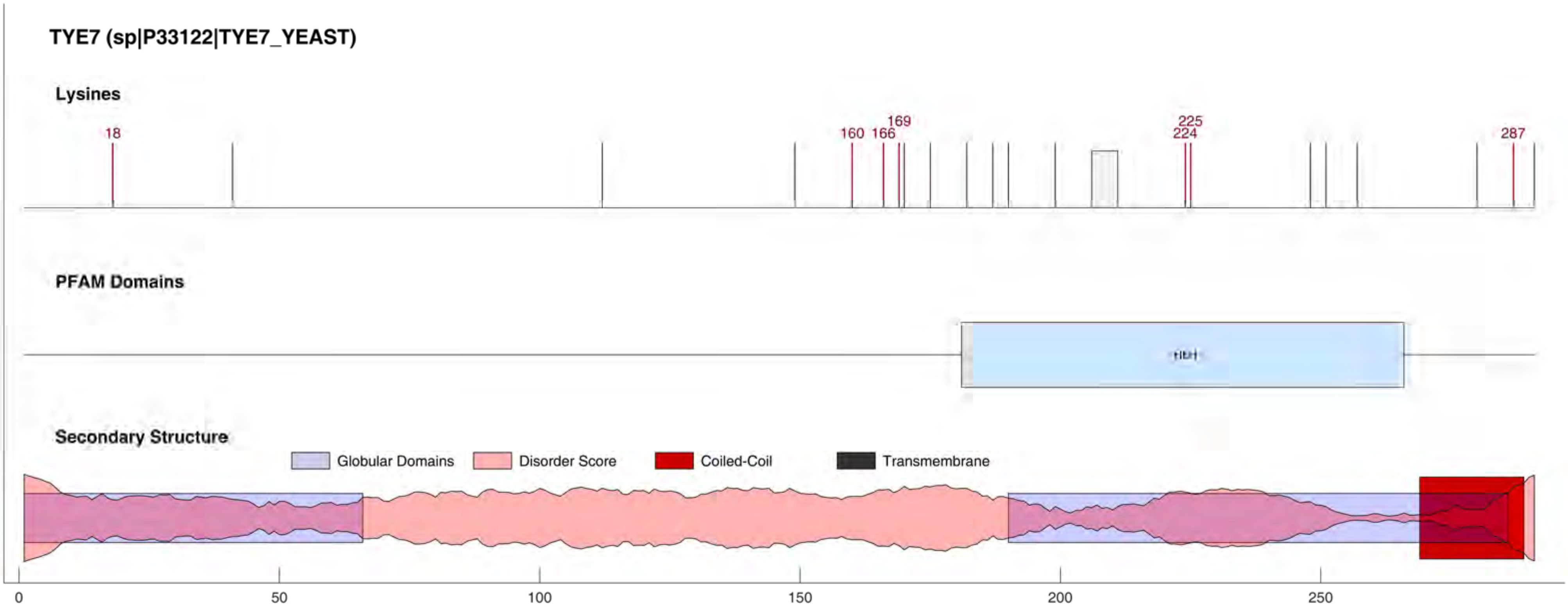

TYE7 (sp|P33122|TYE7_YEAST)
Lysines
18
160
166
169
224
225
287
PFAM Domains
HLH
Secondary Structure
Globular Domains
Disorder Score
Coiled-Coil
Transmembrane
0
50
100
150
200
250

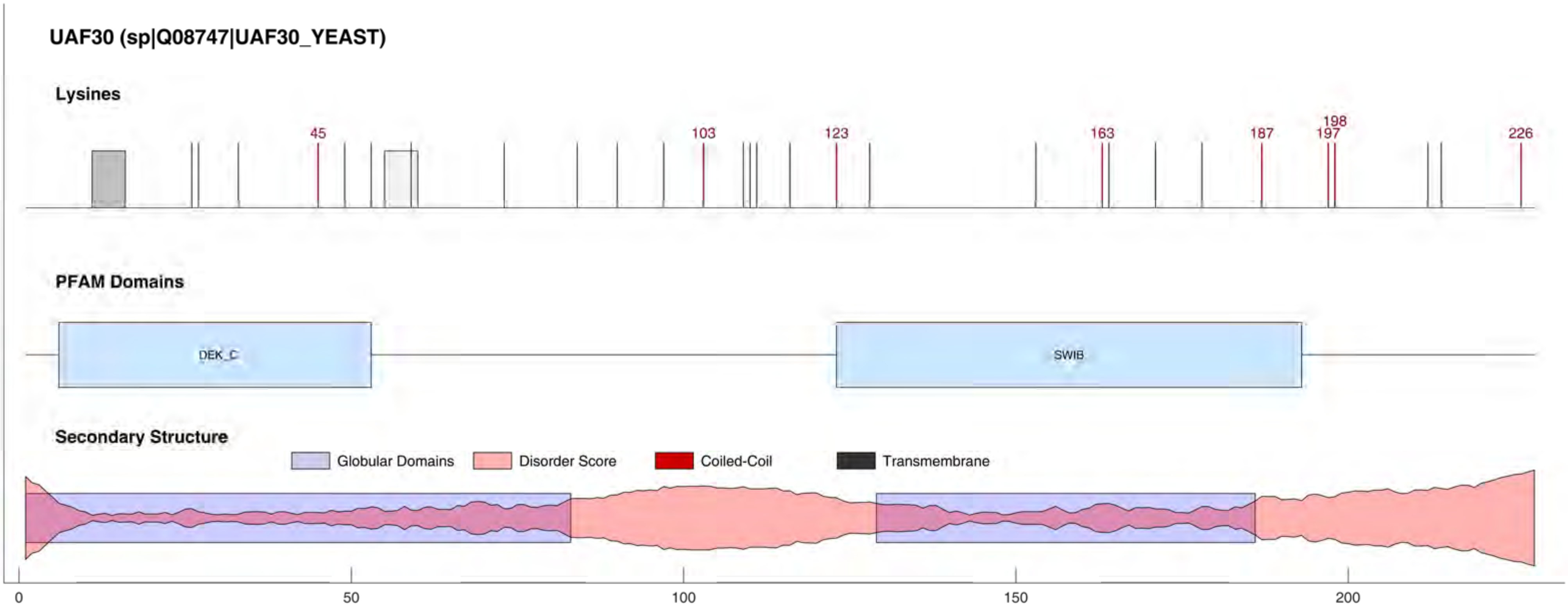
UAF30 (sp|Q08747|UAF30_YEAST)
Lysines
45
103
123
163
187
198
197
226
PFAM Domains
DEK_C
SWIB
Secondary Structure
Globular Domains
Disorder Score
Coiled-Coil
Transmembrane
0
50
100
150
200

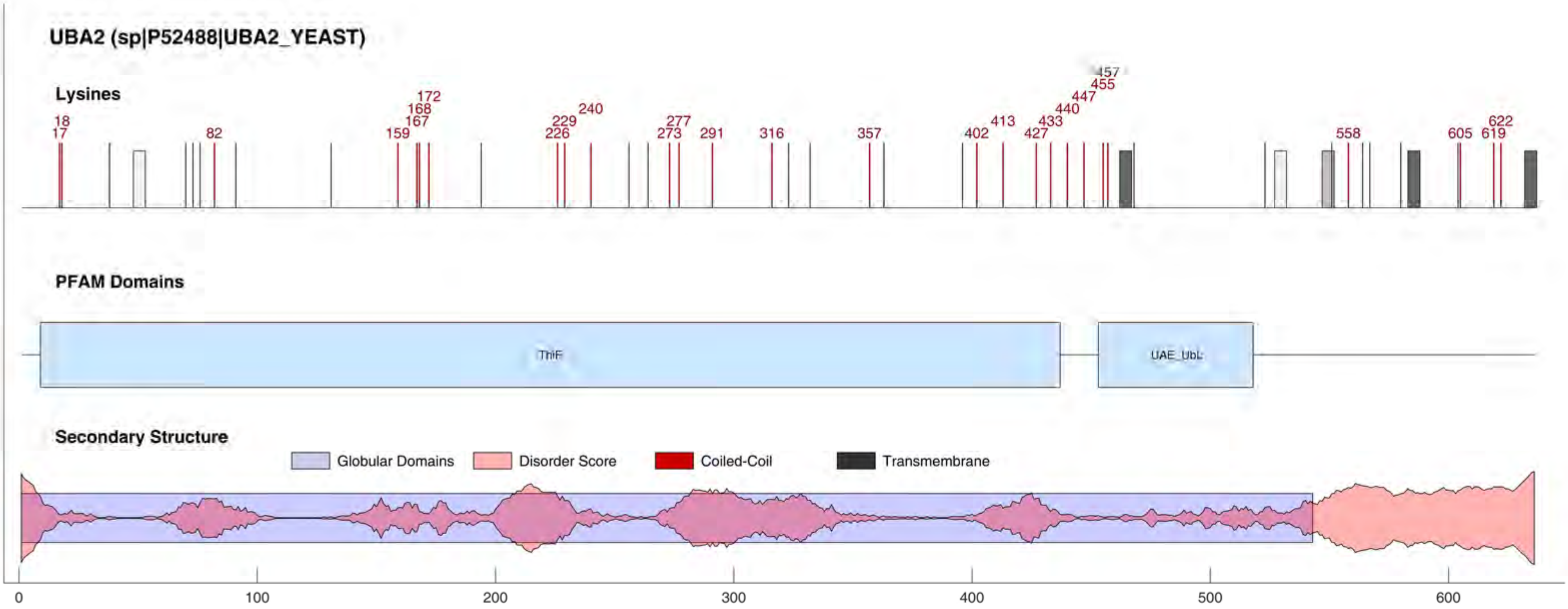

UBA2 (sp|P52488|UBA2_YEAST)
Lysines
17
18
82
159
167
168
172
226
229
240
273
277
291
316
357
402
413
427
433
440
447
455
457
558
605
619
622
PFAM Domains
ThiF
UAE_UbL
Secondary Structure
Globular Domains
Disorder Score
Coiled-Coil
Transmembrane
0
100
200
300
400
500
600

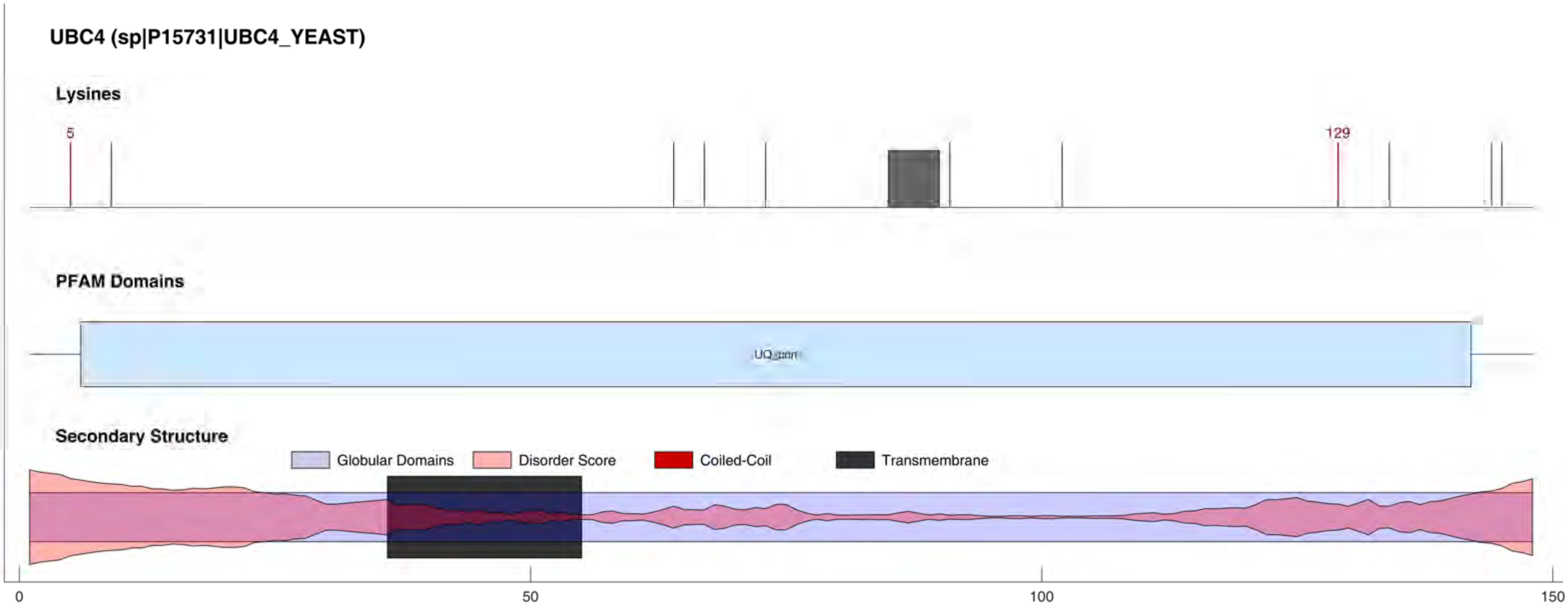
UBC4 (sp|P15731|UBC4_YEAST)
Lysines
5
129
PFAM Domains
Secondary Structure
Globular Domains
Disorder Score
Coiled-Coil
Transmembrane
0
50
100
150

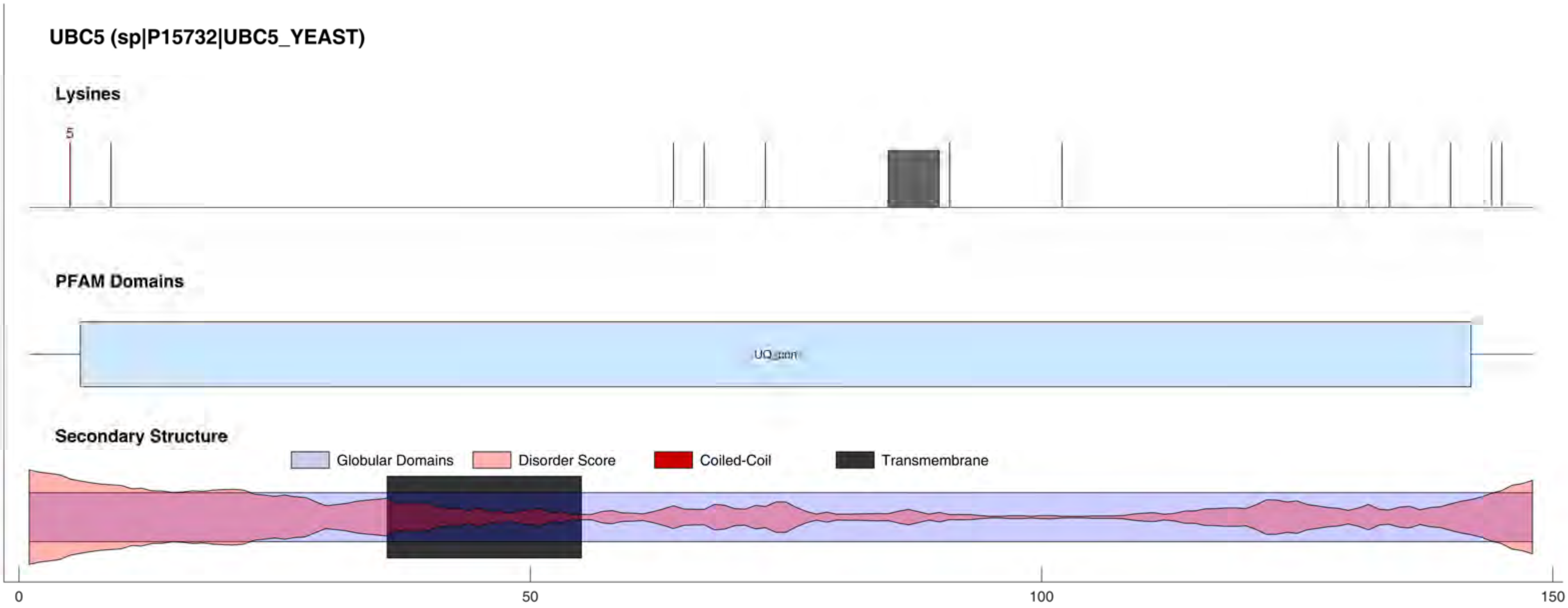
UBC5 (sp|P15732|UBC5_YEAST)
Lysines
5
PFAM Domains
UQ_con
Secondary Structure
Globular Domains
Disorder Score
Coiled-Coil
Transmembrane
0
50
100
150

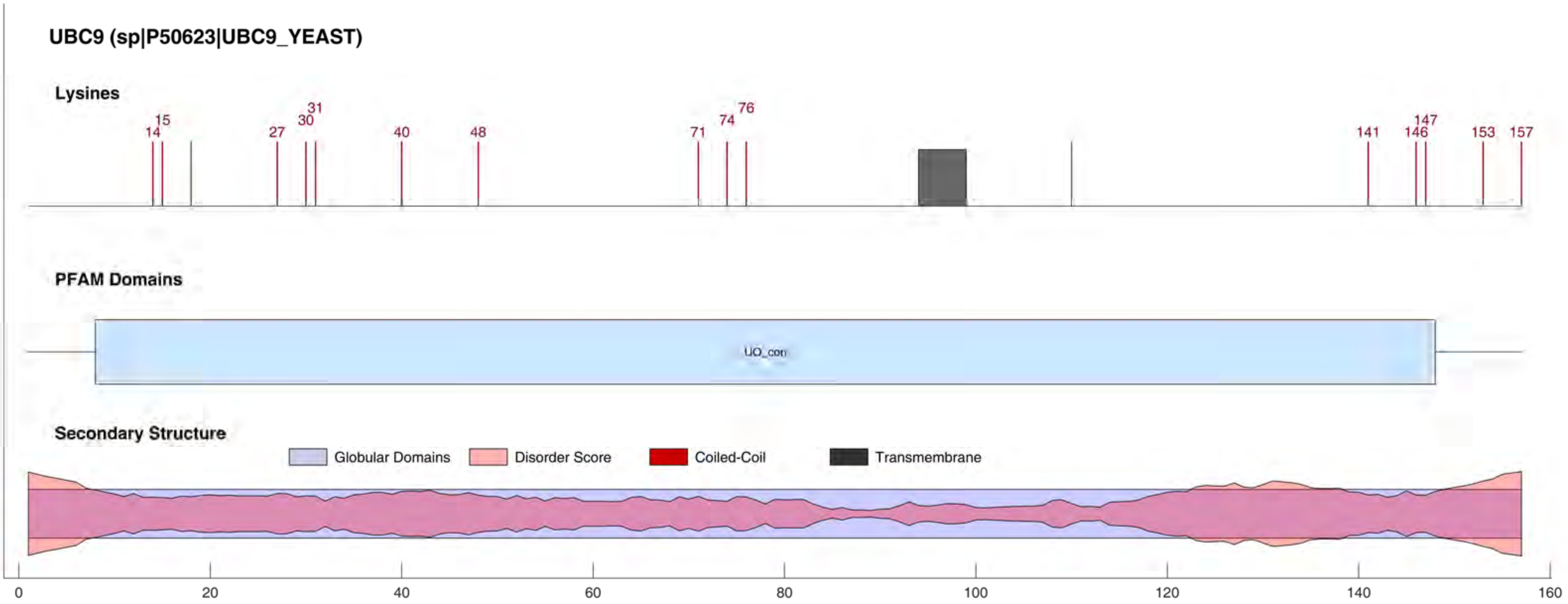
UBC9 (sp|P50623|UBC9_YEAST)
Lysines
14
15
27
30
31
40
48
71
74
76
141
146
147
153
157
PFAM Domains
UQ_con
Secondary Structure
Globular Domains
Disorder Score
Coiled-Coil
Transmembrane
0
20
40
60
80
100
120
140
160

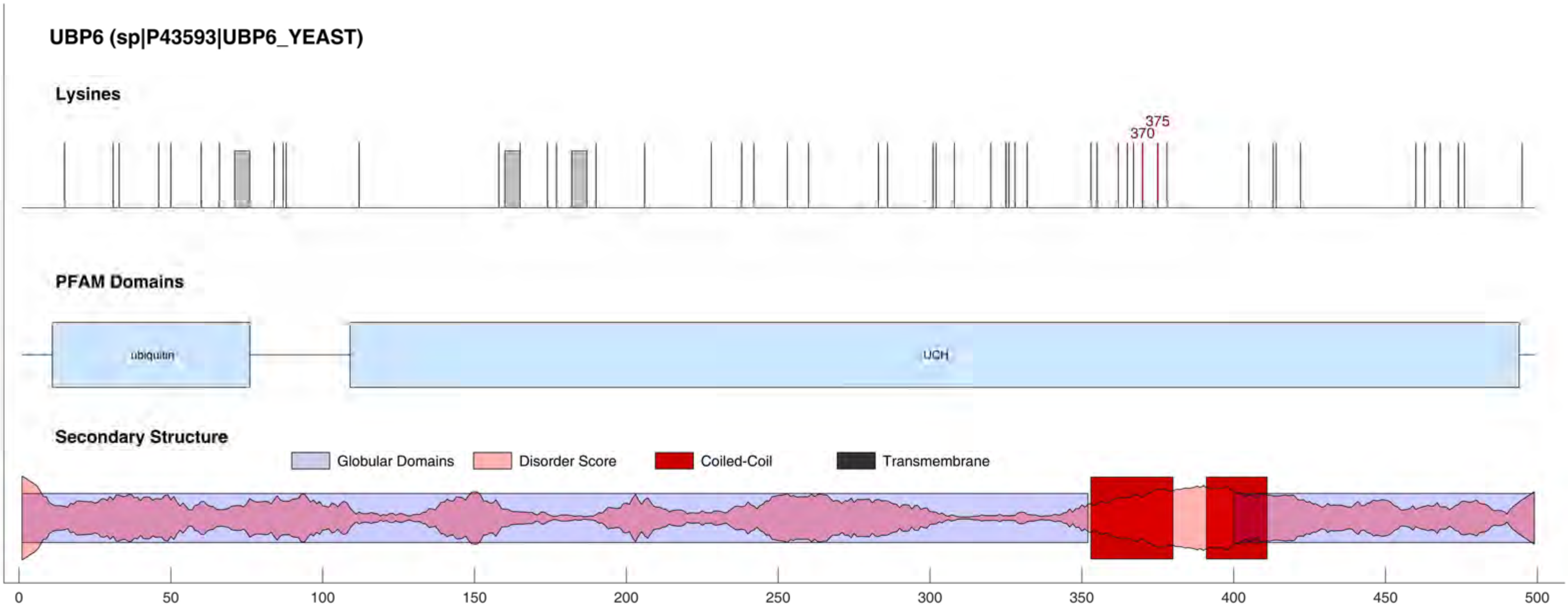
UBP6 (sp|P43593|UBP6_YEAST)
Lysines
375
370
PFAM Domains
ubiquitin
UCH
Secondary Structure
Globular Domains
Disorder Score
Coiled-Coil
Transmembrane
0
50
100
150
200
250
300
350
400
450
500

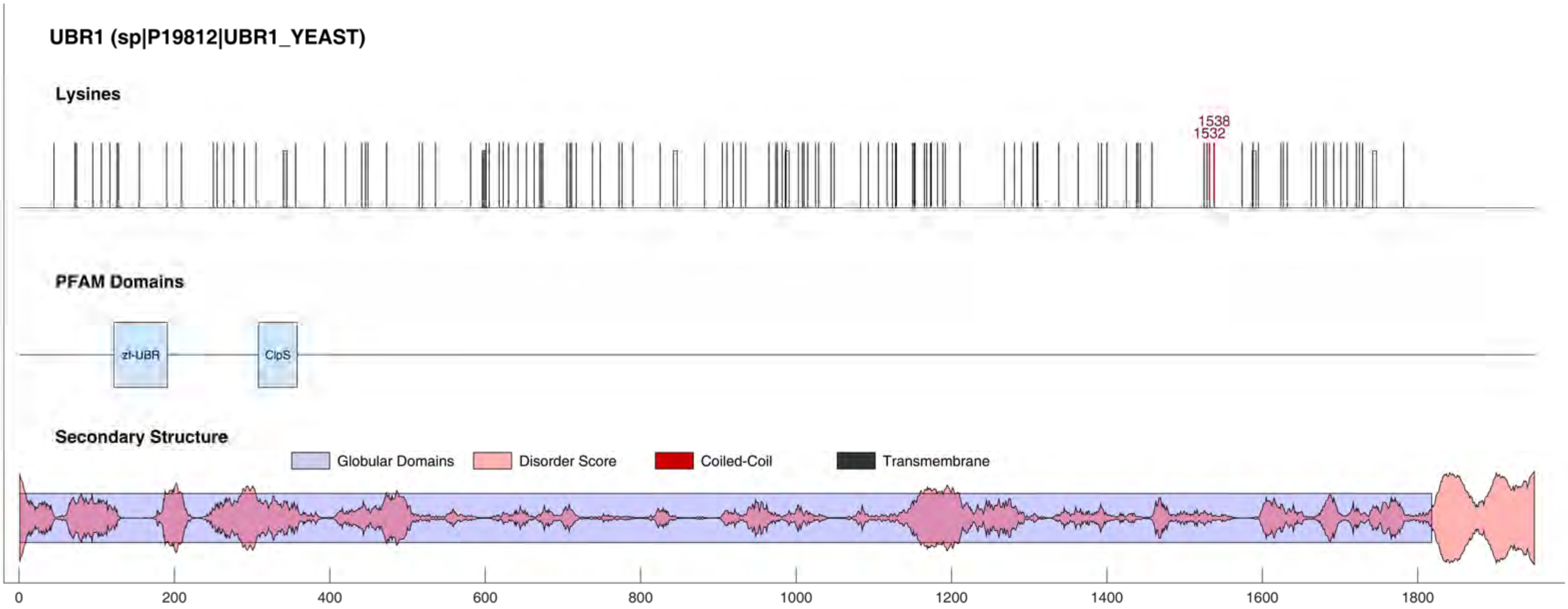

UBR1 (sp|P19812|UBR1_YEAST)
Lysines
1538
1532
PFAM Domains
zf-UBR
ClpS
Secondary Structure
Globular Domains
Disorder Score
Coiled-Coil
Transmembrane
0
200
400
600
800
1000
1200
1400
1600
1800

#### UFD1 (sp|P53044|UFD1_YEAST)

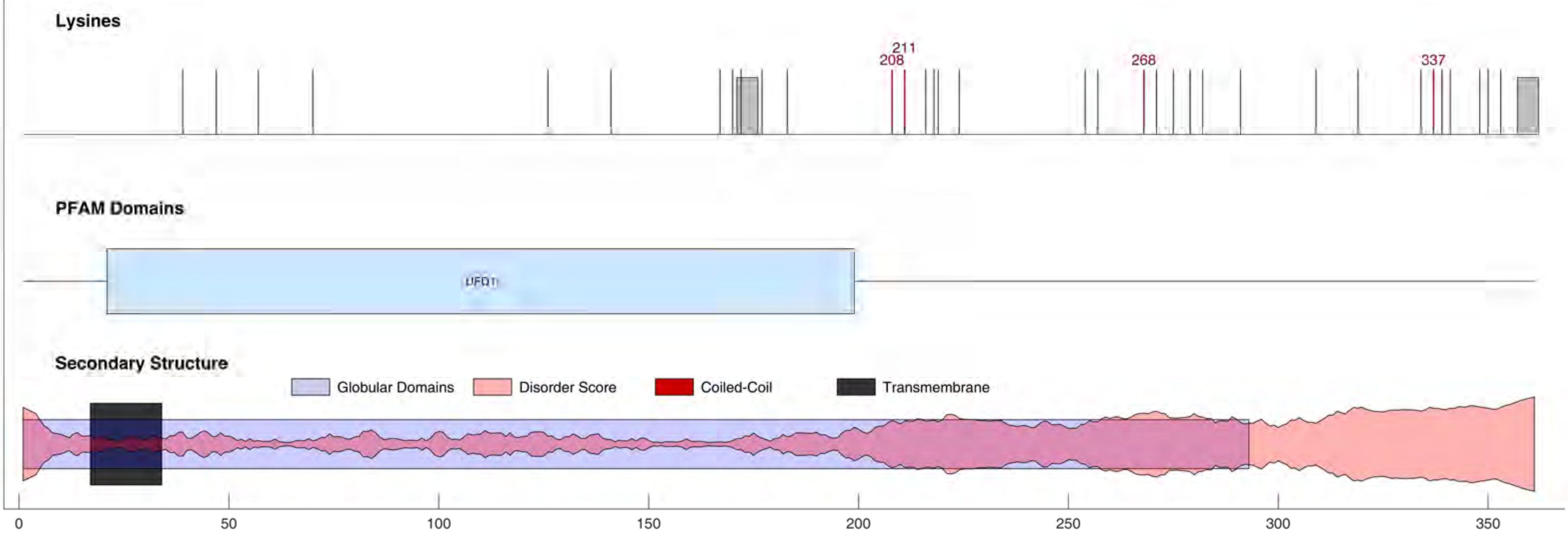

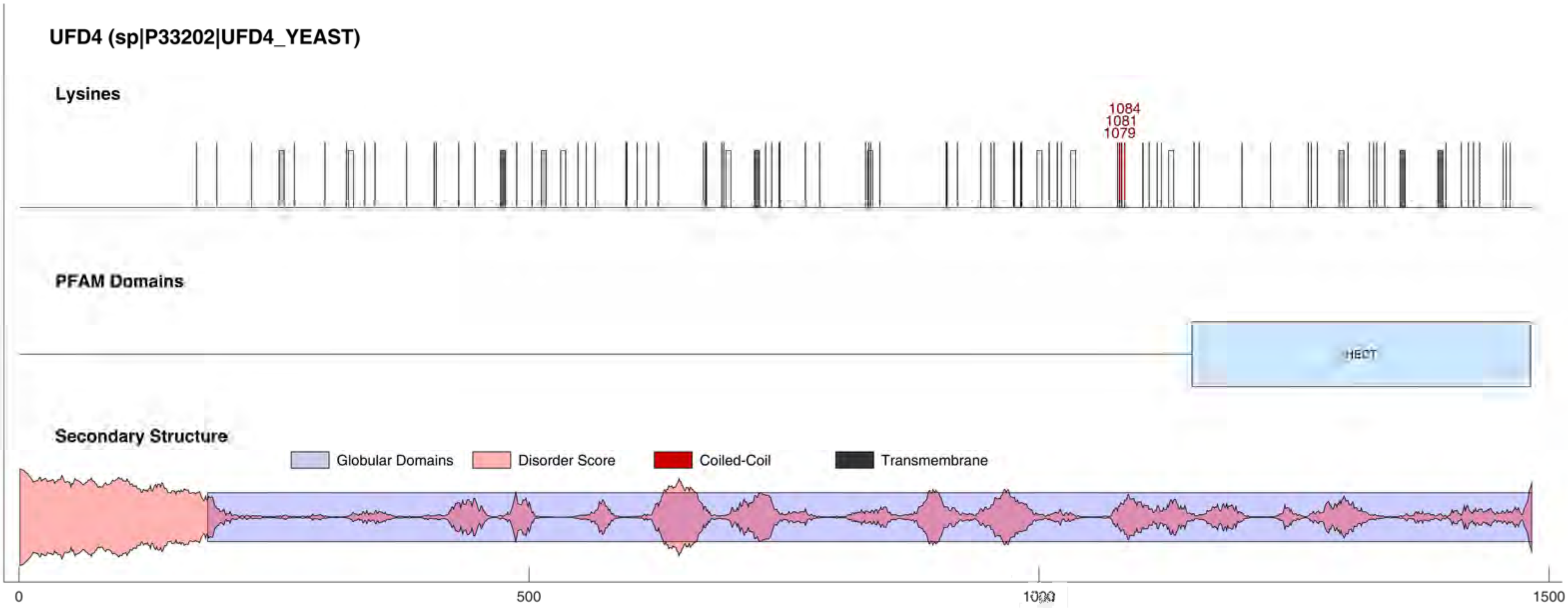
UFD4 (sp|P33202|UFD4_YEAST)
Lysines
1084
1081
1079
PFAM Domains
HECT
Secondary Structure
Globular Domains
Disorder Score
Coiled-Coil
Transmembrane
0
500
1000
1500

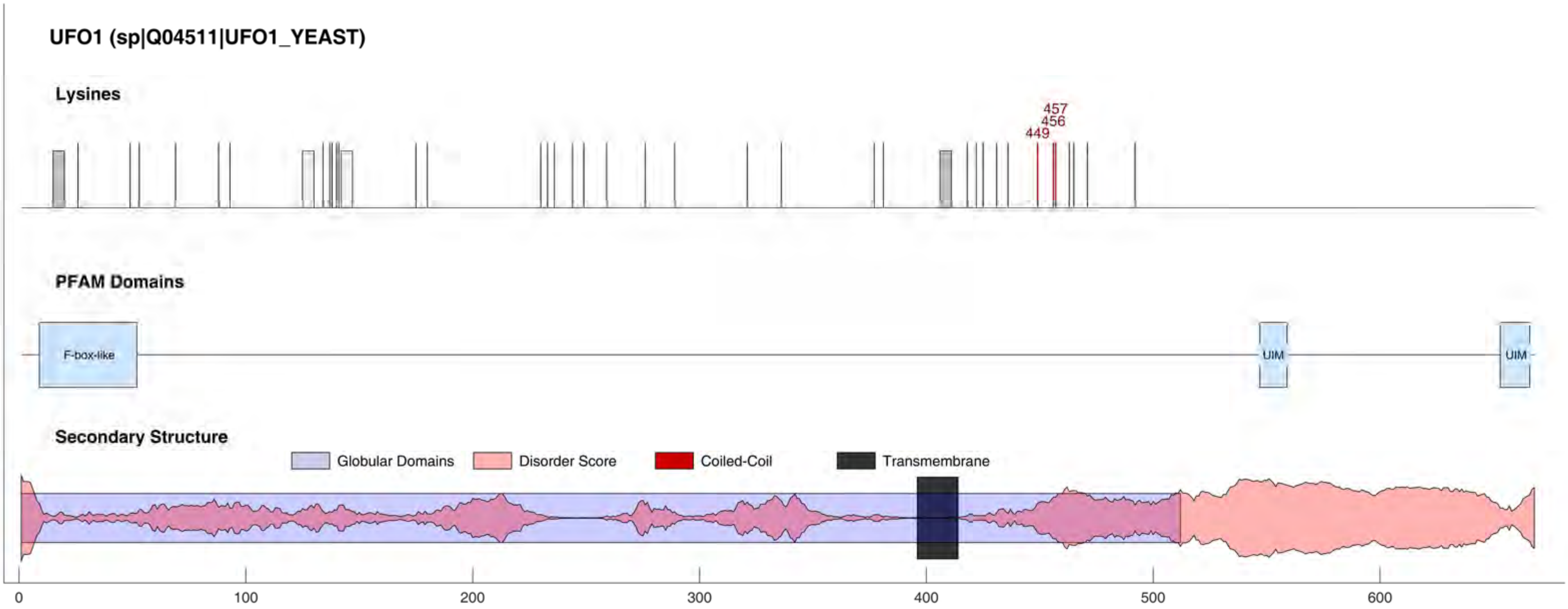

UFO1 (sp|Q04511|UFO1_YEAST)
Lysines
457
456
449
PFAM Domains
F-box-like
UIM
UIM
Secondary Structure
Globular Domains
Disorder Score
Coiled-Coil
Transmembrane
0
100
200
300
400
500
600

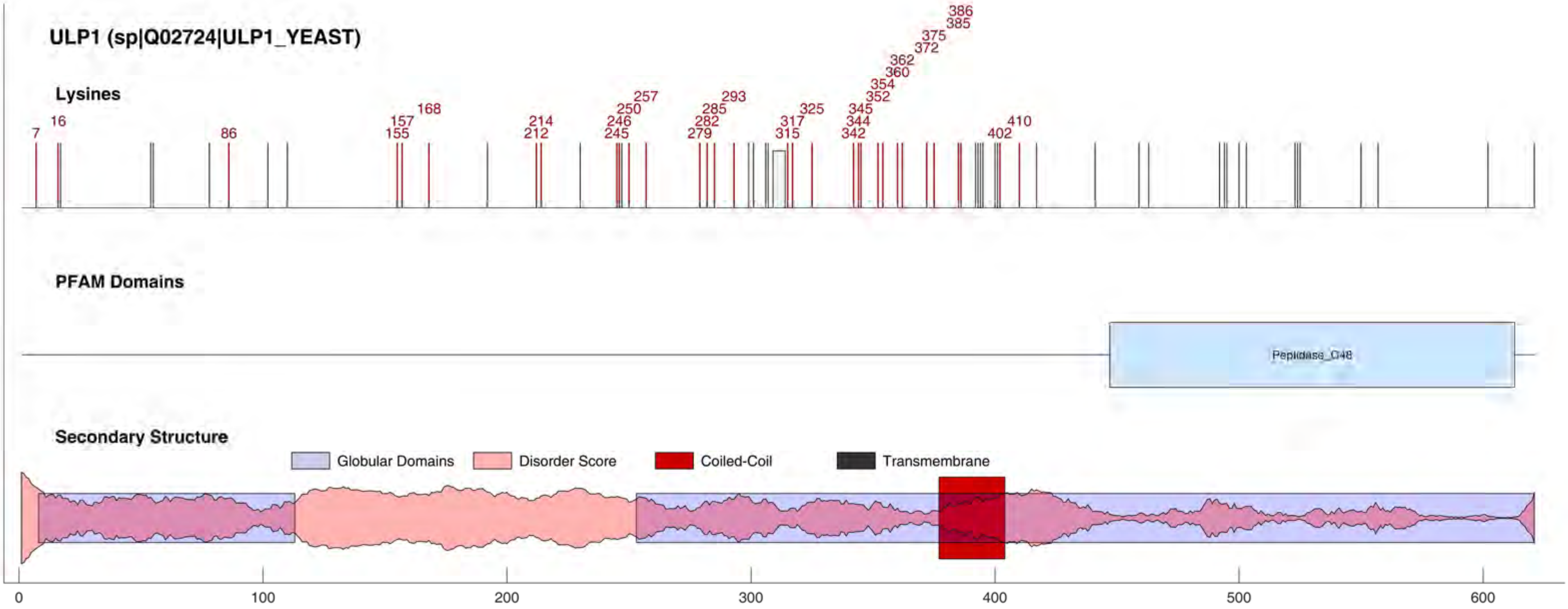
ULP1 (sp|Q02724|ULP1_YEAST)
Lysines
7
16
86
155
157
168
212
214
245
246
250
257
279
282
285
293
315
317
325
342
344
345
352
354
360
362
372
375
385
386
402
410
PFAM Domains
Peptidase_C48
Secondary Structure
Globular Domains
Disorder Score
Coiled-Coil
Transmembrane
0
100
200
300
400
500
600

#### ULS1 (sp|Q08562|ULS1_YEAST)

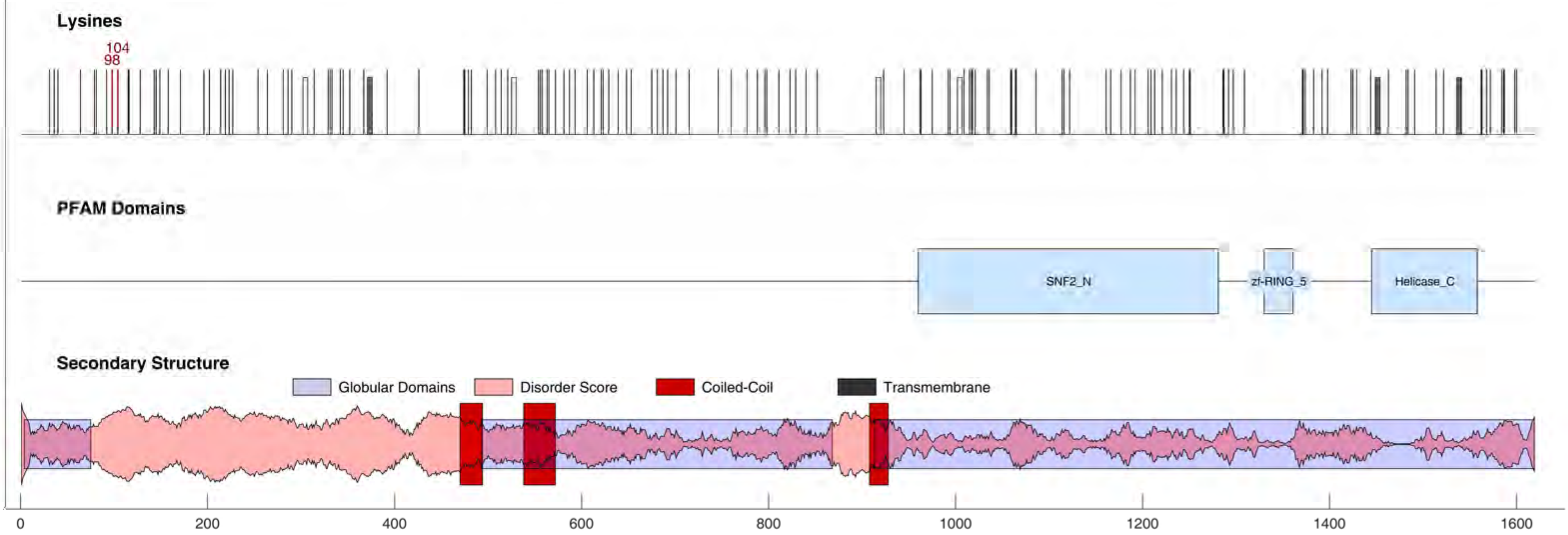

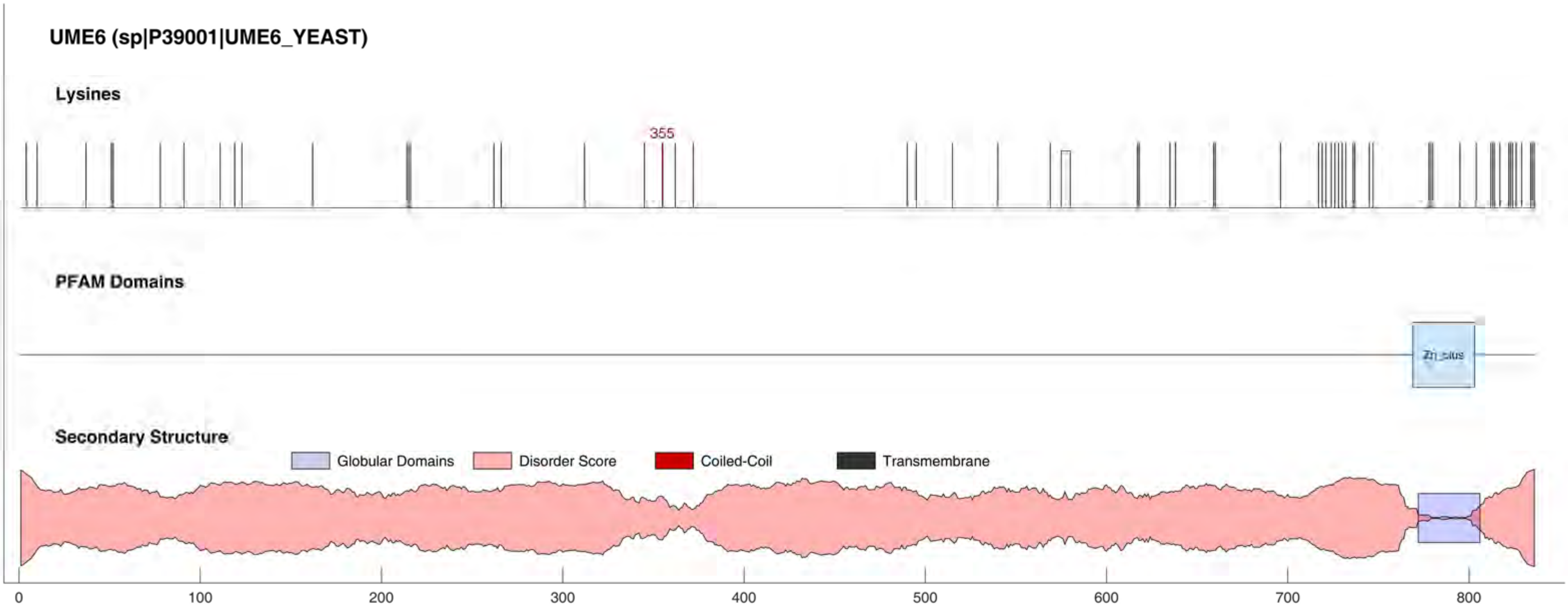
UME6 (sp|P39001|UME6_YEAST)
Lysines
355
PFAM Domains
Zn_clus
Secondary Structure
Globular Domains
Disorder Score
Coiled-Coil
Transmembrane
0
100
200
300
400
500
600
700
800

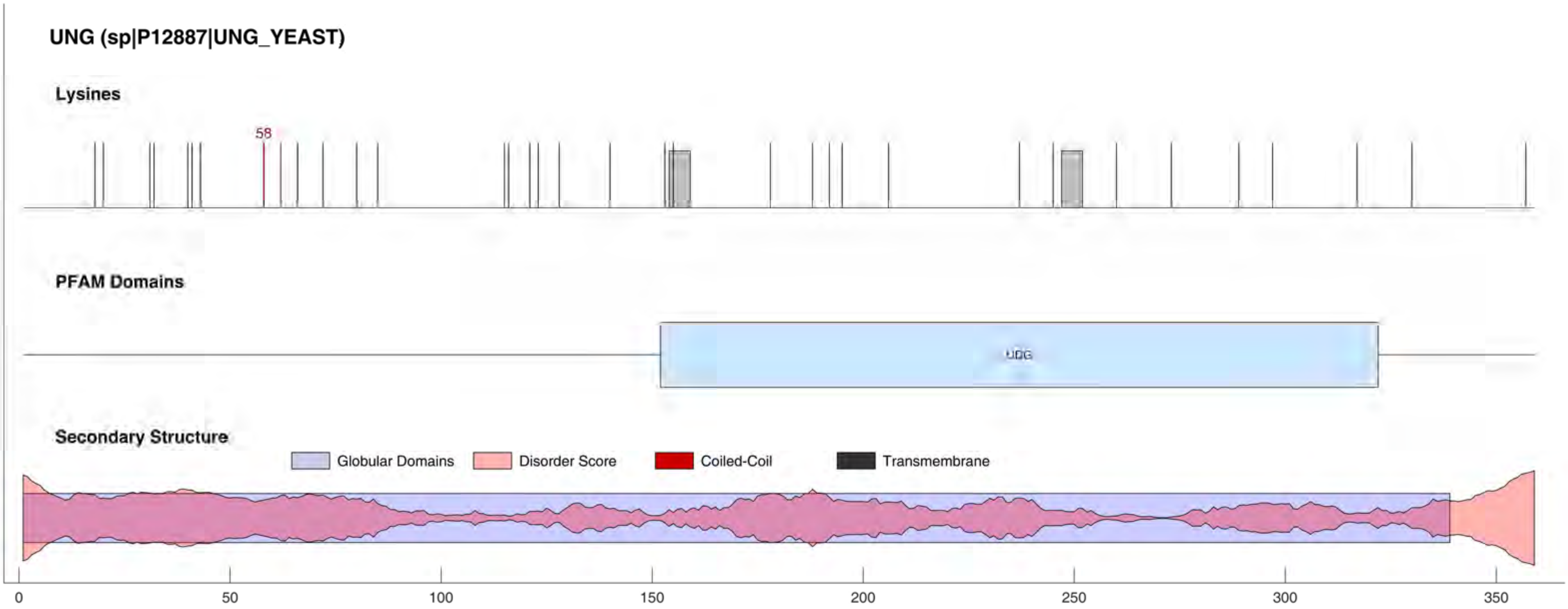
UNG (sp|P12887|UNG_YEAST)
Lysines
58
PFAM Domains
UDG
Secondary Structure
Globular Domains
Disorder Score
Coiled-Coil
Transmembrane
0
50
100
150
200
250
300
350

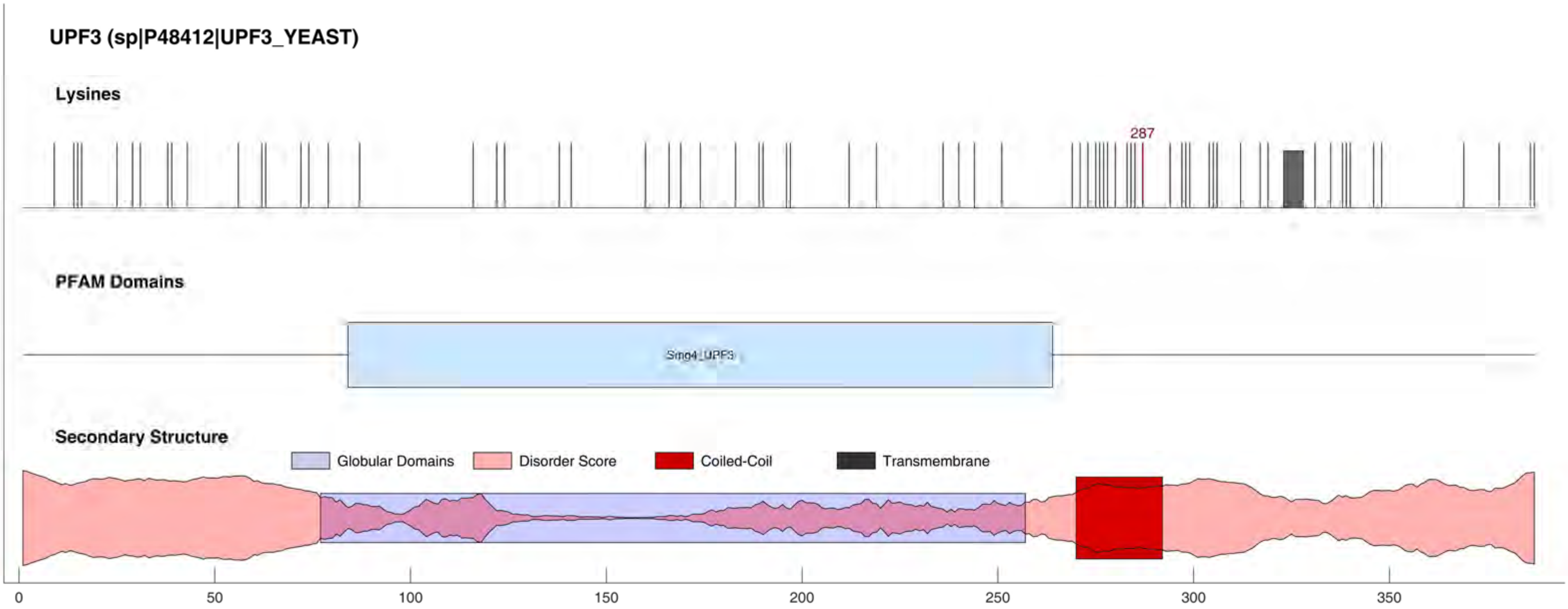
UPF3 (sp|P48412|UPF3_YEAST)
Lysines
287
PFAM Domains
Smg4_UPF3
Secondary Structure
Globular Domains
Disorder Score
Coiled-Coil
Transmembrane
0
50
100
150
200
250
300
350

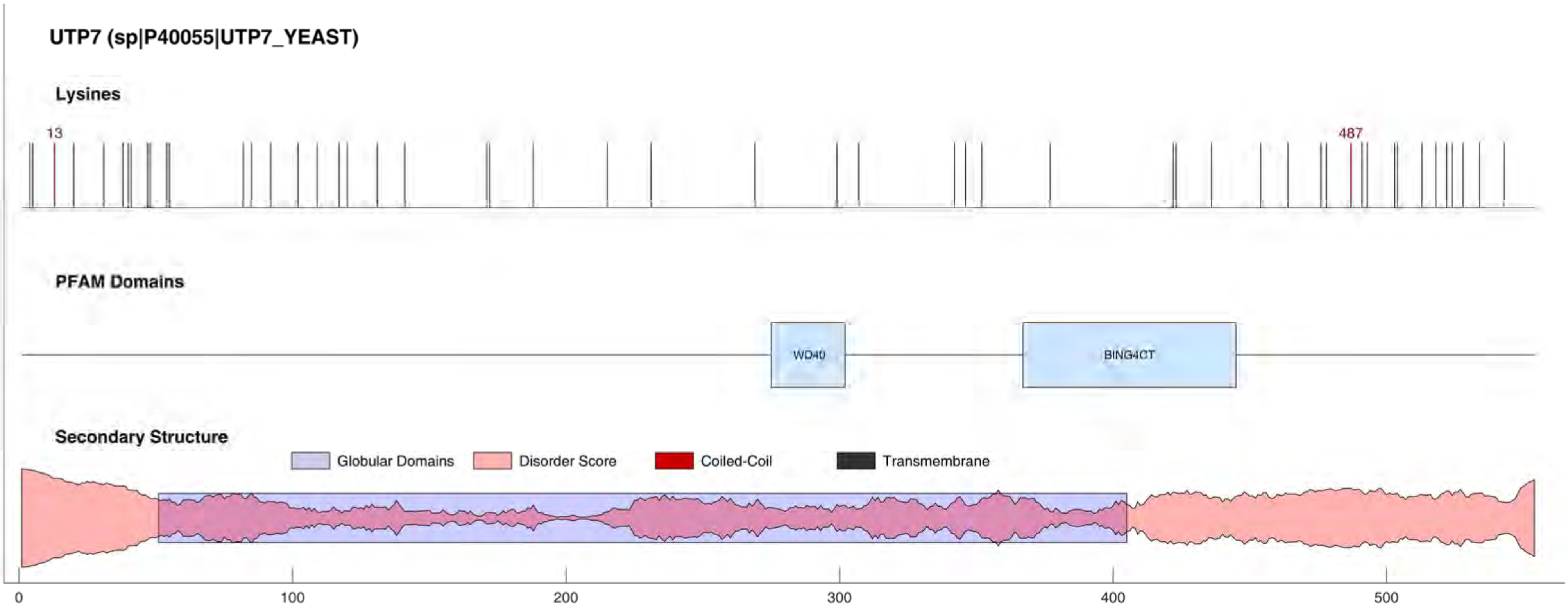
UTP7 (sp|P40055|UTP7_YEAST)
Lysines
13
487
PFAM Domains
WD40
BING4CT
Secondary Structure
Globular Domains
Disorder Score
Coiled-Coil
Transmembrane
0
100
200
300
400
500

#### UTP8 (sp|P53276|UTP8_YEAST)

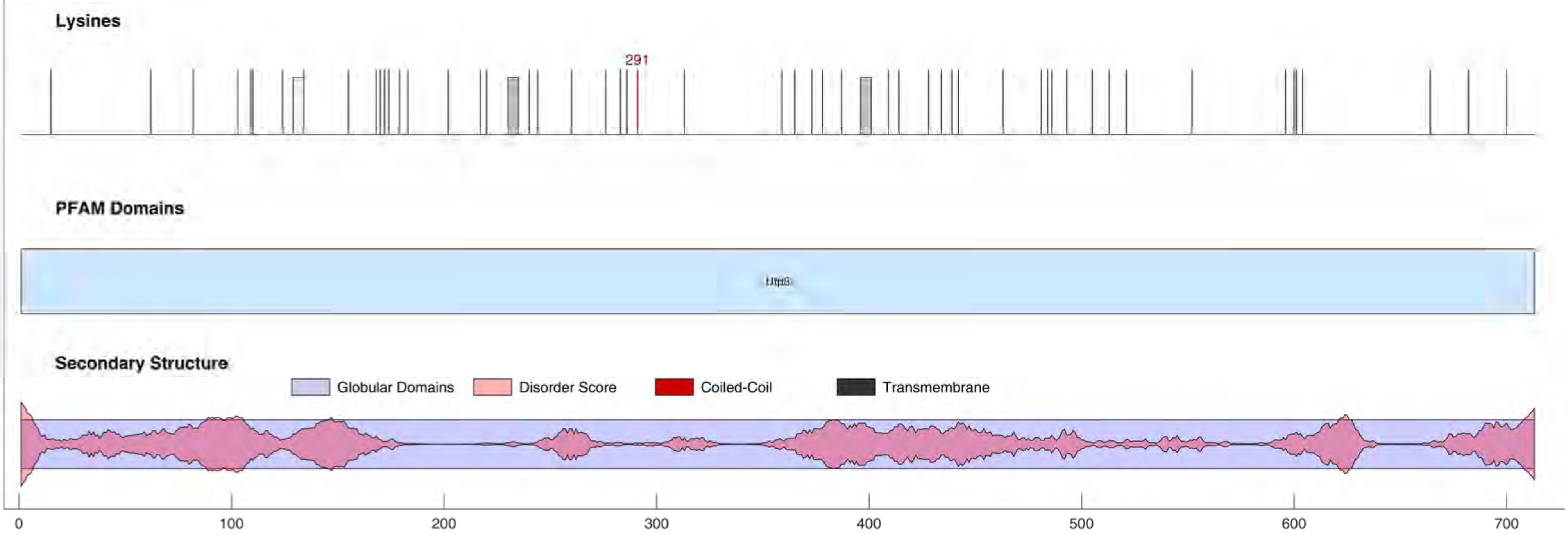

#### UTP11 (sp|P34247|UTP11_YEAST)

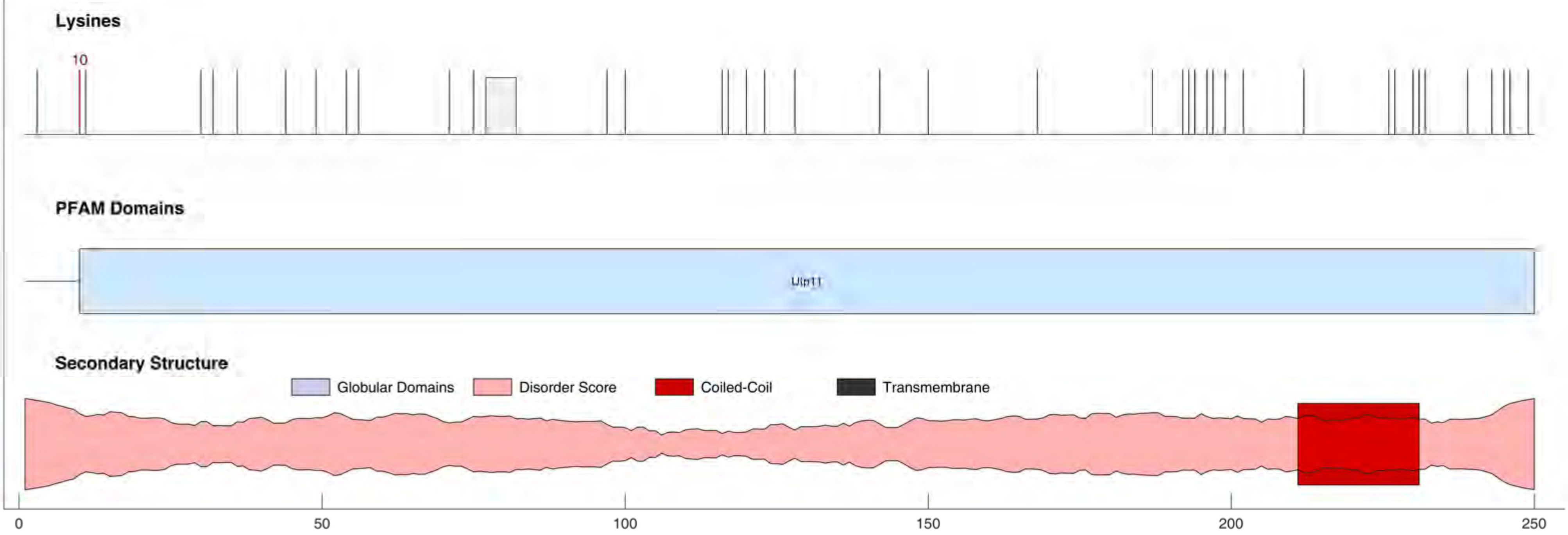

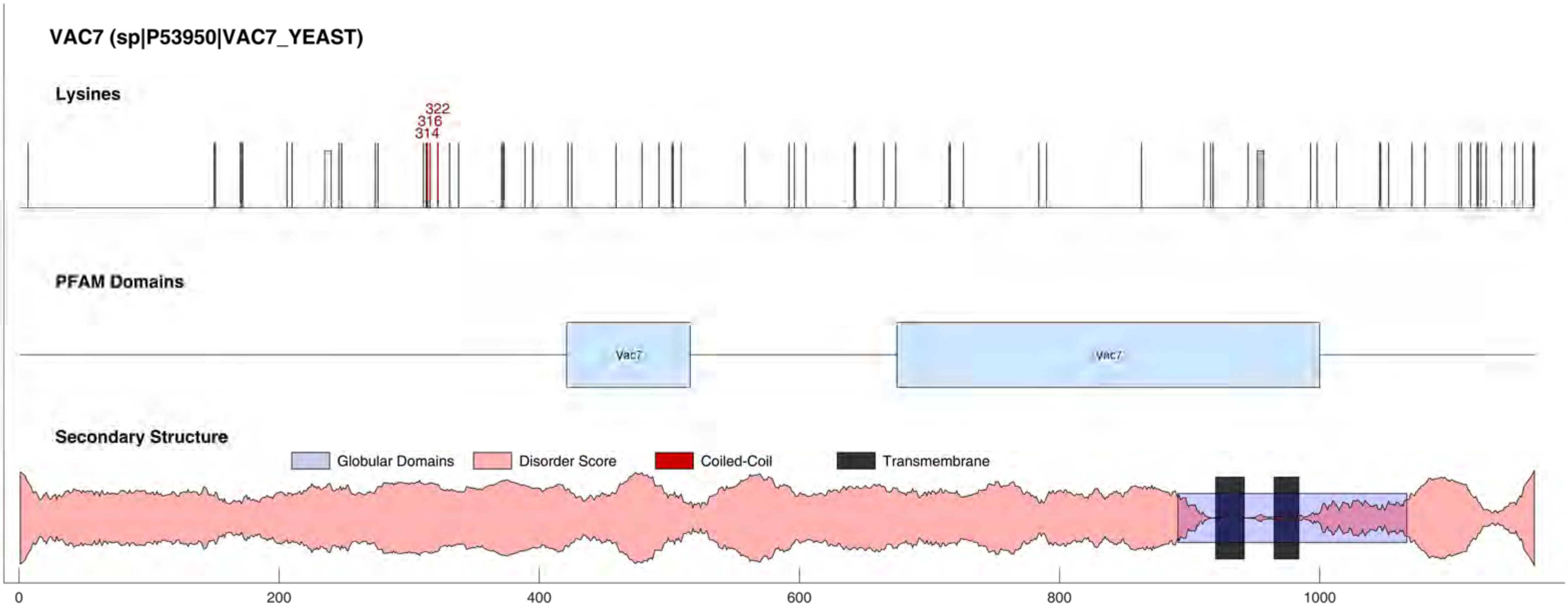

VAC7 (sp|P53950|VAC7_YEAST)
Lysines
322
316
314
PFAM Domains
Vac7
Vac7
Secondary Structure
Globular Domains
Disorder Score
Coiled-Coil
Transmembrane
0
200
400
600
800
1000

#### VAC14 (sp|Q06708|VAC14_YEAST)

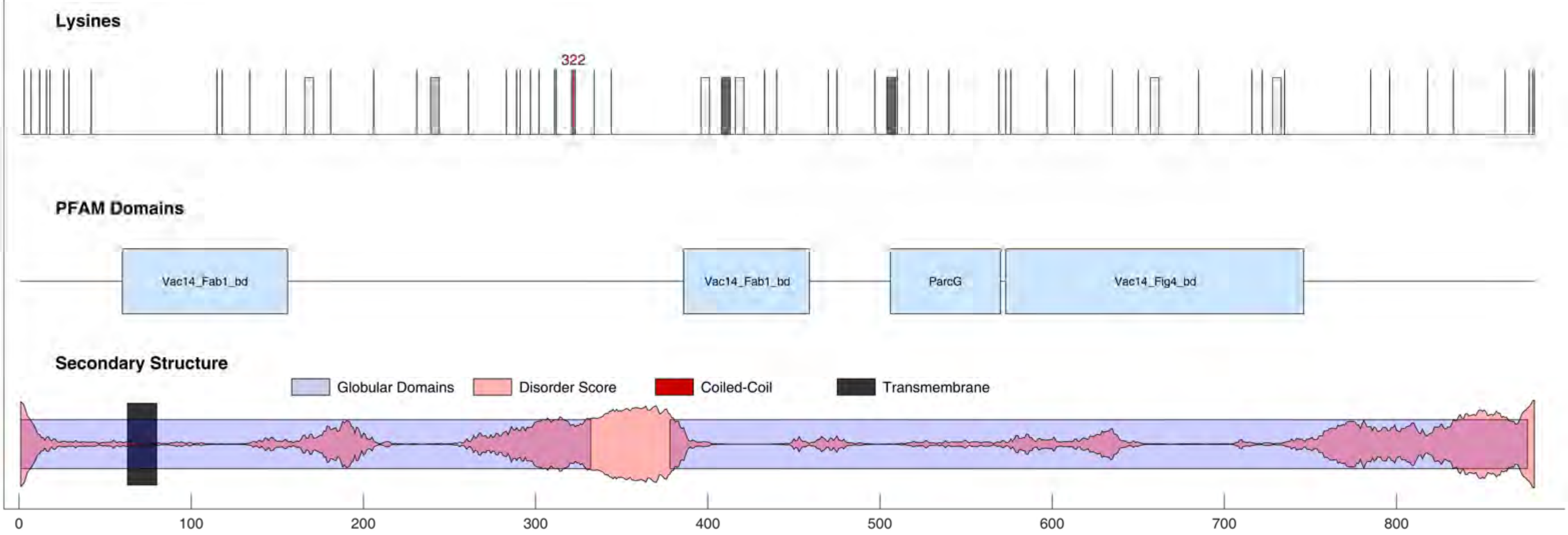

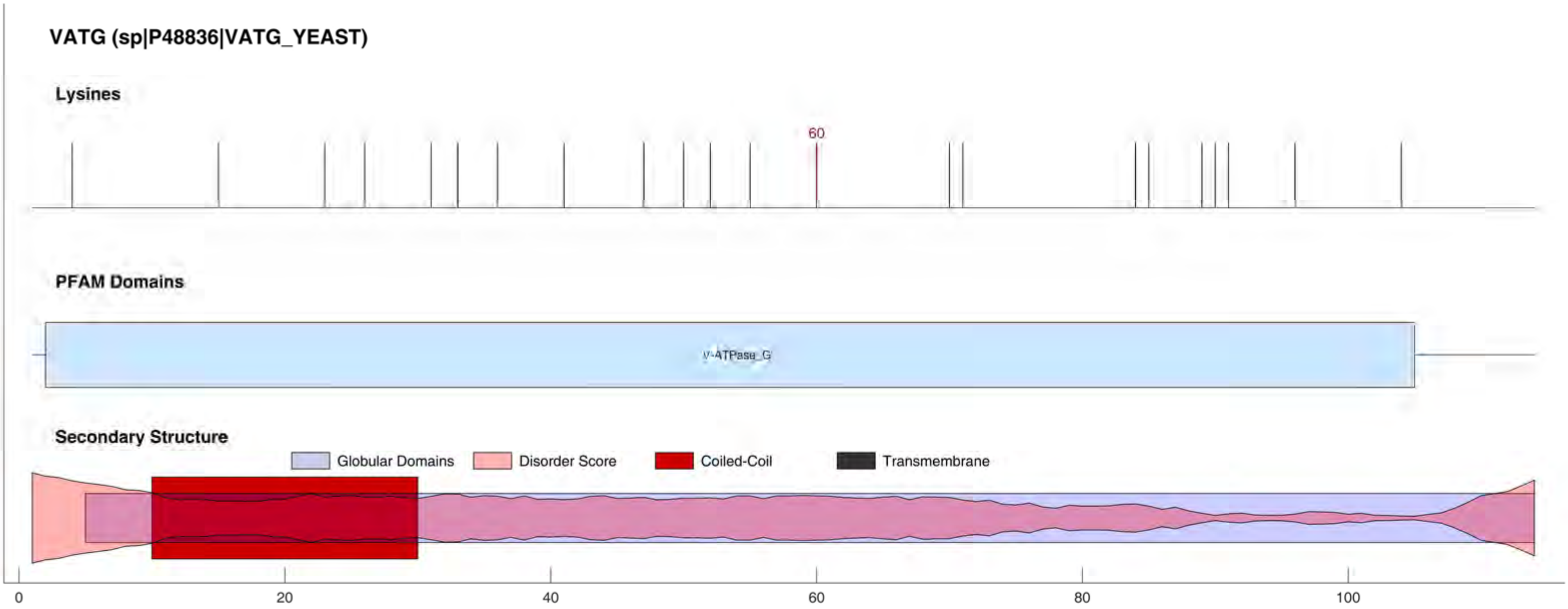
VATG (sp|P48836|VATG_YEAST)
Lysines
60
PFAM Domains
V-ATPase_G
Secondary Structure
Globular Domains
Disorder Score
Coiled-Coil
Transmembrane
0
20
40
60
80
100

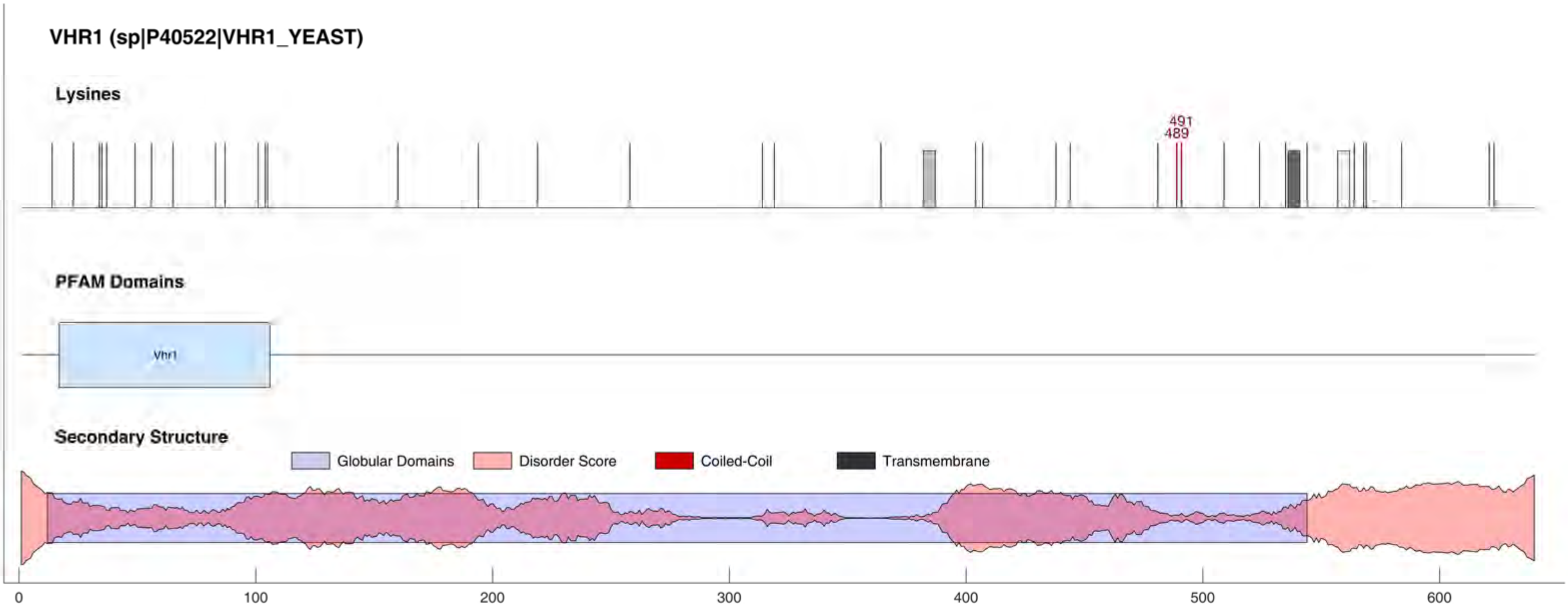
VHR1 (sp|P40522|VHR1_YEAST)
Lysines
491
489
PFAM Domains
Vhr1
Secondary Structure
Globular Domains
Disorder Score
Coiled-Coil
Transmembrane
0
100
200
300
400
500
600

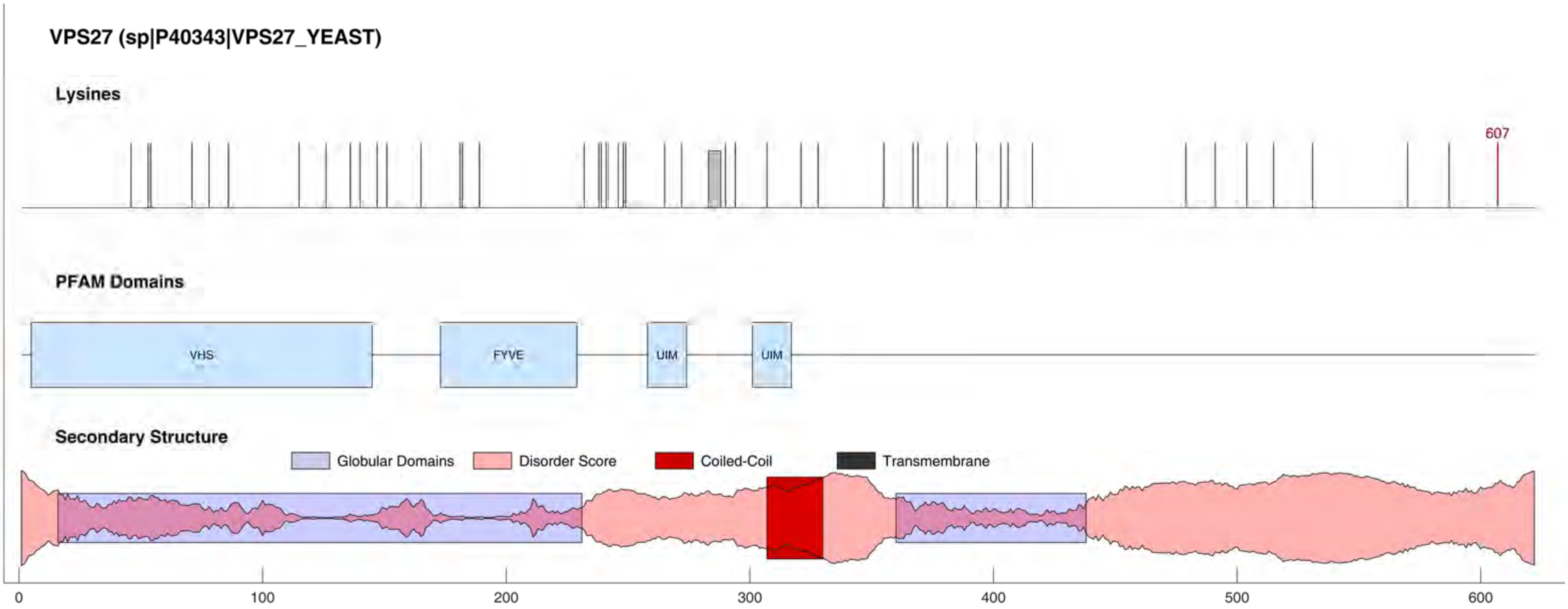
VPS27 (sp|P40343|VPS27_YEAST)
Lysines
607
PFAM Domains
VHS
FYVE
UIM
UIM
Secondary Structure
Globular Domains
Disorder Score
Coiled-Coil
Transmembrane
0
100
200
300
400
500
600

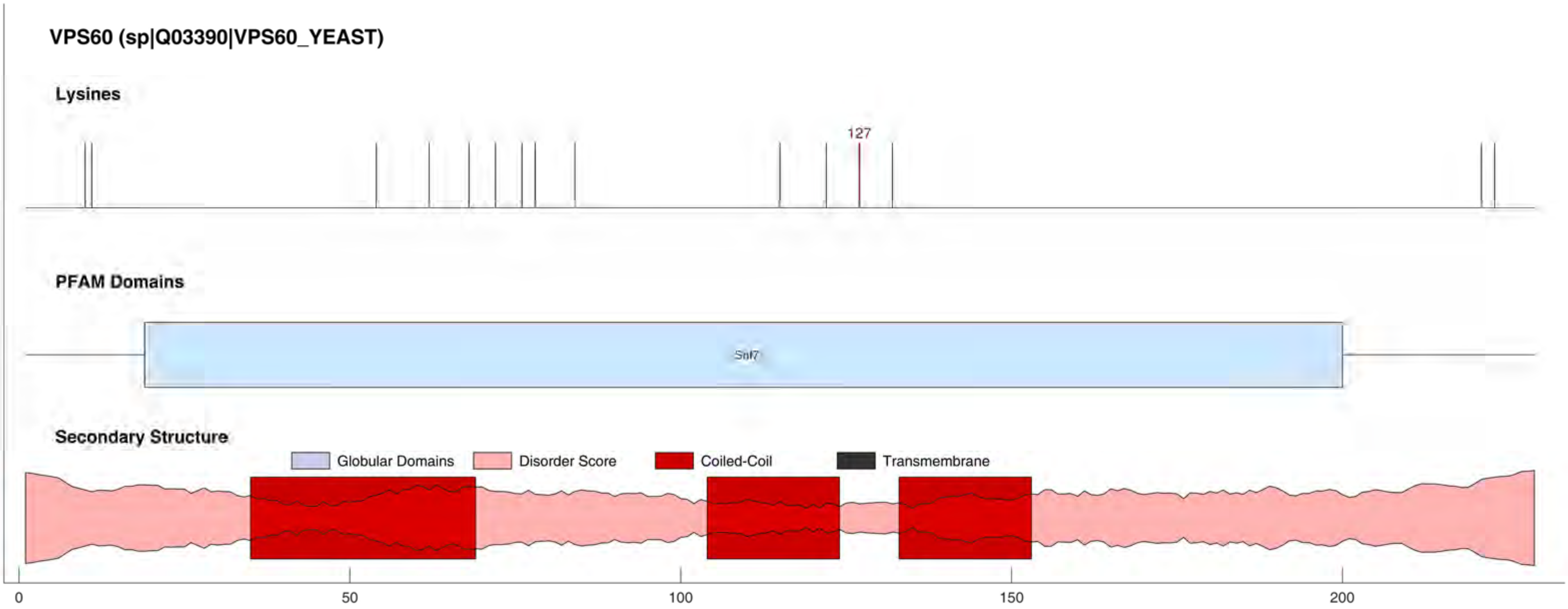

VPS60 (sp|Q03390|VPS60_YEAST)
Lysines
127
PFAM Domains
Snf7
Secondary Structure
Globular Domains
Disorder Score
Coiled-Coil
Transmembrane
0
50
100
150
200

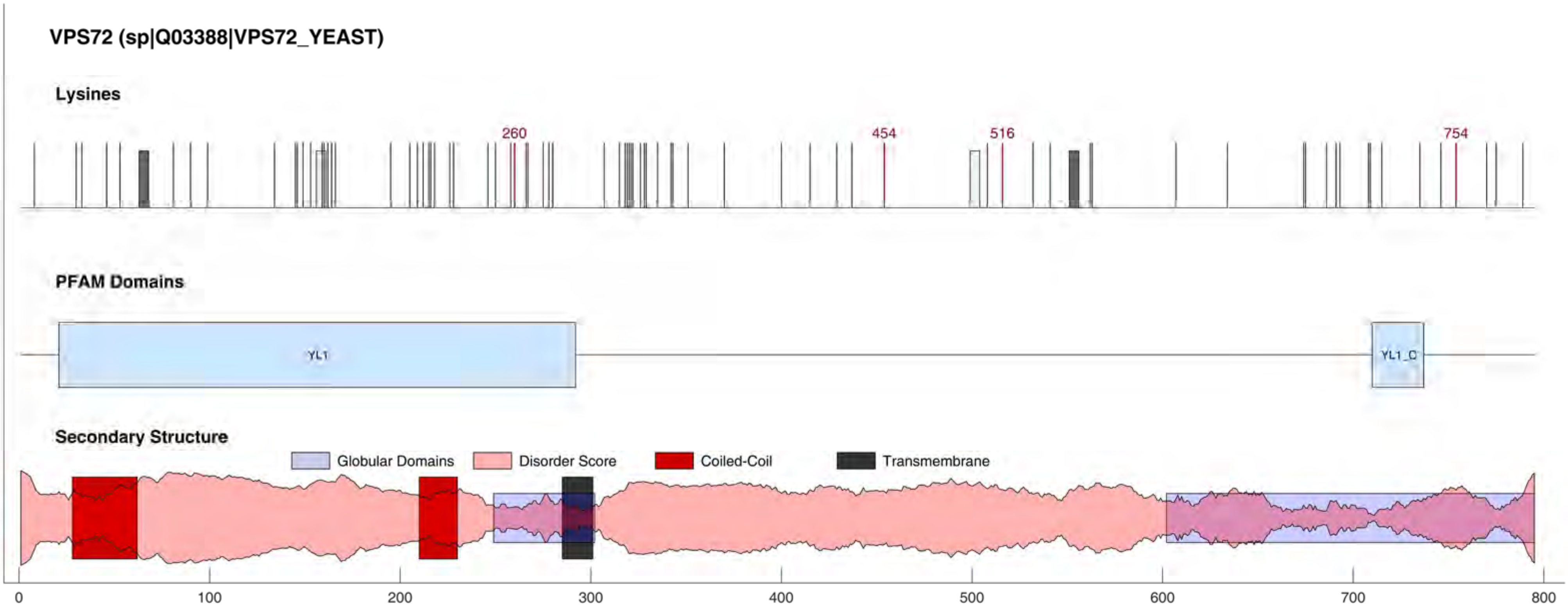

VPS72 (sp|Q03388|VPS72_YEAST)
Lysines
260
454
516
754
PFAM Domains
YL1
YL1_C
Secondary Structure
Globular Domains
Disorder Score
Coiled-Coil
Transmembrane
0
100
200
300
400
500
600
700
800

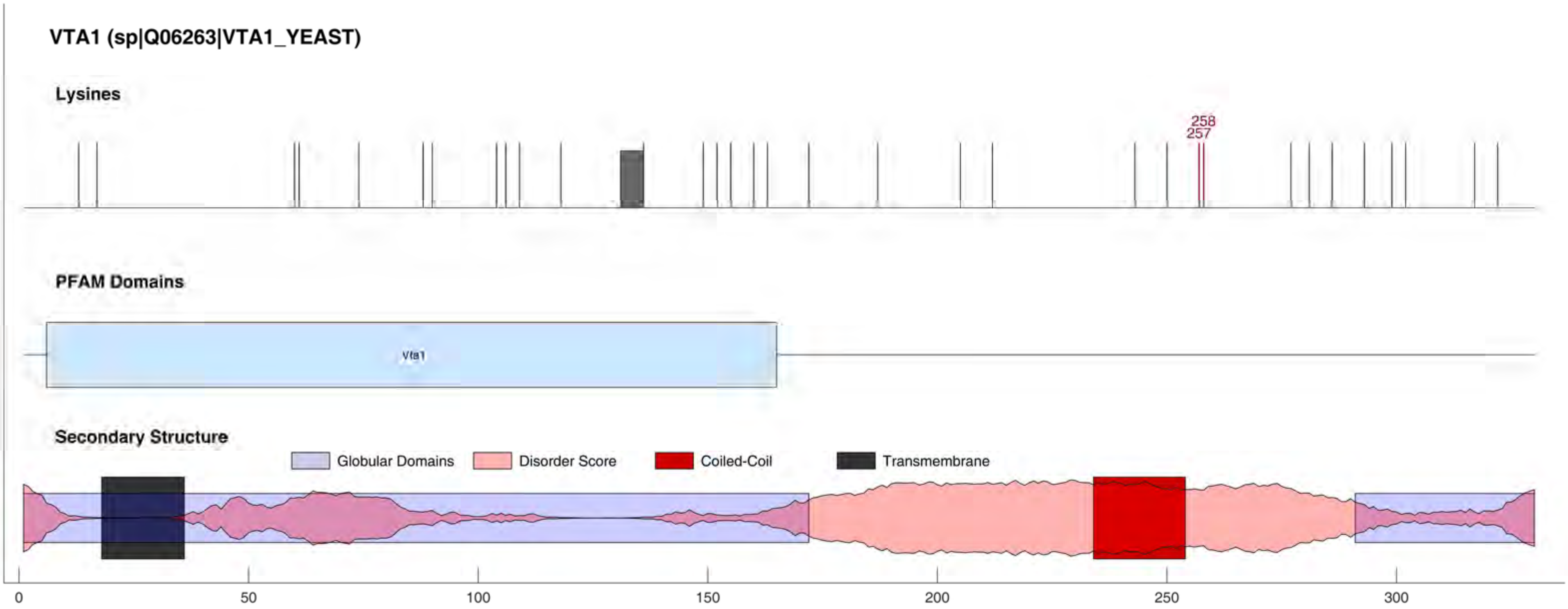
VTA1 (sp|Q06263|VTA1_YEAST)
Lysines
258
257
PFAM Domains
Vta1
Secondary Structure
Globular Domains
Disorder Score
Coiled-Coil
Transmembrane
0
50
100
150
200
250
300

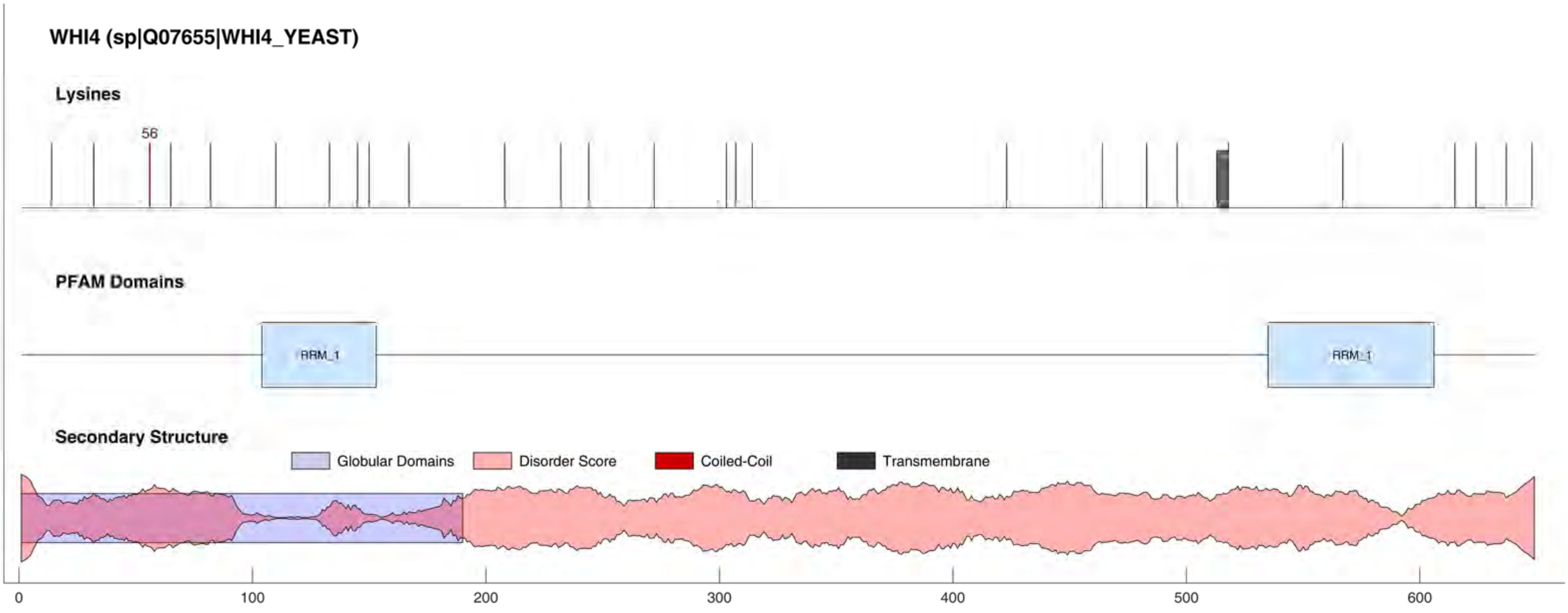
WHI4 (sp|Q07655|WHI4_YEAST)
Lysines
56
PFAM Domains
RRM_1
RRM_1
Secondary Structure
Globular Domains
Disorder Score
Coiled-Coil
Transmembrane
0
100
200
300
400
500
600

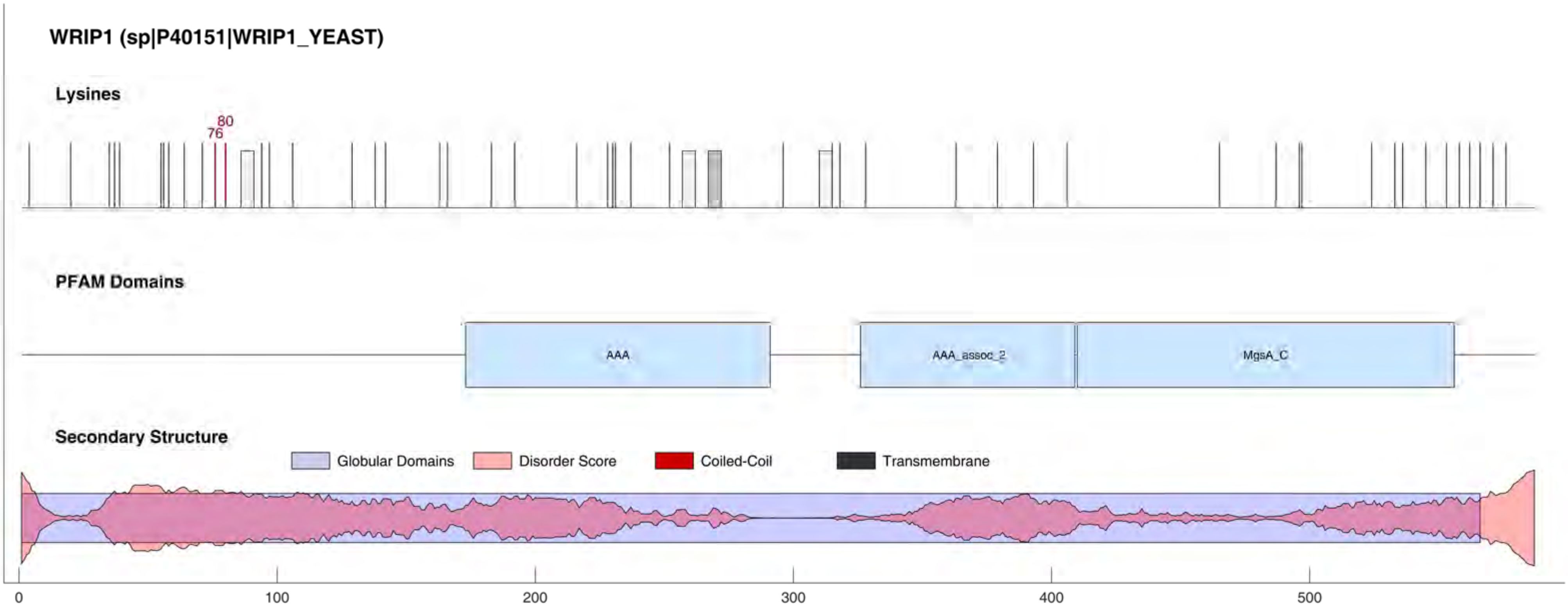
WRIP1 (sp|P40151|WRIP1_YEAST)
Lysines
80
76
PFAM Domains
AAA
AAA_assoc_2
MgsA_C
Secondary Structure
Globular Domains
Disorder Score
Coiled-Coil
Transmembrane
0
100
200
300
400
500

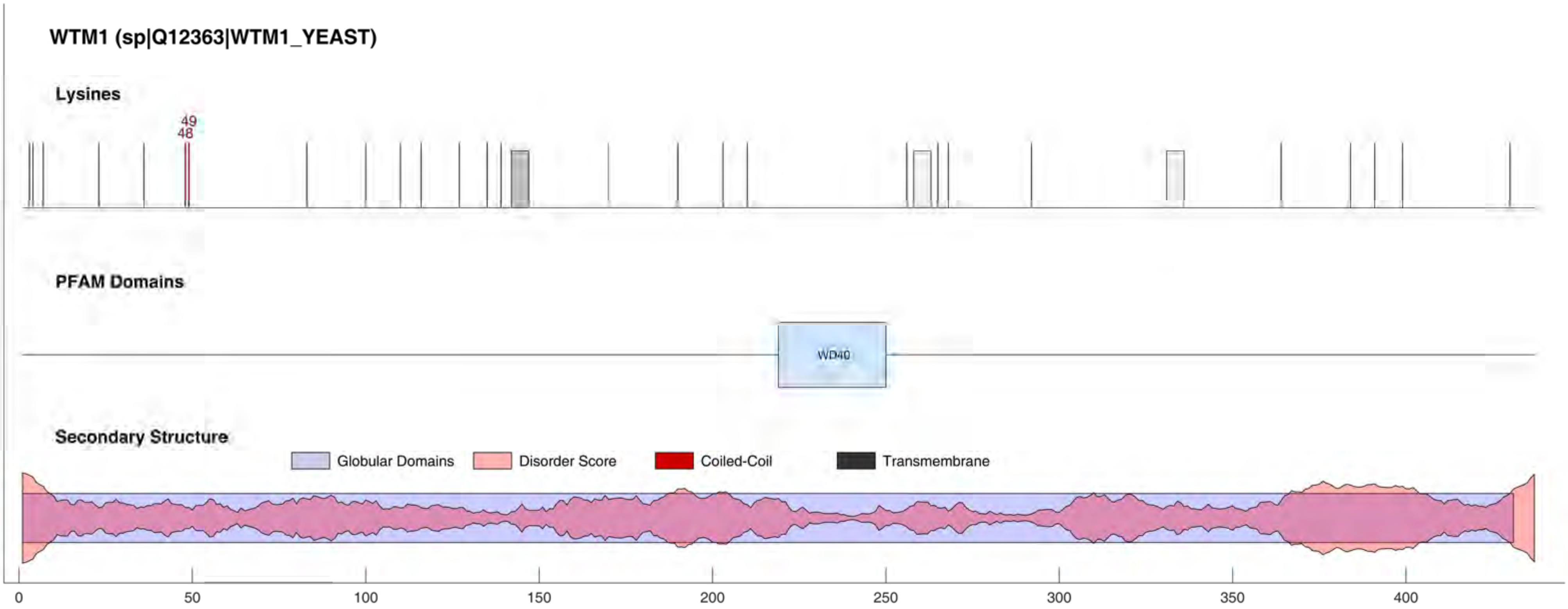

WTM1 (sp|Q12363|WTM1_YEAST)
Lysines
49
48
PFAM Domains
WD40
Secondary Structure
Globular Domains
Disorder Score
Coiled-Coil
Transmembrane
0
50
100
150
200
250
300
350
400

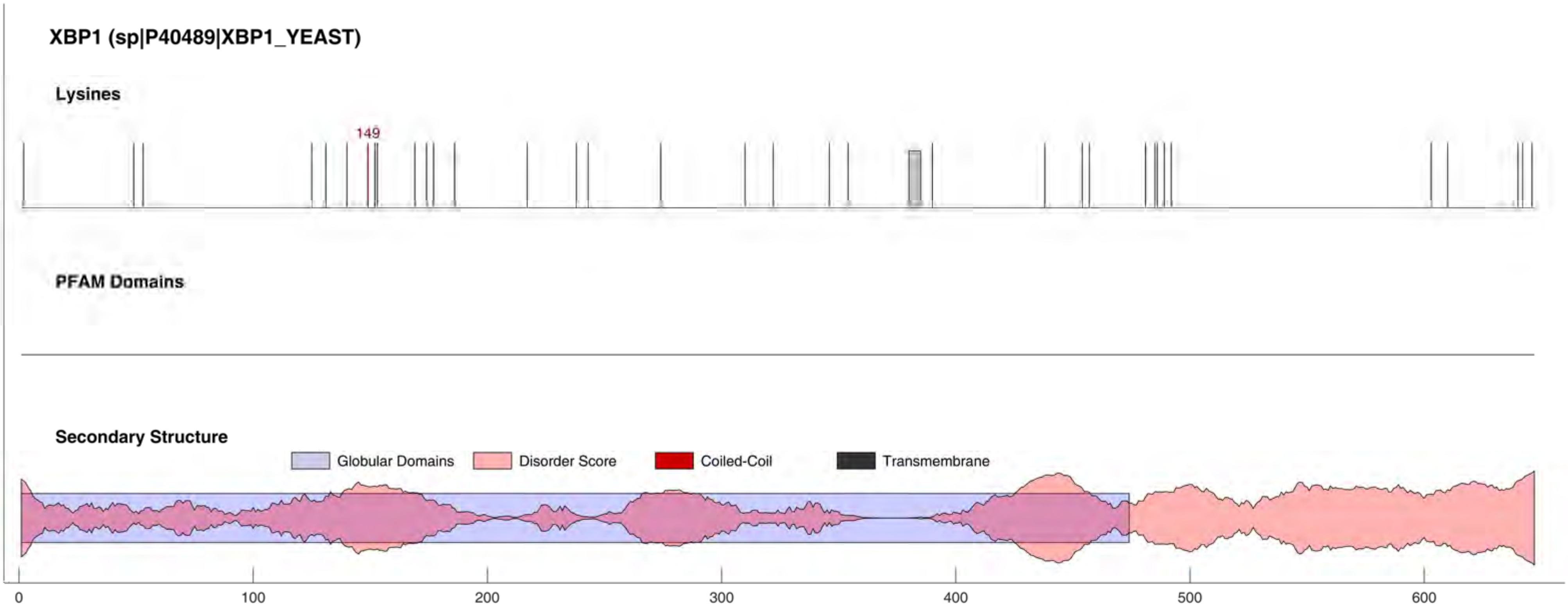
XBP1 (sp|P40489|XBP1_YEAST)
Lysines
149
PFAM Domains
Secondary Structure
Globular Domains
Disorder Score
Coiled-Coil
Transmembrane
0
100
200
300
400
500
600

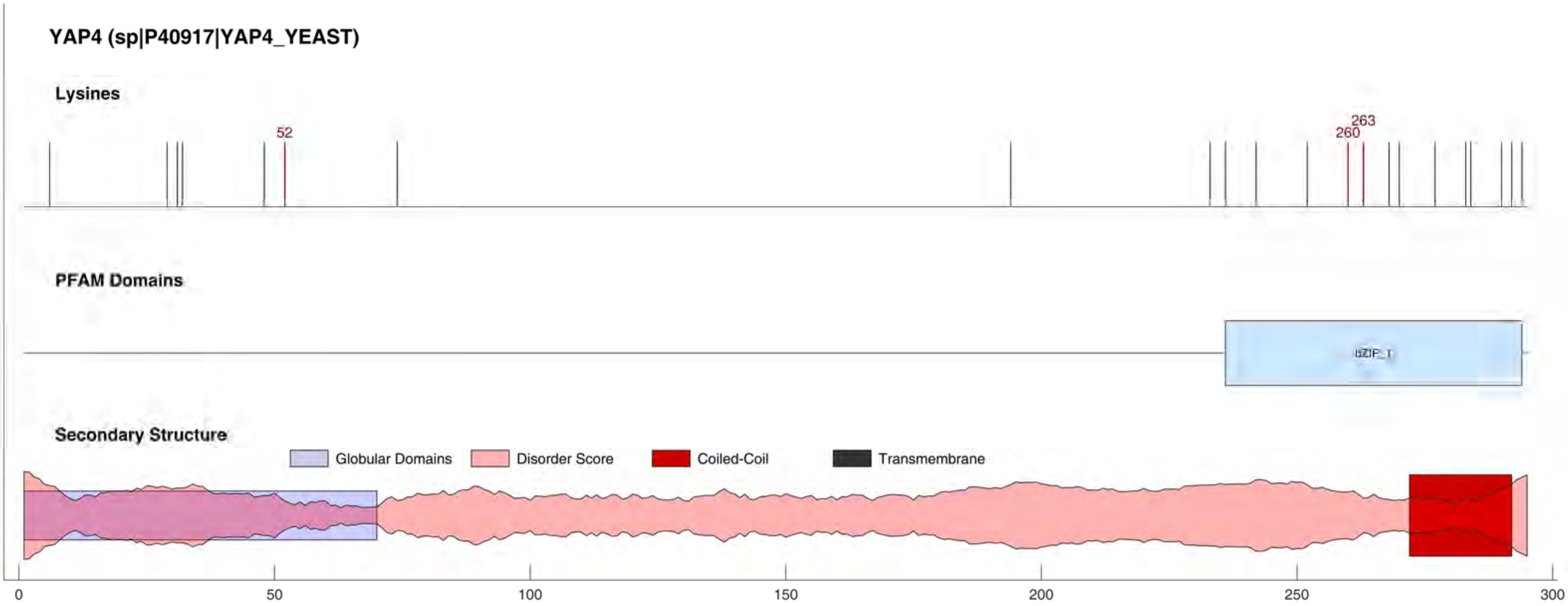
YAP4 (sp|P40917|YAP4_YEAST)
Lysines
52
263
260
PFAM Domains
bZIP_1
Secondary Structure
Globular Domains
Disorder Score
Coiled-Coil
Transmembrane
0
50
100
150
200
250
300

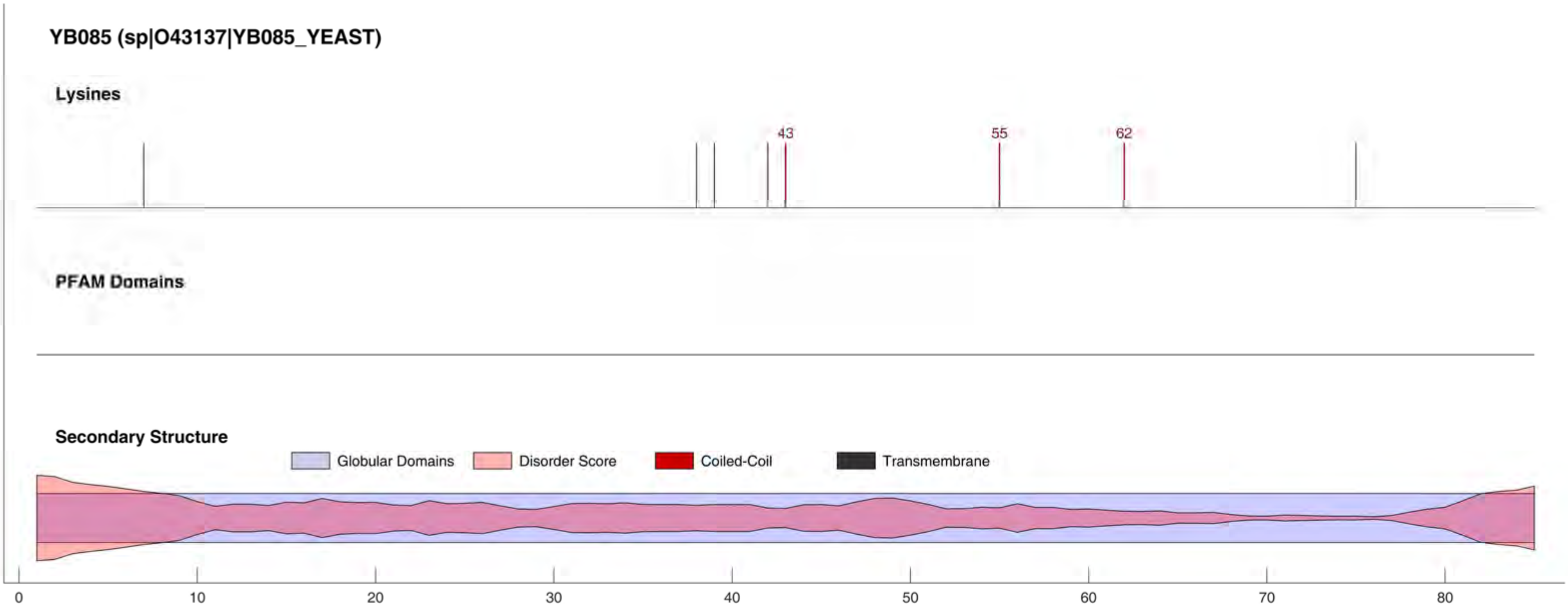
YB085 (sp|O43137|YB085_YEAST)
Lysines
43
55
62
PFAM Domains
Secondary Structure
Globular Domains
Disorder Score
Coiled-Coil
Transmembrane
0
10
20
30
40
50
60
70
80

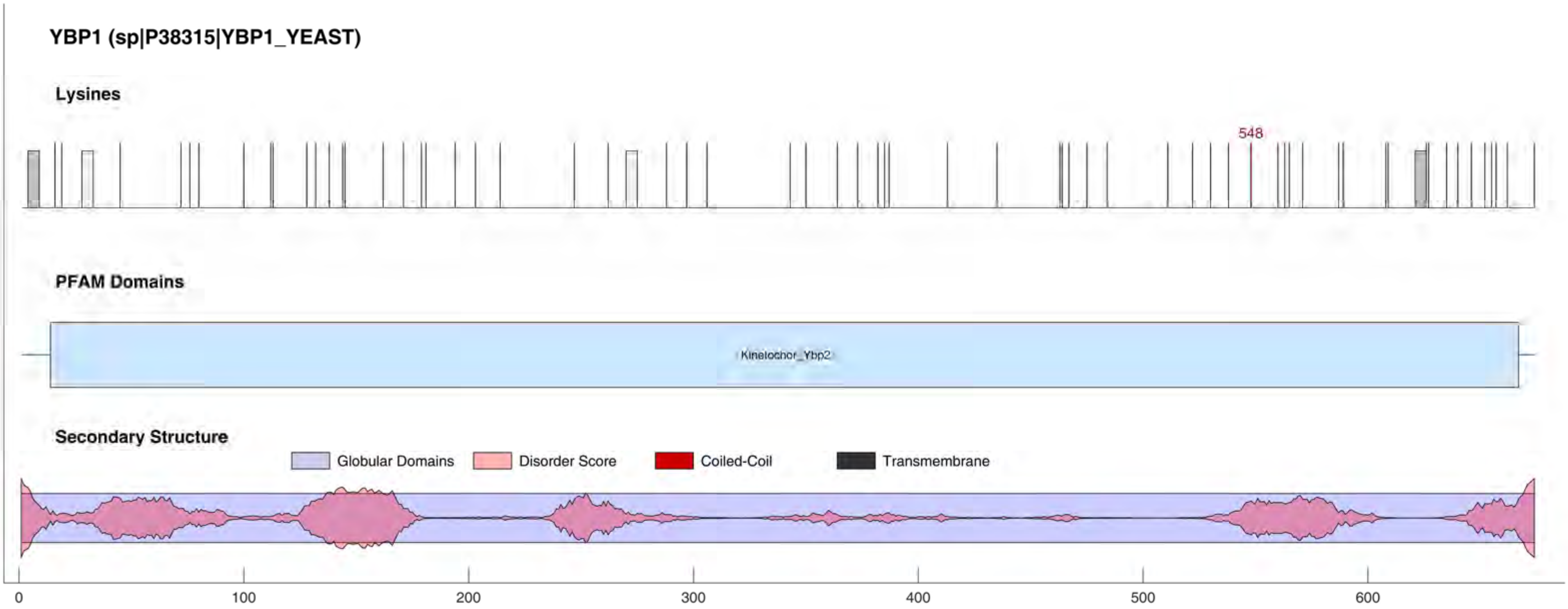

YBP1 (sp|P38315|YBP1_YEAST)
Lysines
548
PFAM Domains
Kinetochor_Ybp2
Secondary Structure
Globular Domains
Disorder Score
Coiled-Coil
Transmembrane
0
100
200
300
400
500
600

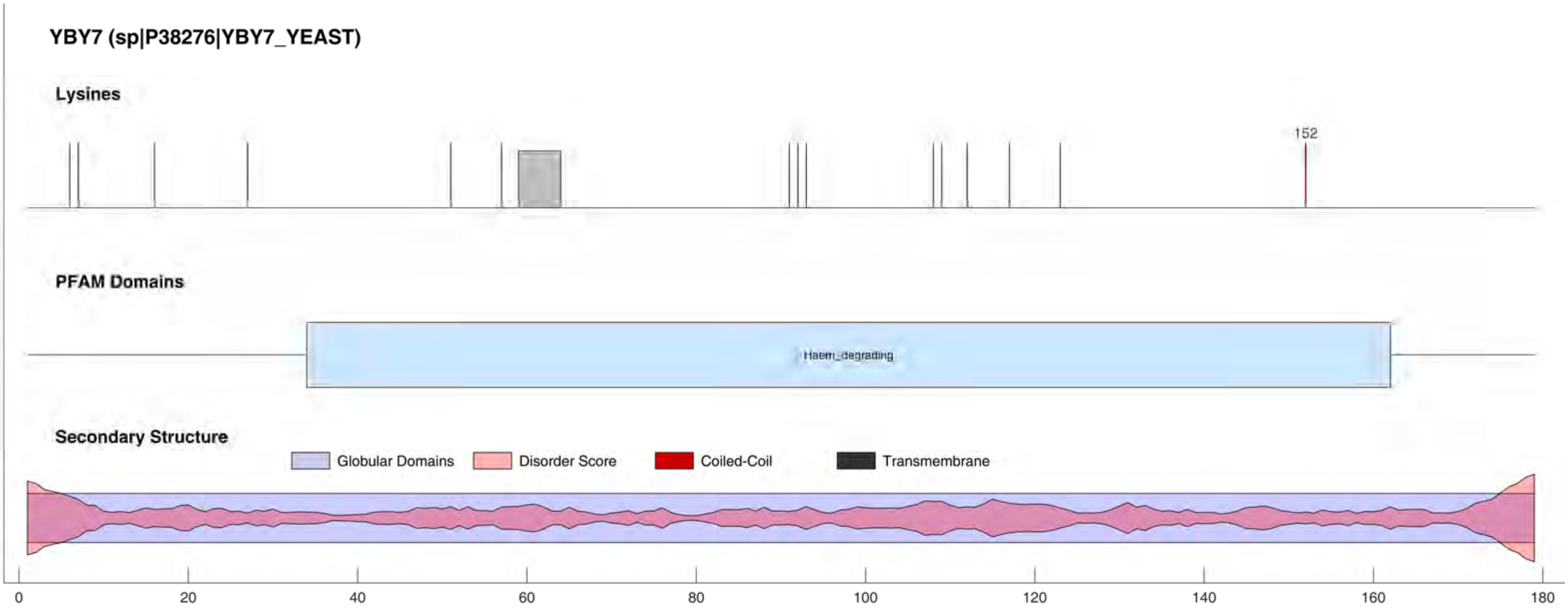
YBY7 (sp|P38276|YBY7_YEAST)
Lysines
152
PFAM Domains
Haem_degrading
Secondary Structure
Globular Domains
Disorder Score
Coiled-Coil
Transmembrane
0
20
40
60
80
100
120
140
160
180

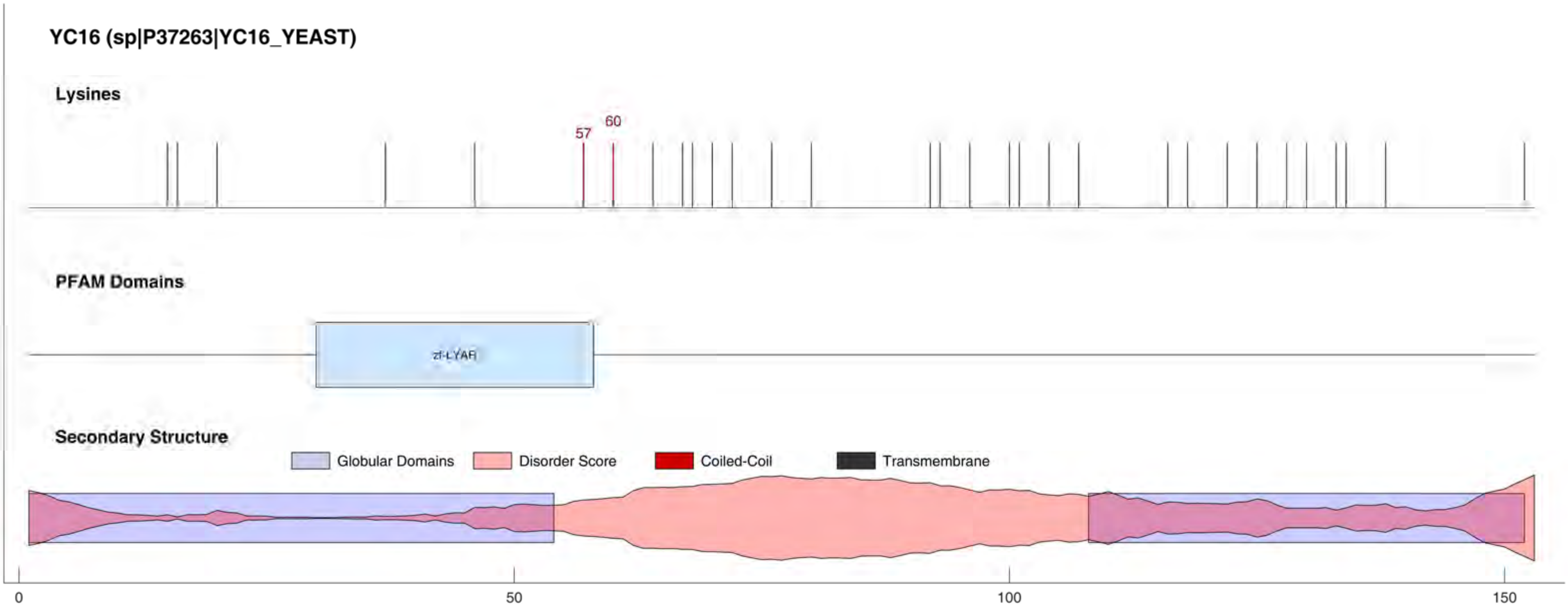
YC16 (sp|P37263|YC16_YEAST)
Lysines
57
60
PFAM Domains
zf-LYAR
Secondary Structure
Globular Domains
Disorder Score
Coiled-Coil
Transmembrane
0
50
100
150

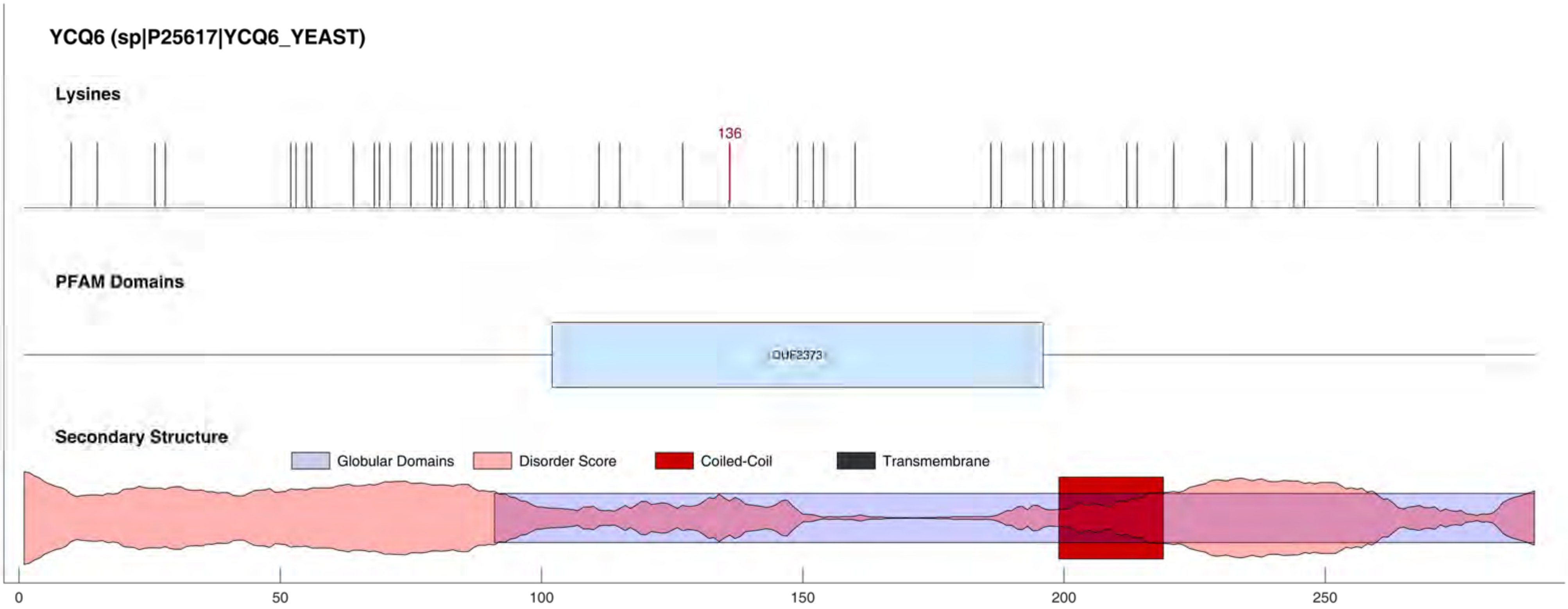
YCQ6 (sp|P25617|YCQ6_YEAST)
Lysines
136
PFAM Domains
DUF2373
Secondary Structure
Globular Domains
Disorder Score
Coiled-Coil
Transmembrane
0
50
100
150
200
250

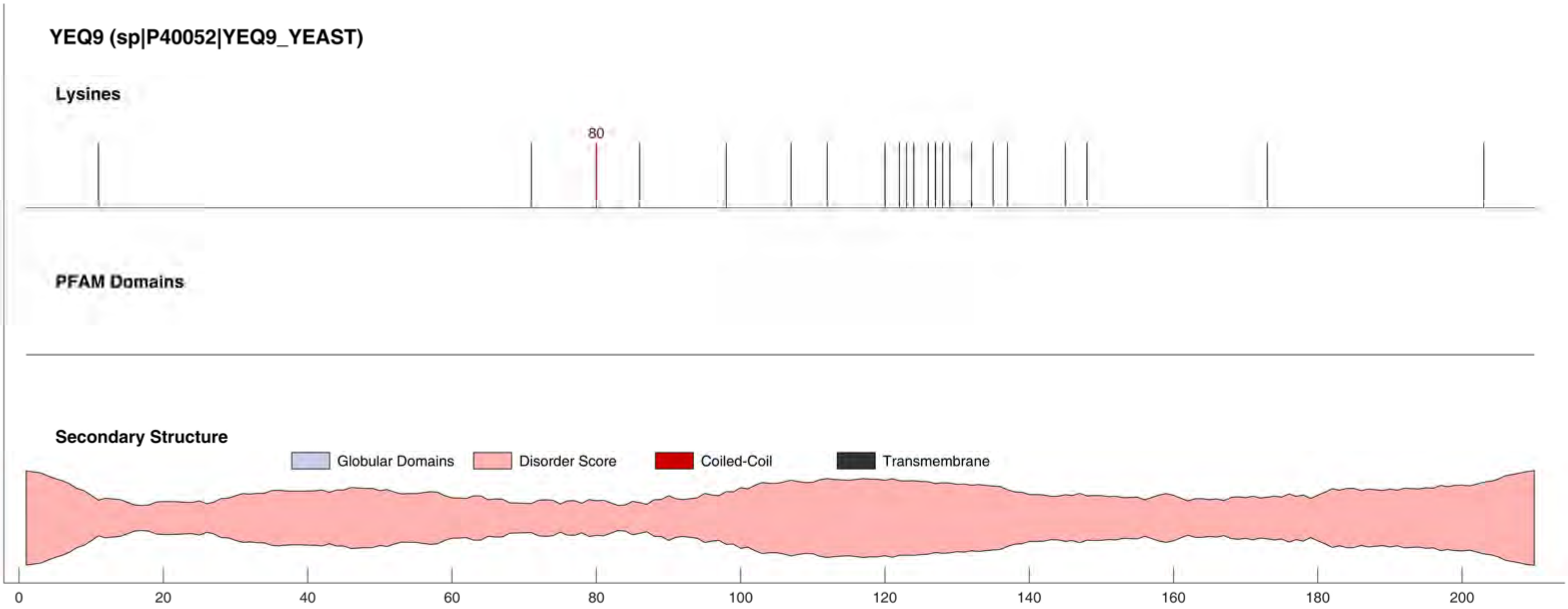

YEQ9 (sp|P40052|YEQ9_YEAST)
Lysines
80
PFAM Domains
Secondary Structure
Globular Domains
Disorder Score
Coiled-Coil
Transmembrane
0
20
40
60
80
100
120
140
160
180
200

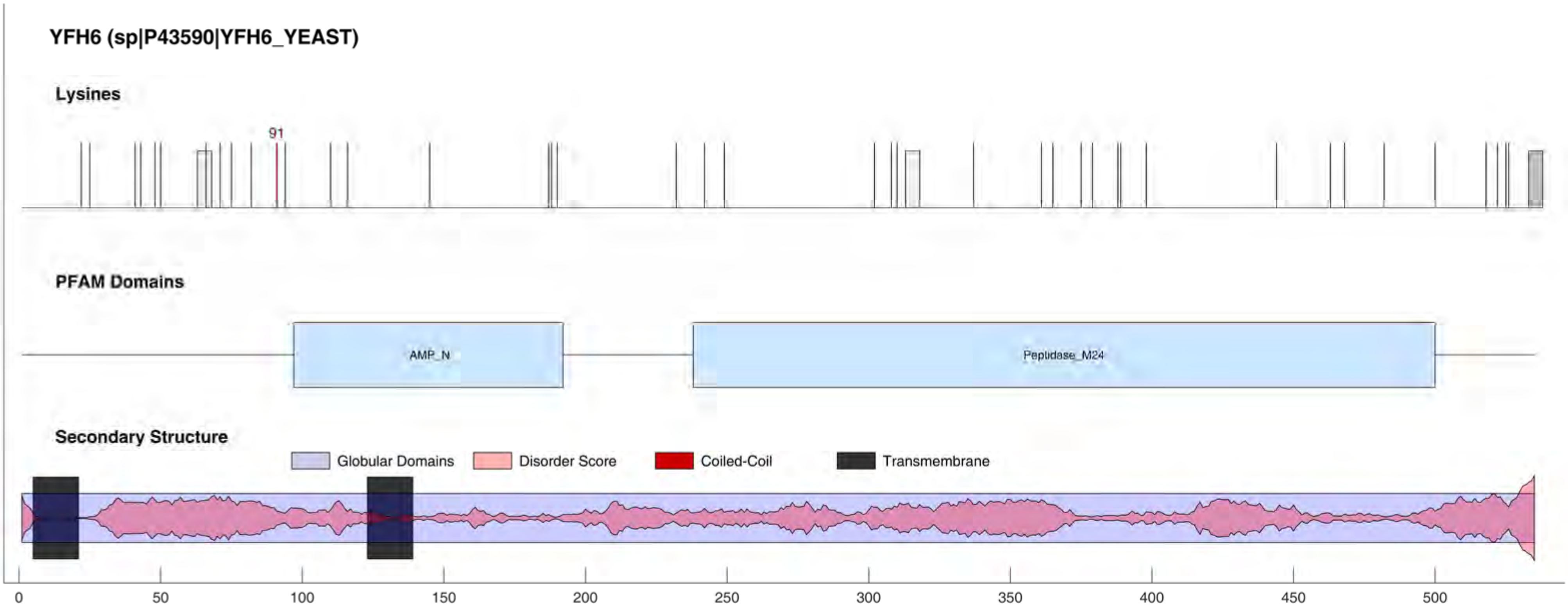
YFH6 (sp|P43590|YFH6_YEAST)
Lysines
91
PFAM Domains
AMP_N
Peptidase_M24
Secondary Structure
Globular Domains
Disorder Score
Coiled-Coil
Transmembrane
0
50
100
150
200
250
300
350
400
450
500

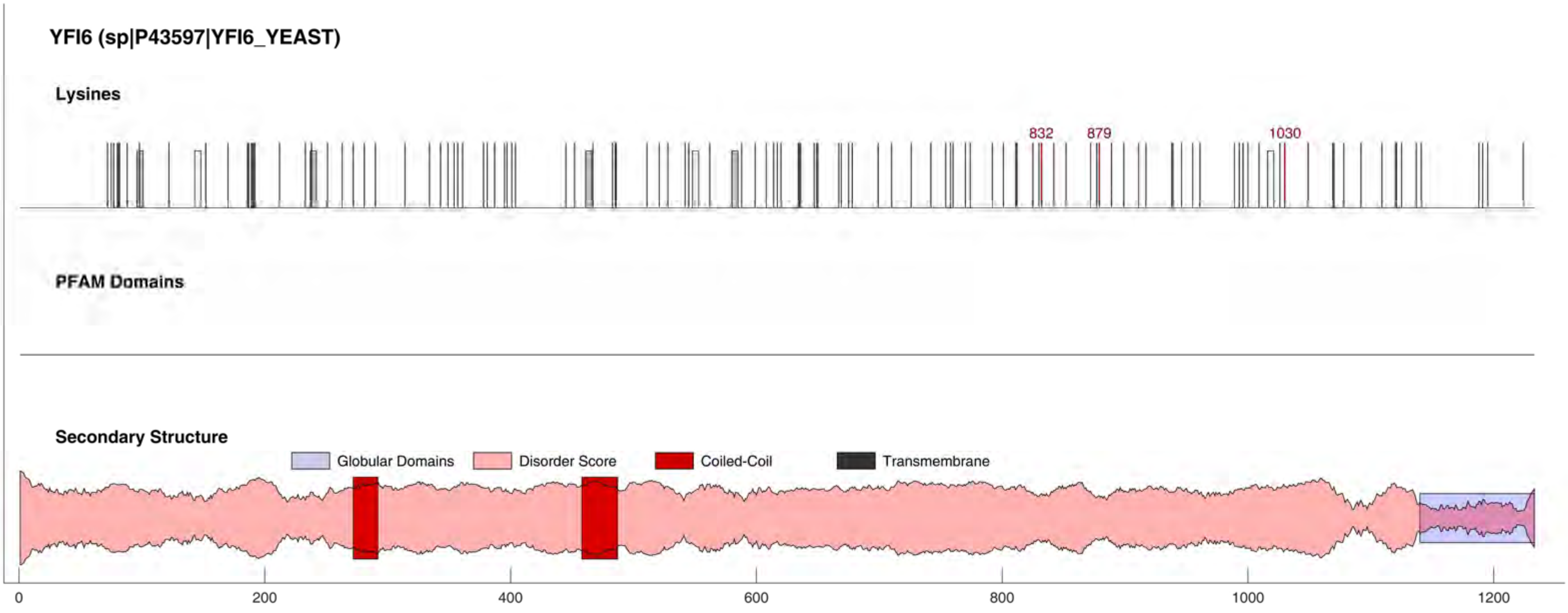
YFI6 (sp|P43597|YFI6_YEAST)
Lysines
832
879
1030
PFAM Domains
Secondary Structure
Globular Domains
Disorder Score
Coiled-Coil
Transmembrane
0
200
400
600
800
1000
1200

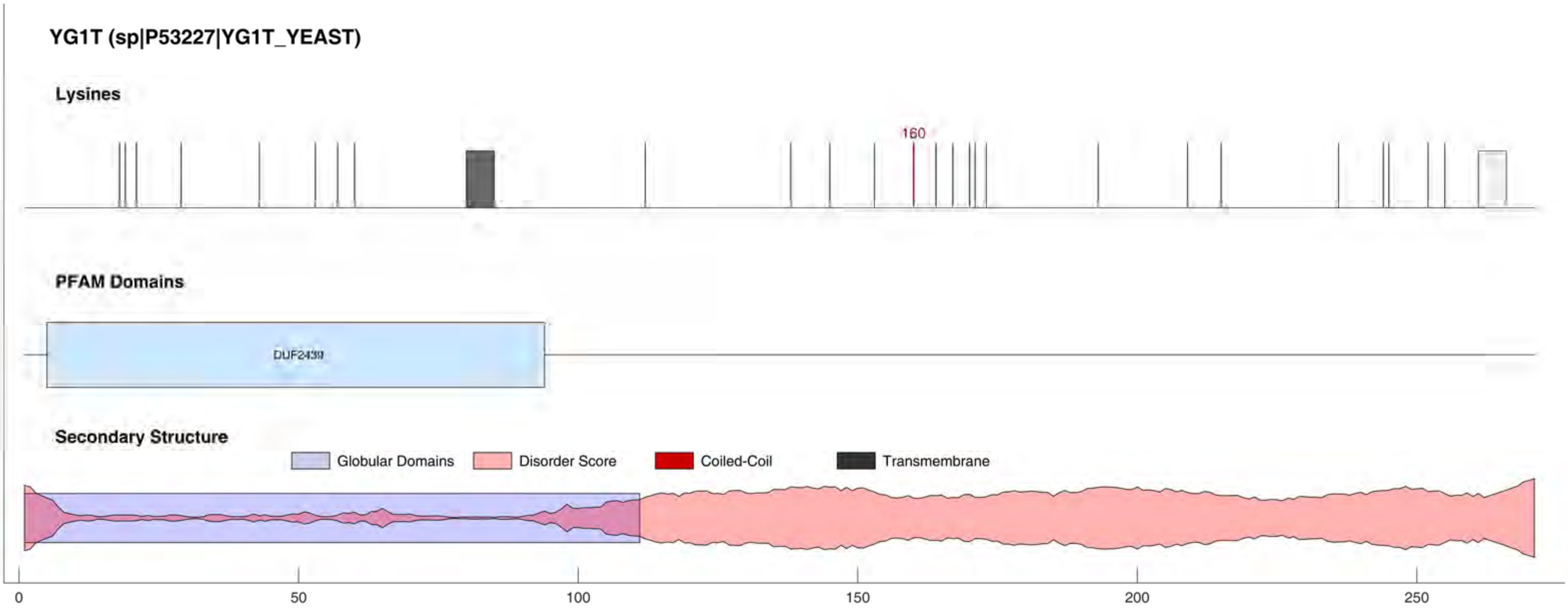

YG1T (sp|P53227|YG1T_YEAST)
Lysines
160
PFAM Domains
DUF2439
Secondary Structure
Globular Domains
Disorder Score
Coiled-Coil
Transmembrane
0
50
100
150
200
250

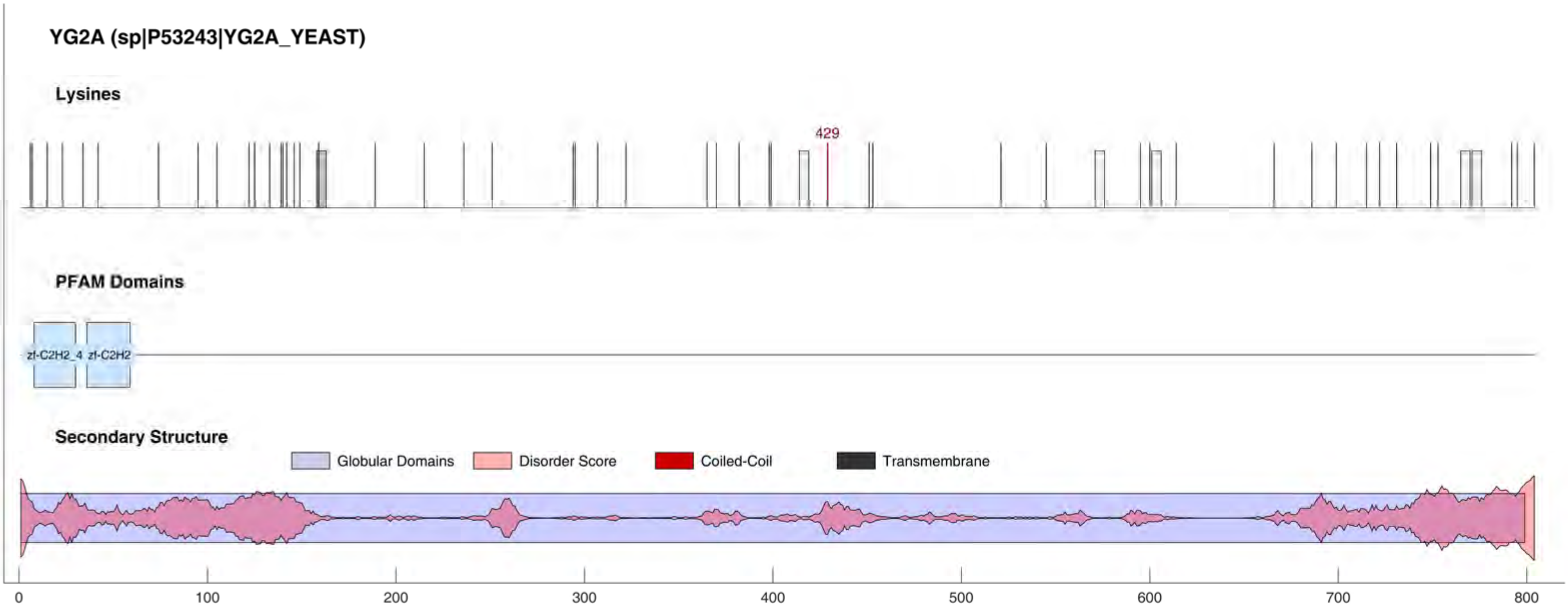

YG2A (sp|P53243|YG2A_YEAST)
Lysines
429
PFAM Domains
zf-C2H2_4
zf-C2H2
Secondary Structure
Globular Domains
Disorder Score
Coiled-Coil
Transmembrane
0
100
200
300
400
500
600
700
800

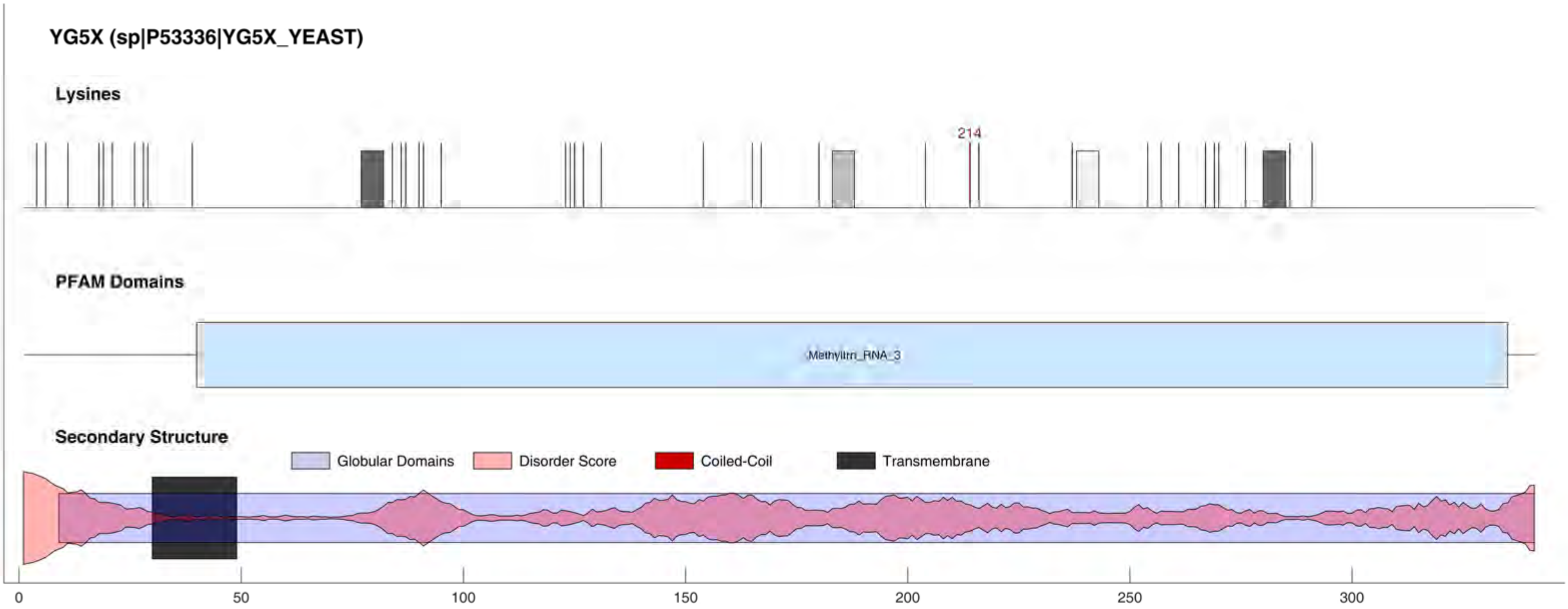
YG5X (sp|P53336|YG5X_YEAST)
Lysines
214
PFAM Domains
Methyltrn_RNA_3
Secondary Structure
Globular Domains
Disorder Score
Coiled-Coil
Transmembrane
0
50
100
150
200
250
300

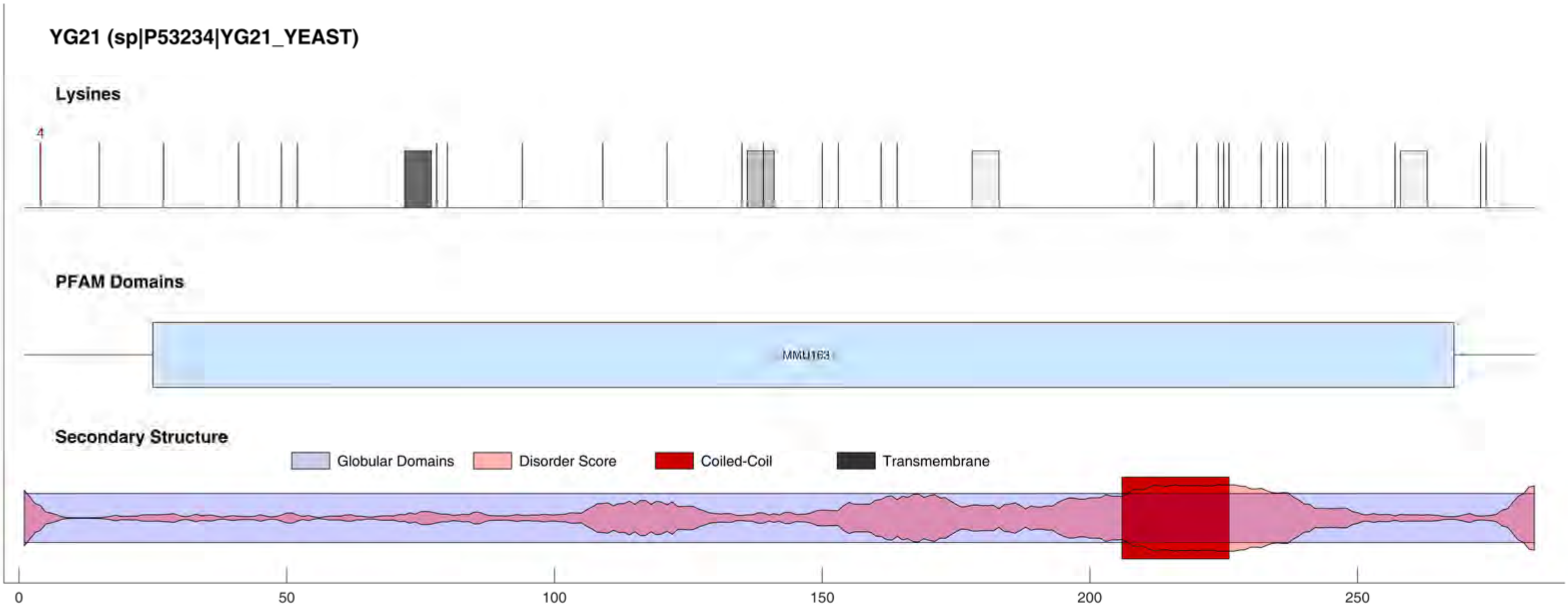

YG21 (sp|P53234|YG21_YEAST)
Lysines
4
PFAM Domains
MMU163
Secondary Structure
Globular Domains
Disorder Score
Coiled-Coil
Transmembrane
0
50
100
150
200
250

#### YG51 (sp|P50089|YG51_YEAST)

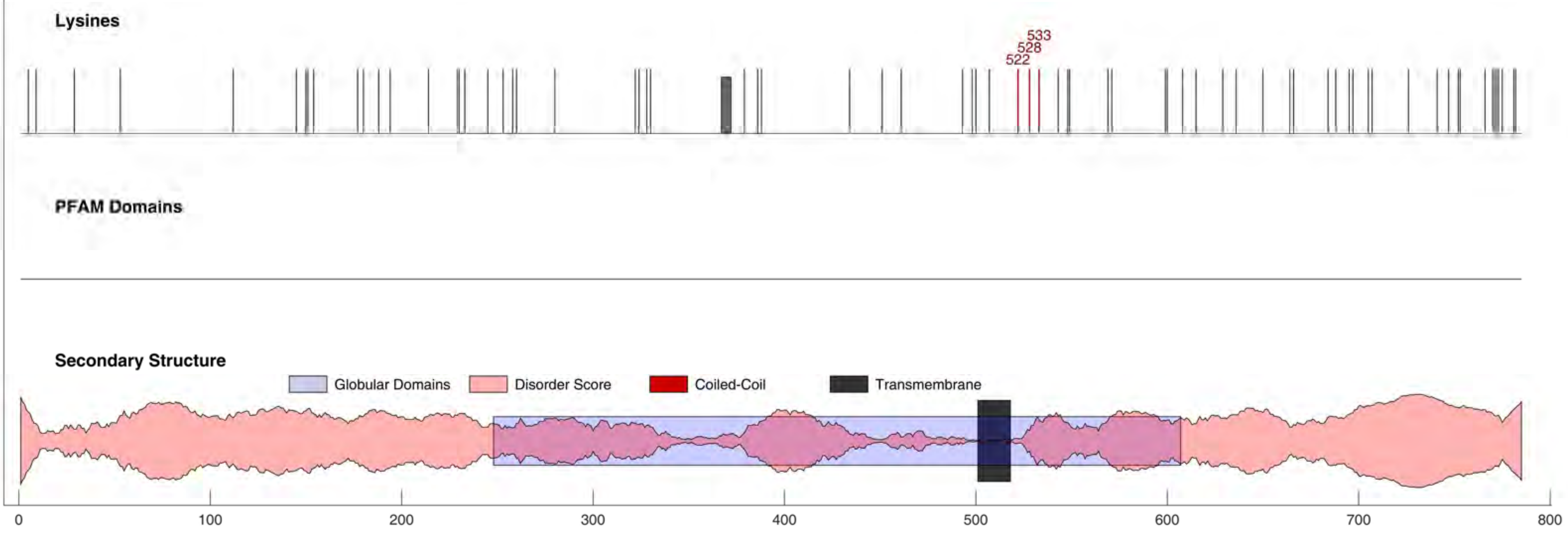

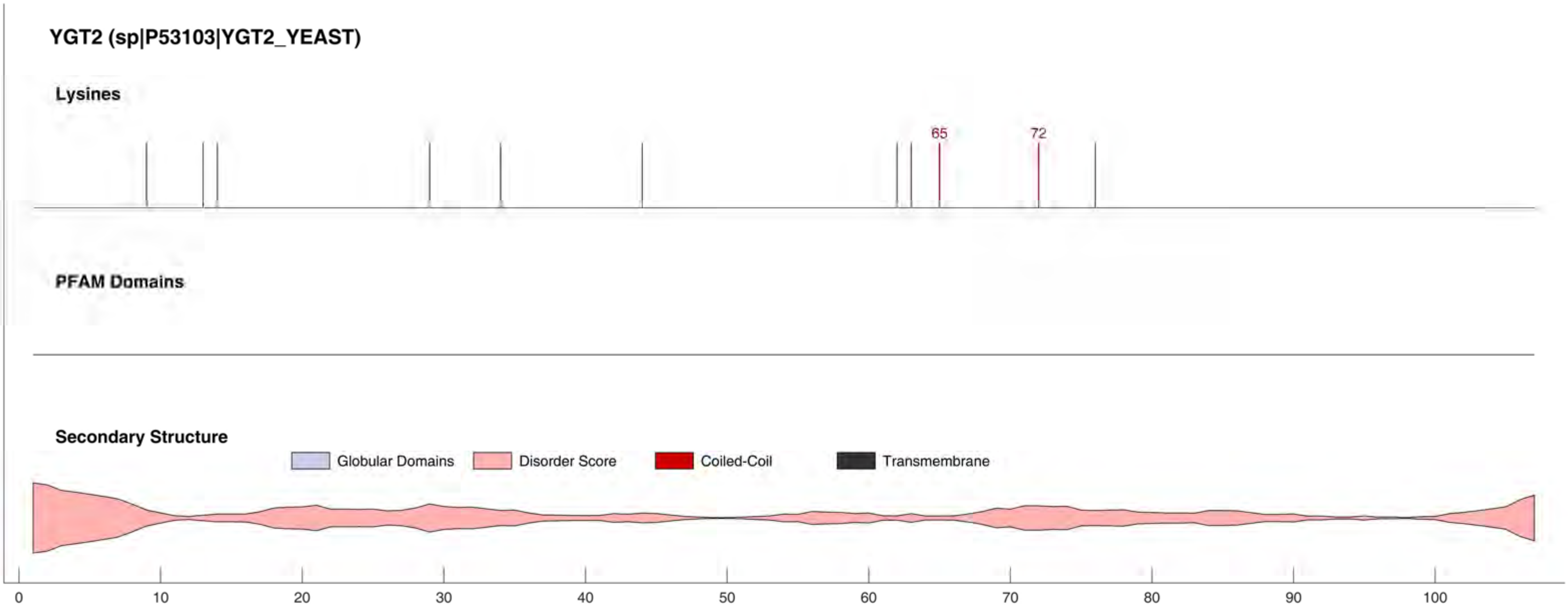

YGT2 (sp|P53103|YGT2_YEAST)
Lysines
65
72
PFAM Domains
Secondary Structure
Globular Domains
Disorder Score
Coiled-Coil
Transmembrane
0
10
20
30
40
50
60
70
80
90
100

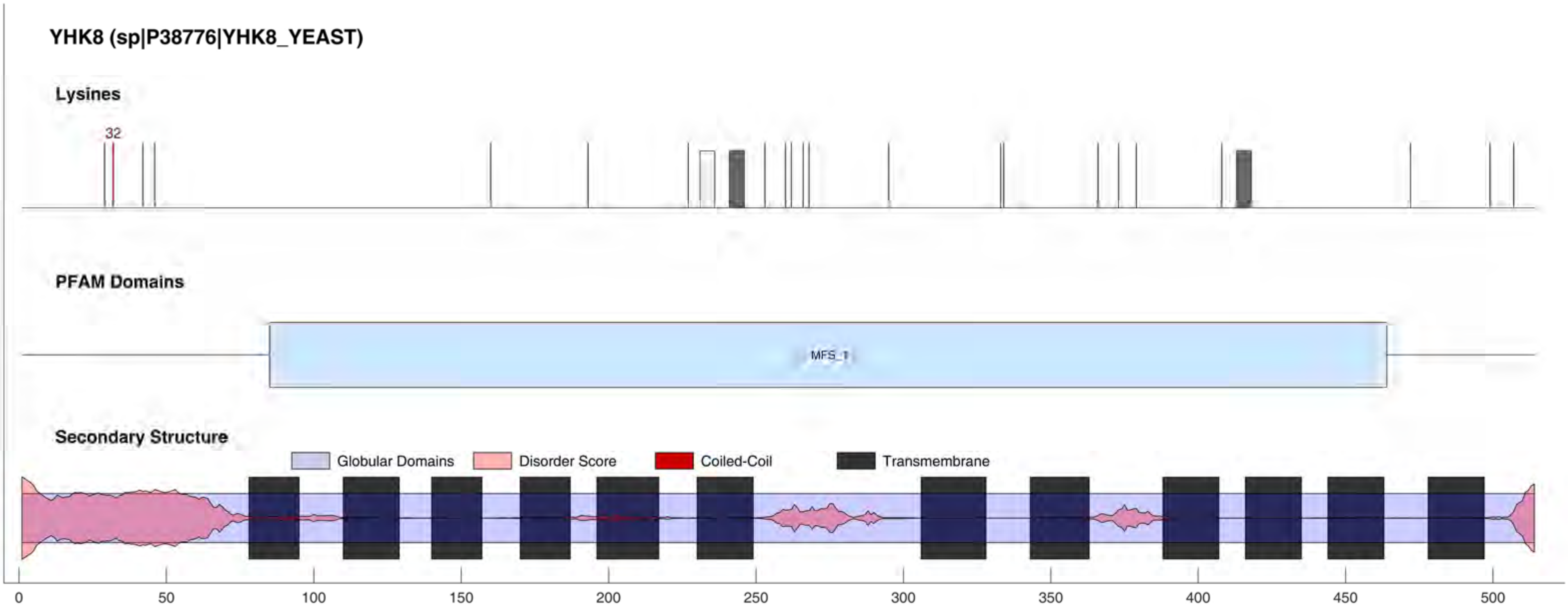

YHK8 (sp|P38776|YHK8_YEAST)
Lysines
32
PFAM Domains
MFS_1
Secondary Structure
Globular Domains
Disorder Score
Coiled-Coil
Transmembrane
0
50
100
150
200
250
300
350
400
450
500

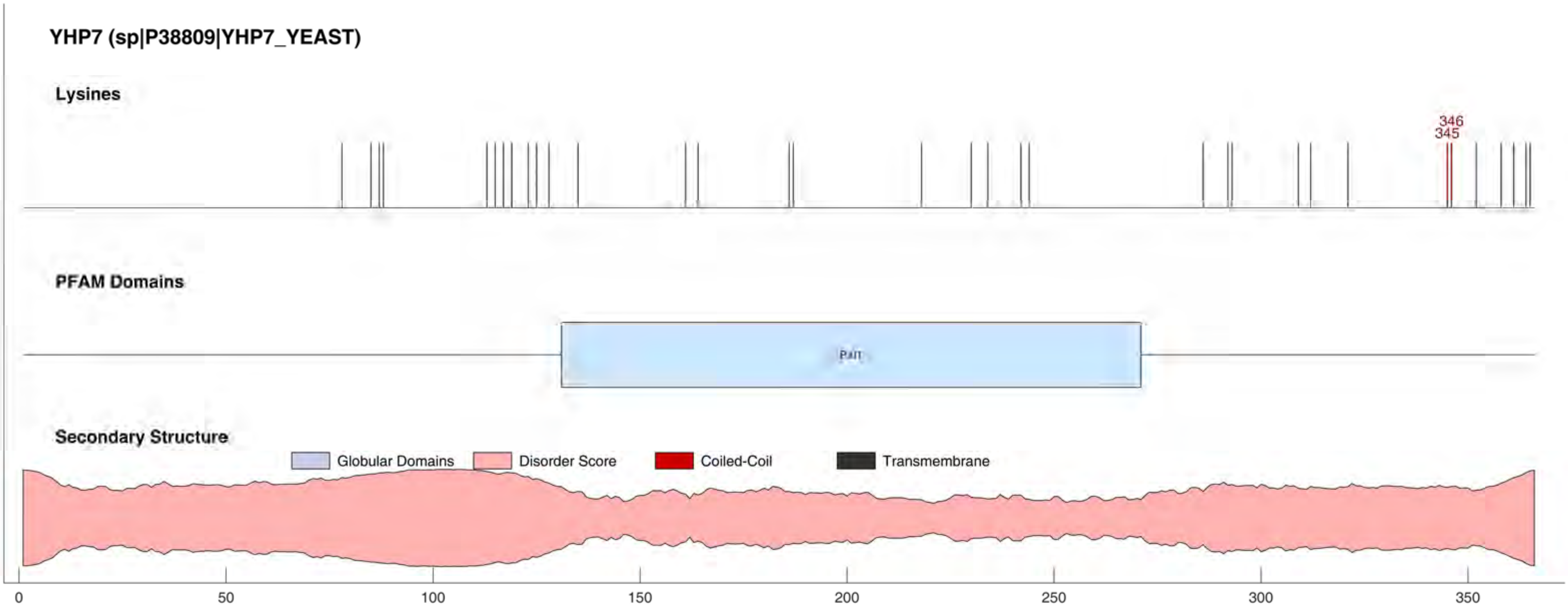
YHP7 (sp|P38809|YHP7_YEAST)
Lysines
346
345
PFAM Domains
Pat
Secondary Structure
Globular Domains
Disorder Score
Coiled-Coil
Transmembrane
0
50
100
150
200
250
300
350

#### YIC4 (sp|P40543|YIC4_YEAST)

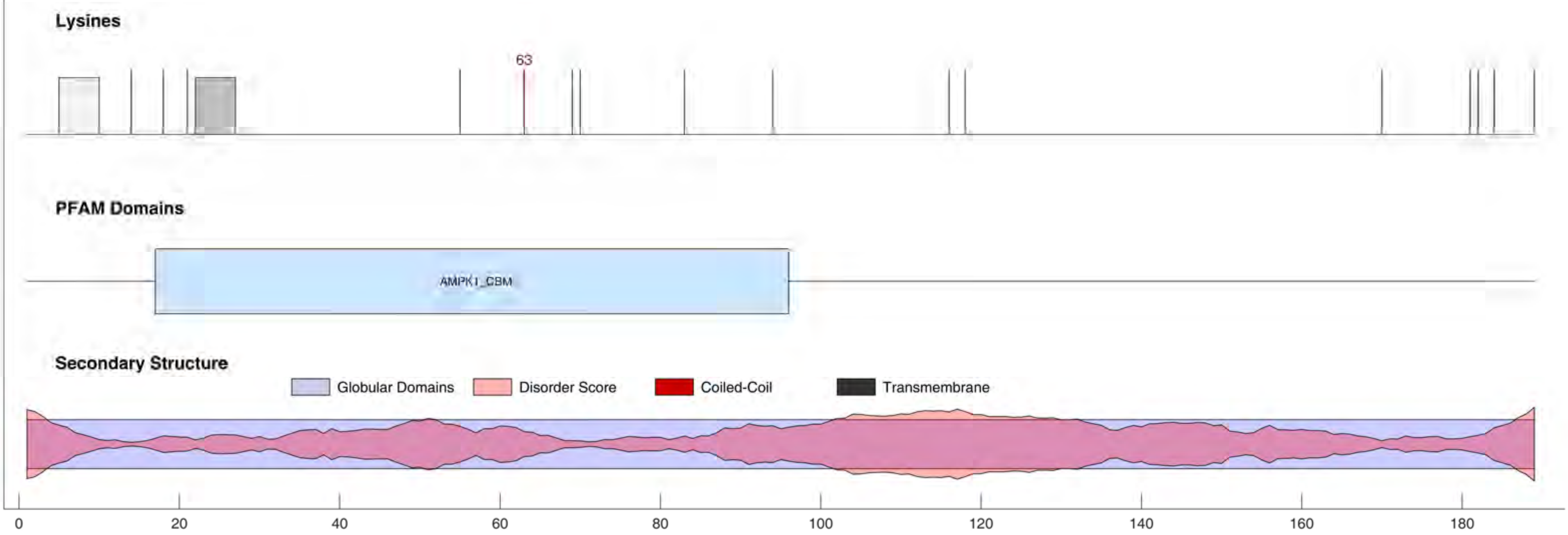

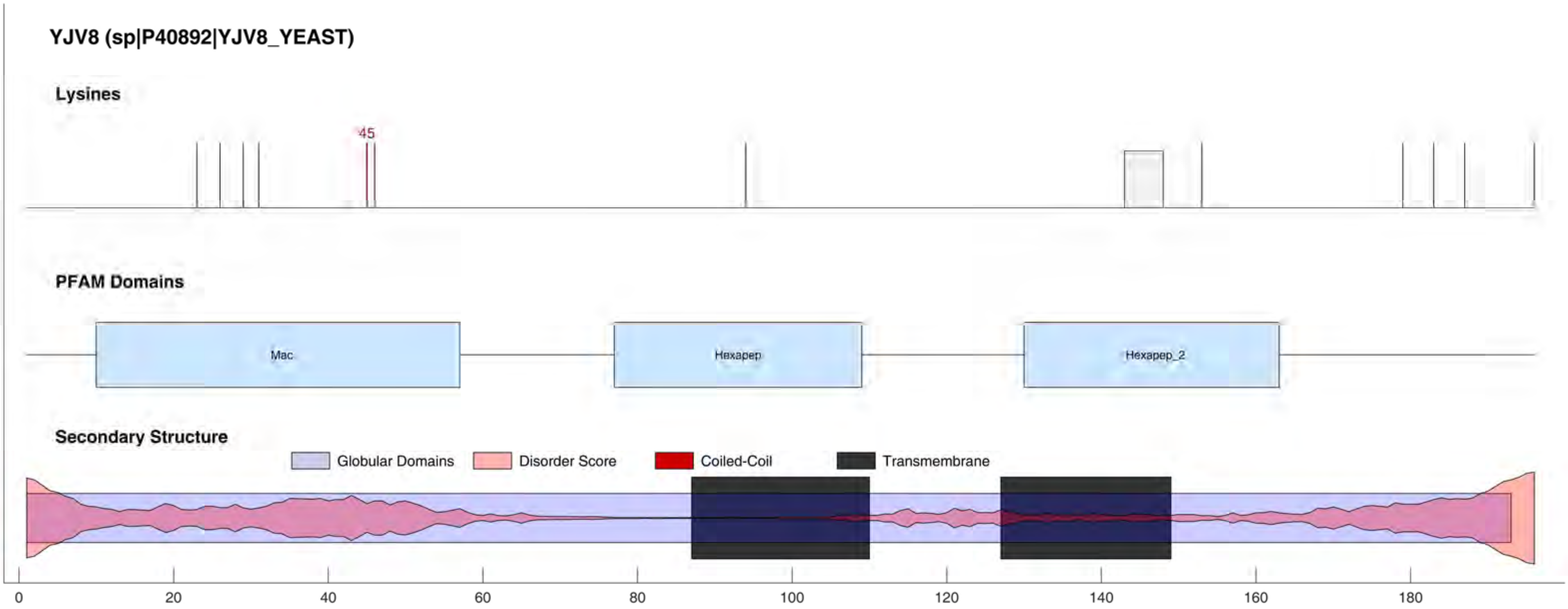

YJV8 (sp|P40892|YJV8_YEAST)
Lysines
45
PFAM Domains
Mac
Hexapep
Hexapep_2
Secondary Structure
Globular Domains
Disorder Score
Coiled-Coil
Transmembrane
0
20
40
60
80
100
120
140
160
180

#### YKZ1 (sp|Q02209|YKZ1_YEAST)

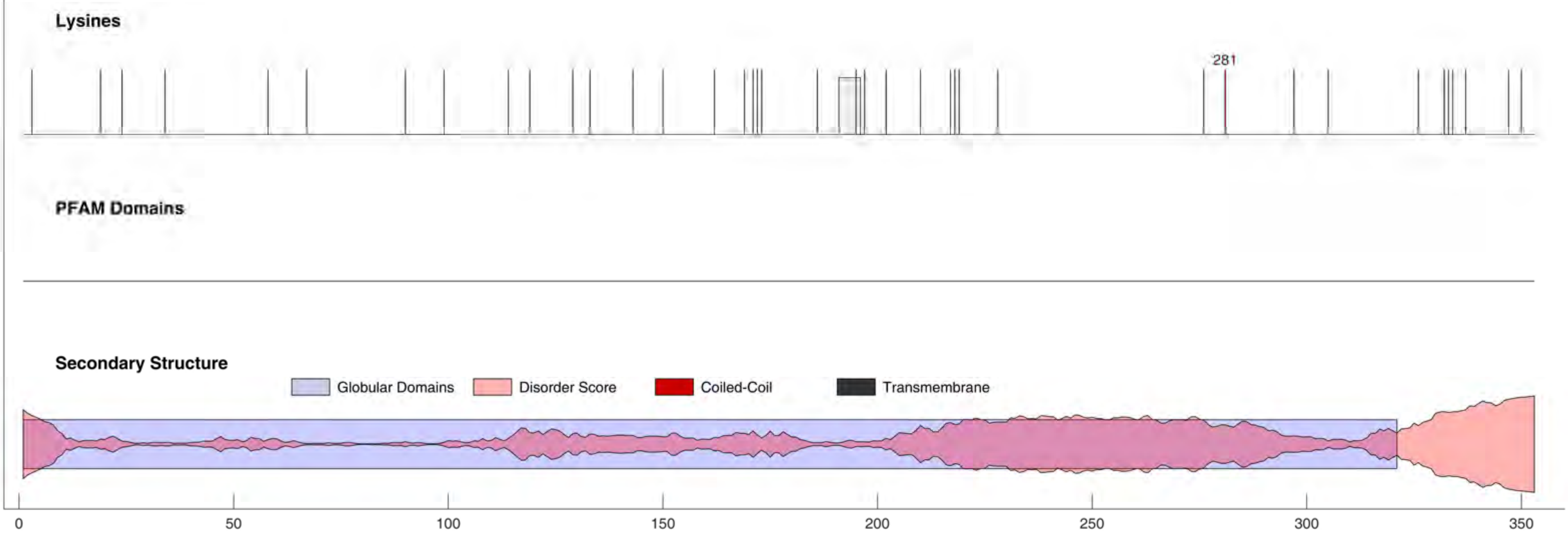

YL054 (sp|Q12244|YL054_YEAST)

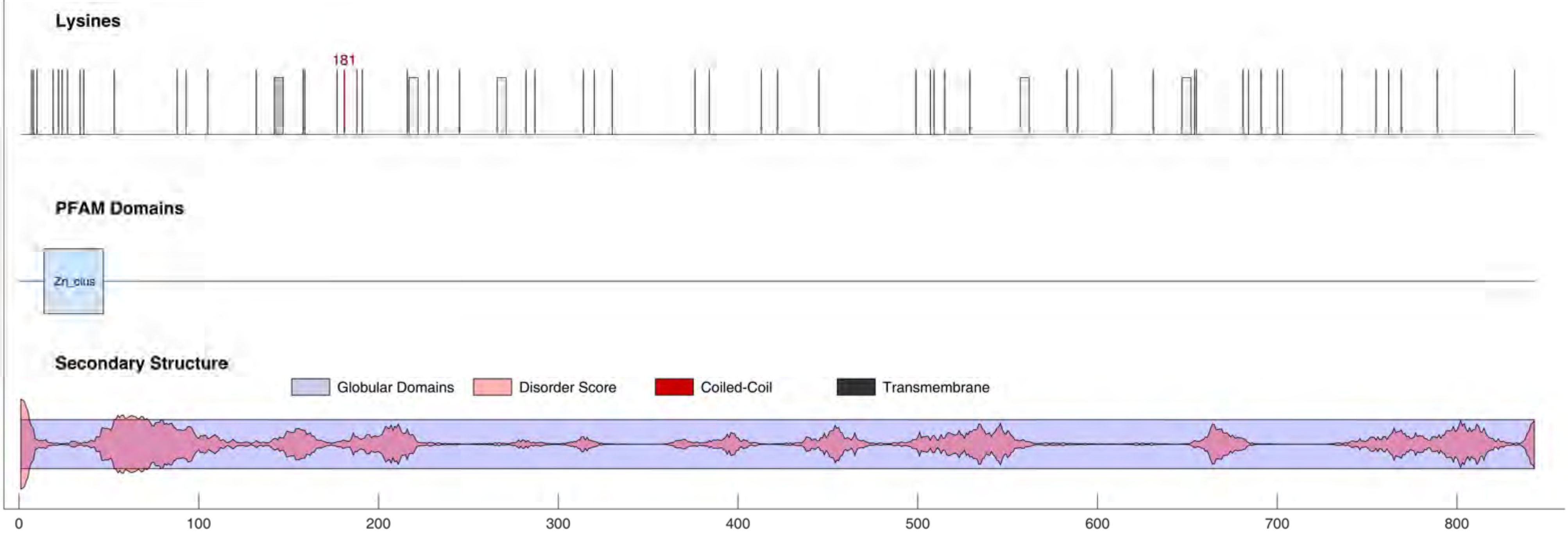

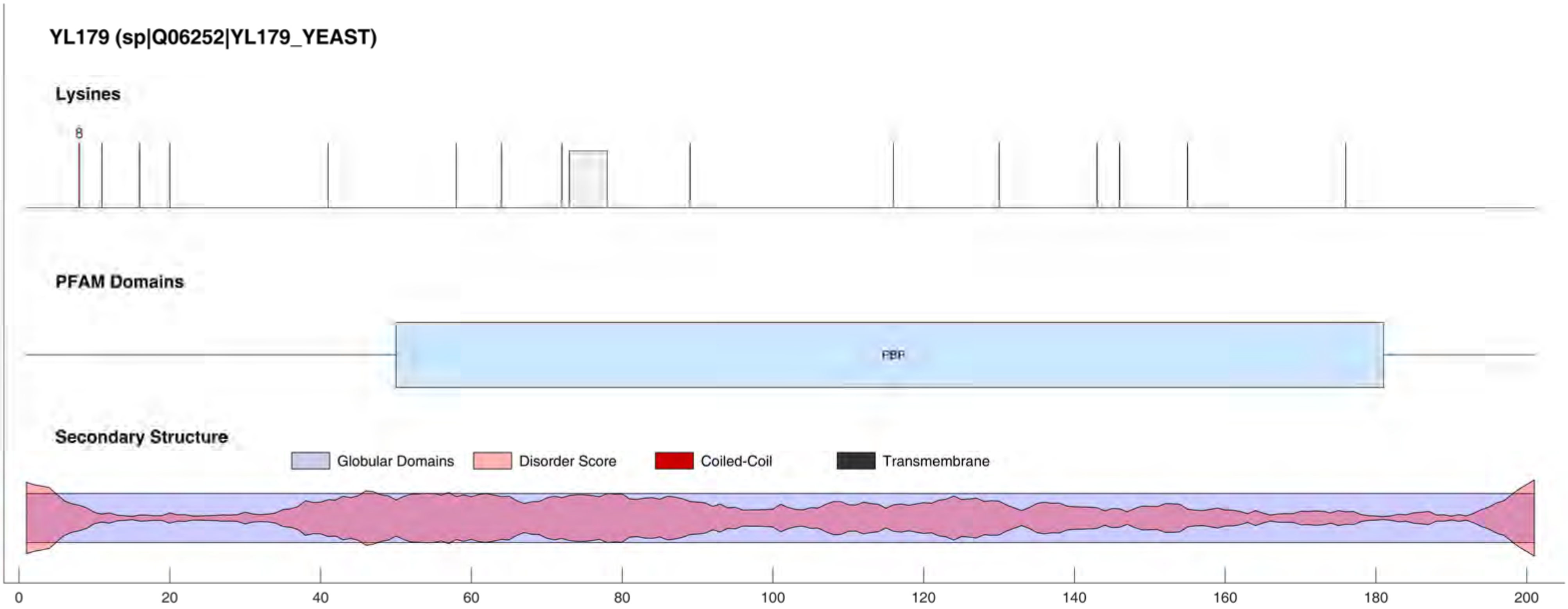

YL179 (sp|Q06252|YL179_YEAST)
Lysines
8
PFAM Domains
PBP
Secondary Structure
Globular Domains
Disorder Score
Coiled-Coil
Transmembrane
0
20
40
60
80
100
120
140
160
180
200

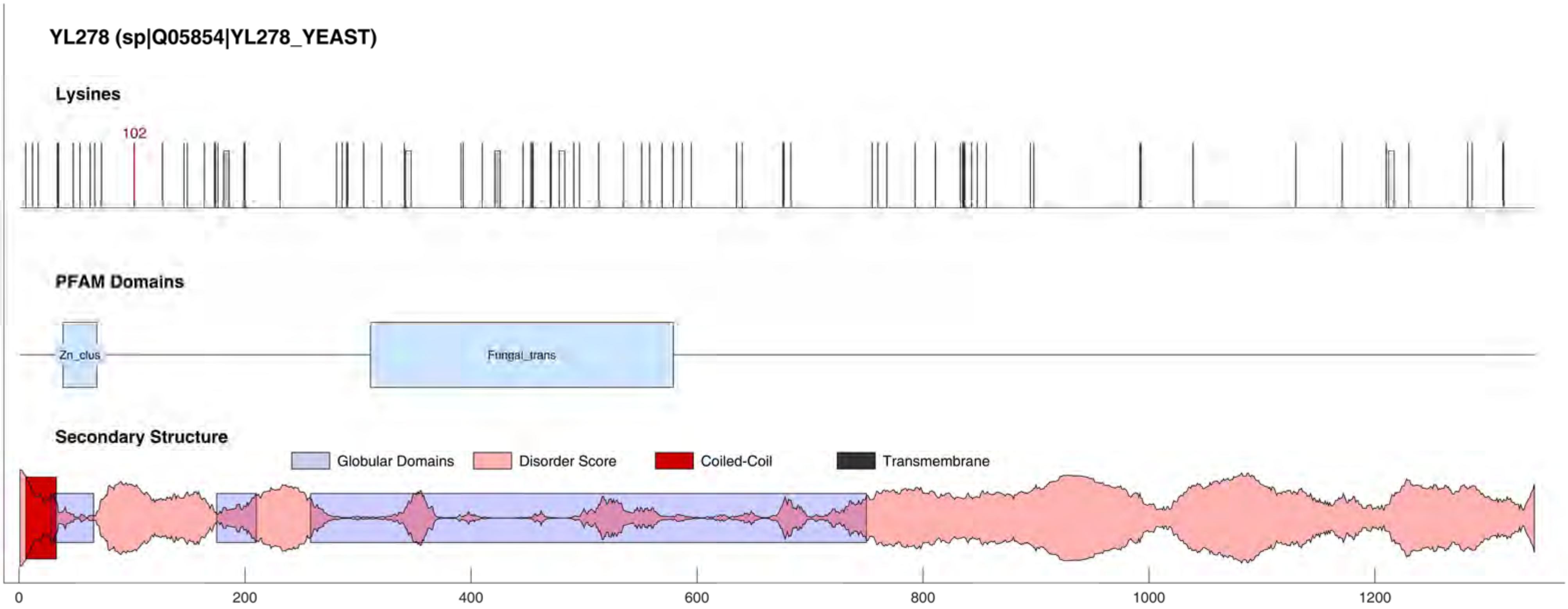
YL278 (sp|Q05854|YL278_YEAST)
Lysines
102
PFAM Domains
Zn_clus
Fungal_trans
Secondary Structure
Globular Domains
Disorder Score
Coiled-Coil
Transmembrane
0
200
400
600
800
1000
1200

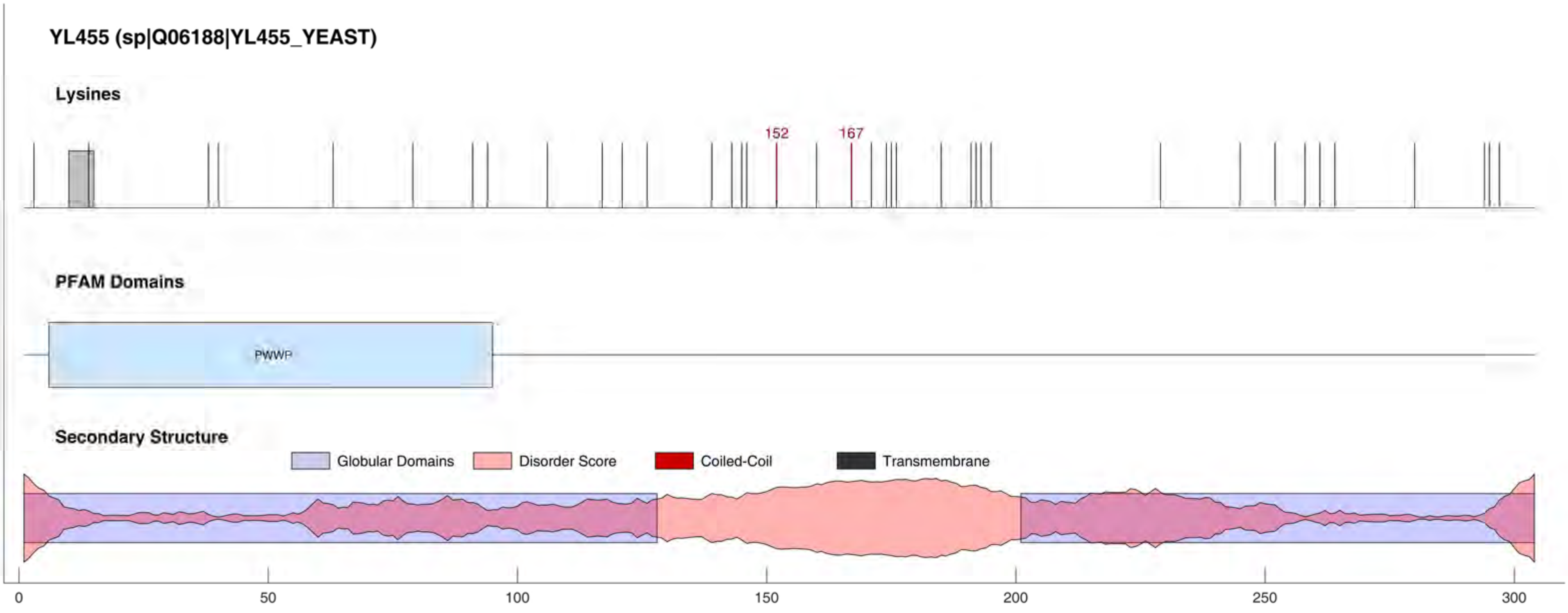

YL455 (sp|Q06188|YL455_YEAST)
Lysines
152
167
PFAM Domains
PWWP
Secondary Structure
Globular Domains
Disorder Score
Coiled-Coil
Transmembrane
0
50
100
150
200
250
300

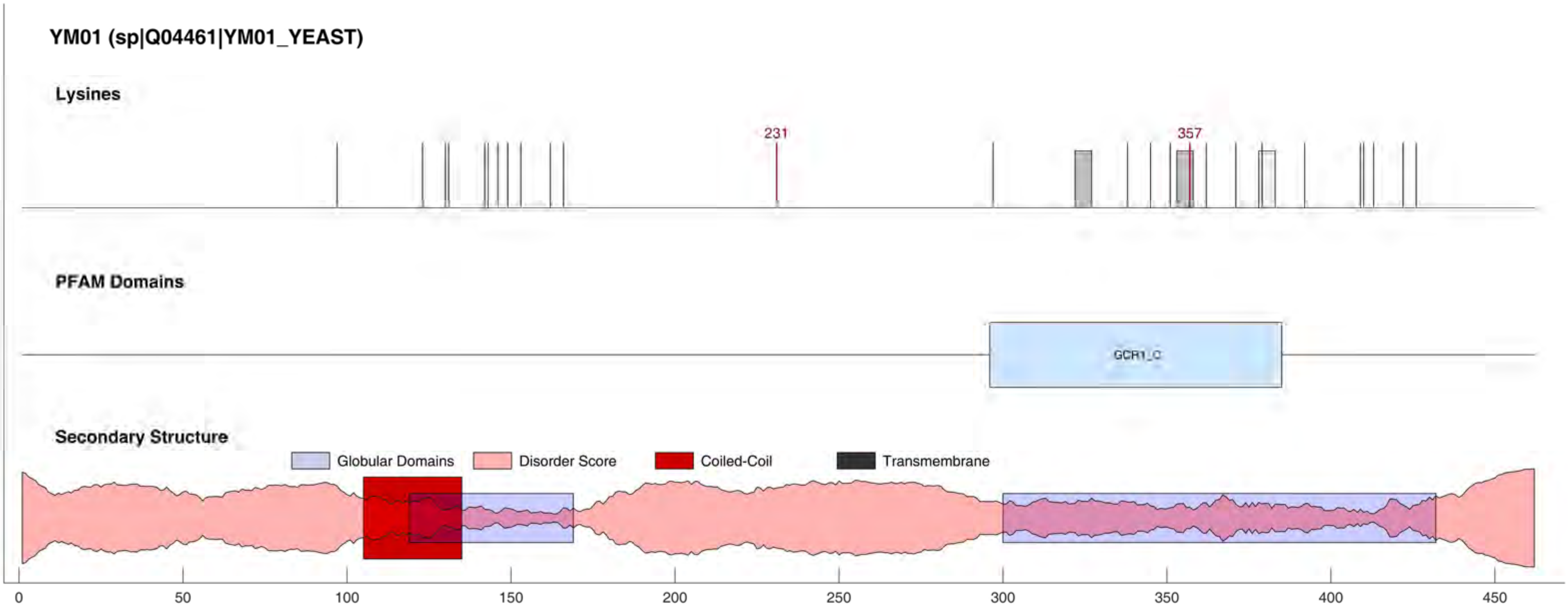
YM01 (sp|Q04461|YM01_YEAST)
Lysines
231
357
PFAM Domains
GCR1_C
Secondary Structure
Globular Domains
Disorder Score
Coiled-Coil
Transmembrane
0
50
100
150
200
250
300
350
400
450

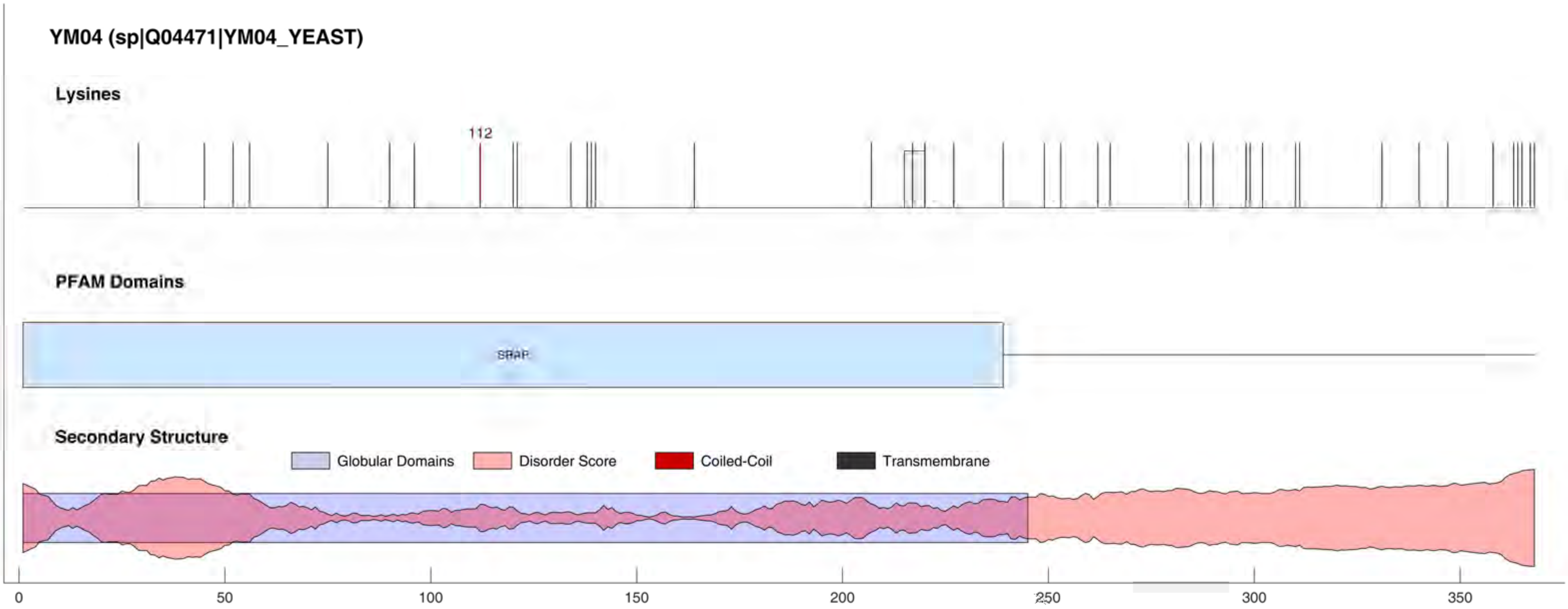

YM04 (sp|Q04471|YM04_YEAST)
Lysines
112
PFAM Domains
SRAP
Secondary Structure
Globular Domains
Disorder Score
Coiled-Coil
Transmembrane
0
50
100
150
200
250
300
350

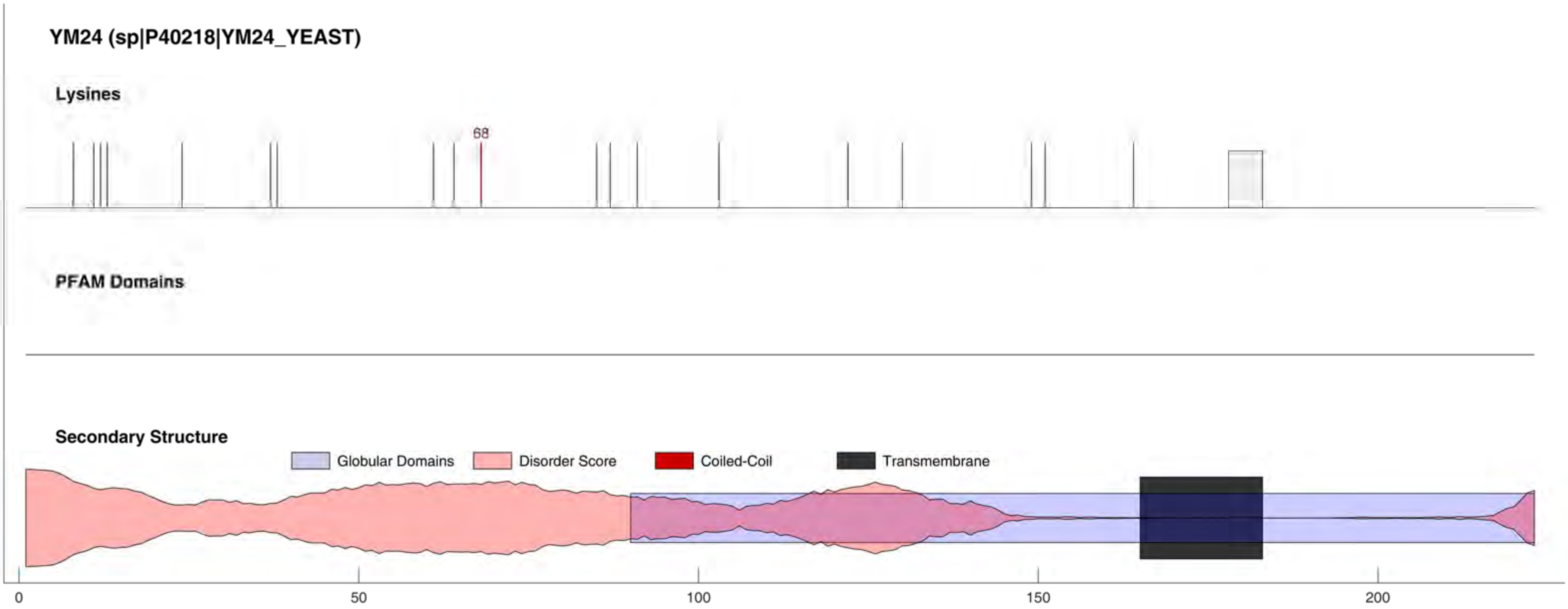
YM24 (sp|P40218|YM24_YEAST)
Lysines
68
PFAM Domains
Secondary Structure
Globular Domains
Disorder Score
Coiled-Coil
Transmembrane
0
50
100
150
200

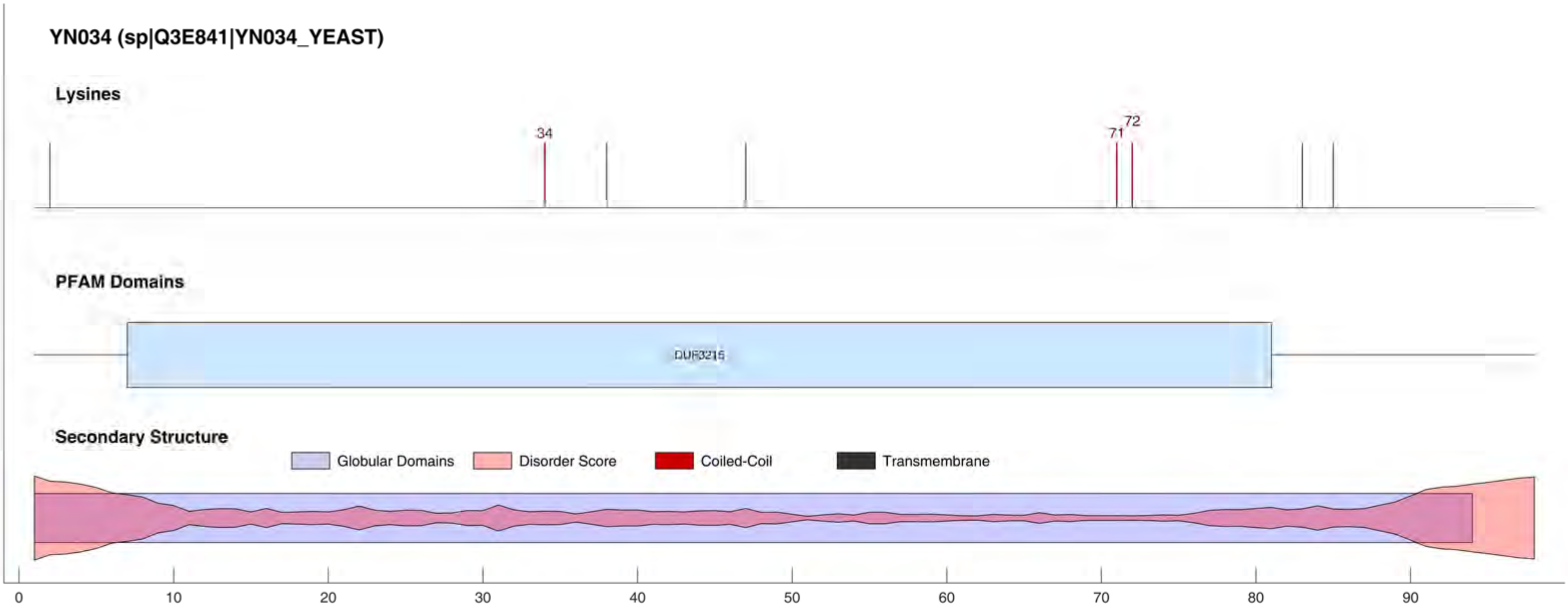
YN034 (sp|Q3E841|YN034_YEAST)
Lysines
34
71
72
PFAM Domains
DUF3215
Secondary Structure
Globular Domains
Disorder Score
Coiled-Coil
Transmembrane
0
10
20
30
40
50
60
70
80
90

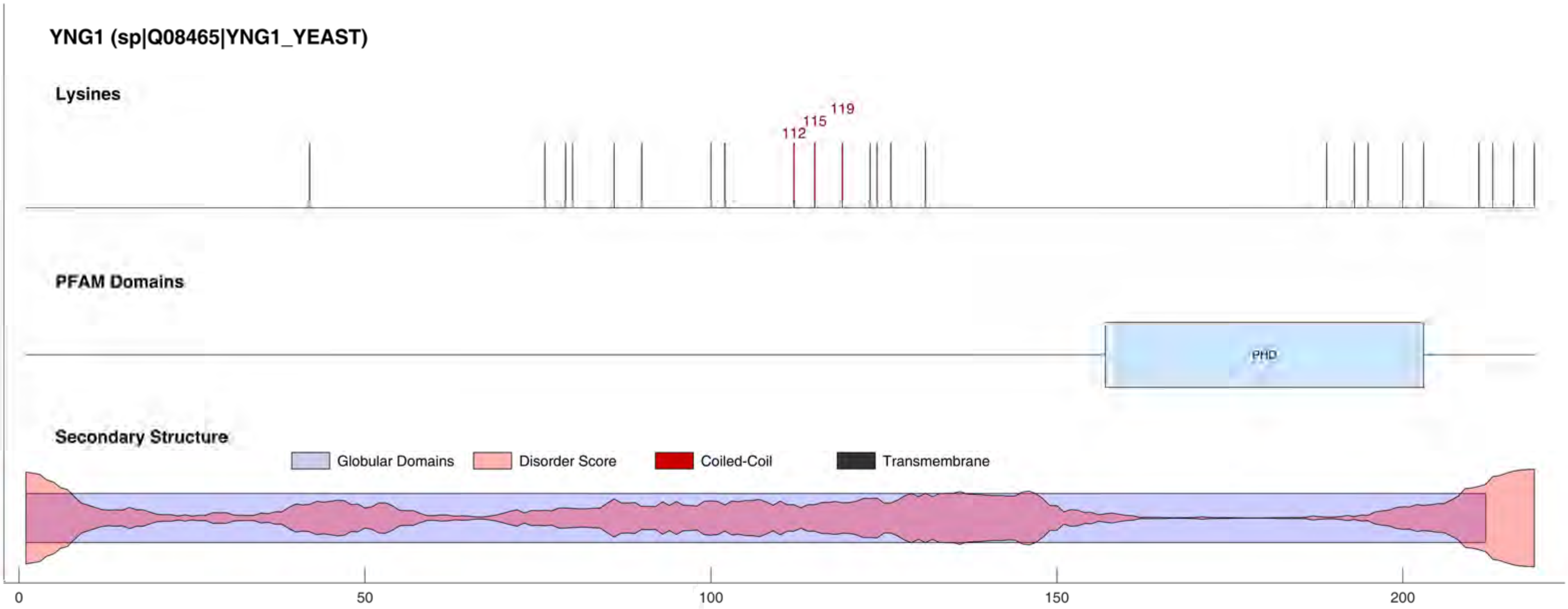
YNG1 (sp|Q08465|YNG1_YEAST)
Lysines
112
115
119
PFAM Domains
PHD
Secondary Structure
Globular Domains
Disorder Score
Coiled-Coil
Transmembrane
0
50
100
150
200

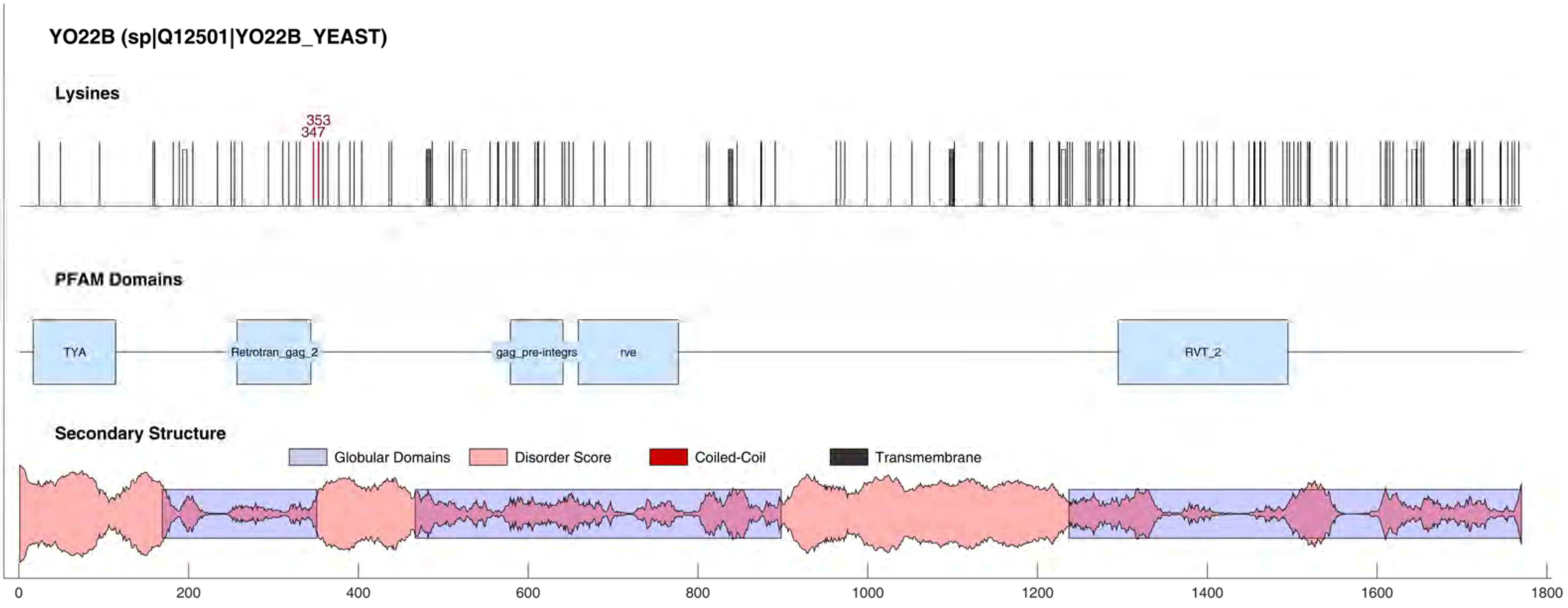
YO22B (sp|Q12501|YO22B_YEAST)
Lysines
353
347
PFAM Domains
TYA
Retrotran_gag_2
gag_pre-integrs
rve
RVT_2
Secondary Structure
Globular Domains
Disorder Score
Coiled-Coil
Transmembrane
0
200
400
600
800
1000
1200
1400
1600
1800

#### YP022 (sp|Q12139|YP022_YEAST)

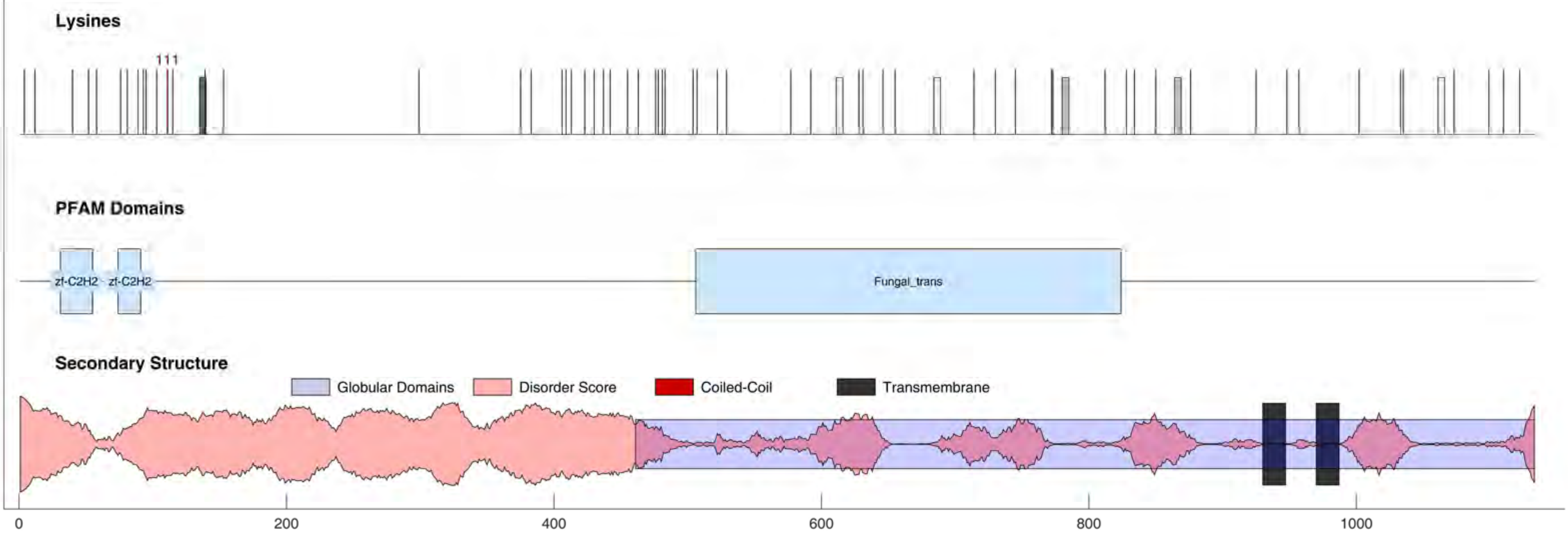

#### YP174 (sp|Q06616|YP174_YEAST)

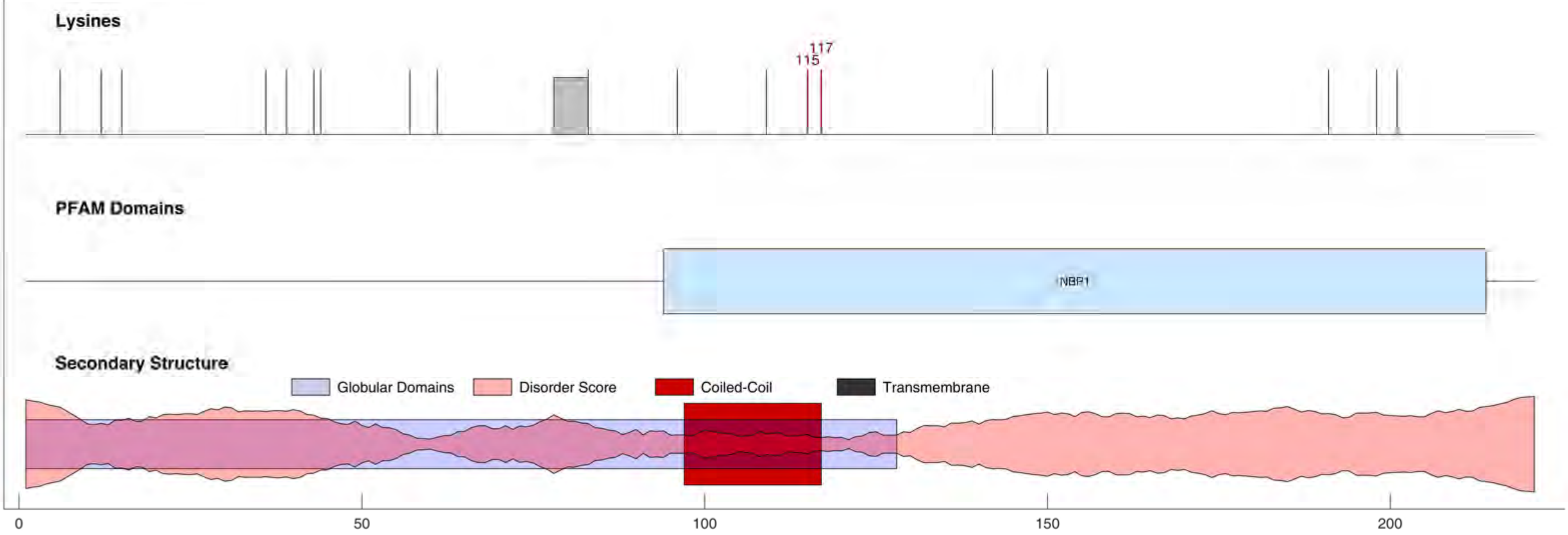

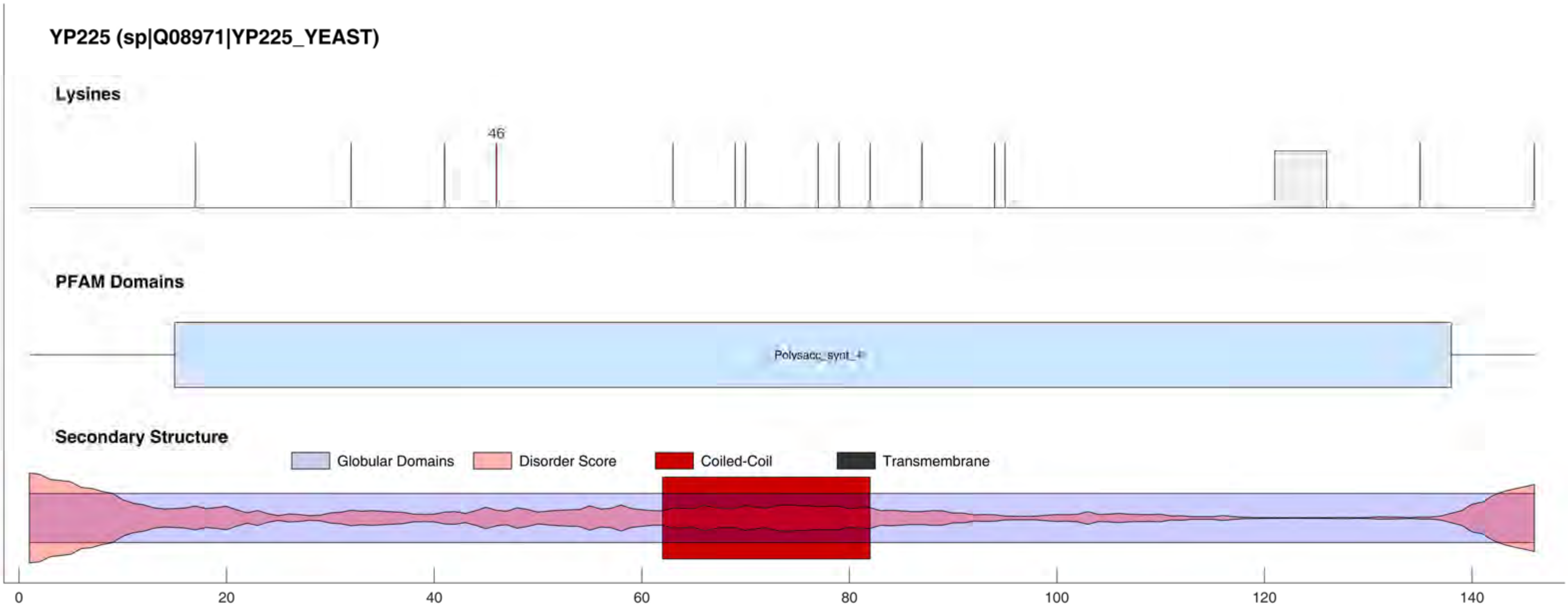
YP225 (sp|Q08971|YP225_YEAST)
Lysines
46
PFAM Domains
Polysacc_synt_4
Secondary Structure
Globular Domains
Disorder Score
Coiled-Coil
Transmembrane
0
20
40
60
80
100
120
140

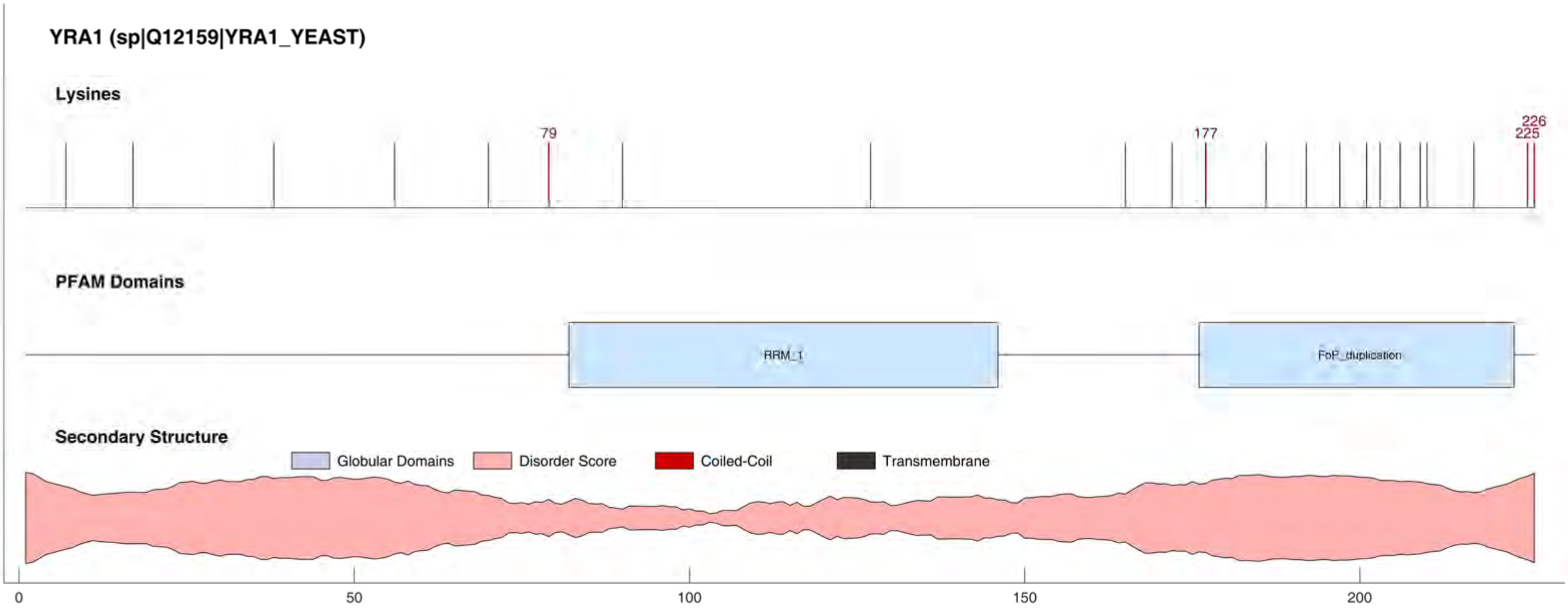

YRA1 (sp|Q12159|YRA1_YEAST)
Lysines
79
177
226
225
PFAM Domains
RRM_1
FoP_duplication
Secondary Structure
Globular Domains
Disorder Score
Coiled-Coil
Transmembrane
0
50
100
150
200

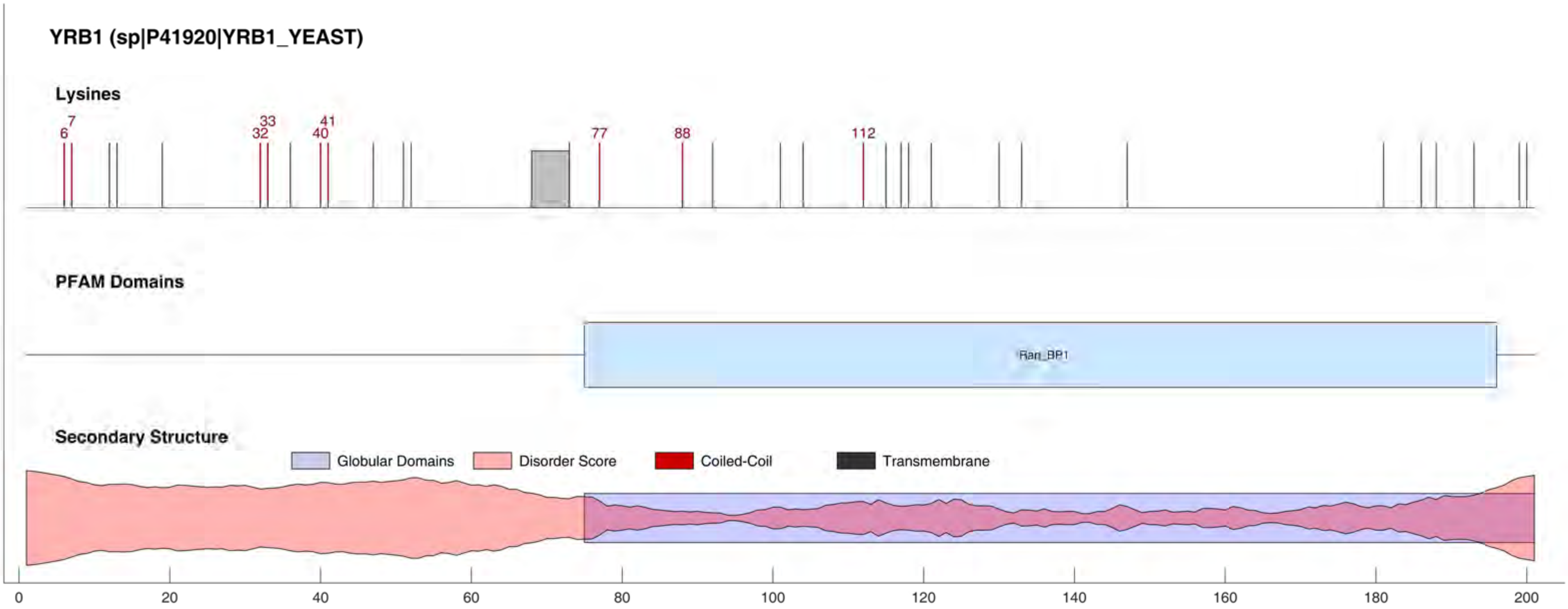
YRB1 (sp|P41920|YRB1_YEAST)
Lysines
6
7
32
33
40
41
77
88
112
PFAM Domains
Ran_BP1
Secondary Structure
Globular Domains
Disorder Score
Coiled-Coil
Transmembrane
0
20
40
60
80
100
120
140
160
180
200

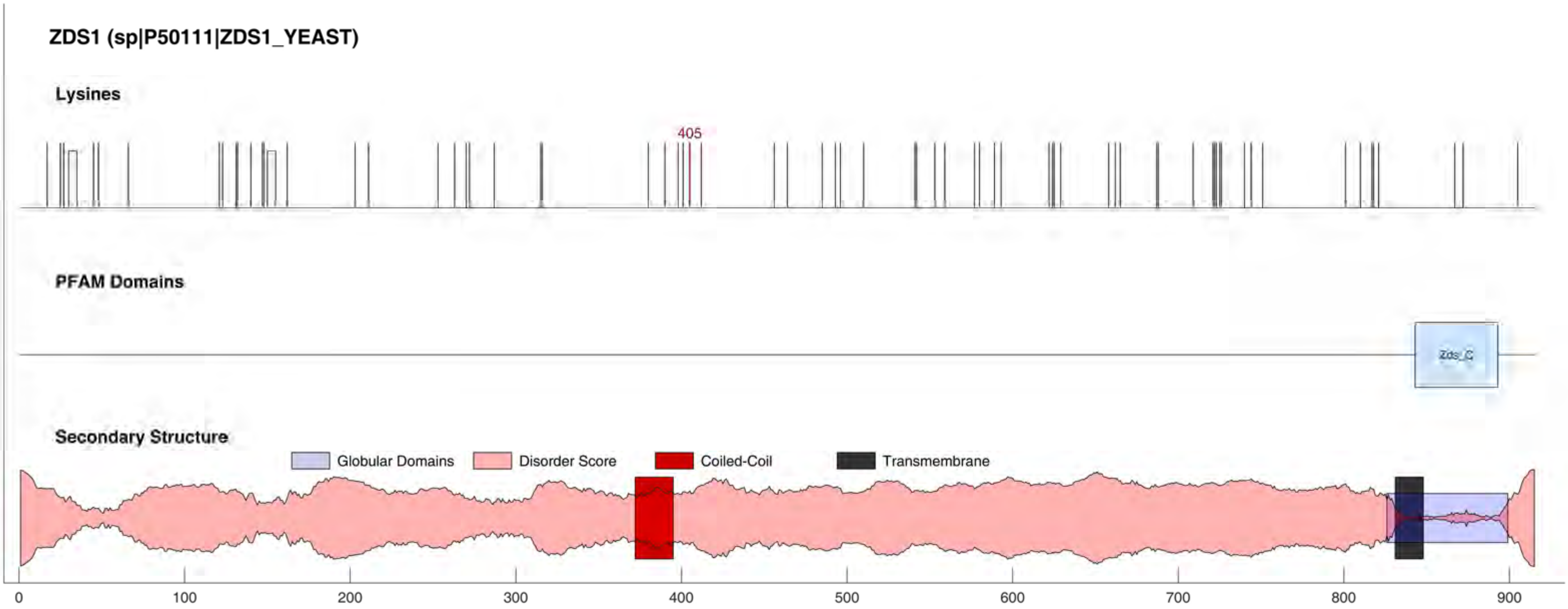
ZDS1 (sp|P50111|ZDS1_YEAST)
Lysines
405
PFAM Domains
Zds_C
Secondary Structure
Globular Domains
Disorder Score
Coiled-Coil
Transmembrane
0
100
200
300
400
500
600
700
800
900

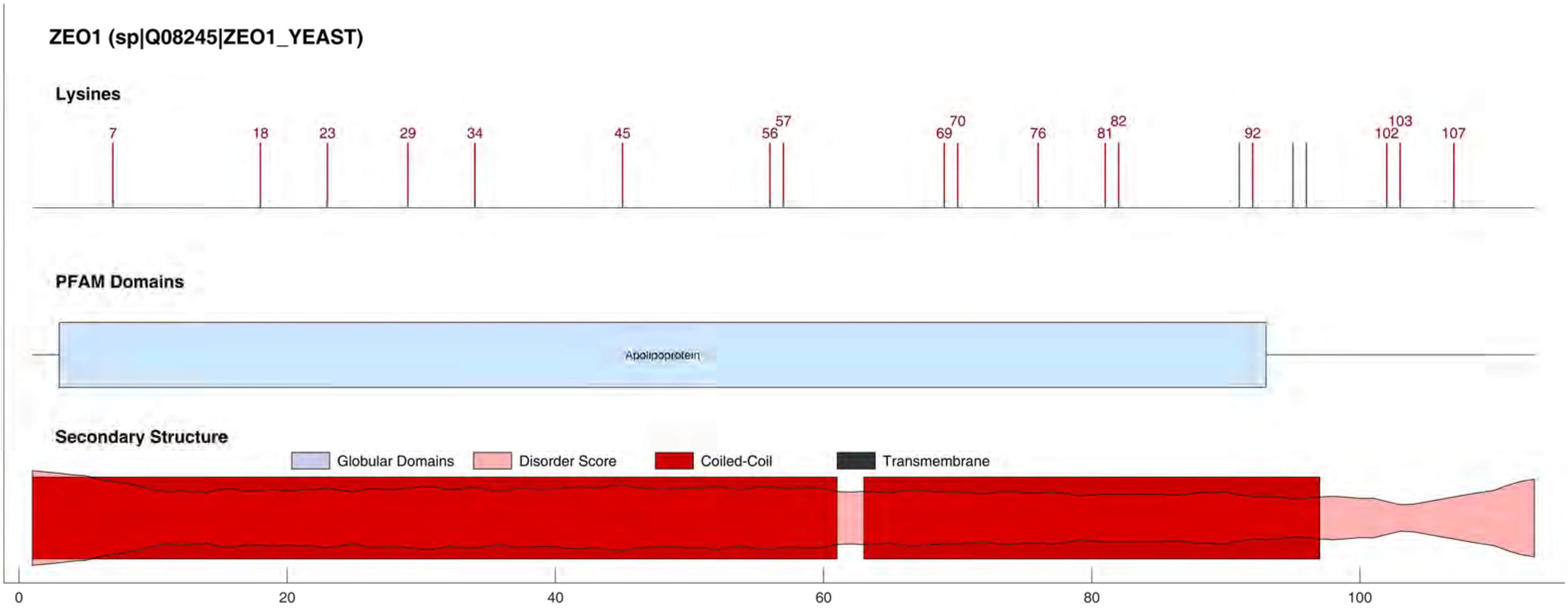
ZEO1 (sp|Q08245|ZEO1_YEAST)
Lysines
7
18
23
29
34
45
56
57
69
70
76
81
82
92
102
103
107
PFAM Domains
Apolipoprotein
Secondary Structure
Globular Domains
Disorder Score
Coiled-Coil
Transmembrane
0
20
40
60
80
100

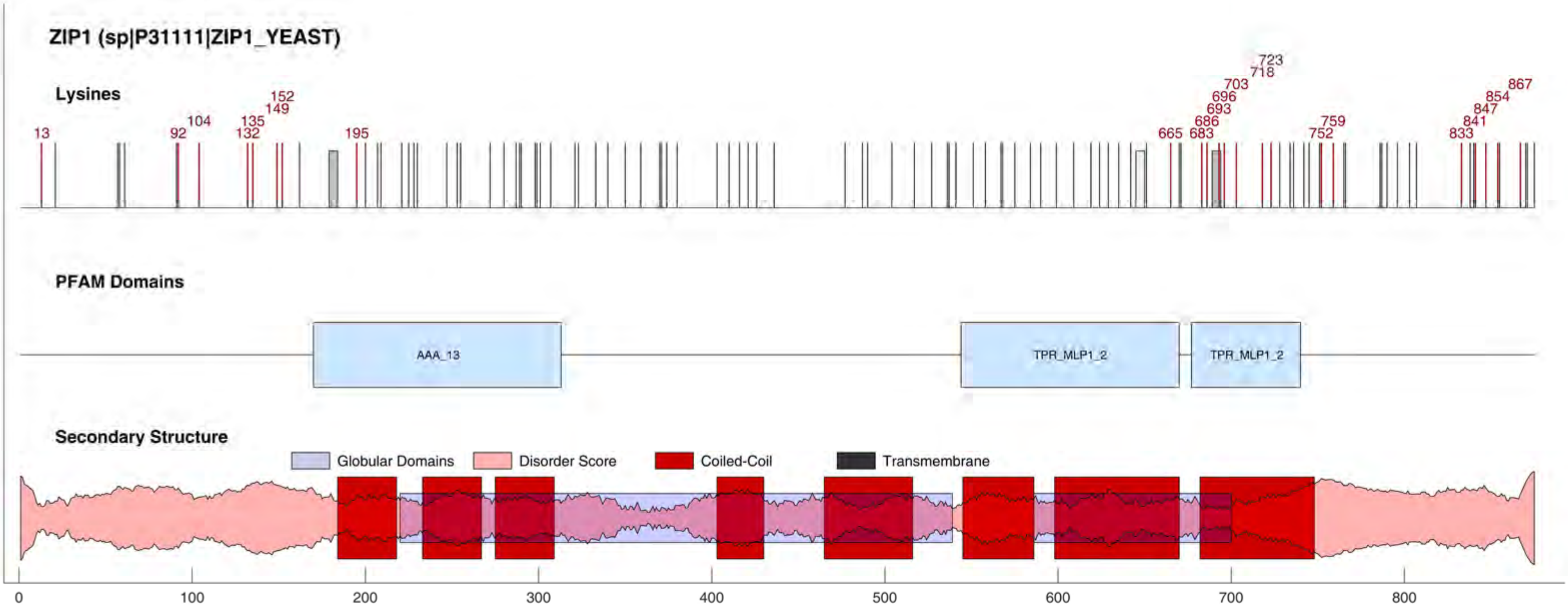
ZIP1 (sp|P31111|ZIP1_YEAST)
Lysines
13
92
104
132
135
149
152
195
665
683
686
693
696
703
718
723
752
759
833
841
847
854
867
PFAM Domains
AAA_13
TPR_MLP1_2
TPR_MLP1_2
Secondary Structure
Globular Domains
Disorder Score
Coiled-Coil
Transmembrane
0
100
200
300
400
500
600
700
800

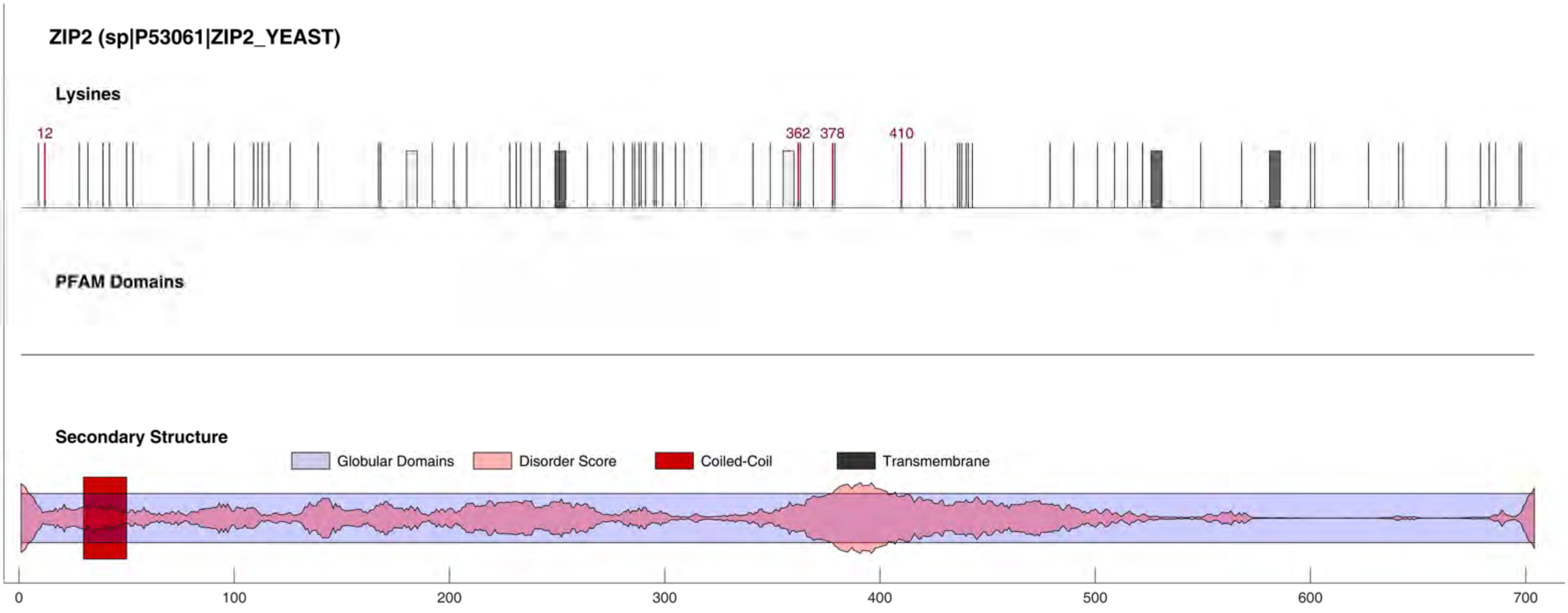
ZIP2 (sp|P53061|ZIP2_YEAST)
Lysines
12
362
378
410
PFAM Domains
Secondary Structure
Globular Domains
Disorder Score
Coiled-Coil
Transmembrane
0
100
200
300
400
500
600
700

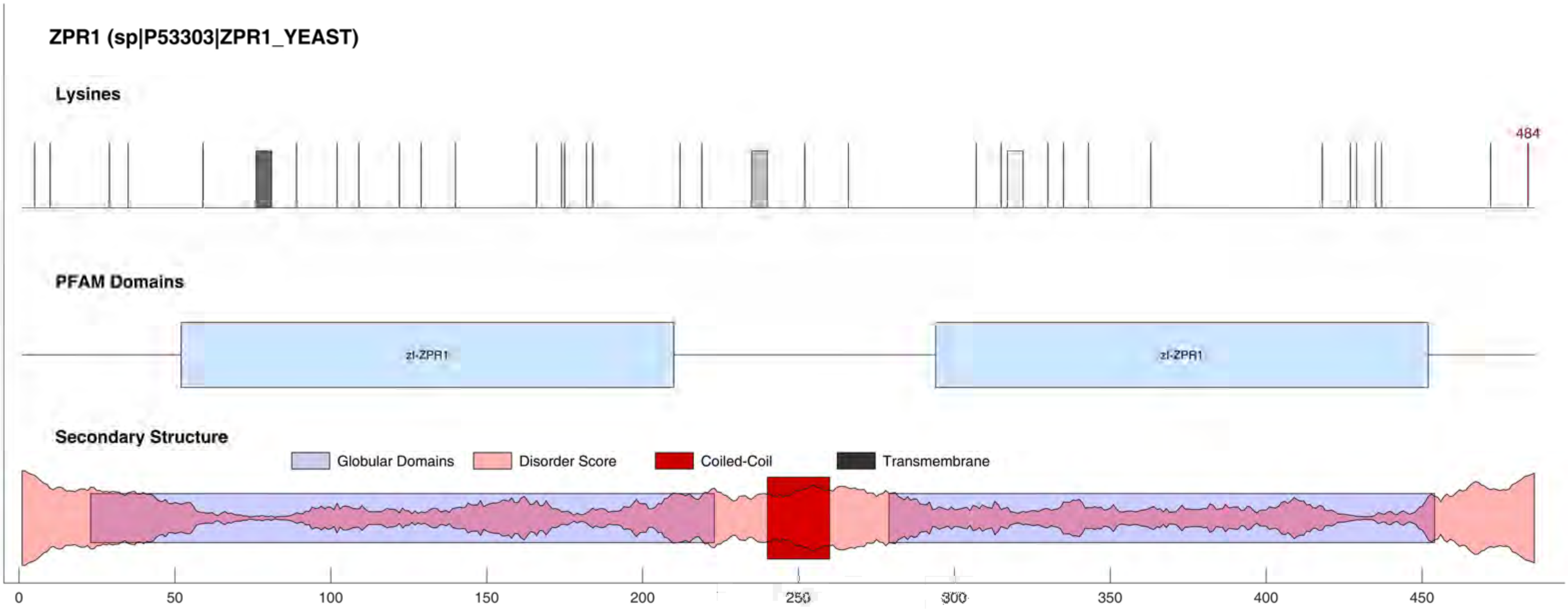

ZPR1 (sp|P53303|ZPR1_YEAST)
Lysines
484
PFAM Domains
zf-ZPR1
zf-ZPR1
Secondary Structure
Globular Domains
Disorder Score
Coiled-Coil
Transmembrane
0
50
100
150
200
250
300
350
400
450

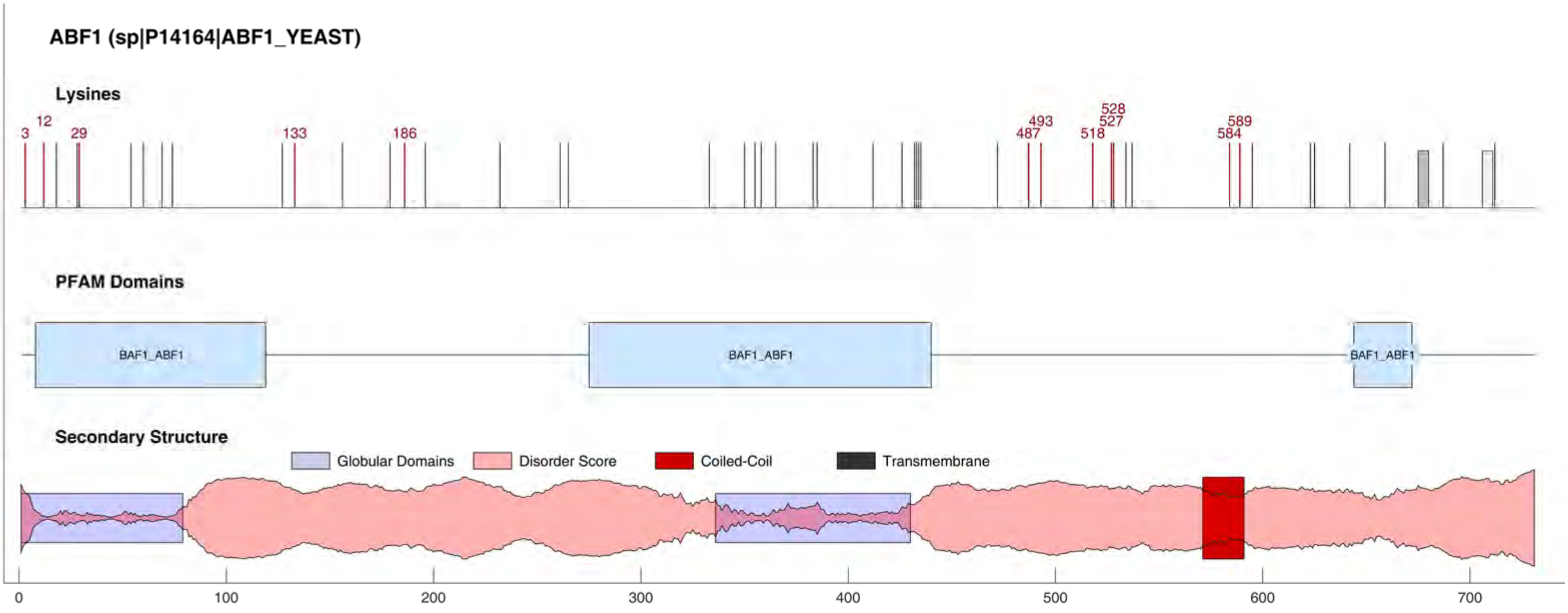
ABF1 (sp|P14164|ABF1_YEAST)
Lysines
3
12
29
133
186
487
493
518
527
528
584
589
PFAM Domains
BAF1_ABF1
BAF1_ABF1
BAF1_ABF1
Secondary Structure
Globular Domains
Disorder Score
Coiled-Coil
Transmembrane
0
100
200
300
400
500
600
700

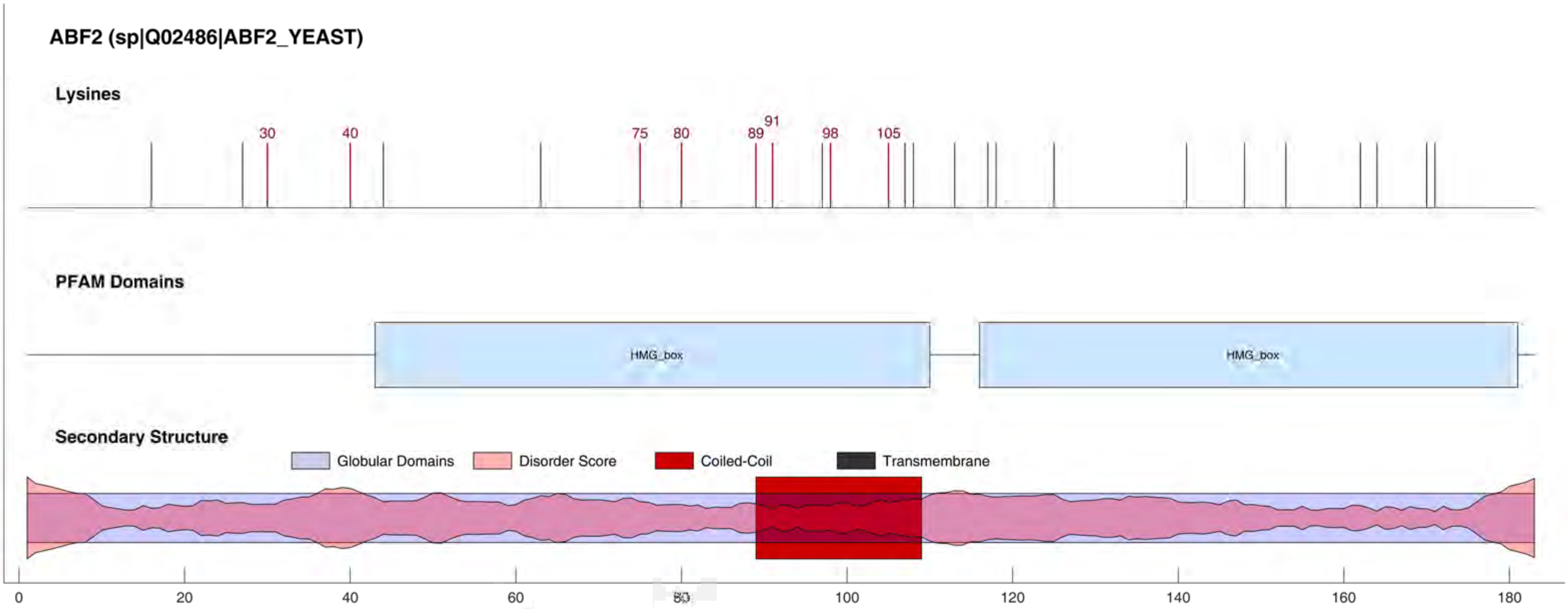

ABF2 (sp|Q02486|ABF2_YEAST)
Lysines
30
40
75
80
89
91
98
105
PFAM Domains
HMG_box
HMG_box
Secondary Structure
Globular Domains
Disorder Score
Coiled-Coil
Transmembrane
0
20
40
60
80
100
120
140
160
180

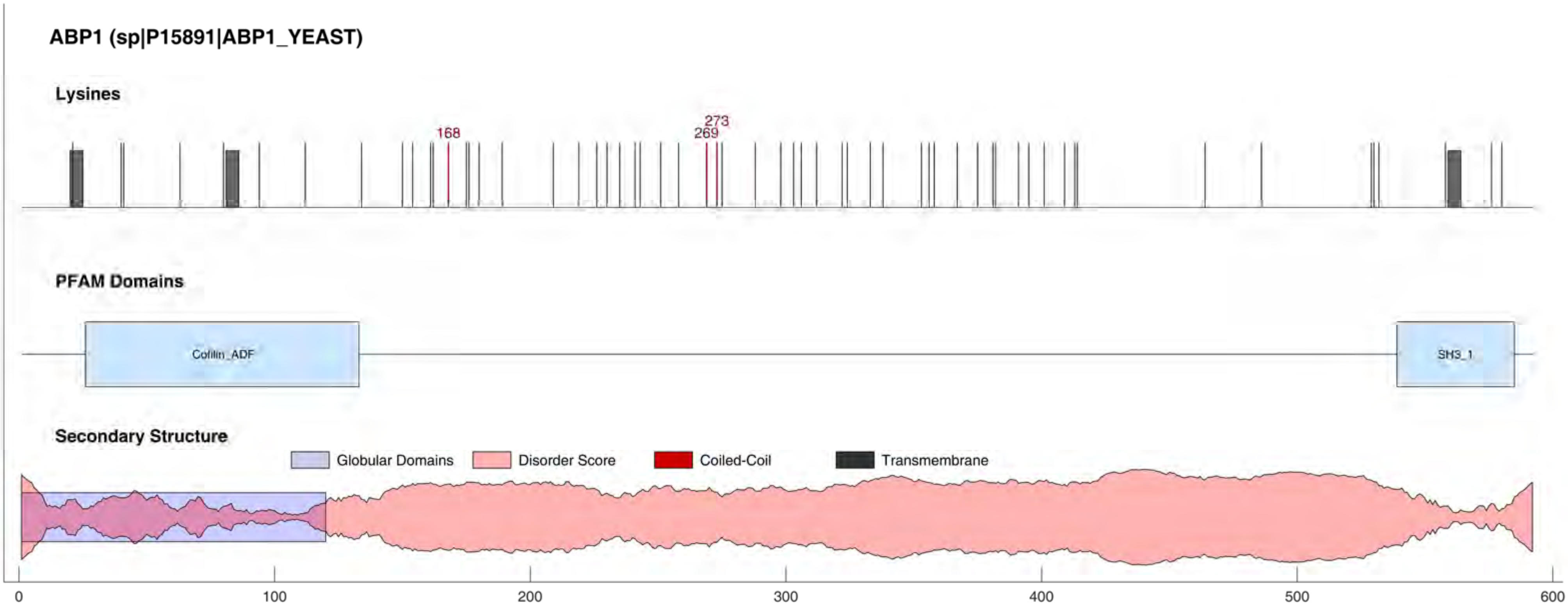

ABP1 (sp|P15891|ABP1_YEAST)
Lysines
168
273
269
PFAM Domains
Cofilin_ADF
SH3_1
Secondary Structure
Globular Domains
Disorder Score
Coiled-Coil
Transmembrane
0
100
200
300
400
500
600

#### ACA2 (sp|P40535|ACA2_YEAST)

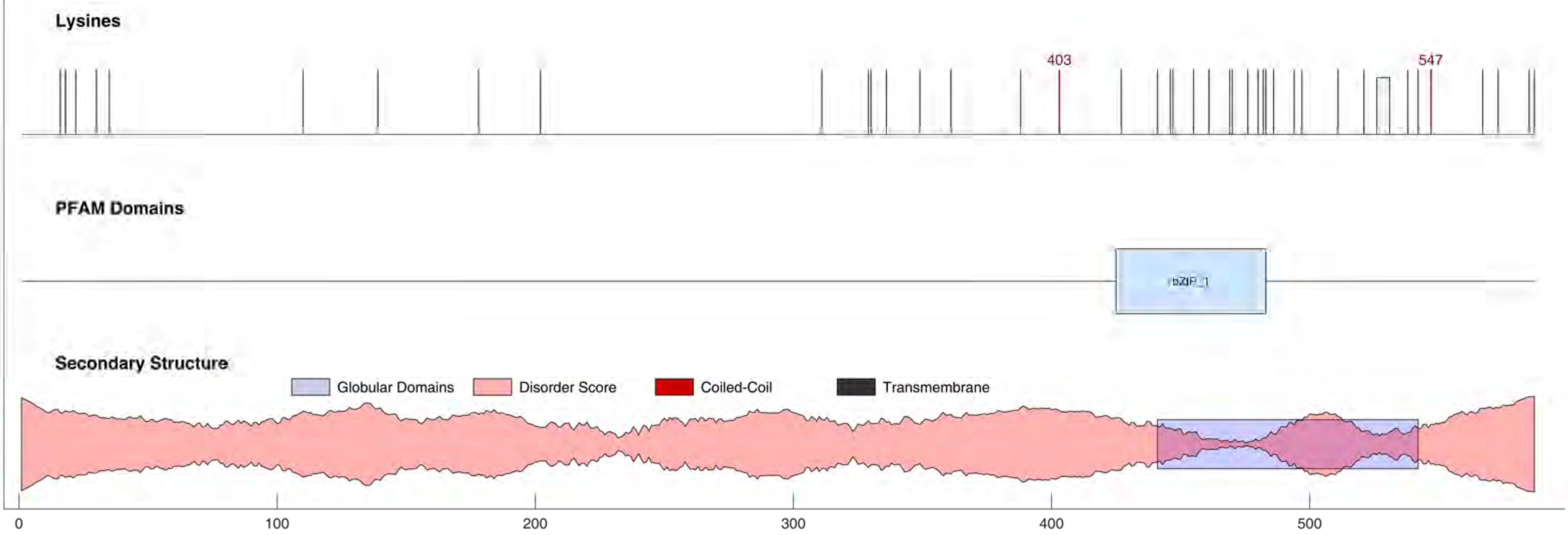

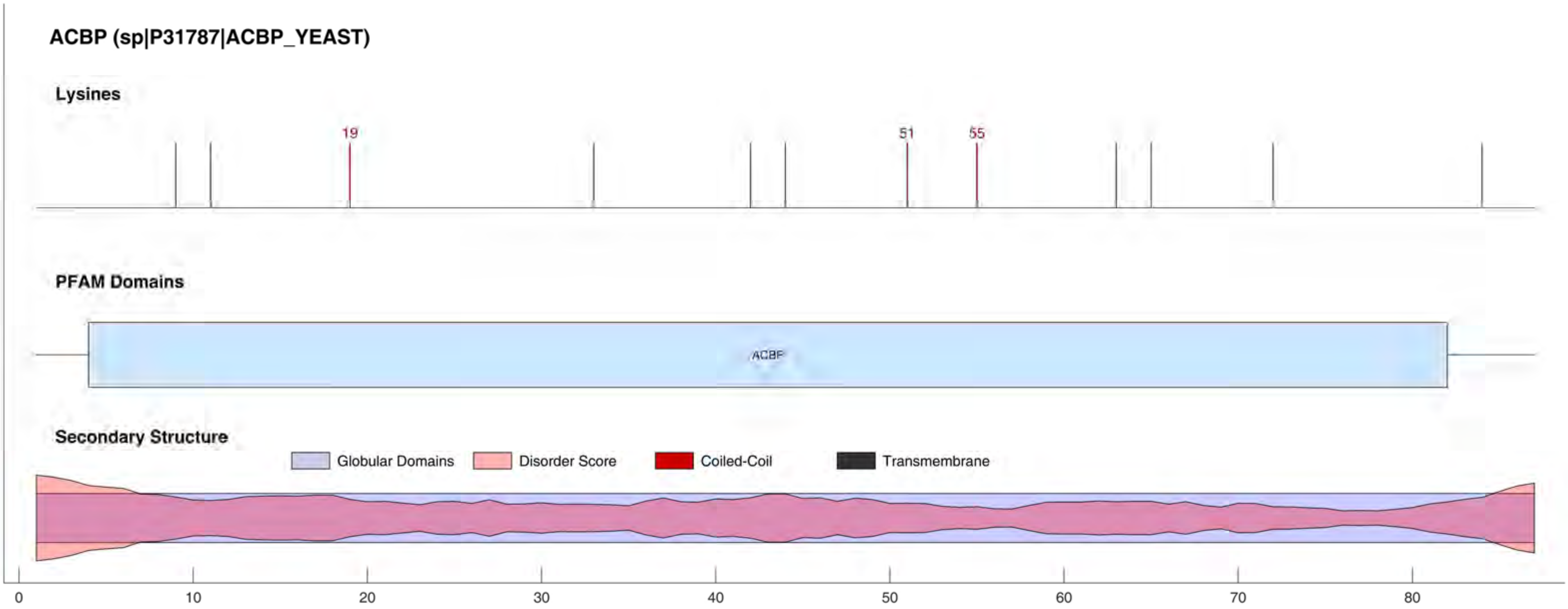
ACBP (sp|P31787|ACBP_YEAST)
Lysines
19
51
55
PFAM Domains
ACBP
Secondary Structure
Globular Domains
Disorder Score
Coiled-Coil
Transmembrane
0
10
20
30
40
50
60
70
80

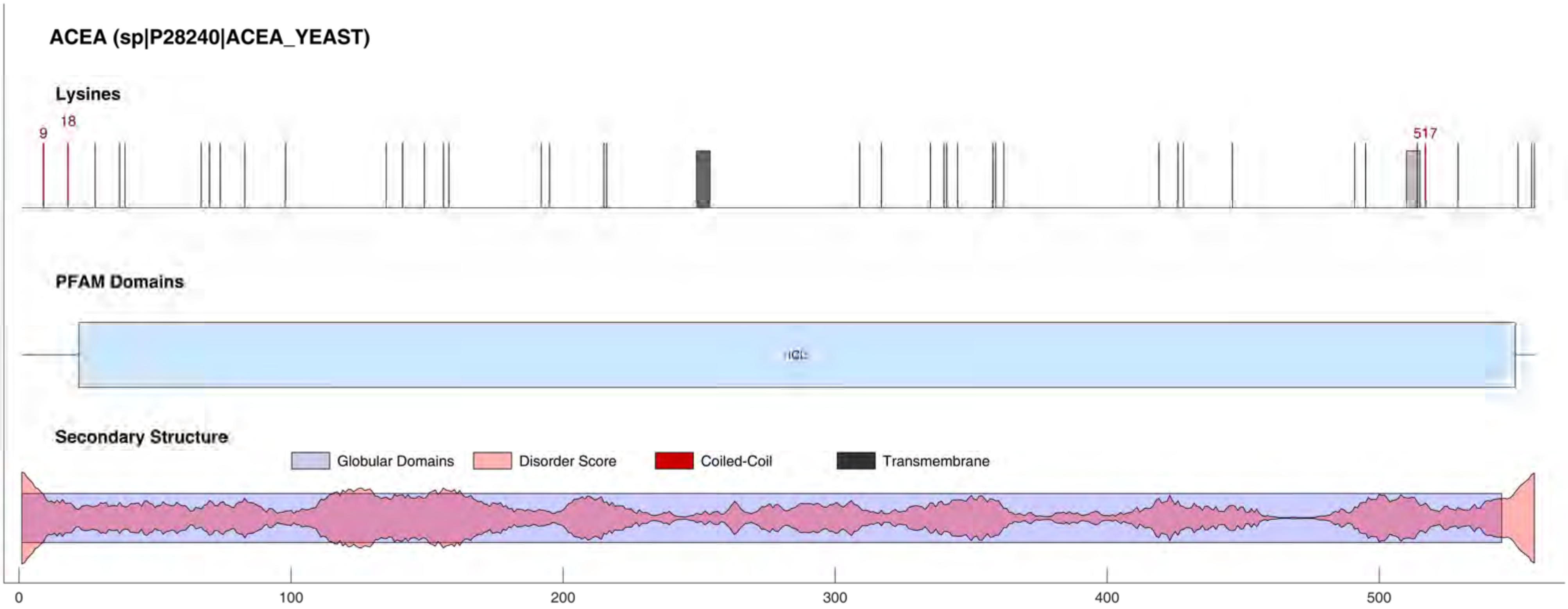
ACEA (sp|P28240|ACEA_YEAST)
Lysines
9
18
517
PFAM Domains
ICL
Secondary Structure
Globular Domains
Disorder Score
Coiled-Coil
Transmembrane
0
100
200
300
400
500

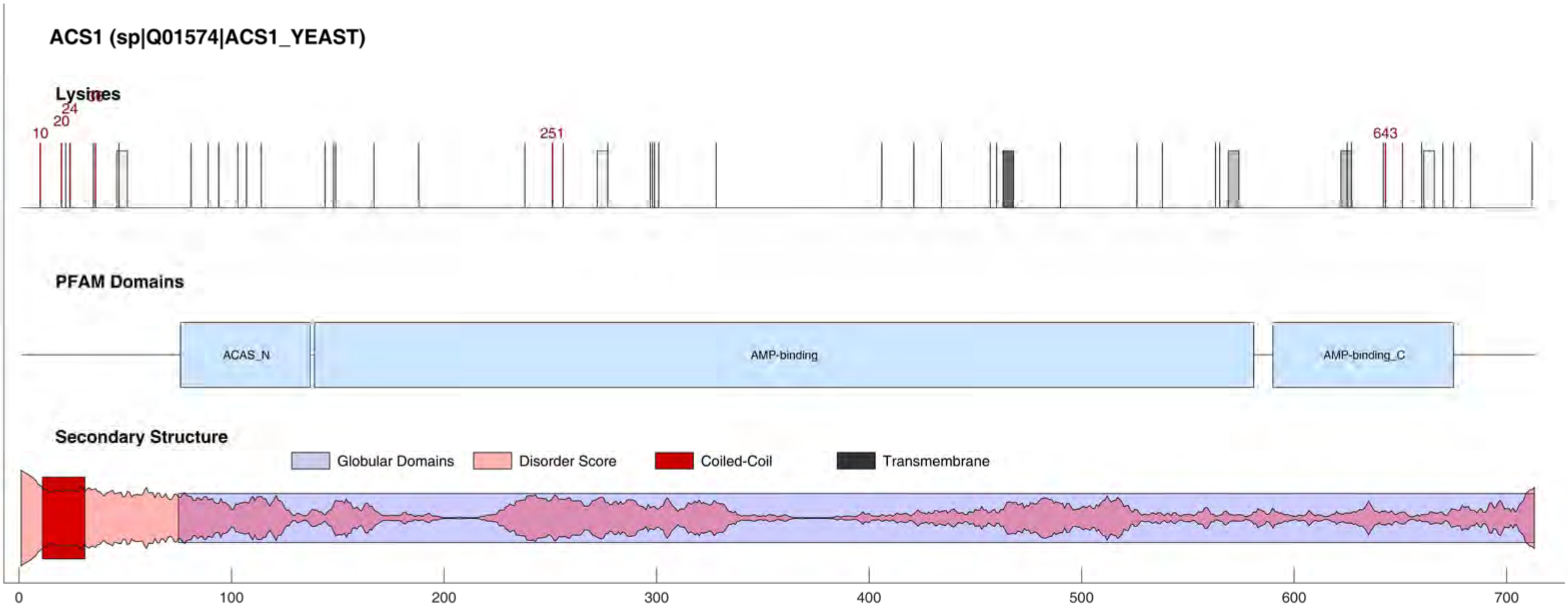
ACS1 (sp|Q01574|ACS1_YEAST)
Lysines
10
20
24
251
643
PFAM Domains
ACAS_N
AMP-binding
AMP-binding_C
Secondary Structure
Globular Domains
Disorder Score
Coiled-Coil
Transmembrane
0
100
200
300
400
500
600
700

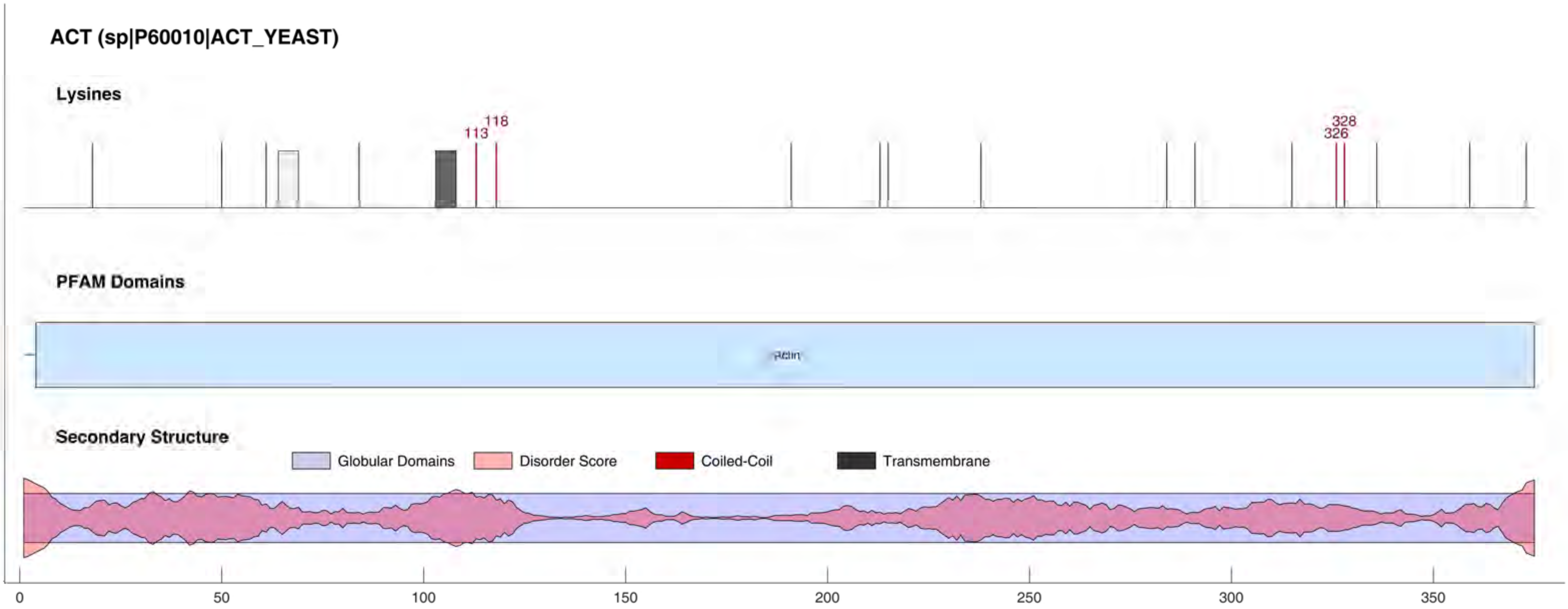
ACT (sp|P60010|ACT_YEAST)
Lysines
118
113
328
326
PFAM Domains
Secondary Structure
Globular Domains
Disorder Score
Coiled-Coil
Transmembrane
0
50
100
150
200
250
300
350
